# Cellular reprogramming for successful CNS axon regeneration is driven by a temporally changing cast of transcription factors

**DOI:** 10.1101/638734

**Authors:** Sumona P. Dhara, Andrea Rau, Michael J. Flister, Nicole M. Recka, Michael D. Laiosa, Paul L. Auer, Ava J. Udvadia

## Abstract

In contrast to mammals, adult fish display a remarkable ability to fully regenerate central nervous system (CNS) axons, enabling functional recovery from CNS injury. Both fish and mammals normally undergo a developmental downregulation of axon growth activity as neurons mature. Fish are able to undergo damage-induced “reprogramming” through re-expression of genes necessary for axon growth and guidance, however, the gene regulatory mechanisms remain unknown. Here we present the first comprehensive analysis of gene regulatory reprogramming in zebrafish retinal ganglion cells at specific time points along the axon regeneration continuum from early growth to target reinnervation. Our analyses reveal a regeneration program characterized by sequential activation of stage-specific pathways, regulated by a temporally changing cast of transcription factors that bind to stably accessible DNA regulatory regions. Strikingly, we also find a discrete set of regulatory regions that change in accessibility, consistent with higher-order changes in chromatin organization that mark (1) the beginning of regenerative axon growth in the optic nerve, and (2) the re-establishment of synaptic connections in the brain. Together, these data provide valuable insight into the regulatory logic driving successful vertebrate CNS axon regeneration, revealing key gene regulatory candidates for therapeutic development.

## Introduction

Damage to nerves in the CNS as a result of disease or injury most often results in a permanent loss of function in humans. The loss of function stems from the failure of adult mammalian CNS neurons to support regenerative axon growth. In adult mammalian retinal ganglion cell (RGC) neurons, genetic and pharmacologic manipulations of neuron-intrinsic pathways have shown promise in activating a regenerative state after optic nerve injury ^1–8^. However, even under these growth-enhanced conditions, regeneration mostly occurs only in a subset of RGCs ^2^, with the majority of axons rarely growing beyond a few millimeters. Furthermore, the regenerating axons frequently grow in an undirected manner, resulting in the “regenerated” axons terminating growth far from their appropriate brain targets ^5,8,9^. As such, there remains a gap in our understanding of the genetic programs that drive RGC axon regeneration culminating in target re-innervation and recovery of visual function.

In contrast to mammals, zebrafish display a remarkable ability to spontaneously regenerate RGC axons. As opposed to regenerating axons in the growth-enhanced mouse models, regenerating axons in the zebrafish optic nerve successfully navigate across the chiasm and reinnervate target neurons in the optic tectum ^10,11^, ultimately leading to functional recovery ^12^. It is well known that proteins regulating RGC axon growth and guidance in the developing visual system are highly conserved across vertebrate species, and are transcriptionally down-regulated once the mature circuitry has been established ^13,14^. In addition, like mammals, optic nerve injury in adult zebrafish induces the expression of axon growth attenuators such as *socs3* in RGCs ^6,15^, suggesting that expression of such negative regulators of axon growth is a normal part of the regenerative program. One major difference in response to CNS axon injury between mammals and fish is the re-expression of a genetic program that promotes axon growth and guidance ^16,17^. However, we and others have found that the regulation of axon growth-associated genes differs between development and regeneration ^18–20^. Thus, the difference between mammals and vertebrate species capable of optic nerve regeneration is their ability to reprogram adult RGCs for axon growth in response to optic nerve injury.

It has been established that specific regeneration-associated gene expression changes in the adult zebrafish retina begin within the first day after optic nerve crush and persist through the re-innervation of the optic tectum ^21^. What is less clear are the genome-wide changes in expression within the RGCs over time as they first grow toward their intermediate target of the optic chiasm and then navigate toward their principal brain target, the optic tectum. Here we present the first comprehensive analysis of gene expression changes (RNA-seq) and DNA regulatory element accessibility (ATAC-seq) in zebrafish RGCs at specific time points along the axon regeneration continuum from early growth to target reinnervation. Our analysis reveals that successful CNS axon regeneration is regulated by stage-specific gene regulatory modules, and punctuated by regeneration-associated changes in chromatin accessibility at stages corresponding to axonogenesis and synaptogenesis. Together, these data suggest candidates for gene regulatory targets for promoting successful vertebrate CNS axon regeneration.

## Results

### Stage-specific temporal changes in regeneration-associated gene expression

We hypothesized that regeneration-associated gene expression changes in axotomized RGCs would follow a temporal pattern corresponding to the changing requirements of axons as they grow through different environments leading from retina to optic tectum. The timing of successful axon regeneration after optic nerve crush in zebrafish is well characterized ^10,11,22^. To achieve a comprehensive picture of the genetic programming driving successful vertebrate CNS axon regeneration, we used RNA-seq to identify changes in gene expression that accompanied axon growth in regenerating RGCs at critical time points after optic nerve injury. We specifically examined how transcript expression in naïve retinas compared with that in retinas dissected from fish at 2, 4, 7, and 12 days post-injury (dpi). Based on the previously established regeneration chronologies, our chosen time-points (Fig. 1A) correspond to following stages of optic nerve regeneration: (1) axon growth past the site of injury toward the midline, (2) axon guidance across the midline, (3) selection of axon targets within the brain, and (4) synaptogenesis in the optic tectum ^10,11,22^.

**Fig. 1.**
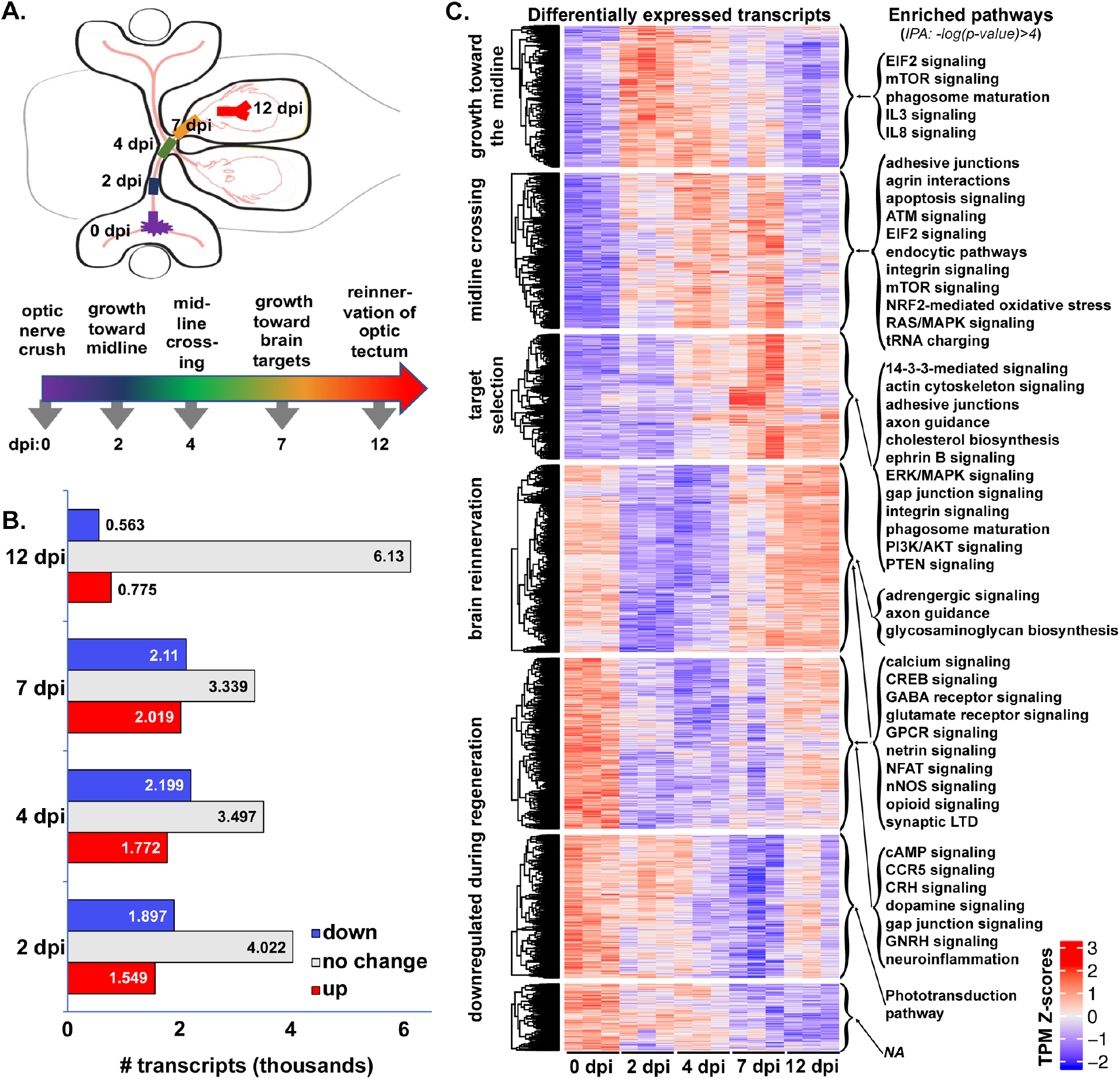
Regeneration-associated genes display stage-specific expression after optic nerve injury. (A) Schematic depicting stages of optic nerve regeneration from 0 – 12 days post-injury (dpi). (B) RNA-seq results generated from dissected retinas of adult zebrafish at 7-9 months of age. Transcripts from retinas dissected 2-12 days post optic nerve crush were compared with those from uninjured animals. Together, 7,080 transcripts were expressed at either higher (red = upregulated) or lower (blue = downregulated) levels in injured retina at 2, 4, 7, and/or 12 dpi when compared to control retinas (0 dpi), adjusted p values <0.05. (C) Temporally clustered transcripts demonstrate stage-specific enrichment of canonical pathways. Expression heatmaps produced from Z-scores (calculated using transcripts per million [TPM] estimates) of 4,614 differentially-expressed transcripts with adjusted p values <0.01. Mean normalized transcript counts were generated for each transcript across all samples at all time points. Each row represents a single transcript where each biological sample was compared to the mean, with red and blue indicating standard deviation above and below the mean, respectively. Each column represents one of the three biological replicates for each time point. Genes within each cluster were analyzed by Ingenuity Pathway Analysis (IPA) using *–*log(*p-value*)>4, to identify pathways expressed in a regeneration stage-specific manner. *NA*, not applicable, indicates that no pathways we detected that met our significance threshold.

Over the course of the first two weeks after optic nerve injury, we identified thousands of transcripts that displayed regeneration-associated changes in expression with respect to the baseline established from uninjured control retinas (Fig. 1B). Specifically, we found that 7,480 transcripts, roughly 19% of the retinal transcriptome, were differentially expressed in at least one time point (Table S1). At time points corresponding to periods of regenerative axon growth from retina to brain, 2-7 dpi, approximately three to four thousand transcripts were differentially expressed at either higher or lower levels in injured retina when compared to control retinas. At 12 dpi, a time point corresponding to the re-establishment of synaptic connections in the tectum, the number of differentially expressed transcripts was lower, but still exceeded 1,000. These results reveal substantial changes in gene regulatory programming over the course of regeneration.

To visualize the temporal patterns of transcript expression over the full course of axon regeneration from retina to brain, we identified the timing of peak expression for each of the differentially expressed transcripts. This was achieved by calculating the Z-score for normalized transcript counts of individual transcripts in each sample, comparing against the mean derived from all replicates across all time points. Transcripts were then clustered into seven groups based on temporal patterning of Z-scores using the K-means algorithm (Fig. 1C, *Differentially expressed transcripts;* Table S2). As hypothesized, we detected distinct clusters of transcripts that were upregulated in response to injury that displayed peak expression at early, intermediate and late time points during regeneration. We also observed transcripts that were expressed at their highest levels in the uninjured retina and down-regulated early, midway, and later in the regenerative time course. Finally, we observed transcripts expressed in uninjured retina that were down-regulated early in regeneration, but displayed peak expression at the latest stage when regenerating fibers are in the process of synaptogenesis. The temporal patterning of the differentially regulated genes signifies a dynamic program of gene regulation that changes over the course of regeneration.

Given the temporal dynamics in gene expression associated with different stages of regeneration, we queried the data for evidence of stage-specific processes that drive successful regeneration. To identify canonical pathways represented by differentially expressed transcripts in each cluster, we conducted Ingenuity Pathway Analysis (Qiagen, Redwood City, CA). Enriched pathways, based on high stringency criteria (−log(*p*-value) > 4), were detected for six out of the seven temporal clusters (Fig. 1C, *Enriched pathways*; Table S3). We found little overlap in enriched pathways between the first three clusters, consistent with the idea of distinct processes active at different stages of regeneration.

Many of the enriched pathways at each time point were consistent with pathways previously associated with axon regeneration and/or developmental wiring of the nervous system. For example, the enrichment of pathways involved in mTOR and cytokine signaling early in regeneration is consistent with results in mammalian models in which activation of these pathways enhances axon growth and survival of RGCs after optic nerve crush ^4,6,7^. Interestingly, PTEN signaling pathway genes are enriched at an intermediate timepoint, consistent with the down-regulation of mTOR observed as a normal part of the regenerative program in zebrafish ^23^. At timepoints associated with midline crossing and target selection, we also found an enrichment of pathways involved in cytoskeletal regulation, cell-cell adhesion, and cell-substrate interactions, as would be expected during axon growth and guidance. Simultaneously we see pathways associated with neurotransmitter receptor signaling and G-protein coupled receptor signaling that are downregulated during stages of axon growth and guidance, but are upregulated during synaptogenesis. It should be noted that genes that are down regulated in response to injury and re-expressed at later stages of regeneration may be contributing to the recently characterized RGC dendritic remodeling that is necessary for successful regeneration ^22^. We also observed downregulation of neuroinflammatory pathways early in regeneration, which may contribute to successful axon regeneration in zebrafish. Local inflammatory responses have previously been demonstrated to accelerate axon growth, suggesting that there are specific inflammatory pathways that promote regeneration and others that are inhibitory. Taken together, these results demonstrate that post-injury transcriptional programming dictates the timing of stage-specific processes associated with successful CNS axon regeneration.

### Changes in transcription factor expression, not chromatin accessibility, are associated with temporal patterning of regeneration-associated transcripts

In order to understand the regulatory system that guides the post-injury transcriptional program, we identified putative regulatory DNA elements (promoters, enhancers and insulators) in RGCs at different stages of regeneration. Assay for Transposase Accessible Chromatin sequencing (ATAC-seq) ^24^, was employed to assess chromatin accessibility in nuclei isolated from RGCs. Transgenic zebrafish expressing GFP under the regulation of the *gap43* promoter/enhancer ^25^ enabled the specific isolation of RGCs from dissociated retinal cells at specific post-injury time points using fluorescence activated cell sorting (FACS).

We detected over 40,000 peaklets representing consensus regions of accessible chromatin in RGCs. The majority of peaks were located in intergenic regions, upstream or downstream from annotated genes (Fig. 2A). Of the peaks located within genes, the vast majority were found within non-coding sequences (Fig. 2A). Approximately 80% of accessible chromatin peaklets were found distal to any annotated genes (Fig. S1), consistent with potential function as enhancers or insulators. Correspondingly, the remaining approximately 20% of peaklets overlapped with 5’ UTRs or were located within 1kb of transcriptional start sites, consistent with potential function as promoters (Fig. S1). These results are consistent with previous findings associating chromatin accessible regions with proximal and distal gene regulatory elements ^26^.

**Fig. 2.**
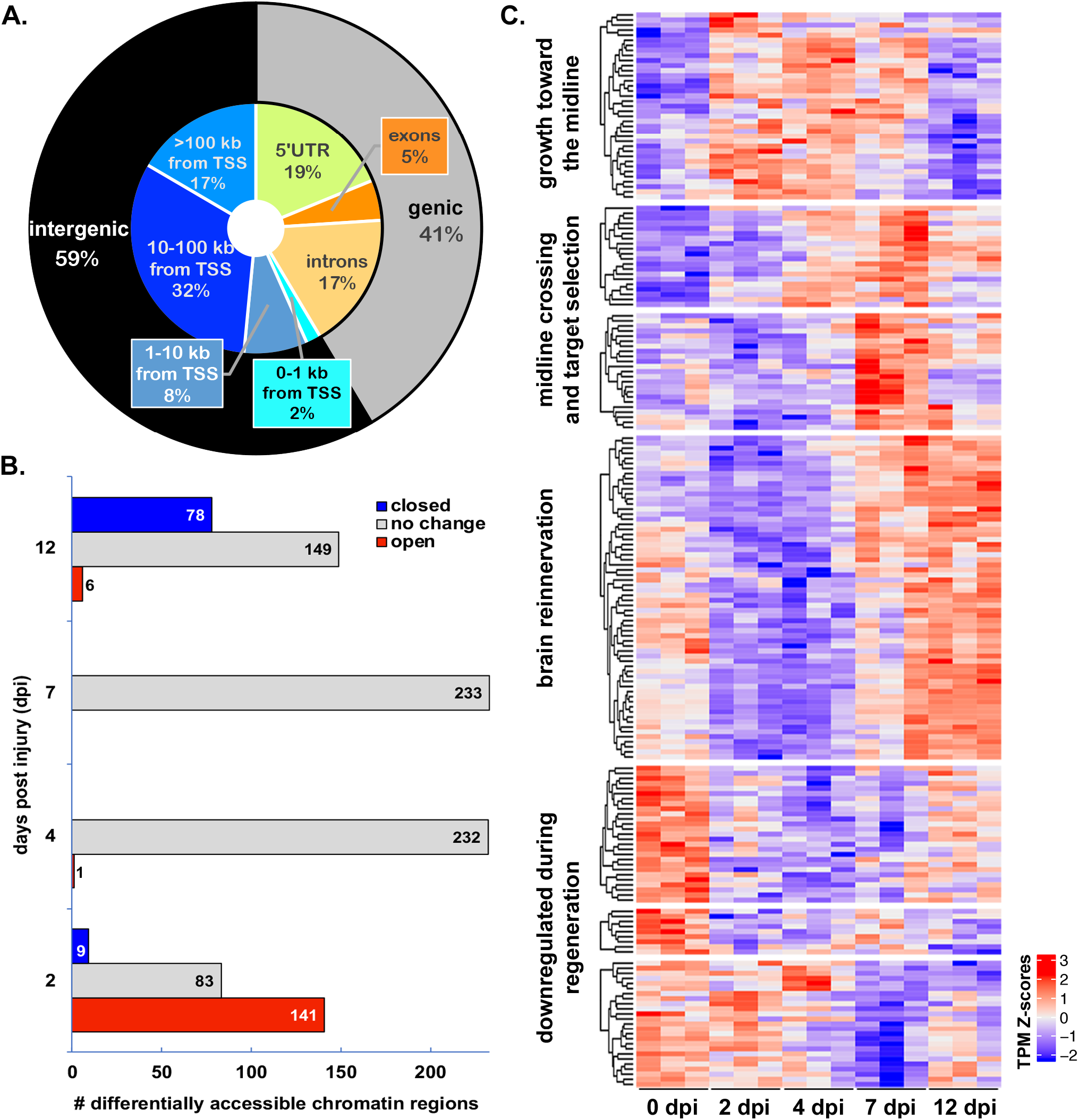
Regeneration stage-specific gene expression changes are correlated with temporal changes in transcription factor expression rather than changes in chromatin accessibility. (A) Genomic distribution of 42,198 high confidence regions of accessible chromatin, identified using ATAC-seq on samples of RGCs isolated from control and regenerating retinas using fluorescence activated cell sorting (FACS). High confidence regions consist of 500 bp sequences surrounding peaklet summits with *p*-value < 10^−10^. The genic region includes 5’-untranslated regions (UTR), exons and introns. The intergenic region consists of sequences found between annotated genes. (B) Most transcriptionally accessible regions do not change over the course of regeneration when compared to uninjured control (0 dpi). Differentially accessible regions (233 total) were observed primarily at early (2 dpi) and late (12 dpi) stages of regeneration. (C) Temporal progression of transcription factors characterizes the distinct stages of regeneration. The expression heatmap of 205 differentially expressed transcripts represents 159 unique transcription factor (TF) genes. TSS, transcriptional start site.

Surprisingly, overall chromatin accessibility changed very little in response to optic nerve injury. In fact, only 233 consensus regions of accessible chromatin (0.5%) were differentially accessible in injured RGCs compared to control RGCs (Fig. 2B; Table S4). All but one of the differentially accessible peaklets were found at 2 dpi or 12 dpi. Most of the differentially accessible peaklets at 2 dpi were differentially open, while most of those at 12 dpi were differentially closed. Furthermore, the overlap between differentially accessible peaklets between the two time points consisted of only two peaklets that were differentially open at both 2 and 12 dpi. Together these results suggest that the accessibility of DNA regulatory elements in RGCs is relatively constant, even under conditions that result in dynamic changes within the transcriptome. Thus, we predicted that the availability of the transcription factors that bind to RGC DNA regulatory elements, rather than the accessibility of the elements, must change over the course of regeneration.

To test the hypothesis that transcription factor expression is differentially regulated at different stages of regeneration we identified transcription factors that displayed differential expression at any point over the regeneration time course. We achieved this by cross-referencing our transcriptomic data to a recently compiled list of human transcription factors ^27^ and used a global differential (likelihood ratio test, LRT) to compile a list of transcription factor-encoding transcripts associated with regeneration. We discovered 265 definitive transcription factor encoding genes associated with 339 transcripts that were differentially expressed at one or more post-injury time points (Fig. S2, Table S5). We further refined this list to 205 transcripts corresponding to 159 transcription factor encoding genes with defined DNA recognition motifs available in the JASPAR ^28^ or CIS-BP ^29^ databases. As predicted, we found that differentially expressed transcription factors display temporal clustering similar to that of the regeneration-associated differentially expressed transcripts at large (Fig. 2C; Table S6).

Thirty transcription factor families are represented among the transcription factors that were differentially expressed during regeneration. Over half of the differentially expressed transcription factors fall into four families (basic leucine zipper, basic helix-loop-helix, 2-Cys-2-His zinc finger, and homeodomain). Each of these four families of transcription factors included representatives with peak expression early, middle, and late in the regenerative process (Fig. 3), as well as those whose expression was down-regulated during regeneration (Figs S3-S6). Within a given transcription factor family, there are frequently binding motif similarities shared between members. However, since transcription factors in these families may function as homo- and/or heterodimers, their binding site affinity and transactivation ability may vary. Thus, a changing cast of transcription factors would have the power to regulate the temporal progression of regeneration-associated gene expression through a common set of accessible regulatory regions.

**Fig. 3.**
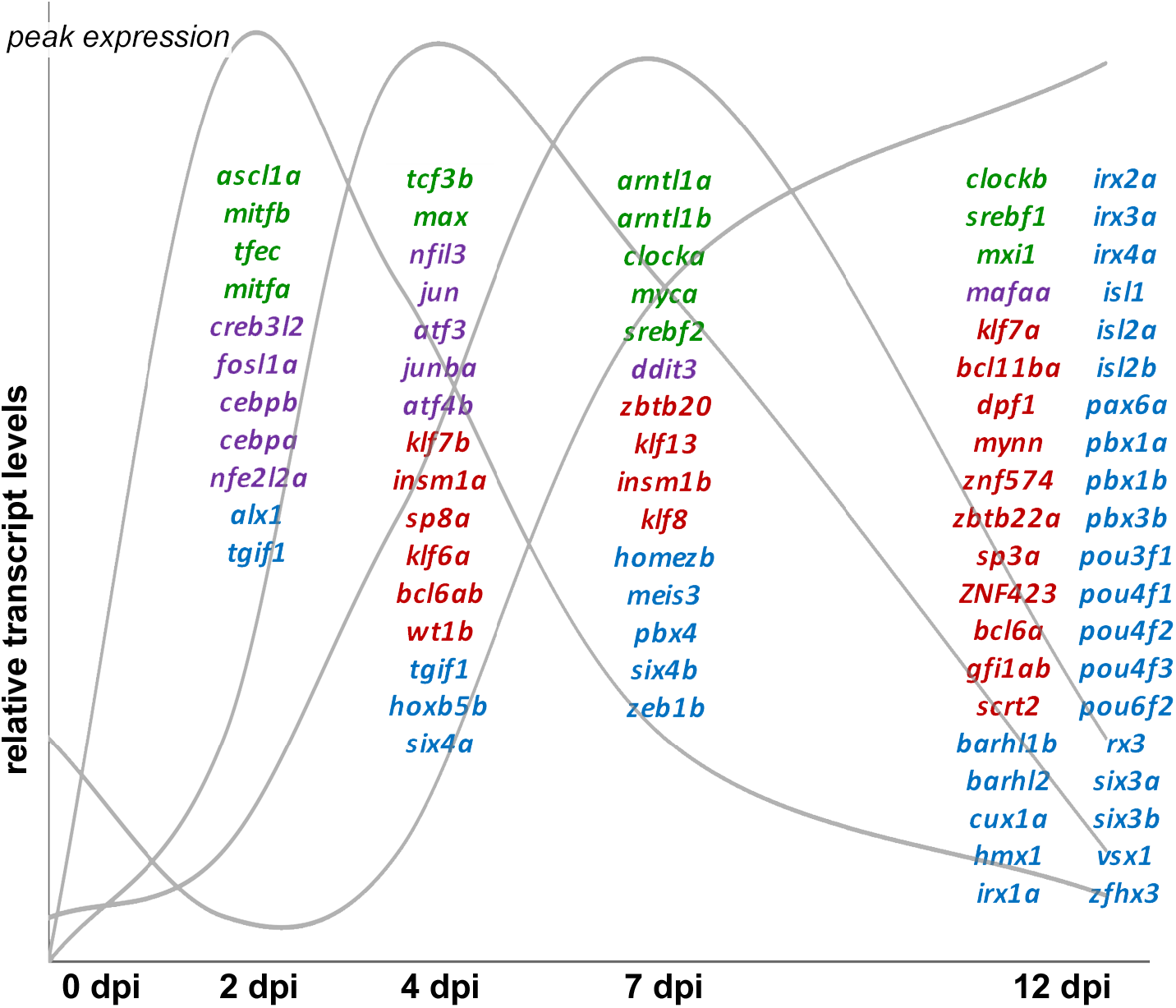
The majority of regeneration-associated transcription factors fall within four transcription factor families. Select transcription factor-encoding transcripts from the helix-loop-helix (green), leucine zipper (purple), C2H2 zinc finger (red), and homeodomain families (blue) transcription factor families grouped according to their peak expression (schematically represented by grey curves). Although transcription factors from each of the families are represented at different stages in regeneration, leucine zipper transcription factors appear more prominent early in regeneration and homeodomain transcription factors are most prevalent later in regeneration.

### Binding sites for differentially expressed transcription factors are enriched in inferred RGC regulatory elements detected by chromatin accessibility

To evaluate the potential for temporally expressed transcription factors to regulate stage-specific regeneration-associated gene transcription, we used motif enrichment analysis ^30^. For each cluster of temporally expressed transcripts we identified potential regulatory elements in the form of accessible chromatin peaklets that were proximal (peaklet center within ±1kb from transcription start site) or distal (peaklet center within ±100 kb from transcription start site, but not proximal) to the associated gene. Proximal and distal peaklet sequences were then queried for enrichment of binding motifs of transcription factors with similar temporal expression profiles (Table S7). For transcripts whose expression is upregulated early in regeneration (Fig. 1C, *growth toward the midline* cluster), we found that motifs for 17 out of the 31 transcription factors queried were enriched in the surrounding regions of accessible chromatin (Fig. 4A; Table S7). Motifs for the zinc finger transcription factors, KLF6 and WT1, were detected in over 70% of both proximal and distal elements. Including KLF6, a number of previously identified regeneration-associated transcription factors are both upregulated early in optic nerve regeneration (Fig. 2C) and have binding sites that are enriched within putative regulatory elements surrounding genes that are also upregulated early in regeneration (Fig. 4A, Fig. 1C). Most notable among these are ASCL1 and the basic leucine zipper (bZIP) factors JUN, ATF3, FOSL1, and JUNB. In addition, other transcription factors whose binding sites are enriched include those that have previously been associated with axon growth in developing CNS neurons, such as WT1 and TCF3 ^31,32^.

**Fig. 4.**
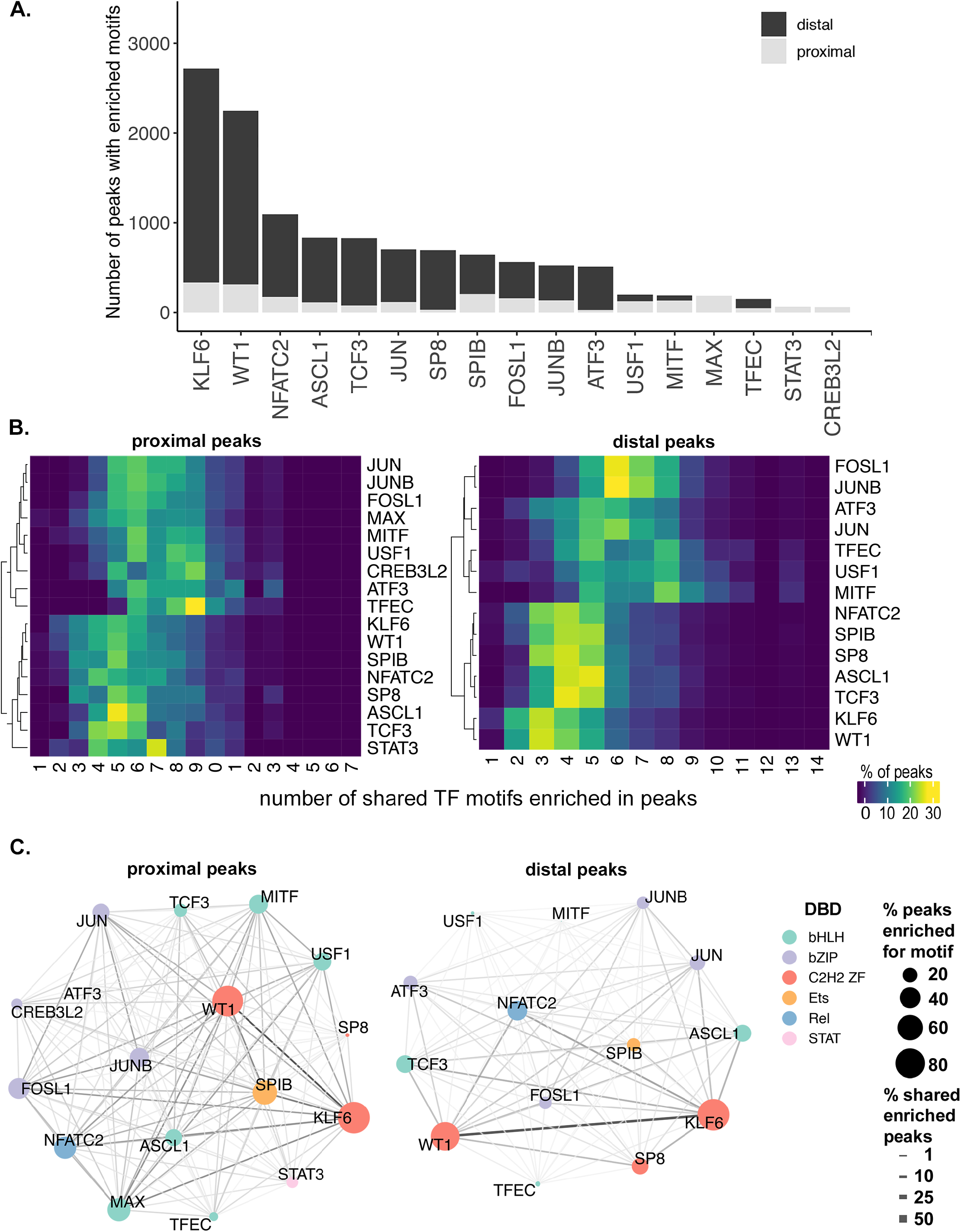
Potential regulatory interactions between regeneration-associated transcription factors and putative promoters and enhancers. (A) Stacked bar graph of number of peaks enriched with transcription factor (TF) motif in the proximal (gray) and distal (black) sequences for each TF in the cluster axon growth towards midline (cluster 1 in Fig.1). (B) Heat map of shared TF motif enrichment in cluster1 accessible peaks. X-axis = number of shared TF motifs enriched in peaks, Y-axis = TF. Heatmap colors based on % of total peaks enriched for given TF. (C) Pair-wise co-occurrence of TF motifs found in the proximal and distal accessible regions surrounding the differentially expressed transcripts of cluster 1. Node color corresponds to TF family based on DNA-binding domain (DBD). bHLH, basic helix-loop-helix; bZIP, basic leucine zipper; C2H2 ZF, two cys, two his zinc finger; Ets, E26 transformation-specific; Rel, member of NF-kB family; STAT, signal transducer and activator of transcription. Node size corresponds to percentage of peaks enriched for given TF. Edge thickness corresponds to % shared enriched peaks.

Promoters and enhancers rarely function in response to binding of single transcription factors. This is due both to the propensity of transcription factors within the same family to dimerize, and the existence of multiple transcription factor binding sites within a given promoter or enhancer. To identify potential interactions between regeneration-associated transcription factors, we quantified the frequency with which binding sites for differentially expressed transcription factors co-occurred within putative regulatory elements. For example, although KLF6 and WT1 were the most prevalent binding sites within both proximal and distal peaklets, the co-occurrence rate with binding sites with other transcription factors present in this time window was among the lowest (Fig. 4B). Within distal peaklets, KLF6 and WT1 binding sites frequently co-occurred with those of 2-4 other factors (Fig. 4B, *distal peaks*). By comparison, binding motifs for FOSL1 and JUNB displayed a high frequency of co-occurrence with binding sites for 6-7 other transcription factors (Fig. 4B, *distal peaks*). A similar trend was observed among proximal peaklets, although the differences were less marked (Fig. 4B, *proximal peaks*). These differences suggest the possibility that binding site composition and potential transcription factor interactions may be distinguishing characteristics of regeneration-associated promoters and enhancers.

We further quantified the frequency of specific co-occurring binding sites. The co-occurrence of members of the same family was expected due to similarities in recognition sequence specificity. This was observed most clearly in the frequency of shared enriched peaklets between KLF6 and WT1 binding sites, and between the bZIP factors (Fig. 4C). However, we also observed frequent co-occurrence of binding sites between members of different transcription families, most notably ASCL1 and NFATC2. These results suggest numerous complex regulatory mechanisms that fine-tune the expression of regeneration-associated genes including: (a) multiple within-family interactions that influence binding site specificity and affinity, as well as transcriptional activity; and (b) a variety of higher order multi-family complexes that may physically and/or functionally interact.

### Regeneration-associated regulatory sequences target *jun* expression

Although chromatin accessibility remains mostly stable in RGCs in response to optic nerve injury, we detected a small number of DNA elements in which chromatin accessibility changed in regenerating neurons compared to controls. Intriguingly, the few differentially accessible chromatin peaklets were not evenly distributed across the time points. Instead, most of the elements that became more accessible in response to injury did so at the earliest time point (2 dpi). This led us to postulate a role for these elements in triggering the regenerative growth program. We hypothesized that transcription factors regulated by such elements would not only regulate biological processes necessary for initiation of axon growth, but would also contribute to the regulation of downstream transcription factors that in turn regulate subsequent phases of regeneration.

In order to identify potential transcription factor targets of the regeneration-associated regulatory elements, we ranked the differentially expressed transcription factor genes (Fig. 2C) on the basis of their proximity to the differentially accessible chromatin regions. Surprisingly, *jun* was the only regeneration-associated transcription factor encoding gene that was located within at least 100 kb of a differentially accessible chromatin region. In fact, the *jun* gene is flanked by three peaklets that are differentially open at 2 dpi with respect to controls (Fig. 5A). One peaklet is centered at 148 bp upstream of the transcription start site within the putative promoter, and the two remaining peaklets are located distally, approximately 3.6 kb downstream of the transcription start site. Motif enrichment analysis of these sequences identified a motif for the JUN family of transcription factors, suggesting the potential for autoregulation. We also scanned these sequences ^33^ for motifs of other regeneration-associated transcription factors whose expression peaks early in regeneration (Fig. 2C, Fig. 3). In addition to JUN binding sites, we found high-scoring matches for KLF6, WT1, SP8, SPIB, FOSL1, JUNB, ATF3 and STAT3 motifs within these sequences. Although JUN has long been associated with axon regeneration in both CNS and PNS ^34–40^, mechanisms regulating its transcription in response to axonal injury are not well understood. Our results indicate that differential activation of JUN early in regeneration is potentially a consequence of increased accessibility of promoter and enhancer sequences to injury-induced transcription factors.

**Fig. 5.**
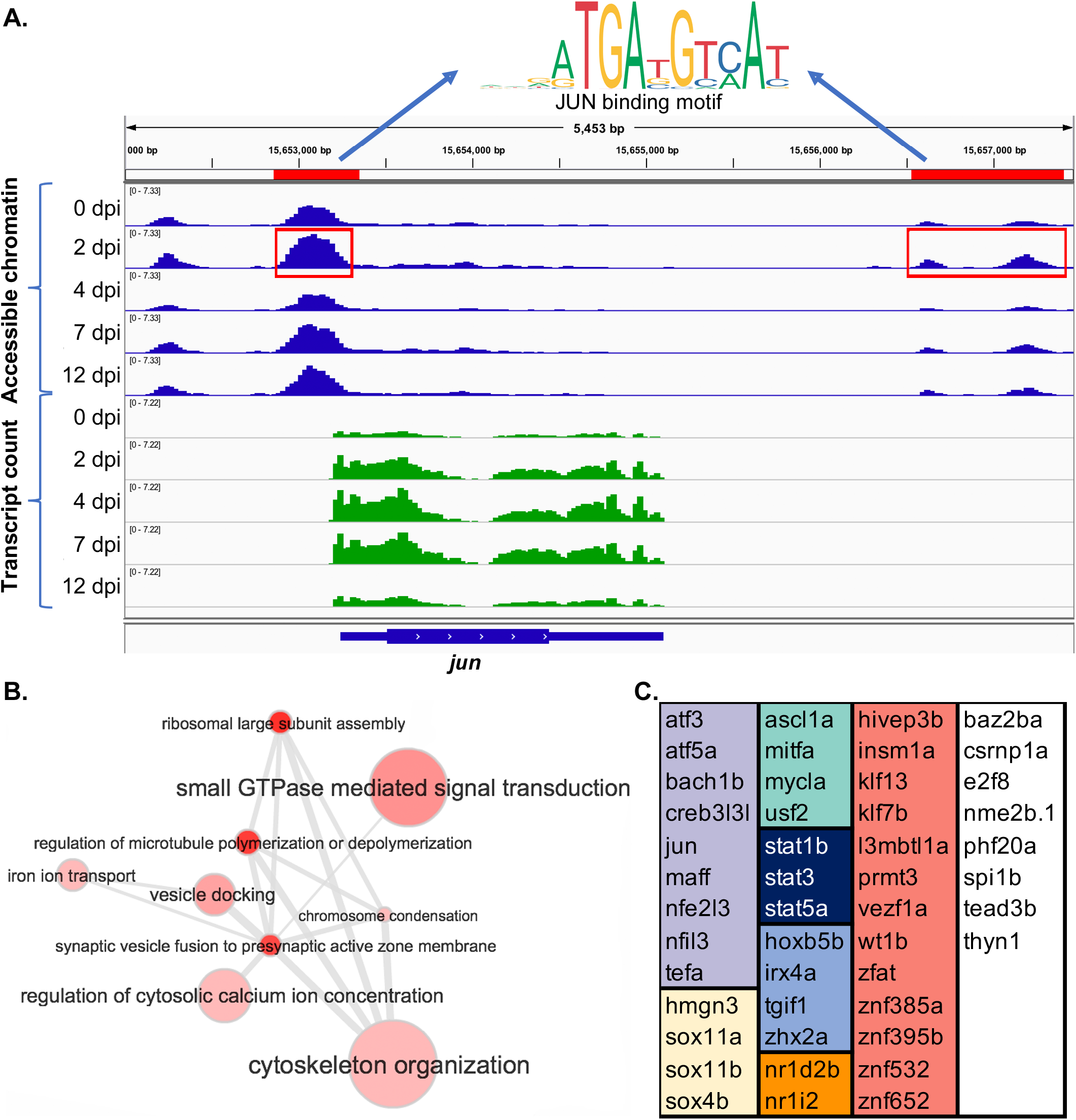
*Jun* is a potential regulatory target of regeneration-associated promoters and enhancers. (A) IGV browser screenshot displaying accessible chromatin sequence pileups (*blue*) and expressed sequence pileups (*green*) surrounding *jun* gene at 0, 2, 4, 7, and 12 dpi. Red bars and boxes indicate sequence determined to be differentially open at 2 dpi compared to control (0 dpi). Motif finding of sequences highlighted in red identified motifs corresponding to the JUN binding site. (B) Gene ontology (GO) analysis of inferred targets of JUN suggests roles in calcium regulation, microtubule dynamics, and translation. GO terms corresponding to JUN target genes were summarized, clustered and visualized using REVIGO ^36^. Node size corresponds to GO term frequency. Similar GO terms are linked by edges whose thickness corresponds to degree of similarity. (C) Inferred targets of JUN include 47 regeneration-associated TF genes. TF genes are grouped by transcription factor family: *violet*, leucine zipper; *yellow*, HMG/sox; *green*, helix-loop-helix; *dark blue*, STAT; *light blue*, hox; *orange*, nuclear receptor; *red*, C2H2 zinc finger; *white*, includes members of the AT hook, E2F, Ets, MBD, and TEA families as well as factors with uncategorized DNA binding domains.

We next analyzed the putative target genes of JUN for specific functional roles in the regenerative process. Putative JUN targets were identified based on our previous motif analysis (Fig. 3). Gene ontology analysis revealed a number of enriched biological processes consistent with a role for JUN in initiating axon regeneration (Table S8). The most significantly enriched terms include those involved in the regulation of microtubule dynamics and organization of the cytoskeleton, as well as those associated with small GTPase mediated signal transduction (Rho and Rab), and calcium regulation (Fig. 5B). The biological processes associated with potential JUN targets are a distinct subset of those associated with the larger list of all regeneration-associated genes in the same temporal clusters (Table S9). In the list of inferred JUN transcriptional targets, we also identified 47 regeneration-associated transcription factor encoding genes (Fig. 5C). This list includes 60% of the regeneration-associated transcription factors found in the first two temporal clusters (Fig. 1C). Thus, JUN has the potential for promoting and sustaining the regenerative program through activation of multiple downstream regeneration-associated transcription factors. Together the putative JUN targets suggest a central role for JUN in initiating and supporting successful CNS axon growth.

## Discussion

We have conducted the first combined temporal analysis of chromatin accessibility and transcriptomic changes that accompanies successful optic nerve regeneration. By identifying accessible regulatory elements, coupled with stage-specific transcription factor availability and downstream targets, these results provide a roadmap to the gene regulatory networks governing successful optic nerve regeneration. A major conclusion of this study is that temporally distinct functional modules are regulated by a dynamic cast of regeneration-associated transcription factors binding to regulatory elements that are accessible in naïve and regenerating RGCs. Many of the transcription factors have previously established roles in axon regeneration, while the roles of many more remain to be functionally validated. More than half of these regeneration-associated transcription factors fall into four transcription factor families, each with 25 or more members whose peak expression varies in a stage-specific manner. This is the first study that establishes a temporal hierarchy for regeneration-associated transcription factors based on expression patterns of transcription factors, target genes, and binding site accessibility. Future studies that use mass spectrometry approaches to identify stage-specific binding complexes could determine how the relative stoichiometry of individual factors at different stages of regeneration impacts complex formation and transcriptional activity.

Interestingly, the number of regeneration-associated changes in regulatory element accessibility are more than an order of magnitude less frequent in number than regeneration-associated changes in gene expression. Furthermore, the differentially accessible elements are almost exclusively found at 2 dpi and 12 dpi, our earliest and latest timepoints, respectively. Based on the timing and limited number of differentially accessible elements, we postulated a role for these elements in triggering transcriptional programs for axon regrowth, and synaptogenesis. We hypothesize at least two mechanisms by which this could occur: (i) The elements may be responsible for initiating the expression of key regeneration-associated transcription factors, which would subsequently regulate other factors in the hierarchy; and (ii) The elements may serve to shift the higher order chromatin structure to reposition enhancers and promoters for the transitions necessary in adult RGCs to reinitiate programs for axonogenesis (2 dpi), and and synaptogenesis (12 dpi).

Our data contain evidence supporting both hypotheses. Supporting the first hypothesis, we find that the gene encoding transcription factor JUN is flanked by promoter and distal enhancer elements that increase in accessibility during axon regeneration. In fact, JUN involvement in such epigenetic activation of pro-regenerative genes in response to axon injury was suggested in a recent review ^41^. Supporting our second hypothesis, we find that roughly half the differentially-accessible regions at both early and late times are enriched in CTCF binding sites. CTCF is a transcription factor that has recently been implicated in mediating chromatin looping and marking the boundaries of topologically associating domains ^42^. A logical next step would be to functionally validate interactions between the predicted *jun* promoter and enhancers, as well as additional long-range regulatory interactions between stably and differentially accessible elements using chromatin capture and genome editing technologies.

The combination of gene expression and genomic accessibility over time provides a powerful model of axon regeneration-associated gene regulatory networks. As with any modeling approach, empirical testing and improved technology are expected to refine and strengthen the predictive power of the model. For example, current methods for associating enhancers with target genes will improve with the functional testing of these types of interactions as described above. Another potential caveat of these studies is that the transcriptomic analysis were carried out on whole retinas, rather than FACS-sorted RGCs that were utilized in the chromatin accessibility. Because RGCs are the only cells directly injured by optic nerve crush, a comparison of gene expression between injured and uninjured retina should primarily reflect changes in RGCs. Yet, it is possible that some of the transcriptomic changes detected may be occurring in other cells within the injured retina, such as infiltrating microglia or retinal neurons connected to RGCs. However, we find good concordance between our transcriptomic results and microarray studies based on RNA extracted from RGCs isolated by laser capture microdissection ^17^. In addition, our temporal approach using RNA-seq has greatly expanded detection of regeneration-associated changes in gene expression over previous studies. We anticipate that functional testing of the networks predicted by this model will substantially expand our understanding of the molecular mechanisms governing successful optic nerve regeneration.

In summary, these data provide a roadmap for the identification of key combinations of transcription factors necessary to reprogram adult RGCs for optic nerve regeneration. Similar approaches have been employed to discover transcription factors necessary for direct reprogramming of somatic cells to produce motor neurons ^43^. However, to our knowledge, this is the first gene regulatory network analysis of optic nerve regeneration that couples temporal analysis of gene expression with the identification of putative regulatory interactions based on chromatin accessibility and transcription factor expression. We expect that these findings may be applicable to neurons in other regions of the CNS undergoing regenerative axon growth, such as the spinal cord and brain. In order to facilitate comparisons between our data and those derived from other regenerative models, we have created an interactive web application (*Regeneration Rosetta;* http://ls-shiny-prod.uwm.edu/rosetta/) ^44^. This app enables users to upload a gene list of interest from any Ensembl-supported species and determine how those genes are expressed and potentially regulated in the context of optic nerve regeneration. Such cross-species and cross-system analyses will facilitate identification of novel pathways specifically associated with successful CNS regeneration.

## Materials and Methods

Detailed methods can be found in supplementary information.

### Zebrafish husbandry and optic nerve injury

Zebrafish husbandry and all experimental procedures were approved by the *Institutional Animal Care and Use Committee* (IACUC) and carried out in accordance with the relevant guidelines and regulations thereof. The Tg (Tru.gap43:egfp) mil1 (aka fgap43:egfp) transgenic line and Ekkwill wild type strain were maintained as previously described ^25^. Optic nerve crush (ONC) lesions were performed on adult zebrafish, 7-9 months of age, as previously described ^21^.

### RNA-seq data generation

RNA was extracted and purified from retinas dissected from naïve (0 dpi) and regenerating adult fish at 2, 4, 7, and 12 dpi. RNA was extracted using the RNeasy Micro kit (Qiagen) and concentrated using the RNA Clean & Concentrator kit (Zymo). Three replicates were obtained for each time point. Whole retinas were used for transcriptomic studies because the quality of the RNA was much higher from whole retina. cDNA libraries were generated for each RNA sample using Tru-Seq Stranded Total & mRNA Sample Prep Kits, (Illumina 20020595). Each cDNA library was indexed for multiplexing and subsequently sequenced on four lanes of the Illumina Hiseq2000. Libraries were sequenced at 50 bp, 30–40 million paired-end reads/sample.

After merging technical replicates of RNA-seq samples across lanes, adapter sequences were trimmed (TrimGalore, v0.4.4), sequence quality was validated (FastQC, v0.11.5), and transcript abundance was estimated using 500 bootstrap samples (Kallisto (v0.42.4) ^45^.

### ATAC-seq data generation

Chromatin was extracted from nuclei isolated from purified populations of RGCs. Retinas were dissected from naïve and regenerating fgap43:egfp fish as described above, and enzymatically dissociated for FACS. Cell suspensions were pooled from multiple retinas at each time point (0 dpi, 8-10 retina; 2, 4, 7, and 12 dpi, 4-6 retinas), and sorted for RGCs expressing GFP using a Becton Dickinson FACSAria™ III sorter. Although levels of endogenous Gap43 are low in the naïve adult retina, the GFP transgene tends to be long lived and is detectable by FACS at levels that are above background but below the level of visual detection. This enabled us to sort RGCs from naïve retinas since new RGCs that continue to be added throughout the life of the animal and low-levels of GFP persist in RGCs even after these new RGCs have established connections in the tectum. However, we required twice the number of naïve retinas compared with injured retinas to accumulate the 50,000 RGCs required for constructing the ATAC-seq libraries. Three replicates of pooled cells were collected for each time point. ATAC-seq libraries were prepared using the Tn5 transposase system (Nextera DNA library kit, Illumina, FC-121–1030) as previously described ^46^, and purified using DNA Clean & Concentrator kit (SKU: ZD5205, Zymo).

Prior to running the full sequence, sequencing depth was estimated and one sample was below the cut-off criteria and therefore omitted. The remaining fourteen samples were indexed for multiplexing and subsequently sequenced on four lanes of the Illumina Hiseq2000Data at 50 bp reads to obtain approximately 25 million paired-end reads/samples.

After merging technical replicates of ATAC-seq samples across lanes, adapter sequences were trimmed (TrimGalore, v0.4.4) and sequence quality was validated (FastQC, v0.11.5). Reads were aligned to GRCz10 as well as the transgene sequence (BWA-MEM (v0.7.9a-r786) ^47^). Duplicate and multiple mapped reads were removed using samtools (v1.6) ^48^. After concatenating aligned reads across all replicates and time points, MACS2 ^49^ (v2.1.1.20160309) was used to call peaks from aligned reads. Only peaks with a p-value < 10-10 were retained for subsequent analyses. For each remaining subpeak summit, a 500bp “peaklet” interval was defined using GenomicRanges (v1.30.3) ^50^. We refer to these peaklets as consensus regions of accessible chromatin. Open chromatin in each replicate of each time point was then quantified using DiffBind (v2.6.6, default parameters) by counting the number of overlapping reads for each retained peaklet.

We used motif analysis to determine potential binding sites of differentially expressed transcription factors within regions of accessible chromatin identified by ATAC-seq. Motif enrichment and discovery was carried out within the MEME Suite of motif-based sequence analysis tools (version 5.0.3) ^51^.

### Statistical analysis of RNA-seq and ATAC-seq data

Following pseudoalignment and quantification of transcripts, differentially expressed transcripts were identified using Sleuth (v0.29.0) ^52^, either comparing each time point post injury (2, 4, 7, 12 dpi) to the initial time point (0dpi) using a Wald test statistic or using a likelihood ratio test (LRT) to compare the full model with a time factor versus the null model, controlling the false discovery rate (FDR) at 5% ^53^. Expression heatmaps (based on Z-scores calculated using log transcripts per million [TPM] estimates) were produced using ComplexHeatmap ^54^ (v1.17.1), where transcript clusters were identified using the K-means algorithm, and hierarchical clustering (Euclidean distance, complete linkage) was used to cluster rows. The Ingenuity Pathway Analysis tool (Qiagen, Redwood City, CA, USA) was used to analyze enrichment of molecular and functional gene networks within the differentially expressed gene sets (FDR<0.05) at each time point after injury (2, 4, 7, 12 dpi) compared with the initial time point (0dpi).

After quantifying peaklet accessibility, DESeq2 (v1.18.1) ^55^ was used to identify differentially accessible peaklets in an analogous manner to the RNA-seq analysis described above. Peaklets with FDR-controlled p-values < 0.05 in one of the four comparisons were considered to be differentially accessible. ChIPpeakAnno (v3.12.7) ^56^, the danRer10.refGene UCSC annotation package (v3.4.2), and AnnotationHub (v2.10.1) were used to annotate peaklets with genes. Specifically, non-exonic (i.e., not overlapping exons by more than 50bp) peaklets overlapping a transcription start site (TSS) or within 1kb of a TSS were considered to represent proximal peaks, whereas those greater than 1kb but less than 100kb of a TSS were considered distal peaks. All statistical analyses were performed in R (v3.4.3). Integrative Genome Viewer (IGV) was used to visualize RNA-seq and ATAC-seq alignments ^57^.

## Acknowledgements

We thank the <u>UWM High Performance Computing (HPC) Service</u> for computing resources used in this work. We are grateful to Heather Leskinen and Sarah Sarich for comments on the manuscript and edits to the material and methods. We wish to thank the University of Wisconsin Biotechnology Center DNA Sequencing Facility, Madison for providing RNA and ATAC sample sequencing facilities. We gratefully acknowledge support from the Research Growth Initiative Grant RGI (to A.J.U. and P.L.A.), University of Wisconsin-Milwaukee. A.R. was supported by the AgreenSkills+ fellowship program, which received funding from the EU’s Seventh Framework Program under grant agreement number FP7-609398 (AgreenSkills+ contract). S.D. was supported by the Clifford H. Mortimer Award, University of Wisconsin-Milwaukee

## Author Contributions

AJU was responsible for the conception and design of the work. SPD, NMR, MDL and AJU contributed to tissue processing and sample collection for nucleic acid libraries. AR and PLA were responsible for the bioinformatics and statistical analyses of the RNA-seq and ATAC-seq data. MJF conducted the pathway analysis. SPD, AR, PLA and AJU contributed to the interpretation of the data, preparation of the figures, and writing of the manuscript. All authors reviewed the manuscript.

## Data availability

Raw sequencing files for the RNA-seq and ATAC-seq data will be submitted upon publication to the Sequence Read Archive (SRA). All scripts used to process and analyze the RNA-seq and ATAC-seq data may be found at https://github.com/andreamrau/OpticRegen_2019. The *Regeneration Rosetta* app may be accessed at http://ls-shiny-prod.uwm.edu/rosetta/, and all associated source files for creating the app may be found at https://github.com/andreamrau/rosetta.

## References

1 Belin, S. et al. Injury-induced decline of intrinsic regenerative ability revealed by quantitative proteomics. Neuron 86, 1000–1014, doi:10.1016/j.neuron.2015.03.060 (2015).

2 Duan, X. et al. Subtype-specific regeneration of retinal ganglion cells following axotomy: effects of osteopontin and mTOR signaling. Neuron 85, 1244–1256, doi:10.1016/j.neuron.2015.02.017 (2015).

3 Leibinger, M. et al. Boosting central nervous system axon regeneration by circumventing limitations of natural cytokine signaling. Mol Ther 24, 1712–1725, doi:10.1038/mt.2016.102 (2016).

4 Park, K. K. et al. Promoting Axon Regeneration in the Adult CNS by Modulation of the PTEN/mTOR Pathway. Science 322, 963–966, doi:10.1126/science.1161566 (2008).

5 Pernet, V. et al. Long-distance axonal regeneration induced by CNTF gene transfer is impaired by axonal misguidance in the injured adult optic nerve. Neurobiol Dis 51, 202–213, doi:10.1016/j.nbd.2012.11,011 (2013).

6 Smith, P. D. et al. SOCS3 deletion promotes optic nerve regeneration in vivo. Neuron 64, 617–623, doi:10.1016/j.neuron.2009.11.021 (2009).

7 Sun, F. et al. Sustained axon regeneration induced by co-deletion of PTEN and SOCS3. Nature 480, 372–U125, doi:10.1038/nature10594 (2011).

8 Trakhtenberg, E. F. et al. Zinc chelation and Klf9 knockdown cooperatively promote axon regeneration after optic nerve injury. Exp Neurol 300, 22–29, doi:10.1016/j.expneurol.2017.10.025 (2018).

9 Luo, X. T. et al. Three-dimensional evaluation of retinal ganglion cell axon regeneration and pathfinding in whole mouse tissue after injury. Exp Neurol 247, 653–662, doi:10.1016/j.expneurol.2013.03.001 (2013).

10 Diekmann, H., Kalbhen, P. & Fischer, D. Characterization of optic nerve regeneration using transgenic zebrafish. Front Cell Neurosci 9, doi:ARTN 118, 10.3389/fncel.2015.00118 (2015).

11 Lemmens, K. et al. Matrix metalloproteinases as promising regulators of axonal regrowth in the injured adult zebrafish retinotectal system. Journal of Comparative Neurology 524, 1472–1493, doi:10.1002/cne.23920 (2016).

12 Kaneda, M. et al. Changes of phospho-growth-associated protein 43 (phospho-GAP43) in the zebrafish retina after optic nerve injury: a long-term observation. Neurosci Res 61, 281–288, doi:10.1016/j.neures.2008.03.008 (2008).

13 Erskine, L. & Herrera, E. Connecting the retina to the brain. ASN Neuro 6, doi:10.1177/1759091414562107 (2014).

14 Skene, J. H. Axonal growth-associated proteins. Annu Rev Neurosci 12, 127–156, doi:10.1146/annurev.ne.12.030189.001015 (1989).

15 Elsaeidi, F., Bemben, M. A., Zhao, X. F. & Goldman, D. Jak/Stat signaling stimulates zebrafish optic nerve regeneration and overcomes the inhibitory actions of Socs3 and Sfpq. J Neurosci 34, 2632–2644, doi:10.1523/JNEUROSCI.3898-13.2014 (2014).

16 Bernhardt, R. R., Tongiorgi, E., Anzini, P. & Schachner, M. Increased expression of specific recognition molecules by retinal ganglion cells and by optic pathway glia accompanies the successful regeneration of retinal axons in adult zebrafish. J Comp Neurol 376, 253–264, doi:10.1002/(SICI)1096-9861(19961209)376:2<253::AID-CNE7>3.0.CO;2-2 (1996).

17 Veldman, M. B., Bemben, M. A., Thompson, R. C. & Goldman, D. Gene expression analysis of zebrafish retinal ganglion cells during optic nerve regeneration identifies KLF6a and KLF7a as important regulators of axon regeneration. Dev Biol 312, 596–612, doi:10.1016/j.ydbio.2007.09.019 (2007).

18 Kusik, B. W., Hammond, D. R. & Udvadia, A. J. Transcriptional regulatory regions of gap43 needed in developing and regenerating retinal ganglion cells. Dev Dyn 239, 482–495, doi:10.1002/dvdy.22190 (2010).

19 Senut, M. C., Gulati-Leekha, A. & Goldman, D. An element in the alpha1-tubulin promoter is necessary for retinal expression during optic nerve regeneration but not after eye injury in the adult zebrafish. J Neurosci 24, 7663–7673, doi:10.1523/JNEUROSCI.2281-04.2004 (2004).

20 Udvadia, A. J., Koster, R. W. & Skene, J. H. GAP-43 promoter elements in transgenic zebrafish reveal a difference in signals for axon growth during CNS development and regeneration. Development 128, 1175–1182 (2001).

21 Bormann, P., Zumsteg, V. M., Roth, L. W. A. & Reinhard, E. Target contact regulates GAP-43 and alpha-tubulin mRNA levels in regenerating retinal ganglion cells. J Neurosci Res 52, 405–419, doi:10.1002/(Sici)1097-4547(19980515)52:4<405::Aid-Jnr4>3.0.Co;2-D (1998).

22 Beckers, A. et al. An Antagonistic Axon-Dendrite Interplay Enables Efficient Neuronal Repair in the Adult Zebrafish Central Nervous System. Mol Neurobiol 56, 3175–3192, doi:10.1007/s12035-018-1292-5 (2019).

23 Diekmann, H., Kalbhen, P. & Fischer, D. Active mechanistic target of rapamycin plays an ancillary rather than essential role in zebrafish CNS axon regeneration. Front Cell Neurosci 9, doi:ARTN 251, 10.3389/fncel.2015.00251 (2015).

24 Buenrostro, J. D., Giresi, P. G., Zaba, L. C., Chang, H. Y. & Greenleaf, W. J. Transposition of native chromatin for fast and sensitive epigenomic profiling of open chromatin, DNA-binding proteins and nucleosome position. Nat Methods 10, 1213–1218, doi:10.1038/nmeth.2688 (2013).

25 Udvadia, A. J. 3.6 kb Genomic sequence from Takifugu capable of promoting axon growth-associated gene expression in developing and regenerating zebrafish neurons. Gene Expr Patterns 8, 382–388, doi:10.1016/j.gep.2008.05.002 (2008).

26 Daugherty, A. C. et al. Chromatin accessibility dynamics reveal novel functional enhancers in C. elegans. Genome Res 27, 2096–2107, doi:10.1101/gr.226233.117 (2017).

27 Lambert, S. A. et al. The Human Transcription Factors. Cell 172, 650–665, doi:10.1016/j.cell.2018.01.029 (2018).

28 Khan, A. et al. JASPAR 2018: update of the open-access database of transcription factor binding profiles and its web framework. Nucleic Acids Res 46, D260–D266, doi:10.1093/nar/gkx1126 (2018).

29 Weirauch, M. T. et al. Determination and Inference of Eukaryotic Transcription Factor Sequence Specificity. Cell 158, 1431–1443, doi:10.1016/j.cell.2014.08.009 (2014).

30 McLeay, R. C. & Bailey, T. L. Motif Enrichment Analysis: a unified framework and an evaluation on ChIP data. BMC Bioinformatics 11, 165, doi:10.1186/1471-2105-11-165 (2010).

31 Miao, Q. et al. Tcf3 promotes cell migration and wound repair through regulation of lipocalin 2. Nat Commun 5, 4088, doi:10.1038/ncomms5088 (2014).

32 Wagner, K. D. et al. The Wilms’ tumor gene Wt1 is required for normal development of the retina. EMBO J 21, 1398–1405, doi:10.1093/emboj/21.6.1398 (2002).

33 Grant, C. E., Bailey, T. L. & Noble, W. S. FIMO: scanning for occurrences of a given motif. Bioinformatics 27, 1017–1018, doi:10.1093/bioinformatics/btr064 (2011).

34 Chandran, V. et al. A Systems-Level Analysis of the Peripheral Nerve Intrinsic Axonal Growth Program. Neuron 89, 956–970, doi:10.1016/j.neuron.2016.01.034 (2016).

35 Herdegen, T. et al. Expression of JUN, KROX, and CREB transcription factors in goldfish and rat retinal ganglion cells following optic nerve lesion is related to axonal sprouting. J Neurobiol 24, 528–543, doi:10.1002/neu.480240410 (1993).

36 Herdegen, T., Tolle, T. R., Bravo, R., Zieglgansberger, W. & Zimmermann, M. Sequential expression of JUN B, JUN D and FOS B proteins in rat spinal neurons: cascade of transcriptional operations during nociception. Neurosci Lett 129, 221–224 (1991).

37 Herman, P. E. et al. Highly conserved molecular pathways, including Wnt signaling, promote functional recovery from spinal cord injury in lampreys. Sci Rep 8, 742, doi:10.1038/s41598-017-18757-1 (2018).

38 Kenney, A. M. & Kocsis, J. D. Peripheral axotomy induces long-term c-Jun amino-terminal kinase-1 activation and activator protein-1 binding activity by c-Jun and junD in adult rat dorsal root ganglia In vivo. J Neurosci 18, 1318–1328 (1998).

39 Leah, J. D., Herdegen, T. & Bravo, R. Selective expression of Jun proteins following axotomy and axonal transport block in peripheral nerves in the rat: evidence for a role in the regeneration process. Brain Res 566, 198–207, doi:10.1016/0006-8993(91)91699-2 (1991).

40 Ruff, C. A. et al. Neuronal c-Jun is required for successful axonal regeneration, but the effects of phosphorylation of its N-terminus are moderate. Journal of neurochemistry 121, 607–618, doi:10.1111/j.1471-4159.2012.07706.x (2012).

41 Mahar, M. & Cavalli, V. Intrinsic mechanisms of neuronal axon regeneration. Nat Rev Neurosci 19, 323–337, doi:10.1038/s41583-018-0001-8 (2018).

42 Dixon, J. R. et al. Topological domains in mammalian genomes identified by analysis of chromatin interactions. Nature 485, 376–380, doi:10.1038/nature11082 (2012).

43 Velasco, S. et al. A Multi-step Transcriptional and Chromatin State Cascade Underlies Motor Neuron Programming from Embryonic Stem Cells. Cell Stem Cell 20, 205–217 e208, doi:10.1016/j.stem.2016.11.006 (2017).

44 Rau, A., Dhara, S. P., Udvadia, A. J. & Auer, P. L. Regeneration Rosetta: An interactive we application to explore regeneration-associated gene expression and chromatin accessibility. bioRxiv, doi:https://doi.org/10.1101/632018 (2019).

45 Bray, N. L., Pimentel, H., Melsted, P. & Pachter, L. Near-optimal probabilistic RNA-seq quantification. Nature Biotechnology 34, 525–527, doi:10.1038/nbt.3519 (2016).

46 Buenrostro, J. D., Wu, B., Chang, H. Y. & Greenleaf, W. J. ATAC-seq: A Method for Assaying Chromatin Accessibility Genome-Wide. Curr Protoc Mol Biol 109, 21 29 21–29, doi:10.1002/0471142727.mb2129s109 (2015).

47 Li, H. & Durbin, R. Fast and accurate short read alignment with Burrows-Wheeler transform. Bioinformatics 25, 1754–1760, doi:10.1093/bioinformatics/btp324 (2009).

48 Li, H. et al. The Sequence Alignment/Map format and SAMtools. Bioinformatics 25, 2078–2079, doi:10.1093/bioinformatics/btp352 (2009).

49 Zhang, Y. et al. Model-based Analysis of ChIP-Seq (MACS). Genome Biology 9, doi:ARTN R137; 10.1186/gb-2008-9-9-r137 (2008).

50 Lawrence, M. et al. Software for Computing and Annotating Genomic Ranges. Plos Comput Biol 9, doi:ARTN e1003118; 10.1371/journal.pcbi.1003118 (2013).

51 Bailey, T. L., Johnson, J., Grant, C. E. & Noble, W. S. The MEME Suite. Nucleic Acids Res 43, W39–49, doi:10.1093/nar/gkv416 (2015).

52 Pimentel, H., Bray, N. L., Puente, S., Melsted, P. & Pachter, L. Differential analysis of RNA-seq incorporating quantification uncertainty. Nat Methods 14, 687-+, doi:10.1038/nmeth.4324 (2017).

53 Benjamini, Y. & Hochberg, Y. Controlling the False Discovery Rate - a Practical and Powerful Approach to Multiple Testing. J R Stat Soc B 57, 289–300 (1995).

54 Gu, Z., Eils, R. & Schlesner, M. Complex heatmaps reveal patterns and correlations in multidimensional genomic data. Bioinformatics 32, 2847–2849, doi:10.1093/bioinformatics/btw313 (2016).

55 Love, M. I., Huber, W. & Anders, S. Moderated estimation of fold change and dispersion for RNA-seq data with DESeq2. Genome Biol 15, 550, doi:10.1186/s13059-014-0550-8 (2014).

56 Zhu, L. J. et al. ChIPpeakAnno: a Bioconductor package to annotate ChIP-seq and ChIP-chip data. BMC Bioinformatics 11, 237, doi:10.1186/1471-2105-11-237 (2010).

57 Robinson, J. T. et al. Integrative genomics viewer. Nat Biotechnol 29, 24–26, doi:10.1038/nbt.1754 (2011).

58 Supek, F., Bosnjak, M., Skunca, N. & Smuc, T. REVIGO summarizes and visualizes long lists of gene ontology terms. PLoS One 6, e21800, doi:10.1371/journal.pone.0021800 (2011).

