## Supplementary Information for "Cellular reprogramming for successful CNS axon regeneration is driven by a temporally changing cast of transcription factors"

1. Detailed Materials and Methods, SI references
2. Figure S1. Accessible chromatin is mainly located distal to annotated genes.
3. Figure S2. Differentially expressed transcripts that encode transcription factors display regeneration stage-specific temporal patterning
4. Figure S3. Temporal patterning of differentially expressed leucine zipper (bZIP) family transcription factors.
5. Figure S4. Temporal patterning of differentially expressed basic helix-loop-helix (bHLH) family transcription factors.
6. Figure S5. Temporal patterning of differentially expressed 2 cys-2 his zinc finger (C2H2ZF) family transcription factors.
7. Figure S6. Temporal patterning of differentially expressed homeodomain family transcription factors.
8. Figure S7. Injury-induced eGFP expression in retinal ganglion cells can be visualized through the lens of intact animals.
9. Figure S8. FASTA sequence of transgene.
10. Table S1. Transcripts differentially expressed compared to controls (0 dpi) in at least one time point (5% FDR).
11. Table S2. Differentially expressed transcripts (1% FDR) clustered based on temporal expression pattern.
12. Table S3. Ingenuity Pathway Analysis of temporally clustered, differentially expressed genes.
13. Table S4. Differentially accessible sequences with distance to nearest annotated genes and nearest differentially-expressed genes.
14. Table S5. Differentially expressed transcripts that encode transcription factors with and without known motifs.
15. Table S6. Differentially expressed transcripts that encode transcription factors with known motifs.
16. Table S7. Transcription factors that are differentially expressed during regeneration.
17. Table S8. Gene ontology (GO) analysis of putative Jun transcriptional targets.
18. Table S9. Gene ontology analysis of regeneration-associated genes with peak expression during initial axon growth toward the midline.
19. Table S10. RNA sample quality control.
20. Table S11. ATAC-seq library sample quality control
21. Table S12. PCR primers for ATAC-seq libraries based on Nextera indices

***Detailed Materials and Methods***

### Zebrafish husbandry and maintenance

Zebrafish husbandry and all experimental procedures were approved by the *Institutional Animal Care and Use Committee* (IACUC). Zebrafish colonies were maintained as previously described [1]. Adult fish were housed in recirculating rack systems (Aquatic Habitats, Apopka, FL) at 28.5°C on a 14-hour light, 10-hour dark cycle, and fed twice daily with Adult Zebrafish Complete Diet (VWR, West Chester, PA) and once daily with brine shrimp (*Artemia*). A wild type strain (Ekkwill, EK) was used as a negative control for FACS. A transgenic reporter strain constructed on the EK background, Tg (*Tru.gap43:egfp*) mil1, or *fgap43:egfp* [2] was used for all other experiments.

### Zebrafish optic nerve injury

Optic nerve crush (ONC) lesions were performed on adult zebrafish, 7-9 months of age, as previously described [3]. Briefly, fish were anesthetized in 0.46 mg/mL tricaine (Argent Chemical Labs, Redmond, WA) in 30% Danieau [4]. The left optic nerve of anesthetized fish was exposed and crushed for 10 seconds using Dumont #5 forceps. The intact retina of an uninjured fish served as the unoperated control (0dpi). Fish were sacrificed 0, 2, 4, 7, or 12 days post injury (dpi) and retinas were dissected. Prior to dissection, left and right eyes of the fish were examined under Nikon eclipse TE2000-U fluorescence microscope for GFP fluorescence as an indicator of regeneration-induced transcriptional activity. GFP was easily detected through the lens in eyes which had previously undergone optic nerve crush (Fig. S7).

### RNA-seq data generation

*RNA Isolation*. RNA was extracted and purified from retinas dissected from naïve (0 dpi) and regenerating adult fish at 2, 4, 7, and 12 dpi. Three biological replicates of RNA were obtained for each time point. To prevent RNA degradation, dissected retinas were immediately immersed in an RNA stabilization reagent (RLT buffer of RNeasy Micro Kit, Cat No./ID 74004, Qiagen; Valencia, CA). The retinas were homogenized (sterile Fisherbrand™ RNase-Free Disposable Pellet Pestles Cat No.12-141-368) and filtered (Cat No./ID: 79654, Qiagen). Total RNA was extracted from the homogenized mixture according to manufacturer instructions (RNeasy Micro Kit, Cat No./ID 74004, Qiagen). Total RNA concentration and purity were quantified with NanoDrop ND2000 spectrophotometer (Thermo Scientific) and QuBit fluorometer (Invitrogen), respectively. RNA integrity was quantified using 2100 Bioanalyzer (Agilent High Sensitivity RNA 6000 Pico Reagents Cat No. 5067-1514). For each biological replicate, 3-6 retinas were pooled, cleaned and concentrated (RNA Clean & Concentrator kit, SKU R1013, Zymo Research) to obtain 1 ug total RNA. Total RNA concentration, purity, and integrity of pooled samples was determined as described above. Only samples with RNA integrity numbers (RIN) ranging from 7.7 to 8.2 were used for sequencing (Table S10).

*Library preparation and sequencing*. cDNA libraries (n=3 for 0, 2, 4, 7, and 12 dpi) were generated using Tru-Seq Stranded Total & mRNA Sample Prep Kits, (Illumina 20020595) at University of Wisconsin-Madison Biotechnology Center (UWBC). Each cDNA library was indexed for multiplexing and subsequently sequenced on four lanes of the Illumina Hiseq2000 device, UWBC. Libraries were sequenced at 50 bp, 30–40 million paired-end reads/sample, on Illumina HiSeq 2500 at UWBC, Madison.

*Bioinformatic analysis*. After merging technical replicates of RNA-seq samples across lanes, TrimGalore (v0.4.4, --stringency 3 -q 20) was used in paired-end mode to trim adaptor sequences,

FastQC (v0.11.5) was used to validate sequence quality, and Kallisto [5] (v0.42.4) was used to build an index on a FASTA file consisting of the zebrafish transcriptome (GRCz10 Danio rerio genome assembly, Ensembl release 84 annotation) and the transgene sequence (Fig. S8). Kallisto was subsequently used to quantify transcript abundances using 500 bootstrap samples.

**ATAC-seq data generation**

*Cell sorting.* RGCs were collected from dissociated regenerating and control retinas, at each time point using fluorescent activated cell sorting (FACS). Zebrafish were sacrificed, and retinas were dissected and immersed in ice cold PBS with no calcium and magnesium. To one well of a 24-well plate, a single retina (divided into 8 uniform pieces) was added to 500 μL Accumax (STEMCELL Technologies). Each retina was chemically digested for 70 mins with agitation on a nutator. Reactions were quenched in heat inactivated fetal calf serum (HI-FCS; Gemini Bio-Products) in DMEM/F12 (Gibco) and lysates were mechanically dissociated by gentle pipetting. Undigested fragments were removed, and the cell suspension was pelleted at 200 g for 3 min. Supernatant was aspirated and the pellet re-suspended in 100 μl fresh quenching buffer (DMEM/F12 + 20% FCS). Cell suspensions from multiple retinas at each time point were pooled (0 dpi, 8-10 retina; 2, 4, 7, and 12 dpi, 4-6 retinas). Pooled cell suspension were filtered (Falcon, Cat No. 352235), and sorted for RGCs expressing GFP using a Becton Dickinson FACSAria™ III sorter fitted with the 100 μM nozzle. Negative control cells were used to set gates to separate GFP positive (GFP+) from GFP negative (GFP−) fractions. We collected 50,000 FACS-sorted GFP+ cells per sample that were immediately used for chromatin isolation as described below. Three biologically distinct replicates of pooled cells were collected for each time point.

*Library preparation.* ATAC-seq libraries were prepared using the Tn5 transposase system (Nextera DNA library kit, Illumina, FC-121–1030) as previously described [6], and purified using DNA Clean & Concentrator kit (SKU: ZD5205, Zymo).The purified samples were assessed for quality as described above (NanoDrop ND2000 spectrophotometer and QuBit fluorometer) and appropriate nucleosomal laddering was determined by 2100 Bioanalyzer (Agilent High Sensitivity DNA Kit, Cat No. 5067-4626) (Table S11). For PCR amplification and qPCR side reactions, PerfeCTa SYBR Green FastMix Quanta (VWR Cat No. 101414-150) was used (Table S12 for PCR Primers).

*Sequencing.* Prior to running the full sequence, Mi-seq was used to estimate sequencing depth. One sample was below the cut-off criteria and therefore omitted. The remaining fourteen samples were indexed for multiplexing and subsequently sequenced on four lanes of the Illumina Hiseq2000, UWBC. Data were sequenced at 50 bp to obtain approximately 25 million paired-end reads/samples.

*Bioinformatic analysis.* After merging technical replicates of ATAC-seq samples across lanes, TrimGalore (v0.4.4, --stringency 3 -q 20) was used in paired-end mode to trim adaptor sequences and FastQC (v0.11.5) was used to validate sequence quality. BWA-MEM [7] (v0.7.9a-r786) was used to align reads to the zebrafish genome (GRCz10) and transgene sequence. Duplicate and multiple mapped reads were removed using samtools (v1.6) [8]. After concatenating aligned reads across all replicates and time points, MACS2 [9] (v2.1.1.20160309, --no-model -g 1.37e+09 --keep-dup all --call-summits) was used in paired-end mode without shifting model to call peaks from aligned reads, and summits of deconvoluted subpeaks were identified. Only peaks with a p-value $< 10^{-10}$ were retained for subsequent analyses. For each remaining subpeak summit, a 500bp "peaklet" interval was defined using [summit - 250bp, summit + 249bp] using GenomicRanges (v1.30.3) [10]. We refer to these peaklets as consensus regions of accessible chromatin. Open chromatin in each replicate of each time point was then quantified using DiffBind (v2.6.6, default parameters) by counting the number of overlapping reads for each retained peaklet.

**Statistical analysis of RNA-seq and ATAC-seq data**

Following pseudoalignment and quantification of transcripts, differentially expressed transcripts were identified using Sleuth (v0.29.0) [11]. Specifically, a full model, including a factor for each time point after injury (2, 4, 7, 12 dpi), was estimated for each transcript, and a Wald test was calculated for each coefficient to identify significant differences with the initial time point (0dpi). After controlling the false discovery rate (FDR) at 5% within each comparison using the Benjamini-Hochberg [12] approach, differentially expressed transcripts with respect to the baseline were identified for each post-injury time point. Beta values from the model were used as a biased estimator of log-fold change. Expression heatmaps (based on Z-scores calculated using either log fold-changes or log transcripts per million [TPM] estimates) were produced using ComplexHeatmap [13] (v1.17.1), where transcript clusters were identified using the K-means algorithm, and hierarchical clustering (Euclidean distance, complete linkage) was used to cluster rows. The Ingenuity Pathway Analysis tool (Qiagen, Redwood City, CA, USA) was used to analyze enrichment of molecular and functional gene networks within the differentially expressed gene sets (FDR<0.05) at each time point after injury (2, 4, 7, 12 dpi) compared with the initial time point (0dpi).

After quantifying peaklet accessibility, DESeq2 (v1.18.1) [14] was used to identify differentially accessible peaklets in an analogous manner to the RNA-seq analysis described above. As before, a full generalized linear model including a factor for each post-injury time point was estimated for each peaklet, and a Wald test was calculated for each coefficient to determine significant differences in accessibility compared to the baseline. Peaklets with FDR-controlled p-values < 0.05 in one of the four comparisons were considered to be differentially accessible. ChIPpeakAnno (v3.12.7) [14,15], the TxDb.Drerio.UCSC.danRer10.refGene UCSC annotation package (v3.4.2), and AnnotationHub (v2.10.1) were used to annotate peaklets with genes. Specifically, non-exonic (i.e., not overlapping exons by more than 50bp) peaklets overlapping a transcription start site (TSS) or within 1kb of a TSS were considered to represent proximal peaks, whereas those greater than 1kb but less than 100kb of a TSS were considered to represent distal peaks. All statistical analyses were performed in R (v3.4.3). Integrative Genome Viewer (IGV) was used to visualize RNA-seq and ATAC-seq alignments [16].

We used motif analysis to determine potential binding sites of our differentially expressed transcription factors within regions of accessible chromatin identified by ATAC-seq. Motif enrichment and discovery was carried out using various applications within the MEME Suite of motif-based sequence analysis tools (version 5.0.4) [17]. We compiled a user-supplied file for motifs corresponding to the transcription factors we identified as differentially expressed (Fig. 2), for which there were existing motifs in JASPAR or CIS-BP databases (Table S7). Motifs were formatted to MEME Motif format as specified with the MEME suite applications (meme-suite.org). The Analysis of Motif Enrichment (AME) tool [18] was used to determine motif enrichment in accessible chromatin surrounding genes in each temporal cluster (Fig. 1C). FASTA files of 500 bp peaklet sequences located proximal (≤1 kb from transcription start site) or distal (within > 1 kb, but ≤100 kb, from transcription start site) from differentially expressed genes were used in conjunction with our user-supplied motif file. Analysis was run using average odds score sequence scoring and Fisher's exact test for motif enrichment. The AME tool was also used to identify motifs within the differentially accessible chromatin regions surrounding the *jun* gene (Fig. 4), using both our user supplied motif file and the built in motif file for Eukaryotic DNA, Vertebrates (*in vivo* and *in silico*). The Find Individual Motif Occurrences (FIMO) tool [19] was used to scan for additional binding sites with the putative *jun* promoter and enhancers sequences using our user-supplied motif list. AME and FIMO analysis were both run using default parameter settings.

***References cited in Detailed Materials and Methods***


1 Westerfield, M. *The Zebrafish Book: A Guide for the Laboratory Use of Zebrafish (Danio rerio)*. (1997).

2 Udvadia, A. J. 3.6 kb Genomic sequence from Takifugu capable of promoting axon growth-associated gene expression in developing and regenerating zebrafish neurons. *Gene Expr Patterns* **8**, 382-388, doi:10.1016/j.gep.2008.05.002 (2008).

3 Bormann, P., Zumsteg, V. M., Roth, L. W. A. & Reinhard, E. Target contact regulates GAP-43 and alpha-tubulin mRNA levels in regenerating retinal ganglion cells. *J Neurosci Res* **52**, 405-419, doi:Doi 10.1002/(Sici)1097-4547(19980515)52:4<405::Aid-Jnr4>3.0.Co;2-D (1998).

4 Manoli, M. & Driever, W. Fluorescence-activated cell sorting (FACS) of fluorescently tagged cells from zebrafish larvae for RNA isolation. *Cold Spring Harb Protoc* **2012**, doi:10.1101/pdb.prot069633 (2012).

5 Bray, N. L., Pimentel, H., Melsted, P. & Pachter, L. Near-optimal probabilistic RNA-seq quantification. *Nature Biotechnology* **34**, 525-527, doi:10.1038/nbt.3519 (2016).

6 Buenrostro, J. D., Wu, B., Chang, H. Y. & Greenleaf, W. J. ATAC-seq: A Method for Assaying Chromatin Accessibility Genome-Wide. *Curr Protoc Mol Biol* **109**, 21 29 21-29, doi:10.1002/0471142727.mb2129s109 (2015).

7 Li, H. & Durbin, R. Fast and accurate short read alignment with Burrows-Wheeler transform. *Bioinformatics* **25**, 1754-1760, doi:10.1093/bioinformatics/btp324 (2009).

8 Li, H., Handsaker, B., Wysoker, A., Fennell, T., Ruan, J., Homer, N., Marth, G., Abecasis, G., Durbin, R. & Proc, G. P. D. The Sequence Alignment/Map format and SAMtools. *Bioinformatics* **25**, 2078-2079, doi:10.1093/bioinformatics/btp352 (2009).

9 Zhang, Y., Liu, T., Meyer, C. A., Eeckhoute, J., Johnson, D. S., Bernstein, B. E., Nussbaum, C., Myers, R. M., Brown, M., Li, W. & Liu, X. S. Model-based Analysis of ChIP-Seq (MACS). *Genome Biology* **9**, doi:ARTN R137; 10.1186/gb-2008-9-9-r137 (2008).

10 Lawrence, M., Huber, W., Pages, H., Aboyoun, P., Carlson, M., Gentleman, R., Morgan, M. T. & Carey, V. J. Software for Computing and Annotating Genomic Ranges. *Plos Comput Biol* **9**, doi:ARTN e1003118; 10.1371/journal.pcbi.1003118 (2013).

11 Pimentel, H., Bray, N. L., Puente, S., Melsted, P. & Pachter, L. Differential analysis of RNA-seq incorporating quantification uncertainty. *Nat Methods* **14**, 687-+, doi:10.1038/nmeth.4324 (2017).

12 Benjamini, Y. & Hochberg, Y. Controlling the False Discovery Rate - a Practical and Powerful Approach to Multiple Testing. *J R Stat Soc B* **57**, 289-300 (1995).

13 Gu, Z., Eils, R. & Schlesner, M. Complex heatmaps reveal patterns and correlations in multidimensional genomic data. *Bioinformatics* **32**, 2847-2849, doi:10.1093/bioinformatics/btw313 (2016).

14 Love, M. I., Huber, W. & Anders, S. Moderated estimation of fold change and dispersion for RNA-seq data with DESeq2. *Genome Biol* **15**, 550, doi:10.1186/s13059-014-0550-8 (2014).

15 Zhu, L. J., Gazin, C., Lawson, N. D., Pages, H., Lin, S. M., Lapointe, D. S. & Green, M. R. ChIPpeakAnno: a Bioconductor package to annotate ChIP-seq and ChIP-chip data. *BMC Bioinformatics* **11**, 237, doi:10.1186/1471-2105-11-237 (2010).

16 Robinson, J. T., Thorvaldsdottir, H., Winckler, W., Guttman, M., Lander, E. S., Getz, G. & Mesirov, J. P. Integrative genomics viewer. *Nat Biotechnol* **29**, 24-26, doi:10.1038/nbt.1754 (2011).

17 Bailey, T. L., Johnson, J., Grant, C. E. & Noble, W. S. The MEME Suite. *Nucleic Acids Res* **43**, W39-49, doi:10.1093/nar/gkv416 (2015).

18 McLeay, R. C. & Bailey, T. L. Motif Enrichment Analysis: a unified framework and an evaluation on ChIP data. *BMC Bioinformatics* **11**, 165, doi:10.1186/1471-2105-11-165 (2010).

19 Grant, C. E., Bailey, T. L. & Noble, W. S. FIMO: scanning for occurrences of a given motif. *Bioinformatics* **27**, 1017-1018, doi:10.1093/bioinformatics/btr064 (2011).

### Supplementary Figures

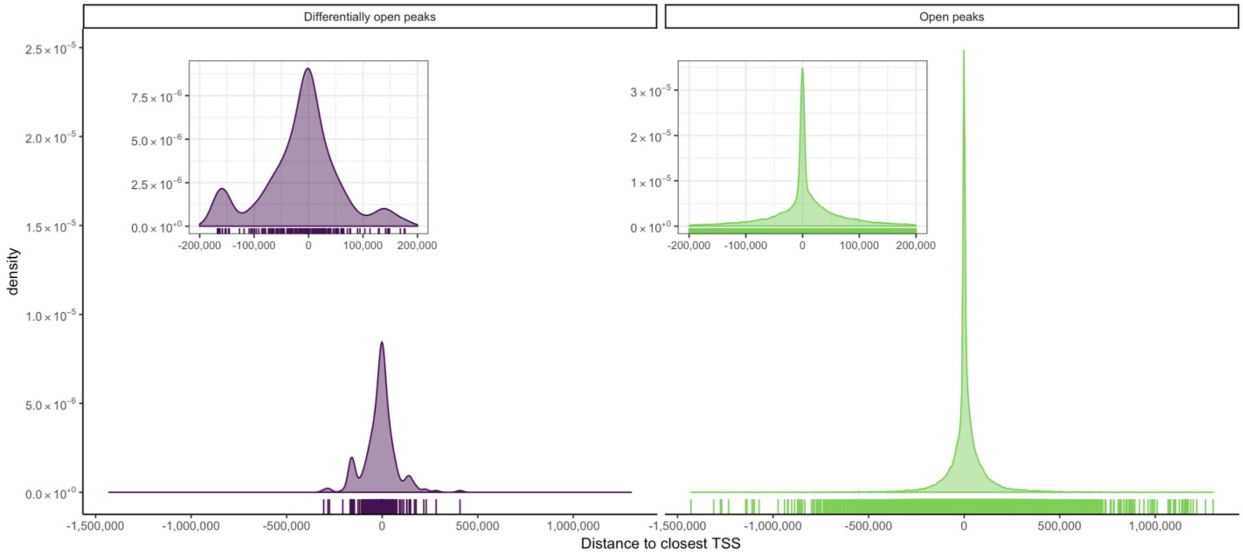


**Figure S1. Accessible chromatin is mainly located distal to annotated genes.** Distribution of accessible chromatin peaklets was plotted to determine the distance of the peaklet center to the nearest transcriptional start site (TSS). There were 42,198 high confidence open peaklets (green, right) identified ($p < 10^{-10}$). Accessibility of 233 peaklets changed in at least one time point compared to controls were considered differentially open (purple, left). Insets show magnified view of distribution of sequences within 200,000 kb of the TSS. There was a mostly equal distribution of sequences located upstream and downstream of the TSS, 20% found within 1 kb of the TSS.

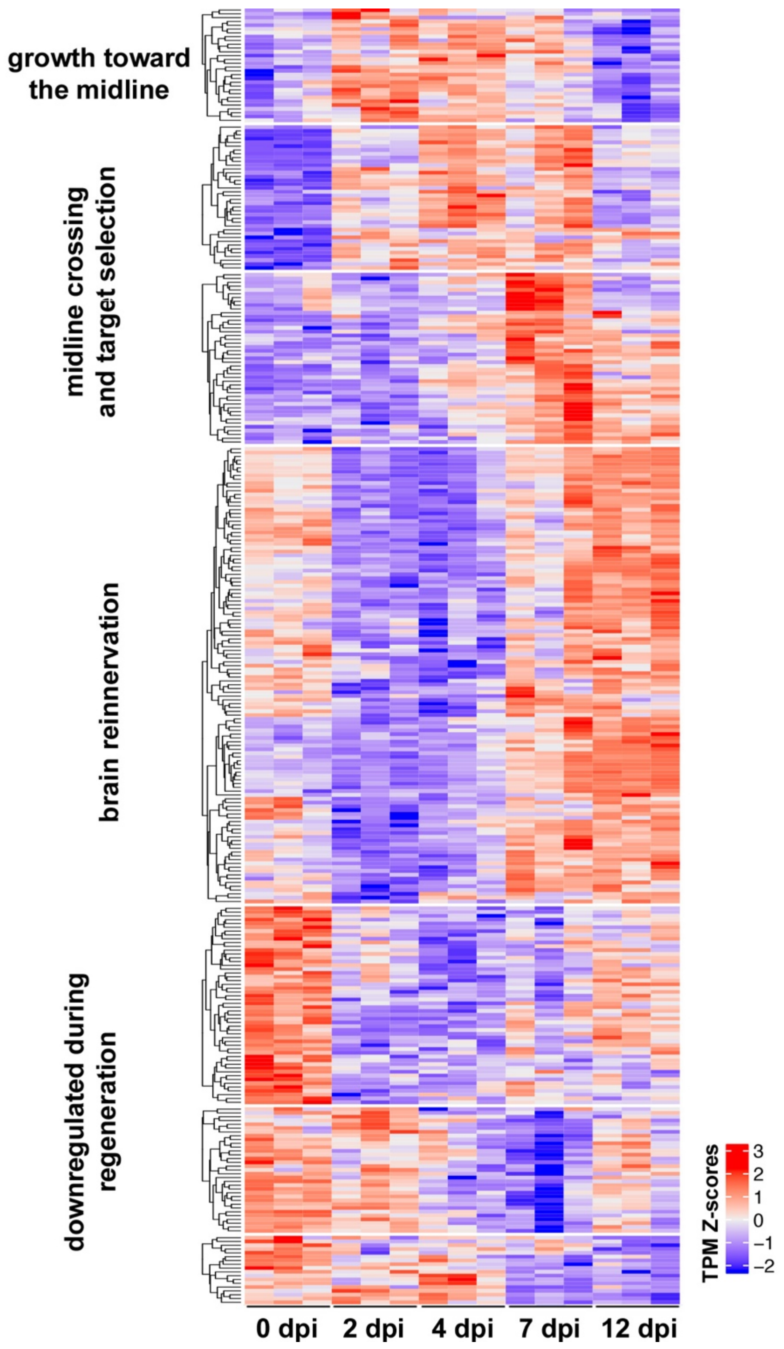


**Figure S2. Differentially expressed transcripts that encode transcription factors display regeneration stage-specific temporal patterning.** The expression heatmap of 339 differentially expressed transcripts represents 265 unique transcription factor (TF) genes. Unlike in Fig. 2C, this heatmap includes TFs with and without known DNA binding motifs.

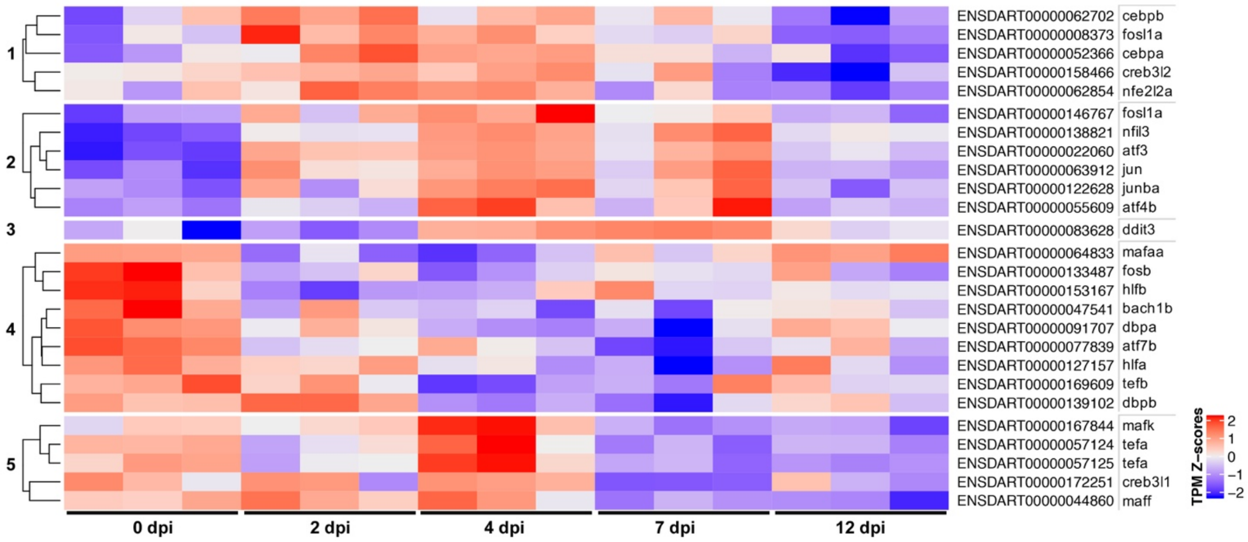


**Figure S3. Temporal patterning of differentially expressed leucine zipper (bZIP) family transcription factors.**

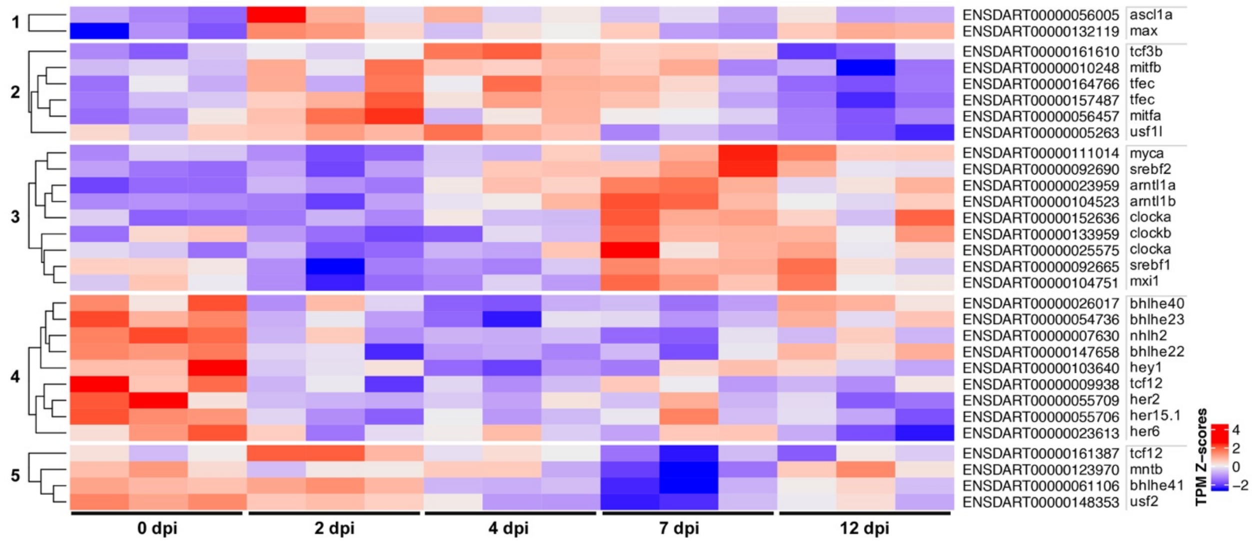


**Figure S4. Temporal patterning of differentially expressed basic helix-loop-helix (bHLH) family transcription factors.**

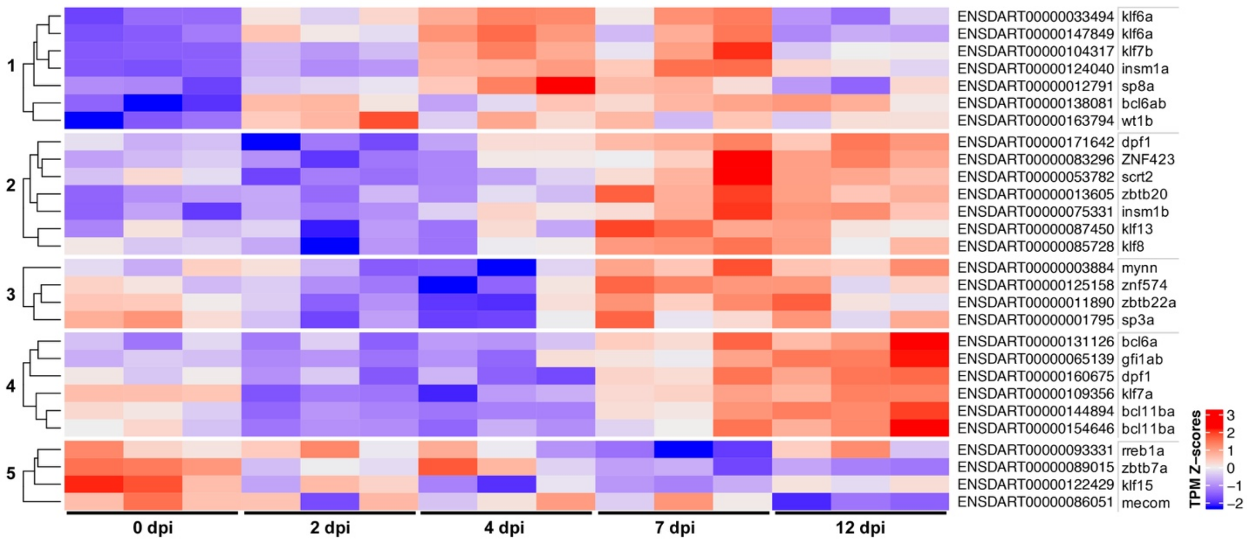


**Figure S5. Temporal patterning of differentially expressed 2 cys-2 his zinc finger (C2H2ZF) family transcription factors.**

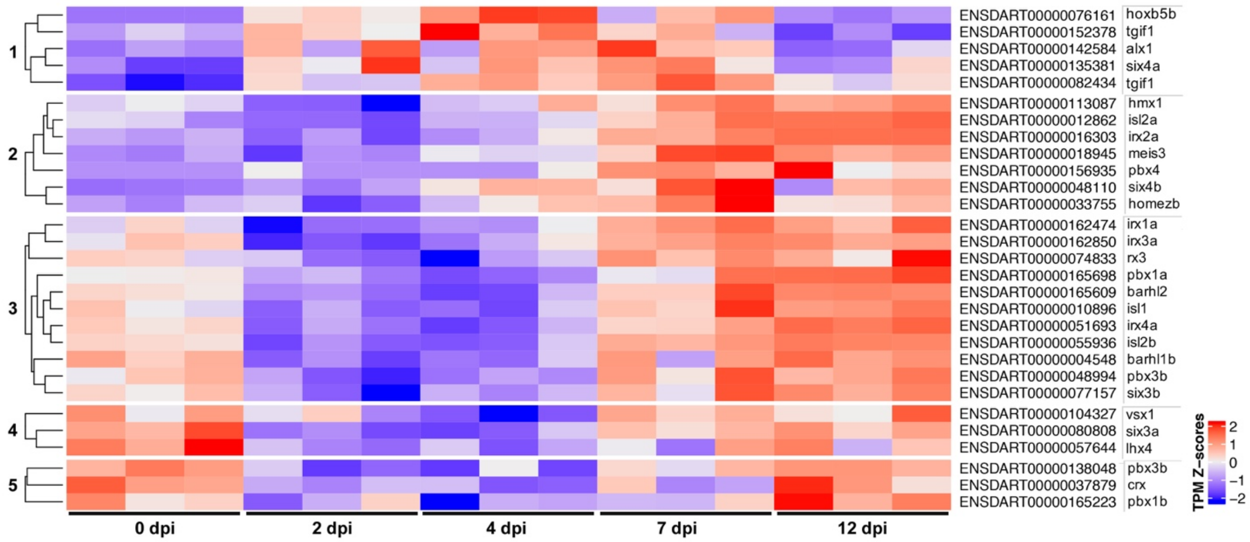


**Figure S6. Temporal patterning of differentially expressed homeodomain family transcription factors.**

>99999 TRANSGENE_contig
tcgactgtgtggcttccgtttgatccacaattgataattaaagggctggggtgaaacacggatcctcgtcttgtctcgtgtcttaactg
cttcttttccattcatacttgtaggttttaatatttcactctgagctggagcaattatcattcctgaatcatctgaactgccatggcaaca
gcttgctgatgtcagccaggagaataaataaatgaaacccttgaaacatctgccttttgctgagaattgtggttacaacttatcaata
caactaaccactgatggaatgtaactaaatccctgtgaatgatttccacatctgattagtttcattttatattttacttccactctgcgat
gtttttaaactgaccagtgttactttgcttgcaaaggaagaacatattggagttttattcacatttgcaaacacttaaacacaaaattt
agttcccagacttttgcagcaagttatactagtttaacaggttggcccttaaagtacttttgtcttctgctacttcctgatcgaaatacg
tttaatccgttttatttcacaatcggttgatgtttgtatccgagatgatacagaagtaaaactttttaaaagatttttaaaagatttttt
tacaagcgtctaatccggccccagaatcacaatgctccagcaaaagctgcagtcaagtctccaaatcttgctgaatatgtatcatcg
tccatattcatcaggtcaaatttgtctattgttgaatatttactagactaaagagtaaatgcattcaaaattctgagttggtagaggag
caaaacttctctgcttcagatttcaacaaaaggaatcactcatttactcgattactccacaccaacgttgctaggatacctgagaaat
tgcccccaaaaacctacattcaagtagaattttcatctcagacgcattcggaatattggtgaagctgctcataaactctggaaaaac
ctctgaaatgcgttcggcacttgctcacctccacttgccgggtgaggaggaccgtgcacggctgttccaggtctcctcgcaggagat
gatgcatcagtgttctgcactgcagacaaacgtggcgggactaacagccattcagacatcagactctaagctcttaatcctacagtc
agatttgatcgtggtgagacgaaaaccttcaaattgcacttgatacatatacccaagcatgtggcagcgctttgttgccatagcaac
aacaactagagttgtcttctgcgttgactcctgcctcagctgctgtggttcacccggtgtcacctgcagactctaagttgatgtgtgcg
gtgactctaagaatgctctcattactgacaatcctggaaatttaattaaaaaatactatcactttgtttcctggtgtacgaggacacg
gacgtgcgcaggttagcttagcttggtagtttactgttagtgtcaaactgatggagtaaaagactgtaacagttaattaatgagggtg
ttaaaaggtcacagggttgatgtgcaacatttctggtgcgctcggatggttctttctcaaaagtatcctattttcgagacagagtgcca
atgaagaccttacacagggtgttgtcaggttgtacctgacagcttctcaatacagcgcgacactatcgatccgtgcaatcactacag
cggcagattctacccctttactggaaagtccatgtccagatctggacatggacttagcatatgctaatgtgttccagagcagttgatg
gagtttgcatacacagttctgtgtatctgggcaaacaaggcgtccaatcatctcttcatttagagtgttgaagattgcaaaaatctgt
aattggtgaatagatcagcacgtgttaatgaaacggataatttaatttatggctggttgccattactcactatattgtagagcaatga
tttggtatctgtcaggatttgcagtgtagtgaaaatgaagcgtaacagttggagaaatactacttaagttttttttttttccaagattta
gctgtgattgatgtttcccctatcttccgtaatcaactaatagaagaaatttaattcaatctggcagcctctgaaaaacttacatcccc
tatttaagggcttcaaaaactgctcggtgaaaaactacccccagaaacaatgacttcagaacttggatgatctgaaacaatgaaa
cgcgaacagtggatttcatctgcatgattgtattaaattaaattaaacccctgagaagaaggtgtgtaaaggacaagcagggcagc
ggcagcggcagtggcaccggcagcctgcatatttccacgatgtgtgtcatccaggccagttcagaccatgatgtcacagtggcgctg
acgtattctgcattcaagtaacgatgaccaccaggaaggctgtggctcagcaggctgcttaatatcgcccatccatccatccatccat
ccatccatccatccatccatccatccatccatccatccatccatccatccatctgtagcatctgctgctgttaatgcgggggtgg
ggacactcagacaccagtttgtgttcctgtgtgtgtgcgtgtgtattatggaaaacaaatacggagtgtgggtgttttgcatggacat
gcacctgccatgctctgtaaatactgatggtgagacagtgccgggggggtgagggcaggggtgtgctcattaaaccacactgtgtg
tcaatgtgggcaggaaaaatgggggagggggagagaaacagagtcagttcttcttgtctgccacacccggccctcttgcctcttttc
ttctctctccctattcctctctctctctctctcttcctattcctctttctctttccctcgctctctcttactcctcctcctctgctcttctggtc
tatctgctggtgtttccatgagaacagagtgagctgcgtttcgggggggagtagtggtagtgagggtgatgtggcttaaatgcgcgcg
cgcgtgtgtgagtgtgcacgcacgtgtgtgtgtgtggcatcatcatctttctcacattgctcctggcagtgagatgctgctggctgagc
caaaaaccatctttgtgacatcgccgtggcaatttggaaacactcattaaagacttggaaagtgaaaagcgagggcaacatatgta
tatggctgcacgcttgatgaaaggatacgatgatacagatgagtggataaagcaaattggaggtacccttgcagctattaagggag
aacctttttttagggtggggccgcatcttgaagtcaggcagagagacgaggcggggaggagggagggagggagggagggaggga
gggaggggtttaaagggtgggcttacagtcattgtcagtgagagcgagagaaaaagagagagaggagcaactgaagaaggagt
gcaagcagactgggatgagtaaaagcactgggcagcttagctgttgatacagcctgaaagagagaaagagagagagagagagt
gtgtgtgtgtgcgcgagagcgtgtgagagagaagtgaaaaattgaggagcaaactcaagggaagaaagaagcagagcatcaaa
agggaggatagaaccaacttgaaccatgctgtgctgcataagaagaaccaaaccggtcgccaccatggtgagcaagggcgagga
gctgttcaccggggtggtgcccatcctggtcgagctggacggcgacgtaaacggccacaagttcagcgtgtccggcgagggcgag

ggcgatgccacctacggcaagctgaccctgaagttcatctgcaccaccggcaagctgcccgtgccctggcccaccctcgtgaccac
cctgacctacggcgtgcagtgcttcagccgctaccccgaccacatgaagcagcacgacttcttcaagtccgccatgcccgaaggcta
cgtccaggagcgcaccatcttcttcaaggacgacggcaactacaagacccgcgccgaggtgaagttcgagggcgacaccctggtg
aaccgcatcgagctgaagggcatcgacttcaaggaggacggcaacatcctggggcacaagctggagtacaactacaacagccac
aacgtctatatcatggccgacaagcagaagaacggcatcaaggtgaacttcaagatccgccacaacatcgaggacggcagcgtg
cagctcgccgaccactaccagcagaacacccccatcggcgacggccccgtgctgctgcccgacaaccactacctgagcacccagt
ccgccctgagcaaagaccccaacgagaagcgcgatcacatggtcctgctggagttcgtgaccgccgccgggatcactctcggcatg
gacgagctgtacaagtaaggatccactagtgatgcagatccccggatctttgtgaaggaaccttacttctgtggtgtgacataattg
gacaaactacctacagagatttaaagctctaaggtaaatataaaatttttaagtgtataatgtgttaaactactgattctaattgtttg
tgtattttagattccaacctatggaactgatgaatgggagcagtggtggaatgcctttaatgaggaaaacctgttttgctcagaaga
aatgccatctagtgatgatgaggctactgctgactctcaacattctactcctccaaaaaagaagagaaaggtagaagaccccaagg
actttccttcagaattgctaagttttttgagtcatgctgtgtttagtaatagaactcttgcttgctttgctatttacaccacaaaggaaa
aagctgcactgctatacaagaaaattatggaaaaatattctgtaacctttataagtaggcataacagttataatcataacatactgtt
ttttcttactccacacaggcatagagtgtctgctattaataactatgctcaaaaattgtgtacctttagctttttaatttgtaaaggggtt
aataaggaatatttgatgtatagtgccttgactagagatcataatcagccataccacatttgtagaggttttacttgctttaaaaaac
ctcccacacctccccctgaacctgaaacataaaatgaatgcaattgttgttgttaacttgtttattgcagcttataatggttacaaata
aagcaatagcatcacaaatttcacaaataaagcatttttttcactgcattctagttgtggtttgtccaaactcatcaatgtatcttatca
tgtctggatctgcatattctatagtgtcacctaaatctgc

**Fig. S7. Sequence of GAP43-GFP transgene.** Transgene sequence in FATSA format. GAP43-GFP fusion protein coding sequences highlighted in yellow.

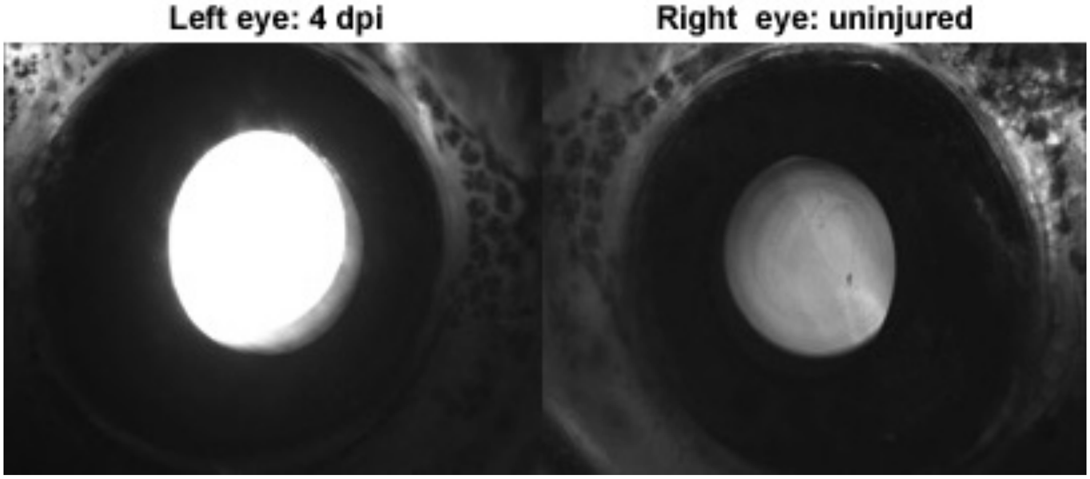


**Figure S8. Injury-induced eGFP expression in retinal ganglion cells can be visualized through the lens of intact animals.** Optic nerve injury can be validated based GFP expression, which is easily visualized through the lens. Example of injury and naïve eyes from the same fish 4 dpi.

**Table S1. Transcripts differentially expressed compared to controls (0 dpi) in at least one time point (5% FDR).**

| Ensembl transcript ID | zebrafish gene symbol | LFC: 2dpi-0dpi | LFC: 4dpi-0dpi | LFC: 7dpi-0dpi | LFC: 12dpi-0dpi |
|---|---|---|---|---|---|
| ENSDART00000000069 | slc9a3r1a | -0.019828621 | -0.165096913 | -0.579587833 | -0.318451026 |
| ENSDART00000000192 | ptpn4b | -0.253123991 | -0.355280061 | -0.416236309 | -0.192007 |
| ENSDART00000000280 | stat1b | -0.005066017 | 0.035352481 | 0.819259146 | 0.412054224 |
| ENSDART00000000486 | cntn2 | 0.015280519 | 0.733067848 | 1.819345027 | 1.713832915 |
| ENSDART00000000678 | opn1mw4 | -0.668549106 | -0.384195773 | -0.152811605 | -0.428104315 |
| ENSDART00000000744 | sypl2b | 0.279822836 | 0.157132955 | 0.007001172 | -0.094022715 |
| ENSDART00000000804 | slc8a1b | -0.216070414 | -0.730017061 | -0.564883211 | -0.309532287 |
| ENSDART00000000876 | nr4a1 | -2.262372442 | -2.399082592 | -2.145415905 | -1.543723254 |
| ENSDART00000001313 | rimbp2 | -0.29201819 | -0.480722019 | -0.187717263 | 0.029819479 |
| ENSDART00000001444 | g2e3 | 0.958887749 | 1.066626105 | 0.575974767 | 0.288448068 |
| ENSDART00000001678 | adam8a | 1.934717518 | 1.802449426 | 1.368408045 | -0.21633234 |
| ENSDART00000001795 | sp3a | -0.269877105 | -0.305437708 | -0.047306155 | -0.033496274 |
| ENSDART00000001805 | csmd2 | -0.646636229 | -0.876947398 | -0.300952212 | -0.072675683 |
| ENSDART00000001861 | slc6a22.1 | 0.506403045 | 0.610928969 | 0.211334169 | -0.287297819 |
| ENSDART00000001907 | slc16a3 | -0.228617593 | -0.852511558 | -0.452127784 | -0.215137636 |
| ENSDART00000002027 | gulp1a | -0.229497305 | -0.154479555 | -0.273130036 | -0.186901702 |
| ENSDART00000002029 | fkbp8 | -0.081020923 | -0.107047698 | -0.329759128 | -0.094897987 |
| ENSDART00000002186 | uck2a | 0.69693934 | 0.733529046 | 0.658757388 | 0.439621207 |
| ENSDART00000002250 | hs6st2 | 0.156001692 | 0.361544481 | 0.514379395 | 0.208727247 |
| ENSDART00000002309 | mafba | 0.316562739 | 0.132730939 | 0.096554959 | 0.286187946 |
| ENSDART00000002393 | rundc3aa | -0.355283185 | 0.971542583 | 1.209230568 | 0.944629469 |
| ENSDART00000002453 | acsl4b | 0.682559347 | 0.484852434 | 0.056581739 | -0.183902113 |
| ENSDART00000002469 | hspa4b | 0.156626095 | 0.21868197 | 0.285696466 | 0.103348348 |
| ENSDART00000002501 | ip6k2a | -0.189409699 | -0.526120261 | -0.315517036 | -0.114843314 |
| ENSDART00000002556 | mrto4 | 0.589078781 | 0.447556222 | 0.419365915 | 0.058126759 |
| ENSDART00000002595 | rpl21 | 0.36985839 | 0.3662658 | 0.201825483 | -0.004056179 |
| ENSDART00000002641 | kif26aa | 0.255726279 | 0.47696992 | 0.58781469 | 0.342072456 |
| ENSDART00000002684 | ddx26b | -0.289572133 | -0.160183273 | -0.164172394 | -0.052122897 |
| ENSDART00000002691 | tspan7b | -0.507607245 | -0.383440036 | -0.192864148 | 0.037733122 |
| ENSDART00000002741 | itprip | 0.724094373 | 0.694272578 | 0.397322653 | -0.062298334 |
| ENSDART00000002908 | olfm1a | -0.612571688 | -0.410284408 | -0.115653646 | 0.220963821 |
| ENSDART00000002932 | marcksb | 1.016829958 | 1.677411962 | 1.758104803 | 1.229873931 |
| ENSDART00000002945 | NPC2 (1 of many) | 3.182180208 | 2.136779184 | 1.408480393 | 0.09501531 |
| ENSDART00000002961 | rcor2 | -0.353333584 | 0.726682254 | 0.762799057 | 0.664353552 |
| ENSDART00000003001 | rpl23a | 0.412203534 | 0.396855245 | 0.244635942 | -0.052563059 |
| ENSDART00000003008 | gad1b | -0.375298425 | -0.380219055 | -0.456528647 | -0.347996362 |
| ENSDART00000003042 | mdkb | -0.457982371 | -0.488604373 | -0.629508021 | -0.485178616 |
| ENSDART00000003066 | cyth1a | 0.591135613 | 0.827117646 | 0.654961234 | 0.137500638 |
| ENSDART00000003076 | usp28 | 1.676441425 | 4.659547988 | 4.384020448 | 3.839785573 |
| ENSDART00000003170 | mid1ip1l | -0.359873821 | -0.329776238 | -0.564188118 | -0.321447922 |
| ENSDART00000003193 | gpr183a | 0.785395164 | 0.62770375 | 0.383370825 | -0.244463733 |
| ENSDART00000003248 | nek2 | 0.657099139 | 0.725246979 | 0.481556646 | 0.191253132 |
| ENSDART00000003278 | tacr3l | -0.098648497 | -0.084970679 | -0.345458842 | -0.125397678 |
| ENSDART00000003293 | mid2 | -0.366666589 | -0.252968542 | -0.080242025 | -0.101833487 |
| ENSDART00000003296 | sars | 0.182171112 | 0.390636242 | 0.505424837 | 0.283972917 |
| ENSDART00000003303 | rnf13 | 4.71068286 | 4.417862481 | 3.703911232 | 4.352869871 |
| ENSDART00000003314 | nusap1 | 0.825744448 | 0.908244509 | 0.647266522 | 0.3629471 |
| ENSDART00000003335 | snx12 | -0.102084433 | -0.181980514 | -0.47779818 | -0.16493026 |

| | | | | | |
|---|---|---|---|---|---|
| ENSDART00000003346 | pdcd2l | -0.007609279 | -0.153705238 | -0.388070334 | -0.362770652 |
| ENSDART00000003465 | gipc2 | -0.142551263 | -0.216969239 | -0.344609382 | -0.269869981 |
| ENSDART00000003475 | ppef1 | -3.70073529 | -0.21405279 | -0.165262449 | -0.400509129 |
| ENSDART00000003517 | trmt61a | 0.394655787 | 0.457245144 | 0.184403345 | 0.19719299 |
| ENSDART00000003548 | znf385a | -0.544762633 | -0.635515273 | -0.316063497 | -0.128464507 |
| ENSDART00000003550 | nmnat2 | -0.104759046 | 0.216048796 | 0.282752276 | 0.230034653 |
| ENSDART00000003612 | | -0.145765054 | -0.499207076 | -0.35686816 | -0.607333409 |
| ENSDART00000003648 | wdr3 | 0.482670527 | 0.422648714 | 0.349517488 | 0.089241222 |
| ENSDART00000003690 | acana | 0.770985542 | 0.283939017 | 0.058793221 | 0.105884754 |
| ENSDART00000003736 | anos1b | -0.470963792 | -0.442368787 | -0.444507654 | -0.318716779 |
| ENSDART00000003745 | vim | 0.110317427 | 0.81438402 | 0.975636783 | 0.646819018 |
| ENSDART00000003752 | cct3 | 0.348496547 | 0.407130663 | 0.410390239 | 0.168454536 |
| ENSDART00000003790 | pwp1 | 0.32756693 | 0.411455433 | 0.339342319 | 0.085530698 |
| ENSDART00000003825 | cplx2l | -1.121107993 | -0.523528158 | 0.430342309 | 0.533514858 |
| ENSDART00000003891 | jupa | 0.359241944 | 1.297843964 | 1.552619484 | 0.724987275 |
| ENSDART00000003913 | ifih1 | 0.480118819 | 0.196152277 | 1.199192931 | 0.158891974 |
| ENSDART00000003939 | syngr1a | 0.053450312 | -0.149997984 | -0.371497084 | -0.066149387 |
| ENSDART00000003947 | flot2a | 0.262220916 | 0.241361404 | 0.622111227 | 0.24129723 |
| ENSDART00000003998 | ewsr1b | 0.136063726 | 0.084077199 | 0.439371598 | 0.241694854 |
| ENSDART00000004034 | hpca | -0.538662149 | -0.391929378 | -0.242660916 | -0.195867715 |
| ENSDART00000004043 | enpp4 | -0.15333394 | -0.216742775 | -0.388554121 | -0.343087399 |
| ENSDART00000004065 | zgc:91909 | 0.594004329 | 0.47164071 | 0.350756766 | 0.105236702 |
| ENSDART00000004075 | uqcc2 | -0.167833305 | -0.222493804 | -0.340331706 | -0.224913678 |
| ENSDART00000004109 | gng3 | -0.129761127 | 0.224238645 | 0.384018688 | 0.329285885 |
| ENSDART00000004200 | sarm1 | 0.137263218 | 0.386089559 | 0.382312262 | 0.28972731 |
| ENSDART00000004238 | rpl7a | 0.295188189 | 0.336446742 | 0.303958011 | 0.013338352 |
| ENSDART00000004241 | inhbaa | 0.208890493 | -0.32024378 | -0.370155553 | -0.220482209 |
| ENSDART00000004392 | fkbp9 | 0.758370209 | 0.583896419 | 0.427971915 | -0.289954899 |
| ENSDART00000004416 | lrp1ba | -0.680071532 | -0.933155051 | -0.315048731 | 0.19015716 |
| ENSDART00000004420 | rab4a | -0.104588348 | -0.223085036 | -0.330744686 | -0.154422823 |
| ENSDART00000004423 | iars | 0.363891152 | 0.394414274 | 0.599518689 | 0.155539422 |
| ENSDART00000004474 | mapre1b | 0.531914947 | 0.849936108 | 0.970422656 | 0.515027849 |
| ENSDART00000004521 | arih2 | -0.124662771 | -0.251962223 | -0.001375347 | 0.051084605 |
| ENSDART00000004548 | barhl1b | -2.472825095 | -1.41656094 | -0.268019579 | 0.230183047 |
| ENSDART00000004550 | rnf145a | -0.045034762 | -0.327456151 | -0.349649694 | 0.141940758 |
| ENSDART00000004588 | asic1a | 0.159371955 | 0.158599541 | 0.248620425 | 0.288433907 |
| ENSDART00000004601 | laptm4a | -0.20461666 | -0.134693111 | -0.288435619 | -0.251769605 |
| ENSDART00000004622 | sf3b4 | 0.222493415 | 0.286225027 | 0.527062646 | 0.243398323 |
| ENSDART00000004626 | sec62 | -0.335700502 | -0.25907023 | -0.14783183 | -0.155829549 |
| ENSDART00000004656 | GCA | 0.318444393 | 0.283040087 | 0.060810751 | -0.123164086 |
| ENSDART00000004664 | tram1 | 0.410276471 | 0.318086482 | 0.103470746 | -0.163408872 |
| ENSDART00000004717 | igf1 | -0.688444818 | -0.421253017 | -0.11318285 | 0.107184312 |
| ENSDART00000004740 | rab34b | 1.81284078 | 2.340901968 | 1.779523446 | 1.639598632 |
| ENSDART00000004780 | man2b1 | 0.352578138 | 0.195228445 | 0.062298599 | -0.189062834 |
| ENSDART00000004903 | rdh10b | -0.385872948 | -0.19159484 | -0.565090954 | -0.486336464 |
| ENSDART00000005053 | slc12a4 | -0.135764977 | -0.039802706 | -0.399029267 | -0.999098872 |
| ENSDART00000005086 | atp1a1b | -0.434454878 | -0.290341007 | -0.096388542 | -0.096973501 |
| ENSDART00000005105 | psme1 | 0.846890997 | 0.915850539 | 1.159835522 | 0.124970843 |
| ENSDART00000005119 | eif3i | 0.208433038 | 0.349717288 | 0.244101269 | 0.053458955 |
| ENSDART00000005143 | mycb | 0.44444506 | 0.5301099 | 0.532628488 | 0.161240961 |
| ENSDART00000005191 | uqcrb | -0.157513362 | -0.342943294 | -0.33904791 | -0.120275765 |

| | | | | | |
|---|---|---|---|---|---|
| ENSDART00000005299 | hsd17b12a | -0.345584522 | -0.261321345 | -0.169519073 | -0.263954396 |
| ENSDART00000005337 | rimkla | -0.133054455 | 0.30447461 | 0.568720788 | 0.413212633 |
| ENSDART00000005366 | tpd52l2b | 0.128420316 | 0.410964244 | 0.59220548 | 0.186138167 |
| ENSDART00000005381 | zgc:110269 | 0.510417857 | 0.424502267 | 0.418900744 | 0.314541955 |
| ENSDART00000005382 | gadd45bb | 0.573666184 | 0.621043614 | 0.188615995 | 0.24843391 |
| ENSDART00000005453 | chd4a | 0.299643374 | 0.423708585 | 0.655765013 | 0.391160008 |
| ENSDART00000005479 | chm | -0.006745339 | 0.178224285 | 0.286548011 | 0.253225199 |
| ENSDART00000005496 | kctd9b | 0.161429839 | 0.406047159 | 0.541753399 | 0.319048316 |
| ENSDART00000005547 | gnb3b | -0.185553204 | -0.343974986 | -0.508555347 | -0.13419791 |
| ENSDART00000005568 | pdlim3b | 0.923424522 | 1.668936507 | 1.372862264 | 1.014758632 |
| ENSDART00000005573 | tmem237b | 0.093826402 | -0.187405072 | -0.456194249 | -0.210768209 |
| ENSDART00000005590 | churc1 | -0.030881068 | -0.24423848 | -0.319974325 | -0.284115834 |
| ENSDART00000005593 | casp3a | 0.498469533 | 1.198942327 | 1.332280554 | 0.912417093 |
| ENSDART00000005609 | kifap3a | 0.203573738 | 0.628570592 | 0.743430197 | 0.605809471 |
| ENSDART00000005616 | rnpep | 0.523525834 | 0.750055071 | 0.77634383 | 0.462027268 |
| ENSDART00000005638 | stxbp1b | -0.193317072 | -0.160131943 | -0.312017598 | -0.116900269 |
| ENSDART00000005720 | stat1a | 0.035880276 | 0.045195542 | 0.615137639 | 0.020671748 |
| ENSDART00000005724 | ncanb | -0.308688969 | -0.586976696 | -0.410319756 | -0.132925594 |
| ENSDART00000005733 | tma16 | -0.013223994 | -0.144098168 | -0.425909208 | -0.168461973 |
| ENSDART00000005738 | slitrk2 | -0.40615183 | -0.499760308 | -0.343598039 | -0.061449053 |
| ENSDART00000005740 | mef2aa | -3.741432455 | -2.723175321 | -1.742061709 | -1.012535024 |
| ENSDART00000005784 | itgb1bp1 | 0.440140968 | 0.384181368 | 0.282182476 | -0.012450945 |
| ENSDART00000005842 | fgf1a | -0.526483511 | -0.462706743 | -0.482028176 | -0.1224057 |
| ENSDART00000005847 | nav3 | 0.773382381 | 1.009593057 | 0.901548272 | 0.783565613 |
| ENSDART00000005869 | rpp14 | -0.113270836 | -0.030182103 | -0.321668148 | -0.171027578 |
| ENSDART00000005929 | ppp3ca | -0.536938929 | -0.189476468 | 0.016656951 | 0.377870736 |
| ENSDART00000005944 | rpl5a | 0.381988038 | 0.550810639 | 0.338076702 | 0.013320357 |
| ENSDART00000005957 | lrit1a | -0.250152343 | -0.282879742 | -0.392873609 | -0.351289311 |
| ENSDART00000005989 | dffb | 0.378338656 | 0.455339874 | 0.443055793 | 0.165200782 |
| ENSDART00000006058 | eif2s1a | 0.414016542 | 0.357076684 | 0.28859634 | 0.112415797 |
| ENSDART00000006061 | tcea3 | -0.180343963 | -0.073708986 | -0.351054612 | -0.319778395 |
| ENSDART00000006085 | cbl | 0.165119386 | 0.1775588 | 0.379174787 | 0.206196339 |
| ENSDART00000006132 | cfl1 | 0.199336875 | 0.296057395 | 0.303504194 | 0.166132927 |
| ENSDART00000006211 | prkcba | -0.217161161 | -0.38326462 | -0.338691874 | -0.182528175 |
| ENSDART00000006290 | plekhb2 | -0.293609185 | -0.084638835 | -0.118110885 | -0.124049463 |
| ENSDART00000006380 | tbx3a | -0.244591708 | -0.352646993 | -0.144447405 | 0.06961498 |
| ENSDART00000006381 | psen2 | 0.338086739 | 0.204848015 | 0.067000942 | -0.024014416 |
| ENSDART00000006417 | pgm1 | -0.367653769 | -0.345273975 | -0.348210636 | -0.086795004 |
| ENSDART00000006435 | gpr27 | -0.291018325 | -0.294873264 | -0.134468137 | 0.025589784 |
| ENSDART00000006474 | glra4b | -0.56482245 | -0.497893783 | 0.107334542 | 0.502166045 |
| ENSDART00000006489 | acsl4a | 0.116872844 | 1.158097412 | 1.477099844 | 0.59458295 |
| ENSDART00000006513 | pdhb | -0.272240562 | -0.336402892 | -0.245891496 | -0.069634702 |
| ENSDART00000006602 | pde4a | -0.284660623 | -0.353015908 | -0.053068214 | 0.108985293 |
| ENSDART00000006604 | clpp | 0.129850837 | 0.170330862 | 0.350507109 | 0.146117928 |
| ENSDART00000006612 | tbr1b | -1.166029945 | -1.008207372 | 0.617040877 | 1.169470893 |
| ENSDART00000006619 | rbpms2b | -0.996572205 | -0.617915271 | -0.145310289 | 0.143289699 |
| ENSDART00000006724 | smarcd3b | -0.214336649 | 0.06424201 | 0.499562006 | 0.504466248 |
| ENSDART00000006778 | acat2 | -0.122422954 | 0.489045334 | 0.688039864 | 0.280940334 |
| ENSDART00000006802 | cct7 | 0.251283975 | 0.283261629 | 0.291096583 | 0.139424973 |
| ENSDART00000006843 | cacng1a | 0.923524414 | 3.482209925 | 3.70002605 | 2.994326832 |
| ENSDART00000006908 | itgb3b | 1.52546911 | 1.915133846 | 1.613887964 | 0.668451164 |

| | | | | | |
|---|---|---|---|---|---|
| ENSDART00000006927 | use1 | -0.314698173 | -0.230168127 | -0.342013543 | -0.20782604 |
| ENSDART00000006990 | elovl5 | -0.296553838 | -0.148251934 | 0.039955151 | -0.062523987 |
| ENSDART00000007021 | atp6v1ba | -0.168038972 | -0.268033099 | -0.367112615 | -0.152327119 |
| ENSDART00000007103 | nuak1a | -0.384666887 | -0.409555272 | 0.121768678 | 0.275010156 |
| ENSDART00000007122 | guca1b | -0.373174817 | -0.243090423 | -0.518795688 | -0.641476933 |
| ENSDART00000007204 | ddx49 | 0.428211201 | 0.38977455 | 0.371001972 | 0.147828916 |
| ENSDART00000007208 | lrrc4bb | -0.502121898 | -0.528156819 | 0.019633372 | 0.067706598 |
| ENSDART00000007231 | psmb1 | 0.336843399 | 0.41477024 | 0.441532829 | 0.103495117 |
| ENSDART00000007271 | mtfr1l | -0.183189215 | -0.389013192 | -0.122809179 | -0.106769923 |
| ENSDART00000007293 | tcap | 1.354758323 | 0.76915174 | 1.088235317 | 0.716372996 |
| ENSDART00000007308 | wnt10a | -1.246823676 | -1.58888825 | -0.322358389 | 0.209755977 |
| ENSDART00000007401 | MAP3K13 | -0.231843501 | -0.363439198 | -0.331818795 | -0.269324908 |
| ENSDART00000007512 | pole3 | 0.11728021 | 0.33533824 | 0.217028028 | 0.042569097 |
| ENSDART00000007522 | anos1a | -0.500742525 | -0.407287946 | -0.526812004 | -0.211300468 |
| ENSDART00000007531 | slit2 | -0.281096395 | -0.311205466 | -0.174560483 | 0.038385548 |
| ENSDART00000007584 | snap25a | -0.520469397 | -0.457109198 | -0.208714288 | -0.117586032 |
| ENSDART00000007587 | gtf2h2 | 0.253100515 | 0.350059727 | 0.329257684 | 0.204207466 |
| ENSDART00000007624 | plch2a | -0.294880531 | -0.004127951 | 0.226839247 | 0.19499931 |
| ENSDART00000007630 | nhlh2 | -0.28794611 | -0.324673306 | -0.385652145 | -0.261092419 |
| ENSDART00000007642 | zgc:110239 | 0.831334701 | 0.383173307 | 0.0742642 | -0.168536089 |
| ENSDART00000007778 | grik1a | -0.423674769 | -0.190804556 | -0.271114709 | -0.0631136 |
| ENSDART00000007789 | idh1 | 0.209391072 | 0.491026617 | 0.59658334 | 0.190896441 |
| ENSDART00000007797 | slc30a4 | 0.328612859 | 0.21837447 | 0.122260678 | 0.141372312 |
| ENSDART00000007806 | zbtb16a | -0.634450415 | -0.731920804 | 0.190781828 | -0.364381887 |
| ENSDART00000007827 | spra | 0.429147243 | 0.5653912 | 0.349944557 | -0.12347627 |
| ENSDART00000007856 | fkbp16 | -0.168302487 | -0.129290801 | -0.313457803 | -0.012446152 |
| ENSDART00000007857 | mettl2a | 0.337109376 | 0.448868897 | 0.378403392 | 0.127678255 |
| ENSDART00000007961 | nt5c2l1 | 0.559923749 | 0.945665914 | 1.614610333 | 0.555210778 |
| ENSDART00000007972 | dlgap4b | -0.346497047 | -0.558852799 | -0.093125576 | -0.08715572 |
| ENSDART00000008010 | pdk2a | -0.070635847 | 0.148907691 | -0.31523766 | -0.171233724 |
| ENSDART00000008038 | sulf2a | 0.420520728 | 1.274767381 | 2.360438199 | 2.491815243 |
| ENSDART00000008058 | aak1a | -0.071484403 | -0.294378944 | -0.458052722 | -0.070668535 |
| ENSDART00000008152 | sgk2b | 1.195404929 | 0.789404089 | 0.784574894 | 0.645661759 |
| ENSDART00000008287 | pgam1a | -0.270559913 | -0.416576315 | -0.324017463 | -0.168254728 |
| ENSDART00000008302 | insra | -0.066772744 | -0.178139649 | -0.319689592 | -0.098012672 |
| ENSDART00000008326 | pon2 | 0.005358778 | -0.010211923 | -0.591626082 | -0.490299784 |
| ENSDART00000008373 | fosl1a | 0.78271954 | 0.570182098 | 0.226810387 | -0.332448721 |
| ENSDART00000008402 | sart3 | -0.246563675 | -0.213758299 | -0.263959387 | -0.22518745 |
| ENSDART00000008594 | tmem178 | -0.626672748 | -0.469011917 | -0.272377327 | 0.052520712 |
| ENSDART00000008607 | ttyh2l | 0.127438148 | -0.071060271 | -0.346686037 | -0.327410338 |
| ENSDART00000008638 | rgma | -0.505076193 | -0.28541733 | -0.181405309 | 0.127564695 |
| ENSDART00000008663 | adam10b | 0.007023444 | 0.111335196 | 0.430661155 | 0.328928973 |
| ENSDART00000008711 | gys1 | -0.668450567 | -0.607630136 | -0.28295897 | -0.009086585 |
| ENSDART00000008785 | anp32a | -0.194673883 | -0.086461888 | -0.275568041 | -0.184432261 |
| ENSDART00000008807 | rpl12 | 0.258765274 | 0.397825613 | 0.190712058 | -0.053203791 |
| ENSDART00000008840 | otofa | -0.788283647 | -0.975123326 | -0.33101193 | -0.149384457 |
| ENSDART00000008854 | wsb1 | 0.135165037 | 0.298497884 | 0.131347407 | -0.114776528 |
| ENSDART00000008906 | znf503 | -0.367880648 | -0.389528031 | 0.161538025 | 0.184463 |
| ENSDART00000008986 | atp6v1e1a | 0.082323928 | -0.133398426 | -0.212735096 | -0.280773093 |
| ENSDART00000009164 | esco2 | 1.573729566 | 0.878542321 | 0.358323204 | 0.160150267 |
| ENSDART00000009178 | impdh2 | 0.589155818 | 0.398351191 | 0.511972012 | 0.456829015 |

| | | | | | |
|---|---|---|---|---|---|
| ENSDART00000009194 | aimp2 | 0.035867833 | 0.204745928 | 0.516008081 | 0.307277143 |
| ENSDART00000009241 | rpl35 | 0.440738233 | 0.45418276 | 0.310964715 | 0.055658665 |
| ENSDART00000009277 | tuba1a | 0.466164103 | 1.394077049 | 1.773814961 | 1.31379942 |
| ENSDART00000009337 | eno1a | -0.396364992 | -0.428345122 | -0.288646494 | -0.130833414 |
| ENSDART00000009343 | pyroxd2 | 0.778556587 | 0.730253969 | 0.295998209 | 0.188824159 |
| ENSDART00000009393 | col1a1a | 0.640444957 | 1.804362836 | 2.839654111 | 1.436438571 |
| ENSDART00000009477 | cct8 | 0.2739469 | 0.4225965 | 0.299361505 | 0.113502813 |
| ENSDART00000009484 | cct6a | 0.227309436 | 0.27374384 | 0.264898076 | 0.101996724 |
| ENSDART00000009549 | rhag | 0.97898574 | 1.27206332 | 1.678759018 | 0.617102069 |
| ENSDART00000009569 | slc12a5b | -0.543274129 | -0.639831062 | -0.420973005 | -0.072437192 |
| ENSDART00000009609 | eif5a | 0.382861639 | 0.513300938 | 0.466153608 | 0.171038659 |
| ENSDART00000009653 | kcna1b | -0.51649998 | -0.917921415 | -0.602149268 | -0.420615578 |
| ENSDART00000009689 | mhc1uba | 0.3733109 | 0.635354786 | 0.756757815 | 0.245538965 |
| ENSDART00000009691 | scml4 | -0.348233268 | -0.325522885 | -0.113421585 | 0.115178638 |
| ENSDART00000009740 | smad7 | -0.082405078 | -0.352637537 | -0.492507075 | -0.066549718 |
| ENSDART00000009777 | glra3 | -0.385455571 | -0.554614528 | -0.516083394 | -0.22262256 |
| ENSDART00000009888 | casq1b | 2.284244029 | 1.995698495 | 4.521796612 | 3.078320888 |
| ENSDART00000009892 | gabbr1a | -0.618262887 | -0.683782394 | 0.013008531 | 0.484024166 |
| ENSDART00000009938 | tcf12 | -0.385864267 | -0.351608927 | -0.283149928 | -0.323464659 |
| ENSDART00000009952 | zfand5a | -0.376361025 | -0.47427013 | -1.180526792 | -1.063292908 |
| ENSDART00000010041 | dhfr | 0.095253657 | -0.024940068 | -0.334824981 | -0.34089932 |
| ENSDART00000010046 | rhpn2 | -0.531285649 | -0.280012714 | -0.367908087 | -0.481376216 |
| ENSDART00000010104 | crtap | 0.225199403 | 0.115819527 | -0.25953471 | -0.588755148 |
| ENSDART00000010119 | eef1a2 | -1.586399577 | -1.404358062 | -0.455450934 | 0.174130308 |
| ENSDART00000010140 | igf2bp3 | 0.357904839 | 0.436869624 | 1.38352682 | 1.118120985 |
| ENSDART00000010144 | pvalb2 | 0.379417352 | 4.690875982 | 5.447468716 | 3.629116223 |
| ENSDART00000010199 | fam219ab | -0.329176512 | -0.152550548 | -0.14317011 | 0.01066054 |
| ENSDART00000010246 | ugt1ab | 1.578212697 | 1.169819697 | 0.509084599 | -0.046131609 |
| ENSDART00000010256 | eif3m | 0.301540845 | 0.415293475 | 0.295465222 | 0.073592881 |
| ENSDART00000010257 | fam73a | -0.203941778 | -0.403469011 | -0.199714259 | -0.045216291 |
| ENSDART00000010261 | pno1 | 0.396735226 | 0.379415162 | 0.20506239 | -0.042887698 |
| ENSDART00000010271 | aida | 0.207262438 | 0.531449642 | 0.462716258 | 0.148174666 |
| ENSDART00000010274 | dpysl5a | 0.119382655 | 1.052869734 | 1.414498731 | 1.230769727 |
| ENSDART00000010282 | cbr1l | 0.612622623 | 0.498855932 | 0.466697421 | -0.088158588 |
| ENSDART00000010378 | myo3b | 0.121341422 | -0.396947151 | -0.677290933 | -0.232688387 |
| ENSDART00000010420 | actr1 | 0.079833108 | 0.201277948 | 0.258101559 | 0.148775132 |
| ENSDART00000010452 | zgc:91860 | -0.346186809 | -0.437676548 | -0.463741509 | -0.203120402 |
| ENSDART00000010495 | znrf1 | 0.045008698 | 0.17288769 | 0.379569247 | 0.252907322 |
| ENSDART00000010512 | zgc:92907 | -0.132831847 | -0.132364384 | -0.338242819 | -0.200606823 |
| ENSDART00000010647 | rcc2 | 0.312248055 | 0.54892666 | 0.398800486 | 0.1537421 |
| ENSDART00000010683 | impa1 | 0.486043053 | 0.448191747 | 0.382482237 | 0.291162031 |
| ENSDART00000010757 | rgmb | -0.151394227 | 0.132378035 | 0.469172545 | 0.35352068 |
| ENSDART00000010824 | acy1 | -0.16534468 | -0.281016974 | -0.38333817 | -0.31990568 |
| ENSDART00000010982 | fgf13a | -0.282182655 | -0.227308636 | -0.321295112 | -0.222652121 |
| ENSDART00000010997 | tpm3 | 0.446911874 | 0.508209148 | 0.705423057 | 0.171806011 |
| ENSDART00000011004 | mfsd5 | -0.162600193 | -0.289194121 | -0.512584457 | -0.472173717 |
| ENSDART00000011052 | eif3d | 0.194265429 | 0.254685579 | 0.234360887 | -0.025541379 |
| ENSDART00000011135 | kita | -0.1368269 | -0.266279657 | -0.502022945 | -0.592221276 |
| ENSDART00000011143 | myd88 | 0.15184495 | 0.21846972 | -0.215204616 | -0.645828486 |
| ENSDART00000011149 | fam185a | -0.225870361 | -0.305876809 | -0.396323932 | -0.356813487 |
| ENSDART00000011224 | itga10 | 2.439207816 | 3.888845597 | 4.302533523 | 3.055176776 |

| | | | | | |
|---|---|---|---|---|---|
| ENSDART00000011229 | sub1b | -0.082257398 | -0.179031151 | -0.405374566 | -0.271111897 |
| ENSDART00000011251 | rpl3 | 0.32045382 | 0.346921833 | 0.271797845 | 0.014406427 |
| ENSDART00000011258 | npl | 0.373272645 | 0.54061543 | 0.468726439 | 0.170404917 |
| ENSDART00000011283 | cnr1 | 0.295670673 | 0.19447276 | 0.12958679 | 0.047335468 |
| ENSDART00000011287 | aqp4 | -0.584037248 | -0.688442177 | -0.670108654 | -0.488547149 |
| ENSDART00000011305 | dpp3 | 0.260304454 | 0.228408409 | 0.405249148 | 0.162562787 |
| ENSDART00000011317 | ntm | -0.235410973 | -0.452806913 | -0.580240928 | -0.252707536 |
| ENSDART00000011362 | arrdc2 | 0.681126697 | 0.66886629 | 0.476399009 | 0.092022718 |
| ENSDART00000011398 | si:ch73-335l21.1 | -0.01193014 | -0.263037651 | -0.302583762 | -0.048051484 |
| ENSDART00000011447 | sae1 | 0.282597281 | 0.674186287 | 0.734155697 | 0.424524708 |
| ENSDART00000011453 | sypb | -0.342744234 | -0.453708067 | -0.33513289 | -0.074863584 |
| ENSDART00000011456 | tsg101b | 0.363139745 | 0.1150057 | 0.061390783 | -0.157671493 |
| ENSDART00000011519 | slc6a1l | 0.105782424 | -0.162202998 | -0.466558128 | -0.662799909 |
| ENSDART00000011568 | syngr3a | -0.064295047 | 0.234968681 | 0.63147157 | 0.633880469 |
| ENSDART00000011570 | zgc:101716 | 0.810996291 | 0.904458728 | 0.54949556 | 0.44631829 |
| ENSDART00000011627 | irx7 | -0.27830639 | -0.509115666 | -0.233145912 | -0.150983114 |
| ENSDART00000011691 | baxa | 0.500319714 | 0.752407609 | 0.690306826 | 0.30333895 |
| ENSDART00000011699 | nono | 0.031722902 | 0.329157035 | 0.240624596 | 0.057167441 |
| ENSDART00000011863 | hdlbpa | 0.44689202 | 0.322223522 | 0.244841165 | -0.050194137 |
| ENSDART00000011865 | sec23b | -0.409405473 | 0.102005828 | 0.534469651 | 0.138588714 |
| ENSDART00000011878 | eif4a1b | 0.134279506 | 0.344064551 | 0.309281202 | 0.284267369 |
| ENSDART00000011936 | ccdc106a | -0.452156269 | -0.26003137 | -0.330808493 | -0.261299438 |
| ENSDART00000012023 | faimb | -0.164145301 | -0.164693039 | -0.391621682 | -0.153828799 |
| ENSDART00000012119 | zgc:110366 | -0.07598566 | 0.059679618 | -0.31484475 | -0.244737795 |
| ENSDART00000012164 | tmod2 | 0.222306993 | 0.411676025 | 0.560901639 | 0.297325519 |
| ENSDART00000012229 | fkbp1b | -0.199888531 | -0.218085861 | -0.490853212 | -0.25173597 |
| ENSDART00000012247 | dhcr24 | 2.392110544 | 2.697063662 | 3.119814675 | 2.653468904 |
| ENSDART00000012256 | tnni2a.3 | 0.628164776 | 3.762159614 | 4.672246791 | 3.241031276 |
| ENSDART00000012357 | sav1 | 0.272575283 | 0.33526037 | 0.280505115 | -0.039530435 |
| ENSDART00000012376 | gabrr1 | -0.444264456 | -0.314539223 | -0.677749333 | -0.339326443 |
| ENSDART00000012391 | cabp1a | -0.440359676 | -0.517676368 | -0.117661538 | 0.040314239 |
| ENSDART00000012450 | dvl2 | 0.778001668 | 0.759654351 | 0.507462592 | -0.16358587 |
| ENSDART00000012478 | mmadhc | -0.097599671 | -0.214733382 | -0.311326383 | -0.186284094 |
| ENSDART00000012546 | ctbp2a | -0.410954451 | -0.209595307 | -0.078015176 | -0.146761341 |
| ENSDART00000012580 | | -0.21861763 | 0.154213857 | -0.169850923 | -1.246152902 |
| ENSDART00000012673 | gnb3a | -0.611271729 | -0.360057677 | -0.265096652 | -0.439594606 |
| ENSDART00000012677 | OTUD7A | -0.320444813 | -0.33972382 | -0.130240255 | -0.027710623 |
| ENSDART00000012686 | dnase1l4.1 | 0.654985436 | 1.279054377 | 1.240763186 | 0.531962605 |
| ENSDART00000012718 | fabp11b | 1.386159817 | 1.224465105 | 0.518278623 | -0.419802831 |
| ENSDART00000012791 | sp8a | 1.57251014 | 2.331511764 | 2.00847442 | 1.039704847 |
| ENSDART00000012822 | CR354540.1 | -0.197702519 | 0.015464017 | -0.410213127 | -0.063874026 |
| ENSDART00000012859 | psma6b | 0.220061705 | 0.322650996 | 0.197619136 | 0.013143764 |
| ENSDART00000012862 | isl2a | -0.663813396 | 0.011664777 | 0.746201858 | 1.074694305 |
| ENSDART00000012938 | phgdh | -0.315221747 | -0.241109028 | -0.448879252 | -0.249449553 |
| ENSDART00000012940 | grm2b | -0.52717467 | -0.642099924 | -0.570224924 | -0.520457538 |
| ENSDART00000013003 | tfap2b | -0.299930992 | -0.578681033 | -0.25971747 | -0.279510711 |
| ENSDART00000013066 | ercc3 | 0.273698243 | 0.1312947 | 0.187090251 | 0.076904615 |
| ENSDART00000013117 | syt5b | -0.473395784 | -0.439637374 | -0.372750952 | -0.393185839 |
| ENSDART00000013228 | cacna1aa | -0.425294008 | -0.48842278 | 0.10428355 | 0.42170802 |
| ENSDART00000013229 | gnaq | 0.258587746 | 0.466151888 | 0.597563156 | 0.37656513 |
| ENSDART00000013263 | ugdh | 0.441443931 | 0.390023192 | 0.086037142 | -0.108373731 |

| | | | | | |
|---|---|---|---|---|---|
| ENSDART00000013311 | grm6a | -0.718441456 | -0.89272341 | -0.019297194 | 0.441642652 |
| ENSDART00000013360 | ppp1r3cb | -0.357586146 | -0.346668382 | -0.405594995 | -0.235559662 |
| ENSDART00000013409 | prmt3 | 0.583250552 | 0.486757337 | 0.420985526 | -0.028812313 |
| ENSDART00000013411 | cahz | -0.282657507 | -0.146895201 | -0.382063619 | -0.133810291 |
| ENSDART00000013449 | CHST13 | 0.129876823 | 0.161538207 | -0.188391866 | -0.557482268 |
| ENSDART00000013575 | bzw1a | 0.288880361 | 0.296312188 | 0.255308329 | 0.039925909 |
| ENSDART00000013588 | klhl41b | 0.067761781 | 2.964864958 | 3.322998807 | 2.346652378 |
| ENSDART00000013605 | zbtb20 | 0.021191929 | 0.070670156 | 0.462529396 | 0.345025042 |
| ENSDART00000013690 | rplp2l | 0.526551673 | 0.441805947 | 0.587852857 | 0.20929518 |
| ENSDART00000013781 | mcm6 | 1.053827708 | 1.005302691 | 0.681665045 | 0.184026433 |
| ENSDART00000013785 | insig1 | -0.148803263 | 0.843708665 | 1.082868772 | 0.366540247 |
| ENSDART00000013797 | asb8 | -0.250585187 | -0.305587852 | -0.166685061 | -0.071337431 |
| ENSDART00000013835 | bloc1s1 | 0.447187715 | 0.435712256 | 0.356793883 | 0.203995374 |
| ENSDART00000013839 | tmbim4 | -0.093382601 | -0.138429676 | -0.303111951 | -0.191824583 |
| ENSDART00000013961 | mycla | 0.091841991 | 0.503579515 | 0.855574629 | 0.294011787 |
| ENSDART00000014021 | slc25a39 | -0.175830036 | -0.233948732 | -0.630252381 | -0.299226836 |
| ENSDART00000014031 | dpf2 | 0.062076591 | -0.192206303 | -0.356876024 | -0.045146421 |
| ENSDART00000014036 | optn | -0.037971093 | -0.094029875 | -0.163689177 | -0.30174806 |
| ENSDART00000014049 | wdr36 | 0.429321608 | 0.423447996 | 0.351992847 | 0.181701695 |
| ENSDART00000014058 | zgc:100829 | 0.836226257 | 1.366715788 | 1.414785917 | 0.344237354 |
| ENSDART00000014095 | rap2c | 0.031678102 | 0.116919511 | 0.30035611 | 0.141485249 |
| ENSDART00000014098 | ggctb | -0.310700478 | -0.20595478 | -0.456981273 | -1.259991115 |
| ENSDART00000014168 | zfp36l1b | -0.068733676 | -0.015026144 | -0.139788147 | -0.387873136 |
| ENSDART00000014183 | colgalt2 | -0.33250641 | -0.46357994 | -0.396923695 | -0.603253205 |
| ENSDART00000014207 | myl1 | 0.550709909 | 1.927654149 | 2.148710994 | 1.508189135 |
| ENSDART00000014274 | glcea | -0.30114601 | -0.147144768 | -0.347048688 | -0.226291229 |
| ENSDART00000014306 | mpp5a | -0.106349761 | -0.16109327 | -0.308554572 | -0.109175977 |
| ENSDART00000014568 | urod | 0.384025692 | 0.230369566 | 0.194017408 | -0.024966445 |
| ENSDART00000014632 | katnb1 | -0.1660519 | -0.366507385 | -0.386560357 | -0.02476169 |
| ENSDART00000014668 | pcsk1 | -0.444651661 | -0.49391188 | -0.327964124 | -0.406864512 |
| ENSDART00000014726 | tp53i11b | -0.378677803 | -0.267454179 | -0.193372925 | -0.148965425 |
| ENSDART00000014729 | arpc1a | 0.133812581 | 0.244815919 | 0.290121908 | 0.113999438 |
| ENSDART00000014806 | npas2 | -0.341343772 | 0.009000027 | 0.555800516 | 0.156695025 |
| ENSDART00000014843 | bdnf | -0.572929745 | -0.434797241 | -0.517838864 | -0.969054184 |
| ENSDART00000014871 | akr7a3 | 0.445881137 | 0.133128629 | 0.063605049 | 0.058812872 |
| ENSDART00000014877 | robo2 | -0.381239115 | -0.949448522 | 0.586346925 | 0.554614257 |
| ENSDART00000014897 | srgap1b | -0.562468132 | -0.53652608 | -0.21984937 | 0.24230424 |
| ENSDART00000014922 | arhgap22 | 0.337853511 | 0.496203573 | 0.771443869 | 0.664153347 |
| ENSDART00000014983 | zgc:153867 | 0.36977967 | 0.600215172 | 0.463408621 | 0.079627153 |
| ENSDART00000015034 | blvrb | 0.700108065 | 0.61517331 | 0.539502173 | -0.031641171 |
| ENSDART00000015040 | hrasb | -0.035400047 | 0.468215746 | 0.71708775 | 0.509094476 |
| ENSDART00000015081 | COX5B (1 of many) | -0.161517675 | -0.197504434 | -0.340992957 | -0.152202035 |
| ENSDART00000015092 | col1a1b | 0.334762626 | 1.456833944 | 2.512390696 | 1.131086568 |
| ENSDART00000015095 | uts1 | -0.304915972 | -0.432455801 | -0.651436618 | -0.693973757 |
| ENSDART00000015103 | hps3 | 0.398719216 | 0.34180849 | 0.31802597 | 0.20234989 |
| ENSDART00000015193 | chmp4bb | 0.349511966 | 0.500219693 | 0.425249872 | 0.076172611 |
| ENSDART00000015279 | rtn4rl1a | -0.620130019 | -0.765846328 | -0.205912996 | 0.07424375 |
| ENSDART00000015286 | ankrd13b | -0.294042347 | 0.149598935 | 0.484140903 | 0.605167909 |
| ENSDART00000015333 | gbx2 | -0.202064704 | -0.378779754 | -0.261598975 | -0.176372699 |
| ENSDART00000015374 | cyb5r1 | 0.481284363 | 0.621601719 | 0.549959374 | 0.29311095 |
| ENSDART00000015401 | ercc6l | 1.183315334 | 0.605525038 | 0.299117061 | -0.215859916 |

| | | | | | |
|---|---|---|---|---|---|
| ENSDART00000015418 | irf2bpl | -0.517127602 | -0.432851187 | -0.112506589 | -0.180160421 |
| ENSDART00000015628 | klhl24b | 0.257620397 | 0.582600724 | 0.358856753 | 0.009507758 |
| ENSDART00000015629 | stxbp1a | -0.407999477 | -0.254916105 | -0.074938098 | 0.048338374 |
| ENSDART00000015632 | nkain1 | 0.221220236 | 0.815235139 | 1.176280245 | 0.965286214 |
| ENSDART00000015710 | snrkb | -0.234771599 | -0.444259361 | -0.383002363 | -0.24868656 |
| ENSDART00000015732 | mylz3 | 0.011883193 | 1.89110665 | 2.420669252 | 1.523105093 |
| ENSDART00000015755 | rasl11b | -0.21929532 | -0.630137444 | -0.720127948 | -0.404206999 |
| ENSDART00000015777 | abce1 | 0.253644034 | 0.246716072 | 0.263123796 | 0.067670414 |
| ENSDART00000015827 | tnr | -0.348895518 | -0.471355706 | -0.275641147 | -0.100175719 |
| ENSDART00000015841 | gstt1b | 0.844147349 | 0.559916631 | 0.300223231 | -0.404935077 |
| ENSDART00000015951 | bsg | -0.039046362 | -0.213267965 | -0.489263219 | -0.308938953 |
| ENSDART00000015956 | efna1b | -0.412330789 | -0.211680079 | -0.214205067 | -0.214663513 |
| ENSDART00000015979 | farsb | 0.087018819 | 0.265294461 | 0.405710239 | 0.211888538 |
| ENSDART00000016053 | rnf144aa | -0.119412737 | -0.323472384 | -0.040822296 | 0.138732642 |
| ENSDART00000016057 | ctnnal1 | 0.110311248 | 0.013228232 | -0.224558056 | -0.622388694 |
| ENSDART00000016099 | CASKIN2 | -0.196661177 | -0.318487444 | -0.498724654 | -0.242993173 |
| ENSDART00000016112 | capns1b | 0.258538261 | 0.719472725 | 0.61845775 | 0.248329876 |
| ENSDART00000016135 | nfe2l3 | 0.030439041 | 0.342226134 | 0.430712572 | 0.086440986 |
| ENSDART00000016143 | SPIN4 (1 of many) | -0.27419945 | -0.33038212 | -0.171096922 | -0.003336972 |
| ENSDART00000016181 | ndrg3a | -0.359622862 | -0.274997666 | -0.226177976 | -0.143111599 |
| ENSDART00000016283 | psmd11b | 0.064194443 | 0.317482468 | 0.345168865 | 0.113362384 |
| ENSDART00000016303 | irx2a | -0.381386746 | 0.064410059 | 0.752093904 | 0.989751717 |
| ENSDART00000016350 | pgam1b | -0.297181524 | -0.257905953 | 0.060817132 | 0.142368471 |
| ENSDART00000016360 | si:ch73-199e17.1 | 2.690029092 | 3.363503415 | 3.69470671 | 3.529091333 |
| ENSDART00000016370 | dirc2 | 0.134905484 | 0.442850685 | 0.565287267 | 0.464631841 |
| ENSDART00000016464 | dcps | 0.045608002 | 0.377363095 | -0.076988765 | 0.017970252 |
| ENSDART00000016535 | kcns3a | -1.299949965 | -1.227726419 | -0.464780559 | 0.150117362 |
| ENSDART00000016591 | fgf6a | 0.502410209 | 1.375431955 | 1.297903392 | 1.026892284 |
| ENSDART00000016597 | nfkbiab | 0.207474067 | 0.316720279 | 0.269395099 | 0.411969915 |
| ENSDART00000016602 | cdh23 | -0.199390412 | -0.313772685 | -0.367682037 | -0.223939401 |
| ENSDART00000016628 | fam129bb | 0.861865647 | 1.020541987 | 0.89470832 | 0.182764632 |
| ENSDART00000016710 | scrn3 | -0.247629441 | -0.171036354 | 0.44401532 | -0.052873668 |
| ENSDART00000016791 | eif3c | -0.057805139 | -0.126818292 | -0.360850099 | 0.005995136 |
| ENSDART00000016803 | grpel1 | -0.106418203 | -0.12256904 | -0.467582595 | -0.298130125 |
| ENSDART00000016814 | fmnl2a | 0.496433192 | 0.213976057 | -0.143298216 | -0.634969289 |
| ENSDART00000016860 | ppp2r1bb | 0.018499492 | 0.284521267 | 0.321334616 | 0.193996267 |
| ENSDART00000016864 | slc35f6 | 0.392001078 | 0.280281029 | 0.187768856 | 0.16995993 |
| ENSDART00000016890 | eif6 | 0.391815405 | 0.223439355 | 0.155230229 | 0.056706473 |
| ENSDART00000016916 | gria4b | -0.27171917 | -0.41554348 | -0.145968175 | 0.021630803 |
| ENSDART00000016946 | glud1a | 0.489439701 | 0.519599049 | 0.659106358 | 0.391317016 |
| ENSDART00000016983 | spon1a | -0.566333419 | -0.54864414 | -0.425205045 | -0.214533594 |
| ENSDART00000017148 | gclc | 0.213009245 | 0.302558434 | 0.032586299 | -0.083839473 |
| ENSDART00000017153 | hps4 | 0.574215829 | 0.524013089 | 0.377950963 | 0.050651634 |
| ENSDART00000017176 | dkc1 | 0.45463167 | 0.261033913 | 0.276931433 | 0.02149911 |
| ENSDART00000017202 | kcnk1b | -0.464281499 | -0.11026216 | 0.124659295 | 0.114555904 |
| ENSDART00000017229 | ncam1a | -0.359842421 | -0.143522661 | 0.045009487 | 0.122759016 |
| ENSDART00000017230 | snrpc | 0.173923367 | 0.335809723 | 0.464722714 | 0.188235196 |
| ENSDART00000017259 | fgf13a | -0.403933655 | -0.684393894 | -0.638256176 | -0.336496098 |
| ENSDART00000017292 | stxbp5l | -0.151460711 | -0.273207638 | -0.026036945 | 0.086181343 |
| ENSDART00000017299 | tdg.1 | 0.102853523 | 0.194463285 | 0.348171907 | 0.196594799 |
| ENSDART00000017309 | ca16b | -0.395585068 | -0.492018229 | -0.260071918 | -0.046674115 |

| | | | | | |
|---|---|---|---|---|---|
| ENSDART00000017359 | sfpq | 0.205055054 | 0.247501694 | 0.177806588 | 0.262507487 |
| ENSDART00000017413 | zmynd10 | 1.112930172 | 1.692414467 | 1.985637202 | 1.515304432 |
| ENSDART00000017422 | tbc1d17 | 0.129347814 | 0.49736446 | 0.343125168 | -0.01149421 |
| ENSDART00000017424 | ptmaa | -0.012891551 | -0.178423729 | -0.469624362 | -0.475147609 |
| ENSDART00000017485 | sf3b6 | -0.19155437 | -0.273140548 | -0.376775786 | -0.432460535 |
| ENSDART00000017551 | slc6a1b | -0.387372781 | -0.441576322 | -0.34448646 | -0.23699282 |
| ENSDART00000017593 | tmem237a | 0.016738535 | -0.334661164 | -0.543525535 | -0.214177149 |
| ENSDART00000017599 | rem1 | -0.151492945 | -0.383680013 | -0.100323724 | -0.098680467 |
| ENSDART00000017619 | impdh1a | -0.178621661 | -0.394468991 | -0.26888728 | -0.032885741 |
| ENSDART00000017646 | atp6v0a1a | 1.074304865 | 0.491570366 | 0.543801777 | 0.26961956 |
| ENSDART00000017679 | ppp2r2ca | -0.437371106 | -0.275733569 | 0.22042039 | 0.361372826 |
| ENSDART00000017695 | foxd3 | 1.678381818 | 3.49818723 | 3.736655174 | 1.826274371 |
| ENSDART00000017763 | CABZ01071180.1 | -0.429470778 | -0.866508186 | -0.658991937 | -0.375481088 |
| ENSDART00000017774 | cacng5a | -0.685385238 | -0.470575642 | -0.348446742 | -0.159039188 |
| ENSDART00000017829 | HELZ2 (1 of many) | 0.30258536 | 0.160661865 | 1.299699291 | -0.234524229 |
| ENSDART00000018047 | zgc:112294 | -0.129085452 | -0.359907528 | -0.863079854 | -0.32905376 |
| ENSDART00000018054 | trh | -0.369432982 | -0.400024601 | -0.784012545 | -0.865751572 |
| ENSDART00000018117 | ppp1r14aa | -0.10674343 | -0.179038105 | -0.425069279 | -0.232436913 |
| ENSDART00000018150 | neurod6b | 0.484226112 | 1.695956927 | 1.935318885 | 1.555476315 |
| ENSDART00000018155 | adss | -0.479311511 | -0.372005861 | -0.294412527 | -0.197085254 |
| ENSDART00000018159 | FOXK2 (1 of many) | 0.025048904 | 0.121059845 | 0.321917677 | 0.256498575 |
| ENSDART00000018163 | irf2bp1 | -0.194489303 | -0.25009473 | -0.19907539 | -0.124082211 |
| ENSDART00000018228 | gsk3b | -0.075471253 | 0.063596815 | 0.240256058 | 0.249724511 |
| ENSDART00000018261 | akr1b1 | 1.00051996 | 0.934286328 | 0.670184322 | 0.161867768 |
| ENSDART00000018304 | mcm3 | 0.755786584 | 0.39595981 | 0.332438259 | 0.197655199 |
| ENSDART00000018347 | cab39l1 | -0.259625217 | -0.162823278 | -0.312896931 | -0.294075261 |
| ENSDART00000018351 | zgc:65851 | -1.669295979 | -1.346342078 | -0.126748353 | 0.319422369 |
| ENSDART00000018408 | anxa13l | 0.767695981 | 1.876785961 | 1.969202275 | 1.362090979 |
| ENSDART00000018461 | vmp1 | 0.430900752 | 0.323245721 | 0.184562524 | 0.049764543 |
| ENSDART00000018475 | snrpd3 | 0.421865611 | 0.381193487 | 0.380446926 | 0.20019783 |
| ENSDART00000018498 | helz2 | -0.063748361 | -0.192163226 | 1.151139617 | 0.013207133 |
| ENSDART00000018501 | opn4.1 | -0.494086134 | -0.180456491 | -0.369290026 | -0.100666897 |
| ENSDART00000018523 | ahcy | 0.45208938 | 0.606696476 | 0.316876579 | -0.072835379 |
| ENSDART00000018528 | | -0.854763044 | -1.028413156 | -0.142975543 | -0.344918955 |
| ENSDART00000018603 | tbx4 | -0.302508569 | -0.188111077 | -0.131947864 | 0.000377602 |
| ENSDART00000018625 | napab | 0.495528529 | 0.395594582 | 0.072464634 | -0.31666363 |
| ENSDART00000018654 | rnd1b | -0.711104866 | -0.668463449 | -0.311306501 | -0.458881777 |
| ENSDART00000018676 | cyp3c1 | 0.69067592 | 0.432985427 | -0.038151099 | -0.472952311 |
| ENSDART00000018685 | syt9a | 0.383050894 | 0.997731574 | 0.678563906 | 0.37205351 |
| ENSDART00000018686 | rrp15 | 0.358996202 | 0.261032346 | 0.197604486 | -0.018179208 |
| ENSDART00000018735 | dnaja2l | -0.20745831 | 0.012628814 | 0.350147509 | 0.311991137 |
| ENSDART00000018743 | phf20a | 0.312029532 | 0.419745206 | 0.325238046 | 0.318832686 |
| ENSDART00000018792 | spag7 | -0.393898515 | -0.535366384 | -0.519915196 | -0.305218132 |
| ENSDART00000018886 | ghra | -0.12746236 | -0.080311259 | -0.403757478 | -0.551820215 |
| ENSDART00000018945 | meis3 | -0.263611366 | 0.389783427 | 1.01804147 | 0.898863275 |
| ENSDART00000018972 | zgc:92818 | 0.012004445 | -0.040419062 | -0.784852086 | -0.38349701 |
| ENSDART00000019003 | psmd10 | 0.510870261 | 0.455922769 | 0.143258659 | 0.040691699 |
| ENSDART00000019029 | atp6v1h | -0.249435289 | -0.255917009 | -0.243059404 | -0.047539915 |
| ENSDART00000019045 | ebp | -0.186359255 | 0.947675675 | 1.363611199 | 0.55228711 |
| ENSDART00000019053 | faima | -0.147906168 | -0.161120397 | -0.359996788 | -0.164587924 |
| ENSDART00000019140 | rorab | -0.499015892 | -0.249117065 | -0.271786747 | 0.004902877 |

| | | | | | |
|---|---|---|---|---|---|
| ENSDART00000019149 | rpl7 | 0.429749031 | 0.41708455 | 0.235093103 | -0.015027688 |
| ENSDART00000019165 | apaf1 | 0.51528421 | 0.965000853 | 0.888709667 | 0.68935409 |
| ENSDART00000019199 | rab39ba | -0.125963686 | 0.3041991 | 0.44382958 | 0.206562916 |
| ENSDART00000019294 | si:dkeyp-75b4.9 | -0.789027106 | -0.841731007 | -0.221050872 | 0.359721563 |
| ENSDART00000019330 | ech1 | 0.344696401 | 0.425292318 | -0.031478205 | -0.016576624 |
| ENSDART00000019521 | dip2ba | 0.352814519 | 0.501689217 | 0.539560646 | 0.272762208 |
| ENSDART00000019573 | zgc:65894 | -0.648921378 | -0.373996254 | 0.357740851 | 0.502243837 |
| ENSDART00000019595 | | -2.247733873 | -0.556417246 | -3.574262962 | -0.309310232 |
| ENSDART00000019617 | rsad2 | -0.056061565 | -0.489318095 | 2.739834278 | -0.378754222 |
| ENSDART00000019647 | psmc2 | 0.088369453 | 0.252149794 | 0.191509771 | 0.022895546 |
| ENSDART00000019658 | nacad | -0.254606168 | 0.44010123 | 0.744298293 | 0.29398429 |
| ENSDART00000019698 | anxa5b | 1.162500386 | 1.67744918 | 1.46667303 | 0.432101083 |
| ENSDART00000019706 | phc2b | -0.309131033 | 0.04672236 | 0.268644424 | 0.076203283 |
| ENSDART00000019748 | lin7a | -0.334365216 | -0.254636287 | -0.400398823 | -0.216397551 |
| ENSDART00000019750 | wdr5 | 0.151689533 | 0.300718903 | 0.396766724 | 0.199844693 |
| ENSDART00000019766 | tgfb3 | -0.14127581 | -0.182290633 | -0.363811528 | -0.271524635 |
| ENSDART00000019770 | gpm6ba | -0.26452505 | -0.240754685 | -0.112562472 | 0.004364713 |
| ENSDART00000019818 | ric8b | 0.219425238 | 0.304273846 | 0.411142498 | 0.146708414 |
| ENSDART00000019905 | fndc4b | -0.870561937 | -0.746602431 | -0.973680622 | -0.402474596 |
| ENSDART00000019907 | unc119.1 | -0.258282731 | -0.126927399 | -0.208777989 | -0.147908262 |
| ENSDART00000019925 | GNB4 | 0.24039241 | 0.43881347 | 0.966289057 | 1.074168682 |
| ENSDART00000019936 | prkacab | -0.083728046 | -0.043118374 | 0.244894467 | 0.164419526 |
| ENSDART00000019937 | gadd45ga | 0.98351774 | 1.304995817 | 0.165178221 | 0.462881948 |
| ENSDART00000019949 | ndrg2 | -0.366275522 | -0.294111898 | -0.252300979 | -0.169375948 |
| ENSDART00000020017 | aldh3b1 | 0.019655473 | 0.632122683 | 0.647018435 | 0.316141972 |
| ENSDART00000020048 | gsna | 0.451913636 | 0.79824491 | 0.782008443 | 0.357949724 |
| ENSDART00000020054 | opcml | -0.412711216 | -0.540582472 | -0.347997034 | -0.057254512 |
| ENSDART00000020084 | hsp90ab1 | 0.266636397 | 0.434039186 | 0.537534154 | 0.384889667 |
| ENSDART00000020096 | fgf13b | -3.880462854 | -2.396384616 | -1.469971299 | 0.217849124 |
| ENSDART00000020122 | ywhah | -0.243048068 | -0.041853394 | 0.44002204 | 0.422069749 |
| ENSDART00000020153 | adck3 | -0.178069281 | -0.150101974 | -0.272732357 | -0.02457601 |
| ENSDART00000020167 | slc16a9a | 0.895506163 | 0.735954298 | 0.591907512 | -0.466816888 |
| ENSDART00000020168 | kctd5a | 0.075119561 | 0.1276866 | 0.301730152 | 0.250326391 |
| ENSDART00000020174 | dynll2b | -0.376289244 | -0.003912141 | -0.039237308 | 0.00732876 |
| ENSDART00000020183 | fam102bb | -0.149766416 | -0.311804442 | -0.169646826 | -0.06993891 |
| ENSDART00000020249 | dusp5 | -0.842223428 | -0.88819591 | -1.001072274 | -0.721604289 |
| ENSDART00000020252 | pdia6 | 0.374033372 | 0.009098395 | 0.091775115 | 0.023604955 |
| ENSDART00000020256 | lgsn | 1.361702081 | 2.686172063 | 1.394569806 | 0.349584846 |
| ENSDART00000020296 | nadl1.2 | -0.369828112 | -0.124790776 | 0.28925914 | 0.362042383 |
| ENSDART00000020311 | rpl27 | 0.430086554 | 0.381494789 | 0.182914397 | -0.034120883 |
| ENSDART00000020342 | sgsm3 | -0.132410481 | -0.068536613 | 0.176019735 | 0.283913195 |
| ENSDART00000020497 | snx13 | -0.143691651 | -0.273195754 | -0.089112423 | 0.016117988 |
| ENSDART00000020541 | lipf | 0.528006329 | 0.723139512 | 0.390796686 | -0.03933789 |
| ENSDART00000020569 | creld1b | -0.396401062 | -0.06913614 | 0.015512001 | 0.011512871 |
| ENSDART00000020621 | mapk4 | -0.137443479 | 0.345820304 | 0.800085717 | 0.474998646 |
| ENSDART00000020638 | rcan1a | 0.548382891 | 0.378475469 | 0.383543152 | 0.544764217 |
| ENSDART00000020646 | chrm1b | 0.963286522 | 0.623965705 | 0.291783612 | 0.576118226 |
| ENSDART00000020655 | psma5 | 0.424960543 | 0.53028483 | 0.342202642 | 0.090736505 |
| ENSDART00000020665 | sgtb | -0.247959729 | -0.080915193 | -0.275270124 | -0.210502305 |
| ENSDART00000020741 | aldoaa | -0.416493866 | -0.444493146 | -0.351916406 | -0.154405211 |
| ENSDART00000020810 | sdcbp2 | 0.429533724 | 0.242026939 | 0.181097661 | -0.126536592 |

| | | | | | |
|---|---|---|---|---|---|
| ENSDART00000020824 | pank1b | 0.041844369 | 0.140368278 | 0.263424196 | 0.005383312 |
| ENSDART00000020908 | zc4h2 | -0.133072632 | 0.229085423 | 0.337553441 | 0.264734015 |
| ENSDART00000020970 | pgm2 | 0.182379442 | 0.224260514 | 0.427220163 | 0.369397058 |
| ENSDART00000020999 | angptl1a | -0.447558553 | -0.389827093 | -0.516507166 | -1.101603289 |
| ENSDART00000021037 | hspa4a | 0.42074258 | 0.381012047 | 0.243421438 | 0.132816774 |
| ENSDART00000021062 | slc9a8 | 0.308335488 | -0.627436336 | -3.684031993 | -1.491743311 |
| ENSDART00000021069 | rpl38 | 0.561397297 | 0.414839383 | 0.25425723 | 0.049186317 |
| ENSDART00000021083 | calm2a | -0.300457039 | -0.171937069 | 0.044961812 | 0.17764614 |
| ENSDART00000021092 | snx27b | -0.196827884 | -0.228935762 | -0.254873921 | -0.135355059 |
| ENSDART00000021121 | stx5al | -0.079551416 | -0.239231712 | -0.554538789 | -0.564757113 |
| ENSDART00000021168 | rxrga | 0.040179684 | -0.194377683 | -0.381512289 | -0.377392107 |
| ENSDART00000021213 | cpne2 | -0.275625296 | -0.394159316 | -0.352439432 | -0.230978713 |
| ENSDART00000021231 | slmapb | -0.557559725 | -0.496297491 | -0.195575587 | 0.049672527 |
| ENSDART00000021260 | sept8b | -0.246039495 | -0.120094514 | -0.518439508 | -0.250786208 |
| ENSDART00000021299 | nmd3 | 0.353895501 | 0.176055483 | 0.10015969 | 0.031379837 |
| ENSDART00000021341 | kif3ca | 0.127548246 | 0.488322423 | 0.730937873 | 0.492487971 |
| ENSDART00000021346 | arl3l2 | -0.339022452 | -0.304754125 | -0.490852638 | -0.247921956 |
| ENSDART00000021417 | p2rx3a | 0.048222852 | -0.166951272 | -1.130394212 | -0.103958246 |
| ENSDART00000021491 | csnk1db | -0.098564588 | -0.355768317 | -0.198945325 | -0.067100917 |
| ENSDART00000021559 | coro1b | 0.006396016 | -0.122229179 | -0.278308819 | -0.121710131 |
| ENSDART00000021596 | rxrbb | -0.183103481 | -0.256930789 | -0.18551154 | -0.16195495 |
| ENSDART00000021604 | gins4 | 0.598069286 | 0.116671159 | -0.218567942 | -0.173827642 |
| ENSDART00000021605 | LRRC4C | -0.456880748 | -0.587108705 | -0.299621227 | -0.131466817 |
| ENSDART00000021609 | gad2 | -0.376538118 | -0.375877531 | -0.287892544 | -0.231277899 |
| ENSDART00000021666 | rtca | 0.450183742 | 0.82121323 | 0.721903733 | 0.20677388 |
| ENSDART00000021693 | ank2a | 0.088061869 | 0.283890991 | 0.488712819 | 0.41975176 |
| ENSDART00000021788 | pbk | 1.438031033 | 0.973567544 | 0.667760742 | 0.195406357 |
| ENSDART00000021798 | fabp11a | 3.578286828 | 1.692122399 | 1.073895347 | -0.266988673 |
| ENSDART00000021950 | mthfd1b | 1.089156306 | 0.599589593 | 0.18588711 | -0.096895618 |
| ENSDART00000021976 | nsa2 | 0.087749249 | 0.255476165 | 0.234966715 | 0.01893274 |
| ENSDART00000022010 | hivep2b | -0.404683257 | -0.428225913 | -0.085969125 | 0.061919909 |
| ENSDART00000022042 | scn8aa | -0.64882134 | -0.631929626 | 0.169338069 | 0.591046475 |
| ENSDART00000022044 | dct | 0.864997441 | 0.322109214 | -0.10561397 | -0.63566776 |
| ENSDART00000022051 | gins1 | -0.049060202 | -0.07395231 | -0.409999594 | -0.285702931 |
| ENSDART00000022060 | atf3 | 2.958540036 | 3.348918067 | 2.852908333 | 1.811501419 |
| ENSDART00000022270 | arhgap33 | 0.060975731 | 0.144058893 | 0.360327899 | 0.271763038 |
| ENSDART00000022290 | mdh1aa | -0.386096556 | -0.387245145 | -0.361974171 | -0.162628891 |
| ENSDART00000022307 | atic | 1.012801143 | 0.518350872 | 0.289570342 | -0.11925608 |
| ENSDART00000022356 | ppp1r7 | -0.289189989 | -0.228515822 | -0.279744824 | -0.100278848 |
| ENSDART00000022393 | si:dkeyp-57f11.2 | 0.163453067 | -0.122007101 | -0.459561011 | 0.032532197 |
| ENSDART00000022499 | psmb3 | 0.223119041 | 0.273754026 | 0.150624654 | -0.009410181 |
| ENSDART00000022533 | kcnj2a | -0.207812042 | -0.209993339 | -0.395014972 | -1.14439113 |
| ENSDART00000022549 | atp1b3a | -0.438514495 | -0.341784188 | -0.389598734 | -0.186751495 |
| ENSDART00000022562 | rhogb | 0.97771224 | 0.454579503 | 0.287206454 | -0.277565993 |
| ENSDART00000022579 | GABRG3 | -0.524427233 | -0.895243712 | -0.602147004 | -0.152848624 |
| ENSDART00000022581 | rab22a | -0.072513798 | -0.08434934 | -0.299709625 | -0.230388715 |
| ENSDART00000022586 | lrrc40 | 0.466457292 | 0.391920358 | 0.288821604 | -0.015697706 |
| ENSDART00000022625 | nrarpb | -0.295040721 | -0.303522559 | -0.284578601 | -0.228026779 |
| ENSDART00000022634 | acp2 | 0.356430058 | 0.295051752 | 0.030540215 | -0.026401831 |
| ENSDART00000022646 | cnot4b | 0.11528648 | 0.170971297 | 0.271245163 | 0.121803983 |
| ENSDART00000022660 | si:ch211-195b15.7 | -0.190059614 | -0.249688476 | -0.324813294 | -0.157998073 |

| | | | | | |
|---|---|---|---|---|---|
| ENSDART00000022663 | | -0.241007108 | -3.679203505 | -0.288618865 | -1.771039097 |
| ENSDART00000022688 | tob1b | -0.393522131 | -0.302261171 | -0.612204875 | -0.372656833 |
| ENSDART00000022694 | ehd3 | -0.278899876 | -0.241188601 | -0.097281852 | -0.017987416 |
| ENSDART00000022729 | unm_sa808 | -0.546569433 | -0.564171517 | -0.267551607 | 0.046437921 |
| ENSDART00000022765 | riok1 | 0.429965951 | 0.477495547 | 0.301061387 | 0.021181676 |
| ENSDART00000022768 | ak5 | -0.529999471 | -0.599207091 | -0.317508372 | 0.131326788 |
| ENSDART00000022866 | pisd | -0.555411298 | -0.214837183 | -0.086260422 | -0.186182517 |
| ENSDART00000022909 | klhl18 | -0.153412936 | 0.073357396 | 0.397749012 | 0.249762723 |
| ENSDART00000022963 | cdc14aa | -0.366168568 | -0.36782201 | -0.106140735 | 0.082726559 |
| ENSDART00000022976 | kctd16b | -0.982447595 | -1.281976718 | -0.566394426 | 0.110166911 |
| ENSDART00000022998 | ANO2 (1 of many) | 0.005158406 | -0.146342409 | -0.548931715 | -0.196350546 |
| ENSDART00000023038 | dacha | -0.511894205 | -0.418991667 | -0.004522839 | 0.112876213 |
| ENSDART00000023054 | atl3 | 0.317687991 | 0.347306029 | 0.121176286 | -0.186993768 |
| ENSDART00000023089 | acadvl | 0.315023187 | 0.311610409 | 0.17259027 | -0.0260318 |
| ENSDART00000023123 | nup88 | 0.221684741 | 0.406336851 | 0.406160624 | 0.250704697 |
| ENSDART00000023156 | eef1a1l2 | 0.512554224 | 0.741741761 | 0.709552788 | 0.28897731 |
| ENSDART00000023206 | plk2b | -0.591714767 | -0.133232909 | -0.517936071 | -0.57159861 |
| ENSDART00000023210 | trim13 | -0.37704188 | -0.363410115 | -0.204332616 | 0.075535016 |
| ENSDART00000023278 | fads2 | -0.280275959 | 0.609167416 | 1.126645574 | 0.470269511 |
| ENSDART00000023463 | uap1l1 | 0.706648736 | 0.512468414 | 0.225773419 | -0.015907861 |
| ENSDART00000023509 | ska2 | -0.036357965 | -0.092147723 | -0.294463498 | -0.176013793 |
| ENSDART00000023547 | anxa11b | 0.050301812 | -0.087626764 | -0.004152721 | -0.398004466 |
| ENSDART00000023550 | hsp90aa1.2 | -0.115909172 | -0.134363979 | -0.322775345 | -0.097362577 |
| ENSDART00000023562 | CABZ01041610.1 | -1.012987922 | -1.132303657 | -0.526387248 | -0.210938966 |
| ENSDART00000023588 | guca1a | 0.527717677 | -0.52699324 | -1.299472888 | -0.244316084 |
| ENSDART00000023613 | her6 | -0.335042933 | -0.234567638 | -0.250120727 | -0.59712361 |
| ENSDART00000023686 | ankrd33ab | -0.23872945 | -0.400254403 | -1.062972805 | -0.551724897 |
| ENSDART00000023709 | ptp4a2b | 0.271575512 | 0.562448581 | 0.634906429 | 0.086969331 |
| ENSDART00000023763 | wdcp | -0.038191062 | -0.371104976 | -0.391285873 | -0.034783387 |
| ENSDART00000023779 | vcp | 0.230561184 | 0.246570266 | 0.325389872 | 0.137878127 |
| ENSDART00000023806 | zgc:110319 | -0.195130618 | -0.122813174 | -0.337219496 | -0.098833156 |
| ENSDART00000023831 | cry5 | -0.290776024 | 0.507969364 | -0.405324234 | -0.228415974 |
| ENSDART00000023833 | eif2s3 | 0.172990624 | 0.349255612 | 0.311895175 | 0.045890851 |
| ENSDART00000023926 | eif3eb | -0.055777547 | -0.145564801 | -0.286794414 | -0.128155341 |
| ENSDART00000023944 | lmnl3 | -0.134674249 | 0.538751787 | 0.920098245 | 0.528719758 |
| ENSDART00000023959 | arntl1a | 0.340904931 | 1.030627189 | 1.45255612 | 0.975945139 |
| ENSDART00000024034 | gpsm2 | 0.06632826 | -0.389217225 | -0.522197566 | -0.062079955 |
| ENSDART00000024082 | psmb6 | 0.195043333 | 0.259353241 | 0.359595812 | 0.192042468 |
| ENSDART00000024135 | tubb2 | -0.097026879 | 1.085064176 | 1.27895467 | 1.027950073 |
| ENSDART00000024136 | gngt2a | 0.175894019 | 0.078473268 | -0.788482627 | -0.299650763 |
| ENSDART00000024194 | kif11 | 1.409119378 | 1.269759488 | 0.810392576 | 0.401186663 |
| ENSDART00000024208 | nutf2l | 0.951686549 | 0.855710394 | 0.537014601 | -0.220899011 |
| ENSDART00000024287 | zgc:165604 | -0.009089907 | -0.228105246 | -0.581619742 | -0.405549266 |
| ENSDART00000024296 | egln1b | -0.145407238 | -0.296649941 | -0.383343901 | -0.196291651 |
| ENSDART00000024304 | per3 | -0.478794008 | -0.23341732 | -0.352073188 | -0.024889246 |
| ENSDART00000024309 | rb1 | 0.736743325 | 0.968558289 | 0.720111001 | 0.238176467 |
| ENSDART00000024313 | rnf150b | -0.152451949 | -0.254343985 | -0.250476516 | -0.120482055 |
| ENSDART00000024316 | mcm5 | 1.292009744 | 0.913737369 | 0.774968175 | 0.275487022 |
| ENSDART00000024320 | ybx1 | 0.257630768 | 0.252218726 | 0.414084055 | 0.18414263 |
| ENSDART00000024328 | slc34a2a | -1.294382652 | -0.588426598 | -0.664087867 | -1.114341851 |
| ENSDART00000024331 | glsb | 0.029280722 | -0.254060441 | -0.66092512 | -0.487127171 |

| | | | | | |
|---|---|---|---|---|---|
| ENSDART00000024354 | csad | 0.826762252 | 0.889896753 | 0.709970739 | 0.637298928 |
| ENSDART00000024415 | epas1a | 0.419060345 | 0.571698023 | 0.827681076 | -0.059626191 |
| ENSDART00000024528 | emc7 | -0.102957427 | -0.142334415 | -0.248955976 | -0.189343286 |
| ENSDART00000024615 | rnpepl1 | 0.274953884 | 0.219791217 | 0.571525784 | 0.173823915 |
| ENSDART00000024619 | gorasp1a | -0.145027203 | -0.174354089 | -0.499227113 | -0.186036756 |
| ENSDART00000024662 | plppr3a | 1.050789931 | 1.738581828 | 1.609411361 | 1.082826483 |
| ENSDART00000024720 | si:ch211-282j22.3 | 0.265772408 | 0.220333711 | 0.187153701 | 0.074808124 |
| ENSDART00000024778 | robo3 | -0.327155095 | -0.251732012 | -0.124129181 | 0.058627004 |
| ENSDART00000024832 | stat5a | -0.212236095 | -0.408877789 | -0.375060924 | -0.185569558 |
| ENSDART00000024858 | chchd10 | -0.331794878 | -0.431239071 | -0.4916218 | -0.229989739 |
| ENSDART00000024872 | creb3l3l | 0.316205692 | 0.628018553 | 0.525420097 | 0.254183225 |
| ENSDART00000025031 | pou4f1 | -0.37458842 | -0.169318799 | 0.774588614 | 1.160187758 |
| ENSDART00000025046 | ppp1caa | 0.04938914 | -0.175273742 | -0.396680839 | -0.049267892 |
| ENSDART00000025096 | larp1b | 0.593217694 | 0.576647108 | 0.472990072 | 0.196836578 |
| ENSDART00000025198 | mettl21a | -0.380455489 | -0.566803653 | -0.570464602 | -0.564292758 |
| ENSDART00000025229 | adi1 | 0.582007509 | 0.274560712 | -0.200617197 | -0.21352575 |
| ENSDART00000025256 | igfbp2b | -0.742463234 | -0.107964006 | 0.616733017 | 0.74050304 |
| ENSDART00000025326 | csnk1da | -0.048665312 | -0.227600133 | -0.273972646 | -0.14458728 |
| ENSDART00000025385 | cers2a | 0.59368354 | 0.657003513 | 0.583941604 | 0.192392368 |
| ENSDART00000025414 | slc2a1a | 0.059681786 | -0.155207235 | -0.469818878 | -0.137194664 |
| ENSDART00000025466 | slc18a2 | -0.576881609 | -0.281211576 | -0.258781682 | -0.379077041 |
| ENSDART00000025487 | icmt | -0.205683018 | -0.185657781 | -0.262148423 | -0.151909752 |
| ENSDART00000025494 | hprt1l | -0.364550689 | -0.385937863 | -0.083973479 | -0.129534044 |
| ENSDART00000025496 | rras | 0.483455604 | 0.817924325 | 0.502791397 | -0.193021858 |
| ENSDART00000025501 | snap23.1 | 0.353906863 | 0.265231797 | -0.028292671 | -0.317654401 |
| ENSDART00000025535 | sept5a | 0.521581645 | 1.371610673 | 0.945407012 | 0.566204111 |
| ENSDART00000025550 | top1mt | 0.816108961 | 1.133963099 | 0.883225094 | 0.407533529 |
| ENSDART00000025573 | CABZ01071723.1 | -0.48632791 | -0.607491421 | -0.117657015 | 0.102596694 |
| ENSDART00000025583 | fgf8a | -0.058331217 | -0.819626176 | 0.05604093 | 0.237151994 |
| ENSDART00000025620 | ppiaa | 0.369681961 | 0.563046067 | 0.555154111 | 0.174901331 |
| ENSDART00000025698 | zgc:153311 | -0.134647599 | -0.156342149 | -0.388760407 | -1.773822225 |
| ENSDART00000025782 | nup93 | 0.17632244 | 0.241668195 | 0.293793434 | 0.152368608 |
| ENSDART00000025852 | tnni2b.1 | -0.577234038 | 4.833234821 | 5.215670785 | 3.899605393 |
| ENSDART00000025860 | si:dkey-247m21.3 | 0.550902269 | 0.531374836 | -0.221903085 | -0.681264404 |
| ENSDART00000025877 | cldn12 | -0.148592994 | -0.140436414 | -0.371702065 | -0.263635262 |
| ENSDART00000025912 | si:dkey-32n7.4 | 0.676729111 | 0.251567535 | 0.290364244 | 0.056388056 |
| ENSDART00000025962 | gyg1a | -0.132586486 | -0.231554909 | -0.320101499 | -0.261622046 |
| ENSDART00000025997 | dip2cb | -0.352423277 | -0.192868324 | 0.083423296 | 0.161945092 |
| ENSDART00000026085 | ptges | -0.133597623 | -0.186036004 | -0.449634161 | -0.455296325 |
| ENSDART00000026145 | AMOTL1 | 1.0586494 | 0.570711762 | 0.741896115 | 0.335943589 |
| ENSDART00000026152 | asap2a | -0.416411014 | -0.084267369 | -0.025927141 | 0.001189582 |
| ENSDART00000026174 | dgkh | -0.6307495 | -1.007774241 | -0.588261808 | -0.107017931 |
| ENSDART00000026178 | kif4 | 0.630775369 | 0.400037801 | 0.195646982 | 0.069714171 |
| ENSDART00000026180 | fabp7a | 0.745585804 | 2.003621538 | 1.857780253 | 1.020257385 |
| ENSDART00000026303 | rasd1 | -0.723056439 | -0.693453912 | -0.473472236 | -0.755311511 |
| ENSDART00000026316 | sema3gb | -0.148485986 | -0.183297493 | -0.3621349 | -0.465943338 |
| ENSDART00000026339 | gtpbp4 | 0.113641949 | 0.327640711 | 0.222548232 | 0.071359586 |
| ENSDART00000026378 | slc25a6 | -0.475274891 | -0.471356749 | -0.229323143 | 0.038491163 |
| ENSDART00000026401 | TMEM178B (1 of many) | -0.387053719 | -0.214500403 | -0.049460199 | 0.011667684 |
| ENSDART00000026409 | cct4 | 0.459151422 | 0.538338734 | 0.621039126 | 0.313903974 |
| ENSDART00000026492 | flncb | 1.767414623 | 1.156524725 | 0.931280558 | 0.481368808 |

| | | | | | |
|---|---|---|---|---|---|
| ENSDART00000026692 | ubtd1a | -0.10869502 | -0.342281946 | -0.985605367 | -0.704345293 |
| ENSDART00000026765 | slc18a3a | -0.342520426 | -0.454539851 | -0.359489571 | -0.310297658 |
| ENSDART00000026766 | aldocb | -0.483629021 | -0.540701781 | -0.406696036 | -0.135568984 |
| ENSDART00000026800 | kifap3b | 0.203180282 | 0.588115089 | 0.640508934 | 0.450843691 |
| ENSDART00000026814 | ptp4a1 | 0.419787699 | 0.591512581 | 0.636748721 | 0.300535035 |
| ENSDART00000026865 | l3mbtl1a | -0.339631378 | -0.338161449 | -0.153879513 | -0.055902641 |
| ENSDART00000026924 | dnah7 | -0.347974455 | -0.178245432 | -0.682598609 | -0.174885691 |
| ENSDART00000026992 | sox4a | 0.168735758 | 0.553498244 | 0.871364123 | 0.630648218 |
| ENSDART00000027050 | cnga3b | -0.139313288 | -0.277577971 | -1.175814494 | -0.697938349 |
| ENSDART00000027115 | nob1 | -0.149963351 | -0.185404709 | -0.224057031 | -0.30485959 |
| ENSDART00000027158 | psmd3 | 0.079773681 | 0.202742016 | 0.46801517 | 0.212038089 |
| ENSDART00000027274 | efna3a | -0.422282495 | -0.289576322 | -0.118198025 | 0.092530837 |
| ENSDART00000027345 | tmem59l | -0.559867163 | -0.384910477 | -0.005343137 | 0.202377241 |
| ENSDART00000027379 | bicral | 0.180958951 | 0.201052129 | 0.429766407 | 0.252075745 |
| ENSDART00000027393 | ckmt1 | 0.722688207 | 0.598583347 | 0.435615456 | 0.204934783 |
| ENSDART00000027398 | kcna2a | -1.558634612 | -1.413069345 | -0.244650937 | 0.296687437 |
| ENSDART00000027417 | zgc:171704 | 0.227240571 | -0.665719796 | -1.161293822 | -1.438167938 |
| ENSDART00000027454 | si:ch211-207i1.2 | -0.470697042 | -0.923472632 | -1.21436549 | -0.514660002 |
| ENSDART00000027463 | hmx4 | -0.340009377 | -0.236809718 | -0.106768059 | 0.00156649 |
| ENSDART00000027465 | cacna2d4b | -0.370016882 | -0.141565087 | -0.247577188 | 0.068079685 |
| ENSDART00000027466 | fam63b | -0.160774327 | 0.127626681 | 0.377768257 | 0.305765252 |
| ENSDART00000027532 | mapkapk2a | -0.086338753 | -0.192582349 | -0.252851985 | -0.027631364 |
| ENSDART00000027598 | tpm3 | 0.692886998 | 2.316316343 | 2.715886041 | 1.570432549 |
| ENSDART00000027616 | eif4g2a | 0.333990776 | 0.435258045 | 0.644700731 | 0.350291836 |
| ENSDART00000027689 | amph | -0.260662658 | -0.30969295 | -0.237665092 | 0.01178982 |
| ENSDART00000027718 | fxr2 | 0.070260526 | 0.251047986 | 0.370057736 | 0.114198992 |
| ENSDART00000027758 | rtn1b | -0.626628578 | 0.207141964 | 0.617559397 | 0.433726227 |
| ENSDART00000027957 | hmgcl | -0.03513095 | -0.160364915 | -0.261992968 | -0.251204537 |
| ENSDART00000028003 | ankrd22 | 0.462762131 | 0.494960479 | 0.738379791 | 0.088098985 |
| ENSDART00000028033 | emc8 | -0.03555204 | -0.206564635 | -0.268269465 | -0.211006035 |
| ENSDART00000028048 | necap1 | -0.117195999 | -0.341344035 | -0.478671947 | -0.252467934 |
| ENSDART00000028090 | eif2ak1 | 0.218229206 | 0.400243646 | 0.398265404 | 0.179590234 |
| ENSDART00000028108 | ddc | -0.192852254 | -0.317464474 | -0.303298772 | 0.040886808 |
| ENSDART00000028219 | pvalb4 | -1.549055704 | 2.87360786 | 4.516864647 | 3.395003162 |
| ENSDART00000028225 | mao | -0.257909806 | -0.164169194 | -0.255680521 | -0.167259868 |
| ENSDART00000028285 | pgbd5 | -0.471148265 | -0.169774685 | 0.263561856 | 0.22277577 |
| ENSDART00000028338 | scamp5a | -0.358452184 | -0.354788931 | -0.092970627 | 0.058203665 |
| ENSDART00000028390 | fgf12a | -0.792701948 | -0.713373555 | -0.153423419 | 0.185286633 |
| ENSDART00000028417 | lrit2 | -0.281592969 | -0.383251603 | -0.165960328 | 0.039848719 |
| ENSDART00000028500 | nxn | -0.356492001 | -0.304319321 | -0.337794036 | -0.002781587 |
| ENSDART00000028607 | chd6 | 0.00488027 | -0.079639497 | 0.533277243 | 0.378096362 |
| ENSDART00000028787 | ahr1b | -0.27962376 | -0.18218298 | -0.129040126 | 0.07690852 |
| ENSDART00000028883 | gna13b | 0.442397382 | 0.38652983 | 0.274274635 | -0.034947864 |
| ENSDART00000028895 | negr1 | -0.362135851 | -0.256191375 | -0.15121136 | -0.084875523 |
| ENSDART00000028946 | tpd52l2a | 0.038391394 | -0.134825728 | -0.275007322 | -0.131471646 |
| ENSDART00000028960 | ndufa2 | -0.147139307 | -0.190704061 | -0.363806149 | -0.139491855 |
| ENSDART00000028997 | myo9ab | 0.179745305 | 0.101321956 | 0.280102899 | 0.331620625 |
| ENSDART00000029121 | usp5 | 0.186533139 | 0.264124698 | 0.696851044 | 0.548402162 |
| ENSDART00000029133 | snu13b | 0.760760317 | 0.56984098 | 0.429270049 | -0.029914173 |
| ENSDART00000029380 | bnip4 | 0.50246171 | 0.517948631 | -0.062532971 | -0.337429982 |
| ENSDART00000029387 | ppan | 0.365110957 | 0.289310564 | 0.234072786 | 0.008675203 |

| | | | | | |
|---|---|---|---|---|---|
| ENSDART00000029457 | sh2d3ca | -0.605491696 | -0.301886794 | 0.051143961 | 0.055429229 |
| ENSDART00000029459 | gipr | -0.120368109 | -0.334116608 | -0.559985722 | -0.423861096 |
| ENSDART00000029492 | cmtm7 | 0.705299486 | 0.845221281 | 0.768474344 | -0.28587683 |
| ENSDART00000029528 | mospd2 | 0.309594516 | 0.158328804 | 0.007644114 | -0.042795417 |
| ENSDART00000029646 | rplp1 | 0.363694976 | 0.319088557 | 0.167660064 | 0.013197371 |
| ENSDART00000029703 | kcnh1a | -0.531746722 | -1.366001544 | -0.565478925 | 0.09193536 |
| ENSDART00000029774 | tmem55bb | -0.03621571 | -0.196356812 | -0.337053328 | -0.094153869 |
| ENSDART00000029843 | vezf1a | 0.660507751 | 0.776194217 | 0.884683465 | 0.428803142 |
| ENSDART00000029946 | ube2b | -0.262228145 | -0.117966704 | -0.377845713 | -0.176574464 |
| ENSDART00000029981 | ppp3cb | -0.453839737 | -0.543662557 | -0.385030051 | 0.007551117 |
| ENSDART00000030125 | znhit3 | 0.760210037 | 0.691966223 | 0.487369205 | 0.388320697 |
| ENSDART00000030205 | bnip3lb | 0.087436849 | 0.295010863 | 0.335325724 | 0.120618456 |
| ENSDART00000030211 | gmfb | -0.040702268 | 0.286017295 | 0.157496122 | 0.031502906 |
| ENSDART00000030213 | mapk1 | 0.367982033 | 0.335025068 | 0.254642258 | -0.112923758 |
| ENSDART00000030409 | asap1b | -0.387126717 | -0.32480613 | -0.179905202 | -0.016544261 |
| ENSDART00000030509 | glra4a | -0.54562871 | -0.440353116 | -0.570533638 | -0.182903567 |
| ENSDART00000030579 | crhbp | 0.109353477 | -0.053663997 | -0.382440888 | -0.742296052 |
| ENSDART00000030691 | clic4 | 0.290171682 | 0.490608957 | 0.279069846 | 0.046035564 |
| ENSDART00000030773 | foxo3a | -0.522674186 | -0.346284342 | -0.025883071 | 0.086783781 |
| ENSDART00000030794 | tmem169a | 0.073168182 | 0.288108809 | 0.520379557 | 0.751344779 |
| ENSDART00000030811 | cables2b | -0.198257533 | -0.480718843 | -0.449759807 | -0.060172476 |
| ENSDART00000030885 | uckl1a | 0.183203205 | -0.212616494 | -0.705475026 | -0.013040484 |
| ENSDART00000030887 | slc45a2 | 0.835000405 | 0.559834035 | -0.13177777 | -0.758141187 |
| ENSDART00000030890 | hmox1a | 2.623035746 | 1.043671484 | 1.096486903 | 0.061370069 |
| ENSDART00000030920 | gid8a | -0.116648875 | -0.277182294 | -0.355798783 | -0.233597587 |
| ENSDART00000030995 | umps | 0.416416432 | 0.256033809 | 0.30736318 | -0.006312473 |
| ENSDART00000031047 | cd63 | 1.025048213 | 0.521327717 | 0.067995805 | -0.605090068 |
| ENSDART00000031091 | vsnl1a | -0.845008079 | -0.77079299 | -0.390375819 | 0.072718281 |
| ENSDART00000031121 | VDAC3 (1 of many) | -0.281245453 | -0.237325849 | -0.158924417 | 0.113945733 |
| ENSDART00000031139 | slc24a4b | -0.756345109 | -1.188291659 | -0.474432561 | -0.06484409 |
| ENSDART00000031165 | enoph1 | 0.127295774 | 0.248943534 | 0.342353563 | 0.2531466 |
| ENSDART00000031167 | tfap2d | -0.967562253 | -0.629795841 | 0.638492839 | 1.047009841 |
| ENSDART00000031234 | stxbp2 | 0.406820773 | 0.260992395 | 0.143612276 | -0.234256086 |
| ENSDART00000031265 | rtn4r | -0.753039937 | -0.836146377 | -0.430012914 | -0.160533624 |
| ENSDART00000031390 | caskin1 | -0.735732711 | -0.907721451 | -0.315045095 | -0.088633322 |
| ENSDART00000031425 | zgc:55582 | -0.13184915 | -0.205452993 | -0.25748443 | -0.094044364 |
| ENSDART00000031426 | skilb | -0.234186907 | -0.553407151 | -0.565936684 | -0.420869627 |
| ENSDART00000031470 | pafah1b1b | -0.06625793 | 0.19164974 | 0.422888398 | 0.388967525 |
| ENSDART00000031498 | ccna2 | 1.569664461 | 1.378683751 | 0.895010724 | 0.581055828 |
| ENSDART00000031546 | chrna6 | -0.487671995 | -0.420962215 | 0.210734418 | 0.46609085 |
| ENSDART00000031638 | slc48a1a | -0.121591299 | -0.156663294 | -0.340136476 | -0.182759405 |
| ENSDART00000031650 | hsp70l | -0.846526969 | -1.057845455 | -1.357523269 | -1.215078404 |
| ENSDART00000031727 | vamp8 | 0.086516161 | -0.020853386 | -0.34297459 | -0.531441391 |
| ENSDART00000031937 | diras1a | -0.474540918 | -0.418703997 | -0.179232074 | -0.012563291 |
| ENSDART00000032161 | galnt14 | -0.529482639 | -0.362873656 | -0.264119496 | -0.01842007 |
| ENSDART00000032212 | fynrk | -0.162448847 | -0.18539887 | -0.276020606 | -0.108651002 |
| ENSDART00000032275 | atp6v1c1a | -0.172326217 | -0.24874129 | -0.293288363 | -0.174129603 |
| ENSDART00000032290 | esyt1a | 0.414133861 | 0.186544426 | 0.071532791 | -0.03493067 |
| ENSDART00000032322 | abcg2c | 0.455571157 | 0.699504596 | 0.641510999 | 0.335341634 |
| ENSDART00000032324 | hddc3 | 0.491004501 | 0.803369277 | 0.623276564 | 0.474326204 |
| ENSDART00000032331 | gmppab | 0.298836739 | 0.452114759 | 0.362408461 | -0.028707454 |

| | | | | | |
|---|---|---|---|---|---|
| ENSDART00000032392 | dhdhl | 0.557297889 | 0.381488116 | 0.091728361 | -0.116661732 |
| ENSDART00000032393 | ginm1 | -0.054454012 | -0.092963972 | -0.327748186 | -0.224389181 |
| ENSDART00000032459 | aqp1a.1 | -0.275584647 | -0.402261221 | -0.531838868 | -0.410567004 |
| ENSDART00000032498 | tspan36 | 1.455763612 | 0.805559086 | 0.192569945 | -0.715366149 |
| ENSDART00000032502 | nebl | -0.405575453 | -0.348273521 | -0.638746976 | -0.69843843 |
| ENSDART00000032540 | usp14 | 0.236991779 | 0.380670741 | 0.291281165 | 0.061189525 |
| ENSDART00000032547 | lect2l | 1.708944579 | 1.070031122 | 0.348802096 | -0.787093126 |
| ENSDART00000032603 | tspo | 0.47255795 | 0.736846252 | 0.52533392 | 0.11686825 |
| ENSDART00000032695 | asic4a | -0.405928595 | -0.689262684 | -0.527033258 | -0.378352566 |
| ENSDART00000032821 | cyth1b | 0.161176597 | 0.057433607 | -0.079026086 | -0.299717291 |
| ENSDART00000032844 | plekha6 | -0.171611787 | -0.149520127 | -0.254818235 | 0.001528605 |
| ENSDART00000032857 | mapk11 | 0.353170314 | 0.603154564 | 0.6388701 | 0.458511735 |
| ENSDART00000032899 | clpxa | -0.125001004 | -0.138480195 | -0.243789891 | -0.138582094 |
| ENSDART00000032963 | apoob | -0.330795 | -0.271281795 | -0.230754946 | 0.035824011 |
| ENSDART00000033053 | dennd5b | -0.288493162 | -0.362789161 | -0.132763243 | 0.070545146 |
| ENSDART00000033248 | fam107b | -0.588148397 | -0.439287754 | -0.362004691 | -0.311865235 |
| ENSDART00000033316 | vangl2 | 0.207091244 | 0.447705982 | 0.579933447 | 0.26125294 |
| ENSDART00000033325 | slc25a24 | -0.429859875 | -0.309149544 | -0.52291133 | -0.286046915 |
| ENSDART00000033361 | ttyh3b | -0.3186879 | -0.037617104 | 0.269211477 | 0.394612295 |
| ENSDART00000033362 | gatad2b | -0.40244942 | -0.286045422 | -0.214483951 | -0.084688348 |
| ENSDART00000033386 | ocstamp | 2.954317372 | 2.548446344 | 2.458244526 | 1.328553982 |
| ENSDART00000033479 | si:ch211-129c21.1 | 1.539581327 | 2.000726167 | 1.978342783 | 1.084446492 |
| ENSDART00000033494 | klf6a | 1.084550888 | 1.797692613 | 1.625562666 | 0.460444617 |
| ENSDART00000033545 | | 0.299034458 | 0.290865093 | 0.474990251 | 0.248453432 |
| ENSDART00000033566 | smad1 | 0.486585209 | 1.249927606 | 1.264727704 | 0.536891999 |
| ENSDART00000033574 | slc24a5 | 0.460212086 | 0.054768672 | -0.306746704 | -1.141417252 |
| ENSDART00000033657 | grm6b | -0.379593806 | -0.511743403 | -0.159835887 | 0.100793439 |
| ENSDART00000033663 | rps21 | 0.311890725 | 0.366197651 | 0.105076559 | -0.038309008 |
| ENSDART00000033713 | arpc1b | 0.908662023 | 0.412162066 | 0.131006833 | -0.30826968 |
| ENSDART00000033724 | fabp3 | 0.043357927 | 0.829727862 | 0.980692721 | 0.446444515 |
| ENSDART00000033746 | gins2 | 1.156804675 | 0.565959042 | -0.013280867 | 0.106301442 |
| ENSDART00000033761 | glb1 | 0.347409105 | 0.187375641 | 0.031649312 | 0.073918812 |
| ENSDART00000033980 | lims1 | 0.497849916 | 0.369061975 | 0.247283344 | -0.174205881 |
| ENSDART00000034004 | faf1 | 0.052082526 | 0.093691656 | 0.281166911 | 0.24525998 |
| ENSDART00000034216 | dync1h1 | 0.15631552 | 0.099035755 | 0.755817679 | 0.554910929 |
| ENSDART00000034248 | rab32a | 0.364170748 | 0.270180665 | -0.161567701 | -0.659765413 |
| ENSDART00000034313 | gas2a | -0.351038034 | -0.291680692 | -0.523203366 | -0.413177021 |
| ENSDART00000034377 | cpa5 | 2.043914839 | 0.12107472 | 0.19928898 | 0.161423339 |
| ENSDART00000034421 | cdk14 | -0.37810724 | -0.738234108 | -0.33600573 | -0.064327168 |
| ENSDART00000034432 | susd4 | -0.32348116 | 0.086685706 | 0.47906988 | 0.317031782 |
| ENSDART00000034441 | tcp11l2 | 0.020775808 | -0.208761169 | -0.796222831 | -0.248557177 |
| ENSDART00000034523 | tars | 0.184292328 | 0.408838041 | 0.43347383 | 0.219690833 |
| ENSDART00000034638 | ccdc28a | -0.059055282 | -0.067802136 | -0.368600818 | -0.274995246 |
| ENSDART00000034705 | ntmt1 | -0.020729663 | -0.187405157 | -0.44452011 | -0.329716724 |
| ENSDART00000034737 | cpne8 | -0.307817154 | -0.810818352 | -0.660194536 | 0.005050706 |
| ENSDART00000034784 | adcyap1b | 1.680397515 | 2.015053329 | 1.852069794 | 1.09666998 |
| ENSDART00000034790 | pcp4l1 | -0.408039344 | -0.390896541 | -0.213514281 | -0.035791454 |
| ENSDART00000034829 | rrp12 | 0.494057186 | 0.423927766 | 0.372142811 | 0.099318986 |
| ENSDART00000034834 | ppfia2 | -0.326502158 | -0.151128851 | -0.013552631 | 0.061359529 |
| ENSDART00000034849 | grin1b | -0.325252553 | -0.609188282 | -0.185270574 | 0.122354295 |
| ENSDART00000034850 | dbi | 1.011739121 | 0.830016329 | 0.708552366 | 0.209278197 |

| | | | | | |
|---|---|---|---|---|---|
| ENSDART00000034883 | mcf2a | -0.149015362 | -0.337699782 | -0.173487291 | 0.057006752 |
| ENSDART00000034914 | pvalb3 | -0.375270417 | 3.922610155 | 4.415489459 | 3.238850832 |
| ENSDART00000034935 | desi2 | 0.073850834 | 0.377607475 | 0.427284588 | 0.193399467 |
| ENSDART00000035031 | sgk1 | 0.036456511 | -0.228295468 | -0.323414575 | -0.393803665 |
| ENSDART00000035067 | abhd2a | -0.280282686 | -0.122469223 | -0.369410687 | -0.218167862 |
| ENSDART00000035093 | col9a2 | 0.360349924 | 0.265960604 | -0.052354611 | -0.707154609 |
| ENSDART00000035150 | spast | 0.060265858 | 0.143278561 | 0.351254395 | 0.195874089 |
| ENSDART00000035152 | kif26ab | 0.198116066 | 0.127768449 | 0.42622306 | 0.434736458 |
| ENSDART00000035239 | nek1 | -0.350009624 | -0.188792207 | -0.078432875 | 0.138948692 |
| ENSDART00000035245 | spire2 | 0.223570717 | 0.667119992 | 0.874236758 | 0.626613708 |
| ENSDART00000035409 | zc2hc1a | 0.100393521 | 0.424633369 | 0.532716266 | 0.492034687 |
| ENSDART00000035447 | mtmr9 | -0.272610717 | -0.235611081 | 0.139540941 | 0.187819723 |
| ENSDART00000035538 | ptp4a3 | -0.088675269 | 0.257809374 | 0.053091443 | -0.336323707 |
| ENSDART00000035628 | srsf10a | -0.231452965 | -0.136268467 | -0.25871265 | -0.135010634 |
| ENSDART00000035670 | polr2eb | 0.35113115 | 0.294169174 | 0.063963778 | -0.045691283 |
| ENSDART00000035676 | bnip3la | -0.292353372 | -0.099807874 | -0.059866968 | -0.067415369 |
| ENSDART00000035710 | lin7c | -0.082526521 | -0.252501014 | -0.262742844 | -0.186356266 |
| ENSDART00000035737 | slc11a2 | 0.286882779 | 0.187123756 | 0.119307344 | -0.030269778 |
| ENSDART00000035739 | tmem134 | 0.246242014 | 0.369929781 | 0.082611565 | -0.161334554 |
| ENSDART00000035899 | pkp2 | 0.428534184 | 1.016588803 | 1.016153499 | 0.312913104 |
| ENSDART00000035907 | sec24d | 0.279449898 | 0.329558363 | 0.384805326 | -0.077598974 |
| ENSDART00000035944 | clic5a | 2.554130845 | 1.311062136 | 1.150965478 | 0.646568675 |
| ENSDART00000036015 | ryr1b | -0.116788998 | -0.165312146 | 0.372637833 | 0.693396813 |
| ENSDART00000036050 | rs1a | -0.391328986 | -0.392177441 | -0.469736828 | -0.271292581 |
| ENSDART00000036153 | ccdc3a | 0.286306507 | 0.061698184 | -0.1803174 | -0.442994145 |
| ENSDART00000036240 | cers4b | -0.360584189 | -0.184519104 | -0.081616234 | -0.015625515 |
| ENSDART00000036373 | cfap206 | 0.026049802 | 0.014543106 | -0.421468514 | -0.063922477 |
| ENSDART00000036421 | chek2 | 0.623199787 | 0.358982062 | 0.455251589 | 0.098448549 |
| ENSDART00000036472 | zgc:110852 | -0.482143709 | -0.509026678 | -0.613205563 | -0.094495965 |
| ENSDART00000036513 | trib3 | 0.704680301 | 0.933969628 | 0.780405513 | 0.230790744 |
| ENSDART00000036531 | gnai1 | 0.25941591 | 0.265271449 | 0.201009078 | -0.026939111 |
| ENSDART00000036581 | cdk2 | 1.227545183 | 1.116666759 | 0.903071292 | 0.284538183 |
| ENSDART00000036649 | sfxn2 | 0.32661292 | 0.494950572 | 0.442482013 | 0.063629138 |
| ENSDART00000036668 | psmc1a | 0.311771761 | 0.423109618 | 0.307731242 | 0.107467578 |
| ENSDART00000036680 | ptgr1 | 0.476212613 | 0.619935312 | 0.420689871 | 0.024158985 |
| ENSDART00000036703 | pfdn2 | 0.369409429 | 0.460943577 | 0.431067405 | 0.197162016 |
| ENSDART00000036718 | eif4e1c | 0.366528838 | 0.391471958 | 0.37708707 | 0.13997088 |
| ENSDART00000036729 | spi1b | 0.910663058 | 0.717004711 | 0.095052493 | -0.117318513 |
| ENSDART00000036760 | tppp2 | -0.225112914 | 0.608288105 | 0.560408419 | 0.667346525 |
| ENSDART00000036797 | uchl1 | 1.004812771 | 1.916965558 | 1.938952257 | 1.446591845 |
| ENSDART00000036854 | glcci1 | -0.216167779 | -0.276312685 | -0.306037916 | -0.0692944 |
| ENSDART00000036891 | rabac1 | 0.184715811 | 0.277575159 | 0.08241236 | -0.087331685 |
| ENSDART00000036926 | vangl1 | -0.329590873 | -0.033233324 | 0.383553313 | 0.3451937 |
| ENSDART00000036939 | gadd45ba | 1.15110753 | 1.253886794 | 1.06146692 | 0.82843487 |
| ENSDART00000036997 | camk2n1a | -0.377223565 | -0.40737699 | -0.242898936 | -0.23919335 |
| ENSDART00000037007 | tpi1a | -0.595344218 | -0.412573356 | -0.319655345 | -0.136192131 |
| ENSDART00000037036 | cnppd1 | -0.251371332 | -0.185078308 | -0.27690601 | -0.219995028 |
| ENSDART00000037065 | sccpdhb | -0.135500672 | -0.263394229 | -0.35527789 | -0.147169147 |
| ENSDART00000037109 | srpk1a | -0.158297297 | -0.128401953 | -0.285156041 | -0.13534624 |
| ENSDART00000037126 | eno2 | -0.315372643 | -0.196767149 | 0.300750061 | 0.433471501 |
| ENSDART00000037195 | kif26bb | 1.206892082 | 1.55094827 | 1.874551064 | 1.519778039 |

| | | | | | |
|---|---|---|---|---|---|
| ENSDART00000037224 | cst14a.2 | 0.549538616 | 0.553010386 | 0.34149596 | -0.057482802 |
| ENSDART00000037265 | olfm1b | -0.595000689 | -1.108211527 | -0.18099665 | 0.18250463 |
| ENSDART00000037371 | ppp1r13ba | -0.171240808 | -0.158646598 | -0.283609683 | -0.120899195 |
| ENSDART00000037698 | uck2b | 0.641974401 | 1.119646558 | 1.077673009 | 0.598617402 |
| ENSDART00000037709 | nol11 | 0.254424392 | 0.434300513 | 0.299474854 | 0.102573384 |
| ENSDART00000037846 | focad | 0.435964232 | 0.512370823 | 0.456134064 | 0.112317494 |
| ENSDART00000037848 | dpp6b | -0.469010511 | -0.191315433 | -0.026922326 | -0.081540125 |
| ENSDART00000037850 | dync1li2 | 0.268558729 | 0.496891146 | 0.585345447 | 0.309833954 |
| ENSDART00000037879 | crx | -0.254014844 | -0.310425293 | -0.22793154 | -0.020194538 |
| ENSDART00000037922 | slc6a8 | -0.298142259 | -0.526146924 | -0.491663812 | -0.167137822 |
| ENSDART00000038202 | cndp2 | 1.63370988 | 0.725493655 | 0.197899383 | -0.10985912 |
| ENSDART00000038290 | crhb | -0.533237668 | -0.584543834 | -0.601107267 | -0.51443145 |
| ENSDART00000038294 | tp53inp1 | -0.610161281 | 0.029061275 | -0.033185195 | -0.028504863 |
| ENSDART00000038301 | gnpda2 | 0.314221876 | 0.240267665 | 0.128522583 | -0.033807169 |
| ENSDART00000038310 | ormdl3 | -0.300841314 | -0.124106092 | -0.535837175 | -0.413488994 |
| ENSDART00000038330 | khsrp | -0.050330793 | 0.071549867 | 0.273984704 | 0.161173774 |
| ENSDART00000038391 | pkz | 0.411244814 | 0.129537507 | 1.595328762 | 0.021626224 |
| ENSDART00000038495 | ctnnb1 | 0.20653709 | 0.248644759 | 0.214149858 | 0.090949105 |
| ENSDART00000038505 | rprmb | -0.046727426 | -0.153123084 | -0.26580673 | 0.05838964 |
| ENSDART00000038648 | ptbp2b | 0.207623817 | 0.150446749 | 0.253639385 | 0.163738179 |
| ENSDART00000038674 | tmem230a | -0.312877262 | -0.360088684 | -0.444507782 | -0.485519275 |
| ENSDART00000038696 | flvcr2b | 1.475160998 | 0.41510575 | 0.393658072 | -0.063620422 |
| ENSDART00000038740 | galnt9 | -0.705450847 | -0.542663541 | -0.098391024 | 0.177029625 |
| ENSDART00000038888 | hsdl2 | 0.331360296 | 0.293802861 | 0.248668235 | 0.085382052 |
| ENSDART00000038924 | sult1st1 | 0.0423301 | -0.129133641 | -0.497646461 | -0.249352869 |
| ENSDART00000038990 | jak1 | 0.414591598 | 0.220201928 | 0.070657217 | -0.102548961 |
| ENSDART00000039043 | rgs7bpb | -0.383267221 | -0.448718374 | -0.120132554 | 0.14756534 |
| ENSDART00000039080 | wasb | 0.537774473 | 0.660070537 | 0.394408091 | -0.045801603 |
| ENSDART00000039161 | hdhd2 | 0.366788589 | 0.057115496 | -0.197391924 | -0.146678217 |
| ENSDART00000039206 | rps23 | 0.388349294 | 0.322594165 | 0.070172062 | -0.095059659 |
| ENSDART00000039277 | lhfpl3 | -0.493635642 | -0.41740998 | -0.318633213 | -0.189800628 |
| ENSDART00000039295 | lrrfip1a | 0.201172846 | -0.052744603 | -0.60255637 | -0.234396047 |
| ENSDART00000039312 | ipo4 | 0.472349209 | 0.340323729 | 0.32654819 | 0.07839591 |
| ENSDART00000039399 | cavin2a | -0.454722967 | -0.463350309 | -0.727916614 | -0.482344794 |
| ENSDART00000039443 | tuba8l4 | 0.45016266 | 0.547239574 | 0.701695743 | 0.194541839 |
| ENSDART00000039466 | rnf25 | 0.190959954 | 0.415197213 | 0.378723734 | 0.235880676 |
| ENSDART00000039485 | gabarapl2 | -0.111002032 | -0.162035427 | -0.331707696 | -0.213225245 |
| ENSDART00000039551 | mef2ca | 2.857314883 | 3.343180737 | 3.275011062 | 3.48649726 |
| ENSDART00000039571 | camk2a | -0.838498146 | -0.905374301 | -0.593078838 | -0.446882891 |
| ENSDART00000039585 | klhl36 | 0.23105636 | 0.314614252 | 0.405624735 | 0.264885345 |
| ENSDART00000039693 | pgp | -0.344619279 | -0.551044992 | -0.227681437 | -0.38433532 |
| ENSDART00000039746 | epb41b | 0.556721113 | 0.962382373 | 1.378686301 | 0.45048608 |
| ENSDART00000039788 | uqcrq | -0.201371674 | -0.242426303 | -0.394323978 | -0.159681057 |
| ENSDART00000039865 | sdhdb | -0.123077363 | -0.199973196 | -0.304203762 | -0.144165352 |
| ENSDART00000039868 | usp4 | -0.25602575 | -0.139457748 | -0.15513491 | -0.170681969 |
| ENSDART00000039987 | pgm3 | 0.351341795 | 0.131342573 | -0.075397144 | -0.106723338 |
| ENSDART00000040035 | ccdc80l1 | -0.195911171 | -0.175911639 | -0.396207214 | -0.331422513 |
| ENSDART00000040049 | camk2d2 | -0.441100303 | -0.359209869 | -0.184359733 | 0.062709499 |
| ENSDART00000040066 | adam9 | 0.358589887 | 0.583904956 | 0.549640677 | 0.256288209 |
| ENSDART00000040086 | pacsin1a | -0.311462824 | -0.286213879 | -0.425986821 | -0.031983095 |
| ENSDART00000040116 | tnrc5 | -0.132560784 | -0.181210054 | -0.368693498 | -0.22290886 |

| | | | | | |
|---|---|---|---|---|---|
| ENSDART00000040184 | tenm1 | -0.25843006 | -0.314319075 | -0.175496746 | -0.005818614 |
| ENSDART00000040275 | kcnj11l | -0.790164361 | -0.507671216 | -0.85342057 | -0.51870362 |
| ENSDART00000040278 | efna2a | -0.820985512 | -0.748851824 | -0.238813858 | 0.346525856 |
| ENSDART00000040334 | pik3r3b | -0.289525561 | -0.103810065 | 0.221174364 | 0.296766414 |
| ENSDART00000040346 | efr3ba | -0.15754636 | -0.295932904 | -0.334562517 | -0.212090743 |
| ENSDART00000040434 | asah1b | 0.563509259 | 0.420868789 | -0.056225927 | -0.32971968 |
| ENSDART00000040456 | cdc42bpab | -0.484798472 | -0.258449967 | -0.177779388 | 0.139329121 |
| ENSDART00000040500 | tspan9a | -0.374305977 | -0.544164723 | -0.123124308 | -0.153915406 |
| ENSDART00000040502 | trpc5a | -0.568729425 | -1.014519921 | -0.522696272 | -0.18638518 |
| ENSDART00000040537 | gjd1a | -0.421902855 | -0.214222628 | 0.04546907 | 0.177131603 |
| ENSDART00000040542 | arhgef12a | -0.286992716 | -0.169562154 | -0.197347098 | -0.111099085 |
| ENSDART00000040557 | CRIP2 (1 of many) | -0.132882678 | 0.562285777 | 0.768710088 | 0.441120913 |
| ENSDART00000040669 | sphkap | -0.388395859 | -0.454597188 | 0.021563144 | 0.051812339 |
| ENSDART00000040672 | mecp2 | -0.242382035 | -0.413395534 | -0.149398689 | -0.148226048 |
| ENSDART00000040701 | ing5a | 0.004576245 | -0.184827807 | -0.360745549 | -0.284929321 |
| ENSDART00000040708 | caprin2 | -0.196766553 | 0.78279717 | 0.834337788 | 0.695156181 |
| ENSDART00000040771 | rpl34 | 0.291435523 | 0.406434571 | 0.222752448 | -0.006604457 |
| ENSDART00000040804 | praf2 | 0.166225557 | 0.427661756 | 0.402173801 | 0.274020381 |
| ENSDART00000040827 | ncaph2 | 0.654906877 | 0.777407912 | 0.63027477 | 0.226543553 |
| ENSDART00000040900 | baxb | 0.66837016 | 0.786823818 | 0.515591205 | 0.094501002 |
| ENSDART00000041007 | stmn1b | -0.180266883 | 0.501280237 | 0.34630752 | 0.107820579 |
| ENSDART00000041114 | psmb2 | 0.343581501 | 0.272779677 | 0.256301247 | -0.006269993 |
| ENSDART00000041191 | gyg2 | 0.909324979 | 1.617533024 | 1.086480874 | 0.519765295 |
| ENSDART00000041257 | gsto2 | 0.760127953 | 0.383922047 | -0.10799465 | -0.396095115 |
| ENSDART00000041279 | tubb4b | 0.433747505 | 0.468816243 | 0.626770783 | 0.445102188 |
| ENSDART00000041388 | cacng2a | -0.088843708 | -0.391878721 | -0.181623249 | -0.021159404 |
| ENSDART00000041417 | camk1b | -0.259356662 | -0.222277099 | -0.408590245 | -0.120593834 |
| ENSDART00000041443 | igsf21a | -0.224595347 | -0.278632007 | -0.419145006 | -0.220026937 |
| ENSDART00000041468 | ap1ar | 0.253231094 | 0.535778421 | 0.657902256 | 0.098067269 |
| ENSDART00000041503 | slc4a4a | -0.308542931 | -0.316588825 | -0.297229141 | -0.158105481 |
| ENSDART00000041504 | tescb | 0.44458368 | -0.282095189 | 0.067591772 | -0.116025945 |
| ENSDART00000041707 | unc119a | -0.21008147 | -0.190962771 | -0.5453967 | -0.153347487 |
| ENSDART00000041714 | atp6v0a1b | -0.327889206 | -0.302694525 | -0.070700098 | 0.131627227 |
| ENSDART00000041728 | cyp26a1 | -0.110572523 | -0.30661175 | -0.074827314 | 0.003208979 |
| ENSDART00000041740 | ubl7a | -0.21096997 | -0.119372658 | 0.298633763 | 0.574535573 |
| ENSDART00000041751 | ercc1 | 0.342208194 | 0.265119266 | 0.084902107 | 0.028919147 |
| ENSDART00000041800 | epha8 | -0.01929649 | -0.346405332 | 0.164805877 | 0.748357168 |
| ENSDART00000041805 | metrn | -0.185458155 | -0.302026239 | -0.151541546 | -0.276934548 |
| ENSDART00000041820 | lingo1a | -0.3685918 | -0.267056385 | -0.206204269 | -0.004686852 |
| ENSDART00000041861 | syt1a | -0.326937152 | -0.373717465 | -0.346529457 | -0.191055943 |
| ENSDART00000041869 | grin1a | -0.314741638 | -0.554584424 | -0.238417502 | 0.088185899 |
| ENSDART00000041877 | csrnp1a | 1.299872398 | 1.369159722 | 1.350463811 | 0.717287101 |
| ENSDART00000041992 | dhrs12 | 0.21282475 | 0.60945999 | -0.222288782 | -0.248821854 |
| ENSDART00000042083 | gria4a | -0.246573777 | -0.413439839 | -0.038291419 | 0.265641743 |
| ENSDART00000042123 | cx52.6 | -0.622226461 | -0.079417665 | -0.692157173 | -0.611068819 |
| ENSDART00000042134 | dock7 | 0.063646619 | 0.209946461 | 0.48316367 | 0.274325565 |
| ENSDART00000042162 | tm7sf2 | 0.1050645 | 0.758038931 | 0.952561873 | 0.4137868 |
| ENSDART00000042189 | pdk2b | 0.734399082 | 1.569570791 | 1.007918384 | 0.211070696 |
| ENSDART00000042194 | cers4a | -0.343157696 | -0.266390398 | -0.236875341 | -0.142108033 |
| ENSDART00000042200 | aldoab | 0.726673418 | 1.529219824 | 2.318485768 | 1.809025522 |
| ENSDART00000042218 | pafah1b1a | 0.049322983 | 0.127134272 | 0.313096774 | 0.220746894 |

| | | | | | |
|---|---|---|---|---|---|
| ENSDART00000042250 | rap1b | 0.339690926 | 0.041784403 | 0.019413921 | -0.132436477 |
| ENSDART00000042255 | rab6bb | 0.249992007 | 1.135776594 | 1.498200114 | 1.307756366 |
| ENSDART00000042276 | nxph1 | -0.320960824 | -0.220089718 | -0.50268354 | -0.230788559 |
| ENSDART00000042297 | kdelc1 | 0.328425656 | 0.736886183 | 0.437422887 | 0.039597286 |
| ENSDART00000042307 | fam60a | -0.454119179 | -0.522927402 | -0.624149133 | -1.020018948 |
| ENSDART00000042386 | unm_sa1261 | -0.368338169 | -0.259611498 | -0.383565927 | -0.183992374 |
| ENSDART00000042481 | phf23a | -0.371442433 | -0.230558793 | -0.302265692 | -0.28776115 |
| ENSDART00000042572 | ablim1b | 0.023130185 | -0.084172657 | -0.818233794 | -0.360713717 |
| ENSDART00000042599 | dennd6aa | -0.0532753 | -0.029202145 | -0.371955611 | -0.136277272 |
| ENSDART00000042624 | arap3 | -0.123633155 | -0.147189694 | -0.365577073 | -0.154745368 |
| ENSDART00000042683 | cadpsb | -0.417914059 | -0.394489719 | 0.149869704 | 0.381243736 |
| ENSDART00000042963 | chst11 | 0.296547318 | 0.476599287 | 0.638913095 | 0.263644905 |
| ENSDART00000042972 | srpk1b | -0.590887937 | -0.595753709 | -0.259675811 | -0.090086557 |
| ENSDART00000042984 | epha6 | -4.335346398 | -2.736601299 | -1.023821359 | 0.479123476 |
| ENSDART00000043058 | TENM2 | -0.559723778 | -0.553182426 | -0.524601572 | -0.11140967 |
| ENSDART00000043076 | ppdpfb | -0.038298907 | 0.045116804 | -0.329828604 | -0.321738012 |
| ENSDART00000043091 | iqsec1b | -0.102282724 | -0.271543671 | -0.319516795 | -0.119256647 |
| ENSDART00000043173 | rpl18 | 0.389153851 | 0.347510342 | 0.235541148 | 0.00423952 |
| ENSDART00000043180 | gria3b | -0.628974157 | -0.919920549 | -0.567970281 | 0.039022237 |
| ENSDART00000043226 | guca1c | -0.531402053 | -0.370893475 | -0.804093243 | -0.754648782 |
| ENSDART00000043312 | srsf5a | -0.406457659 | -0.349179029 | 0.276259838 | -0.003669678 |
| ENSDART00000043429 | jph2 | -0.203422393 | 1.998676475 | 2.481431759 | 1.87668434 |
| ENSDART00000043455 | smad3b | -0.137393915 | -0.114661975 | -0.243963828 | -0.226543078 |
| ENSDART00000043492 | trappc6bl | -0.12751818 | -0.21743347 | -0.476970052 | -0.302538631 |
| ENSDART00000043507 | ciarta | -0.034891069 | -0.156171585 | -0.423543489 | -0.111653268 |
| ENSDART00000043651 | dnal1 | -0.182764591 | -0.191705301 | -0.306394992 | -0.190228735 |
| ENSDART00000043666 | hs1bp3 | 1.060777617 | 0.463082325 | 0.168627927 | -0.515305865 |
| ENSDART00000043678 | apobec2b | 1.815363093 | 0.931239114 | 0.794076282 | 0.812995775 |
| ENSDART00000043801 | cabp5b | -0.44296443 | -0.237302409 | -0.884658288 | -0.372660272 |
| ENSDART00000043823 | osbpl10b | -0.615302858 | -0.415836047 | -0.293972471 | 0.154234814 |
| ENSDART00000043855 | dclk2a | -0.104441255 | -0.395008642 | -0.62766162 | -0.184810585 |
| ENSDART00000043857 | irx5a | -0.492824084 | -0.561776224 | 0.072886283 | 0.234832625 |
| ENSDART00000043924 | mpp6b | -0.45376092 | -0.273823891 | -0.178237916 | -0.157850955 |
| ENSDART00000043932 | atp2a1 | 1.138744452 | 4.285725335 | 5.558096993 | 3.994336477 |
| ENSDART00000043933 | ndufb7 | -0.183005381 | -0.27475981 | -0.256491733 | -0.145691426 |
| ENSDART00000043945 | | -0.527606114 | -0.345502165 | -0.38556129 | -0.085040536 |
| ENSDART00000043953 | mfsd2b | -0.226277331 | -0.20295362 | -0.451440771 | 0.020277702 |
| ENSDART00000044000 | plxna3 | 0.522678231 | 0.518528602 | 0.862237135 | 0.748340859 |
| ENSDART00000044009 | scdb | -0.293214367 | -0.066392815 | 0.084209837 | 0.174747579 |
| ENSDART00000044057 | sept3 | -0.17670354 | 0.196164293 | 0.478730018 | 0.43476862 |
| ENSDART00000044150 | dnajc9 | -0.130894557 | -0.15046904 | -0.343680246 | -0.373143883 |
| ENSDART00000044154 | tnnt2c | 0.256126919 | 0.781011759 | 0.616305237 | 0.377618007 |
| ENSDART00000044157 | scn4ab | 1.170816112 | 2.050777967 | 2.7725392 | 2.001159499 |
| ENSDART00000044208 | lmo1 | -0.34602538 | -0.891618988 | -0.018405226 | 0.102622885 |
| ENSDART00000044238 | zgc:92066 | 1.060411872 | 1.124461317 | 0.621339815 | 0.051820002 |
| ENSDART00000044241 | KCNQ2 (1 of many) | -0.793205862 | -0.43808803 | 0.589879296 | 0.84351704 |
| ENSDART00000044264 | mmp14b | 0.997152397 | 1.115840894 | 0.915921565 | 0.094130452 |
| ENSDART00000044276 | dip2bb | -0.182549034 | -0.085935994 | 0.236910154 | 0.379645319 |
| ENSDART00000044294 | fryb | -0.196859444 | -0.540115494 | -0.186847147 | -0.085192364 |
| ENSDART00000044314 | itgav | 0.188708464 | -0.009103257 | -0.190172369 | -0.475418475 |
| ENSDART00000044328 | acss1 | 0.948368639 | 0.660526303 | 0.176495325 | -0.460639914 |

| | | | | | |
|---|---|---|---|---|---|
| ENSDART00000044371 | tox | -0.382184105 | -0.377592563 | -0.085511133 | 0.05287602 |
| ENSDART00000044423 | magi1b | -0.521004582 | -0.785194432 | 0.042021364 | 0.52355612 |
| ENSDART00000044426 | si:dkey-240h12.4 | 0.077516947 | -0.039107616 | -0.22879164 | -0.507759534 |
| ENSDART00000044453 | ano5a | -0.027551778 | 0.223592536 | 0.548330121 | 0.427829524 |
| ENSDART00000044647 | ppil3 | 0.391623501 | 0.578322707 | 0.226885679 | 0.10080052 |
| ENSDART00000044658 | letmd1 | 0.311235989 | 0.320526266 | 0.39411459 | 0.349216193 |
| ENSDART00000044678 | GABRA2 (1 of many) | -0.18505593 | -0.452041699 | -0.058625322 | 0.015850219 |
| ENSDART00000044733 | NPBWR2 | -0.533926323 | -0.49881296 | -0.385102988 | -0.276547602 |
| ENSDART00000044735 | gria1b | -0.281408167 | -0.37749512 | -0.41239512 | -0.179683873 |
| ENSDART00000044860 | maff | 0.057535561 | 0.039208376 | -0.366680053 | -0.441007369 |
| ENSDART00000044896 | camk2d2 | -0.471410347 | -0.378784897 | -0.323522916 | -0.120871605 |
| ENSDART00000044949 | syt16 | -0.001457693 | -0.053066568 | 0.72824965 | 0.634879457 |
| ENSDART00000044963 | loxl4 | 1.867896709 | 1.134004715 | 1.5146614 | 0.310989698 |
| ENSDART00000044986 | rnd1a | -0.894483212 | -0.275620607 | -0.461458 | -0.242681728 |
| ENSDART00000045071 | foxk2 | 0.01041623 | 0.131070865 | 0.347129546 | 0.265521213 |
| ENSDART00000045086 | prkceb | -0.397671937 | -0.107653926 | 0.08012307 | 0.031299867 |
| ENSDART00000045126 | lama5 | -0.044158589 | 0.007727115 | -0.324426294 | -0.496259172 |
| ENSDART00000045232 | mtss1la | -0.698215115 | -0.447289487 | 0.119673501 | -0.022641935 |
| ENSDART00000045284 | rpl37 | 0.494352175 | 0.447787745 | 0.229606145 | -0.010646089 |
| ENSDART00000045299 | ola1 | 0.514191862 | 0.666266592 | 0.533008533 | 0.271611648 |
| ENSDART00000045303 | tmprss9 | -0.116725282 | -0.194770382 | -0.365140883 | -0.064254559 |
| ENSDART00000045374 | smad3a | -0.063329624 | -0.157712113 | -0.303211349 | -0.108940629 |
| ENSDART00000045391 | srgap2 | 0.042169336 | 0.118632088 | 0.377794988 | 0.307955564 |
| ENSDART00000045397 | stx11b.1 | 1.953998203 | 1.413835145 | 1.924246083 | 1.02039349 |
| ENSDART00000045410 | thy1 | 0.906274294 | 2.922050483 | 2.917431913 | 2.440516966 |
| ENSDART00000045479 | syt4 | 0.199773705 | 0.561264928 | 0.42766733 | -0.021116223 |
| ENSDART00000045555 | rab41 | -0.452536385 | -0.052299617 | 0.3824053 | 0.576200707 |
| ENSDART00000045616 | gabbr1b | -0.290531763 | -0.52893984 | -0.416723119 | -0.192345482 |
| ENSDART00000045628 | irx6a | -0.424070635 | -0.239579413 | 0.17038084 | 0.276181325 |
| ENSDART00000045659 | tcp11l1 | 0.225446633 | 0.445852163 | 0.278888616 | 0.19328127 |
| ENSDART00000045675 | slc52a2 | 0.16071928 | 0.405611618 | 0.292146022 | 0.046169521 |
| ENSDART00000045682 | rrp36 | 0.38981283 | 0.603671455 | 0.338055221 | 0.296271067 |
| ENSDART00000045684 | porcn | -0.472251654 | -0.321238385 | -0.136837542 | 0.064329115 |
| ENSDART00000045697 | zgc:56493 | 0.323838871 | 0.24043595 | -0.000448434 | -0.189381569 |
| ENSDART00000045757 | march5l | 3.527750979 | 3.782483659 | 2.945706162 | 1.186881806 |
| ENSDART00000045842 | rcan3 | -0.531033223 | -0.615136042 | -0.34287969 | -0.068102336 |
| ENSDART00000045861 | slc43a2a | -0.304475968 | -0.378976343 | 0.111239312 | 0.188992557 |
| ENSDART00000045888 | tkta | -0.540178562 | -0.404491163 | -0.145350137 | 0.055411831 |
| ENSDART00000045933 | sh3glb1b | 0.424660866 | 0.274153114 | 0.044458674 | -0.153303197 |
| ENSDART00000046004 | wnt2bb | -0.582117328 | -0.289636407 | -0.865141829 | -0.874797573 |
| ENSDART00000046050 | pcbd1 | 0.519297333 | 0.526198373 | -0.092439376 | -0.055017497 |
| ENSDART00000046066 | capn1a | -0.174433704 | -0.141559786 | -0.304523077 | -0.134358035 |
| ENSDART00000046115 | mfsd2aa | 0.413173515 | 1.356876257 | 1.539393726 | 0.598141045 |
| ENSDART00000046209 | acbd7 | 0.172970527 | 0.805855312 | 0.630012733 | 0.123911744 |
| ENSDART00000046211 | lnx2a | -0.592157981 | -0.322543056 | -0.148137638 | -0.128059395 |
| ENSDART00000046218 | flnca | 6.980277858 | 6.302871335 | 6.08375751 | 5.531037957 |
| ENSDART00000046253 | prkcq | 0.856641094 | 1.199706901 | 0.196829623 | -0.396081861 |
| ENSDART00000046268 | pmelb | 0.894230709 | 0.185058494 | -0.384535315 | -0.811085128 |
| ENSDART00000046360 | rhoua | -0.234114921 | -0.11992892 | -0.459682602 | -0.396381281 |
| ENSDART00000046438 | kcnk2b | -1.164610191 | -1.614380279 | -0.460863171 | 0.258871322 |
| ENSDART00000046498 | sema3fa | -0.424113863 | -0.318396861 | -0.293026202 | -0.101692261 |

| | | | | | |
|---|---|---|---|---|---|
| ENSDART00000046530 | rab42a | -0.542354056 | -0.349553972 | -0.104901238 | 0.132258211 |
| ENSDART00000046542 | igf1rb | -0.265329623 | -0.432225627 | -0.135218314 | -0.147354602 |
| ENSDART00000046587 | ap2m1a | 0.393397352 | 0.521859633 | 0.500225377 | 0.048547949 |
| ENSDART00000046626 | PRKAR2B | -0.181202107 | -0.272867538 | -0.191341626 | -0.012099353 |
| ENSDART00000046663 | camta1b | -0.175908238 | -0.543108898 | -0.289499765 | -0.001152787 |
| ENSDART00000046678 | pak2b | 0.067537468 | 0.265866932 | 0.367938524 | 0.197303091 |
| ENSDART00000046689 | tmed3 | 0.310928608 | 0.094278442 | -0.151364778 | -0.47701209 |
| ENSDART00000046712 | zgc:86609 | 0.278113439 | 0.188615214 | -0.290177028 | -0.454070036 |
| ENSDART00000046716 | cited2 | -0.252125812 | -0.225219779 | -0.427941506 | -0.423861644 |
| ENSDART00000046764 | gmds | 0.16546148 | 0.334907039 | 0.47465655 | 0.2924672 |
| ENSDART00000046922 | rab13 | 0.426602975 | 0.215892939 | 0.022612864 | -0.308555932 |
| ENSDART00000046933 | sult1st5 | 0.218426293 | 0.869949792 | 0.854237199 | 0.702293309 |
| ENSDART00000046934 | coq9 | -0.280335275 | -0.284319265 | -0.255484353 | -0.03877329 |
| ENSDART00000046951 | ptpn11b | -0.12533711 | -0.212829251 | -0.391615008 | -0.186327808 |
| ENSDART00000046973 | capza1a | 0.117659515 | 0.282132124 | 0.303069026 | 0.119362421 |
| ENSDART00000046995 | txn2 | -1.624875893 | -1.011052145 | -0.305336636 | -0.0755861 |
| ENSDART00000047020 | casp9 | 0.406413734 | 0.845063479 | 0.618979742 | 0.291728295 |
| ENSDART00000047069 | tyms | 1.498823911 | 1.293125646 | 0.841397173 | 0.487713483 |
| ENSDART00000047073 | oxsr1a | 0.179223329 | 0.268680956 | 0.376160209 | 0.212719894 |
| ENSDART00000047082 | gdap1l1 | -0.237605119 | 0.285676986 | 0.669438597 | 0.59731396 |
| ENSDART00000047126 | clcn4 | -0.325495856 | -0.290597126 | 0.018460992 | 0.082569002 |
| ENSDART00000047143 | specc1 | 0.250947915 | 0.397558549 | 0.627196335 | 0.309975843 |
| ENSDART00000047175 | ssr4 | 0.274556908 | 0.120119155 | 0.036407988 | -0.174453213 |
| ENSDART00000047191 | glb1l | 1.329711044 | 0.552831622 | 0.213363411 | -0.364178392 |
| ENSDART00000047362 | msra | -0.031803809 | -0.196438308 | -0.422002476 | -0.295991402 |
| ENSDART00000047378 | sst3 | -0.489863502 | -0.562162931 | -0.649979789 | -0.586308909 |
| ENSDART00000047399 | mmp24 | -0.337136574 | -0.281370104 | 0.467116638 | 0.594580786 |
| ENSDART00000047409 | myh14 | 0.151853201 | 0.372826224 | 0.643058289 | 0.555183408 |
| ENSDART00000047416 | slc4a8 | -0.393460236 | -0.539636776 | -0.542467324 | -0.330988764 |
| ENSDART00000047541 | bach1b | -0.343772395 | -0.510091534 | -0.468889885 | -0.333515696 |
| ENSDART00000047569 | igf2b | -0.538912754 | -0.46572679 | -0.194582206 | 0.102619195 |
| ENSDART00000047662 | ppp1r13bb | -0.080629431 | -0.11550057 | -0.271468608 | 0.046859797 |
| ENSDART00000047728 | melk | 0.808255278 | 0.366287521 | 0.23278244 | 0.524042022 |
| ENSDART00000047857 | orc3 | 3.881796907 | 3.620009487 | 3.715059064 | 3.478890874 |
| ENSDART00000047954 | ugcg | -0.094752728 | 0.184472082 | 0.264381156 | 0.087288534 |
| ENSDART00000048036 | gem | -0.746612275 | -0.695957849 | -0.479539177 | -0.330372498 |
| ENSDART00000048050 | ITGB1BP2 | -0.056534701 | 0.311580562 | 0.724894589 | 0.983194822 |
| ENSDART00000048073 | zgc:171775 | 0.625715462 | 0.657465213 | 0.200858631 | -0.066721925 |
| ENSDART00000048107 | FP102018.1 | 0.402516182 | 0.222629184 | -0.14999501 | -0.581967671 |
| ENSDART00000048110 | six4b | 0.757934328 | 2.322346312 | 2.776043581 | 1.88012445 |
| ENSDART00000048365 | syt6b | -0.461087231 | -0.40037241 | -0.458496339 | -0.153902769 |
| ENSDART00000048383 | creld2 | 0.499343204 | 0.520824125 | 0.482480278 | 0.212076819 |
| ENSDART00000048432 | dlg4a | -0.380839229 | -0.483618741 | -0.31591813 | -0.034480778 |
| ENSDART00000048599 | rps19 | 0.313599405 | 0.28541891 | 0.281484325 | -0.024061243 |
| ENSDART00000048707 | srgap1b | -0.90586475 | -0.695392849 | -0.011664453 | 0.331175331 |
| ENSDART00000048775 | mbd3b | -0.291847963 | -0.361950456 | -0.358913558 | -0.222238531 |
| ENSDART00000048819 | rassf2a | -0.204366055 | -0.399787028 | -0.377471637 | -0.118487631 |
| ENSDART00000048853 | ube2d1a | 0.133955448 | 0.280395319 | 0.282059322 | 0.156573319 |
| ENSDART00000048855 | mtus1b | -0.449558058 | -0.427741832 | 0.166302333 | 0.000701193 |
| ENSDART00000048866 | ipmkb | -0.299703051 | -0.158343907 | 0.084121168 | 0.000422995 |
| ENSDART00000048871 | desi1a | -0.247171878 | -0.259133346 | 0.047182486 | 0.101740307 |

| | | | | | |
|---|---|---|---|---|---|
| ENSDART00000048890 | slc22a2 | 0.30008182 | 0.486824781 | 0.442015061 | 0.075671897 |
| ENSDART00000048893 | pcbp3 | -0.37198127 | -0.476664366 | -0.345261768 | -0.190932807 |
| ENSDART00000048940 | vill | -0.011902914 | -0.216832622 | -0.570278713 | -0.337359605 |
| ENSDART00000048977 | abcf1 | 0.295994115 | 0.413143633 | 0.435605861 | 0.158915281 |
| ENSDART00000048994 | pbx3b | -0.545498331 | -0.428509616 | 0.129533791 | 0.149586816 |
| ENSDART00000049036 | zgc:92275 | -0.447578375 | -0.84153571 | -0.52710535 | -0.386325614 |
| ENSDART00000049075 | add3a | -0.25124279 | -0.167771329 | -0.330721354 | -0.242087591 |
| ENSDART00000049099 | trip13 | 1.738467479 | 1.594316711 | 0.735690402 | 0.585616546 |
| ENSDART00000049135 | si:dkey-261m9.12 | 0.781337004 | 0.750536245 | 0.608174972 | 0.211334904 |
| ENSDART00000049154 | pthlha | -0.57822331 | -0.570371268 | -0.522569114 | -0.481173816 |
| ENSDART00000049177 | rab6ba | -0.532131764 | -0.115244957 | 0.417564602 | 0.557714845 |
| ENSDART00000049194 | gpr37b | -0.562015213 | 0.012994785 | 0.671207906 | 0.757619967 |
| ENSDART00000049240 | tob1a | -0.553621268 | -0.528794995 | -0.629368997 | -0.384085742 |
| ENSDART00000049264 | sdr16c5b | 0.496225849 | 0.925512272 | 0.840394736 | 0.492626723 |
| ENSDART00000049291 | gria3a | 0.136141556 | -3.339713475 | 0.061156993 | 0.119617746 |
| ENSDART00000049368 | atat1 | -0.456100083 | -0.161841688 | -0.10743096 | 0.240512647 |
| ENSDART00000049373 | cmtr1 | 3.529734054 | 4.85424569 | 5.483410402 | 3.408663479 |
| ENSDART00000049425 | sec61a1l | 0.040274537 | 0.042437444 | 0.248643876 | 0.074434633 |
| ENSDART00000049434 | scamp4 | 0.767657025 | 0.801831806 | 0.544244726 | 0.399234701 |
| ENSDART00000049437 | cdc42bpb | 0.021859432 | -0.018472292 | 0.242347511 | 0.210611193 |
| ENSDART00000049462 | rab15 | -0.425089968 | -0.185335259 | 0.21479615 | 0.289096284 |
| ENSDART00000049464 | fermt2 | 0.28468341 | 0.302053037 | 0.328009634 | 0.040193822 |
| ENSDART00000049465 | slc19a1 | 0.32288456 | 0.126425631 | -0.165912503 | -0.040050165 |
| ENSDART00000049572 | ncapd3 | 0.694925235 | 0.6794902 | 0.488650623 | 0.149625384 |
| ENSDART00000049589 | col11a1b | 0.748320952 | 1.159222727 | 1.621624084 | 0.283450272 |
| ENSDART00000049633 | zgc:110006 | -0.212812767 | -0.199160747 | -0.43210422 | -0.304919838 |
| ENSDART00000049676 | depdc1a | 1.209441584 | 0.891467797 | 0.780861798 | 0.190374501 |
| ENSDART00000049684 | bag2 | 0.817498026 | 0.419391857 | 0.164101637 | 0.068852338 |
| ENSDART00000049722 | pfdn5 | 0.271437948 | 0.37616378 | 0.12572137 | -0.064637497 |
| ENSDART00000049793 | gstm.1 | -0.113008537 | -0.081297777 | -0.307261435 | -0.386339749 |
| ENSDART00000049836 | bgnb | -0.474953019 | -0.067188313 | -0.175252125 | -0.757542027 |
| ENSDART00000049885 | si:dkey-172j4.3 | -0.146868316 | -0.089398841 | -0.317501099 | -0.03734398 |
| ENSDART00000049900 | tagln2 | 0.588048217 | 1.077774727 | 1.105142914 | 0.190917953 |
| ENSDART00000049992 | syt9b | -0.672988403 | -0.530192885 | 0.095421952 | 0.328394957 |
| ENSDART00000050018 | cnksr1 | 0.043033394 | -0.349786212 | -0.053406139 | -0.094004348 |
| ENSDART00000050037 | chrnb3b | -0.414911692 | -0.004263798 | 0.444800143 | 0.950014126 |
| ENSDART00000050077 | sdcbp | -0.585127298 | -0.412438306 | -0.175403243 | -0.033038502 |
| ENSDART00000050140 | CABZ01088365.1 | -0.08566459 | -0.329955925 | -0.093026456 | -0.10850944 |
| ENSDART00000050202 | rcan3 | -0.240590006 | -0.307380747 | -0.423427653 | -0.130410568 |
| ENSDART00000050217 | efna1a | -0.276270159 | -0.087399972 | -0.269604188 | -0.23920109 |
| ENSDART00000050230 | tspan3a | -0.260999172 | -0.146975695 | -0.364155673 | -0.280149491 |
| ENSDART00000050271 | hexb | 0.237342774 | -0.182676311 | -0.489143261 | -0.233593029 |
| ENSDART00000050303 | b3gat2 | -0.439065182 | -0.488507748 | -0.270928467 | 0.095807034 |
| ENSDART00000050308 | calm1b | -0.359535363 | -0.241046962 | -0.074464366 | 0.015414816 |
| ENSDART00000050311 | rltpr | -0.433814029 | -0.24205503 | 0.339832334 | 0.433192877 |
| ENSDART00000050332 | gna12a | 0.472300359 | 0.391620605 | 0.3757936 | 0.086884817 |
| ENSDART00000050352 | si:ch211-87m7.2 | 0.324289445 | 0.10825652 | 0.093171262 | -0.186818955 |
| ENSDART00000050399 | npc2 | 0.729567353 | 0.287798394 | -0.063690935 | -0.310097689 |
| ENSDART00000050445 | trim2a | -0.427473932 | -0.288810529 | -0.127175492 | 0.041370255 |
| ENSDART00000050559 | sh3rf1 | -0.402839399 | -0.313691549 | -0.2479685 | -0.17125224 |
| ENSDART00000050750 | rrm2b | -0.324758311 | -0.221575036 | -0.394470152 | -0.266012642 |

| | | | | | |
|---|---|---|---|---|---|
| ENSDART00000050753 | cd36 | -0.281884107 | 0.047747268 | 0.431169618 | 0.458008288 |
| ENSDART00000050762 | phactr3b | -0.629592387 | -0.363049766 | -0.266715209 | -0.083768273 |
| ENSDART00000050847 | GLDC | 0.384969982 | 0.265449522 | 0.198357794 | 0.108750757 |
| ENSDART00000050863 | zgc:101858 | 0.531270185 | 0.546733132 | 0.428408309 | 0.278322374 |
| ENSDART00000050898 | ncf1 | 0.6701936 | 0.170526908 | -0.157127601 | -0.640736559 |
| ENSDART00000050910 | orai2 | -0.105799627 | 0.023857148 | 0.452924674 | 0.120793773 |
| ENSDART00000051182 | arhgap4b | -0.41225839 | -0.320516515 | -0.225939723 | -0.055826953 |
| ENSDART00000051197 | c10h21orf59 | -0.175790498 | -0.1331203 | -0.516409592 | -0.442586367 |
| ENSDART00000051231 | gnb2 | -0.235263329 | 0.124417594 | 0.575032632 | 0.495502612 |
| ENSDART00000051234 | tnika | 0.106489865 | 0.189584243 | 0.309030435 | 0.511149463 |
| ENSDART00000051357 | zmat5 | -0.126081267 | -0.164432805 | -0.376616148 | -0.245219161 |
| ENSDART00000051392 | spns3 | 1.320734766 | 0.750958136 | 0.798555508 | -0.096372125 |
| ENSDART00000051491 | sfrp1a | 0.324084954 | 0.211296795 | -0.122014524 | -1.0775933 |
| ENSDART00000051515 | zgc:110329 | 0.175997647 | 0.392691176 | 0.340182733 | 0.206652443 |
| ENSDART00000051516 | tacr1a | -0.234436772 | -0.252170036 | -0.371966451 | -0.377528767 |
| ENSDART00000051518 | rasa1a | -0.264162997 | -0.276154428 | 0.166590332 | 0.366548962 |
| ENSDART00000051546 | rps6ka3a | -0.063638799 | 0.031915539 | -0.376231824 | -0.261540289 |
| ENSDART00000051552 | mpdu1a | -0.142610001 | -0.050886814 | -0.417815071 | -0.558529014 |
| ENSDART00000051556 | abca1b | 0.962705135 | 0.587027009 | 0.239926413 | -0.279230387 |
| ENSDART00000051566 | zgc:101016 | 0.072400267 | -0.278957969 | -0.822407258 | -0.192676389 |
| ENSDART00000051614 | tchp | 0.132551453 | 0.24050278 | 0.418776194 | 0.215450614 |
| ENSDART00000051621 | pgam5 | 0.243607349 | 0.33552261 | 0.455304621 | 0.149179919 |
| ENSDART00000051644 | coq5 | -0.310043454 | -0.139390844 | -0.270196538 | -0.205912752 |
| ENSDART00000051655 | snrnp27 | -0.149284802 | -0.196914421 | -0.340354698 | -0.296345743 |
| ENSDART00000051664 | ypel1 | -0.285276636 | -0.074600808 | -0.32874479 | -0.098702346 |
| ENSDART00000051666 | ppm1f | 0.089738986 | 0.277096017 | 0.445138137 | 0.206148798 |
| ENSDART00000051693 | irx4a | -1.596922819 | -2.014349322 | 0.164610846 | 0.924604512 |
| ENSDART00000051697 | evla | -0.055749697 | 0.354222037 | 0.437976697 | 0.322467243 |
| ENSDART00000051723 | si:ch211-193k19.1 | -0.524075663 | -0.333228258 | -0.259244771 | -0.088981638 |
| ENSDART00000051763 | rps3a | 0.36528781 | 0.402890251 | 0.287932185 | 0.011460462 |
| ENSDART00000051792 | sema3aa | 0.043541661 | -0.100781063 | -0.302534685 | -0.247434431 |
| ENSDART00000051807 | lasp1 | 0.121443442 | 0.384838028 | 0.393366972 | 0.231292626 |
| ENSDART00000051906 | ube2c | 1.239012745 | 1.032037835 | 0.553625781 | 0.164264884 |
| ENSDART00000051919 | n6amt1 | 0.421272913 | 0.445198021 | 0.366479977 | 0.202998784 |
| ENSDART00000051948 | si:dkey-17m8.2 | -2.249269734 | -0.678959768 | -1.034434829 | -1.318620097 |
| ENSDART00000051974 | drd4b | -0.419693377 | 0.024027774 | -0.191636302 | -0.159413451 |
| ENSDART00000052029 | cart3 | -0.527262177 | -0.518524777 | -0.63363087 | -0.567396556 |
| ENSDART00000052061 | cnn2 | 0.810381891 | 0.55886321 | 0.442877218 | -0.431652883 |
| ENSDART00000052065 | si:rp71-39b20.4 | -0.313076871 | -0.443642109 | -0.50831344 | -0.2481846 |
| ENSDART00000052067 | insl3 | -0.34446258 | -0.603612124 | -0.679111871 | -0.291897986 |
| ENSDART00000052082 | rpl30 | 0.377785904 | 0.342931306 | 0.098750304 | -0.086094728 |
| ENSDART00000052083 | fjx1 | -0.061315226 | -0.667932816 | -0.149604151 | 0.07387804 |
| ENSDART00000052090 | fuca1.2 | 0.607737239 | 0.489644017 | 0.400376939 | 0.091450373 |
| ENSDART00000052104 | fuca1.1 | 0.706031916 | 0.625980807 | 0.554044121 | 0.174374129 |
| ENSDART00000052113 | lingo1b | -0.337763041 | -0.148396867 | -0.080123582 | -0.033848116 |
| ENSDART00000052124 | fam49al | -0.409057809 | -0.177161374 | 0.563238983 | 0.47605027 |
| ENSDART00000052126 | yars | 0.350203533 | 0.365550787 | 0.323606131 | 0.105596802 |
| ENSDART00000052168 | hrh3 | 0.375251408 | 0.91270689 | 1.213693931 | 0.380712944 |
| ENSDART00000052256 | sumo3b | -0.129514175 | -0.087060285 | -0.253552673 | -0.198560199 |
| ENSDART00000052307 | arrdc3b | -0.385234832 | -0.285495364 | -0.380608917 | -0.283766566 |
| ENSDART00000052318 | mdka | -0.324471926 | -0.37363799 | -0.54050807 | -0.370260207 |

| | | | | | |
|---|---|---|---|---|---|
| ENSDART00000052322 | zgc:110699 | -0.266905411 | -0.248249923 | -0.482205646 | -0.223841905 |
| ENSDART00000052331 | rps20 | 0.422414342 | 0.403711723 | 0.215205833 | -0.059084102 |
| ENSDART00000052346 | gnao1b | -0.534691372 | -0.39418636 | -0.143300564 | 0.218777626 |
| ENSDART00000052351 | cnep1r1 | -0.133092087 | -0.135980199 | -0.291785579 | -0.071751039 |
| ENSDART00000052385 | tph1b | -0.370644209 | -0.494882963 | -0.236758451 | -0.433606661 |
| ENSDART00000052397 | pias1a | -0.248631388 | -0.263511268 | -0.331035933 | -0.095610748 |
| ENSDART00000052421 | txnipa | 1.054359781 | 1.031010785 | 1.048664605 | 0.430054853 |
| ENSDART00000052423 | spry2 | -0.373633249 | -0.274093568 | -0.317163264 | -0.534996547 |
| ENSDART00000052503 | nudcd1 | 0.220268862 | 0.211574095 | 0.29643349 | 0.134588484 |
| ENSDART00000052511 | hnrnpa0l | 0.098563881 | 0.214926285 | 0.391405743 | 0.211573162 |
| ENSDART00000052537 | zgc:163107 | 0.254281338 | 0.227112479 | 0.142188905 | 0.117476144 |
| ENSDART00000052539 | myo1ea | 0.574953247 | 0.619444947 | 0.654028338 | 0.173056841 |
| ENSDART00000052541 | ccnb2 | 1.168966788 | 1.017487756 | 0.69127691 | 0.504285143 |
| ENSDART00000052620 | npy | -0.456788991 | -0.449905237 | -0.529362885 | -0.310086201 |
| ENSDART00000052638 | slc27a2a | 0.420886529 | 0.920976528 | 0.833136576 | 0.001602661 |
| ENSDART00000052656 | rras2 | 0.556656504 | 0.801505793 | 0.534247735 | 0.118931735 |
| ENSDART00000052703 | nucb2b | -0.072514611 | -0.317049949 | -0.490530665 | -0.459714357 |
| ENSDART00000052730 | rps13 | 0.392793291 | 0.407259594 | 0.231403052 | 0.009148185 |
| ENSDART00000052749 | nod1 | 0.321629747 | 0.111770638 | -0.064002417 | -0.693750338 |
| ENSDART00000052761 | rpl39 | 0.335691979 | 0.34926929 | 0.1141905 | -0.000371665 |
| ENSDART00000052802 | calb2b | -1.334486374 | -1.407708939 | -0.475837952 | 0.218368357 |
| ENSDART00000052838 | acta1a | 0.35046601 | 2.371192607 | 2.304510866 | 0.91139897 |
| ENSDART00000052871 | pop7 | -0.057197275 | 0.295354721 | 0.463638671 | 0.27819074 |
| ENSDART00000052912 | pcdh20 | -0.38541457 | -0.110151808 | -0.462368809 | -0.238274084 |
| ENSDART00000052915 | ash1l | 0.173309311 | 0.167145004 | 0.317504355 | 0.12008578 |
| ENSDART00000052917 | slc3a2a | 0.509465627 | 0.333845604 | -0.024881296 | -0.285712584 |
| ENSDART00000052989 | ache | -0.369275035 | -0.306496087 | -0.342496448 | -0.099646921 |
| ENSDART00000053001 | tcn2 | 0.058701765 | -0.020005828 | -0.457202856 | -0.602127411 |
| ENSDART00000053003 | hexim1 | -0.295110864 | -0.174423589 | 0.055276892 | 0.067518179 |
| ENSDART00000053095 | rhbdf1a | 0.708583914 | 0.695350284 | 0.352388623 | 0.004181603 |
| ENSDART00000053120 | gpr185b | -0.57955621 | -1.320827874 | -0.56412655 | -0.654076001 |
| ENSDART00000053126 | aanat1 | -0.126925707 | -0.471091345 | -0.287328219 | -0.040926683 |
| ENSDART00000053139 | atp6v0cb | -0.239525606 | -0.317576827 | -0.287566582 | -0.021198281 |
| ENSDART00000053187 | stk35l | -0.090868048 | -0.135202146 | -0.364753171 | -0.24725243 |
| ENSDART00000053240 | cab39l | 0.313790093 | 0.462769475 | 0.441611468 | 0.290026864 |
| ENSDART00000053267 | hnrnpa1b | 0.306954561 | 0.087684612 | 0.037367395 | -0.002841599 |
| ENSDART00000053284 | bcl9 | -0.057531558 | 0.216352129 | 0.432379429 | 0.260365529 |
| ENSDART00000053285 | ndufa7 | -0.051624088 | -0.210448889 | -0.329202666 | -0.117246353 |
| ENSDART00000053304 | si:ch211-114n24.6 | 0.627425485 | 0.596987541 | 0.770158215 | 0.21603311 |
| ENSDART00000053310 | tmem18 | -0.091641733 | -0.057752607 | -0.353980434 | -0.264448619 |
| ENSDART00000053325 | tomm40l | 0.351677862 | 0.400415779 | 0.406067875 | 0.134025152 |
| ENSDART00000053367 | hmgn3 | -0.174648563 | -0.165335262 | -0.310027237 | -0.181225286 |
| ENSDART00000053380 | hax1 | -0.143446597 | -0.149458858 | -0.363330115 | -0.294300868 |
| ENSDART00000053405 | scamp2 | 0.377844511 | 0.106568039 | 0.060099408 | -0.324550719 |
| ENSDART00000053463 | mgll | -0.18197879 | -0.150318561 | -0.320930933 | -0.180307684 |
| ENSDART00000053494 | anks4b | -0.562445249 | -0.270911364 | -0.580434775 | -0.289114333 |
| ENSDART00000053750 | acsl2 | -0.784570064 | -0.626588121 | -0.673780261 | -0.449722468 |
| ENSDART00000053761 | bms1 | 0.415838377 | 0.296751152 | 0.241962107 | 0.100266391 |
| ENSDART00000053773 | lsm6 | -0.104076864 | -0.094751595 | -0.350900229 | -0.413521857 |
| ENSDART00000053806 | gab1 | 2.484308926 | 2.134716604 | 2.429792903 | 1.630078037 |
| ENSDART00000053834 | psmc6 | 0.096829198 | 0.261162402 | 0.176391614 | 0.055545263 |

| | | | | | |
|---|---|---|---|---|---|
| ENSDART00000053841 | ddhd1b | -0.251051379 | -0.26914625 | -0.031733869 | 0.07838534 |
| ENSDART00000053860 | saxo2 | -0.675755032 | -0.618170257 | -0.767927477 | -1.172976969 |
| ENSDART00000053869 | slc44a2 | 0.03477078 | 0.96666703 | 1.3453814 | 0.899279007 |
| ENSDART00000053916 | mtnr1ab | -0.333808774 | -0.363646472 | -0.291230509 | -0.411800208 |
| ENSDART00000053925 | mtmr7a | 0.252266702 | 0.492232945 | 0.532543286 | 0.441567176 |
| ENSDART00000053932 | cbsa | 1.196219296 | 0.806157212 | 0.553945426 | 0.279319174 |
| ENSDART00000054007 | slc8a4b | -0.011734485 | -0.299949751 | -0.451416872 | -0.315264472 |
| ENSDART00000054020 | hivep3b | 0.041000622 | -0.364922266 | -0.147208921 | -0.149490959 |
| ENSDART00000054026 | rcc1 | 0.376288807 | 0.340916908 | 0.360562186 | 0.188805345 |
| ENSDART00000054062 | nek12 | 0.979847079 | 2.121821413 | 1.72566276 | 1.104924836 |
| ENSDART00000054070 | surf2 | 0.311577894 | 0.338934517 | 0.34052692 | 0.125736963 |
| ENSDART00000054071 | | 0.085813894 | -0.09956216 | 0.306019728 | 0.602458638 |
| ENSDART00000054078 | rpa2 | 0.380370186 | 0.444560906 | 0.363788059 | 0.096462711 |
| ENSDART00000054137 | igfbp5b | -0.49960111 | -0.321727345 | 0.242454388 | 0.365417817 |
| ENSDART00000054175 | smad5 | 0.268560343 | 0.329305064 | 0.367006369 | -0.092635665 |
| ENSDART00000054202 | si:ch211-145b13.5 | -0.57317034 | 0.068003489 | -0.485735579 | -0.183705508 |
| ENSDART00000054209 | cdkn2a/b | -0.651631236 | -0.494220791 | -0.672480074 | -0.361655459 |
| ENSDART00000054243 | dpf2l | 0.196658591 | 0.403359284 | 0.517168503 | 0.26603871 |
| ENSDART00000054322 | cnrip1b | -0.129671726 | -0.251461369 | -0.057872634 | -0.024935203 |
| ENSDART00000054386 | qdprb1 | -0.130677564 | -0.146959168 | -0.522581238 | -0.268035824 |
| ENSDART00000054408 | gsg1l | -0.338795387 | -0.334881942 | -0.642215658 | -0.428166696 |
| ENSDART00000054452 | dlgap1b | -0.55212867 | -0.830902632 | -0.186012386 | 0.281181047 |
| ENSDART00000054462 | smim19 | -0.090105146 | -0.174385509 | -0.37090077 | -0.289350438 |
| ENSDART00000054472 | tll1 | -0.241128673 | -0.208136 | -0.494254448 | -0.202017577 |
| ENSDART00000054552 | cdh8 | -0.482196874 | -0.526834841 | -0.273874965 | 0.0505098 |
| ENSDART00000054574 | polr1e | 0.39788071 | 0.231113321 | 0.09078837 | -0.014285707 |
| ENSDART00000054581 | march1 | -0.100945889 | -0.377180996 | -0.379666436 | -0.107576973 |
| ENSDART00000054664 | tnnc1b | -0.083935925 | 3.176389417 | 3.430200861 | 1.884280525 |
| ENSDART00000054674 | mtnr1aa | -0.46530953 | -0.508091587 | -0.367618326 | -0.477932737 |
| ENSDART00000054687 | il1rapl2 | -0.293064258 | -0.530563685 | -0.194225592 | 0.06661817 |
| ENSDART00000054689 | atoh8 | -0.386924835 | -0.23478989 | -0.297672036 | -0.53086166 |
| ENSDART00000054691 | uba1 | 0.10035699 | 0.287144805 | 0.420196482 | 0.237625478 |
| ENSDART00000054735 | SYNPR (1 of many) | -0.204530212 | -0.189982465 | -0.242945057 | -0.369440481 |
| ENSDART00000054736 | bhlhe23 | -0.369485488 | -0.514619457 | -0.390416516 | -0.190127988 |
| ENSDART00000054760 | zgc:162144 | -0.154581168 | -0.586579914 | -0.580706302 | -0.278802593 |
| ENSDART00000054790 | zmp:0000001069 | -0.970537582 | -1.254323928 | -0.841864806 | -0.072822598 |
| ENSDART00000054833 | rgs11 | -0.582556718 | -0.789821251 | -0.427532995 | 0.127688375 |
| ENSDART00000054837 | ap1s2 | -0.004289143 | 0.194115514 | 0.293597748 | 0.151563344 |
| ENSDART00000054849 | pls3 | 0.365005671 | 0.852391802 | 1.00799598 | 0.660436059 |
| ENSDART00000054867 | aup1 | -0.021467306 | -0.053295505 | -0.309019838 | -0.217937602 |
| ENSDART00000054876 | npm1b | 0.13266464 | 0.298387409 | 0.130756001 | 0.083140409 |
| ENSDART00000054877 | fgf24 | -0.371366592 | -0.422467709 | -0.342651042 | -0.342396761 |
| ENSDART00000054987 | actb1 | 0.719668079 | 0.845173501 | 0.884967639 | 0.398548729 |
| ENSDART00000054989 | fscn1b | -0.529299226 | -0.417661269 | 0.066759573 | 0.494675778 |
| ENSDART00000055019 | ndufa4 | -0.243803126 | -0.403243102 | -0.444723985 | -0.066106721 |
| ENSDART00000055038 | rybpa | -0.189184893 | -0.138456944 | 0.268105644 | 0.373370341 |
| ENSDART00000055071 | nptx2a | -3.565846773 | -2.700342636 | -0.646259822 | -0.789148036 |
| ENSDART00000055134 | ogfr | -0.383562939 | -0.286066309 | -0.21229445 | -0.172842524 |
| ENSDART00000055139 | col9a3 | 0.340683061 | 0.20859259 | -0.192921312 | -1.195377277 |
| ENSDART00000055152 | taf11 | -0.482193927 | -0.885413263 | -0.244432703 | -0.649425725 |
| ENSDART00000055160 | il11a | 1.677180571 | 0.610178512 | -0.274837581 | -0.887218575 |

| | | | | | |
|---|---|---|---|---|---|
| ENSDART00000055171 | grapa | 0.991444674 | 1.026501701 | 0.907734856 | 0.689609359 |
| ENSDART00000055186 | atp5j2 | -0.168904246 | -0.309514947 | -0.449394402 | -0.201772036 |
| ENSDART00000055253 | filip1l | 3.442073683 | 3.796340861 | 4.199635374 | 3.015929109 |
| ENSDART00000055262 | cdk5r1a | -0.345552765 | -0.416237328 | -0.217352163 | -0.195826689 |
| ENSDART00000055264 | CA10 (1 of many) | -0.387267589 | -0.307777286 | -0.264643025 | -0.127280824 |
| ENSDART00000055269 | gng13b | -0.70656822 | -0.547882014 | 0.352208471 | 0.531882213 |
| ENSDART00000055287 | zgc:109934 | 0.762332695 | 0.487859158 | -0.045189444 | -0.423101752 |
| ENSDART00000055325 | psmb7 | 3.620821341 | 3.668842367 | 3.694371073 | 3.321198364 |
| ENSDART00000055328 | nek6 | 0.491226657 | 0.428884947 | 0.11516767 | -0.052210357 |
| ENSDART00000055336 | dennd1a | 0.280502184 | 0.250804932 | 0.331143912 | 0.195537503 |
| ENSDART00000055340 | fus | -0.011490396 | 0.383733399 | 0.662780299 | 0.416441518 |
| ENSDART00000055380 | tubb5 | 2.169554793 | 3.899556169 | 4.316772272 | 3.57490014 |
| ENSDART00000055395 | osr2 | 0.396980684 | 1.020974716 | 1.320896999 | 1.066319399 |
| ENSDART00000055428 | cbx7a | 0.780269243 | 0.615351279 | 0.524687311 | -0.270346219 |
| ENSDART00000055465 | si:ch211-149k23.9 | -0.23663044 | -0.394558494 | -0.331258304 | -0.323817823 |
| ENSDART00000055473 | grb2b | -0.073731957 | 0.038177184 | -0.265388265 | -0.174944785 |
| ENSDART00000055487 | chmp3 | -0.028426397 | -0.094862832 | -0.316592568 | -0.291442829 |
| ENSDART00000055492 | ddx5 | 0.105976659 | 0.207326629 | 0.32877191 | 0.140872666 |
| ENSDART00000055567 | gnrhr4 | -0.356419162 | -0.425772903 | -0.482362822 | -0.623317801 |
| ENSDART00000055607 | pdgfbb | -0.474199721 | -0.401424866 | -0.415940684 | -0.433592391 |
| ENSDART00000055609 | atf4b | 0.138344185 | 0.51758353 | 0.353617358 | 0.087935611 |
| ENSDART00000055611 | isca2 | -0.127743819 | -0.187288303 | -0.524298231 | -0.307747827 |
| ENSDART00000055694 | cdab | -0.082751258 | -0.099425929 | -0.28235988 | -0.295865587 |
| ENSDART00000055706 | her15.1 | -1.483608056 | -1.154441068 | -0.698110478 | -1.436427969 |
| ENSDART00000055709 | her2 | -0.984524994 | -0.833253569 | -0.717745512 | -1.233522642 |
| ENSDART00000055710 | aldh4a1 | -0.225772345 | -0.209769264 | -0.373180163 | -0.178717825 |
| ENSDART00000055756 | tbc1d12a | -0.033893992 | 0.095679716 | 0.342628218 | 0.232107662 |
| ENSDART00000055779 | ggact.2 | -0.390632373 | -0.197002813 | -0.565987089 | -0.546417063 |
| ENSDART00000055780 | jpt2 | 0.208785852 | 0.245567055 | 0.12691728 | -0.082734531 |
| ENSDART00000055817 | PIGG | -0.041747749 | -0.262998575 | -0.390921965 | -0.198616638 |
| ENSDART00000055890 | znf385c | -0.397413591 | -0.444425008 | -0.419820845 | -0.297765515 |
| ENSDART00000055913 | hist2h2l | -0.328500488 | -0.315748168 | -0.27355808 | -0.150594371 |
| ENSDART00000055932 | pigh | -0.149859895 | -0.076712594 | -0.279647992 | -0.164515657 |
| ENSDART00000055936 | isl2b | -1.527471977 | -1.312125436 | 0.470803449 | 0.853726063 |
| ENSDART00000055995 | sagb | -0.512319213 | -0.180320177 | -0.444781708 | -0.186196646 |
| ENSDART00000056005 | ascl1a | 1.787445127 | 1.283325532 | 0.749862779 | 0.792492648 |
| ENSDART00000056035 | PMM1 | -0.318342084 | -0.457172695 | -0.421149458 | -0.159828821 |
| ENSDART00000056081 | sulf1 | 1.721344224 | 3.792219559 | 4.448503543 | 2.727690528 |
| ENSDART00000056138 | igsf8 | -0.41438699 | -0.276734989 | -0.097715498 | 0.052342087 |
| ENSDART00000056213 | pik3r1 | -0.34639661 | -0.306212461 | -0.436349985 | -0.247937167 |
| ENSDART00000056254 | stap2a | 0.934389037 | 1.208596726 | 1.287672321 | 0.743017194 |
| ENSDART00000056278 | SLC25A22 (1 of many) | -0.288459944 | -0.379459616 | 0.002234165 | 0.21084669 |
| ENSDART00000056286 | h1f0 | -0.326358805 | 0.273543898 | 0.626230484 | 0.593578182 |
| ENSDART00000056294 | pitrm1 | 0.368247543 | 0.389541704 | 0.328836098 | 0.178268037 |
| ENSDART00000056295 | psap | 0.394315497 | 0.070717853 | -0.202183441 | -0.455675198 |
| ENSDART00000056305 | fzd8b | -0.95322142 | -0.49703101 | 0.067617805 | 0.048955399 |
| ENSDART00000056328 | elovl4b | -0.472600721 | -0.194626997 | -0.10087467 | 0.110456628 |
| ENSDART00000056333 | CU929150.1 | 0.302438567 | 0.370614966 | 0.439965996 | 0.209083279 |
| ENSDART00000056369 | cadm2a | -0.177754586 | -0.327047473 | -0.068598077 | -0.03429359 |
| ENSDART00000056376 | tmem55ba | 0.03327073 | 0.191352043 | 0.318689436 | 0.072412325 |
| ENSDART00000056420 | alas2 | 0.719442022 | 0.937854985 | 1.477738462 | 0.179843085 |

| | | | | | |
|---|---|---|---|---|---|
| ENSDART00000056457 | mitfa | 0.929754118 | 0.38849893 | 0.30836176 | -0.311729197 |
| ENSDART00000056460 | gbp1 | 3.785035011 | 3.151116027 | 3.74676295 | 2.092352698 |
| ENSDART00000056466 | camk2d1 | -0.163681889 | -0.265767151 | -0.291660383 | 0.000303403 |
| ENSDART00000056514 | gng7 | -0.676363728 | -0.720108736 | -0.535048991 | -0.288646977 |
| ENSDART00000056522 | skila | -0.013727131 | -0.304210156 | -0.124205506 | -0.010199259 |
| ENSDART00000056540 | casq1a | 0.340240931 | 1.131481384 | 1.513535686 | 1.324229104 |
| ENSDART00000056544 | tox4a | 0.131870382 | 0.296259606 | 0.350040714 | 0.249645538 |
| ENSDART00000056577 | RBP1 | 0.71451345 | 0.476305531 | 0.296937751 | 0.089894332 |
| ENSDART00000056639 | faim2a | -0.345615546 | -0.285647372 | -0.169343793 | 0.000937504 |
| ENSDART00000056671 | brinp2 | -0.564206507 | -0.55260615 | -0.353392227 | 0.031594069 |
| ENSDART00000056686 | mrc1b | 0.993000172 | 0.810204425 | 0.489266441 | -0.068136928 |
| ENSDART00000056712 | etfdh | 0.23563688 | 0.425716128 | 0.377386118 | 0.136575345 |
| ENSDART00000056721 | ldhd | -0.445248015 | -0.387879357 | -0.435006793 | -0.179899207 |
| ENSDART00000056734 | setd7 | -0.044426299 | -0.050437779 | -0.332150365 | -0.191170985 |
| ENSDART00000056735 | rgs20 | -0.263112929 | -0.317514258 | -0.646869905 | -0.425880318 |
| ENSDART00000056795 | hectd3 | 0.187585997 | 0.34309988 | 0.435708832 | 0.360997812 |
| ENSDART00000056810 | drd1b | -0.453660686 | -0.522513213 | -0.435699921 | -0.247093083 |
| ENSDART00000056865 | ctnnbip1 | 0.431885199 | 0.971175617 | 1.010655462 | 0.599546466 |
| ENSDART00000056885 | CU929046.1 | -0.462145697 | -0.331087748 | 0.285634599 | 0.389113052 |
| ENSDART00000056893 | pdcd7 | 0.040703622 | 0.122107923 | 0.35136997 | 0.096989832 |
| ENSDART00000056927 | egln1a | -0.373304639 | -0.331810821 | -0.548972472 | -0.202283187 |
| ENSDART00000056939 | zgc:85858 | -0.254903494 | -0.357469282 | -0.615277126 | -0.499477333 |
| ENSDART00000056963 | stk25b | -0.084642109 | -0.27353983 | -0.240916991 | -0.092979316 |
| ENSDART00000056987 | marcksl1a | 0.199644536 | 0.480185443 | 0.561780108 | 0.295794028 |
| ENSDART00000056996 | sfrp5 | 0.077859934 | 0.01622227 | -0.244583159 | -1.042810978 |
| ENSDART00000057095 | si:dkey-24p1.1 | -0.474380833 | -0.459390323 | -0.572876221 | -0.481612744 |
| ENSDART00000057124 | tefa | -0.17959692 | 0.074135039 | -0.330628318 | -0.262330789 |
| ENSDART00000057125 | tefa | -0.190705988 | 0.094171259 | -0.306461854 | -0.301060608 |
| ENSDART00000057159 | cacnb1 | 1.304600132 | 2.011299334 | 2.307200837 | 1.559835487 |
| ENSDART00000057174 | arpc5a | 0.695733745 | 0.614813351 | 0.637681436 | 0.535387536 |
| ENSDART00000057258 | slc12a5a | -0.776929597 | -2.750292921 | -0.618148385 | -0.058229169 |
| ENSDART00000057299 | st6galnac5a | -0.846236683 | -0.849009548 | -0.253708637 | -0.007082793 |
| ENSDART00000057318 | dusp8b | 0.80659762 | 0.962100249 | 0.680537568 | 0.386784732 |
| ENSDART00000057320 | zgc:171579 | -1.053157551 | -0.622493525 | -1.271148863 | -1.344481218 |
| ENSDART00000057325 | cacng4a | -0.326351391 | -0.28139538 | -0.566189708 | -0.15243738 |
| ENSDART00000057369 | igfbp5a | -0.503493701 | -0.12557648 | 0.011432007 | -0.035083178 |
| ENSDART00000057377 | arg2 | -0.266719608 | -0.373190344 | -0.501724428 | -0.206121282 |
| ENSDART00000057422 | pacsin1a | -0.590114235 | -0.673330755 | -0.518895915 | -0.206589128 |
| ENSDART00000057439 | parla | -0.763945698 | -0.13237506 | -0.4980372 | -0.373141189 |
| ENSDART00000057458 | CABZ01088149.1 | 0.207455616 | 0.356974308 | 0.107522223 | 0.159078208 |
| ENSDART00000057519 | zgc:194209 | 0.014343488 | -0.12072846 | -0.449908342 | -0.453116382 |
| ENSDART00000057553 | ch25hl1.1 | 0.477648419 | 0.50568744 | -0.255154291 | -1.200576847 |
| ENSDART00000057565 | sdhaf4 | -0.101594286 | -0.172555564 | -0.390218832 | -0.338776854 |
| ENSDART00000057584 | slc1a4 | 0.291595567 | 1.233259249 | 1.405286425 | 1.089173123 |
| ENSDART00000057638 | hk1 | -0.255062585 | -0.314354089 | -0.268977492 | -0.110279892 |
| ENSDART00000057644 | lhx4 | -0.267668007 | -0.268543602 | -0.190125613 | -0.119751289 |
| ENSDART00000057645 | qsox1 | 0.191462539 | 0.115464338 | -0.177011398 | -0.490873756 |
| ENSDART00000057689 | bag3 | -0.537315577 | -0.216588912 | -0.658528557 | -0.795247472 |
| ENSDART00000057710 | ccdc85a | -0.175789149 | -0.226891363 | -0.323862394 | -0.128238988 |
| ENSDART00000057865 | ier3ip1 | -0.032004312 | -0.136273649 | -0.345681209 | -0.364367077 |
| ENSDART00000057910 | nrgna | -0.338737739 | -0.336045781 | -0.223139332 | 0.191467698 |

| | | | | | |
|---|---|---|---|---|---|
| ENSDART00000057918 | si:ch211-147h1.4 | 0.058919084 | -0.340722361 | -0.680330981 | -0.211102009 |
| ENSDART00000057957 | itm2cb | -0.410626815 | -0.396999775 | -0.657592832 | -0.426965619 |
| ENSDART00000058093 | ldlrap1b | -0.019793679 | -0.383158495 | -0.468203159 | -0.036502488 |
| ENSDART00000058147 | dync2li1 | -0.035887703 | -0.047873047 | -0.361961236 | -0.288662687 |
| ENSDART00000058255 | bbs5 | -0.153890395 | -0.351139568 | -0.380364475 | -0.12338798 |
| ENSDART00000058258 | gng5 | 0.433032497 | 0.476321934 | 0.436326472 | 0.078774872 |
| ENSDART00000058277 | znf800b | 0.363600272 | 0.378301813 | 0.156623376 | -0.047842936 |
| ENSDART00000058324 | rpz4 | 0.754354311 | 0.61115296 | 0.469864295 | -0.17226809 |
| ENSDART00000058339 | ap3s2 | -0.205243617 | -0.191498501 | -0.306412801 | -0.176311273 |
| ENSDART00000058346 | c1qbp | 0.293247509 | 0.073830319 | 0.050211764 | 0.059638874 |
| ENSDART00000058370 | arhgap32b | -0.032405011 | 0.136784851 | 0.285715572 | 0.090147098 |
| ENSDART00000058384 | gapdhs | -0.42821082 | -0.374069856 | -0.238011978 | -0.087569823 |
| ENSDART00000058415 | zmp:0000001075 | -0.428240629 | -0.582010832 | -0.289943093 | -0.116590577 |
| ENSDART00000058424 | fam46ba | -0.297360056 | -0.251077248 | -0.176640513 | -0.109784644 |
| ENSDART00000058466 | fgfbp2a | 0.160228453 | 0.394614467 | 0.900155936 | 0.480238405 |
| ENSDART00000058470 | pik3r1 | -0.272621258 | -0.391826992 | -0.565271638 | -0.177820971 |
| ENSDART00000058484 | cnn3b | -0.281927631 | -0.203420372 | -0.230407516 | -0.341119299 |
| ENSDART00000058485 | rai14 | 0.377402754 | 0.592516996 | 0.371359301 | 0.056635681 |
| ENSDART00000058574 | | 3.231975259 | 2.942133944 | 3.360676782 | 3.717330017 |
| ENSDART00000058605 | scpep1 | 0.659557613 | 0.259300703 | 0.157418507 | -0.140654612 |
| ENSDART00000058628 | ccsapb | -0.109779777 | -0.249497929 | -0.321372328 | -0.06107705 |
| ENSDART00000058665 | kif20bb | 0.867797834 | 0.730592963 | 0.371647572 | 0.126469391 |
| ENSDART00000058667 | RDH13 (1 of many) | -0.138024992 | -0.086121321 | -0.690305733 | -0.540517763 |
| ENSDART00000058685 | zfpm2a | -0.080421638 | -0.032548615 | 0.512175645 | 0.757319343 |
| ENSDART00000058706 | fosaa | -1.574082598 | -1.225695241 | -1.240461281 | -1.154909276 |
| ENSDART00000058736 | grm4 | -0.475368296 | -0.757934708 | -0.299132614 | 0.099413724 |
| ENSDART00000058737 | cdc42l | 0.49085446 | 0.397672444 | 0.316872224 | -0.053147981 |
| ENSDART00000058773 | rgs16 | -0.222100312 | -0.210502322 | -0.307458084 | -0.014937374 |
| ENSDART00000058774 | havcr1 | 1.378230425 | 0.803624411 | 0.332422006 | 0.11017803 |
| ENSDART00000058785 | fam210ab | -0.074078069 | -0.284264661 | -0.891770658 | -0.393378695 |
| ENSDART00000058789 | qdpra | 0.319429593 | 0.277303758 | -0.201013706 | -0.346583237 |
| ENSDART00000058829 | scrt1b | -0.466509454 | -0.564612343 | -0.054278048 | 0.250358448 |
| ENSDART00000058843 | krcp | 1.066051129 | 1.030166402 | 0.671597771 | 0.484235506 |
| ENSDART00000058876 | kpnb3 | 0.233004274 | 0.111806623 | 0.257484325 | 0.090678098 |
| ENSDART00000058877 | rap2ab | -0.786170126 | -0.835369537 | -0.021239814 | 0.42494962 |
| ENSDART00000058936 | scamp5b | -0.278094691 | -0.317619737 | -0.403892079 | -0.134676622 |
| ENSDART00000058955 | arl6ip1 | -0.023708036 | -0.021671154 | -0.307639798 | -0.304733223 |
| ENSDART00000058965 | apoeb | -0.08755429 | -0.343001804 | -0.600108968 | -0.53348823 |
| ENSDART00000059001 | C5AR1 | 0.862718576 | 0.211750471 | 0.205995335 | 0.011542683 |
| ENSDART00000059003 | rx2 | -0.299942042 | -0.32670005 | -0.113463003 | 0.138106865 |
| ENSDART00000059013 | sec61b | 0.564454151 | 0.292288284 | 0.352954607 | 0.073391744 |
| ENSDART00000059179 | nptxra | -0.572157626 | -0.495039684 | -0.356374901 | -0.015079326 |
| ENSDART00000059228 | vil1 | -0.324461409 | -0.136026611 | -0.08718702 | 0.103993243 |
| ENSDART00000059369 | phykpl | 0.561796588 | 0.786946988 | 0.37392556 | 0.380622564 |
| ENSDART00000059402 | n6amt2 | 0.433448701 | 0.441036405 | 0.122223886 | -0.017461279 |
| ENSDART00000059425 | CU570684.1 | 3.082478322 | 1.756945445 | 2.993403253 | 2.236315825 |
| ENSDART00000059446 | znf385b | -0.260906147 | -0.315001382 | -0.161772411 | -0.110285684 |
| ENSDART00000059476 | psmg1 | 0.316645081 | 0.323743917 | 0.365248072 | -0.009289235 |
| ENSDART00000059478 | lrrc32 | -0.083635057 | -0.196945688 | -0.286292563 | -0.79945825 |
| ENSDART00000059489 | prmt8b | -0.385132635 | -0.375733065 | -0.525036565 | 0.065426538 |
| ENSDART00000059550 | lrrc51 | -1.204791639 | -0.640574252 | -1.049736423 | -1.0433072 |

| | | | | | |
|---|---|---|---|---|---|
| ENSDART00000059586 | spegb | 0.60633626 | 1.046592342 | 1.773590673 | 1.534849845 |
| ENSDART00000059619 | fkbp14 | 0.418122006 | 0.409098495 | -0.000966617 | 0.084647379 |
| ENSDART00000059631 | BX936415.1 | -0.988751505 | -1.218737364 | -0.343900834 | 0.242173034 |
| ENSDART00000059667 | wdr75 | 0.34735847 | 0.29669502 | 0.189255306 | 0.039773826 |
| ENSDART00000059732 | id1 | -0.232839225 | -0.281205165 | -0.284745791 | -0.436615034 |
| ENSDART00000059756 | ralba | 0.050810042 | 0.323679093 | 0.529596851 | 0.259207353 |
| ENSDART00000059841 | si:ch211-257p13.3 | -0.713606094 | -0.675183925 | 0.07538087 | 0.401418787 |
| ENSDART00000059869 | adra2a | -1.042718178 | -0.710015995 | -0.232479367 | 0.220270649 |
| ENSDART00000059955 | ildr1b | 0.550873694 | 0.893755801 | 0.988897575 | 0.559320969 |
| ENSDART00000059984 | deptor | -0.356805464 | -0.334367455 | -0.324987791 | -0.320690594 |
| ENSDART00000060001 | pnp6 | -0.139557872 | -0.321502493 | -0.386186078 | -0.030509602 |
| ENSDART00000060005 | rpl32 | 0.348697332 | 0.358910178 | 0.175532175 | 0.008609589 |
| ENSDART00000060015 | chka | -0.144459192 | -0.274568124 | -0.456317122 | -0.171918553 |
| ENSDART00000060049 | hspa13 | -0.254728034 | -0.327282661 | -0.28629746 | -0.272283731 |
| ENSDART00000060051 | fgf14 | -0.493332703 | -0.385375543 | -0.154886482 | -0.098910242 |
| ENSDART00000060056 | tpi1b | -0.330637537 | -0.371028005 | -0.31545087 | -0.122811105 |
| ENSDART00000060160 | calb2a | -1.240040436 | -1.237007367 | -0.232768326 | 0.397108945 |
| ENSDART00000060162 | hspb1 | 0.08558143 | -0.277493669 | -0.503626412 | -0.55244605 |
| ENSDART00000060174 | jagn1a | -0.416430339 | -0.038483688 | -0.018273384 | -0.314381472 |
| ENSDART00000060181 | zgc:114174 | 0.760260394 | 1.239485919 | 1.043412191 | 0.428827471 |
| ENSDART00000060184 | chka | -0.101617548 | -0.321325171 | -0.420229119 | -0.136541498 |
| ENSDART00000060193 | thap3 | 0.219009088 | 0.189913345 | 0.546438828 | 0.599142566 |
| ENSDART00000060251 | wdr18 | 0.292919064 | 0.371056299 | 0.311893069 | 0.087148131 |
| ENSDART00000060255 | blmh | 0.318055181 | 0.435968503 | 0.441694486 | 0.152219233 |
| ENSDART00000060259 | wnt2 | -0.011888956 | 0.307798324 | -0.019010839 | -1.701804028 |
| ENSDART00000060302 | ddb2 | -0.150084227 | 0.300264829 | -0.452176446 | -0.393222799 |
| ENSDART00000060304 | dhrs13a.3 | -0.00917183 | 0.036729289 | -0.280144118 | -0.407439325 |
| ENSDART00000060321 | RAMP1 | -0.401526014 | -0.370225897 | -0.385056936 | -0.044799362 |
| ENSDART00000060356 | dgkh | -0.649006381 | -0.810232451 | -0.629585196 | -0.212393687 |
| ENSDART00000060363 | rpl4 | 0.373964359 | 0.396853604 | 0.212646351 | -0.016195336 |
| ENSDART00000060425 | ftr53 | 0.363344395 | 0.103507635 | 1.747305289 | -0.30385669 |
| ENSDART00000060444 | rps29 | 0.369322836 | 0.397792199 | 0.13443893 | -0.022749507 |
| ENSDART00000060532 | zgc:110796 | -0.139659209 | -0.284517331 | -0.33969202 | -0.16468656 |
| ENSDART00000060561 | csdc2a | -0.084744858 | 0.609682067 | 0.574951056 | 0.285883083 |
| ENSDART00000060576 | myoz1a | -0.659756236 | 2.254391903 | 2.759775555 | 1.48758278 |
| ENSDART00000060577 | tmem33 | 0.319754096 | 0.307056024 | 0.247854771 | 0.133996431 |
| ENSDART00000060625 | lgi3 | -0.698992604 | -0.869112792 | -0.10986795 | 0.409134057 |
| ENSDART00000060702 | rmdn3 | -0.094193374 | -0.184889649 | -0.247613016 | -0.093111552 |
| ENSDART00000060710 | adgrg11 | 2.568700304 | 0.61302127 | 2.440739962 | 2.102266876 |
| ENSDART00000060714 | atp6ap1a | -0.204397719 | -0.266136404 | -0.201574338 | -0.008870212 |
| ENSDART00000060718 | taz | 0.896308653 | 4.991849973 | 1.010024269 | 2.916260354 |
| ENSDART00000060745 | uba52 | 0.32327466 | 0.351581625 | 0.155851402 | -0.01111286 |
| ENSDART00000060765 | nppb | 4.501656565 | 4.501350371 | 1.812509246 | -0.191856821 |
| ENSDART00000060766 | rab11a | 0.179968247 | 0.479437298 | 0.674000894 | 0.468733557 |
| ENSDART00000060773 | taar12b | 2.538886261 | 1.95415769 | 1.735717491 | 1.72259581 |
| ENSDART00000060812 | adcyap1b | 1.704872196 | 2.272116909 | 1.908485885 | 0.966573225 |
| ENSDART00000060865 | rasal1b | 1.120636523 | 0.769242819 | 0.458217171 | 0.392656742 |
| ENSDART00000060898 | mrps28 | -0.371192118 | -0.496136281 | -0.011483012 | -0.046106248 |
| ENSDART00000060910 | pimr138 | 0.646921071 | 1.324373168 | 1.239639246 | 0.98486272 |
| ENSDART00000060919 | qars | 0.348349072 | 0.542908344 | 0.574420452 | 0.306598142 |
| ENSDART00000060938 | snx9b | 0.471862308 | 0.584797813 | 0.101490869 | -0.223444858 |

| | | | | | |
|---|---|---|---|---|---|
| ENSDART00000060946 | sgsm1b | -0.173061984 | -0.355540157 | -0.215401885 | -0.063998625 |
| ENSDART00000060949 | zfpm1 | 0.286528567 | 0.561608339 | 0.375954127 | 0.217874539 |
| ENSDART00000061000 | bbs2 | -0.085680202 | -0.169473458 | -0.253361227 | -0.025608691 |
| ENSDART00000061001 | gnb2l1 | 0.316385721 | 0.345297586 | 0.431111492 | 0.12899342 |
| ENSDART00000061007 | mt2 | -0.024087588 | -0.437804286 | -0.890429121 | -0.686083783 |
| ENSDART00000061106 | bhlhe41 | 0.043056173 | -0.423617996 | -0.754959095 | -0.246095493 |
| ENSDART00000061117 | rrbp1b | -0.094086695 | 0.04423759 | -0.324207933 | -0.60750175 |
| ENSDART00000061141 | cep85l | -0.248281327 | -0.136664261 | 0.313311334 | 0.132594592 |
| ENSDART00000061149 | TUBB4A (1 of many) | 0.971780187 | 0.499035917 | 0.545475118 | 0.962590028 |
| ENSDART00000061156 | raph1a | 0.024271858 | 0.223289034 | 0.290402674 | 0.224778316 |
| ENSDART00000061196 | | 0.755433358 | 1.498415514 | 1.365853078 | 0.846252469 |
| ENSDART00000061261 | cx43 | -0.064274202 | -0.132171143 | -0.392515504 | -0.502400185 |
| ENSDART00000061265 | rnf141 | -0.12379903 | -0.215585747 | -0.342006011 | -0.212187704 |
| ENSDART00000061417 | si:ch211-245h14.1 | 0.537591116 | 0.459433616 | 1.387404638 | -0.014689392 |
| ENSDART00000061435 | hsbp1b | -0.02310974 | -0.083751559 | -0.317854229 | -0.256774332 |
| ENSDART00000061470 | mtss1la | -0.342228031 | 0.053961203 | 0.113929451 | 0.018495568 |
| ENSDART00000061497 | OLFM4 (1 of many) | -0.269705239 | -0.443597761 | -0.460075072 | -0.13609298 |
| ENSDART00000061499 | cxcr4b | 1.134597457 | 1.176774484 | 0.975229504 | 0.34365956 |
| ENSDART00000061523 | il17a/f3 | 0.228114616 | 0.421937628 | 0.621201213 | 0.769032572 |
| ENSDART00000061555 | si:ch211-63o20.7 | 1.506216579 | 0.813490393 | 0.865397342 | 0.102228691 |
| ENSDART00000061633 | zgc:171971 | -0.243215545 | -0.195809688 | -0.289637739 | -0.171793375 |
| ENSDART00000061653 | pebp1 | -0.450366142 | -0.339729586 | -0.205596712 | 0.038256249 |
| ENSDART00000061736 | si:dkey-4e7.3 | 0.80946555 | 0.660586871 | 0.14634027 | 0.03058403 |
| ENSDART00000061745 | inpp4ab | -0.337306716 | -0.94773917 | -0.831022148 | -0.394471236 |
| ENSDART00000061886 | sema3ab | -0.067887191 | -0.37217703 | -0.138233187 | 0.034639157 |
| ENSDART00000061926 | stx11b.2 | 3.447906519 | 2.671571656 | 2.785952697 | 1.680276674 |
| ENSDART00000061955 | myl13 | 0.400602979 | 3.991209963 | 4.290518336 | 2.830575983 |
| ENSDART00000062003 | efnb3b | -0.314139927 | -0.178367777 | 0.04557076 | 0.250001858 |
| ENSDART00000062066 | si:dkey-177p2.6 | -0.250914133 | -0.251912097 | -0.422547135 | -0.291399654 |
| ENSDART00000062073 | BCR (1 of many) | 0.11359461 | -0.080987241 | -0.284063713 | -0.288348423 |
| ENSDART00000062143 | zgc:77650 | -0.109887255 | -0.133378465 | -0.376117658 | -0.331935654 |
| ENSDART00000062150 | zgc:77752 | -0.401367403 | -0.150322453 | -0.162634546 | -0.096457711 |
| ENSDART00000062181 | RALGDS | -0.386705371 | -0.180287867 | -0.473841885 | -0.479084169 |
| ENSDART00000062185 | rab40b | -0.04766047 | 0.014828604 | -0.304246935 | -0.082657522 |
| ENSDART00000062220 | gstt1a | 1.233095877 | 1.492566267 | 1.464760691 | 0.555034442 |
| ENSDART00000062229 | p2rx7 | 0.280693894 | 0.946609864 | 0.955265868 | 0.573409065 |
| ENSDART00000062257 | slc39a1 | 0.740781667 | 0.522946282 | 0.350404642 | 0.245154327 |
| ENSDART00000062360 | nup205 | 0.380838378 | 0.174863772 | 0.543980624 | 0.366347355 |
| ENSDART00000062383 | ywhaqa | -0.013489826 | 0.266644821 | 0.309888274 | 0.200982907 |
| ENSDART00000062402 | tpd52l1 | -0.180007689 | -0.341056198 | -0.772074089 | -0.368132956 |
| ENSDART00000062403 | tmem9 | -0.368514764 | -0.231056659 | -0.311288279 | -0.066743785 |
| ENSDART00000062518 | gstr | -0.173225499 | -0.241367808 | -0.232576695 | -0.339957813 |
| ENSDART00000062551 | cyp51 | 0.018772193 | 0.850105191 | 1.080881624 | 0.554137375 |
| ENSDART00000062552 | wtap | -0.338047264 | -0.227200913 | -0.06961558 | -0.130824995 |
| ENSDART00000062556 | sod2 | -0.183508685 | -0.287491323 | -0.424543766 | -0.237833299 |
| ENSDART00000062560 | zgc:77784 | 0.130172136 | 0.171187774 | 0.614861949 | 0.368542156 |
| ENSDART00000062576 | thyn1 | 0.902996379 | 0.741810201 | 0.792519989 | 0.219218083 |
| ENSDART00000062587 | klf2a | 0.563534905 | 1.202186524 | 1.977026531 | 0.968508443 |
| ENSDART00000062603 | cadm1b | -0.182938767 | -0.538377364 | -0.267052947 | -0.164493851 |
| ENSDART00000062633 | s1pr1 | -0.300458433 | -0.0458986 | -0.017104561 | -0.199608488 |
| ENSDART00000062671 | tuba8l | 1.253108099 | 0.545680562 | 0.42270127 | -0.336104805 |

| | | | | | |
|---|---|---|---|---|---|
| ENSDART00000062697 | gfra2a | -0.450482933 | -0.554254218 | -0.069449306 | -0.130685043 |
| ENSDART00000062704 | plaa | 0.222653649 | 0.333324552 | 0.491597491 | 0.251933193 |
| ENSDART00000062727 | stx6 | -0.088840172 | -0.113781331 | -0.247667952 | -0.166950515 |
| ENSDART00000062736 | coasy | -0.31852882 | -0.407402551 | -0.404610538 | -0.370618301 |
| ENSDART00000062761 | cnstb | -0.361465575 | -0.117860324 | 0.271149791 | 0.383194937 |
| ENSDART00000062845 | mmp9 | 2.08287012 | 1.917473556 | 1.298502519 | -0.06470518 |
| ENSDART00000062850 | agps | 0.085764498 | 0.319290218 | 0.455652831 | 0.366743535 |
| ENSDART00000062874 | atp1b3b | -0.415562805 | -0.319287363 | -0.078895256 | 0.137689897 |
| ENSDART00000062887 | disp2 | -0.441631902 | -0.296469901 | 0.444242177 | 0.781646932 |
| ENSDART00000062908 | rpl7l1 | 0.414752526 | 0.336818546 | 0.137726222 | -0.109457063 |
| ENSDART00000062931 | abracl | 0.878200021 | 0.73529721 | 0.710676136 | 0.289919909 |
| ENSDART00000062935 | heca | -0.371152131 | -0.569989768 | -0.691744059 | -0.51577268 |
| ENSDART00000062983 | rpl10a | 0.388498934 | 0.402298329 | 0.17374506 | -0.059011317 |
| ENSDART00000063008 | mfng | 0.785568878 | 0.314558937 | 0.02546364 | -0.36422945 |
| ENSDART00000063071 | dgcr2 | -0.097084868 | -0.122785849 | -0.277264164 | -0.131322908 |
| ENSDART00000063081 | cyp2ad3 | 0.42839208 | 0.705423845 | 0.252750259 | 0.171232166 |
| ENSDART00000063107 | cyp2p7 | 0.910849098 | 1.165275545 | 0.616791402 | -0.023701278 |
| ENSDART00000063151 | napga | 0.319825484 | 0.237345685 | 0.317095517 | 0.164414652 |
| ENSDART00000063251 | ctsz | 1.117383701 | 0.530773826 | 0.448185866 | 0.142841944 |
| ENSDART00000063337 | cdca8 | 2.552618125 | 1.809029687 | 2.257850367 | 1.156797598 |
| ENSDART00000063357 | ccnb1 | 1.564145903 | 1.350801576 | 0.91228963 | 0.49209964 |
| ENSDART00000063359 | ucp3 | 0.449516929 | 0.816201403 | 1.070983817 | 0.155240874 |
| ENSDART00000063418 | nsun5 | 0.551107756 | 0.480901258 | 0.442024494 | 0.196344317 |
| ENSDART00000063478 | nfil3-2 | -0.375816946 | 0.376110137 | 0.458578211 | 0.007201456 |
| ENSDART00000063551 | ppm1e | -0.4375512 | -0.67599947 | -0.28655858 | -0.160579724 |
| ENSDART00000063564 | nmu | -0.490206998 | -0.305049777 | -0.445855788 | -0.16147485 |
| ENSDART00000063625 | gpx3 | 0.093413749 | 0.050792899 | -0.431389541 | -0.288399492 |
| ENSDART00000063648 | nck2b | -0.186136113 | -0.294499599 | 0.095653682 | 0.163593769 |
| ENSDART00000063703 | si:dkey-71h2.2 | -0.547517928 | -0.696070326 | -0.446988451 | -0.222080286 |
| ENSDART00000063704 | crip3 | -0.295432006 | -0.264072284 | -0.393391872 | -0.087537117 |
| ENSDART00000063706 | fndc4a | -0.898991439 | -1.194204845 | -0.829294946 | -0.104966589 |
| ENSDART00000063714 | rapgef6 | -0.363328977 | -0.303671651 | 0.266966568 | 0.457046047 |
| ENSDART00000063725 | xkr6b | -0.377201403 | -0.64365216 | 0.065079126 | -0.045638513 |
| ENSDART00000063764 | si:dkey-5n18.1 | 1.420288455 | 0.932190706 | 0.716992196 | -0.355341421 |
| ENSDART00000063779 | efhd1 | -0.527832136 | -0.620416333 | -0.244685479 | -0.230757866 |
| ENSDART00000063781 | gpr55a | 3.100202316 | 2.329994693 | 2.002857636 | 2.112952219 |
| ENSDART00000063783 | itm2ca | -0.362486731 | -0.282712517 | -0.07258841 | 0.153576522 |
| ENSDART00000063786 | cab39 | -0.24368294 | -0.258313334 | -0.267466597 | -0.001027932 |
| ENSDART00000063804 | wu:fj39g12 | -1.205265642 | -1.354971991 | -1.517691105 | -1.318916155 |
| ENSDART00000063816 | kcnk3a | -0.460164643 | -0.659361375 | -0.48206205 | -0.175491486 |
| ENSDART00000063817 | ndufb11 | -0.172664849 | -0.275426505 | -0.329160032 | -0.102461652 |
| ENSDART00000063825 | sprn | -0.451586162 | -0.272184034 | -0.06406637 | 0.092302795 |
| ENSDART00000063832 | rbbp8 | -0.238022634 | -0.227071096 | 0.03579094 | -0.764989117 |
| ENSDART00000063835 | otx5 | -0.207695123 | -0.226833273 | -0.417236259 | -0.126675877 |
| ENSDART00000063870 | rpl11 | 0.4298733 | 0.509956948 | 0.251365855 | 0.019450433 |
| ENSDART00000063874 | vamp4 | 0.706049434 | 0.688342777 | 0.412447235 | 0.06985511 |
| ENSDART00000063912 | jun | 1.166146477 | 1.429328618 | 1.29995447 | 0.480560375 |
| ENSDART00000063938 | mast1a | 0.148358744 | 0.555318149 | 1.032634027 | 1.008672267 |
| ENSDART00000063944 | tmem30ab | -0.052321591 | -0.186777496 | -0.351757869 | -0.167369044 |
| ENSDART00000063950 | psmc1b | 0.214901771 | 0.338782243 | 0.27453656 | 0.103637979 |
| ENSDART00000063953 | zgc:65997 | 0.11562047 | 0.484476273 | -0.11592626 | -0.024456549 |

| | | | | | |
|---|---|---|---|---|---|
| ENSDART00000064012 | ca4a | -0.265180291 | -0.250252223 | -0.144355238 | 0.018302985 |
| ENSDART00000064017 | rapgef1a | 0.181570205 | 0.344922552 | 0.212193731 | 0.084768062 |
| ENSDART00000064032 | eif4ebp1 | 0.426193835 | 0.395196044 | 0.352513449 | 0.114800609 |
| ENSDART00000064067 | ehbp1 | 0.439358276 | 0.535882532 | 0.636938582 | 0.35957066 |
| ENSDART00000064111 | faub | 0.024926044 | -0.061766338 | -0.402186419 | -0.29870419 |
| ENSDART00000064112 | glrx5 | -0.15774683 | -0.2561987 | -0.313570229 | -0.138985623 |
| ENSDART00000064113 | abt1 | 0.598506286 | 0.454995795 | 0.386684419 | 0.116646594 |
| ENSDART00000064130 | BEGAIN | -0.586123435 | -0.609324776 | -0.400772811 | -0.09647247 |
| ENSDART00000064241 | nrxn3a | -0.593329637 | -0.86602502 | -0.45366796 | -0.193651725 |
| ENSDART00000064311 | arhgdia | 0.49745023 | 0.520698759 | 0.529544756 | 0.127344369 |
| ENSDART00000064375 | tmem244 | -0.163876969 | -0.255273585 | -0.331911996 | -0.04221405 |
| ENSDART00000064376 | sod1 | -0.248477917 | -0.212122084 | -0.427881094 | -0.390078569 |
| ENSDART00000064403 | nptnb | -0.282191976 | -0.43563203 | -0.351497687 | -0.147485108 |
| ENSDART00000064462 | psma6l | 0.604857346 | 0.425092318 | 0.840213456 | -0.035624292 |
| ENSDART00000064468 | crmp1 | -0.244499065 | -0.053767261 | 0.158838362 | 0.344619203 |
| ENSDART00000064509 | stmn4l | 2.57043941 | 3.576220537 | 3.295095816 | 2.31833518 |
| ENSDART00000064511 | il17a/f1 | 1.319538931 | 1.072672417 | 0.769667562 | 1.083526716 |
| ENSDART00000064581 | kcnip3b | -1.323762559 | -2.171156238 | -0.883181569 | 0.0425137 |
| ENSDART00000064657 | stx11a | 1.187636483 | 0.704469765 | 0.33181163 | -0.135694961 |
| ENSDART00000064662 | rassf2b | -0.5330742 | -0.904631137 | -0.740838414 | -0.317057253 |
| ENSDART00000064666 | prnpb | 1.397742128 | 1.699664466 | 1.642053833 | 0.642245778 |
| ENSDART00000064672 | | -1.010331454 | -0.756315191 | -0.126164495 | 0.278123254 |
| ENSDART00000064700 | fuca2 | 0.436466876 | 0.137195427 | -0.005101419 | -0.171105064 |
| ENSDART00000064738 | atpif1b | -0.12884804 | -0.286001583 | -0.530012586 | -0.266598201 |
| ENSDART00000064739 | rpl13a | 0.411158528 | 0.410041852 | 0.123606951 | -0.078504495 |
| ENSDART00000064789 | txn | 1.421742139 | 1.973125 | 1.39538042 | 0.274434823 |
| ENSDART00000064798 | aspn | 0.856747155 | 1.499777714 | 1.699781764 | 0.582997672 |
| ENSDART00000064805 | cenpp | 0.292664259 | 0.48497191 | 0.107661064 | -0.056951786 |
| ENSDART00000064826 | mov10a | 0.329642907 | 0.141820564 | 0.774895549 | -0.030140661 |
| ENSDART00000064833 | mafaa | -0.957052225 | -1.143383392 | -0.379735148 | 0.067354966 |
| ENSDART00000064842 | padi2 | -0.110475283 | -0.200123181 | -0.394420172 | -0.195593685 |
| ENSDART00000064860 | rbms1a | -0.125610759 | 0.088275426 | 0.5443086 | 0.313591986 |
| ENSDART00000064866 | prkab1a | -0.370484412 | -0.353103565 | -0.349388814 | -0.328698623 |
| ENSDART00000064878 | gxylt2 | -0.088947943 | -0.180342387 | -0.380560197 | -0.388684923 |
| ENSDART00000064902 | ssbp4 | -0.349245142 | -0.241492171 | 0.147485085 | 0.214650497 |
| ENSDART00000064913 | fto | -0.098234146 | -0.428383355 | -0.257568512 | -0.239748235 |
| ENSDART00000064968 | rasgef1bb | 0.10103678 | 0.334291276 | 0.155368522 | -0.094945106 |
| ENSDART00000065057 | itgb7 | 1.200094562 | 0.885337917 | 0.497721229 | 0.006866552 |
| ENSDART00000065097 | dpysl3 | -0.247117182 | 0.32883927 | 0.696704704 | 0.442557487 |
| ENSDART00000065132 | zgc:171740 | -0.488981736 | -0.378281723 | 0.139236952 | 0.2855891 |
| ENSDART00000065143 | unc119b | -0.086346577 | -0.282642438 | -0.651558775 | -0.265879719 |
| ENSDART00000065159 | zgc:158291 | -1.165646756 | -0.784542684 | 0.443310678 | 0.751149417 |
| ENSDART00000065183 | cldn2 | 0.624880574 | 0.284479914 | -0.382379938 | -0.379228488 |
| ENSDART00000065208 | nop16 | 0.539941841 | 0.465464987 | 0.330311237 | 0.01578578 |
| ENSDART00000065228 | csmd1a | -0.138553555 | -0.429039185 | -0.109478646 | 0.118637902 |
| ENSDART00000065264 | cdca5 | 0.52809349 | 0.795495163 | 0.547641137 | 0.341339334 |
| ENSDART00000065337 | kif20a | 1.518648505 | 1.334799108 | 0.992824147 | 0.572892998 |
| ENSDART00000065356 | desmb | 0.665915236 | 0.890084527 | 0.800282347 | 0.503228927 |
| ENSDART00000065361 | etv5b | -0.645686026 | -0.347502384 | -0.293939032 | -0.317696675 |
| ENSDART00000065366 | st6gal1 | -0.520593433 | -0.711298232 | -0.60540202 | -0.072634403 |
| ENSDART00000065372 | kcnj3b | -0.605628976 | -0.778191304 | -0.104344748 | 0.360074992 |

| | | | | | |
|---|---|---|---|---|---|
| ENSDART00000065373 | eef1b2 | 0.250630086 | 0.293877839 | 0.677481758 | 0.365569434 |
| ENSDART00000065380 | camk1ga | -0.057459289 | -0.334433278 | -0.359469575 | -0.058443268 |
| ENSDART00000065397 | fkbp2 | 0.018660881 | -0.129009813 | -0.324870267 | -0.155859325 |
| ENSDART00000065420 | pacs1a | -0.146340142 | -0.310676632 | -0.118017371 | -0.142794888 |
| ENSDART00000065467 | dedd1 | -0.00326722 | -0.100306912 | -0.35251663 | -0.041277003 |
| ENSDART00000065495 | emp2 | 0.342641107 | 0.530182619 | 0.397727804 | 0.205643264 |
| ENSDART00000065500 | abcc4 | 0.484659612 | 0.306343643 | -0.287697763 | -0.373355455 |
| ENSDART00000065507 | plppr2b | -0.22717931 | -0.353326082 | -0.20008884 | 0.028481334 |
| ENSDART00000065551 | zak | 1.359635484 | 2.260289477 | 1.621487921 | 0.736191985 |
| ENSDART00000065563 | ccdc90b | 0.430098539 | 0.770932327 | 0.278586215 | 0.111905478 |
| ENSDART00000065567 | guca1d | -0.53307102 | -0.365098729 | -0.535522544 | -0.275955769 |
| ENSDART00000065599 | cadm1a | -0.54681119 | -0.778847768 | -0.187580903 | 0.116530415 |
| ENSDART00000065600 | sc5d | -0.12560715 | 0.619273704 | 0.802648489 | 0.437253012 |
| ENSDART00000065664 | dusp4 | -0.466715295 | -0.312689089 | -0.30035434 | -0.446313481 |
| ENSDART00000065674 | fybb | 0.68375018 | 0.334314613 | 0.079534024 | -0.242430054 |
| ENSDART00000065728 | nrsn1 | 0.207141455 | 0.540557286 | 0.398847193 | 0.186515655 |
| ENSDART00000065755 | gpn3 | -0.172157475 | -0.155442383 | -0.381057864 | -0.19097226 |
| ENSDART00000065805 | tspan10 | 0.90782785 | 0.406843364 | -0.022479662 | -0.576522773 |
| ENSDART00000065807 | kctd13 | 0.261042411 | 0.352189581 | 0.50608049 | 0.336398415 |
| ENSDART00000065817 | pou5f3 | 1.443769101 | 1.494882152 | 0.928756649 | 0.660724256 |
| ENSDART00000065818 | fut7 | 0.972435403 | 0.738351 | 0.320621995 | -0.576532021 |
| ENSDART00000065853 | dhrs3b | -0.214859587 | -0.196724114 | -0.346466706 | -0.463603398 |
| ENSDART00000065929 | hs6st3b | -0.878348455 | -0.942691234 | -0.255367675 | 0.253670478 |
| ENSDART00000066177 | tuba2 | -0.341980556 | 0.290211936 | 0.695639518 | 0.704284087 |
| ENSDART00000066192 | glra2 | -0.5075286 | -0.700662657 | -0.376893165 | 0.421840665 |
| ENSDART00000066198 | rab9a | 0.076150809 | 0.228460008 | 0.322859875 | 0.206383991 |
| ENSDART00000066230 | ARL3 (1 of many) | -0.707746447 | -0.419158458 | -0.489675877 | -0.206340441 |
| ENSDART00000066256 | vti1a | -0.147124368 | -0.216073091 | -0.331785573 | -0.195609352 |
| ENSDART00000066259 | kcnk1a | -0.289802216 | -0.227947241 | -0.67177 | -0.392395516 |
| ENSDART00000066269 | arl4d | -0.431926082 | -0.429953919 | -0.294676093 | -0.224713874 |
| ENSDART00000066288 | spata20 | -0.009504348 | -0.162892766 | -0.35319246 | -0.173028944 |
| ENSDART00000066290 | UTS2R | -0.720663714 | -0.982454784 | -0.512077429 | -0.545080013 |
| ENSDART00000066294 | cdk5r1b | -0.511562763 | -0.495240798 | 0.085364893 | 0.188592346 |
| ENSDART00000066372 | id4 | -0.719733282 | -0.518590484 | 0.441847928 | 0.655214602 |
| ENSDART00000066373 | vdac1 | -0.316380085 | -0.269111014 | -0.266508034 | -0.031245506 |
| ENSDART00000066380 | ca7 | -0.236035328 | -0.017625204 | -0.765902599 | -0.026455411 |
| ENSDART00000066382 | aqp8a.1 | -0.164652843 | -0.325293072 | -0.493634593 | -0.109118528 |
| ENSDART00000066385 | hbz | 0.819612283 | 1.064127771 | 1.5494854 | 0.440895582 |
| ENSDART00000066386 | shisa9a | -0.462438855 | -0.347473048 | -0.310604934 | -0.078066291 |
| ENSDART00000066389 | tmem184ba | -0.204152722 | -0.333415893 | -0.147371729 | -0.097737372 |
| ENSDART00000066391 | csnk1e | 0.01172114 | 0.595315231 | 0.792117229 | 0.433182971 |
| ENSDART00000066411 | dlgap5 | 1.041688171 | 0.895823709 | 0.540002804 | 0.33349173 |
| ENSDART00000066471 | adam8b | 1.018895271 | 1.190253553 | 1.122506289 | 0.517972783 |
| ENSDART00000066477 | dkk1b | -0.674554346 | -1.187574562 | -0.676297256 | -1.09065776 |
| ENSDART00000066506 | cox6b1 | 0.122510297 | 0.115648463 | 0.536625345 | 0.628910558 |
| ENSDART00000066590 | rdh12l | 0.806755691 | 0.842269039 | 0.848807771 | 0.120108035 |
| ENSDART00000066623 | st8sia3 | -0.182339116 | -0.349623634 | -0.081530336 | -0.065322773 |
| ENSDART00000066625 | smpx | 0.108625243 | 0.797810146 | 1.548262684 | 1.15135764 |
| ENSDART00000066655 | mybl1 | -0.255810306 | -0.487313899 | -0.72749049 | -0.668391364 |
| ENSDART00000066703 | rdh10a | -0.287660525 | -0.158056129 | -0.356819377 | -0.262167994 |
| ENSDART00000066733 | gpr22b | -0.9723222 | -1.512791135 | -0.598895258 | -0.577277225 |

| | | | | | |
|---|---|---|---|---|---|
| ENSDART00000066760 | cct5 | 0.325276039 | 0.436851681 | 0.379245954 | 0.163513221 |
| ENSDART00000066765 | bmi1a | 3.316489868 | 2.947911885 | 2.359801458 | 2.244842944 |
| ENSDART00000066778 | acad11 | 0.796145744 | 0.566674022 | 0.216041695 | -0.190350168 |
| ENSDART00000066784 | fam49bb | 0.049871707 | -0.095277519 | -0.294897796 | -0.151467176 |
| ENSDART00000066794 | otulina | 0.468513579 | 0.440716075 | 0.351949179 | 0.239625569 |
| ENSDART00000066839 | slc35g2b | -0.671846523 | -0.750415444 | -0.309110396 | 0.04424778 |
| ENSDART00000066895 | rassf8b | -0.275713104 | -0.148423362 | -0.148246743 | -0.166873018 |
| ENSDART00000066896 | syt1a | -0.255336633 | -0.349305824 | -0.330024338 | -0.178637525 |
| ENSDART00000066963 | atp6v1f | -0.089791813 | -0.169571399 | -0.304106913 | -0.234245815 |
| ENSDART00000066975 | impdh1b | 0.195914466 | 0.339716801 | 0.56808711 | 0.477613078 |
| ENSDART00000066997 | dram1 | 0.801491354 | 0.536766251 | 0.146394385 | -0.606440034 |
| ENSDART00000066999 | ccdc53 | 0.300412133 | 0.290286375 | 0.140411091 | 0.00256916 |
| ENSDART00000067005 | bcat1 | -0.064708558 | -0.22197643 | -0.301840187 | -0.097404482 |
| ENSDART00000067053 | vta1 | 0.2430424 | 0.457630564 | 0.367763165 | 0.149685321 |
| ENSDART00000067059 | fam19a5b | -0.282093441 | -0.382553277 | -0.381033174 | -0.505286252 |
| ENSDART00000067066 | parp6b | -0.443166413 | 0.101161009 | 0.402103535 | 0.516387314 |
| ENSDART00000067078 | plekhg5a | -0.147306721 | -0.274518319 | -0.555343847 | -0.255866929 |
| ENSDART00000067082 | clta | -0.242881724 | -0.298079508 | -0.220247727 | -0.021275929 |
| ENSDART00000067147 | ANKRD50 | 0.122102324 | 0.471943767 | 0.546358337 | 0.270036712 |
| ENSDART00000067168 | pdzrn4 | 0.330865082 | 1.464493171 | 1.433744497 | 0.793719225 |
| ENSDART00000067190 | tspan9b | -0.605960388 | -0.476716761 | -0.251844268 | -0.266924971 |
| ENSDART00000067193 | adm2a | 0.4356689 | 0.184728608 | -0.002686414 | -1.106691874 |
| ENSDART00000067211 | gpr37l1b | -0.390917116 | -0.36533905 | -0.206436073 | -0.113142613 |
| ENSDART00000067239 | guca1g | -0.979413995 | -1.042352228 | -1.597708164 | -1.275542792 |
| ENSDART00000067258 | syt10 | -0.290010761 | -3.233857576 | -0.084108756 | -0.40009033 |
| ENSDART00000067312 | si:dkey-106n21.1 | 0.584969753 | 0.102146183 | -0.031661382 | 0.069325035 |
| ENSDART00000067324 | mfge8b | -0.041376155 | -0.13672935 | -0.391535511 | -0.353629136 |
| ENSDART00000067327 | abhd2b | 0.159640353 | 0.698443956 | 0.611964225 | 0.750739323 |
| ENSDART00000067362 | cart2 | -0.749355761 | -0.384853037 | -0.719287733 | -0.625188951 |
| ENSDART00000067427 | yeats4 | 0.136322543 | 0.363843408 | 0.318943581 | 0.037728114 |
| ENSDART00000067434 | nudt4b | -0.510387585 | -0.468486348 | -0.329729048 | -0.412952232 |
| ENSDART00000067446 | slc38a4 | -0.251589327 | -0.331078479 | -0.606665517 | -0.538910759 |
| ENSDART00000067448 | acat1 | -0.217922513 | -0.251881532 | -0.396031362 | -0.209577052 |
| ENSDART00000067461 | si:ch211-152c2.3 | -0.474479692 | -0.343258906 | -0.010850784 | 0.158497108 |
| ENSDART00000067478 | pkp3a | 0.648319477 | 0.895152082 | 2.33162898 | 0.940793267 |
| ENSDART00000067500 | si:dkey-280e21.3 | 0.421024272 | 0.822626751 | 0.979586691 | 0.632877181 |
| ENSDART00000067510 | crabp1a | -0.186487239 | -0.41656224 | -0.606281006 | -0.420918895 |
| ENSDART00000067512 | psma4 | 0.210284594 | 0.22504456 | 0.324917099 | 0.119970251 |
| ENSDART00000067514 | rbpms2a | -1.175610868 | -1.156235961 | 0.045281157 | 0.593799851 |
| ENSDART00000067531 | syn2a | -0.50370548 | -0.414636803 | -0.227904446 | 0.132370635 |
| ENSDART00000067537 | elovl6l | 0.404046488 | 1.053095893 | 1.924314262 | 1.109723769 |
| ENSDART00000067542 | kcnk10b | 2.685681832 | 3.478119742 | 3.471523284 | 2.49042133 |
| ENSDART00000067594 | cst14b.1 | 1.003408293 | 1.132156972 | 0.601894798 | 0.102871452 |
| ENSDART00000067599 | angptl2a | 1.671461404 | 1.166310219 | 0.762115684 | 0.407368061 |
| ENSDART00000067637 | dstyk | -0.298992266 | -0.18939574 | -0.18546883 | -0.220617575 |
| ENSDART00000067678 | zgc:110339 | 0.949255799 | 0.562708107 | 0.125395868 | -0.585210787 |
| ENSDART00000067733 | zgc:77838 | -0.192172111 | -0.379046905 | -0.170230593 | -0.131538824 |
| ENSDART00000067741 | cacng6b | -2.498204907 | -1.377375045 | -1.107680653 | -1.191335449 |
| ENSDART00000067762 | MYO1D | -0.207118869 | -0.403579734 | -0.286614596 | -0.314212256 |
| ENSDART00000067764 | stk17a | -0.185636748 | -0.198053553 | -0.515379898 | -0.253038095 |
| ENSDART00000067776 | rab10 | 0.291742266 | 0.421774457 | 0.459760968 | 0.189497347 |

| | | | | | |
|---|---|---|---|---|---|
| ENSDART00000073405 | zgc:173552 | 0.695539535 | 0.568218274 | 1.113411469 | 0.383673074 |
| ENSDART00000073452 | si:ch211-113a14.12 | 0.801955974 | 0.847093718 | 0.855214481 | 0.503313711 |
| ENSDART00000073462 | rplp0 | 0.408607572 | 0.593255771 | 0.47280039 | 0.165347026 |
| ENSDART00000073500 | ptprz1a | -0.320567664 | -0.32912405 | -0.255947625 | -0.47902768 |
| ENSDART00000073511 | hyal6 | -0.14022603 | -0.303368807 | -0.413603406 | -0.552708783 |
| ENSDART00000073564 | tes | 0.818084026 | 1.011033966 | 0.902633694 | 0.089277909 |
| ENSDART00000073583 | islr2 | 0.313523936 | 1.12325114 | 1.332162091 | 1.179316934 |
| ENSDART00000073588 | kcnj11 | -0.465815764 | -0.671487298 | -0.425150531 | -0.293255153 |
| ENSDART00000073617 | opn4xa | -0.377358091 | -0.474333233 | -0.655013246 | -0.269116892 |
| ENSDART00000073634 | slc30a5 | -0.005479497 | -0.126675199 | -0.24346742 | -0.137800332 |
| ENSDART00000073694 | smu1b | -0.023572641 | -0.696487436 | -0.934479138 | -0.393819626 |
| ENSDART00000073705 | abcf1 | 0.284347738 | 0.522430424 | 0.46970431 | 0.327069748 |
| ENSDART00000073726 | cav2 | -0.135895339 | 0.080426407 | -0.366803726 | -0.546607126 |
| ENSDART00000073735 | rrad | 0.475668516 | 0.908053908 | 0.772751604 | 0.279273357 |
| ENSDART00000073846 | si:ch211-122f10.4 | 0.620706192 | 0.26696454 | 0.219379439 | -0.011733355 |
| ENSDART00000073861 | gabarapb | -0.042475311 | -0.021227012 | -0.297506188 | -0.271766658 |
| ENSDART00000073903 | crtc3 | -0.232649231 | -0.295606215 | -0.176321155 | -0.096709861 |
| ENSDART00000073919 | kcnc1b | -0.758440974 | -0.784989596 | -0.33092052 | 0.322314215 |
| ENSDART00000073932 | | 0.121194935 | -0.376988949 | -0.788016576 | -0.265683315 |
| ENSDART00000073936 | acvr1bb | -0.291123202 | -0.390021531 | -0.349875902 | -0.253586872 |
| ENSDART00000073950 | olfm1a | -0.852827208 | -0.587563633 | 0.013637392 | 0.435090193 |
| ENSDART00000073970 | uap1 | 0.430311379 | 0.939111864 | 0.868311508 | 0.206630788 |
| ENSDART00000073981 | eif2s1b | 0.229006291 | 0.299741922 | 0.510349465 | 0.185458194 |
| ENSDART00000073985 | rbfox2 | -0.063885036 | 0.156691707 | 0.429046907 | 0.332565675 |
| ENSDART00000074010 | ubald1b | 0.486188172 | 0.523878559 | 0.246068485 | 0.014830353 |
| ENSDART00000074036 | rcvrna | -0.553376367 | -0.06531313 | -0.11758335 | -0.07626787 |
| ENSDART00000074070 | aff2 | -0.074830038 | -0.283904996 | -0.159789643 | -0.147467133 |
| ENSDART00000074099 | cabp2b | -0.664167129 | -0.641383476 | -0.676380934 | -0.590070868 |
| ENSDART00000074100 | osgn1 | 0.255332714 | 0.013078065 | -0.613186841 | -0.487873824 |
| ENSDART00000074117 | aspa | -0.032383232 | -0.304124913 | -0.397629458 | -0.358898864 |
| ENSDART00000074161 | slc4a2b | 0.481685785 | 0.057728735 | -0.200942778 | -0.247347753 |
| ENSDART00000074212 | scrib | 0.155467238 | 0.226272134 | 0.329350211 | 0.126740521 |
| ENSDART00000074317 | GSK3B (1 of many) | 0.32408852 | 0.570538304 | 0.615748631 | 0.310967098 |
| ENSDART00000074362 | pcdh18b | -0.440881831 | -0.447646282 | -0.227758863 | 0.176890552 |
| ENSDART00000074380 | tsga10 | -0.358572407 | -0.32335115 | -0.582555062 | -0.41305108 |
| ENSDART00000074384 | stx4 | 0.365157378 | 0.335603422 | 0.150714004 | -0.050227712 |
| ENSDART00000074400 | tia1 | 0.181056891 | 0.310121029 | 0.46787956 | 0.221267203 |
| ENSDART00000074438 | cenpi | 0.547415003 | 0.708397465 | 0.401862169 | 0.054393549 |
| ENSDART00000074458 | ptpmt1 | -0.138560892 | -0.279060294 | -0.535667209 | -0.518833125 |
| ENSDART00000074543 | hs3st4 | -0.453389419 | -0.572006435 | -0.310659668 | 0.002760951 |
| ENSDART00000074609 | suclg1 | -0.204210404 | -0.293159601 | -0.171956898 | 0.002164837 |
| ENSDART00000074678 | chrnb3a | -0.683036397 | -0.585209438 | -0.069659384 | 0.207417172 |
| ENSDART00000074685 | glrbb | -0.387255204 | -0.421576892 | -0.370807969 | -0.094075713 |
| ENSDART00000074689 | eif5b | 0.118742051 | 0.204941132 | 0.292461428 | 0.069777195 |
| ENSDART00000074698 | opn3 | -0.589287895 | -0.235981184 | 0.224725485 | -0.032745726 |
| ENSDART00000074718 | spire1b | -0.446651583 | -0.478750814 | -0.155393912 | -0.07838085 |
| ENSDART00000074786 | ctso | 0.33157854 | 0.154941794 | -0.070268612 | -0.012066877 |
| ENSDART00000074838 | kcnk3b | -0.808453975 | -0.995140212 | -0.30053552 | -0.689829672 |
| ENSDART00000074924 | mbnl1 | 0.32889023 | 0.277718787 | 0.642145523 | 0.211541009 |
| ENSDART00000074936 | gabrr2a | -0.375623148 | -0.534637844 | -0.431940406 | -0.359950744 |
| ENSDART00000074950 | slc25a28 | -0.022495638 | -0.038026223 | -0.243484898 | -0.120098609 |

| | | | | | |
|---|---|---|---|---|---|
| ENSDART00000074959 | slc35f1 | -0.294905966 | -0.396956867 | -0.318533983 | -0.137876179 |
| ENSDART00000074960 | cd22 | 0.815723271 | 0.646940316 | 0.539427268 | -0.232061943 |
| ENSDART00000074979 | rnft2 | 0.06664962 | 0.449659345 | 0.588779493 | 0.279626887 |
| ENSDART00000074997 | CU302436.3 | -0.857583363 | -0.990723788 | -0.187854306 | 0.199234775 |
| ENSDART00000075009 | eif2s2 | 0.324008789 | 0.419532633 | 0.338652657 | 0.07614456 |
| ENSDART00000075028 | rps11 | 0.42100484 | 0.441088802 | 0.128015907 | -0.080449898 |
| ENSDART00000075039 | gosr2 | 0.009731507 | 0.24829488 | 0.212269525 | -0.090945666 |
| ENSDART00000075070 | hsf2 | -0.066635095 | -0.276496416 | -0.474097609 | -0.186498896 |
| ENSDART00000075092 | pkib | -0.08506216 | -0.264513554 | -0.401500984 | -0.212860417 |
| ENSDART00000075112 | clvs2 | 0.165980729 | 0.346379383 | 0.75563827 | 0.535875063 |
| ENSDART00000075116 | si:dkey-148a17.6 | 0.841894804 | 0.387365318 | 0.191323415 | -0.035438886 |
| ENSDART00000075123 | pcp4a | -0.598752351 | -0.632528063 | -0.40265776 | -0.022235336 |
| ENSDART00000075129 | lrrc47 | 0.095574868 | 0.246973338 | 0.327055852 | 0.135309168 |
| ENSDART00000075150 | bmp4 | -0.321654174 | -0.192558055 | -0.354985644 | -0.520317573 |
| ENSDART00000075172 | cttnbp2nla | -0.399030632 | -0.078691774 | -0.193018514 | -0.275655386 |
| ENSDART00000075184 | snx1a | 0.267188596 | 0.126701463 | 0.120486891 | -0.033463537 |
| ENSDART00000075187 | pdzd11 | 1.031277997 | 1.70516213 | 1.574957324 | 1.135180778 |
| ENSDART00000075223 | cox7a2a | -0.206130471 | -0.297851957 | -0.422850723 | -0.097231315 |
| ENSDART00000075260 | inab | -0.558694355 | 0.275313434 | 1.165237079 | 1.111612471 |
| ENSDART00000075262 | cad | 0.41294438 | 0.156970425 | 0.074105835 | 0.04769287 |
| ENSDART00000075278 | atp1b4 | -0.758712842 | -0.43098592 | -0.240547748 | -0.077197984 |
| ENSDART00000075286 | slc2a15b | 1.023992215 | 0.671159354 | 0.170031714 | -0.73573625 |
| ENSDART00000075299 | zgc:153911 | 1.082638325 | 2.093301655 | 1.816780233 | 1.137984438 |
| ENSDART00000075320 | nampta | -0.083138724 | -0.057980927 | 0.268248216 | -0.062095237 |
| ENSDART00000075331 | insm1b | 0.180818368 | 0.558805202 | 0.852549514 | 0.856981121 |
| ENSDART00000075340 | eef1a1b | -0.66677551 | -0.473919347 | -0.344351157 | 0.106841151 |
| ENSDART00000075351 | zgc:112285 | 0.930923442 | 1.110746674 | 0.609701543 | 0.79082813 |
| ENSDART00000075398 | cilp | 1.006018993 | 0.43809517 | 0.394851989 | 0.084259634 |
| ENSDART00000075400 | eif3jb | 0.284407943 | 0.11040548 | -0.086691777 | -0.1231685 |
| ENSDART00000075421 | sord | -0.008165085 | -0.251228216 | -0.437538687 | -0.226988953 |
| ENSDART00000075465 | mylpfa | -0.627780558 | 3.258441537 | 4.096966032 | 2.542734101 |
| ENSDART00000075491 | pop5 | 0.444559136 | 0.7451635 | 0.571282976 | 0.537107281 |
| ENSDART00000075495 | rpl23 | 0.390673252 | 0.341184618 | 0.264108815 | -0.005521244 |
| ENSDART00000075499 | si:dkey-283b1.7 | -0.753135255 | -0.256638623 | -0.866666345 | -0.691414461 |
| ENSDART00000075510 | ngb | -0.534714702 | -0.310443771 | -0.511019459 | -0.235764625 |
| ENSDART00000075513 | aqp9b | -0.181865878 | -0.358464178 | -0.54013493 | -0.322928595 |
| ENSDART00000075519 | aldh1a2 | -0.136704985 | -0.161016939 | -0.528203419 | -0.420522124 |
| ENSDART00000075551 | adhfe1 | 0.155892684 | -0.065430004 | -0.440766413 | -0.267456669 |
| ENSDART00000075663 | cracr2b | 0.080855847 | 0.057965052 | -0.927246946 | -0.842015409 |
| ENSDART00000075743 | eprs | 0.181617497 | 0.208060281 | 0.357036503 | 0.248273691 |
| ENSDART00000075749 | ppp2r2ca | -0.492014929 | -0.268923214 | 0.328067214 | 0.425314748 |
| ENSDART00000075808 | apbb3 | -0.025817957 | 0.161567226 | 0.465828121 | 0.45985711 |
| ENSDART00000075889 | trub1 | 0.429771719 | 0.538562347 | 0.550227428 | 0.249034481 |
| ENSDART00000075902 | klhl43 | 1.602362948 | 3.319039146 | 3.627754288 | 3.174379945 |
| ENSDART00000075903 | crlf3 | -0.351458679 | -0.119437595 | 0.002747455 | -0.299343819 |
| ENSDART00000075918 | pcmtd2 | -0.427264647 | -0.257111837 | -0.149394946 | 0.062644737 |
| ENSDART00000075927 | rad50 | 0.535614901 | 0.654138999 | 0.420227838 | 0.362493118 |
| ENSDART00000075935 | vtnb | -0.377853158 | -0.359520721 | -0.399371834 | -0.233041975 |
| ENSDART00000075940 | mtnr1ba | -0.423445838 | -1.139676296 | -0.754844062 | -0.657118397 |
| ENSDART00000075974 | ism2b | -0.752954546 | -0.582959852 | -0.548810475 | -0.377147389 |
| ENSDART00000075993 | crtc1b | -0.280354395 | -0.577500657 | -0.194615785 | -0.207111517 |

| | | | | | |
|---|---|---|---|---|---|
| ENSDART00000076004 | tmem62 | -0.08960985 | 0.130455024 | 0.308433272 | 0.432472028 |
| ENSDART00000076009 | hadhab | 0.274869752 | 0.225585159 | 0.168828199 | 0.114580965 |
| ENSDART00000076030 | fbl | 0.33280558 | 0.192834191 | 0.116054515 | 0.005015438 |
| ENSDART00000076066 | lin37 | -0.271630989 | -0.17922011 | -0.496068109 | -0.578879729 |
| ENSDART00000076082 | fetub | 1.519329398 | 0.709563768 | 0.228067904 | 0.727615025 |
| ENSDART00000076083 | cdc42ep2 | -0.005463107 | 0.666894523 | 0.327900404 | 0.083241845 |
| ENSDART00000076157 | rab24 | 0.15781613 | 0.389364959 | 0.281581252 | 0.130911228 |
| ENSDART00000076160 | mustn1a | 1.455614739 | 1.360374319 | 1.112954668 | 0.977466313 |
| ENSDART00000076161 | hoxb5b | 2.652791903 | 3.351280664 | 2.618897772 | 0.864721852 |
| ENSDART00000076215 | CR735102.1 | -0.360410997 | -0.514821634 | -0.137026978 | -0.257442941 |
| ENSDART00000076238 | rbm41 | -0.118558432 | -0.076666908 | -0.261877568 | -0.209896281 |
| ENSDART00000076333 | pgk1 | -0.277163813 | -0.302384935 | -0.297237462 | -0.087607565 |
| ENSDART00000076399 | nat10 | 0.304377518 | 0.292344426 | 0.160797024 | -0.000572845 |
| ENSDART00000076417 | cdkn1bb | 0.050164178 | 0.266815944 | 0.205910633 | -0.014851012 |
| ENSDART00000076423 | sobpa | -0.461445043 | -0.18351314 | 0.23019679 | 0.473568961 |
| ENSDART00000076483 | zgc:77151 | 0.249827469 | 0.539125177 | 0.736975565 | 0.447972658 |
| ENSDART00000076496 | stk33 | -1.544458095 | -0.103215934 | -2.939032674 | -0.830432799 |
| ENSDART00000076502 | rerglb | 0.481268892 | 0.304392452 | -0.423368722 | -1.032950672 |
| ENSDART00000076506 | wisp1a | 0.075916017 | 0.908100257 | 1.435223818 | 0.542654062 |
| ENSDART00000076518 | sla1 | 0.79177335 | 0.326196979 | 0.029071124 | -0.326507352 |
| ENSDART00000076554 | si:zfos-464b6.2 | 0.5351721 | 0.505133501 | 0.952773319 | 0.026200547 |
| ENSDART00000076571 | rtn1a | 0.31833938 | 1.022959786 | 0.981624341 | 0.706878795 |
| ENSDART00000076574 | rtn1a | -0.132166015 | 0.374408359 | 0.606191338 | 0.476970779 |
| ENSDART00000076600 | rpe65c | 1.983711129 | 2.589375601 | 1.902404689 | 1.435704768 |
| ENSDART00000076636 | fzd2 | 0.563199333 | 0.424474998 | 0.171148118 | -0.301884624 |
| ENSDART00000076648 | clip3 | -0.386141796 | -0.253171682 | 0.072987847 | 0.103065331 |
| ENSDART00000076786 | clec19a | 0.036182435 | 0.047143104 | -0.366633197 | -0.838257146 |
| ENSDART00000076815 | | 2.456176486 | 3.15180058 | 2.959048559 | 2.414794476 |
| ENSDART00000076925 | ITPRIPL2 | 0.939030216 | 0.593569322 | 0.667833004 | -0.090802943 |
| ENSDART00000076929 | prkg2 | -0.322610383 | -0.221985796 | -0.523406711 | -0.304941411 |
| ENSDART00000076938 | pogza | 0.098614093 | 0.259761887 | 0.429750084 | 0.224276825 |
| ENSDART00000076946 | PDE4DIP | -0.575491599 | -0.478988093 | -0.210265098 | -0.157443712 |
| ENSDART00000076997 | lmo4b | -0.24164689 | -0.249534589 | -0.123100185 | -0.051398466 |
| ENSDART00000077008 | alox5ap | 0.122223757 | -0.082901221 | -0.550940123 | -0.70030614 |
| ENSDART00000077047 | btr09 | 0.55744156 | 0.619797073 | 1.272153942 | -0.048548492 |
| ENSDART00000077080 | PTP4A3 (1 of many) | 0.077833133 | 0.004217905 | -0.463336754 | -0.666773375 |
| ENSDART00000077087 | id3 | 0.047452456 | 0.423524517 | 0.285867988 | -0.296627095 |
| ENSDART00000077157 | six3b | -0.273810969 | -0.17356253 | 0.033591817 | 0.098241195 |
| ENSDART00000077185 | dgat1b | 0.79138422 | 0.60487817 | 0.419420549 | -0.346047138 |
| ENSDART00000077197 | tmsb | 1.972985822 | 3.598018383 | 3.544381625 | 2.894174576 |
| ENSDART00000077215 | ppp2r5b | 0.185911454 | 0.229029828 | 0.615811771 | 0.611674376 |
| ENSDART00000077216 | astn1 | -0.280184115 | -0.378831195 | -0.109389357 | 0.054556021 |
| ENSDART00000077222 | ldlrad4b | -0.327819788 | -0.475734059 | -0.025614815 | 0.028486052 |
| ENSDART00000077259 | ebna1bp2 | 0.420936188 | 0.344892308 | 0.188849483 | 0.008127131 |
| ENSDART00000077386 | prss16 | 0.714972584 | 0.32483027 | 0.071612523 | -0.240875958 |
| ENSDART00000077406 | cdh27 | -0.746226456 | -0.635554214 | -0.974185768 | -0.424771371 |
| ENSDART00000077411 | cxcl12b | -0.236947042 | -0.2488008 | -0.593903488 | -0.559818867 |
| ENSDART00000077418 | ctsba | 1.550723066 | 1.033464808 | 0.97579905 | 0.475659932 |
| ENSDART00000077420 | dtnbp1a | 0.285624415 | 0.193705819 | 0.051855359 | -0.043275565 |
| ENSDART00000077445 | pim3 | -0.017745548 | 0.425023548 | -0.018053873 | 0.016773558 |
| ENSDART00000077459 | smyd2a | 0.109333535 | 0.326579335 | 0.423001533 | 0.365696401 |

| | | | | | |
|---|---|---|---|---|---|
| ENSDART00000077462 | slc6a9 | -0.157728483 | -0.388200876 | -0.444419631 | -0.487528264 |
| ENSDART00000077476 | prox1a | -0.192871011 | -0.360465718 | -0.389825586 | -0.065307919 |
| ENSDART00000077484 | zhx2a | 0.117201305 | 0.356185793 | 0.299966576 | 0.024327323 |
| ENSDART00000077511 | ccr9a | 1.513300322 | 1.48326526 | 0.981085195 | 0.082724583 |
| ENSDART00000077538 | kpna2 | 3.376600286 | 3.247024938 | 2.263140647 | 0.702563762 |
| ENSDART00000077539 | tuba1c | -0.211130957 | 0.180769664 | 0.840172721 | 0.834422327 |
| ENSDART00000077545 | slc7a7 | 1.304066715 | 0.45830859 | 0.136139102 | 0.056436424 |
| ENSDART00000077582 | pitpnm3 | -0.256916567 | -0.419244886 | -0.147473339 | -0.136698838 |
| ENSDART00000077619 | b3gat1b | -0.266663197 | -0.223974588 | -0.393683459 | -0.125347427 |
| ENSDART00000077635 | si:ch211-103n10.5 | 0.712451842 | 0.944946244 | 1.075959446 | -0.046690849 |
| ENSDART00000077662 | myl6 | 0.570434868 | 0.203215664 | -0.213132641 | -0.482895856 |
| ENSDART00000077664 | atp2b1a | -0.283300071 | -0.323367416 | -0.218529441 | -0.07251725 |
| ENSDART00000077707 | llph | 0.612018484 | 0.617804437 | 0.382050706 | 0.042759152 |
| ENSDART00000077715 | si:dkey-25e12.3 | 1.07190149 | 0.953044464 | 0.701373655 | 0.117439727 |
| ENSDART00000077724 | gnb5b | -0.216593155 | -0.279473354 | -0.485943487 | -0.092161444 |
| ENSDART00000077783 | hmox2a | 0.068185569 | 0.120478633 | -0.326668325 | -0.242069945 |
| ENSDART00000077805 | gria2a | -0.63140941 | -0.839709321 | -0.362303249 | -0.053629448 |
| ENSDART00000077809 | cyp26c1 | -0.796873586 | -0.763455549 | -0.70044496 | -0.582141171 |
| ENSDART00000077823 | lrit3a | -0.481897882 | -0.48516269 | -0.481656788 | -0.455604814 |
| ENSDART00000077834 | rps27.2 | 0.365621497 | 0.39961171 | 0.113670294 | -0.081169937 |
| ENSDART00000077836 | mmp20b | -0.526034453 | -0.625086024 | -0.345110188 | -0.027134291 |
| ENSDART00000077839 | atf7b | -0.232485852 | -0.196508033 | -0.391624214 | -0.206346003 |
| ENSDART00000077868 | si:ch211-212k18.7 | 1.166604646 | 0.658609076 | 0.758814708 | -0.422210365 |
| ENSDART00000077895 | rnf7 | -0.220302586 | -0.252047578 | -0.316298741 | -0.185486087 |
| ENSDART00000077898 | grk7b | -0.582244813 | -0.358943116 | -0.402813205 | 0.000611673 |
| ENSDART00000077951 | pcolce2b | -0.119540012 | -0.232226227 | -0.447813206 | -1.286575617 |
| ENSDART00000077998 | cgna | 0.243171731 | 0.392675733 | 0.184983165 | 0.011488923 |
| ENSDART00000078014 | poldip2 | 0.012504595 | -0.191194187 | -0.45353852 | -0.221096745 |
| ENSDART00000078018 | ruvbl2 | 0.355681647 | 0.304270351 | 0.334910735 | 0.092604809 |
| ENSDART00000078024 | crk | -0.263270816 | -0.200000869 | -0.157496558 | -0.11349953 |
| ENSDART00000078033 | GPT (1 of many) | -0.491126295 | -0.482727977 | -0.247455346 | -0.055451846 |
| ENSDART00000078037 | aclyb | -0.084913872 | -0.379038696 | -0.658479637 | -0.284307838 |
| ENSDART00000078053 | tbcb | 0.21865146 | 0.428683048 | 0.432538452 | 0.172774297 |
| ENSDART00000078072 | akap12b | 2.975823835 | 3.328543122 | 3.653370868 | 2.600734247 |
| ENSDART00000078079 | pcnxl2 | -0.518593139 | -0.325199094 | 0.020646332 | 0.403616211 |
| ENSDART00000078115 | sdhda | -0.602799581 | -0.148637611 | 0.027660422 | 0.175615096 |
| ENSDART00000078137 | ankrd54 | 0.051281727 | 0.23783078 | 0.382868701 | 0.32135773 |
| ENSDART00000078148 | smc1a | -3.399991944 | -0.069804776 | -1.248223526 | -0.569589215 |
| ENSDART00000078156 | srm | 0.127220715 | 0.22303682 | 0.345313656 | 0.085476496 |
| ENSDART00000078181 | SLC3A2 (1 of many) | -0.23816401 | -0.122270906 | -0.56307324 | -1.283472558 |
| ENSDART00000078187 | foxo4 | -0.056772502 | -0.136599728 | -0.36455832 | -0.08188156 |
| ENSDART00000078192 | cnpy4 | 0.065580495 | 0.06657667 | -0.1806698 | -0.737660818 |
| ENSDART00000078202 | phka2 | -0.128194416 | -0.258622872 | -0.071035307 | 0.033146819 |
| ENSDART00000078226 | mtnr1bb | -0.285812097 | -0.289316068 | -0.599481549 | -0.276891324 |
| ENSDART00000078232 | cdh10a | -0.495858439 | -0.491434243 | -0.331011431 | 0.033804718 |
| ENSDART00000078249 | kcnc3a | -0.908848026 | -1.288714191 | -0.644595688 | 0.010477706 |
| ENSDART00000078256 | dopey2 | -0.056439943 | -0.30342326 | -0.284521914 | -0.1564256 |
| ENSDART00000078266 | rsl1d1 | 0.351365949 | 0.392552672 | 0.175928217 | -0.070353732 |
| ENSDART00000078277 | msmo1 | 0.087631699 | 1.109674669 | 1.47165822 | 0.852850391 |
| ENSDART00000078283 | tex10 | 0.186689066 | 0.248875437 | 0.343572259 | 0.154977674 |
| ENSDART00000078304 | lzic | 0.119250866 | 0.44378754 | 0.377529349 | 0.128075007 |

| | | | | | |
|---|---|---|---|---|---|
| ENSDART00000078305 | zswim5 | -0.126655043 | 0.076098312 | 0.245252624 | 0.223511201 |
| ENSDART00000078306 | arhgef2 | 0.429610468 | 0.807274797 | 0.854804837 | 0.533632211 |
| ENSDART00000078311 | zgc:154093 | -0.170397489 | -0.310689773 | -0.421136702 | -0.191616401 |
| ENSDART00000078316 | nipa2 | 0.247262018 | 0.436255727 | 0.4433152 | 0.003440759 |
| ENSDART00000078325 | slc16a8 | 2.36549078 | 0.51323664 | -0.815797835 | -1.896302174 |
| ENSDART00000078334 | celsr3 | 0.469496099 | 0.512562938 | 0.74195357 | 0.599858827 |
| ENSDART00000078336 | klc3 | -0.15725421 | -0.267140148 | -0.672772286 | -0.331888789 |
| ENSDART00000078352 | tspan14 | 0.358298965 | 0.408510067 | 0.238594355 | -0.012844708 |
| ENSDART00000078412 | rps8a | 0.416303691 | 0.456350868 | 0.419741526 | 0.122907452 |
| ENSDART00000078438 | | 0.22834743 | 0.583517385 | 0.218267131 | 0.08549295 |
| ENSDART00000078449 | itih6 | -0.23325998 | -0.04381884 | -0.494745518 | -1.555031348 |
| ENSDART00000078491 | mov10b.2 | 0.18890489 | 0.151487185 | 1.352539948 | -0.244846261 |
| ENSDART00000078494 | l3mbtl2 | 1.071567754 | 1.041266281 | 0.642956308 | 0.316295602 |
| ENSDART00000078522 | eef1g | 0.462831941 | 0.651007925 | 0.567355432 | 0.209607784 |
| ENSDART00000078529 | kin | -1.437736057 | -0.295663012 | -0.493580296 | -4.734617185 |
| ENSDART00000078533 | kcnd3 | -0.299800885 | -0.32147822 | -0.292734142 | -0.136341479 |
| ENSDART00000078535 | gtf2h5 | 0.788173845 | 1.2490535 | 0.921738211 | 0.911998292 |
| ENSDART00000078543 | syt11b | 0.975892204 | 1.6420197 | 1.925407845 | 1.338618886 |
| ENSDART00000078561 | SPTBN4 | -0.239897718 | -0.33102434 | 0.293671093 | 0.445086753 |
| ENSDART00000078594 | tyrp1b | 0.662132615 | 0.257868212 | -0.309681889 | -0.980797447 |
| ENSDART00000078596 | hspd1 | 0.512169965 | 0.358816938 | 0.201006014 | 0.20720259 |
| ENSDART00000078611 | jac2 | 3.404844888 | 3.429750736 | 2.921387124 | 1.113891823 |
| ENSDART00000078630 | nme7 | 0.176767548 | 0.421131107 | 0.35919241 | 0.2453347 |
| ENSDART00000078634 | eif5a2 | 0.493280465 | 0.485481598 | 0.418411017 | 0.051860815 |
| ENSDART00000078642 | vps37b | 0.195137227 | 0.217160121 | 0.327187369 | 0.074828156 |
| ENSDART00000078647 | si:ch211-201h21.5 | 1.148616031 | 0.689283012 | 0.40532037 | 0.049879214 |
| ENSDART00000078652 | camk2g2 | -0.278366483 | -0.285453125 | -0.22783561 | -0.037372706 |
| ENSDART00000078694 | TMEM179B (1 of many) | 0.551851512 | 0.299347782 | 0.026749154 | -0.302916803 |
| ENSDART00000078708 | zgc:162952 | -0.249018938 | -0.407117986 | -0.404487357 | -0.328471526 |
| ENSDART00000078723 | dixdc1a | 0.324551864 | 0.395166532 | 0.459267183 | 0.274985584 |
| ENSDART00000078771 | SBK1 | -0.390774004 | 0.13684565 | 0.923517231 | 0.784697628 |
| ENSDART00000078792 | ahcyl1 | 0.058902937 | 0.000957907 | -0.341276653 | -0.275160426 |
| ENSDART00000078795 | ahcyl1 | -0.360916144 | -0.460734164 | -0.414619806 | -0.201451999 |
| ENSDART00000078838 | rab3aa | -0.225658115 | -0.303506805 | -0.374028659 | -0.138563861 |
| ENSDART00000078843 | SLC22A7 (1 of many) | -0.18908726 | -0.089234859 | -0.388754878 | -1.485871042 |
| ENSDART00000078856 | dlg3 | -0.31041276 | -0.361138693 | -0.191257864 | 0.041004354 |
| ENSDART00000078858 | si:ch73-86n18.1 | 0.106229924 | -0.155735751 | -0.58770094 | -0.854457693 |
| ENSDART00000078866 | ifi30 | 1.574791131 | 0.896478525 | 0.373397245 | -0.020268978 |
| ENSDART00000078877 | sncga | -0.059233705 | 0.307631997 | 0.27775793 | 0.434885626 |
| ENSDART00000078908 | usp1 | -0.163356677 | -0.229514103 | -0.329514373 | -0.251997637 |
| ENSDART00000078916 | smim7 | -0.217464923 | -0.109941617 | -0.350272436 | -0.25934653 |
| ENSDART00000078949 | afap1l1b | 1.061182692 | 1.814819842 | 1.643265844 | 0.877712611 |
| ENSDART00000078953 | afap1l1b | 0.880715307 | 1.833091326 | 1.686561879 | 0.852421188 |
| ENSDART00000078982 | vat1 | 0.139786944 | 0.289776545 | 0.306149934 | -0.025026234 |
| ENSDART00000079019 | spsb1 | 0.176345745 | 0.177736786 | 0.29021019 | 0.163667209 |
| ENSDART00000079035 | rap1gap | -0.57187807 | -0.701331328 | -0.299987324 | -0.289210087 |
| ENSDART00000079046 | cpamd8 | -0.122112474 | -0.085213564 | -0.191819718 | -0.872911162 |
| ENSDART00000079050 | nutf2 | 0.337010875 | 0.380281227 | 0.325036311 | 0.156803484 |
| ENSDART00000079092 | si:dkey-261i16.5 | -0.526813533 | -0.563690076 | -0.369398724 | 0.061616577 |
| ENSDART00000079104 | ndufs6 | -0.146008815 | -0.377028896 | -0.443194118 | -0.327081615 |
| ENSDART00000079112 | calca | -0.439449486 | -0.32190484 | -0.601830596 | -0.728154291 |

| | | | | | |
|---|---|---|---|---|---|
| ENSDART00000079138 | ptenb | -0.337433521 | -0.277401842 | -0.145004066 | -0.034401548 |
| ENSDART00000079144 | ptenb | -0.303152595 | -0.238217531 | -0.001915727 | -0.031063744 |
| ENSDART00000079173 | lepr | 1.173965688 | 1.129708824 | 0.920018098 | 0.6672678 |
| ENSDART00000079202 | abcb8 | -0.111089791 | -0.249333097 | -0.256171214 | -0.098593711 |
| ENSDART00000079222 | chaf1b | 1.242408116 | 0.820706616 | 0.642546621 | 0.111426229 |
| ENSDART00000079235 | cd99l2 | -0.167082345 | 0.448795434 | 0.91020128 | 0.864265064 |
| ENSDART00000079283 | tmeff1b | 0.060962109 | 0.679659218 | 0.971148806 | 0.712213835 |
| ENSDART00000079310 | manba | 0.495609624 | 0.109254813 | 0.005925635 | -0.12409296 |
| ENSDART00000079341 | plch1 | -0.457888719 | -0.43639001 | -0.345543016 | -0.155469708 |
| ENSDART00000079364 | snapc2 | -0.030032776 | -0.185097862 | -0.560264081 | -0.400163167 |
| ENSDART00000079397 | ryk | -0.107777448 | 0.087264815 | -0.11559458 | -0.531439469 |
| ENSDART00000079431 | rtn2b | 0.025228511 | -0.123165337 | -0.319794883 | -0.05842346 |
| ENSDART00000079443 | GABRA2 (1 of many) | -0.623319426 | -0.542833879 | -0.157704672 | 0.138036393 |
| ENSDART00000079454 | vamp2 | -0.197978353 | -0.366012835 | -0.326357547 | -0.1060827 |
| ENSDART00000079497 | emg1 | 0.201493898 | 0.380905959 | 0.206415684 | 0.036355374 |
| ENSDART00000079518 | dnajc1 | 0.043021447 | -0.028027956 | -0.123370167 | -0.287218054 |
| ENSDART00000079528 | ilk | 0.338564211 | 0.350049719 | 0.346485507 | 0.032055843 |
| ENSDART00000079536 | nek7 | 0.300813238 | 0.263862785 | 0.126120434 | -0.061245789 |
| ENSDART00000079549 | tpte | 0.809167451 | 0.638911968 | 0.260848622 | -0.250321254 |
| ENSDART00000079559 | ifit16 | 0.716638095 | 0.325468397 | 2.311988055 | -0.283531995 |
| ENSDART00000079563 | fas | 0.786193523 | 0.954741877 | 1.62170637 | 0.201092535 |
| ENSDART00000079591 | dpm1 | -0.022510061 | -0.080956932 | -0.275574355 | -0.162526678 |
| ENSDART00000079597 | vps36 | -0.205754956 | -0.373190829 | -0.317459289 | -0.134792851 |
| ENSDART00000079629 | ppm1nb | -0.279926594 | -0.345062767 | -0.462173038 | -0.171806172 |
| ENSDART00000079656 | tvp23b | -0.013268416 | -0.155098046 | -0.309054243 | -0.268412353 |
| ENSDART00000079686 | zmp:0000001103 | 0.143973249 | -0.889323937 | -0.759747318 | -0.487193938 |
| ENSDART00000079695 | zwilch | 1.159509442 | 0.995361025 | 0.861360163 | 0.105564769 |
| ENSDART00000079711 | slc25a1a | 0.025053389 | 0.916416559 | 1.283110285 | 0.925187833 |
| ENSDART00000079716 | hpf1 | -0.113759988 | -0.292000236 | -0.336217157 | -0.156517495 |
| ENSDART00000079778 | ifit8 | 1.432186723 | 0.097505141 | 4.166007884 | 0.96250098 |
| ENSDART00000079803 | nmt1b | 0.899927422 | 1.156945789 | 1.408075075 | 0.459974747 |
| ENSDART00000079810 | si:dkey-222b8.4 | -0.076664011 | -0.231174428 | -0.530241167 | -0.088965105 |
| ENSDART00000079840 | rorca | -0.65562057 | -0.064052995 | 0.161319682 | 0.103804992 |
| ENSDART00000079843 | nceh1b.2 | 0.690825571 | 0.217907848 | -0.110601617 | -0.426325911 |
| ENSDART00000079866 | slc30a9 | -0.061810697 | -0.211371943 | -0.307919743 | -0.172076689 |
| ENSDART00000079879 | si:dkey-91i10.3 | -0.580141681 | -0.553279281 | -0.678150475 | -0.52463511 |
| ENSDART00000079884 | alox5a | 0.143901368 | -0.192427785 | -0.334678776 | -0.585419844 |
| ENSDART00000079984 | rpl22l1 | 0.678692893 | 0.584241724 | 0.331994973 | -0.011632328 |
| ENSDART00000080014 | rps8b | -0.261943359 | -0.309301007 | -0.481979191 | -0.409711249 |
| ENSDART00000080016 | ppp3ccb | -0.214631103 | -0.298468254 | 0.05278016 | 0.038022829 |
| ENSDART00000080033 | SSBP2 (1 of many) | 0.123939988 | 0.461173497 | 0.600467157 | 0.3572706 |
| ENSDART00000080042 | rab33a | -0.045588719 | 0.341497657 | 0.630733663 | 0.503197845 |
| ENSDART00000080064 | CR847953.1 | 0.050009503 | -0.040451977 | 0.259802571 | 0.438904414 |
| ENSDART00000080079 | slc44a5b | 0.101790496 | -0.11939972 | -0.347901034 | 0.093736244 |
| ENSDART00000080100 | slc24a2 | -0.273129774 | -0.420453101 | -0.415034704 | -0.05535122 |
| ENSDART00000080129 | ston2 | -0.17811412 | -0.063813163 | -0.270400144 | -0.092351554 |
| ENSDART00000080135 | gfpt1 | 0.380664344 | 0.49511154 | 0.544262456 | 0.337010461 |
| ENSDART00000080256 | nefma | 0.572074143 | 2.009260502 | 2.281701505 | 1.529767044 |
| ENSDART00000080289 | prrc2c | 0.060093623 | 0.047472332 | 0.306363857 | 0.189281148 |
| ENSDART00000080313 | arl4ca | -0.176267156 | -0.177242747 | -0.285935967 | -0.135129968 |
| ENSDART00000080328 | nf1a | -0.041622168 | -0.338478743 | -0.243156878 | -0.085905935 |

| | | | | | |
|---|---|---|---|---|---|
| ENSDART00000080339 | galm | 0.868912331 | 0.484945087 | -0.111697277 | -0.429352779 |
| ENSDART00000080342 | josd2 | 0.375878923 | 0.743043487 | 0.777511515 | 0.447917768 |
| ENSDART00000080351 | dhx57 | 0.296631931 | 0.310315824 | 0.343123119 | 0.20162921 |
| ENSDART00000080377 | aldoca | 2.758856293 | 1.548001605 | 2.09362575 | 1.794239524 |
| ENSDART00000080385 | SLC9A3R2 (1 of many) | -0.157518802 | -0.380020166 | -0.549234439 | -0.123708468 |
| ENSDART00000080389 | fam13a | 0.342651645 | -0.065971229 | -0.119440295 | 0.116639155 |
| ENSDART00000080414 | sbk3 | -0.999751639 | -0.617799332 | -0.160630934 | -0.723965523 |
| ENSDART00000080423 | ctsd | 0.448869492 | 0.173368574 | -0.412435712 | -0.637183839 |
| ENSDART00000080430 | gfra2b | -0.62088032 | -0.576945607 | -0.392707521 | -0.144958503 |
| ENSDART00000080465 | hells | 0.812911201 | 0.936234984 | 0.557478835 | 0.182957003 |
| ENSDART00000080486 | ywhag1 | -1.02392497 | -0.686234935 | -0.010129168 | 0.2177797 |
| ENSDART00000080523 | itgae.2 | 1.067269567 | 0.812178684 | 0.361968863 | 0.003662922 |
| ENSDART00000080549 | lyz | 1.762095695 | 1.25446765 | 1.569214818 | 0.577231805 |
| ENSDART00000080602 | map7d2b | 0.243598476 | 0.831190471 | 1.239754477 | 0.898189276 |
| ENSDART00000080628 | arpc3 | 0.270913011 | 0.099552933 | 0.079917753 | -0.036372289 |
| ENSDART00000080664 | zgc:86709 | 1.428698865 | 3.355720442 | 2.498933956 | 1.732893579 |
| ENSDART00000080673 | syt11a | -0.340996032 | -0.110491382 | 0.046613391 | 0.116186716 |
| ENSDART00000080679 | ccdc136b | -1.000484865 | -0.898166187 | -0.625061619 | -0.323992777 |
| ENSDART00000080712 | slc43a3b | 2.109023631 | 1.075213126 | 0.724428131 | -0.154263604 |
| ENSDART00000080771 | slc6a16a | -0.167322319 | -0.266319768 | -0.350765511 | -0.319541774 |
| ENSDART00000080789 | trpc4a | -0.390087102 | -0.398102567 | -0.462339329 | -0.409517574 |
| ENSDART00000080808 | six3a | -0.437705937 | -0.430575558 | -0.140593988 | -0.07114268 |
| ENSDART00000080829 | hspa14 | 0.672662829 | 0.694717457 | 0.8195912 | 0.418167344 |
| ENSDART00000080854 | stat3 | 0.856010749 | 0.444560766 | 0.379796349 | -0.326021696 |
| ENSDART00000080864 | magt1 | 0.441285986 | 0.302582629 | 0.047526992 | -0.172355248 |
| ENSDART00000080875 | b3gnt3 | 0.44585097 | 0.568958183 | 0.901392395 | 0.139618312 |
| ENSDART00000080900 | cfap57 | 2.52486974 | 2.757508228 | 2.941219811 | 1.994632033 |
| ENSDART00000080904 | sardh | 0.759429129 | 0.915608159 | 0.345777903 | -0.12435587 |
| ENSDART00000080919 | rpl36a | 0.405736348 | 0.485454355 | 0.225930563 | -0.002722727 |
| ENSDART00000080927 | snap25b | -0.418114286 | -0.385475041 | -0.102955296 | 0.076752829 |
| ENSDART00000081039 | | 2.046984335 | 2.965673996 | 2.312859948 | -1.109370855 |
| ENSDART00000081059 | rps6kb1b | 0.156747941 | 0.167680861 | 0.344534054 | 0.174989656 |
| ENSDART00000081092 | si:dkeyp-77h1.4 | -0.383994825 | 0.24121529 | 0.967657356 | 0.94823494 |
| ENSDART00000081129 | cdk15 | -0.731126626 | -0.682923022 | -0.544281357 | -0.253398129 |
| ENSDART00000081140 | CT990561.1 | -0.348057632 | -0.380510005 | -0.109063338 | 0.180807023 |
| ENSDART00000081154 | prpf18 | -0.15179636 | -0.149318949 | -0.443970585 | -0.330213785 |
| ENSDART00000081170 | cux1a | -0.092953946 | -0.2283391 | 0.138990685 | 0.296890549 |
| ENSDART00000081183 | enc3 | 0.409273368 | 0.447706974 | 0.259987961 | 0.244016846 |
| ENSDART00000081204 | acot9.1 | 0.172924803 | 0.462969941 | 0.48161826 | 0.340925129 |
| ENSDART00000081214 | selt1a | -0.31898529 | -0.284714031 | -0.513594635 | -0.412445291 |
| ENSDART00000081223 | krt5 | 2.400508858 | 4.729984226 | 4.862312842 | 5.813921541 |
| ENSDART00000081228 | abhd14a | -0.111769944 | -0.123001167 | -0.255304369 | -0.088239573 |
| ENSDART00000081272 | gcn1 | 0.168820856 | 0.100284898 | 0.363305756 | 0.2232516 |
| ENSDART00000081290 | rnaset2l | 0.733135995 | 0.630785596 | 0.575081946 | 0.019045787 |
| ENSDART00000081323 | abhd17c | -0.03998334 | 0.119899277 | 0.261541408 | 0.19734068 |
| ENSDART00000081325 | dynll1 | -0.057380796 | -0.1118203 | -0.516198015 | -0.112248637 |
| ENSDART00000081326 | prpf40a | 0.136343368 | 0.189733856 | 0.260275336 | 0.074340716 |
| ENSDART00000081338 | slc9a5 | 0.890559011 | 1.045395859 | 1.329125407 | 1.036960607 |
| ENSDART00000081343 | plk1 | 1.547304713 | 1.292689538 | 1.02080622 | 0.322979307 |
| ENSDART00000081359 | zgc:110425 | 0.523386438 | 0.743109297 | 0.819385742 | 0.340090119 |
| ENSDART00000081411 | pole | 0.480999213 | 0.458730193 | 0.174065324 | -0.07089485 |

| | | | | | |
|---|---|---|---|---|---|
| ENSDART00000081432 | sprb | -0.390584563 | -0.260880914 | -0.194039821 | -0.19414991 |
| ENSDART00000081447 | fbxo30a | 0.072897711 | 0.325740781 | 0.333966964 | 0.186898423 |
| ENSDART00000081468 | ccdc79 | 0.331555315 | 0.505423262 | 0.318137438 | -0.103372006 |
| ENSDART00000081510 | celf4 | -0.384169548 | -0.376019008 | -0.02531238 | 0.147771891 |
| ENSDART00000081546 | trim46b | 0.101540064 | 0.068781295 | 0.327282891 | 0.456457337 |
| ENSDART00000081568 | tcf19l | 0.712871286 | 0.397385898 | 0.351815803 | 0.227291981 |
| ENSDART00000081601 | cept1a | 0.150762086 | 0.296977372 | 0.445683893 | 0.275050142 |
| ENSDART00000081611 | cgnb | -0.200219306 | -0.438466186 | -1.067503957 | -0.17391972 |
| ENSDART00000081620 | vax2 | -0.290535146 | -0.478626837 | -0.10258237 | -0.009557703 |
| ENSDART00000081646 | glrx | 0.838457918 | 1.280146822 | 1.066830025 | 0.395822945 |
| ENSDART00000081761 | bin1b | -0.023096846 | -0.153451242 | -0.459181763 | -0.132136814 |
| ENSDART00000081781 | PLEKHG3 | -0.076813883 | -0.173391418 | -0.382548311 | -0.034437681 |
| ENSDART00000081794 | rasgrf2a | -0.063040785 | -0.093088949 | -0.266637358 | 0.015786604 |
| ENSDART00000081797 | sash1b | -0.368433045 | -0.365106826 | -0.357216383 | -0.178826864 |
| ENSDART00000081811 | zgc:112052 | 0.311036284 | 0.397766463 | 0.340201557 | 0.120599155 |
| ENSDART00000081832 | ptpdc1b | 0.49240902 | 0.580136174 | 0.651643292 | 0.340422756 |
| ENSDART00000081870 | pcdh2ab6 | 0.090888519 | 0.246515279 | 0.356189629 | 0.303363778 |
| ENSDART00000081926 | CU856539.1 | -0.250902601 | -0.402416425 | -0.31567263 | -0.091643419 |
| ENSDART00000081946 | zgc:112332 | -0.051059621 | 0.027246103 | -0.289131429 | -0.652053172 |
| ENSDART00000081951 | stx1b | -0.290947433 | -0.255213869 | 0.04983789 | 0.170348415 |
| ENSDART00000081966 | rtn4a | 0.42137623 | 0.811004738 | 0.55995797 | 0.082545275 |
| ENSDART00000081978 | KCNJ6 | -0.364130454 | -0.389723131 | -0.355694518 | -0.126078935 |
| ENSDART00000081985 | pim2 | -0.011194591 | -0.040743933 | -0.388705404 | -0.753111264 |
| ENSDART00000081990 | strip2 | -0.530994895 | -0.582233551 | -0.263527962 | -0.22210065 |
| ENSDART00000082011 | lim2.2 | 1.304610538 | 0.978295786 | 0.86763989 | 0.856876261 |
| ENSDART00000082012 | gsk3aa | -0.41333003 | -0.370003942 | -0.288947953 | -0.203102626 |
| ENSDART00000082050 | zgc:174904 | 1.977093547 | 1.430607855 | 1.439069833 | 0.228460651 |
| ENSDART00000082063 | fam114a1 | 0.586474654 | 0.396933414 | 0.041198489 | -0.237340023 |
| ENSDART00000082066 | atpv0e2 | -0.268294846 | -0.377237748 | -0.604078331 | -0.39998077 |
| ENSDART00000082080 | jupb | -0.499893077 | -0.526312414 | -0.258393629 | -0.100168242 |
| ENSDART00000082082 | gars | 0.395042441 | 0.664200633 | 0.717176297 | 0.392291214 |
| ENSDART00000082097 | PRSS35 (1 of many) | 0.904416504 | 0.725234092 | 1.067628772 | 0.791677307 |
| ENSDART00000082142 | EFEMP1 (1 of many) | -1.76159347 | -0.849874708 | -0.584181957 | -1.080912576 |
| ENSDART00000082151 | uqcrh | -0.212671975 | -0.358890471 | -0.337939592 | -0.281374001 |
| ENSDART00000082223 | tax1bp3 | 0.85130321 | 1.256170012 | 1.075265217 | 0.22088107 |
| ENSDART00000082264 | pxdc1b | 0.801924492 | 1.033443039 | 0.634881962 | 0.139425975 |
| ENSDART00000082301 | myrip | -0.137038516 | -0.238384017 | 0.084718693 | 0.41131665 |
| ENSDART00000082346 | tfap2a | -0.238141907 | -0.307957291 | -0.242159487 | -0.230339237 |
| ENSDART00000082368 | marco | 0.980427549 | 0.782180823 | 1.00834904 | 0.091028138 |
| ENSDART00000082434 | tgif1 | 0.822910382 | 1.229522702 | 1.441745222 | 0.912580325 |
| ENSDART00000082438 | dlgap2a | -0.366533513 | -0.540400426 | -0.257659814 | -0.037294062 |
| ENSDART00000082458 | sarnp | 0.25573821 | 0.275816383 | 0.16902753 | 0.007612146 |
| ENSDART00000082471 | mfap2 | 0.61383585 | 0.779035446 | 0.354750417 | 0.055843134 |
| ENSDART00000082517 | rab43 | 0.357940531 | 0.212689136 | -0.148032038 | -0.068856369 |
| ENSDART00000082523 | impa2 | -0.208830066 | -0.398914852 | -0.498863304 | -0.267214166 |
| ENSDART00000082604 | galnt18b | -0.214631898 | -0.351577935 | -0.246678211 | -0.218706176 |
| ENSDART00000082620 | dysf | 1.140285476 | 2.000991026 | 2.040057287 | 1.067021482 |
| ENSDART00000082622 | fsd1l | -0.046506283 | -0.340407639 | -0.41095417 | -0.201162325 |
| ENSDART00000082698 | kank4 | -0.273638259 | -0.263464734 | -0.200789629 | -0.103573896 |
| ENSDART00000082715 | camsap3 | 0.302702799 | 0.347512275 | 0.333666179 | -0.022433363 |
| ENSDART00000082745 | EMB | -0.651102521 | -0.703010006 | -0.074268357 | 0.192993872 |

| | | | | | |
|---|---|---|---|---|---|
| ENSDART00000082821 | rims4 | -0.331865117 | -0.050822604 | 0.125345094 | 0.041526523 |
| ENSDART00000082830 | KIAA0895L | 0.216380627 | 0.852873359 | 0.835551488 | 0.401334264 |
| ENSDART00000082842 | JPH3 (1 of many) | -0.540363483 | -0.227235849 | 0.34555956 | 0.43863208 |
| ENSDART00000082937 | fscn2a | -0.244556215 | -0.436229092 | -0.571250568 | -0.199107259 |
| ENSDART00000082944 | dock6 | 0.129469289 | 0.283374744 | 0.379764071 | 0.290753034 |
| ENSDART00000082983 | clip2 | 0.69105479 | 0.899482735 | 1.166942455 | 0.710683744 |
| ENSDART00000083002 | map1aa | 0.023824463 | 0.057494175 | 0.8302085 | 0.470304447 |
| ENSDART00000083010 | acad9 | 0.301381925 | 0.15641426 | -0.079322034 | -0.023998781 |
| ENSDART00000083033 | sik1 | 1.146093805 | 0.631897288 | 0.512350737 | 1.123188354 |
| ENSDART00000083040 | hs3st2 | -1.059711916 | -1.469328404 | -0.190665179 | 0.30791737 |
| ENSDART00000083063 | tal1 | 0.195096671 | 0.315620662 | 0.697094431 | 0.015122988 |
| ENSDART00000083066 | asphd2 | -0.487940941 | -0.316881347 | -0.126036033 | 0.003992278 |
| ENSDART00000083085 | mtmr14 | 0.202842573 | 0.147255325 | 0.298709035 | 0.120380705 |
| ENSDART00000083100 | | -0.514905306 | -0.64333228 | -0.306814422 | -0.178642671 |
| ENSDART00000083126 | cidec | 3.092638419 | 2.402632883 | 1.147285193 | -0.231828568 |
| ENSDART00000083212 | fscn1a | -0.172521555 | 0.403697769 | 0.865067524 | 0.635846942 |
| ENSDART00000083294 | nol6 | 0.354535252 | 0.232360208 | 0.196448331 | 0.109158776 |
| ENSDART00000083359 | sec14l8 | -0.359432343 | -0.324487355 | -0.138175947 | -0.2576341 |
| ENSDART00000083367 | prcp | -0.058674879 | -0.090308681 | -0.37460188 | -0.414832363 |
| ENSDART00000083394 | si:dkey-106c17.3 | -0.374020521 | -0.387399838 | -0.436274502 | -0.115892618 |
| ENSDART00000083407 | b4galnt4a | -0.141610106 | -0.312267637 | -0.129787335 | -0.08625756 |
| ENSDART00000083416 | gabrd | -0.516993342 | -0.41812722 | -0.415766575 | -0.292897654 |
| ENSDART00000083427 | slc25a29 | -0.338331576 | -0.636979798 | -0.361185623 | -0.191915619 |
| ENSDART00000083449 | dub | 0.532370465 | 0.813595698 | 0.783551631 | -0.001039848 |
| ENSDART00000083453 | slc32a1 | -0.393731273 | -0.378330473 | -0.304655077 | -0.16825712 |
| ENSDART00000083467 | parp8 | 0.336619688 | 0.399001121 | 0.399118225 | 0.307424215 |
| ENSDART00000083569 | oaz2b | -0.447559352 | -0.669337328 | -0.007273423 | 0.348858251 |
| ENSDART00000083572 | zgc:136864 | 0.242428375 | 0.324173418 | -0.028258929 | -0.082397484 |
| ENSDART00000083605 | tbc1d25 | 0.094218097 | 0.226970926 | 0.332123969 | 0.140604089 |
| ENSDART00000083628 | ddit3 | -0.053032847 | 0.505000026 | 0.598711637 | 0.25299651 |
| ENSDART00000083670 | CABZ01041604.1 | -0.549505735 | -0.427354636 | -0.238335089 | -0.140824262 |
| ENSDART00000083684 | pappab | -0.696202319 | -0.797866787 | -0.223755037 | -0.122490212 |
| ENSDART00000083731 | trpv1 | 1.343483137 | 1.058081357 | 0.730442906 | -0.443240884 |
| ENSDART00000083788 | CU633832.1 | -0.745373175 | -0.699527481 | -0.49975457 | -0.170614876 |
| ENSDART00000083797 | TBC1D9B | 0.273448063 | 0.469124554 | 0.581828184 | 0.40676136 |
| ENSDART00000083830 | sdc4 | 0.356242427 | 0.466156404 | 0.190102023 | -0.370187936 |
| ENSDART00000083890 | usp24 | 0.219267617 | 0.201082864 | 0.325420731 | 0.216599954 |
| ENSDART00000084007 | TULP2 | -0.184892554 | -0.234742374 | -0.492430755 | -0.167640019 |
| ENSDART00000084011 | cplx4a | -0.220118176 | -0.343953082 | -0.589002173 | -0.338126511 |
| ENSDART00000084014 | cplx1 | -0.741807214 | -0.862001833 | -0.211893904 | 0.449532765 |
| ENSDART00000084024 | sv2c | -0.354928985 | -0.65582646 | -0.477522685 | -0.220826118 |
| ENSDART00000084035 | znf532 | -0.199573234 | -0.524026601 | -0.06012509 | 0.154210784 |
| ENSDART00000084055 | fzd7a | -0.350921417 | -0.318886271 | -0.299513467 | -0.86880486 |
| ENSDART00000084069 | iqgap2 | 1.059547107 | 0.55852141 | 0.220571876 | -0.354231411 |
| ENSDART00000084119 | si:ch1073-44g3.1 | -0.384776638 | -0.202587046 | 0.01646479 | 0.134112994 |
| ENSDART00000084131 | fam160b2 | 0.036570039 | -0.082247362 | -0.429951116 | -0.30533058 |
| ENSDART00000084135 | ubtd1a | -0.22864625 | -0.383233305 | -0.947657938 | -0.688823144 |
| ENSDART00000084184 | aimp1 | 0.209084138 | 0.262090446 | 0.396284966 | 0.096096937 |
| ENSDART00000084238 | CABZ01057411.1 | -0.328182832 | -0.451030622 | -0.21027367 | 0.014791503 |
| ENSDART00000084264 | adcy2a | -0.574108823 | -0.494423944 | 0.049230433 | 0.113035845 |
| ENSDART00000084353 | tbc1d10ab | -0.174770874 | -0.349891521 | -0.619429045 | -0.244339697 |

| | | | | | |
|---|---|---|---|---|---|
| ENSDART00000084354 | cpeb3 | -0.1650884 | -0.603027317 | 0.172363697 | 0.2398868 |
| ENSDART00000084355 | zgc:165481 | -0.665978496 | -0.883055142 | -0.199024252 | -0.082238068 |
| ENSDART00000084373 | frmd4bb | 0.582631667 | 0.828834738 | 0.835563839 | 0.412330382 |
| ENSDART00000084378 | crb2a | -0.010590407 | -0.348182237 | -0.479892445 | -0.295243873 |
| ENSDART00000084381 | sybu | -0.763177411 | -0.608942869 | -0.49552138 | -0.188366829 |
| ENSDART00000084416 | ablim1a | -0.251488969 | -0.08542034 | 0.289459971 | 0.335228062 |
| ENSDART00000084417 | timm17b | -0.134748392 | -0.15578708 | -0.404232794 | -0.18557959 |
| ENSDART00000084448 | psmd11a | 0.167797415 | 0.349500372 | 0.318678727 | 0.030124317 |
| ENSDART00000084512 | pkn1a | 0.199755386 | 0.283744748 | 0.437490279 | 0.320601704 |
| ENSDART00000084517 | vcpip1 | 0.020173722 | 0.114070565 | 0.246341011 | 0.321118019 |
| ENSDART00000084530 | coro2ba | -0.558160234 | -0.686474796 | -0.264666518 | -0.204619183 |
| ENSDART00000084598 | vimp | -0.003252442 | -0.076004385 | -0.324006724 | -0.304272734 |
| ENSDART00000084714 | zranb1a | -0.285770869 | -0.28077285 | -0.104830495 | 0.014952891 |
| ENSDART00000084729 | pecam1 | 0.670854861 | 0.415048291 | 0.182847943 | 0.022819591 |
| ENSDART00000084730 | zgc:162160 | -0.758204424 | -0.82447566 | -0.164736838 | 0.207922006 |
| ENSDART00000084771 | pde9a | 0.836076081 | 1.443202889 | 1.33955445 | 1.070879094 |
| ENSDART00000084792 | prosc | -0.109541047 | -0.121094609 | -0.31852643 | -0.202300262 |
| ENSDART00000084803 | asic2 | -0.588195088 | -0.477145292 | -0.317477641 | -0.106481463 |
| ENSDART00000084806 | slc4a10b | -0.257067272 | -0.281842585 | -0.215225145 | -0.174870087 |
| ENSDART00000084819 | arhgap35b | 0.101645005 | 0.291379647 | 0.535032879 | 0.462050203 |
| ENSDART00000084861 | cish | -0.446878588 | -0.18032932 | -1.508647694 | -1.506136587 |
| ENSDART00000084890 | si:ch211-284e13.4 | -0.124688162 | -0.262617592 | 0.237393976 | 0.369141192 |
| ENSDART00000084965 | cep104 | -0.196545786 | -0.225873142 | -0.456327943 | -0.208454995 |
| ENSDART00000085121 | sdk2b | -0.222566721 | -0.365719278 | -0.147657392 | -0.012041014 |
| ENSDART00000085135 | tbl1x | 0.180884774 | 0.566948084 | 0.5040067 | 0.049402086 |
| ENSDART00000085142 | map1b | -0.213270056 | 0.210763047 | 0.526451874 | 0.312155026 |
| ENSDART00000085210 | cacna1ha | -0.413195046 | -0.405391107 | -0.236177845 | 0.006546496 |
| ENSDART00000085230 | atl1 | -0.283512177 | 0.138626411 | 0.619353197 | 0.442930265 |
| ENSDART00000085252 | pqlc3 | -0.117366098 | -0.27675705 | -0.43476147 | -0.253093536 |
| ENSDART00000085253 | mid1 | -0.357661223 | -0.330080838 | -0.002902011 | 0.078651096 |
| ENSDART00000085263 | SELENOI | 0.036305982 | 0.149651758 | 0.460720877 | 0.164646013 |
| ENSDART00000085277 | pfkmb | 0.010653815 | -0.414332737 | -0.69351419 | -0.144457554 |
| ENSDART00000085284 | pfkla | -0.114418267 | -0.338020067 | -0.177396742 | 0.008723864 |
| ENSDART00000085294 | tnfrsf9a | -0.468498688 | -0.352709139 | -0.665957795 | -1.403982945 |
| ENSDART00000085309 | dpcd | 0.675929667 | 0.665840567 | 0.458693669 | 0.16198721 |
| ENSDART00000085319 | sos2 | 0.062155314 | 0.173737983 | 0.464042198 | 0.208637802 |
| ENSDART00000085370 | kcnq5b | -0.41441932 | -0.216130879 | -0.15503054 | -0.008181026 |
| ENSDART00000085388 | bmp3 | -0.146774305 | 1.240822845 | 1.494378971 | 0.416428855 |
| ENSDART00000085438 | rps6ka5 | -0.245466951 | -0.387017016 | -0.309352273 | -0.119154274 |
| ENSDART00000085442 | mut | -0.094284216 | -0.181855717 | -0.336054518 | -0.342951233 |
| ENSDART00000085453 | cluhb | -0.484145834 | -0.430773206 | 0.678836708 | -0.017745757 |
| ENSDART00000085472 | grm2a | -0.506007863 | -0.654883229 | -0.268518199 | 0.087304206 |
| ENSDART00000085522 | hspb6 | -0.63697673 | -0.382982575 | -0.524822831 | -0.392317705 |
| ENSDART00000085528 | zgc:158659 | 0.601601674 | 0.738184978 | 0.82446023 | 0.805817257 |
| ENSDART00000085565 | capn15 | -0.382220369 | -0.426351921 | -0.021041552 | 0.033591442 |
| ENSDART00000085573 | rgs7bpa | -0.750985181 | -0.701592527 | -0.219110071 | -0.146877341 |
| ENSDART00000085612 | pcdh7b | -0.228836495 | -0.682608646 | -0.209101131 | 0.321031694 |
| ENSDART00000085675 | clstn2 | -0.244638643 | -0.37840099 | -0.228974377 | 0.094704371 |
| ENSDART00000085684 | ttll11 | -0.16810055 | 0.020727161 | 0.348458395 | 0.408691778 |
| ENSDART00000085693 | gpm6bb | -0.236628216 | -0.325741908 | -0.400571921 | -0.218025638 |
| ENSDART00000085716 | mtmr10 | 0.168573884 | -0.120117023 | -0.772748173 | -0.357648365 |

| | | | | | |
|---|---|---|---|---|---|
| ENSDART00000085719 | si:ch211-10a23.2 | 0.141906082 | 0.927202925 | 1.15606105 | 0.59243708 |
| ENSDART00000085743 | AL935194.1 | -0.774745358 | -0.68914542 | -0.32626184 | 0.039334462 |
| ENSDART00000085764 | PLOD3 | 0.414167142 | 0.202506542 | 0.107388816 | -0.10168282 |
| ENSDART00000085894 | pgm5 | 1.697021259 | 1.783911903 | 2.232651815 | 1.757015772 |
| ENSDART00000085993 | pxnb | 0.457322476 | 0.299419771 | 0.2051328 | -0.178681091 |
| ENSDART00000086051 | mecom | -0.550304416 | -0.298852875 | -0.293768303 | -1.334615988 |
| ENSDART00000086117 | kcnab2b | -0.362026581 | -0.574025944 | -0.45188395 | -0.145196155 |
| ENSDART00000086131 | pik3c2a | -0.071329777 | -0.019579026 | -0.128057314 | -0.294437789 |
| ENSDART00000086176 | nckap1 | -0.438983658 | -0.533344195 | -0.439721489 | -0.241138066 |
| ENSDART00000086181 | cabp7b | -0.513778995 | -0.579329622 | -0.46209057 | -0.104843884 |
| ENSDART00000086263 | mettl7a | -0.420009286 | -0.158826062 | -0.791334734 | -0.334080113 |
| ENSDART00000086281 | mavs | 3.81695815 | 3.176539113 | 3.535647274 | 1.650162791 |
| ENSDART00000086301 | irge4 | 0.875091617 | 0.681146857 | 1.737500297 | 0.803175592 |
| ENSDART00000086333 | jarid2a | -0.109318405 | 0.11715985 | 0.373609719 | 0.35216542 |
| ENSDART00000086409 | dync1i1 | -0.021724393 | 0.427961112 | 0.551451734 | 0.438467635 |
| ENSDART00000086434 | tmcc2 | 0.022092982 | 0.338134833 | 0.850580716 | 0.840264518 |
| ENSDART00000086495 | zgc:154077 | -0.133678102 | -0.278559633 | -0.319223928 | -0.190822097 |
| ENSDART00000086537 | | 0.422133332 | 0.379254853 | 0.651471983 | 0.472158305 |
| ENSDART00000086617 | gabbr2 | -0.577324776 | -0.926125922 | -0.433004274 | 0.052999603 |
| ENSDART00000086619 | prkca | -0.184045571 | -0.418285198 | -0.5357326 | -0.310736719 |
| ENSDART00000086664 | trpm1a | 2.715802028 | 4.516032094 | 3.238851669 | 3.187690164 |
| ENSDART00000086720 | nfasca | -0.417931887 | -0.576283691 | -0.313335716 | -0.150054847 |
| ENSDART00000086753 | dapk2a | 0.129419785 | -0.339935803 | -1.219284243 | -0.369104907 |
| ENSDART00000086797 | adgrl3.1 | -0.114053477 | -0.068166121 | 0.465121947 | 0.57391191 |
| ENSDART00000086867 | tapt1b | -0.153549174 | -0.259614839 | -0.218992302 | -0.161727037 |
| ENSDART00000086905 | nrn1lb | -0.455174799 | -0.417502528 | -0.690990621 | -0.656036806 |
| ENSDART00000086946 | mov10b.1 | 0.41644893 | 0.305142124 | 1.89370555 | 0.04709269 |
| ENSDART00000086952 | st14a | -0.283184933 | 0.988901668 | 1.786647129 | 1.271482733 |
| ENSDART00000086994 | nat15 | -0.144704703 | -0.149678764 | -0.337458722 | -0.152039739 |
| ENSDART00000087070 | abcc5 | 0.063506972 | -0.209331316 | -0.331966374 | -0.070006315 |
| ENSDART00000087097 | ogfrl1 | -0.189021726 | -0.350288897 | -0.29146473 | 0.000580163 |
| ENSDART00000087105 | myom2a | 2.177324505 | 3.608086766 | 3.587707417 | 2.779461648 |
| ENSDART00000087107 | eif4g1a | 0.252655247 | -0.051267066 | 0.413833734 | 0.330964133 |
| ENSDART00000087112 | pfdn4 | 0.448340647 | 0.452924674 | 0.218263471 | -0.022743355 |
| ENSDART00000087114 | alg5 | 0.317519293 | 0.464842808 | 0.220188706 | 0.059168575 |
| ENSDART00000087115 | rims1b | -0.155072254 | 0.041830315 | 0.378907446 | 0.287621284 |
| ENSDART00000087118 | xylt1 | -0.272537601 | -0.39007956 | -0.067871098 | 0.053209242 |
| ENSDART00000087148 | cbln4 | -0.802680954 | -0.769511703 | -0.146403482 | 0.304907793 |
| ENSDART00000087191 | mark4a | -0.415170271 | -0.372344968 | 0.042679549 | 0.108705644 |
| ENSDART00000087196 | zgc:153240 | -0.381380177 | -0.392146036 | -0.277166885 | -0.130863015 |
| ENSDART00000087204 | dusp3a | -0.327203373 | -0.053264752 | 0.592884224 | 0.622404395 |
| ENSDART00000087280 | cacnb3a | -0.323388935 | -0.465191463 | 0.268224123 | 0.533679905 |
| ENSDART00000087295 | ppp1r9a | -0.056007418 | -0.599550993 | -0.15522086 | -0.017719681 |
| ENSDART00000087300 | gabrb3 | -0.106002759 | -0.319333142 | -0.574890474 | -0.11105969 |
| ENSDART00000087311 | oca2 | 0.449332393 | 0.338475194 | -0.09076694 | -0.174077968 |
| ENSDART00000087329 | znf438 | -0.159611183 | -0.256202272 | -0.311134609 | -0.110177372 |
| ENSDART00000087339 | cdon | -0.358409006 | -0.160970657 | -0.495687672 | -0.865355598 |
| ENSDART00000087426 | bcl11aa | -0.367520296 | -0.334252656 | 0.232372171 | 0.511638417 |
| ENSDART00000087441 | GFOD1 | -0.421601041 | -0.219799435 | -0.160457067 | 0.058971995 |
| ENSDART00000087449 | CR847968.1 | -0.323687626 | -0.307881439 | 0.124728655 | 0.148402357 |
| ENSDART00000087450 | klf13 | -0.22166629 | -0.059571459 | 0.622443971 | 0.288396102 |

| | | | | | |
|---|---|---|---|---|---|
| ENSDART00000087565 | eva1a | -0.378976356 | -0.384921525 | -0.250389955 | -0.168669364 |
| ENSDART00000087570 | BRSK2 (1 of many) | -0.035673495 | -0.289130617 | -0.495870531 | -0.115059951 |
| ENSDART00000087586 | c2cd4a | -0.882018106 | -1.482044104 | -0.368555759 | 0.13043866 |
| ENSDART00000087624 | arhgef1b | 4.493077515 | 3.634214641 | 2.657484566 | 1.969740568 |
| ENSDART00000087643 | tesk2 | -0.250604524 | -0.077411225 | -0.349635246 | -0.097357509 |
| ENSDART00000087654 | adcy6a | -0.242190723 | -0.49624198 | -0.145481294 | -0.053244608 |
| ENSDART00000087726 | igf2bp1 | 0.052821083 | 0.330444102 | 0.755149975 | 0.389599697 |
| ENSDART00000087857 | unc5db | -0.464548804 | -0.771046574 | -0.267052184 | 0.0901494 |
| ENSDART00000087884 | ccdc85b | -0.309651273 | -0.084106622 | -0.027008739 | -0.094025951 |
| ENSDART00000087991 | fndc3bb | 0.509196457 | 0.583942244 | 0.486760061 | 0.158477497 |
| ENSDART00000088026 | prmt5 | 0.184159659 | 0.212388642 | 0.353449866 | 0.122961111 |
| ENSDART00000088027 | ssx2ipb | -0.254273665 | -0.300537366 | -0.659689526 | -0.365610708 |
| ENSDART00000088033 | trpm3 | -1.238410284 | -1.468940768 | -0.88192323 | -0.200606191 |
| ENSDART00000088042 | myo10l3 | 0.440398704 | 0.46516816 | 0.325219038 | -0.023584336 |
| ENSDART00000088093 | sipa1l2 | -0.401627573 | -0.319202181 | 0.366701976 | 0.056822769 |
| ENSDART00000088141 | ankrd34bb | 1.379198636 | 2.420328938 | 2.155206475 | 1.17838067 |
| ENSDART00000088146 | ensab | -0.270112525 | -0.281607362 | -0.344659286 | -0.084313447 |
| ENSDART00000088159 | nrxn1a | -0.55423697 | -0.307440551 | 0.007663974 | -0.036725507 |
| ENSDART00000088178 | nrxn1a | -0.424942944 | -0.357890595 | -0.01594532 | 0.150075885 |
| ENSDART00000088179 | nrxn3a | -0.554627027 | -0.919560714 | 0.023875656 | 0.130979956 |
| ENSDART00000088199 | zgc:162707 | -0.481388546 | -0.51007365 | -0.179950315 | -0.039369231 |
| ENSDART00000088240 | sypb | -0.232689126 | -0.445901517 | -0.375392134 | -0.149058325 |
| ENSDART00000088249 | hcn4l | -0.487910369 | -0.340719397 | 0.02727434 | 0.039415999 |
| ENSDART00000088270 | yjefn3 | -0.490178998 | -0.350279664 | -0.399921095 | -0.213170988 |
| ENSDART00000088290 | raph1b | 0.812305175 | 0.733830465 | 0.671238907 | 0.330964135 |
| ENSDART00000088336 | setdb1a | -0.249989255 | -0.274821833 | -0.108228373 | -0.04389578 |
| ENSDART00000088342 | cytip | 0.001918005 | -0.070156162 | -0.471206178 | -0.369058853 |
| ENSDART00000088364 | kif1aa | -0.06544832 | 0.34113245 | 0.707853532 | 0.648494399 |
| ENSDART00000088488 | opa3 | -0.282490417 | -0.240544978 | -0.173330249 | -0.094645025 |
| ENSDART00000088513 | gnl1 | 0.102638572 | 0.086645512 | -0.243843993 | -0.223952657 |
| ENSDART00000088569 | nyx | -0.258999398 | -0.197132955 | -0.26271962 | -0.211182996 |
| ENSDART00000088603 | UNC13A | -0.361581411 | -0.491389738 | -0.199237568 | 0.353821 |
| ENSDART00000088639 | wscd2 | -0.261498814 | -0.268790529 | -0.287619442 | -0.123420729 |
| ENSDART00000088643 | col15a1b | -0.287399924 | -0.285830353 | -0.360508672 | -0.35081124 |
| ENSDART00000088653 | prss12 | -0.641371677 | 0.068962263 | 0.215098805 | 0.069158882 |
| ENSDART00000088687 | rxfp3.2b | -0.990302103 | -0.900399551 | -0.191860941 | 0.37665355 |
| ENSDART00000088690 | lman2 | -0.00791489 | -0.046317548 | -0.243638306 | -0.186515184 |
| ENSDART00000088818 | fhod3b | -0.341939339 | -0.263444238 | -0.179945034 | 0.005787251 |
| ENSDART00000088833 | si:ch73-233f7.1 | -1.214678988 | -0.552820604 | -0.229157577 | 0.532052231 |
| ENSDART00000088881 | git2a | -0.368814104 | -0.568589271 | -0.278482365 | -0.109167728 |
| ENSDART00000088908 | srgap1a | -0.173831935 | -0.499543132 | -0.183148374 | 0.021193005 |
| ENSDART00000088973 | sytl2a | 0.735131572 | 0.479031428 | -0.049007315 | -0.597552798 |
| ENSDART00000089012 | kif1ab | -0.037675888 | 0.080069758 | 0.360995087 | 0.379282318 |
| ENSDART00000089015 | zbtb7a | -0.478666207 | -0.203919761 | -0.738823143 | -0.713706626 |
| ENSDART00000089033 | lingo3a | -0.421566075 | -0.428623374 | -0.267183911 | -0.077777224 |
| ENSDART00000089042 | kcnh4b | -0.293363138 | -0.305961812 | -0.46205676 | -0.207459851 |
| ENSDART00000089076 | dot1l | 0.094529933 | 0.165735937 | 0.448240264 | 0.463978672 |
| ENSDART00000089079 | mpnd | 0.049906779 | 0.784892334 | 0.951903848 | 0.691834891 |
| ENSDART00000089126 | trhde.2 | -0.409075032 | -0.405058445 | -0.383617798 | -0.218104015 |
| ENSDART00000089133 | rufy2 | -0.198896291 | 0.087437671 | 0.267347782 | 0.283743397 |
| ENSDART00000089141 | fsd1 | 0.140911421 | 0.537063785 | 0.66216846 | 0.489473142 |

| | | | | | |
|---|---|---|---|---|---|
| ENSDART00000089158 | hmha1a | 1.15366088 | 0.773436896 | 0.705540127 | -0.035689576 |
| ENSDART00000089161 | CCDC181 | -0.238708752 | -0.276570098 | -0.364394143 | -0.212879757 |
| ENSDART00000089246 | elmod1 | -0.367956253 | -0.511095234 | 0.063835683 | 0.117723407 |
| ENSDART00000089325 | mief1 | -0.272432924 | -0.240984001 | -0.06454421 | 0.031387691 |
| ENSDART00000089339 | dph7 | -0.317220664 | -0.267955931 | -0.402220665 | -0.245139947 |
| ENSDART00000089342 | cfap126 | -0.196227391 | -0.233968621 | -0.449699376 | -0.526724945 |
| ENSDART00000089408 | shdb | -0.379456562 | -0.060183647 | 0.137084622 | 0.164920585 |
| ENSDART00000089442 | klhl5 | -0.428861339 | -0.43123859 | -0.099854083 | -0.052484669 |
| ENSDART00000089445 | agap1 | 0.111728293 | 0.12652166 | 0.321279752 | 0.223152778 |
| ENSDART00000089488 | sytl5 | -0.2422586 | -0.383469303 | 0.0006109 | 0.143303123 |
| ENSDART00000089526 | otc | -0.228950714 | -0.50691522 | -0.941930968 | -0.587900491 |
| ENSDART00000089540 | sacm1la | -0.121729942 | -0.218997931 | -0.253159012 | -0.069593401 |
| ENSDART00000089549 | fam65a | 0.138661993 | 0.299682143 | 0.16216271 | 0.248418418 |
| ENSDART00000089574 | tub | -0.338880988 | -0.401453479 | -0.101730157 | -0.038037712 |
| ENSDART00000089577 | cacnb4b | -0.344473142 | 0.166216688 | 0.479684005 | 0.375262058 |
| ENSDART00000089699 | prrt1 | -0.603201784 | -0.702306626 | -0.280611263 | -0.047540659 |
| ENSDART00000089748 | rorb | -0.330969943 | -0.539847879 | -0.425468683 | -0.138099942 |
| ENSDART00000089867 | ppp2r2cb | -0.452836704 | -0.44451904 | -0.189867818 | -0.082146078 |
| ENSDART00000089923 | znf652 | -0.30560984 | -0.322535846 | -0.127141181 | -0.092457617 |
| ENSDART00000089961 | sik2a | -0.349078539 | -0.249421195 | -0.402456418 | 0.05469833 |
| ENSDART00000089967 | cacna1bb | -0.202836628 | -0.60611593 | -0.329318476 | -0.282649478 |
| ENSDART00000089968 | rasl10a | -0.525764509 | -0.884552715 | -0.504672449 | -0.419223822 |
| ENSDART00000089992 | hmgn7 | -0.150988875 | -0.15001014 | -0.251989002 | -0.139708395 |
| ENSDART00000089999 | b4galt3 | -0.174643035 | -0.177020263 | -0.279641129 | -0.298606277 |
| ENSDART00000090010 | phex | -0.386748819 | -0.231869791 | -0.214849122 | -0.613717457 |
| ENSDART00000090019 | zeb2b | -0.694941899 | -0.428168304 | -0.296660745 | 0.024845807 |
| ENSDART00000090079 | synm | -0.113622285 | -0.399027729 | -0.70355851 | -0.251171145 |
| ENSDART00000090174 | dock9b | 0.356010204 | 0.810416379 | 0.943578275 | 0.58912554 |
| ENSDART00000090191 | flcn | 0.595641364 | 0.372738928 | 0.082710169 | 0.079174241 |
| ENSDART00000090221 | cdc42ep5 | 0.276662145 | 0.611781241 | 0.632201471 | -0.020036411 |
| ENSDART00000090235 | nfixb | -0.570962215 | -0.312719979 | -0.289246101 | -0.209164384 |
| ENSDART00000090252 | atp2b3a | -0.293678426 | -0.382019678 | -0.276234519 | -0.034568861 |
| ENSDART00000090266 | gpd2 | 0.305577979 | 0.440771229 | 0.397599751 | 0.146739419 |
| ENSDART00000090292 | ctnnd2b | -0.150466015 | -0.062584536 | 0.302491723 | 0.338163791 |
| ENSDART00000090306 | xpr1a | -0.158299732 | -0.301120383 | 0.179396195 | 0.424625304 |
| ENSDART00000090335 | hipk2 | -0.356266924 | -0.261702878 | -0.185362868 | -0.057591831 |
| ENSDART00000090397 | kiaa1549la | -0.446832614 | -0.575803117 | -0.232865535 | 0.10265337 |
| ENSDART00000090406 | dock11 | 0.704689621 | 0.535408523 | 0.615472635 | 0.363574103 |
| ENSDART00000090483 | cplx3a | -0.167017721 | -0.181539522 | -0.397723314 | -0.182035461 |
| ENSDART00000090484 | tecpr1a | -0.079583722 | -0.244227266 | -0.30265493 | -0.135390116 |
| ENSDART00000090521 | ankle2 | 0.59590492 | 0.614380049 | 0.713135872 | 0.291410446 |
| ENSDART00000090528 | rhoca | 0.570199757 | 0.738551724 | 0.41277501 | -0.074138712 |
| ENSDART00000090534 | ulk1a | 0.04466428 | 0.367530579 | 0.126656159 | 0.034855767 |
| ENSDART00000090548 | ASTE1 | 0.186068328 | 0.308540808 | 0.373512724 | 0.303370928 |
| ENSDART00000090580 | si:dkey-215k6.1 | -0.353341621 | -0.406611115 | -0.330167398 | -0.116132596 |
| ENSDART00000090596 | fgf12b | -0.346267349 | -0.034488788 | 0.704892235 | 0.660670447 |
| ENSDART00000090611 | sh3gl2a | -0.3620314 | -0.426828792 | -0.627861098 | -0.256916064 |
| ENSDART00000090669 | pleca | -0.254130273 | -0.426440212 | -0.236918819 | -0.12154508 |
| ENSDART00000090689 | brat1 | 0.263072231 | 0.449900152 | 0.263104838 | 0.157050832 |
| ENSDART00000090709 | coq7 | -0.258539203 | -0.273311663 | -0.220704286 | -0.134757577 |
| ENSDART00000090711 | ltc4s | 0.402288552 | 0.209272555 | 0.13658761 | 0.029040107 |

| | | | | | |
|---|---|---|---|---|---|
| ENSDART00000090748 | pcdh1g9 | 0.151679401 | 0.129768355 | 0.681696549 | 0.559518569 |
| ENSDART00000090757 | kat2b | -0.251752686 | -0.26045996 | -0.262277009 | -0.210249532 |
| ENSDART00000090771 | cyth1a | 1.102348748 | 1.012723243 | 0.839881021 | 0.442796833 |
| ENSDART00000090844 | zgc:153018 | -0.414733046 | -0.080390611 | -0.214556251 | -0.032786964 |
| ENSDART00000090864 | lmod3 | 0.210035398 | 1.148748445 | 1.650370875 | 1.365208896 |
| ENSDART00000090874 | kcnh7 | 0.089634054 | 0.066677874 | 0.800119794 | 0.739010611 |
| ENSDART00000090883 | gpnmb | 1.78024327 | 1.792109343 | 1.484400673 | -0.264951498 |
| ENSDART00000091004 | pcdh1a | -0.630879138 | -0.627674521 | -0.227167517 | 0.17418403 |
| ENSDART00000091017 | pkn1b | 0.119108789 | 0.041947386 | 0.390816496 | 0.137505167 |
| ENSDART00000091021 | col10a1a | 0.454586287 | 3.186498396 | 3.689025905 | 2.153861861 |
| ENSDART00000091124 | aifm3 | -0.238811768 | -0.279253276 | -0.598306352 | -0.861256088 |
| ENSDART00000091140 | snx21 | 0.278697916 | 0.637246869 | 0.128859029 | 0.534229621 |
| ENSDART00000091151 | nell2b | -0.408104365 | -0.533629775 | -0.525750199 | -0.236120482 |
| ENSDART00000091156 | tacc1 | -0.166984032 | -0.162668339 | -0.396514006 | -0.330157066 |
| ENSDART00000091158 | irg1l | 1.781534333 | 2.6026377 | 1.369996198 | -0.413577442 |
| ENSDART00000091183 | erfl3 | -0.064022984 | -0.222171088 | -0.380622737 | -0.084018803 |
| ENSDART00000091205 | sdk1b | -0.11409829 | -0.274173881 | -0.261923413 | -0.147107702 |
| ENSDART00000091241 | si:ch73-22o12.1 | -0.053659471 | 0.086463787 | 0.414689493 | 0.463272551 |
| ENSDART00000091252 | spata13 | 0.108055641 | -0.023595371 | -0.309799561 | -0.91906317 |
| ENSDART00000091271 | prkg2l | -0.404320096 | -0.412762377 | -0.544649422 | -0.030408858 |
| ENSDART00000091331 | prodha | -0.295055194 | -0.294720047 | -0.215860993 | -0.495616796 |
| ENSDART00000091351 | gk5 | 0.51312557 | 0.289440127 | 0.273716274 | -0.08764101 |
| ENSDART00000091409 | smarcad1a | 0.370815487 | 0.374702329 | 0.447117461 | 0.290143851 |
| ENSDART00000091416 | cntn3a.1 | -0.478191534 | -0.24621663 | -0.434975173 | -0.073764559 |
| ENSDART00000091452 | TULP2 | -0.183522253 | -0.233522919 | -0.588056807 | -0.14814607 |
| ENSDART00000091472 | kcnv2b | -0.549604515 | -0.416623158 | -0.289225353 | -0.099467789 |
| ENSDART00000091489 | ppp1r9bb | -0.563706295 | -0.346664158 | -0.080794638 | 0.037246954 |
| ENSDART00000091508 | FAM83G | 0.624622597 | 0.848983587 | 0.795841019 | 0.137374674 |
| ENSDART00000091532 | ndnf | -0.379105533 | -0.357674977 | -0.332342823 | -0.25490154 |
| ENSDART00000091584 | zgc:158785 | -0.030208367 | -0.597188759 | -0.577152486 | -0.343586348 |
| ENSDART00000091599 | sbf1 | -0.602535014 | -0.60076227 | 0.11831949 | 0.138965635 |
| ENSDART00000091612 | dab2ipa | -0.32378487 | -0.312766229 | -0.160534626 | -0.215968275 |
| ENSDART00000091615 | iffo1a | -0.255558776 | -0.319129649 | -0.402938235 | -0.1114809 |
| ENSDART00000091620 | atp8a1 | -0.02813322 | -0.262306637 | -0.342910163 | -0.163047911 |
| ENSDART00000091644 | abi1b | 0.181595456 | 0.359192672 | 0.565695572 | 0.32038751 |
| ENSDART00000091662 | noc2l | 0.778877814 | 0.364993344 | 0.393512667 | -0.105586454 |
| ENSDART00000091664 | apc2 | -0.58155429 | -0.22615315 | 0.136652181 | 0.006999355 |
| ENSDART00000091683 | alkbh5 | -0.400075016 | -0.448742364 | -0.161106699 | -0.178285375 |
| ENSDART00000091707 | dbpa | -0.158262702 | -0.327851995 | -0.357551076 | -0.129469939 |
| ENSDART00000091726 | fam78ba | -0.542782218 | -0.34207093 | 0.322203074 | 0.384590985 |
| ENSDART00000091727 | ntrk3a | -0.313031117 | -0.882724333 | -0.168396888 | 0.114845125 |
| ENSDART00000091729 | mlc1 | -0.413445193 | -0.441446959 | -0.597560289 | -0.400243567 |
| ENSDART00000091780 | rc3h2 | -0.185144786 | -0.339077629 | -0.231755111 | -0.132461257 |
| ENSDART00000091818 | tulp4b | -0.531343855 | -0.277019809 | 0.128892269 | 0.551121944 |
| ENSDART00000091899 | ccm2l | 0.082170149 | -0.344528731 | -0.637200318 | -0.089681617 |
| ENSDART00000091901 | psmd14 | 0.227724961 | 0.262600592 | 0.258422041 | 0.051950672 |
| ENSDART00000091923 | slc4a10a | 0.0431458 | -0.481021076 | -0.208884142 | 0.00832231 |
| ENSDART00000091932 | gusb | -0.012758115 | -0.102829729 | -0.25264355 | -0.30752198 |
| ENSDART00000091955 | nrxn2b | -0.363238969 | -0.38605701 | -0.167716958 | 0.046918181 |
| ENSDART00000092013 | tmtc1 | -0.379958165 | -0.299353758 | -0.293525457 | -0.123093715 |
| ENSDART00000092050 | stab1 | 0.830009942 | 0.662655195 | 0.610953846 | 0.245694157 |

| | | | | | |
|---|---|---|---|---|---|
| ENSDART00000092051 | CABZ01081780.1 | -0.497076796 | -0.683547714 | -0.447049069 | -0.155316145 |
| ENSDART00000092114 | ERBB4 (1 of many) | -0.124937285 | -0.553537291 | -0.521885412 | -0.338309306 |
| ENSDART00000092164 | prmt2 | 0.865577917 | 0.607769301 | 0.137454327 | 0.149744108 |
| ENSDART00000092182 | ppm1la | -0.411282562 | -0.137536548 | 0.156346356 | 0.193635075 |
| ENSDART00000092183 | lrrc3b | -0.469734111 | -0.850621377 | -0.496353665 | -0.22903801 |
| ENSDART00000092239 | lix1l | -0.01751945 | 0.310063812 | 0.327664672 | 0.131969727 |
| ENSDART00000092250 | btbd11a | -0.250819977 | -0.42148278 | -0.144893414 | 0.093892491 |
| ENSDART00000092257 | PLD5 | -0.97635635 | -0.828911686 | 0.040708181 | 0.36058098 |
| ENSDART00000092270 | r3hdm4 | 0.109330042 | 0.341009284 | 0.430965659 | 0.322117566 |
| ENSDART00000092290 | pcdh9 | -0.617638559 | -1.059395061 | -0.632995564 | 0.015160294 |
| ENSDART00000092356 | neto1 | -0.404606374 | -0.302098716 | -0.383296194 | -0.272667962 |
| ENSDART00000092357 | sgsm2 | -0.379055867 | -0.375730796 | -0.097265456 | -0.053118339 |
| ENSDART00000092381 | pcloa | -0.033919449 | -0.560249847 | -0.672386944 | -0.219295292 |
| ENSDART00000092389 | nup210 | 0.743287563 | 0.315006844 | 0.505674911 | 0.088834448 |
| ENSDART00000092406 | apba2a | 0.032423751 | 0.148342908 | 0.468800105 | 0.354794985 |
| ENSDART00000092416 | rabl2 | -0.237808365 | -0.250658742 | -0.360133928 | -0.144079351 |
| ENSDART00000092435 | mgat4a | 0.252074722 | 0.11851857 | 0.178053587 | 0.019110648 |
| ENSDART00000092493 | ptprt | -0.175495642 | -0.444255461 | -0.308001112 | -0.058909837 |
| ENSDART00000092524 | rasa3 | -0.348887401 | -0.411266738 | -0.465367898 | -0.079002088 |
| ENSDART00000092646 | lrrc73 | -0.475821333 | -0.175234523 | 0.140632325 | 0.202367883 |
| ENSDART00000092647 | cers1 | -0.35221666 | -0.292832737 | -0.390981877 | -0.135220693 |
| ENSDART00000092665 | srebf1 | -0.417658309 | -0.274360919 | 0.114636817 | 0.022214824 |
| ENSDART00000092690 | srebf2 | -0.007030288 | 0.441941306 | 0.773565631 | 0.436893785 |
| ENSDART00000092691 | pea15 | -0.578361557 | -0.742039167 | -0.342285902 | -0.180997926 |
| ENSDART00000092884 | lrrc58b | 0.353547699 | 0.596933578 | 0.36096561 | 0.06120599 |
| ENSDART00000092948 | peli1b | 0.088983174 | 0.130189996 | 0.342697853 | 0.254903056 |
| ENSDART00000093000 | PLEKHB1 | -0.366311468 | -0.143931789 | -0.403311135 | -0.393296015 |
| ENSDART00000093003 | syt7a | -0.241216791 | -0.606859779 | -0.288317918 | -0.007740028 |
| ENSDART00000093005 | DGKI | -0.31222183 | -0.268945726 | 0.173367169 | 0.253167333 |
| ENSDART00000093093 | coro2bb | -0.315281988 | -0.251087256 | -0.471013192 | -0.151819548 |
| ENSDART00000093149 | ddx21 | 0.146331643 | 0.244276574 | 0.410170406 | 0.14971067 |
| ENSDART00000093155 | hpse | 0.712086441 | 0.364284216 | 0.211412521 | -0.486650265 |
| ENSDART00000093163 | galnt11 | -0.148415291 | -0.19498093 | -0.27912109 | -0.157806623 |
| ENSDART00000093166 | nrxn1b | 0.047890691 | -0.072856775 | -0.26075298 | -0.08007638 |
| ENSDART00000093193 | CABZ01090021.1 | 0.615976841 | 0.876382917 | 0.705233449 | 0.369811976 |
| ENSDART00000093199 | tead3b | 1.081516431 | 0.86043279 | 0.478204597 | -0.215799128 |
| ENSDART00000093236 | TULP3 | -0.257622072 | -0.196815312 | -0.318097587 | -0.153320475 |
| ENSDART00000093279 | spi1b | 0.891535687 | 0.563855895 | 0.100587929 | -0.531177734 |
| ENSDART00000093304 | nrm | 0.448051883 | 0.19826204 | 0.059665608 | 0.061320945 |
| ENSDART00000093310 | celf5a | -0.268154664 | -0.455004312 | 0.014815598 | 0.070405312 |
| ENSDART00000093331 | rreb1a | -0.024901133 | -0.155149005 | -0.517234878 | -0.050769944 |
| ENSDART00000097176 | CABZ01090749.1 | 1.277443607 | 2.06888261 | 1.727457254 | 0.676889483 |
| ENSDART00000097194 | serinc5 | 0.184853776 | 0.289193819 | 0.374387724 | 0.244648582 |
| ENSDART00000097198 | sc:d217 | 2.007573379 | 2.429847079 | 2.207258628 | 1.807569646 |
| ENSDART00000097248 | aldh2.2 | 0.244379894 | -0.20748537 | -0.365189715 | -0.463727898 |
| ENSDART00000097249 | aldh2.2 | -1.049924711 | -1.051244298 | -1.854940324 | -3.57887137 |
| ENSDART00000097330 | dnm1b | -0.51283412 | -0.480365631 | 0.101619237 | 0.531582232 |
| ENSDART00000097338 | napaa | -0.21867082 | -0.198889828 | -0.441263528 | -0.268359175 |
| ENSDART00000097359 | dnajc25 | 0.258588311 | 0.413120409 | 0.370076595 | 0.197964188 |
| ENSDART00000097460 | hmgcra | -0.174557714 | 0.058201434 | 0.38988663 | 0.431312576 |
| ENSDART00000097466 | fam169aa | -0.25560307 | -0.389662586 | -0.539552997 | -0.303655 |

| | | | | | |
|---|---|---|---|---|---|
| ENSDART00000097670 | ggcx | -0.008080389 | -0.071755289 | -0.320769539 | -0.236029667 |
| ENSDART00000097685 | LINGO3 (1 of many) | -0.608919806 | -0.847171007 | -0.317394062 | -0.126526617 |
| ENSDART00000097695 | cntnap3 | -0.160493697 | -0.529313782 | -0.088115665 | 0.248028839 |
| ENSDART00000097731 | valopa | -0.109885721 | -0.121152137 | -0.343824948 | -0.490387529 |
| ENSDART00000097738 | panx1b | 0.147981258 | 0.33056409 | 0.233249696 | 0.121977096 |
| ENSDART00000097770 | gc3 | -0.462816548 | -0.433977737 | -0.284956472 | -0.124147363 |
| ENSDART00000097792 | tnikb | 0.209649579 | 0.123927361 | 0.351444411 | 0.3418539 |
| ENSDART00000097822 | atp1b2b | -0.302706184 | -0.327970639 | -0.363261939 | -0.05554983 |
| ENSDART00000097934 | cdh4 | -0.709680135 | -0.297434282 | -0.207521755 | 0.118311374 |
| ENSDART00000097935 | si:dkey-226m8.10 | -0.692305988 | -0.817395111 | -0.44855289 | 0.050175879 |
| ENSDART00000097939 | zcchc2 | -0.095224914 | -0.152148503 | 0.35105086 | 0.119260586 |
| ENSDART00000098038 | dscamb | -0.314235441 | -0.377418781 | -0.108405688 | -0.031209686 |
| ENSDART00000098045 | gas1b | -0.225161666 | -0.17230943 | -0.331649886 | -0.52065088 |
| ENSDART00000098057 | ftr19 | 0.135898929 | 0.299355243 | 0.876453981 | -0.013001669 |
| ENSDART00000098058 | ftr22 | 0.119426338 | -0.017860927 | 0.367700006 | -0.042033613 |
| ENSDART00000098072 | myhz1.1 | -1.045546706 | 2.305130821 | 3.388084443 | 2.053099835 |
| ENSDART00000098082 | GJD2 (1 of many) | -0.795483498 | -0.746707283 | 0.031300804 | 0.456345915 |
| ENSDART00000098173 | vapal | -0.239328654 | -0.269836839 | -0.058452226 | -0.037670486 |
| ENSDART00000098209 | sirt1 | 0.078955516 | 0.10137907 | 0.304759128 | 0.182124247 |
| ENSDART00000098263 | kctd9a | -0.029411295 | -0.237424393 | -0.359070767 | -0.190010537 |
| ENSDART00000098284 | ftr14 | 0.428599203 | 0.304151818 | 1.741077228 | -0.09285898 |
| ENSDART00000098285 | atf5a | 0.558593719 | 0.754458044 | 0.445954377 | 0.160619888 |
| ENSDART00000098311 | KCNJ4 | -0.486217707 | -0.434414335 | -0.027097558 | 0.056787682 |
| ENSDART00000098361 | nmba | -1.096273595 | -1.192632984 | -0.904325972 | -1.052265102 |
| ENSDART00000098424 | trib2 | -0.755876906 | -0.434835466 | -0.414057266 | -0.366344874 |
| ENSDART00000098545 | tmem150aa | 0.458575321 | 0.166363456 | 0.137813682 | -0.033806431 |
| ENSDART00000098567 | CRAT (1 of many) | -0.211452678 | -0.154808037 | -0.361957852 | -0.341818431 |
| ENSDART00000098571 | CD53 | 0.954510613 | 0.498572956 | 0.147686328 | -0.376083033 |
| ENSDART00000098575 | trim110 | -0.538539785 | -0.074004136 | -0.613819609 | -0.257802554 |
| ENSDART00000098590 | cyb561a3a | 0.500971443 | 0.776880119 | 0.690384128 | 0.607288952 |
| ENSDART00000098616 | rwdd | 0.47601582 | 0.122947158 | 0.180975352 | 0.051928757 |
| ENSDART00000098627 | pros1 | -0.136816155 | -0.05658809 | -0.154307635 | -0.425593287 |
| ENSDART00000098639 | cntn5 | -0.259764888 | -0.493723855 | -0.530432667 | -0.172558087 |
| ENSDART00000098643 | tomm5 | 0.01929525 | -0.165199909 | -0.247037767 | -0.265103531 |
| ENSDART00000098648 | gc2 | -0.195652835 | -0.378422178 | -0.402398473 | -0.153861921 |
| ENSDART00000098667 | camk2b1 | -0.575252799 | -0.701444666 | -0.414382214 | 0.077165391 |
| ENSDART00000098668 | abcc8b | -0.217525237 | -0.529877646 | -0.409188595 | -0.288785069 |
| ENSDART00000098673 | ptx3a | -0.211528365 | -0.387085724 | -0.417540192 | -0.397382448 |
| ENSDART00000098727 | svopa | -0.499297278 | -0.394465259 | 0.04106692 | 0.031976449 |
| ENSDART00000098750 | pdlim5b | 0.340621907 | 0.643165703 | 0.410131138 | 0.075261902 |
| ENSDART00000098840 | ralgps1 | -0.525377749 | -0.528705894 | -0.140236147 | 0.156796103 |
| ENSDART00000098859 | neurod6a | -0.523659161 | -0.308368621 | 0.583701544 | 0.979244768 |
| ENSDART00000098970 | lin28a | 3.476899763 | 1.864207832 | 0.92280153 | 0.893269961 |
| ENSDART00000098982 | h3f3b.1 | -0.123835393 | -0.18991532 | -0.279701511 | -0.152040805 |
| ENSDART00000099003 | plscr3b | 0.864146968 | 0.903022775 | 0.644348755 | -0.059087289 |
| ENSDART00000099019 | tmem91 | -0.661722152 | -0.699132453 | -0.1920109 | -0.057277853 |
| ENSDART00000099049 | | -0.472159773 | -0.420532241 | -0.250637325 | -0.02437969 |
| ENSDART00000099056 | gpx4a | -0.526913186 | -0.311025548 | -0.820323342 | -0.895800055 |
| ENSDART00000099089 | ggcx | -0.059935848 | -0.149142859 | -0.333945915 | -0.254577798 |
| ENSDART00000099102 | sept5a | 0.488119217 | 0.804836681 | 0.729175054 | 0.142689277 |
| ENSDART00000099138 | ncf2 | 0.590402106 | 0.352177279 | 0.071326562 | -0.071022128 |

| | | | | | |
|---|---|---|---|---|---|
| ENSDART00000099180 | elovl8a | -0.351208319 | -0.254455279 | -0.702580214 | -0.196165774 |
| ENSDART00000099192 | htr5ab | -0.290360215 | -0.577686019 | -0.280786496 | -0.015762839 |
| ENSDART00000099202 | igsf11 | -0.183727476 | -0.31314564 | -0.202768681 | -0.036422441 |
| ENSDART00000099208 | asph | -0.288641006 | -0.271409423 | -0.460024196 | -0.387402543 |
| ENSDART00000099235 | rnf44 | -0.234703995 | -0.37643632 | -0.256248531 | 0.012942846 |
| ENSDART00000099244 | CDHR2 | -0.127835174 | -0.277679177 | -0.519160965 | -0.069385603 |
| ENSDART00000099248 | rabggtb | 0.075526469 | 0.438084725 | 0.330428907 | 0.205346366 |
| ENSDART00000099283 | dalrd3 | -0.272196688 | -0.367831508 | -0.108158836 | 0.07067938 |
| ENSDART00000099325 | si:dkey-27p18.5 | -0.821017867 | -0.681399645 | -0.140207831 | 0.113491694 |
| ENSDART00000099389 | dnlz | 0.363796601 | 0.422259676 | 0.108203262 | 0.027839921 |
| ENSDART00000099392 | irgq2 | -0.080759981 | -0.21229475 | -0.382096269 | -0.042741954 |
| ENSDART00000099476 | fam174b | -0.249603276 | -0.126773683 | -0.376574062 | -0.35637543 |
| ENSDART00000099501 | masp1 | 0.656652704 | -0.018219479 | -0.317919643 | -0.733675787 |
| ENSDART00000099528 | spry4 | -1.005535524 | -0.789062042 | -0.760146662 | -1.480463024 |
| ENSDART00000099532 | vmhc | -0.018141777 | 3.134710049 | 3.519069106 | 1.786644694 |
| ENSDART00000099566 | si:ch211-244o22.2 | 0.576141582 | 0.625409237 | 0.697152634 | 0.270183175 |
| ENSDART00000099568 | gpr137bb | 0.063339268 | 0.018950684 | 0.393403815 | 0.411006 |
| ENSDART00000099607 | slc6a17 | -0.493487825 | -0.358976735 | -0.156955324 | 0.010739948 |
| ENSDART00000099690 | fam129ab | 1.024572276 | 1.489725431 | 1.470862577 | 0.737761428 |
| ENSDART00000099764 | zgc:153031 | 0.216426658 | 0.538240257 | 0.500334626 | 0.476962119 |
| ENSDART00000099769 | ccdc22 | 0.325453994 | 0.346355629 | 0.367424682 | 0.130188245 |
| ENSDART00000099839 | map2k2b | 4.411409189 | 4.089988152 | 4.160217635 | 3.732217772 |
| ENSDART00000099849 | arntl2 | 0.982355145 | 1.78511579 | 1.609893892 | 1.255639962 |
| ENSDART00000099869 | slc17a7b | -0.359089552 | -0.531552161 | -0.360552754 | 0.022409885 |
| ENSDART00000099872 | slc17a6b | -0.645728405 | -0.242319052 | 0.108083805 | 0.273480994 |
| ENSDART00000099891 | atp5ib | -0.202173954 | -0.194749139 | -0.288932188 | -0.060278853 |
| ENSDART00000099934 | kcnc1a | -0.987934197 | -1.510396524 | -0.387995788 | 0.170469595 |
| ENSDART00000099947 | samsn1a | 0.989297054 | 0.26224771 | 0.078735734 | -0.489522484 |
| ENSDART00000099977 | mex3c | 0.161233198 | 0.169039948 | 0.509657606 | 0.094152947 |
| ENSDART00000099978 | pdlim4 | 0.618309518 | 0.542997573 | 0.115667653 | -0.045450127 |
| ENSDART00000099994 | hspa8 | 0.258700063 | 0.242999424 | 0.401354403 | 0.332565903 |
| ENSDART00000100000 | gabra1 | -0.401815194 | -0.497271272 | -0.174154846 | 0.00256896 |
| ENSDART00000100022 | H2AFX (1 of many) | -0.512580891 | -0.39296931 | -0.220510282 | -0.111022185 |
| ENSDART00000100074 | pbx3a | -0.314908702 | -0.751877287 | -0.359257757 | -0.077459645 |
| ENSDART00000100103 | acss2l | -0.342716857 | -0.308956489 | -0.115247207 | 0.030743328 |
| ENSDART00000100110 | pak6b | -0.707774149 | -0.488636508 | 0.157100943 | -0.07651745 |
| ENSDART00000100117 | znf143b | -4.095841085 | -0.108331644 | -1.7298033 | -2.907687665 |
| ENSDART00000100131 | si:ch211-242e8.1 | 0.043404066 | -0.450710823 | -0.466087638 | 0.073034701 |
| ENSDART00000100145 | lgals9l1 | 1.730493466 | 0.73620688 | 0.17444521 | -0.331961456 |
| ENSDART00000100156 | agpat4 | 1.221504112 | 1.6039826 | 1.49974957 | 0.799345848 |
| ENSDART00000100181 | sall3b | -0.170454734 | -0.209016072 | -0.285639823 | -0.232809645 |
| ENSDART00000100194 | msi2b | -0.174290948 | -0.286012891 | -0.448717518 | -0.171818835 |
| ENSDART00000100223 | zgc:91860 | -0.412727201 | -0.311203152 | -0.280203316 | -0.243367869 |
| ENSDART00000100234 | col2a1a | -0.262380123 | -0.070947069 | -0.203936285 | -0.943179815 |
| ENSDART00000100241 | haao | -0.290824489 | -0.384992406 | -0.679902583 | -0.875375738 |
| ENSDART00000100286 | fgfr4 | -0.349487842 | -0.279142199 | -0.557520873 | -0.83628708 |
| ENSDART00000100287 | grk7a | -0.29429259 | -0.34179807 | -0.766271921 | -0.26689268 |
| ENSDART00000100290 | napbb | -0.403034353 | -0.312472409 | -0.341096083 | -0.127811116 |
| ENSDART00000100310 | dbn1 | 0.014767176 | 0.221674025 | 0.550020524 | 0.324902581 |
| ENSDART00000100320 | dnmt3aa | -0.203167185 | -0.31036097 | -0.115665251 | -0.0614106 |
| ENSDART00000100322 | kcnh5b | -0.532935362 | -0.444322241 | -0.174846578 | 0.219995697 |

| | | | | | |
|---|---|---|---|---|---|
| ENSDART00000100327 | nptx1l | -0.348659958 | -0.433920996 | -0.393251042 | -0.286427391 |
| ENSDART00000100332 | fgf12b | -0.220479512 | 0.029676005 | 0.518232207 | 0.657400596 |
| ENSDART00000100386 | mstnb | -0.232479836 | -0.19741782 | -0.546065599 | -0.487217319 |
| ENSDART00000100401 | aars | 0.036948074 | 0.225050642 | 0.3787132 | 0.301138312 |
| ENSDART00000100415 | map3k7cl | 1.690116609 | 2.517738065 | 2.168584787 | 1.392466275 |
| ENSDART00000100438 | rab38b | 0.552254991 | 0.331427199 | 0.26198981 | -0.020729573 |
| ENSDART00000100444 | fam19a5a | -0.63242802 | -0.536601669 | -0.265992198 | 0.12320759 |
| ENSDART00000100453 | cerk | -0.133196976 | -0.125062952 | -0.291644205 | -0.1718331 |
| ENSDART00000100458 | si:dkey-73n10.1 | 1.648347098 | 0.766263068 | 0.204715566 | -0.361349683 |
| ENSDART00000100473 | PLIN3 | 1.275027857 | 0.887739595 | 0.535269477 | -0.111777778 |
| ENSDART00000100596 | pcdh1a6 | -0.483816366 | -0.257584823 | -0.06041051 | 0.00168094 |
| ENSDART00000100605 | ttc32 | -0.362267617 | -0.196641209 | -0.477779827 | -0.443477008 |
| ENSDART00000100619 | zgc:158803 | 0.343521186 | 0.388812734 | 0.518620429 | 0.45576524 |
| ENSDART00000100622 | slc15a2 | 1.330001469 | 0.871350782 | 0.119367987 | -0.187130808 |
| ENSDART00000100639 | chrm4a | -0.541979173 | -0.464947812 | -0.239039654 | -0.216630506 |
| ENSDART00000100658 | esrra | -0.571475474 | -0.395296451 | 0.169003635 | 0.212368184 |
| ENSDART00000100667 | skia | -0.180939747 | -0.108825609 | -0.328725059 | -0.076853639 |
| ENSDART00000100681 | ncam2 | -0.42007768 | -0.748629002 | -0.327699832 | -0.08364547 |
| ENSDART00000100743 | cttnbp2 | -0.40355339 | -0.391763552 | 0.009167842 | 0.081098 |
| ENSDART00000100762 | inpp4ab | -0.155677368 | -0.41981961 | -0.539565797 | -0.358079101 |
| ENSDART00000100798 | trip6 | -0.01587524 | -0.010041711 | 0.309937262 | 0.254192292 |
| ENSDART00000100813 | rps24 | 0.369574152 | 0.327820695 | 0.143165091 | -0.049601165 |
| ENSDART00000100869 | ppp3r1b | -0.102347685 | -0.214447853 | -0.31663745 | -0.211814655 |
| ENSDART00000100877 | zgc:153142 | -0.183846915 | -0.253996232 | -0.437661908 | -0.081274008 |
| ENSDART00000100885 | nrn1la | -0.520024314 | -0.30927413 | -0.690569972 | -0.461401545 |
| ENSDART00000100898 | tnfsf12 | 0.387181383 | 0.213965186 | 0.122654268 | 0.108692776 |
| ENSDART00000101014 | cx32.2 | 1.233652332 | 0.399790197 | -0.050586016 | -0.568712262 |
| ENSDART00000101037 | nhp2 | 0.465910387 | 0.391566586 | 0.240162003 | -0.008355829 |
| ENSDART00000101038 | tmie | 0.304653896 | 1.158765103 | 1.299122704 | 1.103412502 |
| ENSDART00000101044 | hsbp1a | 0.110662102 | 0.30138061 | 0.346862926 | 0.268898796 |
| ENSDART00000101070 | dachd | -0.296270747 | -0.351582482 | -0.130791564 | -0.009393292 |
| ENSDART00000101097 | acp6 | 0.210112925 | 0.333975464 | 0.275264967 | 0.086639765 |
| ENSDART00000101124 | rnaseka | 1.077486687 | 0.59165486 | 0.038549521 | -0.354475184 |
| ENSDART00000101134 | khdrbs2 | -0.661640499 | -0.469301828 | -0.265343985 | -0.029039929 |
| ENSDART00000101142 | chsy3 | 0.236575675 | 0.054760791 | -0.253204551 | -0.378519013 |
| ENSDART00000101143 | mhc1zea | 0.57323499 | 0.526487775 | 0.424884066 | 0.07882824 |
| ENSDART00000101204 | alcamb | 1.334306533 | 2.490758317 | 2.513428399 | 1.826617796 |
| ENSDART00000101208 | abhd11 | -0.157962098 | -0.321289806 | -0.312791151 | -0.149912405 |
| ENSDART00000101219 | mettl27 | 0.439044633 | 0.662527528 | 0.547984428 | 0.178832583 |
| ENSDART00000101231 | syt7b | -0.85705254 | -2.266705376 | -0.529986528 | 0.010282974 |
| ENSDART00000101265 | pik3c3 | 0.317460591 | 0.079259824 | 0.186108571 | 0.079398024 |
| ENSDART00000101282 | bcr | 0.172989029 | 0.500540591 | 0.708792261 | 0.385177654 |
| ENSDART00000101292 | si:dkey-238c7.16 | 0.35472996 | 0.602144814 | 0.450575535 | -0.163119012 |
| ENSDART00000101319 | zgc:162396 | 0.344606585 | 0.405244483 | 0.227818995 | 0.060602938 |
| ENSDART00000101394 | | -0.473756662 | -0.114672377 | -0.629243212 | -2.05184474 |
| ENSDART00000101477 | emp1 | -0.241905322 | -0.037268798 | 0.203715295 | 0.377786296 |
| ENSDART00000101513 | ppp6r2b | -0.18278033 | -0.275899098 | -0.242929069 | -0.103103109 |
| ENSDART00000101530 | | -0.360327211 | -0.388582556 | -0.376333338 | -0.16940017 |
| ENSDART00000101537 | mex3b | -0.20648165 | 0.480244705 | 0.713185315 | 0.514833911 |
| ENSDART00000101576 | tmem230b | 0.001436175 | -0.176978383 | -0.445323711 | -0.253116022 |
| ENSDART00000101577 | lrrfip1a | 0.656945169 | 0.402220137 | -0.125928686 | -0.548022342 |

| | | | | | |
|---|---|---|---|---|---|
| ENSDART00000101603 | kidins220b | -0.01985848 | -0.087358266 | 0.173253678 | 0.374974923 |
| ENSDART00000101627 | IGLON5 | -0.423175903 | -0.742308046 | -0.313681293 | 0.102626627 |
| ENSDART00000101631 | satb1b | -0.360112936 | -0.384116906 | 0.134197196 | 0.481946288 |
| ENSDART00000101653 | CU639469.1 | -0.529716959 | -0.505372118 | -0.050085906 | 0.195515782 |
| ENSDART00000101658 | ppp1r1b | -0.639757199 | -0.267805539 | -0.098160681 | -0.228099263 |
| ENSDART00000101698 | rpz3 | 0.288382306 | 0.347423497 | 0.479161923 | 0.327971533 |
| ENSDART00000101707 | dhx40 | 1.733776583 | 1.072360854 | 0.976040329 | 1.006521884 |
| ENSDART00000101789 | flot2b | 0.047673595 | 0.464453756 | 0.590584487 | 0.461504068 |
| ENSDART00000101943 | rragca | 0.302260739 | 0.264945177 | 0.40523597 | 0.261316897 |
| ENSDART00000101948 | GJA9 (1 of many) | -0.638779519 | -0.432316335 | -0.423080448 | -0.46097286 |
| ENSDART00000101974 | erh | 0.03890153 | -0.044950443 | -0.279975387 | -0.23272597 |
| ENSDART00000101982 | irg1 | 0.318182673 | 0.173060394 | -0.011525449 | -0.063563032 |
| ENSDART00000101985 | zgc:162944 | 0.341402057 | 0.426793325 | 0.183161821 | -0.332319729 |
| ENSDART00000102011 | G3BP2 (1 of many) | 0.183053703 | 0.292940696 | 0.414353657 | 0.391197687 |
| ENSDART00000102062 | timp2b | 0.538603085 | 0.084002119 | -0.039686845 | -0.313946909 |
| ENSDART00000102075 | rxrba | -5.032133145 | -2.347844506 | -3.410287183 | -0.630106553 |
| ENSDART00000102111 | tpt1 | 0.26125953 | 0.357793849 | 0.090393072 | -0.094022362 |
| ENSDART00000102125 | schip1 | -0.065634739 | -0.008431238 | -0.356995513 | -0.237294266 |
| ENSDART00000102148 | ddx3b | -0.056845376 | -0.031755759 | 0.246518412 | 0.169198996 |
| ENSDART00000102212 | tdp2a | 1.019634105 | 0.831370615 | 2.830207945 | 0.310574015 |
| ENSDART00000102214 | ndufv3 | -0.094484605 | -0.143659913 | -0.265916135 | -0.00411612 |
| ENSDART00000102260 | si:dkey-222f8.3 | 0.869989363 | 1.191058069 | 0.864781519 | 0.175715859 |
| ENSDART00000102279 | lingo2b | -0.319370755 | 0.274609796 | 0.545692974 | 0.078594547 |
| ENSDART00000102305 | cspg5a | -0.238742988 | 0.181941504 | 0.751580581 | 0.677972326 |
| ENSDART00000102368 | grin1a | -0.57807298 | -0.694484674 | -0.374619925 | 0.025284051 |
| ENSDART00000102384 | sesn2 | 0.221656214 | 0.707312613 | 0.831938385 | 0.650662976 |
| ENSDART00000102411 | dctn1b | 0.519457248 | 0.529061234 | 0.697567556 | 0.548863629 |
| ENSDART00000102419 | igf2bp2a | 0.598022036 | 0.736011973 | 0.87621145 | 0.419702221 |
| ENSDART00000102431 | CU571081.1 | 0.38462754 | 0.22357069 | 0.170924196 | -0.045496543 |
| ENSDART00000102434 | ehhadh | 0.70922182 | 0.453174195 | 0.063745289 | -0.562849237 |
| ENSDART00000102445 | clasp1a | -0.444419251 | -0.511465397 | -0.218813787 | -0.372606163 |
| ENSDART00000102455 | gucy1a3 | -0.52073984 | -0.506169014 | -0.27571082 | 0.101070807 |
| ENSDART00000102459 | rbp2a | 3.492314469 | 2.902874566 | 1.90821276 | 1.340524369 |
| ENSDART00000102461 | rgs8 | -0.536939066 | -0.575929959 | -0.463779418 | -0.057720257 |
| ENSDART00000102520 | palm1a | 0.447081887 | 0.96607967 | 1.214937327 | 0.924772733 |
| ENSDART00000102539 | st8sia5 | 0.153973405 | 0.592846857 | 1.061043515 | 0.777183018 |
| ENSDART00000102559 | zgc:122979 | -0.659472503 | -0.379774871 | -0.282725672 | -0.191125017 |
| ENSDART00000102562 | ankrd10b | -0.585231519 | -0.124121182 | 0.139197554 | 0.179030577 |
| ENSDART00000102567 | si:ch211-1o7.3 | 0.865836952 | 0.255817573 | 0.19284358 | -0.410198985 |
| ENSDART00000102665 | aste1a | 0.478587949 | 0.328068557 | 2.324198666 | 0.218780766 |
| ENSDART00000102672 | nck2a | 0.205356684 | 0.423877669 | 0.371996615 | 0.147611944 |
| ENSDART00000102681 | pnp5a | 0.202802193 | 0.499095293 | 0.029150263 | -0.463436134 |
| ENSDART00000102712 | tgm2a | 0.064390317 | 0.421175856 | 0.197637738 | 0.730671181 |
| ENSDART00000102715 | tuba8l3 | 1.032352614 | 1.505768874 | 1.918452626 | 1.697934809 |
| ENSDART00000102767 | fbxo9 | -0.10612083 | -0.188227567 | -0.252741487 | -0.179227033 |
| ENSDART00000102782 | gria2a | -0.472482927 | -0.595061919 | -0.203521695 | 0.050111807 |
| ENSDART00000102788 | epha7 | -0.240787156 | -0.359871475 | -0.404301639 | -0.038479801 |
| ENSDART00000102790 | glrba | -0.477875363 | -0.33299196 | -0.36632817 | -0.172063586 |
| ENSDART00000102791 | klhl31 | 1.919909064 | 3.64974722 | 3.354681727 | 2.995697572 |
| ENSDART00000102843 | src | 0.333102086 | 0.263030754 | 0.362117443 | 0.2047457 |
| ENSDART00000102846 | si:dkey-23a23.2 | 0.194667046 | 0.692432605 | 0.589344711 | 0.460081053 |

| | | | | | |
|---|---|---|---|---|---|
| ENSDART00000102868 | etnk2 | -0.119773861 | 0.158574082 | 0.41890552 | 0.249052751 |
| ENSDART00000102881 | fam43b | -0.413239873 | -0.283572592 | -0.358392622 | -0.273570929 |
| ENSDART00000102898 | zgc:158258 | -0.205000336 | -0.263168834 | -0.54350319 | -0.257007642 |
| ENSDART00000102903 | dmd | -0.080183076 | -0.276170831 | -0.483186786 | -0.073725636 |
| ENSDART00000102913 | cyp2v1 | 0.603838095 | 0.398439801 | -0.002186897 | -0.384510352 |
| ENSDART00000102952 | suz12a | 0.614499433 | 0.610664915 | 0.72051821 | 0.430761521 |
| ENSDART00000102969 | spock3 | -0.0552616 | -0.214726057 | -0.592493725 | -0.285702337 |
| ENSDART00000102981 | col8a1a | 1.700212586 | 0.444548046 | 0.772515926 | 0.395334934 |
| ENSDART00000103016 | zgc:173552 | 1.605722523 | 1.309975449 | 1.476680982 | 1.096088035 |
| ENSDART00000103043 | nsfa | -0.472055845 | -0.418381067 | -0.313792962 | -0.037512478 |
| ENSDART00000103070 | cdk17 | 0.018864851 | -0.23502559 | -0.366641807 | -0.137520544 |
| ENSDART00000103076 | arl8bb | 0.705280948 | 0.466975375 | 0.584557674 | 0.096505859 |
| ENSDART00000103151 | dlgap3 | -0.304185948 | -0.430889474 | -0.103996234 | 0.188728565 |
| ENSDART00000103267 | fam212ab | -0.276107898 | -0.473876533 | -0.614056911 | -0.421775209 |
| ENSDART00000103293 | ndufa5 | -0.136898371 | -0.38263765 | -0.527827076 | -0.351287298 |
| ENSDART00000103352 | vps18 | 0.165861852 | 0.202619769 | 0.265495404 | 0.159870348 |
| ENSDART00000103365 | ociad1 | -0.01475249 | -0.099734574 | -0.308153486 | -0.083780451 |
| ENSDART00000103368 | rpl22 | 0.33350855 | 0.471741899 | 0.142215466 | -0.053336675 |
| ENSDART00000103385 | slc25a22 | -0.5158616 | -0.319809073 | -0.784039688 | -0.881615638 |
| ENSDART00000103405 | gch1 | -0.102362632 | -0.124839141 | -0.313487955 | 0.056848423 |
| ENSDART00000103407 | tmem245 | -0.353429601 | -0.331655767 | -0.089201446 | 0.066435984 |
| ENSDART00000103448 | tbx18 | 1.78643571 | 2.579858492 | 2.64743129 | 1.939376924 |
| ENSDART00000103450 | lactbl1b | -0.310148031 | -0.346227281 | -0.611014657 | -0.245438949 |
| ENSDART00000103463 | dnajc2 | 0.193851448 | 0.30912042 | 0.441518195 | 0.223806707 |
| ENSDART00000103467 | zgc:77650 | 0.216349316 | 0.315934081 | 0.173077104 | -0.055613825 |
| ENSDART00000103471 | khdrbs1b | -0.296913345 | -0.271356827 | -0.067247312 | 0.118087564 |
| ENSDART00000103474 | tspan13b | -0.348967086 | -0.119531637 | 0.084177739 | 0.198302304 |
| ENSDART00000103487 | zgc:195001 | -0.335765581 | -0.344851478 | -0.707965666 | -0.31393888 |
| ENSDART00000103491 | rbp7b | 2.051254968 | 1.285649693 | 0.558375641 | -0.074104197 |
| ENSDART00000103526 | bicc1b | 0.362058268 | 0.640014631 | 0.968410254 | 0.365236005 |
| ENSDART00000103532 | kcnh5a | -0.481841968 | -0.667651405 | -0.403869416 | -0.116032422 |
| ENSDART00000103549 | skib | -0.170611371 | -0.408422527 | -0.193614247 | -0.134954723 |
| ENSDART00000103586 | hdac9b | -0.63802499 | -0.480501706 | -0.6152094 | -0.590506452 |
| ENSDART00000103588 | mxa | 0.174383297 | -0.012993595 | 1.447594397 | 0.019311963 |
| ENSDART00000103602 | lgals2a | 1.886236295 | 1.628346862 | 1.325259297 | 0.487532368 |
| ENSDART00000103622 | irf7 | -0.049332142 | -0.2878598 | 1.533925917 | -0.275107455 |
| ENSDART00000103626 | mief2 | -0.089955279 | -0.188542146 | -0.285241779 | -0.158709862 |
| ENSDART00000103628 | btbd6a | -0.278201553 | -0.439382236 | -0.547351526 | -0.157769223 |
| ENSDART00000103639 | arf3a | 0.291954045 | 0.68170241 | 0.689772164 | 0.341898272 |
| ENSDART00000103640 | hey1 | -0.385659584 | -0.747117397 | -0.392306381 | -0.416692063 |
| ENSDART00000103646 | kcng2 | -0.538883844 | -0.646084515 | -0.39646557 | 0.008271529 |
| ENSDART00000103660 | clcn7 | 0.412276024 | 0.162279088 | -0.074638152 | -0.273644307 |
| ENSDART00000103704 | nap1l4a | 0.303924114 | 0.290352176 | 0.138900988 | 0.027879829 |
| ENSDART00000103750 | fam131bb | -0.555314971 | -0.519589863 | -0.143114917 | 0.156539518 |
| ENSDART00000103753 | fn1a | 2.705855593 | 3.095500526 | 2.8206804 | 2.414837328 |
| ENSDART00000103754 | fn1b | 2.49600075 | 3.130597747 | 1.885882871 | -0.148711913 |
| ENSDART00000103755 | fn1b | 4.126396153 | 4.640072685 | 4.590004259 | 2.904538972 |
| ENSDART00000103785 | ggact.3 | -1.209900002 | -0.367470534 | -1.050388105 | -0.531293993 |
| ENSDART00000103795 | ggact.1 | 0.007439004 | -0.050660849 | -0.406911538 | -0.573615848 |
| ENSDART00000103815 | stmn2a | 0.140137036 | 1.176009379 | 1.538144395 | 1.355356024 |
| ENSDART00000103831 | ano10b | 0.06580331 | 0.43549934 | 0.640553067 | 0.460128123 |

| | | | | | |
|---|---|---|---|---|---|
| ENSDART00000103878 | vhl | -0.19558438 | -0.426476861 | -0.33011461 | -0.126927905 |
| ENSDART00000103894 | cacng8b | -0.589374624 | -0.680390231 | -0.379441318 | -0.105621743 |
| ENSDART00000103911 | CABZ01077217.1 | 0.385992878 | 0.554652123 | 0.464993777 | 0.2375762 |
| ENSDART00000103922 | atat1 | 0.145026536 | 0.665460259 | 0.937418253 | 0.683690675 |
| ENSDART00000103940 | mgat1b | -0.294073166 | -0.13896492 | -0.278393093 | -0.224242354 |
| ENSDART00000103957 | exoc3l1 | 0.634582138 | 1.052789014 | 1.334761372 | 0.478611156 |
| ENSDART00000103969 | magi1b | -0.261202469 | -0.221356443 | 0.045512769 | 0.18273538 |
| ENSDART00000103980 | crip2 | -0.517107368 | -0.364195263 | -0.546778441 | -0.267901424 |
| ENSDART00000103982 | nfe2l1a | 0.652859432 | 0.675562602 | 0.38648833 | -0.18398816 |
| ENSDART00000103992 | ttll3 | -0.156492299 | -0.149773154 | -0.299786209 | -0.201934967 |
| ENSDART00000104008 | dtnbp1b | -0.424284075 | -0.457257716 | -0.424822836 | -0.199214695 |
| ENSDART00000104009 | slc30a1b | 0.469731317 | 0.847749615 | 0.68298124 | 0.247669117 |
| ENSDART00000104027 | rims3 | -0.345163808 | -0.70584123 | -0.314103014 | -0.01029762 |
| ENSDART00000104043 | dkk3b | 0.124905707 | -0.440538945 | -0.566305744 | -1.06504734 |
| ENSDART00000104058 | ncalda | 0.174416215 | 0.325917683 | 0.093263441 | -0.133323566 |
| ENSDART00000104135 | gabra5 | -0.398056951 | -0.510180941 | 0.262181183 | 0.751061332 |
| ENSDART00000104140 | dyrk4 | 0.302696779 | -0.345665012 | -0.79115152 | 0.018427723 |
| ENSDART00000104188 | igsf21b | -0.476667576 | -0.274511883 | -0.073843861 | 0.013583331 |
| ENSDART00000104234 | gjd2b | -0.226334837 | -0.451484904 | -0.506629051 | -0.204050045 |
| ENSDART00000104257 | lpin1 | -0.263973453 | -0.162876624 | -0.357103753 | -0.3626544 |
| ENSDART00000104279 | znf516 | -0.151066862 | -0.30868422 | -0.587761132 | -0.246031248 |
| ENSDART00000104289 | rab3ab | -0.744433679 | -0.710299004 | -0.400971594 | -0.021261155 |
| ENSDART00000104293 | pex5la | -0.678333589 | -0.710207201 | -0.577955484 | -0.480904778 |
| ENSDART00000104298 | ndufb5 | -0.141095269 | -0.212773777 | -0.353488244 | -0.122828248 |
| ENSDART00000104299 | cnp | 0.814960421 | 1.713687269 | 2.20652149 | 1.77606709 |
| ENSDART00000104307 | eml1 | 0.215532654 | -0.343903678 | -0.604837308 | -0.271121591 |
| ENSDART00000104317 | klf7b | 0.389436838 | 1.156291561 | 1.140404424 | 0.652703479 |
| ENSDART00000104322 | uchl3 | 0.256651569 | 0.374538377 | 0.322263704 | 0.147277389 |
| ENSDART00000104327 | vsx1 | -0.248525466 | -0.619641572 | -0.034766195 | -0.028538746 |
| ENSDART00000104336 | tnnc2 | 0.703924131 | 2.455705453 | 2.232729877 | 1.561721699 |
| ENSDART00000104338 | pitpnbl | 0.001600172 | 0.291216057 | 0.176136036 | 0.011592783 |
| ENSDART00000104353 | atp2b1b | -0.132002912 | -0.283527357 | -0.37922998 | -0.020226682 |
| ENSDART00000104361 | si:dkey-56d12.4 | 0.671595911 | 0.775149092 | 0.509958528 | 0.270735073 |
| ENSDART00000104364 | rps15 | 0.338617661 | 0.309197703 | 0.255669723 | -0.012990535 |
| ENSDART00000104423 | rbbp9 | 0.243640055 | 0.525148359 | 0.333645697 | 0.149282905 |
| ENSDART00000104475 | tmsb4x | 0.344214142 | 0.313013147 | 0.275125575 | 0.128255357 |
| ENSDART00000104478 | fxr1 | 0.018053797 | 0.098113697 | 0.377704225 | 0.105513892 |
| ENSDART00000104481 | slc17a7a | -0.380361502 | -0.377656856 | -0.259723773 | -0.186490981 |
| ENSDART00000104487 | cox4i2 | 0.027176247 | -0.218042175 | -0.688187338 | -0.440124172 |
| ENSDART00000104496 | dusp6 | 0.185513151 | -0.01084107 | 0.164485744 | 0.322445555 |
| ENSDART00000104519 | stat3 | 0.6458865 | 0.459098575 | 0.438501049 | 0.201812376 |
| ENSDART00000104520 | cnbpb | -0.203757271 | -0.252097684 | -0.248932139 | -0.261165322 |
| ENSDART00000104523 | arntl1b | -0.158144123 | 0.59291526 | 1.113237486 | 0.52944604 |
| ENSDART00000104525 | kmt2a | 0.050001667 | 0.172870774 | 0.277996699 | 0.044684436 |
| ENSDART00000104536 | chsy1 | 0.021777196 | -0.175630111 | -0.644756219 | -0.079001152 |
| ENSDART00000104545 | avpr2ab | -0.127228111 | -0.325246207 | -0.456539735 | -1.288721053 |
| ENSDART00000104576 | psmd13 | 0.182320821 | 0.32294371 | 0.367344771 | 0.118433419 |
| ENSDART00000104592 | pm20d1.2 | -0.022497234 | -0.114097556 | -0.391118153 | -0.365853437 |
| ENSDART00000104612 | hmx3a | -0.260599707 | -0.468562174 | -0.112652566 | -0.078307838 |
| ENSDART00000104616 | lepr | 0.762949533 | 0.720319043 | 0.625239528 | 0.526452567 |
| ENSDART00000104637 | si:ch211-81a5.8 | 0.707959832 | 0.064083353 | -0.292232881 | -0.006669302 |

| | | | | | |
|---|---|---|---|---|---|
| ENSDART00000104657 | mrps21 | -0.148781558 | -0.231390216 | -0.424136936 | -0.351598574 |
| ENSDART00000104673 | camk1gb | -0.378218392 | -0.417447419 | 0.009394779 | -0.088307876 |
| ENSDART00000104674 | camk1db | -0.318172003 | -0.339895825 | -0.49572727 | -0.229789777 |
| ENSDART00000104687 | pfn2 | 0.330558178 | 0.38737784 | 0.240819292 | -0.058450663 |
| ENSDART00000104708 | abhd12 | -0.134054463 | -0.304895231 | -0.501902497 | -0.262631127 |
| ENSDART00000104712 | hmg20a | 0.003064442 | 0.120221197 | 0.366113134 | 0.275609671 |
| ENSDART00000104720 | si:ch1073-385f13.3 | 0.258133799 | 0.237616235 | 1.745458762 | -0.119672944 |
| ENSDART00000104722 | cdk5r2a | -0.371112934 | -0.317321269 | -0.137896092 | -0.229143186 |
| ENSDART00000104728 | ssuh2.2 | 0.252481495 | 0.103488209 | 1.66757315 | -0.204612589 |
| ENSDART00000104730 | ifit14 | 0.457651023 | 0.403391968 | 2.518119653 | 0.537556431 |
| ENSDART00000104750 | mllt11 | -0.266364682 | 0.400357279 | 0.342752209 | 0.193353613 |
| ENSDART00000104751 | mxi1 | -0.384587031 | -0.210909301 | 0.22179917 | 0.104111801 |
| ENSDART00000104791 | zgc:153722 | 0.356828804 | 0.479757966 | 0.536710875 | 0.133740974 |
| ENSDART00000104828 | gtpbp2b | -0.421398738 | -0.385176207 | -0.273058228 | -0.083654585 |
| ENSDART00000104835 | hps5 | 0.972059444 | 0.583168765 | 0.57443346 | -0.161192354 |
| ENSDART00000104845 | tapbp.1 | 3.172722647 | 3.246630579 | 3.542395564 | 2.955685304 |
| ENSDART00000104866 | ldhbb | -0.207883318 | -0.435615938 | -0.519651289 | -0.183614515 |
| ENSDART00000104895 | rgs7a | -0.76773786 | -0.54211529 | -0.053731925 | 0.298916153 |
| ENSDART00000104933 | eepd1 | -0.181346781 | -0.362247495 | -0.01782264 | 0.011720383 |
| ENSDART00000104950 | atp1a3a | -0.469887861 | -0.514207298 | -0.064097938 | 0.181102676 |
| ENSDART00000104999 | ccdc85ca | -0.136040811 | -0.297438948 | -0.211938794 | -0.169428565 |
| ENSDART00000105120 | tmem170b | -0.08312029 | 0.138452609 | 0.437118287 | 0.263377377 |
| ENSDART00000105174 | stm | 3.414215545 | 0.041843024 | -0.026881567 | -0.531574221 |
| ENSDART00000105179 | zgc:158254 | -0.229665852 | -0.301892648 | -0.223905123 | -0.049497491 |
| ENSDART00000105215 | emilin1b | 1.074845253 | 1.078203948 | 0.684747395 | -0.035555204 |
| ENSDART00000105323 | zgc:162255 | -0.250927131 | -0.470845634 | -0.44137647 | -0.557741796 |
| ENSDART00000105399 | cbarpb | 0.427437854 | 0.525694666 | 0.646908744 | 0.015511077 |
| ENSDART00000105404 | cirbpb | 0.178847577 | 0.103859165 | 0.267594995 | 0.007629823 |
| ENSDART00000105405 | cirbpb | 0.001082658 | 0.185977938 | 0.46521688 | -0.018730312 |
| ENSDART00000105477 | lrrtm2 | -0.540265042 | -0.543880814 | -0.470126158 | -0.070951006 |
| ENSDART00000105484 | si:ch211-216b21.2 | -0.960990997 | -0.619204359 | -0.135410073 | 0.236742995 |
| ENSDART00000105485 | si:dkey-78l4.14 | -0.103336903 | -0.24562657 | -0.381336593 | -0.194197896 |
| ENSDART00000105503 | rab3gap2 | 2.422276289 | 3.562211475 | 1.182834682 | 2.012188356 |
| ENSDART00000105545 | arl3 | -0.028403306 | -0.158533169 | -0.38830015 | -0.193573844 |
| ENSDART00000105561 | si:dkey-42p8.3 | -0.126414242 | -0.359710173 | -0.388006213 | -0.105772793 |
| ENSDART00000105588 | pcdh1b | -0.246217756 | -0.446101289 | -0.039720059 | 0.09005245 |
| ENSDART00000105597 | si:ch211-129c21.1 | 3.291282195 | 4.994402871 | 3.669154828 | 0.685912998 |
| ENSDART00000105602 | elovl6 | -0.283618771 | -0.171237628 | -0.388002017 | -0.174233768 |
| ENSDART00000105608 | prkcea | -0.17407648 | -0.282581987 | -0.238186172 | -0.104981942 |
| ENSDART00000105659 | mhc1zja | 0.668521736 | 0.771454181 | 0.462558446 | 0.066622239 |
| ENSDART00000105667 | si:dkey-121a11.3 | -0.369671493 | -0.58967553 | -0.842073302 | -0.52980672 |
| ENSDART00000105681 | cdc14ab | -0.104501589 | -0.239167071 | -0.39502699 | -0.259112662 |
| ENSDART00000105694 | adprm | 0.302167589 | 0.521514455 | 0.908630272 | 0.271324554 |
| ENSDART00000105743 | ntng1a | -0.262343031 | -0.343435025 | -0.06227485 | 0.248931143 |
| ENSDART00000105749 | nrros | 0.925801893 | 0.55337814 | 0.265308462 | -0.351361566 |
| ENSDART00000105753 | olfm3a | -0.365975807 | -0.465135675 | -0.139223772 | 0.043379292 |
| ENSDART00000105767 | fhl1a | 0.562168934 | 1.496722819 | 1.339494009 | 0.93265064 |
| ENSDART00000105774 | ek1 | -0.422572147 | -0.621215995 | -0.219935537 | -0.023014864 |
| ENSDART00000105813 | krt222 | -0.522679169 | -0.470823148 | -0.650543043 | -0.842913393 |
| ENSDART00000105818 | guk1a | 0.053300944 | 0.162526517 | 0.245626369 | 0.043825221 |
| ENSDART00000105848 | si:dkey-42i9.7 | 0.176745483 | 0.033035183 | 1.603344014 | -0.243326761 |

| | | | | | |
|---|---|---|---|---|---|
| ENSDART00000105854 | josd1 | -0.238139004 | 0.044983845 | 0.263174427 | 0.210087455 |
| ENSDART00000105866 | si:ch73-213k20.5 | -0.515754761 | -0.874847543 | -0.892891327 | -0.392973916 |
| ENSDART00000105873 | cry4 | -0.404336974 | 0.122755184 | 0.671823255 | 0.515326773 |
| ENSDART00000105878 | fblim1 | 1.656130522 | 1.752691305 | 1.013125802 | 0.616877976 |
| ENSDART00000105896 | atp2a1l | -1.028985284 | 2.5381413 | 3.515981153 | 2.247616466 |
| ENSDART00000105898 | TSTD1 | 0.614724746 | 0.98570488 | 0.724640457 | 0.337367855 |
| ENSDART00000105903 | lsm5 | 0.135971292 | -0.078983757 | -0.333257253 | -0.331997667 |
| ENSDART00000105932 | si:dkeyp-110e4.11 | -0.775118699 | -0.662012678 | -0.111191342 | 0.348802036 |
| ENSDART00000105942 | si:dkey-253d23.3 | 0.99314399 | 1.287434296 | 0.804068321 | 0.742353386 |
| ENSDART00000105967 | cacna2d1a | -0.039509635 | -0.246396203 | -0.304011671 | -0.176720638 |
| ENSDART00000105974 | pvalb9 | -0.398960737 | -0.428084025 | -0.618672138 | -0.278532192 |
| ENSDART00000105988 | si:ch211-156p11.1 | 0.766312966 | 0.897687307 | 0.740122106 | 0.441957966 |
| ENSDART00000106048 | ctnnd1 | 0.632970644 | 0.745690204 | 0.779606908 | 0.274224061 |
| ENSDART00000106081 | BX511034.2 | 0.124750693 | 0.347355362 | 0.433795514 | 0.117013159 |
| ENSDART00000106096 | drd1a | -0.279465728 | -0.508847304 | -0.2349577 | -0.295766526 |
| ENSDART00000106120 | ywhag2 | 0.097311782 | 0.334968358 | 0.584303472 | 0.612363287 |
| ENSDART00000106141 | tma7 | 0.110821999 | -0.104393908 | -0.337001227 | -0.221432499 |
| ENSDART00000106152 | top2a | 1.478970046 | 0.796399019 | 1.005046229 | 0.716311641 |
| ENSDART00000106166 | rx1 | -1.001905061 | -0.358282402 | 0.191592029 | -0.031411117 |
| ENSDART00000106172 | rac1a | 0.326396929 | 0.503439113 | 0.328932342 | 0.053159748 |
| ENSDART00000106186 | prex2 | -0.375401587 | -0.276106515 | -0.274390577 | -0.295066887 |
| ENSDART00000106198 | zgc:66433 | 0.123524201 | -0.419463467 | -0.600279473 | -0.27606853 |
| ENSDART00000106229 | zgc:113886 | -1.108782489 | -3.592108749 | -0.501312332 | -0.527928714 |
| ENSDART00000106260 | ankha | 0.083956215 | -0.292178887 | -0.396040739 | -0.042168676 |
| ENSDART00000106566 | CABZ01030107.1 | 2.449267078 | 1.581540517 | 1.3239843 | 0.803203173 |
| ENSDART00000106619 | nrn1a | -1.225032386 | -1.272422646 | -0.361032349 | 0.032061464 |
| ENSDART00000106680 | rai1 | -0.196842736 | -0.463425257 | -0.402954237 | -0.12043583 |
| ENSDART00000106690 | pcdh2g7 | 0.533636914 | 0.291763532 | 0.532651571 | 0.481339584 |
| ENSDART00000106704 | pcdh2ab1 | -0.111727697 | 0.203588847 | 0.448801307 | 0.505664817 |
| ENSDART00000108471 | ipo7 | 0.142918554 | 0.150506384 | 0.285667017 | 0.157707046 |
| ENSDART00000108493 | gareml | 2.608537205 | 3.376838547 | 3.284344596 | 2.814340641 |
| ENSDART00000108496 | clec11a | -1.064574084 | -0.301046329 | -0.192981843 | -0.515534742 |
| ENSDART00000108507 | CSMD3 | -0.201722796 | -0.640032982 | -0.257154173 | 0.075603806 |
| ENSDART00000108535 | gabrb1 | -1.10767094 | -1.249599593 | -0.238929019 | 0.417524871 |
| ENSDART00000108574 | gramd1bb | -0.232214076 | -0.405839915 | -0.354417408 | -0.144413075 |
| ENSDART00000108581 | si:dkey-17m8.1 | 0.789679514 | 0.574820222 | 0.300394764 | -0.045841382 |
| ENSDART00000108596 | zgc:162150 | 0.993292495 | 1.420859127 | 1.257973066 | 0.026357222 |
| ENSDART00000108629 | lrch1 | -0.183622444 | -0.188824595 | -0.180380725 | -0.322184411 |
| ENSDART00000108655 | RHOBTB2 (1 of many) | -0.407905552 | -0.427681874 | -0.180479384 | -0.131684888 |
| ENSDART00000108729 | adamtsl2 | -0.842317462 | -0.70962753 | 0.040474933 | 0.841307736 |
| ENSDART00000108736 | adam12 | -0.319460786 | -0.316431452 | 0.445307645 | 0.87790832 |
| ENSDART00000108796 | stox2b | 0.143615187 | 0.758742406 | 1.120462456 | 0.794663921 |
| ENSDART00000108804 | brinp1 | -0.191216427 | -0.292751478 | -0.180680558 | 0.046514423 |
| ENSDART00000108808 | frem2a | -0.381822233 | -0.539670066 | 0.662155423 | 0.974116851 |
| ENSDART00000108814 | nrip2 | -0.213998541 | -0.302896779 | -0.276048404 | -0.173620142 |
| ENSDART00000108818 | arhgap32a | -0.576365369 | -0.434660146 | -0.258610983 | 0.095307278 |
| ENSDART00000108831 | espnla | 0.888146475 | 0.74973908 | 0.792131149 | 0.719500985 |
| ENSDART00000108916 | SIPA1 | -0.25031723 | -0.283192607 | 0.116871701 | 0.232178471 |
| ENSDART00000108928 | pnpla8 | -0.24833123 | -0.178721384 | -0.052341018 | -0.031698962 |
| ENSDART00000108943 | shroom2a | 0.03299057 | -0.127271939 | -0.499446332 | -0.229050052 |
| ENSDART00000108959 | RNF208 | -0.615021133 | -0.757500788 | -0.372077015 | -0.309696014 |

| | | | | | |
|---|---|---|---|---|---|
| ENSDART00000108963 | CABZ01102109.1 | 0.394501993 | 0.771748701 | 0.296144389 | 0.075524582 |
| ENSDART00000108989 | adamts14 | -0.185318243 | -0.40325002 | -0.417840329 | -0.182643804 |
| ENSDART00000108990 | pex5lb | -0.323313621 | -0.539021494 | -0.34249137 | -0.198103063 |
| ENSDART00000108995 | tex2 | 0.515345319 | 0.756490823 | 0.434164856 | 0.106684566 |
| ENSDART00000109014 | GJC1 | -0.38880893 | -0.552238053 | -0.31502822 | -0.300328618 |
| ENSDART00000109017 | usp31 | -0.203091269 | -0.25315355 | -0.034862909 | 0.058179546 |
| ENSDART00000109023 | fmnl3 | -0.186687955 | -0.312847961 | -0.533074314 | -0.280720309 |
| ENSDART00000109029 | map6a | -0.330392405 | -0.354970167 | -0.034172445 | 0.05340091 |
| ENSDART00000109031 | gpr26 | -0.593199232 | -0.45461936 | -0.805315901 | -0.585023693 |
| ENSDART00000109037 | cdk5r2b | -0.48847507 | -0.44246865 | -0.592226601 | -0.418268347 |
| ENSDART00000109040 | gpr158a | -0.331785981 | -0.632662732 | -0.236983197 | 0.105308258 |
| ENSDART00000109044 | grm8a | -0.470031444 | -0.746629562 | -0.459544685 | -0.031467357 |
| ENSDART00000109062 | nckap5l | -0.29308537 | -0.083948409 | 0.248864543 | 0.105880197 |
| ENSDART00000109065 | ccng1 | 0.498527324 | 0.814436551 | 0.266438219 | -0.054038693 |
| ENSDART00000109070 | gigyf1b | -0.072846918 | -0.184795854 | -0.274652224 | -0.080632892 |
| ENSDART00000109091 | mettl1 | 1.65076712 | 2.831574782 | 2.963448936 | 2.721211005 |
| ENSDART00000109099 | ttn.2 | -4.072510837 | 1.236801213 | 2.302382204 | 1.171450021 |
| ENSDART00000109124 | rspo2 | 0.077521406 | 0.271943255 | -0.228972258 | -0.854369599 |
| ENSDART00000109138 | hbegfa | 0.956645779 | 1.991576979 | 1.27153829 | 0.330229075 |
| ENSDART00000109147 | brsk2b | -0.075048445 | -0.279504807 | -0.192446814 | 0.052331491 |
| ENSDART00000109171 | rce1a | -0.218282831 | -0.358569107 | -0.271597779 | -0.119151709 |
| ENSDART00000109224 | CCKBR (1 of many) | -0.56783642 | -0.51463677 | -0.498556224 | -0.286398879 |
| ENSDART00000109235 | bicd2 | 0.048329883 | -0.336297195 | -0.602745642 | -0.296802706 |
| ENSDART00000109243 | sema4bb | -0.591195487 | -0.421350422 | -0.25044186 | -0.118505649 |
| ENSDART00000109249 | cenpo | 3.029141821 | 1.947037559 | 2.256164745 | 0.98054484 |
| ENSDART00000109252 | nudt5 | 0.602324776 | 0.61383313 | 0.580277921 | 0.203918347 |
| ENSDART00000109257 | RASGRF1 | -0.570969407 | -0.754386918 | -0.426488142 | -0.098297531 |
| ENSDART00000109259 | smarcc1a | 0.060519576 | 0.104503692 | 0.246223049 | 0.143325093 |
| ENSDART00000109288 | myo16 | 0.115485168 | 0.089369719 | 0.360073499 | 0.534996054 |
| ENSDART00000109308 | si:ch73-60h1.1 | -0.222101504 | -0.315217201 | -0.340339223 | -0.206722759 |
| ENSDART00000109314 | arhgap10 | -0.214061464 | -0.171864796 | -0.398074347 | -0.037252057 |
| ENSDART00000109319 | arf1 | 0.284053831 | 0.377774948 | 0.307132687 | 0.034662092 |
| ENSDART00000109356 | klf7a | -0.7063233 | -0.659659661 | -0.044315321 | 0.157900055 |
| ENSDART00000109415 | blzf1 | -4.190621432 | -2.580757916 | 0.161069597 | -0.394790047 |
| ENSDART00000109416 | smc2 | 1.321753383 | 1.108242465 | 0.857262832 | 0.308883411 |
| ENSDART00000109420 | kcnip1b | -0.712749849 | -0.701262469 | -0.816924162 | -0.561177799 |
| ENSDART00000109425 | nkd3 | 3.023582344 | 2.734865433 | 2.70294964 | 2.317338999 |
| ENSDART00000109432 | cercam | 0.726258015 | 0.540687546 | 0.072936326 | -0.232601995 |
| ENSDART00000109440 | adamts9 | -0.162680626 | -0.21784206 | -0.326391297 | -0.145144869 |
| ENSDART00000109452 | sbf1 | -0.62469158 | -0.583723087 | 0.106179087 | 0.127879066 |
| ENSDART00000109464 | g0s2 | 2.993222235 | 1.499538045 | 0.588261277 | 0.527162957 |
| ENSDART00000109485 | gal3st3 | -0.139388785 | -0.282505485 | -0.388230748 | -0.163229068 |
| ENSDART00000109486 | gprc5ba | -0.628238673 | -0.569388482 | -0.508289237 | -0.226450363 |
| ENSDART00000109497 | tbxas1 | 0.141369479 | -0.217979555 | -0.576549291 | -0.268338462 |
| ENSDART00000109507 | RAP1GDS1 | -0.399835735 | -0.351308804 | 0.076766568 | 0.119897865 |
| ENSDART00000109511 | si:ch211-186j3.6 | -0.164289524 | -0.357536961 | -0.081167433 | 0.02827477 |
| ENSDART00000109528 | adgre5b.3 | 0.525818313 | 0.538353042 | 0.246499359 | -0.175742704 |
| ENSDART00000109535 | gmps | 0.181071124 | 0.257853381 | 0.376587367 | 0.112990813 |
| ENSDART00000109537 | snphb | -0.466057723 | -0.636087511 | -0.318006337 | -0.040021342 |
| ENSDART00000109546 | unc5b | -0.076752071 | -0.101810869 | -0.423560106 | -0.292578205 |
| ENSDART00000109552 | baz2ba | 0.627489244 | 0.525669422 | 0.712410774 | 0.393101067 |

| | | | | | |
|---|---|---|---|---|---|
| ENSDART00000109567 | nhsa | -0.05015222 | -0.331046598 | -0.348641258 | -0.190382203 |
| ENSDART00000109568 | pip4k2ab | -0.31785183 | -0.37156543 | -0.209120418 | 0.064881056 |
| ENSDART00000109570 | | -0.315520739 | -0.355339215 | -1.137398742 | -0.355379016 |
| ENSDART00000109573 | akap6 | 0.927127599 | 1.241947706 | 1.04133516 | 0.789730231 |
| ENSDART00000109581 | abtb1 | -0.353930406 | -0.557673054 | -0.197302182 | -0.044180273 |
| ENSDART00000109604 | palmdb | -0.228743892 | -0.477529612 | -0.779896236 | -0.307691368 |
| ENSDART00000109615 | tmem255a | 0.198392818 | 0.721353886 | 0.92551731 | 0.71998369 |
| ENSDART00000109698 | nacc1b | -0.541808458 | -0.507646097 | -0.425509734 | -0.201090141 |
| ENSDART00000109714 | usp53b | -0.259276756 | -0.08547014 | -0.255057866 | -0.127997302 |
| ENSDART00000109732 | amigo1 | -0.336374535 | -0.536025942 | -0.442802906 | -0.174075192 |
| ENSDART00000109750 | rap2aa | 0.293487605 | 0.482823125 | 0.662590656 | 0.396980886 |
| ENSDART00000109752 | serpinh1a | 0.578805227 | 0.380412306 | 0.080680913 | -0.569755212 |
| ENSDART00000109759 | tmx2a | 2.647169883 | 3.21425121 | 2.659532867 | 1.569734674 |
| ENSDART00000109807 | EMILIN3 | 0.322021142 | 0.194240185 | 0.000409902 | -0.0946206 |
| ENSDART00000109822 | hps1 | 0.431913269 | 0.483030578 | 0.349219151 | -0.038818149 |
| ENSDART00000109831 | BX088653.1 | 0.96716269 | 0.47589947 | 0.338658984 | -0.013815104 |
| ENSDART00000109833 | zmat3 | -0.462380241 | -0.737159777 | -0.151865097 | 0.090633241 |
| ENSDART00000109853 | zmp:0000000801 | -0.4485216 | -0.462342097 | -0.503236916 | -0.960013508 |
| ENSDART00000109872 | cln6a | 0.07921086 | -0.018544414 | -0.27320282 | -0.083540486 |
| ENSDART00000109876 | sun1 | -0.255190881 | -0.339472925 | -0.466317308 | -0.169389668 |
| ENSDART00000109973 | tp53bp1 | 0.061139928 | 0.539512942 | 0.657824217 | 0.222950524 |
| ENSDART00000109990 | abhd15a | -0.466192052 | -0.318128979 | -0.316342009 | -0.301397533 |
| ENSDART00000110004 | frmpd3 | -0.530320903 | -0.511652794 | -0.387982186 | 0.071931622 |
| ENSDART00000110005 | cdhr1a | -0.099288941 | -0.157780229 | -0.305559135 | -0.101115232 |
| ENSDART00000110016 | ube2ql1 | -0.274241146 | -0.513839036 | -0.485900432 | -0.131799131 |
| ENSDART00000110033 | abhd8b | -0.069420963 | -0.144372686 | -0.35601389 | -0.098305508 |
| ENSDART00000110040 | sox11a | 1.573346229 | 1.723237215 | 1.23195588 | 0.781584101 |
| ENSDART00000110041 | lrrc38a | -0.167375954 | -0.314913504 | -0.608219505 | -0.293673506 |
| ENSDART00000110061 | spock2 | -0.746309396 | -0.624301954 | -0.158480844 | 0.116789254 |
| ENSDART00000110064 | plaua | 1.293433748 | 1.730730518 | 1.0162544 | 0.507140212 |
| ENSDART00000110069 | CABZ01075131.1 | -0.117445692 | -0.062527677 | 0.307574487 | 0.429328576 |
| ENSDART00000110076 | gas1a | -0.414278752 | -0.328067296 | -0.317423655 | -0.582681301 |
| ENSDART00000110080 | aatka | -0.15444172 | -0.531359629 | -0.399426039 | -0.066680489 |
| ENSDART00000110092 | mdga1 | -0.307586742 | -0.496736277 | -0.237358341 | -0.048338544 |
| ENSDART00000110100 | hint3 | 0.280351117 | 0.473623985 | 0.484346588 | 0.216760955 |
| ENSDART00000110126 | cacng3b | -0.394325343 | -0.354117689 | -0.012321564 | 0.185829622 |
| ENSDART00000110136 | cntnap5b | -0.584561492 | -0.553235513 | -0.330365654 | 0.070638148 |
| ENSDART00000110204 | pikfyve | 0.281293639 | 0.125670374 | 0.334758542 | 0.135267095 |
| ENSDART00000110224 | mon1bb | -0.275995109 | -0.242724773 | -0.120595351 | -0.023350049 |
| ENSDART00000110241 | cnksr2a | -0.309766586 | -0.445156957 | -0.12266438 | -0.140501824 |
| ENSDART00000110270 | pwp2h | 0.562435944 | 0.598132459 | 0.373677258 | 0.062993981 |
| ENSDART00000110278 | rassf9 | -0.213149801 | -0.230795795 | -0.114749715 | -0.801660509 |
| ENSDART00000110279 | si:dkey-183c6.8 | -0.455316373 | -0.174866477 | 0.555526708 | 0.790514442 |
| ENSDART00000110331 | PTGFRN (1 of many) | -0.468065048 | -0.642664771 | -0.293257997 | -0.171150885 |
| ENSDART00000110370 | nrcama | 0.101341326 | 0.125318979 | 0.288644016 | 0.298095665 |
| ENSDART00000110377 | zgc:194398 | -0.335442291 | -0.298995581 | -0.514027337 | -0.216616028 |
| ENSDART00000110383 | map7a | -0.540958501 | -0.669475763 | -0.490700501 | -0.102714716 |
| ENSDART00000110409 | frmd5 | -0.554751162 | -0.59840736 | -0.384150671 | 0.071106799 |
| ENSDART00000110411 | cerkl | -0.014858902 | -0.169171069 | -0.284528511 | -0.188701157 |
| ENSDART00000110416 | eif2b5 | 0.075177684 | 0.238615258 | 0.255578096 | 0.114861074 |
| ENSDART00000110424 | si:ch211-153b23.4 | 0.636187514 | 1.795807227 | 1.178477605 | -0.329338933 |

| | | | | | |
|---|---|---|---|---|---|
| ENSDART00000110431 | si:ch73-280o22.2 | -0.128270202 | -0.267395445 | -0.366965125 | -0.13375727 |
| ENSDART00000110432 | kazna | -0.433352738 | -0.292283864 | -0.217890679 | -0.318143053 |
| ENSDART00000110447 | slitrk3b | -0.707318961 | -0.905108657 | -0.279729387 | -0.217159944 |
| ENSDART00000110458 | fam117ab | 0.691322185 | 0.501468791 | 0.171633838 | -0.275806067 |
| ENSDART00000110478 | zgc:174906 | 0.44828184 | 1.063594358 | 0.857934554 | 0.434989632 |
| ENSDART00000110497 | tmem63a | 0.205919412 | 0.504153398 | 0.414725435 | 0.171827856 |
| ENSDART00000110503 | adam11 | -0.835498408 | -0.874358127 | 0.055154913 | 0.615136144 |
| ENSDART00000110512 | ybx1 | 0.187678935 | 0.306636132 | 0.543417204 | 0.157462635 |
| ENSDART00000110522 | tulp1a | -0.068953976 | -0.228376734 | -0.263123945 | -0.07738281 |
| ENSDART00000110529 | bub1bb | 1.321570387 | 1.337769105 | 1.007493385 | 0.59867765 |
| ENSDART00000110544 | znf219 | -0.234403747 | -0.210413301 | -0.211690001 | -0.32688667 |
| ENSDART00000110547 | ANKRD34A | -0.459666228 | -0.579840169 | -0.409587984 | -0.250229823 |
| ENSDART00000110571 | fam150a | -0.089437558 | -0.463371816 | -0.643350488 | -0.328581938 |
| ENSDART00000110588 | cdh26.1 | 1.937401648 | 2.777818936 | 3.097440102 | 2.219591598 |
| ENSDART00000110590 | VSTM2B | -0.57346103 | -0.627600936 | -0.34222959 | -0.040178972 |
| ENSDART00000110595 | si:dkey-188g12.1 | -0.689058181 | -0.354561593 | -0.987489008 | -0.761086303 |
| ENSDART00000110606 | abhd16a | 0.034068276 | -0.290276411 | -0.385494504 | -0.249441823 |
| ENSDART00000110622 | si:ch211-222l21.1 | 0.830701135 | 0.950120833 | 0.646058632 | 0.069324807 |
| ENSDART00000110627 | EPB41L1 | 0.536765101 | 1.012243023 | 0.739789984 | 0.259906496 |
| ENSDART00000110633 | cdon | -0.443468077 | -0.07594048 | -0.652877368 | -0.650457598 |
| ENSDART00000110679 | lrfn5b | -0.958299909 | -0.447121499 | 0.131848444 | 0.317540452 |
| ENSDART00000110691 | wnt6b | 2.452564896 | 1.637817496 | 0.779324842 | 0.681102996 |
| ENSDART00000110696 | mxb | -0.714840475 | -2.62990329 | 3.629462021 | -0.04990443 |
| ENSDART00000110734 | FAM163A | -0.371556039 | -0.423320211 | -0.272260914 | 0.021575603 |
| ENSDART00000110751 | tiam1a | -0.473694473 | -0.373415926 | 0.157725394 | 0.265421677 |
| ENSDART00000110777 | eef1db | 0.19406492 | 0.18930318 | 0.407362874 | -0.014164442 |
| ENSDART00000110789 | esyt2b | -0.389945342 | -0.213760019 | -0.33183342 | -0.035634516 |
| ENSDART00000110804 | hspb15 | 0.646957051 | 1.106828293 | 1.50955981 | 0.885055615 |
| ENSDART00000110814 | fam83b | -0.195583845 | -0.26819738 | -0.2682294 | -0.101487813 |
| ENSDART00000110821 | ttc19 | -0.356526006 | -0.209564592 | -0.274076967 | -0.065932254 |
| ENSDART00000110824 | wdr17 | -0.247986884 | -0.387190174 | -0.468364816 | -0.181828903 |
| ENSDART00000110835 | myo18aa | -0.29501199 | -0.246090679 | -0.193185897 | -0.027559028 |
| ENSDART00000110842 | rps2 | 0.387589328 | 0.42480999 | 0.452974598 | 0.136625115 |
| ENSDART00000110848 | HBZ (1 of many) | 0.826283934 | 0.913969919 | 1.466570031 | 0.062329776 |
| ENSDART00000110854 | elk1 | -0.317422178 | -0.365267467 | -0.144808984 | -0.160403171 |
| ENSDART00000110866 | cln5 | 0.673394988 | 0.327334357 | 0.152385216 | -0.174612372 |
| ENSDART00000110879 | magi2b | -0.318025437 | -0.557603079 | -0.245924365 | -0.15150652 |
| ENSDART00000110883 | akap12b | 0.700660963 | 1.064613646 | 0.897645503 | 0.160893575 |
| ENSDART00000110884 | zmiz1a | -0.245374026 | -0.039847801 | 0.243895684 | 0.299607928 |
| ENSDART00000110923 | cnksr2b | -0.347259219 | -0.644524762 | -0.33094872 | 0.06260613 |
| ENSDART00000110935 | si:ch1073-59l16.1 | 1.356882908 | 0.676384004 | 0.249167454 | 0.069788375 |
| ENSDART00000110964 | bag6 | 0.132774646 | 0.150367292 | 0.354477336 | 0.199439987 |
| ENSDART00000110974 | zdhhc12b | -0.321463982 | -0.336778233 | -0.359206693 | -0.209775575 |
| ENSDART00000110976 | col19a1 | -0.113310455 | -0.554338462 | -0.336015886 | -0.135833904 |
| ENSDART00000110994 | sqlea | 0.510664466 | 1.875874264 | 2.415595993 | 1.866281513 |
| ENSDART00000111002 | si:ch211-74f19.2 | 0.463826256 | 0.294740419 | -0.053551592 | -0.452436201 |
| ENSDART00000111021 | si:dkeyp-14d3.1 | -0.147889699 | -0.589075251 | -0.655456899 | -0.257575305 |
| ENSDART00000111025 | boc | -0.443385595 | -0.283925154 | -0.350248278 | -0.500405723 |
| ENSDART00000111055 | kcna4 | -0.237309303 | -0.448087799 | -0.436530246 | -0.275701844 |
| ENSDART00000111058 | mpx | 0.612551173 | 0.415979564 | 0.081520104 | -3.505775593 |
| ENSDART00000111080 | adgrb3 | -0.692483265 | -0.604853517 | -0.34586076 | -0.181234915 |

| | | | | | |
|---|---|---|---|---|---|
| ENSDART00000111088 | depdc5 | 0.225009609 | 0.230516632 | 0.305363773 | 0.172290661 |
| ENSDART00000111108 | CABZ01071972.1 | -0.309847536 | -0.540379608 | -0.256707009 | -0.057294184 |
| ENSDART00000111111 | nlgn4b | -0.558022607 | -0.771924705 | -0.300759795 | -0.167214796 |
| ENSDART00000111131 | elfn1b | -1.012125284 | -0.755320746 | -0.301892543 | 0.27076153 |
| ENSDART00000111140 | rpl29 | 0.336664553 | 0.356246951 | 0.105865493 | -0.061220703 |
| ENSDART00000111146 | si:ch211-26b3.4 | -0.414256231 | -0.420417531 | -0.178854692 | 0.040407002 |
| ENSDART00000111156 | pdp1 | -0.641430349 | -0.338633085 | -0.014499951 | 0.034332362 |
| ENSDART00000111165 | best2 | -0.15773889 | -0.249582303 | -0.643700083 | -1.151697377 |
| ENSDART00000111189 | gar1 | 0.542751892 | 0.313383695 | 0.267097115 | 0.089542944 |
| ENSDART00000111203 | tagln3a | -0.530904757 | -0.489741378 | -0.221881281 | -0.037658487 |
| ENSDART00000111213 | ubash3bb | -0.454832578 | -0.295198237 | 0.117674424 | 0.358244958 |
| ENSDART00000111234 | gdpd5a | -0.267734676 | -0.323349922 | -0.30731905 | -0.184332435 |
| ENSDART00000111246 | cadm3 | -0.068241621 | -0.322804819 | -0.138950978 | -0.087560594 |
| ENSDART00000111261 | cdh24b | -0.501530291 | -0.712397312 | -0.362295258 | -0.168930493 |
| ENSDART00000111271 | CABZ01080074.2 | 0.091182245 | -0.411344825 | -0.551440399 | -0.297376381 |
| ENSDART00000111278 | sorcs2 | -0.061568527 | 0.240045859 | 0.726875042 | 0.744539967 |
| ENSDART00000111301 | luzp2 | -0.629816567 | -0.548346899 | 0.040106065 | 0.088332731 |
| ENSDART00000111303 | rfx7 | 0.075232989 | 0.085195342 | 0.459783838 | 0.727409902 |
| ENSDART00000111308 | bbs9 | -0.095835492 | -0.30952763 | -0.39815573 | -0.067617558 |
| ENSDART00000111321 | zgc:152830 | 0.702895787 | 0.346388089 | 0.27465484 | 0.003141495 |
| ENSDART00000111323 | AMIGO3 (1 of many) | -0.274114526 | -0.487248865 | -0.174614423 | -0.21591611 |
| ENSDART00000111324 | zgc:193807 | 2.638791453 | 3.489955154 | 3.039693941 | 3.396234279 |
| ENSDART00000111343 | lmf2a | -0.264313993 | -0.256081583 | -0.425329386 | -0.228346273 |
| ENSDART00000111374 | ndufs7 | -0.171726296 | -0.309298972 | -0.568241927 | -0.290550046 |
| ENSDART00000111400 | kel | 0.291873396 | 0.332474665 | 0.393429393 | 0.121747313 |
| ENSDART00000111435 | hdac5 | -0.290855518 | -0.361511948 | -0.278953131 | -0.082259238 |
| ENSDART00000111438 | mgea5 | -0.307275123 | -0.024811963 | 0.160874404 | 0.159343333 |
| ENSDART00000111444 | ppip5k1a | -0.365205244 | -0.30690644 | -0.053146194 | 0.037826946 |
| ENSDART00000111453 | myo19 | -0.517639897 | -0.807225171 | -0.660718539 | -0.325322766 |
| ENSDART00000111454 | UBA6 | -0.095626975 | 0.120665542 | 0.496662318 | 0.328619297 |
| ENSDART00000111456 | rbm10 | -0.2435311 | -0.244375112 | -0.319750026 | -0.111931812 |
| ENSDART00000111475 | pcdh7a | 0.109274008 | 0.268985487 | 0.465377244 | 0.354118271 |
| ENSDART00000111480 | bcorl1 | 0.013917394 | 0.173170994 | 0.344992278 | 0.232054818 |
| ENSDART00000111506 | lrrc75ba | -0.649545281 | -0.408918781 | -0.064365089 | 0.060080923 |
| ENSDART00000111509 | CABZ01072096.1 | -0.19271783 | -0.37213696 | -0.269302922 | -0.183488656 |
| ENSDART00000111531 | epdl1 | 1.357192284 | 1.365774088 | 0.753975689 | 0.235060778 |
| ENSDART00000111535 | elfn1a | -0.273088167 | -0.336173663 | -0.097567542 | 0.139029668 |
| ENSDART00000111536 | rmdn2 | -0.153020266 | -0.402861584 | -0.346146607 | -0.222410217 |
| ENSDART00000111539 | efcc1 | -0.101491995 | 0.211154978 | -0.621378561 | -0.272291709 |
| ENSDART00000111555 | fbxo10 | -0.524035405 | 0.01323878 | -0.15955536 | 0.008552029 |
| ENSDART00000111561 | zmp:0000000735 | 0.220285812 | 0.110420777 | 1.549961519 | 0.06406158 |
| ENSDART00000111571 | rap1gap2b | -0.543266541 | -0.656699315 | -0.515603594 | -0.222139298 |
| ENSDART00000111625 | ninl | -0.236310585 | -0.326810382 | -0.437556672 | -0.154099243 |
| ENSDART00000111636 | DTX4 (1 of many) | -1.011696111 | -1.148570348 | -0.809918586 | -0.692193186 |
| ENSDART00000111639 | rereb | -0.168137061 | -0.390988508 | -0.080335114 | 0.086629083 |
| ENSDART00000111641 | morn4 | 0.104472779 | 0.38320597 | 0.615725038 | 0.556227667 |
| ENSDART00000111642 | brinp3a.1 | -0.710663483 | -0.69660402 | -0.30207639 | -0.147819903 |
| ENSDART00000111656 | gpr78a | -0.727925057 | -0.963113505 | -0.708360358 | -0.204121228 |
| ENSDART00000111666 | npdc1a | -0.090316422 | -0.266587539 | -0.30135563 | -0.100212297 |
| ENSDART00000111671 | mri1 | 1.003264298 | 0.870246845 | 1.443034807 | 1.04475049 |
| ENSDART00000111680 | iqsec2b | -0.255745447 | -0.305278608 | -0.149665745 | -0.145530664 |

| | | | | | |
|---|---|---|---|---|---|
| ENSDART00000111688 | zgc:109934 | 0.61674072 | 0.212807687 | -0.164219088 | -0.482167701 |
| ENSDART00000111706 | si:dkey-108k21.14 | 0.724790653 | 1.05338345 | 1.058888091 | 0.439899526 |
| ENSDART00000111707 | cacnb1 | 1.693120335 | 2.349827744 | 2.415610432 | 1.652036859 |
| ENSDART00000111717 | fscn2b | -0.120313044 | -0.274854028 | -0.483029563 | -0.127989104 |
| ENSDART00000111724 | hist1h4l | 0.596018889 | 0.814003935 | 0.407366731 | 0.206389719 |
| ENSDART00000111748 | pcdhb | 0.251557806 | 0.39162054 | 0.85552916 | 0.748922663 |
| ENSDART00000111753 | vmp1 | 3.174699109 | 3.199538621 | 2.234258919 | 1.662616904 |
| ENSDART00000111758 | si:ch211-165d12.4 | 0.498072983 | 0.272355872 | -0.037512583 | -0.064496932 |
| ENSDART00000111759 | zmp:0000000794 | -0.428447438 | -0.466968786 | -0.39807432 | -0.23111401 |
| ENSDART00000111767 | si:ch211-39k3.2 | -0.094085509 | -0.095643794 | -0.600534706 | -0.396103136 |
| ENSDART00000111799 | HEPACAM (1 of many) | -0.300776869 | -0.301240305 | -0.377807505 | -0.329578876 |
| ENSDART00000111806 | cenpe | 1.006157891 | 0.692740621 | 0.695639634 | 0.175054418 |
| ENSDART00000111823 | GRIK3 | -0.298812703 | -0.633195231 | -0.313411843 | -0.167732562 |
| ENSDART00000111841 | BX248501.1 | -0.587056195 | -1.220156186 | -0.870300177 | -0.776464497 |
| ENSDART00000111842 | sall1a | -0.3702182 | -0.245271728 | -0.043742586 | 0.007554245 |
| ENSDART00000111880 | PDE2A | -0.027201032 | -0.255047431 | -0.265146028 | -0.145249945 |
| ENSDART00000111905 | BX649498.1 | -0.175767299 | -0.206924623 | -0.494153858 | -0.425985895 |
| ENSDART00000111923 | ajuba | 0.639088613 | 1.058968297 | 0.866425943 | 0.103913512 |
| ENSDART00000111948 | sez6l2 | -0.5142749 | -0.140866125 | 0.323444655 | 0.419947542 |
| ENSDART00000111966 | arhgef10lb | 0.250651161 | 0.391053857 | 0.470093443 | 0.290981355 |
| ENSDART00000111982 | sgce | -0.25805142 | -0.148921708 | -0.040980656 | -0.017549451 |
| ENSDART00000111993 | si:dkey-19b23.15 | -0.211537347 | -0.357802737 | -0.511679441 | -0.364770311 |
| ENSDART00000112003 | adgrb1a | -0.32135813 | -0.631154823 | -0.317944877 | 0.097147786 |
| ENSDART00000112028 | si:ch211-180f4.1 | 2.108953042 | 1.952679055 | 1.909638244 | 1.352861352 |
| ENSDART00000112032 | ARHGAP22 (1 of many) | 0.396989069 | 0.894459004 | 1.264783984 | 1.011214691 |
| ENSDART00000112075 | | 0.743523521 | 0.909798042 | 1.147105824 | 0.978802209 |
| ENSDART00000112079 | tlcd2 | 0.75377096 | 1.111252223 | 1.10627033 | 0.372244628 |
| ENSDART00000112106 | fam155a | -0.397020883 | -0.419375873 | -0.258307362 | -0.051593808 |
| ENSDART00000112116 | ctss2.1 | 1.048314865 | 0.112315631 | 0.133571721 | -0.538227279 |
| ENSDART00000112127 | dnase2b | 0.959684518 | 0.954914684 | 0.226944152 | -0.0439051 |
| ENSDART00000112152 | gas7a | -0.397395182 | -0.5126218 | -0.479089603 | -0.475145138 |
| ENSDART00000112155 | rrm2 | 1.131329801 | 1.215867796 | 0.761186289 | 0.339828406 |
| ENSDART00000112156 | si:ch211-253b8.5 | -0.087783514 | -0.192500909 | -0.49330541 | -0.306875488 |
| ENSDART00000112166 | fam60al | -0.06751471 | 0.448215095 | 0.685621255 | 0.373624581 |
| ENSDART00000112170 | unm_hu7912 | -0.404814593 | -0.573062573 | -0.192411138 | 0.071188401 |
| ENSDART00000112183 | cep170b | 0.178654359 | -0.17509023 | -0.407988469 | -0.254556637 |
| ENSDART00000112201 | mat2aa | 0.26207582 | 0.205858664 | 0.104925379 | 0.050770536 |
| ENSDART00000112211 | si:ch211-264f5.2 | 0.548989898 | 0.369154347 | -0.087668194 | -1.075571179 |
| ENSDART00000112227 | BRINP3 (1 of many) | -0.375889079 | -0.310381341 | -0.391723747 | -0.183098379 |
| ENSDART00000112243 | crlf1a | 1.589627089 | 0.916105309 | 0.558277873 | -0.020827681 |
| ENSDART00000112270 | tanc2a | -0.071755517 | -0.057687988 | 0.274000197 | 0.309857736 |
| ENSDART00000112287 | gas2l1 | -0.30204098 | -0.119087794 | 0.361891535 | 0.246536224 |
| ENSDART00000112296 | si:ch211-152n14.4 | -3.416151322 | -1.061691573 | -0.948841593 | -2.14718796 |
| ENSDART00000112299 | gpr37a | -0.365323083 | -0.2884934 | -0.319868677 | -0.150652035 |
| ENSDART00000112301 | trhde.1 | -0.264986832 | -0.332160295 | -0.182975887 | -0.056935825 |
| ENSDART00000112312 | lrch2 | 0.101952948 | 0.206054969 | 0.497048672 | 0.404162087 |
| ENSDART00000112313 | wdr43 | 2.967338106 | 2.621161906 | 3.405726008 | 2.58425126 |
| ENSDART00000112330 | rassf7a | -0.0890413 | -0.097498745 | -0.456633564 | -0.220562537 |
| ENSDART00000112333 | cnnm2b | -0.413171039 | -0.296279679 | -0.539640998 | -0.529524127 |
| ENSDART00000112370 | ano11 | -1.080551268 | -1.39069517 | -0.608655336 | 0.046438031 |
| ENSDART00000112407 | | -0.278472028 | -0.07282662 | 0.029836447 | -0.030627475 |

| | | | | | |
|---|---|---|---|---|---|
| ENSDART00000112414 | rapgef5a | -0.533059964 | -0.605979704 | -0.027243653 | 0.007460247 |
| ENSDART00000112438 | si:ch73-335m24.5 | -0.12977304 | -0.024649938 | 0.356284768 | 0.481799029 |
| ENSDART00000112441 | tlr7 | -0.49393668 | -0.654080017 | -0.57399465 | -0.158959649 |
| ENSDART00000112445 | tns2a | 0.230856175 | 0.361046363 | 0.629539855 | -0.041949691 |
| ENSDART00000112460 | dolk | 0.234400054 | 0.274257795 | 0.504999751 | 0.504335897 |
| ENSDART00000112470 | ccdc88b | 0.86030789 | 0.601824136 | 0.501988839 | -0.28013627 |
| ENSDART00000112484 | nlgn2b | -0.707313853 | -0.645741453 | -0.193542068 | 0.120518294 |
| ENSDART00000112493 | syt6a | -0.290141292 | -0.351590983 | -0.39435722 | -0.242348488 |
| ENSDART00000112495 | zgc:110340 | 0.095667849 | -0.009447766 | 0.241257465 | 0.332404774 |
| ENSDART00000112529 | znf319a | -0.271600655 | -0.228932764 | -0.01869534 | 0.01981331 |
| ENSDART00000112546 | pkn3 | -0.415654116 | -0.225897422 | -0.124759806 | -0.079978695 |
| ENSDART00000112550 | mapk9 | 0.429625805 | 0.878911558 | 0.999507915 | 0.700660398 |
| ENSDART00000112555 | spc25 | 0.851542956 | 0.93394939 | 0.260782437 | -0.118635043 |
| ENSDART00000112567 | ppfia4 | -0.303784132 | -0.279792224 | 0.037275964 | 0.113653516 |
| ENSDART00000112579 | scg2b | -0.360046965 | -0.210825369 | -0.351760362 | -0.440018453 |
| ENSDART00000112589 | efna2b | -0.324619544 | 0.075961218 | 0.539972471 | 0.527283417 |
| ENSDART00000112598 | otud4 | 0.000834797 | 0.118841749 | 0.673957038 | 0.019221004 |
| ENSDART00000112646 | CNNM1 | -0.428104026 | -0.544400388 | -0.592385966 | -0.267977747 |
| ENSDART00000112653 | wdfy4 | 0.841684139 | 0.566530892 | 0.08064366 | -0.41519717 |
| ENSDART00000112655 | nfasca | -0.501256457 | -0.441261682 | -0.289256026 | -0.116172993 |
| ENSDART00000112659 | ipp | 0.043576539 | 0.16998945 | 0.371873925 | 0.242338779 |
| ENSDART00000112671 | bicd1a | -0.391567366 | -0.28876668 | -0.040397452 | 0.195532163 |
| ENSDART00000112694 | fam171a2b | -0.001835186 | 0.12088862 | 0.541529951 | 0.657778888 |
| ENSDART00000112698 | RNF14 (1 of many) | 0.142790619 | 0.542852416 | 0.218671557 | 0.088301354 |
| ENSDART00000112711 | kcnab1b | -0.291029319 | -0.341790858 | -0.379711216 | -0.329063836 |
| ENSDART00000112728 | tmem175 | -0.092618029 | 0.36107371 | 0.426939782 | 0.221686367 |
| ENSDART00000112735 | fam19a1a | -0.689638571 | -0.889095001 | -0.657230812 | -0.483951597 |
| ENSDART00000112743 | nmbb | -0.255890888 | -1.177861162 | -0.767211345 | -0.814299265 |
| ENSDART00000112768 | ubald1a | -0.163342676 | -0.260286427 | -0.366999537 | -0.147535942 |
| ENSDART00000112789 | AKAP13 (1 of many) | 0.092610458 | -0.131036675 | -0.325814147 | -0.202546922 |
| ENSDART00000112845 | zgc:195173 | -0.156651959 | -0.182049019 | -0.580426063 | -0.462296521 |
| ENSDART00000112856 | TNC (1 of many) | -0.33689758 | -0.285587425 | -0.51132075 | -1.127367477 |
| ENSDART00000112883 | fmnl2b | -0.095811604 | 0.162122972 | 0.557976718 | 0.595265469 |
| ENSDART00000112895 | FNDC10 | -0.46103084 | -0.039905964 | 0.379016706 | 0.57822448 |
| ENSDART00000112900 | si:ch211-150i13.1 | -0.166104961 | -0.246781433 | -0.279012086 | -0.114516059 |
| ENSDART00000112909 | tex2l | -0.590392514 | -0.678584931 | -0.756484503 | -0.42899031 |
| ENSDART00000112926 | adora1b | 0.387978003 | 0.609753862 | 0.576471225 | 0.261995134 |
| ENSDART00000112956 | slc9a6b | -0.347273739 | -0.284992324 | -0.155606241 | -0.015875628 |
| ENSDART00000112959 | ankrd1a | 2.226071129 | 3.68265785 | 3.832168007 | 2.790435602 |
| ENSDART00000112967 | rilp | -0.073950343 | -0.174960248 | -0.511756745 | -0.145692497 |
| ENSDART00000112987 | stk32a | -0.326877709 | -0.547605605 | -0.241724362 | 0.046529539 |
| ENSDART00000113004 | CABZ01068356.1 | -0.41749624 | -0.503262627 | -0.597219984 | -0.132734415 |
| ENSDART00000113058 | gpsm2 | -0.158548143 | -0.583640034 | -0.403831438 | -0.106682634 |
| ENSDART00000113077 | FERMT3 (1 of many) | 0.719070873 | 0.278758571 | 0.151005206 | -0.238328508 |
| ENSDART00000113081 | gpr158b | -0.777838913 | -0.859733549 | -0.109236002 | 0.425659764 |
| ENSDART00000113087 | hmx1 | -0.389387565 | 0.021377927 | 0.234018679 | 0.270888749 |
| ENSDART00000113089 | slc4a11 | -0.230777933 | -0.44932083 | -0.596234565 | -0.271629672 |
| ENSDART00000113092 | abhd10b | 0.399222781 | 0.423215001 | 0.172340204 | 0.123654926 |
| ENSDART00000113093 | gba | 0.473554928 | 0.246228233 | -0.028007284 | -0.118288896 |
| ENSDART00000113097 | hsd17b7 | -0.488876774 | 0.314769091 | 0.517151777 | 0.148788215 |
| ENSDART00000113098 | hbaa1 | 0.802810656 | 0.919517173 | 1.438314563 | 0.102144076 |

| | | | | | |
|---|---|---|---|---|---|
| ENSDART00000113101 | smarcad1b | 0.481835623 | 0.651517984 | 0.685097173 | 0.491069329 |
| ENSDART00000113112 | wscd1b | 0.189205134 | 0.487630626 | 0.094129579 | 0.021242458 |
| ENSDART00000113135 | msl1a | -0.136399644 | -0.187672093 | -0.188075662 | -0.302625769 |
| ENSDART00000113142 | phldb1b | 0.143763621 | 0.070606141 | 0.367437708 | 0.295748576 |
| ENSDART00000113162 | pi4kaa | -0.145989753 | -0.358429295 | -0.235575632 | -0.007193852 |
| ENSDART00000113166 | si:ch1073-159d7.7 | 2.749837608 | 3.675596621 | 2.897443721 | 1.850742921 |
| ENSDART00000113171 | nlgn1 | -0.287475058 | -0.532070389 | -0.095767589 | 0.228264127 |
| ENSDART00000113193 | cbln2a | -0.405462517 | -0.494893494 | -0.545170143 | -0.282281473 |
| ENSDART00000113196 | si:ch211-157b11.14 | -0.299973945 | -0.39773998 | -0.291303207 | -0.069868393 |
| ENSDART00000113197 | prok1 | -0.431580361 | -0.477066939 | -0.390033973 | -0.689815046 |
| ENSDART00000113241 | tmem163a | 0.329551441 | 0.273820345 | 0.385569684 | 0.333416277 |
| ENSDART00000113246 | si:ch211-102c2.4 | 0.606357076 | 0.357270403 | 0.232028808 | -0.115778826 |
| ENSDART00000113248 | b3galt1b | -0.287109392 | -0.341327689 | -0.370081074 | -0.245129506 |
| ENSDART00000113259 | TARBP1 | 0.600840202 | 0.638996908 | 0.05884907 | -0.019773553 |
| ENSDART00000113280 | FRMD5 (1 of many) | -0.439670364 | -0.372637017 | -0.247112316 | -0.218940122 |
| ENSDART00000113301 | si:dkeyp-41f9.3 | -0.138197975 | -0.385879672 | -0.585370915 | -0.421818221 |
| ENSDART00000113311 | col9a1a | 0.34651107 | 0.351090918 | -0.438999012 | -0.868499411 |
| ENSDART00000113332 | nav2a | 0.038264794 | -0.31474256 | -0.162187636 | 0.073256894 |
| ENSDART00000113342 | tox3 | -0.349601618 | -0.245853476 | -0.159173384 | -0.016959002 |
| ENSDART00000113347 | | -0.143628386 | 0.120543612 | 0.968351441 | 1.05709332 |
| ENSDART00000113351 | ncapd2 | 0.642337818 | 0.531864694 | 0.423422126 | 0.276399068 |
| ENSDART00000113356 | crtc1a | -0.294987163 | -0.303249696 | -0.252865857 | -0.15461469 |
| ENSDART00000113358 | wdtc1 | -0.236133684 | -0.265438127 | -0.168580048 | -0.081863191 |
| ENSDART00000113362 | ip6k1 | -0.059673843 | -0.326658711 | -0.456047384 | -0.047258149 |
| ENSDART00000113376 | fam13b | -0.247096949 | -0.341873515 | -0.575011995 | -0.179280023 |
| ENSDART00000113384 | lyrm4 | -0.082514668 | -0.167513155 | -0.434528553 | -0.231621665 |
| ENSDART00000113405 | inpp5d | 0.955185108 | 0.470897855 | 0.050793817 | -0.341647614 |
| ENSDART00000113414 | kirrel3a | -0.106976766 | -0.45848865 | -0.133882402 | 0.030163053 |
| ENSDART00000113418 | igsf3 | 0.097020516 | 0.19405944 | 0.607620623 | 0.491282072 |
| ENSDART00000113441 | mylk5 | 0.7890096 | 0.412195956 | -0.130414718 | -0.622741882 |
| ENSDART00000113448 | WDR31 | -0.320489538 | -0.191593939 | -0.444041238 | -0.334593236 |
| ENSDART00000113449 | LEPROT | 0.490278812 | 0.508314674 | 0.201181129 | 0.107244197 |
| ENSDART00000113454 | chgb | -0.341137392 | -0.331982228 | -0.294276744 | -0.206705528 |
| ENSDART00000113486 | mxra5a | 0.309757309 | 0.386526989 | 0.398748931 | 0.219293393 |
| ENSDART00000113493 | tmem14ca | -0.005514111 | -0.126364666 | -0.303791848 | -0.180226659 |
| ENSDART00000113494 | MB21D2 (1 of many) | -0.150231845 | 0.017615171 | 0.437179993 | 0.383202322 |
| ENSDART00000113502 | si:dkey-84j12.1 | -0.363232597 | -0.10430598 | 0.492332906 | 0.665079551 |
| ENSDART00000113511 | lzts1 | -1.206756297 | -0.713910989 | -0.061680884 | 0.474255612 |
| ENSDART00000113524 | fam20ca | -0.141135887 | -0.526654581 | -0.212681972 | 0.069156499 |
| ENSDART00000113532 | CABZ01053588.1 | 0.66169386 | 1.399901635 | 1.305166578 | 0.721799042 |
| ENSDART00000113550 | si:dkey-16p21.7 | -0.102214974 | 0.203772916 | 0.542107134 | 0.32053481 |
| ENSDART00000113551 | trim2b | -0.50417548 | -0.207383112 | -0.112448816 | 0.043074002 |
| ENSDART00000113574 | anln | 1.428520356 | 0.903846206 | 0.598652735 | 0.284032371 |
| ENSDART00000113589 | gpx4b | 0.633021831 | -0.10742897 | -0.312510701 | -0.292370818 |
| ENSDART00000113643 | fbxo25 | 0.400029831 | 0.002394742 | -0.303777301 | 0.069383699 |
| ENSDART00000113649 | impg1a | -0.10562067 | -0.311339049 | -0.59002435 | -0.135488682 |
| ENSDART00000113655 | tmem26a | 0.744638282 | 1.01684072 | 1.017300963 | 0.48692919 |
| ENSDART00000113661 | kif16bb | -0.079672799 | -0.046364361 | -0.755779075 | -0.266178188 |
| ENSDART00000113673 | scpp5 | 1.321767426 | 0.530715467 | 0.931995129 | 0.748355856 |
| ENSDART00000113699 | hgfb | 1.149006949 | 1.556415854 | 1.569424226 | 0.45979101 |
| ENSDART00000113732 | orc6 | 1.104098282 | 1.496929933 | 0.974585154 | 0.388395497 |

| | | | | | |
|---|---|---|---|---|---|
| ENSDART00000113734 | chrdl2 | -0.47545279 | -0.359333121 | -0.57438139 | -0.417914769 |
| ENSDART00000113752 | si:dkey-6i22.5 | 1.012202134 | 1.421252013 | 0.919204178 | 0.319242336 |
| ENSDART00000113773 | cdc42bpab | -0.428966416 | -0.296406465 | 0.003528598 | 0.157857062 |
| ENSDART00000113789 | zgc:198419 | 1.388791656 | 0.463187117 | 0.161859067 | -0.160692863 |
| ENSDART00000113796 | cacnb3b | -1.378695476 | -1.717513295 | -0.09050578 | 0.468842834 |
| ENSDART00000113799 | si:ch211-132b12.7 | 0.091573373 | -0.269264305 | -0.847675231 | -0.371595689 |
| ENSDART00000113805 | nfatc2b | -0.301956989 | -0.356397992 | -0.30237662 | -0.2051156 |
| ENSDART00000113829 | cdca7b | 1.046232255 | 1.184963787 | 0.646928123 | 0.587090661 |
| ENSDART00000113842 | dpep2 | -0.005800421 | 0.049668681 | -0.202900206 | -0.709132578 |
| ENSDART00000113847 | chpfa | -0.17880495 | -0.243387936 | -0.273328364 | -0.205137329 |
| ENSDART00000113853 | prokr1a | -0.220461946 | -0.109814734 | -0.599222392 | -0.17974305 |
| ENSDART00000113859 | crispld1a | 0.006653123 | -0.124334074 | -0.47278833 | -1.109435686 |
| ENSDART00000113864 | faxca | -0.531664025 | -0.191322961 | -0.005943785 | 0.107092327 |
| ENSDART00000113890 | rabep2 | -0.293276158 | -0.227772629 | -0.308410671 | -0.161791109 |
| ENSDART00000113912 | ntm | -0.178976918 | -0.363265579 | -0.513893339 | -0.211182567 |
| ENSDART00000113924 | lrrc7 | -0.537461037 | -0.585108204 | -0.303180727 | -0.024642369 |
| ENSDART00000113936 | SAMD8 | 0.425621262 | 0.457128542 | 0.374121986 | 0.140204511 |
| ENSDART00000113954 | si:ch211-113e8.11 | -0.149005557 | -0.166754978 | -0.25646894 | -0.094366468 |
| ENSDART00000113985 | mctp1a | -0.392181593 | -0.510456941 | -0.185251742 | 0.106694132 |
| ENSDART00000114000 | zgc:101663 | 0.831482698 | 0.254442158 | 0.060051833 | 0.091105397 |
| ENSDART00000114010 | slitrk3a | -0.361859007 | -0.532282315 | -0.225491041 | 0.106517685 |
| ENSDART00000114023 | hspb9 | 2.157513115 | 1.55136025 | 0.742718373 | 1.461470885 |
| ENSDART00000114024 | fam107b | 0.899175294 | 1.079533729 | 0.969523488 | 0.434748246 |
| ENSDART00000114026 | lingo2a | -0.52437969 | -3.569137229 | -2.642152541 | -0.423360666 |
| ENSDART00000114069 | cnnm4b | -0.449604106 | -0.276862048 | -0.121356457 | -0.173176154 |
| ENSDART00000114078 | si:dkey-193c22.2 | 0.250019316 | -0.054913753 | -0.32754917 | -0.367609802 |
| ENSDART00000114081 | sms | -0.273814009 | -0.150000435 | 0.170943482 | 0.265559933 |
| ENSDART00000114083 | mcf2la | -0.277595318 | -0.392280599 | -0.224516886 | 0.025969597 |
| ENSDART00000114098 | dkk2 | -0.775548216 | -0.461837556 | -0.163285316 | -0.107943795 |
| ENSDART00000114099 | BX545917.1 | -0.259517692 | -0.573524763 | -0.511746601 | -0.397812833 |
| ENSDART00000114117 | necab1 | -0.468499392 | -0.405674747 | -0.46743135 | -0.222378141 |
| ENSDART00000114118 | CU639468.1 | -0.073130118 | -0.36036163 | -0.387238652 | -0.163804242 |
| ENSDART00000114134 | wrb | -0.123951898 | -0.095408141 | -0.313002762 | -0.1622087 |
| ENSDART00000114162 | ftr02 | -0.105106717 | -0.079876083 | -0.28693813 | -0.123365826 |
| ENSDART00000114168 | itga6a | 1.998266436 | 2.016280588 | 1.463165074 | 0.79739173 |
| ENSDART00000114169 | bnip1a | -0.010894413 | -0.404734866 | -0.484887345 | -0.670176287 |
| ENSDART00000114172 | pde6d | -0.201449776 | -0.351770435 | -0.511138243 | -0.216254622 |
| ENSDART00000114182 | fyco1b | 0.10137681 | -0.156409007 | -0.465781329 | -0.01243848 |
| ENSDART00000114226 | grm6b | -0.535654703 | -0.571067618 | -0.480335418 | -0.370568762 |
| ENSDART00000114263 | zgc:194242 | -0.119842829 | -0.109342365 | -0.31411889 | -0.410626816 |
| ENSDART00000114267 | map6b | -0.567896811 | -0.403955591 | 0.492985119 | 0.85461741 |
| ENSDART00000114272 | nop2 | 0.471489342 | 0.531037709 | 0.420888693 | 0.177047326 |
| ENSDART00000114282 | rpsa | 0.494276409 | 0.531800028 | 0.113194341 | -0.004402994 |
| ENSDART00000114293 | areg | -0.232855169 | -0.639973524 | -1.102041863 | -0.924035384 |
| ENSDART00000114301 | si:ch211-152p11.4 | -0.510547269 | -0.515838482 | -0.89019375 | -0.843865894 |
| ENSDART00000114319 | si:ch211-76l23.7 | -0.274578898 | -0.196596974 | -0.473622321 | -1.111584541 |
| ENSDART00000114322 | rusc1 | -0.055969033 | -0.296840251 | -0.124744669 | -0.048299568 |
| ENSDART00000114336 | FAM189B | -0.002256766 | 0.039346929 | 0.240983806 | 0.310705552 |
| ENSDART00000114383 | mesdc1 | 0.277986788 | 0.306020697 | 0.430972974 | 0.195133048 |
| ENSDART00000114410 | wdr62 | 0.584622073 | 0.50588924 | 0.329377091 | 0.089655627 |
| ENSDART00000114432 | ntng2a | -0.38146173 | -0.278136767 | -0.282818481 | 0.289761799 |

| | | | | | |
|---|---|---|---|---|---|
| ENSDART00000114442 | fndc5b | -0.498984664 | -0.658119927 | -0.45669978 | -0.18817484 |
| ENSDART00000114448 | CHST8 | -0.44164787 | -0.464005415 | 0.047368929 | 0.426791372 |
| ENSDART00000114497 | si:dkey-204l11.1 | 0.312589581 | 0.828199841 | 0.99726607 | 0.068811015 |
| ENSDART00000114533 | mn1a | -0.109536494 | -0.207833595 | -0.685216226 | -0.20481073 |
| ENSDART00000114611 | sipa1l1 | -0.211106005 | -0.264425665 | -0.102896881 | -0.110763007 |
| ENSDART00000114659 | maml3 | -0.010023722 | -0.260659863 | -0.229491996 | 0.012935978 |
| ENSDART00000114660 | opn8b | -0.615308217 | -0.695080703 | -0.641485332 | -0.112308149 |
| ENSDART00000114663 | b4galt2 | -0.500896606 | -0.53488094 | -0.237386402 | -0.192514996 |
| ENSDART00000114676 | gstm.2 | 0.102339801 | -0.045272058 | -0.466318631 | -0.673317054 |
| ENSDART00000114677 | si:ch73-62l21.1 | -0.323827666 | -0.531978643 | -0.239685061 | 0.043755156 |
| ENSDART00000114678 | nanos1 | -0.186501948 | -0.191589001 | -0.393479142 | -1.039695471 |
| ENSDART00000114705 | gprc5bb | -0.132340316 | -0.338695197 | -0.459834622 | -0.091865364 |
| ENSDART00000114711 | sorcs1 | -0.736062495 | -0.837966489 | -0.594713883 | -0.208228471 |
| ENSDART00000114719 | frmpd1a | -0.29565145 | -0.338039747 | -0.498795038 | -0.059125706 |
| ENSDART00000114723 | rapgefl1 | -0.431713581 | -0.430345279 | -0.085299154 | -0.113952199 |
| ENSDART00000114746 | lrrc58a | -0.133537399 | 0.014548786 | -0.622798481 | -0.259773873 |
| ENSDART00000114748 | vip | -0.375915262 | -0.449108971 | -0.514462327 | -0.417228927 |
| ENSDART00000114750 | gabrb2 | -0.56997023 | -0.610789713 | -0.475013901 | -0.067415538 |
| ENSDART00000114774 | ptpn5 | -0.403695841 | -0.145979825 | 0.358989634 | 0.339107007 |
| ENSDART00000114800 | INAVA (1 of many) | 0.074314999 | 0.343046405 | 0.508626667 | 0.501928464 |
| ENSDART00000114888 | picalmb | -0.141762949 | -0.21097576 | -0.321137175 | -0.075858488 |
| ENSDART00000114909 | cuedc1a | -0.291201492 | -0.290842243 | -0.025622448 | 0.12561667 |
| ENSDART00000114919 | si:dkey-85k7.12 | 0.382806755 | -0.226494664 | 2.101494432 | -0.132340955 |
| ENSDART00000114954 | rapgef5b | -0.637101425 | -0.566396383 | -0.284127257 | 0.00714868 |
| ENSDART00000114959 | cdh24a | -0.51153845 | -0.93145045 | -0.229616461 | 0.228723625 |
| ENSDART00000114964 | zgc:165573 | 0.691787708 | 0.943542464 | 0.634593348 | -0.039695615 |
| ENSDART00000114975 | slc5a7a | -0.574231808 | -0.735930483 | -0.690375373 | -0.752661657 |
| ENSDART00000115023 | PARG | 0.013352577 | 0.067569536 | 0.312910159 | 0.18617429 |
| ENSDART00000115027 | tmem151bb | -0.856524713 | -0.708298022 | 0.291447963 | 0.495918048 |
| ENSDART00000115030 | dip2a | -0.094281248 | 0.006396805 | 0.484131414 | 0.408709873 |
| ENSDART00000115049 | mych | 2.666296724 | 2.42380403 | 1.76502541 | 0.819436082 |
| ENSDART00000115058 | pbxip1a | -0.476509078 | -0.442420659 | -0.523364891 | -0.310557938 |
| ENSDART00000115088 | birc5a | 0.588066272 | 0.489725837 | 0.133434985 | 0.421614791 |
| ENSDART00000115089 | zgc:171482 | -0.437080228 | -0.507022457 | -0.285662093 | 0.021798354 |
| ENSDART00000115118 | tns1a | -0.409165389 | -0.800200161 | -0.457588882 | -0.074958883 |
| ENSDART00000115128 | usp2b | -0.251717928 | -0.321635103 | -0.314073339 | -0.302633188 |
| ENSDART00000115130 | apbb1 | -0.260426658 | -0.249126495 | -0.138007572 | -0.113696464 |
| ENSDART00000115138 | rapgef4 | -0.335306767 | -0.551146979 | -0.593390846 | -0.144257686 |
| ENSDART00000115141 | cacnb3b | -1.10918155 | -1.536403603 | -0.203181521 | 0.418821499 |
| ENSDART00000115157 | tuba4l | 0.217024219 | 0.564563401 | 0.525644709 | 0.157922416 |
| ENSDART00000115161 | reps2 | -0.084468471 | -0.327140094 | -0.166340372 | -0.127803804 |
| ENSDART00000115221 | adat2 | 0.066031062 | -1.495132242 | -2.125701316 | -0.144340549 |
| ENSDART00000115224 | pvrl1a | -0.001527 | -0.239159739 | -0.512986688 | -0.121351808 |
| ENSDART00000115244 | MEX3A | 1.222333965 | 1.709642669 | 1.900449694 | 1.354844663 |
| ENSDART00000115255 | mcf2l2 | 0.795464349 | 0.848343644 | 0.628078884 | 0.002922432 |
| ENSDART00000115260 | si:ch211-113g11.6 | -0.558167113 | -1.193520332 | -0.037233796 | 0.533309069 |
| ENSDART00000115278 | cx47.1 | -0.445735222 | -0.554690047 | -0.399294096 | -0.241214414 |
| ENSDART00000115330 | ppp1r14c | -0.176056212 | -0.351217013 | -0.000962023 | -0.249969624 |
| ENSDART00000115343 | atp8b4 | -0.126263785 | -0.363733117 | -0.385610912 | -0.00148052 |
| ENSDART00000115354 | si:dkey-188i13.10 | 0.511265672 | 0.230301163 | 1.128651443 | -0.074861923 |
| ENSDART00000115356 | rabif | -0.104235333 | -0.157538528 | -0.394672329 | -0.327606577 |

| | | | | | |
|---|---|---|---|---|---|
| ENSDART00000115365 | rassf10a | -0.525931221 | -0.509933384 | -0.256813636 | -0.202027084 |
| ENSDART00000115370 | mettl22 | -0.250044414 | -0.320129908 | -0.572576042 | -0.437533988 |
| ENSDART00000115398 | arid5a | 0.65326128 | 0.28174461 | 0.163458034 | -0.140420456 |
| ENSDART00000115403 | NAV1 (1 of many) | 0.648363377 | 0.712988904 | 0.734499876 | 0.497918559 |
| ENSDART00000115417 | si:ch211-197l9.2 | 0.463829162 | 0.238718738 | 0.779021857 | 0.564789152 |
| ENSDART00000115550 | RNase_MRP | -0.088007656 | -0.32868786 | -0.789978682 | -0.744862923 |
| ENSDART00000115759 | | 2.750406791 | 2.822768519 | 2.52417923 | 3.741762635 |
| ENSDART00000115901 | SNORA71 | 0.179219354 | 0.724792108 | 0.150624912 | 0.373922197 |
| ENSDART00000115914 | SNORA71 | 0.179219354 | 0.724792108 | 0.150624912 | 0.373922197 |
| ENSDART00000116021 | | -0.209375753 | 0.199058885 | -0.850598268 | -0.180561216 |
| ENSDART00000116415 | | -0.367308157 | -0.297245306 | -0.576085374 | -0.866023911 |
| ENSDART00000117636 | dre-mir-21-1 | 1.553092638 | 1.563315667 | 0.906878174 | 0.77260601 |
| ENSDART00000117680 | dre-mir-21-2 | 3.366495632 | 3.222859121 | 2.199417673 | 1.472943682 |
| ENSDART00000118106 | SNORA57 | 0.429083089 | 0.942452098 | 0.631445196 | 0.725057114 |
| ENSDART00000118384 | SNORA16 | 0.565934898 | 0.9072012 | 0.499756229 | 0.494901316 |
| ENSDART00000118961 | 5S_rRNA | -0.375589637 | -0.40645143 | -0.614838385 | -0.339165991 |
| ENSDART00000119160 | SCARNA6 | 0.120700111 | -0.063996796 | -0.81140797 | -1.067895541 |
| ENSDART00000119311 | SNORA53 | -0.615548348 | -0.821029313 | -0.724467957 | -1.224866543 |
| ENSDART00000121226 | FO704882.1 | -0.227266262 | -0.409349874 | -0.661498137 | -0.983304677 |
| ENSDART00000121457 | lbh | -0.556077202 | -0.419617758 | -0.340252817 | -0.354555634 |
| ENSDART00000121460 | prdm8b | -0.383342673 | -0.361617984 | -0.117709411 | 0.067402978 |
| ENSDART00000121476 | asns | 0.135112284 | 0.218356853 | 0.383559102 | 0.26633883 |
| ENSDART00000121489 | mybl2b | 0.480424926 | 0.360866643 | 0.136903159 | 0.039795348 |
| ENSDART00000121496 | gpr153 | -0.22286471 | -0.331603234 | -0.24476212 | -0.082544732 |
| ENSDART00000121503 | cplx3b | -0.427974445 | -0.393867199 | -0.470572672 | -0.340193244 |
| ENSDART00000121531 | mat2aa | 0.194654747 | 0.17321442 | 0.334936873 | 0.070313151 |
| ENSDART00000121545 | brms1 | -0.198338884 | -0.120854433 | -0.325603911 | -0.103470514 |
| ENSDART00000121598 | phf10 | -0.237213765 | -0.309795055 | -0.326418269 | -0.213703564 |
| ENSDART00000121647 | PRMT8 (1 of many) | 0.693543627 | 0.209410281 | 0.412670468 | 0.652513913 |
| ENSDART00000121675 | angptl1a | -0.785726694 | -0.739578659 | -0.711456 | -1.277545302 |
| ENSDART00000121684 | nat8l | -0.712971152 | -0.758652511 | -0.423007329 | -0.36098028 |
| ENSDART00000121708 | pcsk1nl | -0.536353064 | -0.421583117 | -0.086670577 | -0.06684806 |
| ENSDART00000121714 | gnptab | 0.326534887 | 0.246690992 | 0.1944971 | 0.101206309 |
| ENSDART00000121716 | FAM107A | -0.353665396 | -0.22740498 | -0.416694229 | -0.314761483 |
| ENSDART00000121722 | si:dkey-274m17.3 | -0.27733184 | -0.305292621 | -0.626644892 | -0.318061843 |
| ENSDART00000121731 | | -0.190786213 | -0.244913354 | -0.339998812 | -0.082783766 |
| ENSDART00000121756 | sybu | -0.180478431 | -0.141687617 | -0.791670055 | -0.226701204 |
| ENSDART00000121817 | fbln7 | 0.143978228 | 0.156275351 | -0.420940219 | -1.064416584 |
| ENSDART00000121822 | | 0.766367421 | 1.508595661 | 1.280597751 | 1.015709274 |
| ENSDART00000121823 | syngr3b | -0.148548395 | -0.252198032 | -0.105567779 | 0.074385745 |
| ENSDART00000121826 | bean1 | -0.425046714 | -0.600778843 | -0.413029577 | -0.180932047 |
| ENSDART00000121837 | efs | 0.29165502 | 0.378275961 | 0.750399498 | 0.505660243 |
| ENSDART00000121861 | prph | 1.238160677 | 2.697856064 | 2.616616233 | 2.213149799 |
| ENSDART00000121864 | slc27a6 | -0.486568844 | -0.183000547 | -0.356047466 | -0.257017832 |
| ENSDART00000121866 | desi1b | 0.626366318 | 0.579237123 | 0.318409588 | -0.099655493 |
| ENSDART00000121867 | eif3c | 0.151751793 | 0.171944002 | 0.307946358 | 0.118389215 |
| ENSDART00000121872 | mast3b | -0.277342703 | -0.556828614 | -0.261994844 | -0.168334337 |
| ENSDART00000121874 | nfasca | -0.536181619 | -0.366142839 | -0.19557692 | -0.058427118 |
| ENSDART00000121886 | hdr | 3.187835864 | 3.220765325 | 3.322516101 | 2.611876044 |
| ENSDART00000121913 | kctd12b | -0.424186634 | -0.205906213 | -0.149727383 | -0.132895958 |
| ENSDART00000121952 | h2afy2 | -0.217094069 | -0.401231455 | -0.429907398 | -0.128476741 |

| | | | | | |
|---|---|---|---|---|---|
| ENSDART00000121981 | smarce1 | 0.060968412 | 0.140256121 | 0.341840216 | 0.044607056 |
| ENSDART00000121984 | ssbp3b | -0.092095029 | 0.005321722 | 0.831022729 | 0.629434393 |
| ENSDART00000121989 | cald1b | -0.204161416 | -0.080240225 | -0.306263705 | -1.232197212 |
| ENSDART00000121998 | grwd1 | 0.591308312 | 0.087252775 | -0.045976929 | -0.167650618 |
| ENSDART00000122015 | fam169ab | -0.221722765 | -0.165161236 | -0.4562862 | -0.136486601 |
| ENSDART00000122037 | rbfox1 | -0.419101241 | -0.392215043 | 0.102043249 | 0.360818016 |
| ENSDART00000122041 | nrcama | -0.49531831 | -0.707479994 | -0.094699246 | 0.009754423 |
| ENSDART00000122059 | scoca | -0.009226259 | -0.04540809 | -0.349436614 | -0.115738429 |
| ENSDART00000122074 | slc1a7b | -0.35845108 | -0.359766371 | -0.277402725 | -0.101652869 |
| ENSDART00000122081 | sybu | -0.707212841 | -0.524661917 | -0.600466578 | -0.221891711 |
| ENSDART00000122099 | dynll2a | -0.463212519 | -0.169502826 | -0.013591394 | 0.001709702 |
| ENSDART00000122101 | tbx2b | -0.056468629 | 0.017248499 | 0.19422425 | 0.382938949 |
| ENSDART00000122102 | wee1 | 1.082199278 | 1.291191801 | 0.972164457 | 0.241130696 |
| ENSDART00000122115 | si:ch211-153j24.3 | -1.030489347 | -0.968888956 | -1.068398619 | -1.338359032 |
| ENSDART00000122133 | mkrn2os.1 | 0.117189294 | 0.106487037 | 0.442579902 | 0.198380082 |
| ENSDART00000122159 | si:dkey-118k5.3 | 0.43932377 | 0.244452571 | 0.265050848 | -0.163246789 |
| ENSDART00000122170 | smc5 | 0.002327719 | -0.01045929 | 0.55091638 | 0.063311494 |
| ENSDART00000122238 | tyr | 1.0676401 | 0.617097515 | 0.165901971 | -0.685666733 |
| ENSDART00000122288 | ctsla | 0.76727009 | 0.372627978 | 0.369896487 | -0.200637952 |
| ENSDART00000122305 | Metazoa_SRP | 1.452964427 | 1.874874084 | 1.874716847 | 1.224958179 |
| ENSDART00000122307 | gcsha | -0.027923504 | -0.123933497 | -0.256124703 | -0.163875624 |
| ENSDART00000122321 | slc33a1 | 0.868257035 | 0.839034235 | 0.725290511 | 0.201428784 |
| ENSDART00000122348 | gad1b | -0.426563629 | -0.260162283 | -0.435212228 | -0.267475156 |
| ENSDART00000122359 | si:dkey-164f24.2 | -0.316246733 | -0.171154186 | -0.508268616 | -0.361228089 |
| ENSDART00000122389 | elmod1 | -0.62893484 | -0.673357812 | -0.159560027 | 0.32261476 |
| ENSDART00000122407 | cdk1 | 1.677931921 | 1.534655447 | 0.824563588 | 0.384011667 |
| ENSDART00000122429 | klf15 | -0.341190152 | -0.616050077 | -0.578895446 | -0.337948098 |
| ENSDART00000122433 | trim33l | 0.38510745 | 0.766516023 | 0.671400212 | 0.185347257 |
| ENSDART00000122439 | si:dkey-238d18.4 | 0.670270269 | 0.479235347 | 0.13889457 | -0.356907703 |
| ENSDART00000122454 | citb | -0.619522503 | -1.363556906 | -0.83905374 | -0.294671991 |
| ENSDART00000122519 | st8sia6 | -0.258839848 | -0.191246071 | -0.329756561 | -0.383108642 |
| ENSDART00000122564 | trmt10a | 0.324659997 | 0.317173694 | 0.413928399 | 0.18195893 |
| ENSDART00000122566 | mxa | -0.032262098 | -0.382486532 | 3.249286652 | -1.622639363 |
| ENSDART00000122574 | ppp1r3aa | -0.153287429 | -0.684598529 | 0.022675976 | 0.569399664 |
| ENSDART00000122601 | pgam1b | -0.261329131 | -0.223010936 | -0.252995896 | -0.069406148 |
| ENSDART00000122605 | atxn7l1 | -0.290852975 | -0.415130691 | -0.145082777 | -0.005809908 |
| ENSDART00000122617 | si:ch211-24o10.6 | -0.23677423 | -0.709531228 | 3.09570127 | -0.222670409 |
| ENSDART00000122628 | junba | 0.280813702 | 0.583374691 | 0.41181471 | 0.073998168 |
| ENSDART00000122632 | si:ch211-173d10.4 | -0.251574462 | -0.294229692 | -0.256969485 | -0.062618481 |
| ENSDART00000122654 | rnd1a | -0.834540827 | -0.662552714 | -0.915729348 | -0.640521788 |
| ENSDART00000122665 | mta3 | -0.481842523 | -0.542700403 | 0.031188286 | 0.042011127 |
| ENSDART00000122681 | CR848841.1 | 0.726082565 | 1.593466942 | 1.807590182 | 1.450515621 |
| ENSDART00000122682 | cabp1b | -0.294760944 | -0.547217913 | -0.530251845 | -0.289128709 |
| ENSDART00000122692 | CABZ01112317.1 | 0.655801165 | 0.399271741 | 0.354973485 | -0.117162708 |
| ENSDART00000122696 | hnrnpub | 0.216889898 | 0.1794005 | 0.293328342 | 0.140752076 |
| ENSDART00000122700 | tenm3 | 0.157516404 | 0.267363723 | 0.981020214 | 1.113087141 |
| ENSDART00000122716 | PDE1C | -0.224024171 | -0.483764938 | -0.500061121 | -0.40054702 |
| ENSDART00000122742 | rorcb | -0.773981248 | 0.298893511 | 0.803866913 | 0.643975269 |
| ENSDART00000122747 | tmem30aa | -0.229416393 | -0.299526834 | -0.201671762 | -0.059032826 |
| ENSDART00000122768 | CU984579.1 | 0.668419525 | 0.524624014 | 0.112892825 | -0.531459653 |
| ENSDART00000122796 | rgl3a | -0.364392925 | -0.529093083 | -0.585369163 | -0.370045273 |

| | | | | | |
|---|---|---|---|---|---|
| ENSDART00000122803 | usp9 | 0.139967851 | 0.094990222 | 0.369664537 | 0.232490119 |
| ENSDART00000122829 | pik3r1 | -0.369164024 | -0.336790822 | -0.454949598 | -0.226248156 |
| ENSDART00000122863 | ptgs1 | -0.31160045 | -0.175184364 | -0.407394512 | -0.554967944 |
| ENSDART00000122889 | myhz1.3 | 0.065159767 | 3.686062072 | 4.139837605 | 2.765333679 |
| ENSDART00000122891 | tmem97 | 0.209697033 | 0.237651064 | 0.43725776 | 0.414256644 |
| ENSDART00000122898 | phf21b | 0.630758244 | -0.049111481 | 0.173874589 | 0.290793166 |
| ENSDART00000122905 | rac3b | -0.501948401 | -0.363291367 | -0.120049924 | 0.166932504 |
| ENSDART00000122924 | CABZ01089151.1 | 0.66835416 | 0.574150269 | 1.297855118 | 0.40982894 |
| ENSDART00000122929 | bbs4 | -0.14534671 | -0.308002271 | -0.496501405 | -0.252523436 |
| ENSDART00000122930 | scg3 | -0.159257317 | -0.25593578 | -0.408715103 | -0.119340458 |
| ENSDART00000122945 | noc2l | 0.680401135 | 0.443128484 | 0.389864365 | 0.196289408 |
| ENSDART00000122953 | gnl3 | 0.534095037 | 0.261409226 | 0.267291075 | 0.063642911 |
| ENSDART00000122966 | hapln1a | -5.328261829 | -0.041096146 | -1.913311737 | -3.425165696 |
| ENSDART00000123002 | nme4 | 3.640965481 | 3.577167408 | 3.045307609 | 2.652336979 |
| ENSDART00000123040 | pfkpb | -0.26169445 | -0.241526904 | -0.415842652 | -0.206785479 |
| ENSDART00000123063 | fgf18a | 0.196370642 | 0.632761225 | 0.668396752 | 0.413800422 |
| ENSDART00000123081 | zgc:173552 | 0.884992881 | 1.080371934 | 1.302685665 | 0.777637365 |
| ENSDART00000123096 | marcksa | 1.85087225 | 2.605359702 | 2.620602183 | 2.013819083 |
| ENSDART00000123117 | ggh | 0.485870455 | 0.424307993 | 0.067357919 | -0.14640404 |
| ENSDART00000123136 | FP236812.4 | 0.346817394 | 0.374427758 | 0.928731714 | 0.426600457 |
| ENSDART00000123152 | cpeb4a | -0.291020391 | -0.445618867 | -0.40169983 | -0.306227449 |
| ENSDART00000123203 | vbp1 | 0.24089733 | 0.440645383 | 0.41864886 | 0.239214051 |
| ENSDART00000123254 | dpp9 | 0.199548032 | 0.283576523 | 0.332661933 | 0.196698648 |
| ENSDART00000123263 | phf20a | 3.204242437 | 4.684552705 | 5.028378766 | 3.758919167 |
| ENSDART00000123278 | pikfyve | 0.216525391 | 0.503566219 | -0.050873981 | 0.098099496 |
| ENSDART00000123282 | si:dkey-7i4.1 | -0.131885839 | -0.288104676 | -0.328002164 | -0.299371464 |
| ENSDART00000123338 | kif3a | 0.128582487 | 0.488909245 | 0.490973182 | 0.361344362 |
| ENSDART00000123350 | sptlc2a | 0.577620333 | 0.063030792 | -0.107114764 | -0.718762785 |
| ENSDART00000123359 | nlgn1 | -0.188766802 | -0.577714477 | -0.350080443 | 0.032932651 |
| ENSDART00000123360 | susd5 | -0.191753051 | -0.73208098 | -0.853142314 | -0.526763596 |
| ENSDART00000123364 | vegfba | 0.24629132 | 0.079711009 | -0.059768872 | -0.046839233 |
| ENSDART00000123380 | KCNIP4 | -0.549899676 | -0.507571644 | -0.455586059 | -0.106289449 |
| ENSDART00000123381 | si:ch211-117k10.3 | -0.057115592 | -0.184208007 | -0.285329447 | -0.136081942 |
| ENSDART00000123392 | rab11bb | -0.044482336 | 0.125801068 | 0.511028514 | 0.477170112 |
| ENSDART00000123407 | pdzd7b | -0.257572727 | -0.396050586 | -0.283495799 | -0.108489618 |
| ENSDART00000123409 | si:ch1073-296d18.1 | 0.174260872 | 0.155958113 | 0.452524339 | -0.02334577 |
| ENSDART00000123450 | KCNB2 (1 of many) | -0.271043423 | -0.632402343 | -0.828557597 | -0.168958818 |
| ENSDART00000123473 | Metazoa_SRP | 0.899034068 | 1.683872421 | 0.662894374 | -0.48789514 |
| ENSDART00000123479 | | -0.276038029 | -0.380729607 | -0.780200096 | -0.278407445 |
| ENSDART00000123505 | rab11fip4a | -0.154164113 | -0.404988879 | -0.479622862 | -0.179766383 |
| ENSDART00000123506 | si:dkeyp-115e12.6 | 0.611372618 | 1.198993764 | 1.526686816 | 0.680010003 |
| ENSDART00000123518 | tuba1b | 0.343846477 | 0.74266829 | 1.086564506 | 0.883480954 |
| ENSDART00000123519 | ube2e1 | 0.178513765 | 0.355426845 | 0.339753979 | 0.0793807 |
| ENSDART00000123534 | cacna1g | -0.255311686 | -0.53660486 | -0.469326401 | -0.279013734 |
| ENSDART00000123544 | tex2 | 0.532623113 | 0.735572961 | 0.212010938 | -0.135781649 |
| ENSDART00000123559 | znf521 | 1.710422257 | 2.255410725 | 1.850379402 | 0.814044199 |
| ENSDART00000123568 | pmela | 0.634272858 | 0.294790377 | -0.243906899 | -1.030971827 |
| ENSDART00000123590 | si:ch211-91p5.3 | 0.174508046 | -0.021173312 | 0.906803099 | -0.35666968 |
| ENSDART00000123607 | cyfip2 | 0.234079405 | 0.294840867 | 0.917405124 | 0.783474958 |
| ENSDART00000123634 | CABZ01076616.1 | -0.026473104 | -0.096132504 | -0.342132107 | -0.122009843 |
| ENSDART00000123648 | pcdh1a3 | -0.182322903 | 0.28663259 | 1.138716215 | 1.048655391 |

| | | | | | |
|---|---|---|---|---|---|
| ENSDART00000123713 | gria1b | -0.296227004 | -0.446226017 | -0.161357303 | 0.142828716 |
| ENSDART00000123797 | pcp4b | -0.598256523 | -0.489822031 | -0.568163011 | -0.081816834 |
| ENSDART00000123820 | tmem135 | -0.104475369 | -0.143585965 | -0.370543783 | -0.122253976 |
| ENSDART00000123834 | exoc7 | 0.170713952 | 0.17934497 | 0.358341658 | 0.328688768 |
| ENSDART00000123839 | arrb1 | -0.44263739 | -0.489445552 | -0.281663489 | 0.083496974 |
| ENSDART00000123844 | si:dkey-77f5.3 | -4.277427291 | -0.495308507 | -0.670171681 | -0.698447042 |
| ENSDART00000123851 | fut8b | -0.379186004 | -0.673089848 | -0.195485297 | 0.141395943 |
| ENSDART00000123868 | dthd1 | -0.700427011 | -0.013611963 | -0.073941373 | -0.012612052 |
| ENSDART00000123878 | ppiaa | 0.40248836 | 0.711363911 | 0.687992513 | 0.279031014 |
| ENSDART00000123887 | MAPK8IP1 (1 of many) | -0.420573616 | -0.373470966 | -0.247368538 | -0.089797668 |
| ENSDART00000123917 | tmem47 | -0.036122446 | 0.257386953 | 0.324417161 | 0.235522978 |
| ENSDART00000123950 | PRIMA1 | -0.578840887 | -0.634067967 | -0.550475282 | -0.159964437 |
| ENSDART00000123966 | chrng | 1.138454485 | 1.750205793 | 1.604781052 | 0.687837172 |
| ENSDART00000123970 | mntb | -0.115861458 | -0.101461538 | -0.360694649 | -0.01567692 |
| ENSDART00000123972 | smarca5 | 0.189313952 | 0.223635359 | 0.34449924 | 0.204975788 |
| ENSDART00000123999 | SLC27A2 (1 of many) | -1.114337124 | -0.48467376 | -0.495436914 | -0.504796953 |
| ENSDART00000124040 | insm1a | 0.50962443 | 1.551047216 | 1.738068624 | 1.104992333 |
| ENSDART00000124075 | mtss1lb | -0.362597648 | -0.323844879 | 0.066492818 | 0.131710049 |
| ENSDART00000124085 | usp18 | 0.058285571 | 0.001224867 | 0.967078356 | -0.263223461 |
| ENSDART00000124112 | pou4f2 | -1.368026557 | -1.308261738 | 0.344262724 | 0.908586045 |
| ENSDART00000124140 | zgc:101566 | -0.242404826 | -0.290163127 | -0.636219456 | -0.416307006 |
| ENSDART00000124176 | gfpt1 | 0.432149516 | 0.480766327 | 0.474022237 | 0.242457652 |
| ENSDART00000124179 | hist1h4l | 0.531035238 | 0.617056656 | 0.928084772 | 0.625350282 |
| ENSDART00000124217 | map1sb | -0.319538816 | -0.066329928 | 0.070606275 | 0.217460247 |
| ENSDART00000124220 | nptxrb | -0.324541852 | -0.000122151 | 0.687818829 | 0.894343302 |
| ENSDART00000124242 | lamp2 | 0.063109036 | 0.101381463 | -0.092849158 | -0.275072982 |
| ENSDART00000124244 | rtn3 | -0.261781297 | -0.269593684 | -0.376187628 | -0.121031259 |
| ENSDART00000124290 | MFN1 | -0.108610875 | -0.185612855 | -0.471855683 | -0.083062873 |
| ENSDART00000124298 | klhl29 | -0.069964723 | -0.409007114 | -0.117673983 | 0.023652831 |
| ENSDART00000124300 | rhoab | 0.400265647 | 0.097339191 | -0.031332977 | -0.120716407 |
| ENSDART00000124306 | | 0.869134686 | 1.333529821 | 1.120901364 | 0.868931063 |
| ENSDART00000124327 | havcr2 | 0.988496388 | 0.35234238 | 0.213092908 | 0.061810802 |
| ENSDART00000124329 | cbx8a | -0.13552578 | -0.23364206 | -0.31740894 | -0.19855692 |
| ENSDART00000124331 | si:ch1073-174d20.1 | -0.948877018 | -0.570444361 | 0.133772468 | 0.53293925 |
| ENSDART00000124333 | adcy5 | -0.349730091 | -0.163331233 | -0.158561989 | -0.605303683 |
| ENSDART00000124335 | smim4 | -0.115055175 | -0.371982424 | -0.338416379 | -0.248500803 |
| ENSDART00000124346 | fn1a | 1.458758212 | 2.264590948 | 2.662817487 | 2.142590168 |
| ENSDART00000124367 | si:cabz01074946.1 | 1.457343255 | 1.182560392 | 0.713827232 | 0.020183118 |
| ENSDART00000124439 | cpn1 | 0.590321112 | 0.454790314 | -0.048032667 | -0.505920429 |
| ENSDART00000124440 | CABZ01029822.1 | 0.125194338 | 0.049659662 | 0.670452016 | 0.712162088 |
| ENSDART00000124455 | fbxl15 | -0.435034118 | -0.27455581 | -0.042060966 | 0.121857574 |
| ENSDART00000124480 | adgrl1a | -0.174945207 | -0.479872966 | -0.185768365 | 0.06038548 |
| ENSDART00000124485 | pcdh2ab8 | 0.093884444 | 0.182265582 | 0.466276394 | 0.290720698 |
| ENSDART00000124499 | hprt1 | 0.192517236 | 0.178234353 | 0.277835111 | 0.149971609 |
| ENSDART00000124534 | mbnl2 | -0.407254222 | 0.078580914 | -0.044925118 | 0.120266311 |
| ENSDART00000124562 | zgc:152977 | -0.424781995 | -0.464146326 | -0.707134675 | -0.367623104 |
| ENSDART00000124563 | | 3.317882457 | 3.087315331 | 3.467692067 | 2.846513731 |
| ENSDART00000124582 | napbb | -0.403034353 | -0.312472409 | -0.341096083 | -0.127811116 |
| ENSDART00000124653 | dhrs4 | 0.898977368 | 0.567421347 | 0.389421075 | 0.098869507 |
| ENSDART00000124660 | ascc3 | 0.285037378 | 0.269547515 | 0.334772964 | 0.241795126 |
| ENSDART00000124662 | rtn4a | -0.503868889 | -0.627866078 | -0.609450822 | -0.190620657 |

| | | | | | |
|---|---|---|---|---|---|
| ENSDART00000124663 | npc2 | 1.578129482 | 0.684715762 | 0.389649636 | 0.031719264 |
| ENSDART00000124676 | sv2ba | -0.120324523 | -0.401787287 | -0.561843381 | -0.191682717 |
| ENSDART00000124708 | gabra6b | -0.347504052 | -0.445856563 | -0.426704654 | -0.236544239 |
| ENSDART00000124710 | dlg5a | -0.356924394 | -0.265850728 | -0.130233283 | 0.096340819 |
| ENSDART00000124716 | si:dkeyp-121d4.3 | 0.246129368 | 0.104493361 | 0.091668358 | 0.120636876 |
| ENSDART00000124748 | b2ml | -0.109461573 | -0.036064969 | -0.083861942 | -0.342769695 |
| ENSDART00000124751 | kcnip3b | -1.29606503 | -1.383048631 | -0.490749215 | -0.039127508 |
| ENSDART00000124762 | hsp70.1 | -0.061931986 | -0.652822267 | -0.746588848 | -0.837604985 |
| ENSDART00000124773 | ppid | 0.236087087 | 0.322094513 | 0.445941037 | 0.285911702 |
| ENSDART00000124800 | fam212aa | -0.670414346 | -0.372526126 | -0.229531445 | -0.379904435 |
| ENSDART00000124809 | acsbg2 | 0.494562304 | 1.915512609 | 1.979085784 | 0.982887956 |
| ENSDART00000124827 | lgi2a | -0.389885576 | -0.462014195 | -0.106597692 | 0.145509175 |
| ENSDART00000124833 | pdcd11 | 0.337926893 | 0.21951318 | 0.267537583 | 0.067942864 |
| ENSDART00000124843 | mtss1la | -0.820086229 | 0.028014444 | 0.433717014 | 0.003181165 |
| ENSDART00000124868 | lpl | 0.342987533 | -0.032282276 | -0.168396519 | -1.276175937 |
| ENSDART00000124876 | VSTM2B | -0.453041699 | -0.513162057 | -0.416504873 | -0.137923272 |
| ENSDART00000124925 | si:ch211-235e9.8 | -0.615429137 | -1.514463249 | -0.238786067 | 0.039474733 |
| ENSDART00000124945 | ubap1lb | 0.21207026 | -0.004652141 | -0.773302877 | -0.344881822 |
| ENSDART00000124963 | pkig | 0.251389387 | 0.584439023 | 0.334726577 | 0.10933368 |
| ENSDART00000124968 | rpn2 | 0.26137974 | 0.104649464 | 0.042914045 | -0.061671941 |
| ENSDART00000124991 | lmcd1 | 2.103120162 | 2.716646733 | 2.223480288 | 1.685964001 |
| ENSDART00000124998 | rtn2a | 0.520408534 | 1.345034307 | 1.147677305 | 0.715797789 |
| ENSDART00000125019 | dhcr24 | -0.273134055 | -0.003097208 | 0.45273437 | 0.079225052 |
| ENSDART00000125039 | six6b | -0.391804698 | -0.26201342 | -0.503562642 | -0.356304334 |
| ENSDART00000125045 | dscama | -0.36310557 | -0.486996676 | -0.080151729 | 0.074065642 |
| ENSDART00000125058 | nipsnap3a | 0.560274837 | 0.34716726 | 0.187463486 | 0.300128027 |
| ENSDART00000125074 | kcnab2b | -0.745923366 | -1.091983107 | -0.420367552 | 0.000775062 |
| ENSDART00000125097 | si:dkey-126g1.7 | 0.825737488 | 0.388125468 | -0.092297047 | -0.324389582 |
| ENSDART00000125116 | tnfaip8l3 | -0.29259694 | -0.291950956 | -0.382018775 | -0.337072666 |
| ENSDART00000125174 | nr1i2 | -0.056601149 | -0.085618729 | -0.264588441 | -0.326132843 |
| ENSDART00000125178 | eif5a | 0.322397945 | 0.394023524 | 0.161340886 | -0.011483865 |
| ENSDART00000125203 | hopx | 0.434700645 | 0.383648473 | 0.145485294 | -0.121364202 |
| ENSDART00000125281 | ngfra | -0.140591626 | 0.234351775 | 0.618758909 | 0.552257695 |
| ENSDART00000125284 | nlgn2a | -0.210046545 | -0.366872132 | 0.090424981 | 0.163948503 |
| ENSDART00000125299 | plk2a | -0.143584568 | -0.170521478 | -0.297471862 | -0.183865098 |
| ENSDART00000125302 | fbn2b | 0.974537401 | 0.789475875 | 1.035854452 | 0.178462064 |
| ENSDART00000125344 | skilb | -0.241964121 | -0.627384358 | -0.493621211 | -0.368838752 |
| ENSDART00000125348 | id2b | -0.501394328 | 0.114793842 | 0.840457629 | 0.257023029 |
| ENSDART00000125349 | bada | 0.528966306 | 1.266997252 | 1.043776037 | 0.694034948 |
| ENSDART00000125371 | mknk1 | 0.126961076 | 0.659607681 | 0.524109906 | -0.03336174 |
| ENSDART00000125381 | gig2o | 0.334408908 | 0.020553966 | 0.877629524 | -0.325957124 |
| ENSDART00000125397 | kri1 | -0.318299605 | -0.278223015 | -0.439932558 | -0.332258967 |
| ENSDART00000125430 | pprc1 | 1.538620876 | 1.109491459 | 0.874206496 | 0.220876028 |
| ENSDART00000125432 | esrrd | -0.825323711 | -0.251243419 | -0.11920169 | -0.154136212 |
| ENSDART00000125440 | hist1h4l | 0.76925705 | 0.664322574 | 0.48261672 | 0.298833148 |
| ENSDART00000125450 | gpc1a | -0.40101998 | -0.317779496 | 0.568583174 | 0.472068591 |
| ENSDART00000125466 | alpi.2 | -0.05537717 | -0.356245119 | -0.658920603 | -0.289506352 |
| ENSDART00000125468 | apodb | 0.705398759 | 0.818661135 | 1.046059167 | 0.463604749 |
| ENSDART00000125472 | dnmt3bb.3 | 2.081331813 | 1.555687378 | 1.386584326 | 1.824843519 |
| ENSDART00000125531 | plppr5a | -0.485992962 | -0.468686144 | -0.487868254 | -0.311270154 |
| ENSDART00000125536 | appb | -0.473539223 | -0.363304525 | 0.075744005 | 0.226987466 |

| | | | | | |
|---|---|---|---|---|---|
| ENSDART00000125561 | atp6ap2 | -0.203601896 | -0.343976801 | -0.371684174 | -0.265876436 |
| ENSDART00000125563 | homer3b | -0.366344126 | -0.72350131 | -0.2232749 | 0.112901448 |
| ENSDART00000125590 | adam22 | -0.13203003 | 0.438612977 | 0.776065364 | 0.829224171 |
| ENSDART00000125594 | zgc:194312 | 0.282121707 | 0.314684279 | 0.206211766 | 0.077189618 |
| ENSDART00000125617 | smoc1 | -0.728822769 | -0.318137722 | -0.435938883 | -0.283225846 |
| ENSDART00000125619 | thop1 | -0.701072314 | 0.174567303 | -0.277028493 | 0.035731797 |
| ENSDART00000125638 | tbx5b | -0.961612456 | -0.494056529 | -0.317150736 | -0.11065371 |
| ENSDART00000125674 | pcdh2ab9 | 0.154453003 | 0.05176964 | 0.233264506 | 0.354320756 |
| ENSDART00000125691 | sntb1 | -0.296403963 | -0.433378655 | -0.538134366 | -0.484556917 |
| ENSDART00000125733 | chst1 | -1.779723555 | -0.640531075 | -0.420086911 | -0.327548455 |
| ENSDART00000125753 | rpl22 | 0.364344172 | 0.362576681 | 0.16140506 | -0.121749389 |
| ENSDART00000125759 | si:dkey-22o12.2 | -0.433356276 | -0.386370833 | -0.195481295 | -0.118191078 |
| ENSDART00000125766 | camkvb | -0.518737667 | -0.560273381 | -0.364168693 | -0.104855004 |
| ENSDART00000125767 | metap2a | -0.473789427 | -0.340839673 | -0.148249289 | -0.006647252 |
| ENSDART00000125824 | tmem198a | -0.201432593 | -0.312524143 | -0.133551129 | -0.103080035 |
| ENSDART00000125835 | si:dkey-16p6.1 | 0.273905491 | 0.038923635 | 0.125549673 | 0.126712455 |
| ENSDART00000125864 | camkk1b | -0.169636507 | -0.512096721 | -0.591294824 | -0.14296099 |
| ENSDART00000125900 | hs3st3b1b | -0.2288214 | -0.294028167 | -0.23141286 | 0.023426889 |
| ENSDART00000125923 | cxcl14 | -0.090480689 | -0.043715373 | -0.409912244 | -0.49269465 |
| ENSDART00000125925 | tead1a | 0.393334189 | 0.562637179 | 0.380089957 | 0.085328359 |
| ENSDART00000125971 | gpc1b | 0.246742539 | 0.417837609 | 0.415208578 | 0.061504531 |
| ENSDART00000125979 | tshz3a | -0.321412605 | -0.323547538 | -0.394821529 | -0.291421679 |
| ENSDART00000126005 | efna5a | -0.170916 | 0.110640262 | 0.306287547 | 0.169125563 |
| ENSDART00000126022 | rarab | -0.000236754 | 0.039022735 | 0.1518943 | 0.316882235 |
| ENSDART00000126029 | dhx58 | 0.173247162 | -0.097510604 | 2.186112068 | 0.050096401 |
| ENSDART00000126038 | tcp1 | 0.327351769 | 0.30721299 | 0.45795341 | 0.229539742 |
| ENSDART00000126041 | ano10a | 0.295415156 | 0.394412522 | 0.244334431 | 0.10037004 |
| ENSDART00000126044 | FO904898.3 | 0.246202383 | 0.067501502 | -0.306505539 | -0.628858511 |
| ENSDART00000126076 | | -0.016912473 | -0.350302068 | -0.401962349 | -0.001679825 |
| ENSDART00000126084 | ctdspl3 | -0.310396194 | -0.278185547 | -0.408791973 | -0.234117517 |
| ENSDART00000126092 | irs4a | -0.078713301 | -0.053650022 | 0.329231719 | 0.37127028 |
| ENSDART00000126148 | zmp:0000000912 | 0.164571908 | 0.167838799 | 1.993878391 | 0.463546785 |
| ENSDART00000126172 | fn1b | 1.985622531 | 1.85759146 | 1.646183965 | 1.167188621 |
| ENSDART00000126196 | gucy2f | -0.393875584 | -0.283587572 | -0.315272826 | -0.060630266 |
| ENSDART00000126198 | lrrfip1a | 0.470310797 | 0.07807124 | -0.268319723 | -0.389271494 |
| ENSDART00000126205 | st6gal1 | -0.371204022 | -0.447413159 | -0.30616801 | 0.05897999 |
| ENSDART00000126229 | rps7 | 0.396514179 | 0.373862291 | 0.136780142 | -0.080022773 |
| ENSDART00000126253 | fam124b | -0.802465606 | 0.105938283 | 0.208441414 | 0.082147827 |
| ENSDART00000126259 | zgc:110434 | 0.950260296 | 1.108697637 | 1.201811754 | 0.695021733 |
| ENSDART00000126260 | anxa6 | 0.678382414 | 0.731151807 | 0.839317026 | 0.481712468 |
| ENSDART00000126282 | nr1d1 | -0.1014969 | -0.191656676 | -0.494430716 | -0.13667305 |
| ENSDART00000126299 | gdi1 | -0.113565456 | 0.226089497 | 0.711067572 | 0.636383115 |
| ENSDART00000126300 | shisa8b | -0.373408846 | -0.464508355 | -0.412634146 | -0.258826308 |
| ENSDART00000126301 | prkab1b | 0.061924737 | -0.091377696 | -0.203476199 | -0.507302968 |
| ENSDART00000126314 | pcdh8 | -0.25826413 | -0.454008614 | -0.231067015 | -0.062706758 |
| ENSDART00000126397 | tspan4a | -0.347274376 | -0.217924211 | -0.29861217 | -0.115670743 |
| ENSDART00000126414 | eml1 | 0.475872706 | 0.625227433 | 0.675164018 | 0.217118522 |
| ENSDART00000126417 | bri3 | 0.500014173 | 0.414528928 | 0.11265957 | -0.17961569 |
| ENSDART00000126444 | mmp13a | 0.046684026 | 3.040517608 | 1.846123489 | -1.625378496 |
| ENSDART00000126460 | il34 | 0.203388167 | 0.052375422 | -0.136564829 | -0.587275531 |
| ENSDART00000126470 | zgc:85777 | 0.313383122 | 0.223125376 | -0.098199946 | -0.16516057 |

| | | | | | |
|---|---|---|---|---|---|
| ENSDART00000126474 | si:ch211-265o23.1 | 1.117262816 | 1.407297939 | 1.180818753 | 0.875131455 |
| ENSDART00000126542 | ppp1r14ba | 0.329398236 | 1.23193061 | 1.655895613 | 1.416335991 |
| ENSDART00000126559 | rap1gap2a | -0.313212246 | -0.347504992 | -0.446138134 | -0.231468434 |
| ENSDART00000126565 | si:ch211-150g13.3 | -0.477986184 | -0.41948879 | 0.219835784 | 0.507153945 |
| ENSDART00000126578 | si:ch211-200e2.1 | -0.112922065 | -0.175836986 | -0.289153296 | -0.021149659 |
| ENSDART00000126588 | rac3a | -0.258496234 | 0.207987177 | 0.575325222 | 0.581016326 |
| ENSDART00000126621 | pfdn1 | 0.222765458 | 0.317529573 | 0.180735937 | 0.092770718 |
| ENSDART00000126634 | si:dkey-237m9.2 | 2.393787497 | 2.476689815 | 2.575358161 | 0.828914545 |
| ENSDART00000126640 | RHOBTB3 | -0.934122492 | -0.733675156 | -0.213973967 | -0.180952113 |
| ENSDART00000126661 | rhobtb1 | -0.165202139 | -0.331565823 | -0.522361525 | -0.299461153 |
| ENSDART00000126681 | tardbpl | 0.019310972 | 0.344433444 | 0.212149787 | 0.045304759 |
| ENSDART00000126705 | eif3ea | 0.116700572 | 0.376861812 | 0.239268123 | 0.089153507 |
| ENSDART00000126739 | tspan5a | -0.28156215 | -0.24229244 | -0.35515584 | -0.059588327 |
| ENSDART00000126744 | nrg3b | -0.484773132 | -0.477903604 | -0.212248622 | 0.011007147 |
| ENSDART00000126766 | erbb4b | -0.378985194 | -0.566156911 | -0.016423806 | 0.214915468 |
| ENSDART00000126768 | cc2d2a | 0.207920961 | 0.256447903 | 0.267293619 | 0.323639423 |
| ENSDART00000126780 | rnaset2 | 0.561676509 | 0.243610345 | -0.050562112 | -0.093659569 |
| ENSDART00000126826 | tmem59 | 0.0843517 | -0.210535259 | -0.365503975 | -0.184419203 |
| ENSDART00000126842 | flj11011l | -0.120546184 | -0.161649138 | -0.295902143 | -0.182318736 |
| ENSDART00000126845 | ldb1a | -0.250500011 | -0.460439315 | -0.246081535 | -0.230460175 |
| ENSDART00000126866 | ccser2b | 0.139738782 | 0.390945754 | 0.269322283 | 0.164198289 |
| ENSDART00000126870 | ndufa4l2b | -0.364506533 | -0.363911454 | -0.576856062 | -0.429052776 |
| ENSDART00000126873 | myhz2 | -2.902978535 | 4.975016398 | 5.62978424 | 4.2721239 |
| ENSDART00000126891 | sept2 | 0.631165801 | 0.418693285 | 0.081387841 | -0.156566045 |
| ENSDART00000126897 | lamtor5 | 0.078015812 | -0.030543248 | -0.309612502 | -0.278777082 |
| ENSDART00000126916 | mob2b | 0.26836075 | -0.004670217 | -0.398977994 | -0.072830738 |
| ENSDART00000126928 | si:dkey-73n8.3 | -0.354837143 | 0.244926012 | -3.843926683 | -0.176933271 |
| ENSDART00000126945 | adgrb3 | -0.310129925 | -0.752937386 | -0.198981749 | -0.020103794 |
| ENSDART00000126963 | ucp3 | -1.126828953 | 2.695519216 | 3.149953006 | 2.002864913 |
| ENSDART00000126966 | esrrga | -0.248566713 | -0.265310961 | -0.321308408 | -0.070523791 |
| ENSDART00000126978 | acadl | 0.315070362 | 0.266918423 | -0.034640916 | -0.111553388 |
| ENSDART00000126979 | ubxn1 | 0.239657791 | 0.430777506 | 0.361799544 | 0.043577807 |
| ENSDART00000126991 | ascl1b | 0.538037435 | 0.289776062 | 1.627554047 | -0.154081652 |
| ENSDART00000127009 | | 0.669374866 | 0.368365497 | 0.110306036 | -0.47074427 |
| ENSDART00000127015 | srrm4 | -0.1898883 | -0.282207317 | -0.262672915 | -0.106216594 |
| ENSDART00000127047 | tusc5a | -1.04192916 | -0.556828029 | 0.377559818 | 0.604205052 |
| ENSDART00000127050 | nrxn3b | -0.323579219 | -0.438342846 | -0.422724255 | 0.054279865 |
| ENSDART00000127062 | egr3 | -1.907539966 | -1.048685041 | -1.405658043 | -1.087307214 |
| ENSDART00000127073 | lrrc75bb | -0.380397254 | -0.264539972 | -0.318155891 | -0.090026751 |
| ENSDART00000127093 | CR749162.1 | -1.06520168 | -0.24719149 | -0.825954556 | -0.565925266 |
| ENSDART00000127099 | nr2e3 | 0.281603482 | 0.002911001 | -0.560613973 | -0.033774044 |
| ENSDART00000127131 | mtmr10 | 0.05904867 | -0.240178 | -0.607174209 | -0.120957306 |
| ENSDART00000127136 | slco5a1b | -0.145489914 | -0.411558687 | 0.007460079 | -0.021875568 |
| ENSDART00000127144 | cacna2d2a | -0.206435603 | -0.394190367 | -0.167540436 | 0.000520565 |
| ENSDART00000127157 | hlfa | -0.089881453 | -0.212609643 | -0.346930267 | -0.147102401 |
| ENSDART00000127162 | si:dkey-165a24.9 | -0.154532095 | -0.18408156 | -0.261561109 | -0.381673579 |
| ENSDART00000127167 | fam217b | -0.253255723 | -0.39768374 | -0.389011868 | -0.208058743 |
| ENSDART00000127173 | clu | 0.051800584 | 0.063760613 | -0.14536086 | -0.386244567 |
| ENSDART00000127214 | tmtc4 | -0.125359905 | 0.4024138 | 0.47318852 | 0.32908352 |
| ENSDART00000127216 | fam161a | -0.082566753 | -0.243407902 | -0.44855597 | -0.155296925 |
| ENSDART00000127219 | FAM155A (1 of many) | -0.247003441 | -0.575960705 | -0.390291652 | -0.269667553 |

| | | | | | |
|---|---|---|---|---|---|
| ENSDART00000127236 | syt13 | -0.555582206 | -0.558950845 | -0.738809627 | -0.549910893 |
| ENSDART00000127239 | si:ch211-237i5.4 | -0.435620687 | -0.33667742 | 0.058425883 | -0.022982403 |
| ENSDART00000127274 | sema7a | -0.198678021 | -0.353988759 | -0.478218296 | -0.427230313 |
| ENSDART00000127277 | pum3 | 0.522067564 | 0.375606403 | 0.43721492 | 0.160017235 |
| ENSDART00000127286 | si:ch73-140j24.4 | -0.467040467 | -0.569237917 | -0.172853738 | 0.202142114 |
| ENSDART00000127318 | frmpd1b | -0.135556639 | -0.245654095 | -0.299228813 | -0.069164239 |
| ENSDART00000127331 | cdc37l1 | -0.115815783 | -0.102908381 | -0.388723888 | -0.245749713 |
| ENSDART00000127350 | enpp1 | 1.121109181 | 0.817391101 | 0.578924138 | 0.065420517 |
| ENSDART00000127353 | nr1d2b | -0.29103008 | -0.243688493 | -0.346507775 | -0.100908835 |
| ENSDART00000127355 | CR855274.1 | -1.80396628 | -1.678561981 | -0.998732969 | -1.157402562 |
| ENSDART00000127396 | ppm1lb | -0.618109529 | -0.371965944 | 0.29932303 | 0.461142505 |
| ENSDART00000127397 | utp15 | 0.476513751 | 0.260847638 | 0.257137466 | 0.080743511 |
| ENSDART00000127398 | CR753886.1 | -0.159141993 | -0.332294072 | -0.331878243 | -0.244595408 |
| ENSDART00000127420 | MDP1 | 1.927039626 | 2.737884833 | 2.522148059 | 1.176014094 |
| ENSDART00000127426 | ERBB4 (1 of many) | -0.100134914 | -0.701677799 | -0.360350675 | -0.397963746 |
| ENSDART00000127428 | peli3 | -0.233448958 | -0.50115077 | -0.383840384 | -0.13792159 |
| ENSDART00000127467 | pygmb | -0.413201582 | -0.432147367 | -0.560055907 | -0.134010249 |
| ENSDART00000127496 | loxl1 | 0.412088952 | 0.133411325 | -0.193765804 | -0.987510306 |
| ENSDART00000127502 | cdkl5 | -0.227663486 | -0.408317592 | -0.318577534 | 0.048062851 |
| ENSDART00000127504 | trabd2a | -0.34035671 | -0.379312956 | -0.491238318 | -0.073447564 |
| ENSDART00000127506 | si:ch73-380n15.2 | -0.654968439 | -0.466259045 | -0.336236181 | -0.320196074 |
| ENSDART00000127508 | thnsl2 | 0.534241071 | 0.753527133 | 0.557550473 | 0.458352807 |
| ENSDART00000127513 | znf385d | -0.506377127 | -0.828614957 | -0.387596815 | -0.346943547 |
| ENSDART00000127536 | cbx7b | -0.401352835 | -0.168978702 | -0.391659396 | -0.597817617 |
| ENSDART00000127568 | ankha | -0.068226585 | -0.359480026 | -0.686453521 | -0.222767571 |
| ENSDART00000127581 | trim23 | -0.305887304 | -0.304885837 | -0.114854094 | -0.109504112 |
| ENSDART00000127654 | tagapb | -0.160683989 | -0.122717735 | -0.284237949 | -0.235548963 |
| ENSDART00000127656 | cln6b | -0.363769389 | -0.208045101 | -0.481848131 | -0.19203643 |
| ENSDART00000127673 | si:ch73-362m14.2 | -0.314971012 | -0.45829899 | -0.590197547 | -0.155316982 |
| ENSDART00000127693 | vps4b | 0.2667443 | 0.328628912 | 0.300618305 | 0.02818525 |
| ENSDART00000127695 | rpl5b | 0.469645013 | 0.524119827 | 0.418266352 | 0.096930554 |
| ENSDART00000127699 | si:ch73-335l21.4 | 1.682684641 | 2.104012275 | 1.803809975 | 0.730936452 |
| ENSDART00000127717 | FO704622.1 | 0.358399682 | 0.167598174 | 1.49594356 | -0.017716595 |
| ENSDART00000127719 | kdm6ba | 0.187905295 | -0.040637976 | 0.338835017 | 0.306420124 |
| ENSDART00000127753 | adam15 | 0.405103987 | 0.059018909 | -0.196195339 | -0.053874543 |
| ENSDART00000127769 | plekhg7 | -0.272021337 | -0.179975164 | -0.292604049 | -0.294852391 |
| ENSDART00000127795 | tsku | 0.321336842 | 0.111241615 | -0.27306537 | -0.619712406 |
| ENSDART00000127834 | zeb2a | -0.12592194 | -0.244794968 | -0.205587566 | -0.079770723 |
| ENSDART00000127854 | ccni | 3.365173646 | 3.708027214 | 3.915718651 | 4.117994882 |
| ENSDART00000127859 | trim63b | 0.3316873 | 0.607011237 | 0.873528277 | 0.725631694 |
| ENSDART00000127936 | prdm2a | -0.397627804 | -0.350024919 | -0.182979057 | 0.028300134 |
| ENSDART00000127971 | TMEM184B (1 of many) | 0.084681099 | 0.512647767 | 0.678333061 | 0.362195553 |
| ENSDART00000127974 | lrrk1 | 0.237699588 | 0.043061851 | 0.529534709 | 0.581549893 |
| ENSDART00000127977 | CABZ01016547.1 | 2.8163175 | 2.999575494 | 2.826245437 | 2.227407522 |
| ENSDART00000127981 | rgs9b | -0.268928405 | -0.181867875 | -0.388691411 | -0.111378496 |
| ENSDART00000127982 | suox | 0.48894768 | 0.565232546 | 0.460591806 | 0.13079859 |
| ENSDART00000128011 | tnfsf10l3 | -0.207145482 | -0.351713028 | -0.904110327 | -0.837055548 |
| ENSDART00000128014 | dnah2 | -0.407226928 | -0.498892771 | -0.604204962 | -0.453086208 |
| ENSDART00000128024 | wdsub1 | -2.995723015 | 0.554879586 | 0.200121371 | 0.371936521 |
| ENSDART00000128049 | hunk | 0.851411376 | 0.241049411 | 0.130293149 | 0.977016396 |
| ENSDART00000128050 | UTP14C | 0.741190051 | 0.667056074 | 0.681031837 | 0.432356782 |

| | | | | | |
|---|---|---|---|---|---|
| ENSDART00000128053 | tmem121b | -0.393762063 | -0.457536323 | -0.455856107 | -0.077381852 |
| ENSDART00000128127 | slc38a3b | -0.302545777 | -0.299963192 | -0.361680425 | -0.612800784 |
| ENSDART00000128149 | fam32a | -0.07314666 | -0.18816053 | -0.368263865 | -0.361807904 |
| ENSDART00000128174 | foxp1b | 0.060807795 | 0.283454627 | 0.694455914 | 0.748409918 |
| ENSDART00000128187 | mbnl2 | -0.194930621 | -0.139118187 | -0.302093023 | -0.22924185 |
| ENSDART00000128193 | rnf152 | 0.402113699 | 0.480129081 | 0.225745135 | 0.017035201 |
| ENSDART00000128198 | scn12aa | -0.298887564 | -0.309751416 | -0.23551982 | -0.15404656 |
| ENSDART00000128223 | CHCHD5 | -0.405489857 | -0.5400056 | -0.416697607 | -0.210022955 |
| ENSDART00000128224 | FO904873.1 | -0.267293261 | -0.580808884 | -0.319669262 | -0.131136583 |
| ENSDART00000128226 | slc6a11b | -0.388946235 | -0.51506191 | -0.246391039 | 0.364300065 |
| ENSDART00000128246 | polr1c | 0.52692078 | 0.422008081 | 0.272814737 | 0.11358433 |
| ENSDART00000128247 | lsm3 | -0.046938555 | -0.089126233 | -0.195168175 | -0.315381754 |
| ENSDART00000128271 | si:dkey-114c15.7 | -0.608254547 | -0.511435956 | -0.075108717 | 0.021034155 |
| ENSDART00000128281 | cntn1a | 0.147578203 | 0.564199709 | 0.636356286 | 0.425015924 |
| ENSDART00000128284 | nudt9 | 0.256265084 | 0.958235099 | 0.875663921 | 0.386996856 |
| ENSDART00000128302 | stk35 | -0.274652044 | -0.340334252 | -0.095489621 | 0.00314331 |
| ENSDART00000128346 | si:ch211-108c6.2 | 1.285191812 | 1.464423055 | 1.491945601 | 0.337219819 |
| ENSDART00000128350 | rad51d | 0.854501715 | 0.522915635 | 0.508614026 | 0.238331303 |
| ENSDART00000128457 | ephb1 | 0.374973875 | 0.620484905 | 1.032855939 | 0.805207117 |
| ENSDART00000128488 | e2f8 | 3.852661131 | -0.929008264 | 3.252279997 | 2.01178299 |
| ENSDART00000128497 | nat14 | -0.333855923 | -0.175587715 | -0.007501498 | 0.001822083 |
| ENSDART00000128535 | dmbx1a | -3.533682655 | -2.208339716 | -0.232250672 | -0.456601031 |
| ENSDART00000128550 | palb2 | 0.912196863 | 1.171643148 | 0.786370898 | 0.077481484 |
| ENSDART00000128613 | rps15a | 0.315521947 | 0.355667698 | 0.260468714 | -0.049823228 |
| ENSDART00000128624 | sgk494a | 0.606573335 | 0.92915728 | 0.662849036 | 0.240608511 |
| ENSDART00000128659 | pcdh2ab10 | 0.122813964 | 0.17598472 | 0.479887931 | 0.226887021 |
| ENSDART00000128673 | CABZ01078944.1 | 0.788095742 | 0.425650182 | 0.277053304 | -0.16966963 |
| ENSDART00000128681 | DYRK3 | 0.346870394 | 0.024160089 | 0.051669126 | 0.414421342 |
| ENSDART00000128690 | agap3 | -0.218144691 | -0.339443752 | 0.012538674 | 0.008782978 |
| ENSDART00000128696 | CABZ01035108.1 | -0.30766878 | -0.382099096 | -0.769557831 | -0.298631085 |
| ENSDART00000128698 | rnf34b | -0.122706526 | -0.390373676 | -0.658840165 | -0.257572429 |
| ENSDART00000128705 | ndrg3a | -0.684036937 | -0.148509308 | 0.416765356 | 0.500717941 |
| ENSDART00000128715 | kif3cb | 0.538226215 | 1.727920879 | 1.708926367 | 1.162753729 |
| ENSDART00000128721 | rcvrn3 | -0.045382046 | -0.143627209 | -0.579607896 | -0.393091134 |
| ENSDART00000128722 | RGS9BP | -0.328845343 | -0.481472863 | -0.560905879 | -0.19840165 |
| ENSDART00000128742 | CABZ01118678.1 | 0.234931395 | 0.748418816 | 0.975879993 | 0.726138214 |
| ENSDART00000128743 | tmem214 | 0.301583498 | 0.153317199 | 0.10691428 | 0.053655055 |
| ENSDART00000128760 | pacs2 | -0.00152054 | 0.063293894 | 0.531537781 | 0.626023304 |
| ENSDART00000128762 | dedd1 | -0.117364287 | -0.167908679 | -0.255046257 | -0.080222107 |
| ENSDART00000128763 | si:ch211-204c21.1 | 2.511635154 | 2.527602305 | 2.787303506 | 2.348876075 |
| ENSDART00000128784 | vmhcl | 0.489881537 | 1.799102022 | 2.075276497 | 1.004457616 |
| ENSDART00000128786 | pyya | 0.897714064 | 0.120108152 | -0.15783225 | -0.181132487 |
| ENSDART00000128794 | gpm6ab | -0.753205656 | -0.206704843 | 0.388986514 | 0.547688547 |
| ENSDART00000128807 | gpr22a | -0.50401992 | -0.32421881 | -0.337086094 | -0.059033497 |
| ENSDART00000128839 | fez1 | -0.463224643 | -0.323499809 | -0.160691051 | 0.039189596 |
| ENSDART00000128859 | prlra | -0.153277376 | -0.232075961 | -0.477332275 | -0.11182176 |
| ENSDART00000128894 | dchs1b | 0.298598944 | 0.508598812 | 0.623831745 | 0.178131833 |
| ENSDART00000128917 | rps4x | 0.379768803 | 0.358465257 | 0.299276991 | 0.035010431 |
| ENSDART00000128931 | igf2bp2a | 1.122085018 | 1.168437537 | 1.475101165 | 0.642256664 |
| ENSDART00000128959 | zgc:113223 | -1.993126278 | -0.750526857 | -1.059101824 | -0.610323514 |
| ENSDART00000128965 | bcar1 | 0.218283788 | 0.254312241 | 0.324642744 | 0.284302999 |

| | | | | | |
|---|---|---|---|---|---|
| ENSDART00000128969 | slit1a | -1.459038277 | -0.917332072 | -0.293864838 | 0.107599131 |
| ENSDART00000128971 | gm2a | 0.278528878 | 0.08350244 | -0.093671452 | -0.160146628 |
| ENSDART00000128975 | slc25a32a | 0.048183005 | -0.130483176 | -0.493409396 | -0.216291506 |
| ENSDART00000128978 | CR391998.1 | 1.262775633 | 1.767194773 | 1.40829971 | 2.265912659 |
| ENSDART00000128981 | | -2.281844224 | 0.709538363 | 0.831386018 | 0.078800359 |
| ENSDART00000129032 | CR847543.1 | 0.382490935 | 0.466481864 | 0.639639363 | 0.274235342 |
| ENSDART00000129033 | si:ch211-232b12.5 | 0.004752407 | 0.255447225 | 0.704679929 | 0.201922932 |
| ENSDART00000129035 | prdx5 | -0.081081909 | -0.174815089 | -0.303582143 | -0.169217832 |
| ENSDART00000129055 | nrg2b | -0.525587506 | -0.415280837 | -0.153878493 | 0.206094594 |
| ENSDART00000129058 | rheb | -0.430188972 | -0.312082399 | -0.301953935 | -0.04976624 |
| ENSDART00000129062 | ikbke | 0.350594041 | 0.34624224 | 0.094974339 | -0.516125523 |
| ENSDART00000129151 | strbp | -0.586891224 | -0.661538944 | -0.132603114 | 0.173676112 |
| ENSDART00000129155 | vmp1 | 3.054036254 | 2.972129495 | 2.202758828 | 1.232955319 |
| ENSDART00000129156 | adcy7 | -0.081832737 | -0.232314587 | -0.506962474 | -0.282923342 |
| ENSDART00000129167 | mapre1b | -0.039417049 | 0.272931214 | 0.675389418 | 0.445435787 |
| ENSDART00000129189 | slc25a38b | -0.383549453 | -0.49018808 | -0.499376221 | -0.30515595 |
| ENSDART00000129202 | DDX17 | -0.231356945 | -0.275266303 | -0.31249155 | -0.24318473 |
| ENSDART00000129210 | cry1ba | -0.306284135 | 0.315253029 | 0.635836839 | 0.449599168 |
| ENSDART00000129214 | COX7A2 (1 of many) | -0.085157083 | -0.364093354 | -0.62455009 | -0.475756885 |
| ENSDART00000129248 | clstn1 | -0.435055335 | -0.404808779 | -0.072836605 | 0.145009995 |
| ENSDART00000129254 | zgc:173552 | 0.679613651 | 0.919867905 | 0.987455267 | 0.613846632 |
| ENSDART00000129268 | fads2 | -0.454418186 | 0.562318002 | 0.686613427 | 0.310070087 |
| ENSDART00000129273 | rps6ka1 | -0.302406528 | -0.323918919 | -0.248745254 | -0.049749754 |
| ENSDART00000129308 | ACVR1C | -0.083617591 | -0.230914728 | -0.630759756 | -0.255735924 |
| ENSDART00000129362 | eef1a1l2 | 2.651521704 | 2.343527866 | 2.861864827 | 2.05813724 |
| ENSDART00000129365 | kcnb2 | -0.416222923 | -0.447467239 | -0.257551256 | -0.183749417 |
| ENSDART00000129380 | ppp5c | 0.718884884 | 0.082309464 | 0.464958134 | 0.512508903 |
| ENSDART00000129407 | plin2 | 0.385951592 | 0.445429516 | 0.411868628 | -0.015563455 |
| ENSDART00000129423 | slc37a1 | 0.300742512 | 0.561138843 | 0.611837666 | 0.153755959 |
| ENSDART00000129427 | vkorc1 | -0.08593935 | -0.003065091 | -0.36339301 | -0.313479955 |
| ENSDART00000129458 | fasn | -0.115300867 | 0.324414338 | 0.375920468 | 0.257409388 |
| ENSDART00000129459 | spata5l1 | 0.084981287 | 0.159429104 | 0.443203325 | 0.272680023 |
| ENSDART00000129471 | gucy1b3 | -0.32002288 | -0.161099136 | -0.090567082 | 0.034491801 |
| ENSDART00000129474 | ogmb | -0.463469684 | -0.477885606 | -0.227590281 | -0.024188229 |
| ENSDART00000129490 | gb:bc139872 | -0.183429225 | -0.314918476 | -0.330428185 | -0.307368048 |
| ENSDART00000129498 | mef2d | -0.188451231 | -0.381445449 | -0.263610552 | -0.197973462 |
| ENSDART00000129501 | tmem235 | -0.584898252 | -0.649073781 | -0.768355336 | -0.056926947 |
| ENSDART00000129511 | robo2 | -0.431501059 | -0.512946091 | -0.075617078 | 0.250070953 |
| ENSDART00000129521 | cpsf6 | -0.125055538 | -0.280935092 | -0.125914075 | -0.005682478 |
| ENSDART00000129530 | rps27a | 0.327713738 | 0.311901912 | 0.216145511 | -0.053626939 |
| ENSDART00000129559 | btbd3b | -0.741839183 | -0.720572279 | -0.364214548 | 0.15940566 |
| ENSDART00000129569 | gle1 | 2.578232986 | 2.574994626 | 4.293223748 | 4.445278491 |
| ENSDART00000129593 | znf217 | 0.260126556 | 0.54491492 | 0.40143457 | 0.121556273 |
| ENSDART00000129597 | sorbs3 | 0.016863628 | 0.072993125 | -0.314882072 | -0.546339357 |
| ENSDART00000129617 | atp6ap1b | 0.602272356 | 0.200096368 | -0.075510461 | -0.466117163 |
| ENSDART00000129643 | aplnrb | -0.038448881 | 0.28030901 | 0.821411842 | 0.34582741 |
| ENSDART00000129668 | fam57bb | -0.325481107 | -0.111394328 | 0.301848072 | 0.107888881 |
| ENSDART00000129674 | slc44a2 | 0.253178883 | 0.658899779 | 0.831363022 | 0.428394942 |
| ENSDART00000129679 | abcd4 | -0.064830814 | -0.217311868 | -0.258015712 | -0.43583137 |
| ENSDART00000129704 | abhd17aa | -0.089448965 | -0.057797962 | 0.474531723 | 0.44635249 |
| ENSDART00000129710 | crtac1a | -0.277382094 | 0.025987329 | -0.337990061 | -0.608225673 |

| | | | | | |
|---|---|---|---|---|---|
| ENSDART00000129744 | grin1a | -0.756444509 | -0.702815529 | -0.204482345 | 0.004357228 |
| ENSDART00000129749 | psmb7 | 0.370284184 | 0.518350984 | 0.484691258 | 0.174219127 |
| ENSDART00000129751 | si:ch211-105f12.2 | -0.375204291 | -0.254543259 | -0.612224773 | -0.289815804 |
| ENSDART00000129772 | gck | 2.162290839 | 3.39609934 | 3.505078852 | 1.699355805 |
| ENSDART00000129777 | lrp11 | -0.275801225 | -0.119366993 | -0.12158768 | 0.035259399 |
| ENSDART00000129819 | si:ch211-151p13.8 | -0.747854488 | -0.665734365 | -0.328239319 | -0.007548317 |
| ENSDART00000129834 | nat16 | -0.654571013 | -0.560511117 | -0.238004357 | -0.212203603 |
| ENSDART00000129866 | pcsk2 | -0.608807617 | -0.601188495 | -0.114458443 | 0.092104737 |
| ENSDART00000129895 | palm1b | 0.197850318 | 0.594118975 | 0.532496272 | 0.226704229 |
| ENSDART00000129898 | ctsf | -3.184979516 | -6.647256647 | -2.226327525 | 0.123813808 |
| ENSDART00000129903 | mfsd10 | 0.819604016 | 0.72451413 | 0.409438046 | 0.004032617 |
| ENSDART00000129910 | tmem240a | -0.628384792 | -0.334786051 | 0.214415995 | 0.163191044 |
| ENSDART00000129916 | tmem69 | -0.236861508 | -0.141159535 | -0.401818194 | -0.186294868 |
| ENSDART00000129919 | TIFA | 0.580552682 | 0.759337474 | 0.369383589 | -0.101109572 |
| ENSDART00000129924 | hmgb3a | -0.441161001 | -0.289637174 | 0.049371785 | 0.069315625 |
| ENSDART00000129963 | rbp1 | 1.30707657 | 0.844804596 | -0.023243783 | -0.809227723 |
| ENSDART00000129975 | scamp4 | 0.678060482 | 0.570045099 | 0.569688168 | 0.357822663 |
| ENSDART00000130007 | GABRA2 (1 of many) | -0.583670012 | -0.533882123 | -0.170964807 | 0.218833255 |
| ENSDART00000130093 | c10h21orf59 | -0.163395921 | -0.484168269 | -0.87971069 | -0.649928123 |
| ENSDART00000130103 | kctd16a | -0.864638484 | -0.862435538 | 0.056246994 | 0.316558654 |
| ENSDART00000130105 | psma3 | 0.177380698 | 0.356390892 | 0.223933012 | 0.001457342 |
| ENSDART00000130125 | gnao1a | -0.213616659 | -0.159180808 | 0.167107834 | 0.384555308 |
| ENSDART00000130131 | sult6b1 | 0.679202914 | 0.436031956 | 0.32900324 | -0.176606639 |
| ENSDART00000130142 | ube2w | 0.018587872 | -0.041621436 | -0.37919808 | -0.07627037 |
| ENSDART00000130171 | ncl | -0.573496849 | -3.495808243 | 0.185293047 | 0.359494588 |
| ENSDART00000130215 | TMEM179 (1 of many) | -0.219747105 | -0.315705451 | -0.194562459 | -0.061125473 |
| ENSDART00000130264 | rpz | -0.422979934 | -0.28718615 | -0.432905099 | -0.242626605 |
| ENSDART00000130273 | traf3ip2b | 0.459447337 | 1.01424616 | 0.933831804 | 0.205410709 |
| ENSDART00000130277 | rpl9 | 0.669628298 | 0.786807801 | 0.580702605 | 0.163431583 |
| ENSDART00000130297 | atg16l2 | 0.527142084 | 0.480414213 | 0.279348799 | 0.207971893 |
| ENSDART00000130310 | epb41l3a | -0.373985055 | -0.254132145 | 0.194766582 | 0.26124696 |
| ENSDART00000130317 | arhgap20 | -0.299393415 | -0.261693015 | -0.196202201 | -0.493979065 |
| ENSDART00000130343 | parp12a | 0.340672377 | 0.108363497 | 1.07497332 | -0.29758817 |
| ENSDART00000130353 | fdps | 0.408185548 | 0.643895342 | 0.766681528 | 0.202680939 |
| ENSDART00000130366 | si:dkey-21c1.4 | -0.116466944 | -0.239339578 | -0.392905857 | -0.311667635 |
| ENSDART00000130375 | rap1gap | -0.163733322 | -0.471249398 | -0.158019615 | -0.014828203 |
| ENSDART00000130388 | gfra3 | -0.144665825 | -0.360556971 | -0.224595855 | -0.144682167 |
| ENSDART00000130393 | tlr7 | -0.687566787 | -0.954352742 | -0.823581986 | -0.270839916 |
| ENSDART00000130397 | akt2 | 0.652260614 | -3.897305009 | -1.351826378 | -1.626684296 |
| ENSDART00000130413 | smc4 | 1.589552868 | 1.367681745 | 1.270952692 | 0.499256923 |
| ENSDART00000130453 | ahcyl2 | -0.228112194 | -0.250899226 | -0.220486764 | -0.265846948 |
| ENSDART00000130472 | pkhd1l1 | 0.896390148 | 0.750907757 | 1.185805195 | 0.60737198 |
| ENSDART00000130494 | moxd1 | 0.536918295 | 0.137560293 | -0.344113602 | -0.82692019 |
| ENSDART00000130537 | spag1a | -0.063362594 | -0.300759627 | -0.420912866 | -0.152669564 |
| ENSDART00000130546 | maptb | 0.390340472 | 1.128769701 | 1.393133544 | 1.028418073 |
| ENSDART00000130554 | isg15 | 0.469884875 | 0.09681696 | 3.101380991 | 0.025074196 |
| ENSDART00000130569 | st8sia5 | -0.025137728 | 0.545403948 | 0.761405173 | 0.616910757 |
| ENSDART00000130573 | atp6ap1la | -0.419066331 | -0.227150493 | -0.139216735 | 0.064701891 |
| ENSDART00000130594 | afap1 | 0.283339001 | 0.479700622 | 0.53264953 | 0.107820725 |
| ENSDART00000130601 | CABZ01020840.1 | 1.211268037 | 0.754343565 | 0.351699193 | -0.297623866 |
| ENSDART00000130604 | ssuh2.4 | 0.995807077 | 0.517001744 | 0.13232339 | -0.007643053 |

| | | | | | |
|---|---|---|---|---|---|
| ENSDART00000130623 | dus4l | 0.445376751 | 0.468587727 | 0.304661728 | 0.017366201 |
| ENSDART00000130626 | rars | 0.486608727 | 0.578942242 | 0.46060205 | 0.253410521 |
| ENSDART00000130632 | hif1al2 | 0.724834586 | 1.00924352 | 1.339085575 | 0.665259415 |
| ENSDART00000130648 | scarb2c | 0.530119641 | 0.364624401 | 0.255215167 | -0.022903408 |
| ENSDART00000130655 | csnk1a1 | -0.038868211 | -0.151998306 | -0.302247752 | -0.150662513 |
| ENSDART00000130675 | pi4k2a | 0.32642219 | 0.46314623 | 0.446740681 | 0.229275132 |
| ENSDART00000130692 | cry1aa | -0.303120324 | -0.112884884 | -0.294756028 | -0.171760667 |
| ENSDART00000130697 | runx1 | 0.52419199 | 0.991442797 | 1.34044624 | 0.449640925 |
| ENSDART00000130712 | pcdh2aa15 | -0.110623458 | -0.061292827 | 0.28564543 | 0.306014091 |
| ENSDART00000130722 | cacnb2b | -0.144539412 | -0.25934202 | -0.052732887 | 0.085080318 |
| ENSDART00000130741 | fam20cb | -0.286292768 | -0.244788679 | -0.108258226 | -0.149995213 |
| ENSDART00000130780 | clstn1 | -0.461252674 | -0.35014 | -0.010361021 | 0.286350898 |
| ENSDART00000130781 | eif4a2 | -0.090677214 | -0.087646154 | -0.280941023 | -0.172536563 |
| ENSDART00000130815 | e2f3 | 0.222958128 | 0.350714695 | 0.469146338 | 0.300980509 |
| ENSDART00000130818 | ptprz1b | -0.275892935 | -0.352115142 | -0.541325565 | -0.202924453 |
| ENSDART00000130820 | hbegfb | 1.26577708 | 2.42087461 | 1.934889729 | 0.952474054 |
| ENSDART00000130828 | cnrip1a | -0.043463049 | 0.089234147 | 0.284898375 | 0.266071808 |
| ENSDART00000130880 | si:ch73-269m14.2 | -0.24405591 | -0.090252599 | -0.043271301 | -0.190219309 |
| ENSDART00000130881 | gabrr2a | -0.366410023 | -0.492281459 | -0.409834 | -0.272101123 |
| ENSDART00000130891 | camk4 | -0.554565795 | -0.502410128 | -0.607057653 | -0.157488548 |
| ENSDART00000130911 | cmklr1 | 1.293061027 | 1.016453103 | 0.21085461 | 0.074156746 |
| ENSDART00000130977 | mtcl1 | -0.152016911 | -0.472594538 | -0.24239267 | -0.004366794 |
| ENSDART00000131010 | zgc:114175 | -0.532140281 | -0.400405891 | 0.129904669 | 0.109706695 |
| ENSDART00000131014 | sept7a | 0.353546424 | 0.539836173 | 0.262138094 | 0.154511745 |
| ENSDART00000131019 | acadvl | 0.465985818 | 0.491501242 | 0.281421395 | 0.167335906 |
| ENSDART00000131027 | TMEM216 | -0.171407049 | -0.12306379 | -0.494331831 | -0.331123076 |
| ENSDART00000131030 | gorasp1a | -0.03820825 | -0.156177406 | -0.398312611 | -0.044697109 |
| ENSDART00000131041 | doc2b | -0.197976996 | -0.14606408 | -0.273923734 | -0.068935148 |
| ENSDART00000131072 | rpz2 | 0.192971155 | 0.376625546 | 0.204804322 | 0.050600303 |
| ENSDART00000131075 | dusp23b | -0.349967426 | -0.590231643 | 0.026826234 | -0.132825162 |
| ENSDART00000131101 | alpl | -0.065559311 | -0.084665618 | -0.114820296 | -0.355758107 |
| ENSDART00000131110 | pigx | -0.041181166 | -0.247067458 | -0.407100449 | -0.233879595 |
| ENSDART00000131126 | bcl6a | -0.096649799 | -0.044696983 | 0.295011601 | 0.436349263 |
| ENSDART00000131134 | si:ch211-206a7.2 | -0.226522125 | -0.472315068 | -0.371394997 | -0.101330709 |
| ENSDART00000131137 | arhgef16 | 0.581874028 | 0.364359378 | 0.072001305 | -0.274180473 |
| ENSDART00000131143 | sept5b | -0.480326507 | -0.332562488 | 0.283118689 | 0.526750102 |
| ENSDART00000131161 | sema3c | -0.186853445 | 0.134603767 | 0.50487559 | 0.558561927 |
| ENSDART00000131177 | adcy1b | -0.090441849 | -0.360182616 | -0.355106619 | 0.00963355 |
| ENSDART00000131204 | asap1b | -0.546891821 | -0.455253903 | -0.26913051 | -0.092257648 |
| ENSDART00000131206 | march1 | -0.087431315 | -0.336892252 | -0.48450384 | -0.18585378 |
| ENSDART00000131217 | ippk | 0.31853123 | 0.426685761 | 0.198265799 | 0.012388457 |
| ENSDART00000131241 | CD53 | 1.613639378 | 1.480348694 | 1.150180939 | 0.900907437 |
| ENSDART00000131286 | erlin2 | -0.000462383 | -0.183644452 | -0.434826094 | -0.105981059 |
| ENSDART00000131288 | adap2 | 0.075508361 | 0.577403234 | 0.52972951 | 0.102379235 |
| ENSDART00000131319 | ctps1a | 0.387589066 | 0.225454863 | 0.161786406 | -0.016402726 |
| ENSDART00000131323 | si:dkey-193c22.1 | -0.375872197 | -0.187390024 | -0.399004983 | -0.194977152 |
| ENSDART00000131339 | capns1a | 0.087047749 | 0.358949459 | 0.212677567 | 0.044194443 |
| ENSDART00000131349 | mob2b | 0.337219743 | -0.092777728 | -0.774689204 | -0.397735432 |
| ENSDART00000131354 | si:ch211-235f12.2 | -0.417898946 | 0.007918255 | -0.299865948 | -0.220993476 |
| ENSDART00000131355 | cntn3a.1 | -0.54935923 | -0.424970108 | -0.640814852 | -0.044155876 |
| ENSDART00000131361 | kcnip3b | -0.353450144 | -1.035812483 | -0.692091767 | -0.071599751 |

| | | | | | |
|---|---|---|---|---|---|
| ENSDART00000131400 | CR792453.3 | 2.47650448 | 0.829837833 | -0.069994818 | -0.405416724 |
| ENSDART00000131431 | sulf2a | 0.379800676 | 0.895838573 | 1.99747824 | 2.182238849 |
| ENSDART00000131443 | dpm2 | -0.116130222 | -0.034113196 | -0.2501536 | -0.189683686 |
| ENSDART00000131477 | si:dkeyp-72g9.4 | -1.377600786 | -1.267958825 | -0.607564413 | -0.337098864 |
| ENSDART00000131478 | kcnab1a | -0.495628945 | -0.276495434 | -0.300290472 | -0.132892856 |
| ENSDART00000131506 | cdh6 | -0.336043184 | -0.313603925 | -0.315311457 | -0.136816058 |
| ENSDART00000131508 | col9a1b | -0.262030114 | 0.179891299 | -0.305486246 | -1.812812408 |
| ENSDART00000131517 | si:ch211-232m10.6 | -0.306074293 | -0.295884968 | -0.441790198 | -0.281067097 |
| ENSDART00000131557 | mvdb | 0.053284375 | 1.694163577 | 1.512604373 | 0.78016058 |
| ENSDART00000131558 | zgc:103438 | 0.587891191 | 0.925990315 | 0.240143957 | -0.080675365 |
| ENSDART00000131579 | ptprdb | -0.516674931 | -0.642051878 | 0.273217974 | 0.528486925 |
| ENSDART00000131582 | cbx3b | -0.219219 | -0.174464705 | -0.354832647 | -0.341719477 |
| ENSDART00000131610 | serbp1a | 0.184686975 | 0.28885198 | 0.237091318 | 0.011396501 |
| ENSDART00000131616 | atp5f1 | -0.440716151 | -0.423675229 | -0.427802237 | -0.218465008 |
| ENSDART00000131627 | kcnn3 | -0.085165392 | -0.23467025 | -0.387311243 | -0.108636046 |
| ENSDART00000131632 | ptk6a | -0.239390363 | 0.61942439 | 1.358814463 | 0.292212059 |
| ENSDART00000131649 | sulf2b | -0.232056185 | -0.450328943 | -0.252313155 | 0.036414893 |
| ENSDART00000131661 | rps9 | 0.491211514 | 0.443239408 | 0.342502062 | -0.078353052 |
| ENSDART00000131694 | selenbp1 | 0.440307718 | 0.23934875 | 0.219921774 | -0.162883702 |
| ENSDART00000131714 | mbnl2 | -0.406601788 | -0.032543807 | -0.091713267 | 0.092369986 |
| ENSDART00000131721 | fam65b | -0.06007994 | 0.061772028 | 0.37892142 | 0.335909594 |
| ENSDART00000131724 | pld3 | 0.801498613 | 0.654598157 | 0.377873753 | 0.229417497 |
| ENSDART00000131731 | mef2ca | 0.234535922 | -0.250149626 | -0.381098175 | -0.047609857 |
| ENSDART00000131736 | kcnq2a | -0.749729315 | -0.34933354 | 0.575885203 | 0.723193716 |
| ENSDART00000131768 | abi1a | 0.352423076 | 0.342648792 | 0.429769283 | 0.194498692 |
| ENSDART00000131802 | si:dkey-204f11.64 | 0.501807484 | 0.364460777 | 0.066146426 | -0.103454568 |
| ENSDART00000131846 | osbpl11 | 0.131220058 | 0.159712436 | 0.340514183 | 0.184103052 |
| ENSDART00000131848 | BX004774.2 | 0.065629354 | -0.435237015 | -1.040450944 | -0.818745042 |
| ENSDART00000131860 | rpl10 | 0.591996417 | 0.137912232 | 0.041524333 | -0.247459563 |
| ENSDART00000131861 | tmsb2 | 0.228394728 | 0.766531981 | 0.776266217 | 0.553413207 |
| ENSDART00000131866 | trhra | -0.77311942 | -0.482590954 | -0.518111078 | -0.348001173 |
| ENSDART00000131880 | ftr34 | 0.638978186 | 0.318226461 | 1.201576119 | 0.041863892 |
| ENSDART00000131922 | ucp3 | 0.966573752 | 1.664449524 | 1.416927563 | 1.012896527 |
| ENSDART00000131940 | si:dkey-30c15.10 | 1.656397572 | 1.745533662 | 1.503033451 | 0.676482739 |
| ENSDART00000131944 | tagln3b | 0.184610888 | 0.818287372 | 1.023242059 | 0.555937355 |
| ENSDART00000131954 | irf2a | -0.009751778 | -0.124292592 | -0.35526394 | -0.165453436 |
| ENSDART00000131956 | si:dkey-33c12.14 | 2.548697019 | 3.072973071 | 2.718694487 | 2.992696235 |
| ENSDART00000131966 | AL929520.1 | 1.864305398 | 1.102319083 | 0.530092783 | 0.456970098 |
| ENSDART00000131973 | lrmp | -0.070199137 | -0.469517154 | -1.098854311 | -0.136597848 |
| ENSDART00000131976 | adipor1a | -0.018834429 | -0.17416639 | -0.271837866 | -0.17995065 |
| ENSDART00000131983 | myhz1.2 | -0.851007348 | 2.593036141 | 3.254881434 | 2.63776379 |
| ENSDART00000131994 | mthfd1l | 2.796349329 | 3.017388421 | 3.058988321 | 0.924891143 |
| ENSDART00000132027 | spock2 | -0.61060119 | -0.435727551 | -0.163124474 | 0.204372495 |
| ENSDART00000132119 | max | 0.339991432 | 0.197083488 | 0.14057043 | 0.318686745 |
| ENSDART00000132123 | thsd7ba | -0.490674171 | -0.551076996 | -0.028899921 | -0.007032812 |
| ENSDART00000132146 | hsd17b7 | 2.880332402 | 3.535090877 | 2.510243514 | 2.078786507 |
| ENSDART00000132156 | cirbpb | 0.302166347 | 0.411233461 | 0.214244406 | 0.367929369 |
| ENSDART00000132158 | ptpn18 | 0.848975716 | 0.700592324 | 0.349323957 | -0.024370763 |
| ENSDART00000132170 | dlg1 | -0.324614076 | 0.23335851 | 0.829812481 | 1.031575976 |
| ENSDART00000132171 | CALHM1 (1 of many) | -0.125733999 | -0.221900772 | -0.83821175 | -0.699866376 |
| ENSDART00000132175 | nadl1.1 | 0.709405923 | 0.881560036 | 1.238414697 | 1.046443747 |

| | | | | | |
|---|---|---|---|---|---|
| ENSDART00000132189 | mogs | 0.278316469 | 0.095835936 | 0.119934101 | 0.001358604 |
| ENSDART00000132206 | si:dkey-242g16.2 | 0.572333551 | 0.964289734 | 1.608250495 | 0.992663233 |
| ENSDART00000132216 | paox1 | 0.706860126 | 0.311306251 | -0.007602467 | -0.279227462 |
| ENSDART00000132235 | klhl24b | 0.278493921 | 0.515923362 | 0.500638596 | 0.185998384 |
| ENSDART00000132278 | rnf144ab | -0.480742338 | -0.392838261 | -0.157627213 | -0.112649454 |
| ENSDART00000132294 | fut8a | -0.131220853 | -0.116575506 | 0.125377151 | 0.342318079 |
| ENSDART00000132296 | klhl17 | -0.721040801 | -0.498955013 | 0.192513289 | 0.489858233 |
| ENSDART00000132297 | adam9 | 0.312303854 | 0.492550355 | 0.565368245 | 0.224580764 |
| ENSDART00000132342 | kiaa1549lb | -0.130776123 | -0.485143208 | -0.118054479 | 0.086040071 |
| ENSDART00000132365 | b3gnt5a | 0.764122726 | 0.855145882 | 0.650888001 | 0.125992099 |
| ENSDART00000132366 | emp1 | -1.811345528 | -0.544421651 | -0.327114312 | 0.270404939 |
| ENSDART00000132367 | si:ch211-235f12.2 | 0.051407266 | -0.037706651 | -0.368292836 | -0.27129379 |
| ENSDART00000132384 | si:dkey-222b8.4 | -0.294618002 | -0.481620656 | -0.505454645 | -0.268964873 |
| ENSDART00000132386 | si:dkey-21e2.15 | 2.037905679 | 4.749110822 | 3.364621679 | 1.867440085 |
| ENSDART00000132387 | pvalb6 | -0.612280143 | -0.883743713 | -0.424014361 | -0.028285642 |
| ENSDART00000132390 | ankrd33ba | -0.590428215 | -0.156095959 | 0.097976154 | 0.018978411 |
| ENSDART00000132411 | si:ch1073-184j22.2 | -0.765110378 | -1.0433591 | -0.590491476 | -0.153408732 |
| ENSDART00000132426 | stmn4 | 0.67103814 | 1.650930846 | 1.637640616 | 1.211182541 |
| ENSDART00000132437 | mknk2a | 1.021696261 | 1.216138852 | 0.746540318 | 1.080690265 |
| ENSDART00000132471 | csmd2 | -0.459001483 | -0.911607393 | -0.371722747 | 0.062349627 |
| ENSDART00000132491 | wdr83os | -0.018771998 | -0.11821754 | -0.249438444 | -0.184919889 |
| ENSDART00000132522 | syt13 | -0.475976654 | -0.458152818 | -0.514134367 | -0.475986362 |
| ENSDART00000132542 | bdh2 | 0.716106023 | 0.289356437 | -0.133918448 | -0.268013639 |
| ENSDART00000132543 | klhdc8b | -0.423366405 | -0.574887213 | -0.139148992 | -0.034877401 |
| ENSDART00000132545 | slc25a24 | -0.397809038 | -0.303980258 | -0.481236788 | -0.168683615 |
| ENSDART00000132582 | map4l | -0.700979622 | -0.493129551 | -0.385069171 | -0.268022699 |
| ENSDART00000132583 | nrxn2a | -0.296541609 | -0.578930697 | -0.085512446 | 0.138469202 |
| ENSDART00000132591 | si:ch211-191i18.4 | 0.099974449 | -0.068457558 | -0.479286849 | -0.192940747 |
| ENSDART00000132605 | shc2 | 0.131499648 | 0.254257685 | 0.501707163 | 0.375914832 |
| ENSDART00000132637 | rab34a | 0.251888735 | 0.316357959 | 0.030371007 | -1.217256649 |
| ENSDART00000132658 | si:ch211-194p6.12 | 0.494361641 | 0.293067303 | 0.277540839 | 0.243465173 |
| ENSDART00000132659 | INAVA (1 of many) | -0.598820951 | -0.566244118 | -0.753512572 | -0.519207877 |
| ENSDART00000132660 | polr3gla | 0.823813849 | 0.325249077 | 0.007723106 | -0.29164484 |
| ENSDART00000132663 | si:ch211-163b2.4 | 0.347936584 | 0.995221531 | 0.769857122 | 0.372703166 |
| ENSDART00000132664 | hmbsa | 1.719683433 | 3.421951592 | 1.469413717 | 1.652766494 |
| ENSDART00000132667 | si:ch211-213a13.2 | 1.006426879 | 1.056394216 | 0.902670878 | 0.63305431 |
| ENSDART00000132685 | kcnn1b | -1.625205585 | -1.495755896 | -0.13719571 | 0.28532188 |
| ENSDART00000132691 | adgrb3 | -0.413275118 | -0.450913383 | -0.19602447 | 0.067293319 |
| ENSDART00000132696 | nckap1 | 0.516973126 | 0.759090684 | 0.845846182 | 0.627500278 |
| ENSDART00000132710 | pimr127 | -0.279310009 | -0.946546987 | -0.201892971 | -0.041573331 |
| ENSDART00000132732 | tmem86b | 0.107129941 | 0.17748496 | 0.221749507 | 0.409811812 |
| ENSDART00000132740 | ppp3cca | -0.073909682 | -0.313026697 | -0.310470542 | -0.074974696 |
| ENSDART00000132742 | apoc1 | 0.900470905 | 0.457797087 | -0.066389631 | 0.150028097 |
| ENSDART00000132744 | ppcdc | 0.019198695 | -0.177358944 | -0.471740583 | -0.31681582 |
| ENSDART00000132761 | arhgdig | -3.192123378 | 0.713824666 | 1.228568907 | 0.922752193 |
| ENSDART00000132767 | pxdc1a | -0.518773739 | -0.198850875 | -0.436310873 | -0.094674576 |
| ENSDART00000132781 | si:dkey-78l4.1 | 3.429067431 | 4.682405989 | 3.946041907 | 2.798505508 |
| ENSDART00000132788 | uap1 | 1.271261725 | 1.549537095 | 1.061485612 | 0.525207108 |
| ENSDART00000132827 | slc29a4 | -0.39428721 | -0.406514798 | -0.162001645 | -0.399843403 |
| ENSDART00000132829 | inadl | -0.343759067 | -0.387749769 | -0.321781319 | -0.246918999 |
| ENSDART00000132830 | stoml3b | 2.594772138 | 1.896323147 | 1.645470476 | 1.013599137 |

| | | | | | |
|---|---|---|---|---|---|
| ENSDART00000132838 | ank1a | -1.15020902 | -1.58246422 | -0.789622662 | -0.12782961 |
| ENSDART00000132884 | cxxc5a | -0.136575965 | 0.119118699 | 0.415415664 | 0.166435977 |
| ENSDART00000132911 | vwc2l | -0.573360674 | -0.839884971 | -0.564408735 | 0.020047377 |
| ENSDART00000132915 | serpine3 | -0.093377161 | -0.220423221 | -0.800809367 | -1.077921671 |
| ENSDART00000132930 | pcbp3 | -0.89548835 | -0.955377646 | -0.326630607 | 0.075971616 |
| ENSDART00000132941 | si:ch1073-287p18.1 | -0.223138733 | -0.188273 | -0.453602691 | -0.273864822 |
| ENSDART00000132953 | gsto2 | 2.493368999 | 2.571702083 | 1.494039414 | 1.444027796 |
| ENSDART00000132981 | si:dkey-126g1.9 | 0.817837135 | 2.40599851 | 2.663801596 | 3.155063025 |
| ENSDART00000132982 | leng9 | 3.031473639 | 2.780356944 | 2.241298703 | 1.634482288 |
| ENSDART00000132995 | ak3 | 0.087221034 | 0.050666468 | -0.189471235 | -0.418943401 |
| ENSDART00000133028 | mus81 | -0.28108498 | -0.562830421 | -0.307741383 | -0.072240912 |
| ENSDART00000133030 | psmd1 | 3.699340083 | 3.88833786 | 2.358920441 | 3.148062929 |
| ENSDART00000133035 | syt5a | -0.188232763 | -0.271065673 | -0.434073976 | -0.186984299 |
| ENSDART00000133036 | elavl3 | 0.009017018 | 0.132987196 | 0.394658714 | 0.30975228 |
| ENSDART00000133039 | tac3a | -0.2410584 | -0.489343566 | -0.63037654 | -0.413025711 |
| ENSDART00000133042 | lrriq1 | 1.127502736 | 0.596946573 | 0.146972775 | 0.204949423 |
| ENSDART00000133059 | cited4a | -0.002602779 | 0.055649701 | -0.293136781 | -0.030683128 |
| ENSDART00000133079 | c3a.5 | 1.938352782 | 2.924341422 | 2.291243352 | 0.500346416 |
| ENSDART00000133092 | st6galnac5a | -0.798487612 | -0.408716424 | -0.156113795 | 0.220161008 |
| ENSDART00000133099 | zgc:123105 | -0.457591599 | -0.29947281 | -0.397070368 | -0.177964487 |
| ENSDART00000133109 | rps28 | 0.456862734 | 0.459372965 | 0.307002895 | 0.056264372 |
| ENSDART00000133114 | bend5 | -0.067900889 | -0.281239203 | 0.001023992 | -0.054776456 |
| ENSDART00000133121 | bub1 | 1.470451257 | 1.232808742 | 1.262010591 | 0.843166609 |
| ENSDART00000133131 | uba1 | 0.147030579 | 0.239356262 | 0.529136799 | 0.293288779 |
| ENSDART00000133134 | efemp2b | -0.009444426 | 0.019713651 | -0.293460329 | -0.603842237 |
| ENSDART00000133143 | gpd1b | -0.326643231 | -0.124586783 | -0.280591098 | -0.372586222 |
| ENSDART00000133167 | si:dkey-98f17.5 | 0.605842267 | 0.54718235 | 0.319480313 | 0.041690788 |
| ENSDART00000133172 | palm1a | -0.485644015 | -0.226508351 | 0.746051517 | 0.735100737 |
| ENSDART00000133176 | si:dkey-171o17.8 | -0.335115287 | -0.40576097 | -0.392381982 | -0.176105305 |
| ENSDART00000133183 | parp8 | 0.666223987 | 0.351288157 | 0.707539986 | 0.438871753 |
| ENSDART00000133186 | ntng1a | -0.121155505 | -0.544453536 | -0.029519061 | 0.307964269 |
| ENSDART00000133194 | tbc1d1 | 2.782559818 | 3.186152585 | 2.750502559 | 1.318506947 |
| ENSDART00000133220 | CR788254.2 | -0.013907422 | -0.231082485 | -0.739605077 | -0.487191429 |
| ENSDART00000133223 | COQ10A | -0.381890626 | -0.12435669 | 0.152677483 | 0.108453147 |
| ENSDART00000133242 | syne2b | 0.14436825 | -0.456480635 | -0.844135801 | -0.206154738 |
| ENSDART00000133267 | grip2b | -0.249633664 | -0.394997267 | -0.432273427 | -0.409837225 |
| ENSDART00000133302 | pfkfb4a | 0.027105161 | -0.087436901 | -0.335656317 | -0.089798414 |
| ENSDART00000133304 | npm1a | 1.159038348 | 0.761096985 | 0.663684243 | 0.261874194 |
| ENSDART00000133325 | nell2b | -0.450880659 | -0.507896343 | -0.516937003 | -0.181021434 |
| ENSDART00000133330 | fam214b | -0.307417712 | -0.32820559 | -0.047756819 | 0.003374146 |
| ENSDART00000133348 | si:ch211-240g9.1 | -0.327991939 | -0.379613386 | -0.287166398 | -0.033274341 |
| ENSDART00000133354 | irx4b | -1.675681332 | -1.90655223 | 0.291488221 | 1.156903659 |
| ENSDART00000133375 | si:dkey-30c15.17 | 0.981163755 | 0.773615181 | 0.451221931 | -0.42308326 |
| ENSDART00000133384 | nr2f1a | -0.318698493 | -0.194424161 | -0.079355488 | -0.084390635 |
| ENSDART00000133385 | lrp1ba | -0.614935587 | -0.929322267 | -0.360685611 | 0.193952764 |
| ENSDART00000133404 | rps24 | 0.584478639 | 0.636503255 | 0.16891132 | -0.309462265 |
| ENSDART00000133419 | CR847953.1 | 0.226198142 | -0.018883626 | 0.274295718 | 0.480234099 |
| ENSDART00000133440 | prdm2a | -0.400160361 | -0.315344703 | -0.135326537 | 0.100769097 |
| ENSDART00000133449 | FO818659.1 | -0.258028224 | -0.214758594 | -0.076374423 | 0.03227851 |
| ENSDART00000133466 | CR847503.1 | -0.023677254 | 0.459363169 | 0.719903439 | 0.430626169 |
| ENSDART00000133473 | CU207343.1 | 0.284146912 | 1.104685019 | 1.557001573 | 0.994606199 |

| | | | | | |
|---|---|---|---|---|---|
| ENSDART00000133485 | BX004774.1 | 0.155994197 | -0.204649087 | -0.333501142 | -0.428721476 |
| ENSDART00000133487 | fosb | -0.882572207 | -1.273781011 | -0.772506712 | -0.914864674 |
| ENSDART00000133496 | CU467110.1 | -0.265746647 | -0.189978673 | -0.362219245 | -0.151144337 |
| ENSDART00000133497 | si:dkey-250k15.4 | 0.495799772 | 1.080770342 | 1.149092422 | 0.612727116 |
| ENSDART00000133504 | dab2ipb | -0.309610082 | -0.535236809 | -0.062134515 | 0.075905091 |
| ENSDART00000133512 | fosl1b | 2.958627889 | 2.652355797 | 1.322528527 | 0.313643003 |
| ENSDART00000133517 | si:dkey-228b2.5 | 0.6930874 | 0.410660128 | 0.386710557 | 0.222079502 |
| ENSDART00000133550 | fkbpl | 0.247078783 | 0.644847847 | 0.380481334 | 0.08075265 |
| ENSDART00000133583 | si:ch211-117m20.4 | -0.073043862 | -0.13648785 | -0.437013406 | -0.146443055 |
| ENSDART00000133587 | wsb2 | -0.218297067 | -0.234838526 | -0.321740633 | -0.105339309 |
| ENSDART00000133613 | AL954694.1 | -0.189304139 | -0.336191594 | -0.52116431 | -0.341767633 |
| ENSDART00000133628 | ankib1a | 0.478883806 | 0.425404685 | 0.598914996 | 0.306729273 |
| ENSDART00000133644 | magi2a | -0.426844627 | -0.586214992 | -0.344780913 | 0.065236587 |
| ENSDART00000133659 | ncapg | 0.759144886 | 0.927669912 | 0.449671959 | 0.278225274 |
| ENSDART00000133661 | tasp1 | -0.129454637 | -0.177592146 | -0.299309238 | -0.129141605 |
| ENSDART00000133699 | drd4b | -0.558425031 | 0.210961997 | -0.355610364 | -0.126077055 |
| ENSDART00000133701 | slc6a1a | -0.417718543 | -0.444992936 | -0.335539494 | -0.221449117 |
| ENSDART00000133707 | st3gal3a | -0.191118701 | -0.289597986 | -0.435587316 | -0.218434667 |
| ENSDART00000133708 | arhgef4 | 0.196005293 | 0.61534617 | 0.621912538 | 0.36146972 |
| ENSDART00000133718 | ddx5 | 0.680973492 | 0.739800314 | 0.555100275 | 0.545762532 |
| ENSDART00000133721 | si:ch211-156j22.4 | 0.531762631 | 0.492062657 | 0.618453608 | 0.322450987 |
| ENSDART00000133735 | cacng7b | -0.52911361 | -0.351880161 | -0.311790544 | -0.261905914 |
| ENSDART00000133743 | camkk1a | -0.520213698 | -0.784879077 | -0.322164805 | -0.513359014 |
| ENSDART00000133775 | pdlim3b | 2.202802348 | 2.787120555 | 2.295385553 | 1.784176005 |
| ENSDART00000133786 | si:ch211-196l7.4 | -0.265117484 | -0.278594168 | -0.287454952 | -0.249634822 |
| ENSDART00000133802 | inpp5jb | -0.26851081 | -0.522364408 | -0.244805361 | -0.012462884 |
| ENSDART00000133823 | arvcfb | 0.059276527 | -0.645309269 | -0.357027269 | -0.368113028 |
| ENSDART00000133851 | nyap2b | -0.344969094 | 0.36816036 | 0.857787268 | 0.825724613 |
| ENSDART00000133867 | arfip2a | -0.237325336 | -0.320204428 | -0.424850477 | -0.153128941 |
| ENSDART00000133869 | kcnk1a | -0.389144165 | -0.38978007 | -0.667699511 | -0.286776021 |
| ENSDART00000133891 | FP015789.1 | -0.349747936 | -0.468679484 | -0.676764109 | -0.710268862 |
| ENSDART00000133902 | clstn1 | -0.274744137 | -0.313184588 | -0.391319406 | -0.194558837 |
| ENSDART00000133917 | lrrtm1 | -0.267345102 | -0.23914052 | -0.165843211 | -0.135383045 |
| ENSDART00000133922 | atp2b3a | -0.330787062 | -0.407621044 | -0.300317219 | -0.080866095 |
| ENSDART00000133934 | rps9 | 0.75844303 | 0.709617891 | 0.420926612 | 0.393006212 |
| ENSDART00000133946 | si:dkey-147f3.8 | 1.477973714 | 1.681506394 | 1.170700784 | -0.068341199 |
| ENSDART00000133954 | mhc1uka | 4.819339023 | 5.453978456 | 5.536236268 | 3.849327613 |
| ENSDART00000133956 | phactr3a | -0.466148377 | -0.040906797 | 0.400008422 | 0.572153181 |
| ENSDART00000133968 | camkmt | 0.353110068 | 0.455754701 | 0.544139309 | 0.380701512 |
| ENSDART00000133981 | bco2a | -0.093189558 | -0.108753759 | -0.2707511 | -0.08847471 |
| ENSDART00000133985 | kcnd1 | -1.187342293 | -1.392518344 | -0.747443213 | -0.002522665 |
| ENSDART00000134035 | cntnap5b | -1.01200416 | -1.154440105 | -0.403471674 | 0.139198923 |
| ENSDART00000134044 | vdac3 | -0.155270833 | 0.146773494 | 0.378912201 | 0.402649801 |
| ENSDART00000134047 | cyp2p10 | -0.641341489 | -0.436063229 | -0.777629241 | -0.630838584 |
| ENSDART00000134052 | si:dkey-17e16.8 | 0.112444287 | 0.152961651 | 0.336755528 | 0.189687147 |
| ENSDART00000134053 | usp2a | -0.375529791 | -0.273036858 | -0.320691469 | -0.159380591 |
| ENSDART00000134054 | si:dkey-13i19.8 | 0.097967988 | -0.010643615 | -0.259041425 | -0.27695026 |
| ENSDART00000134064 | mxd1 | -0.125104323 | -0.265030232 | -0.047451672 | 0.092856449 |
| ENSDART00000134083 | arhgap21b | -0.613534268 | -0.758748481 | -1.557344733 | -0.550383994 |
| ENSDART00000134087 | col14a1a | -0.156423731 | -0.247987955 | -0.214368462 | -0.368307818 |
| ENSDART00000134094 | zgc:162193 | 0.951212203 | 1.108680472 | 1.240038838 | 0.640044887 |

| | | | | | |
|---|---|---|---|---|---|
| ENSDART00000134128 | impg1a | -0.245363296 | -0.207318324 | -0.487433197 | -0.127440305 |
| ENSDART00000134146 | mtss1 | 0.331132861 | 0.244842152 | 0.229292017 | -0.102446838 |
| ENSDART00000134186 | cplx4b | -0.392479495 | -0.543085477 | -0.51593407 | -0.267696813 |
| ENSDART00000134189 | npm1a | 1.523433306 | 1.012610127 | 1.305980875 | 0.439289484 |
| ENSDART00000134190 | sort1b | -0.233086943 | -0.116629656 | -0.586941063 | -0.191255985 |
| ENSDART00000134202 | ccka | -0.373763887 | -0.443030746 | -0.635815918 | -0.444990754 |
| ENSDART00000134206 | hspb8 | 0.868767381 | 0.59923642 | 0.380053431 | 0.101288698 |
| ENSDART00000134241 | olfm1a | -1.215716601 | -1.195013518 | 0.048027409 | 0.32030838 |
| ENSDART00000134242 | zfand5a | 0.011715993 | -0.114565058 | -0.599260538 | -0.310476512 |
| ENSDART00000134257 | znf1027 | 2.316081338 | 1.234819597 | 2.431130405 | 1.907826667 |
| ENSDART00000134261 | dnm1b | -0.433650537 | -0.451510325 | 0.138870574 | 0.465815695 |
| ENSDART00000134269 | fkbp1aa | 0.162085973 | 0.545601006 | 0.44355879 | 0.374542557 |
| ENSDART00000134307 | dclk1a | -1.448196855 | -0.41548572 | -0.010990365 | -0.586401585 |
| ENSDART00000134393 | arhgef9a | -0.866173468 | -1.122671917 | -0.27973916 | -0.154746662 |
| ENSDART00000134399 | slkb | -0.338857379 | -0.632619838 | -0.770465733 | -0.089077643 |
| ENSDART00000134404 | si:ch211-158d24.4 | 0.479480538 | 0.856462378 | 0.502566478 | -0.11398241 |
| ENSDART00000134431 | pcbp3 | -0.497499314 | -0.525712825 | -0.323624866 | -0.140513929 |
| ENSDART00000134455 | zgc:100906 | -0.319125396 | 0.101796228 | 0.299790493 | 0.225508495 |
| ENSDART00000134470 | fhdc1 | 0.136576688 | 0.100413162 | -0.280286506 | -0.662790087 |
| ENSDART00000134472 | taok3b | -0.264981976 | -0.491201835 | -0.021652579 | 0.079259902 |
| ENSDART00000134475 | gdap1 | -0.459369959 | -0.408869462 | -0.283212336 | -0.008346648 |
| ENSDART00000134489 | htr2cl1 | -0.503209957 | -0.409297224 | -0.43743207 | -0.200430987 |
| ENSDART00000134506 | si:dkey-196j8.2 | -0.894031142 | -1.351407426 | -0.577940716 | -0.262796642 |
| ENSDART00000134514 | smarca4a | 0.020398578 | 0.110031025 | 0.339984244 | 0.255390677 |
| ENSDART00000134518 | cpa6 | -0.382405278 | -0.418058887 | -0.443147645 | -0.210306107 |
| ENSDART00000134542 | csde1 | 0.186004618 | 0.310572638 | 0.257821474 | 0.010065768 |
| ENSDART00000134560 | slc45a1 | -0.495253985 | -0.301829281 | 0.42188175 | 0.485995526 |
| ENSDART00000134564 | srsf5b | -0.27445076 | -0.31745813 | -0.113899297 | -0.080138136 |
| ENSDART00000134571 | rpl13a | 0.370507733 | 0.272503899 | 0.102928018 | -0.095604932 |
| ENSDART00000134602 | gria2b | -0.15956996 | -0.75241939 | -0.264702392 | 0.253919663 |
| ENSDART00000134608 | phyhiplb | -0.442372787 | -0.312061243 | -0.158725579 | -0.176785812 |
| ENSDART00000134635 | si:dkey-9i23.15 | 0.362598675 | 0.484654675 | 0.418405681 | -0.053498182 |
| ENSDART00000134649 | CNGA1 (1 of many) | -0.31835112 | -0.262022337 | -0.573519962 | -0.315754252 |
| ENSDART00000134658 | slc44a1a | 0.707497042 | 0.591443803 | 0.629657706 | 0.651062281 |
| ENSDART00000134661 | gstcd | -0.989748846 | -0.61234101 | -0.452964111 | -0.446444607 |
| ENSDART00000134697 | pik3ip1 | -0.252912627 | -0.195771257 | -0.573002676 | -0.212029641 |
| ENSDART00000134715 | elmod1 | -1.115600356 | -1.115324879 | -0.105456948 | 0.485174818 |
| ENSDART00000134717 | tspan18a | 0.042661269 | 0.285439514 | 0.614193732 | 0.731823089 |
| ENSDART00000134722 | bcl2l13 | -0.594192147 | -0.544774096 | -0.184876076 | -0.124896832 |
| ENSDART00000134727 | TNNC2 (1 of many) | -0.059692967 | 2.582271594 | 3.518368083 | 2.376076569 |
| ENSDART00000134729 | BX936308.1 | -0.246065702 | 0.716060471 | 1.529870623 | 1.562652796 |
| ENSDART00000134763 | zgc:112056 | 0.091717396 | -0.08997535 | -0.362094913 | -0.168303392 |
| ENSDART00000134773 | zfand5a | -0.170709233 | -0.221425066 | -0.980962271 | -0.517481995 |
| ENSDART00000134787 | elavl4 | 1.446773831 | 1.526430349 | 1.657041106 | 1.070021064 |
| ENSDART00000134801 | ccdc92 | 0.127931369 | 0.677319918 | 0.797788944 | 0.697201329 |
| ENSDART00000134813 | rx1 | -1.006947632 | -0.71282108 | -0.035273805 | -0.106549447 |
| ENSDART00000134816 | slc2a3a | -0.406786727 | -0.444029943 | -0.052958142 | 0.129218418 |
| ENSDART00000134819 | slc5a6a | 0.44313334 | -0.037657655 | -0.159985236 | -0.577627715 |
| ENSDART00000134832 | rbpms2b | -1.576691045 | -2.099790887 | -0.701820765 | 0.187979586 |
| ENSDART00000134843 | sst6 | 1.638903894 | 2.735989277 | 2.458183626 | 2.301951228 |
| ENSDART00000134855 | efr3bb | -0.263384976 | -0.381748192 | -0.1453091 | 0.002074194 |

| | | | | | |
|---|---|---|---|---|---|
| ENSDART00000134864 | gigyf1b | -0.032906085 | -0.141419247 | -0.36943011 | -0.170431533 |
| ENSDART00000134868 | zgc:113314 | 1.966628384 | 1.545280953 | 2.058380505 | 1.451511877 |
| ENSDART00000134870 | si:ch1073-170o4.1 | 0.916491768 | 1.072144968 | 0.377684395 | 0.341437061 |
| ENSDART00000134890 | dusp11 | 0.272279441 | 0.221032047 | 0.049148172 | 0.030930206 |
| ENSDART00000134922 | kctd7 | -0.261930929 | -0.25302271 | -0.255500491 | -0.12435787 |
| ENSDART00000134924 | actb2 | 0.443688406 | 0.512853066 | 0.501918371 | 0.442049165 |
| ENSDART00000134934 | pag1 | -0.161885621 | -0.157851391 | -0.203479947 | -0.293661261 |
| ENSDART00000134941 | prc1b | 1.557351387 | 1.026365943 | 0.57100162 | 0.47167284 |
| ENSDART00000134969 | rnd2 | -0.341329865 | -0.337414256 | -0.483308665 | -0.257319117 |
| ENSDART00000134973 | pcnxl2 | -0.772294262 | -0.54715302 | -0.00593982 | 0.313205165 |
| ENSDART00000135029 | BX927398.3 | 0.541935951 | 0.347132897 | 0.430074452 | 0.28825753 |
| ENSDART00000135032 | si:dkey-57a22.11 | 1.01281435 | 1.486077838 | 0.644934957 | 0.560278535 |
| ENSDART00000135052 | slc8a4a | -0.449869312 | -0.676457859 | -0.284767037 | 0.064060964 |
| ENSDART00000135076 | scrn3 | 0.148271997 | -2.654959907 | 1.093269557 | 0.033366542 |
| ENSDART00000135093 | ola1 | 0.415909116 | 0.691871408 | 0.432580712 | -0.033470782 |
| ENSDART00000135118 | efr3ba | -0.168363836 | -0.303586815 | -0.356914459 | -0.264543553 |
| ENSDART00000135125 | serp2 | 4.40213E-05 | 0.384051284 | 0.40863701 | 0.667539121 |
| ENSDART00000135133 | FRMD5 (1 of many) | -0.238198919 | -0.442975354 | -0.623723948 | -0.410224766 |
| ENSDART00000135147 | si:ch211-218d20.15 | -0.111515958 | 0.398515484 | 0.749278715 | 0.887643411 |
| ENSDART00000135164 | pdcd6ip | 0.194711079 | 0.272768586 | 0.208605784 | 0.054641806 |
| ENSDART00000135184 | pafah1b3 | 0.785314803 | 0.490699437 | 0.476757912 | 0.524529338 |
| ENSDART00000135206 | ppp6r2b | -0.281142732 | -0.364406934 | -0.331381396 | -0.106942863 |
| ENSDART00000135230 | chd7 | 0.169349437 | 0.346809261 | 0.458651128 | 0.318074781 |
| ENSDART00000135238 | dok2 | 1.480525707 | 0.841647962 | 0.697333529 | 0.138703296 |
| ENSDART00000135248 | anxa13l | 2.875521493 | 3.590436866 | 3.752911284 | 3.4448321 |
| ENSDART00000135256 | ubtd2 | 0.043462174 | 0.213810574 | 0.405778219 | 0.255726061 |
| ENSDART00000135284 | vsnl1a | -0.802778535 | -0.782046001 | -0.520311896 | -0.04073792 |
| ENSDART00000135285 | clk2a | 0.251961136 | 0.113407524 | 0.334958884 | 0.51104172 |
| ENSDART00000135337 | zgc:64189 | 0.108656283 | -0.42063095 | -0.437081395 | -0.295412247 |
| ENSDART00000135356 | nabp1a | 0.371235634 | 0.392313666 | 0.26759026 | 0.229305948 |
| ENSDART00000135357 | si:ch211-220f16.2 | 3.87916704 | 3.441079612 | 2.990059648 | 3.744387905 |
| ENSDART00000135374 | ptpn21 | -0.060719081 | -0.163907746 | -0.321158981 | -0.096688483 |
| ENSDART00000135381 | six4a | 2.521510715 | 2.3671408 | 2.587075649 | 1.691172374 |
| ENSDART00000135399 | phf11 | -0.105908519 | 0.331215825 | 2.01559672 | 0.351931859 |
| ENSDART00000135403 | nuak2 | -0.203159013 | -0.012266437 | -0.34684763 | -0.520334493 |
| ENSDART00000135415 | dysf | 3.146286589 | 3.204629318 | 3.124686608 | 3.126558821 |
| ENSDART00000135431 | rabgap1l | 1.914692926 | 2.028305841 | 2.310641205 | 1.631726628 |
| ENSDART00000135436 | jtb | -0.083914092 | -0.13827795 | -0.410566521 | -0.208466854 |
| ENSDART00000135472 | zgc:86764 | 1.170468948 | 0.97326651 | 0.792803713 | 0.492808604 |
| ENSDART00000135475 | cadm2a | -0.589549243 | -0.825400299 | -0.33568035 | 0.065563209 |
| ENSDART00000135479 | fstl5 | -0.187703631 | -0.312045142 | -0.413295205 | -0.11029682 |
| ENSDART00000135481 | si:ch211-67n3.9 | -0.264301646 | -0.482759797 | -0.356089399 | -0.148916978 |
| ENSDART00000135510 | syne2b | -0.011968335 | -0.47360345 | -0.918119469 | -0.289310189 |
| ENSDART00000135513 | rgs6 | -0.368608418 | -0.140264684 | -0.010741291 | 0.084818744 |
| ENSDART00000135517 | cenpf | 1.006357007 | 0.930294184 | 0.598168756 | 0.318387864 |
| ENSDART00000135556 | lonrf1 | -0.330548829 | 0.12250287 | -0.445965292 | -0.332349033 |
| ENSDART00000135569 | gale | 0.396996119 | 0.386168327 | 0.425741481 | 0.058303999 |
| ENSDART00000135583 | mlpha | 2.247424425 | 4.049714869 | 4.15888721 | 1.709523081 |
| ENSDART00000135587 | cd9a | -0.312163633 | -0.264335113 | 0.189621918 | 0.343999271 |
| ENSDART00000135595 | mhc1zaa | 0.19156418 | 1.036766333 | 0.760670357 | 0.331702215 |
| ENSDART00000135599 | lypla2 | -0.359594871 | -0.446096937 | -0.182980597 | -0.086611879 |

| | | | | | |
|---|---|---|---|---|---|
| ENSDART00000135602 | si:ch73-290k24.6 | -0.456455854 | -0.524968049 | -0.128888416 | -0.002715321 |
| ENSDART00000135606 | TMEM269 | -0.06366918 | -0.165749005 | -0.266297774 | -0.18037018 |
| ENSDART00000135608 | gabrp | -2.860562828 | -1.443836066 | -0.572564003 | -0.537301716 |
| ENSDART00000135624 | FAM163A | -0.714036121 | -0.427639458 | -0.142310427 | 0.163241634 |
| ENSDART00000135631 | myl6 | 0.462877699 | 0.806124657 | 0.938068157 | 0.529055927 |
| ENSDART00000135661 | slc20a1a | 1.716116201 | 1.644628019 | 1.747440585 | 0.991048186 |
| ENSDART00000135690 | lactbl1a | -0.207283355 | -0.397072585 | -0.513557165 | -0.348206521 |
| ENSDART00000135692 | UNC13A | -0.209420413 | -0.513653337 | -0.217224189 | 0.093101705 |
| ENSDART00000135702 | rgs11 | -0.576481143 | -0.793334647 | -0.383725367 | 0.048951277 |
| ENSDART00000135715 | fam184a | -0.344643213 | -0.481218539 | -0.282456424 | 0.057932244 |
| ENSDART00000135719 | si:ch211-171h4.6 | 1.187005549 | 0.373648016 | 0.271137881 | -0.125827965 |
| ENSDART00000135730 | lrmp | -0.028204227 | -0.338623204 | -0.590263694 | -0.109841967 |
| ENSDART00000135738 | zgc:194990 | 0.067005172 | -0.387370648 | -0.731552205 | -0.192679748 |
| ENSDART00000135756 | kirrel3a | -0.197416496 | -0.816919597 | -0.348887933 | -0.004164348 |
| ENSDART00000135764 | si:dkey-70p6.1 | -0.54777588 | -0.724047584 | 0.062614493 | 0.292769909 |
| ENSDART00000135768 | lrp1ba | -0.685782746 | -0.945707299 | -0.285013804 | 0.195853428 |
| ENSDART00000135781 | plcg1 | 0.270370473 | 0.251405726 | 0.297242892 | 0.117629016 |
| ENSDART00000135794 | sepn1 | 0.371752242 | 0.285638022 | 0.016731978 | -0.094169263 |
| ENSDART00000135811 | atp2b3a | -0.361290503 | -0.436904447 | -0.166639435 | 0.060424957 |
| ENSDART00000135821 | mfsd6b | 0.610408738 | 1.551908888 | 1.729395905 | 1.289342978 |
| ENSDART00000135860 | rnf41l | -0.353446179 | -0.473576771 | -0.616364395 | -0.289592085 |
| ENSDART00000135888 | rrp9 | 0.649352382 | 0.396806051 | 0.419459571 | 0.067410678 |
| ENSDART00000135897 | tmeff2a | -0.507501095 | -0.570530323 | -0.389897465 | -0.071849112 |
| ENSDART00000135919 | amd1 | -0.131714213 | -0.276160556 | -0.537901531 | -0.163925689 |
| ENSDART00000135965 | plch1 | -0.574356968 | -0.567600235 | -0.421755163 | -0.154028312 |
| ENSDART00000135973 | nfyal | 0.202128253 | 0.401502491 | 0.256237772 | 0.346886156 |
| ENSDART00000135989 | cep170aa | 0.062401976 | 0.336049788 | 0.4713762 | 0.353973795 |
| ENSDART00000136028 | CR753844.1 | 0.308802389 | 0.230251216 | 0.146560512 | 0.144442899 |
| ENSDART00000136049 | syt10 | -0.382437751 | -0.322146795 | -0.465356993 | -0.383104059 |
| ENSDART00000136084 | mfhas1 | -0.229093837 | -0.123648609 | -0.428888075 | -0.346098599 |
| ENSDART00000136091 | c1ql2 | -0.439731401 | -0.550972938 | -0.293500773 | -0.243598849 |
| ENSDART00000136103 | ubap2a | 0.173955465 | 0.276885256 | 0.421817829 | 0.278699527 |
| ENSDART00000136123 | kdm6bb | -0.32427763 | -0.244056677 | -0.170049978 | -0.028150527 |
| ENSDART00000136128 | FP016056.1 | -0.005351848 | -0.270043208 | -0.447280202 | -0.270195958 |
| ENSDART00000136141 | kcnq3 | -0.157652998 | 0.519381195 | 1.385651499 | 1.123132968 |
| ENSDART00000136152 | calua | 0.347598327 | 0.562237059 | 0.505429013 | 0.302262295 |
| ENSDART00000136154 | gpr158a | -0.522434846 | -0.680761714 | -0.064607745 | -0.081447185 |
| ENSDART00000136161 | faxcb | -0.355434657 | -0.26288196 | 0.155305377 | 0.179364353 |
| ENSDART00000136182 | si:ch211-222n4.2 | 2.334818577 | 2.498262776 | 2.056155354 | 0.938824321 |
| ENSDART00000136187 | bloc1s4 | 0.27836522 | 0.479128543 | 0.397988078 | 0.159673351 |
| ENSDART00000136190 | si:ch211-133l5.7 | 0.046455816 | -0.284643729 | -0.455348095 | -0.512893839 |
| ENSDART00000136200 | cbln2b | -0.48916567 | -0.352468034 | -0.442788192 | -0.14816736 |
| ENSDART00000136208 | ncam1b | -0.292790209 | -0.308372802 | -0.421218735 | -0.19466435 |
| ENSDART00000136213 | bzw2 | 0.202381787 | -0.263816931 | -0.545867534 | -0.324585711 |
| ENSDART00000136233 | TENM3 | -0.571502608 | -0.593065001 | 0.242250375 | 0.436842415 |
| ENSDART00000136239 | kcnv2a | -0.144268598 | -0.378520976 | -0.127025179 | 0.043873086 |
| ENSDART00000136255 | otofa | -0.67545742 | -0.815938232 | -0.602392654 | -0.103624276 |
| ENSDART00000136267 | slc35f3b | -0.474511888 | -0.87524613 | 0.142496146 | 0.506913479 |
| ENSDART00000136269 | rimbp2 | -0.196271668 | -0.410552217 | -0.160743106 | 0.064316711 |
| ENSDART00000136296 | CR847543.2 | 0.7189738 | 0.800339311 | 1.432214548 | 0.183639146 |
| ENSDART00000136314 | cthl | 0.026768092 | 0.498239754 | -0.095502024 | -0.417996724 |

| | | | | | |
|---|---|---|---|---|---|
| ENSDART00000136377 | si:dkey-239h2.3 | -0.060598158 | -0.178560319 | -0.404445305 | -0.188298872 |
| ENSDART00000136378 | mb | 0.060967695 | -0.006783267 | -0.887462271 | -1.204052153 |
| ENSDART00000136382 | gadd45ba | 1.037707985 | 1.186135766 | 0.645465362 | 0.508629 |
| ENSDART00000136383 | cd99 | 0.67248236 | 1.035021667 | 0.666974643 | -0.164650894 |
| ENSDART00000136386 | si:ch211-210c8.7 | 0.436864275 | 0.595310791 | 0.386350533 | 0.024375719 |
| ENSDART00000136390 | hcn2b | -0.288020893 | -0.351077892 | -0.312560003 | -0.049583355 |
| ENSDART00000136418 | sf3a1 | 0.245658224 | 0.479306215 | 0.291480298 | 0.207121133 |
| ENSDART00000136439 | bcl7a | 0.000990732 | 0.374886928 | 0.452965996 | 0.230358497 |
| ENSDART00000136447 | si:dkey-161j23.7 | -0.523562694 | -0.013224352 | -0.081189145 | 0.161013961 |
| ENSDART00000136472 | trim3b | -0.264051102 | -0.238034524 | -0.31432798 | -0.179272082 |
| ENSDART00000136488 | zmat4a | -0.690962776 | -0.637104526 | -0.28999313 | -0.088561559 |
| ENSDART00000136492 | si:ch211-158d24.4 | 0.620322497 | 0.916930916 | 0.610223614 | 0.170743477 |
| ENSDART00000136497 | CR318624.1 | -0.290569983 | -0.906819744 | -0.441990313 | 0.029701559 |
| ENSDART00000136523 | CU467861.1 | 1.460403168 | 1.124457561 | 1.294733731 | 1.217332143 |
| ENSDART00000136538 | cish | -0.292880454 | -0.45612089 | -0.971812918 | -1.16494411 |
| ENSDART00000136543 | tnni4b.1 | 1.504503498 | 0.982879899 | 0.857598266 | -0.145526765 |
| ENSDART00000136547 | rpl10a | 0.572905103 | 0.588248568 | 0.30592141 | 0.14058962 |
| ENSDART00000136559 | ano8a | -0.477178876 | -0.572871399 | -0.340924619 | -0.093243478 |
| ENSDART00000136578 | syn1 | -0.141221338 | -0.191909782 | 0.363564902 | 0.481540125 |
| ENSDART00000136592 | ckbb | -0.985379774 | -1.268435214 | -0.989267529 | -0.749991465 |
| ENSDART00000136647 | ext1c | -0.283149695 | -0.275883116 | -0.465218527 | -0.113484828 |
| ENSDART00000136649 | si:dkey-92i17.2 | 0.27365425 | 1.100188551 | 1.105138404 | 0.404606112 |
| ENSDART00000136653 | ccdc106b | 0.176566138 | -0.096095881 | -1.050174868 | -0.31850308 |
| ENSDART00000136654 | grapb | -0.34841027 | -0.721289328 | -0.501261917 | -0.353877694 |
| ENSDART00000136655 | ndufa3 | -0.198867993 | -0.196758365 | -0.355261116 | -0.136756127 |
| ENSDART00000136692 | ccnl1a | 0.306138381 | 0.080789654 | -0.032909692 | 0.080389121 |
| ENSDART00000136695 | yrk | 1.103276689 | 0.393392175 | 0.380351314 | -0.354474005 |
| ENSDART00000136700 | hunk | 1.085007384 | 0.795931304 | 0.468549793 | 1.046395882 |
| ENSDART00000136707 | mical1 | -0.443299702 | -0.458390885 | 0.146469401 | 0.374257303 |
| ENSDART00000136708 | rbpms2b | -1.024689498 | -0.80090545 | -0.103078191 | 0.159293341 |
| ENSDART00000136722 | cdkn1a | 1.083504495 | 0.97819224 | 0.258587523 | 0.060123909 |
| ENSDART00000136726 | magi3b | -0.381646177 | -0.461208385 | -0.226303301 | -0.080158554 |
| ENSDART00000136729 | ebf1b | -2.79991744 | -3.068779235 | -0.720206417 | 0.313112936 |
| ENSDART00000136733 | glra1 | -0.259893278 | -0.432961012 | -0.409765299 | -0.248913141 |
| ENSDART00000136744 | foxq2 | -0.278235678 | -0.492834499 | -0.366505779 | -0.190958275 |
| ENSDART00000136754 | cdh13 | -0.587587145 | -0.627840519 | -0.172128457 | 0.142910694 |
| ENSDART00000136759 | prdm13 | -0.225066145 | -0.413438688 | -0.129520305 | 0.036146962 |
| ENSDART00000136761 | prkx | 0.698491961 | 1.03323314 | 0.943847516 | 0.577218336 |
| ENSDART00000136770 | si:dkey-246e1.3 | -0.264481758 | -0.308595158 | -0.554881445 | -0.406535962 |
| ENSDART00000136771 | dnajc5ga | -0.246041931 | -0.267357646 | -0.352933326 | -0.221144382 |
| ENSDART00000136779 | entpd5b | 0.819562392 | 0.580844217 | 0.174817773 | -0.04896424 |
| ENSDART00000136782 | si:dkey-24c2.7 | 4.902793471 | 4.654641885 | 4.451812012 | 3.610899339 |
| ENSDART00000136789 | tjp2a | -0.247782376 | -0.511487885 | -0.433268095 | -0.195886379 |
| ENSDART00000136797 | btbd10a | -0.201172343 | -0.244946837 | 0.724827561 | 0.837222426 |
| ENSDART00000136805 | rbm10 | -4.539752918 | -3.15582054 | -1.19980751 | -2.046490722 |
| ENSDART00000136830 | stmn4l | 1.323269884 | 1.682347243 | 1.639725976 | 1.096832109 |
| ENSDART00000136835 | gpr84 | 0.70622544 | 0.183343809 | -0.153052934 | -0.427967202 |
| ENSDART00000136836 | rab11fip3 | -0.285287519 | -0.130906101 | 0.115401989 | 0.056439488 |
| ENSDART00000136853 | fam163b | -0.426911899 | -0.715978676 | -0.456698496 | -0.087825906 |
| ENSDART00000136865 | cfp | 0.329536618 | 0.225136408 | -0.193120136 | -0.891420593 |
| ENSDART00000136873 | clasp2 | 0.301439566 | 0.245236713 | 0.309865242 | 0.356365378 |

| | | | | | |
|---|---|---|---|---|---|
| ENSDART00000136910 | armc1l | 0.626427947 | 0.775572599 | 0.634581641 | 0.292392529 |
| ENSDART00000136914 | tpd52l2a | -0.012405325 | -0.254912754 | -0.378255822 | -0.151435535 |
| ENSDART00000136943 | si:dkey-21e2.13 | 2.273396315 | 3.247336732 | 1.860650754 | 0.467276759 |
| ENSDART00000136977 | wdr11 | -0.244389043 | -0.296442513 | -0.19757682 | -0.033240657 |
| ENSDART00000137019 | cmklr1 | 1.293423342 | 0.826870113 | 0.435422926 | 0.179038361 |
| ENSDART00000137021 | cabp4 | -0.36160439 | -0.600623701 | -0.526609212 | -0.179412794 |
| ENSDART00000137026 | fam208ab | -0.664193623 | -0.40965157 | -0.455043185 | -0.262800574 |
| ENSDART00000137036 | ppiaa | 0.323927062 | 0.685997539 | 0.775560135 | 0.372712842 |
| ENSDART00000137037 | iqsec3b | -0.405090433 | -0.576572578 | -0.30180595 | 0.030413937 |
| ENSDART00000137038 | asic2 | -0.373156822 | -0.72567768 | -0.446804249 | -0.152680828 |
| ENSDART00000137047 | scoca | -0.178388629 | -0.216665672 | -0.49140437 | -0.218641566 |
| ENSDART00000137061 | klc1a | -0.086604178 | 0.023703263 | 0.276488629 | 0.211488076 |
| ENSDART00000137083 | impg2a | -0.438459919 | -0.214037195 | -0.28206161 | -0.025553958 |
| ENSDART00000137098 | impdh1a | -0.236861779 | -0.534821929 | -0.182618892 | 0.088190256 |
| ENSDART00000137110 | sepw1 | 0.165066135 | 0.642417047 | 0.502149684 | 0.242826446 |
| ENSDART00000137131 | dnah7 | -0.559601026 | -0.233046623 | -0.366131677 | -0.080127176 |
| ENSDART00000137135 | mei4 | -0.382006481 | -0.256759987 | -1.043399236 | -0.457749266 |
| ENSDART00000137185 | cnga3b | -0.145832564 | -0.301563517 | -1.136000666 | -0.708190865 |
| ENSDART00000137203 | ponzr6 | 2.803770034 | 2.268583293 | 2.342420518 | 1.869529587 |
| ENSDART00000137212 | dip2ca | 0.131800934 | -0.026475424 | 0.308770191 | 0.394512532 |
| ENSDART00000137214 | gpm6bb | -0.410929898 | -0.351705205 | -0.352631557 | -0.244368267 |
| ENSDART00000137228 | si:rp71-80o10.4 | -0.005305454 | 1.094390245 | 1.175846782 | 0.577915575 |
| ENSDART00000137236 | hpcal1 | -0.065446201 | -0.487196869 | -0.639863396 | -0.431847778 |
| ENSDART00000137262 | vdac3 | -0.257205324 | -0.141410703 | -0.061503392 | 0.052713477 |
| ENSDART00000137287 | pebp1 | -0.410574808 | -0.266170824 | -0.343257245 | -0.128567003 |
| ENSDART00000137291 | METTL18 | -0.209734106 | 0.269511362 | -2.440335414 | -0.139098945 |
| ENSDART00000137292 | tmem106bb | -0.265825137 | -0.29210537 | -0.411370487 | -0.25270276 |
| ENSDART00000137293 | dia1b | -0.377972932 | -0.365130624 | -0.231479469 | -0.228625621 |
| ENSDART00000137308 | plxnb1a | 0.257511262 | 0.267128747 | 0.167145068 | 0.075399376 |
| ENSDART00000137309 | gemin5 | 0.29381973 | 0.279160353 | 0.398962684 | 0.210137956 |
| ENSDART00000137315 | rdh12 | -0.392438907 | -0.330126959 | 0.208728813 | 0.139463991 |
| ENSDART00000137325 | mvp | 1.415505945 | 1.485945809 | 1.461661333 | 0.448751577 |
| ENSDART00000137332 | si:ch211-132g1.3 | -2.704437973 | -0.017410436 | -0.395377626 | -0.276512676 |
| ENSDART00000137353 | klhl17 | -0.368352542 | -0.314849207 | -0.039056928 | 0.155772163 |
| ENSDART00000137355 | tet3 | -0.245273707 | -0.407219331 | -0.050664281 | -0.06569376 |
| ENSDART00000137364 | si:dkey-33c12.3 | 0.174067673 | -0.534549963 | -0.496919129 | 0.189169246 |
| ENSDART00000137371 | atp8b3 | -0.113445648 | -0.348083251 | -0.436781891 | -0.110584084 |
| ENSDART00000137373 | si:ch211-266i6.3 | 1.973293451 | 1.253222026 | 0.772689981 | 0.242956342 |
| ENSDART00000137381 | khdrbs1b | -0.12744256 | -0.621948205 | -0.491970892 | 0.090347508 |
| ENSDART00000137391 | celsr3 | 0.73044715 | 0.817745044 | 0.845582062 | 0.571451225 |
| ENSDART00000137418 | BX004774.2 | -0.041779011 | -0.239619277 | -0.711634051 | -0.543815741 |
| ENSDART00000137424 | c1qtnf4 | -0.387372103 | -0.312978581 | -0.242907328 | -0.243606392 |
| ENSDART00000137426 | gcgb | -0.226806487 | -0.599969225 | -0.659481129 | -0.843993634 |
| ENSDART00000137443 | ANK1 (1 of many) | -0.657062014 | -0.810024547 | -0.456200062 | -0.254232075 |
| ENSDART00000137471 | MFAP4 (1 of many) | 1.677720699 | 2.807166131 | 2.128226179 | 1.146528193 |
| ENSDART00000137485 | tpd52l1 | 0.045987753 | -0.292323377 | -0.626535278 | -0.266196407 |
| ENSDART00000137486 | klhl21 | 0.151379043 | 0.263264305 | 0.384374563 | 0.339611926 |
| ENSDART00000137505 | vamp1 | -0.431791589 | -0.393372103 | -0.433651273 | -0.142134375 |
| ENSDART00000137561 | ucp3 | 2.475646127 | 3.272328187 | 3.19387131 | 0.905806223 |
| ENSDART00000137572 | mpc1 | -0.077665143 | -0.162227554 | -0.29007527 | -0.035755645 |
| ENSDART00000137620 | si:ch73-46j18.5 | -0.071928647 | 0.074859157 | 0.409828499 | 0.45946174 |

| | | | | | |
|---|---|---|---|---|---|
| ENSDART00000137624 | eef2a.2 | 2.619194541 | 2.67605773 | 3.551783602 | 3.00375243 |
| ENSDART00000137628 | mthfd1l | 3.69201835 | 0.840641874 | 3.245529647 | 1.343967089 |
| ENSDART00000137633 | si:ch211-194e15.5 | -0.00297372 | -0.215384681 | -0.930894546 | -0.593142383 |
| ENSDART00000137638 | si:ch73-25f10.6 | -0.304397619 | -0.080258192 | -0.006488078 | 0.003923632 |
| ENSDART00000137648 | si:dkeyp-123h10.2 | -0.55097741 | -0.596976893 | -0.447594093 | -0.275058949 |
| ENSDART00000137659 | cmpk2 | 0.149738862 | -0.572975919 | 2.540871599 | -0.412773402 |
| ENSDART00000137679 | sash1a | 0.328343986 | -0.118753926 | -0.077897758 | -0.030798 |
| ENSDART00000137680 | BX663503.1 | -0.232606929 | -0.407355209 | -0.255389687 | -0.079162056 |
| ENSDART00000137728 | st6gal1 | -0.677667407 | -0.699420985 | -0.736040173 | -0.083670402 |
| ENSDART00000137756 | hecw2a | -0.728856656 | -1.003560302 | -0.22088311 | 0.383324077 |
| ENSDART00000137757 | irf9 | 0.327252488 | 0.410367721 | 0.769408328 | 0.032820811 |
| ENSDART00000137782 | yif1a | 0.379643142 | 0.667454267 | 0.241343175 | 0.280143891 |
| ENSDART00000137795 | ssrp1a | 0.416335187 | 0.470997147 | 0.449691903 | 0.161896374 |
| ENSDART00000137799 | trim66 | -0.46849178 | -0.260779632 | 0.306613928 | 0.711500351 |
| ENSDART00000137817 | rap1gap2a | -0.351211604 | -0.330279371 | -0.569882036 | -0.258477103 |
| ENSDART00000137848 | palmda | -0.364561265 | -0.248835039 | -0.231952958 | -0.127331926 |
| ENSDART00000137851 | tfr1b | 0.020013224 | -0.000851268 | 0.44833745 | 0.43610072 |
| ENSDART00000137858 | alpk2 | 0.667374473 | 0.189664169 | 0.047615444 | -0.048884335 |
| ENSDART00000137889 | ank1b | -0.447220804 | -0.737203628 | 0.205089912 | 0.418795119 |
| ENSDART00000137896 | fibpb | -0.042394017 | -0.144008268 | -0.458312875 | -0.18906893 |
| ENSDART00000137899 | tmem9 | -0.365732748 | -0.460492017 | -0.592427521 | -0.282793654 |
| ENSDART00000137900 | grin2ab | -0.452993272 | -0.616161489 | 0.576592075 | 1.050321835 |
| ENSDART00000137903 | BX640512.2 | -0.158278671 | -0.01949183 | 1.915495669 | -0.120027797 |
| ENSDART00000137918 | bbc3 | 0.101268325 | 0.767629492 | 0.853650102 | 0.865966102 |
| ENSDART00000137920 | si:ch211-169p10.1 | -0.466773792 | -0.281022003 | 0.027119426 | -0.105594514 |
| ENSDART00000137967 | diras1b | -0.821870921 | -0.072822932 | 0.659174512 | 0.085565822 |
| ENSDART00000137973 | grna | 1.322360375 | 0.965948715 | 0.567189356 | -0.062803655 |
| ENSDART00000137984 | SBNO2 (1 of many) | 0.66540874 | 0.705510404 | 0.683617676 | -0.079102756 |
| ENSDART00000138026 | gabra5 | -0.35782795 | -0.794620157 | 0.43219928 | 0.956240287 |
| ENSDART00000138045 | whsc1l1 | -0.152220762 | 0.158744769 | 0.494234043 | 0.50708853 |
| ENSDART00000138048 | pbx3b | -0.599655595 | -0.602159282 | -0.228775798 | -0.019503135 |
| ENSDART00000138070 | mb | -0.127824671 | -0.447974107 | -0.919218534 | -1.274065689 |
| ENSDART00000138081 | bcl6ab | 0.948060654 | 0.690527167 | 0.988530239 | 1.012261919 |
| ENSDART00000138094 | eef1b2 | 0.3943229 | 0.586163757 | 0.433468435 | 0.083803882 |
| ENSDART00000138116 | surf6 | 0.641305655 | 0.652953357 | 0.565146387 | 0.291720746 |
| ENSDART00000138120 | pbdc1 | 0.312703559 | 0.29678566 | 0.127166498 | 0.014012212 |
| ENSDART00000138139 | zgc:153157 | -0.590921601 | -0.369202687 | -0.339560998 | -0.377004198 |
| ENSDART00000138140 | SLC4A5 (1 of many) | -0.322967614 | -0.430634585 | -0.36463151 | -0.15311364 |
| ENSDART00000138143 | si:ch211-233m11.1 | 0.782546411 | 0.773895723 | 0.974137465 | 0.285762089 |
| ENSDART00000138174 | slc16a12b | -0.100392011 | 0.567507799 | 1.765497005 | 0.866245296 |
| ENSDART00000138185 | gad1b | -0.304744145 | -0.418178899 | -0.358818296 | -0.231950915 |
| ENSDART00000138192 | tnnt3b | -0.439749688 | 3.945748378 | 4.736531723 | 2.568867245 |
| ENSDART00000138210 | ubac1 | -0.339032913 | -0.702323176 | -0.152684805 | 0.046765242 |
| ENSDART00000138216 | CR855311.1 | 1.84161592 | 1.501667665 | 0.707831526 | -0.102637826 |
| ENSDART00000138223 | si:ch211-71m22.1 | 0.850777626 | 0.641874022 | 1.554600154 | -0.004827675 |
| ENSDART00000138232 | arl13a | -0.123428499 | -0.297389057 | -0.827109894 | -0.393133456 |
| ENSDART00000138240 | tpt1 | 2.855871394 | 2.728090369 | 3.227118826 | 0.916859453 |
| ENSDART00000138246 | EMB | -0.957103253 | -1.089886746 | -0.283796575 | 0.121320475 |
| ENSDART00000138270 | copz2 | -0.133761169 | -0.127273242 | -0.35316272 | -0.169760508 |
| ENSDART00000138289 | prex1 | -0.517788451 | -0.174645698 | -0.909102985 | -0.603337454 |
| ENSDART00000138294 | rab41 | -0.246606124 | -0.179576466 | 0.274617961 | 0.438688085 |

| | | | | | |
|---|---|---|---|---|---|
| ENSDART00000138303 | si:dkey-23c22.9 | 0.616317614 | 0.5640973 | 0.527774625 | 0.331889659 |
| ENSDART00000138308 | gpr186 | -1.473279373 | -0.953004046 | -1.169373422 | -1.314238919 |
| ENSDART00000138312 | zbtb18 | -0.233659879 | -0.456610703 | -0.155318743 | 0.01760237 |
| ENSDART00000138321 | parp12b | 0.255975647 | 0.107710029 | 0.742615255 | 0.386793554 |
| ENSDART00000138328 | xpo6 | 0.06358993 | 0.090933498 | 0.285770037 | 0.126601033 |
| ENSDART00000138334 | ppp2r2ab | -0.498478384 | -0.353284175 | -0.096797126 | -0.036101553 |
| ENSDART00000138350 | rps12 | 0.592717981 | 0.744811164 | 0.572850481 | 0.279364485 |
| ENSDART00000138351 | cth | 1.012814265 | 1.438428325 | 1.066275824 | 0.495805428 |
| ENSDART00000138404 | si:ch211-199g17.2 | 1.114605772 | 0.829720741 | 1.042136455 | 0.129449804 |
| ENSDART00000138412 | acbd5a | 0.434518843 | 0.610473848 | 0.985396736 | 0.824271669 |
| ENSDART00000138415 | rnf207b | 0.16648554 | 0.461805711 | 0.480213875 | 0.411411872 |
| ENSDART00000138432 | cul2 | 0.103145244 | 0.093005828 | 0.308783998 | 0.205589181 |
| ENSDART00000138475 | sorbs2a | -0.319831096 | -0.27540368 | -0.20838001 | -0.137765232 |
| ENSDART00000138477 | BX571945.1 | -0.597559877 | -0.845904193 | -0.62766334 | -0.501592231 |
| ENSDART00000138527 | rab20 | 0.632157851 | 0.410164386 | 0.278506263 | 0.009483031 |
| ENSDART00000138540 | ttyh1 | -0.563796304 | -0.437755604 | -0.092270149 | 0.165178407 |
| ENSDART00000138541 | rpgrip1 | -0.096214383 | -0.271667846 | -0.617065899 | -0.286110235 |
| ENSDART00000138606 | las1l | 0.170918093 | 0.302723725 | 0.361920168 | 0.174466647 |
| ENSDART00000138621 | zgc:153867 | 3.188427096 | 3.682050782 | 3.111570491 | 2.809604178 |
| ENSDART00000138633 | cd99 | 1.201301736 | 1.818711588 | 1.295470825 | 0.581149232 |
| ENSDART00000138658 | si:ch211-117m20.4 | -0.301116828 | -0.255854166 | -0.65213458 | -0.329202684 |
| ENSDART00000138661 | tspan11 | -0.312645804 | -0.301551652 | -0.855364566 | -0.684847545 |
| ENSDART00000138687 | ppp1r9a | -0.410087357 | -0.337226328 | -0.257566135 | -0.006942521 |
| ENSDART00000138695 | ank1b | -0.498943842 | -0.553539638 | -0.03799849 | 0.33423854 |
| ENSDART00000138703 | trpc7a | -0.333459575 | -0.723289936 | -0.168085186 | -0.375723791 |
| ENSDART00000138714 | sult5a1 | 2.445310663 | 3.836816007 | 1.972331729 | 2.072167418 |
| ENSDART00000138719 | cyp3c4 | -2.572466969 | -4.228419984 | 0.354827699 | -0.36271005 |
| ENSDART00000138733 | cdh12a | -0.402719414 | -0.467471363 | -0.247597318 | -0.111519505 |
| ENSDART00000138747 | urb2 | 0.51682098 | 0.426109616 | 0.379169845 | 0.148916896 |
| ENSDART00000138755 | dclk1a | -0.464513032 | -0.326162873 | -0.241704881 | -0.01162875 |
| ENSDART00000138759 | clta | -0.330987711 | -0.247185854 | 1.110067577 | 0.98649153 |
| ENSDART00000138769 | cdh24b | -0.290981343 | -0.436100905 | -0.352222227 | -0.098902248 |
| ENSDART00000138793 | si:ch211-203k16.3 | -0.114803625 | -0.341143223 | -0.894722004 | -0.064858026 |
| ENSDART00000138820 | lrtm2a | -0.205456304 | -0.473378158 | -0.285055681 | -0.252340958 |
| ENSDART00000138821 | nfil3 | 1.376309909 | 2.121391468 | 1.99718084 | 1.39511595 |
| ENSDART00000138822 | SSBP4 (1 of many) | -0.268549166 | -0.417362634 | -0.271649078 | -0.060566364 |
| ENSDART00000138834 | mab21l3 | -0.591171056 | -0.506133859 | -0.165242719 | -0.104286764 |
| ENSDART00000138845 | BX571724.1 | -0.626541209 | -0.809565722 | -0.893218772 | -2.725413714 |
| ENSDART00000138850 | cacna2d2b | -0.226630579 | -0.433395437 | -0.06209639 | 0.11961013 |
| ENSDART00000138866 | pbxip1a | -0.45481847 | -0.358362517 | -0.587330976 | -0.331438505 |
| ENSDART00000138878 | ndufaf5 | -0.068766975 | -0.169285384 | -0.3114844 | -0.186576816 |
| ENSDART00000138890 | kcnd2 | -0.399063976 | -0.347742148 | -0.33423415 | -0.044907554 |
| ENSDART00000138895 | igsf9ba | -0.084504285 | -0.116086258 | 0.293952477 | 0.264801901 |
| ENSDART00000138915 | cirbpb | 0.297824374 | 0.314736859 | 0.24846932 | 0.269446209 |
| ENSDART00000138922 | elovl8a | -0.299886352 | -0.255106336 | -0.59316691 | -0.339691401 |
| ENSDART00000138954 | plekha1b | -0.288171631 | -0.266395551 | 0.185168153 | 0.476938782 |
| ENSDART00000138963 | rcan1a | 0.723709877 | 0.557125789 | 0.817932454 | 0.892580049 |
| ENSDART00000138995 | vps9d1 | 0.170379692 | -2.63011314 | -0.343991992 | -0.469718482 |
| ENSDART00000138996 | si:ch211-214p13.9 | 1.396990228 | 1.323446174 | 0.865221946 | -0.306555509 |
| ENSDART00000139002 | dus2 | 0.373595906 | 0.357863942 | 0.47349873 | 0.230284081 |
| ENSDART00000139033 | lrp1bb | -0.300129828 | -0.380241873 | -0.239929923 | -0.108887683 |

| | | | | | |
|---|---|---|---|---|---|
| ENSDART00000139035 | si:ch211-215c18.3 | -1.205532305 | -0.512285433 | 0.437440349 | 0.984798646 |
| ENSDART00000139038 | CU467861.1 | 1.207959792 | 0.605422627 | 0.760051253 | 0.598045855 |
| ENSDART00000139042 | si:ch211-132b12.7 | 0.072730857 | -0.322841214 | -0.765244522 | -0.309126323 |
| ENSDART00000139048 | glsb | -0.011350798 | -0.181323189 | -0.40931902 | -0.142239159 |
| ENSDART00000139069 | decr1 | 0.368869625 | 0.410230162 | -0.003538468 | -0.070408167 |
| ENSDART00000139087 | lgmn | 0.782730519 | 0.338051823 | 0.153128345 | -0.09953272 |
| ENSDART00000139102 | dbpb | 0.113164671 | -0.364559303 | -0.423196921 | -0.137052116 |
| ENSDART00000139124 | ckba | 1.776728636 | 1.213212461 | 1.201789834 | 0.178980804 |
| ENSDART00000139132 | steap3 | 0.144648286 | 0.301641021 | 0.828461526 | -0.036698505 |
| ENSDART00000139151 | si:ch211-193k19.1 | 1.470035909 | 0.356445905 | 0.995091982 | 1.049578611 |
| ENSDART00000139174 | fam46c | -0.16066992 | -0.125475027 | -0.118453643 | -0.616919987 |
| ENSDART00000139176 | eif2s3 | 0.381558608 | 0.49753559 | 0.407153714 | 0.243011405 |
| ENSDART00000139178 | slc25a14 | -0.405048408 | -0.151867564 | -0.020130795 | 0.082212943 |
| ENSDART00000139191 | trpc1 | -0.495478264 | -0.440776327 | -0.289497882 | -0.094213915 |
| ENSDART00000139196 | kif1ab | -0.338072705 | -0.653527626 | -0.314444638 | -0.193117382 |
| ENSDART00000139237 | ret | -0.448934402 | -0.273383642 | -0.43725639 | -0.287562193 |
| ENSDART00000139241 | rnf38 | 0.363207367 | 0.385948281 | 0.537494883 | 0.39766179 |
| ENSDART00000139259 | zfand5b | 0.677389122 | 0.588881516 | 0.462655957 | 0.625250498 |
| ENSDART00000139265 | si:ch211-12e13.12 | -0.4797238 | -0.27645528 | 0.604278273 | 0.83384752 |
| ENSDART00000139279 | rbpms2b | -0.774960547 | -0.830410704 | 0.12323811 | 0.508122144 |
| ENSDART00000139299 | zgc:162928 | 0.559235647 | 0.580335587 | 1.258831581 | 1.081287075 |
| ENSDART00000139310 | bcat2 | -0.180902608 | -0.445596686 | -0.579108461 | -0.343900677 |
| ENSDART00000139324 | si:dkeyp-41f9.3 | -0.154282935 | -0.395797654 | -0.452635792 | -0.310260961 |
| ENSDART00000139329 | cry-dash | -0.104880347 | 0.380813844 | -0.32837694 | -0.194368371 |
| ENSDART00000139339 | mrpl34 | -0.322878188 | -0.414873817 | -0.404846888 | -0.143871621 |
| ENSDART00000139368 | atp1b1a | 1.877087153 | 1.808227818 | 1.283859651 | 0.966677715 |
| ENSDART00000139387 | atp1b1b | -0.121937317 | 0.340318468 | 0.40085255 | 0.510906972 |
| ENSDART00000139392 | rras | 2.928045636 | 3.065800127 | 2.84735064 | 1.624630606 |
| ENSDART00000139412 | SMIM18 | 0.730335352 | 0.956017708 | 0.99931328 | 0.671209714 |
| ENSDART00000139448 | GALNTL6 | -0.494238764 | -0.383984121 | -0.129586789 | 0.079582323 |
| ENSDART00000139449 | si:dkey-147f3.4 | 1.865073313 | 2.098559635 | 1.808022762 | 0.571602171 |
| ENSDART00000139452 | pde4d | 0.066848851 | -0.561165148 | -0.294908352 | 0.123175575 |
| ENSDART00000139454 | pde4cb | 0.616688899 | 0.67005432 | 0.986689732 | 0.473877974 |
| ENSDART00000139461 | spock3 | -0.122253969 | -0.220033244 | -0.389207603 | -0.090203867 |
| ENSDART00000139464 | CU467861.1 | 0.786254771 | 0.561892643 | 0.40560384 | 0.357004849 |
| ENSDART00000139475 | samd10b | -0.569533963 | -0.524261904 | -0.125238267 | -0.046566236 |
| ENSDART00000139477 | trim110 | -0.21821102 | -0.289321848 | -0.373728133 | -0.159590708 |
| ENSDART00000139479 | slc12a9 | 0.435405363 | 0.483188456 | 0.53485188 | 0.300178427 |
| ENSDART00000139493 | olfm1b | -0.400757951 | -0.50188464 | -0.148663949 | 0.07973127 |
| ENSDART00000139514 | ano11 | -1.063704055 | -1.17363318 | -0.628817004 | -0.055546726 |
| ENSDART00000139517 | bbs4 | -0.163398089 | -0.230989705 | -0.424869507 | -0.049448344 |
| ENSDART00000139560 | ppp3r1a | -0.205912354 | -0.312380575 | -0.194151002 | 0.010760735 |
| ENSDART00000139565 | L3MBTL4 | -0.137141942 | -0.164863308 | -0.259223674 | -0.097063711 |
| ENSDART00000139568 | nsmfa | -0.224590353 | -0.433156538 | -0.157442583 | -0.141070436 |
| ENSDART00000139569 | zgc:92658 | -0.993075003 | -0.842810673 | -0.747118722 | -0.361693255 |
| ENSDART00000139596 | adarb2 | -0.297400744 | -0.411863485 | -0.163774062 | -0.140762569 |
| ENSDART00000139608 | upf3a | -0.436308546 | -0.475063889 | -0.33694709 | -0.215591369 |
| ENSDART00000139623 | si:dkey-85k15.6 | 1.245336652 | 2.138651628 | 1.967890086 | 1.386101724 |
| ENSDART00000139633 | pip4k2ab | -0.336773688 | -0.358059228 | -0.306910553 | -0.048531465 |
| ENSDART00000139644 | lin7b | -0.266884847 | 0.193113139 | -0.021741296 | 0.002335697 |
| ENSDART00000139659 | si:dkey-222f2.7 | 1.108120911 | 0.818052369 | 0.396214124 | -0.100365974 |

| | | | | | |
|---|---|---|---|---|---|
| ENSDART00000139668 | sept8a | 1.681949125 | 2.926818092 | 4.206471175 | 2.971436039 |
| ENSDART00000139677 | si:ch211-274p24.4 | -0.523882432 | -0.343944487 | -0.011557753 | 0.069892959 |
| ENSDART00000139682 | eps8l3b | 1.489223017 | 0.930088019 | 0.068279636 | -0.057094698 |
| ENSDART00000139684 | lypla2 | -0.275250904 | -0.278688664 | -0.279438301 | -0.171299302 |
| ENSDART00000139701 | zfand5a | -0.357581428 | -0.278083252 | -0.905903537 | -0.782247372 |
| ENSDART00000139702 | casp3a | 0.559260909 | 1.157211853 | 1.445414898 | 1.128298777 |
| ENSDART00000139715 | dpm3 | -0.11194386 | -0.125527265 | -0.41579896 | -0.188378465 |
| ENSDART00000139722 | dmtn | 1.105433633 | 1.577455563 | 1.872515003 | 1.217364091 |
| ENSDART00000139727 | si:dkey-14o18.2 | -0.196190231 | -0.516815682 | -0.462096519 | -0.093362239 |
| ENSDART00000139763 | stat5a | 3.442434617 | 3.964844196 | 3.417693165 | 3.322184683 |
| ENSDART00000139793 | BX324164.1 | -0.743832764 | -0.340035488 | -0.551790628 | -0.137670464 |
| ENSDART00000139795 | syt2a | -0.704240682 | -0.905957314 | -0.438675098 | 0.054000022 |
| ENSDART00000139796 | BX276103.2 | 0.569427954 | 0.391382554 | 0.350830467 | 0.475593369 |
| ENSDART00000139834 | mybpha | -0.819804571 | 2.682359397 | 3.592112028 | 2.697092568 |
| ENSDART00000139839 | rbmx | 0.438924108 | 0.178102249 | 0.089667681 | 0.22023097 |
| ENSDART00000139841 | gabrb1 | -0.439523493 | -0.431067027 | -0.18481289 | 0.263500104 |
| ENSDART00000139863 | dcbld2 | 0.210663919 | 0.284613411 | 0.515685864 | 0.299926066 |
| ENSDART00000139872 | oip5-as1 | -0.165940131 | -0.248925583 | -0.345111453 | -0.229952519 |
| ENSDART00000139892 | elfn2a | 0.520234542 | 0.439147218 | 0.642970325 | 0.553931258 |
| ENSDART00000139893 | kcnma1a | -0.580697526 | -0.766998113 | -0.101431021 | -0.110885596 |
| ENSDART00000139900 | diaph3 | 0.052388836 | -0.258521684 | -0.529199016 | -0.186043701 |
| ENSDART00000139917 | myha | 0.315818594 | 4.091377824 | 5.948901319 | 4.44901869 |
| ENSDART00000139937 | frmpd1a | -0.391315372 | -0.461157908 | -0.442405648 | -0.041073617 |
| ENSDART00000139947 | rapgef1a | 0.201307435 | 0.3482683 | 0.17911742 | 0.096155984 |
| ENSDART00000139954 | BX005364.1 | -0.374788931 | -0.616534875 | -0.121001872 | 0.068005074 |
| ENSDART00000139964 | rps27.2 | 1.536999183 | 2.160860082 | 1.306857449 | 1.796586172 |
| ENSDART00000139994 | bbx | -0.232049516 | -0.386470684 | -0.08432371 | -0.161403736 |
| ENSDART00000140026 | smarcd3a | -0.62517615 | -0.573651045 | 0.020278487 | 0.08791531 |
| ENSDART00000140028 | lrrc4.1 | -0.491138395 | -0.422849217 | -0.374271959 | -0.216352385 |
| ENSDART00000140032 | si:dkeyp-69c1.9 | 1.662700986 | 3.288206015 | 1.780235269 | 1.584891446 |
| ENSDART00000140058 | zgc:153867 | -0.784474297 | -0.55629273 | 0.467767396 | 0.765384352 |
| ENSDART00000140086 | SUN2 | 0.64389802 | 0.258658486 | -0.305248612 | -0.209793481 |
| ENSDART00000140089 | cngb1a | -0.120444883 | -0.269286081 | -0.242009625 | -0.11168715 |
| ENSDART00000140133 | ldhbb | -1.001178594 | -1.13585969 | -1.022737795 | -0.597954004 |
| ENSDART00000140136 | soga3b | -0.191550416 | -0.105037683 | 0.304311303 | 0.351545117 |
| ENSDART00000140145 | ajap1 | -0.237572949 | -0.464416488 | -0.62912768 | -0.262821389 |
| ENSDART00000140153 | mri1 | 1.105039291 | 1.085223643 | 1.409079861 | 0.961058569 |
| ENSDART00000140161 | necab2 | -0.820451686 | -0.597146212 | 0.232199679 | 0.59216621 |
| ENSDART00000140201 | cct8 | -0.644125552 | -3.075796442 | 0.532907513 | 0.591603801 |
| ENSDART00000140213 | npas1 | -0.414273238 | -0.495063512 | -0.247065017 | -0.227188437 |
| ENSDART00000140220 | si:ch211-168k14.2 | -0.180662482 | -0.408562234 | -0.217555172 | -0.18329178 |
| ENSDART00000140226 | CYTH2 | 0.074864847 | 0.258157093 | 0.345085089 | 0.148251292 |
| ENSDART00000140230 | zgc:123068 | 0.653257046 | 0.043755773 | 2.027002482 | 0.024521261 |
| ENSDART00000140253 | arhgef10lb | 0.447026855 | 0.470705895 | 0.502550702 | 0.280062014 |
| ENSDART00000140255 | si:dkey-12h9.6 | 0.01002873 | 0.192183301 | 0.434559619 | 0.43853246 |
| ENSDART00000140284 | zbtb16a | 2.407975491 | 2.022869932 | 2.341949009 | 2.679474288 |
| ENSDART00000140301 | TTC9 | 0.099722728 | 0.582192427 | 0.813799221 | 0.513650171 |
| ENSDART00000140302 | naca | 0.278198217 | -2.435431446 | -4.196452356 | -2.112057753 |
| ENSDART00000140365 | slc12a5b | -0.506920714 | -0.535232712 | -0.246533716 | 0.015331251 |
| ENSDART00000140375 | zfyve27 | -0.214217108 | -0.191210439 | -0.353015645 | -0.21311838 |
| ENSDART00000140422 | slmapb | -0.858203956 | -0.622061933 | -0.375625924 | 0.17086689 |

| | | | | | |
|---|---|---|---|---|---|
| ENSDART00000140430 | ssx2ipb | -4.013140014 | 0.070172399 | -2.728976212 | -1.746786541 |
| ENSDART00000140436 | map2 | -0.405015536 | -0.195771527 | -0.230781013 | 0.041022881 |
| ENSDART00000140448 | c1ql4a | -0.329031661 | -0.450164426 | -0.498485743 | -0.359733933 |
| ENSDART00000140469 | deptor | -0.370280506 | -0.334232129 | -0.271445752 | -0.25182283 |
| ENSDART00000140476 | atp6v0a2a | -0.389741082 | 0.011917076 | 0.174652763 | 0.265809184 |
| ENSDART00000140481 | nfu1 | -0.046571941 | -0.14924591 | -0.367699445 | -0.24132739 |
| ENSDART00000140490 | MFAP4 (1 of many) | 1.29105804 | 2.228445007 | 1.795046858 | 1.318796638 |
| ENSDART00000140523 | mrc1b | 0.693910087 | 0.60918831 | 0.270261416 | -0.09789336 |
| ENSDART00000140537 | dhrs11a | 0.251388963 | 0.647205288 | 0.714640117 | 0.4437028 |
| ENSDART00000140540 | si:ch211-241e1.3 | 0.461435917 | 0.712590644 | 1.616566418 | 0.894218174 |
| ENSDART00000140553 | frmpd1a | -0.279082186 | -0.31970424 | -0.46726564 | -0.019879501 |
| ENSDART00000140575 | map4l | -0.647123698 | -0.438384786 | -0.184748643 | -0.024632978 |
| ENSDART00000140586 | col1a1a | 0.136049595 | 0.737681947 | 1.130025782 | 0.5982928 |
| ENSDART00000140615 | cry1ba | -0.093526328 | 0.597915555 | 0.380272469 | 0.502019284 |
| ENSDART00000140632 | bcl2l13 | -0.49849375 | -0.354616708 | -0.335413941 | -0.223271321 |
| ENSDART00000140645 | si:ch211-69e5.1 | 0.001617087 | -0.376948151 | -0.916309926 | -0.180025883 |
| ENSDART00000140650 | ankrd6b | -0.31619699 | -0.471630097 | -0.512160686 | -0.029088129 |
| ENSDART00000140667 | spna2 | 0.558956985 | 0.663728247 | 0.276626138 | 0.347556663 |
| ENSDART00000140694 | eif4a1a | 0.811760097 | 0.762478852 | 0.622630963 | 0.570594418 |
| ENSDART00000140695 | cdca7a | 1.06377599 | 0.897895687 | 0.374788915 | 0.044135144 |
| ENSDART00000140711 | tbc1d5 | 0.254578098 | 0.244282662 | 0.119026019 | -0.013937387 |
| ENSDART00000140718 | sdhc | -0.135426633 | -0.207907336 | -0.390001522 | -0.128998947 |
| ENSDART00000140736 | si:dkey-42i9.6 | 0.573524661 | 0.712949488 | 0.985257189 | 0.193370375 |
| ENSDART00000140739 | lrrc4bb | -0.565179268 | -0.464872302 | -0.152447222 | -0.079782637 |
| ENSDART00000140764 | si:ch211-262h13.3 | 0.50717215 | 0.501857896 | 0.472590221 | 0.360293055 |
| ENSDART00000140776 | CR391940.1 | 0.57109507 | 0.234979352 | 0.004616304 | -0.046528801 |
| ENSDART00000140779 | fundc1 | -1.365145935 | -0.380101108 | -0.950161557 | -1.079798871 |
| ENSDART00000140782 | proza | -0.315906897 | -0.22277663 | -0.304920909 | -0.22580936 |
| ENSDART00000140786 | qtrtd1 | 0.520156295 | 0.318813771 | 0.348267514 | 0.13612068 |
| ENSDART00000140788 | gbp1 | 3.129821614 | 2.838243862 | 2.757243294 | 2.490411238 |
| ENSDART00000140819 | rgmd | -0.726396179 | -0.581866824 | -0.277540481 | -0.271416627 |
| ENSDART00000140827 | bcl11ab | -0.293093875 | -0.330078462 | 0.796150714 | 1.216507448 |
| ENSDART00000140828 | CU179758.1 | -0.432947507 | -0.14249918 | -0.435392677 | -0.381420161 |
| ENSDART00000140852 | rgcc | 1.458728484 | 1.527927815 | 1.077034908 | 0.967514776 |
| ENSDART00000140874 | seph | 0.619876107 | 0.651122471 | 0.195858296 | -0.082542729 |
| ENSDART00000140875 | zgc:123010 | 0.555608218 | 0.491589354 | 0.22352835 | -0.077508349 |
| ENSDART00000140897 | larp6 | 0.136488111 | 0.20186899 | 0.305132975 | 0.028670036 |
| ENSDART00000140908 | gse1 | 0.266945251 | 0.781532641 | 1.123032019 | 0.70398488 |
| ENSDART00000140920 | prom1b | 0.88825196 | 0.404698565 | 0.393910982 | 0.372363663 |
| ENSDART00000140940 | cluhb | -1.005907144 | -1.058846619 | 1.005581701 | -0.029667711 |
| ENSDART00000140970 | si:ch73-52p7.1 | 1.046367332 | 0.798731839 | 0.265928239 | -0.108652338 |
| ENSDART00000140995 | mfsd2b | -0.430905256 | -0.255202968 | -0.49728314 | -0.134050367 |
| ENSDART00000141030 | pde4ca | -0.308097694 | -0.281644648 | -0.391359015 | -0.416436171 |
| ENSDART00000141031 | BX004774.2 | -0.096197487 | -0.455215086 | -0.947274727 | -0.721728098 |
| ENSDART00000141062 | man2b1 | 2.731854161 | 3.054858723 | 3.461869821 | 2.8427063 |
| ENSDART00000141068 | sox11b | 1.611838536 | 2.177304965 | 2.010624009 | 1.194765222 |
| ENSDART00000141086 | robo2 | -0.237929535 | -0.432116468 | -0.02855362 | 0.131332743 |
| ENSDART00000141087 | crema | 0.064696008 | -0.100227063 | -0.477732232 | -0.233875873 |
| ENSDART00000141103 | gfra4a | -0.446789167 | -0.472299982 | -0.170582645 | 0.04427624 |
| ENSDART00000141107 | cd99 | 2.277092862 | 3.207492883 | 2.696592348 | 2.108764519 |
| ENSDART00000141110 | rpl10a | 0.638320568 | 0.343299632 | 0.070055535 | 0.288835787 |

| | | | | | |
|---|---|---|---|---|---|
| ENSDART00000141132 | ppp6r2b | -0.175401955 | -0.328399574 | -0.337567841 | -0.076173198 |
| ENSDART00000141142 | mvp | 1.187603651 | 1.369710798 | 0.919014152 | 0.045597494 |
| ENSDART00000141157 | ablim3 | -0.090063728 | -0.09247473 | -0.384522771 | 0.048083962 |
| ENSDART00000141173 | elavl4 | 2.4638469 | 2.569363943 | 2.090923257 | 1.273250901 |
| ENSDART00000141177 | slc24a4a | -0.524899894 | -0.691002096 | -0.394086808 | -0.038902918 |
| ENSDART00000141188 | st3gal3b | 0.577420785 | 0.203550656 | -0.28232291 | -0.213031688 |
| ENSDART00000141193 | clu | -0.012066925 | 0.064405046 | -0.155611703 | -0.357238215 |
| ENSDART00000141194 | mpp4a | -0.333593633 | -0.165171491 | -0.10082691 | 0.099007973 |
| ENSDART00000141224 | rpl4 | 0.608116905 | 0.375484686 | 0.195287178 | 0.211394215 |
| ENSDART00000141227 | anxa13l | -1.29005108 | 2.955855077 | 2.234819741 | 2.935945321 |
| ENSDART00000141231 | si:dkey-18a10.3 | -0.291486428 | -0.180620296 | -0.412941733 | -0.186503605 |
| ENSDART00000141237 | si:dkeyp-113d7.10 | -0.138953924 | 0.055420663 | 0.3781204 | 0.217616879 |
| ENSDART00000141238 | sdk1b | -0.289169472 | -0.436527814 | -0.423787237 | -0.252263885 |
| ENSDART00000141263 | ENKD1 | -0.178779434 | -0.320320795 | -0.370791775 | -0.057231507 |
| ENSDART00000141265 | znf106b | 0.649691237 | 0.737898777 | 0.38852404 | 0.011375685 |
| ENSDART00000141278 | si:ch211-14c7.2 | 0.247729936 | 0.220022891 | 0.550240269 | 0.375370237 |
| ENSDART00000141328 | pip5kl1 | 0.719167688 | 0.681103792 | 0.759572826 | 0.613914474 |
| ENSDART00000141338 | enc1 | 0.132252305 | 0.206249243 | 0.415692262 | 0.451357063 |
| ENSDART00000141340 | si:dkey-183n20.15 | -0.434296772 | -0.549401302 | -1.33032419 | -0.748989345 |
| ENSDART00000141345 | si:ch211-133l5.7 | 0.067098833 | -0.266407921 | -0.458017208 | -0.557765253 |
| ENSDART00000141367 | si:ch211-238n5.4 | -0.027130317 | 0.13157615 | 0.368504617 | 0.417326096 |
| ENSDART00000141373 | si:rp71-36a1.1 | 0.482287147 | 0.869957532 | 0.833507441 | 0.171228281 |
| ENSDART00000141382 | rab34b | 1.020539104 | 1.398669915 | 1.148932644 | 0.356641036 |
| ENSDART00000141385 | si:dkey-27p18.3 | 0.644114707 | 0.569712036 | 1.095314507 | 0.257528776 |
| ENSDART00000141387 | mtus2a | -0.19505707 | -0.297253066 | -0.281123654 | -0.028898398 |
| ENSDART00000141397 | ube2e1 | 0.445794988 | 0.509089566 | 0.636760169 | 0.02502865 |
| ENSDART00000141401 | arhgap12b | -0.092120969 | -0.196587501 | -0.494285673 | -0.272796913 |
| ENSDART00000141421 | flna | 2.437429319 | 2.513695725 | 2.398098837 | 2.160679529 |
| ENSDART00000141437 | sdc2 | 0.001568967 | 0.007072686 | -0.196613876 | -0.36025161 |
| ENSDART00000141444 | abcg4a | -0.060112999 | 0.269266665 | 0.721083006 | 0.688157505 |
| ENSDART00000141446 | coro1cb | -0.332235831 | 0.115994662 | 0.453423489 | 0.231540088 |
| ENSDART00000141487 | pvrl3b | -0.333456043 | -0.273917158 | -0.488003853 | -0.734425178 |
| ENSDART00000141493 | ANO2 (1 of many) | 0.056182997 | -0.29662332 | -0.7863422 | -0.307277237 |
| ENSDART00000141553 | pfkfb1 | -0.184104594 | -0.132769648 | -0.647913533 | -0.249279861 |
| ENSDART00000141569 | tpt1 | 0.218299383 | 0.390841103 | 0.22789951 | -0.0744535 |
| ENSDART00000141578 | shmt1 | 2.367581981 | 4.511004025 | 4.354684857 | -7.25195E-16 |
| ENSDART00000141579 | fbxo10 | 0.546896982 | 0.414684656 | 0.492652565 | 0.452194062 |
| ENSDART00000141596 | gpat3 | 0.765600139 | 0.644414234 | 0.654748666 | 0.141764441 |
| ENSDART00000141597 | lrrfip1b | -0.336053735 | 0.307524713 | 0.270509711 | 0.2738567 |
| ENSDART00000141606 | commd8 | 0.496320125 | 0.646713153 | 0.422272129 | 0.302789387 |
| ENSDART00000141625 | si:ch211-156p11.1 | 0.900660828 | 1.143855847 | 0.822477469 | 0.791150698 |
| ENSDART00000141626 | gtf2h5 | 0.582335915 | 0.792355335 | 0.589249078 | 0.547281799 |
| ENSDART00000141629 | nrxn2a | 0.086318651 | -0.075949898 | 0.321968103 | 0.300507297 |
| ENSDART00000141634 | grid2 | -0.260805209 | -0.581973695 | -0.28493372 | -0.055775802 |
| ENSDART00000141657 | anxa4 | 0.870599992 | 0.703638401 | 0.563849009 | -0.254549515 |
| ENSDART00000141671 | socs2 | -0.321775893 | -0.091083829 | -1.001751456 | -1.186845404 |
| ENSDART00000141676 | rpl12 | 0.410901536 | 0.227715625 | 0.271640237 | 0.155083239 |
| ENSDART00000141691 | opn7d | -0.282482677 | -0.654185182 | -0.202456799 | -0.407261262 |
| ENSDART00000141697 | mthfd1a | 0.286969677 | 0.500882111 | 0.266896386 | -0.002369285 |
| ENSDART00000141714 | nupr1 | 1.321058122 | 1.103287011 | 0.703763985 | 0.083130869 |
| ENSDART00000141716 | inpp4b | -0.094098482 | -0.269093973 | -0.37484544 | 0.075565559 |

| | | | | | |
|---|---|---|---|---|---|
| ENSDART00000141734 | hivep2a | -0.218148846 | -0.337091836 | -0.481607769 | -0.327040294 |
| ENSDART00000141750 | tnnt3b | -1.89549374 | 3.616832877 | 4.325157956 | 3.265344737 |
| ENSDART00000141752 | asphd1 | -0.390703626 | -0.359372328 | -0.291984856 | 0.011586147 |
| ENSDART00000141779 | fndc5a | -1.597130885 | -1.939288998 | -0.836516829 | 0.004414498 |
| ENSDART00000141791 | si:ch211-284e13.6 | -0.25009242 | -0.144926208 | 0.53594874 | 0.219211266 |
| ENSDART00000141831 | WDR31 | -0.216001124 | -0.160387595 | -0.391138756 | -0.241605138 |
| ENSDART00000141835 | tspan18b | -0.306678314 | -0.201727051 | 0.109445214 | 0.302434848 |
| ENSDART00000141837 | esd | 0.782720008 | 0.801148933 | 0.794217077 | 0.347191174 |
| ENSDART00000141841 | psmb7 | 2.547438262 | 1.548911002 | 3.335199455 | 2.703257309 |
| ENSDART00000141868 | si:dkey-184a18.5 | 0.011848086 | 0.007989242 | -0.315131621 | -0.067508448 |
| ENSDART00000141876 | ftr30 | 0.380222613 | 0.277973219 | 1.408194744 | 0.300573986 |
| ENSDART00000141877 | kcnk10a | 1.57307368 | 1.979146889 | 1.661354893 | 0.924431526 |
| ENSDART00000141917 | ank1a | -1.167872922 | -1.059135645 | -0.600851313 | -0.041118915 |
| ENSDART00000141920 | si:ch211-224l10.4 | -1.330425532 | -0.617989075 | -0.986788428 | -0.796603249 |
| ENSDART00000141941 | zfpm2a | 3.791965294 | 3.402723828 | 3.788725401 | 3.249505128 |
| ENSDART00000141955 | mcf2b | -0.173106859 | -0.26422462 | -0.398065805 | 0.016363494 |
| ENSDART00000141958 | cdh24a | -0.328998631 | -0.596758779 | -0.216164592 | 0.127161048 |
| ENSDART00000141974 | acsl3b | -0.154671373 | 0.25230778 | 0.484219802 | 0.119435773 |
| ENSDART00000141977 | herc3 | -0.166490009 | -0.257660784 | -0.135253579 | -0.013930049 |
| ENSDART00000141981 | loxhd1a | 0.329259022 | 0.257582221 | 0.952686722 | 1.148017375 |
| ENSDART00000142010 | ncam1b | -0.299560612 | -0.317330716 | -0.349579067 | -0.156694047 |
| ENSDART00000142013 | penka | -0.474299096 | -0.531588241 | -0.656476493 | -0.472875583 |
| ENSDART00000142026 | si:dkey-167i21.2 | -0.765729811 | -0.21936881 | -0.933028715 | -0.2803887 |
| ENSDART00000142061 | si:dkey-32e23.4 | -0.369490073 | -0.293305321 | 0.04108125 | 0.132687244 |
| ENSDART00000142087 | foxp2 | -0.522019738 | -0.4194261 | 0.065656049 | 0.27497631 |
| ENSDART00000142088 | si:ch211-218c6.8 | 1.058750612 | 0.593589729 | 0.534639469 | 0.071525268 |
| ENSDART00000142104 | cep162 | 0.409390298 | 0.286359079 | 0.288581786 | 0.16808044 |
| ENSDART00000142112 | vma21 | -0.082665202 | -0.082534524 | -0.180302218 | -0.34518071 |
| ENSDART00000142122 | cplx2l | -1.429903189 | -0.89072761 | 0.133226492 | 0.114783163 |
| ENSDART00000142129 | tacr2 | -0.694450519 | -0.993924228 | -0.826737427 | -0.21380073 |
| ENSDART00000142140 | ppfia2 | -0.37587085 | -0.48411577 | 0.218087957 | 0.20796301 |
| ENSDART00000142141 | grin2aa | -0.499492643 | -0.789969038 | -0.312539264 | 0.08079802 |
| ENSDART00000142146 | dmtn | 0.637268571 | 0.800533621 | 0.832554824 | 0.713529971 |
| ENSDART00000142155 | myh14 | 0.116679043 | 0.189831513 | 0.30854982 | 0.263648695 |
| ENSDART00000142157 | arl15a | -0.037401347 | -0.185890543 | -0.49114445 | -0.348659966 |
| ENSDART00000142158 | hars | 0.313357085 | 0.416680865 | 0.429639073 | 0.153346223 |
| ENSDART00000142164 | ccl38.1 | 0.183884553 | 0.311556871 | -0.574993202 | -1.074418737 |
| ENSDART00000142171 | syt6a | -0.424476082 | -0.649670345 | -0.435293945 | -0.206511692 |
| ENSDART00000142196 | fam167ab | -0.920556961 | -0.832834787 | -1.167437341 | -1.267796279 |
| ENSDART00000142223 | hivep3b | -0.312228644 | -0.441058379 | -0.334712198 | -0.18438466 |
| ENSDART00000142239 | CR450716.1 | -0.604404157 | -0.186424277 | -0.082867047 | -0.199369979 |
| ENSDART00000142336 | hipk3b | -0.202639972 | -0.175487886 | -0.276049793 | -0.081101671 |
| ENSDART00000142356 | smyd2a | 2.425657932 | 2.779397529 | 2.178998127 | 1.672270997 |
| ENSDART00000142377 | MEGF9 (1 of many) | -0.489828408 | -0.163053003 | 0.463102805 | 0.22210349 |
| ENSDART00000142389 | cracr2ab | 1.038814867 | 1.254276576 | 0.507748466 | -0.396339594 |
| ENSDART00000142397 | samd12 | -0.674160268 | -0.821942052 | -0.657095298 | -0.360701575 |
| ENSDART00000142413 | BX897741.1 | -0.396693848 | -0.590528792 | -0.411359985 | -0.422415067 |
| ENSDART00000142422 | tgm2l | 1.901086414 | 0.665689502 | 0.275506536 | 0.14002471 |
| ENSDART00000142428 | josd2 | 0.235661569 | 0.659033561 | 0.722639048 | 0.359420457 |
| ENSDART00000142450 | zgc:153219 | 0.356379183 | 0.700229042 | 0.242962571 | 0.037010709 |
| ENSDART00000142454 | si:dkey-121a11.3 | -0.218109071 | -0.329830559 | -0.334040533 | -0.225452629 |

| | | | | | |
|---|---|---|---|---|---|
| ENSDART00000142462 | tshz3a | -1.120926636 | -3.365499709 | -0.84112851 | -0.654584115 |
| ENSDART00000142467 | si:ch211-199g17.2 | 2.796012298 | 2.177853709 | 2.464842477 | 1.654064057 |
| ENSDART00000142489 | slco5a1b | -0.38890353 | -0.329924623 | -0.120653024 | -0.00920305 |
| ENSDART00000142492 | eno1b | -0.238975644 | -0.29896654 | -0.354383484 | -0.060042567 |
| ENSDART00000142524 | chrnb2a | -0.869143617 | -0.99350722 | -0.040693199 | 0.426386933 |
| ENSDART00000142529 | si:dkey-286j15.1 | 0.703675816 | 0.544043539 | 0.896839213 | 0.54754086 |
| ENSDART00000142543 | si:dkey-206f10.1 | -0.205833854 | -0.070068511 | -0.826534181 | -0.287245824 |
| ENSDART00000142546 | cdc42l2 | 0.502232145 | 0.904679926 | 0.447404351 | 0.132174294 |
| ENSDART00000142560 | pdxkb | 0.184442367 | 0.163391714 | 0.33929775 | 0.221566535 |
| ENSDART00000142569 | specc1lb | 0.378697075 | 0.547073638 | 0.617855803 | 0.275260653 |
| ENSDART00000142572 | sparc | -0.204187388 | -0.23572234 | -0.326346891 | -0.277892827 |
| ENSDART00000142584 | alx1 | 1.361674144 | 1.279023188 | 1.723053191 | -0.115335088 |
| ENSDART00000142605 | sorbs2b | -0.090898219 | -0.353328393 | -0.576838002 | -0.278787769 |
| ENSDART00000142634 | FP067425.1 | 1.971749592 | 2.170812908 | 1.958026148 | 1.009935297 |
| ENSDART00000142648 | gramd1a | -0.23770878 | -0.291013375 | -0.237165881 | -0.156534014 |
| ENSDART00000142653 | grm1a | -0.469462455 | -0.467735907 | -0.088400916 | 0.178280203 |
| ENSDART00000142654 | BX324123.2 | -0.377689742 | -0.458973085 | -0.444383379 | -0.275709525 |
| ENSDART00000142661 | osbpl2a | -0.307460163 | -0.305018775 | -0.245046053 | -0.17275397 |
| ENSDART00000142665 | kctd15a | -0.244770563 | -0.321970355 | -0.179500858 | -0.083583768 |
| ENSDART00000142678 | slc10a1 | -0.780280747 | -0.374232851 | -0.51558614 | -0.22594401 |
| ENSDART00000142687 | cnn2 | 0.635956839 | -0.518048624 | 0.612837411 | -3.662416494 |
| ENSDART00000142691 | trim35-10 | 1.464216907 | 1.327065997 | 0.977518865 | 0.007627538 |
| ENSDART00000142692 | eef1b2 | 0.349947783 | 0.50162657 | 0.39446346 | 0.093426844 |
| ENSDART00000142726 | creg2 | -0.073456763 | -0.046645374 | -0.204358474 | -0.271881041 |
| ENSDART00000142731 | rce1b | -0.103631801 | -0.326325127 | -0.391135117 | -0.061782364 |
| ENSDART00000142735 | si:ch211-13f8.1 | -0.324319655 | -0.241821619 | -0.588481479 | -0.554389586 |
| ENSDART00000142748 | UNC13A | -0.222825276 | -0.520698317 | -0.335972829 | 0.134560543 |
| ENSDART00000142772 | eif4a1a | 0.755438851 | 0.594665203 | 0.351260825 | 0.661749697 |
| ENSDART00000142778 | BX569784.1 | -0.298918438 | -0.277434239 | -0.189295135 | -0.216741666 |
| ENSDART00000142806 | stxbp5a | -0.250986358 | -0.382479123 | -0.333968332 | -0.143471828 |
| ENSDART00000142810 | gabra3 | -0.405359169 | -0.863749916 | -0.692169745 | -0.647424845 |
| ENSDART00000142815 | fdps | 0.434008031 | 0.982096797 | 0.955228312 | 0.608495312 |
| ENSDART00000142837 | enpp5 | -0.408425134 | -0.376571863 | -0.363824939 | -0.214479077 |
| ENSDART00000142851 | itih6 | -0.217828278 | -0.246312318 | -4.361209214 | -4.055941297 |
| ENSDART00000142873 | BX901918.1 | 0.845880305 | 0.530715844 | 1.008998061 | 0.591786557 |
| ENSDART00000142880 | BX294379.1 | 0.237830465 | 0.561801634 | 0.415821035 | 0.460224197 |
| ENSDART00000142884 | gngt2a | -0.004065507 | -0.183032589 | -0.673226204 | -0.483998962 |
| ENSDART00000142919 | ssh1b | 0.196097873 | 0.173857746 | 0.465900153 | 0.503515831 |
| ENSDART00000142920 | zgc:162150 | 1.709110448 | 2.426589338 | 1.780018708 | 1.501264432 |
| ENSDART00000142922 | slc17a5 | 0.2420628 | 0.323011659 | 0.210848621 | 0.174231112 |
| ENSDART00000142975 | si:ch1073-70f20.1 | -0.443514404 | -0.337920446 | -0.430300513 | -0.297428401 |
| ENSDART00000142976 | kcnip3a | -0.515340349 | -0.554488278 | -0.50826115 | -0.165828333 |
| ENSDART00000143000 | rnf220b | -0.004804151 | -0.254916502 | -0.367501204 | -0.153255414 |
| ENSDART00000143048 | lmo7b | -0.147694355 | -0.241574624 | -0.520231272 | -0.160787429 |
| ENSDART00000143056 | zyx | 0.416244119 | 0.475600395 | 0.593066786 | 0.15882784 |
| ENSDART00000143066 | ube2e1 | 0.330371306 | 0.357877825 | 0.149635317 | 0.181174607 |
| ENSDART00000143083 | CT573139.1 | -0.081020454 | -0.307026797 | -0.341094511 | -0.086434359 |
| ENSDART00000143120 | si:dkey-147f3.4 | 1.419591527 | 3.82528139 | 2.470866243 | 3.546981594 |
| ENSDART00000143152 | cpne5a | -0.664046359 | -0.825602355 | -0.316663455 | 0.050239809 |
| ENSDART00000143165 | tsc22d1 | -0.098364585 | -0.21923408 | -0.39849137 | 0.017570634 |
| ENSDART00000143199 | wasf3a | -0.216537889 | -0.280788628 | -0.398676762 | -0.269991874 |

| | | | | | |
|---|---|---|---|---|---|
| ENSDART00000143200 | kcnma1a | -0.226894558 | -0.334282966 | -0.480968248 | -0.397036214 |
| ENSDART00000143203 | parp2 | 0.638789429 | 0.581199479 | 0.559257592 | 0.245299499 |
| ENSDART00000143208 | etnk1 | 0.29866149 | 0.19770258 | 0.103751026 | 0.382131244 |
| ENSDART00000143245 | nptnb | -0.0813348 | -0.106325128 | -0.477835951 | -0.112902967 |
| ENSDART00000143286 | lnx1 | -0.74422542 | -0.772813048 | -0.069056217 | 0.13158415 |
| ENSDART00000143291 | RPL41 | 0.451636798 | 0.464258672 | 0.193261311 | 0.034762109 |
| ENSDART00000143376 | rab3b | -0.460716785 | -0.436622412 | 0.05069835 | 0.173014647 |
| ENSDART00000143382 | il1rapl2 | -0.355933029 | -0.71045348 | -0.38678508 | -0.068375826 |
| ENSDART00000143410 | EMB | -1.826210099 | -1.592080235 | -0.486327336 | 0.178400158 |
| ENSDART00000143413 | si:ch211-170d8.2 | 2.110280665 | 1.653225126 | 1.38414074 | 0.820720018 |
| ENSDART00000143434 | ubash3ba | -0.070569577 | 0.008399701 | 0.263717646 | 0.11493286 |
| ENSDART00000143457 | cndp2 | 0.393197537 | 0.458029638 | 0.473219597 | 0.244423454 |
| ENSDART00000143503 | usp21 | -0.183997919 | -0.332937745 | -0.642449016 | -0.356635014 |
| ENSDART00000143519 | impdh1a | -0.362024165 | -0.511295198 | -0.258726495 | -0.069176235 |
| ENSDART00000143554 | ampd2b | -0.328956412 | -0.382404642 | -0.421715131 | -0.209511328 |
| ENSDART00000143562 | gbp2 | 0.680404258 | 0.927169543 | 1.353668195 | 0.527828388 |
| ENSDART00000143573 | ncor1 | 0.450093687 | 0.292142618 | 0.666194454 | 0.343803309 |
| ENSDART00000143582 | si:ch211-236p22.1 | -0.593497691 | -1.183006832 | -0.39659144 | -0.180756629 |
| ENSDART00000143587 | ern2 | 0.504450294 | 0.214096186 | 0.104332211 | 0.050090305 |
| ENSDART00000143618 | birc7 | 0.867487648 | 0.469428911 | 0.548081783 | 0.007263478 |
| ENSDART00000143625 | map7d2b | 0.025364857 | -1.606530561 | -4.429840485 | -0.88152256 |
| ENSDART00000143662 | ankrd33aa | -0.118189532 | -0.449119978 | -0.599045615 | 0.004982045 |
| ENSDART00000143697 | si:dkeyp-69e1.8 | 0.277815064 | 0.526114648 | 0.670410394 | 0.498924627 |
| ENSDART00000143703 | flrt3 | -0.315208699 | -0.338016477 | -0.216002726 | -0.160149445 |
| ENSDART00000143723 | idi1 | 0.080932725 | 0.304370095 | 0.372374682 | 0.127047939 |
| ENSDART00000143741 | fkbp5 | 1.764298053 | 1.865156046 | 1.631976013 | 1.871443577 |
| ENSDART00000143743 | si:ch211-14a17.10 | 0.56355315 | 0.97468243 | 0.596767311 | 0.343957389 |
| ENSDART00000143761 | cadm4 | -0.396807548 | -0.426573725 | -0.041424036 | -0.105152637 |
| ENSDART00000143766 | tbc1d4 | 0.551251852 | 0.35059591 | 0.424872318 | 0.224396247 |
| ENSDART00000143784 | doc2d | -0.282456786 | -0.449882775 | -0.341326778 | 0.036564808 |
| ENSDART00000143787 | tnr | -0.299938525 | -0.646994379 | -0.19048806 | -0.113023917 |
| ENSDART00000143793 | C3 (1 of many) | 3.237329668 | 4.023510735 | 3.503827488 | 2.024562274 |
| ENSDART00000143819 | VASH1 | 0.033504925 | 0.324442057 | 0.595830734 | 0.534870749 |
| ENSDART00000143829 | nxnl1 | -0.184554354 | -0.325799787 | -0.486032926 | -0.185522737 |
| ENSDART00000143840 | gstk1 | 0.057517576 | -0.140839441 | -0.201267582 | -0.834065155 |
| ENSDART00000143846 | si:ch211-198c19.3 | 0.524345776 | 0.185185856 | -0.297806816 | -0.062260629 |
| ENSDART00000143860 | si:ch211-154e10.1 | -0.109941804 | -0.36125539 | -0.451271764 | -0.06288628 |
| ENSDART00000143867 | syngap1b | -0.339959547 | -0.514759454 | -0.488418852 | -0.182310837 |
| ENSDART00000143874 | akna | -0.124615674 | -0.323808831 | -0.482322372 | -0.23639171 |
| ENSDART00000143875 | BX323074.2 | 0.32505808 | -0.029313176 | -0.099305279 | -0.178101813 |
| ENSDART00000143878 | arhgef9a | -0.465269949 | -0.707003765 | -0.492066921 | -0.306386806 |
| ENSDART00000143887 | taar12g | 2.133495532 | 1.949189055 | 1.339108526 | 1.854618081 |
| ENSDART00000143909 | tbc1d9 | -0.237612035 | -0.281483225 | -0.04707848 | 0.049665559 |
| ENSDART00000143911 | si:dkey-105e17.1 | -0.529795865 | -0.492017338 | -0.300236892 | -0.145540976 |
| ENSDART00000143919 | gabbr1b | -0.121288245 | -0.415278076 | -0.279488234 | -0.175446101 |
| ENSDART00000143928 | rwdd1 | 0.331616231 | 0.265135658 | 0.236169655 | 0.064492205 |
| ENSDART00000143938 | sema3ga | 0.034230149 | -0.448431565 | 0.102011548 | 0.447499442 |
| ENSDART00000143963 | pcloa | -0.078913048 | -0.544095469 | -0.414092766 | -0.012445983 |
| ENSDART00000143969 | BX247868.1 | -1.179119979 | -1.430780681 | -0.233116942 | 0.342359274 |
| ENSDART00000143999 | tnnt3b | 0.546685869 | 4.375596584 | 5.084902064 | 3.992962258 |
| ENSDART00000144046 | rapgef4 | -0.382423404 | -0.346599548 | -0.357063132 | -0.146328348 |

| | | | | | |
|---|---|---|---|---|---|
| ENSDART00000144048 | cecr1b | 1.174703015 | 1.015529557 | 0.446419815 | -0.23634795 |
| ENSDART00000144050 | | -0.572877113 | -0.701246998 | -0.196460214 | -0.040487474 |
| ENSDART00000144053 | si:dkey-21e2.10 | 0.342164757 | 1.862494203 | 1.068542572 | -0.153388265 |
| ENSDART00000144067 | cabp5a | -0.477283062 | -0.255392436 | -0.467866686 | -0.403996112 |
| ENSDART00000144068 | zgc:101851 | 0.417625916 | 0.309307199 | 0.017931645 | -0.07898028 |
| ENSDART00000144139 | elavl4 | 1.756586356 | 1.578077782 | 0.938796608 | 0.66956976 |
| ENSDART00000144175 | ccdc136b | -0.408583735 | -0.440451575 | -0.345784282 | -0.125541648 |
| ENSDART00000144186 | pou6f2 | -0.474463335 | -0.465571046 | 0.092941552 | 0.436081081 |
| ENSDART00000144215 | pcnxl2 | -0.697249288 | -0.448197444 | 0.040412938 | 0.382373048 |
| ENSDART00000144216 | gal3st1b | -0.491885196 | -0.717325904 | -0.58217274 | -0.190076988 |
| ENSDART00000144237 | gig2p | 0.374493634 | 0.174814468 | 1.411754829 | -0.127638364 |
| ENSDART00000144238 | cry1bb | -2.336873942 | 0.347073563 | -0.04859154 | -0.311120285 |
| ENSDART00000144246 | tbxas1 | -0.011108915 | 0.001179893 | -0.320953776 | -0.371393575 |
| ENSDART00000144255 | magixb | -0.339150091 | -0.441125122 | -0.366876369 | -0.172380242 |
| ENSDART00000144261 | fryl | -0.221060382 | -0.265751165 | -0.423351874 | -0.131445521 |
| ENSDART00000144285 | si:ch211-196g2.4 | -0.287314781 | -0.275198401 | -0.33300774 | -0.198378078 |
| ENSDART00000144297 | ubac2 | -0.266999312 | -0.798888734 | -0.707434326 | -0.564642121 |
| ENSDART00000144303 | AK6 | 0.166979081 | -0.045967241 | -0.30997345 | -0.290616599 |
| ENSDART00000144308 | si:dkey-11o15.5 | 1.026391135 | 2.84679961 | 3.809819934 | 2.718686222 |
| ENSDART00000144335 | cyp27c1 | 1.539730469 | 0.927858614 | 0.30747539 | -0.356140532 |
| ENSDART00000144342 | ppp1r14ba | 0.017178827 | 0.781919568 | 1.244415225 | 1.189050173 |
| ENSDART00000144346 | cdkn1bb | 0.579581082 | 0.718938042 | 0.058805886 | -0.088669944 |
| ENSDART00000144351 | malt1 | -0.740504685 | -0.746576787 | -0.600575844 | -0.384172626 |
| ENSDART00000144353 | calcrla | -0.265245102 | -0.29868229 | -0.249979568 | -0.195320595 |
| ENSDART00000144361 | ank1b | -0.731673078 | -0.825040806 | 0.086374477 | 0.310723345 |
| ENSDART00000144448 | pcbp3 | -0.386917442 | -0.448839957 | -0.257435258 | -0.084182982 |
| ENSDART00000144469 | adora2b | -0.015759069 | -0.127966141 | -0.39792302 | -0.033165164 |
| ENSDART00000144500 | si:ch211-235f1.3 | 2.218077738 | 0.39200338 | 2.195588647 | 1.510701941 |
| ENSDART00000144503 | ran | -0.075551405 | 0.448221073 | 0.249018928 | 0.118064827 |
| ENSDART00000144516 | lonrf1l | 0.003516441 | 0.629397595 | -0.101368349 | 0.117101343 |
| ENSDART00000144544 | col1a2 | 0.443294043 | 1.698639717 | 2.716305218 | 1.282115721 |
| ENSDART00000144545 | rc3h1a | 0.1127689 | 0.086123881 | 0.323320533 | 0.192355814 |
| ENSDART00000144555 | si:dkey-84o3.3 | 0.84569976 | 1.063530072 | 1.082186309 | 0.816110648 |
| ENSDART00000144592 | sik2a | -0.05071398 | -0.135507163 | -0.424760133 | 0.101311014 |
| ENSDART00000144600 | bin1a | -0.647409952 | -0.602565457 | -0.5226216 | -0.498459302 |
| ENSDART00000144608 | plcl1 | -0.399605613 | -0.241387332 | -0.124693789 | 0.02448912 |
| ENSDART00000144647 | bcat1 | -0.053055163 | -0.144301718 | -0.580079153 | -0.061442763 |
| ENSDART00000144681 | anxa13l | 0.688342481 | 1.628944564 | 2.065542903 | 1.40538163 |
| ENSDART00000144702 | clint1a | 0.287820278 | 0.14751069 | 0.132691317 | -0.0310018 |
| ENSDART00000144711 | tmem154 | 0.48356725 | 0.317274745 | 0.352966647 | 0.032904309 |
| ENSDART00000144714 | HTRA2 | 0.502271196 | 0.745024882 | 0.567017564 | 0.566812931 |
| ENSDART00000144737 | srsf3b | 0.361810279 | 0.317106803 | 0.304841622 | 0.541582711 |
| ENSDART00000144750 | tnni2b.1 | 0.190106792 | 2.993607369 | 3.517771751 | 2.049682842 |
| ENSDART00000144766 | nars | 0.166803353 | 0.38562143 | 0.598983821 | 0.308867409 |
| ENSDART00000144767 | gabra1 | -0.494710008 | -0.525246082 | -0.558533509 | -0.303902197 |
| ENSDART00000144770 | cast | 1.895732545 | 2.316221594 | 1.433447785 | 1.629878019 |
| ENSDART00000144802 | dfnb31b | -0.206535765 | -0.346317212 | -0.484783097 | -0.018383317 |
| ENSDART00000144804 | mxra7 | -0.254910086 | -0.321688262 | -0.500299705 | -0.323281006 |
| ENSDART00000144813 | march8 | -0.52367029 | -0.417394306 | -0.174509938 | -0.172830735 |
| ENSDART00000144814 | si:dkey-31f5.8 | 0.429768416 | 1.027387879 | 0.82502917 | 0.369456203 |
| ENSDART00000144817 | cfl1 | 0.341949415 | 0.252829572 | 0.49073898 | 0.258667203 |

| | | | | | |
|---|---|---|---|---|---|
| ENSDART00000144841 | si:dkey-242g16.2 | 0.827348141 | 1.743182823 | 1.970186443 | 1.411913803 |
| ENSDART00000144842 | dctn2 | -0.498329292 | -3.353631529 | 0.161761115 | 0.105336326 |
| ENSDART00000144862 | ptpmt1 | 3.64529742 | 2.311630713 | 2.226765633 | 3.455463537 |
| ENSDART00000144886 | dicp1.1 | 1.748045166 | 0.589930553 | 0.172712572 | 0.178536852 |
| ENSDART00000144890 | bokb | -0.312629523 | -0.278040732 | -0.730243904 | -0.348692502 |
| ENSDART00000144891 | gpnmb | 2.526942309 | 2.443053131 | 2.053909603 | -0.779431544 |
| ENSDART00000144894 | bcl11ba | -0.656989822 | -0.679320488 | 0.181338486 | 0.623847694 |
| ENSDART00000144925 | snx27a | -0.201299797 | -0.367903189 | -0.251100705 | -0.13350684 |
| ENSDART00000144946 | adcyap1b | 2.385755846 | 2.421463563 | 2.507590258 | 1.873754889 |
| ENSDART00000144966 | nt5c2l1 | 0.621464058 | 1.214449784 | 1.836289861 | 0.635815132 |
| ENSDART00000144970 | meis2b | -0.299148131 | 0.020280124 | 0.107132127 | 0.297996513 |
| ENSDART00000144986 | otud5a | 0.068487051 | 0.299156316 | 0.307343363 | 0.221359831 |
| ENSDART00000144993 | tbc1d14 | 0.120746825 | 0.349052884 | 0.289914122 | 0.097064778 |
| ENSDART00000144995 | MFSD3 | -0.419691404 | -0.388696945 | -0.281399212 | -0.247251894 |
| ENSDART00000145018 | gprc5bb | -0.280944825 | -0.565589689 | -0.320436299 | -0.038626462 |
| ENSDART00000145019 | akr1a1b | 0.370317087 | 0.266617428 | 0.13791003 | -0.009346531 |
| ENSDART00000145022 | CU459186.8 | 2.820516441 | 4.510118733 | 2.721368195 | 2.385722915 |
| ENSDART00000145035 | saga | -0.094875139 | -0.251791894 | -0.561974573 | -0.363496236 |
| ENSDART00000145055 | pnrc2 | -0.148748622 | -0.240065036 | -0.476375745 | -0.443408468 |
| ENSDART00000145065 | | 4.897906418 | 2.757476262 | 4.60015098 | 5.019792283 |
| ENSDART00000145095 | celsr3 | 0.648145926 | 0.648607509 | 0.829081729 | 0.619308918 |
| ENSDART00000145096 | fam219ab | -0.366547695 | -0.337752539 | -0.191344563 | -0.015845187 |
| ENSDART00000145103 | cntfr | 0.307964555 | 0.391102809 | 0.626651223 | 0.264745691 |
| ENSDART00000145107 | agfg1b | -0.053414539 | 0.037458177 | 0.272453618 | 0.201504327 |
| ENSDART00000145108 | MFF (1 of many) | -0.387524941 | -0.341112286 | -0.251149344 | -0.013627184 |
| ENSDART00000145111 | parp4 | 1.113854381 | 0.990369473 | 0.690636927 | 0.186507426 |
| ENSDART00000145114 | ptprsa | -0.133739633 | -0.353332073 | -0.057919239 | 0.041724305 |
| ENSDART00000145124 | dact3a | 0.112856383 | 0.770301052 | 1.625107526 | 0.355230326 |
| ENSDART00000145170 | smox | 0.719497557 | 0.099508547 | -0.032495938 | 0.130866421 |
| ENSDART00000145184 | srrd | -3.467209407 | 0.181528862 | -0.326150144 | -0.189288253 |
| ENSDART00000145198 | steap4 | 0.551058744 | 0.178525002 | -0.419104103 | -0.970957834 |
| ENSDART00000145210 | ankle2 | 0.203906028 | 0.433259544 | 0.357710257 | 0.326696668 |
| ENSDART00000145215 | grin2ca | -0.197290629 | -0.356995819 | -0.467710351 | -0.053225483 |
| ENSDART00000145220 | arvcfa | -0.706915106 | -0.825791875 | -0.448746575 | -0.046016635 |
| ENSDART00000145226 | si:ch211-233h19.2 | -0.372287835 | -0.175787817 | -0.260726103 | -0.131614613 |
| ENSDART00000145230 | eef1db | 0.569560585 | 0.627404265 | 1.018116547 | 0.789247315 |
| ENSDART00000145258 | glsb | -0.010345949 | -0.147193332 | -0.750348982 | -0.227873062 |
| ENSDART00000145269 | ostm1 | 0.549197851 | 0.4592474 | 0.036533403 | -0.163457465 |
| ENSDART00000145275 | si:dkey-147f3.8 | 1.443349889 | 1.631277345 | 1.233548809 | -0.107779207 |
| ENSDART00000145282 | sepp1a | -0.462981641 | -0.396376841 | -0.662591735 | -0.92136404 |
| ENSDART00000145294 | FEZ2 (1 of many) | 0.025283475 | -0.037737708 | -0.286085795 | -0.169439894 |
| ENSDART00000145309 | exoc3 | 2.170059758 | 1.622886137 | 1.185093243 | 1.97278521 |
| ENSDART00000145330 | apoc1 | 1.228570608 | 0.722853476 | 0.218571174 | 0.363892889 |
| ENSDART00000145331 | zgc:92066 | 3.539715592 | 2.15686966 | 1.135379743 | 1.013519624 |
| ENSDART00000145342 | cbln1 | -0.285689309 | -0.291683977 | -0.350650667 | -0.126158592 |
| ENSDART00000145349 | bcat1 | -0.398195711 | -0.070627262 | -0.574724119 | -0.164567732 |
| ENSDART00000145364 | sybu | -0.525383816 | -0.345930455 | -0.451355227 | -0.158271762 |
| ENSDART00000145377 | rpl38 | 0.331388676 | 0.460944412 | 0.086514771 | -0.0487643 |
| ENSDART00000145379 | fblim1 | 0.305913455 | 0.545649791 | 0.383397127 | 0.135782412 |
| ENSDART00000145387 | shank2 | -0.271198688 | -0.514736215 | -0.289482704 | -0.011228429 |
| ENSDART00000145413 | si:ch211-253p2.2 | -0.612329709 | -0.583952587 | -0.545467055 | -0.273784221 |

| | | | | | |
|---|---|---|---|---|---|
| ENSDART00000145428 | si:ch211-51e12.7 | -0.292178538 | -0.197575677 | -0.034168177 | 0.070057863 |
| ENSDART00000145434 | gpr52 | -0.413465096 | -0.575440094 | -0.276023467 | -0.017738845 |
| ENSDART00000145436 | atp2a2a | -0.880456774 | -0.552067998 | -0.067905189 | -0.332284195 |
| ENSDART00000145449 | rpl10 | 0.64701527 | 0.814914535 | 0.479616309 | 0.351438297 |
| ENSDART00000145486 | si:ch211-243a20.3 | 1.596806687 | 1.581489321 | 1.736665736 | -0.106318155 |
| ENSDART00000145489 | ano11 | -1.065264341 | -1.355279668 | -0.34443399 | 0.075205433 |
| ENSDART00000145494 | kcnq3 | -0.150620266 | 0.575528836 | 1.458464727 | 1.126016522 |
| ENSDART00000145545 | dclk1a | -0.21717792 | -0.314020567 | -0.407893866 | -0.042242501 |
| ENSDART00000145558 | si:ch211-237l4.6 | -0.183280044 | -0.231325216 | -0.614649778 | -0.510830341 |
| ENSDART00000145562 | robo4 | 1.739515239 | 1.001047952 | 0.857159879 | -0.094789579 |
| ENSDART00000145605 | camk2d1 | 0.045519482 | 0.325117172 | 0.151136219 | 0.629830606 |
| ENSDART00000145615 | cacna1i | -0.38207214 | -0.804209146 | -0.404812647 | -0.085276411 |
| ENSDART00000145616 | adarb1b | -0.549552658 | -0.593167659 | -0.056719481 | 0.082749855 |
| ENSDART00000145668 | si:ch211-236p5.2 | -0.579324415 | -0.253977811 | -0.665119908 | -0.600847942 |
| ENSDART00000145673 | si:ch211-126i22.5 | 0.167835794 | 0.336202454 | 0.573573931 | 0.33326684 |
| ENSDART00000145681 | atp6v1h | 0.62737841 | 0.200274483 | 0.109012982 | -0.483227692 |
| ENSDART00000145691 | PTP4A3 (1 of many) | 0.061273139 | -0.037655485 | -0.389148491 | -0.598282045 |
| ENSDART00000145705 | ablim1a | -0.310864127 | 0.043751002 | 0.292255753 | 0.332622511 |
| ENSDART00000145728 | sdhc | -0.035025769 | -0.113602993 | -0.325388824 | -0.192371035 |
| ENSDART00000145743 | si:dkey-222h21.3 | 2.082143417 | 2.044714298 | 1.752169227 | 1.309410865 |
| ENSDART00000145762 | dnaaf1 | -0.407466904 | -0.372899622 | -0.590562671 | -0.565268948 |
| ENSDART00000145775 | mpp4a | -0.462462268 | -0.206618846 | -0.102159961 | 0.056169031 |
| ENSDART00000145777 | adgrb3 | -0.131394962 | -0.455026919 | -0.13584478 | 0.067074927 |
| ENSDART00000145778 | eif4g1a | 0.287822409 | 0.058653961 | 0.329719916 | 0.260115281 |
| ENSDART00000145782 | ccl34a.3 | 1.066630706 | 1.262530626 | 1.391273396 | 1.026476376 |
| ENSDART00000145789 | fbxo41 | -0.304859 | -0.365689262 | -0.001299255 | 0.043794304 |
| ENSDART00000145809 | prkag3a | -4.153185554 | -1.353616268 | -0.923701192 | -0.043754788 |
| ENSDART00000145834 | si:ch211-233a24.2 | -0.054480296 | -0.065552868 | -0.373189 | -0.049333125 |
| ENSDART00000145835 | tmx3 | -0.913440216 | 0.029499449 | 0.308787653 | 0.34181057 |
| ENSDART00000145852 | arpc5b | 3.848588728 | 2.885756027 | 4.201851599 | 3.883565062 |
| ENSDART00000145859 | slc25a23b | -0.271106084 | -0.480607261 | -0.132697812 | 0.164389769 |
| ENSDART00000145862 | gramd1ba | -0.165891466 | -0.265579133 | -0.22242289 | -0.116668883 |
| ENSDART00000145875 | ANKRD34C | -0.466770544 | -0.643453831 | -0.660279878 | 0.01171991 |
| ENSDART00000145886 | osbpl3b | 0.13475307 | 0.290875362 | 0.450588606 | 0.199350453 |
| ENSDART00000145894 | zgc:195245 | -0.325743953 | -0.471221888 | -0.661613866 | -0.246097731 |
| ENSDART00000145907 | gc3 | -0.385628346 | -0.300281403 | -0.1220973 | 0.029888955 |
| ENSDART00000145916 | lingo4a | -0.858684072 | -0.829176543 | -0.734684966 | -0.796722528 |
| ENSDART00000145929 | slc24a2 | -0.262244646 | -0.388750168 | -0.249062022 | 0.013032324 |
| ENSDART00000145933 | si:rp71-68n21.9 | 0.834432396 | 0.464946482 | 0.375058394 | -0.112544064 |
| ENSDART00000145956 | ptpn23a | 0.252886396 | 0.192353563 | 0.35477406 | 0.187678789 |
| ENSDART00000145959 | sik2a | -0.340525942 | -0.474784074 | -0.456194757 | 0.011390214 |
| ENSDART00000145970 | si:dkey-83k24.5 | 0.413467032 | 0.037399564 | -0.039197609 | -0.146263052 |
| ENSDART00000145979 | col19a1 | -0.158274269 | -0.337487778 | -0.291137211 | -0.179712512 |
| ENSDART00000145981 | mmachc | -0.180799553 | -0.336507302 | -0.603391198 | -0.368222803 |
| ENSDART00000145983 | cdkn1a | 1.243824395 | 0.764194636 | 0.498342294 | 0.064303597 |
| ENSDART00000145997 | uap1 | 1.529889024 | 1.202439925 | 1.069222792 | 0.723659511 |
| ENSDART00000145999 | prx | 1.586883428 | 3.204392125 | 2.633047079 | 1.399238803 |
| ENSDART00000146005 | gtpbp1 | 0.461505187 | 0.53679144 | 0.551879705 | 0.330029037 |
| ENSDART00000146007 | usf2 | -0.027075797 | -0.165338495 | -0.358464486 | -0.103064043 |
| ENSDART00000146008 | camkvl | -0.251523364 | -0.299024254 | -0.462517021 | -0.14005057 |
| ENSDART00000146055 | ptchd4 | -0.534340079 | -0.556417786 | 0.177414073 | 0.199962898 |

| | | | | | |
|---|---|---|---|---|---|
| ENSDART00000146070 | arvcfb | -0.070894502 | -0.208057535 | -0.298756488 | -0.110396791 |
| ENSDART00000146101 | mcoln1a | 0.490011182 | 0.372261102 | 0.240356077 | 0.080050098 |
| ENSDART00000146106 | nfil3-5 | 0.032256315 | 1.067374086 | 1.300867348 | 1.052274713 |
| ENSDART00000146107 | ywhae1 | -0.007595908 | 0.375656099 | 0.665999115 | 0.632685234 |
| ENSDART00000146113 | arhgef9b | -0.6095368 | -0.587211831 | -0.441711189 | -0.048741645 |
| ENSDART00000146114 | si:ch211-139g16.8 | 1.21401658 | 0.728240858 | 0.247635929 | -0.180282803 |
| ENSDART00000146120 | luzp2 | -0.491780393 | -0.439244253 | 0.071649151 | 0.105029542 |
| ENSDART00000146132 | si:ch211-254n4.3 | -0.407187074 | -0.522964952 | -0.215771966 | 0.042955183 |
| ENSDART00000146180 | csrnp2 | 0.237147162 | 0.374829692 | 0.697707124 | 0.402573932 |
| ENSDART00000146210 | si:ch211-214p13.9 | 1.80985471 | 2.473449699 | 1.675655513 | -0.45037085 |
| ENSDART00000146222 | BX548073.2 | 0.396874686 | 0.19924291 | 0.197044037 | 0.232842702 |
| ENSDART00000146227 | slc24a3 | -0.627503774 | -0.581196675 | -0.447342274 | -0.279205796 |
| ENSDART00000146247 | zc2hc1a | 0.181680565 | 0.536623189 | 0.750568991 | 0.668481158 |
| ENSDART00000146284 | kcnh3 | -0.880077839 | -1.089282019 | -0.771605014 | -0.433474862 |
| ENSDART00000146315 | stx5al | -0.219102203 | -0.145337646 | -0.303930137 | -0.044513676 |
| ENSDART00000146321 | fibcd1 | -0.340172452 | -0.594838596 | -0.369380512 | -0.071609744 |
| ENSDART00000146327 | sgk1 | -0.407553517 | -0.497547741 | -0.442772363 | -0.391494952 |
| ENSDART00000146348 | shisa7a | -0.257432452 | -0.343813852 | -0.064533631 | 0.047373088 |
| ENSDART00000146371 | olfm1b | -0.300182439 | -0.072903502 | 0.253479105 | 0.553473031 |
| ENSDART00000146373 | nasp | 3.16018099 | 3.265920313 | 2.837245417 | 1.580807285 |
| ENSDART00000146380 | map6d1 | -0.622612519 | -0.482084178 | -0.312516702 | -0.212427733 |
| ENSDART00000146394 | macf1a | 0.292220827 | 0.266325023 | 0.282692477 | 0.251543817 |
| ENSDART00000146400 | st8sia5 | 0.352731191 | 0.917371468 | 1.067350167 | 0.852063315 |
| ENSDART00000146411 | sst6 | 0.736642137 | 1.700467827 | 1.549366173 | 1.261691667 |
| ENSDART00000146422 | samd12 | -0.651610173 | -1.082672704 | -0.694390946 | -0.535477884 |
| ENSDART00000146424 | bin1a | -0.478145779 | -0.444599263 | -0.2729609 | -0.137965745 |
| ENSDART00000146434 | zgc:153115 | 0.499570303 | 0.817313835 | 0.902072926 | 0.441195032 |
| ENSDART00000146455 | si:dkey-245p14.7 | 1.534786293 | 2.054169347 | 1.459641722 | 1.163785428 |
| ENSDART00000146463 | samd11 | -0.429238253 | 0.041148803 | 0.036400645 | 0.142792336 |
| ENSDART00000146472 | kcng4a | -0.174962319 | -0.077658154 | -0.410630934 | -0.442662471 |
| ENSDART00000146508 | ldlrad4b | -0.645866164 | -0.626215777 | -0.091198282 | -0.040482632 |
| ENSDART00000146517 | dpysl2b | 0.345677828 | 1.022911519 | 1.287775094 | 0.946528154 |
| ENSDART00000146525 | chtopb | 3.459860751 | 4.15390826 | 1.273403713 | 1.821621278 |
| ENSDART00000146530 | epha6 | -0.320688922 | -0.528894361 | -0.185025461 | 0.089827402 |
| ENSDART00000146553 | pfkpb | -0.397983333 | -0.578737682 | -0.358519605 | -0.195354249 |
| ENSDART00000146563 | magixa | 0.075133988 | 0.355594726 | 0.57567448 | 0.363486006 |
| ENSDART00000146564 | si:ch211-198c19.3 | 1.407263699 | 0.462763747 | -0.033079511 | -0.600140762 |
| ENSDART00000146567 | cenpk | 2.135920979 | 1.866039444 | 1.584297235 | 1.144121982 |
| ENSDART00000146611 | cxcr3.3 | 0.415816929 | 0.611232638 | 0.663522207 | -0.008061564 |
| ENSDART00000146632 | rassf2a | 0.029735353 | -0.579515238 | -0.449210112 | -0.253532753 |
| ENSDART00000146641 | adgrb2 | -0.339356043 | -0.358957699 | -0.00479776 | 0.052341655 |
| ENSDART00000146648 | cd99 | 0.568325087 | 1.118199036 | 0.745908705 | -0.085222667 |
| ENSDART00000146671 | lgals3a | 0.211860458 | 0.266423347 | 0.163466771 | 0.057807983 |
| ENSDART00000146704 | pax6b | -0.61310857 | -0.673866221 | -0.260691596 | -0.192950711 |
| ENSDART00000146708 | adar | -0.042581428 | -0.004444765 | 0.40535861 | 0.024953366 |
| ENSDART00000146712 | ccdc85a | -0.396306457 | -0.805958759 | -0.536293039 | -0.356615708 |
| ENSDART00000146715 | st6galnac3 | -0.320119064 | -0.207968819 | 0.143950361 | 0.08219826 |
| ENSDART00000146742 | ewsr1b | 0.415208463 | 0.221707988 | 0.300205339 | 0.348462361 |
| ENSDART00000146744 | nfkbiab | 0.787570548 | 0.783257317 | 0.468593447 | 0.633863732 |
| ENSDART00000146767 | fosl1a | 0.841007094 | 1.285901651 | 0.733422298 | 0.173089455 |
| ENSDART00000146775 | kank3 | 2.938783957 | 2.713493484 | 2.4846528 | 0.885288567 |

| | | | | | |
|---|---|---|---|---|---|
| ENSDART00000146776 | tspan2b | 0.412627689 | 1.44398444 | 1.603529521 | 0.88938646 |
| ENSDART00000146785 | tmem163b | -0.68540579 | -0.770110571 | -0.115396875 | 0.356833449 |
| ENSDART00000146808 | si:ch73-184c24.1 | 1.14465672 | 1.950542793 | 2.864785673 | 1.868117236 |
| ENSDART00000146872 | emid1 | 0.014325328 | 0.393704295 | 0.644359098 | 0.692252594 |
| ENSDART00000146873 | tmem183a | 2.267285842 | 2.498042527 | 2.600977396 | 2.807645399 |
| ENSDART00000146892 | abcg2d | 0.141142908 | -0.029073312 | -0.499348256 | -1.743973024 |
| ENSDART00000146914 | mpp6a | -0.247758591 | -0.21742389 | -0.181144549 | -0.11922972 |
| ENSDART00000146930 | pdk3a | -0.237508559 | -0.287205966 | -0.210198002 | -0.212117205 |
| ENSDART00000146956 | fibpb | 0.004335049 | -0.126807206 | -0.29094593 | -0.318585256 |
| ENSDART00000146962 | STX3 (1 of many) | -0.081644241 | -0.246049482 | -0.328837917 | -0.149895857 |
| ENSDART00000146991 | si:ch211-127d4.3 | -0.09259617 | -0.379853865 | -0.53001099 | -0.593912213 |
| ENSDART00000147001 | si:dkeyp-72h1.1 | -0.667988087 | -0.713655804 | -0.172938308 | 0.232582151 |
| ENSDART00000147004 | oxsr1b | -0.068072204 | -0.071587779 | -0.30197654 | -0.095099951 |
| ENSDART00000147009 | si:dkey-251i10.2 | 2.257544115 | 2.36261429 | 1.933731915 | 0.997576718 |
| ENSDART00000147019 | si:ch211-235o23.1 | 0.394717538 | 0.700152429 | 0.448882118 | 0.273599764 |
| ENSDART00000147068 | olfml2ba | -0.192176717 | 0.036009408 | -0.274084705 | -0.587185898 |
| ENSDART00000147105 | tdh | -0.645059373 | -0.498416454 | -0.202501341 | -0.068176647 |
| ENSDART00000147110 | tubb6 | 1.236908798 | 0.772404142 | 0.732906531 | -0.307326846 |
| ENSDART00000147137 | si:dkey-161j23.6 | -1.245759967 | -0.352718506 | 0.050703213 | 0.033267232 |
| ENSDART00000147175 | si:dkeyp-69c1.7 | -0.274009215 | -0.118199859 | 0.157998072 | 0.359473526 |
| ENSDART00000147188 | trpm3 | -0.625119804 | -0.737554924 | -0.108574358 | 0.285413653 |
| ENSDART00000147201 | slc5a7a | -0.334790717 | -0.440092664 | -0.539424926 | -0.607892581 |
| ENSDART00000147217 | atrnl1a | -0.191475884 | -0.379227232 | -0.255587915 | -0.309192132 |
| ENSDART00000147218 | efna3a | -0.679676128 | -0.850928012 | -0.177104508 | -0.025688419 |
| ENSDART00000147246 | MPP4 (1 of many) | -0.296462508 | -0.373112158 | -0.170989023 | -0.099561343 |
| ENSDART00000147264 | fam213b | 0.418860252 | 0.552271711 | 0.440362956 | 0.212029386 |
| ENSDART00000147265 | rpl8 | 1.326651746 | 0.800643986 | 0.962132043 | 0.631592879 |
| ENSDART00000147275 | plat | 0.870075522 | 0.84761039 | 0.802236758 | 0.134643658 |
| ENSDART00000147285 | tomm70a | -0.111704054 | -0.295896007 | -0.277787909 | -0.179585993 |
| ENSDART00000147308 | amigo1 | -0.366406631 | -0.580638611 | -0.462226604 | -0.20619589 |
| ENSDART00000147318 | zgc:123105 | -0.365732482 | -0.633059491 | -0.048801121 | 0.05490191 |
| ENSDART00000147341 | tubb4b | 0.566193096 | 0.992023395 | -0.136369873 | 0.130190834 |
| ENSDART00000147343 | dnase2 | 0.744692712 | 0.455061161 | 0.199303414 | -0.152365099 |
| ENSDART00000147368 | osbp2 | -0.137018463 | -0.22987814 | -0.404444797 | -0.125629393 |
| ENSDART00000147377 | si:ch211-218c6.8 | 0.580687503 | 0.496233306 | 0.169999239 | -0.049284637 |
| ENSDART00000147394 | rcvrna | -0.601511252 | -0.165338563 | -0.351313293 | -0.435988789 |
| ENSDART00000147422 | arhgap23b | -0.106264079 | -0.33507693 | -0.670558797 | -0.201207875 |
| ENSDART00000147439 | atp6v0b | -0.143289001 | -0.224190834 | -0.266752197 | -0.126455259 |
| ENSDART00000147461 | pih1d1 | 2.042300587 | 3.359045894 | 1.936071133 | 3.025420547 |
| ENSDART00000147464 | ryr3 | -0.823352364 | -0.716491782 | 0.05941278 | 0.608108075 |
| ENSDART00000147473 | pcsk1 | -0.408472737 | -0.345735504 | -0.351470668 | -0.372482226 |
| ENSDART00000147474 | stmn2b | 0.410261377 | 1.543598694 | 2.044768785 | 1.667647819 |
| ENSDART00000147481 | phf11 | 2.519105146 | 2.056646778 | 4.198159431 | 2.169035406 |
| ENSDART00000147483 | ephb6 | -0.346946617 | -0.195722816 | 0.204471928 | 0.396713027 |
| ENSDART00000147496 | wsb1 | 0.50240599 | 0.657726247 | 0.306985059 | 0.144649209 |
| ENSDART00000147502 | grhprb | 0.278026338 | -0.217638113 | -0.512645749 | -0.092734666 |
| ENSDART00000147504 | bzw2 | 0.031120244 | -0.513458671 | -0.643056049 | -0.325765553 |
| ENSDART00000147516 | si:ch73-343l4.8 | 0.629933021 | 0.124466643 | -0.147547456 | -2.286690979 |
| ENSDART00000147518 | fam89b | 0.288965136 | 0.576511697 | 0.598868415 | 0.448283091 |
| ENSDART00000147527 | si:dkey-9i23.14 | 2.675666068 | 2.038792867 | 2.055161538 | 1.544748993 |
| ENSDART00000147581 | celf5b | -0.452125975 | -0.583204528 | -0.153939994 | -0.143297487 |

| | | | | | |
|---|---|---|---|---|---|
| ENSDART00000147582 | agmo | -0.172065266 | -0.060434285 | -0.409289881 | -0.408628944 |
| ENSDART00000147609 | shc1 | -0.131375988 | 0.113051171 | 0.56388659 | 0.681815315 |
| ENSDART00000147617 | bcl2l1 | 0.692333613 | 0.690280735 | 0.392732217 | 0.224132994 |
| ENSDART00000147625 | pik3r3b | -0.257748291 | 0.021181096 | 0.371087748 | 0.412468691 |
| ENSDART00000147635 | gpd1c | 0.454472166 | 0.231690448 | 0.167995006 | -0.010446218 |
| ENSDART00000147658 | bhlhe22 | -0.302235414 | -0.31899187 | -0.305987775 | -0.102192149 |
| ENSDART00000147682 | atp1a1a.2 | 1.135451283 | 0.290531702 | 0.256841542 | -0.140895848 |
| ENSDART00000147683 | inpp4b | -0.101056224 | -0.522680445 | -0.598179821 | -0.221205188 |
| ENSDART00000147686 | si:dkey-222p3.1 | -0.111715584 | -0.482884964 | -0.578818209 | -0.433340731 |
| ENSDART00000147694 | tm4sf4 | -0.310160873 | 0.021226995 | -0.077203568 | -0.031986481 |
| ENSDART00000147699 | mical3b | 0.112525626 | 0.108456465 | 0.226887131 | 0.765699718 |
| ENSDART00000147729 | rps24 | 0.371025561 | 0.416364277 | 0.295416172 | 0.004577514 |
| ENSDART00000147731 | BX927329.1 | 0.853591712 | 1.100390597 | 0.929772221 | 0.285035676 |
| ENSDART00000147742 | fgfr1a | -0.118664773 | -0.073758578 | -0.167833486 | -0.603740653 |
| ENSDART00000147779 | sh3glb2b | -0.245646809 | -0.296043128 | -0.126401752 | -0.042735049 |
| ENSDART00000147787 | znf512 | 1.743634906 | 1.038845919 | 2.753652323 | 1.346161863 |
| ENSDART00000147789 | slco2a1 | -0.167945299 | -0.15402624 | -0.306762825 | -0.440433916 |
| ENSDART00000147793 | gcdha | -0.253694486 | -0.123690535 | -0.220775945 | -0.080919569 |
| ENSDART00000147794 | nt5c2l1 | 2.1295515 | 2.541950381 | 3.263642166 | 1.818207515 |
| ENSDART00000147799 | ndrg3b | -0.751369578 | -0.184479497 | 0.170135881 | -0.060187322 |
| ENSDART00000147820 | carkd | -0.239894355 | -0.307342094 | -0.0807657 | -0.048288884 |
| ENSDART00000147821 | spsb4a | 0.371275002 | 0.33311003 | 0.599280348 | 0.314031744 |
| ENSDART00000147826 | jade3 | 0.261388203 | 0.282176502 | 0.335957333 | 0.193451186 |
| ENSDART00000147831 | ACAP2 (1 of many) | -0.265239588 | -0.086744823 | 0.171263201 | 0.342225898 |
| ENSDART00000147849 | klf6a | 1.130982604 | 1.870693821 | 1.403514717 | 0.445060502 |
| ENSDART00000147851 | eef1b2 | 0.252066551 | 0.849118612 | 0.755935234 | 0.422914839 |
| ENSDART00000147854 | mfap5 | -0.362752481 | -0.424642186 | -0.795916329 | -0.74190928 |
| ENSDART00000147868 | vwc2 | -0.776498099 | -0.767193994 | -0.55435848 | -0.286994867 |
| ENSDART00000147884 | syne2b | 0.046598878 | -0.495566729 | -1.074400394 | -0.335549303 |
| ENSDART00000147889 | kat5a | 0.445126528 | 0.474310239 | 0.435051995 | 0.434365446 |
| ENSDART00000147902 | megf11 | -0.496471177 | -0.600303312 | -0.359789074 | -0.208412389 |
| ENSDART00000147903 | lrit1a | -0.179060864 | -0.15522738 | -0.479781004 | -0.346169989 |
| ENSDART00000147905 | fam213aa | -0.253972183 | -0.158419429 | -0.284170369 | -0.196877631 |
| ENSDART00000147938 | heg1 | -1.97027775 | -0.730181455 | -5.193199021 | -2.593528186 |
| ENSDART00000147963 | bco2a | -0.106052135 | -0.070354665 | -0.433217487 | -0.165630718 |
| ENSDART00000147968 | rbpms2b | -0.96188315 | -0.47610893 | 0.419978504 | 0.426103029 |
| ENSDART00000147991 | g3bp1 | 3.532775357 | 2.102820417 | 2.620663227 | 2.000522751 |
| ENSDART00000147998 | plxna2 | -0.136413264 | -0.347150332 | -0.077394761 | 0.05783618 |
| ENSDART00000148011 | cby1 | 0.256002046 | 0.39067073 | 0.214338805 | 0.125714094 |
| ENSDART00000148038 | lrfn4b | -0.35705614 | -0.30916456 | 0.014604429 | 0.153474614 |
| ENSDART00000148039 | snx19a | -0.086070054 | -0.238060141 | -0.431834798 | -0.166151211 |
| ENSDART00000148055 | si:ch211-67e16.11 | -0.532227459 | -0.535880084 | -0.161015502 | 0.246252593 |
| ENSDART00000148066 | znf395b | -0.110739145 | -0.366906743 | -0.460042969 | -0.183105049 |
| ENSDART00000148073 | shisa7b | -0.212233425 | -0.392255987 | -0.298336106 | -0.168085028 |
| ENSDART00000148076 | fbxw7 | -0.313125013 | -0.183670552 | -0.007156474 | -0.059168192 |
| ENSDART00000148080 | rnf19b | 0.061412453 | -0.676245271 | -0.33437989 | -0.030752067 |
| ENSDART00000148106 | mef2aa | -0.081330082 | -0.55524128 | -0.609035118 | -0.185252531 |
| ENSDART00000148107 | SRCIN1 | -0.358492187 | -0.594267639 | -0.130298738 | 0.102935385 |
| ENSDART00000148120 | ephb2a | -0.204590684 | -0.196760275 | 0.359222548 | 0.385663441 |
| ENSDART00000148125 | fam21c | 0.420817307 | 0.137516364 | 0.034339244 | -0.014273742 |
| ENSDART00000148138 | lrrc8aa | 0.370743632 | 0.558987897 | 0.552524282 | 0.096775057 |

| | | | | | |
|---|---|---|---|---|---|
| ENSDART00000148154 | stx2a | 0.73746977 | 1.316523952 | 0.880682553 | 0.300142462 |
| ENSDART00000148163 | frmd4bb | 0.943172358 | 1.400649402 | 1.507648518 | 0.548522517 |
| ENSDART00000148175 | si:ch1073-440b2.1 | -0.235967394 | -0.243649374 | -0.449710659 | -0.316169237 |
| ENSDART00000148181 | inpp4aa | -0.260407808 | -0.050773424 | 0.267315477 | 0.333876837 |
| ENSDART00000148199 | stxbp1a | -0.541285454 | -0.573795052 | -0.292066803 | -0.020658402 |
| ENSDART00000148213 | yipf6 | -0.115338541 | -0.114709124 | -0.248877885 | -0.220567506 |
| ENSDART00000148236 | tmem179b | 0.928914935 | 1.103132699 | 0.9233627 | 0.146290121 |
| ENSDART00000148246 | pole4 | -0.080211224 | -0.333184545 | -0.567162517 | -0.350297708 |
| ENSDART00000148249 | plppr4b | -0.24256353 | -0.068311603 | 1.098973668 | 1.360337893 |
| ENSDART00000148278 | klf11a | -0.030586225 | 0.176099641 | 0.85687578 | 0.762070868 |
| ENSDART00000148280 | prr18 | 0.103510486 | 0.905956065 | 1.016662607 | 0.752186459 |
| ENSDART00000148296 | tnni1d | 0.552273501 | 5.279876604 | 5.419924358 | 3.332159231 |
| ENSDART00000148304 | myo1f | 0.706981048 | 0.452539664 | 0.150931707 | -0.296211435 |
| ENSDART00000148306 | map2 | -0.504312708 | -0.348446647 | -0.171538441 | 0.068561288 |
| ENSDART00000148327 | adamtsl7 | 2.218956465 | 1.395778664 | 0.846103004 | 0.026559812 |
| ENSDART00000148353 | usf2 | -0.107987138 | -0.265954127 | -0.405201274 | -0.200197809 |
| ENSDART00000148388 | smox | 1.14663E-15 | 4.65884204 | 4.212605499 | 2.256263175 |
| ENSDART00000148398 | MRAS (1 of many) | 1.275583164 | 1.112018386 | 0.557077896 | 0.217843634 |
| ENSDART00000148405 | si:ch211-113g11.6 | -0.63339569 | -0.861477951 | -0.083811842 | 0.259008105 |
| ENSDART00000148422 | BX649485.3 | 1.236550789 | 0.98767472 | 0.358686087 | -0.350942213 |
| ENSDART00000148432 | ptp4a3 | 0.27125556 | 0.468426427 | 0.608500351 | -0.013500628 |
| ENSDART00000148447 | hnrnpaba | 0.222094219 | 0.226002223 | 0.267110054 | 0.234088848 |
| ENSDART00000148456 | abcb4 | -0.551564057 | -0.079040965 | -0.313501093 | -1.124934266 |
| ENSDART00000148459 | cbx1b | -2.930250472 | -0.688512699 | 0.434984356 | 0.566379272 |
| ENSDART00000148465 | slc1a1 | -0.751249091 | -0.22594806 | 0.456533505 | 0.753777978 |
| ENSDART00000148475 | cspg5a | 0.090085539 | 0.726711302 | 1.793409667 | 1.632985142 |
| ENSDART00000148531 | arhgef25b | -0.052778378 | -0.311697766 | -0.416645212 | -0.115394554 |
| ENSDART00000148536 | GK3P | -0.308304163 | -0.269790809 | -0.586409039 | -0.39358576 |
| ENSDART00000148537 | roraa | -0.160828233 | -0.221872095 | -0.45696721 | -0.083100906 |
| ENSDART00000148540 | fech | 0.007306354 | 0.381217581 | -0.12547162 | -0.024425403 |
| ENSDART00000148564 | si:ch211-236p22.1 | -0.699364761 | -0.998683731 | -0.566423622 | -0.06995253 |
| ENSDART00000148570 | atp2b1b | -0.232205009 | -0.177768331 | -0.430678176 | 0.039634464 |
| ENSDART00000148576 | pygmb | -0.380093922 | -0.425025096 | -0.447938233 | -0.092571282 |
| ENSDART00000148580 | BX005108.2 | -0.474486107 | -0.575204555 | -0.246024325 | -0.012613897 |
| ENSDART00000148582 | smpd1 | 0.404585024 | 0.130204031 | 0.022544887 | -0.113079466 |
| ENSDART00000148590 | coro2ba | -0.3608902 | -0.079968694 | -0.141782791 | 0.048751465 |
| ENSDART00000148601 | insig1 | -0.03765091 | 0.839125818 | 0.622258944 | 0.165134159 |
| ENSDART00000148639 | ubxn2a | -0.292754646 | -0.267419804 | -0.368360861 | -0.228619688 |
| ENSDART00000148643 | phkb | -0.170217144 | -0.13328883 | -0.2898938 | -0.026363851 |
| ENSDART00000148648 | rps25 | 0.412929704 | 0.39629643 | 0.197445263 | 0.016589956 |
| ENSDART00000148653 | tmem145 | -0.454635973 | -0.21216688 | -0.078241248 | 0.035128268 |
| ENSDART00000148659 | clip2 | 0.500760868 | 0.904821279 | 1.156870185 | 0.693646301 |
| ENSDART00000148661 | stat2 | 0.11967996 | -0.016653911 | 1.354000143 | 0.023510931 |
| ENSDART00000148662 | kcnc3a | -0.800710128 | -1.106624714 | -0.368118634 | 0.013489304 |
| ENSDART00000148737 | btk | 1.027482823 | 0.714086499 | 0.58431037 | 0.237640489 |
| ENSDART00000148741 | met | 0.874910977 | 0.435904294 | -0.125027531 | -0.62235308 |
| ENSDART00000148764 | ednrbb | 1.564011968 | 2.847115487 | 1.8915508 | 0.999631285 |
| ENSDART00000148766 | dpf2 | 1.058462293 | 0.640887505 | 0.623810049 | 0.633616815 |
| ENSDART00000148786 | exoc5 | 0.05617118 | 0.166256406 | 0.294936832 | 0.214091952 |
| ENSDART00000148789 | wfs1b | 0.104249443 | 0.302506651 | 0.278674006 | 0.047812088 |
| ENSDART00000148794 | jupb | -0.404857846 | -0.638395215 | -0.40436567 | -0.262479222 |

| | | | | | |
|---|---|---|---|---|---|
| ENSDART00000148795 | dpf2 | 0.12341877 | -0.325727004 | -0.660147455 | -0.013247322 |
| ENSDART00000148802 | nexn | -0.381565638 | -0.122933428 | -0.397541694 | -0.196186804 |
| ENSDART00000148822 | adgrg1 | 0.746405238 | 1.031170771 | 0.764826924 | 0.143644812 |
| ENSDART00000148831 | fgf11b | -0.269342716 | -0.341549392 | -0.474527478 | -0.280873475 |
| ENSDART00000148834 | si:ch1073-155h21.2 | -0.200452217 | -0.231957247 | -0.32585922 | -0.162282019 |
| ENSDART00000148845 | BX649485.4 | 2.24411597 | 1.39694109 | 1.224587369 | -0.358910932 |
| ENSDART00000148894 | serpinb14 | 0.554435408 | 1.134239409 | 1.417283925 | 0.684075923 |
| ENSDART00000148900 | kcnc2 | -0.819802341 | -0.972756522 | -0.275729768 | 0.337871594 |
| ENSDART00000148927 | tgfb2 | 0.452467242 | 0.589301589 | 0.923363405 | 0.590115295 |
| ENSDART00000148930 | lingo4b | -0.229730557 | -0.308881186 | -0.199125951 | -0.339156928 |
| ENSDART00000148958 | fam150a | -0.081185595 | -0.392189222 | -0.650641125 | -0.329645149 |
| ENSDART00000148963 | BX649485.1 | 1.148936794 | 0.907522814 | 0.409280883 | -0.023295659 |
| ENSDART00000148971 | ctbp2a | -0.618298464 | -0.594645977 | -0.414478241 | -0.261581238 |
| ENSDART00000148975 | CU657977.1 | 0.934819878 | 0.853574256 | 0.944102682 | 0.804503627 |
| ENSDART00000148982 | bin1b | 0.009431427 | -0.214307092 | -0.49686004 | -0.050411548 |
| ENSDART00000148984 | rpl31 | 0.518485066 | 0.61867131 | 0.274282849 | 0.246502602 |
| ENSDART00000148993 | hdac6 | 0.150136767 | 0.327518494 | 0.405976253 | 0.20477563 |
| ENSDART00000148997 | nos1apa | -0.375839925 | -0.17136424 | -0.288037974 | -0.136490158 |
| ENSDART00000149019 | itpr1a | 0.453945089 | 0.40663332 | 0.191730219 | 0.048560775 |
| ENSDART00000149024 | btbd7 | -0.242016986 | -0.307624283 | -0.194506419 | -0.161017473 |
| ENSDART00000149029 | sagb | -0.046274749 | -0.449576755 | -0.950069775 | -0.043925247 |
| ENSDART00000149032 | oat | -0.368235431 | -0.40939103 | -0.17630927 | -0.009811502 |
| ENSDART00000149041 | reep1 | -0.249037096 | 0.218537351 | 0.449912583 | 0.370752516 |
| ENSDART00000149043 | CR855300.1 | 0.458598352 | 0.519077258 | 0.374073731 | 0.109291075 |
| ENSDART00000149074 | sall1b | 1.974229239 | 2.883794957 | 2.506155499 | 1.735121972 |
| ENSDART00000149079 | sagb | -0.468101765 | -0.230897218 | -0.419750319 | -0.182809331 |
| ENSDART00000149089 | anxa11b | 0.040867541 | 0.398676842 | 0.148460283 | -0.067955816 |
| ENSDART00000149115 | zgc:158659 | 1.901261478 | 2.561373529 | 3.048076346 | 2.366410956 |
| ENSDART00000149121 | clcf1 | 1.771744054 | 1.594454703 | 1.360720614 | 0.466339861 |
| ENSDART00000149151 | sptbn2 | -0.256411956 | -0.316484744 | -0.145496042 | 0.017178704 |
| ENSDART00000149158 | dmxl2 | -0.239475556 | -0.38571718 | -0.12120231 | 0.150747303 |
| ENSDART00000149169 | ssr2 | 0.474946343 | 0.309006192 | 0.206925318 | -0.029893306 |
| ENSDART00000149171 | srp9 | -0.011162136 | -0.135869324 | -0.296581397 | -0.280358452 |
| ENSDART00000149175 | pou4f3 | -1.520089487 | -1.231379123 | 0.357488642 | 0.743891028 |
| ENSDART00000149198 | epas1a | 0.235051634 | 0.343796351 | 0.502704685 | -0.080436042 |
| ENSDART00000149200 | anxa11b | 1.743372734 | 3.631513258 | 2.108721032 | 3.508584927 |
| ENSDART00000149204 | si:dkey-85n7.7 | 2.603000794 | 2.822199237 | 2.219318015 | 0.958882963 |
| ENSDART00000149218 | elk3 | 0.538927451 | 0.404579774 | 0.375543991 | 0.048753146 |
| ENSDART00000149222 | pglyrp2 | -0.019258678 | -0.204426336 | -0.652532049 | -2.674820692 |
| ENSDART00000149237 | xirp2a | 0.190889632 | 0.253883647 | 0.37076536 | 0.19112506 |
| ENSDART00000149260 | nt5c3a | 0.06053373 | -0.254595189 | -0.368052769 | -0.218176136 |
| ENSDART00000149267 | kel | 0.933657877 | 1.291347217 | 1.534062377 | 0.13083457 |
| ENSDART00000149276 | lzts3b | -0.642406039 | -0.470964243 | -0.143114993 | -0.054979203 |
| ENSDART00000149283 | zgc:112001 | -0.707617666 | -0.492468931 | 0.006563399 | 0.231921747 |
| ENSDART00000149297 | slc20a2 | -0.247675699 | -0.319353289 | -0.309560982 | -0.222732695 |
| ENSDART00000149299 | cers2b | 0.489876545 | 0.297388144 | 0.375525704 | 0.021430111 |
| ENSDART00000149311 | gpr63 | -0.396864855 | -0.311816928 | -0.030936535 | 0.061966221 |
| ENSDART00000149316 | rasa1a | -0.4434975 | -0.604862985 | 0.040513048 | 0.244834334 |
| ENSDART00000149320 | arl8a | 0.200543842 | 0.321313695 | 0.387648659 | 0.1981557 |
| ENSDART00000149335 | kcna1a | -0.92291358 | -0.966385056 | -0.49989391 | -0.001249229 |
| ENSDART00000149339 | si:ch211-175f12.2 | -0.297380973 | -0.332932663 | -0.389357991 | -0.204952824 |

| | | | | | |
|---|---|---|---|---|---|
| ENSDART00000149350 | fkbp4 | -0.198226552 | -0.122860425 | -0.266664215 | -0.086727437 |
| ENSDART00000149351 | scd | -0.095125933 | 1.291914984 | 1.511394876 | 0.539585936 |
| ENSDART00000149352 | si:ch211-163l21.11 | -0.266781177 | -0.351609906 | -0.560319486 | -0.211979525 |
| ENSDART00000149367 | si:ch73-290k24.5 | -0.213839599 | -0.466705261 | -0.018322662 | 0.130528894 |
| ENSDART00000149411 | atxn1b | -0.356760798 | -0.298446699 | -0.334747495 | -0.251003673 |
| ENSDART00000149425 | SLC25A22 (1 of many) | -0.428786809 | -0.626535356 | -0.125118952 | 0.12464047 |
| ENSDART00000149429 | boka | -0.220682727 | -0.305502243 | -0.692638969 | -0.393597854 |
| ENSDART00000149431 | sema4aa | 1.058121853 | 0.676658919 | 0.239884836 | -0.1730827 |
| ENSDART00000149443 | pcdh19 | -0.547618951 | -0.594468642 | -0.345755383 | 0.11053464 |
| ENSDART00000149468 | rps10 | 2.202930165 | 1.872426465 | 0.292067044 | 0.38759709 |
| ENSDART00000149478 | galns | 1.09483472 | 0.726913129 | 0.227424057 | 0.052897193 |
| ENSDART00000149486 | def6b | 4.188632994 | 4.544630212 | 4.117698226 | 0.985190768 |
| ENSDART00000149510 | f13a1b | 0.823651803 | 0.467086429 | 0.602200924 | 0.078453199 |
| ENSDART00000149512 | CU467655.1 | -0.320227651 | -0.180971804 | -0.440965145 | -0.207148298 |
| ENSDART00000149518 | SLC25A22 (1 of many) | -1.000880918 | -1.544420441 | -0.824653844 | -0.438490591 |
| ENSDART00000149546 | myo15aa | -0.410128868 | -0.287434444 | -0.355050844 | -0.094306045 |
| ENSDART00000149552 | FP325120.1 | -0.456523138 | -0.25356238 | -0.125718739 | 0.024031302 |
| ENSDART00000149553 | hmgcra | -0.282814218 | -0.006298507 | 0.4014174 | 0.446278745 |
| ENSDART00000149564 | CU467655.1 | -0.291119215 | -0.167904608 | -0.380163625 | -0.141898613 |
| ENSDART00000149569 | CR848047.1 | -0.094108393 | -0.299800905 | -0.352835774 | -0.193405328 |
| ENSDART00000149574 | ca14 | -0.121794187 | -0.18625006 | -0.561537732 | -0.325393218 |
| ENSDART00000149584 | gng12a | 0.603189677 | 0.302272198 | -0.104854769 | -0.377654322 |
| ENSDART00000149594 | ank3b | -0.174131144 | -0.266770414 | 0.69684285 | 0.78513869 |
| ENSDART00000149600 | tpp1 | 0.822885392 | 0.356076016 | 0.047488091 | -0.289304028 |
| ENSDART00000149608 | nexn | -0.437154987 | -0.289194529 | -0.346094746 | -0.307861675 |
| ENSDART00000149618 | mid1 | -0.406831126 | -0.160364287 | 0.131354752 | 0.136351442 |
| ENSDART00000149634 | atp6ap2 | -0.108949021 | -0.143807338 | -0.457436101 | -0.199272375 |
| ENSDART00000149652 | AL928685.4 | -0.221690272 | -0.265792933 | -0.407449839 | -0.293257945 |
| ENSDART00000149655 | adam8b | 1.224022341 | 1.219775985 | 1.033392521 | 0.588329024 |
| ENSDART00000149659 | ezh1 | 0.045526141 | 0.005041587 | -0.251417621 | -0.327846848 |
| ENSDART00000149666 | dock3 | -0.065088232 | -0.425598201 | -0.009057049 | 0.161364952 |
| ENSDART00000149684 | si:dkeyp-52c3.2 | 2.125770421 | 2.315774699 | 2.351460593 | 2.938761267 |
| ENSDART00000149685 | gnao1a | -0.290173142 | -0.321971232 | 0.118032615 | 0.301423091 |
| ENSDART00000149689 | mob1bb | -0.2705299 | -0.18213785 | -0.463988827 | -0.361129298 |
| ENSDART00000149711 | slc4a2b | 0.638486491 | 0.276788243 | -0.283420926 | -0.322786037 |
| ENSDART00000149730 | stox2a | 0.026475093 | 0.234017219 | 0.506304198 | 0.292160739 |
| ENSDART00000149740 | si:ch211-149b19.2 | -0.851560484 | -0.688459713 | 0.123433485 | 0.442881876 |
| ENSDART00000149760 | triobpa | 0.630702727 | 0.141628128 | 0.231366625 | 0.068551496 |
| ENSDART00000149768 | kif21a | -0.021643242 | 0.361851849 | 0.686688714 | 0.546979573 |
| ENSDART00000149777 | setx | 0.020956021 | 0.009824369 | 0.394668282 | 0.077312111 |
| ENSDART00000149784 | scn3b | 0.168579716 | 0.346498596 | 0.172197933 | 0.640507294 |
| ENSDART00000149786 | dhrsx | 4.71086866 | 4.438400236 | 3.335716316 | 3.374642973 |
| ENSDART00000149787 | col4a3bpa | 0.193295875 | 0.430904088 | 0.63143914 | 0.354005578 |
| ENSDART00000149816 | hspb1 | -0.566005876 | -0.156876752 | -0.219080206 | -0.313669761 |
| ENSDART00000149821 | ecsit | 4.865284679 | 4.410737197 | 4.242321149 | 4.574026911 |
| ENSDART00000149833 | cnga3a | -0.072993636 | -0.371949035 | -0.58833358 | 0.07281763 |
| ENSDART00000149859 | gldn | -0.670802099 | -0.540937826 | -0.034323087 | -0.157128438 |
| ENSDART00000149871 | cx40.8 | 0.501427104 | 0.583813885 | 0.280966778 | -0.058317019 |
| ENSDART00000149878 | otud4 | 0.242358714 | 0.196542793 | 0.87579795 | 0.138730183 |
| ENSDART00000149882 | si:ch211-117c9.1 | 1.22256875 | 0.270055012 | 0.422501128 | 0.414018825 |
| ENSDART00000149910 | abl1 | 0.087815124 | 0.252042838 | 0.385819934 | 0.051793038 |

| | | | | | |
|---|---|---|---|---|---|
| ENSDART00000149913 | b4galt1 | 0.757613392 | 0.563165908 | 0.39069258 | 0.030742735 |
| ENSDART00000149914 | aatka | -0.270082025 | -0.541170375 | -0.440872523 | -0.175726567 |
| ENSDART00000149935 | nt5c3a | 0.05115392 | -0.273408486 | -0.497911915 | -0.317644766 |
| ENSDART00000149947 | bcl2b | -0.663523207 | -0.706570637 | -0.112082166 | -0.007869569 |
| ENSDART00000149951 | BX666064.1 | 1.493495344 | 0.978793976 | 0.565382246 | -0.392061768 |
| ENSDART00000149956 | apc | 0.195183663 | 0.26527048 | 0.385323133 | 0.453680151 |
| ENSDART00000149960 | fasn | 2.288734609 | 2.375010827 | 1.541557565 | 1.919894603 |
| ENSDART00000149966 | b3gat3 | -0.190835062 | -0.286049968 | -0.133094624 | -0.134465202 |
| ENSDART00000149978 | fabp7a | 0.33770709 | 0.764013846 | 0.558825405 | 0.488886506 |
| ENSDART00000149992 | si:dkey-231j24.3 | -0.792591394 | -0.543368341 | 0.065178142 | 0.204406436 |
| ENSDART00000149993 | pygmb | -0.443639651 | -0.459929444 | -0.472379886 | -0.083989126 |
| ENSDART00000150013 | cd276 | 1.481977913 | 1.919249755 | 1.718653495 | 0.833090259 |
| ENSDART00000150023 | slc16a1a | -2.911491812 | 0.586509109 | -0.148102592 | -0.093926934 |
| ENSDART00000150028 | onecutl | -0.126745973 | -0.253663984 | -0.285061154 | -0.103777564 |
| ENSDART00000150033 | bin1b | 0.041335049 | -0.185764451 | -0.523044089 | -0.09192537 |
| ENSDART00000150036 | adam23a | -0.62166293 | -0.515362417 | 0.016147076 | 0.303932596 |
| ENSDART00000150043 | dhrsx | -0.370311342 | -3.592572721 | -0.142322612 | -0.462525038 |
| ENSDART00000150045 | adamts18 | -0.187059316 | -0.292648437 | -0.628764119 | 0.030985495 |
| ENSDART00000150050 | hnrnpaba | 0.395350897 | 0.441973379 | 0.591388417 | 0.200313414 |
| ENSDART00000150064 | cdc14b | -0.488634498 | -0.055933719 | -0.080964497 | 0.027342931 |
| ENSDART00000150068 | sox6 | -0.111707858 | -0.047191704 | 0.355284385 | 0.700337568 |
| ENSDART00000150072 | si:dkey-85n7.8 | 0.988462904 | 1.496111016 | 1.983172292 | 0.611120113 |
| ENSDART00000150088 | BX649485.2 | 1.751466958 | 1.112873174 | 0.6867673 | -0.722119651 |
| ENSDART00000150116 | alk | -0.197441993 | -0.291262237 | -0.113299878 | -0.034468735 |
| ENSDART00000150128 | foxp1b | 0.347545443 | 0.556370195 | 1.342974258 | 1.269825931 |
| ENSDART00000150146 | smarcd3b | -0.318612181 | -0.278781268 | 0.723080439 | 0.640333876 |
| ENSDART00000150148 | slc7a6 | 0.22923155 | 0.720543979 | 1.05061766 | 1.171565717 |
| ENSDART00000150149 | tdp1 | 0.245251359 | 0.536780821 | 0.276916415 | 0.174119987 |
| ENSDART00000150184 | gpatch4 | 0.624444079 | 0.471783034 | 0.550844363 | 0.152733798 |
| ENSDART00000150190 | ggctb | -0.381240597 | -0.099058914 | -0.562063682 | -1.136274784 |
| ENSDART00000150201 | MRAS (1 of many) | 1.00558208 | 0.977546999 | 0.676765678 | 0.026394977 |
| ENSDART00000150228 | chata | -0.431975216 | -0.532525907 | -0.558690348 | -0.371717362 |
| ENSDART00000150353 | mdh1aa | -0.527537579 | -0.6816935 | -0.262947 | -0.041325795 |
| ENSDART00000150362 | si:ch211-197k17.3 | -0.145582952 | 0.205377076 | 0.711396584 | 0.681320306 |
| ENSDART00000150376 | ldb1a | -0.16522649 | -0.276396303 | -0.2812216 | -0.239874431 |
| ENSDART00000150386 | si:ch73-6k14.2 | -0.397499259 | -0.321996107 | -1.565555621 | -1.132350175 |
| ENSDART00000150389 | tmem168a | 0.525344796 | 0.286469995 | 0.221730459 | 0.080011152 |
| ENSDART00000150391 | si:ch211-214j24.14 | 2.502222784 | 2.698584631 | 2.367333489 | 0.935046128 |
| ENSDART00000150400 | tbc1d2b | -4.044163911 | -1.453975239 | -0.803224358 | -1.31483616 |
| ENSDART00000150430 | pane1 | 0.044208151 | -3.417207412 | -0.655487552 | -0.439296687 |
| ENSDART00000150484 | BX465862.4 | 1.071580051 | 0.886976814 | 0.477743025 | 0.134467836 |
| ENSDART00000150553 | fam19a5a | -0.538194047 | -0.928835726 | -0.2762266 | 0.015308786 |
| ENSDART00000150580 | BX649307.1 | 0.257402348 | 0.657324095 | 0.496054556 | 0.285668011 |
| ENSDART00000150651 | BX465862.4 | 0.896323656 | 0.956165673 | 0.709439713 | 0.378868912 |
| ENSDART00000150692 | | 0.448027135 | 0.604850915 | 0.68748119 | 0.482541087 |
| ENSDART00000150751 | CR293532.1 | 1.967646607 | 1.446997774 | 2.118831868 | 1.197447208 |
| ENSDART00000150789 | USMG5 | -0.285153636 | -0.258859464 | -0.457141251 | -0.093210696 |
| ENSDART00000150839 | fam57bb | -0.627521697 | -0.387137362 | 0.25664183 | 0.060693872 |
| ENSDART00000150863 | nrp2a | 0.543372402 | 0.197731133 | 0.582446066 | 0.760458596 |
| ENSDART00000150919 | si:ch211-191i18.2 | 1.045780222 | 1.362401557 | 0.586788909 | 0.260330898 |
| ENSDART00000150927 | ubl7b | 0.617142901 | 0.651681168 | 0.305601173 | 0.171994365 |

| | | | | | |
|---|---|---|---|---|---|
| ENSDART00000150932 | lpcat1 | 1.824153615 | 2.726791026 | 2.775021546 | 2.58536861 |
| ENSDART00000150934 | alg6 | 0.502789869 | 0.534691837 | 0.441461289 | 0.13055149 |
| ENSDART00000150940 | si:ch211-193k19.2 | 1.599024227 | 0.310661889 | 0.753508784 | 0.054490277 |
| ENSDART00000150949 | nfyc | -0.535274529 | -0.301463031 | -0.212807295 | 0.074596031 |
| ENSDART00000150955 | bmpr2a | -0.667543016 | -0.436786519 | -0.136898455 | 0.053141678 |
| ENSDART00000150958 | lrrc4ba | -0.155064877 | -0.413501981 | -0.171991106 | -0.022578699 |
| ENSDART00000150961 | wdr91 | 4.358324282 | 4.633846856 | 4.480446443 | 4.649897846 |
| ENSDART00000150975 | smad7 | -0.404377575 | -0.51744405 | -0.473359841 | -0.49979754 |
| ENSDART00000151013 | ensaa | -0.338295629 | -0.414678183 | -0.638369138 | -0.341312384 |
| ENSDART00000151022 | si:ch211-191i18.4 | -0.318627462 | -0.467051418 | -0.723047751 | -0.473341501 |
| ENSDART00000151044 | baz2ba | 2.371956879 | 3.808025242 | 3.802702579 | 3.968575995 |
| ENSDART00000151084 | uqcrb | -0.019553677 | -0.080328791 | -0.732409977 | -0.191112562 |
| ENSDART00000151099 | cacng5a | -0.404305845 | -0.269968357 | -0.142737848 | 0.07277718 |
| ENSDART00000151127 | thraa | -0.798583389 | -0.802364861 | -0.351487338 | -0.589241182 |
| ENSDART00000151143 | rbfox1 | -0.676737106 | -0.648326228 | 0.275380809 | 0.524086709 |
| ENSDART00000151158 | ncaldb | 0.047289673 | -1.437087464 | -0.332259167 | 0.054087527 |
| ENSDART00000151189 | sept12 | -0.01197373 | -0.348568497 | -0.258432997 | -0.257716236 |
| ENSDART00000151195 | march7 | -0.18277274 | 0.240052097 | 0.74148711 | 0.799803253 |
| ENSDART00000151200 | mmp17a | -0.370431866 | -1.068890772 | -0.706081815 | -0.179291377 |
| ENSDART00000151203 | rgs9a | -0.190499153 | -0.256482975 | -0.462262315 | -0.086082277 |
| ENSDART00000151205 | aldh1l1 | 0.651327821 | 0.835468293 | 0.448149629 | 0.158439741 |
| ENSDART00000151229 | kcnj5 | -0.403592435 | -0.680204447 | -0.664541848 | -0.399506515 |
| ENSDART00000151235 | rcvrn3 | -0.033671108 | -0.144752489 | -0.498884135 | -0.315369716 |
| ENSDART00000151237 | oxnad1 | 0.332718432 | 0.237746325 | 0.070395116 | 0.076154746 |
| ENSDART00000151246 | gnb3b | -0.446677088 | -0.413500571 | -0.183093758 | -0.072200588 |
| ENSDART00000151247 | scn1lab | 0.293159534 | 0.684635422 | 0.628869811 | 0.299982725 |
| ENSDART00000151252 | myo10l1 | -0.09485067 | -0.323915115 | 0.494165668 | 0.855040148 |
| ENSDART00000151282 | si:ch73-248e21.5 | 0.830287534 | 0.642657587 | 0.149520253 | -0.453672012 |
| ENSDART00000151299 | scn8ab | -0.339946061 | -0.372438378 | 0.28752079 | 0.526264643 |
| ENSDART00000151305 | slc16a6b | -0.420275567 | -0.158970487 | -0.073625291 | -0.053092638 |
| ENSDART00000151311 | GPR161 (1 of many) | 0.101386436 | 0.588870162 | 0.204324291 | -0.302118942 |
| ENSDART00000151322 | KRT18 (1 of many) | 3.074943635 | 2.466392918 | 2.650579676 | 1.467715456 |
| ENSDART00000151342 | arhgap42a | 0.69255684 | 0.716882566 | 0.356486042 | 0.299890281 |
| ENSDART00000151358 | rgl3a | -0.536133566 | -0.674550597 | -0.735844208 | -0.535493428 |
| ENSDART00000151383 | eif1b | 0.629753576 | 0.386164956 | 0.580421795 | 0.441741802 |
| ENSDART00000151386 | CU659670.1 | -0.061309281 | -0.197320939 | -0.510180812 | -0.129307388 |
| ENSDART00000151390 | si:ch211-195b13.1 | 0.447846827 | 0.275025282 | 0.031728831 | 0.543393224 |
| ENSDART00000151393 | zak | 0.559981256 | 0.606971645 | 0.291742344 | 0.078029866 |
| ENSDART00000151399 | pak7 | -0.095169867 | -0.142909818 | -0.239638425 | 0.065715442 |
| ENSDART00000151404 | abat | -0.288767469 | -0.303619722 | -0.197451372 | -0.147078748 |
| ENSDART00000151419 | tgfa | -0.147139604 | -0.1316737 | -0.368157435 | -0.244071655 |
| ENSDART00000151423 | c1ql3b | -0.428342943 | -0.592751887 | -0.458841393 | -0.097254864 |
| ENSDART00000151448 | vamp4 | 0.36274105 | 0.703182259 | 0.708668654 | 0.04883985 |
| ENSDART00000151467 | slc44a2 | 1.227786397 | 2.405127088 | 2.406151956 | 1.882399418 |
| ENSDART00000151489 | LRRC2 | -0.220896871 | -0.332799828 | -0.436191804 | -0.38527929 |
| ENSDART00000151514 | wipi1 | -0.487231741 | -0.283410935 | -0.040753743 | -0.07881031 |
| ENSDART00000151547 | pnpla7a | 0.300123054 | 0.241347051 | 0.10241403 | -0.203931018 |
| ENSDART00000151550 | usp8 | 0.146844734 | 0.224801555 | 0.32026745 | 0.237867483 |
| ENSDART00000151567 | sgk494b | -0.371209064 | -0.282108451 | -0.446431854 | -0.117181288 |
| ENSDART00000151571 | ahdc1 | -0.753523402 | -0.774930454 | -0.265860844 | -0.025283973 |
| ENSDART00000151586 | ftr56 | 2.21645788 | 0.879044173 | 3.24618964 | 0.72203566 |

| | | | | | |
|---|---|---|---|---|---|
| ENSDART00000151611 | BX537277.4 | -0.440906854 | -0.477467469 | -0.367060961 | -0.356985579 |
| ENSDART00000151613 | lhfpl2a | 2.211015019 | 2.8976425 | 2.404140419 | 1.864383361 |
| ENSDART00000151627 | si:ch73-362m14.2 | -1.029772234 | -1.504293907 | -1.467613461 | -0.562650292 |
| ENSDART00000151634 | tnrc6c1 | -0.074330004 | -0.222991449 | -0.321878312 | -0.006293329 |
| ENSDART00000151648 | ntng2b | -0.44606387 | -0.406664301 | -0.175135105 | 0.111487092 |
| ENSDART00000151661 | wipf1b | 0.584481191 | 0.31541115 | 0.189596046 | -0.173471027 |
| ENSDART00000151670 | fbxl16 | -0.100117774 | -0.155204519 | -0.627708744 | -0.129502503 |
| ENSDART00000151672 | si:ch211-276a17.5 | -0.716987346 | -0.427751363 | -0.201345778 | -0.037618143 |
| ENSDART00000151674 | pcdh9 | -0.204552278 | -0.440082629 | -0.022873178 | 0.221939437 |
| ENSDART00000151679 | si:ch73-233k15.2 | 0.305634484 | 0.260016753 | 1.072404438 | -0.272816623 |
| ENSDART00000151685 | lhfpl4a | -0.450299252 | -0.419390296 | -0.093062661 | 0.17106537 |
| ENSDART00000151698 | syt3 | -0.446756628 | -0.536044497 | -0.128625894 | 0.106551311 |
| ENSDART00000151717 | march7 | 0.010098896 | 0.251138543 | 0.521642437 | 0.401235935 |
| ENSDART00000151718 | tns1a | -0.41721923 | -0.880763022 | -0.123281901 | 0.108356531 |
| ENSDART00000151739 | pax10 | -0.427677153 | -0.286784205 | -0.176549765 | -0.047893912 |
| ENSDART00000151743 | tmem237b | 0.052429896 | -0.216668621 | -0.455661391 | -0.118191943 |
| ENSDART00000151785 | klc4 | 0.265672724 | 0.701188452 | 0.757625768 | 0.469102543 |
| ENSDART00000151805 | vamp4 | 0.722926228 | 0.32988105 | 0.08556674 | 0.068052265 |
| ENSDART00000151808 | rcvrn3 | -0.028816037 | -0.156203513 | -0.580402071 | -0.415774036 |
| ENSDART00000151822 | tuba8l4 | 0.798817947 | 0.734886282 | 0.324378588 | 0.272246902 |
| ENSDART00000151842 | lrrn3a | -0.579197629 | -0.548770022 | -0.143313453 | 0.249550285 |
| ENSDART00000151853 | btbd10a | -0.220795221 | -0.048228521 | 0.169228662 | 0.338815973 |
| ENSDART00000151874 | gtf2h5 | -0.186830286 | -0.721090531 | -0.840818906 | -0.885273426 |
| ENSDART00000151878 | rab3gap2 | 0.026444427 | 0.181039642 | 0.242597677 | 0.247371128 |
| ENSDART00000151881 | rp1l1b | -0.468794546 | -0.617233135 | -0.603918539 | -0.270450371 |
| ENSDART00000151892 | BX248324.2 | 1.389473856 | 2.137590244 | 1.979774015 | 1.113747327 |
| ENSDART00000151899 | ldlrb | -0.013482381 | 0.739364147 | 1.30137615 | 0.56832803 |
| ENSDART00000151904 | foxn3 | -0.402174688 | -0.533042351 | -0.141780798 | 0.022724604 |
| ENSDART00000151915 | ptchd1 | -0.713611001 | -0.66822522 | 0.042208909 | 0.328774722 |
| ENSDART00000151920 | si:ch211-140m22.7 | -0.371308279 | -0.473110833 | -0.103450206 | 0.052557108 |
| ENSDART00000151921 | dnmt3ab | -0.131014471 | -0.104325638 | 0.10563751 | 0.340310238 |
| ENSDART00000151970 | si:ch211-248l17.3 | -0.372087768 | -0.337523565 | 0.171433041 | 0.720267147 |
| ENSDART00000151999 | ttc27 | 0.537657382 | 0.63628465 | 0.751878141 | 0.471568333 |
| ENSDART00000152000 | hectd1 | 2.43305833 | 3.547853034 | -1.454489893 | -0.017509384 |
| ENSDART00000152005 | ppp1r13ba | -0.032736325 | -0.117362615 | -0.454836475 | -0.009028338 |
| ENSDART00000152006 | si:ch211-236p5.3 | -1.010175109 | -0.375171908 | -0.903663548 | -0.553145976 |
| ENSDART00000152011 | DOCK4 (1 of many) | -0.107148853 | -0.150156526 | 0.321338768 | 0.540658105 |
| ENSDART00000152019 | si:ch211-234p6.5 | -0.203738294 | -0.079554548 | -0.244044589 | -0.027716123 |
| ENSDART00000152068 | AL590149.1 | -0.322191556 | -0.083241136 | -0.42500156 | -0.283301232 |
| ENSDART00000152069 | chst2b | -0.329630849 | -0.401663244 | -0.360860216 | -0.363893389 |
| ENSDART00000152122 | si:ch211-176g6.2 | 1.234754318 | 1.008584835 | 0.458934208 | -0.407921233 |
| ENSDART00000152128 | MAZ (1 of many) | -0.18246323 | -0.447481895 | -0.170629979 | -0.120255425 |
| ENSDART00000152145 | si:ch211-207i1.2 | -1.105453902 | -1.700204988 | -2.013950338 | -1.361589232 |
| ENSDART00000152152 | si:ch73-24k9.2 | 1.048518423 | 0.665615029 | 1.113264445 | -0.254081843 |
| ENSDART00000152172 | BX248418.1 | -0.267252802 | -0.476860817 | -0.659952516 | -0.030823789 |
| ENSDART00000152181 | mapta | -0.246945515 | 0.298203412 | 0.945051693 | 0.777575562 |
| ENSDART00000152195 | si:ch211-113a14.18 | 0.940862394 | 0.989824089 | 1.044244017 | 0.604507986 |
| ENSDART00000152240 | si:ch1073-174d20.2 | 0.11047675 | 0.254355415 | 0.357239942 | 0.147146143 |
| ENSDART00000152253 | kcnj13 | -0.175334575 | -0.22735068 | -0.506366898 | -0.552648836 |
| ENSDART00000152269 | syncrip | 0.263965684 | 0.255402715 | 0.319958135 | 0.269096167 |
| ENSDART00000152275 | BX908780.3 | -0.123347867 | -0.42152381 | -0.357247701 | -0.183442147 |

| | | | | | |
|---|---|---|---|---|---|
| ENSDART00000152277 | si:ch73-168d20.1 | 0.909762162 | 0.836471756 | 1.203009656 | 0.352187699 |
| ENSDART00000152284 | si:ch73-301j1.1 | -0.68804704 | -0.624114046 | 1.993753311 | -0.48476616 |
| ENSDART00000152295 | FAM83G | 1.150380037 | 1.206829236 | 1.079415548 | 0.591155843 |
| ENSDART00000152299 | si:dkey-25o16.4 | 1.175283487 | 1.143366275 | 0.842863117 | 0.368204805 |
| ENSDART00000152318 | si:ch211-286b5.2 | 1.976393579 | 3.213645724 | 2.557166791 | 1.560923939 |
| ENSDART00000152371 | usp2a | -0.286860533 | -0.506577502 | -0.337595735 | 0.001938619 |
| ENSDART00000152376 | CLSTN2 (1 of many) | -0.314646809 | -0.323760148 | -0.422303692 | -0.021838531 |
| ENSDART00000152378 | tgif1 | 0.194631245 | 0.419803811 | 0.14379228 | -0.205607529 |
| ENSDART00000152388 | CR354435.3 | 1.276581803 | 1.114721557 | 1.265047914 | 0.436655827 |
| ENSDART00000152400 | dbf4b | 0.645863235 | 0.809166025 | 0.868170031 | 0.806915507 |
| ENSDART00000152409 | mpp3b | -0.399859226 | -0.270912664 | -0.49906829 | -0.114613724 |
| ENSDART00000152428 | arhgap32a | -0.870901224 | -0.197122132 | -0.387452332 | -0.355800105 |
| ENSDART00000152429 | si:ch211-22i13.2 | -0.050442481 | -0.183983476 | -0.310965899 | -0.270852228 |
| ENSDART00000152431 | grid1b | -0.386738283 | -0.602313736 | -0.348261702 | 0.022251996 |
| ENSDART00000152432 | lratb.2 | 0.819163408 | 1.604350772 | -0.35089707 | -0.825333131 |
| ENSDART00000152478 | stambpl1 | 0.081881747 | 0.120249323 | 0.496702134 | 0.282826765 |
| ENSDART00000152479 | si:dkey-58j15.11 | 0.30322392 | 0.88586109 | 0.224237301 | -0.316952944 |
| ENSDART00000152489 | irf3 | -0.152502705 | 0.050924712 | 1.421424509 | -0.375867622 |
| ENSDART00000152510 | ccser2b | 0.726161978 | 0.836833026 | 1.118702985 | 0.610749876 |
| ENSDART00000152513 | arfip2a | -0.209731462 | -0.427433763 | -0.319460479 | -0.125960154 |
| ENSDART00000152520 | sh3bgr | -0.276595165 | -0.180114366 | -0.559026228 | -0.573707548 |
| ENSDART00000152531 | chrna10a | -0.39848157 | -0.348304481 | -0.425468269 | -0.274269467 |
| ENSDART00000152535 | CR846082.2 | -0.517845666 | -0.356698278 | -0.420655711 | -0.212748903 |
| ENSDART00000152556 | FAM13C | -0.458278075 | -0.375744323 | -0.341944162 | -0.232139437 |
| ENSDART00000152562 | eif3f | 0.257721359 | 0.386526337 | 0.235538886 | 0.038968621 |
| ENSDART00000152567 | frmpd2 | -0.319245931 | -0.373961894 | -0.203551264 | -0.037181194 |
| ENSDART00000152580 | nlgn4a | -0.383612645 | -0.391621152 | -0.368243921 | -0.24740854 |
| ENSDART00000152603 | egln2 | -0.344914112 | -0.184325703 | -0.029073268 | 0.070142585 |
| ENSDART00000152636 | clocka | 0.008631834 | 0.196030905 | 0.593132542 | 0.426491394 |
| ENSDART00000152641 | dnm3a | 1.808565272 | 2.497351665 | 2.011092537 | 0.57584064 |
| ENSDART00000152666 | slc25a23a | -0.288940584 | -0.374696227 | 0.31705403 | 0.139123262 |
| ENSDART00000152672 | zgc:152977 | -0.296261868 | -0.305183243 | -0.595435728 | -0.424752212 |
| ENSDART00000152678 | ifit10 | 0.50194473 | 0.092674262 | 4.442632616 | 1.169305685 |
| ENSDART00000152680 | pyyb | 2.62463081 | 4.023644413 | 3.815932945 | 2.663962921 |
| ENSDART00000152689 | oplah | -0.018001137 | -0.164313071 | -0.461321001 | -0.221934453 |
| ENSDART00000152699 | si:ch211-286b5.9 | 1.056007406 | 0.325301277 | 0.188178466 | -0.341462888 |
| ENSDART00000152703 | BX901920.1 | -0.098796288 | 0.57736287 | 0.482412674 | 0.325371495 |
| ENSDART00000152720 | MARK4 (1 of many) | -0.106851349 | -0.112526868 | -0.273643684 | -0.009380005 |
| ENSDART00000152766 | CR354435.4 | 0.814174745 | 0.989990518 | 0.663039704 | 0.339445782 |
| ENSDART00000152768 | si:ch211-207i1.2 | -0.201357893 | -0.51029626 | -0.665018804 | -0.121583536 |
| ENSDART00000152780 | tex2l | -0.378255597 | -0.430378354 | -0.394491857 | -0.23990429 |
| ENSDART00000152788 | mki67 | 1.022403776 | 0.934711217 | 0.634468641 | 0.246495311 |
| ENSDART00000152789 | etaa1 | 1.060104568 | 1.006586924 | 0.636303316 | 0.452429723 |
| ENSDART00000152844 | cdc42ep1a | -0.176432004 | -0.234011448 | -0.309447838 | -0.303047875 |
| ENSDART00000152855 | prdx1 | 2.014399342 | 2.912277881 | 2.532639813 | 2.043057891 |
| ENSDART00000152858 | rcan2 | -0.344037465 | -0.451039286 | -0.208970338 | -0.321499331 |
| ENSDART00000152879 | ablim2 | -0.320970088 | -0.240571783 | -0.134413966 | 0.04563774 |
| ENSDART00000152892 | slc16a8 | 2.182950408 | 0.789162193 | -0.768133939 | -1.779969995 |
| ENSDART00000152907 | si:ch1073-184j22.2 | 0.289101439 | 0.219661264 | 0.052983655 | 0.072807111 |
| ENSDART00000152924 | stxbp4 | 0.009805274 | -0.210366163 | 0.385226099 | 0.296155125 |
| ENSDART00000152936 | gabrz | -0.611503657 | -0.756211778 | -0.185809647 | 0.356886347 |

| | | | | | |
|---|---|---|---|---|---|
| ENSDART00000152942 | jcada | -0.053928395 | -0.076278472 | -0.332447962 | -0.296311136 |
| ENSDART00000152954 | rrm2 | 3.254915605 | 2.753529904 | 1.974728566 | 1.135298044 |
| ENSDART00000152965 | si:dkeyp-104f11.9 | -0.448853405 | -0.428655614 | -0.583739797 | -0.149977842 |
| ENSDART00000152968 | si:dkey-21c1.1 | 0.134479312 | -0.180316529 | -0.580606373 | -0.202289334 |
| ENSDART00000152973 | RNF157 | -0.237155894 | -0.35583425 | -0.110722293 | 0.097030733 |
| ENSDART00000153013 | eef1db | 0.150654853 | 0.305331563 | 0.633860114 | 0.458033815 |
| ENSDART00000153022 | AL935194.1 | -0.839883385 | -0.952071296 | -0.323659944 | -0.119202215 |
| ENSDART00000153086 | kctd17 | -0.109500243 | -0.169719356 | -0.312398256 | -0.16098927 |
| ENSDART00000153087 | bmf2 | 0.285623957 | 0.323465948 | 0.348774195 | 0.627801645 |
| ENSDART00000153124 | CU570682.1 | -0.274154453 | -0.462025745 | -0.49154679 | -0.26836126 |
| ENSDART00000153134 | mark3b | -0.033481129 | -0.166637764 | 0.542361776 | 0.482050901 |
| ENSDART00000153146 | rbfox3a | -0.546449898 | -0.43963695 | 0.174623158 | 0.484406984 |
| ENSDART00000153167 | hlfb | -0.470839666 | -0.326611575 | -0.235752692 | -0.264047019 |
| ENSDART00000153177 | mkl2a | -0.042089487 | -0.340254508 | -0.226976022 | -0.084520833 |
| ENSDART00000153187 | thrab | -0.362356661 | -0.551240408 | -0.403932333 | -0.192068079 |
| ENSDART00000153190 | vash2 | -0.078136066 | -0.226923996 | -0.395295378 | -0.160874636 |
| ENSDART00000153194 | si:dkey-38p12.3 | 0.25264745 | 0.485169484 | 0.572803819 | 0.07332987 |
| ENSDART00000153199 | afap1 | 0.840574953 | 1.106233667 | 1.438882155 | 1.005659187 |
| ENSDART00000153200 | si:ch211-194k22.8 | 0.210290351 | 0.451170205 | 0.37501933 | 0.225583201 |
| ENSDART00000153212 | adgra1a | 0.27394507 | 0.382231602 | 0.735064793 | 0.705504548 |
| ENSDART00000153217 | eef1db | 0.666551397 | 1.483640099 | 2.024242417 | 1.323561336 |
| ENSDART00000153225 | GUCY2C (1 of many) | 3.255008425 | 3.866902647 | 2.201523204 | 2.781639432 |
| ENSDART00000153227 | nfe2l1b | 0.688085235 | 1.301360444 | 1.138008541 | 0.729999096 |
| ENSDART00000153245 | stxbp4 | -0.144159334 | -0.445073444 | 0.187773192 | 0.216239919 |
| ENSDART00000153247 | si:ch211-120g10.1 | -0.520249419 | -0.414424265 | -0.454570577 | -0.107523144 |
| ENSDART00000153248 | chadlb | 1.217222776 | 0.847984599 | 0.639882396 | -0.038384888 |
| ENSDART00000153256 | cacna1g | -0.53708851 | -0.717133052 | -0.415346006 | -0.243050395 |
| ENSDART00000153284 | tbkbp1 | -0.504212532 | -0.489626186 | -0.240468452 | 0.072471561 |
| ENSDART00000153289 | nos1apb | -0.207119032 | 0.371227589 | 0.722590517 | 0.685079944 |
| ENSDART00000153296 | nsfb | 0.330739865 | 0.349896221 | -0.030123095 | -0.208569864 |
| ENSDART00000153306 | hcar1-4 | 0.883818254 | 1.413387657 | 1.160612678 | 0.305432292 |
| ENSDART00000153307 | si:ch73-344o19.1 | 0.586530995 | 0.605191095 | 0.75799445 | 0.479323327 |
| ENSDART00000153308 | dcun1d4 | -0.403892315 | -0.460237446 | -0.667559321 | -0.876458812 |
| ENSDART00000153311 | syndig1l | -0.402504141 | -0.415168554 | -0.17721556 | 0.113761296 |
| ENSDART00000153317 | abracl | 3.781753376 | 3.851435778 | 3.391861813 | 0.497752701 |
| ENSDART00000153334 | serinc1 | 0.574524524 | 0.605164909 | 0.10821359 | 0.438731193 |
| ENSDART00000153339 | adam17b | -0.104549731 | -0.262233446 | -0.335652873 | -0.112647107 |
| ENSDART00000153355 | grin2cb | -0.524459194 | -0.47344108 | -0.357891852 | 0.017542632 |
| ENSDART00000153375 | CU571074.1 | -0.192846631 | -0.331910339 | -0.069344033 | 0.142583214 |
| ENSDART00000153377 | eef1db | 0.533867077 | 0.950085432 | 1.385417409 | 1.0336151 |
| ENSDART00000153390 | gpam | -0.267998677 | -0.071527709 | 0.102371515 | 0.045213236 |
| ENSDART00000153391 | otofa | -0.790883428 | -1.357026899 | -0.581227909 | -0.190433713 |
| ENSDART00000153442 | rxfp2a | -0.191591927 | -0.398184153 | -0.707111514 | -0.312025768 |
| ENSDART00000153445 | exoc7 | 3.557809232 | 3.85646058 | 3.295092173 | 3.102150432 |
| ENSDART00000153452 | hsp90ab1 | 0.284157203 | 0.639252574 | 0.548907136 | 0.495783326 |
| ENSDART00000153491 | si:dkey-238d18.5 | 0.983293911 | 0.529719125 | 0.777356871 | -0.049086963 |
| ENSDART00000153510 | si:ch73-160l8.2 | 0.056068509 | 0.128148598 | 1.161251796 | 0.352706117 |
| ENSDART00000153514 | si:ch211-121j5.4 | -0.438676742 | -0.189457468 | -0.187818697 | -0.117536785 |
| ENSDART00000153562 | ppfia3 | -0.022654827 | 0.232304266 | 0.485406891 | 0.444415736 |
| ENSDART00000153570 | AL808129.1 | -0.250097653 | -0.308065347 | -0.243803188 | 0.04714878 |
| ENSDART00000153591 | si:dkey-56f14.7 | -0.35474859 | -0.1685547 | -0.219665386 | -0.050831688 |

| | | | | | |
|---|---|---|---|---|---|
| ENSDART00000153619 | thsd7ab | -0.196544647 | -0.395072863 | -0.388102838 | -0.013196959 |
| ENSDART00000153654 | piezo2a.2 | 0.718362747 | 0.545453706 | 0.495090248 | 0.182038228 |
| ENSDART00000153657 | trim3a | 0.068235605 | 0.356875561 | 0.585605002 | 0.467600874 |
| ENSDART00000153666 | AL732567.1 | 0.845331402 | 1.050170733 | 0.86587331 | 0.828206988 |
| ENSDART00000153673 | kcnb1 | -0.632196562 | -0.836186932 | -0.481100002 | -0.109364665 |
| ENSDART00000153700 | BX088707.1 | 0.029406694 | 0.3559423 | 0.787563577 | 0.447534651 |
| ENSDART00000153704 | anks1ab | -0.440221806 | -0.75898091 | -0.133742531 | 0.137005152 |
| ENSDART00000153730 | CU856222.1 | -0.553584022 | -0.481855955 | -0.007954528 | 0.343506731 |
| ENSDART00000153731 | si:dkey-79d12.5 | 1.619937595 | 1.685958359 | 0.880018827 | 0.052586274 |
| ENSDART00000153743 | rp1l1a | -0.216164399 | -0.344666036 | -0.324542341 | -0.024409204 |
| ENSDART00000153751 | rnf146 | 2.626965217 | 2.377858038 | 3.541283251 | 3.121825847 |
| ENSDART00000153773 | cipca | -0.044231438 | -0.31334692 | -0.44667945 | -0.17971329 |
| ENSDART00000153824 | napba | -0.634929885 | -0.324273151 | -0.096308603 | 0.10005339 |
| ENSDART00000153828 | atp2b3b | -0.474522045 | -0.66566516 | -0.285016921 | 0.075707306 |
| ENSDART00000153834 | sypl1 | 0.749942367 | 0.493446597 | 0.028135514 | -0.603815485 |
| ENSDART00000153841 | lmcd1 | 1.804357101 | 2.873087613 | 2.523453656 | 2.002290777 |
| ENSDART00000153847 | si:dkey-7i4.24 | 4.195698891 | 4.052966176 | 3.712547417 | 1.148430087 |
| ENSDART00000153897 | ndnfl | -0.781136652 | -1.071097643 | -0.053351937 | 0.577095222 |
| ENSDART00000153904 | si:dkey-108k21.7 | -0.206298886 | -0.326824637 | -0.265415499 | -0.00099298 |
| ENSDART00000153925 | fut9a | -0.379466906 | -0.312742341 | -0.333889875 | -0.027131911 |
| ENSDART00000153937 | prkd1 | -0.431650217 | -0.382419593 | -0.495097396 | -0.310742086 |
| ENSDART00000153944 | grm8a | -0.44814853 | -0.63071211 | -0.403185415 | 0.043396209 |
| ENSDART00000153950 | si:dkey-106n21.1 | 1.634202521 | 0.388901976 | 0.409369194 | 0.514423363 |
| ENSDART00000153951 | myo18ab | 0.15536019 | 0.13591756 | 0.249484283 | 0.108775298 |
| ENSDART00000153959 | pdcl | -0.115849049 | -0.178115822 | -0.269080801 | -0.124407546 |
| ENSDART00000153968 | apba2a | -0.094717367 | 0.193224107 | 0.542818077 | 0.184287162 |
| ENSDART00000153990 | CR848791.2 | 3.062868799 | 2.217074852 | 2.048805323 | 1.916392428 |
| ENSDART00000153992 | AL844141.1 | 0.512134791 | 0.207788846 | -0.514504406 | -0.130928693 |
| ENSDART00000154039 | cntnap2a | -0.318660467 | -0.524836596 | -0.23230439 | 0.054653452 |
| ENSDART00000154045 | glg1a | 0.211468857 | 0.293330382 | 0.40088818 | 0.351496464 |
| ENSDART00000154066 | isg20 | 0.340483696 | 0.28200181 | 0.116846361 | -0.121466791 |
| ENSDART00000154077 | arhgap29a | -0.13604217 | -0.133932896 | -0.339961978 | -0.305768852 |
| ENSDART00000154085 | gipr | -0.077819916 | -0.236822644 | -0.445528895 | -0.377678023 |
| ENSDART00000154109 | tp53inp2 | 0.254956915 | 0.876540848 | 0.694070175 | 0.684041374 |
| ENSDART00000154132 | nptx2a | -0.534082272 | -0.549346998 | -0.399917367 | -0.21269489 |
| ENSDART00000154141 | BX649398.2 | 0.278828233 | 0.21540553 | -0.861481785 | -0.785329833 |
| ENSDART00000154148 | camk1a | -0.513931344 | -0.665527159 | -0.499415187 | -0.020106333 |
| ENSDART00000154187 | fat3a | -0.16861177 | -0.487078475 | -0.285722941 | -0.100061338 |
| ENSDART00000154212 | BX323824.1 | 0.386508588 | 0.636151192 | 0.398585293 | 0.423381311 |
| ENSDART00000154217 | tapbpl | -0.790387635 | -1.110046917 | -0.234167162 | 0.243016032 |
| ENSDART00000154228 | sgip1a | -0.210460002 | -0.370824353 | -0.364544583 | -0.177769156 |
| ENSDART00000154238 | AL928908.3 | -0.284715118 | -0.465774511 | -0.246230328 | -0.209644064 |
| ENSDART00000154253 | VAMP1 (1 of many) | -0.590038106 | -0.991557836 | -0.153916319 | 0.292977881 |
| ENSDART00000154256 | gabbr1a | -0.701390177 | -1.072254559 | -0.049084095 | 0.380533583 |
| ENSDART00000154282 | CT737162.2 | 2.40869789 | 3.049681827 | 2.839672212 | 2.555397461 |
| ENSDART00000154295 | RPS17 (1 of many) | 0.397556507 | 0.446477566 | 0.193480989 | 0.136840294 |
| ENSDART00000154307 | hsp90aa1.2 | -0.273233427 | -0.388821318 | -0.732231951 | -0.738083556 |
| ENSDART00000154314 | prph | 1.502464936 | 3.204214666 | 3.264208069 | 2.453842104 |
| ENSDART00000154316 | birc6-as2 | -0.437572334 | -0.469057905 | -0.275420253 | -0.034926922 |
| ENSDART00000154321 | BX908750.1 | 0.337703917 | 0.710327992 | 0.54373662 | 0.188876538 |
| ENSDART00000154324 | CR848791.3 | 3.13610848 | 2.257927159 | 1.961382093 | 1.531269093 |

| | | | | | |
|---|---|---|---|---|---|
| ENSDART00000154329 | plppr3b | -0.697577837 | -0.505300644 | -0.575363423 | -0.078498654 |
| ENSDART00000154333 | camkva | -0.731914874 | -0.620456244 | -0.460647205 | -0.026116956 |
| ENSDART00000154359 | ppil1 | 0.446724137 | 0.384000004 | 0.201188176 | -0.044104474 |
| ENSDART00000154363 | chst7 | -0.08877204 | -0.106063572 | -0.492144158 | -0.182223467 |
| ENSDART00000154375 | CR812464.1 | -0.471069407 | -0.288977412 | -0.485957331 | -0.299311124 |
| ENSDART00000154377 | HIST1H2BM (1 of many) | 0.871205458 | 1.144896653 | 0.89956001 | 0.544514919 |
| ENSDART00000154385 | nhsl1a | -0.136848286 | -0.0387183 | -0.42116689 | -0.266423729 |
| ENSDART00000154393 | helz2 | -0.348999688 | -0.710690348 | 2.383802665 | -0.188541475 |
| ENSDART00000154400 | tmem121a | -0.218470186 | -0.361690646 | -0.304266051 | -0.077189842 |
| ENSDART00000154405 | rac1b | 0.253264939 | 0.144566871 | 0.063714363 | 0.127687084 |
| ENSDART00000154418 | relb | -0.043359819 | 0.083083005 | 0.26390054 | -0.071618881 |
| ENSDART00000154429 | gnai2b | -0.171950228 | -0.218087414 | 0.308490155 | 0.376283024 |
| ENSDART00000154434 | frmd4ba | 0.373950657 | 0.549595304 | 0.324851185 | -0.019839546 |
| ENSDART00000154437 | dyrk1ab | -0.405599591 | -0.293606249 | -0.153631672 | -0.200586701 |
| ENSDART00000154438 | ltb4r | 0.406290343 | 0.081638761 | 0.101793584 | -0.225121243 |
| ENSDART00000154460 | FP243382.1 | 2.715231463 | 2.420271029 | 2.173535574 | 1.735635064 |
| ENSDART00000154495 | si:ch73-195i19.3 | 0.285619034 | 0.153921873 | 0.078118908 | 0.066673852 |
| ENSDART00000154501 | cntnap1 | -0.715173489 | -0.84270145 | -0.359147123 | 0.122603648 |
| ENSDART00000154503 | syncripl | 0.163572635 | 0.201195284 | 0.244411469 | 0.118655043 |
| ENSDART00000154519 | nrxn3a | -0.161675514 | -0.445079711 | -0.329497924 | -0.050382407 |
| ENSDART00000154539 | slc17a7b | -0.194590302 | -0.417792811 | -0.270877884 | 0.095165091 |
| ENSDART00000154543 | prelid3a | 0.578261204 | 0.829307037 | 0.712329147 | 0.291766611 |
| ENSDART00000154573 | prob1 | 1.935993422 | 2.091368151 | 1.752154811 | 1.442205958 |
| ENSDART00000154627 | eogt | -0.136366673 | -0.382886458 | -0.560103122 | -0.203176718 |
| ENSDART00000154628 | fam101b | 0.450957894 | 0.982590486 | 0.845377619 | -0.490650185 |
| ENSDART00000154633 | chst15 | -0.234696766 | -0.27064598 | -0.214752082 | -0.155581795 |
| ENSDART00000154638 | kif26ab | 2.382609513 | 1.668373417 | 1.704066313 | 1.906262158 |
| ENSDART00000154652 | irgq1 | 0.050480218 | 0.069496236 | 1.163101864 | 0.075649547 |
| ENSDART00000154667 | SORCS3 | -0.864119649 | -0.84034251 | -0.370097899 | 0.056888282 |
| ENSDART00000154679 | ypel2b | -0.575722509 | -0.671947843 | -0.397167717 | -0.03983803 |
| ENSDART00000154682 | CR847531.1 | -0.541034447 | -0.676553431 | -0.554824044 | -0.149530967 |
| ENSDART00000154719 | hip1 | 0.155954348 | 0.357624902 | 0.536540352 | 0.421858349 |
| ENSDART00000154726 | MYADM (1 of many) | 1.393641215 | 1.121324137 | 1.000193747 | 0.38217935 |
| ENSDART00000154730 | tsc22d1 | 0.125146937 | 0.071380598 | 0.346511952 | 0.250641998 |
| ENSDART00000154732 | ptprt | -0.128635692 | -0.441586772 | -0.224155866 | -0.045463061 |
| ENSDART00000154748 | sv2bb | -0.593359268 | -0.537099772 | -0.285465428 | 0.087161334 |
| ENSDART00000154754 | grik1b | -0.541985174 | -0.220416154 | -0.114773141 | -0.128231092 |
| ENSDART00000154760 | CU302436.3 | -0.931829111 | -0.859047685 | -0.631190961 | -0.184130931 |
| ENSDART00000154778 | map1ab | -0.607759134 | -0.711103734 | -0.339973576 | -0.303457691 |
| ENSDART00000154783 | BX511311.5 | -0.420893047 | -0.406058694 | -0.342057398 | -0.24848029 |
| ENSDART00000154801 | itgae.2 | 0.922252153 | 0.655667188 | 0.379341087 | -0.144710616 |
| ENSDART00000154809 | CU467861.1 | 0.653813027 | 0.40784225 | 0.192026805 | 0.151191997 |
| ENSDART00000154811 | shank1 | -0.390024258 | -0.720748027 | -0.113172669 | 0.256359072 |
| ENSDART00000154846 | CR381618.1 | 0.115176903 | 0.900181702 | 1.102625706 | 0.253393379 |
| ENSDART00000154851 | zgc:153405 | 0.317950752 | 1.076463864 | 0.674378682 | 0.394781222 |
| ENSDART00000154860 | CU682780.1 | -0.134139035 | -0.255412373 | -0.351732915 | -0.064350576 |
| ENSDART00000154869 | cacna1ha | -0.561980799 | -0.554587419 | -0.200655166 | 0.115415453 |
| ENSDART00000154880 | arhgef33 | 0.636994605 | 0.687196577 | 0.646579762 | 0.300121216 |
| ENSDART00000154885 | si:ch211-167j9.4 | 1.022951831 | 0.721088624 | 0.864353518 | 0.647190994 |
| ENSDART00000154897 | srsf10a | -0.118782385 | -0.22214156 | -0.484269556 | -0.079154163 |
| ENSDART00000154916 | p4htm | -0.322717782 | -0.242652877 | -0.246255125 | -0.141036056 |

| | | | | | |
|---|---|---|---|---|---|
| ENSDART00000154917 | ciartb | -0.07665894 | -0.17887759 | -0.516654864 | -0.219843882 |
| ENSDART00000154933 | si:dkey-13p1.4 | -2.565488296 | -3.472379023 | -0.405040918 | 0.505631615 |
| ENSDART00000154945 | si:dkey-7j14.6 | 0.563979024 | 0.172625977 | 0.010511104 | -0.696238166 |
| ENSDART00000154961 | btbd10b | -0.035347687 | 0.05017631 | 0.315377826 | 0.177817418 |
| ENSDART00000154978 | si:ch211-284f22.3 | 0.111486547 | 0.17394927 | 0.442535968 | 0.352663532 |
| ENSDART00000154985 | ddx3b | -0.099315614 | -0.086363684 | 0.465639279 | 0.388612698 |
| ENSDART00000154991 | b4galnt1b | -0.249570746 | -0.126095261 | 0.190575038 | 0.316262718 |
| ENSDART00000155033 | srpk2 | -0.213188431 | 0.160547857 | 0.269227711 | 0.471530406 |
| ENSDART00000155038 | si:dkeyp-100a5.4 | -0.19105809 | -0.704501455 | -0.315530312 | -0.360932845 |
| ENSDART00000155051 | mych | 1.81452306 | 1.847457308 | 1.423042212 | 0.692601562 |
| ENSDART00000155057 | capns1b | 0.743122931 | 1.233401578 | 0.848245138 | 0.67484109 |
| ENSDART00000155066 | atf5b | 0.303223783 | 0.449779577 | 0.245269838 | -0.118029433 |
| ENSDART00000155078 | CU466278.1 | -0.496260032 | -0.275828398 | -0.937323619 | -0.494833793 |
| ENSDART00000155088 | ankrd33ab | -0.091123111 | -0.280595773 | -2.122583018 | -0.908625438 |
| ENSDART00000155094 | si:ch211-286c4.6 | 0.931010056 | 1.14507062 | 1.165043802 | 1.805632584 |
| ENSDART00000155108 | si:ch211-153l6.6 | -0.396073461 | -0.324628235 | -0.330118035 | -0.333690214 |
| ENSDART00000155116 | im:7152348 | -0.17038115 | -0.180140684 | -0.567414199 | -0.647722466 |
| ENSDART00000155128 | meis3 | 1.719560674 | 1.869294835 | 2.789896441 | 2.118152659 |
| ENSDART00000155178 | man1b1b | -0.197923135 | -0.298790741 | -0.488552512 | -0.408314648 |
| ENSDART00000155184 | lcorl | -0.199231763 | -0.444480785 | -0.032192793 | -0.024389475 |
| ENSDART00000155188 | ccnjl | 0.284030133 | 0.391162848 | 0.492680219 | 0.242842213 |
| ENSDART00000155190 | maptb | 1.096239137 | 1.804532446 | 2.232737802 | 1.717568221 |
| ENSDART00000155203 | ttbk1b | -0.886183357 | -0.409260354 | 0.093887166 | 0.234308172 |
| ENSDART00000155212 | msi2b | -0.193481816 | -0.280091063 | -0.40298969 | -0.141186615 |
| ENSDART00000155227 | si:dkey-248g15.2 | -0.512231666 | -0.490597761 | -0.50523481 | -0.410257249 |
| ENSDART00000155229 | rxfp3.3b | -0.899836922 | -0.961252446 | -0.631060044 | -0.38629797 |
| ENSDART00000155243 | BX571762.2 | -0.58394173 | -0.66558831 | -0.494788274 | -0.139509269 |
| ENSDART00000155256 | VWA5A (1 of many) | 0.207030506 | 0.487713239 | 0.641785484 | 0.122484219 |
| ENSDART00000155262 | nek1 | -0.289218155 | -0.162294169 | 0.032473831 | 0.21974529 |
| ENSDART00000155267 | si:ch211-149e23.4 | 0.305598533 | 0.378888393 | 0.64366915 | 0.422691793 |
| ENSDART00000155268 | pik3cb | 0.672881324 | 0.55380807 | 0.246761254 | 0.040522215 |
| ENSDART00000155273 | FP102016.1 | 0.672121642 | 0.238521805 | 0.194526989 | 0.041260629 |
| ENSDART00000155280 | GDPGP1 | 0.576074599 | 0.656458452 | 0.674683139 | 0.299654628 |
| ENSDART00000155306 | CR925728.1 | 1.81197689 | 3.280738703 | 3.024711765 | 2.003163607 |
| ENSDART00000155313 | hspa4l | -0.318813042 | -0.280574999 | -0.269018374 | 0.033454147 |
| ENSDART00000155320 | atxn7l1 | -0.301256075 | -0.435278991 | -0.214418032 | -0.09758885 |
| ENSDART00000155324 | pdpk1a | -0.082713691 | -0.135052421 | -0.317218269 | -0.180842854 |
| ENSDART00000155327 | ip6k2b | 0.635904907 | 0.734729077 | 0.476402387 | 0.497162385 |
| ENSDART00000155344 | VWA5A (1 of many) | 0.009320645 | 0.148549206 | 0.649252702 | 0.190646512 |
| ENSDART00000155346 | tbc1d24 | -0.444989153 | -0.328605075 | -0.340554332 | -0.035933185 |
| ENSDART00000155368 | si:ch73-95l15.3 | 1.516126674 | 2.634881995 | 2.849448463 | 2.215584039 |
| ENSDART00000155397 | soat2 | -0.618559497 | -0.31486995 | -0.194806138 | -0.125519338 |
| ENSDART00000155427 | reep6 | 0.890805905 | 0.284631883 | 0.369468446 | 0.541597773 |
| ENSDART00000155440 | eif3ba | 0.265595321 | 0.439061322 | 0.461693724 | 0.229141991 |
| ENSDART00000155445 | si:dkey-7j14.6 | 4.555762034 | 3.357060696 | 4.122496376 | 1.366436667 |
| ENSDART00000155456 | gria4a | -0.214950555 | -0.367205609 | -0.037830298 | 0.237179158 |
| ENSDART00000155458 | ubap1la | -0.102679722 | -0.709773263 | -0.512828077 | -0.333708251 |
| ENSDART00000155464 | ftr73 | 0.436406361 | 0.364310996 | 2.049125852 | 0.40110848 |
| ENSDART00000155472 | nudt8 | -0.430833003 | -0.309991274 | -0.295400218 | -0.125942696 |
| ENSDART00000155480 | ptprnb | -0.501074747 | -0.734898939 | -0.55867421 | -0.198202123 |
| ENSDART00000155494 | CR384099.1 | -0.702495416 | -0.755760921 | -0.229440848 | 0.245420937 |

| | | | | | |
|---|---|---|---|---|---|
| ENSDART00000155496 | gal3st2 | -0.347287801 | -0.358994611 | -0.502209455 | 0.011977936 |
| ENSDART00000155499 | CU571319.1 | 0.914674175 | 1.459254054 | 0.740447631 | 0.310619615 |
| ENSDART00000155502 | sez6b | -0.382545228 | -0.653854879 | -0.149982953 | 0.168281376 |
| ENSDART00000155519 | si:ch211-139d20.3 | 0.778333948 | 0.572606648 | 0.403486279 | 0.486914049 |
| ENSDART00000155525 | si:rp71-45k5.4 | 0.233215264 | 0.362150294 | 0.25819678 | 0.006995576 |
| ENSDART00000155537 | VWA5A (1 of many) | 1.206080865 | 0.987004598 | 1.27402251 | 0.722325665 |
| ENSDART00000155539 | CR786577.1 | -0.296712547 | -0.710798753 | -0.341045728 | 0.032686751 |
| ENSDART00000155553 | CR931780.1 | 1.845884645 | 1.39773807 | 0.761518659 | 0.588325484 |
| ENSDART00000155554 | frs3 | -0.288569277 | -0.348098053 | 0.092392931 | 0.139305366 |
| ENSDART00000155563 | abca5 | -0.240041791 | -0.35443471 | -0.098648169 | 0.068480985 |
| ENSDART00000155580 | ccdc32 | -0.137851793 | -0.182759184 | -0.331717766 | -0.143672128 |
| ENSDART00000155592 | pcdh17 | -0.283490769 | -0.422406754 | -0.268768577 | 0.186259856 |
| ENSDART00000155595 | zgc:174263 | -0.246339018 | -0.142647436 | -0.250696029 | -0.144411513 |
| ENSDART00000155610 | tdg.2 | 0.198987795 | 0.279928785 | 0.39077816 | 0.364649362 |
| ENSDART00000155658 | rapgefl1 | -0.418362292 | -0.397338899 | 0.000660896 | -0.001869225 |
| ENSDART00000155666 | znf576.1 | -0.372721752 | -0.23410815 | -0.134923201 | -0.013577883 |
| ENSDART00000155683 | CU855758.2 | -0.197771803 | -0.264936874 | -0.616643588 | -0.19367265 |
| ENSDART00000155723 | si:ch211-24o10.6 | -0.255651916 | 0.580162906 | 3.993132482 | 0.654326546 |
| ENSDART00000155751 | nuak1b | 0.785416985 | 1.722603254 | 1.683891996 | 1.302083807 |
| ENSDART00000155755 | CU469362.1 | 0.467928162 | 0.778645014 | 0.342312822 | -0.038725012 |
| ENSDART00000155757 | si:dkey-16p21.8 | -0.383053569 | -0.315569666 | -0.222717784 | 0.028079211 |
| ENSDART00000155764 | slc25a18 | -0.081957182 | -0.293577862 | -0.279616954 | -0.236786045 |
| ENSDART00000155786 | adgrl1a | -0.190147203 | -0.479741899 | -0.292479007 | 0.056973675 |
| ENSDART00000155808 | SRMS | -0.518504687 | -0.21831974 | 0.182924623 | 0.347888554 |
| ENSDART00000155817 | spsb3b | 0.397854128 | -0.261249457 | 0.795238945 | 1.076488289 |
| ENSDART00000155821 | kcnt2 | -1.064051569 | -1.353278896 | -1.209879321 | -0.444718597 |
| ENSDART00000155838 | luzp1 | -0.38633981 | -0.211709262 | -0.146984907 | -0.314685818 |
| ENSDART00000155839 | CR759918.1 | -0.263859971 | -0.262088445 | -0.379170984 | -0.118091172 |
| ENSDART00000155840 | nyap2a | -0.526663005 | -0.654892136 | 0.27374848 | 0.170536216 |
| ENSDART00000155843 | cntnap2a | -0.263075507 | -0.343897302 | 0.676718804 | 0.99202798 |
| ENSDART00000155844 | nr1d4b | 0.850133899 | 0.016728696 | -0.171313869 | 0.165802614 |
| ENSDART00000155848 | CU929046.2 | -0.462340311 | -0.419607593 | -0.082173547 | -0.018364502 |
| ENSDART00000155853 | gprc5ba | -0.568480744 | -0.680176078 | -0.476445621 | -0.089942946 |
| ENSDART00000155859 | larp4ab | -0.193577441 | -0.330790279 | -0.534105504 | -0.201385609 |
| ENSDART00000155864 | CU929070.1 | 1.274116455 | 1.096698631 | 0.870265354 | 0.298589514 |
| ENSDART00000155865 | tmtops2a | -0.317973372 | -0.37404155 | -0.204729003 | -0.178609254 |
| ENSDART00000155866 | zbtb16b | -0.434457287 | -0.595844526 | 0.044569606 | 0.015262284 |
| ENSDART00000155902 | frya | -0.208619052 | -0.429125005 | -0.269458135 | -0.004719121 |
| ENSDART00000155935 | clstn3 | -0.324952015 | -0.256429293 | 0.203826088 | 0.321978764 |
| ENSDART00000155947 | | -0.00158363 | 0.212586024 | 0.278540905 | 0.169021899 |
| ENSDART00000155949 | grm8b | -0.426657348 | -0.505637682 | -0.163630603 | 0.069819734 |
| ENSDART00000155956 | ssx2ipa | -0.036908945 | -0.252732501 | -0.388773092 | -0.156731785 |
| ENSDART00000155973 | si:ch211-282b22.1 | 0.150167669 | -0.093228424 | -0.31353999 | -0.332567216 |
| ENSDART00000155992 | tjp1b | -0.305505503 | -0.430213518 | -0.153805225 | 0.023043416 |
| ENSDART00000156002 | tmem206 | 0.793441462 | 0.286165316 | 0.167291274 | -0.130249565 |
| ENSDART00000156008 | adcy1b | -0.126277791 | -0.148680925 | -0.963041725 | -0.495906031 |
| ENSDART00000156012 | si:dkey-35i13.1 | -1.182557557 | -0.949918392 | -0.338171836 | -0.056213778 |
| ENSDART00000156019 | phlpp1 | -0.378155747 | -0.257656641 | -0.150573304 | -0.126584813 |
| ENSDART00000156034 | VWA5A (1 of many) | 1.04711946 | 1.090403168 | 1.260861512 | 0.694741705 |
| ENSDART00000156038 | epn2 | 0.165862475 | 0.434471463 | 0.333526121 | 0.113707323 |
| ENSDART00000156051 | dhx32b | -0.034036848 | -0.386771049 | -0.600165189 | -0.131769991 |

| | | | | | |
|---|---|---|---|---|---|
| ENSDART00000156062 | calml4b | -0.12658352 | -0.200610116 | -0.571249563 | -0.361192591 |
| ENSDART00000156167 | si:ch73-182a11.2 | 0.438099394 | 0.613388756 | 0.677790276 | 0.474745988 |
| ENSDART00000156169 | ube2o | -0.089939623 | -0.17956897 | -0.346977584 | -0.218736801 |
| ENSDART00000156179 | sema7a | -0.273960356 | -0.559537035 | -0.479585038 | -0.478067901 |
| ENSDART00000156211 | adcyap1r1b | -0.120918148 | -0.186017 | -0.540439336 | -0.277725276 |
| ENSDART00000156224 | dhx33 | 0.472474316 | 0.556602191 | 0.639276654 | 0.492917502 |
| ENSDART00000156247 | CU468915.1 | -0.147323171 | -0.162953088 | -0.662599642 | -0.420048082 |
| ENSDART00000156251 | ehbp1 | 0.342339175 | 0.353753636 | 0.38894242 | 0.04618254 |
| ENSDART00000156255 | ago1 | 0.402935641 | 0.576022371 | 0.656640241 | 0.478601065 |
| ENSDART00000156256 | angpt4 | -0.447666401 | -0.531623063 | -0.584960614 | -0.899124282 |
| ENSDART00000156257 | wnk1a | -0.012583832 | 0.164942132 | 0.328941429 | 0.287930853 |
| ENSDART00000156258 | pfkpb | -0.354296164 | -0.444812712 | -0.357490287 | -0.175370256 |
| ENSDART00000156280 | si:ch211-195b15.7 | -0.343611395 | -0.333351498 | -0.20091334 | -0.160599804 |
| ENSDART00000156295 | myhb | -2.03199835 | 4.211906798 | 4.339398917 | 2.940685436 |
| ENSDART00000156306 | ptgir | 0.787352004 | 0.130407388 | 0.033259328 | -0.478389399 |
| ENSDART00000156340 | si:ch211-167b20.8 | -0.291865267 | -0.289158159 | -0.304691219 | -0.111151074 |
| ENSDART00000156341 | fam117ba | -0.500099563 | -0.265235629 | 0.202099115 | 0.368226992 |
| ENSDART00000156351 | zfat | -0.386268069 | -0.362611932 | -0.387180541 | -0.301395116 |
| ENSDART00000156365 | BX248121.1 | -0.069755444 | -0.2828198 | -0.35442143 | -0.173325265 |
| ENSDART00000156375 | kcnq2b | -0.284091354 | -0.197722227 | 0.27148365 | 0.512045099 |
| ENSDART00000156380 | CR848757.1 | -0.431298815 | -0.377014166 | 0.404962464 | 0.865655337 |
| ENSDART00000156393 | AL772314.1 | -0.272422966 | -0.157258449 | 0.218420805 | 0.368190657 |
| ENSDART00000156403 | gfpt1 | 0.402370916 | 0.732684838 | 0.696897001 | 0.375214811 |
| ENSDART00000156411 | fam98a | 0.369288002 | 0.399859986 | 0.477248543 | 0.233728964 |
| ENSDART00000156415 | dock9b | 0.187525252 | 0.556466903 | 0.665071238 | 0.341730571 |
| ENSDART00000156418 | itpk1b | -0.567310725 | -0.6808899 | -0.20043499 | -0.215676742 |
| ENSDART00000156428 | hephl1a | 0.427057323 | 0.215282086 | 0.277986668 | -0.033310258 |
| ENSDART00000156437 | BX248501.1 | -0.044614101 | -0.492506227 | -0.236687215 | -0.012438349 |
| ENSDART00000156438 | tmem138 | -0.131418974 | -0.291816206 | -0.603920899 | -0.471900931 |
| ENSDART00000156447 | ap2a1 | -0.168026562 | -0.400633541 | -0.149036642 | -0.01932092 |
| ENSDART00000156450 | wnk4b | 0.377998578 | 0.587708612 | 1.132797895 | 0.554283794 |
| ENSDART00000156473 | CR627483.1 | -0.446874417 | -0.944979531 | -0.551426346 | -0.288797786 |
| ENSDART00000156492 | tiam2a | -0.024981814 | -0.236489206 | -0.397107093 | -0.120407246 |
| ENSDART00000156499 | prkcdb | 0.617876805 | 0.366145269 | 0.057590279 | -0.230105501 |
| ENSDART00000156509 | fam150bb | -0.200699582 | -0.03489316 | -0.310837396 | -0.503245808 |
| ENSDART00000156516 | tmem74b | -0.490393884 | -0.38109679 | -0.101020414 | 0.194211555 |
| ENSDART00000156517 | GSE1 | -0.323310165 | -0.383945273 | -0.036135566 | -0.046797605 |
| ENSDART00000156527 | crhr1 | -0.121975919 | -0.246139891 | -0.376184792 | -0.120567191 |
| ENSDART00000156535 | vav3b | -0.155549402 | 0.298871549 | 0.871724785 | 0.959064908 |
| ENSDART00000156546 | plcxd1 | 0.33889602 | 0.161133476 | 0.572020307 | 0.368311625 |
| ENSDART00000156568 | CR387996.1 | -0.565799723 | -0.534585499 | -0.855400153 | -0.72349809 |
| ENSDART00000156570 | slc6a17 | -0.448593137 | -0.394227085 | -0.032159276 | 0.074756396 |
| ENSDART00000156574 | si:ch211-270g19.5 | -0.160521866 | -0.305818991 | -0.352315858 | -0.223463705 |
| ENSDART00000156575 | dicp3.3 | 1.874714519 | 0.882148407 | 0.706359774 | -0.079481712 |
| ENSDART00000156577 | si:ch211-130m23.5 | -0.007378589 | 0.400588111 | 0.725075783 | 0.606240178 |
| ENSDART00000156580 | hspa12a | -0.664613601 | -0.584311718 | 0.126225265 | 0.380065995 |
| ENSDART00000156584 | msi2b | -0.183156983 | -0.402159154 | -0.705145609 | -0.389308076 |
| ENSDART00000156607 | CT737162.1 | 0.778225783 | 0.597499699 | 0.53495134 | 0.647915122 |
| ENSDART00000156608 | ap2a1 | -0.16265507 | -0.313644183 | -0.332880329 | -0.023511806 |
| ENSDART00000156613 | CR848683.1 | -0.419259508 | -0.300093589 | -0.534359803 | -0.243066507 |
| ENSDART00000156615 | amigo3 | -0.174881747 | -0.348495554 | -0.328942663 | 0.002458748 |

| | | | | | |
|---|---|---|---|---|---|
| ENSDART00000156668 | BX569793.1 | -0.298302127 | -0.566890135 | -0.349641739 | -0.271668844 |
| ENSDART00000156690 | ftr88 | 0.478641753 | 0.356185618 | 1.950902073 | 0.248066147 |
| ENSDART00000156705 | si:dkeyp-72g9.4 | -0.743143803 | -0.697928004 | -0.454613917 | -0.077239927 |
| ENSDART00000156741 | BX663503.2 | -0.41351136 | -0.477289266 | -0.358517546 | -0.050787842 |
| ENSDART00000156757 | CU633486.1 | -0.302968456 | -0.206059 | -0.531920994 | -0.165463059 |
| ENSDART00000156760 | grik4 | -0.347451056 | -0.634709172 | -0.373537369 | -0.211041818 |
| ENSDART00000156765 | soga3a | 0.515035881 | 0.806381678 | 1.137120698 | 0.812578505 |
| ENSDART00000156774 | dph6 | 0.485952337 | 0.860504803 | 0.608054536 | 0.313680056 |
| ENSDART00000156792 | kif1aa | -0.016449727 | 0.304055121 | 0.524314954 | 0.462526752 |
| ENSDART00000156795 | zcchc8 | 0.52372258 | 0.428501893 | 0.558201403 | 0.582679631 |
| ENSDART00000156806 | rps6kl1 | 0.03495508 | 0.325476415 | 0.37135553 | 0.249571068 |
| ENSDART00000156820 | si:dkey-7j14.5 | -0.971869146 | -0.757334904 | 0.08249331 | 0.800607437 |
| ENSDART00000156822 | cdip1 | -0.168452851 | -0.279910749 | -0.292336463 | -0.062208262 |
| ENSDART00000156831 | si:dkeyp-47f9.4 | -0.215799622 | -0.290204625 | -0.428110244 | -0.249285093 |
| ENSDART00000156832 | kif13ba | 0.259733256 | 0.195728022 | 0.255457371 | 0.044129384 |
| ENSDART00000156853 | si:ch211-284e13.14 | 0.249769449 | 0.06508988 | 0.403860054 | 0.279589235 |
| ENSDART00000156864 | CR376745.2 | -0.465037351 | -0.777375633 | -0.748299595 | -0.606506499 |
| ENSDART00000156877 | prss56 | -1.154037343 | -0.806544973 | -0.688753663 | -0.465249106 |
| ENSDART00000156885 | EML5 | -0.115526296 | -0.315705485 | -0.106202597 | 0.094534493 |
| ENSDART00000156913 | BX927184.1 | 1.282107066 | 1.77927465 | 1.802027966 | 1.376990986 |
| ENSDART00000156918 | si:dkey-16p21.8 | -0.278111877 | -0.271868386 | -0.501824009 | -0.172182623 |
| ENSDART00000156923 | plekhd1 | -0.640317774 | -0.396125765 | -0.428766409 | -0.43244988 |
| ENSDART00000156928 | nrxn3a | 0.020925857 | -0.341808434 | -0.239366202 | 0.052782357 |
| ENSDART00000156935 | pbx4 | 0.960146191 | 0.964981474 | 3.716226941 | 3.606625876 |
| ENSDART00000156965 | osbpl5 | -0.452639629 | -0.228065482 | 0.296781507 | 0.346469129 |
| ENSDART00000156967 | itcha | -0.048958922 | -0.233534262 | -0.334119802 | -0.057332431 |
| ENSDART00000156970 | si:ch73-265h17.5 | 1.250341969 | 1.481314496 | 1.76830523 | 0.8438116 |
| ENSDART00000156972 | slc12a7a | 0.533547881 | 0.750475895 | 0.543271513 | 0.05077876 |
| ENSDART00000156982 | si:ch211-195b15.8 | -0.203622983 | -0.158924419 | -0.050797883 | 0.351311882 |
| ENSDART00000156995 | sorl1 | -0.129978292 | -0.268802979 | -0.097227509 | 0.102143372 |
| ENSDART00000157018 | stmn3 | -1.008205118 | -1.406964646 | -0.116902659 | 0.621837811 |
| ENSDART00000157043 | rtn4r | -0.358605066 | -0.3593825 | -0.41712234 | -0.136427457 |
| ENSDART00000157045 | CU861453.1 | -0.184297698 | -0.238197074 | -0.240198755 | -0.095792528 |
| ENSDART00000157050 | thoc5 | -3.378273157 | -0.016679798 | 0.227931101 | 0.074505779 |
| ENSDART00000157058 | si:dkey-276j7.1 | -0.114061291 | 0.35744712 | 0.794710779 | 0.605587625 |
| ENSDART00000157066 | ppp1r16b | -0.047627869 | -0.115324195 | -0.408403631 | -0.014573035 |
| ENSDART00000157067 | BX510324.1 | -0.110028536 | -3.040813753 | -0.681076994 | -0.348601244 |
| ENSDART00000157116 | cyp8b3 | -4.272126467 | -1.470280769 | 0.08019487 | -1.373424813 |
| ENSDART00000157122 | elmsan1b | 0.072375655 | 0.138875939 | -0.375191445 | -0.292342976 |
| ENSDART00000157124 | CT033825.1 | -0.560424235 | -0.809708864 | -0.680777647 | -0.144372819 |
| ENSDART00000157125 | vsnl1b | -0.832964446 | -0.803942924 | -0.29707501 | 0.109485311 |
| ENSDART00000157127 | lmcd1 | 1.160892936 | 2.271204052 | 2.063825472 | 1.282095985 |
| ENSDART00000157129 | slc12a5a | -0.482400538 | -0.74419089 | -0.401518291 | -0.07447909 |
| ENSDART00000157137 | HIST1H2BM (1 of many) | 0.980872833 | 1.453339934 | 1.015669985 | 0.930663607 |
| ENSDART00000157139 | tmbim1a | 0.963253296 | 1.499596205 | 1.306783479 | 0.549513539 |
| ENSDART00000157171 | si:ch211-168d23.3 | -0.276296251 | -0.552181073 | -0.115547916 | -0.061680907 |
| ENSDART00000157190 | ostn | 0.371505716 | 0.628806712 | 0.229636498 | 0.167623814 |
| ENSDART00000157195 | ncam2 | -0.590792178 | -0.719154845 | -0.40764277 | 0.002619526 |
| ENSDART00000157199 | mkl2b | 0.238133435 | 0.28120222 | 0.324920076 | 0.149313819 |
| ENSDART00000157237 | si:dkey-56f14.7 | -0.377561364 | -0.203140811 | -0.202521092 | -0.065412084 |
| ENSDART00000157258 | BX004991.1 | -0.288834986 | -0.59132286 | -0.471054454 | -0.432666578 |

| | | | | | |
|---|---|---|---|---|---|
| ENSDART00000157264 | RAP1GAP (1 of many) | -0.397821605 | -0.344805586 | -0.195900343 | -0.114570879 |
| ENSDART00000157265 | trim13 | -0.831738752 | -0.32341984 | -0.102064486 | -0.071474505 |
| ENSDART00000157302 | si:dkey-7j14.5 | -0.795010515 | -0.583416344 | 0.327377123 | 0.951280694 |
| ENSDART00000157330 | necab3 | -0.311593638 | -0.396628677 | -0.274835563 | -0.023399638 |
| ENSDART00000157338 | gstz1 | -0.391223456 | -0.14748942 | -0.590184545 | -0.319150237 |
| ENSDART00000157350 | CT573344.1 | 0.592860846 | 1.013860007 | 1.275401288 | 0.850112465 |
| ENSDART00000157378 | hdac5 | -0.360172826 | -0.414916304 | -0.488703112 | -0.152605137 |
| ENSDART00000157407 | nacad | -0.349281215 | 0.532726398 | 0.824860187 | 0.457523633 |
| ENSDART00000157414 | plxna4 | -0.256858488 | -0.391015088 | -0.285048615 | -0.106219383 |
| ENSDART00000157415 | acvr2ab | -0.152487638 | -0.319222405 | -0.146638904 | -0.027512421 |
| ENSDART00000157416 | si:dkey-23n7.10 | 0.164308792 | 0.543803953 | 1.528490128 | 1.278412625 |
| ENSDART00000157428 | pik3r3b | -0.522784766 | -0.44851499 | -0.152608244 | 0.028231585 |
| ENSDART00000157437 | park2 | -0.135617767 | -0.306295495 | -0.445890787 | -0.195890902 |
| ENSDART00000157441 | filip1l | 1.025025125 | 2.361056221 | 1.959563135 | 1.214723721 |
| ENSDART00000157449 | clstn2 | -0.339354151 | -0.60478304 | 0.06475081 | 0.303731917 |
| ENSDART00000157454 | ncam1a | -0.363879899 | -0.114400367 | 0.050687773 | 0.064981174 |
| ENSDART00000157464 | adkb | 0.004906369 | 0.081747651 | 0.332869381 | 0.17633498 |
| ENSDART00000157475 | coro7 | 0.174942149 | 0.053392758 | 0.483358888 | 0.70362148 |
| ENSDART00000157479 | nr4a1 | -1.499367308 | -4.285014891 | -2.179617587 | -2.01791928 |
| ENSDART00000157487 | tfec | 0.534318298 | 0.423012313 | 0.205933849 | -0.313187686 |
| ENSDART00000157515 | FO704641.1 | -0.135830256 | -0.41280018 | -0.451790561 | -0.015210245 |
| ENSDART00000157518 | hspa12a | -0.767789618 | -0.701014156 | 0.093464144 | 0.364648116 |
| ENSDART00000157535 | unc13c | -0.230182256 | -0.566319347 | -0.291866986 | -0.102048139 |
| ENSDART00000157538 | npm3 | 0.706470742 | 0.616423608 | 0.411598819 | -0.103962898 |
| ENSDART00000157546 | | -4.297830881 | -2.011048515 | -0.106888147 | -0.169461052 |
| ENSDART00000157559 | si:dkey-183n20.15 | 2.324560868 | 2.818869593 | 3.029863009 | 3.039600704 |
| ENSDART00000157563 | noctb | -0.197344301 | -0.486083216 | -0.268092319 | 0.128030661 |
| ENSDART00000157570 | asmt | -0.467907647 | -0.512776687 | -0.801096178 | -0.257600065 |
| ENSDART00000157581 | dctn2 | 0.246810516 | 0.408963324 | 0.394068831 | 0.222953154 |
| ENSDART00000157591 | CT583642.1 | 0.946563077 | 0.733732925 | 0.580155557 | -0.143517637 |
| ENSDART00000157597 | Metazoa_SRP | 3.784983496 | 4.068975573 | 4.489534416 | 3.689887905 |
| ENSDART00000157604 | si:dkey-19c16.12 | 0.518457933 | 0.406892634 | 0.690003002 | 0.445775315 |
| ENSDART00000157625 | mapkapk3 | 0.008165693 | -0.129188005 | -0.419061818 | -0.281468347 |
| ENSDART00000157636 | Metazoa_SRP | 2.626880551 | 2.386640401 | 1.529298522 | 0.72995056 |
| ENSDART00000157643 | si:dkey-88e18.8 | 0.426205845 | 0.069942358 | 0.168991341 | 0.195596151 |
| ENSDART00000157659 | camta1b | -0.574265168 | -0.690891494 | -0.316064045 | -0.339000781 |
| ENSDART00000157661 | pdlim5a | -0.256112858 | -0.374018749 | -0.32171501 | -0.127659508 |
| ENSDART00000157663 | reln | -0.012850615 | -0.065342924 | 0.75286937 | 0.74278543 |
| ENSDART00000157678 | nop53 | 0.221649254 | 0.357738319 | 0.319808875 | 0.087032288 |
| ENSDART00000157682 | kat2a | 0.094563424 | 0.264392397 | 0.483214757 | 0.33498107 |
| ENSDART00000157697 | sec22bb | -0.209339329 | -0.24617122 | -0.297558263 | -0.212223069 |
| ENSDART00000157711 | ap2m1b | -0.212778909 | -0.504780227 | -0.48236783 | -0.089136332 |
| ENSDART00000157726 | fam20a | 0.506263561 | 0.217850689 | -0.118035628 | -0.139373167 |
| ENSDART00000157728 | gdap1l1 | -0.078507744 | 0.185505427 | 0.335339207 | 0.32984222 |
| ENSDART00000157735 | cdc42ep4a | 0.06354285 | 0.460607642 | 0.576300047 | 0.396817817 |
| ENSDART00000157755 | cds1 | -0.167833858 | -0.281185983 | -0.514312915 | -0.176231692 |
| ENSDART00000157769 | pcdh1gc5 | -0.403524411 | -0.226721593 | 0.040583925 | 0.029815654 |
| ENSDART00000157771 | sumo2b | 0.072691072 | 0.240484774 | 0.205807834 | 0.084424554 |
| ENSDART00000157775 | CR381700.1 | 0.888966397 | 0.292933465 | 0.058270274 | 0.51311949 |
| ENSDART00000157788 | CABZ01031894.1 | -0.75935498 | -0.746140313 | -0.275189537 | -0.221209558 |
| ENSDART00000157815 | elmo1 | -0.27246831 | -0.132899842 | -0.049572106 | 0.087486375 |

| | | | | | |
|---|---|---|---|---|---|
| ENSDART00000157817 | lrrfip1b | -0.412751624 | 0.086992217 | 0.168101768 | 0.241775802 |
| ENSDART00000157820 | mfsd6a | 0.403998194 | 0.775291086 | 0.690391907 | -0.020489453 |
| ENSDART00000157824 | slc6a15 | 0.370043272 | 0.932121448 | 1.025205824 | 0.470213629 |
| ENSDART00000157826 | mpp1 | 0.283246867 | 0.228442011 | 0.159131554 | 0.043281733 |
| ENSDART00000157852 | crtac1b | -0.401370563 | -0.59197436 | -0.358347223 | -0.176621049 |
| ENSDART00000157855 | ebf3a | -0.765871102 | -0.436598509 | 0.580406091 | 0.942511549 |
| ENSDART00000157871 | mgst3a | 0.496607086 | 0.574379254 | 0.622130807 | 0.504216816 |
| ENSDART00000157873 | hacd1 | 0.841608261 | 0.46729923 | 0.326975211 | 0.16703571 |
| ENSDART00000157877 | CABZ01092282.1 | -0.281901821 | -0.460522749 | 0.165109465 | 0.263636132 |
| ENSDART00000157890 | tcf7l1b | -4.555123239 | -4.673271453 | -1.829433189 | -0.732249154 |
| ENSDART00000157894 | apaf1 | 0.69901776 | 0.613857472 | 0.804406673 | 0.575412751 |
| ENSDART00000157899 | dad1 | 0.443099375 | 0.390242441 | 0.192203328 | -0.014224626 |
| ENSDART00000157900 | chmp1b | 0.92760321 | 0.512046614 | 0.591242771 | 0.272231533 |
| ENSDART00000157907 | rpl34 | 0.216382556 | 0.50724745 | 0.251806906 | 0.167915072 |
| ENSDART00000157919 | med24 | 0.290079054 | 0.266724064 | 0.247847379 | 0.11623123 |
| ENSDART00000157920 | BX664622.3 | 0.309964523 | 0.2387983 | 0.211520656 | 0.43648414 |
| ENSDART00000157937 | supt3h | 0.425159053 | 0.217206475 | 0.313741902 | -0.075091294 |
| ENSDART00000157943 | TENM2 | -0.301363214 | -0.846881931 | -0.344978545 | 0.012452 |
| ENSDART00000157944 | CR848788.1 | 0.572199002 | 0.285271614 | 0.143233659 | 0.192859067 |
| ENSDART00000157958 | nav3 | 0.289857115 | 0.341415382 | 0.725222646 | 0.845169813 |
| ENSDART00000157976 | hif1al2 | 0.549135047 | 0.585055165 | 1.624530088 | 0.686809008 |
| ENSDART00000157979 | plppr5b | -0.141986763 | -0.474240998 | -0.437275413 | -0.217953336 |
| ENSDART00000157998 | cngb3.1 | 0.075294731 | -0.249736043 | -0.588641013 | -0.198039383 |
| ENSDART00000158016 | flnca | 5.435651542 | 4.853809922 | 4.737861308 | 3.940836376 |
| ENSDART00000158017 | GUCA1A (1 of many) | -1.245659292 | -0.578230856 | -1.064205686 | -1.242533296 |
| ENSDART00000158021 | | -0.273860764 | -0.2373593 | -0.233720455 | -0.186840477 |
| ENSDART00000158036 | cript | -0.080102736 | -0.07683769 | -0.280131139 | -0.196435314 |
| ENSDART00000158042 | mdga2a | -0.261819905 | -0.306879785 | -0.151540127 | 0.022043669 |
| ENSDART00000158052 | rnf130 | -0.205769239 | 0.102328211 | 0.30768262 | 0.182366646 |
| ENSDART00000158055 | kcnh6a | -0.288242124 | -0.323669014 | -0.559331748 | -0.259833446 |
| ENSDART00000158072 | gpc2 | 0.147140386 | 0.761040446 | 1.023252339 | 0.612693915 |
| ENSDART00000158077 | arr3b | -0.13602706 | -0.148118881 | -0.527427286 | -0.215489656 |
| ENSDART00000158098 | gpr132b | -0.335128962 | -0.097182083 | 0.890467313 | 0.848088338 |
| ENSDART00000158123 | magi1b | -0.699965885 | -0.960349882 | -0.3166938 | 0.663491004 |
| ENSDART00000158147 | calr3b | 0.659493637 | 0.343153236 | 0.419640601 | 0.12486596 |
| ENSDART00000158154 | dlg2 | -0.258379843 | -0.32848175 | -0.404306744 | -0.064632311 |
| ENSDART00000158169 | cacna2d2a | -0.273970943 | -0.601526504 | -0.288906256 | -0.138856816 |
| ENSDART00000158183 | | -0.077181373 | -0.541229915 | -0.5215371 | -0.066571696 |
| ENSDART00000158202 | CABZ01078449.1 | -0.388078647 | -0.185889564 | -2.923902226 | 0.109460724 |
| ENSDART00000158204 | ntrk3a | -0.448230073 | -0.669954401 | -0.450935682 | -0.114414424 |
| ENSDART00000158212 | zgc:92161 | 0.66498591 | 0.968065208 | 0.600651955 | -0.278084019 |
| ENSDART00000158213 | akap12b | -1.00724688 | -1.146388756 | -0.599808188 | 0.009805796 |
| ENSDART00000158245 | si:ch73-1a9.3 | -0.234948586 | -0.211979293 | -0.351222857 | -0.141776696 |
| ENSDART00000158259 | lrtm1 | -0.264679565 | -0.523836662 | -0.478235847 | -0.361630227 |
| ENSDART00000158260 | c1galt1a | 0.184684125 | 0.138397385 | -0.286164045 | -0.509177074 |
| ENSDART00000158272 | dclk1b | 0.134331666 | 0.528656936 | 0.571389243 | 0.299537528 |
| ENSDART00000158289 | taok1b | -0.136932712 | -0.168799291 | -0.390684145 | -0.122927658 |
| ENSDART00000158290 | rpl24 | 0.677207663 | 0.41367136 | 0.306049933 | 0.275882655 |
| ENSDART00000158294 | ppp2r2cb | -0.427479199 | -0.698226267 | -0.466339082 | -0.260646218 |
| ENSDART00000158301 | CR749168.4 | -0.284512048 | -0.514337485 | -0.519239658 | -0.283197555 |
| ENSDART00000158310 | sat2b | -0.409697091 | -0.182271077 | -0.610869777 | -0.543847015 |

| | | | | | |
|---|---|---|---|---|---|
| ENSDART00000158311 | klhl7 | 0.128707755 | 0.106332897 | 0.351073118 | 0.278667069 |
| ENSDART00000158338 | CNNM1 | -0.362878072 | -0.490740729 | -0.646651234 | -0.29213891 |
| ENSDART00000158344 | apbb2b | -0.312721901 | -0.306004033 | 0.116860588 | 0.307202209 |
| ENSDART00000158348 | zmiz1b | 0.167782576 | 0.103403196 | 0.408056208 | 0.722421278 |
| ENSDART00000158354 | adarb2 | -0.457769313 | -0.257870403 | -0.020766954 | -0.013045223 |
| ENSDART00000158355 | bsk146 | -0.973617487 | -0.7163723 | -0.116416923 | 0.102394026 |
| ENSDART00000158358 | stk40 | -0.305579471 | -0.394234001 | -0.179882029 | -0.355392322 |
| ENSDART00000158364 | si:dkey-112e17.1 | -0.140789793 | -0.187774176 | -0.188421319 | -0.472049113 |
| ENSDART00000158373 | epc1b | 0.106120055 | 0.059882061 | 0.624967071 | 0.299679159 |
| ENSDART00000158380 | si:dkey-35m8.1 | -0.462314219 | -0.595244504 | 0.018350172 | 0.199682571 |
| ENSDART00000158395 | dclk2a | -0.152411223 | -0.291700953 | -0.710292221 | -0.218291454 |
| ENSDART00000158429 | tmem42b | -0.433937172 | -0.428108528 | -0.048957595 | 0.127616146 |
| ENSDART00000158436 | lypla1 | 0.03667991 | -0.201169357 | -0.480610097 | -0.044568257 |
| ENSDART00000158439 | ece2b | -0.235035459 | -0.256980434 | -0.233206993 | -0.167380402 |
| ENSDART00000158440 | arhgap12b | -0.039244537 | -0.138433678 | -0.355110347 | -0.091493676 |
| ENSDART00000158466 | creb3l2 | 0.073267533 | 0.091531466 | -0.07087733 | -0.32121815 |
| ENSDART00000158489 | zgc:77058 | -0.495737975 | -0.228595493 | 0.141443601 | 0.440592752 |
| ENSDART00000158500 | atp8b2 | -0.533001796 | -0.553651375 | -0.17260282 | 0.179842571 |
| ENSDART00000158515 | | -0.601694587 | -0.246037227 | -0.439227399 | -0.290329932 |
| ENSDART00000158532 | gnb2 | -0.185937539 | 0.268112673 | 0.644184639 | 0.701859814 |
| ENSDART00000158533 | nup50 | 0.24455835 | 0.550142856 | 0.78129311 | 0.523300254 |
| ENSDART00000158540 | PRKCA (1 of many) | -0.338994629 | -0.305121271 | -0.237610526 | -0.160203254 |
| ENSDART00000158555 | camsap2a | -0.296024961 | -0.235576155 | 0.04836376 | 0.162312505 |
| ENSDART00000158563 | wu:fj29h11 | 0.059907274 | -0.166860874 | 1.78519307 | 0.017403627 |
| ENSDART00000158564 | CU681842.1 | -0.1327123 | -0.276414972 | -0.289623815 | -0.090007903 |
| ENSDART00000158581 | spna2 | 0.480311119 | 0.377153274 | 0.267336877 | -0.085977531 |
| ENSDART00000158587 | casd1 | -0.21622109 | -0.133405356 | -0.572868553 | -0.255662771 |
| ENSDART00000158598 | si:ch211-232b12.5 | -0.304573383 | 0.212713124 | 0.452361247 | 0.038090441 |
| ENSDART00000158618 | cpne4b | -1.041271467 | -0.75854166 | 0.438341768 | 1.045239509 |
| ENSDART00000158646 | dpp6a | -0.408023938 | -0.433201282 | -0.219888215 | -0.008677852 |
| ENSDART00000158671 | sult4a1 | -0.462609111 | -0.368663113 | -0.083169387 | 0.19862288 |
| ENSDART00000158686 | si:dkey-14k9.2 | -0.49897987 | -0.263068549 | 0.75542148 | 1.074830065 |
| ENSDART00000158692 | CABZ01112401.1 | 0.113542035 | -0.344191992 | -0.461207823 | -0.256682679 |
| ENSDART00000158696 | mhc1zda | -0.851268531 | -0.246667696 | -0.324999925 | -0.385360573 |
| ENSDART00000158721 | | 1.164230248 | 1.13549021 | 1.093110731 | 1.576844966 |
| ENSDART00000158723 | sncb | -0.257429657 | -0.430720868 | -0.069889176 | 0.08182323 |
| ENSDART00000158733 | si:dkey-33i11.1 | 1.813729642 | 1.289849367 | 0.876112899 | 0.279282769 |
| ENSDART00000158734 | CABZ01088933.1 | -0.385251814 | -0.457284358 | 0.104668245 | 0.198079674 |
| ENSDART00000158739 | Metazoa_SRP | 1.431390172 | 1.797550169 | 1.401344471 | 0.830250287 |
| ENSDART00000158743 | zgc:113348 | 0.011068258 | 0.221603341 | 0.278365427 | 0.216955383 |
| ENSDART00000158744 | BX927336.3 | 4.021248005 | 3.307131569 | 3.405842268 | 3.528289574 |
| ENSDART00000158766 | CR848025.1 | -0.596230295 | -0.433122151 | -0.989806185 | -0.729213035 |
| ENSDART00000158777 | cirbpa | 0.225987764 | 0.314210398 | 0.241597292 | 0.01082888 |
| ENSDART00000158780 | mapk8ip1a | 0.350641915 | 0.585959364 | 0.286490226 | -0.139620155 |
| ENSDART00000158808 | cnn3a | 0.586268263 | 0.320248819 | -0.106889335 | -0.483977637 |
| ENSDART00000158810 | fkbp3 | -0.588686525 | -0.386876086 | -0.522309858 | -0.719247198 |
| ENSDART00000158817 | letm1 | -0.280849194 | -0.370052113 | -0.172062949 | -0.075482878 |
| ENSDART00000158820 | golga7bb | -0.441549718 | -0.650879189 | -0.491036037 | -0.164497612 |
| ENSDART00000158825 | actc1b | 0.015935591 | 2.858633574 | 3.729694641 | 2.583924646 |
| ENSDART00000158826 | CABZ01058650.1 | 0.277651195 | 0.407733859 | 0.060586967 | -0.150969128 |
| ENSDART00000158837 | dync1h1 | 0.179298196 | 0.363338197 | 0.829945857 | 0.598133568 |

| | | | | | |
|---|---|---|---|---|---|
| ENSDART00000158854 | tpd52l1 | -0.063123021 | -0.257733521 | -0.65813611 | -0.260747428 |
| ENSDART00000158917 | abhd10a | -0.199951388 | -0.281143798 | -0.472212357 | -0.246558895 |
| ENSDART00000158919 | lpar1 | -0.789836287 | -0.638879508 | -0.710773837 | -0.465534721 |
| ENSDART00000158936 | grin1a | -0.976292061 | -1.003442352 | -0.317903979 | 0.15163936 |
| ENSDART00000158962 | dpf2 | 0.023344289 | -0.160855631 | -0.651177712 | -0.029134577 |
| ENSDART00000158977 | si:ch73-266f23.1 | 0.491629493 | 0.41114477 | 0.471295394 | 0.297543095 |
| ENSDART00000159004 | si:ch73-233f7.8 | 0.057778797 | 0.28613349 | 0.517419704 | 0.568556694 |
| ENSDART00000159016 | psmd1 | 0.220320535 | 0.390334714 | 0.382386975 | 0.213567973 |
| ENSDART00000159024 | btr29 | 2.93548853 | 3.916483422 | 3.440636421 | 1.361130093 |
| ENSDART00000159034 | rsu1 | 0.959727483 | 0.959427671 | 0.950095712 | 0.424675541 |
| ENSDART00000159061 | si:ch211-185a18.2 | -0.270017947 | -0.418794122 | -0.535787315 | -0.21407671 |
| ENSDART00000159062 | paqr7a | -0.415028281 | -0.250472931 | -0.476345463 | -0.551526055 |
| ENSDART00000159087 | drd2b | -0.654775306 | -0.766659414 | -0.766987138 | -0.309487495 |
| ENSDART00000159103 | nkd3 | 3.293495606 | 1.842810424 | 2.875712797 | 0.7112483 |
| ENSDART00000159104 | atp1b2a | -0.488650552 | -0.417675156 | -0.129746035 | 0.084213141 |
| ENSDART00000159112 | si:ch211-285j22.3 | -0.41628689 | -0.545704947 | -0.599123468 | -0.263727805 |
| ENSDART00000159123 | btbd17b | -0.172708316 | -0.362184438 | -0.675325658 | -0.245697841 |
| ENSDART00000159142 | si:dkey-56m15.2 | 0.312405009 | 0.342224899 | 0.497281083 | 0.253644828 |
| ENSDART00000159160 | impdh1b | 0.016028158 | -0.054860403 | 0.596376969 | 0.34908306 |
| ENSDART00000159163 | c2cd2l | -0.400916824 | -0.28913261 | -0.285175907 | -0.160469096 |
| ENSDART00000159165 | b3galt2 | -0.421223634 | -0.395995108 | -0.436113716 | -0.390524412 |
| ENSDART00000159177 | si:dkey-61n16.5 | -0.004555446 | 0.268605493 | -0.72825811 | -0.687838681 |
| ENSDART00000159178 | hspa8 | -1.810725395 | -0.905134344 | -6.819365873 | -1.947204217 |
| ENSDART00000159184 | erc2 | 0.268311207 | -0.09915171 | 0.379637611 | 0.748745363 |
| ENSDART00000159246 | ctsa | 0.554249571 | 0.498653237 | 0.219459997 | 0.055975716 |
| ENSDART00000159249 | oxr1a | -0.748854688 | -0.518408458 | -0.301236362 | -0.153948362 |
| ENSDART00000159252 | abca3b | 0.181429492 | 0.428459166 | 0.640212151 | 0.464148107 |
| ENSDART00000159256 | kif5c | 0.534952361 | 0.603334839 | 0.411520272 | 0.273079865 |
| ENSDART00000159259 | capn1b | 0.6323058 | 1.402210713 | 1.384902028 | 1.097440242 |
| ENSDART00000159274 | sf3b3 | 0.246778314 | 0.098942964 | 0.083691016 | -0.027346389 |
| ENSDART00000159291 | podxl2 | -0.45367951 | -0.207714244 | 0.333702352 | 0.497299671 |
| ENSDART00000159296 | si:ch73-138e16.4 | -0.371496737 | -0.627266426 | -0.52708555 | -0.509641971 |
| ENSDART00000159300 | mcm7 | 0.714704002 | 0.831682391 | 0.874291792 | 0.339164006 |
| ENSDART00000159312 | pim1 | 0.541604371 | 0.710042231 | 0.310719343 | -0.194529837 |
| ENSDART00000159316 | mvda | 0.177821409 | 1.524901044 | 1.830059782 | 0.927790622 |
| ENSDART00000159328 | FO704782.1 | 0.142074859 | 0.061082305 | 0.804157236 | 0.053378695 |
| ENSDART00000159330 | pcyox1 | 0.406833689 | 0.21789945 | 0.044242143 | -0.158070725 |
| ENSDART00000159340 | ctnnd1 | 0.771872624 | 0.982959313 | 0.944642837 | 0.226155974 |
| ENSDART00000159368 | myo19 | -0.061571264 | -0.562765185 | -0.263758975 | -0.061266965 |
| ENSDART00000159372 | eif3ha | 0.499618724 | 0.599836819 | 0.638052337 | 0.263915251 |
| ENSDART00000159390 | FO704846.1 | -0.36021367 | -0.403087033 | 0.444915596 | 0.998151126 |
| ENSDART00000159409 | unc45a | 0.063679601 | 0.06008134 | 0.330631234 | 0.129605203 |
| ENSDART00000159433 | BX571691.1 | 2.919338685 | 1.742268402 | 2.797274384 | 2.742997875 |
| ENSDART00000159434 | tmem254 | 0.529951039 | 0.26685766 | 0.074245289 | -0.042910999 |
| ENSDART00000159435 | pcdh2g28 | 0.000816386 | 0.116171 | 0.335030381 | 0.350715644 |
| ENSDART00000159438 | zgc:162730 | 0.775020432 | 0.781916155 | 0.788293058 | 0.232021834 |
| ENSDART00000159450 | ctnnd1 | 0.19135693 | 0.714895836 | -0.026058241 | 0.026164146 |
| ENSDART00000159454 | CABZ01044277.1 | 0.109016927 | 0.066686129 | 0.335359626 | 0.279394938 |
| ENSDART00000159466 | FUT9 (1 of many) | -0.32488192 | -0.508163292 | -0.746293394 | -0.756603469 |
| ENSDART00000159486 | anapc11 | -0.082411238 | -0.111278818 | -0.383958603 | -0.347049397 |
| ENSDART00000159491 | ddx46 | 0.204487031 | 0.180157737 | 0.416223492 | 0.263114106 |

| | | | | | |
|---|---|---|---|---|---|
| ENSDART00000159493 | baiap2a | -0.408759075 | -0.584635172 | -0.205977447 | -0.074404323 |
| ENSDART00000159519 | SIKE1 | -0.137869817 | -0.17784406 | -0.297014412 | -0.12625892 |
| ENSDART00000159520 | dusp1 | 1.00607749 | 1.105172193 | 0.739133795 | 1.131411014 |
| ENSDART00000159523 | BX248501.1 | -0.802081931 | -1.281225474 | -1.178391388 | -0.841643515 |
| ENSDART00000159534 | leng9 | 0.34368633 | 0.401471128 | 0.132926162 | 0.109702136 |
| ENSDART00000159543 | si:ch211-284e13.6 | -2.69508904 | -0.200744398 | -0.117258809 | -0.389858773 |
| ENSDART00000159581 | CT583723.1 | -0.203564175 | -0.159816156 | -0.458224986 | -0.16588667 |
| ENSDART00000159588 | rnf24 | -0.199572807 | -0.336306858 | -0.290748993 | -0.299485771 |
| ENSDART00000159589 | rtn4rl1b | -0.618605611 | -0.334861623 | -0.03219779 | 0.419833076 |
| ENSDART00000159601 | serpini1 | -0.44709139 | -0.357709199 | -0.198408132 | 0.076823185 |
| ENSDART00000159604 | mbd3b | -0.404570692 | -0.312781624 | -0.318828454 | -0.248466452 |
| ENSDART00000159608 | fstl5 | -0.296329015 | -0.522282128 | -0.291629941 | -0.144161607 |
| ENSDART00000159614 | r3hdm1 | -0.174685122 | -0.341121434 | 0.099641284 | 0.042466817 |
| ENSDART00000159620 | | 0.106577262 | 0.152035698 | 0.604289635 | 0.634717201 |
| ENSDART00000159627 | scn4ba | -1.651719386 | -1.747876787 | -0.730948262 | -0.207716251 |
| ENSDART00000159629 | mcm2 | 0.995796878 | 0.693727511 | 0.639304212 | 0.157223466 |
| ENSDART00000159634 | b3glcta | -0.092602693 | -0.194400961 | -0.44155461 | -0.530900199 |
| ENSDART00000159642 | DPP10 | -0.369468467 | -0.43082527 | -0.361060788 | -0.270120898 |
| ENSDART00000159652 | sox4b | 0.767109202 | 1.080655119 | 1.389017231 | 0.976257694 |
| ENSDART00000159654 | CR847844.2 | 0.025394661 | -0.340096605 | -0.548288987 | -0.346027376 |
| ENSDART00000159670 | abtb1 | -0.10462458 | -0.155472174 | -0.372376774 | -0.233981977 |
| ENSDART00000159673 | dync1i2b | 0.025252757 | 0.176262587 | 0.413723725 | 0.300516643 |
| ENSDART00000159690 | gpr75 | -0.483974147 | -0.377125661 | 0.037666989 | 0.123067934 |
| ENSDART00000159693 | NYAP1 | -0.548984567 | -0.516611795 | -0.021366781 | 0.000605996 |
| ENSDART00000159697 | si:rp71-5o12.3 | 0.531788358 | 0.397969004 | 0.570994446 | 0.299640856 |
| ENSDART00000159716 | si:ch73-236c18.6 | 0.962463693 | 0.259951761 | 0.066014607 | 0.162933451 |
| ENSDART00000159725 | mtm1 | -0.155968464 | -0.279489824 | -0.277642503 | -0.186816665 |
| ENSDART00000159727 | pcdh11 | -0.294947297 | -0.365638864 | -0.089140913 | 0.17419873 |
| ENSDART00000159732 | si:dkey-203a12.8 | 3.015691266 | 3.637172924 | 3.104587293 | 1.713663998 |
| ENSDART00000159733 | CABZ01063757.1 | 0.129910694 | -0.203491407 | -0.855433405 | -0.227320323 |
| ENSDART00000159745 | PCP4L1 (1 of many) | -0.632453626 | -0.700162017 | -0.354659005 | -0.133179733 |
| ENSDART00000159747 | rab11fip4b | -0.416792935 | -0.701212044 | -0.335764895 | -0.067364991 |
| ENSDART00000159752 | sort1a | -0.095510148 | -0.060577332 | -0.362899297 | -0.357946523 |
| ENSDART00000159769 | pam | -0.468861179 | -0.403502349 | -0.282541701 | -0.241769832 |
| ENSDART00000159782 | agtpbp1 | -0.044706055 | 0.173624177 | 0.473061984 | 0.407831162 |
| ENSDART00000159805 | CABZ01085924.1 | -0.392684277 | -0.456709342 | 0.004929566 | 0.18758642 |
| ENSDART00000159845 | epb41a | -0.285946023 | -0.193154687 | -0.17771991 | -0.091337383 |
| ENSDART00000159853 | FQ311890.1 | -0.276177481 | -0.432407927 | -0.032078728 | 0.27665545 |
| ENSDART00000159859 | tmem132e | -0.259691326 | -0.228995571 | -0.113717547 | -0.037626683 |
| ENSDART00000159871 | ptk2aa | 0.224270244 | 0.259268875 | 0.303476245 | 0.275768955 |
| ENSDART00000159899 | mapk10 | -0.350531159 | -0.285511604 | -0.104624207 | 0.054397304 |
| ENSDART00000159908 | camk1db | -0.548571048 | -0.478436516 | -0.873630088 | -0.641909911 |
| ENSDART00000159916 | tns1a | -0.270694335 | -0.694531751 | -0.372872148 | -0.014128978 |
| ENSDART00000159928 | si:dkey-274m17.3 | -0.105974275 | -0.260400887 | -0.454854053 | -0.384734441 |
| ENSDART00000159942 | kcnc3b | -0.794025304 | -1.361623092 | -0.389946342 | 0.009706478 |
| ENSDART00000159949 | wdr1 | 0.362540326 | 0.308247591 | 0.564916756 | 0.170183845 |
| ENSDART00000159950 | si:ch1073-450f2.1 | -0.285454261 | -0.252056938 | -0.418320337 | -0.16839092 |
| ENSDART00000159952 | sema5bb | 0.4270075 | 0.43121825 | 0.280410439 | 0.178648412 |
| ENSDART00000159956 | ndufa4l2a | 0.563891792 | 0.528048098 | 0.326797956 | 0.131990505 |
| ENSDART00000159958 | hcn1 | -0.047230014 | -0.465327973 | -0.263546106 | -0.045423137 |
| ENSDART00000159965 | si:ch211-255i3.4 | 0.121150704 | 0.407179882 | 0.241107117 | 0.125992518 |

| | | | | | |
|---|---|---|---|---|---|
| ENSDART00000159987 | si:dkey-65b13.13 | -0.209652039 | -0.259703439 | -0.637719895 | -0.234430743 |
| ENSDART00000159989 | cpne7 | -0.2792921 | -0.302878059 | -0.342117017 | -0.194332814 |
| ENSDART00000159997 | cacna1ab | -0.118996824 | -0.488974274 | -0.025554874 | 0.331301032 |
| ENSDART00000160000 | spdl1 | 1.145158953 | 1.158182543 | 1.166185329 | 0.390123322 |
| ENSDART00000160006 | Metazoa_SRP | 1.639017988 | 2.129447988 | 1.830106321 | 0.946987666 |
| ENSDART00000160017 | CABZ01058261.1 | 0.903038959 | 0.866986019 | 0.595792077 | 0.158138964 |
| ENSDART00000160018 | DUSP26 | -0.660306321 | -0.709450564 | -0.129656337 | 0.007184288 |
| ENSDART00000160048 | | 1.164230248 | 1.13549021 | 1.093110731 | 1.576844966 |
| ENSDART00000160055 | CABZ01048399.1 | 2.04373243 | 1.562490919 | 0.966755902 | 1.237958862 |
| ENSDART00000160073 | sntg2 | -0.084902345 | -0.387083384 | -0.20960703 | -0.013813106 |
| ENSDART00000160107 | CU639413.1 | 0.54252914 | 0.409259037 | 1.419043143 | 0.130740048 |
| ENSDART00000160115 | per1a | 0.041503641 | -0.272737679 | -0.476359218 | -0.15653808 |
| ENSDART00000160116 | racgap1 | 0.647984241 | 0.386764067 | 0.354192772 | 0.100327502 |
| ENSDART00000160134 | hibadha | -0.191259291 | -0.221221329 | -0.647202373 | -0.244609156 |
| ENSDART00000160138 | sobpa | -0.321225255 | -0.895188795 | -0.069068022 | 0.312612038 |
| ENSDART00000160154 | si:dkey-238i5.2 | -0.200329967 | 0.657530632 | 1.151706484 | 1.107326351 |
| ENSDART00000160155 | PDCL3 | 1.150151141 | 1.512943629 | 1.437587964 | 2.391657524 |
| ENSDART00000160175 | SYNDIG1 | -0.028604943 | -0.23293463 | -0.464850771 | -0.153309965 |
| ENSDART00000160177 | kcnb2 | 4.745895577 | 4.144383899 | 5.457366505 | 5.666212289 |
| ENSDART00000160189 | hip1rb | -0.298878525 | -0.25639736 | 0.130568312 | 0.08288489 |
| ENSDART00000160201 | tnfrsf11b | 2.52634883 | 1.22900254 | 0.873090756 | -0.039240801 |
| ENSDART00000160206 | rap2b | 0.536023906 | 0.628551975 | 0.691733134 | 0.267820563 |
| ENSDART00000160209 | si:ch211-204a13.2 | -0.408114948 | -0.445711734 | -0.270878027 | 0.146816503 |
| ENSDART00000160220 | kif5aa | -0.17024292 | 0.126849007 | 0.479028265 | 0.345007215 |
| ENSDART00000160236 | fras1 | -0.444181856 | -0.638830186 | -0.533612869 | -0.242512624 |
| ENSDART00000160242 | TYMP | -0.668893964 | -0.374517624 | -0.757876583 | -0.434642219 |
| ENSDART00000160245 | si:ch73-236c18.7 | 2.124440879 | 1.318007353 | 1.287829039 | 0.437471879 |
| ENSDART00000160284 | pclob | -0.164621492 | -0.643566861 | -0.569524507 | -0.247739035 |
| ENSDART00000160288 | FP236812.6 | 0.238382616 | 0.543365019 | 0.866982901 | 0.48841948 |
| ENSDART00000160297 | vcana | 1.601498468 | 0.665214133 | 0.454293537 | 0.486706931 |
| ENSDART00000160305 | si:zfos-943e10.1 | -0.732433247 | -0.018482138 | -0.2347001 | -0.520119378 |
| ENSDART00000160308 | pde3a | -0.225066293 | -0.312371173 | 0.037986462 | 0.082811204 |
| ENSDART00000160324 | CABZ01102047.1 | -0.357465059 | -0.236339235 | 0.433910345 | 0.432628055 |
| ENSDART00000160328 | zgc:109949 | -0.362552375 | -0.659300829 | -0.471802707 | -0.430381288 |
| ENSDART00000160331 | atp6v0e1 | 0.536560194 | 0.109862681 | -0.220128717 | -0.336777163 |
| ENSDART00000160337 | si:ch73-299h12.8 | -0.912707831 | -0.414120081 | -0.53567861 | -0.405510308 |
| ENSDART00000160350 | IQSEC1 (1 of many) | -0.73025949 | -0.724161984 | -0.227945297 | -0.158150827 |
| ENSDART00000160362 | EGFLAM | -0.160131743 | -0.324587558 | -0.48719254 | -0.200185739 |
| ENSDART00000160400 | gchfr | -0.373470826 | -0.390793782 | -0.620398999 | -0.352985133 |
| ENSDART00000160401 | tmem63ba | -0.025633015 | 0.10439182 | 0.273009198 | 0.198949087 |
| ENSDART00000160409 | arf6a | 0.10280988 | -0.030227439 | 0.312347712 | -0.095328656 |
| ENSDART00000160424 | ftr36 | -0.057998525 | -0.261050684 | 0.030750272 | -0.246695716 |
| ENSDART00000160425 | LAMP1 | -0.291224845 | -0.711008477 | -0.768820578 | -0.726430297 |
| ENSDART00000160431 | dhrs13l1 | -0.306310555 | -0.206975564 | -0.316329966 | -0.293060295 |
| ENSDART00000160439 | CABZ01037298.1 | -0.12483018 | -0.201319697 | -0.520978425 | -0.240731645 |
| ENSDART00000160444 | si:dkey-95j14.1 | -0.172108125 | 0.025439256 | 0.48271623 | 0.286285124 |
| ENSDART00000160450 | CU571069.1 | -0.391785076 | -0.855515025 | -0.904047805 | -0.612021184 |
| ENSDART00000160464 | pcdh1gb2 | -0.570850707 | -0.31567681 | -0.173184848 | -0.165810233 |
| ENSDART00000160465 | opn6a | -0.331059122 | -0.451170972 | -0.507697352 | -0.105376089 |
| ENSDART00000160466 | ndrg4 | -0.797536516 | -0.447807808 | -0.265830716 | -0.169347229 |
| ENSDART00000160468 | ubl3a | -0.157200321 | -0.291255908 | -0.684230474 | -0.436379037 |

| | | | | | |
|---|---|---|---|---|---|
| ENSDART00000160469 | si:ch211-271b14.1 | -0.816113506 | -0.956386746 | -0.419124903 | -0.031051269 |
| ENSDART00000160492 | alad | 0.30171604 | 0.443307609 | 0.248348623 | 0.158762163 |
| ENSDART00000160495 | si:dkey-184p18.2 | 0.380936601 | -0.937356475 | -0.364392869 | -0.267264411 |
| ENSDART00000160503 | slc15a4 | 0.534058917 | 0.262741482 | 0.05558204 | -0.165911292 |
| ENSDART00000160509 | trip10a | 5.462743185 | 3.19366481 | 2.422825884 | 1.530105247 |
| ENSDART00000160527 | prokr1a | 0.020123926 | -0.217659905 | -0.314012019 | -0.217346505 |
| ENSDART00000160536 | gnsb | 1.380175579 | 0.91424754 | 0.488823056 | 0.00723105 |
| ENSDART00000160538 | arhgap21a | -0.416942655 | -0.374699279 | -1.045101409 | -0.721453587 |
| ENSDART00000160542 | cabp2a | -0.310654407 | -0.386778791 | -0.172317372 | -0.340396034 |
| ENSDART00000160555 | sema6e | -0.284106568 | -0.324928323 | -0.250029521 | -0.168860793 |
| ENSDART00000160562 | impg2b | 0.024461351 | -0.26080833 | -0.706384589 | -0.340668028 |
| ENSDART00000160564 | cyyr1 | -0.193986492 | -0.219795811 | -0.518063496 | -0.536169671 |
| ENSDART00000160566 | PPEF2 | 0.06510909 | 0.104690545 | -0.262166795 | 0.034536141 |
| ENSDART00000160571 | mxe | 0.057431509 | -0.237146132 | 2.037062599 | -0.095948656 |
| ENSDART00000160575 | CU459186.1 | 0.73831154 | 1.064084865 | 0.743119809 | 0.540430362 |
| ENSDART00000160582 | creb5a | 2.028867303 | 1.691967909 | 1.27021295 | 0.836790625 |
| ENSDART00000160588 | si:dkeyp-86d6.2 | -0.052465168 | -0.307686486 | -0.441266657 | -0.083441391 |
| ENSDART00000160600 | cecr5 | -0.047380691 | 0.036242917 | 0.854821049 | 0.34511846 |
| ENSDART00000160604 | ftsj3 | 0.856590556 | 0.695492085 | 0.825371198 | 0.428239045 |
| ENSDART00000160606 | mat2aa | 0.169445123 | 0.256103943 | 0.254062393 | 0.177493443 |
| ENSDART00000160612 | bin3 | 0.276069755 | 0.185522197 | -0.128294303 | -0.167255765 |
| ENSDART00000160614 | tmem106ba | -0.216998722 | -0.275060394 | -0.388680799 | -0.295077411 |
| ENSDART00000160652 | iqsec3a | -0.564945044 | -0.609250827 | -0.334399494 | -0.124185292 |
| ENSDART00000160662 | CABZ01056578.1 | -0.678580777 | -0.38089804 | -0.298190661 | -0.130306976 |
| ENSDART00000160663 | tnk2b | 0.180874765 | 0.228576581 | 0.365115453 | 0.173216656 |
| ENSDART00000160672 | MYH7 (1 of many) | 0.452458943 | 2.61830504 | 2.948579403 | 1.474866684 |
| ENSDART00000160700 | dnm1l | -0.370704077 | -0.348232697 | -0.495814147 | -0.141373029 |
| ENSDART00000160707 | smx5 | -0.006041688 | -0.141079427 | -0.427667941 | -0.488718653 |
| ENSDART00000160712 | dusp1 | 0.749410253 | 0.773951106 | 0.584904324 | 1.278120339 |
| ENSDART00000160715 | pik3r3a | -0.538136397 | -0.043164195 | -0.249012844 | -0.109162985 |
| ENSDART00000160738 | arhgap11a | 2.137805097 | 2.112857321 | 2.062351639 | 2.227173194 |
| ENSDART00000160742 | clmn | -0.341601235 | -0.332225059 | -0.489517534 | -0.288919784 |
| ENSDART00000160753 | GANAB (1 of many) | 0.004666803 | -0.095016348 | -0.32432452 | -0.161685037 |
| ENSDART00000160763 | Metazoa_SRP | 1.465611255 | 3.00506243 | 2.307455794 | 0.729030238 |
| ENSDART00000160766 | pim1 | 0.465870139 | 0.649909313 | 0.413306503 | 0.082988542 |
| ENSDART00000160767 | zgc:153845 | -0.366926311 | -0.362052716 | -0.223963795 | -0.098406698 |
| ENSDART00000160783 | rorcb | -0.657257878 | 0.184912673 | 0.679992874 | 0.484278813 |
| ENSDART00000160789 | SDK2 (1 of many) | -0.531586731 | -0.76443238 | -0.126364502 | 0.194536592 |
| ENSDART00000160809 | agap1 | -0.12094537 | -0.11171461 | 0.245949776 | 0.322867446 |
| ENSDART00000160811 | b3gat1a | -0.509506437 | -0.522178413 | -0.230803048 | 0.152925891 |
| ENSDART00000160829 | mhc1zda | -0.410179214 | -0.276613879 | -0.418038837 | -0.322538004 |
| ENSDART00000160840 | prkar1b | -0.31036091 | -0.161481279 | -0.113380497 | -0.010356954 |
| ENSDART00000160841 | si:ch211-248g20.5 | -0.153486145 | -0.342330243 | -0.525669956 | -0.178412263 |
| ENSDART00000160847 | hgh1 | 0.505719048 | 0.435629442 | 0.388567431 | 0.068708352 |
| ENSDART00000160850 | inpp5e | -0.436095235 | -0.00283574 | 0.156525888 | 0.020455167 |
| ENSDART00000160866 | cry2 | -0.114688269 | -0.279009524 | -0.604836383 | -0.332518261 |
| ENSDART00000160881 | pcdh1g30 | -0.53895262 | -0.083348601 | 0.159825551 | 0.241016465 |
| ENSDART00000160885 | stbd1 | -0.425801769 | -0.465748469 | -0.195945725 | 0.034690374 |
| ENSDART00000160901 | FMNL1 (1 of many) | 1.814810207 | 1.982271858 | 2.471309982 | 2.117776194 |
| ENSDART00000160902 | naga | 0.641123507 | 0.233762511 | 0.291444137 | -0.017668678 |
| ENSDART00000160927 | cers1 | -0.519571193 | -0.339170129 | -0.296459142 | -0.066758323 |

| | | | | | |
|---|---|---|---|---|---|
| ENSDART00000160933 | si:ch211-153l6.6 | -0.192271617 | -0.190332633 | -0.289802511 | -0.209468422 |
| ENSDART00000160950 | dyrk1b | -0.419283008 | -0.231908493 | 0.058730046 | 0.190507005 |
| ENSDART00000160955 | slc38a3b | -0.475545538 | -0.367921084 | -0.471714594 | -0.850494481 |
| ENSDART00000160956 | oaz1b | -0.133855172 | -0.165932835 | -0.380106363 | -0.375655267 |
| ENSDART00000160979 | jpt1b | 0.682460724 | 1.366219793 | 1.515562314 | 0.876953288 |
| ENSDART00000161002 | kbtbd11 | -0.286857265 | -0.122332373 | -0.245379055 | -0.228313034 |
| ENSDART00000161003 | cpne9 | -1.056006542 | -1.193405404 | -0.117014879 | 0.290018934 |
| ENSDART00000161007 | ets2 | -0.669649632 | -0.198034695 | -0.011258584 | -0.233709301 |
| ENSDART00000161008 | filip1a | 1.277613198 | 1.48763933 | 0.952639764 | 1.12798471 |
| ENSDART00000161017 | eif3s6ip | 0.313830903 | 0.293421495 | 0.276651786 | 0.120270233 |
| ENSDART00000161040 | U1 | 0.811168663 | 0.450901632 | 0.418592021 | 0.580227266 |
| ENSDART00000161048 | psmf1 | -0.002898908 | -0.064926331 | 0.496331974 | -0.135580746 |
| ENSDART00000161056 | zgc:63587 | 0.468520933 | 0.333532949 | 0.066863333 | -0.028483781 |
| ENSDART00000161059 | tusc2a | 0.068801861 | -0.121312821 | -0.423616648 | -0.242714442 |
| ENSDART00000161063 | heatr1 | 0.383217268 | 0.256645773 | 0.173900631 | 0.113179939 |
| ENSDART00000161081 | satb1a | -0.172286726 | 0.059256047 | 0.306569639 | 0.367466245 |
| ENSDART00000161087 | efna3b | -0.321733474 | -0.009174863 | 0.104028903 | 0.249289152 |
| ENSDART00000161100 | FAT3 (1 of many) | -0.16621971 | -0.375379053 | -0.163291758 | -0.002633554 |
| ENSDART00000161101 | lyrm5a | -0.096487275 | -0.208413777 | -0.302906059 | -0.144622611 |
| ENSDART00000161107 | pcdh1g9 | 0.063645152 | 0.386867409 | 0.439855094 | 0.563716536 |
| ENSDART00000161115 | CR382281.1 | -1.148281272 | -0.70950654 | -0.315399276 | 0.210191426 |
| ENSDART00000161117 | | 0.194355392 | 0.331000906 | 0.496659366 | 0.20108058 |
| ENSDART00000161121 | oxct1a | -0.18384202 | -0.521566935 | -0.230939432 | -0.260776637 |
| ENSDART00000161137 | scn1bb | -0.455645009 | -0.546610376 | 0.183837534 | 0.407926687 |
| ENSDART00000161153 | GAN | -0.076211127 | 0.283155249 | 0.536837618 | 0.420868128 |
| ENSDART00000161156 | kcnj9 | -0.647175508 | -0.597438502 | -0.398278727 | -0.23455974 |
| ENSDART00000161168 | si:ch211-232d10.1 | -0.094530829 | -0.188391079 | -0.421762057 | -0.370881661 |
| ENSDART00000161184 | si:dkey-238f9.1 | -0.217284848 | -0.396117732 | -0.370650729 | 0.049289618 |
| ENSDART00000161216 | dync1i2a | 1.005522301 | 0.520541958 | 0.468549497 | -0.022454144 |
| ENSDART00000161230 | arhgef12b | 0.167990134 | 0.325221884 | 0.334885818 | 0.114051067 |
| ENSDART00000161240 | zgc:92161 | 0.584076871 | 0.663364233 | 0.486030672 | 0.10810748 |
| ENSDART00000161244 | CR293509.1 | 0.407469749 | 1.465267423 | 1.274347742 | 0.975210426 |
| ENSDART00000161250 | grin1b | -0.242810387 | -0.349766254 | -0.246157691 | 0.067747546 |
| ENSDART00000161261 | lmbrd2a | -0.185598793 | -0.153033552 | -0.360733538 | -0.113007589 |
| ENSDART00000161264 | ntrk3b | -0.768053413 | -0.724776652 | -0.30616772 | 0.183034577 |
| ENSDART00000161266 | CU928046.1 | -0.122276569 | -0.195535625 | -0.336584046 | -0.188368371 |
| ENSDART00000161272 | dclk1a | -0.780154894 | -0.444639927 | -0.375433616 | -0.486681237 |
| ENSDART00000161297 | lrrn3b | -0.369968613 | -0.470809526 | -0.36045804 | -0.046069156 |
| ENSDART00000161322 | slc29a1a | 0.535023852 | 0.636573744 | 0.365586045 | -0.115892928 |
| ENSDART00000161330 | col27a1a | 0.210670367 | 0.547410859 | 0.646435326 | 0.255665234 |
| ENSDART00000161347 | CT583728.4 | 1.170122423 | 1.14685482 | 1.328877257 | 0.769737347 |
| ENSDART00000161377 | cdk2ap2 | 0.241317874 | 0.341753789 | 0.416424679 | 0.144968334 |
| ENSDART00000161389 | CR388042.1 | 0.279433291 | 0.140072251 | 0.112517442 | 0.232646376 |
| ENSDART00000161397 | atg13 | -0.137451879 | -0.18705676 | -0.405447689 | -0.269252338 |
| ENSDART00000161407 | tnika | -0.172678134 | 0.105967748 | 0.465016741 | 0.501980476 |
| ENSDART00000161414 | meis2a | -0.352024462 | -0.259692709 | -0.258698057 | -0.07620107 |
| ENSDART00000161421 | zgc:194627 | 1.998498644 | 1.298544665 | 0.773438314 | 0.249109157 |
| ENSDART00000161436 | pcdh7b | -0.687665671 | -0.773740554 | -0.245502692 | 0.12030382 |
| ENSDART00000161440 | limch1b | -0.008873535 | -0.193340096 | -0.430561872 | -0.157445684 |
| ENSDART00000161445 | fkrp | -0.298106015 | -0.219330579 | -0.124679519 | -0.039794435 |
| ENSDART00000161456 | zgc:153924 | -0.044959851 | 0.322360252 | 0.292539556 | -0.042756925 |

| | | | | | |
|---|---|---|---|---|---|
| ENSDART00000161460 | pmt | 2.538801298 | 3.7270331 | 2.780124527 | 1.870914111 |
| ENSDART00000161505 | mgrn1a | -0.282029057 | -0.374993034 | -0.219589284 | -0.246471373 |
| ENSDART00000161507 | zgc:153426 | -1.880708803 | -0.117657621 | 0.865737326 | 1.072245785 |
| ENSDART00000161514 | CABZ01079986.1 | -0.184749182 | 1.474076268 | 0.73277167 | -0.443845367 |
| ENSDART00000161525 | zgc:171704 | 0.267822787 | -0.368878081 | -1.0813007 | -1.211102226 |
| ENSDART00000161532 | zfyve9b | -0.607365928 | -0.871459147 | -0.090833114 | 0.138377783 |
| ENSDART00000161533 | gfra1b | -0.396684985 | -0.507239408 | -0.32341327 | -0.285757426 |
| ENSDART00000161561 | ANK1 (1 of many) | -0.90876402 | -0.980007711 | -0.299110873 | -0.111310605 |
| ENSDART00000161566 | b4galt6 | 0.155163411 | 0.348743299 | 0.367166825 | 0.307075635 |
| ENSDART00000161567 | KCNT1 (1 of many) | -0.564631347 | -0.618941642 | -0.212804629 | 0.112863315 |
| ENSDART00000161582 | ubfd1 | 0.213892452 | 0.378867256 | 0.328612523 | 0.082100895 |
| ENSDART00000161586 | flot2a | 0.149914225 | 0.375737842 | 0.414499477 | 0.21470497 |
| ENSDART00000161597 | si:ch211-262i1.3 | 0.726810222 | 0.918966234 | 1.284521547 | 0.120004192 |
| ENSDART00000161613 | SLC7A1 (1 of many) | 0.777752401 | 1.137604367 | 0.924750387 | 0.450737498 |
| ENSDART00000161646 | sema6bb | -0.22063759 | -0.362707192 | -0.160882628 | 0.032299092 |
| ENSDART00000161647 | fnbp1b | 0.397270086 | 0.395681626 | 0.254962279 | -0.147831954 |
| ENSDART00000161650 | atp2a2b | 0.018261356 | 0.090439762 | 0.265163492 | 0.020190481 |
| ENSDART00000161652 | si:dkeyp-72e1.9 | -0.109528412 | -0.231594686 | -0.432617543 | -0.203491956 |
| ENSDART00000161662 | ttbk1a | -0.363444816 | 0.177390115 | 0.591497043 | 0.421229884 |
| ENSDART00000161673 | CR450729.3 | -0.466290317 | -0.130444226 | -0.527062063 | -0.184859299 |
| ENSDART00000161693 | cetn2 | -0.42442122 | 0.157217987 | 0.592113306 | 0.77779027 |
| ENSDART00000161698 | spock1 | -0.400537237 | -0.522818259 | -0.36383883 | -0.3494177 |
| ENSDART00000161700 | st3gal2 | -0.330022994 | -0.187434223 | 0.066957229 | 0.392836726 |
| ENSDART00000161704 | CNDP1 | -0.617638491 | -0.321066179 | -0.670749127 | -2.338544776 |
| ENSDART00000161708 | si:ch73-29l19.1 | -0.183985257 | -0.352166832 | -0.310452263 | -0.095145467 |
| ENSDART00000161725 | map7d1a | -0.696007823 | -0.432281688 | 0.042931769 | 0.13512701 |
| ENSDART00000161735 | flrt1a | -0.150942569 | -0.355176624 | -0.261636355 | -0.097731299 |
| ENSDART00000161755 | CU681842.1 | -0.157328162 | -0.245310878 | -0.348302227 | -0.161138988 |
| ENSDART00000161770 | cpne9 | -0.832265602 | -0.611063661 | -0.201518154 | 0.170760147 |
| ENSDART00000161773 | KCNJ11 (1 of many) | -0.422252277 | -0.360338989 | -0.607728772 | -0.332578335 |
| ENSDART00000161798 | si:ch211-188f17.1 | -0.190236489 | -0.379885925 | -0.177844783 | -0.0015826 |
| ENSDART00000161807 | gnl2 | 0.286549103 | 0.322764935 | 0.293655806 | 0.087475664 |
| ENSDART00000161818 | prr33 | 0.089254338 | 1.155971891 | 0.95184285 | 0.668590241 |
| ENSDART00000161823 | ubl3a | -0.089124518 | -0.239047703 | -0.471864878 | -0.290785384 |
| ENSDART00000161828 | si:rp71-1h20.9 | -1.646379844 | -1.190834107 | -1.032421214 | -1.471047696 |
| ENSDART00000161840 | CT574575.1 | -0.29915223 | -0.372107241 | -0.764083264 | -0.340151107 |
| ENSDART00000161854 | pls3 | 0.235895362 | 0.649174918 | 0.495504879 | 0.31420242 |
| ENSDART00000161869 | dysf | 1.077492273 | 1.30463907 | 1.440346007 | 1.017975966 |
| ENSDART00000161882 | bzw2 | 0.083435949 | -0.31200966 | -0.5490146 | -0.324311149 |
| ENSDART00000161892 | vdra | -0.26830787 | -0.422747599 | -0.12129137 | -0.077310915 |
| ENSDART00000161906 | vwa11 | 1.529478256 | 0.755786632 | 0.896680083 | 0.546980022 |
| ENSDART00000161908 | nsun2 | 4.199768382 | 2.926785137 | 4.095496425 | 2.988717061 |
| ENSDART00000161925 | rundc3aa | -0.101671945 | 0.461428265 | 0.452450402 | 0.467812596 |
| ENSDART00000161930 | strn4 | -0.158687938 | -0.097490818 | 0.286103327 | 0.194201646 |
| ENSDART00000161932 | npr3 | 0.255355781 | 0.183557252 | -0.357151542 | -1.063769364 |
| ENSDART00000161938 | si:dkeyp-9d4.3 | -0.230392301 | -0.461066854 | -0.432261332 | -0.276495229 |
| ENSDART00000161967 | ctsc | 1.279775059 | 0.583635254 | 0.329059668 | -0.157215155 |
| ENSDART00000161978 | BX649556.1 | 0.438096957 | 0.234757098 | 0.251734491 | 0.204407619 |
| ENSDART00000161982 | SCARNA6 | 0.10205479 | -0.170611192 | -0.840119974 | -1.131956342 |
| ENSDART00000161992 | PAQR9 | -0.193753966 | -0.524789375 | -0.118624882 | 0.042415903 |
| ENSDART00000162002 | grm1b | -0.568603213 | -0.750999293 | -0.21404532 | -0.094504234 |

| | | | | | |
|---|---|---|---|---|---|
| ENSDART00000162007 | Metazoa_SRP | 0.903949119 | 2.303055473 | 1.702580761 | 0.513671528 |
| ENSDART00000162010 | zgc:92140 | -0.256831407 | -0.245054913 | -0.345912399 | -0.346929099 |
| ENSDART00000162018 | CABZ01002768.2 | 0.265184254 | 0.128771486 | 0.086197325 | 0.254354536 |
| ENSDART00000162023 | plppr2a | 0.019417247 | -0.576626688 | 0.082766084 | 0.317388761 |
| ENSDART00000162026 | ninl | -0.053184172 | -0.137296281 | -0.336798097 | -0.200288689 |
| ENSDART00000162046 | sez6a | -0.383252568 | -0.442703172 | -0.383098336 | -0.145401001 |
| ENSDART00000162055 | mark1 | -0.053725407 | 0.099849875 | 0.364172927 | 0.499618468 |
| ENSDART00000162063 | ppifb | -0.192515925 | -0.861096392 | -0.286677305 | -0.512048401 |
| ENSDART00000162070 | tmem9 | -0.208225931 | -0.361514875 | -0.511868133 | -0.393713678 |
| ENSDART00000162097 | CABZ01044157.1 | 1.031550352 | 0.670727071 | 0.212982548 | -0.249292553 |
| ENSDART00000162119 | ewsr1a | -3.071838226 | 0.182908872 | -0.16348899 | 0.256173861 |
| ENSDART00000162133 | tenm4 | -0.287462387 | -0.412164757 | -0.184969779 | 0.049525758 |
| ENSDART00000162139 | si:ch211-244c8.4 | -0.176471944 | 0.210261822 | 0.365856922 | 0.312900704 |
| ENSDART00000162145 | FQ377629.1 | -0.12898611 | -0.128745299 | 0.191405016 | 0.357155255 |
| ENSDART00000162200 | STX3 (1 of many) | -0.155323726 | -0.36938348 | -0.41294698 | -0.132327196 |
| ENSDART00000162218 | col10a1b | 0.45003474 | 2.713575183 | 3.448261467 | 1.87125716 |
| ENSDART00000162222 | SHC3 | -0.255397657 | -0.359792108 | -0.178043549 | -0.323533675 |
| ENSDART00000162228 | grip2a | -0.209580983 | -0.468643062 | -0.36247099 | -0.147880595 |
| ENSDART00000162277 | grip1 | -0.456620328 | -0.2169797 | 0.146006741 | 0.157130316 |
| ENSDART00000162278 | CABZ01053748.1 | -0.203505904 | -0.497916501 | -0.162237791 | 0.002359168 |
| ENSDART00000162282 | dicp3.1 | 0.839763954 | 0.379787637 | 0.004488715 | -0.034973807 |
| ENSDART00000162290 | GARNL3 | 0.087177538 | 0.188445106 | 0.266999217 | 0.231469459 |
| ENSDART00000162294 | utrn | -0.256734894 | -0.284908669 | -0.969336134 | -0.369539596 |
| ENSDART00000162296 | cfp | 0.329536618 | 0.225136408 | -0.193120136 | -0.891420593 |
| ENSDART00000162302 | bnip3 | -0.278521269 | -0.244198471 | -0.204837653 | -0.079960627 |
| ENSDART00000162324 | kalrnb | -0.363230198 | -0.20882241 | 0.367737091 | 0.490925401 |
| ENSDART00000162331 | si:dkey-22o22.2 | -0.434953721 | -0.633123757 | -0.287732073 | -0.108064019 |
| ENSDART00000162359 | sorbs1 | -0.104497249 | -0.20837462 | -0.418151878 | -0.317718588 |
| ENSDART00000162377 | FP085414.1 | 0.809712454 | 1.275862162 | 1.389373351 | 0.791945088 |
| ENSDART00000162381 | dkk3a | -0.551126502 | -0.361561955 | -0.324385943 | 0.074151181 |
| ENSDART00000162387 | zbtb38 | -0.362042869 | -0.285747988 | -0.199364681 | -0.27814593 |
| ENSDART00000162399 | tacc3 | 1.052644038 | 0.862999188 | 0.469743238 | -0.177095944 |
| ENSDART00000162403 | napgb | -0.42647327 | -0.557407706 | -0.383093342 | -0.343712014 |
| ENSDART00000162421 | si:ch73-233f7.7 | 0.22834555 | 0.078820055 | 0.428518248 | 0.477794315 |
| ENSDART00000162459 | scarb2a | 0.204566318 | 0.711810949 | 0.821282796 | 0.534817262 |
| ENSDART00000162472 | VWA5A (1 of many) | 0.543940251 | 0.262980075 | 0.410699646 | -0.136546959 |
| ENSDART00000162474 | irx1a | -1.132797992 | -0.349993381 | 0.581252213 | 0.55366573 |
| ENSDART00000162482 | zgc:152863 | 0.539805357 | 0.660295595 | 0.432850415 | 0.095615261 |
| ENSDART00000162485 | pax6a | -0.383059238 | -0.321515523 | -0.107334897 | 0.077147684 |
| ENSDART00000162493 | kif5ab | -0.358862317 | -0.407894033 | -0.585721315 | -0.34361757 |
| ENSDART00000162501 | abca4a | -0.194244085 | -0.198091298 | -0.350966144 | -0.228681741 |
| ENSDART00000162520 | SHC3 | -0.325188551 | -0.499304719 | 0.001359259 | -0.194280026 |
| ENSDART00000162527 | si:ch211-160f23.7 | -0.331311011 | -0.409364601 | -0.344819459 | -0.114250304 |
| ENSDART00000162529 | zgc:172139 | 0.398363371 | 0.414482162 | 0.745277151 | 0.505205904 |
| ENSDART00000162539 | VWA5A (1 of many) | 1.505543925 | 2.134948516 | 2.442684472 | 2.675816303 |
| ENSDART00000162540 | Metazoa_SRP | 1.640394936 | 1.730362994 | 1.214251464 | 0.854214156 |
| ENSDART00000162541 | thbs4a | 1.94439423 | 3.279546426 | 3.089397424 | 2.286354886 |
| ENSDART00000162555 | dpyda.1 | 0.585872258 | 0.381337517 | 0.027895003 | -0.679828257 |
| ENSDART00000162564 | CT027980.1 | -0.351165538 | -0.178285414 | -0.533647461 | -0.500417084 |
| ENSDART00000162566 | h2afvb | 0.462673668 | 0.293160887 | 0.459514152 | 0.229847552 |
| ENSDART00000162587 | chd5 | -0.534129477 | -1.024077887 | -0.70517204 | -0.958580386 |

| | | | | | |
|---|---|---|---|---|---|
| ENSDART00000162595 | camk2g1 | -0.045798893 | -0.36296086 | -0.180481912 | 0.032536235 |
| ENSDART00000162601 | scn2b | -0.529498392 | -0.64570335 | -0.359290818 | 0.039638437 |
| ENSDART00000162607 | CR361561.2 | 0.410555114 | 0.246904987 | 0.207590099 | 0.517188645 |
| ENSDART00000162617 | ppp2cb | 0.050413288 | 0.287682373 | 0.42325509 | 0.191699941 |
| ENSDART00000162622 | FP103009.1 | -0.352224047 | -0.048475545 | 0.149997583 | 0.073530546 |
| ENSDART00000162637 | CABZ01054392.2 | -0.811315315 | 0.030869752 | -0.070916412 | -0.071296331 |
| ENSDART00000162664 | 5S_rRNA | -0.438943374 | -1.186726175 | -0.920371372 | -0.548334869 |
| ENSDART00000162668 | cremb | 4.310209888 | 4.052771 | 3.333577791 | 2.240090056 |
| ENSDART00000162669 | slc4a5 | -0.13383932 | -0.22542524 | -0.430682825 | -0.287279235 |
| ENSDART00000162670 | slc8a1b | -0.269852688 | -0.529256481 | -0.096682296 | 0.048355338 |
| ENSDART00000162675 | trim2b | -0.409757052 | -0.631223683 | -0.07653124 | 0.093922127 |
| ENSDART00000162683 | trappc9 | -0.370182614 | -0.24071592 | -0.273499446 | -0.133452933 |
| ENSDART00000162696 | cacnb4b | -0.474781259 | -0.004559691 | 0.081636688 | 0.060425945 |
| ENSDART00000162710 | fgf13b | 0.123584158 | 0.587591002 | 0.690906211 | 0.488537905 |
| ENSDART00000162711 | cnksr2b | -0.20949159 | -0.28214605 | -0.418907028 | -0.082150791 |
| ENSDART00000162714 | pcdh10b | -0.340908137 | -0.430434424 | -0.069086695 | 0.178421467 |
| ENSDART00000162722 | zgc:153867 | 0.467799327 | 0.468299246 | 0.579069745 | 0.022677056 |
| ENSDART00000162732 | | -0.497798966 | -0.517625334 | -0.441909965 | -0.132483146 |
| ENSDART00000162761 | CU984600.2 | 0.225517614 | -0.442281881 | 2.62985535 | 0.023562363 |
| ENSDART00000162799 | crb3a | 0.362169262 | 0.040586113 | -0.086039525 | -0.014272311 |
| ENSDART00000162804 | KCNT1 (1 of many) | -0.406408577 | -0.48233323 | -0.378259831 | 0.110845983 |
| ENSDART00000162827 | si:dkey-92j12.5 | -0.013228451 | -0.265522895 | -0.385437174 | -0.080127062 |
| ENSDART00000162838 | agrn | 0.183430669 | 0.084636207 | 0.451853257 | 0.572854924 |
| ENSDART00000162850 | irx3a | -0.694816328 | -0.301932125 | 0.187764734 | 0.208405543 |
| ENSDART00000162855 | pcdh1g13 | 0.288424517 | 0.288939089 | 0.455966507 | 0.614698967 |
| ENSDART00000162857 | nr4a3 | -1.482016444 | -1.152680943 | -0.606031454 | -1.193862677 |
| ENSDART00000162858 | lmbrd1 | -0.243655467 | -0.23291492 | -0.332336473 | -0.114530821 |
| ENSDART00000162868 | PCMTD2 (1 of many) | -0.184710465 | -0.224010966 | -0.277853434 | -0.160151685 |
| ENSDART00000162875 | rogdi | -0.224891588 | -0.27706391 | -0.36191309 | -0.159630193 |
| ENSDART00000162886 | il1rapl1b | -0.535131005 | -0.60836138 | -0.536159373 | -0.086785386 |
| ENSDART00000162897 | Metazoa_SRP | 1.178659737 | 1.5924133 | 1.384383775 | 1.197556527 |
| ENSDART00000162915 | dclk2b | 0.022821815 | 0.633802822 | 1.144009661 | 0.962163073 |
| ENSDART00000162916 | si:ch211-177d9.1 | -0.561077771 | -0.777629972 | -0.458457407 | -0.043076344 |
| ENSDART00000162924 | SCARNA2 | -0.051299469 | -0.033570311 | -0.355924265 | -0.225455165 |
| ENSDART00000162940 | mob2b | 0.020755037 | -0.2829937 | -0.541864256 | -0.204837959 |
| ENSDART00000162958 | si:ch211-260e23.9 | 0.345171245 | 0.431307401 | 0.079575678 | 0.236719343 |
| ENSDART00000162970 | znf1001 | 2.835681681 | 3.195296273 | 3.608291375 | 4.001715823 |
| ENSDART00000162984 | cacnb2a | -0.567328475 | -0.858575333 | 0.120461387 | 0.550245867 |
| ENSDART00000163006 | magi1b | -0.558734199 | -0.633598853 | 0.045195354 | 0.572492467 |
| ENSDART00000163018 | bet1l | 0.723011674 | 0.444641169 | 0.411413339 | 0.530095318 |
| ENSDART00000163023 | sypb | -0.076679322 | -0.067070783 | -0.475180992 | -0.188875227 |
| ENSDART00000163025 | slc37a1 | 0.146947624 | 0.663221495 | 0.752146696 | 0.093340063 |
| ENSDART00000163026 | igf2bp3 | 1.596757122 | 2.364080903 | 3.304264113 | 3.343316319 |
| ENSDART00000163032 | Metazoa_SRP | 1.221075219 | 2.596056056 | 2.04902232 | 1.755997125 |
| ENSDART00000163039 | fgfrl1b | -0.263701101 | -0.339844216 | -0.261709078 | -0.264179912 |
| ENSDART00000163042 | si:dkey-16p6.2 | 0.770521119 | 0.592959463 | 0.681550899 | 0.554585181 |
| ENSDART00000163057 | FP101882.1 | -0.031169748 | -0.097998763 | -0.346688618 | -0.474056029 |
| ENSDART00000163075 | zgc:173552 | 0.365012264 | 0.677919016 | 1.138040868 | 0.74357363 |
| ENSDART00000163077 | pcdh1g22 | -0.379602696 | -0.085601773 | 0.347287318 | 0.30809701 |
| ENSDART00000163089 | BX649411.2 | 0.422281392 | 0.001934371 | -0.524390062 | -0.335409314 |
| ENSDART00000163093 | lrp12 | -0.07747456 | -0.233717712 | -0.371900586 | -0.129203418 |

| | | | | | |
|---|---|---|---|---|---|
| ENSDART00000163096 | gpr22b | -1.106969479 | -1.366483735 | -0.599957035 | 0.051341663 |
| ENSDART00000163106 | cntnap5l | -0.230959418 | -0.661142608 | -0.469168415 | -0.076363144 |
| ENSDART00000163132 | rab11fip5a | -0.667316443 | -0.571914956 | -0.021672662 | 0.188839381 |
| ENSDART00000163137 | elovl1a | 1.359201374 | 2.32562135 | 1.99000125 | 0.940099891 |
| ENSDART00000163149 | slco1e1 | 0.176388599 | 0.147707318 | -0.389938785 | -1.215116068 |
| ENSDART00000163174 | si:ch73-42p12.2 | 0.635401366 | 0.972834844 | 0.669544171 | 0.150769788 |
| ENSDART00000163182 | tpm2 | -0.110471418 | -0.35380494 | -0.587423123 | -0.759315105 |
| ENSDART00000163216 | prune2 | -0.414992243 | -0.329741076 | -0.094156508 | 0.003976732 |
| ENSDART00000163238 | stom | -0.421228902 | -0.189947439 | -0.023059486 | -0.114725703 |
| ENSDART00000163243 | dpysl4 | 0.526851432 | 0.969802663 | 1.139409914 | 0.808513895 |
| ENSDART00000163250 | mef2cb | 0.361224616 | -0.300957895 | -0.57227951 | -0.080214844 |
| ENSDART00000163305 | fxyd6l | -0.209100518 | -0.188713869 | -0.620315584 | -0.44198331 |
| ENSDART00000163310 | exosc4 | -1.250232551 | -1.071141483 | -0.30340673 | -0.290327378 |
| ENSDART00000163327 | rpl36 | 0.36660589 | 0.401460063 | 0.099733249 | -0.024083137 |
| ENSDART00000163337 | wu:fj30f06 | 0.562927267 | 0.339612631 | 0.922878619 | 0.481097928 |
| ENSDART00000163352 | impg2b | -0.050961847 | -0.309849144 | -0.745225088 | -0.302394708 |
| ENSDART00000163353 | lrp2a | -0.727729459 | -1.488145344 | -2.965914229 | -0.433516037 |
| ENSDART00000163355 | gria2a | -0.37007276 | -0.57694131 | -0.293854127 | 0.240567837 |
| ENSDART00000163367 | TFR2 | 0.892525381 | 1.842645951 | 2.060867557 | 1.063290775 |
| ENSDART00000163370 | si:rp71-36a1.2 | 0.777729217 | 1.232972386 | 1.065047677 | 0.579071639 |
| ENSDART00000163380 | si:dkey-22o22.2 | -0.500713087 | -0.482673389 | -0.426086075 | -0.212897339 |
| ENSDART00000163391 | si:ch211-149k12.3 | 0.507081525 | 0.32718269 | 0.316220639 | 0.200966933 |
| ENSDART00000163394 | dgke | -0.154264184 | -0.257555682 | -0.346312178 | -0.3002364 |
| ENSDART00000163398 | pla2g15 | 0.617752366 | 0.443695085 | 0.159097428 | 0.08811481 |
| ENSDART00000163407 | slc20a1a | 0.970510209 | 0.576996952 | 0.406775549 | -0.419444699 |
| ENSDART00000163441 | limk1a | 1.134897491 | 0.940120128 | 1.036684953 | 0.715200814 |
| ENSDART00000163445 | zgc:109982 | 0.178949347 | -0.334063834 | -0.837985378 | -0.307869575 |
| ENSDART00000163463 | | 0.439033688 | 0.673311945 | 0.275086265 | 0.11827711 |
| ENSDART00000163491 | reep6 | -1.146193953 | -0.500723418 | -0.964434532 | -0.007386228 |
| ENSDART00000163516 | BX927374.1 | -0.474896067 | -0.084073015 | -0.358098021 | -0.301541481 |
| ENSDART00000163519 | CABZ01050166.1 | -0.484053035 | -0.532447227 | -0.143157501 | 0.191505756 |
| ENSDART00000163521 | si:ch1073-83n3.2 | -0.135028227 | -0.352956778 | -0.389044273 | -0.105210001 |
| ENSDART00000163523 | sez6l | -0.314616095 | -0.257996779 | -0.399166894 | -0.159678617 |
| ENSDART00000163526 | mansc1 | 0.060022776 | 0.316271416 | 0.600176682 | 0.64715281 |
| ENSDART00000163529 | FAM126B | -0.519426512 | -0.370293712 | 0.022402011 | 0.032038169 |
| ENSDART00000163531 | CU929294.1 | -0.283562448 | -0.225727022 | -0.174388489 | -0.430860144 |
| ENSDART00000163539 | scrt1a | -0.576475687 | -0.187048661 | 0.063865515 | 0.398969662 |
| ENSDART00000163565 | plcl2 | -0.007453137 | -0.119106713 | -0.251934099 | -0.322673112 |
| ENSDART00000163568 | kcnc1a | -0.672590813 | -0.97944076 | -0.78875743 | -0.1747006 |
| ENSDART00000163577 | fnbp1b | 0.973038753 | 0.902992088 | 0.678793665 | 0.275094435 |
| ENSDART00000163582 | desi1b | 0.334416001 | 0.338516182 | 0.11270838 | -0.034533924 |
| ENSDART00000163585 | csf1ra | 0.499534778 | 0.258389647 | 0.175653195 | -0.286108065 |
| ENSDART00000163597 | CABZ01029366.1 | -0.380235128 | -0.253799187 | -0.219703335 | -0.156720283 |
| ENSDART00000163609 | mrps16 | -0.312251198 | -0.078513731 | -0.202471055 | -0.117952357 |
| ENSDART00000163612 | gabrb4 | -0.445185281 | -0.610684009 | -0.211332691 | 0.127493005 |
| ENSDART00000163616 | homer1b | -0.487597805 | -0.239525084 | -0.239441952 | -0.083482271 |
| ENSDART00000163622 | kirrel3l | -0.615947415 | -0.493751402 | -0.207879997 | 0.127527609 |
| ENSDART00000163635 | erbb4b | -0.423431984 | -0.772483288 | -0.24962409 | 0.07826798 |
| ENSDART00000163656 | CAMK2N1 | -0.350066285 | -0.376343666 | -0.297112769 | 0.007693675 |
| ENSDART00000163668 | si:ch1073-392o20.2 | 0.543875349 | 0.345011244 | 0.082952674 | -0.090250505 |
| ENSDART00000163669 | cplx2 | -0.402497387 | -0.424820216 | -0.158797592 | 0.046498934 |

| | | | | | |
|---|---|---|---|---|---|
| ENSDART00000163677 | | -0.639905731 | -0.179601491 | 0.106036319 | 0.34272584 |
| ENSDART00000163680 | hmgcs1 | 0.250779539 | 0.717953967 | 2.106714123 | 1.525845318 |
| ENSDART00000163724 | SLA (1 of many) | 1.207342807 | 0.416124608 | 0.383103847 | 0.193096581 |
| ENSDART00000163728 | dlg1l | -0.038220111 | -0.173015233 | -0.306246457 | -0.06107777 |
| ENSDART00000163741 | pwwp2b | 0.088274726 | 0.335331052 | 0.428666425 | 0.318380002 |
| ENSDART00000163753 | kcnc2 | -0.67860628 | -0.849972581 | -0.373023023 | -0.287955655 |
| ENSDART00000163791 | acss2l | -0.694063426 | -0.500876422 | -0.115488475 | -0.045115535 |
| ENSDART00000163793 | slitrk6 | -0.404346501 | -0.384982132 | -0.24261489 | -0.144542921 |
| ENSDART00000163794 | wt1b | 0.454982734 | 0.330694147 | 0.317167947 | 0.300420298 |
| ENSDART00000163822 | glula | 0.714593131 | 0.414537751 | 0.09156337 | 0.359301323 |
| ENSDART00000163867 | gnb1b | -0.511340157 | -0.236747213 | 0.117718298 | 0.111335872 |
| ENSDART00000163870 | si:dkey-51d8.3 | 0.628993426 | 0.565032034 | 0.497692882 | 0.102165836 |
| ENSDART00000163882 | si:zfos-932h1.2 | 0.498808647 | 0.465594286 | 0.502820055 | 0.062918681 |
| ENSDART00000163892 | ldha | -0.444310122 | -0.296598203 | -0.299673411 | -0.138453042 |
| ENSDART00000163897 | lgi1b | -0.358188713 | -0.274223364 | -0.009680681 | 0.361584537 |
| ENSDART00000163903 | kcna2b | -1.041609083 | -1.150732339 | -0.278604653 | 0.240702332 |
| ENSDART00000163908 | rnasekb | -0.236636314 | -0.265732137 | -0.418465587 | -0.246757364 |
| ENSDART00000163909 | sepw1 | 1.561335076 | 4.635912393 | 4.23829642 | 2.739511684 |
| ENSDART00000163930 | znfx1 | 0.194042843 | 0.097168789 | 1.523223234 | 0.201252376 |
| ENSDART00000163935 | med30 | -0.198525489 | -0.227576976 | -0.505162548 | -0.348410732 |
| ENSDART00000163951 | plppr1 | -0.393500272 | -0.316896098 | -0.148613826 | 0.078198855 |
| ENSDART00000163952 | zgc:110045 | -0.36736834 | -0.55642828 | -0.201308362 | -0.286026212 |
| ENSDART00000163965 | brdt | -0.081678017 | -0.160102967 | -0.300823041 | -0.066034281 |
| ENSDART00000163976 | CABZ01069287.1 | 0.247964945 | -0.272161916 | -0.664208233 | -0.362164102 |
| ENSDART00000163998 | rps6ka3a | 0.01131102 | -0.11493059 | -0.372925255 | -0.200625555 |
| ENSDART00000164015 | zgc:66483 | -0.233929761 | -0.339147918 | -0.342271085 | -0.145997916 |
| ENSDART00000164016 | kif16ba | 0.136253596 | 0.020087235 | 0.515201001 | 0.413972136 |
| ENSDART00000164038 | SAMD14 | -0.096477793 | 0.217514922 | 0.565397199 | 0.769428937 |
| ENSDART00000164044 | CR339041.2 | 1.41667761 | 2.386461772 | 3.081137029 | 2.622909743 |
| ENSDART00000164055 | cap2 | 0.292123593 | 0.450008213 | 0.705800498 | 0.580755871 |
| ENSDART00000164067 | coq4 | -0.217054082 | -0.203935963 | -0.344175456 | -0.073829043 |
| ENSDART00000164086 | slc25a42 | -0.002109364 | -0.020496378 | -0.265228332 | -0.112125599 |
| ENSDART00000164095 | scpp8 | 1.980889265 | 1.287952222 | 0.276966611 | -1.560257402 |
| ENSDART00000164097 | DRAP1 (1 of many) | -0.372264093 | -0.35663103 | -0.955802153 | -1.03240984 |
| ENSDART00000164102 | cirbpa | 0.225987764 | 0.314210398 | 0.241597292 | 0.01082888 |
| ENSDART00000164107 | mex3b | 2.977404608 | 3.048404514 | 3.190626187 | 2.69620274 |
| ENSDART00000164112 | si:dkey-191g9.7 | -0.2514272 | -0.278147864 | -0.225636717 | -0.211416033 |
| ENSDART00000164113 | cpeb1a | 1.336706075 | 1.190402311 | 1.006023313 | 0.300918459 |
| ENSDART00000164114 | grb2a | -0.50918281 | -0.254702947 | 0.293231191 | 0.639895729 |
| ENSDART00000164121 | mboat2b | -0.365653488 | -0.229675996 | -0.118902893 | -0.108744854 |
| ENSDART00000164129 | asic4b | -0.92148029 | -1.144921239 | -0.577321683 | -0.304835302 |
| ENSDART00000164139 | nhsb | -0.370661862 | -0.378323584 | -0.044398497 | -0.049874132 |
| ENSDART00000164141 | dmd | -0.285273333 | -3.746442585 | -0.310755288 | -0.137191678 |
| ENSDART00000164149 | si:ch211-272n13.3 | -0.267513209 | -0.389121708 | -0.597494594 | -0.154083189 |
| ENSDART00000164160 | acss1 | 0.721950379 | 0.415946373 | -0.236554542 | -0.337079704 |
| ENSDART00000164161 | osbpl1a | 0.351337908 | 0.311134875 | -0.016374257 | -0.257205353 |
| ENSDART00000164163 | si:dkey-202l22.3 | -0.676171429 | -1.127391316 | 0.216542828 | 0.638796234 |
| ENSDART00000164175 | si:dkey-19c16.12 | 0.421774029 | 0.293774627 | 0.297672563 | 0.254989754 |
| ENSDART00000164178 | prrt2 | -0.547052118 | -0.283537507 | 0.297793332 | 0.499767197 |
| ENSDART00000164190 | ksr2 | -0.449890716 | -0.587945856 | -0.131638865 | 0.158651331 |
| ENSDART00000164198 | si:dkey-102m7.3 | 0.949739085 | 1.411170301 | 0.641678924 | 0.378323778 |

| | | | | | |
|---|---|---|---|---|---|
| ENSDART00000164204 | ubac2 | 3.978074982 | 4.357970342 | 4.255380594 | 4.157698498 |
| ENSDART00000164207 | impdh2 | 0.582365155 | 0.675083436 | 1.006708284 | 0.417204588 |
| ENSDART00000164210 | SI | 1.002064064 | 1.522108486 | 0.992417222 | 0.446482404 |
| ENSDART00000164218 | AKAP13 (1 of many) | 0.310415201 | 0.00930034 | -0.071769787 | 0.052155489 |
| ENSDART00000164282 | si:zfos-1011f11.1 | 1.204368625 | 1.598509532 | 2.613199623 | 1.667018794 |
| ENSDART00000164298 | RNF122 | 0.037145597 | 0.153249015 | 0.432199178 | 0.405886045 |
| ENSDART00000164305 | dicp1.1 | 0.970381914 | 0.35730766 | 0.127672979 | 0.007978514 |
| ENSDART00000164311 | mfsd2aa | 0.188250738 | 0.92622451 | 1.277563791 | 0.357720797 |
| ENSDART00000164326 | si:ch73-119p20.1 | -0.36329758 | -0.388299747 | 0.050179382 | 0.305653209 |
| ENSDART00000164328 | mical3b | 0.181716224 | -0.808395946 | -0.697708655 | -4.388355255 |
| ENSDART00000164353 | lsg1 | 0.428728295 | -3.620309582 | -3.671353201 | -0.354626047 |
| ENSDART00000164359 | rpl24 | 0.373088629 | 0.328659722 | 0.253068013 | 0.037812576 |
| ENSDART00000164361 | gcgra | -0.404304946 | -0.520606663 | -0.179928762 | -0.009148827 |
| ENSDART00000164390 | chmp1a | 0.019722961 | 0.342209851 | 0.202368709 | 0.087357775 |
| ENSDART00000164392 | lrrc20 | 0.934799406 | 1.852088866 | 1.534338919 | 1.07983234 |
| ENSDART00000164440 | si:ch211-195b11.3 | 0.887240489 | 0.373309939 | 0.100891486 | -0.191092546 |
| ENSDART00000164454 | plbd1 | 0.523653463 | 0.489772378 | 0.050282092 | -0.147404491 |
| ENSDART00000164456 | ftr50 | 0.637566129 | 0.360052404 | 0.566672785 | 0.193605767 |
| ENSDART00000164471 | CABZ01084230.2 | 0.442580477 | 0.587031258 | 0.401618345 | 0.289449509 |
| ENSDART00000164473 | MAP7 | -0.064645672 | -0.140135551 | -0.331730838 | -0.206670905 |
| ENSDART00000164484 | dtnba | 0.785988532 | 0.671033087 | 0.445356623 | 0.140878127 |
| ENSDART00000164506 | dlg1l | -0.401430092 | -0.615035177 | -0.528577315 | -0.282785588 |
| ENSDART00000164543 | kalrna | 0.833553496 | 1.079150473 | 0.963779963 | 0.661385395 |
| ENSDART00000164563 | elmo1 | -0.350954029 | -0.242629157 | -0.047540163 | 0.212201861 |
| ENSDART00000164566 | akt3a | 0.045255643 | -0.432188612 | -0.498284714 | -0.321504061 |
| ENSDART00000164581 | galr2b | -0.842297369 | -0.875318855 | -0.561173438 | -0.495794792 |
| ENSDART00000164585 | mvda | -0.034972011 | 1.767545823 | 1.673143467 | 0.929274175 |
| ENSDART00000164597 | si:ch73-127m5.1 | -0.712165198 | -0.239448184 | 0.59846121 | 0.690834997 |
| ENSDART00000164609 | si:ch211-126j24.1 | -0.283466546 | -0.277532239 | -0.146779223 | -0.04114819 |
| ENSDART00000164612 | MYH7 (1 of many) | 0.128311883 | 3.589180748 | 4.208970977 | 3.051349831 |
| ENSDART00000164621 | ndrg4 | -0.368994826 | -0.243279912 | -0.209623349 | -0.144971204 |
| ENSDART00000164623 | ptn | -0.285740219 | -0.301123559 | -0.395585979 | -0.169574914 |
| ENSDART00000164647 | slc12a2 | -0.231168401 | 0.356367716 | 0.84043135 | 0.654792579 |
| ENSDART00000164650 | pdk3b | -0.133813888 | -0.219494437 | -0.474289524 | -0.225858079 |
| ENSDART00000164653 | chl1b | 0.395134275 | 0.182602935 | 0.205655131 | 0.15670367 |
| ENSDART00000164658 | si:ch211-225h24.2 | 0.868137568 | 0.716622061 | 0.306228611 | -0.52736976 |
| ENSDART00000164663 | aclya | -0.268277102 | 0.066826112 | 0.215179278 | 0.06328981 |
| ENSDART00000164670 | frem2b | -0.279819948 | -0.158818402 | -0.300273376 | -0.936234223 |
| ENSDART00000164692 | dctn4 | -0.108443255 | 0.034030376 | 0.494831818 | 0.293587926 |
| ENSDART00000164693 | CT583672.1 | -0.40975231 | -0.465896033 | -0.295925202 | -0.161007007 |
| ENSDART00000164695 | arl3l1 | -0.208201897 | -0.292220291 | -0.610724074 | -0.376046469 |
| ENSDART00000164700 | sptbn1 | -0.093232375 | 0.007130054 | 0.336225009 | 0.189868537 |
| ENSDART00000164711 | NFATC2 (1 of many) | 0.774520656 | 0.760515741 | 0.734404519 | 0.027133082 |
| ENSDART00000164712 | cobll1b | -0.06356256 | -0.113578182 | -0.259720894 | -0.442027325 |
| ENSDART00000164726 | si:dkey-200c24.1 | -0.33778778 | -0.588766562 | -0.504539854 | -0.492073189 |
| ENSDART00000164729 | SBSPON | -0.213183167 | -0.151008123 | -0.380923154 | -0.573582751 |
| ENSDART00000164733 | sept15 | -0.353566412 | -0.328984592 | -0.353599969 | -0.269459941 |
| ENSDART00000164759 | cntn4 | -0.356168225 | -0.879631419 | -0.511604399 | -0.608850913 |
| ENSDART00000164771 | ppfibp1a | 0.181979482 | 0.06403475 | 0.045243081 | -0.766863631 |
| ENSDART00000164773 | CABZ01072036.1 | -0.106672866 | -0.494009284 | -0.613761036 | -0.476277849 |
| ENSDART00000164791 | fkbp3 | -0.300438451 | -0.231059822 | -0.287267966 | -0.464270989 |

| | | | | | |
|---|---|---|---|---|---|
| ENSDART00000164792 | cfap74 | -0.15899724 | -0.307564837 | -0.482816939 | -0.375147114 |
| ENSDART00000164805 | camk2b2 | -1.031859946 | -0.873779607 | -0.527330652 | -0.617230087 |
| ENSDART00000164809 | si:ch73-233f7.5 | 0.260792098 | 0.492638431 | 0.828767158 | 0.768289306 |
| ENSDART00000164810 | ano2 | 0.191725223 | -0.262322186 | -0.177000117 | -0.045593369 |
| ENSDART00000164816 | cnr1 | 0.971414159 | 0.97686046 | 0.907214473 | -0.258209562 |
| ENSDART00000164844 | rictora | -0.308713132 | -0.348726405 | -0.242009414 | -0.189747444 |
| ENSDART00000164853 | cnp | 1.119904973 | 1.074260191 | 2.53120974 | 1.692544703 |
| ENSDART00000164855 | crebl2 | -0.127345071 | -0.2122205 | -0.432997567 | -0.264872881 |
| ENSDART00000164879 | slc8a2b | -0.298961891 | -0.426517442 | -0.175774312 | 0.049460205 |
| ENSDART00000164890 | pxdc1a | -0.553855303 | -0.268569574 | -0.515208528 | -0.163588555 |
| ENSDART00000164891 | trim25 | 1.17805401 | 1.473587562 | 2.17710407 | 1.406205223 |
| ENSDART00000164902 | ca9 | 0.158296773 | 0.100302393 | -0.407653731 | -0.624825129 |
| ENSDART00000164904 | nrp1a | 0.238338546 | 0.288203349 | 0.415887875 | 0.403021552 |
| ENSDART00000164928 | mab21l1 | -0.236289538 | -0.264159199 | -0.014378888 | 0.140356353 |
| ENSDART00000164979 | ARHGAP44 (1 of many) | -0.298736282 | -0.611440452 | -0.190082373 | -0.04950856 |
| ENSDART00000164982 | cdh4 | -0.415355609 | -0.489911794 | 0.041403227 | 0.334639192 |
| ENSDART00000164983 | anapc15 | -0.036185378 | -0.198619926 | -0.408385072 | -0.233599439 |
| ENSDART00000164988 | bod1l1 | 0.085003082 | 0.094605568 | 0.323418829 | 0.18061312 |
| ENSDART00000164989 | si:ch211-121j5.4 | -2.541662275 | -0.597024815 | -0.157115726 | -0.615748868 |
| ENSDART00000165000 | zgc:136767 | 1.074177299 | 1.362099519 | 1.806009349 | 1.052151095 |
| ENSDART00000165002 | aqp11 | -0.052004039 | -0.030290396 | -0.282964979 | -0.265886582 |
| ENSDART00000165004 | gria3b | -0.410652366 | -0.638891942 | -0.274205904 | 0.132979921 |
| ENSDART00000165006 | hpca | -0.709011759 | -0.754814005 | -0.136865422 | -0.15729929 |
| ENSDART00000165018 | cdc42se2 | -0.053295614 | -0.137825766 | -0.478383249 | -0.349419762 |
| ENSDART00000165021 | ndrg2 | -0.572041099 | -0.272188031 | 0.107233459 | 0.07090542 |
| ENSDART00000165030 | si:dkeyp-9d4.3 | -0.136094032 | -0.353586884 | -0.269174482 | -0.139574898 |
| ENSDART00000165031 | nr2f6b | 0.621146984 | 0.406827879 | 0.54917158 | 0.373267529 |
| ENSDART00000165049 | impg2b | -0.010489295 | -0.310856751 | -0.727501593 | -0.334825186 |
| ENSDART00000165058 | rims2a | -0.759006031 | -0.607036719 | -0.257997898 | -0.111766909 |
| ENSDART00000165065 | uqcr10 | -0.222678585 | -0.318669731 | -0.412855116 | -0.201529958 |
| ENSDART00000165066 | eif2ak2 | 0.145136508 | 0.290335878 | 1.008511718 | 0.199274727 |
| ENSDART00000165082 | ppp1r1b | -0.592550768 | -0.433249399 | 0.132918309 | 0.038849346 |
| ENSDART00000165097 | unm_sa1614 | -0.093711156 | -0.459181727 | -0.424406225 | -0.126303047 |
| ENSDART00000165108 | jph3 | -0.254573235 | -0.426555215 | 0.040987467 | 0.08534723 |
| ENSDART00000165115 | adcy3a | -0.345722559 | -0.440118918 | -0.380340005 | -0.11083624 |
| ENSDART00000165120 | purab | -0.246648902 | -0.188751457 | -0.157104845 | -0.035763175 |
| ENSDART00000165124 | si:ch73-213k20.5 | -0.366310901 | -0.428402723 | -0.397554725 | -0.148030091 |
| ENSDART00000165141 | elavl4 | 1.008784658 | 0.753903544 | 1.037731482 | 0.21557706 |
| ENSDART00000165147 | MFSD3 | -0.380785464 | -0.635551022 | -0.165628582 | -0.053496107 |
| ENSDART00000165156 | sept15 | -0.316165192 | -0.356420975 | -0.295749689 | -0.213768729 |
| ENSDART00000165158 | iqsec3a | -0.287263936 | -0.339420666 | -0.160997893 | 0.073213089 |
| ENSDART00000165159 | dscaml1 | -0.229621803 | -0.384977161 | -0.055090744 | 0.170582623 |
| ENSDART00000165186 | si:dkey-33i11.9 | 1.335261857 | 1.027980768 | 0.465372707 | 0.114592431 |
| ENSDART00000165195 | CU633762.1 | -0.318810448 | -0.187254152 | -0.150226726 | -0.009770113 |
| ENSDART00000165199 | mapre2 | 0.262487966 | 1.260089875 | 1.425017722 | 0.706527865 |
| ENSDART00000165201 | pacsin3 | -0.207834592 | -0.16294471 | -0.3385586 | -0.115392716 |
| ENSDART00000165207 | fam160a1b | -0.089233769 | -0.189976289 | -0.269606776 | -0.116139265 |
| ENSDART00000165213 | caskb | -0.0150989 | 0.201636443 | 0.432584514 | 0.386918381 |
| ENSDART00000165216 | dph5 | 0.422816122 | 0.515360344 | 0.448987609 | 0.161864292 |
| ENSDART00000165223 | pbx1b | -0.243991197 | -0.377373898 | -0.18693501 | 0.141565589 |
| ENSDART00000165225 | prkar1ab | -0.220219796 | -0.315739009 | -0.190831632 | -0.05456146 |

| | | | | | |
|---|---|---|---|---|---|
| ENSDART00000165228 | kif5aa | -0.119845449 | 0.354985416 | 0.468393446 | 0.466605496 |
| ENSDART00000165230 | map4k4 | -0.058109942 | 0.098582575 | 0.424133469 | 0.341075529 |
| ENSDART00000165290 | cyb5a | 0.19405994 | 0.064877953 | -0.350823438 | -0.576368917 |
| ENSDART00000165292 | nsmfb | -0.267190978 | -0.447188182 | -0.263868703 | -0.115307671 |
| ENSDART00000165308 | me2 | -0.16075229 | -0.327815721 | -0.451710296 | -0.323851706 |
| ENSDART00000165309 | pmt | 1.560743592 | 2.051643174 | 1.965781873 | 1.006007351 |
| ENSDART00000165318 | thsd7bb | -0.173014975 | 0.179455909 | 1.051340912 | 1.022924373 |
| ENSDART00000165326 | adgrl2a | -0.346797592 | -1.122813298 | -2.2772027 | -0.600399475 |
| ENSDART00000165333 | si:ch211-207l14.1 | -0.144575592 | -0.337549275 | -0.714109722 | -0.444634079 |
| ENSDART00000165342 | si:ch211-93f2.1 | -0.158550393 | 0.117683559 | -0.256320731 | -0.510696237 |
| ENSDART00000165370 | nxph2b | -0.350612818 | -0.344729721 | -0.406615862 | -0.05363573 |
| ENSDART00000165400 | slc1a2b | -0.437547383 | -0.373563248 | -0.176045295 | -0.026222695 |
| ENSDART00000165411 | | -0.599585032 | -0.699498667 | -0.398460986 | -0.09881633 |
| ENSDART00000165420 | si:dkey-161j23.5 | -0.418752064 | -0.09080604 | 0.112517016 | 0.202865451 |
| ENSDART00000165423 | abcc8b | -0.169557865 | -0.54863705 | -0.322687745 | -0.280461618 |
| ENSDART00000165425 | aak1a | -0.009756971 | -0.269537246 | -0.43668389 | -0.03607957 |
| ENSDART00000165427 | myt1b | 0.19969921 | 0.396205375 | 0.347986667 | 0.198976782 |
| ENSDART00000165433 | rgs2 | 0.779425284 | 0.385417705 | 0.555491331 | -0.186899017 |
| ENSDART00000165437 | si:dkey-26m3.3 | 0.220558358 | 0.221546162 | 0.282286139 | 0.194663631 |
| ENSDART00000165443 | zgc:153615 | -0.585165411 | -0.400257189 | -0.265863973 | -0.020005469 |
| ENSDART00000165448 | rims2b | -0.321612136 | -0.490793078 | -0.295423658 | 0.030315799 |
| ENSDART00000165453 | epb41l3a | -0.480811799 | 0.184033911 | 0.640683351 | 0.633983333 |
| ENSDART00000165454 | CU915762.1 | 0.350263787 | 0.108189136 | -0.057165889 | 0.160983657 |
| ENSDART00000165472 | snx2 | 0.34033168 | -0.040635813 | 0.394688984 | 0.105073801 |
| ENSDART00000165479 | has3 | 0.356722134 | 0.209377149 | 0.150402808 | 0.132473433 |
| ENSDART00000165484 | jpt1a | 0.363755224 | 1.1796439 | 1.211655345 | 0.942198247 |
| ENSDART00000165486 | rps18 | 0.252319024 | 0.42379942 | 0.281645226 | -0.028370317 |
| ENSDART00000165491 | BX294383.1 | 0.352047768 | 0.065820831 | 0.027639622 | 0.019046581 |
| ENSDART00000165499 | rps17 | 0.504096498 | 0.637791027 | 0.385574489 | 0.024446186 |
| ENSDART00000165541 | map7d1b | 0.274410575 | 0.322558005 | 0.501220001 | 0.438496507 |
| ENSDART00000165542 | si:dkeyp-72e1.9 | 0.115590472 | -0.201422352 | -0.774050016 | -0.351374892 |
| ENSDART00000165547 | CABZ01045062.1 | -0.214219167 | -0.28679968 | -0.112596272 | 0.066374108 |
| ENSDART00000165548 | ap2m1a | -0.100883554 | -0.260493779 | -0.290981943 | -0.160008883 |
| ENSDART00000165557 | BX088524.3 | -0.275785171 | -0.331787981 | -0.469187171 | -0.127928431 |
| ENSDART00000165570 | rgs3a | -0.252183908 | -0.25277418 | -0.390798302 | -0.347427277 |
| ENSDART00000165572 | zgc:162193 | 0.654407013 | 0.606984165 | 0.583589232 | 0.369009896 |
| ENSDART00000165594 | CT030188.1 | 0.949647945 | 1.541943795 | 1.047931527 | 0.898450066 |
| ENSDART00000165609 | barhl2 | -0.31272019 | -0.374018735 | 0.126371762 | 0.224852149 |
| ENSDART00000165628 | il4r.1 | 2.521820035 | 2.797213431 | 2.312532052 | 1.674117298 |
| ENSDART00000165638 | pax10 | -0.459647113 | -0.417992165 | -0.200180992 | -0.120970793 |
| ENSDART00000165654 | atp1b2a | -0.572829136 | -0.493634763 | -0.065881564 | 0.075479325 |
| ENSDART00000165656 | mxd3 | 0.030444632 | 0.928898979 | 1.014066924 | 0.787179484 |
| ENSDART00000165659 | epb41l3a | -0.249067696 | -0.637410719 | -0.072556493 | -0.093775549 |
| ENSDART00000165680 | ntn4 | 0.240096357 | 0.070794746 | -0.41629651 | -0.558332982 |
| ENSDART00000165698 | pbx1a | -0.205212048 | -0.330154536 | 0.047550564 | 0.321002436 |
| ENSDART00000165710 | gpbp1l1 | -0.164201122 | -0.182753901 | -0.290062625 | -0.125147623 |
| ENSDART00000165715 | BX284638.1 | 0.544886732 | 0.81514457 | 1.035521411 | 0.49952411 |
| ENSDART00000165735 | mdm4 | 0.677008086 | 0.457427602 | 0.365768719 | 0.56900659 |
| ENSDART00000165743 | gabrg2 | -0.320626657 | -0.470320689 | -0.258833602 | 0.01857369 |
| ENSDART00000165744 | kif1b | -0.018685925 | -0.126032816 | 0.340319561 | 0.347744455 |
| ENSDART00000165757 | pax6b | -1.030621281 | -0.736504391 | -0.510180699 | -0.112500923 |

| | | | | | |
|---|---|---|---|---|---|
| ENSDART00000165774 | pax6a | -0.382942423 | -0.218376905 | 0.094924321 | 0.282463268 |
| ENSDART00000165775 | nlrc3 | 2.263794952 | 2.415644973 | 1.694313933 | 0.888951658 |
| ENSDART00000165785 | pcdh10a | -0.2607204 | -0.385620436 | -0.040787194 | 0.144409091 |
| ENSDART00000165824 | setdb1b | 0.086359323 | 0.217956862 | 0.320141564 | 0.188338727 |
| ENSDART00000165835 | ddx54 | 0.300757648 | 0.157122409 | 0.421066488 | 0.21493331 |
| ENSDART00000165864 | acox3 | -0.089913641 | -0.178492527 | -0.263049665 | -0.100985253 |
| ENSDART00000165875 | csnk1g1 | 0.004513388 | 0.131360351 | 0.256793811 | 0.293233915 |
| ENSDART00000165877 | purg | -0.159238984 | -0.087175961 | 1.048436006 | 0.831059287 |
| ENSDART00000165883 | opn9 | -0.186957867 | -0.12243997 | -0.333236797 | -0.10321317 |
| ENSDART00000165887 | flvcr1 | -0.333518999 | -0.053749594 | 0.204875088 | 0.015969559 |
| ENSDART00000165898 | gbe1b | -0.311269447 | -0.395300505 | -0.303280984 | -0.17462698 |
| ENSDART00000165903 | slmapa | -5.974312545 | -3.688098031 | -4.583524263 | -0.943141088 |
| ENSDART00000165912 | si:ch73-380n15.2 | -0.521676191 | -0.515528635 | -0.335328887 | -0.345110245 |
| ENSDART00000165920 | nucb1 | 0.353439048 | 0.296422234 | 0.314532166 | -0.093049018 |
| ENSDART00000165932 | sik1 | 1.156253383 | 0.676084919 | 0.687880039 | 1.28219737 |
| ENSDART00000165938 | CU466240.1 | 0.428025361 | 0.165679287 | 0.139754855 | 0.298251046 |
| ENSDART00000165943 | fam102aa | 0.068136883 | 0.160885629 | -0.464613828 | -0.136577362 |
| ENSDART00000165949 | fahd2a | 0.346683888 | 0.20577135 | -0.135781443 | -0.151237654 |
| ENSDART00000165955 | zhx3 | -0.011624436 | -0.109014479 | 0.013312322 | 0.351264725 |
| ENSDART00000165973 | npnt | -0.201530974 | -0.119421961 | -0.271951683 | -0.242671185 |
| ENSDART00000165974 | agla | -0.253110547 | -0.164938699 | -0.089751197 | 0.189815825 |
| ENSDART00000165979 | sncgb | -0.522103986 | -0.516181722 | -0.379782599 | -0.140861461 |
| ENSDART00000165987 | DST | 0.207553536 | 0.19383587 | 0.337156934 | 0.233508941 |
| ENSDART00000165991 | lect1 | -3.06408831 | -0.430314435 | 0.807667083 | 0.659541685 |
| ENSDART00000165993 | f3a | 1.175718098 | 1.946056838 | 1.259834727 | 0.15077318 |
| ENSDART00000166025 | irs2a | -0.31567938 | -0.330763757 | -0.333387068 | -0.269837586 |
| ENSDART00000166027 | trpc1 | -0.303725684 | -0.42864481 | -0.334157939 | -0.231516525 |
| ENSDART00000166028 | mcf2la | -0.298550829 | -0.194248433 | -0.224063256 | -0.02772208 |
| ENSDART00000166040 | sh3bp5b | 0.359354612 | 1.105816025 | 1.101214783 | 0.69319763 |
| ENSDART00000166042 | vipr2 | -0.341756427 | -0.277003119 | -0.379732317 | -0.038627099 |
| ENSDART00000166058 | cmah | 1.523029746 | 4.3199312 | 4.414697946 | 2.901009315 |
| ENSDART00000166086 | CABZ01055522.1 | -0.167914645 | -0.1533494 | -0.033990591 | 0.313482965 |
| ENSDART00000166101 | tlr22 | 0.744049575 | 0.528297207 | 0.210656038 | -0.353063092 |
| ENSDART00000166105 | frem1a | -0.140882698 | -0.093532592 | -0.525495905 | -1.018483252 |
| ENSDART00000166110 | itga4 | 0.891623018 | 0.709388891 | 0.523610434 | -0.253853847 |
| ENSDART00000166114 | sema3ab | -0.726704016 | -0.305486009 | -0.172433457 | -0.053903823 |
| ENSDART00000166120 | si:ch211-212d10.1 | 1.398436882 | 1.472487792 | 1.228480368 | 0.254562361 |
| ENSDART00000166135 | zbtb47b | -0.402709337 | -0.409852132 | -0.474957603 | -0.394147628 |
| ENSDART00000166148 | gabra1 | -0.578656795 | -0.788703398 | -0.59715214 | -0.072809071 |
| ENSDART00000166152 | DAB2 (1 of many) | 0.967097608 | 0.419929333 | 0.22593558 | -0.403858057 |
| ENSDART00000166174 | PARP12 | 0.236086028 | 0.041618587 | 1.350827793 | 0.15615804 |
| ENSDART00000166175 | zgc:171534 | 0.978558952 | 0.553291836 | 0.353063819 | -0.014246545 |
| ENSDART00000166177 | impg2b | 0.006257085 | -0.447154942 | -0.735592553 | -0.255871486 |
| ENSDART00000166192 | pik3r5 | 0.977110656 | 0.482953861 | 0.341342874 | -0.116377345 |
| ENSDART00000166209 | wu:fb44b02 | 0.715207055 | 0.457109376 | 0.301906263 | -0.147536436 |
| ENSDART00000166213 | lcorl | -0.114650844 | -0.153895276 | -0.40050268 | -0.037010303 |
| ENSDART00000166224 | smyhc2 | 1.194842223 | 5.367086788 | 6.852910143 | 4.718916943 |
| ENSDART00000166241 | inpp5b | -0.146961161 | -0.217731387 | -0.46875304 | -0.117265313 |
| ENSDART00000166242 | | 0.20194611 | 0.241162188 | 0.512911525 | 0.246398136 |
| ENSDART00000166246 | ssuh2rs1 | 0.375757195 | 0.261052668 | 0.200946491 | -0.016238014 |
| ENSDART00000166254 | gpn2 | 0.085397213 | 0.349689217 | 0.008215986 | -0.050534057 |

| | | | | | |
|---|---|---|---|---|---|
| ENSDART00000166259 | wars | 0.30804143 | 0.479087994 | 0.526413933 | 0.245904587 |
| ENSDART00000166268 | YTHDC2 | 0.015267861 | -0.127181664 | 0.395706164 | 0.317645369 |
| ENSDART00000166274 | phlda1 | -0.962333929 | -0.775569749 | 0.008452009 | 0.660576028 |
| ENSDART00000166308 | cib2 | -0.434731659 | -0.335262372 | -0.470437684 | -0.240772493 |
| ENSDART00000166313 | thrb | -0.193382381 | -0.415286369 | -0.400030228 | -0.136855946 |
| ENSDART00000166317 | mtus1b | -0.441732874 | -0.537161422 | 0.305687832 | 0.223661346 |
| ENSDART00000166324 | ctnnd1 | 0.460487451 | 0.641759187 | 1.004918197 | 0.346719852 |
| ENSDART00000166341 | AL954715.1 | 0.715525501 | 0.583886882 | 0.488572453 | 0.64332223 |
| ENSDART00000166351 | nkrf | -0.332043429 | -0.213944917 | -0.117297764 | -0.03436019 |
| ENSDART00000166372 | si:ch211-222l21.1 | 0.890920299 | 0.959321743 | 0.886208086 | 0.413211736 |
| ENSDART00000166374 | si:dkey-31f5.11 | -0.246069029 | -0.257528931 | -0.445065012 | -0.224168767 |
| ENSDART00000166387 | prickle2a | -0.236957906 | -0.307926948 | -0.968400315 | -0.433283323 |
| ENSDART00000166388 | il6st | 0.326944598 | 0.248050944 | 0.233084281 | -0.066517896 |
| ENSDART00000166395 | fcer1gl | 0.672549379 | 0.370716598 | -0.127334281 | -0.409142814 |
| ENSDART00000166432 | slc2a8 | 0.33290564 | 0.367124099 | 0.471066456 | 0.250147846 |
| ENSDART00000166435 | si:ch73-28h20.1 | -0.187153724 | -0.333881358 | -0.353210004 | -0.031764093 |
| ENSDART00000166449 | pik3r3a | -0.698577911 | -0.3007013 | -0.248394691 | -0.547713549 |
| ENSDART00000166463 | cnot6b | -0.099106983 | -0.163429858 | -0.300924183 | -0.105283807 |
| ENSDART00000166470 | CU855947.1 | -0.432778868 | -0.336734093 | -0.219445916 | -0.079633712 |
| ENSDART00000166496 | cat | 0.249947463 | 0.551180864 | 0.073389354 | -0.18327225 |
| ENSDART00000166502 | satb2 | -0.540440017 | -0.158642442 | 1.216799418 | 1.496340928 |
| ENSDART00000166504 | sod3b | 1.107778201 | 3.874418447 | 1.285382343 | 3.248105088 |
| ENSDART00000166508 | fdft1 | -0.467410589 | -0.45021858 | 0.057865823 | -0.125768039 |
| ENSDART00000166515 | si:dkeyp-57d7.4 | -0.099733783 | -0.492859873 | -1.003446185 | -0.404973513 |
| ENSDART00000166518 | ptprfb | 0.067979508 | -0.359782965 | 0.005118486 | 0.040886074 |
| ENSDART00000166527 | ftr87 | -0.180960216 | -0.104077306 | -0.471569991 | -0.343688512 |
| ENSDART00000166531 | pcdh15b | -0.215971807 | -0.468745045 | -0.336826359 | 0.084344716 |
| ENSDART00000166533 | zgc:123181 | -0.682536387 | -0.078871471 | 0.212564109 | 0.407498801 |
| ENSDART00000166540 | kcnip3b | -0.988549414 | -0.704807394 | -0.278733631 | 0.119291619 |
| ENSDART00000166560 | hpdl | 0.319918962 | 0.542121436 | 0.616518616 | 0.488982996 |
| ENSDART00000166561 | ttc38 | 0.445202505 | 0.288306634 | -0.06639877 | -0.156646921 |
| ENSDART00000166566 | dgkh | -1.438871843 | -1.330275705 | -0.819153793 | -0.068934495 |
| ENSDART00000166575 | ppp3ca | -0.493387865 | -0.574192875 | 0.281204144 | 0.517021392 |
| ENSDART00000166580 | pak1 | 0.254456198 | 0.505135694 | 0.67991388 | 0.523148639 |
| ENSDART00000166587 | rhbdl3 | 1.011972997 | 0.720471419 | 0.019687653 | -0.322773745 |
| ENSDART00000166591 | utp20 | 0.378085419 | 0.246513796 | 0.276819854 | 0.15697444 |
| ENSDART00000166607 | RAPGEF4 (1 of many) | -0.233842979 | 0.163001528 | 0.421327636 | 0.212581301 |
| ENSDART00000166615 | dnm2b | 0.605203252 | 0.413218412 | 0.436607775 | -0.016254619 |
| ENSDART00000166618 | slc43a3a | 0.880026964 | 0.262110574 | 0.534125771 | 0.347202731 |
| ENSDART00000166622 | si:dkey-77g12.1 | 1.256692313 | 2.663525112 | 1.501030031 | -0.081810387 |
| ENSDART00000166632 | si:zfos-741a10.3 | 1.2129476 | 0.572994738 | 0.242126634 | 0.029899239 |
| ENSDART00000166634 | rnf213b | 0.322643691 | -0.072602668 | 2.242716781 | -0.029575481 |
| ENSDART00000166645 | HTR7 (1 of many) | -0.91335189 | -0.965307725 | -0.027473732 | 0.473063601 |
| ENSDART00000166648 | | 0.095481351 | -0.09266769 | 0.456407618 | 0.379239728 |
| ENSDART00000166650 | bsg | -0.333252581 | -0.445374368 | -0.502358398 | -0.336861466 |
| ENSDART00000166659 | lrrn2 | -0.321629641 | -0.404637061 | -0.652990901 | -0.235544369 |
| ENSDART00000166664 | MIDN (1 of many) | 0.609413165 | 0.740169365 | 0.964140975 | 0.212765964 |
| ENSDART00000166681 | frrs1l | -0.4975582 | -0.505651734 | -0.29386319 | -0.136005256 |
| ENSDART00000166690 | gadd45ga | 1.824520476 | 2.195526461 | 0.938941318 | 1.401631316 |
| ENSDART00000166693 | | 0.184413566 | -0.110459993 | 1.581805418 | 0.290855458 |
| ENSDART00000166698 | lrsam1 | 0.035853538 | 0.184404018 | 0.265787714 | 0.064864391 |

| | | | | | |
|---|---|---|---|---|---|
| ENSDART00000166714 | impg2b | -0.072742859 | -0.391473223 | -0.740015258 | -0.360324281 |
| ENSDART00000166724 | myom1b | 0.206144619 | 0.433459222 | 0.498383662 | 0.501410774 |
| ENSDART00000166725 | si:dkey-27j5.9 | 1.657914875 | 0.40212862 | 0.542892045 | 0.141569929 |
| ENSDART00000166730 | slitrk4 | -0.433556166 | -0.537605748 | -0.487948345 | -0.301835783 |
| ENSDART00000166739 | si:zfos-2326c3.2 | 0.356689661 | 0.198077273 | 0.030512595 | -0.103105863 |
| ENSDART00000166755 | tril | 1.481556618 | 0.917097925 | 0.535942919 | 0.363878102 |
| ENSDART00000166756 | mfsd11 | 0.314593735 | 0.230184492 | 0.795700169 | 0.565917277 |
| ENSDART00000166829 | | -0.799969413 | -0.359199835 | -0.712167572 | -2.039500454 |
| ENSDART00000166843 | tom1 | -0.282793851 | -0.146845245 | -0.347923357 | -0.109554429 |
| ENSDART00000166889 | nptna | -0.194507678 | -0.523176341 | -0.288117341 | 0.166715743 |
| ENSDART00000166894 | si:dkey-16p6.1 | -3.041565726 | -0.566863861 | -1.581438015 | -0.461000528 |
| ENSDART00000166904 | AL840638.1 | -0.172150655 | -0.246843461 | -0.35641848 | -0.195598477 |
| ENSDART00000166957 | purba | -0.28509761 | -0.466147452 | -0.146018067 | -0.191067595 |
| ENSDART00000166981 | snx8b | -0.518943385 | -0.495275193 | -0.13677988 | -0.169307474 |
| ENSDART00000166992 | rab6a | 0.863374482 | 0.610837796 | 0.548836905 | 0.594870039 |
| ENSDART00000167001 | cited4b | -0.183655613 | -0.263694381 | -0.243829292 | -0.142076267 |
| ENSDART00000167025 | si:dkey-203a12.7 | 1.830928705 | 3.142170624 | 2.521496169 | 0.613627975 |
| ENSDART00000167032 | tp53rk | 0.299937927 | 0.221076682 | 0.117069539 | 0.10102477 |
| ENSDART00000167035 | mibp2 | 0.52185156 | 0.897496666 | 1.068299106 | 0.608631566 |
| ENSDART00000167040 | pop1 | 0.818813588 | 0.594298576 | 0.588084483 | 0.531688582 |
| ENSDART00000167052 | etv1 | -0.327639473 | -0.469002174 | -0.32895701 | -0.22464313 |
| ENSDART00000167062 | SNORD49 | 0.825973685 | 1.057295844 | 0.696610061 | 0.75374548 |
| ENSDART00000167068 | kifap3b | 0.204750718 | 0.408132617 | 0.569098043 | 0.272936415 |
| ENSDART00000167074 | irf2 | 0.133065149 | 0.222164622 | 0.608761767 | -0.104617341 |
| ENSDART00000167099 | CU459186.5 | 0.808307931 | 1.173553726 | 0.835552808 | 0.510196285 |
| ENSDART00000167117 | si:ch1073-469d17.2 | -0.226355895 | -0.436402154 | -0.568359696 | -0.320345187 |
| ENSDART00000167128 | pargl | 0.159346244 | 0.133314978 | 0.619065571 | -0.065913899 |
| ENSDART00000167132 | isg20l2 | 0.391451776 | 0.35980411 | 0.120415307 | 0.040629954 |
| ENSDART00000167143 | glyr1 | -0.150233074 | -0.208555766 | -0.274679248 | -0.191034867 |
| ENSDART00000167145 | ap1ar | 0.285895558 | 0.368048827 | 0.350688731 | 0.119518374 |
| ENSDART00000167154 | trim9 | -0.216985401 | -0.363840661 | -0.237534447 | -0.020888972 |
| ENSDART00000167164 | MPV17L | -0.408335699 | -0.461800821 | -0.169486828 | -0.082955483 |
| ENSDART00000167177 | ccnf | 1.106246818 | 1.046935028 | 0.737028811 | 0.1071592 |
| ENSDART00000167179 | asf1ba | -0.325814646 | -0.52479315 | -0.773177729 | -0.605632897 |
| ENSDART00000167197 | spata18 | -0.430548228 | -0.388964562 | -0.077452756 | -0.001886549 |
| ENSDART00000167219 | pcdh1g26 | -0.056136761 | 0.084950376 | 0.428942573 | 0.447913813 |
| ENSDART00000167226 | pycr1b | 0.1083633 | 0.252664475 | 0.730734657 | 0.419591074 |
| ENSDART00000167279 | klhl6 | 0.921934193 | 0.563560813 | 0.364277897 | 0.100202901 |
| ENSDART00000167306 | ldb2b | -0.838228207 | -0.415457403 | -0.325734782 | 0.110687477 |
| ENSDART00000167322 | si:ch73-233f7.3 | -0.088252023 | 0.25632847 | 0.562157447 | 0.392412002 |
| ENSDART00000167324 | ebf3a | -0.585714789 | -0.243731213 | 0.563349846 | 0.843370215 |
| ENSDART00000167330 | CU179758.1 | -0.385159517 | -0.273959547 | -0.55992419 | -0.56237529 |
| ENSDART00000167355 | nsfa | -0.252585329 | -0.369691062 | -0.484691854 | -0.137375398 |
| ENSDART00000167359 | dusp27 | 1.665789827 | 1.31173846 | 1.624125812 | 1.13571557 |
| ENSDART00000167388 | vps33a | 0.155328285 | 0.245633599 | 0.343008348 | 0.290252576 |
| ENSDART00000167391 | arhgap21a | -0.385966805 | -0.235029359 | -0.527723306 | -0.308128762 |
| ENSDART00000167423 | si:dkey-242k1.6 | -0.388268724 | -0.337313048 | -0.259026873 | -0.237349722 |
| ENSDART00000167430 | lrrc24 | -0.125265238 | -0.400040145 | -0.093389228 | 0.002539389 |
| ENSDART00000167444 | | -0.313613371 | 0.27829227 | 0.501895099 | 0.307445871 |
| ENSDART00000167449 | sept15 | -0.329936507 | -0.127540958 | -0.290094259 | -0.186050001 |
| ENSDART00000167451 | | -0.147201966 | -0.622186255 | -0.321157937 | -0.196934609 |

| | | | | | |
|---|---|---|---|---|---|
| ENSDART00000167461 | BX897691.1 | 1.652413387 | 2.158428192 | 1.996842668 | 1.949150718 |
| ENSDART00000167464 | GNAZ | -0.341892906 | -0.155068478 | 0.102581405 | 0.079027974 |
| ENSDART00000167468 | prpsap1 | 0.532862733 | 0.121505744 | 0.13944499 | -0.034080691 |
| ENSDART00000167502 | CR855860.1 | -0.377799337 | -0.542912736 | -0.423594114 | -0.115674527 |
| ENSDART00000167506 | scg2a | -0.457333464 | -0.29124415 | -0.144155502 | -0.510044484 |
| ENSDART00000167514 | abca1a | 0.214033988 | 0.130340273 | 0.268396538 | -0.115766571 |
| ENSDART00000167523 | dixdc1b | -0.34575392 | -0.259524065 | -0.191239322 | -0.198083451 |
| ENSDART00000167570 | actr3b | -0.361119463 | -0.449811578 | -0.822009016 | -0.639517901 |
| ENSDART00000167584 | aldh2.1 | -0.136059736 | -0.055428642 | -0.441396328 | -0.687740935 |
| ENSDART00000167612 | rnf34a | -0.125673437 | -0.210904428 | -0.512991238 | -0.24247481 |
| ENSDART00000167613 | hmgcs1 | -0.061825835 | 0.154559892 | 0.893983325 | 0.609306628 |
| ENSDART00000167649 | pik3r3a | -0.582543495 | -0.143058653 | -0.251972796 | -0.459052361 |
| ENSDART00000167660 | pja2 | -0.078344887 | -0.326712885 | -0.382150009 | -0.239022195 |
| ENSDART00000167664 | atxn1a | -0.182914727 | -0.487043114 | -0.17911118 | -0.100136551 |
| ENSDART00000167666 | dnajc21 | 0.569585132 | 0.384796247 | 0.537599141 | 0.160116436 |
| ENSDART00000167667 | FKBP15 (1 of many) | 0.292301754 | 0.132288424 | 0.122963959 | -0.01266455 |
| ENSDART00000167696 | si:ch73-233f7.4 | 0.264747026 | 0.489460093 | 0.625111029 | 0.721718733 |
| ENSDART00000167726 | RYR2 | -0.554726037 | -0.413134766 | 0.099586067 | 0.289722828 |
| ENSDART00000167748 | fbn2b | 1.02333483 | 0.65734038 | 0.922883374 | 0.19244827 |
| ENSDART00000167786 | | 0.801350724 | 0.147008366 | 0.081187044 | 0.322292312 |
| ENSDART00000167818 | atp2b2 | -0.603266889 | -0.926553396 | 0.035397356 | 0.064456748 |
| ENSDART00000167823 | CU651662.1 | 0.332182102 | 0.108414441 | -0.278380094 | -0.999717307 |
| ENSDART00000167824 | timp4.3 | -0.389539203 | -0.013931581 | -0.399789503 | -0.590224284 |
| ENSDART00000167847 | eef1a1l2 | -0.649781637 | -0.501730111 | -0.557800044 | -0.257462283 |
| ENSDART00000167861 | cox4i1l | -0.902856097 | -1.26820767 | -1.375291282 | -0.702334306 |
| ENSDART00000167869 | | -3.349211506 | -1.484996272 | -1.5614357 | -0.542819931 |
| ENSDART00000167873 | baiap2b | -0.183691734 | -0.21009419 | -0.266222434 | -0.177287211 |
| ENSDART00000167879 | Metazoa_SRP | 3.268404605 | 5.790076 | 5.563915837 | 5.137733652 |
| ENSDART00000167881 | | 0.345756532 | 0.215047296 | 0.187007292 | 0.450693648 |
| ENSDART00000167937 | p4hb | 0.35105827 | 0.247669444 | -0.015299318 | -0.184059227 |
| ENSDART00000167947 | CABZ01079480.1 | 0.418778383 | 0.234022686 | 1.236946888 | 0.318728833 |
| ENSDART00000167948 | hcn1 | -0.135796002 | -0.504326543 | -0.514845616 | -0.152463147 |
| ENSDART00000167956 | tspan18a | -0.226187052 | -0.301324705 | -0.40566671 | -0.255395447 |
| ENSDART00000167963 | nrg1 | -0.278608256 | -0.388339035 | -0.420420553 | -0.389329729 |
| ENSDART00000167971 | rps18 | 0.367747716 | 0.388045404 | 0.209564172 | -0.058761163 |
| ENSDART00000167977 | kcnh4b | -0.815041676 | -0.830195363 | -1.200187818 | -0.588368652 |
| ENSDART00000167986 | hadhb | 0.310832473 | 0.238522601 | 0.004510639 | -0.025410475 |
| ENSDART00000167995 | napba | -0.341681302 | -0.334450461 | -0.176497327 | -0.027366162 |
| ENSDART00000168002 | laptm5 | 1.090713638 | 0.609708871 | 0.327355694 | -0.215403696 |
| ENSDART00000168004 | rps18 | 0.800703549 | 0.805799198 | 0.576669388 | 0.538808303 |
| ENSDART00000168028 | si:dkey-225f23.5 | -0.209864951 | -0.328595081 | -0.21349874 | -0.05336952 |
| ENSDART00000168036 | rdh8b | -0.188576333 | -0.341040166 | -0.496499443 | -0.09323063 |
| ENSDART00000168038 | edil3a | -0.585018977 | -0.697545399 | -0.17495912 | 0.33520597 |
| ENSDART00000168083 | Metazoa_SRP | 1.497191597 | 1.645034783 | 1.435853831 | 0.615781715 |
| ENSDART00000168084 | scp2b | -0.093323025 | -0.125270421 | -0.400440129 | -0.264110493 |
| ENSDART00000168089 | cyp27a7 | -0.742372496 | -0.705345887 | -0.793489907 | -0.543448957 |
| ENSDART00000168107 | crb2b | -0.111814011 | -0.354478217 | -0.285757255 | 0.093257061 |
| ENSDART00000168121 | lzts3a | -0.348828928 | -0.543883063 | -0.164874094 | 0.093345065 |
| ENSDART00000168132 | si:ch1073-82l19.1 | -0.151537059 | -0.297036072 | -0.500517261 | -2.428975421 |
| ENSDART00000168155 | si:ch211-194m7.4 | 2.689408284 | 1.434584454 | 0.930290361 | -0.08573457 |
| ENSDART00000168157 | mkrn2os.2 | 0.50354684 | 0.675600001 | 1.137753613 | 0.994948824 |

| | | | | | |
|---|---|---|---|---|---|
| ENSDART00000168160 | pip5k1cb | -0.047296457 | -0.437985038 | -0.478356963 | -0.284131261 |
| ENSDART00000168162 | FQ323156.1 | -0.536894775 | -1.163960845 | -0.494433057 | -0.466334383 |
| ENSDART00000168163 | hnrnpabb | 0.035490772 | 0.022906035 | 0.318157336 | 0.096428682 |
| ENSDART00000168167 | rtcb | 0.060457492 | 0.250042841 | 0.449158496 | 0.303788278 |
| ENSDART00000168170 | ttc39c | -0.270510539 | -0.264194482 | -0.052910688 | -0.079707834 |
| ENSDART00000168188 | mettl1 | 0.565360225 | 0.37588106 | 0.377233069 | 0.251655436 |
| ENSDART00000168201 | golim4a | -0.134763119 | -0.005515007 | -0.245821925 | -1.172515705 |
| ENSDART00000168218 | CABZ01111953.1 | -0.582414072 | -0.415065608 | 0.00894593 | 0.020084563 |
| ENSDART00000168228 | tmem184a | -0.3188994 | -0.465602708 | -0.713259634 | -0.580015552 |
| ENSDART00000168241 | tubb2b | -0.194997441 | 0.285540167 | 0.888424822 | 0.706942534 |
| ENSDART00000168246 | spryd3 | -0.145299627 | -0.212114286 | -0.257834151 | -0.152966783 |
| ENSDART00000168270 | CABZ01048053.1 | -0.515099411 | -0.533680852 | 0.225344787 | 0.362581791 |
| ENSDART00000168276 | scarb1 | -3.646800721 | -2.308559886 | 0.370802474 | 0.243200954 |
| ENSDART00000168277 | boka | -0.236090037 | -0.162724249 | -0.56155479 | -0.509356169 |
| ENSDART00000168278 | si:ch211-276i12.4 | -0.631335043 | -0.270754342 | -0.641142033 | -0.623314164 |
| ENSDART00000168309 | trim25 | 1.908271479 | 1.980136553 | 2.767316449 | 1.926127968 |
| ENSDART00000168310 | acap3b | -0.431660294 | -0.622927148 | -0.474200982 | -0.250587028 |
| ENSDART00000168317 | apba1a | -0.329639415 | -0.090034986 | 0.20399901 | 0.390106315 |
| ENSDART00000168371 | mibp | 1.064831288 | 1.384792727 | 1.720455753 | 0.84273718 |
| ENSDART00000168376 | Metazoa_SRP | 0.77672172 | 1.175457852 | 1.304833035 | 1.047872554 |
| ENSDART00000168377 | thrap3b | 0.204741923 | -0.103859097 | 0.348419918 | 0.136670023 |
| ENSDART00000168396 | FO904869.1 | -0.490668054 | -0.607434677 | -0.194530972 | 0.12309886 |
| ENSDART00000168419 | rsu1 | 0.608707901 | 0.588635147 | 0.668370486 | 0.283061431 |
| ENSDART00000168453 | slc43a2a | -0.36850619 | -0.532753158 | 0.198651765 | 0.248433931 |
| ENSDART00000168472 | creb5a | 2.212193939 | 1.443174599 | 0.809493268 | 0.247644913 |
| ENSDART00000168483 | si:dkey-16p21.7 | 0.12768447 | 0.414452748 | 0.453712173 | 0.342486315 |
| ENSDART00000168497 | eef2b | 0.448731314 | 0.547292163 | 0.7539331 | 0.364287935 |
| ENSDART00000168518 | asic4b | -0.92148029 | -1.144921239 | -0.577321683 | -0.304835302 |
| ENSDART00000168531 | irf2bp2b | -0.128403316 | -0.188582728 | -0.404745176 | -0.138034767 |
| ENSDART00000168534 | NPFFR2 (1 of many) | -0.514269049 | -0.40572501 | -0.533037042 | -0.336902595 |
| ENSDART00000168542 | arf3a | 0.034708072 | 0.783187174 | 0.81852048 | 0.521503928 |
| ENSDART00000168556 | ostf1 | 1.22625882 | 1.436442197 | 0.76765034 | 0.887655813 |
| ENSDART00000168559 | ttf2 | -3.902752419 | 0.990904721 | 1.020977359 | 0.830250106 |
| ENSDART00000168565 | txnbb | 0.43423077 | 0.785550255 | 1.235069922 | 1.025671009 |
| ENSDART00000168568 | CU861477.1 | -0.399340079 | -0.694343635 | -0.183948068 | 0.054030168 |
| ENSDART00000168608 | rnaset2 | 1.085971834 | 0.685727013 | 0.397141355 | 0.382009137 |
| ENSDART00000168610 | si:ch211-283g2.2 | 1.147251625 | 1.274539863 | 1.168500693 | 0.162307024 |
| ENSDART00000168616 | ppa1a | 0.095923853 | -0.258115212 | -0.556184684 | -0.293684673 |
| ENSDART00000168627 | zfyve9a | -0.287315941 | -0.130733792 | -0.157388884 | -0.011640067 |
| ENSDART00000168631 | cacna1hb | -0.565978595 | -0.722362792 | -0.435741841 | -0.029665329 |
| ENSDART00000168633 | CABZ01085700.1 | -0.53401794 | -0.080184443 | 0.667052745 | 0.803861054 |
| ENSDART00000168639 | cry1ab | -0.18716056 | -0.280815639 | -0.387838008 | -0.275846695 |
| ENSDART00000168641 | CT030188.1 | 0.86349998 | 1.431982341 | 1.038679795 | -0.017052584 |
| ENSDART00000168653 | fam110b | -0.266298706 | -0.306955651 | -0.224541228 | -0.085175606 |
| ENSDART00000168683 | lrrc8c | 0.403272924 | 0.771214032 | 0.877257507 | 0.389191376 |
| ENSDART00000168698 | ostf1 | 0.435343641 | 0.254095599 | -0.055556308 | -0.273490651 |
| ENSDART00000168705 | si:ch73-103b11.2 | -0.165771907 | -0.231080784 | -0.260448742 | -0.241729977 |
| ENSDART00000168718 | chrm5a | -0.32077931 | -0.748869105 | -0.317792694 | -0.07500967 |
| ENSDART00000168727 | zgc:64065 | 0.439077495 | 0.173893761 | 0.133684815 | 0.136149811 |
| ENSDART00000168729 | fam196ab | -0.54578901 | -0.555866325 | -0.405243034 | 0.063979844 |
| ENSDART00000168749 | nlgn3a | -0.417078437 | -0.490224096 | -0.240516544 | -0.086053557 |

| | | | | | |
|---|---|---|---|---|---|
| ENSDART00000168750 | smarca1 | -0.162085202 | -0.116464254 | -0.382463649 | -0.380530416 |
| ENSDART00000168754 | cacnb2a | -0.312716955 | -0.298709564 | -0.211334034 | -0.110938146 |
| ENSDART00000168778 | stt3a | 0.3508706 | 0.104695298 | -0.028500788 | -0.125020463 |
| ENSDART00000168799 | ggps1 | 0.829174536 | 0.521929251 | 0.538342651 | 0.450861168 |
| ENSDART00000168805 | kcnk9 | -0.553605245 | -0.952161118 | -0.308244106 | 0.289642563 |
| ENSDART00000168821 | CDH22 | -0.242258311 | -0.624670054 | -0.303879761 | -0.070304381 |
| ENSDART00000168822 | pabpc4 | 1.120879979 | 1.597763229 | 2.131974577 | 1.975353129 |
| ENSDART00000168831 | si:ch73-158p21.3 | 1.161771941 | 0.892729958 | 0.502592433 | 0.147950065 |
| ENSDART00000168840 | rpl35a | 0.357031004 | 0.320334573 | 0.163051637 | -0.036511823 |
| ENSDART00000168849 | si:ch211-147m6.2 | 1.772352788 | 0.845008987 | -0.374275663 | -0.738910487 |
| ENSDART00000168850 | taok1b | -0.044686095 | -0.116318692 | -0.429811181 | -0.132610503 |
| ENSDART00000168851 | rab11fip2 | 0.212709727 | 0.193153756 | 0.330809666 | 0.288597218 |
| ENSDART00000168876 | si:dkey-81h8.1 | -0.18803818 | 0.217329319 | 0.889109393 | 1.046963425 |
| ENSDART00000168885 | nrg1 | -0.78621851 | -0.719452399 | 0.167065606 | 0.289901187 |
| ENSDART00000168898 | hspa8 | 1.373917849 | 0.861861784 | 0.886683158 | 0.865443603 |
| ENSDART00000168899 | pcdh1g33 | -0.406478127 | -0.012838877 | 0.30409891 | 0.306864798 |
| ENSDART00000168902 | Metazoa_SRP | 0.908254207 | 1.60749636 | 1.079432418 | 0.706761496 |
| ENSDART00000168910 | ablim3 | -0.457548186 | -0.818398336 | -0.469699072 | -0.283516024 |
| ENSDART00000168913 | CABZ01113883.1 | 0.283917325 | -0.023921963 | -0.385162746 | -1.507337386 |
| ENSDART00000168918 | pcdh1gc6 | 0.501599317 | 0.826475212 | 1.380988505 | 1.066751642 |
| ENSDART00000168920 | nbeab | -0.251742312 | -0.363594185 | -0.174304201 | -0.037829821 |
| ENSDART00000168922 | Metazoa_SRP | 1.221075219 | 2.596056056 | 2.04902232 | 1.755997125 |
| ENSDART00000168935 | mfsd8 | 0.060271678 | -0.029632781 | -0.242261851 | -0.178722202 |
| ENSDART00000168957 | cdh18a | -0.488858345 | -0.440968125 | -0.25726489 | -0.177287767 |
| ENSDART00000168969 | htr7c | -0.617759768 | -0.791263551 | -0.091828465 | 0.038777953 |
| ENSDART00000168996 | plrdgb | -0.579612528 | -0.647606525 | -0.269512976 | 0.036793619 |
| ENSDART00000169006 | ak2 | 0.70404039 | 0.489848323 | 0.563968118 | 0.137147044 |
| ENSDART00000169023 | scn1lab | 0.230826198 | 0.448869227 | 0.411715364 | 0.200031153 |
| ENSDART00000169026 | jak2a | 0.024039251 | 0.103369971 | 0.328189294 | 0.009690028 |
| ENSDART00000169040 | si:ch211-235e9.8 | -0.436326332 | -0.357257361 | -0.23322888 | -0.084450729 |
| ENSDART00000169052 | elovl1b | 0.945814486 | 1.07228669 | 0.819418014 | 0.365575121 |
| ENSDART00000169060 | si:ch211-212k18.5 | -0.245044911 | -0.301564521 | -0.568174767 | -0.780659219 |
| ENSDART00000169061 | CU138533.1 | 0.641184794 | 1.068185172 | 1.17226183 | 0.773255864 |
| ENSDART00000169081 | mtmr7b | -0.302136082 | -0.184242994 | 0.142322853 | 0.09943817 |
| ENSDART00000169091 | psd2 | -0.329495857 | -0.192167097 | 0.082881422 | 0.17989231 |
| ENSDART00000169105 | pcdh1g29 | -0.285141513 | 0.143203693 | 0.405619816 | 0.452362051 |
| ENSDART00000169108 | kcnj12a | -0.480725794 | -0.616905664 | -0.509410819 | -0.355199716 |
| ENSDART00000169116 | cpne4a | -0.476761925 | -0.590434794 | -0.280550227 | 0.444813194 |
| ENSDART00000169119 | ndrg4 | -0.4692852 | -0.086247828 | 0.285804524 | 0.242386508 |
| ENSDART00000169127 | adgrb3 | -0.338634159 | -0.316792657 | -0.154400064 | 0.089681207 |
| ENSDART00000169129 | ndrg4 | -0.326143464 | -0.61306806 | -0.078381969 | -0.216584442 |
| ENSDART00000169130 | ftr64 | 0.270335706 | 0.025822717 | 1.271602536 | 0.119170332 |
| ENSDART00000169136 | CABZ01007222.1 | -0.363042867 | -0.477045243 | -0.379599521 | -0.116458215 |
| ENSDART00000169165 | CABZ01021450.1 | -0.130594664 | -0.107324732 | -0.381541063 | -0.095206853 |
| ENSDART00000169187 | ptpro | 0.079888282 | 0.61716785 | 1.250280006 | 1.189522062 |
| ENSDART00000169201 | pcdh2g16 | 0.673146228 | 0.480832534 | 0.40403692 | 0.542228011 |
| ENSDART00000169202 | si:ch211-153b23.5 | 4.343063064 | 5.139014825 | 3.462179286 | 1.548865865 |
| ENSDART00000169209 | ptrh1 | 0.68284848 | 0.772674127 | 0.446966304 | 0.22472723 |
| ENSDART00000169210 | CABZ01072550.1 | -0.141044522 | -0.137807967 | -0.329360205 | -0.030428593 |
| ENSDART00000169228 | vat1l | 0.089392965 | -0.09841303 | -0.250056266 | -0.442720544 |
| ENSDART00000169243 | znf384l | -0.366831524 | -0.600767675 | -0.247311028 | 0.205515141 |

| | | | | | |
|---|---|---|---|---|---|
| ENSDART00000169248 | drd4-rs | -0.178692855 | -0.390296441 | -0.526046415 | -0.318223636 |
| ENSDART00000169275 | sdad1 | 0.31633119 | 0.22409068 | 0.190752906 | 0.014420983 |
| ENSDART00000169309 | si:dkey-203a12.2 | 1.554830383 | 2.508546052 | 1.961959042 | 0.546471184 |
| ENSDART00000169322 | zbtb21 | -0.157953464 | -0.169354872 | -0.299279576 | -0.291807058 |
| ENSDART00000169341 | nrp1a | 0.186448987 | 0.415165621 | 0.563193133 | 0.501195382 |
| ENSDART00000169343 | kdf1b | -0.449286765 | -0.274976378 | -0.544184682 | -0.396728296 |
| ENSDART00000169363 | CR678435.1 | -0.089321474 | -0.319658973 | -0.354063558 | -0.259523127 |
| ENSDART00000169365 | Metazoa_SRP | 2.055456458 | 2.688758632 | 2.943606625 | 2.098435714 |
| ENSDART00000169367 | CABZ01061884.1 | -0.298329532 | -0.377577791 | -0.54544765 | -0.245416552 |
| ENSDART00000169375 | znf1151 | 2.128272751 | 1.140055383 | 3.010647967 | 3.121008171 |
| ENSDART00000169376 | PLEKHG5 (1 of many) | -0.003560342 | -0.365335228 | -0.516938003 | -0.057649582 |
| ENSDART00000169380 | slc25a43 | -0.226077688 | 0.088930874 | -0.355915914 | -0.508574161 |
| ENSDART00000169407 | fynb | -0.281673654 | -0.267434526 | -0.200981958 | -0.171531273 |
| ENSDART00000169447 | sorbs1 | 0.299961188 | 0.420748275 | 0.143707838 | 0.107473415 |
| ENSDART00000169452 | CR848841.2 | 1.41519844 | 0.623487161 | 0.225042545 | -0.111771198 |
| ENSDART00000169462 | fam189a1 | -0.165720729 | -0.526215514 | -0.059981823 | -0.273837885 |
| ENSDART00000169477 | tmem176l.1 | 0.297546209 | 0.073954032 | -0.516480584 | -0.864341304 |
| ENSDART00000169484 | amph | -0.176945265 | -0.219628366 | -0.476094931 | -0.135317446 |
| ENSDART00000169511 | ppp2r2bb | -0.347621729 | -0.40572046 | -0.501873649 | -0.158073428 |
| ENSDART00000169514 | cacna1db | -0.154801093 | -0.201285475 | -0.375153121 | -0.07189092 |
| ENSDART00000169531 | traf2b | -0.006517414 | -0.077905282 | -0.353292738 | -0.278704886 |
| ENSDART00000169548 | rgs7b | -0.179117493 | -0.337814246 | -0.235695562 | -0.091038288 |
| ENSDART00000169552 | cdh13 | -0.778914955 | -0.463881372 | -0.06287293 | 0.145633136 |
| ENSDART00000169600 | abhd4 | 0.160082589 | 0.423680439 | -0.10396044 | -0.08495852 |
| ENSDART00000169609 | tefb | -0.156114581 | -0.555141888 | -0.301378906 | -0.24132076 |
| ENSDART00000169614 | nadsyn1 | 0.473419312 | 0.288129879 | -0.022685337 | -0.020046915 |
| ENSDART00000169626 | kcnc1a | -0.804732723 | -1.394467436 | -0.226498418 | 0.073837621 |
| ENSDART00000169633 | col4a3bpb | 0.489929413 | 0.127889926 | -0.192097119 | -0.259910666 |
| ENSDART00000169644 | CU659306.1 | 0.286956046 | 0.461515845 | 0.629089247 | 0.128650576 |
| ENSDART00000169648 | | 0.47490981 | 0.626028668 | 1.036581019 | 0.748938091 |
| ENSDART00000169663 | CT573860.1 | 0.353646224 | 0.562802664 | 0.454063437 | 0.329757181 |
| ENSDART00000169667 | ptpro | -0.163271096 | 0.493460364 | 0.820001756 | 0.696103761 |
| ENSDART00000169682 | hdac4 | -0.093013902 | -0.31519447 | 0.302479135 | 0.371645007 |
| ENSDART00000169687 | dlg1 | -0.200796553 | -0.39742906 | -0.281934456 | -0.007509175 |
| ENSDART00000169698 | zgc:158343 | -0.309656325 | -0.821697026 | -0.621795526 | -0.564704541 |
| ENSDART00000169710 | utp23 | 0.386723475 | 0.25777974 | 0.117505162 | 0.115827548 |
| ENSDART00000169717 | ca5a | 0.91389398 | 0.899982871 | 0.348639788 | 0.672790664 |
| ENSDART00000169723 | HERC5 | -0.128535223 | -0.136551326 | 0.521223162 | -0.158454388 |
| ENSDART00000169733 | klhl23 | -0.117061422 | -0.244154323 | -0.447638501 | -0.20478162 |
| ENSDART00000169746 | ampd2a | -0.446564291 | -0.523779729 | -0.426555345 | -0.117905151 |
| ENSDART00000169764 | syt1a | -0.535633412 | -0.482595993 | -0.403145231 | -0.029612735 |
| ENSDART00000169768 | slc8a3 | -0.133596734 | -0.60366341 | -0.159093114 | 0.187240856 |
| ENSDART00000169789 | grap2b | 0.941263606 | 0.635408975 | 0.416956975 | 0.146969065 |
| ENSDART00000169810 | stub1 | -0.352758303 | -0.24873956 | -0.212157701 | -0.100812458 |
| ENSDART00000169823 | pnpla3 | 0.41574021 | 0.387767111 | 0.775977498 | 0.378524498 |
| ENSDART00000169828 | rlbp1a | -0.278789297 | -0.290042852 | -0.461967142 | -0.22894081 |
| ENSDART00000169834 | CABZ01052431.1 | 0.752252615 | 0.635370532 | 0.523133762 | 0.005708707 |
| ENSDART00000169845 | pargl | 0.069840572 | -0.083042796 | 0.787500652 | -0.250417109 |
| ENSDART00000169852 | RPL37A | 0.3117075 | 0.379509479 | 0.225285723 | 0.045531307 |
| ENSDART00000169853 | slmapa | -0.949896363 | -1.088521821 | -0.936929707 | -0.657167359 |
| ENSDART00000169887 | wu:fu71h07 | 0.98545678 | 0.406691046 | 0.849706316 | 0.386892251 |

| | | | | | |
|---|---|---|---|---|---|
| ENSDART00000169892 | srp54 | 2.340238808 | 3.996874038 | 2.498816008 | 1.178878233 |
| ENSDART00000169895 | zbtb18 | 0.009752394 | -0.192437538 | -0.328439949 | -0.081774144 |
| ENSDART00000169915 | nop58 | 0.425406041 | 0.33755518 | 0.254231429 | 0.051624158 |
| ENSDART00000169916 | dmxl2 | -0.252318602 | -0.45068029 | 0.053959432 | 0.232942265 |
| ENSDART00000169923 | ndel1b | -0.007054921 | -0.080749821 | -0.505244725 | -0.275115017 |
| ENSDART00000169932 | ccdc142 | -0.157730988 | -0.223081294 | -0.3402281 | -0.055715693 |
| ENSDART00000169936 | sox8a | -0.26561801 | -0.150998063 | -0.108428913 | -0.111175355 |
| ENSDART00000169940 | tmem176l.1 | 0.227275106 | 0.228370918 | -0.147494824 | -0.481831855 |
| ENSDART00000169942 | stx3a | -0.199600759 | -0.344637551 | -0.343244623 | -0.200314073 |
| ENSDART00000169943 | CABZ01084447.1 | -0.382296742 | -0.397697491 | -0.572266726 | -0.473542202 |
| ENSDART00000169945 | si:ch73-236c18.7 | 1.628999904 | 1.036261941 | 0.5890347 | 0.798137712 |
| ENSDART00000169946 | | -0.124122046 | -0.322722589 | -0.278305257 | 0.008390957 |
| ENSDART00000169948 | GRM7 | -0.415342332 | -0.376852993 | -0.226859206 | -0.023200649 |
| ENSDART00000169951 | Metazoa_SRP | 1.195279937 | 1.735123002 | 1.057793715 | 0.943058282 |
| ENSDART00000169953 | dctn1b | -0.327369492 | -0.021477792 | 0.065821587 | 0.193833161 |
| ENSDART00000169963 | ryr1b | -0.099451213 | -0.21486392 | 0.409324638 | 0.697954707 |
| ENSDART00000169972 | ndst1b | -0.191299747 | -0.313711004 | -0.152975304 | -0.236354037 |
| ENSDART00000169976 | CABZ01034529.1 | 0.902549952 | 0.806876908 | 0.306228824 | 0.021333022 |
| ENSDART00000169989 | BX649355.1 | 0.986946969 | 0.719472766 | 0.338456492 | 0.762216016 |
| ENSDART00000169993 | stx12 | 0.289245416 | 0.334302649 | 0.162033659 | 0.014864416 |
| ENSDART00000169999 | | 0.330954109 | -0.605210778 | -0.313930686 | -0.099685871 |
| ENSDART00000170012 | mapre3b | -0.205130823 | -0.301066074 | -0.42167523 | -0.136278286 |
| ENSDART00000170021 | | 2.405989626 | 2.77310135 | 2.003596445 | 2.210644738 |
| ENSDART00000170060 | mdh1b | 3.761355509 | 3.246314783 | 3.336555843 | 1.787365183 |
| ENSDART00000170089 | epha4b | -0.586426265 | -0.698463168 | -0.228203192 | -0.016930318 |
| ENSDART00000170095 | CR388166.2 | 0.588171845 | 0.299671021 | 0.452688856 | 0.506464888 |
| ENSDART00000170107 | ppp2r2ba | 0.343342582 | 0.340534473 | 0.3871703 | 0.276562577 |
| ENSDART00000170116 | uap1 | 0.399345456 | 0.560025587 | 0.455223073 | 0.418274301 |
| ENSDART00000170133 | eci1 | 2.097463401 | 3.784028698 | 2.248505252 | 3.097351651 |
| ENSDART00000170165 | cep97 | -4.277597338 | -0.154895502 | -0.337498335 | -0.461056374 |
| ENSDART00000170184 | fgd1 | 0.020796397 | 0.102196495 | 0.582360556 | 0.487921156 |
| ENSDART00000170191 | FQ311879.1 | -0.155204603 | -0.24875676 | -0.288299334 | -0.086604212 |
| ENSDART00000170204 | si:zfos-364h11.2 | 0.504424795 | 0.456776646 | 0.313589859 | -0.247114093 |
| ENSDART00000170208 | limch1b | -0.042803542 | -0.148322889 | -0.621537079 | -0.273084471 |
| ENSDART00000170222 | mibp | 1.202449519 | 1.249757535 | 1.712700336 | 0.61416841 |
| ENSDART00000170228 | sept9b | -0.22300654 | -0.343163525 | -0.367834845 | -0.241839556 |
| ENSDART00000170284 | creb3l3l | 0.444489468 | 0.957040221 | 0.49606022 | 0.414651857 |
| ENSDART00000170285 | def8 | -0.050118345 | -0.13948188 | -0.345534099 | -0.220243255 |
| ENSDART00000170304 | CABZ01039096.1 | 1.863106121 | 1.329668381 | 1.247074235 | 0.639142974 |
| ENSDART00000170324 | tom1 | -0.260627206 | -0.142873643 | -0.174666961 | -0.048298907 |
| ENSDART00000170325 | map7d1b | -0.159995641 | 0.033353383 | 0.380064991 | 0.315967697 |
| ENSDART00000170326 | gadd45ba | 1.741159479 | 1.754335562 | 1.06756424 | 0.870946071 |
| ENSDART00000170341 | apba1a | -0.229826313 | -0.069954525 | 0.548704484 | 0.456234995 |
| ENSDART00000170351 | nbeal1 | -0.047098407 | -0.274555108 | -0.182658383 | -0.033615077 |
| ENSDART00000170362 | CABZ01088490.1 | -0.602203241 | -0.75748097 | -0.591873288 | -0.034015136 |
| ENSDART00000170374 | si:dkey-38l22.2 | 1.50589782 | 1.851640662 | 1.322719067 | 0.692201779 |
| ENSDART00000170376 | mpp7a | -0.022565767 | -0.302686812 | -0.467837523 | -0.093897739 |
| ENSDART00000170381 | CR855389.1 | -0.538290002 | -0.347216121 | -0.566215377 | -0.407287476 |
| ENSDART00000170383 | CABZ01089777.1 | -0.22190804 | -0.652997161 | -0.608886761 | -0.372475629 |
| ENSDART00000170385 | tgm1l1 | 0.820949739 | 0.875506718 | 1.293638142 | 0.305667169 |
| ENSDART00000170396 | apbb2b | -0.997027217 | -0.745617105 | 0.597402109 | 1.180709512 |

| | | | | | |
|---|---|---|---|---|---|
| ENSDART00000170405 | CARTPT (1 of many) | 1.790273718 | 2.206490975 | 2.680185278 | 2.041251968 |
| ENSDART00000170422 | si:dkey-19b23.8 | 0.497209105 | 1.26444289 | 1.061660184 | 0.502612026 |
| ENSDART00000170423 | jakmip3 | -0.243674673 | -0.387799515 | -0.202768181 | 0.090619484 |
| ENSDART00000170433 | tert | 0.747709966 | 0.573674682 | 0.803557653 | 0.065172489 |
| ENSDART00000170441 | CLDN23 | -0.099162778 | -0.365194678 | -0.492897369 | -0.25219875 |
| ENSDART00000170453 | slc20a1b | -0.460714141 | -0.247481668 | -0.017046699 | -0.023964599 |
| ENSDART00000170456 | shroom3 | 0.242207867 | 0.41278364 | 0.171514229 | -0.137113926 |
| ENSDART00000170460 | nrxn3b | -0.2276052 | -0.615850363 | -0.134586161 | -0.003465474 |
| ENSDART00000170466 | gch2 | 0.005943316 | 1.153050374 | 0.394171756 | -0.303921757 |
| ENSDART00000170470 | ghrhra | -0.141377937 | -0.217703208 | -0.411536652 | -0.240637833 |
| ENSDART00000170510 | RIMBP2 (1 of many) | -0.653406892 | -0.568163429 | -0.174788334 | -0.007324529 |
| ENSDART00000170512 | ctnnd1 | 0.874551431 | 1.060254696 | 0.863956713 | 0.322404291 |
| ENSDART00000170518 | anxa1c | -0.336866673 | -0.270774527 | -0.035149981 | -0.070098806 |
| ENSDART00000170526 | CABZ01021220.1 | -0.34731151 | -0.318997758 | -0.303107193 | -0.013605358 |
| ENSDART00000170546 | wdr17 | -0.20142732 | -0.213426713 | -0.433783757 | -0.087288031 |
| ENSDART00000170562 | ece2b | -0.279383738 | -0.308974349 | -0.125983301 | -0.132241867 |
| ENSDART00000170569 | syt12 | -0.747896951 | -0.906145295 | -0.362441574 | -0.047384497 |
| ENSDART00000170571 | dmtn | 0.084482265 | 0.234365609 | 0.453911363 | 0.30033427 |
| ENSDART00000170572 | lepa | 4.559547685 | 4.778234325 | 4.366752593 | 2.765663683 |
| ENSDART00000170575 | nfat5b | -0.001493923 | -0.392122738 | -0.566909919 | -0.288747412 |
| ENSDART00000170583 | march8 | -0.037993536 | -0.377054474 | -0.284528801 | -0.103767971 |
| ENSDART00000170589 | mibp | 1.067201928 | 1.476075399 | 1.501871782 | 0.758874349 |
| ENSDART00000170601 | si:ch1073-228h2.2 | 1.512427811 | 1.187632525 | 1.253650535 | 1.028624188 |
| ENSDART00000170620 | ctxn1 | -1.120908397 | -0.319127099 | 0.144625495 | 0.239695278 |
| ENSDART00000170630 | BX511034.7 | 0.940574459 | 1.49905624 | 1.717428689 | 0.604931976 |
| ENSDART00000170631 | ebf1a | -0.245696473 | -0.326200552 | -0.132641619 | 0.115308885 |
| ENSDART00000170632 | gngt2b | -0.25352879 | -0.46015013 | -0.596390898 | -0.26583618 |
| ENSDART00000170671 | megf8 | 0.369502187 | 0.913401505 | 1.13324158 | 0.463058503 |
| ENSDART00000170675 | col11a1a | -3.564960509 | -0.565137585 | -0.451612456 | -1.057274574 |
| ENSDART00000170680 | ptprdb | -0.822263801 | -1.170454394 | 0.28663796 | 0.697055522 |
| ENSDART00000170684 | btf3 | 0.237794169 | 0.784907824 | 0.656975875 | 0.357635511 |
| ENSDART00000170695 | lrit1b | -0.349980517 | -0.215317539 | -0.688918418 | -0.304938883 |
| ENSDART00000170700 | nacc1b | -0.295710389 | -0.416714615 | 0.013950982 | 0.110349783 |
| ENSDART00000170709 | st8sia1 | -0.358435626 | -0.336878724 | -0.21402261 | -0.093884674 |
| ENSDART00000170752 | tox2 | -0.491067711 | -0.314026775 | -0.014696402 | 0.264527682 |
| ENSDART00000170755 | slc30a8 | 0.607491231 | 0.345740737 | -0.026161917 | -0.397029232 |
| ENSDART00000170757 | kntc1 | 0.719300782 | 0.704073311 | 0.501967558 | 0.060438055 |
| ENSDART00000170758 | tcirg1b | 0.779415513 | 0.215400071 | 0.036968665 | -0.339490039 |
| ENSDART00000170762 | slc44a5b | -0.083921908 | -0.392138039 | -0.663427101 | -0.045247688 |
| ENSDART00000170768 | lect1 | 3.805179034 | 4.274135568 | 2.696697623 | 2.532825072 |
| ENSDART00000170790 | thrb | -0.205328436 | -0.495702017 | -0.443190893 | -0.211013858 |
| ENSDART00000170810 | si:dkey-10p5.7 | 1.115159817 | 1.313044185 | 0.773496009 | 0.334257673 |
| ENSDART00000170827 | ccpg1 | 0.08168444 | -0.121915293 | -0.302033768 | -0.268515452 |
| ENSDART00000170834 | znf1179 | 0.46327857 | 0.626036671 | 0.707498793 | 0.811192005 |
| ENSDART00000170839 | pcdh1g26 | -0.648364026 | -0.34360777 | 0.683964001 | 0.607743239 |
| ENSDART00000170854 | gphnb | -0.471165414 | -0.533169869 | -0.144401896 | 0.220388206 |
| ENSDART00000170856 | smu1b | 0.353209488 | -0.339997639 | -0.798035862 | -0.487523616 |
| ENSDART00000170865 | nme2b.1 | 0.635993303 | 0.679448572 | 0.689008761 | 0.355584206 |
| ENSDART00000170872 | tpm2 | -0.143399476 | 2.621381225 | 3.304047609 | 1.793877447 |
| ENSDART00000170874 | phldb1b | 0.7459988 | 0.426405219 | 0.977480972 | 0.780660286 |
| ENSDART00000170875 | BX663610.1 | -0.528067918 | -0.63401091 | -0.217086155 | -0.025338796 |

| | | | | | |
|---|---|---|---|---|---|
| ENSDART00000170886 | dennd1b | -0.037953425 | -0.315629797 | -0.364933587 | -0.202214668 |
| ENSDART00000170888 | pkma | -0.382873513 | -0.359191464 | -0.202639045 | 0.04775771 |
| ENSDART00000170932 | rims2a | -0.482409185 | -0.599060629 | -0.273352321 | -0.056203652 |
| ENSDART00000170951 | PXDN | 0.459355707 | 0.64772414 | 0.996368549 | 0.498632459 |
| ENSDART00000170952 | pvrl2l | -0.402141005 | -0.731362921 | -0.043297784 | 0.281757642 |
| ENSDART00000170955 | fn3krp | -0.183544623 | -0.156282547 | -0.395153052 | -0.231090781 |
| ENSDART00000170983 | lmnb2 | 0.185291183 | 0.299339057 | 0.275121803 | 0.118770566 |
| ENSDART00000170993 | afmid | -0.391958822 | -0.311185654 | -0.48659874 | -0.071120804 |
| ENSDART00000170998 | tnrc6c2 | -0.257719977 | -0.319066529 | -0.317919253 | -0.242876098 |
| ENSDART00000171003 | CU929447.2 | 0.254824338 | 0.430023304 | 0.328306359 | 0.219409854 |
| ENSDART00000171006 | hpcal4 | -0.435402713 | -0.198074705 | -0.466665783 | -0.338540363 |
| ENSDART00000171013 | tenm4 | -0.557784731 | -0.616815207 | -0.086642376 | -0.101260046 |
| ENSDART00000171014 | ptprfa | -0.231763582 | -0.479834972 | -0.30404021 | -0.129420632 |
| ENSDART00000171021 | rab3ip | 0.020022814 | 0.162794734 | 0.729696081 | 0.694579303 |
| ENSDART00000171041 | tceb3 | -0.063048764 | -0.305499385 | -0.056078786 | -0.065276347 |
| ENSDART00000171058 | pald1a | 0.52269022 | 0.19158808 | 0.108215435 | -0.186659053 |
| ENSDART00000171072 | SEC14L1 | -0.099969512 | -0.17692833 | -0.270780357 | -0.178232632 |
| ENSDART00000171073 | tox2 | -0.713271279 | -0.691854818 | -0.08505138 | -0.233730891 |
| ENSDART00000171090 | sowahd | 0.56962557 | 0.415649109 | 0.184768671 | -0.065367372 |
| ENSDART00000171091 | zeb2b | -0.344020449 | -0.404143226 | -0.222307379 | 0.140992606 |
| ENSDART00000171113 | mibp | 1.815496184 | 1.851942075 | 2.237510566 | 0.882565569 |
| ENSDART00000171114 | si:dkey-18j18.3 | -5.054012109 | -5.054146995 | -1.846245188 | 0.223162792 |
| ENSDART00000171137 | pdia3 | 0.328843285 | 0.182764859 | 0.141869644 | -0.049588298 |
| ENSDART00000171169 | numbl | -0.199436908 | -0.274498759 | 0.034564123 | 0.069477302 |
| ENSDART00000171178 | nlrc3l | 0.061705987 | 0.093907561 | 0.374016309 | -0.032278843 |
| ENSDART00000171179 | si:ch211-241b2.5 | 1.438237825 | 1.596241839 | 1.013176313 | 0.055616895 |
| ENSDART00000171182 | FO704914.1 | -0.372200999 | -0.770608409 | -0.45129487 | -0.014659406 |
| ENSDART00000171188 | si:ch211-165d12.4 | 3.380342846 | 2.16159833 | 1.938428841 | 0.89654506 |
| ENSDART00000171202 | clrn1 | -0.342498627 | -0.174544267 | -0.411984605 | -0.164186516 |
| ENSDART00000171215 | slc6a3 | -0.667608441 | -0.754783262 | -0.446986408 | -0.343120625 |
| ENSDART00000171235 | calua | 0.528206357 | 0.591046444 | 0.528298198 | 0.210887316 |
| ENSDART00000171237 | kcnj2b | -0.406552715 | -0.524353506 | -0.661059283 | -0.393570981 |
| ENSDART00000171244 | ssh2b | -0.244446722 | -0.139215381 | 0.546963377 | 0.2356201 |
| ENSDART00000171252 | man2a1 | 0.336633303 | 0.148045387 | -0.143242682 | -0.284272741 |
| ENSDART00000171270 | ckap2l | 0.621565842 | 0.655006082 | 0.304610509 | 0.072068559 |
| ENSDART00000171288 | magi2b | -0.2674353 | -0.525609031 | -0.078206648 | -0.00532511 |
| ENSDART00000171289 | si:ch73-158p21.2 | 0.432663487 | 0.278579945 | 0.160816438 | 0.272943666 |
| ENSDART00000171306 | stxbp1a | -0.614795684 | -0.539858263 | -0.126841305 | 0.057169963 |
| ENSDART00000171316 | si:dkey-262j3.7 | -0.307347885 | -0.015370213 | 0.260259549 | 0.204989519 |
| ENSDART00000171320 | dcp2 | -0.072540161 | -0.060086595 | -0.320231159 | -0.153613562 |
| ENSDART00000171335 | map4k4 | 0.106316233 | 0.336083411 | 0.398966135 | 0.337674059 |
| ENSDART00000171336 | dyrk4 | 0.12887622 | -0.629074584 | -0.893898922 | 0.00387951 |
| ENSDART00000171343 | FP102192.1 | 1.130120264 | 0.697435719 | 1.760333984 | 0.139610385 |
| ENSDART00000171345 | | 2.015201895 | 1.694676698 | 1.017867776 | 0.711389392 |
| ENSDART00000171354 | nt5c2l1 | 0.533832785 | 0.693578932 | 1.620342483 | 0.507169588 |
| ENSDART00000171359 | asic1b | -0.583045866 | -0.669491195 | -0.186449925 | 0.212036898 |
| ENSDART00000171380 | top1mt | 0.983277478 | 1.316957074 | 1.074308725 | 0.32139594 |
| ENSDART00000171382 | epb41l3a | -0.440465797 | 0.150336785 | 0.618974575 | 0.751984906 |
| ENSDART00000171393 | efr3a | 0.028329226 | -0.20834437 | -0.54511994 | -0.110508442 |
| ENSDART00000171396 | pts | 0.27960755 | 0.015420592 | -0.484687881 | -0.407740441 |
| ENSDART00000171417 | si:ch73-299h12.3 | -0.719381589 | -0.760232748 | -0.826233659 | -0.863901262 |

| | | | | | |
|---|---|---|---|---|---|
| ENSDART00000171426 | pdzph1 | -0.416069913 | -0.411889039 | -0.457121209 | -0.221062124 |
| ENSDART00000171430 | BX908401.1 | 0.558597997 | 0.282396796 | 0.308082083 | 0.228227036 |
| ENSDART00000171433 | tnni1d | 0.40952122 | 3.032354782 | 4.009528497 | 2.823338021 |
| ENSDART00000171456 | rasgrf2b | -0.231105558 | -0.246992498 | -0.342919918 | -0.045608772 |
| ENSDART00000171466 | | -1.418832616 | -2.063774189 | -0.38972739 | -0.735471862 |
| ENSDART00000171490 | PCDH8 | -0.518696681 | -0.628285802 | -0.168709969 | 0.408565003 |
| ENSDART00000171494 | ssbp2 | -0.298958691 | -0.315587827 | -0.028952479 | 0.152638316 |
| ENSDART00000171496 | CDK18 | -0.371495713 | -0.374286185 | -0.433391037 | -0.282164654 |
| ENSDART00000171497 | mfap4 | 2.949855284 | 1.228743784 | 1.366529806 | 0.589539203 |
| ENSDART00000171506 | | -0.792662919 | -0.4993043 | 0.08528346 | 0.180572156 |
| ENSDART00000171523 | CABZ01089030.1 | -0.303576632 | -0.51118876 | -0.528168137 | -0.25789403 |
| ENSDART00000171524 | CABZ01062422.1 | 0.017851166 | 0.325269151 | 0.385529332 | 0.004388505 |
| ENSDART00000171571 | FAM184A (1 of many) | -0.84950254 | -1.068290057 | 0.155587747 | 0.361779982 |
| ENSDART00000171589 | hrasa | 0.543862541 | 0.913653251 | 0.982614467 | 0.767663706 |
| ENSDART00000171594 | mef2aa | -0.649844171 | -0.685014218 | -0.301258241 | -0.003741614 |
| ENSDART00000171624 | Metazoa_SRP | 1.040618727 | 2.218519951 | 2.250692356 | 1.867352594 |
| ENSDART00000171634 | galnt13 | 0.214607397 | -0.183374464 | -0.358548812 | -0.056557988 |
| ENSDART00000171639 | prkacbb | -0.216859972 | -0.333993863 | -0.380485937 | -0.232933535 |
| ENSDART00000171674 | abhd8a | -0.482604068 | -0.462135418 | 0.077091696 | 0.23737745 |
| ENSDART00000171678 | ubb | 0.267730806 | 0.139565922 | 0.302768617 | 0.039108167 |
| ENSDART00000171683 | GABRG1 | -0.455034367 | -0.392373746 | -0.437520168 | -0.270886274 |
| ENSDART00000171686 | si:ch211-225h24.2 | 0.00451198 | -0.128666512 | -0.443683785 | -0.077894977 |
| ENSDART00000171687 | XPO5 | 0.143122457 | 0.173342903 | 0.297288837 | 0.170261463 |
| ENSDART00000171691 | psmc3 | 0.15503219 | 0.267774928 | 0.185121834 | -0.000325126 |
| ENSDART00000171696 | sergef | -0.83692277 | -0.920324583 | -0.711315475 | -0.040508735 |
| ENSDART00000171701 | osmr | 1.90654589 | 2.08952232 | 1.976929325 | 1.246005697 |
| ENSDART00000171704 | soul4 | -0.335441282 | -0.335430337 | -0.104813252 | -0.062286587 |
| ENSDART00000171711 | gpsm1a | -0.376386863 | -0.294884117 | -0.277096957 | -0.079488929 |
| ENSDART00000171728 | FQ323156.1 | -0.3925045 | -0.548422409 | -0.578285827 | -0.21374206 |
| ENSDART00000171743 | sypa | -0.458751426 | -0.455626528 | -0.334063629 | -0.142465032 |
| ENSDART00000171749 | TMEM150A (1 of many) | -0.745537318 | -0.560172793 | -0.666392254 | -0.39318407 |
| ENSDART00000171762 | arhgef11 | -0.293734421 | -0.490220654 | -0.20280274 | 0.049910063 |
| ENSDART00000171777 | syt7b | -0.587310767 | -0.899783938 | -0.612333236 | -0.170082745 |
| ENSDART00000171789 | | 1.544321358 | 1.350890859 | 1.388203792 | 0.786947546 |
| ENSDART00000171806 | ppt1 | 0.388846083 | 0.447401215 | 0.166979487 | -0.03135359 |
| ENSDART00000171811 | grk1b | 2.834746901 | 2.005283407 | 2.325249515 | 3.603901667 |
| ENSDART00000171815 | abcc8b | 0.00284442 | -0.044162256 | -0.758748776 | -0.514944979 |
| ENSDART00000171823 | cdc14ab | -0.17216976 | -0.3147465 | -0.385013143 | -0.20236803 |
| ENSDART00000171824 | si:ch211-227e10.2 | -0.663768919 | -0.251157559 | -0.496668883 | -0.404241953 |
| ENSDART00000171830 | uchl1 | 0.751081517 | 1.035376191 | 0.855000132 | 0.800147781 |
| ENSDART00000171854 | si:ch1073-303d10.1 | -0.079667744 | -0.472299802 | -0.615705243 | -0.228401157 |
| ENSDART00000171867 | fnbp1l | 0.402488238 | 0.925028089 | 0.937640279 | 0.578874881 |
| ENSDART00000171868 | sgut1 | -0.438334002 | -0.252211506 | -0.468859348 | -0.232780566 |
| ENSDART00000171871 | cbfb | 0.41950513 | 0.366762581 | 0.672082991 | 0.461647583 |
| ENSDART00000171882 | CABZ01088567.1 | 0.407931 | 0.822027708 | 0.621405649 | 0.24977578 |
| ENSDART00000171891 | iqsec2a | -0.225046822 | -0.287658971 | -0.015287221 | 0.082198509 |
| ENSDART00000171900 | oit3 | -0.401327729 | -0.441180087 | -0.161045426 | -0.008695935 |
| ENSDART00000171913 | AL773558.1 | -0.541466018 | -0.593947031 | -0.853404048 | -0.564731615 |
| ENSDART00000171915 | FO834898.1 | -0.059070074 | -0.319253855 | -0.196715718 | -0.032975583 |
| ENSDART00000171916 | myom2a | 1.227196118 | 3.513600791 | 3.806244226 | 2.767902474 |
| ENSDART00000171935 | brpf3a | -0.333420883 | -0.26685281 | -0.204490748 | -0.091216638 |

| | | | | | |
|---|---|---|---|---|---|
| ENSDART00000171941 | si:dkeyp-53d3.5 | 2.939427931 | 3.522632269 | 3.599150168 | 3.695003344 |
| ENSDART00000171948 | zgc:172282 | -0.447105187 | -0.64749442 | -0.36124568 | -0.081452365 |
| ENSDART00000171951 | slit2 | -0.546372344 | -0.56571293 | -0.270916548 | -0.035288929 |
| ENSDART00000171965 | galnt18a | -0.106299358 | -0.103906292 | -0.35180024 | -0.003683356 |
| ENSDART00000171975 | si:ch211-230g14.6 | -0.465912508 | -0.525802535 | -0.259029045 | -0.276627158 |
| ENSDART00000171977 | ddx52 | 0.350026309 | 0.214664704 | 0.228079892 | -0.041434316 |
| ENSDART00000171996 | myl12.1 | 0.233861142 | 0.344330245 | 0.260508491 | 0.06338781 |
| ENSDART00000172014 | actr3 | 3.712429453 | 3.456018148 | 3.177836674 | 3.174953878 |
| ENSDART00000172016 | prox1a | -0.34053446 | -0.257376891 | -0.299242567 | -0.073418723 |
| ENSDART00000172019 | CU694219.1 | -0.358471879 | -0.088786066 | -0.423165049 | -1.570028426 |
| ENSDART00000172022 | dpysl4 | 0.270457415 | 0.912051797 | 1.185681882 | 0.874375314 |
| ENSDART00000172045 | zgc:73340 | -0.432558368 | -0.571519732 | -0.606453829 | -0.389736033 |
| ENSDART00000172069 | snrpg | 0.452166641 | 0.481061823 | 0.338564477 | 0.058821342 |
| ENSDART00000172076 | hook1 | -0.040243255 | -0.111929437 | -0.520350092 | -0.090216969 |
| ENSDART00000172080 | ttbk2a | 1.110423963 | 2.886130551 | 3.128605088 | 3.572451241 |
| ENSDART00000172084 | si:ch73-236j9.2 | 0.095305984 | 0.48958329 | 0.505184662 | 0.37060971 |
| ENSDART00000172095 | | -1.418832616 | -2.063774189 | -0.38972739 | -0.735471862 |
| ENSDART00000172102 | elof1 | 0.045486466 | -0.036149641 | -0.262027739 | -0.168895436 |
| ENSDART00000172114 | calm3a | -0.154349916 | -0.209222212 | -0.409530683 | -0.13969668 |
| ENSDART00000172128 | sez6l | -0.457515145 | -0.649880208 | -0.297003963 | -0.214224661 |
| ENSDART00000172135 | sh3pxd2b | 0.68385955 | 0.832942161 | 0.517904326 | 0.090715158 |
| ENSDART00000172149 | sh3rf2 | -0.359308533 | -0.521296357 | -0.501584275 | -0.200324498 |
| ENSDART00000172166 | NDUFC1 | -0.092745693 | -0.325133407 | -0.417118078 | -0.256060161 |
| ENSDART00000172190 | ajap1 | -0.155573908 | -0.463634934 | -0.533953643 | -0.131876486 |
| ENSDART00000172201 | trpv1 | 0.412354182 | 0.082280105 | 0.061779394 | -0.254715035 |
| ENSDART00000172207 | si:ch211-9d9.1 | 1.222547885 | 0.297262625 | 0.271454205 | -2.80261928 |
| ENSDART00000172215 | si:ch211-39i2.2 | 1.017534105 | 1.848603057 | 1.465869245 | 0.571196451 |
| ENSDART00000172218 | nsmfb | -0.786748472 | -1.480880504 | -0.428989171 | -0.523180434 |
| ENSDART00000172232 | sv2a | -0.325501969 | -0.478402861 | -0.246153159 | 0.031695064 |
| ENSDART00000172233 | si:ch73-55i23.1 | 0.636981625 | 0.696972269 | 1.179674288 | 0.207432569 |
| ENSDART00000172241 | nsdhl | 1.591273733 | 3.017393364 | 3.100136405 | 2.442907405 |
| ENSDART00000172251 | creb3l1 | -0.33540662 | 0.193573394 | -4.078076569 | -0.842213331 |
| ENSDART00000172267 | KCNV1 | -0.730291266 | -0.683938113 | -0.423486953 | -0.317970814 |
| ENSDART00000172274 | kifap3a | 2.52903808 | 3.896963711 | 3.83143073 | 1.193936031 |
| ENSDART00000172279 | cavin4a | -0.061511418 | 1.266222675 | 1.14872392 | 0.602916509 |
| ENSDART00000172285 | arf3b | -0.358958449 | -0.189620663 | 0.143810604 | 0.126356011 |
| ENSDART00000172294 | ctps1b | -0.150552652 | -0.426760874 | -0.523567714 | -0.150809326 |
| ENSDART00000172300 | slc38a4 | -0.096535094 | -0.136897206 | -0.74017911 | -0.38118016 |
| ENSDART00000172301 | tk1 | 1.414413476 | 0.945772512 | 1.08664013 | 0.593080412 |
| ENSDART00000172305 | lsm10 | 0.431922906 | 0.307479396 | 0.650924826 | 0.415051794 |
| ENSDART00000172307 | UBE2M | -0.026970956 | 0.061666236 | 0.299772523 | 0.276955817 |
| ENSDART00000172309 | twsg1a | 0.181135796 | 0.092573769 | -0.368073769 | -0.475343856 |
| ENSDART00000172310 | zbtb4 | -0.365422353 | -0.277101341 | -0.237436935 | -0.247791483 |
| ENSDART00000172327 | taok1b | -0.323110832 | -0.519144421 | -0.19684291 | 0.126510211 |
| ENSDART00000172329 | CABZ01084963.1 | -0.076477014 | -0.479017439 | -0.153366972 | -0.215592031 |
| ENSDART00000172335 | cpne3 | 0.169360614 | 0.301780948 | 0.100312932 | -0.018165212 |
| ENSDART00000172336 | cabp2a | -0.40143904 | -0.491969769 | -0.23184904 | -0.50001549 |
| ENSDART00000172337 | rhot1a | -3.878236672 | -5.686158747 | -0.665991717 | -0.822715826 |
| ENSDART00000172338 | cav1 | 0.970717786 | 1.027469024 | 1.102211393 | 0.680419868 |
| ENSDART00000172357 | tmem132a | -0.246648072 | -0.102389273 | -0.224742481 | -0.519178406 |
| ENSDART00000172367 | sgip1b | -0.158702965 | -0.245738955 | -0.177100693 | 0.051555171 |

| | | | | | |
|---|---|---|---|---|---|
| ENSDART00000172372 | Metazoa_SRP | 1.431390172 | 1.797550169 | 1.401344471 | 0.830250287 |
| ENSDART00000172373 | ompa | -0.333379791 | -0.33389964 | -0.383107757 | -0.229559723 |
| ENSDART00000172408 | arhgap11a | 0.695056063 | 0.093403628 | 0.066933123 | 0.263951638 |
| ENSDART00000172409 | il6st | 2.752703405 | 3.785571298 | 4.464867139 | 2.64892811 |
| ENSDART00000172428 | FO704797.2 | 0.17851052 | -0.259187206 | -0.513103189 | -0.127941479 |
| ENSDART00000172430 | si:dkey-203a12.3 | 2.592857339 | 1.584928476 | 2.433004816 | 1.57133529 |
| ENSDART00000172433 | BX323031.2 | 0.315248431 | 0.209012025 | -0.000555571 | 0.219130901 |
| ENSDART00000172437 | dgkab | -0.119408916 | -0.169975589 | -0.468807215 | -0.381241335 |
| ENSDART00000172441 | lima1a | 0.350368785 | 0.588496789 | 0.46589541 | 0.178939257 |
| ENSDART00000172444 | ATG2A | 0.113672015 | 0.132095473 | 0.291990145 | 0.186645397 |
| ENSDART00000172460 | FO904965.1 | -0.329411304 | -0.282535563 | -0.05279435 | 0.008365177 |
| ENSDART00000172465 | gnb1b | -0.429380032 | -0.139749463 | 0.050389396 | 0.071415749 |
| ENSDART00000172496 | add3a | -0.173242509 | -0.314301562 | -0.183893521 | -0.110146899 |
| ENSDART00000172510 | elavl4 | 1.054519465 | 1.174247457 | 1.078840874 | 0.691066079 |
| ENSDART00000172515 | bzw1b | 0.541293913 | 0.477440982 | 0.388001444 | -0.287525957 |
| ENSDART00000172547 | insb | -0.25337952 | -0.120760297 | 0.28326011 | 0.898589828 |
| ENSDART00000172552 | dyrk2 | -1.00865602 | -4.097564958 | -0.48472737 | -0.263450257 |
| ENSDART00000172555 | rps15a | 0.386196694 | 0.41469411 | 0.231657808 | -0.026260131 |
| ENSDART00000172565 | Metazoa_SRP | 1.391513514 | 1.064695711 | 1.082381989 | 0.312800391 |
| ENSDART00000172566 | ptpro | -0.439852276 | 0.14868548 | 0.772757137 | 0.551836031 |
| ENSDART00000172624 | chn1 | -0.429491719 | -0.379515323 | -0.144891996 | -0.16828013 |
| ENSDART00000172634 | ryr3 | -1.057085419 | -1.586723574 | -0.163110923 | 0.331797887 |
| ENSDART00000172638 | gria3a | -0.308989617 | -0.39028425 | -0.157465183 | 0.147319123 |
| ENSDART00000172645 | unc5da | -0.375739047 | -0.576943357 | -0.183677467 | 0.03667029 |
| ENSDART00000172661 | nrgnb | -0.441646104 | -0.659637875 | -0.590118296 | -0.403632798 |
| ENSDART00000172662 | fam96a | -0.029501914 | -0.191990056 | -0.34693921 | -0.272376792 |
| ENSDART00000172664 | kirrel3l | -0.579597094 | -0.334224423 | -0.00269739 | 0.256102934 |
| ENSDART00000172672 | angptl2b | -0.321678324 | -0.001662718 | -0.330394771 | -0.841768465 |
| ENSDART00000172689 | drd4a | -0.269065132 | -0.433761644 | -0.356034882 | -0.090259185 |
| ENSDART00000172703 | trnau1apa | 3.525550379 | 4.03135258 | 2.121100592 | 1.557105834 |
| ENSDART00000172766 | psd3l | -0.547513249 | -0.802641524 | -0.228472689 | 0.098980101 |
| ENSDART00000172775 | tegt | -0.040616632 | 0.038263838 | -0.257883683 | -0.248241903 |
| ENSDART00000172811 | ubap1lb | 0.254294803 | -0.127702895 | -0.996786994 | -0.222342304 |
| ENSDART00000172821 | PHLPP2 | -0.058117589 | -0.547864216 | -0.458033727 | -0.042436898 |
| ENSDART00000172862 | kcnj14 | -0.439592228 | -0.323667837 | -0.482461949 | -0.197270237 |
| ENSDART00000172874 | kcnj10a | -0.114134993 | -0.28596151 | -0.331658021 | -0.188992319 |
| ENSDART00000172892 | CR769769.2 | 0.615702823 | 0.344273677 | 0.373739938 | 0.336791427 |
| ENSDART00000172893 | si:ch211-207g17.2 | 1.081136407 | 0.564546524 | -0.069203481 | -0.244746802 |
| ENSDART00000172899 | hp1bp3 | -0.268836711 | -0.197039259 | -0.254153947 | -0.074858409 |
| ENSDART00000172910 | map1aa | -0.286879932 | 0.224710371 | 0.48760533 | 0.689542868 |
| ENSDART00000172913 | si:ch73-368j24.11 | 0.642626776 | 0.575984051 | 0.848181147 | 0.478362255 |
| ENSDART00000172939 | ptp4a1 | 0.526587293 | 0.493567323 | 0.439467476 | 0.194682023 |
| ENSDART00000172944 | FO681390.1 | -0.061230657 | -0.175829366 | -0.634551101 | -0.148743958 |
| ENSDART00000172948 | TSC22D3 | 0.014403639 | -0.423782958 | -0.297835045 | -0.190699463 |
| ENSDART00000172953 | rpl29 | 0.354363113 | 0.457581567 | 0.206775836 | 0.026974968 |
| ENSDART00000172975 | slc6a1b | 0.350767184 | 0.274634418 | 0.412920252 | 0.546226395 |
| ENSDART00000172988 | si:zfos-80g12.1 | -1.526714169 | -3.752673892 | -1.791649817 | -0.447660424 |
| ENSDART00000173000 | zgc:113184 | 0.45806543 | 0.269621809 | 0.185268488 | 0.0480413 |
| ENSDART00000173011 | GRXCR1 | 0.401804428 | 0.903685752 | 1.561252834 | 1.150514684 |
| ENSDART00000173017 | atp2b2 | -0.485284755 | -0.507921276 | -0.213758166 | 0.050306247 |
| ENSDART00000173020 | utrn | -0.19228303 | -0.092812836 | -0.536357201 | -0.334471005 |

| | | | | | |
|---|---|---|---|---|---|
| ENSDART00000173052 | map7d2b | 0.261433378 | 1.055987677 | 1.048743858 | 0.7752628 |
| ENSDART00000173056 | arap3 | -0.20638853 | -0.271205037 | -0.427111789 | -0.285607029 |
| ENSDART00000173060 | rph3ab | -0.917567753 | -0.595167162 | 0.673778236 | 0.835535825 |
| ENSDART00000173072 | AKAP13 (1 of many) | -0.027726032 | -0.260210303 | -0.309219665 | 0.02079698 |
| ENSDART00000173083 | FO704610.1 | -0.146610059 | -0.222029359 | -0.309017936 | -0.101957892 |
| ENSDART00000173095 | spred3 | -0.542534865 | -0.320111081 | -0.209054264 | -0.626959008 |
| ENSDART00000173098 | gemin6 | 0.181263356 | 0.385271136 | 0.184325919 | 0.274013326 |
| ENSDART00000173108 | gpc3 | -0.979170892 | -0.757011653 | 0.064815285 | 0.413031736 |
| ENSDART00000173109 | nrtn | -0.527610722 | -0.443827628 | -0.46689713 | -0.285016979 |
| ENSDART00000173113 | si:ch211-129p13.1 | -0.464452233 | -0.696136912 | -0.478469763 | -0.206239945 |
| ENSDART00000173119 | pcdh11 | -0.634370446 | -0.732192902 | -0.531501202 | -0.25060062 |
| ENSDART00000173126 | klhl4 | -0.765151123 | -0.487744119 | -0.047554388 | -0.018081928 |
| ENSDART00000173133 | utrn | -0.11206335 | -0.324597481 | -0.223274112 | -0.197393841 |
| ENSDART00000173134 | BX571952.1 | -0.551010684 | -0.633352911 | -0.460069257 | -0.004126181 |
| ENSDART00000173143 | lrp1aa | 0.006114382 | -0.023577911 | -0.261635578 | -0.555062861 |
| ENSDART00000173145 | rpl29 | 0.32197024 | 0.280412475 | 0.068293144 | -0.107729465 |
| ENSDART00000173149 | dgkab | -0.056470672 | -0.094386691 | -0.40550946 | -0.281791004 |
| ENSDART00000173169 | pclob | -0.241926883 | -0.695024268 | -0.666850982 | -0.322920156 |
| ENSDART00000173192 | pcdh1b | -0.448371004 | -0.996454455 | -0.461950735 | 0.179661768 |
| ENSDART00000173195 | zgc:153146 | 2.291177749 | 2.913913042 | 2.827467346 | 2.606942928 |
| ENSDART00000173210 | kcnab2a | -0.633080178 | -0.760216594 | -0.295624932 | -0.02803528 |
| ENSDART00000173222 | gpr101 | -0.442714055 | -0.312795744 | -0.164884489 | -0.01182968 |
| ENSDART00000173228 | slc6a11b | -1.626653664 | -2.205227269 | -0.432079303 | 0.774903159 |
| ENSDART00000173237 | mid2 | -0.115282743 | -0.310001722 | -0.367171215 | -0.341782639 |
| ENSDART00000173257 | si:dkey-97l20.6 | -0.30598121 | -0.315922233 | -0.586659381 | -0.206936738 |
| ENSDART00000173267 | tmem255a | -0.148788206 | 0.632219165 | 0.395055581 | 0.430496671 |
| ENSDART00000173301 | sfxn5b | -0.333126218 | -0.098925577 | -0.061350927 | 0.22562903 |
| ENSDART00000173305 | si:ch1073-456m8.1 | 0.206580067 | 0.587987288 | 0.667691948 | 0.271613596 |
| ENSDART00000173346 | ptp4a1 | 0.667015175 | 0.565109215 | 0.920811931 | 0.492033169 |
| ENSDART00000173386 | cacna1da | -0.09786225 | -0.344688682 | -0.169985603 | 0.048564501 |
| ENSDART00000173398 | pcdh19 | -0.247538527 | -0.756226364 | 0.093604274 | 0.28627895 |
| ENSDART00000173400 | FP236327.1 | -0.107319508 | -0.233503644 | -0.613884847 | -0.314094176 |
| ENSDART00000173421 | lingo2a | -0.727375414 | -0.584905703 | -0.169839849 | 0.100856447 |
| ENSDART00000173423 | pcdh11 | -0.206659469 | -0.430916352 | -0.19368235 | 0.107223081 |
| ENSDART00000173430 | pfklb | -0.150367408 | -0.216221217 | -0.597100494 | -0.348773684 |
| ENSDART00000173436 | si:dkey-280e21.3 | 0.740189589 | 1.373516231 | 1.387529232 | 0.976737711 |
| TRANSGENE | | 3.62113341 | 4.519443337 | 2.76591932 | 1.041497447 |

**Table S2. Differentially expressed transcripts (1% FDR) clustered based on temporal expression pattern.**

| Ensembl ID | zebrafish gene symbol | padj | LFC: 2dpi-0dpi | LFC: 4dpi-0dpi | LFC: 7dpi-0dpi | LFC: 12dpi-0dpi | cluster name |
|---|---|---|---|---|---|---|---|
| ENSDART00000001678 | adam8a | 0.001434846 | 1.934717518 | 1.802449426 | 1.368408045 | -0.21633234 | growth toward the midline |
| ENSDART00000002453 | acsl4b | 0.004676707 | 0.682559347 | 0.484852434 | 0.056581739 | -0.183902113 | growth toward the midline |
| ENSDART00000002556 | mrto4 | 0.019563666 | 0.589078781 | 0.447556222 | 0.419365915 | 0.058126759 | growth toward the midline |
| ENSDART00000002595 | rpl21 | 0.004824695 | 0.36985839 | 0.3662658 | 0.201825483 | -0.004056179 | growth toward the midline |
| ENSDART00000002741 | itprip | 0.000474192 | 0.724094373 | 0.694272578 | 0.397322653 | -0.062298334 | growth toward the midline |
| ENSDART00000002945 | NPC2 (1 of many) | 0.001006824 | 3.182180208 | 2.136779184 | 1.408480393 | 0.09501531 | growth toward the midline |
| ENSDART00000003001 | rpl23a | 0.003882278 | 0.412203534 | 0.396855245 | 0.244635942 | -0.052563059 | growth toward the midline |
| ENSDART00000003314 | nusap1 | 0.035951327 | 0.825744448 | 0.908244509 | 0.647266522 | 0.3629471 | growth toward the midline |
| ENSDART00000004065 | zgc:91909 | 0.00334615 | 0.594004329 | 0.47164071 | 0.350756766 | 0.105236702 | growth toward the midline |
| ENSDART00000004392 | fkbp9 | 0.004270117 | 0.758370209 | 0.583896419 | 0.427971915 | -0.289954899 | growth toward the midline |
| ENSDART00000004664 | tram1 | 0.002818463 | 0.410276471 | 0.318086482 | 0.103470746 | -0.163408872 | growth toward the midline |
| ENSDART00000004780 | man2b1 | 0.002528575 | 0.352578138 | 0.195228445 | 0.062298599 | -0.189062834 | growth toward the midline |
| ENSDART00000006381 | psen2 | 0.01158873 | 0.338086739 | 0.204848015 | 0.067000942 | -0.024014416 | growth toward the midline |
| ENSDART00000007642 | zgc:110239 | 2.33589E-05 | 0.831334701 | 0.383173307 | 0.0742642 | -0.168536089 | growth toward the midline |
| ENSDART00000007827 | spra | 0.013245259 | 0.429147243 | 0.5653912 | 0.349944557 | -0.12347627 | growth toward the midline |
| ENSDART00000008373 | fosl1a | 0.001576361 | 0.78271954 | 0.570182098 | 0.226810387 | -0.332448721 | growth toward the midline |
| ENSDART00000008807 | rpl12 | 0.006094081 | 0.258765274 | 0.397825613 | 0.190712058 | -0.053203791 | growth toward the midline |
| ENSDART00000009164 | esco2 | 0.01613952 | 1.573729566 | 0.878542321 | 0.358323204 | 0.160150267 | growth toward the midline |
| ENSDART00000009241 | rpl35 | 0.001788569 | 0.440738233 | 0.45418276 | 0.310964715 | 0.055658665 | growth toward the midline |
| ENSDART00000009343 | pyroxd2 | 0.025731308 | 0.778556587 | 0.730253969 | 0.295998209 | 0.188824159 | growth toward the midline |
| ENSDART00000010104 | crtap | 0.001353924 | 0.225199403 | 0.115819527 | -0.25953471 | -0.588755148 | growth toward the midline |
| ENSDART00000010246 | ugt1ab | 0.02087678 | 1.578212697 | 1.169819697 | 0.509084599 | -0.046131609 | growth toward the midline |
| ENSDART00000010261 | pno1 | 0.000847723 | 0.396735226 | 0.379415162 | 0.20506239 | -0.042887698 | growth toward the midline |
| ENSDART00000011570 | zgc:101716 | 0.028885428 | 0.810996291 | 0.904458728 | 0.54949556 | 0.44631829 | growth toward the midline |
| ENSDART00000011863 | hdlbpa | 0.002972631 | 0.44689202 | 0.322223522 | 0.244841165 | -0.050194137 | growth toward the midline |
| ENSDART00000013409 | prmt3 | 0.000336326 | 0.583250552 | 0.486757337 | 0.420985526 | -0.028812313 | growth toward the midline |
| ENSDART00000014871 | akr7a3 | 0.017769648 | 0.445881137 | 0.133128629 | 0.063605049 | 0.058812872 | growth toward the midline |
| ENSDART00000015034 | blvrb | 0.000281817 | 0.700108065 | 0.61517331 | 0.539502173 | -0.031641171 | growth toward the midline |
| ENSDART00000015841 | gstt1b | 0.000149381 | 0.844147349 | 0.559916631 | 0.300223231 | -0.404935077 | growth toward the midline |
| ENSDART00000016814 | fmnl2a | 0.0006946 | 0.496433192 | 0.213976057 | -0.143298216 | -0.634969289 | growth toward the midline |
| ENSDART00000016864 | slc35f6 | 0.023157709 | 0.392001078 | 0.280281029 | 0.187768856 | 0.16995993 | growth toward the midline |
| ENSDART00000016890 | eif6 | 0.049349672 | 0.391815405 | 0.223439355 | 0.155230229 | 0.056706473 | growth toward the midline |
| ENSDART00000017153 | hps4 | 0.00377485 | 0.574215829 | 0.524013089 | 0.377950963 | 0.050651634 | growth toward the midline |
| ENSDART00000017176 | dkc1 | 0.026351976 | 0.45463167 | 0.261033913 | 0.276931433 | 0.02149911 | growth toward the midline |
| ENSDART00000018261 | akr1b1 | 0.003041627 | 1.00051996 | 0.934286328 | 0.670184322 | 0.161867768 | growth toward the midline |
| ENSDART00000018304 | mcm3 | 0.010839093 | 0.755786584 | 0.39595981 | 0.332438259 | 0.197655199 | growth toward the midline |
| ENSDART00000018461 | vmp1 | 0.013233841 | 0.430900752 | 0.323245721 | 0.184562524 | 0.049764543 | growth toward the midline |
| ENSDART00000018523 | ahcy | 0.020417188 | 0.45208938 | 0.606696476 | 0.316876579 | -0.072835379 | growth toward the midline |
| ENSDART00000018625 | napab | 0.000529346 | 0.495528529 | 0.395594582 | 0.072464634 | -0.31666363 | growth toward the midline |
| ENSDART00000019003 | psmd10 | 0.013692958 | 0.510870261 | 0.455922769 | 0.143258659 | 0.040691699 | growth toward the midline |
| ENSDART00000019149 | rpl7 | 0.003098732 | 0.429749031 | 0.41708455 | 0.235093103 | -0.015027688 | growth toward the midline |
| ENSDART00000019937 | gadd45ga | 0.013812134 | 0.98351774 | 1.304995817 | 0.165178221 | 0.462881948 | growth toward the midline |
| ENSDART00000020167 | slc16a9a | 0.000935288 | 0.895506163 | 0.735954298 | 0.591907512 | -0.466816888 | growth toward the midline |
| ENSDART00000020252 | pdia6 | 0.043796537 | 0.374033372 | 0.009098395 | 0.091775115 | 0.023604955 | growth toward the midline |
| ENSDART00000020311 | rpl27 | 0.001676296 | 0.430086554 | 0.381494789 | 0.182914397 | -0.034120883 | growth toward the midline |
| ENSDART00000020541 | lipf | 0.000672564 | 0.528006329 | 0.723139512 | 0.390796686 | -0.03933789 | growth toward the midline |
| ENSDART00000020655 | psma5 | 0.023717938 | 0.424960543 | 0.53028483 | 0.342202642 | 0.090736505 | growth toward the midline |
| ENSDART00000020810 | sdcbp2 | 0.003682945 | 0.429533724 | 0.242026939 | 0.181097661 | -0.126536592 | growth toward the midline |
| ENSDART00000021037 | hspa4a | 0.022587526 | 0.42074258 | 0.381012047 | 0.243421438 | 0.132816774 | growth toward the midline |
| ENSDART00000021069 | rpl38 | 0.001331938 | 0.561397297 | 0.414839383 | 0.25425723 | 0.049186317 | growth toward the midline |
| ENSDART00000021299 | nmd3 | 0.014686352 | 0.353895501 | 0.176055483 | 0.10015969 | 0.031379837 | growth toward the midline |
| ENSDART00000021788 | pbk | 0.043154442 | 1.438031033 | 0.973567544 | 0.667760742 | 0.195406357 | growth toward the midline |
| ENSDART00000021798 | fabp11a | 0.000424137 | 3.578286828 | 1.692122399 | 1.073895347 | -0.266988673 | growth toward the midline |
| ENSDART00000021950 | mthfd1b | 0.002507467 | 1.089156306 | 0.599589593 | 0.18588711 | -0.096895618 | growth toward the midline |
| ENSDART00000022307 | atic | 4.11188E-05 | 1.012801143 | 0.518350872 | 0.289570342 | -0.11925608 | growth toward the midline |
| ENSDART00000022562 | rhogb | 0.013807548 | 0.97771224 | 0.454579503 | 0.287206454 | -0.277565993 | growth toward the midline |
| ENSDART00000022586 | lrrc40 | 0.005807456 | 0.466457292 | 0.391920358 | 0.288821604 | -0.015697706 | growth toward the midline |
| ENSDART00000022634 | acp2 | 0.004900248 | 0.356430058 | 0.295051752 | 0.030540215 | -0.026401831 | growth toward the midline |
| ENSDART00000022765 | riok1 | 0.014326582 | 0.429965951 | 0.477495547 | 0.301061387 | 0.021181676 | growth toward the midline |
| ENSDART00000023463 | uap1l1 | 0.015884799 | 0.706648736 | 0.512468414 | 0.225773419 | -0.015907861 | growth toward the midline |
| ENSDART00000024194 | kif11 | 0.007786427 | 1.409119378 | 1.269759488 | 0.810392576 | 0.401186663 | growth toward the midline |
| ENSDART00000024208 | nutf2l | 0.001633797 | 0.951686549 | 0.855710394 | 0.537014601 | -0.220899011 | growth toward the midline |
| ENSDART00000024316 | mcm5 | 0.041407171 | 1.292009744 | 0.913737369 | 0.774968175 | 0.275487022 | growth toward the midline |
| ENSDART00000025096 | larp1b | 0.002618468 | 0.593217694 | 0.576647108 | 0.472990072 | 0.196836578 | growth toward the midline |
| ENSDART00000025229 | adi1 | 0.000305093 | 0.582007509 | 0.274560712 | -0.200617197 | -0.21352575 | growth toward the midline |
| ENSDART00000025912 | si:dkey-32n7.4 | 0.055404088 | 0.676729111 | 0.251567535 | 0.290364244 | 0.056388056 | growth toward the midline |
| ENSDART00000026145 | AMOTL1 | 0.001213585 | 1.0586494 | 0.570711762 | 0.741896115 | 0.335943589 | growth toward the midline |
| ENSDART00000026178 | kif4 | 0.009637478 | 0.630775369 | 0.400037801 | 0.195646982 | 0.069714171 | growth toward the midline |
| ENSDART00000026492 | flncb | 0.000598789 | 1.767414623 | 1.156524725 | 0.931280558 | 0.481368808 | growth toward the midline |
| ENSDART00000027393 | ckmt1 | 0.033377037 | 0.722688207 | 0.598583347 | 0.435615456 | 0.204934783 | growth toward the midline |
| ENSDART00000029133 | snu13b | 0.001170697 | 0.760760317 | 0.56984098 | 0.429270049 | -0.029914173 | growth toward the midline |
| ENSDART00000029380 | bnip4 | 0.000434317 | 0.50246171 | 0.517948631 | -0.062532971 | -0.337429982 | growth toward the midline |

| | | | | | | | |
|---|---|---|---|---|---|---|---|
| ENSDART00000029387 | ppan | 0.025186512 | 0.365110957 | 0.289310564 | 0.234072786 | 0.008675203 | growth toward the midline |
| ENSDART00000030125 | znhit3 | 0.032553766 | 0.760210037 | 0.691966223 | 0.487369205 | 0.388320697 | growth toward the midline |
| ENSDART00000030890 | hmox1a | 0.001147964 | 2.623035746 | 1.043671484 | 1.096486903 | 0.061370069 | growth toward the midline |
| ENSDART00000031047 | cd63 | 0.001899261 | 1.025048213 | 0.521327717 | 0.067995805 | -0.605090068 | growth toward the midline |
| ENSDART00000031498 | ccna2 | 0.001588425 | 1.569664461 | 1.378683751 | 0.895010724 | 0.581055828 | growth toward the midline |
| ENSDART00000032290 | esyt1a | 0.001565956 | 0.414133861 | 0.186544426 | 0.071532791 | -0.03493067 | growth toward the midline |
| ENSDART00000032498 | tspan36 | 2.7529E-05 | 1.455763612 | 0.805559086 | 0.192569945 | -0.715366149 | growth toward the midline |
| ENSDART00000033386 | ocstamp | 0.013948629 | 2.954317372 | 2.548446344 | 2.458244526 | 1.328553982 | growth toward the midline |
| ENSDART00000033574 | slc24a5 | 0.00217038 | 0.460212086 | 0.054768672 | -0.306746704 | -1.141417252 | growth toward the midline |
| ENSDART00000033663 | rps21 | 0.005949363 | 0.311890725 | 0.366197651 | 0.105076559 | -0.038309008 | growth toward the midline |
| ENSDART00000033761 | glb1 | 0.021559058 | 0.347409105 | 0.187375641 | 0.031649312 | 0.073918812 | growth toward the midline |
| ENSDART00000033980 | lims1 | 0.002007275 | 0.497849916 | 0.369061975 | 0.247283344 | -0.174205881 | growth toward the midline |
| ENSDART00000034248 | rab32a | 0.001863533 | 0.364170748 | 0.270180665 | -0.161567701 | -0.659765413 | growth toward the midline |
| ENSDART00000034377 | cpa5 | 0.000100528 | 2.043914839 | 0.12107472 | 0.19928898 | 0.161423339 | growth toward the midline |
| ENSDART00000034829 | rrp12 | 0.00217038 | 0.494057186 | 0.423927766 | 0.372142811 | 0.099318986 | growth toward the midline |
| ENSDART00000034850 | dbi | 0.002568245 | 1.011739121 | 0.830016329 | 0.708552366 | 0.209278197 | growth toward the midline |
| ENSDART00000035093 | col9a2 | 0.002649033 | 0.360349924 | 0.265960604 | -0.052354611 | -0.707154609 | growth toward the midline |
| ENSDART00000035670 | polr2eb | 0.0033491 | 0.35113115 | 0.294169174 | 0.063963778 | -0.045691283 | growth toward the midline |
| ENSDART00000035739 | tmem134 | 0.001247858 | 0.246242014 | 0.369929781 | 0.082611565 | -0.161334554 | growth toward the midline |
| ENSDART00000035944 | clic5a | 0.068646243 | 2.554130845 | 1.311062136 | 1.150965478 | 0.646568675 | growth toward the midline |
| ENSDART00000036729 | spi1b | 0.004358158 | 0.910663058 | 0.717004711 | 0.095052493 | -0.117318513 | growth toward the midline |
| ENSDART00000037224 | cst14a.2 | 0.003240086 | 0.549538616 | 0.553010386 | 0.34149596 | -0.057482802 | growth toward the midline |
| ENSDART00000038202 | cndp2 | 0.001490782 | 1.63370988 | 0.725493655 | 0.197899383 | -0.10985912 | growth toward the midline |
| ENSDART00000038301 | gnpda2 | 0.008531138 | 0.314221876 | 0.240267665 | 0.128522583 | -0.033807169 | growth toward the midline |
| ENSDART00000038696 | flvcr2b | 0.014130752 | 1.475160998 | 0.41510575 | 0.393658072 | -0.063620422 | growth toward the midline |
| ENSDART00000038888 | hsdl2 | 0.018248495 | 0.331360296 | 0.293802861 | 0.248668235 | 0.085382052 | growth toward the midline |
| ENSDART00000038990 | jak1 | 0.006256994 | 0.414591598 | 0.220201928 | 0.070657217 | -0.102548961 | growth toward the midline |
| ENSDART00000039206 | rps23 | 0.004633654 | 0.388349294 | 0.322594165 | 0.070172062 | -0.095059659 | growth toward the midline |
| ENSDART00000039312 | ipo4 | 0.002755913 | 0.472349209 | 0.340323729 | 0.32654819 | 0.07839591 | growth toward the midline |
| ENSDART00000040434 | asah1b | 0.000524004 | 0.563509259 | 0.420868789 | -0.056225927 | -0.32971968 | growth toward the midline |
| ENSDART00000040771 | rpl34 | 0.005202212 | 0.291435523 | 0.406434571 | 0.222752448 | -0.006604457 | growth toward the midline |
| ENSDART00000040827 | ncaph2 | 0.025606413 | 0.654906877 | 0.777407912 | 0.63027477 | 0.226543553 | growth toward the midline |
| ENSDART00000040900 | baxb | 0.005486177 | 0.66837016 | 0.786823818 | 0.515591205 | 0.094501002 | growth toward the midline |
| ENSDART00000041114 | psmb2 | 0.010442784 | 0.343581501 | 0.272779677 | 0.256301247 | -0.006269993 | growth toward the midline |
| ENSDART00000041504 | tescb | 0.004106215 | 0.44458368 | -0.282095189 | 0.067591772 | -0.116025945 | growth toward the midline |
| ENSDART00000041992 | dhrs12 | 0.001102685 | 0.21282475 | 0.60945999 | -0.222288782 | -0.248821854 | growth toward the midline |
| ENSDART00000042250 | rap1b | 0.003665031 | 0.339690926 | 0.041784403 | 0.019413921 | -0.132436477 | growth toward the midline |
| ENSDART00000043173 | rpl18 | 0.005040201 | 0.389153851 | 0.347510342 | 0.235541148 | 0.00423952 | growth toward the midline |
| ENSDART00000043666 | hs1bp3 | 0.003138642 | 1.060777617 | 0.463082325 | 0.168627927 | -0.515305865 | growth toward the midline |
| ENSDART00000043678 | apobec2b | 0.024850978 | 1.815363093 | 0.931239114 | 0.794076282 | 0.812995775 | growth toward the midline |
| ENSDART00000044238 | zgc:92066 | 0.000787471 | 1.060411872 | 1.124461317 | 0.621339815 | 0.051820002 | growth toward the midline |
| ENSDART00000044264 | mmp14b | 0.018216055 | 0.997152397 | 1.115840894 | 0.915921565 | 0.094130452 | growth toward the midline |
| ENSDART00000044314 | itgav | 0.004681738 | 0.188708464 | -0.009103257 | -0.190172369 | -0.475418475 | growth toward the midline |
| ENSDART00000044328 | acss1 | 0.000783264 | 0.948368639 | 0.660526303 | 0.176495325 | -0.460639914 | growth toward the midline |
| ENSDART00000045933 | sh3glb1b | 0.002439392 | 0.424660866 | 0.274153114 | 0.044458674 | -0.153303197 | growth toward the midline |
| ENSDART00000046218 | flnca | 0.000103226 | 6.980277858 | 6.302871335 | 6.08375751 | 5.531037957 | growth toward the midline |
| ENSDART00000046253 | prkcq | 0.008130201 | 0.856641094 | 1.199706901 | 0.196829623 | -0.396081861 | growth toward the midline |
| ENSDART00000046268 | pmelb | 0.000596344 | 0.894230709 | 0.185058494 | -0.384535315 | -0.811085128 | growth toward the midline |
| ENSDART00000047191 | glb1l | 0.017715642 | 1.329711044 | 0.552831622 | 0.213363411 | -0.364178392 | growth toward the midline |
| ENSDART00000047728 | melk | 0.061287033 | 0.808255278 | 0.366287521 | 0.23278244 | 0.524042022 | growth toward the midline |
| ENSDART00000048107 | FP102018.1 | 0.000888975 | 0.402516182 | 0.222629184 | -0.14999501 | -0.581967671 | growth toward the midline |
| ENSDART00000049465 | slc19a1 | 0.0055878 | 0.32288456 | 0.126425631 | -0.165912503 | -0.040050165 | growth toward the midline |
| ENSDART00000050399 | npc2 | 0.002939613 | 0.729567353 | 0.287798394 | -0.063690935 | -0.310097689 | growth toward the midline |
| ENSDART00000050847 | GLDC | 0.017654181 | 0.384969982 | 0.265449522 | 0.198357794 | 0.108750757 | growth toward the midline |
| ENSDART00000051392 | spns3 | 0.001042039 | 1.320734766 | 0.750958136 | 0.798555508 | -0.096372125 | growth toward the midline |
| ENSDART00000051491 | sfrp1a | 0.001342771 | 0.324084954 | 0.211296795 | -0.122014524 | -1.0775933 | growth toward the midline |
| ENSDART00000051556 | abca1b | 0.004394152 | 0.962705135 | 0.587027009 | 0.239926413 | -0.279230387 | growth toward the midline |
| ENSDART00000051763 | rps3a | 0.02114031 | 0.36528781 | 0.402890251 | 0.287932185 | 0.011460462 | growth toward the midline |
| ENSDART00000051906 | ube2c | 0.012497601 | 1.239012745 | 1.032037835 | 0.553625781 | 0.164264884 | growth toward the midline |
| ENSDART00000052082 | rpl30 | 0.00417836 | 0.377785904 | 0.342931306 | 0.098750304 | -0.086094728 | growth toward the midline |
| ENSDART00000052090 | fuca1.2 | 0.012864775 | 0.607737239 | 0.489644017 | 0.400376939 | 0.091450373 | growth toward the midline |
| ENSDART00000052104 | fuca1.1 | 0.006465135 | 0.706031916 | 0.625980807 | 0.554044121 | 0.174374129 | growth toward the midline |
| ENSDART00000052331 | rps20 | 0.004925327 | 0.422414342 | 0.403711723 | 0.215205833 | -0.059084102 | growth toward the midline |
| ENSDART00000052730 | rps13 | 0.00538212 | 0.392793291 | 0.407259594 | 0.231403052 | 0.009148185 | growth toward the midline |
| ENSDART00000052761 | rpl39 | 0.005353377 | 0.335691979 | 0.34926929 | 0.1141905 | -0.000371665 | growth toward the midline |
| ENSDART00000052917 | slc3a2a | 0.001434846 | 0.509465627 | 0.333845604 | -0.024881296 | -0.285712584 | growth toward the midline |
| ENSDART00000053761 | bms1 | 0.015390643 | 0.415838377 | 0.296751152 | 0.241962107 | 0.100266391 | growth toward the midline |
| ENSDART00000053932 | cbsa | 0.000261206 | 1.196219296 | 0.806157212 | 0.553945426 | 0.279319174 | growth toward the midline |
| ENSDART00000055287 | zgc:109934 | 0.00013154 | 0.762332695 | 0.487859158 | -0.045189444 | -0.423101752 | growth toward the midline |
| ENSDART00000055328 | nek6 | 0.003694199 | 0.491226657 | 0.428884947 | 0.11516767 | -0.052210357 | growth toward the midline |
| ENSDART00000055428 | cbx7a | 0.000405977 | 0.780269243 | 0.615351279 | 0.524687311 | -0.270346219 | growth toward the midline |
| ENSDART00000056005 | ascl1a | 0.032093457 | 1.787445127 | 1.283325532 | 0.749862779 | 0.792492648 | growth toward the midline |
| ENSDART00000056457 | mitfa | 0.005353377 | 0.929754118 | 0.38849893 | 0.30836176 | -0.311729197 | growth toward the midline |
| ENSDART00000056460 | gbp1 | 0.045564466 | 3.785035011 | 3.151116027 | 3.74676295 | 2.092352698 | growth toward the midline |
| ENSDART00000056686 | mrc1b | 0.004463264 | 0.993000172 | 0.810204425 | 0.489266441 | -0.068136928 | growth toward the midline |

| | | | | | | | |
|---|---|---|---|---|---|---|---|
| ENSDART00000058324 | rpz4 | 0.00855067 | 0.754354311 | 0.61115296 | 0.469864295 | -0.17226809 | growth toward the midline |
| ENSDART00000058605 | scpep1 | 0.002139835 | 0.659557613 | 0.259300703 | 0.157418507 | -0.140654612 | growth toward the midline |
| ENSDART00000058665 | kif20bb | 0.029064716 | 0.867797834 | 0.730592963 | 0.371647572 | 0.126469391 | growth toward the midline |
| ENSDART00000058774 | havcr1 | 0.002175185 | 1.378230425 | 0.803624411 | 0.332422006 | 0.11017803 | growth toward the midline |
| ENSDART00000059001 | C5AR1 | 0.048093489 | 0.862718576 | 0.211750471 | 0.205995335 | 0.011542683 | growth toward the midline |
| ENSDART00000059013 | sec61b | 0.022498542 | 0.564454151 | 0.292288284 | 0.352954607 | 0.073391744 | growth toward the midline |
| ENSDART00000059667 | wdr75 | 0.022389941 | 0.34735847 | 0.29669502 | 0.189255306 | 0.039773826 | growth toward the midline |
| ENSDART00000060363 | rpl4 | 0.004878958 | 0.373964359 | 0.396853604 | 0.212646351 | -0.016195336 | growth toward the midline |
| ENSDART00000060444 | rps29 | 0.007150368 | 0.369322836 | 0.397792199 | 0.13443893 | -0.022749507 | growth toward the midline |
| ENSDART00000060745 | uba52 | 0.01809863 | 0.32327466 | 0.351581625 | 0.155851402 | -0.01111286 | growth toward the midline |
| ENSDART00000060765 | nppb | 0.001237101 | 4.501656565 | 4.501350371 | 1.812509246 | -0.191856821 | growth toward the midline |
| ENSDART00000060773 | taar12b | 0.025216223 | 2.538886261 | 1.95415769 | 1.735717491 | 1.72259581 | growth toward the midline |
| ENSDART00000060865 | rasal1b | 0.024284969 | 1.120636523 | 0.769242819 | 0.458217171 | 0.392656742 | growth toward the midline |
| ENSDART00000061736 | si:dkey-4e7.3 | 0.017999639 | 0.80946555 | 0.660586871 | 0.14634027 | 0.03058403 | growth toward the midline |
| ENSDART00000061926 | stx11b.2 | 0.006909332 | 3.447906519 | 2.671571656 | 2.785952697 | 1.680276674 | growth toward the midline |
| ENSDART00000062257 | slc39a1 | 0.003541697 | 0.740781667 | 0.522946282 | 0.350404642 | 0.245154327 | growth toward the midline |
| ENSDART00000062576 | thyn1 | 0.006623227 | 0.902996379 | 0.741810201 | 0.792519989 | 0.219218083 | growth toward the midline |
| ENSDART00000062671 | tuba8l | 0.009931768 | 1.253108099 | 0.545680562 | 0.42270127 | -0.336104805 | growth toward the midline |
| ENSDART00000062845 | mmp9 | 0.014615758 | 2.08287012 | 1.917473556 | 1.298502519 | -0.06470518 | growth toward the midline |
| ENSDART00000062931 | abracl | 0.011668571 | 0.878200021 | 0.73529721 | 0.710676136 | 0.289919909 | growth toward the midline |
| ENSDART00000062983 | rpl10a | 0.003138642 | 0.388498934 | 0.402298329 | 0.17374506 | -0.059011317 | growth toward the midline |
| ENSDART00000063107 | cyp2p7 | 0.024411518 | 0.910849098 | 1.165275545 | 0.616791402 | -0.023701278 | growth toward the midline |
| ENSDART00000063251 | ctsz | 3.54236E-05 | 1.117383701 | 0.530773826 | 0.448185866 | 0.142841944 | growth toward the midline |
| ENSDART00000063357 | ccnb1 | 0.012018385 | 1.564145903 | 1.350801576 | 0.91228963 | 0.49209964 | growth toward the midline |
| ENSDART00000063764 | si:dkey-5n18.1 | 0.015077117 | 1.420288455 | 0.932190706 | 0.716992196 | -0.355341421 | growth toward the midline |
| ENSDART00000063874 | vamp4 | 0.003602728 | 0.706049434 | 0.688342777 | 0.412447235 | 0.06985511 | growth toward the midline |
| ENSDART00000063953 | zgc:65997 | 0.015420308 | 0.11562047 | 0.484476273 | -0.11592626 | -0.024456549 | growth toward the midline |
| ENSDART00000064113 | abt1 | 0.009820514 | 0.598506286 | 0.454995795 | 0.386684419 | 0.116646594 | growth toward the midline |
| ENSDART00000064511 | il17a/f1 | 0.023872337 | 1.319538931 | 1.072672417 | 0.769667562 | 1.083526716 | growth toward the midline |
| ENSDART00000064657 | stx11a | 0.006345193 | 1.187636483 | 0.704469765 | 0.33181163 | -0.135694961 | growth toward the midline |
| ENSDART00000064700 | fuca2 | 0.001008722 | 0.436466876 | 0.137195427 | -0.005101419 | -0.171105064 | growth toward the midline |
| ENSDART00000065057 | itgb7 | 0.000118474 | 1.200094562 | 0.885337917 | 0.497721229 | 0.006866552 | growth toward the midline |
| ENSDART00000065183 | cldn2 | 0.000739355 | 0.624880574 | 0.284479914 | -0.382379938 | -0.379228488 | growth toward the midline |
| ENSDART00000065208 | nop16 | 0.021355398 | 0.539941841 | 0.465464987 | 0.330311237 | 0.01578578 | growth toward the midline |
| ENSDART00000065337 | kif20a | 0.001229585 | 1.518648505 | 1.334799108 | 0.992824147 | 0.572892998 | growth toward the midline |
| ENSDART00000065674 | fybb | 0.008250091 | 0.68375018 | 0.334314613 | 0.079534024 | -0.242430054 | growth toward the midline |
| ENSDART00000065805 | tspan10 | 0.003267854 | 0.90782785 | 0.406843364 | -0.022479662 | -0.576522773 | growth toward the midline |
| ENSDART00000065817 | pou5f3 | 0.012966692 | 1.443769101 | 1.494882152 | 0.928756649 | 0.660724256 | growth toward the midline |
| ENSDART00000066411 | dlgap5 | 0.003138642 | 1.041688171 | 0.895823709 | 0.540002804 | 0.33349173 | growth toward the midline |
| ENSDART00000066778 | acad11 | 0.000741396 | 0.796145744 | 0.566674022 | 0.216041695 | -0.190350168 | growth toward the midline |
| ENSDART00000067193 | adm2a | 0.004020103 | 0.4356689 | 0.184728608 | -0.002686414 | -1.106691874 | growth toward the midline |
| ENSDART00000067312 | si:dkey-106n21.1 | 0.003805801 | 0.584969753 | 0.102146183 | -0.031661382 | 0.069325035 | growth toward the midline |
| ENSDART00000067594 | cst14b.1 | 0.003629485 | 1.003408293 | 1.132156972 | 0.601894798 | 0.102871452 | growth toward the midline |
| ENSDART00000067599 | angptl2a | 0.000103226 | 1.671461404 | 1.166310219 | 0.762115684 | 0.407368061 | growth toward the midline |
| ENSDART00000067678 | zgc:110339 | 0.001560042 | 0.949255799 | 0.562708107 | 0.125395868 | -0.585210787 | growth toward the midline |
| ENSDART00000073846 | si:ch211-122f10.4 | 0.006183684 | 0.620706192 | 0.26696454 | 0.219379439 | -0.011733355 | growth toward the midline |
| ENSDART00000074010 | ubald1b | 0.002498008 | 0.486188172 | 0.523878559 | 0.246068485 | 0.014830353 | growth toward the midline |
| ENSDART00000074161 | slc4a2b | 0.000953608 | 0.481685785 | 0.057728735 | -0.200942778 | -0.247347753 | growth toward the midline |
| ENSDART00000074384 | stx4 | 0.006812131 | 0.365157378 | 0.335603422 | 0.150714004 | -0.050227712 | growth toward the midline |
| ENSDART00000074438 | cenpi | 0.023769243 | 0.547415003 | 0.708397465 | 0.401862169 | 0.054393549 | growth toward the midline |
| ENSDART00000075028 | rps11 | 0.001443362 | 0.42100484 | 0.441088802 | 0.128015907 | -0.080449898 | growth toward the midline |
| ENSDART00000075262 | cad | 0.058741505 | 0.41294438 | 0.156970425 | 0.074105835 | 0.04769287 | growth toward the midline |
| ENSDART00000075286 | slc2a15b | 0.000993187 | 1.023992215 | 0.671159354 | 0.170031714 | -0.73573625 | growth toward the midline |
| ENSDART00000075495 | rpl23 | 0.005969755 | 0.390673252 | 0.341184618 | 0.264108815 | -0.005521244 | growth toward the midline |
| ENSDART00000076030 | fbl | 0.017381552 | 0.33280558 | 0.192834191 | 0.116054515 | 0.005015438 | growth toward the midline |
| ENSDART00000076636 | fzd2 | 0.002013265 | 0.563199333 | 0.424474998 | 0.171148118 | -0.301884624 | growth toward the midline |
| ENSDART00000077185 | dgat1b | 0.002147221 | 0.79138422 | 0.60487817 | 0.419420549 | -0.346047138 | growth toward the midline |
| ENSDART00000077259 | ebna1bp2 | 0.007621679 | 0.420936188 | 0.344892308 | 0.188849483 | 0.008127131 | growth toward the midline |
| ENSDART00000077386 | prss16 | 0.000185931 | 0.714972584 | 0.32483027 | 0.071612523 | -0.240875958 | growth toward the midline |
| ENSDART00000077418 | ctsba | 0.016904127 | 1.550723066 | 1.033464808 | 0.97579905 | 0.475659932 | growth toward the midline |
| ENSDART00000077511 | ccr9a | 0.000592988 | 1.513300322 | 1.48326526 | 0.981085195 | 0.082724583 | growth toward the midline |
| ENSDART00000077545 | slc7a7 | 5.02856E-05 | 1.304066715 | 0.45830859 | 0.136139102 | 0.056436424 | growth toward the midline |
| ENSDART00000077707 | llph | 0.006605202 | 0.612018484 | 0.617804437 | 0.382050706 | 0.042759152 | growth toward the midline |
| ENSDART00000077715 | si:dkey-25e12.3 | 0.000947019 | 1.07190149 | 0.953044464 | 0.701373655 | 0.117439727 | growth toward the midline |
| ENSDART00000077834 | rps27.2 | 0.003329747 | 0.365621497 | 0.39961171 | 0.113670294 | -0.081169937 | growth toward the midline |
| ENSDART00000078266 | rsl1d1 | 0.000775831 | 0.351365949 | 0.392552672 | 0.175928217 | -0.070353732 | growth toward the midline |
| ENSDART00000078596 | hspd1 | 0.004163372 | 0.512169965 | 0.358816938 | 0.201006014 | 0.20720259 | growth toward the midline |
| ENSDART00000078634 | eif5a2 | 0.011846009 | 0.493280465 | 0.485481598 | 0.418411017 | 0.051860815 | growth toward the midline |
| ENSDART00000078647 | si:ch211-201h21.5 | 0.024809879 | 1.148616031 | 0.689283012 | 0.40532037 | 0.049879214 | growth toward the midline |
| ENSDART00000078866 | ifi30 | 0.001277839 | 1.574791131 | 0.896478525 | 0.373397245 | -0.020268978 | growth toward the midline |
| ENSDART00000079222 | chaf1b | 0.034090908 | 1.242408116 | 0.820706616 | 0.642546621 | 0.111426229 | growth toward the midline |
| ENSDART00000079310 | manba | 0.003137832 | 0.495609624 | 0.109254813 | 0.005925635 | -0.12409296 | growth toward the midline |
| ENSDART00000079549 | tpte | 0.002192481 | 0.809167451 | 0.638911968 | 0.260848622 | -0.250321254 | growth toward the midline |
| ENSDART00000079695 | zwilch | 0.007014909 | 1.159509442 | 0.995361025 | 0.861360163 | 0.105564769 | growth toward the midline |
| ENSDART00000079984 | rpl22l1 | 0.0003783 | 0.678692893 | 0.584241724 | 0.331994973 | -0.011632328 | growth toward the midline |

| | | | | | | | |
|---|---|---|---|---|---|---|---|
| ENSDART00000080339 | galm | 0.000273326 | 0.868912331 | 0.484945087 | -0.111697277 | -0.429352779 | growth toward the midline |
| ENSDART00000080377 | aldoca | 0.036855016 | 2.758856293 | 1.548001605 | 2.09362575 | 1.794239524 | growth toward the midline |
| ENSDART00000080465 | hells | 0.018490842 | 0.812911201 | 0.936234984 | 0.557478835 | 0.182957003 | growth toward the midline |
| ENSDART00000080523 | itgae.2 | 0.00855067 | 1.067269567 | 0.812178684 | 0.361968863 | 0.003662922 | growth toward the midline |
| ENSDART00000080712 | slc43a3b | 0.00855067 | 2.109023631 | 1.075213126 | 0.724428131 | -0.154263604 | growth toward the midline |
| ENSDART00000080864 | magt1 | 0.006145665 | 0.441285986 | 0.302582629 | 0.047526992 | -0.172355248 | growth toward the midline |
| ENSDART00000080919 | rpl36a | 0.000996442 | 0.405736348 | 0.485454355 | 0.225930563 | -0.002722727 | growth toward the midline |
| ENSDART00000081343 | plk1 | 0.006812131 | 1.547304713 | 1.292689538 | 1.02080622 | 0.322979307 | growth toward the midline |
| ENSDART00000081568 | tcf19l | 0.027985198 | 0.712871286 | 0.397385898 | 0.351815803 | 0.227291981 | growth toward the midline |
| ENSDART00000082011 | lim2.2 | 0.05615844 | 1.304610538 | 0.978295786 | 0.86763989 | 0.856876261 | growth toward the midline |
| ENSDART00000082050 | zgc:174904 | 0.01376164 | 1.977093547 | 1.430607855 | 1.439069833 | 0.228460651 | growth toward the midline |
| ENSDART00000082471 | mfap2 | 0.013128447 | 0.61383585 | 0.779035446 | 0.354750417 | 0.055843134 | growth toward the midline |
| ENSDART00000082517 | rab43 | 0.000993187 | 0.357940531 | 0.212689136 | -0.148032038 | -0.068856369 | growth toward the midline |
| ENSDART00000083033 | sik1 | 0.0147298 | 1.146093805 | 0.631897288 | 0.512350737 | 1.123188354 | growth toward the midline |
| ENSDART00000083126 | cidec | 0.00067643 | 3.092638419 | 2.402632883 | 1.147285193 | -0.231828568 | growth toward the midline |
| ENSDART00000083294 | nol6 | 0.046592019 | 0.354535252 | 0.232360208 | 0.196448331 | 0.109158776 | growth toward the midline |
| ENSDART00000083830 | sdc4 | 0.000487602 | 0.356242427 | 0.466156404 | 0.190102023 | -0.370187936 | growth toward the midline |
| ENSDART00000084069 | iqgap2 | 0.000512675 | 1.059547107 | 0.55852141 | 0.220571876 | -0.354231411 | growth toward the midline |
| ENSDART00000085309 | dpcd | 0.00270716 | 0.675929667 | 0.665840567 | 0.458693669 | 0.16198721 | growth toward the midline |
| ENSDART00000085764 | PLOD3 | 0.005493417 | 0.414167142 | 0.202506542 | 0.107388816 | -0.10168282 | growth toward the midline |
| ENSDART00000087114 | alg5 | 0.020108219 | 0.317519293 | 0.464842808 | 0.220188706 | 0.059168575 | growth toward the midline |
| ENSDART00000087311 | oca2 | 0.000313682 | 0.449332393 | 0.338475194 | -0.09076694 | -0.174077968 | growth toward the midline |
| ENSDART00000088042 | myo10l3 | 0.014447102 | 0.440398704 | 0.46516816 | 0.325219038 | -0.023584336 | growth toward the midline |
| ENSDART00000088973 | sytl2a | 0.000317136 | 0.735131572 | 0.479031428 | -0.049007315 | -0.597552798 | growth toward the midline |
| ENSDART00000089158 | hmha1a | 0.027703711 | 1.15366088 | 0.773436896 | 0.705540127 | -0.035689576 | growth toward the midline |
| ENSDART00000090528 | rhoca | 7.15704E-05 | 0.570199757 | 0.738551724 | 0.41277501 | -0.074138712 | growth toward the midline |
| ENSDART00000090883 | gpnmb | 0.001083951 | 1.78024327 | 1.792109343 | 1.484400673 | -0.264951498 | growth toward the midline |
| ENSDART00000091662 | noc2l | 0.007264334 | 0.778877814 | 0.364993344 | 0.393512667 | -0.105586454 | growth toward the midline |
| ENSDART00000092050 | stab1 | 0.023363274 | 0.830009942 | 0.662655195 | 0.610953846 | 0.245694157 | growth toward the midline |
| ENSDART00000092389 | nup210 | 0.027686308 | 0.743287563 | 0.315006844 | 0.505674911 | 0.088834448 | growth toward the midline |
| ENSDART00000093279 | spi1b | 0.002640644 | 0.891535687 | 0.563855895 | 0.100587929 | -0.531177734 | growth toward the midline |
| ENSDART00000098545 | tmem150aa | 0.010056242 | 0.458575321 | 0.166363456 | 0.137813682 | -0.033806431 | growth toward the midline |
| ENSDART00000098970 | lin28a | 0.050549016 | 3.476899763 | 1.864207832 | 0.92280153 | 0.893269961 | growth toward the midline |
| ENSDART00000099138 | ncf2 | 0.012263789 | 0.590402106 | 0.352177279 | 0.071326562 | -0.071022128 | growth toward the midline |
| ENSDART00000099839 | map2k2b | 0.04077574 | 4.411409189 | 4.089988152 | 4.160217635 | 3.732217772 | growth toward the midline |
| ENSDART00000099947 | samsn1a | 0.007966006 | 0.989297054 | 0.26224771 | 0.078735734 | -0.489522484 | growth toward the midline |
| ENSDART00000099978 | pdlim4 | 0.008900663 | 0.618309518 | 0.542997573 | 0.115667653 | -0.045450127 | growth toward the midline |
| ENSDART00000100145 | lgals9l1 | 0.008814877 | 1.730493466 | 0.73620688 | 0.17444521 | -0.331961456 | growth toward the midline |
| ENSDART00000100458 | si:dkey-73n10.1 | 0.0021138 | 1.648347098 | 0.766263068 | 0.204715566 | -0.361349683 | growth toward the midline |
| ENSDART00000100473 | PLIN3 | 0.014485141 | 1.275027857 | 0.887739595 | 0.535269477 | -0.111777778 | growth toward the midline |
| ENSDART00000100813 | rps24 | 0.004853289 | 0.369574152 | 0.327820695 | 0.143165091 | -0.049601165 | growth toward the midline |
| ENSDART00000101037 | nhp2 | 0.003430859 | 0.465910387 | 0.391566586 | 0.240162003 | -0.008355829 | growth toward the midline |
| ENSDART00000101124 | rnaseka | 0.001851882 | 1.077486687 | 0.59165486 | 0.038549521 | -0.354475184 | growth toward the midline |
| ENSDART00000101143 | mhc1zea | 0.00820374 | 0.57323499 | 0.526487775 | 0.424884066 | 0.07882824 | growth toward the midline |
| ENSDART00000101319 | zgc:162396 | 0.01098705 | 0.344606585 | 0.405244483 | 0.227818995 | 0.060602938 | growth toward the midline |
| ENSDART00000102062 | timp2b | 0.000933245 | 0.538603085 | 0.084002119 | -0.039686845 | -0.313946909 | growth toward the midline |
| ENSDART00000102434 | ehhadh | 0.000776925 | 0.70922182 | 0.453174195 | 0.063745289 | -0.562849237 | growth toward the midline |
| ENSDART00000102459 | rbp2a | 0.000345998 | 3.492314469 | 2.902874566 | 1.90821276 | 1.340524369 | growth toward the midline |
| ENSDART00000102981 | col8a1a | 0.017381552 | 1.700212586 | 0.444548046 | 0.772515926 | 0.395334934 | growth toward the midline |
| ENSDART00000103076 | arl8bb | 0.002876227 | 0.705280948 | 0.466975375 | 0.584557674 | 0.096505859 | growth toward the midline |
| ENSDART00000103368 | rpl22 | 0.006345193 | 0.33350855 | 0.471741899 | 0.142215466 | -0.053336675 | growth toward the midline |
| ENSDART00000103491 | rbp7b | 0.000751152 | 2.051254968 | 1.285649693 | 0.558375641 | -0.074104197 | growth toward the midline |
| ENSDART00000103602 | lgals2a | 0.013513343 | 1.886236295 | 1.628346862 | 1.325259297 | 0.487532368 | growth toward the midline |
| ENSDART00000103660 | clcn7 | 0.002459431 | 0.412276024 | 0.162279088 | -0.074638152 | -0.273644307 | growth toward the midline |
| ENSDART00000104519 | stat3 | 0.004480141 | 0.6458865 | 0.459098575 | 0.438501049 | 0.201812376 | growth toward the midline |
| ENSDART00000104687 | pfn2 | 0.00311878 | 0.330558178 | 0.38737784 | 0.240819292 | -0.058450663 | growth toward the midline |
| ENSDART00000104835 | hps5 | 0.005167494 | 0.972059444 | 0.583168765 | 0.57443346 | -0.161192354 | growth toward the midline |
| ENSDART00000105174 | stm | 0.000739646 | 3.414215545 | 0.041843024 | -0.026881567 | -0.531574221 | growth toward the midline |
| ENSDART00000108581 | si:dkey-17m8.1 | 0.014069453 | 0.789679514 | 0.574820222 | 0.300394764 | -0.045841382 | growth toward the midline |
| ENSDART00000108995 | tex2 | 0.041898142 | 0.515345319 | 0.756490823 | 0.434164856 | 0.106684566 | growth toward the midline |
| ENSDART00000109065 | ccng1 | 0.001754036 | 0.498527324 | 0.814436551 | 0.266438219 | -0.054038693 | growth toward the midline |
| ENSDART00000109416 | smc2 | 0.00879202 | 1.321753383 | 1.108242465 | 0.857262832 | 0.308883411 | growth toward the midline |
| ENSDART00000109432 | cercam | 0.000991767 | 0.726258015 | 0.540687546 | 0.072936326 | -0.232601995 | growth toward the midline |
| ENSDART00000109464 | g0s2 | 8.33897E-05 | 2.993222235 | 1.499538045 | 0.588261277 | 0.527162957 | growth toward the midline |
| ENSDART00000109752 | serpinh1a | 0.000595605 | 0.578805227 | 0.380412306 | 0.080680913 | -0.569755212 | growth toward the midline |
| ENSDART00000109831 | BX088653.1 | 0.022613341 | 0.96716269 | 0.47589947 | 0.338658984 | -0.013815104 | growth toward the midline |
| ENSDART00000110270 | pwp2h | 0.011929276 | 0.562435944 | 0.598132459 | 0.373677258 | 0.062993981 | growth toward the midline |
| ENSDART00000110529 | bub1bb | 0.018115022 | 1.321570387 | 1.337769105 | 1.007493385 | 0.59867765 | growth toward the midline |
| ENSDART00000110866 | cln5 | 0.001248486 | 0.673394988 | 0.327334357 | 0.152385216 | -0.174612372 | growth toward the midline |
| ENSDART00000110935 | si:ch1073-59l16.1 | 0.005875165 | 1.356882908 | 0.676384004 | 0.249167454 | 0.069788375 | growth toward the midline |
| ENSDART00000111002 | si:ch211-74f19.2 | 0.000220872 | 0.463826256 | 0.294740419 | -0.053551592 | -0.452436201 | growth toward the midline |
| ENSDART00000111140 | rpl29 | 0.009681558 | 0.336664553 | 0.356246951 | 0.105865493 | -0.061220703 | growth toward the midline |
| ENSDART00000111321 | zgc:152830 | 0.001021091 | 0.702895787 | 0.346388089 | 0.27465484 | 0.003141495 | growth toward the midline |
| ENSDART00000111531 | epdl1 | 0.000962755 | 1.357192284 | 1.365774088 | 0.753975689 | 0.235060778 | growth toward the midline |
| ENSDART00000111688 | zgc:109934 | 0.000655075 | 0.61674072 | 0.212807687 | -0.164219088 | -0.482167701 | growth toward the midline |

| | | | | | | | |
|---|---|---|---|---|---|---|---|
| ENSDART00000111806 | cenpe | 0.002947105 | 1.006157891 | 0.692740621 | 0.695639634 | 0.175054418 | growth toward the midline |
| ENSDART00000112127 | dnase2b | 0.000645779 | 0.959684518 | 0.954914684 | 0.226944152 | -0.0439051 | growth toward the midline |
| ENSDART00000112155 | rrm2 | 0.028641638 | 1.131329801 | 1.215867796 | 0.761186289 | 0.339828406 | growth toward the midline |
| ENSDART00000112243 | crlf1a | 0.00448838 | 1.589627089 | 0.916105309 | 0.558277873 | -0.020827681 | growth toward the midline |
| ENSDART00000112470 | ccdc88b | 0.005891084 | 0.86030789 | 0.601824136 | 0.501988839 | -0.28013627 | growth toward the midline |
| ENSDART00000112653 | wdfy4 | 0.003815452 | 0.841684139 | 0.566530892 | 0.08064366 | -0.41519717 | growth toward the midline |
| ENSDART00000113093 | gba | 0.00060684 | 0.473554928 | 0.246228233 | -0.028007284 | -0.118288896 | growth toward the midline |
| ENSDART00000113259 | TARBP1 | 0.002686125 | 0.600840202 | 0.638996908 | 0.05884907 | -0.019773553 | growth toward the midline |
| ENSDART00000113405 | inpp5d | 0.005845428 | 0.955185108 | 0.470897855 | 0.050793817 | -0.341647614 | growth toward the midline |
| ENSDART00000113441 | mylk5 | 2.33589E-05 | 0.7890096 | 0.412195956 | -0.130414718 | -0.622741882 | growth toward the midline |
| ENSDART00000113574 | anln | 0.001248576 | 1.428520356 | 0.903846206 | 0.598652735 | 0.284032371 | growth toward the midline |
| ENSDART00000113589 | gpx4b | 0.002390041 | 0.633021831 | -0.10742897 | -0.312510701 | -0.292370818 | growth toward the midline |
| ENSDART00000113673 | scpp5 | 0.009220964 | 1.321767426 | 0.530715467 | 0.931995129 | 0.748355856 | growth toward the midline |
| ENSDART00000113789 | zgc:198419 | 0.012732777 | 1.388791656 | 0.463187117 | 0.161859067 | -0.160692863 | growth toward the midline |
| ENSDART00000114000 | zgc:101663 | 0.003860077 | 0.831482698 | 0.254442158 | 0.060051833 | 0.091105397 | growth toward the midline |
| ENSDART00000114023 | hspb9 | 0.063359837 | 2.157513115 | 1.55136025 | 0.742718373 | 1.461470885 | growth toward the midline |
| ENSDART00000114282 | rpsa | 0.000461528 | 0.494276409 | 0.531800028 | 0.113194341 | -0.004402994 | growth toward the midline |
| ENSDART00000114410 | wdr62 | 0.02432669 | 0.584622073 | 0.50588924 | 0.329377091 | 0.089655627 | growth toward the midline |
| ENSDART00000114964 | zgc:165573 | 0.000117076 | 0.691787708 | 0.943542464 | 0.634593348 | -0.039695615 | growth toward the midline |
| ENSDART00000115049 | mych | 6.48692E-06 | 2.666296724 | 2.42380403 | 1.76502541 | 0.819436082 | growth toward the midline |
| ENSDART00000115255 | mcf2l2 | 0.002771387 | 0.795464349 | 0.848343644 | 0.628078884 | 0.002922432 | growth toward the midline |
| ENSDART00000117636 | dre-mir-21-1 | 0.001938223 | 1.553092638 | 1.563315667 | 0.906878174 | 0.77260601 | growth toward the midline |
| ENSDART00000117680 | dre-mir-21-2 | 0.000180784 | 3.366495632 | 3.222859121 | 2.199417673 | 1.472943682 | growth toward the midline |
| ENSDART00000121998 | grwd1 | 0.006122188 | 0.591308312 | 0.087252775 | -0.045976929 | -0.167650618 | growth toward the midline |
| ENSDART00000122238 | tyr | 0.000536544 | 1.0676401 | 0.617097515 | 0.165901971 | -0.685666733 | growth toward the midline |
| ENSDART00000122288 | ctsla | 0.000770834 | 0.76727009 | 0.372627978 | 0.369896487 | -0.200637952 | growth toward the midline |
| ENSDART00000122321 | slc33a1 | 0.014050993 | 0.868257035 | 0.839034235 | 0.725290511 | 0.201428784 | growth toward the midline |
| ENSDART00000122407 | cdk1 | 0.003977971 | 1.677931921 | 1.534655447 | 0.824563588 | 0.384011667 | growth toward the midline |
| ENSDART00000122692 | CABZ01112317.1 | 0.003971559 | 0.655801165 | 0.399271741 | 0.354973485 | -0.117162708 | growth toward the midline |
| ENSDART00000122768 | CU984579.1 | 0.001277839 | 0.668419525 | 0.524624014 | 0.112892825 | -0.531459653 | growth toward the midline |
| ENSDART00000122898 | phf21b | 0.008936495 | 0.630758244 | -0.049111481 | 0.173874589 | 0.290793166 | growth toward the midline |
| ENSDART00000122945 | noc2l | 0.001509592 | 0.680401135 | 0.443128484 | 0.389864365 | 0.196289408 | growth toward the midline |
| ENSDART00000122953 | gnl3 | 0.004847659 | 0.534095037 | 0.261409226 | 0.267291075 | 0.063642911 | growth toward the midline |
| ENSDART00000123002 | nme4 | 0.004429413 | 3.640965481 | 3.577167408 | 3.045307609 | 2.652336979 | growth toward the midline |
| ENSDART00000123117 | ggh | 0.000844564 | 0.485870455 | 0.424307993 | 0.067357919 | -0.14640404 | growth toward the midline |
| ENSDART00000123473 | Metazoa_SRP | 0.011135896 | 0.899034068 | 1.683872421 | 0.662894374 | -0.48789514 | growth toward the midline |
| ENSDART00000123544 | tex2 | 0.010129858 | 0.532623113 | 0.735572961 | 0.212010938 | -0.135781649 | growth toward the midline |
| ENSDART00000124367 | si:cabz01074946.1 | 0.002461121 | 1.457343255 | 1.182560392 | 0.713827232 | 0.020183118 | growth toward the midline |
| ENSDART00000124663 | npc2 | 0.000902446 | 1.578129482 | 0.684715762 | 0.389649636 | 0.031719264 | growth toward the midline |
| ENSDART00000124833 | pdcd11 | 0.01245587 | 0.337926893 | 0.21951318 | 0.267537583 | 0.067942864 | growth toward the midline |
| ENSDART00000124868 | lpl | 0.000214183 | 0.342987533 | -0.032282276 | -0.168396519 | -1.276175937 | growth toward the midline |
| ENSDART00000125097 | si:dkey-126g1.7 | 0.00111498 | 0.825737488 | 0.388125468 | -0.092297047 | -0.324389582 | growth toward the midline |
| ENSDART00000125178 | eif5a | 0.003534272 | 0.322397945 | 0.394023524 | 0.161340886 | -0.011483865 | growth toward the midline |
| ENSDART00000125430 | pprc1 | 8.24104E-05 | 1.538620876 | 1.109491459 | 0.874206496 | 0.220876028 | growth toward the midline |
| ENSDART00000125753 | rpl22 | 0.00115719 | 0.364344172 | 0.362576681 | 0.16140506 | -0.121749389 | growth toward the midline |
| ENSDART00000126229 | rps7 | 0.004986552 | 0.396514179 | 0.373862291 | 0.136780142 | -0.080022773 | growth toward the midline |
| ENSDART00000126417 | bri3 | 0.002005251 | 0.500014173 | 0.414528928 | 0.11265957 | -0.17961569 | growth toward the midline |
| ENSDART00000126780 | rnaset2 | 0.000371877 | 0.561676509 | 0.243610345 | -0.050562112 | -0.093659569 | growth toward the midline |
| ENSDART00000126978 | acadl | 0.002156175 | 0.315070362 | 0.266918423 | -0.034640916 | -0.111553388 | growth toward the midline |
| ENSDART00000127277 | pum3 | 0.018496255 | 0.522067564 | 0.375606403 | 0.43721492 | 0.160017235 | growth toward the midline |
| ENSDART00000127350 | enpp1 | 0.000451785 | 1.121109181 | 0.817391101 | 0.578924138 | 0.065420517 | growth toward the midline |
| ENSDART00000127496 | loxl1 | 0.000342667 | 0.412088952 | 0.133411325 | -0.193765804 | -0.987510306 | growth toward the midline |
| ENSDART00000127753 | adam15 | 0.005335064 | 0.405103987 | 0.059018909 | -0.196195339 | -0.053874543 | growth toward the midline |
| ENSDART00000127982 | suox | 0.014022061 | 0.48894768 | 0.565232546 | 0.460591806 | 0.13079859 | growth toward the midline |
| ENSDART00000128049 | hunk | 0.016504069 | 0.851411376 | 0.241049411 | 0.130293149 | 0.977016396 | growth toward the midline |
| ENSDART00000128246 | polr1c | 0.013474932 | 0.52692078 | 0.422008081 | 0.272814737 | 0.11358433 | growth toward the midline |
| ENSDART00000128488 | e2f8 | 0.003382652 | 3.852661131 | -0.929008264 | 3.252279997 | 2.01178299 | growth toward the midline |
| ENSDART00000128550 | palb2 | 0.009615324 | 0.912196863 | 1.171643148 | 0.786370898 | 0.077481484 | growth toward the midline |
| ENSDART00000128613 | rps15a | 0.004826213 | 0.315521947 | 0.355667698 | 0.260468714 | -0.049823228 | growth toward the midline |
| ENSDART00000128743 | tmem214 | 0.02086388 | 0.301583498 | 0.153317199 | 0.10691428 | 0.053655055 | growth toward the midline |
| ENSDART00000128786 | pyya | 0.000403881 | 0.897714064 | 0.120108152 | -0.15783225 | -0.181132487 | growth toward the midline |
| ENSDART00000128917 | rps4x | 0.011474173 | 0.379768803 | 0.358465257 | 0.299276991 | 0.035010431 | growth toward the midline |
| ENSDART00000129155 | vmp1 | 1.77791E-05 | 3.054036254 | 2.972129495 | 2.202758828 | 1.232955319 | growth toward the midline |
| ENSDART00000129380 | ppp5c | 0.000402095 | 0.718884884 | 0.082309464 | 0.464958134 | 0.512508903 | growth toward the midline |
| ENSDART00000129617 | atp6ap1b | 0.000657061 | 0.602272356 | 0.200096368 | -0.075510461 | -0.466117163 | growth toward the midline |
| ENSDART00000129903 | mfsd10 | 0.000409421 | 0.819604016 | 0.72451413 | 0.409438046 | 0.004032617 | growth toward the midline |
| ENSDART00000129919 | TIFA | 0.001882493 | 0.580552682 | 0.759337474 | 0.369383589 | -0.101109572 | growth toward the midline |
| ENSDART00000129963 | rbp1 | 0.001296944 | 1.30707657 | 0.844804596 | -0.023243783 | -0.809227723 | growth toward the midline |
| ENSDART00000130131 | sult6b1 | 0.013901532 | 0.679202914 | 0.436031956 | 0.32900324 | -0.176606639 | growth toward the midline |
| ENSDART00000130297 | atg16l2 | 0.036864554 | 0.527142084 | 0.480414213 | 0.279348799 | 0.207971893 | growth toward the midline |
| ENSDART00000130494 | moxd1 | 0.001779714 | 0.536918295 | 0.137560293 | -0.344113602 | -0.82692019 | growth toward the midline |
| ENSDART00000130601 | CABZ01020840.1 | 0.002663982 | 1.211268037 | 0.754343565 | 0.351699193 | -0.297623866 | growth toward the midline |
| ENSDART00000130604 | ssuh2.4 | 0.003590782 | 0.995807077 | 0.517001744 | 0.13232339 | -0.007643053 | growth toward the midline |
| ENSDART00000130648 | scarb2c | 0.030497275 | 0.530119641 | 0.364624401 | 0.255215167 | -0.022903408 | growth toward the midline |
| ENSDART00000130911 | cmklr1 | 0.011725523 | 1.293061027 | 1.016453103 | 0.21085461 | 0.074156746 | growth toward the midline |

| | | | | | | | |
|---|---|---|---|---|---|---|---|
| ENSDART00000131217 | ippk | 0.009000055 | 0.31853123 | 0.426685761 | 0.198265799 | 0.012388457 | growth toward the midline |
| ENSDART00000131241 | CD53 | 0.022148005 | 1.613639378 | 1.480348694 | 1.150180939 | 0.900907437 | growth toward the midline |
| ENSDART00000131319 | ctps1a | 0.008283592 | 0.387589066 | 0.225454863 | 0.161786406 | -0.016402726 | growth toward the midline |
| ENSDART00000131400 | CR792453.3 | 0.001585773 | 2.47650448 | 0.829837833 | -0.069994818 | -0.405416724 | growth toward the midline |
| ENSDART00000131558 | zgc:103438 | 0.001133915 | 0.587891191 | 0.925990315 | 0.240143957 | -0.080675365 | growth toward the midline |
| ENSDART00000131661 | rps9 | 0.01002859 | 0.491211514 | 0.443239408 | 0.342502062 | -0.078353052 | growth toward the midline |
| ENSDART00000131694 | selenbp1 | 0.000844564 | 0.440307718 | 0.23934875 | 0.219921774 | -0.162883702 | growth toward the midline |
| ENSDART00000131724 | pld3 | 0.002957541 | 0.801498613 | 0.654598157 | 0.377873753 | 0.229417497 | growth toward the midline |
| ENSDART00000131802 | si:dkey-204f11.64 | 0.010173835 | 0.501807484 | 0.364460777 | 0.066146426 | -0.103454568 | growth toward the midline |
| ENSDART00000131966 | AL929520.1 | 0.003243686 | 1.864305398 | 1.102319083 | 0.530092783 | 0.456970098 | growth toward the midline |
| ENSDART00000132216 | paox1 | 0.005824405 | 0.706860126 | 0.311306251 | -0.007602467 | -0.279227462 | growth toward the midline |
| ENSDART00000132637 | rab34a | 0.004827977 | 0.251888735 | 0.316357959 | 0.030371007 | -1.217256649 | growth toward the midline |
| ENSDART00000132660 | polr3gla | 0.007082106 | 0.823813849 | 0.325249077 | 0.007723106 | -0.29164484 | growth toward the midline |
| ENSDART00000132742 | apoc1 | 0.007655029 | 0.900470905 | 0.457797087 | -0.066389631 | 0.150028097 | growth toward the midline |
| ENSDART00000132830 | stoml3b | 0.039723869 | 2.594772138 | 1.896323147 | 1.645470476 | 1.013599137 | growth toward the midline |
| ENSDART00000133109 | rps28 | 0.000937145 | 0.456862734 | 0.459372965 | 0.307002895 | 0.056264372 | growth toward the midline |
| ENSDART00000133167 | si:dkey-98f17.5 | 0.012693562 | 0.605842267 | 0.54718235 | 0.319480313 | 0.041690788 | growth toward the midline |
| ENSDART00000133304 | npm1a | 0.000878399 | 1.159038348 | 0.761096985 | 0.663684243 | 0.261874194 | growth toward the midline |
| ENSDART00000133375 | si:dkey-30c15.17 | 0.00181079 | 0.981163755 | 0.773615181 | 0.451221931 | -0.42308326 | growth toward the midline |
| ENSDART00000133404 | rps24 | 0.000717268 | 0.584478639 | 0.636503255 | 0.16891132 | -0.309462265 | growth toward the midline |
| ENSDART00000133512 | fosl1b | 3.29362E-06 | 2.958627889 | 2.652355797 | 1.322528527 | 0.313643003 | growth toward the midline |
| ENSDART00000133517 | si:dkey-228b2.5 | 0.032645202 | 0.6930874 | 0.410660128 | 0.386710557 | 0.222079502 | growth toward the midline |
| ENSDART00000133659 | ncapg | 0.042253127 | 0.759144886 | 0.927669912 | 0.449671959 | 0.278225274 | growth toward the midline |
| ENSDART00000133934 | rps9 | 0.033833708 | 0.75844303 | 0.709617891 | 0.420926612 | 0.393006212 | growth toward the midline |
| ENSDART00000133946 | si:dkey-147f3.8 | 0.003898333 | 1.477973714 | 1.681506394 | 1.170700784 | -0.068341199 | growth toward the midline |
| ENSDART00000134189 | npm1a | 0.015751923 | 1.523433306 | 1.012610127 | 1.305980875 | 0.439289484 | growth toward the midline |
| ENSDART00000134206 | hspb8 | 0.001438131 | 0.868767381 | 0.59923642 | 0.380053431 | 0.101288698 | growth toward the midline |
| ENSDART00000134404 | si:ch211-158d24.4 | 0.024003814 | 0.479480538 | 0.856462378 | 0.502566478 | -0.11398241 | growth toward the midline |
| ENSDART00000134819 | slc5a6a | 2.95145E-05 | 0.44313334 | -0.037657655 | -0.159985236 | -0.577627715 | growth toward the midline |
| ENSDART00000134941 | prc1b | 0.0242075 | 1.557351387 | 1.026365943 | 0.57100162 | 0.47167284 | growth toward the midline |
| ENSDART00000135032 | si:dkey-57a22.11 | 0.026714705 | 1.01281435 | 1.486077838 | 0.644934957 | 0.560278535 | growth toward the midline |
| ENSDART00000135238 | dok2 | 0.013959384 | 1.480525707 | 0.841647962 | 0.697333529 | 0.138703296 | growth toward the midline |
| ENSDART00000135472 | zgc:86764 | 0.036021217 | 1.170468948 | 0.97326651 | 0.792803713 | 0.492808604 | growth toward the midline |
| ENSDART00000135517 | cenpf | 0.003777754 | 1.006357007 | 0.930294184 | 0.598168756 | 0.318387864 | growth toward the midline |
| ENSDART00000135719 | si:ch211-171h4.6 | 0.033435193 | 1.187005549 | 0.373648016 | 0.271137881 | -0.125827965 | growth toward the midline |
| ENSDART00000135888 | rrp9 | 0.002433479 | 0.649352382 | 0.396806051 | 0.419459571 | 0.067410678 | growth toward the midline |
| ENSDART00000136382 | gadd45ba | 0.000960205 | 1.037707985 | 1.186135766 | 0.645465362 | 0.508629 | growth toward the midline |
| ENSDART00000136383 | cd99 | 0.001367004 | 0.67248236 | 1.035021667 | 0.666974643 | -0.164650894 | growth toward the midline |
| ENSDART00000136543 | tnni4b.1 | 0.007654107 | 1.504503498 | 0.982879899 | 0.857598266 | -0.145526765 | growth toward the midline |
| ENSDART00000136722 | cdkn1a | 0.000598084 | 1.083504495 | 0.97819224 | 0.258587523 | 0.060123909 | growth toward the midline |
| ENSDART00000136779 | entpd5b | 0.002650976 | 0.819562392 | 0.580844217 | 0.174817773 | -0.04896424 | growth toward the midline |
| ENSDART00000136835 | gpr84 | 0.003982134 | 0.70622544 | 0.183343809 | -0.153052934 | -0.427967202 | growth toward the midline |
| ENSDART00000136865 | cfp | 0.000230274 | 0.329536618 | 0.225136408 | -0.193120136 | -0.891420593 | growth toward the midline |
| ENSDART00000137019 | cmklr1 | 7.26498E-05 | 1.293423342 | 0.826870113 | 0.435422926 | 0.179038361 | growth toward the midline |
| ENSDART00000137373 | si:ch211-266i6.3 | 0.000114847 | 1.973293451 | 1.253222026 | 0.772689981 | 0.242956342 | growth toward the midline |
| ENSDART00000137858 | alpk2 | 0.024455662 | 0.667374473 | 0.189664169 | 0.047615444 | -0.048884335 | growth toward the midline |
| ENSDART00000137973 | grna | 0.005201666 | 1.322360375 | 0.965948715 | 0.567189356 | -0.062803655 | growth toward the midline |
| ENSDART00000137984 | SBNO2 (1 of many) | 0.001847086 | 0.66540874 | 0.705510404 | 0.683617676 | -0.079102756 | growth toward the midline |
| ENSDART00000138527 | rab20 | 0.009820514 | 0.632157851 | 0.410164386 | 0.278506263 | 0.009483031 | growth toward the midline |
| ENSDART00000138747 | urb2 | 0.019322869 | 0.51682098 | 0.426109616 | 0.379169845 | 0.148916896 | growth toward the midline |
| ENSDART00000138996 | si:ch211-214p13.9 | 0.003686476 | 1.396990228 | 1.323446174 | 0.865221946 | -0.306555509 | growth toward the midline |
| ENSDART00000139087 | lgmn | 2.59942E-05 | 0.782730519 | 0.338051823 | 0.153128345 | -0.09953272 | growth toward the midline |
| ENSDART00000139124 | ckba | 0.004373845 | 1.776728636 | 1.213212461 | 1.201789834 | 0.178980804 | growth toward the midline |
| ENSDART00000139659 | si:dkey-222f2.7 | 0.007791135 | 1.108120911 | 0.818052369 | 0.396214124 | -0.100365974 | growth toward the midline |
| ENSDART00000139839 | rbmx | 0.034811028 | 0.438924108 | 0.178102249 | 0.089667681 | 0.22023097 | growth toward the midline |
| ENSDART00000140086 | SUN2 | 0.001030626 | 0.64389802 | 0.258658486 | -0.305248612 | -0.209793481 | growth toward the midline |
| ENSDART00000140523 | mrc1b | 0.000638552 | 0.693910087 | 0.60918831 | 0.270261416 | -0.09789336 | growth toward the midline |
| ENSDART00000140667 | spna2 | 0.034218044 | 0.558956985 | 0.663728247 | 0.276626138 | 0.347556663 | growth toward the midline |
| ENSDART00000140776 | CR391940.1 | 0.013366808 | 0.57109507 | 0.234979352 | 0.004616304 | -0.046528801 | growth toward the midline |
| ENSDART00000140788 | gbp1 | 0.014906425 | 3.129821614 | 2.838243862 | 2.757243294 | 2.490411238 | growth toward the midline |
| ENSDART00000140874 | seph | 0.005866285 | 0.619876107 | 0.651122471 | 0.195858296 | -0.082542729 | growth toward the midline |
| ENSDART00000140920 | prom1b | 0.039913243 | 0.88825196 | 0.404698565 | 0.393910982 | 0.372363663 | growth toward the midline |
| ENSDART00000140970 | si:ch73-52p7.1 | 0.011521596 | 1.046367332 | 0.798731839 | 0.265928239 | -0.108652338 | growth toward the midline |
| ENSDART00000141142 | mvp | 0.001130894 | 1.187603651 | 1.369710798 | 0.919014152 | 0.045597494 | growth toward the midline |
| ENSDART00000141188 | st3gal3b | 0.000470359 | 0.577420785 | 0.203550656 | -0.28232291 | -0.213031688 | growth toward the midline |
| ENSDART00000141657 | anxa4 | 0.003268346 | 0.870599992 | 0.703638401 | 0.563849009 | -0.254549515 | growth toward the midline |
| ENSDART00000141676 | rpl12 | 0.0451272 | 0.410901536 | 0.227715625 | 0.271640237 | 0.155083239 | growth toward the midline |
| ENSDART00000141714 | nupr1 | 0.022150836 | 1.321058122 | 1.103287011 | 0.703763985 | 0.083130869 | growth toward the midline |
| ENSDART00000142088 | si:ch211-218c6.8 | 0.009300169 | 1.058750612 | 0.593589729 | 0.534639469 | 0.071525268 | growth toward the midline |
| ENSDART00000142104 | cep162 | 0.025259321 | 0.409390298 | 0.286359079 | 0.288581786 | 0.16808044 | growth toward the midline |
| ENSDART00000142389 | cracr2ab | 0.002199978 | 1.038814867 | 1.254276576 | 0.507748466 | -0.396339594 | growth toward the midline |
| ENSDART00000142422 | tgm2l | 0.026925307 | 1.901086414 | 0.665689502 | 0.275506536 | 0.14002471 | growth toward the midline |
| ENSDART00000142467 | si:ch211-199g17.2 | 0.017327178 | 2.796012298 | 2.177853709 | 2.464842477 | 1.654064057 | growth toward the midline |
| ENSDART00000143066 | ube2e1 | 0.020455943 | 0.330371306 | 0.357877825 | 0.149635317 | 0.181174607 | growth toward the midline |
| ENSDART00000143291 | RPL41 | 0.002755913 | 0.451636798 | 0.464258672 | 0.193261311 | 0.034762109 | growth toward the midline |

| | | | | | | | |
|---|---|---|---|---|---|---|---|
| ENSDART00000143587 | ern2 | 0.02452124 | 0.504450294 | 0.214096186 | 0.104332211 | 0.050090305 | growth toward the midline |
| ENSDART00000143618 | birc7 | 0.03238504 | 0.867487648 | 0.469428911 | 0.548081783 | 0.007263478 | growth toward the midline |
| ENSDART00000143766 | tbc1d4 | 0.027561798 | 0.551251852 | 0.35059591 | 0.424872318 | 0.224396247 | growth toward the midline |
| ENSDART00000143846 | si:ch211-198c19.3 | 0.000489924 | 0.524345776 | 0.185185856 | -0.297806816 | -0.062260629 | growth toward the midline |
| ENSDART00000143875 | BX323074.2 | 0.003519185 | 0.32505808 | -0.029313176 | -0.099305279 | -0.178101813 | growth toward the midline |
| ENSDART00000143928 | rwdd1 | 0.018316862 | 0.331616231 | 0.265135658 | 0.236169655 | 0.064492205 | growth toward the midline |
| ENSDART00000144048 | cecr1b | 0.000723654 | 1.174703015 | 1.015529557 | 0.446419815 | -0.23634795 | growth toward the midline |
| ENSDART00000144139 | elavl4 | 0.003742624 | 1.756586356 | 1.578077782 | 0.938796608 | 0.66956976 | growth toward the midline |
| ENSDART00000144711 | tmem154 | 0.034302368 | 0.48356725 | 0.317274745 | 0.352966647 | 0.032904309 | growth toward the midline |
| ENSDART00000144886 | dicp1.1 | 0.024485347 | 1.748045166 | 0.589930553 | 0.172712572 | 0.178536852 | growth toward the midline |
| ENSDART00000144891 | gpnmb | 0.000996442 | 2.526942309 | 2.443053131 | 2.053909603 | -0.779431544 | growth toward the midline |
| ENSDART00000145111 | parp4 | 0.00078525 | 1.113854381 | 0.990369473 | 0.690636927 | 0.186507426 | growth toward the midline |
| ENSDART00000145198 | steap4 | 0.002341904 | 0.551058744 | 0.178525002 | -0.419104103 | -0.970957834 | growth toward the midline |
| ENSDART00000145275 | si:dkey-147f3.8 | 0.000993187 | 1.443349889 | 1.631277345 | 1.233548809 | -0.107779207 | growth toward the midline |
| ENSDART00000145330 | apoc1 | 0.011155618 | 1.228570608 | 0.722853476 | 0.218571174 | 0.363892889 | growth toward the midline |
| ENSDART00000145331 | zgc:92066 | 0.060501903 | 3.539715592 | 2.15686966 | 1.135379743 | 1.013519624 | growth toward the midline |
| ENSDART00000145377 | rpl38 | 0.001017556 | 0.331388676 | 0.460944412 | 0.086514771 | -0.0487643 | growth toward the midline |
| ENSDART00000145562 | robo4 | 0.000595605 | 1.739515239 | 1.001047952 | 0.857159879 | -0.094789579 | growth toward the midline |
| ENSDART00000145681 | atp6v1h | 0.000322013 | 0.62737841 | 0.200274483 | 0.109012982 | -0.483227692 | growth toward the midline |
| ENSDART00000145933 | si:rp71-68n21.9 | 0.016453138 | 0.834432396 | 0.464946482 | 0.375058394 | -0.112544064 | growth toward the midline |
| ENSDART00000145983 | cdkn1a | 0.003897339 | 1.243824395 | 0.764194636 | 0.498342294 | 0.064303597 | growth toward the midline |
| ENSDART00000146101 | mcoln1a | 0.002433479 | 0.490011182 | 0.372261102 | 0.240356077 | 0.080050098 | growth toward the midline |
| ENSDART00000146564 | si:ch211-198c19.3 | 0.000658214 | 1.407263699 | 0.462763747 | -0.033079511 | -0.600140762 | growth toward the midline |
| ENSDART00000147265 | rpl8 | 0.005893002 | 1.326651746 | 0.800643986 | 0.962132043 | 0.631592879 | growth toward the midline |
| ENSDART00000147275 | plat | 0.013807548 | 0.870075522 | 0.84761039 | 0.802236758 | 0.134643658 | growth toward the midline |
| ENSDART00000147341 | tubb4b | 0.0213019 | 0.566193096 | 0.992023395 | -0.136369873 | 0.130190834 | growth toward the midline |
| ENSDART00000147343 | dnase2 | 0.000313682 | 0.744692712 | 0.455061161 | 0.199303414 | -0.152365099 | growth toward the midline |
| ENSDART00000147516 | si:ch73-343l4.8 | 0.002706926 | 0.629933021 | 0.124466643 | -0.147547456 | -2.286690979 | growth toward the midline |
| ENSDART00000147527 | si:dkey-9i23.14 | 0.001146813 | 2.675666068 | 2.038792867 | 2.055161538 | 1.544748993 | growth toward the midline |
| ENSDART00000147617 | bcl2l1 | 0.002976628 | 0.692333613 | 0.690280735 | 0.392732217 | 0.224132994 | growth toward the midline |
| ENSDART00000148125 | fam21c | 0.012864775 | 0.420817307 | 0.137516364 | 0.034339244 | -0.014273742 | growth toward the midline |
| ENSDART00000148327 | adamtsl7 | 0.000283184 | 2.218956465 | 1.395778664 | 0.846103004 | 0.026559812 | growth toward the midline |
| ENSDART00000148398 | MRAS (1 of many) | 0.000781831 | 1.275583164 | 1.112018386 | 0.557077896 | 0.217843634 | growth toward the midline |
| ENSDART00000148582 | smpd1 | 0.002176351 | 0.404585024 | 0.130204031 | 0.022544887 | -0.113079466 | growth toward the midline |
| ENSDART00000148648 | rps25 | 0.003347171 | 0.412929704 | 0.39629643 | 0.197445263 | 0.016589956 | growth toward the midline |
| ENSDART00000148737 | btk | 0.046077182 | 1.027482823 | 0.714086499 | 0.58431037 | 0.237640489 | growth toward the midline |
| ENSDART00000148766 | dpf2 | 0.01541945 | 1.058462293 | 0.640887505 | 0.623810049 | 0.633616815 | growth toward the midline |
| ENSDART00000148845 | BX649485.4 | 0.007338257 | 2.24411597 | 1.39694109 | 1.224587369 | -0.358910932 | growth toward the midline |
| ENSDART00000149019 | itpr1a | 0.025939466 | 0.453945089 | 0.40663332 | 0.191730219 | 0.048560775 | growth toward the midline |
| ENSDART00000149043 | CR855300.1 | 0.019593044 | 0.458598352 | 0.519077258 | 0.374073731 | 0.109291075 | growth toward the midline |
| ENSDART00000149169 | ssr2 | 0.000562043 | 0.474946343 | 0.309006192 | 0.206925318 | -0.029893306 | growth toward the midline |
| ENSDART00000149431 | sema4aa | 0.000470359 | 1.058121853 | 0.676658919 | 0.239884836 | -0.1730827 | growth toward the midline |
| ENSDART00000149468 | rps10 | 0.017275314 | 2.202930165 | 1.872426465 | 0.292067044 | 0.38759709 | growth toward the midline |
| ENSDART00000149478 | galns | 0.01015025 | 1.09483472 | 0.726913129 | 0.227424057 | 0.052897193 | growth toward the midline |
| ENSDART00000149486 | def6b | 0.012263789 | 4.188632994 | 4.544630212 | 4.117698226 | 0.985190768 | growth toward the midline |
| ENSDART00000149600 | tpp1 | 4.91848E-05 | 0.822885392 | 0.356076016 | 0.047488091 | -0.289304028 | growth toward the midline |
| ENSDART00000149711 | slc4a2b | 0.000605314 | 0.638486491 | 0.276788243 | -0.283420926 | -0.322786037 | growth toward the midline |
| ENSDART00000149760 | triobpa | 0.028230031 | 0.630702727 | 0.141628128 | 0.231366625 | 0.068551496 | growth toward the midline |
| ENSDART00000149882 | si:ch211-117c9.1 | 0.076090072 | 1.22256875 | 0.270055012 | 0.422501128 | 0.414018825 | growth toward the midline |
| ENSDART00000149951 | BX666064.1 | 0.012467303 | 1.493495344 | 0.978793976 | 0.565382246 | -0.392061768 | growth toward the midline |
| ENSDART00000150088 | BX649485.2 | 0.007435419 | 1.751466958 | 1.112873174 | 0.6867673 | -0.722119651 | growth toward the midline |
| ENSDART00000150184 | gpatch4 | 0.006624621 | 0.624444079 | 0.471783034 | 0.550844363 | 0.152733798 | growth toward the midline |
| ENSDART00000150201 | MRAS (1 of many) | 0.001127896 | 1.00558208 | 0.977546999 | 0.676765678 | 0.026394977 | growth toward the midline |
| ENSDART00000150389 | tmem168a | 0.066196151 | 0.525344796 | 0.286469995 | 0.221730459 | 0.080011152 | growth toward the midline |
| ENSDART00000150484 | BX465862.4 | 0.000615224 | 1.071580051 | 0.886976814 | 0.477743025 | 0.134467836 | growth toward the midline |
| ENSDART00000150919 | si:ch211-191i18.2 | 0.004498556 | 1.045780222 | 1.362401557 | 0.586788909 | 0.260330898 | growth toward the midline |
| ENSDART00000150934 | alg6 | 0.010983086 | 0.502789869 | 0.534691837 | 0.441461289 | 0.13055149 | growth toward the midline |
| ENSDART00000150940 | si:ch211-193k19.2 | 0.024493238 | 1.599024227 | 0.310661889 | 0.753508784 | 0.054490277 | growth toward the midline |
| ENSDART00000151390 | si:ch211-195b13.1 | 0.006097198 | 0.447846827 | 0.275025282 | 0.031728831 | 0.543393224 | growth toward the midline |
| ENSDART00000151393 | zak | 0.000392364 | 0.559981256 | 0.606971645 | 0.291742344 | 0.078029866 | growth toward the midline |
| ENSDART00000151805 | vamp4 | 0.008628374 | 0.722926228 | 0.32988105 | 0.08556674 | 0.068052265 | growth toward the midline |
| ENSDART00000152122 | si:ch211-176g6.2 | 0.006040855 | 1.234754318 | 1.008584835 | 0.458934208 | -0.407921233 | growth toward the midline |
| ENSDART00000152299 | si:dkey-25o16.4 | 0.014178743 | 1.175283487 | 1.143366275 | 0.842863117 | 0.368204805 | growth toward the midline |
| ENSDART00000152378 | tgif1 | 0.003122803 | 0.194631245 | 0.419803811 | 0.14379228 | -0.205607529 | growth toward the midline |
| ENSDART00000152432 | lratb.2 | 0.001938223 | 0.819163408 | 1.604350772 | -0.35089707 | -0.825333131 | growth toward the midline |
| ENSDART00000152699 | si:ch211-286b5.9 | 0.004853289 | 1.056007406 | 0.325301277 | 0.188178466 | -0.341462888 | growth toward the midline |
| ENSDART00000152766 | CR354435.4 | 0.039457626 | 0.814174745 | 0.989990518 | 0.663039704 | 0.339445782 | growth toward the midline |
| ENSDART00000152788 | mki67 | 0.018439771 | 1.022403776 | 0.934711217 | 0.634468641 | 0.246495311 | growth toward the midline |
| ENSDART00000152892 | slc16a8 | 0.000894529 | 2.182950408 | 0.789162193 | -0.768133939 | -1.779969995 | growth toward the midline |
| ENSDART00000153248 | chadlb | 0.026040756 | 1.217222776 | 0.847984599 | 0.639882396 | -0.038384888 | growth toward the midline |
| ENSDART00000153654 | piezo2a.2 | 0.004108486 | 0.718362747 | 0.545453706 | 0.495090248 | 0.182038228 | growth toward the midline |
| ENSDART00000153731 | si:dkey-79d12.5 | 2.59942E-05 | 1.619937595 | 1.685958359 | 0.880018827 | 0.052586274 | growth toward the midline |
| ENSDART00000153847 | si:dkey-7i4.24 | 0.017660302 | 4.195698891 | 4.052966176 | 3.712547417 | 1.148430087 | growth toward the midline |
| ENSDART00000153950 | si:dkey-106n21.1 | 0.013634701 | 1.634202521 | 0.388901976 | 0.409369194 | 0.514423363 | growth toward the midline |
| ENSDART00000153990 | CR848791.2 | 0.022463379 | 3.062868799 | 2.217074852 | 2.048805323 | 1.916392428 | growth toward the midline |

| | | | | | | | |
|---|---|---|---|---|---|---|---|
| ENSDART00000154066 | isg20 | 0.006815804 | 0.340483696 | 0.28200181 | 0.116846361 | -0.121466791 | growth toward the midline |
| ENSDART00000154295 | RPS17 (1 of many) | 0.002786216 | 0.397556507 | 0.446477566 | 0.193480989 | 0.136840294 | growth toward the midline |
| ENSDART00000154324 | CR848791.3 | 0.00017041 | 3.13610848 | 2.257927159 | 1.961382093 | 1.531269093 | growth toward the midline |
| ENSDART00000154460 | FP243382.1 | 0.01179775 | 2.715231463 | 2.420271029 | 2.173535574 | 1.735635064 | growth toward the midline |
| ENSDART00000154638 | kif26ab | 0.032809386 | 2.382609513 | 1.668373417 | 1.704066313 | 1.906262158 | growth toward the midline |
| ENSDART00000154726 | MYADM (1 of many) | 0.004207729 | 1.393641215 | 1.121324137 | 1.000193747 | 0.38217935 | growth toward the midline |
| ENSDART00000154801 | itgae.2 | 0.002196508 | 0.922252153 | 0.655667188 | 0.379341087 | -0.144710616 | growth toward the midline |
| ENSDART00000154809 | CU467861.1 | 0.013110232 | 0.653813027 | 0.40784225 | 0.192026805 | 0.151191997 | growth toward the midline |
| ENSDART00000155268 | pik3cb | 0.012263789 | 0.672881324 | 0.55380807 | 0.246761254 | 0.040522215 | growth toward the midline |
| ENSDART00000155273 | FP102016.1 | 0.003003196 | 0.672121642 | 0.238521805 | 0.194526989 | 0.041260629 | growth toward the midline |
| ENSDART00000155427 | reep6 | 0.001233735 | 0.890805905 | 0.284631883 | 0.369468446 | 0.541597773 | growth toward the midline |
| ENSDART00000155519 | si:ch211-139d20.3 | 0.030271642 | 0.778333948 | 0.572606648 | 0.403486279 | 0.486914049 | growth toward the midline |
| ENSDART00000155553 | CR931780.1 | 0.003710792 | 1.845884645 | 1.39773807 | 0.761518659 | 0.588325484 | growth toward the midline |
| ENSDART00000155755 | CU469362.1 | 0.001684124 | 0.467928162 | 0.778645014 | 0.342312822 | -0.038725012 | growth toward the midline |
| ENSDART00000155844 | nr1d4b | 0.00277366 | 0.850133899 | 0.016728696 | -0.171313869 | 0.165802614 | growth toward the midline |
| ENSDART00000155864 | CU929070.1 | 0.002360359 | 1.274116455 | 1.096698631 | 0.870265354 | 0.298589514 | growth toward the midline |
| ENSDART00000156002 | tmem206 | 0.007346626 | 0.793441462 | 0.286165316 | 0.167291274 | -0.130249565 | growth toward the midline |
| ENSDART00000156428 | hephl1a | 0.008866484 | 0.427057323 | 0.215282086 | 0.277986668 | -0.033310258 | growth toward the midline |
| ENSDART00000156499 | prkcdb | 0.006467002 | 0.617876805 | 0.366145269 | 0.057590279 | -0.230105501 | growth toward the midline |
| ENSDART00000156575 | dicp3.3 | 0.011810205 | 1.874714519 | 0.882148407 | 0.706359774 | -0.079481712 | growth toward the midline |
| ENSDART00000157190 | ostn | 0.030844025 | 0.371505716 | 0.628806712 | 0.229636498 | 0.167623814 | growth toward the midline |
| ENSDART00000157487 | tfec | 0.001790465 | 0.534318298 | 0.423012313 | 0.205933849 | -0.313187686 | growth toward the midline |
| ENSDART00000157636 | Metazoa_SRP | 4.91295E-05 | 2.626880551 | 2.386640401 | 1.529298522 | 0.72995056 | growth toward the midline |
| ENSDART00000157899 | dad1 | 0.009401721 | 0.443099375 | 0.390242441 | 0.192203328 | -0.014224626 | growth toward the midline |
| ENSDART00000158212 | zgc:92161 | 0.004952764 | 0.66498591 | 0.968065208 | 0.600651955 | -0.278084019 | growth toward the midline |
| ENSDART00000158581 | spna2 | 0.00601663 | 0.480311119 | 0.377153274 | 0.267336877 | -0.085977531 | growth toward the midline |
| ENSDART00000158733 | si:dkey-33i11.1 | 0.004853289 | 1.813729642 | 1.289849367 | 0.876112899 | 0.279282769 | growth toward the midline |
| ENSDART00000158808 | cnn3a | 6.00847E-05 | 0.586268263 | 0.320248819 | -0.106889335 | -0.483977637 | growth toward the midline |
| ENSDART00000159246 | ctsa | 0.004928313 | 0.554249571 | 0.498653237 | 0.219459997 | 0.055975716 | growth toward the midline |
| ENSDART00000159330 | pcyox1 | 0.007509621 | 0.406833689 | 0.21789945 | 0.044242143 | -0.158070725 | growth toward the midline |
| ENSDART00000159433 | BX571691.1 | 0.030614624 | 2.919338685 | 1.742268402 | 2.797274384 | 2.742997875 | growth toward the midline |
| ENSDART00000159434 | tmem254 | 0.019015879 | 0.529951039 | 0.26685766 | 0.074245289 | -0.042910999 | growth toward the midline |
| ENSDART00000159534 | leng9 | 0.002069136 | 0.34368633 | 0.401471128 | 0.132926162 | 0.109702136 | growth toward the midline |
| ENSDART00000159629 | mcm2 | 0.044268959 | 0.995796878 | 0.693727511 | 0.639304212 | 0.157223466 | growth toward the midline |
| ENSDART00000159716 | si:ch73-236c18.6 | 0.022944365 | 0.962463693 | 0.259951761 | 0.066014607 | 0.162933451 | growth toward the midline |
| ENSDART00000160000 | spdl1 | 0.01662249 | 1.145158953 | 1.158182543 | 1.166185329 | 0.390123322 | growth toward the midline |
| ENSDART00000160116 | racgap1 | 0.01613952 | 0.647984241 | 0.386764067 | 0.354192772 | 0.100327502 | growth toward the midline |
| ENSDART00000160201 | tnfrsf11b | 4.91848E-05 | 2.52634883 | 1.22900254 | 0.873090756 | -0.039240801 | growth toward the midline |
| ENSDART00000160245 | si:ch73-236c18.7 | 0.03675533 | 2.124440879 | 1.318007353 | 1.287829039 | 0.437471879 | growth toward the midline |
| ENSDART00000160297 | vcana | 0.04873911 | 1.601498468 | 0.665214133 | 0.454293537 | 0.486706931 | growth toward the midline |
| ENSDART00000160331 | atp6v0e1 | 0.000593122 | 0.536560194 | 0.109862681 | -0.220128717 | -0.336777163 | growth toward the midline |
| ENSDART00000160503 | slc15a4 | 0.005895019 | 0.534058917 | 0.262741482 | 0.05558204 | -0.165911292 | growth toward the midline |
| ENSDART00000160536 | gnsb | 6.00847E-05 | 1.380175579 | 0.91424754 | 0.488823056 | 0.00723105 | growth toward the midline |
| ENSDART00000160582 | creb5a | 4.11188E-05 | 2.028867303 | 1.691967909 | 1.27021295 | 0.836790625 | growth toward the midline |
| ENSDART00000160902 | naga | 0.00155515 | 0.641123507 | 0.233762511 | 0.291444137 | -0.017668678 | growth toward the midline |
| ENSDART00000161040 | U1 | 0.063982053 | 0.811168663 | 0.450901632 | 0.418592021 | 0.580227266 | growth toward the midline |
| ENSDART00000161063 | heatr1 | 0.020490278 | 0.383217268 | 0.256645773 | 0.173900631 | 0.113179939 | growth toward the midline |
| ENSDART00000161216 | dync1i2a | 0.016102005 | 1.005522301 | 0.520541958 | 0.468549497 | -0.022454144 | growth toward the midline |
| ENSDART00000161240 | zgc:92161 | 0.014596701 | 0.584076871 | 0.663364233 | 0.486030672 | 0.10810748 | growth toward the midline |
| ENSDART00000161322 | slc29a1a | 0.003016249 | 0.535023852 | 0.636573744 | 0.365586045 | -0.115892928 | growth toward the midline |
| ENSDART00000161421 | zgc:194627 | 0.005106608 | 1.998498644 | 1.298544665 | 0.773438314 | 0.249109157 | growth toward the midline |
| ENSDART00000161906 | vwa11 | 0.001097466 | 1.529478256 | 0.755786632 | 0.896680083 | 0.546980022 | growth toward the midline |
| ENSDART00000161967 | ctsc | 0.000657061 | 1.279775059 | 0.583635254 | 0.329059668 | -0.157215155 | growth toward the midline |
| ENSDART00000162097 | CABZ01044157.1 | 0.011403904 | 1.031550352 | 0.670727071 | 0.212982548 | -0.249292553 | growth toward the midline |
| ENSDART00000162282 | dicp3.1 | 0.000996442 | 0.839763954 | 0.379787637 | 0.004488715 | -0.034973807 | growth toward the midline |
| ENSDART00000162296 | cfp | 0.000230274 | 0.329536618 | 0.225136408 | -0.193120136 | -0.891420593 | growth toward the midline |
| ENSDART00000162472 | VWA5A (1 of many) | 0.004190237 | 0.543940251 | 0.262980075 | 0.410699646 | -0.136546959 | growth toward the midline |
| ENSDART00000162482 | zgc:152863 | 0.019421916 | 0.539805357 | 0.660295595 | 0.432850415 | 0.095615261 | growth toward the midline |
| ENSDART00000162668 | cremb | 4.56695E-06 | 4.310209888 | 4.052771 | 3.333577791 | 2.240090056 | growth toward the midline |
| ENSDART00000162799 | crb3a | 0.012813136 | 0.362169262 | 0.040586113 | -0.086039525 | -0.014272311 | growth toward the midline |
| ENSDART00000162958 | si:ch211-260e23.9 | 0.01863352 | 0.345171245 | 0.431307401 | 0.079575678 | 0.236719343 | growth toward the midline |
| ENSDART00000163018 | bet1l | 0.049630983 | 0.723011674 | 0.444641169 | 0.411413339 | 0.530095318 | growth toward the midline |
| ENSDART00000163391 | si:ch211-149k12.3 | 0.022976426 | 0.507081525 | 0.32718269 | 0.316220639 | 0.200966933 | growth toward the midline |
| ENSDART00000163398 | pla2g15 | 0.007621679 | 0.617752366 | 0.443695085 | 0.159097428 | 0.08811481 | growth toward the midline |
| ENSDART00000163407 | slc20a1a | 0.005041829 | 0.970510209 | 0.576996952 | 0.406775549 | -0.419444699 | growth toward the midline |
| ENSDART00000163668 | si:ch1073-392o20.2 | 0.002227565 | 0.543875349 | 0.345011244 | 0.082952674 | -0.090250505 | growth toward the midline |
| ENSDART00000163724 | SLA (1 of many) | 0.041839159 | 1.207342807 | 0.416124608 | 0.383103847 | 0.193096581 | growth toward the midline |
| ENSDART00000163794 | wt1b | 0.02934736 | 0.454982734 | 0.330694147 | 0.317167947 | 0.300420298 | growth toward the midline |
| ENSDART00000164113 | cpeb1a | 0.0004696 | 1.336706075 | 1.190402311 | 1.006023313 | 0.300918459 | growth toward the midline |
| ENSDART00000164161 | osbpl1a | 0.000360396 | 0.351337908 | 0.311134875 | -0.016374257 | -0.257205353 | growth toward the midline |
| ENSDART00000164198 | si:dkey-102m7.3 | 0.011741843 | 0.949739085 | 1.411170301 | 0.641678924 | 0.378323778 | growth toward the midline |
| ENSDART00000164305 | dicp1.1 | 0.001163813 | 0.970381914 | 0.35730766 | 0.127672979 | 0.007978514 | growth toward the midline |
| ENSDART00000164440 | si:ch211-195b11.3 | 0.017379805 | 0.887240489 | 0.373309939 | 0.100891486 | -0.191092546 | growth toward the midline |
| ENSDART00000164484 | dtnba | 0.000293873 | 0.785988532 | 0.671033087 | 0.445356623 | 0.140878127 | growth toward the midline |
| ENSDART00000165186 | si:dkey-33i11.9 | 0.01609527 | 1.335261857 | 1.027980768 | 0.465372707 | 0.114592431 | growth toward the midline |

| | | | | | | | |
|---|---|---|---|---|---|---|---|
| ENSDART00000165491 | BX294383.1 | 0.040285619 | 0.352047768 | 0.065820831 | 0.027639622 | 0.019046581 | growth toward the midline |
| ENSDART00000165499 | rps17 | 0.001024575 | 0.504096498 | 0.637791027 | 0.385574489 | 0.024446186 | growth toward the midline |
| ENSDART00000165735 | mdm4 | 0.009204947 | 0.677008086 | 0.457427602 | 0.365768719 | 0.56900659 | growth toward the midline |
| ENSDART00000165932 | sik1 | 0.023071688 | 1.156253383 | 0.676084919 | 0.687880039 | 1.28219737 | growth toward the midline |
| ENSDART00000165938 | CU466240.1 | 0.062849789 | 0.428025361 | 0.165679287 | 0.139754855 | 0.298251046 | growth toward the midline |
| ENSDART00000166152 | DAB2 (1 of many) | 0.00878097 | 0.967097608 | 0.419929333 | 0.22593558 | -0.403858057 | growth toward the midline |
| ENSDART00000166209 | wu:fb44b02 | 0.001191783 | 0.715207055 | 0.457109376 | 0.301906263 | -0.147536436 | growth toward the midline |
| ENSDART00000166388 | il6st | 0.008110834 | 0.326944598 | 0.248050944 | 0.233084281 | -0.066517896 | growth toward the midline |
| ENSDART00000166395 | fcer1gl | 0.000698893 | 0.672549379 | 0.370716598 | -0.127334281 | -0.409142814 | growth toward the midline |
| ENSDART00000166496 | cat | 0.008672693 | 0.249947463 | 0.551180864 | 0.073389354 | -0.18327225 | growth toward the midline |
| ENSDART00000166561 | ttc38 | 0.002390041 | 0.445202505 | 0.288306634 | -0.06639877 | -0.156646921 | growth toward the midline |
| ENSDART00000166591 | utp20 | 0.017280106 | 0.378085419 | 0.246513796 | 0.276819854 | 0.15697444 | growth toward the midline |
| ENSDART00000166632 | si:zfos-741a10.3 | 0.016381744 | 1.2129476 | 0.572994738 | 0.242126634 | 0.029899239 | growth toward the midline |
| ENSDART00000166690 | gadd45ga | 0.000475692 | 1.824520476 | 2.195526461 | 0.938941318 | 1.401631316 | growth toward the midline |
| ENSDART00000166725 | si:dkey-27j5.9 | 0.024411518 | 1.657914875 | 0.40212862 | 0.542892045 | 0.141569929 | growth toward the midline |
| ENSDART00000166755 | tril | 0.013035616 | 1.481556618 | 0.917097925 | 0.535942919 | 0.363878102 | growth toward the midline |
| ENSDART00000167279 | klhl6 | 0.010535127 | 0.921934193 | 0.563560813 | 0.364277897 | 0.100202901 | growth toward the midline |
| ENSDART00000167468 | prpsap1 | 0.0101684 | 0.532862733 | 0.121505744 | 0.13944499 | -0.034080691 | growth toward the midline |
| ENSDART00000167748 | fbn2b | 0.005787591 | 1.02333483 | 0.65734038 | 0.922883374 | 0.19244827 | growth toward the midline |
| ENSDART00000167786 | NA | 0.032175544 | 0.801350724 | 0.147008366 | 0.081187044 | 0.322292312 | growth toward the midline |
| ENSDART00000167937 | p4hb | 0.000670207 | 0.35105827 | 0.247669444 | -0.015299318 | -0.184059227 | growth toward the midline |
| ENSDART00000167971 | rps18 | 0.008672693 | 0.367747716 | 0.388045404 | 0.209564172 | -0.058761163 | growth toward the midline |
| ENSDART00000167986 | hadhb | 0.00721984 | 0.310832473 | 0.238522601 | 0.004510639 | -0.025410475 | growth toward the midline |
| ENSDART00000168002 | laptm5 | 0.00340435 | 1.090713638 | 0.609708871 | 0.327355694 | -0.215403696 | growth toward the midline |
| ENSDART00000168472 | creb5a | 0.024581131 | 2.212193939 | 1.443174599 | 0.809493268 | 0.247644913 | growth toward the midline |
| ENSDART00000168608 | rnaset2 | 0.002456687 | 1.085971834 | 0.685727013 | 0.397141355 | 0.382009137 | growth toward the midline |
| ENSDART00000168727 | zgc:64065 | 0.021156666 | 0.439077495 | 0.173893761 | 0.133684815 | 0.136149811 | growth toward the midline |
| ENSDART00000168831 | si:ch73-158p21.3 | 0.004606351 | 1.161771941 | 0.892729958 | 0.502592433 | 0.147950065 | growth toward the midline |
| ENSDART00000168840 | rpl35a | 0.004477335 | 0.357031004 | 0.320334573 | 0.163051637 | -0.036511823 | growth toward the midline |
| ENSDART00000169006 | ak2 | 0.034654408 | 0.70404039 | 0.489848323 | 0.563968118 | 0.137147044 | growth toward the midline |
| ENSDART00000169202 | si:ch211-153b23.5 | 0.004687334 | 4.343063064 | 5.139014825 | 3.462179286 | 1.548865865 | growth toward the midline |
| ENSDART00000169209 | ptrh1 | 0.000931909 | 0.68284848 | 0.772674127 | 0.446966304 | 0.22472723 | growth toward the midline |
| ENSDART00000169600 | abhd4 | 0.004142854 | 0.160082589 | 0.423680439 | -0.10396044 | -0.08495852 | growth toward the midline |
| ENSDART00000169633 | col4a3bpb | 0.002735847 | 0.489929413 | 0.127889926 | -0.192097119 | -0.259910666 | growth toward the midline |
| ENSDART00000169789 | grap2b | 0.054938882 | 0.941263606 | 0.635408975 | 0.416956975 | 0.146969065 | growth toward the midline |
| ENSDART00000169834 | CABZ01052431.1 | 0.000470359 | 0.752252615 | 0.635370532 | 0.523133762 | 0.005708707 | growth toward the midline |
| ENSDART00000169976 | CABZ01034529.1 | 0.002936628 | 0.902549952 | 0.806876908 | 0.306228824 | 0.021333022 | growth toward the midline |
| ENSDART00000169993 | stx12 | 0.009181211 | 0.289245416 | 0.334302649 | 0.162033659 | 0.014864416 | growth toward the midline |
| ENSDART00000170060 | mdh1b | 0.024987601 | 3.761355509 | 3.246314783 | 3.336555843 | 1.787365183 | growth toward the midline |
| ENSDART00000170304 | CABZ01039096.1 | 0.03168371 | 1.863106121 | 1.329668381 | 1.247074235 | 0.639142974 | growth toward the midline |
| ENSDART00000170326 | gadd45ba | 0.013456921 | 1.741159479 | 1.754335562 | 1.06756424 | 0.870946071 | growth toward the midline |
| ENSDART00000170601 | si:ch1073-228h2.2 | 0.032809386 | 1.512427811 | 1.187632525 | 1.253650535 | 1.028624188 | growth toward the midline |
| ENSDART00000170755 | slc30a8 | 0.000398071 | 0.607491231 | 0.345740737 | -0.026161917 | -0.397029232 | growth toward the midline |
| ENSDART00000170758 | tcirg1b | 0.007786985 | 0.779415513 | 0.215400071 | 0.036968665 | -0.339490039 | growth toward the midline |
| ENSDART00000170810 | si:dkey-10p5.7 | 0.01179775 | 1.115159817 | 1.313044185 | 0.773496009 | 0.334257673 | growth toward the midline |
| ENSDART00000171090 | sowahd | 0.01612441 | 0.56962557 | 0.415649109 | 0.184768671 | -0.065367372 | growth toward the midline |
| ENSDART00000171179 | si:ch211-241b2.5 | 0.005678515 | 1.438237825 | 1.596241839 | 1.013176313 | 0.055616895 | growth toward the midline |
| ENSDART00000171270 | ckap2l | 0.013513343 | 0.621565842 | 0.655006082 | 0.304610509 | 0.072068559 | growth toward the midline |
| ENSDART00000171345 | NA | 0.060649685 | 2.015201895 | 1.694676698 | 1.017867776 | 0.711389392 | growth toward the midline |
| ENSDART00000171430 | BX908401.1 | 0.055863259 | 0.558597997 | 0.282396796 | 0.308082083 | 0.228227036 | growth toward the midline |
| ENSDART00000171789 | NA | 0.00889295 | 1.544321358 | 1.350890859 | 1.388203792 | 0.786947546 | growth toward the midline |
| ENSDART00000172069 | snrpg | 0.014753252 | 0.452166641 | 0.481061823 | 0.338564477 | 0.058821342 | growth toward the midline |
| ENSDART00000172135 | sh3pxd2b | 0.005584581 | 0.68385955 | 0.832942161 | 0.517904326 | 0.090715158 | growth toward the midline |
| ENSDART00000172201 | trpv1 | 0.0044585 | 0.412354182 | 0.082280105 | 0.061779394 | -0.254715035 | growth toward the midline |
| ENSDART00000172207 | si:ch211-9d9.1 | 0.000466856 | 1.222547885 | 0.297262625 | 0.271454205 | -2.80261928 | growth toward the midline |
| ENSDART00000172408 | arhgap11a | 0.030198325 | 0.695056063 | 0.093403628 | 0.066933123 | 0.263951638 | growth toward the midline |
| ENSDART00000172555 | rps15a | 0.004379828 | 0.386196694 | 0.41469411 | 0.231657808 | -0.026260131 | growth toward the midline |
| ENSDART00000172892 | CR769769.2 | 0.02363886 | 0.615702823 | 0.344273677 | 0.373739938 | 0.336791427 | growth toward the midline |
| ENSDART00000172893 | si:ch211-207g17.2 | 0.01138794 | 1.081136407 | 0.564546524 | -0.069203481 | -0.244746802 | growth toward the midline |
| ENSDART00000172953 | rpl29 | 0.008586878 | 0.354363113 | 0.457581567 | 0.206775836 | 0.026974968 | growth toward the midline |
| ENSDART00000173145 | rpl29 | 0.006013085 | 0.32197024 | 0.280412475 | 0.068293144 | -0.107729465 | growth toward the midline |
| TRANSGENE | NA | 7.26665E-10 | 3.62113341 | 4.519443337 | 2.76591932 | 1.041497447 | growth toward the midline |
| ENSDART00000002186 | uck2a | 0.001969989 | 0.69693934 | 0.733529046 | 0.658757388 | 0.439621207 | midline crossing |
| ENSDART00000002250 | hs6st2 | 0.009748008 | 0.156001692 | 0.361544481 | 0.514379395 | 0.208727247 | midline crossing |
| ENSDART00000002641 | kif26aa | 0.003651057 | 0.255726279 | 0.47696992 | 0.58781469 | 0.342072456 | midline crossing |
| ENSDART00000002932 | marcksb | 0.000213603 | 1.016829958 | 1.677411962 | 1.758104803 | 1.229873931 | midline crossing |
| ENSDART00000003066 | cyth1a | 0.000219336 | 0.591135613 | 0.827117646 | 0.654961234 | 0.137500638 | midline crossing |
| ENSDART00000003076 | usp28 | 0.011417724 | 1.676441425 | 4.659547988 | 4.384020448 | 3.839785573 | midline crossing |
| ENSDART00000003296 | sars | 0.039604681 | 0.182171112 | 0.390636242 | 0.505424837 | 0.283972917 | midline crossing |
| ENSDART00000003752 | cct3 | 0.006479841 | 0.348496547 | 0.407130663 | 0.410390239 | 0.168454536 | midline crossing |
| ENSDART00000003790 | pwp1 | 0.01633867 | 0.32756693 | 0.411455433 | 0.339342319 | 0.085530698 | midline crossing |
| ENSDART00000003891 | jupa | 0.020463519 | 0.359241944 | 1.297843964 | 1.552619484 | 0.724987275 | midline crossing |
| ENSDART00000004423 | iars | 0.002440466 | 0.363891152 | 0.394414274 | 0.599518689 | 0.155539422 | midline crossing |
| ENSDART00000004474 | mapre1b | 0.001177059 | 0.531914947 | 0.849936108 | 0.970422656 | 0.515027849 | midline crossing |
| ENSDART00000005119 | eif3i | 0.018217202 | 0.208433038 | 0.349717288 | 0.244101269 | 0.053458955 | midline crossing |

| | | | | | | | |
|---|---|---|---|---|---|---|---|
| ENSDART00000005593 | casp3a | 0.000114847 | 0.498469533 | 1.198942327 | 1.332280554 | 0.912417093 | midline crossing |
| ENSDART00000005616 | rnpep | 0.025134747 | 0.523525834 | 0.750055071 | 0.77634383 | 0.462027268 | midline crossing |
| ENSDART00000005847 | nav3 | 6.75287E-05 | 0.773382381 | 1.009593057 | 0.901548272 | 0.783565613 | midline crossing |
| ENSDART00000005944 | rpl5a | 0.001179374 | 0.381988038 | 0.550810639 | 0.338076702 | 0.013320357 | midline crossing |
| ENSDART00000006058 | eif2s1a | 0.03277385 | 0.414016542 | 0.357076684 | 0.28859634 | 0.112415797 | midline crossing |
| ENSDART00000006908 | itgb3b | 2.33589E-05 | 1.52546911 | 1.915133846 | 1.613887964 | 0.668451164 | midline crossing |
| ENSDART00000007231 | psmb1 | 0.012813136 | 0.336843399 | 0.41477024 | 0.441532829 | 0.103495117 | midline crossing |
| ENSDART00000007789 | idh1 | 0.001120005 | 0.209391072 | 0.491026617 | 0.59658334 | 0.190896441 | midline crossing |
| ENSDART00000007857 | mettl2a | 0.019840997 | 0.337109376 | 0.448868897 | 0.378403392 | 0.127678255 | midline crossing |
| ENSDART00000009477 | cct8 | 0.008761738 | 0.2739469 | 0.4225965 | 0.299361505 | 0.113502813 | midline crossing |
| ENSDART00000009549 | rhag | 0.050085111 | 0.97898574 | 1.27206332 | 1.678759018 | 0.617102069 | midline crossing |
| ENSDART00000009609 | eif5a | 0.00304733 | 0.382861639 | 0.513300938 | 0.466153608 | 0.171038659 | midline crossing |
| ENSDART00000010256 | eif3m | 0.008283592 | 0.301540845 | 0.415293475 | 0.295465222 | 0.073592881 | midline crossing |
| ENSDART00000010271 | aida | 0.002673241 | 0.207262438 | 0.531449642 | 0.462716258 | 0.148174666 | midline crossing |
| ENSDART00000010647 | rcc2 | 0.040200236 | 0.312248055 | 0.54892666 | 0.398800486 | 0.1537421 | midline crossing |
| ENSDART00000010683 | impa1 | 0.024809879 | 0.486043053 | 0.448191747 | 0.382482237 | 0.291162031 | midline crossing |
| ENSDART00000010997 | tpm3 | 0.039972057 | 0.446911874 | 0.508209148 | 0.705423057 | 0.171806011 | midline crossing |
| ENSDART00000011258 | npl | 0.042570446 | 0.373272645 | 0.54061543 | 0.468726439 | 0.170404917 | midline crossing |
| ENSDART00000011447 | sae1 | 0.006887747 | 0.282597281 | 0.674186287 | 0.734155697 | 0.424524708 | midline crossing |
| ENSDART00000011691 | baxa | 0.00197074 | 0.500319714 | 0.752407609 | 0.690306826 | 0.30333895 | midline crossing |
| ENSDART00000011699 | nono | 0.019981901 | 0.031722902 | 0.329157035 | 0.240624596 | 0.057167441 | midline crossing |
| ENSDART00000011878 | eif4a1b | 0.023987365 | 0.134279506 | 0.344064551 | 0.309281202 | 0.284267369 | midline crossing |
| ENSDART00000012164 | tmod2 | 0.019466285 | 0.222306993 | 0.411676025 | 0.560901639 | 0.297325519 | midline crossing |
| ENSDART00000012686 | dnase1l4.1 | 0.01793082 | 0.654985436 | 1.279054377 | 1.240763186 | 0.531962605 | midline crossing |
| ENSDART00000012791 | sp8a | 0.049626195 | 1.57251014 | 2.331511764 | 2.00847442 | 1.039704847 | midline crossing |
| ENSDART00000014049 | wdr36 | 0.014675171 | 0.429321608 | 0.423447996 | 0.351992847 | 0.181701695 | midline crossing |
| ENSDART00000014058 | zgc:100829 | 0.002661298 | 0.836226257 | 1.366715788 | 1.414785917 | 0.344237354 | midline crossing |
| ENSDART00000014983 | zgc:153867 | 0.00612514 | 0.36977967 | 0.600215172 | 0.463408621 | 0.079627153 | midline crossing |
| ENSDART00000015103 | hps3 | 0.030383057 | 0.398719216 | 0.34180849 | 0.31802597 | 0.20234989 | midline crossing |
| ENSDART00000015193 | chmp4bb | 0.001754036 | 0.349511966 | 0.500219693 | 0.425249872 | 0.076172611 | midline crossing |
| ENSDART00000015374 | cyb5r1 | 0.020123282 | 0.481284363 | 0.621601719 | 0.549959374 | 0.29311095 | midline crossing |
| ENSDART00000015628 | klhl24b | 0.021895107 | 0.257620397 | 0.582600724 | 0.358856753 | 0.009507758 | midline crossing |
| ENSDART00000016112 | capns1b | 0.000336326 | 0.258538261 | 0.719472725 | 0.61845775 | 0.248329876 | midline crossing |
| ENSDART00000016135 | nfe2l3 | 0.02142158 | 0.030439041 | 0.342226134 | 0.430712572 | 0.086440986 | midline crossing |
| ENSDART00000016464 | dcps | 0.017642395 | 0.045608002 | 0.377363095 | -0.076988765 | 0.017970252 | midline crossing |
| ENSDART00000016591 | fgf6a | 0.000507398 | 0.502410209 | 1.375431955 | 1.297903392 | 1.026892284 | midline crossing |
| ENSDART00000016628 | fam129bb | 0.003338328 | 0.861865647 | 1.020541987 | 0.89470832 | 0.182764632 | midline crossing |
| ENSDART00000016946 | glud1a | 0.066889451 | 0.489439701 | 0.519599049 | 0.659106358 | 0.391317016 | midline crossing |
| ENSDART00000017230 | snrpc | 0.012958044 | 0.173923367 | 0.335809723 | 0.464722714 | 0.188235196 | midline crossing |
| ENSDART00000017422 | tbc1d17 | 0.000844564 | 0.129347814 | 0.49736446 | 0.343125168 | -0.01149421 | midline crossing |
| ENSDART00000018408 | anxa13l | 5.81566E-05 | 0.767695981 | 1.876785961 | 1.969202275 | 1.362090979 | midline crossing |
| ENSDART00000018685 | syt9a | 0.018500869 | 0.383050894 | 0.997731574 | 0.678563906 | 0.37205351 | midline crossing |
| ENSDART00000018743 | phf20a | 0.027269932 | 0.312029532 | 0.419745206 | 0.325238046 | 0.318832686 | midline crossing |
| ENSDART00000019521 | dip2ba | 0.005277161 | 0.352814519 | 0.501689217 | 0.539560646 | 0.272762208 | midline crossing |
| ENSDART00000019698 | anxa5b | 0.006899426 | 1.162500386 | 1.67744918 | 1.46667303 | 0.432101083 | midline crossing |
| ENSDART00000019750 | wdr5 | 0.067593058 | 0.151689533 | 0.300718903 | 0.396766724 | 0.199844693 | midline crossing |
| ENSDART00000019818 | ric8b | 0.011108721 | 0.219425238 | 0.304273846 | 0.411142498 | 0.146708414 | midline crossing |
| ENSDART00000020048 | gsna | 0.018144005 | 0.451913636 | 0.79824491 | 0.782008443 | 0.357949724 | midline crossing |
| ENSDART00000020084 | hsp90ab1 | 0.015156186 | 0.266636397 | 0.434039186 | 0.537534154 | 0.384889667 | midline crossing |
| ENSDART00000020256 | lgsn | 0.036969899 | 1.361702081 | 2.686172063 | 1.394569806 | 0.349584846 | midline crossing |
| ENSDART00000020638 | rcan1a | 0.04859805 | 0.548382891 | 0.378475469 | 0.383543152 | 0.544764217 | midline crossing |
| ENSDART00000021666 | rtca | 0.004731414 | 0.450183742 | 0.82121323 | 0.721903733 | 0.20677388 | midline crossing |
| ENSDART00000022060 | atf3 | 2.2034E-05 | 2.958540036 | 3.348918067 | 2.852908333 | 1.811501419 | midline crossing |
| ENSDART00000023123 | nup88 | 0.017159406 | 0.221684741 | 0.406336851 | 0.406160624 | 0.250704697 | midline crossing |
| ENSDART00000023156 | eef1a1l2 | 0.002558097 | 0.512554224 | 0.741741761 | 0.709552788 | 0.28897731 | midline crossing |
| ENSDART00000023709 | ptp4a2b | 0.023468821 | 0.271575512 | 0.562448581 | 0.634906429 | 0.086969331 | midline crossing |
| ENSDART00000023779 | vcp | 0.053123832 | 0.230561184 | 0.246570266 | 0.325389872 | 0.137878127 | midline crossing |
| ENSDART00000023833 | eif2s3 | 0.013547844 | 0.172990624 | 0.349255612 | 0.311895175 | 0.045890851 | midline crossing |
| ENSDART00000024309 | rb1 | 0.001317626 | 0.736743325 | 0.968558289 | 0.720111001 | 0.238176467 | midline crossing |
| ENSDART00000024320 | ybx1 | 0.075082889 | 0.257630768 | 0.252218726 | 0.414084055 | 0.18414263 | midline crossing |
| ENSDART00000024662 | plppr3a | 0.000164013 | 1.050789931 | 1.738581828 | 1.609411361 | 1.082826483 | midline crossing |
| ENSDART00000024872 | creb3l3l | 0.003959852 | 0.316205692 | 0.628018553 | 0.525420097 | 0.254183225 | midline crossing |
| ENSDART00000025385 | cers2a | 0.00721984 | 0.59368354 | 0.657003513 | 0.583941604 | 0.192392368 | midline crossing |
| ENSDART00000025496 | rras | 0.010693822 | 0.483455604 | 0.817924325 | 0.502791397 | -0.193021858 | midline crossing |
| ENSDART00000025535 | sept5a | 0.004815134 | 0.521581645 | 1.371610673 | 0.945407012 | 0.566204111 | midline crossing |
| ENSDART00000025550 | top1mt | 0.003833624 | 0.816108961 | 1.133963099 | 0.883225094 | 0.407533529 | midline crossing |
| ENSDART00000025620 | ppiaa | 0.002705637 | 0.369681961 | 0.563046067 | 0.555154111 | 0.174901331 | midline crossing |
| ENSDART00000025782 | nup93 | 0.062586781 | 0.17632244 | 0.241668195 | 0.293793434 | 0.152368608 | midline crossing |
| ENSDART00000026180 | fabp7a | 0.006795032 | 0.745585804 | 2.003621538 | 1.857780253 | 1.020257385 | midline crossing |
| ENSDART00000026339 | gtpbp4 | 0.048096163 | 0.113641949 | 0.327640711 | 0.222548232 | 0.071359586 | midline crossing |
| ENSDART00000026409 | cct4 | 0.00602897 | 0.459151422 | 0.538338734 | 0.621039126 | 0.313903974 | midline crossing |
| ENSDART00000026814 | ptp4a1 | 0.004605292 | 0.419787699 | 0.591512581 | 0.636748721 | 0.300535035 | midline crossing |
| ENSDART00000027616 | eif4g2a | 0.000406473 | 0.333990776 | 0.435258045 | 0.644700731 | 0.350291836 | midline crossing |
| ENSDART00000028090 | eif2ak1 | 0.011517304 | 0.218229206 | 0.400243646 | 0.398265404 | 0.179590234 | midline crossing |
| ENSDART00000029843 | vezf1a | 0.001561003 | 0.660507751 | 0.776194217 | 0.884683465 | 0.428803142 | midline crossing |

| | | | | | | | |
|---|---|---|---|---|---|---|---|
| ENSDART00000030205 | bnip3lb | 0.010960196 | 0.087436849 | 0.295010863 | 0.335325724 | 0.120618456 | midline crossing |
| ENSDART00000030691 | clic4 | 0.028282855 | 0.290171682 | 0.490608957 | 0.279069846 | 0.046035564 | midline crossing |
| ENSDART00000032322 | abcg2c | 0.004813371 | 0.455571157 | 0.699504596 | 0.641510999 | 0.335341634 | midline crossing |
| ENSDART00000032324 | hddc3 | 0.020485853 | 0.491004501 | 0.803369277 | 0.623276564 | 0.474326204 | midline crossing |
| ENSDART00000032331 | gmppab | 0.008531138 | 0.298836739 | 0.452114759 | 0.362408461 | -0.028707454 | midline crossing |
| ENSDART00000032540 | usp14 | 0.04750637 | 0.236991779 | 0.380670741 | 0.291281165 | 0.061189525 | midline crossing |
| ENSDART00000032603 | tspo | 0.008848885 | 0.47255795 | 0.736846252 | 0.52533392 | 0.11686825 | midline crossing |
| ENSDART00000032857 | mapk11 | 0.047290605 | 0.353170314 | 0.603154564 | 0.6388701 | 0.458511735 | midline crossing |
| ENSDART00000033316 | vangl2 | 0.00277587 | 0.207091244 | 0.447705982 | 0.579933447 | 0.26125294 | midline crossing |
| ENSDART00000033479 | si:ch211-129c21.1 | 0.00140081 | 1.539581327 | 2.000726167 | 1.978342783 | 1.084446492 | midline crossing |
| ENSDART00000033494 | klf6a | 0.000170861 | 1.084550888 | 1.797692613 | 1.625562666 | 0.460444617 | midline crossing |
| ENSDART00000033566 | smad1 | 0.004676707 | 0.486585209 | 1.249927606 | 1.264727704 | 0.536891999 | midline crossing |
| ENSDART00000034523 | tars | 0.044608572 | 0.184292328 | 0.408838041 | 0.43347383 | 0.219690833 | midline crossing |
| ENSDART00000034784 | adcyap1b | 0.000144061 | 1.680397515 | 2.015053329 | 1.852069794 | 1.09666998 | midline crossing |
| ENSDART00000034935 | desi2 | 0.009532057 | 0.073850834 | 0.377607475 | 0.427284588 | 0.193399467 | midline crossing |
| ENSDART00000035899 | pkp2 | 0.002199978 | 0.428534184 | 1.016588803 | 1.016153499 | 0.312913104 | midline crossing |
| ENSDART00000036513 | trib3 | 0.00028941 | 0.704680301 | 0.933969628 | 0.780405513 | 0.230790744 | midline crossing |
| ENSDART00000036668 | psmc1a | 0.017576487 | 0.311771761 | 0.423109618 | 0.307731242 | 0.107467578 | midline crossing |
| ENSDART00000036703 | pfdn2 | 0.011267245 | 0.369409429 | 0.460943577 | 0.431067405 | 0.197162016 | midline crossing |
| ENSDART00000036797 | uchl1 | 0.000254175 | 1.004812771 | 1.916965558 | 1.938952257 | 1.446591845 | midline crossing |
| ENSDART00000036939 | gadd45ba | 2.18017E-05 | 1.15110753 | 1.253886794 | 1.06146692 | 0.82843487 | midline crossing |
| ENSDART00000037195 | kif26bb | 6.9873E-05 | 1.206892082 | 1.55094827 | 1.874551064 | 1.519778039 | midline crossing |
| ENSDART00000037698 | uck2b | 6.99374E-05 | 0.641974401 | 1.119646558 | 1.077673009 | 0.598617402 | midline crossing |
| ENSDART00000037709 | nol11 | 0.023203992 | 0.254424392 | 0.434300513 | 0.299474854 | 0.102573384 | midline crossing |
| ENSDART00000037846 | focad | 0.014575529 | 0.435964232 | 0.512370823 | 0.456134064 | 0.112317494 | midline crossing |
| ENSDART00000037850 | dync1li2 | 0.002061124 | 0.268558729 | 0.496891146 | 0.585345447 | 0.309833954 | midline crossing |
| ENSDART00000039443 | tuba8l4 | 0.002876227 | 0.45016266 | 0.547239574 | 0.701695743 | 0.194541839 | midline crossing |
| ENSDART00000039746 | epb41b | 0.061238247 | 0.556721113 | 0.962382373 | 1.378686301 | 0.45048608 | midline crossing |
| ENSDART00000040066 | adam9 | 0.002930263 | 0.358589887 | 0.583904956 | 0.549640677 | 0.256288209 | midline crossing |
| ENSDART00000040804 | praf2 | 0.017079861 | 0.166225557 | 0.427661756 | 0.402173801 | 0.274020381 | midline crossing |
| ENSDART00000041191 | gyg2 | 0.019039254 | 0.909324979 | 1.617533024 | 1.086480874 | 0.519765295 | midline crossing |
| ENSDART00000041279 | tubb4b | 0.017496248 | 0.433747505 | 0.468816243 | 0.626770783 | 0.445102188 | midline crossing |
| ENSDART00000041468 | ap1ar | 0.000902118 | 0.253231094 | 0.535778421 | 0.657902256 | 0.098067269 | midline crossing |
| ENSDART00000041877 | csrnp1a | 0.000166716 | 1.299872398 | 1.369159722 | 1.350463811 | 0.717287101 | midline crossing |
| ENSDART00000042189 | pdk2b | 0.007668959 | 0.734399082 | 1.569570791 | 1.007918384 | 0.211070696 | midline crossing |
| ENSDART00000042297 | kdelc1 | 0.002843439 | 0.328425656 | 0.736886183 | 0.437422887 | 0.039597286 | midline crossing |
| ENSDART00000042963 | chst11 | 0.007952083 | 0.296547318 | 0.476599287 | 0.638913095 | 0.263644905 | midline crossing |
| ENSDART00000044154 | tnnt2c | 0.012339321 | 0.256126919 | 0.781011759 | 0.616305237 | 0.377618007 | midline crossing |
| ENSDART00000045299 | ola1 | 0.002876227 | 0.514191862 | 0.666266592 | 0.533008533 | 0.271611648 | midline crossing |
| ENSDART00000045479 | syt4 | 0.022209173 | 0.199773705 | 0.561264928 | 0.42766733 | -0.021116223 | midline crossing |
| ENSDART00000045682 | rrp36 | 0.024581131 | 0.38981283 | 0.603671455 | 0.338055221 | 0.296271067 | midline crossing |
| ENSDART00000046115 | mfsd2aa | 0.004106312 | 0.413173515 | 1.356876257 | 1.539393726 | 0.598141045 | midline crossing |
| ENSDART00000046209 | acbd7 | 0.004793697 | 0.172970527 | 0.805855312 | 0.630012733 | 0.123911744 | midline crossing |
| ENSDART00000046587 | ap2m1a | 0.018091301 | 0.393397352 | 0.521859633 | 0.500225377 | 0.048547949 | midline crossing |
| ENSDART00000046973 | capza1a | 0.016550715 | 0.117659515 | 0.282132124 | 0.303069026 | 0.119362421 | midline crossing |
| ENSDART00000047020 | casp9 | 0.000847723 | 0.406413734 | 0.845063479 | 0.618979742 | 0.291728295 | midline crossing |
| ENSDART00000047073 | oxsr1a | 0.02511033 | 0.179223329 | 0.268680956 | 0.376160209 | 0.212719894 | midline crossing |
| ENSDART00000048383 | creld2 | 0.024987601 | 0.499343204 | 0.520824125 | 0.482480278 | 0.212076819 | midline crossing |
| ENSDART00000048977 | abcf1 | 0.002096961 | 0.295994115 | 0.413143633 | 0.435605861 | 0.158915281 | midline crossing |
| ENSDART00000049264 | sdr16c5b | 0.002217387 | 0.496225849 | 0.925512272 | 0.840394736 | 0.492626723 | midline crossing |
| ENSDART00000049373 | cmtr1 | 0.067400884 | 3.529734054 | 4.85424569 | 5.483410402 | 3.408663479 | midline crossing |
| ENSDART00000049434 | scamp4 | 0.020544535 | 0.767657025 | 0.801831806 | 0.544244726 | 0.399234701 | midline crossing |
| ENSDART00000049464 | fermt2 | 0.02086388 | 0.28468341 | 0.302053037 | 0.328009634 | 0.040193822 | midline crossing |
| ENSDART00000051621 | pgam5 | 0.017997171 | 0.243607349 | 0.33552261 | 0.455304621 | 0.149179919 | midline crossing |
| ENSDART00000052168 | hrh3 | 0.001356985 | 0.375251408 | 0.91270689 | 1.213693931 | 0.380712944 | midline crossing |
| ENSDART00000052421 | txnipa | 0.000860824 | 1.054359781 | 1.031010785 | 1.048664605 | 0.430054853 | midline crossing |
| ENSDART00000052539 | myo1ea | 0.012462623 | 0.574953247 | 0.619444947 | 0.654028338 | 0.173056841 | midline crossing |
| ENSDART00000052656 | rras2 | 0.005262446 | 0.556656504 | 0.801505793 | 0.534247735 | 0.118931735 | midline crossing |
| ENSDART00000052915 | ash1l | 0.056562408 | 0.173309311 | 0.167145004 | 0.317504355 | 0.12008578 | midline crossing |
| ENSDART00000053240 | cab39l | 0.013803709 | 0.313790093 | 0.462769475 | 0.441611468 | 0.290026864 | midline crossing |
| ENSDART00000053304 | si:ch211-114n24.6 | 0.000776925 | 0.627425485 | 0.596987541 | 0.770158215 | 0.21603311 | midline crossing |
| ENSDART00000053325 | tomm40l | 0.008715601 | 0.351677862 | 0.400415779 | 0.406067875 | 0.134025152 | midline crossing |
| ENSDART00000053925 | mtmr7a | 0.018204569 | 0.252266702 | 0.492232945 | 0.532543286 | 0.441567176 | midline crossing |
| ENSDART00000054062 | nek12 | 0.002113455 | 0.979847079 | 2.121821413 | 1.72566276 | 1.104924836 | midline crossing |
| ENSDART00000054243 | dpf2l | 0.013354221 | 0.196658591 | 0.403359284 | 0.517168503 | 0.26603871 | midline crossing |
| ENSDART00000054849 | pls3 | 0.01571154 | 0.365005671 | 0.852391802 | 1.00799598 | 0.660436059 | midline crossing |
| ENSDART00000054987 | actb1 | 0.002610721 | 0.719668079 | 0.845173501 | 0.884967639 | 0.398548729 | midline crossing |
| ENSDART00000055171 | grapa | 0.005866285 | 0.991444674 | 1.026501701 | 0.907734856 | 0.689609359 | midline crossing |
| ENSDART00000055253 | filip1l | 0.03479782 | 3.442073683 | 3.796340861 | 4.199635374 | 3.015929109 | midline crossing |
| ENSDART00000055325 | psmb7 | 0.009948051 | 3.620821341 | 3.668842367 | 3.694371073 | 3.321198364 | midline crossing |
| ENSDART00000055609 | atf4b | 0.027440234 | 0.138344185 | 0.51758353 | 0.353617358 | 0.087935611 | midline crossing |
| ENSDART00000056254 | stap2a | 0.001692745 | 0.934389037 | 1.208596726 | 1.287672321 | 0.743017194 | midline crossing |
| ENSDART00000056294 | pitrm1 | 0.005065079 | 0.368247543 | 0.389541704 | 0.328836098 | 0.178268037 | midline crossing |
| ENSDART00000056712 | etfdh | 0.011891263 | 0.23563688 | 0.425716128 | 0.377386118 | 0.136575345 | midline crossing |
| ENSDART00000056865 | ctnnbip1 | 0.001123946 | 0.431885199 | 0.971175617 | 1.010655462 | 0.599546466 | midline crossing |

| | | | | | | | |
|---|---|---|---|---|---|---|---|
| ENSDART00000056987 | marcksl1a | 0.024073872 | 0.199644536 | 0.480185443 | 0.561780108 | 0.295794028 | midline crossing |
| ENSDART00000057159 | cacnb1 | 0.000793313 | 1.304600132 | 2.011299334 | 2.307200837 | 1.559835487 | midline crossing |
| ENSDART00000057318 | dusp8b | 0.001038693 | 0.80659762 | 0.962100249 | 0.680537568 | 0.386784732 | midline crossing |
| ENSDART00000058258 | gng5 | 0.005032085 | 0.433032497 | 0.476321934 | 0.436326472 | 0.078774872 | midline crossing |
| ENSDART00000058485 | rai14 | 0.003945154 | 0.377402754 | 0.592516996 | 0.371359301 | 0.056635681 | midline crossing |
| ENSDART00000059955 | ildr1b | 0.057189105 | 0.550873694 | 0.893755801 | 0.988897575 | 0.559320969 | midline crossing |
| ENSDART00000060181 | zgc:114174 | 0.000530705 | 0.760260394 | 1.239485919 | 1.043412191 | 0.428827471 | midline crossing |
| ENSDART00000060255 | blmh | 0.003785955 | 0.318055181 | 0.435968503 | 0.441694486 | 0.152219233 | midline crossing |
| ENSDART00000060812 | adcyap1b | 0.000184957 | 1.704872196 | 2.272116909 | 1.908485885 | 0.966573225 | midline crossing |
| ENSDART00000060910 | pimr138 | 0.000775407 | 0.646921071 | 1.324373168 | 1.239639246 | 0.98486272 | midline crossing |
| ENSDART00000060919 | qars | 0.019842178 | 0.348349072 | 0.542908344 | 0.574420452 | 0.306598142 | midline crossing |
| ENSDART00000061196 | NA | 0.000874891 | 0.755433358 | 1.498415514 | 1.365853078 | 0.846252469 | midline crossing |
| ENSDART00000062704 | plaa | 0.003777754 | 0.222653649 | 0.333324552 | 0.491597491 | 0.251933193 | midline crossing |
| ENSDART00000063359 | ucp3 | 0.038034192 | 0.449516929 | 0.816201403 | 1.070983817 | 0.155240874 | midline crossing |
| ENSDART00000063418 | nsun5 | 0.015044537 | 0.551107756 | 0.480901258 | 0.442024494 | 0.196344317 | midline crossing |
| ENSDART00000063912 | jun | 0.002341904 | 1.166146477 | 1.429328618 | 1.29995447 | 0.480560375 | midline crossing |
| ENSDART00000063950 | psmc1b | 0.032472485 | 0.214901771 | 0.338782243 | 0.27453656 | 0.103637979 | midline crossing |
| ENSDART00000064032 | eif4ebp1 | 0.004205562 | 0.426193835 | 0.395196044 | 0.352513449 | 0.114800609 | midline crossing |
| ENSDART00000064067 | ehbp1 | 0.008004874 | 0.439358276 | 0.535882532 | 0.636938582 | 0.35957066 | midline crossing |
| ENSDART00000064311 | arhgdia | 0.018217202 | 0.49745023 | 0.520698759 | 0.529544756 | 0.127344369 | midline crossing |
| ENSDART00000064462 | psma6l | 0.016647785 | 0.604857346 | 0.425092318 | 0.840213456 | -0.035624292 | midline crossing |
| ENSDART00000064509 | stmn4l | 8.18133E-06 | 2.57043941 | 3.576220537 | 3.295095816 | 2.31833518 | midline crossing |
| ENSDART00000064666 | prnpb | 0.033001298 | 1.397742128 | 1.699664466 | 1.642053833 | 0.642245778 | midline crossing |
| ENSDART00000064789 | txn | 0.000250511 | 1.421742139 | 1.973125 | 1.39538042 | 0.274434823 | midline crossing |
| ENSDART00000064826 | mov10a | 0.000860824 | 0.329642907 | 0.141820564 | 0.774895549 | -0.030140661 | midline crossing |
| ENSDART00000064968 | rasgef1bb | 0.00655086 | 0.10103678 | 0.334291276 | 0.155368522 | -0.094945106 | midline crossing |
| ENSDART00000065563 | ccdc90b | 0.007961928 | 0.430098539 | 0.770932327 | 0.278586215 | 0.111905478 | midline crossing |
| ENSDART00000065728 | nrsn1 | 0.017738289 | 0.207141455 | 0.540557286 | 0.398847193 | 0.186515655 | midline crossing |
| ENSDART00000065807 | kctd13 | 0.056884231 | 0.261042411 | 0.352189581 | 0.50608049 | 0.336398415 | midline crossing |
| ENSDART00000066385 | hbz | 0.013081285 | 0.819612283 | 1.064127771 | 1.5494854 | 0.440895582 | midline crossing |
| ENSDART00000066471 | adam8b | 0.000414404 | 1.018895271 | 1.190253553 | 1.122506289 | 0.517972783 | midline crossing |
| ENSDART00000066760 | cct5 | 0.004847659 | 0.325276039 | 0.436851681 | 0.379245954 | 0.163513221 | midline crossing |
| ENSDART00000067053 | vta1 | 0.003953024 | 0.2430424 | 0.457630564 | 0.367763165 | 0.149685321 | midline crossing |
| ENSDART00000067147 | ANKRD50 | 0.000616782 | 0.122102324 | 0.471943767 | 0.546358337 | 0.270036712 | midline crossing |
| ENSDART00000067168 | pdzrn4 | 0.000390311 | 0.330865082 | 1.464493171 | 1.433744497 | 0.793719225 | midline crossing |
| ENSDART00000067500 | si:dkey-280e21.3 | 0.004246082 | 0.421024272 | 0.822626751 | 0.979586691 | 0.632877181 | midline crossing |
| ENSDART00000067542 | kcnk10b | 6.48692E-06 | 2.685681832 | 3.478119742 | 3.471523284 | 2.49042133 | midline crossing |
| ENSDART00000067776 | rab10 | 0.010011285 | 0.291742266 | 0.421774457 | 0.459760968 | 0.189497347 | midline crossing |
| ENSDART00000073452 | si:ch211-113a14.12 | 0.043318153 | 0.801955974 | 0.847093718 | 0.855214481 | 0.503313711 | midline crossing |
| ENSDART00000073564 | tes | 0.005380857 | 0.818084026 | 1.011033966 | 0.902633694 | 0.089277909 | midline crossing |
| ENSDART00000073705 | abcf1 | 0.050560604 | 0.284347738 | 0.522430424 | 0.46970431 | 0.327069748 | midline crossing |
| ENSDART00000073735 | rrad | 0.003154242 | 0.475668516 | 0.908053908 | 0.772751604 | 0.279273357 | midline crossing |
| ENSDART00000073970 | uap1 | 0.000781013 | 0.430311379 | 0.939111864 | 0.868311508 | 0.206630788 | midline crossing |
| ENSDART00000073981 | eif2s1b | 0.019981901 | 0.229006291 | 0.299741922 | 0.510349465 | 0.185458194 | midline crossing |
| ENSDART00000074317 | GSK3B (1 of many) | 0.040539001 | 0.32408852 | 0.570538304 | 0.615748631 | 0.310967098 | midline crossing |
| ENSDART00000074400 | tia1 | 0.004813371 | 0.181056891 | 0.310121029 | 0.46787956 | 0.221267203 | midline crossing |
| ENSDART00000075009 | eif2s2 | 0.015666457 | 0.324008789 | 0.419532633 | 0.338652657 | 0.07614456 | midline crossing |
| ENSDART00000075129 | lrrc47 | 0.043968276 | 0.095574868 | 0.246973338 | 0.327055852 | 0.135309168 | midline crossing |
| ENSDART00000075187 | pdzd11 | 0.061577067 | 1.031277997 | 1.70516213 | 1.574957324 | 1.135180778 | midline crossing |
| ENSDART00000075299 | zgc:153911 | 1.2193E-05 | 1.082638325 | 2.093301655 | 1.816780233 | 1.137984438 | midline crossing |
| ENSDART00000075351 | zgc:112285 | 0.031816406 | 0.930923442 | 1.110746674 | 0.609701543 | 0.79082813 | midline crossing |
| ENSDART00000075491 | pop5 | 0.007140172 | 0.444559136 | 0.7451635 | 0.571282976 | 0.537107281 | midline crossing |
| ENSDART00000076161 | hoxb5b | 0.013675681 | 2.652791903 | 3.351280664 | 2.618897772 | 0.864721852 | midline crossing |
| ENSDART00000076571 | rtn1a | 0.001728686 | 0.31833938 | 1.022959786 | 0.981624341 | 0.706878795 | midline crossing |
| ENSDART00000076600 | rpe65c | 0.030945673 | 1.983711129 | 2.589375601 | 1.902404689 | 1.435704768 | midline crossing |
| ENSDART00000076815 | NA | 0.020105477 | 2.456176486 | 3.15180058 | 2.959048559 | 2.414794476 | midline crossing |
| ENSDART00000077197 | tmsb | 2.92142E-06 | 1.972985822 | 3.598018383 | 3.544381625 | 2.894174576 | midline crossing |
| ENSDART00000077445 | pim3 | 0.007776904 | -0.017745548 | 0.425023548 | -0.018053873 | 0.016773558 | midline crossing |
| ENSDART00000077484 | zhx2a | 0.005803283 | 0.117201305 | 0.356185793 | 0.299966576 | 0.024327323 | midline crossing |
| ENSDART00000078156 | srm | 0.063878819 | 0.127220715 | 0.22303682 | 0.345313656 | 0.085476496 | midline crossing |
| ENSDART00000078304 | lzic | 0.034279564 | 0.119250866 | 0.44378754 | 0.377529349 | 0.128075007 | midline crossing |
| ENSDART00000078306 | arhgef2 | 0.005059311 | 0.429610468 | 0.807274797 | 0.854804837 | 0.533632211 | midline crossing |
| ENSDART00000078412 | rps8a | 0.006621714 | 0.416303691 | 0.456350868 | 0.419741526 | 0.122907452 | midline crossing |
| ENSDART00000078438 | NA | 0.072487837 | 0.22834743 | 0.583517385 | 0.218267131 | 0.08549295 | midline crossing |
| ENSDART00000078522 | eef1g | 0.00148037 | 0.462831941 | 0.651007925 | 0.567355432 | 0.209607784 | midline crossing |
| ENSDART00000078543 | syt11b | 0.000114847 | 0.975892204 | 1.6420197 | 1.925407845 | 1.338618886 | midline crossing |
| ENSDART00000078611 | jac2 | 0.020202236 | 3.404844888 | 3.429750736 | 2.921387124 | 1.113891823 | midline crossing |
| ENSDART00000078949 | afap1l1b | 6.16509E-05 | 1.061182692 | 1.814819842 | 1.643265844 | 0.877712611 | midline crossing |
| ENSDART00000078953 | afap1l1b | 0.00092561 | 0.880715307 | 1.833091326 | 1.686561879 | 0.852421188 | midline crossing |
| ENSDART00000079050 | nutf2 | 0.041930419 | 0.337010875 | 0.380281227 | 0.325036311 | 0.156803484 | midline crossing |
| ENSDART00000079173 | lepr | 0.000168321 | 1.173965688 | 1.129708824 | 0.920018098 | 0.6672678 | midline crossing |
| ENSDART00000079528 | ilk | 0.003451532 | 0.338564211 | 0.350049719 | 0.346485507 | 0.032055843 | midline crossing |
| ENSDART00000079563 | fas | 0.029614425 | 0.786193523 | 0.954741877 | 1.62170637 | 0.201092535 | midline crossing |
| ENSDART00000079803 | nmt1b | 0.044608572 | 0.899927422 | 1.156945789 | 1.408075075 | 0.459974747 | midline crossing |
| ENSDART00000080342 | josd2 | 0.005001984 | 0.375878923 | 0.743043487 | 0.777511515 | 0.447917768 | midline crossing |

| | | | | | | | |
|---|---|---|---|---|---|---|---|
| ENSDART00000080351 | dhx57 | 0.034302368 | 0.296631931 | 0.310315824 | 0.343123119 | 0.20162921 | midline crossing |
| ENSDART00000080664 | zgc:86709 | 0.020123282 | 1.428698865 | 3.355720442 | 2.498933956 | 1.732893579 | midline crossing |
| ENSDART00000080829 | hspa14 | 0.003158234 | 0.672662829 | 0.694717457 | 0.8195912 | 0.418167344 | midline crossing |
| ENSDART00000080900 | cfap57 | 0.015156186 | 2.52486974 | 2.757508228 | 2.941219811 | 1.994632033 | midline crossing |
| ENSDART00000081338 | slc9a5 | 0.000216813 | 0.890559011 | 1.045395859 | 1.329125407 | 1.036960607 | midline crossing |
| ENSDART00000081468 | ccdc79 | 0.001875423 | 0.331555315 | 0.505423262 | 0.318137438 | -0.103372006 | midline crossing |
| ENSDART00000081646 | glrx | 0.001482107 | 0.838457918 | 1.280146822 | 1.066830025 | 0.395822945 | midline crossing |
| ENSDART00000081832 | ptpdc1b | 0.000675011 | 0.49240902 | 0.580136174 | 0.651643292 | 0.340422756 | midline crossing |
| ENSDART00000081966 | rtn4a | 0.011135713 | 0.42137623 | 0.811004738 | 0.55995797 | 0.082545275 | midline crossing |
| ENSDART00000082082 | gars | 0.009491292 | 0.395042441 | 0.664200633 | 0.717176297 | 0.392291214 | midline crossing |
| ENSDART00000082223 | tax1bp3 | 0.002818463 | 0.85130321 | 1.256170012 | 1.075265217 | 0.22088107 | midline crossing |
| ENSDART00000082264 | pxdc1b | 0.000212316 | 0.801924492 | 1.033443039 | 0.634881962 | 0.139425975 | midline crossing |
| ENSDART00000082434 | tgif1 | 1.75467E-05 | 0.822910382 | 1.229522702 | 1.441745222 | 0.912580325 | midline crossing |
| ENSDART00000082715 | camsap3 | 0.001948843 | 0.302702799 | 0.347512275 | 0.333666179 | -0.022433363 | midline crossing |
| ENSDART00000082830 | KIAA0895L | 0.000166256 | 0.216380627 | 0.852873359 | 0.835551488 | 0.401334264 | midline crossing |
| ENSDART00000082983 | clip2 | 0.000480106 | 0.69105479 | 0.899482735 | 1.166942455 | 0.710683744 | midline crossing |
| ENSDART00000083467 | parp8 | 0.025639797 | 0.336619688 | 0.399001121 | 0.399118225 | 0.307424215 | midline crossing |
| ENSDART00000083890 | usp24 | 0.033976169 | 0.219267617 | 0.201082864 | 0.325420731 | 0.216599954 | midline crossing |
| ENSDART00000084184 | aimp1 | 0.018217202 | 0.209084138 | 0.262090446 | 0.396284966 | 0.096096937 | midline crossing |
| ENSDART00000084373 | frmd4bb | 0.000871881 | 0.582631667 | 0.828834738 | 0.835563839 | 0.412330382 | midline crossing |
| ENSDART00000084771 | pde9a | 0.000212316 | 0.836076081 | 1.443202889 | 1.33955445 | 1.070879094 | midline crossing |
| ENSDART00000085135 | tbl1x | 0.004493561 | 0.180884774 | 0.566948084 | 0.5040067 | 0.049402086 | midline crossing |
| ENSDART00000085528 | zgc:158659 | 0.002794034 | 0.601601674 | 0.738184978 | 0.82446023 | 0.805817257 | midline crossing |
| ENSDART00000085894 | pgm5 | 0.000309757 | 1.697021259 | 1.783911903 | 2.232651815 | 1.757015772 | midline crossing |
| ENSDART00000087991 | fndc3bb | 0.000250511 | 0.509196457 | 0.583942244 | 0.486760061 | 0.158477497 | midline crossing |
| ENSDART00000088141 | ankrd34bb | 7.10385E-05 | 1.379198636 | 2.420328938 | 2.155206475 | 1.17838067 | midline crossing |
| ENSDART00000088290 | raph1b | 0.018433591 | 0.812305175 | 0.733830465 | 0.671238907 | 0.330964135 | midline crossing |
| ENSDART00000090174 | dock9b | 0.004106215 | 0.356010204 | 0.810416379 | 0.943578275 | 0.58912554 | midline crossing |
| ENSDART00000090406 | dock11 | 0.003111385 | 0.704689621 | 0.535408523 | 0.615472635 | 0.363574103 | midline crossing |
| ENSDART00000090771 | cyth1a | 0.00507534 | 1.102348748 | 1.012723243 | 0.839881021 | 0.442796833 | midline crossing |
| ENSDART00000091140 | snx21 | 0.036614379 | 0.278697916 | 0.637246869 | 0.128859029 | 0.534229621 | midline crossing |
| ENSDART00000091409 | smarcad1a | 0.010885977 | 0.370815487 | 0.374702329 | 0.447117461 | 0.290143851 | midline crossing |
| ENSDART00000092884 | lrrc58b | 0.002894799 | 0.353547699 | 0.596933578 | 0.36096561 | 0.06120599 | midline crossing |
| ENSDART00000097176 | CABZ01090749.1 | 0.000169264 | 1.277443607 | 2.06888261 | 1.727457254 | 0.676889483 | midline crossing |
| ENSDART00000097194 | serinc5 | 0.057511608 | 0.184853776 | 0.289193819 | 0.374387724 | 0.244648582 | midline crossing |
| ENSDART00000098285 | atf5a | 0.018496153 | 0.558593719 | 0.754458044 | 0.445954377 | 0.160619888 | midline crossing |
| ENSDART00000098750 | pdlim5b | 0.005948998 | 0.340621907 | 0.643165703 | 0.410131138 | 0.075261902 | midline crossing |
| ENSDART00000099102 | sept5a | 0.003118176 | 0.488119217 | 0.804836681 | 0.729175054 | 0.142689277 | midline crossing |
| ENSDART00000099248 | rabggtb | 0.029966068 | 0.075526469 | 0.438084725 | 0.330428907 | 0.205346366 | midline crossing |
| ENSDART00000099566 | si:ch211-244o22.2 | 0.032262181 | 0.576141582 | 0.625409237 | 0.697152634 | 0.270183175 | midline crossing |
| ENSDART00000099690 | fam129ab | 0.000806652 | 1.024572276 | 1.489725431 | 1.470862577 | 0.737761428 | midline crossing |
| ENSDART00000099764 | zgc:153031 | 0.034733648 | 0.216426658 | 0.538240257 | 0.500334626 | 0.476962119 | midline crossing |
| ENSDART00000099769 | ccdc22 | 0.012893882 | 0.325453994 | 0.346355629 | 0.367424682 | 0.130188245 | midline crossing |
| ENSDART00000099849 | arntl2 | 9.24337E-05 | 0.982355145 | 1.78511579 | 1.609893892 | 1.255639962 | midline crossing |
| ENSDART00000100156 | agpat4 | 0.000784082 | 1.221504112 | 1.6039826 | 1.49974957 | 0.799345848 | midline crossing |
| ENSDART00000100415 | map3k7cl | 0.000292971 | 1.690116609 | 2.517738065 | 2.168584787 | 1.392466275 | midline crossing |
| ENSDART00000101204 | alcamb | 4.11188E-05 | 1.334306533 | 2.490758317 | 2.513428399 | 1.826617796 | midline crossing |
| ENSDART00000101943 | rragca | 0.024838127 | 0.302260739 | 0.264945177 | 0.40523597 | 0.261316897 | midline crossing |
| ENSDART00000102260 | si:dkey-222f8.3 | 0.01219127 | 0.869989363 | 1.191058069 | 0.864781519 | 0.175715859 | midline crossing |
| ENSDART00000102411 | dctn1b | 0.062774652 | 0.519457248 | 0.529061234 | 0.697567556 | 0.548863629 | midline crossing |
| ENSDART00000102419 | igf2bp2a | 0.009024763 | 0.598022036 | 0.736011973 | 0.87621145 | 0.419702221 | midline crossing |
| ENSDART00000102672 | nck2a | 0.019099714 | 0.205356684 | 0.423877669 | 0.371996615 | 0.147611944 | midline crossing |
| ENSDART00000102843 | src | 0.04997602 | 0.333102086 | 0.263030754 | 0.362117443 | 0.2047457 | midline crossing |
| ENSDART00000102952 | suz12a | 0.000445792 | 0.614499433 | 0.610664915 | 0.72051821 | 0.430761521 | midline crossing |
| ENSDART00000103463 | dnajc2 | 0.048389146 | 0.193851448 | 0.30912042 | 0.441518195 | 0.223806707 | midline crossing |
| ENSDART00000103753 | fn1a | 0.044310242 | 2.705855593 | 3.095500526 | 2.8206804 | 2.414837328 | midline crossing |
| ENSDART00000103755 | fn1b | 0.006568275 | 4.126396153 | 4.640072685 | 4.590004259 | 2.904538972 | midline crossing |
| ENSDART00000103911 | CABZ01077217.1 | 0.031270127 | 0.385992878 | 0.554652123 | 0.464993777 | 0.2375762 | midline crossing |
| ENSDART00000104317 | klf7b | 0.000976528 | 0.389436838 | 1.156291561 | 1.140404424 | 0.652703479 | midline crossing |
| ENSDART00000104616 | lepr | 0.000171958 | 0.762949533 | 0.720319043 | 0.625239528 | 0.526452567 | midline crossing |
| ENSDART00000105399 | cbarpb | 0.001847721 | 0.427437854 | 0.525694666 | 0.646908744 | 0.015511077 | midline crossing |
| ENSDART00000105597 | si:ch211-129c21.1 | 0.007788648 | 3.291282195 | 4.994402871 | 3.669154828 | 0.685912998 | midline crossing |
| ENSDART00000105767 | fhl1a | 0.018500869 | 0.562168934 | 1.496722819 | 1.339494009 | 0.93265064 | midline crossing |
| ENSDART00000105898 | TSTD1 | 0.006964515 | 0.614724746 | 0.98570488 | 0.724640457 | 0.337367855 | midline crossing |
| ENSDART00000106048 | ctnnd1 | 0.001938223 | 0.632970644 | 0.745690204 | 0.779606908 | 0.274224061 | midline crossing |
| ENSDART00000106081 | BX511034.2 | 0.013026885 | 0.124750693 | 0.347355362 | 0.433795514 | 0.117013159 | midline crossing |
| ENSDART00000106172 | rac1a | 0.010965837 | 0.326396929 | 0.503439113 | 0.328932342 | 0.053159748 | midline crossing |
| ENSDART00000108493 | gareml | 0.000703816 | 2.608537205 | 3.376838547 | 3.284344596 | 2.814340641 | midline crossing |
| ENSDART00000108596 | zgc:162150 | 0.006040855 | 0.993292495 | 1.420859127 | 1.257973066 | 0.026357222 | midline crossing |
| ENSDART00000108831 | espnla | 0.018765627 | 0.888146475 | 0.74973908 | 0.792131149 | 0.719500985 | midline crossing |
| ENSDART00000108963 | CABZ01102109.1 | 0.0044585 | 0.394501993 | 0.771748701 | 0.296144389 | 0.075524582 | midline crossing |
| ENSDART00000109138 | hbegfa | 6.40327E-05 | 0.956645779 | 1.991576979 | 1.27153829 | 0.330229075 | midline crossing |
| ENSDART00000109252 | nudt5 | 0.003219301 | 0.602324776 | 0.61383313 | 0.580277921 | 0.203918347 | midline crossing |
| ENSDART00000109319 | arf1 | 0.004134049 | 0.284053831 | 0.377774948 | 0.307132687 | 0.034662092 | midline crossing |
| ENSDART00000109425 | nkd3 | 0.010173969 | 3.023582344 | 2.734865433 | 2.70294964 | 2.317338999 | midline crossing |

| | | | | | | | |
|---|---|---|---|---|---|---|---|
| ENSDART00000109535 | gmps | 0.012765568 | 0.181071124 | 0.257853381 | 0.376587367 | 0.112990813 | midline crossing |
| ENSDART00000109552 | baz2ba | 0.004360669 | 0.627489244 | 0.525669422 | 0.712410774 | 0.393101067 | midline crossing |
| ENSDART00000109573 | akap6 | 0.000144061 | 0.927127599 | 1.241947706 | 1.04133516 | 0.789730231 | midline crossing |
| ENSDART00000109759 | tmx2a | 4.58725E-05 | 2.647169883 | 3.21425121 | 2.659532867 | 1.569734674 | midline crossing |
| ENSDART00000110040 | sox11a | 8.488E-05 | 1.573346229 | 1.723237215 | 1.23195588 | 0.781584101 | midline crossing |
| ENSDART00000110064 | plaua | 0.00106989 | 1.293433748 | 1.730730518 | 1.0162544 | 0.507140212 | midline crossing |
| ENSDART00000110478 | zgc:174906 | 0.002541273 | 0.44828184 | 1.063594358 | 0.857934554 | 0.434989632 | midline crossing |
| ENSDART00000110497 | tmem63a | 0.008300402 | 0.205919412 | 0.504153398 | 0.414725435 | 0.171827856 | midline crossing |
| ENSDART00000110622 | si:ch211-222l21.1 | 0.004840707 | 0.830701135 | 0.950120833 | 0.646058632 | 0.069324807 | midline crossing |
| ENSDART00000110627 | EPB41L1 | 0.000414404 | 0.536765101 | 1.012243023 | 0.739789984 | 0.259906496 | midline crossing |
| ENSDART00000110777 | eef1db | 0.022054847 | 0.19406492 | 0.18930318 | 0.407362874 | -0.014164442 | midline crossing |
| ENSDART00000111324 | zgc:193807 | 0.018752626 | 2.638791453 | 3.489955154 | 3.039693941 | 3.396234279 | midline crossing |
| ENSDART00000111706 | si:dkey-108k21.14 | 0.021895107 | 0.724790653 | 1.05338345 | 1.058888091 | 0.439899526 | midline crossing |
| ENSDART00000111707 | cacnb1 | 0.000480969 | 1.693120335 | 2.349827744 | 2.415610432 | 1.652036859 | midline crossing |
| ENSDART00000111923 | ajuba | 0.004299513 | 0.639088613 | 1.058968297 | 0.866425943 | 0.103913512 | midline crossing |
| ENSDART00000111966 | arhgef10lb | 0.003104562 | 0.250651161 | 0.391053857 | 0.470093443 | 0.290981355 | midline crossing |
| ENSDART00000112028 | si:ch211-180f4.1 | 1.23761E-05 | 2.108953042 | 1.952679055 | 1.909638244 | 1.352861352 | midline crossing |
| ENSDART00000112550 | mapk9 | 0.000730604 | 0.429625805 | 0.878911558 | 0.999507915 | 0.700660398 | midline crossing |
| ENSDART00000112698 | RNF14 (1 of many) | 0.001654051 | 0.142790619 | 0.542852416 | 0.218671557 | 0.088301354 | midline crossing |
| ENSDART00000112926 | adora1b | 0.026842776 | 0.387978003 | 0.609753862 | 0.576471225 | 0.261995134 | midline crossing |
| ENSDART00000113101 | smarcad1b | 0.003021903 | 0.481835623 | 0.651517984 | 0.685097173 | 0.491069329 | midline crossing |
| ENSDART00000113241 | tmem163a | 0.011023029 | 0.329551441 | 0.273820345 | 0.385569684 | 0.333416277 | midline crossing |
| ENSDART00000113532 | CABZ01053588.1 | 8.39504E-05 | 0.66169386 | 1.399901635 | 1.305166578 | 0.721799042 | midline crossing |
| ENSDART00000113699 | hgfb | 0.002024233 | 1.149006949 | 1.556415854 | 1.569424226 | 0.45979101 | midline crossing |
| ENSDART00000113732 | orc6 | 0.018651417 | 1.104098282 | 1.496929933 | 0.974585154 | 0.388395497 | midline crossing |
| ENSDART00000113752 | si:dkey-6i22.5 | 0.014624499 | 1.012202134 | 1.421252013 | 0.919204178 | 0.319242336 | midline crossing |
| ENSDART00000114024 | fam107b | 0.002708716 | 0.899175294 | 1.079533729 | 0.969523488 | 0.434748246 | midline crossing |
| ENSDART00000114168 | itga6a | 0.000135341 | 1.998266436 | 2.016280588 | 1.463165074 | 0.79739173 | midline crossing |
| ENSDART00000114272 | nop2 | 0.022328328 | 0.471489342 | 0.531037709 | 0.420888693 | 0.177047326 | midline crossing |
| ENSDART00000114497 | si:dkey-204l11.1 | 0.008595135 | 0.312589581 | 0.828199841 | 0.99726607 | 0.068811015 | midline crossing |
| ENSDART00000115244 | MEX3A | 8.28889E-05 | 1.222333965 | 1.709642669 | 1.900449694 | 1.354844663 | midline crossing |
| ENSDART00000115403 | NAV1 (1 of many) | 8.51245E-05 | 0.648363377 | 0.712988904 | 0.734499876 | 0.497918559 | midline crossing |
| ENSDART00000118384 | SNORA16 | 0.046563471 | 0.565934898 | 0.9072012 | 0.499756229 | 0.494901316 | midline crossing |
| ENSDART00000121861 | prph | 0.000935288 | 1.238160677 | 2.697856064 | 2.616616233 | 2.213149799 | midline crossing |
| ENSDART00000121886 | hdr | 0.013310088 | 3.187835864 | 3.220765325 | 3.322516101 | 2.611876044 | midline crossing |
| ENSDART00000122102 | wee1 | 1.91914E-05 | 1.082199278 | 1.291191801 | 0.972164457 | 0.241130696 | midline crossing |
| ENSDART00000122305 | Metazoa_SRP | 0.002202211 | 1.452964427 | 1.874874084 | 1.874716847 | 1.224958179 | midline crossing |
| ENSDART00000122433 | trim33l | 0.029170799 | 0.38510745 | 0.766516023 | 0.671400212 | 0.185347257 | midline crossing |
| ENSDART00000122628 | junba | 0.016136147 | 0.280813702 | 0.583374691 | 0.41181471 | 0.073998168 | midline crossing |
| ENSDART00000123096 | marcksa | 1.28435E-05 | 1.85087225 | 2.605359702 | 2.620602183 | 2.013819083 | midline crossing |
| ENSDART00000123203 | vbp1 | 0.026626189 | 0.24089733 | 0.440645383 | 0.41864886 | 0.239214051 | midline crossing |
| ENSDART00000123254 | dpp9 | 0.014674041 | 0.199548032 | 0.283576523 | 0.332661933 | 0.196698648 | midline crossing |
| ENSDART00000123263 | phf20a | 0.000701449 | 3.204242437 | 4.684552705 | 5.028378766 | 3.758919167 | midline crossing |
| ENSDART00000123409 | si:ch1073-296d18.1 | 0.016381291 | 0.174260872 | 0.155958113 | 0.452524339 | -0.02334577 | midline crossing |
| ENSDART00000123559 | znf521 | 0.000648976 | 1.710422257 | 2.255410725 | 1.850379402 | 0.814044199 | midline crossing |
| ENSDART00000123878 | ppiaa | 0.001021091 | 0.40248836 | 0.711363911 | 0.687992513 | 0.279031014 | midline crossing |
| ENSDART00000123972 | smarca5 | 0.039476911 | 0.189313952 | 0.223635359 | 0.34449924 | 0.204975788 | midline crossing |
| ENSDART00000124660 | ascc3 | 0.010970596 | 0.285037378 | 0.269547515 | 0.334772964 | 0.241795126 | midline crossing |
| ENSDART00000124809 | acsbg2 | 4.91848E-05 | 0.494562304 | 1.915512609 | 1.979085784 | 0.982887956 | midline crossing |
| ENSDART00000124991 | lmcd1 | 0.002680527 | 2.103120162 | 2.716646733 | 2.223480288 | 1.685964001 | midline crossing |
| ENSDART00000124998 | rtn2a | 0.001225897 | 0.520408534 | 1.345034307 | 1.147677305 | 0.715797789 | midline crossing |
| ENSDART00000125349 | bada | 0.001185711 | 0.528966306 | 1.266997252 | 1.043776037 | 0.694034948 | midline crossing |
| ENSDART00000125371 | mknk1 | 0.001093301 | 0.126961076 | 0.659607681 | 0.524109906 | -0.03336174 | midline crossing |
| ENSDART00000125971 | gpc1b | 0.020961781 | 0.246742539 | 0.417837609 | 0.415208578 | 0.061504531 | midline crossing |
| ENSDART00000126038 | tcp1 | 0.031160253 | 0.327351769 | 0.30721299 | 0.45795341 | 0.229539742 | midline crossing |
| ENSDART00000126041 | ano10a | 0.016565752 | 0.295415156 | 0.394412522 | 0.244334431 | 0.10037004 | midline crossing |
| ENSDART00000126260 | anxa6 | 0.059606535 | 0.678382414 | 0.731151807 | 0.839317026 | 0.481712468 | midline crossing |
| ENSDART00000126414 | eml1 | 0.019102956 | 0.475872706 | 0.625227433 | 0.675164018 | 0.217118522 | midline crossing |
| ENSDART00000126474 | si:ch211-265o23.1 | 0.039039737 | 1.117262816 | 1.407297939 | 1.180818753 | 0.875131455 | midline crossing |
| ENSDART00000126866 | ccser2b | 0.025259321 | 0.139738782 | 0.390945754 | 0.269322283 | 0.164198289 | midline crossing |
| ENSDART00000127420 | MDP1 | 3.63088E-06 | 1.927039626 | 2.737884833 | 2.522148059 | 1.176014094 | midline crossing |
| ENSDART00000127695 | rpl5b | 0.003761148 | 0.469645013 | 0.524119827 | 0.418266352 | 0.096930554 | midline crossing |
| ENSDART00000127699 | si:ch73-335l21.4 | 0.002818463 | 1.682684641 | 2.104012275 | 1.803809975 | 0.730936452 | midline crossing |
| ENSDART00000127977 | CABZ01016547.1 | 0.016879359 | 2.8163175 | 2.999575494 | 2.826245437 | 2.227407522 | midline crossing |
| ENSDART00000128050 | UTP14C | 0.001174125 | 0.741190051 | 0.667056074 | 0.681031837 | 0.432356782 | midline crossing |
| ENSDART00000128284 | nudt9 | 0.002466999 | 0.256265084 | 0.958235099 | 0.875663921 | 0.386996856 | midline crossing |
| ENSDART00000128346 | si:ch211-108c6.2 | 0.021500865 | 1.285191812 | 1.464423055 | 1.491945601 | 0.337219819 | midline crossing |
| ENSDART00000128624 | sgk494a | 0.001650308 | 0.606573335 | 0.92915728 | 0.662849036 | 0.240608511 | midline crossing |
| ENSDART00000128715 | kif3cb | 0.000188131 | 0.538226215 | 1.727920879 | 1.708926367 | 1.162753729 | midline crossing |
| ENSDART00000128894 | dchs1b | 0.040504381 | 0.298598944 | 0.508598812 | 0.623831745 | 0.178131833 | midline crossing |
| ENSDART00000128931 | igf2bp2a | 0.017671888 | 1.122085018 | 1.168437537 | 1.475101165 | 0.642256664 | midline crossing |
| ENSDART00000129362 | eef1a1l2 | 0.00039492 | 2.651521704 | 2.343527866 | 2.861864827 | 2.05813724 | midline crossing |
| ENSDART00000129423 | slc37a1 | 0.001367004 | 0.300742512 | 0.561138843 | 0.611837666 | 0.153755959 | midline crossing |
| ENSDART00000129593 | znf217 | 0.055515409 | 0.260126556 | 0.54491492 | 0.40143457 | 0.121556273 | midline crossing |
| ENSDART00000129674 | slc44a2 | 0.015120147 | 0.253178883 | 0.658899779 | 0.831363022 | 0.428394942 | midline crossing |

| | | | | | | | |
|---|---|---|---|---|---|---|---|
| ENSDART00000129749 | psmb7 | 0.002585852 | 0.370284184 | 0.518350984 | 0.484691258 | 0.174219127 | midline crossing |
| ENSDART00000129772 | gck | 0.004578126 | 2.162290839 | 3.39609934 | 3.505078852 | 1.699355805 | midline crossing |
| ENSDART00000129895 | palm1b | 0.013143068 | 0.197850318 | 0.594118975 | 0.532496272 | 0.226704229 | midline crossing |
| ENSDART00000129975 | scamp4 | 0.001296944 | 0.678060482 | 0.570045099 | 0.569688168 | 0.357822663 | midline crossing |
| ENSDART00000130277 | rpl9 | 0.000286897 | 0.669628298 | 0.786807801 | 0.580702605 | 0.163431583 | midline crossing |
| ENSDART00000130343 | parp12a | 0.011523209 | 0.340672377 | 0.108363497 | 1.07497332 | -0.29758817 | midline crossing |
| ENSDART00000130353 | fdps | 0.049349672 | 0.408185548 | 0.643895342 | 0.766681528 | 0.202680939 | midline crossing |
| ENSDART00000130472 | pkhd1l1 | 0.064070454 | 0.896390148 | 0.750907757 | 1.185805195 | 0.60737198 | midline crossing |
| ENSDART00000130626 | rars | 0.001161091 | 0.486608727 | 0.578942242 | 0.46060205 | 0.253410521 | midline crossing |
| ENSDART00000130675 | pi4k2a | 0.002785677 | 0.32642219 | 0.46314623 | 0.446740681 | 0.229275132 | midline crossing |
| ENSDART00000130815 | e2f3 | 0.079057004 | 0.222958128 | 0.350714695 | 0.469146338 | 0.300980509 | midline crossing |
| ENSDART00000130820 | hbegfb | 0.000370918 | 1.26577708 | 2.42087461 | 1.934889729 | 0.952474054 | midline crossing |
| ENSDART00000131768 | abi1a | 0.011319743 | 0.352423076 | 0.342648792 | 0.429769283 | 0.194498692 | midline crossing |
| ENSDART00000131861 | tmsb2 | 0.01863352 | 0.228394728 | 0.766531981 | 0.776266217 | 0.553413207 | midline crossing |
| ENSDART00000131922 | ucp3 | 0.045293752 | 0.966573752 | 1.664449524 | 1.416927563 | 1.012896527 | midline crossing |
| ENSDART00000131940 | si:dkey-30c15.10 | 0.000543419 | 1.656397572 | 1.745533662 | 1.503033451 | 0.676482739 | midline crossing |
| ENSDART00000131956 | si:dkey-33c12.14 | 0.026803174 | 2.548697019 | 3.072973071 | 2.718694487 | 2.992696235 | midline crossing |
| ENSDART00000132146 | hsd17b7 | 0.043380755 | 2.880332402 | 3.535090877 | 2.510243514 | 2.078786507 | midline crossing |
| ENSDART00000132156 | cirbpb | 0.03016157 | 0.302166347 | 0.411233461 | 0.214244406 | 0.367929369 | midline crossing |
| ENSDART00000132297 | adam9 | 0.003234084 | 0.312303854 | 0.492550355 | 0.565368245 | 0.224580764 | midline crossing |
| ENSDART00000132386 | si:dkey-21e2.15 | 0.014410292 | 2.037905679 | 4.749110822 | 3.364621679 | 1.867440085 | midline crossing |
| ENSDART00000132426 | stmn4 | 0.000135804 | 0.67103814 | 1.650930846 | 1.637640616 | 1.211182541 | midline crossing |
| ENSDART00000132437 | mknk2a | 0.012665885 | 1.021696261 | 1.216138852 | 0.746540318 | 1.080690265 | midline crossing |
| ENSDART00000132663 | si:ch211-163b2.4 | 0.00520796 | 0.347936584 | 0.995221531 | 0.769857122 | 0.372703166 | midline crossing |
| ENSDART00000132696 | nckap1 | 0.003535619 | 0.516973126 | 0.759090684 | 0.845846182 | 0.627500278 | midline crossing |
| ENSDART00000132781 | si:dkey-78l4.1 | 0.00038013 | 3.429067431 | 4.682405989 | 3.946041907 | 2.798505508 | midline crossing |
| ENSDART00000132788 | uap1 | 0.011788375 | 1.271261725 | 1.549537095 | 1.061485612 | 0.525207108 | midline crossing |
| ENSDART00000132982 | leng9 | 0.002204563 | 3.031473639 | 2.780356944 | 2.241298703 | 1.634482288 | midline crossing |
| ENSDART00000133030 | psmd1 | 0.054998036 | 3.699340083 | 3.88833786 | 2.358920441 | 3.148062929 | midline crossing |
| ENSDART00000133183 | parp8 | 0.045069362 | 0.666223987 | 0.351288157 | 0.707539986 | 0.438871753 | midline crossing |
| ENSDART00000133497 | si:dkey-250k15.4 | 0.028695396 | 0.495799772 | 1.080770342 | 1.149092422 | 0.612727116 | midline crossing |
| ENSDART00000133628 | ankib1a | 0.004447566 | 0.478883806 | 0.425404685 | 0.598914996 | 0.306729273 | midline crossing |
| ENSDART00000133708 | arhgef4 | 0.03479782 | 0.196005293 | 0.61534617 | 0.621912538 | 0.36146972 | midline crossing |
| ENSDART00000133775 | pdlim3b | 0.031565534 | 2.202802348 | 2.787120555 | 2.295385553 | 1.784176005 | midline crossing |
| ENSDART00000133954 | mhc1uka | 4.64299E-05 | 4.819339023 | 5.453978456 | 5.536236268 | 3.849327613 | midline crossing |
| ENSDART00000133968 | camkmt | 0.014401489 | 0.353110068 | 0.455754701 | 0.544139309 | 0.380701512 | midline crossing |
| ENSDART00000134094 | zgc:162193 | 0.04742464 | 0.951212203 | 1.108680472 | 1.240038838 | 0.640044887 | midline crossing |
| ENSDART00000134269 | fkbp1aa | 0.020289809 | 0.162085973 | 0.545601006 | 0.44355879 | 0.374542557 | midline crossing |
| ENSDART00000134658 | slc44a1a | 0.029185454 | 0.707497042 | 0.591443803 | 0.629657706 | 0.651062281 | midline crossing |
| ENSDART00000134787 | elavl4 | 0.000177237 | 1.446773831 | 1.526430349 | 1.657041106 | 1.070021064 | midline crossing |
| ENSDART00000134843 | sst6 | 0.005534146 | 1.638903894 | 2.735989277 | 2.458183626 | 2.301951228 | midline crossing |
| ENSDART00000135093 | ola1 | 0.004885993 | 0.415909116 | 0.691871408 | 0.432580712 | -0.033470782 | midline crossing |
| ENSDART00000135356 | nabp1a | 0.057008548 | 0.371235634 | 0.392313666 | 0.26759026 | 0.229305948 | midline crossing |
| ENSDART00000135415 | dysf | 0.021103936 | 3.146286589 | 3.204629318 | 3.124686608 | 3.126558821 | midline crossing |
| ENSDART00000135631 | myl6 | 0.006911648 | 0.462877699 | 0.806124657 | 0.938068157 | 0.529055927 | midline crossing |
| ENSDART00000135661 | slc20a1a | 0.021002038 | 1.716116201 | 1.644628019 | 1.747440585 | 0.991048186 | midline crossing |
| ENSDART00000135973 | nfyal | 0.007974988 | 0.202128253 | 0.401502491 | 0.256237772 | 0.346886156 | midline crossing |
| ENSDART00000136152 | calua | 0.009163406 | 0.347598327 | 0.562237059 | 0.505429013 | 0.302262295 | midline crossing |
| ENSDART00000136182 | si:ch211-222n4.2 | 0.000461528 | 2.334818577 | 2.498262776 | 2.056155354 | 0.938824321 | midline crossing |
| ENSDART00000136187 | bloc1s4 | 0.017000306 | 0.27836522 | 0.479128543 | 0.397988078 | 0.159673351 | midline crossing |
| ENSDART00000136296 | CR847543.2 | 0.012261156 | 0.7189738 | 0.800339311 | 1.432214548 | 0.183639146 | midline crossing |
| ENSDART00000136492 | si:ch211-158d24.4 | 0.020892176 | 0.620322497 | 0.916930916 | 0.610223614 | 0.170743477 | midline crossing |
| ENSDART00000136761 | prkx | 0.002147221 | 0.698491961 | 1.03323314 | 0.943847516 | 0.577218336 | midline crossing |
| ENSDART00000136782 | si:dkey-24c2.7 | 3.29362E-06 | 4.902793471 | 4.654641885 | 4.451812012 | 3.610899339 | midline crossing |
| ENSDART00000136830 | stmn4l | 0.000324566 | 1.323269884 | 1.682347243 | 1.639725976 | 1.096832109 | midline crossing |
| ENSDART00000136873 | clasp2 | 0.012069725 | 0.301439566 | 0.245236713 | 0.309865242 | 0.356365378 | midline crossing |
| ENSDART00000136910 | armc1l | 0.0517318 | 0.626427947 | 0.775572599 | 0.634581641 | 0.292392529 | midline crossing |
| ENSDART00000136943 | si:dkey-21e2.13 | 0.047530035 | 2.273396315 | 3.247336732 | 1.860650754 | 0.467276759 | midline crossing |
| ENSDART00000137036 | ppiaa | 0.007708544 | 0.323927062 | 0.685997539 | 0.775560135 | 0.372712842 | midline crossing |
| ENSDART00000137110 | sepw1 | 0.028322957 | 0.165066135 | 0.642417047 | 0.502149684 | 0.242826446 | midline crossing |
| ENSDART00000137309 | gemin5 | 0.01891028 | 0.29381973 | 0.279160353 | 0.398962684 | 0.210137956 | midline crossing |
| ENSDART00000137325 | mvp | 0.001650308 | 1.415505945 | 1.485945809 | 1.461661333 | 0.448751577 | midline crossing |
| ENSDART00000137471 | MFAP4 (1 of many) | 0.067929911 | 1.677720699 | 2.807166131 | 2.128226179 | 1.146528193 | midline crossing |
| ENSDART00000137782 | yif1a | 0.072487837 | 0.379643142 | 0.667454267 | 0.241343175 | 0.280143891 | midline crossing |
| ENSDART00000137795 | ssrp1a | 0.014296946 | 0.416335187 | 0.470997147 | 0.449691903 | 0.161896374 | midline crossing |
| ENSDART00000138094 | eef1b2 | 0.002299086 | 0.3943229 | 0.586163757 | 0.433468435 | 0.083803882 | midline crossing |
| ENSDART00000138143 | si:ch211-233m11.1 | 0.012369008 | 0.782546411 | 0.773895723 | 0.974137465 | 0.285762089 | midline crossing |
| ENSDART00000138350 | rps12 | 0.000740604 | 0.592717981 | 0.744811164 | 0.572850481 | 0.279364485 | midline crossing |
| ENSDART00000138351 | cth | 0.000144591 | 1.012814265 | 1.438428325 | 1.066275824 | 0.495805428 | midline crossing |
| ENSDART00000138606 | las1l | 0.028216825 | 0.170918093 | 0.302723725 | 0.361920168 | 0.174466647 | midline crossing |
| ENSDART00000138621 | zgc:153867 | 0.004376172 | 3.188427096 | 3.682050782 | 3.111570491 | 2.809604178 | midline crossing |
| ENSDART00000138633 | cd99 | 0.001871569 | 1.201301736 | 1.818711588 | 1.295470825 | 0.581149232 | midline crossing |
| ENSDART00000138821 | nfil3 | 0.000101984 | 1.376309909 | 2.121391468 | 1.99718084 | 1.39511595 | midline crossing |
| ENSDART00000138915 | cirbpb | 0.015289564 | 0.297824374 | 0.314736859 | 0.24846932 | 0.269446209 | midline crossing |
| ENSDART00000139002 | dus2 | 0.040333038 | 0.373595906 | 0.357863942 | 0.47349873 | 0.230284081 | midline crossing |

| | | | | | | | |
|---|---|---|---|---|---|---|---|
| ENSDART00000139176 | eif2s3 | 0.02245917 | 0.381558608 | 0.49753559 | 0.407153714 | 0.243011405 | midline crossing |
| ENSDART00000139241 | rnf38 | 0.0044585 | 0.363207367 | 0.385948281 | 0.537494883 | 0.39766179 | midline crossing |
| ENSDART00000139259 | zfand5b | 0.022389941 | 0.677389122 | 0.588881516 | 0.462655957 | 0.625250498 | midline crossing |
| ENSDART00000139392 | rras | 0.003337326 | 2.928045636 | 3.065800127 | 2.84735064 | 1.624630606 | midline crossing |
| ENSDART00000139412 | SMIM18 | 0.035424549 | 0.730335352 | 0.956017708 | 0.99931328 | 0.671209714 | midline crossing |
| ENSDART00000139449 | si:dkey-147f3.4 | 0.000116387 | 1.865073313 | 2.098559635 | 1.808022762 | 0.571602171 | midline crossing |
| ENSDART00000139454 | pde4cb | 0.055423595 | 0.616688899 | 0.67005432 | 0.986689732 | 0.473877974 | midline crossing |
| ENSDART00000139479 | slc12a9 | 0.004483567 | 0.435405363 | 0.483188456 | 0.53485188 | 0.300178427 | midline crossing |
| ENSDART00000139722 | dmtn | 2.54845E-05 | 1.105433633 | 1.577455563 | 1.872515003 | 1.217364091 | midline crossing |
| ENSDART00000139763 | stat5a | 0.018355029 | 3.442434617 | 3.964844196 | 3.417693165 | 3.322184683 | midline crossing |
| ENSDART00000139892 | elfn2a | 0.064943452 | 0.520234542 | 0.439147218 | 0.642970325 | 0.553931258 | midline crossing |
| ENSDART00000139947 | rapgef1a | 0.043449081 | 0.201307435 | 0.3482683 | 0.17911742 | 0.096155984 | midline crossing |
| ENSDART00000140032 | si:dkeyp-69c1.9 | 0.086002228 | 1.662700986 | 3.288206015 | 1.780235269 | 1.584891446 | midline crossing |
| ENSDART00000140153 | mri1 | 0.000415146 | 1.105039291 | 1.085223643 | 1.409079861 | 0.961058569 | midline crossing |
| ENSDART00000140253 | arhgef10lb | 0.006717057 | 0.447026855 | 0.470705895 | 0.502550702 | 0.280062014 | midline crossing |
| ENSDART00000140490 | MFAP4 (1 of many) | 0.010636065 | 1.29105804 | 2.228445007 | 1.795046858 | 1.318796638 | midline crossing |
| ENSDART00000140537 | dhrs11a | 0.010679495 | 0.251388963 | 0.647205288 | 0.714640117 | 0.4437028 | midline crossing |
| ENSDART00000140736 | si:dkey-42i9.6 | 0.00486688 | 0.573524661 | 0.712949488 | 0.985257189 | 0.193370375 | midline crossing |
| ENSDART00000141068 | sox11b | 8.488E-05 | 1.611838536 | 2.177304965 | 2.010624009 | 1.194765222 | midline crossing |
| ENSDART00000141107 | cd99 | 0.005438298 | 2.277092862 | 3.207492883 | 2.696592348 | 2.108764519 | midline crossing |
| ENSDART00000141173 | elavl4 | 0.000104621 | 2.4638469 | 2.569363943 | 2.090923257 | 1.273250901 | midline crossing |
| ENSDART00000141328 | pip5kl1 | 0.027686308 | 0.719167688 | 0.681103792 | 0.759572826 | 0.613914474 | midline crossing |
| ENSDART00000141382 | rab34b | 0.005807113 | 1.020539104 | 1.398669915 | 1.148932644 | 0.356641036 | midline crossing |
| ENSDART00000141385 | si:dkey-27p18.3 | 0.023714388 | 0.644114707 | 0.569712036 | 1.095314507 | 0.257528776 | midline crossing |
| ENSDART00000141578 | shmt1 | 0.002456687 | 2.367581981 | 4.511004025 | 4.354684857 | -7.25195E-16 | midline crossing |
| ENSDART00000141606 | commd8 | 0.007621679 | 0.496320125 | 0.646713153 | 0.422272129 | 0.302789387 | midline crossing |
| ENSDART00000141625 | si:ch211-156p11.1 | 0.075460344 | 0.900660828 | 1.143855847 | 0.822477469 | 0.791150698 | midline crossing |
| ENSDART00000141837 | esd | 0.031729112 | 0.782720008 | 0.801148933 | 0.794217077 | 0.347191174 | midline crossing |
| ENSDART00000141877 | kcnk10a | 0.000157775 | 1.57307368 | 1.979146889 | 1.661354893 | 0.924431526 | midline crossing |
| ENSDART00000142146 | dmtn | 0.027138478 | 0.637268571 | 0.800533621 | 0.832554824 | 0.713529971 | midline crossing |
| ENSDART00000142356 | smyd2a | 0.012342358 | 2.425657932 | 2.779397529 | 2.178998127 | 1.672270997 | midline crossing |
| ENSDART00000142428 | josd2 | 0.004387455 | 0.235661569 | 0.659033561 | 0.722639048 | 0.359420457 | midline crossing |
| ENSDART00000142569 | specc1lb | 0.004391006 | 0.378697075 | 0.547073638 | 0.617855803 | 0.275260653 | midline crossing |
| ENSDART00000142692 | eef1b2 | 0.008579097 | 0.349947783 | 0.50162657 | 0.39446346 | 0.093426844 | midline crossing |
| ENSDART00000142873 | BX901918.1 | 0.068563448 | 0.845880305 | 0.530715844 | 1.008998061 | 0.591786557 | midline crossing |
| ENSDART00000143056 | zyx | 0.011047999 | 0.416244119 | 0.475600395 | 0.593066786 | 0.15882784 | midline crossing |
| ENSDART00000143457 | cndp2 | 0.016866534 | 0.393197537 | 0.458029638 | 0.473219597 | 0.244423454 | midline crossing |
| ENSDART00000143562 | gbp2 | 0.021682969 | 0.680404258 | 0.927169543 | 1.353668195 | 0.527828388 | midline crossing |
| ENSDART00000143723 | idi1 | 0.030676359 | 0.080932725 | 0.304370095 | 0.372374682 | 0.127047939 | midline crossing |
| ENSDART00000143741 | fkbp5 | 2.85018E-05 | 1.764298053 | 1.865156046 | 1.631976013 | 1.871443577 | midline crossing |
| ENSDART00000143793 | C3 (1 of many) | 0.009758715 | 3.237329668 | 4.023510735 | 3.503827488 | 2.024562274 | midline crossing |
| ENSDART00000144503 | ran | 0.011399481 | -0.075551405 | 0.448221073 | 0.249018928 | 0.118064827 | midline crossing |
| ENSDART00000144516 | lonrf1l | 0.001677265 | 0.003516441 | 0.629397595 | -0.101368349 | 0.117101343 | midline crossing |
| ENSDART00000144814 | si:dkey-31f5.8 | 0.019331363 | 0.429768416 | 1.027387879 | 0.82502917 | 0.369456203 | midline crossing |
| ENSDART00000144817 | cfl1 | 0.008723451 | 0.341949415 | 0.252829572 | 0.49073898 | 0.258667203 | midline crossing |
| ENSDART00000144946 | adcyap1b | 0.016974377 | 2.385755846 | 2.421463563 | 2.507590258 | 1.873754889 | midline crossing |
| ENSDART00000145095 | celsr3 | 0.003711093 | 0.648145926 | 0.648607509 | 0.829081729 | 0.619308918 | midline crossing |
| ENSDART00000145103 | cntfr | 0.071227112 | 0.307964555 | 0.391102809 | 0.626651223 | 0.264745691 | midline crossing |
| ENSDART00000145379 | fblim1 | 0.003095428 | 0.305913455 | 0.545649791 | 0.383397127 | 0.135782412 | midline crossing |
| ENSDART00000145956 | ptpn23a | 0.029718853 | 0.252886396 | 0.192353563 | 0.35477406 | 0.187678789 | midline crossing |
| ENSDART00000145999 | prx | 0.022530244 | 1.586883428 | 3.204392125 | 2.633047079 | 1.399238803 | midline crossing |
| ENSDART00000146005 | gtpbp1 | 0.02305104 | 0.461505187 | 0.53679144 | 0.551879705 | 0.330029037 | midline crossing |
| ENSDART00000146411 | sst6 | 0.000370918 | 0.736642137 | 1.700467827 | 1.549366173 | 1.261691667 | midline crossing |
| ENSDART00000146434 | zgc:153115 | 0.001249772 | 0.499570303 | 0.817313835 | 0.902072926 | 0.441195032 | midline crossing |
| ENSDART00000146611 | cxcr3.3 | 0.009119552 | 0.415816929 | 0.611232638 | 0.663522207 | -0.008061564 | midline crossing |
| ENSDART00000146744 | nfkbiab | 0.003196539 | 0.787570548 | 0.783257317 | 0.468593447 | 0.633863732 | midline crossing |
| ENSDART00000147019 | si:ch211-235o23.1 | 0.001130346 | 0.394717538 | 0.700152429 | 0.448882118 | 0.273599764 | midline crossing |
| ENSDART00000147264 | fam213b | 0.046818285 | 0.418860252 | 0.552271711 | 0.440362956 | 0.212029386 | midline crossing |
| ENSDART00000147518 | fam89b | 0.007332698 | 0.288965136 | 0.576511697 | 0.598868415 | 0.448283091 | midline crossing |
| ENSDART00000147821 | spsb4a | 0.001871569 | 0.371275002 | 0.33311003 | 0.599280348 | 0.314031744 | midline crossing |
| ENSDART00000147849 | klf6a | 0.000870446 | 1.130982604 | 1.870693821 | 1.403514717 | 0.445060502 | midline crossing |
| ENSDART00000147851 | eef1b2 | 0.000349374 | 0.252066551 | 0.849118612 | 0.755935234 | 0.422914839 | midline crossing |
| ENSDART00000148011 | cby1 | 0.052184053 | 0.256002046 | 0.39067073 | 0.214338805 | 0.125714094 | midline crossing |
| ENSDART00000148138 | lrrc8aa | 0.002977128 | 0.370743632 | 0.558987897 | 0.552524282 | 0.096775057 | midline crossing |
| ENSDART00000148154 | stx2a | 0.012799526 | 0.73746977 | 1.316523952 | 0.880682553 | 0.300142462 | midline crossing |
| ENSDART00000148163 | frmd4bb | 0.000286897 | 0.943172358 | 1.400649402 | 1.507648518 | 0.548522517 | midline crossing |
| ENSDART00000148388 | smox | 0.001452556 | 1.14663E-15 | 4.65884204 | 4.212605499 | 2.256263175 | midline crossing |
| ENSDART00000148432 | ptp4a3 | 0.003196078 | 0.27125556 | 0.468426427 | 0.608500351 | -0.013500628 | midline crossing |
| ENSDART00000148601 | insig1 | 0.008603534 | -0.03765091 | 0.839125818 | 0.622258944 | 0.165134159 | midline crossing |
| ENSDART00000148659 | clip2 | 0.000391808 | 0.500760868 | 0.904821279 | 1.156870185 | 0.693646301 | midline crossing |
| ENSDART00000148822 | adgrg1 | 0.015695496 | 0.746405238 | 1.031170771 | 0.764826924 | 0.143644812 | midline crossing |
| ENSDART00000148993 | hdac6 | 0.033606114 | 0.150136767 | 0.327518494 | 0.405976253 | 0.20477563 | midline crossing |
| ENSDART00000149074 | sall1b | 2.18017E-05 | 1.974229239 | 2.883794957 | 2.506155499 | 1.735121972 | midline crossing |
| ENSDART00000149115 | zgc:158659 | 4.64273E-06 | 1.901261478 | 2.561373529 | 3.048076346 | 2.366410956 | midline crossing |
| ENSDART00000149320 | arl8a | 0.011240617 | 0.200543842 | 0.321313695 | 0.387648659 | 0.1981557 | midline crossing |

| | | | | | | | |
|---|---|---|---|---|---|---|---|
| ENSDART00000149910 | abl1 | 0.034395507 | 0.087815124 | 0.252042838 | 0.385819934 | 0.051793038 | midline crossing |
| ENSDART00000149978 | fabp7a | 0.000717715 | 0.33770709 | 0.764013846 | 0.558825405 | 0.488886506 | midline crossing |
| ENSDART00000150013 | cd276 | 1.1419E-05 | 1.481977913 | 1.919249755 | 1.718653495 | 0.833090259 | midline crossing |
| ENSDART00000150149 | tdp1 | 0.042010311 | 0.245251359 | 0.536780821 | 0.276916415 | 0.174119987 | midline crossing |
| ENSDART00000150391 | si:ch211-214j24.14 | 0.000870998 | 2.502222784 | 2.698584631 | 2.367333489 | 0.935046128 | midline crossing |
| ENSDART00000150651 | BX465862.4 | 0.011605032 | 0.896323656 | 0.956165673 | 0.709439713 | 0.378868912 | midline crossing |
| ENSDART00000150692 | NA | 0.001955129 | 0.448027135 | 0.604850915 | 0.68748119 | 0.482541087 | midline crossing |
| ENSDART00000150961 | wdr91 | 0.030491542 | 4.358324282 | 4.633846856 | 4.480446443 | 4.649897846 | midline crossing |
| ENSDART00000151247 | scn1lab | 0.035957782 | 0.293159534 | 0.684635422 | 0.628869811 | 0.299982725 | midline crossing |
| ENSDART00000151448 | vamp4 | 0.002952724 | 0.36274105 | 0.703182259 | 0.708668654 | 0.04883985 | midline crossing |
| ENSDART00000151613 | lhfpl2a | 0.025409101 | 2.211015019 | 2.8976425 | 2.404140419 | 1.864383361 | midline crossing |
| ENSDART00000151679 | si:ch73-233k15.2 | 0.008447718 | 0.305634484 | 0.260016753 | 1.072404438 | -0.272816623 | midline crossing |
| ENSDART00000151785 | klc4 | 0.002952432 | 0.265672724 | 0.701188452 | 0.757625768 | 0.469102543 | midline crossing |
| ENSDART00000151892 | BX248324.2 | 9.23128E-05 | 1.389473856 | 2.137590244 | 1.979774015 | 1.113747327 | midline crossing |
| ENSDART00000151999 | ttc27 | 0.000370918 | 0.537657382 | 0.63628465 | 0.751878141 | 0.471568333 | midline crossing |
| ENSDART00000152240 | si:ch1073-174d20.2 | 0.045564466 | 0.11047675 | 0.254355415 | 0.357239942 | 0.147146143 | midline crossing |
| ENSDART00000152295 | FAM83G | 0.016063659 | 1.150380037 | 1.206829236 | 1.079415548 | 0.591155843 | midline crossing |
| ENSDART00000152318 | si:ch211-286b5.2 | 0.002663982 | 1.976393579 | 3.213645724 | 2.557166791 | 1.560923939 | midline crossing |
| ENSDART00000152510 | ccser2b | 0.001067718 | 0.726161978 | 0.836833026 | 1.118702985 | 0.610749876 | midline crossing |
| ENSDART00000152562 | eif3f | 0.018854609 | 0.257721359 | 0.386526337 | 0.235538886 | 0.038968621 | midline crossing |
| ENSDART00000152641 | dnm3a | 0.000272902 | 1.808565272 | 2.497351665 | 2.011092537 | 0.57584064 | midline crossing |
| ENSDART00000152680 | pyyb | 0.000396404 | 2.62463081 | 4.023644413 | 3.815932945 | 2.663962921 | midline crossing |
| ENSDART00000152855 | prdx1 | 0.045741098 | 2.014399342 | 2.912277881 | 2.532639813 | 2.043057891 | midline crossing |
| ENSDART00000153194 | si:dkey-38p12.3 | 0.015825859 | 0.25264745 | 0.485169484 | 0.572803819 | 0.07332987 | midline crossing |
| ENSDART00000153199 | afap1 | 0.00834805 | 0.840574953 | 1.106233667 | 1.438882155 | 1.005659187 | midline crossing |
| ENSDART00000153200 | si:ch211-194k22.8 | 0.033503027 | 0.210290351 | 0.451170205 | 0.37501933 | 0.225583201 | midline crossing |
| ENSDART00000153225 | GUCY2C (1 of many) | 0.032143116 | 3.255008425 | 3.866902647 | 2.201523204 | 2.781639432 | midline crossing |
| ENSDART00000153227 | nfe2l1b | 0.013323448 | 0.688085235 | 1.301360444 | 1.138008541 | 0.729999096 | midline crossing |
| ENSDART00000153306 | hcar1-4 | 0.004505979 | 0.883818254 | 1.413387657 | 1.160612678 | 0.305432292 | midline crossing |
| ENSDART00000153307 | si:ch73-344o19.1 | 0.006344596 | 0.586530995 | 0.605191095 | 0.75799445 | 0.479323327 | midline crossing |
| ENSDART00000153452 | hsp90ab1 | 0.030379124 | 0.284157203 | 0.639252574 | 0.548907136 | 0.495783326 | midline crossing |
| ENSDART00000153666 | AL732567.1 | 0.002559269 | 0.845331402 | 1.050170733 | 0.86587331 | 0.828206988 | midline crossing |
| ENSDART00000153841 | lmcd1 | 0.012070853 | 1.804357101 | 2.873087613 | 2.523453656 | 2.002290777 | midline crossing |
| ENSDART00000154109 | tp53inp2 | 0.001954005 | 0.254956915 | 0.876540848 | 0.694070175 | 0.684041374 | midline crossing |
| ENSDART00000154212 | BX323824.1 | 0.024957938 | 0.386508588 | 0.636151192 | 0.398585293 | 0.423381311 | midline crossing |
| ENSDART00000154314 | prph | 6.48692E-06 | 1.502464936 | 3.204214666 | 3.264208069 | 2.453842104 | midline crossing |
| ENSDART00000154321 | BX908750.1 | 0.050473914 | 0.337703917 | 0.710327992 | 0.54373662 | 0.188876538 | midline crossing |
| ENSDART00000154377 | HIST1H2BM (1 of many) | 0.051183892 | 0.871205458 | 1.144896653 | 0.89956001 | 0.544514919 | midline crossing |
| ENSDART00000154543 | prelid3a | 0.006027331 | 0.578261204 | 0.829307037 | 0.712329147 | 0.291766611 | midline crossing |
| ENSDART00000154573 | prob1 | 0.001652227 | 1.935993422 | 2.091368151 | 1.752154811 | 1.442205958 | midline crossing |
| ENSDART00000154628 | fam101b | 0.000888975 | 0.450957894 | 0.982590486 | 0.845377619 | -0.490650185 | midline crossing |
| ENSDART00000154846 | CR381618.1 | 0.002605308 | 0.115176903 | 0.900181702 | 1.102625706 | 0.253393379 | midline crossing |
| ENSDART00000154851 | zgc:153405 | 0.027461561 | 0.317950752 | 1.076463864 | 0.674378682 | 0.394781222 | midline crossing |
| ENSDART00000154880 | arhgef33 | 0.01779887 | 0.636994605 | 0.687196577 | 0.646579762 | 0.300121216 | midline crossing |
| ENSDART00000155051 | mych | 0.002016009 | 1.81452306 | 1.847457308 | 1.423042212 | 0.692601562 | midline crossing |
| ENSDART00000155188 | ccnjl | 0.031062271 | 0.284030133 | 0.391162848 | 0.492680219 | 0.242842213 | midline crossing |
| ENSDART00000155256 | VWA5A (1 of many) | 0.021065097 | 0.207030506 | 0.487713239 | 0.641785484 | 0.122484219 | midline crossing |
| ENSDART00000155280 | GDPGP1 | 0.018409732 | 0.576074599 | 0.656458452 | 0.674683139 | 0.299654628 | midline crossing |
| ENSDART00000155440 | eif3ba | 0.00350062 | 0.265595321 | 0.439061322 | 0.461693724 | 0.229141991 | midline crossing |
| ENSDART00000155537 | VWA5A (1 of many) | 0.000220872 | 1.206080865 | 0.987004598 | 1.27402251 | 0.722325665 | midline crossing |
| ENSDART00000155751 | nuak1b | 1.33565E-05 | 0.785416985 | 1.722603254 | 1.683891996 | 1.302083807 | midline crossing |
| ENSDART00000156034 | VWA5A (1 of many) | 8.51245E-05 | 1.04711946 | 1.090403168 | 1.260861512 | 0.694741705 | midline crossing |
| ENSDART00000156038 | epn2 | 0.034236293 | 0.165862475 | 0.434471463 | 0.333526121 | 0.113707323 | midline crossing |
| ENSDART00000156167 | si:ch73-182a11.2 | 0.007880813 | 0.438099394 | 0.613388756 | 0.677790276 | 0.474745988 | midline crossing |
| ENSDART00000156224 | dhx33 | 0.005024347 | 0.472474316 | 0.556602191 | 0.639276654 | 0.492917502 | midline crossing |
| ENSDART00000156251 | ehbp1 | 0.005375255 | 0.342339175 | 0.353753636 | 0.38894242 | 0.04618254 | midline crossing |
| ENSDART00000156255 | ago1 | 0.004702472 | 0.402935641 | 0.576022371 | 0.656640241 | 0.478601065 | midline crossing |
| ENSDART00000156411 | fam98a | 0.014485141 | 0.369288002 | 0.399859986 | 0.477248543 | 0.233728964 | midline crossing |
| ENSDART00000156415 | dock9b | 0.006352299 | 0.187525252 | 0.556466903 | 0.665071238 | 0.341730571 | midline crossing |
| ENSDART00000156774 | dph6 | 0.010921357 | 0.485952337 | 0.860504803 | 0.608054536 | 0.313680056 | midline crossing |
| ENSDART00000156913 | BX927184.1 | 0.000109677 | 1.282107066 | 1.77927465 | 1.802027966 | 1.376990986 | midline crossing |
| ENSDART00000156970 | si:ch73-265h17.5 | 0.019558454 | 1.250341969 | 1.481314496 | 1.76830523 | 0.8438116 | midline crossing |
| ENSDART00000156972 | slc12a7a | 0.016311478 | 0.533547881 | 0.750475895 | 0.543271513 | 0.05077876 | midline crossing |
| ENSDART00000157127 | lmcd1 | 0.008776907 | 1.160892936 | 2.271204052 | 2.063825472 | 1.282095985 | midline crossing |
| ENSDART00000157139 | tmbim1a | 0.004528919 | 0.963253296 | 1.499596205 | 1.306783479 | 0.549513539 | midline crossing |
| ENSDART00000157350 | CT573344.1 | 0.023257205 | 0.592860846 | 1.013860007 | 1.275401288 | 0.850112465 | midline crossing |
| ENSDART00000157441 | filip1l | 0.004552273 | 1.025025125 | 2.361056221 | 1.959563135 | 1.214723721 | midline crossing |
| ENSDART00000157597 | Metazoa_SRP | 0.000305619 | 3.784983496 | 4.068975573 | 4.489534416 | 3.689887905 | midline crossing |
| ENSDART00000157604 | si:dkey-19c16.12 | 0.067756277 | 0.518457933 | 0.406892634 | 0.690003002 | 0.445775315 | midline crossing |
| ENSDART00000157678 | nop53 | 0.020688276 | 0.221649254 | 0.357738319 | 0.319808875 | 0.087032288 | midline crossing |
| ENSDART00000157824 | slc6a15 | 0.001558204 | 0.370043272 | 0.932121448 | 1.025205824 | 0.470213629 | midline crossing |
| ENSDART00000157894 | apaf1 | 0.033808649 | 0.69901776 | 0.613857472 | 0.804406673 | 0.575412751 | midline crossing |
| ENSDART00000157907 | rpl34 | 0.021862757 | 0.216382556 | 0.50724745 | 0.251806906 | 0.167915072 | midline crossing |
| ENSDART00000158016 | flnca | 4.50375E-07 | 5.435651542 | 4.853809922 | 4.737861308 | 3.940836376 | midline crossing |
| ENSDART00000158272 | dclk1b | 0.007797716 | 0.134331666 | 0.528656936 | 0.571389243 | 0.299537528 | midline crossing |

| | | | | | | | |
|---|---|---|---|---|---|---|---|
| ENSDART00000158739 | Metazoa_SRP | 0.027491007 | 1.431390172 | 1.797550169 | 1.401344471 | 0.830250287 | midline crossing |
| ENSDART00000158744 | BX927336.3 | 0.000371354 | 4.021248005 | 3.307131569 | 3.405842268 | 3.528289574 | midline crossing |
| ENSDART00000159024 | btr29 | 0.003508363 | 2.93548853 | 3.916483422 | 3.440636421 | 1.361130093 | midline crossing |
| ENSDART00000159034 | rsu1 | 0.003279742 | 0.959727483 | 0.959427671 | 0.950095712 | 0.424675541 | midline crossing |
| ENSDART00000159256 | kif5c | 0.031850333 | 0.534952361 | 0.603334839 | 0.411520272 | 0.273079865 | midline crossing |
| ENSDART00000159259 | capn1b | 4.91601E-05 | 0.6323058 | 1.402210713 | 1.384902028 | 1.097440242 | midline crossing |
| ENSDART00000159300 | mcm7 | 0.002876227 | 0.714704002 | 0.831682391 | 0.874291792 | 0.339164006 | midline crossing |
| ENSDART00000159340 | ctnnd1 | 0.000135332 | 0.771872624 | 0.982959313 | 0.944642837 | 0.226155974 | midline crossing |
| ENSDART00000159372 | eif3ha | 0.001128218 | 0.499618724 | 0.599836819 | 0.638052337 | 0.263915251 | midline crossing |
| ENSDART00000159652 | sox4b | 0.008897727 | 0.767109202 | 1.080655119 | 1.389017231 | 0.976257694 | midline crossing |
| ENSDART00000159732 | si:dkey-203a12.8 | 0.050696552 | 3.015691266 | 3.637172924 | 3.104587293 | 1.713663998 | midline crossing |
| ENSDART00000159949 | wdr1 | 0.082310414 | 0.362540326 | 0.308247591 | 0.564916756 | 0.170183845 | midline crossing |
| ENSDART00000159965 | si:ch211-255i3.4 | 0.025487547 | 0.121150704 | 0.407179882 | 0.241107117 | 0.125992518 | midline crossing |
| ENSDART00000160006 | Metazoa_SRP | 0.001666219 | 1.639017988 | 2.129447988 | 1.830106321 | 0.946987666 | midline crossing |
| ENSDART00000160206 | rap2b | 0.014591443 | 0.536023906 | 0.628551975 | 0.691733134 | 0.267820563 | midline crossing |
| ENSDART00000160492 | alad | 0.036567119 | 0.30171604 | 0.443307609 | 0.248348623 | 0.158762163 | midline crossing |
| ENSDART00000160575 | CU459186.1 | 0.043586255 | 0.73831154 | 1.064084865 | 0.743119809 | 0.540430362 | midline crossing |
| ENSDART00000160604 | ftsj3 | 0.00161581 | 0.856590556 | 0.695492085 | 0.825371198 | 0.428239045 | midline crossing |
| ENSDART00000160663 | tnk2b | 0.032721257 | 0.180874765 | 0.228576581 | 0.365115453 | 0.173216656 | midline crossing |
| ENSDART00000160766 | pim1 | 0.002635727 | 0.465870139 | 0.649909313 | 0.413306503 | 0.082988542 | midline crossing |
| ENSDART00000160979 | jpt1b | 0.000229447 | 0.682460724 | 1.366219793 | 1.515562314 | 0.876953288 | midline crossing |
| ENSDART00000161017 | eif3s6ip | 0.012456084 | 0.313830903 | 0.293421495 | 0.276651786 | 0.120270233 | midline crossing |
| ENSDART00000161117 | NA | 0.052467991 | 0.194355392 | 0.331000906 | 0.496659366 | 0.20108058 | midline crossing |
| ENSDART00000161244 | CR293509.1 | 5.17084E-05 | 0.407469749 | 1.465267423 | 1.274347742 | 0.975210426 | midline crossing |
| ENSDART00000161330 | col27a1a | 0.012339321 | 0.210670367 | 0.547410859 | 0.646435326 | 0.255665234 | midline crossing |
| ENSDART00000161377 | cdk2ap2 | 0.030614624 | 0.241317874 | 0.341753789 | 0.416424679 | 0.144968334 | midline crossing |
| ENSDART00000161460 | pmt | 0.047763341 | 2.538801298 | 3.7270331 | 2.780124527 | 1.870914111 | midline crossing |
| ENSDART00000161514 | CABZ01079986.1 | 0.000250511 | -0.184749182 | 1.474076268 | 0.73277167 | -0.443845367 | midline crossing |
| ENSDART00000161597 | si:ch211-262i1.3 | 0.034395507 | 0.726810222 | 0.918966234 | 1.284521547 | 0.120004192 | midline crossing |
| ENSDART00000161613 | SLC7A1 (1 of many) | 0.002341037 | 0.777752401 | 1.137604367 | 0.924750387 | 0.450737498 | midline crossing |
| ENSDART00000161869 | dysf | 0.010359104 | 1.077492273 | 1.30463907 | 1.440346007 | 1.017975966 | midline crossing |
| ENSDART00000162007 | Metazoa_SRP | 0.037519793 | 0.903949119 | 2.303055473 | 1.702580761 | 0.513671528 | midline crossing |
| ENSDART00000162377 | FP085414.1 | 0.001772851 | 0.809712454 | 1.275862162 | 1.389373351 | 0.791945088 | midline crossing |
| ENSDART00000162722 | zgc:153867 | 0.001130434 | 0.467799327 | 0.468299246 | 0.579069745 | 0.022677056 | midline crossing |
| ENSDART00000162897 | Metazoa_SRP | 0.0215019 | 1.178659737 | 1.5924133 | 1.384383775 | 1.197556527 | midline crossing |
| ENSDART00000163042 | si:dkey-16p6.2 | 0.048041075 | 0.770521119 | 0.592959463 | 0.681550899 | 0.554585181 | midline crossing |
| ENSDART00000163137 | elovl1a | 0.000445792 | 1.359201374 | 2.32562135 | 1.99000125 | 0.940099891 | midline crossing |
| ENSDART00000163174 | si:ch73-42p12.2 | 6.34429E-05 | 0.635401366 | 0.972834844 | 0.669544171 | 0.150769788 | midline crossing |
| ENSDART00000163243 | dpysl4 | 0.001058776 | 0.526851432 | 0.969802663 | 1.139409914 | 0.808513895 | midline crossing |
| ENSDART00000163367 | TFR2 | 4.96098E-05 | 0.892525381 | 1.842645951 | 2.060867557 | 1.063290775 | midline crossing |
| ENSDART00000163370 | si:rp71-36a1.2 | 0.007804676 | 0.777729217 | 1.232972386 | 1.065047677 | 0.579071639 | midline crossing |
| ENSDART00000163441 | limk1a | 6.80867E-05 | 1.134897491 | 0.940120128 | 1.036684953 | 0.715200814 | midline crossing |
| ENSDART00000164207 | impdh2 | 0.000279675 | 0.582365155 | 0.675083436 | 1.006708284 | 0.417204588 | midline crossing |
| ENSDART00000164210 | SI | 0.000224957 | 1.002064064 | 1.522108486 | 0.992417222 | 0.446482404 | midline crossing |
| ENSDART00000164311 | mfsd2aa | 0.007723957 | 0.188250738 | 0.92622451 | 1.277563791 | 0.357720797 | midline crossing |
| ENSDART00000164392 | lrrc20 | 0.000250511 | 0.934799406 | 1.852088866 | 1.534338919 | 1.07983234 | midline crossing |
| ENSDART00000164471 | CABZ01084230.2 | 0.024883139 | 0.442580477 | 0.587031258 | 0.401618345 | 0.289449509 | midline crossing |
| ENSDART00000164543 | kalrna | 0.002635555 | 0.833553496 | 1.079150473 | 0.963779963 | 0.661385395 | midline crossing |
| ENSDART00000165141 | elavl4 | 0.004826213 | 1.008784658 | 0.753903544 | 1.037731482 | 0.21557706 | midline crossing |
| ENSDART00000165309 | pmt | 0.000144061 | 1.560743592 | 2.051643174 | 1.965781873 | 1.006007351 | midline crossing |
| ENSDART00000165572 | zgc:162193 | 0.029001753 | 0.654407013 | 0.606984165 | 0.583589232 | 0.369009896 | midline crossing |
| ENSDART00000165594 | CT030188.1 | 0.030766044 | 0.949647945 | 1.541943795 | 1.047931527 | 0.898450066 | midline crossing |
| ENSDART00000165628 | il4r.1 | 0.026868362 | 2.521820035 | 2.797213431 | 2.312532052 | 1.674117298 | midline crossing |
| ENSDART00000165715 | BX284638.1 | 0.000261206 | 0.544886732 | 0.81514457 | 1.035521411 | 0.49952411 | midline crossing |
| ENSDART00000165993 | f3a | 0.000100528 | 1.175718098 | 1.946056838 | 1.259834727 | 0.15077318 | midline crossing |
| ENSDART00000166040 | sh3bp5b | 5.24617E-05 | 0.359354612 | 1.105816025 | 1.101214783 | 0.69319763 | midline crossing |
| ENSDART00000166120 | si:ch211-212d10.1 | 0.005272363 | 1.398436882 | 1.472487792 | 1.228480368 | 0.254562361 | midline crossing |
| ENSDART00000166259 | wars | 0.007387753 | 0.30804143 | 0.479087994 | 0.526413933 | 0.245904587 | midline crossing |
| ENSDART00000166324 | ctnnd1 | 0.02267327 | 0.460487451 | 0.641759187 | 1.004918197 | 0.346719852 | midline crossing |
| ENSDART00000166432 | slc2a8 | 0.044776229 | 0.33290564 | 0.367124099 | 0.471066456 | 0.250147846 | midline crossing |
| ENSDART00000166560 | hpdl | 0.029524772 | 0.319918962 | 0.542121436 | 0.616518616 | 0.488982996 | midline crossing |
| ENSDART00000166622 | si:dkey-77g12.1 | 0.030464157 | 1.256692313 | 2.663525112 | 1.501030031 | -0.081810387 | midline crossing |
| ENSDART00000166992 | rab6a | 0.003458178 | 0.863374482 | 0.610837796 | 0.548836905 | 0.594870039 | midline crossing |
| ENSDART00000167035 | mibp2 | 0.000190025 | 0.52185156 | 0.897496666 | 1.068299106 | 0.608631566 | midline crossing |
| ENSDART00000167062 | SNORD49 | 0.030270278 | 0.825973685 | 1.057295844 | 0.696610061 | 0.75374548 | midline crossing |
| ENSDART00000167145 | ap1ar | 0.00427909 | 0.285895558 | 0.368048827 | 0.350688731 | 0.119518374 | midline crossing |
| ENSDART00000167359 | dusp27 | 0.000892328 | 1.665789827 | 1.31173846 | 1.624125812 | 1.13571557 | midline crossing |
| ENSDART00000167461 | BX897691.1 | 0.046057741 | 1.652413387 | 2.158428192 | 1.996862668 | 1.949150718 | midline crossing |
| ENSDART00000167666 | dnajc21 | 0.014542918 | 0.569585132 | 0.384796247 | 0.537599141 | 0.160116436 | midline crossing |
| ENSDART00000167879 | Metazoa_SRP | 0.015884799 | 3.268404605 | 5.790076 | 5.563915837 | 5.137733652 | midline crossing |
| ENSDART00000168083 | Metazoa_SRP | 0.018792355 | 1.497191597 | 1.645034783 | 1.435853831 | 0.615781715 | midline crossing |
| ENSDART00000168371 | mibp | 0.004889638 | 1.064831288 | 1.384792727 | 1.720455753 | 0.84273718 | midline crossing |
| ENSDART00000168376 | Metazoa_SRP | 0.031983637 | 0.77672172 | 1.175457852 | 1.304833035 | 1.047872554 | midline crossing |
| ENSDART00000168419 | rsu1 | 0.004815134 | 0.608707901 | 0.588635147 | 0.668370486 | 0.283061431 | midline crossing |
| ENSDART00000168497 | eef2b | 0.002633576 | 0.448731314 | 0.547292163 | 0.7539331 | 0.364287935 | midline crossing |

| | | | | | | | |
|---|---|---|---|---|---|---|---|
| ENSDART00000168610 | si:ch211-283g2.2 | 0.000871179 | 1.147251625 | 1.274539863 | 1.168500693 | 0.162307024 | midline crossing |
| ENSDART00000168683 | lrrc8c | 0.039756837 | 0.403272924 | 0.771214032 | 0.877257507 | 0.389191376 | midline crossing |
| ENSDART00000168902 | Metazoa_SRP | 0.06078884 | 0.908254207 | 1.60749636 | 1.079432418 | 0.706761496 | midline crossing |
| ENSDART00000169023 | scn1lab | 0.043676774 | 0.230826198 | 0.448869227 | 0.411715364 | 0.200031153 | midline crossing |
| ENSDART00000169052 | elovl1b | 0.016102005 | 0.945814486 | 1.07228669 | 0.819418014 | 0.365575121 | midline crossing |
| ENSDART00000169061 | CU138533.1 | 0.004085991 | 0.641184794 | 1.068185172 | 1.17226183 | 0.773255864 | midline crossing |
| ENSDART00000169663 | CT573860.1 | 0.009406102 | 0.353646224 | 0.562802664 | 0.454063437 | 0.329757181 | midline crossing |
| ENSDART00000170095 | CR388166.2 | 0.047568271 | 0.588171845 | 0.299671021 | 0.452688856 | 0.506464888 | midline crossing |
| ENSDART00000170116 | uap1 | 0.041037763 | 0.399345456 | 0.560025587 | 0.455223073 | 0.418274301 | midline crossing |
| ENSDART00000170222 | mibp | 0.046085697 | 1.202449519 | 1.249757535 | 1.712700336 | 0.61416841 | midline crossing |
| ENSDART00000170284 | creb3l3l | 0.064177665 | 0.444489468 | 0.957040221 | 0.49606022 | 0.414651857 | midline crossing |
| ENSDART00000170374 | si:dkey-38l22.2 | 5.17084E-05 | 1.50589782 | 1.851640662 | 1.322719067 | 0.692201779 | midline crossing |
| ENSDART00000170422 | si:dkey-19b23.8 | 0.000292971 | 0.497209105 | 1.26444289 | 1.061660184 | 0.502612026 | midline crossing |
| ENSDART00000170572 | lepa | 0.00013528 | 4.559547685 | 4.778234325 | 4.366752593 | 2.765663683 | midline crossing |
| ENSDART00000170589 | mibp | 0.028456497 | 1.067201928 | 1.476075399 | 1.501871782 | 0.758874349 | midline crossing |
| ENSDART00000170671 | megf8 | 0.003012872 | 0.369502187 | 0.913401505 | 1.13324158 | 0.463058503 | midline crossing |
| ENSDART00000170684 | btf3 | 0.027344176 | 0.237794169 | 0.784907824 | 0.656975875 | 0.357635511 | midline crossing |
| ENSDART00000170768 | lect1 | 0.055604098 | 3.805179034 | 4.274135568 | 2.696697623 | 2.532825072 | midline crossing |
| ENSDART00000170865 | nme2b.1 | 0.007383951 | 0.635993303 | 0.679448572 | 0.689008761 | 0.355584206 | midline crossing |
| ENSDART00000170983 | lmnb2 | 0.03397627 | 0.185291183 | 0.299339057 | 0.275121803 | 0.118770566 | midline crossing |
| ENSDART00000171113 | mibp | 0.0245375 | 1.815496184 | 1.851942075 | 2.237510566 | 0.882565569 | midline crossing |
| ENSDART00000171235 | calua | 0.003825103 | 0.528206357 | 0.591046444 | 0.528298198 | 0.210887316 | midline crossing |
| ENSDART00000171343 | FP102192.1 | 0.009511897 | 1.130120264 | 0.697435719 | 1.760333984 | 0.139610385 | midline crossing |
| ENSDART00000171380 | top1mt | 0.004985516 | 0.983277478 | 1.316957074 | 1.074308725 | 0.32139594 | midline crossing |
| ENSDART00000171589 | hrasa | 0.008728551 | 0.543862541 | 0.913653251 | 0.982614467 | 0.767663706 | midline crossing |
| ENSDART00000171867 | fnbp1l | 0.000996442 | 0.402488238 | 0.925028089 | 0.937640279 | 0.578874881 | midline crossing |
| ENSDART00000171882 | CABZ01088567.1 | 0.0066372 | 0.407931 | 0.822027708 | 0.621405649 | 0.24977578 | midline crossing |
| ENSDART00000171996 | myl12.1 | 0.017201684 | 0.233861142 | 0.344330245 | 0.260508491 | 0.06338781 | midline crossing |
| ENSDART00000172215 | si:ch211-39i2.2 | 1.36144E-05 | 1.017534105 | 1.848603057 | 1.465869245 | 0.571196451 | midline crossing |
| ENSDART00000172372 | Metazoa_SRP | 0.027491007 | 1.431390172 | 1.797550169 | 1.401344471 | 0.830250287 | midline crossing |
| ENSDART00000172441 | lima1a | 0.014437179 | 0.350368785 | 0.588496789 | 0.46589541 | 0.178939257 | midline crossing |
| ENSDART00000172510 | elavl4 | 0.002554999 | 1.054519465 | 1.174247457 | 1.078840874 | 0.691066079 | midline crossing |
| ENSDART00000173098 | gemin6 | 0.046727574 | 0.181263356 | 0.385271136 | 0.184325919 | 0.274013326 | midline crossing |
| ENSDART00000173195 | zgc:153146 | 0.033291702 | 2.291177749 | 2.913913042 | 2.827467346 | 2.606942928 | midline crossing |
| ENSDART00000173305 | si:ch1073-456m8.1 | 0.012253248 | 0.206580067 | 0.587987288 | 0.667691948 | 0.271613596 | midline crossing |
| ENSDART00000173436 | si:dkey-280e21.3 | 8.39504E-05 | 0.740189589 | 1.373516231 | 1.387529232 | 0.976737711 | midline crossing |
| ENSDART00000000486 | cntn2 | 6.48692E-06 | 0.015280519 | 0.733067848 | 1.819345027 | 1.713832915 | target selection |
| ENSDART00000002393 | rundc3aa | 0.000184957 | -0.355283185 | 0.971542583 | 1.209230568 | 0.944629469 | target selection |
| ENSDART00000002961 | rcor2 | 0.001353924 | -0.353333584 | 0.726682254 | 0.762799057 | 0.664353552 | target selection |
| ENSDART00000003745 | vim | 0.015014385 | 0.110317427 | 0.81438402 | 0.975636783 | 0.646819018 | target selection |
| ENSDART00000003998 | ewsr1b | 0.006990173 | 0.136063726 | 0.084077199 | 0.439371598 | 0.241694854 | target selection |
| ENSDART00000004622 | sf3b4 | 0.02954888 | 0.222493415 | 0.286225027 | 0.527062646 | 0.243398323 | target selection |
| ENSDART00000005337 | rimkla | 0.001648434 | -0.133054455 | 0.30447461 | 0.568720788 | 0.413212633 | target selection |
| ENSDART00000005366 | tpd52l2b | 0.046265136 | 0.128420316 | 0.410964244 | 0.59220548 | 0.186138167 | target selection |
| ENSDART00000005453 | chd4a | 0.006932921 | 0.299643374 | 0.423708585 | 0.655765013 | 0.391160008 | target selection |
| ENSDART00000005496 | kctd9b | 0.010839093 | 0.161429839 | 0.406047159 | 0.541753399 | 0.319048316 | target selection |
| ENSDART00000005609 | kifap3a | 0.003425348 | 0.203573738 | 0.628570592 | 0.743430197 | 0.605809471 | target selection |
| ENSDART00000005720 | stat1a | 0.010250585 | 0.035880276 | 0.045195542 | 0.615137639 | 0.020671748 | target selection |
| ENSDART00000006085 | cbl | 0.034881046 | 0.165119386 | 0.1775588 | 0.379174787 | 0.206196339 | target selection |
| ENSDART00000006489 | acsl4a | 0.000400742 | 0.116872844 | 1.158097412 | 1.477099844 | 0.59458295 | target selection |
| ENSDART00000006724 | smarcd3b | 0.00033575 | -0.214336649 | 0.06424201 | 0.499562006 | 0.504466248 | target selection |
| ENSDART00000006778 | acat2 | 0.003807479 | -0.122422954 | 0.489045334 | 0.688039864 | 0.280940334 | target selection |
| ENSDART00000008038 | sulf2a | 1.1419E-05 | 0.420520728 | 1.274767381 | 2.360438199 | 2.491815243 | target selection |
| ENSDART00000009194 | aimp2 | 0.028420674 | 0.035867833 | 0.204745928 | 0.516008081 | 0.307277143 | target selection |
| ENSDART00000009277 | tuba1a | 0.000180202 | 0.466164103 | 1.394077049 | 1.773814961 | 1.31379942 | target selection |
| ENSDART00000010140 | igf2bp3 | 0.01196819 | 0.357904839 | 0.436869624 | 1.38352682 | 1.118120985 | target selection |
| ENSDART00000010144 | pvalb2 | 0.002147221 | 0.379417352 | 4.690875982 | 5.447468716 | 3.629116223 | target selection |
| ENSDART00000010274 | dpysl5a | 0.000114847 | 0.119382655 | 1.052869734 | 1.414498731 | 1.230769727 | target selection |
| ENSDART00000010495 | znrf1 | 0.011135896 | 0.045008698 | 0.17288769 | 0.379569247 | 0.252907322 | target selection |
| ENSDART00000011224 | itga10 | 0.008920935 | 2.439207816 | 3.888845597 | 4.302533523 | 3.055176776 | target selection |
| ENSDART00000011568 | syngr3a | 0.003678598 | -0.064295047 | 0.234968681 | 0.63147157 | 0.633880469 | target selection |
| ENSDART00000011865 | sec23b | 0.002343787 | -0.409405473 | 0.102005828 | 0.534469651 | 0.138588714 | target selection |
| ENSDART00000012247 | dhcr24 | 0.030292962 | 2.392110544 | 2.697063662 | 3.119814675 | 2.653468904 | target selection |
| ENSDART00000012256 | tnni2a.3 | 0.004454925 | 0.628164776 | 3.762159614 | 4.672246791 | 3.241031276 | target selection |
| ENSDART00000013605 | zbtb20 | 0.00075553 | 0.021191929 | 0.070670156 | 0.462529396 | 0.345025042 | target selection |
| ENSDART00000013785 | insig1 | 0.003330818 | -0.148803263 | 0.843708665 | 1.082868772 | 0.366540247 | target selection |
| ENSDART00000013961 | mycla | 0.015752787 | 0.091841991 | 0.503579515 | 0.855574629 | 0.294011787 | target selection |
| ENSDART00000014806 | npas2 | 0.005305184 | -0.341343772 | 0.009000027 | 0.555800516 | 0.156695025 | target selection |
| ENSDART00000014922 | arhgap22 | 0.004853289 | 0.337853511 | 0.496203573 | 0.771443869 | 0.664153347 | target selection |
| ENSDART00000015040 | hrasb | 0.001609185 | -0.035400047 | 0.468215746 | 0.71708775 | 0.509094476 | target selection |
| ENSDART00000015286 | ankrd13b | 0.001076713 | -0.294042347 | 0.149598935 | 0.484140903 | 0.605167909 | target selection |
| ENSDART00000015632 | nkain1 | 0.000894529 | 0.221220236 | 0.815235139 | 1.176280245 | 0.965286214 | target selection |
| ENSDART00000015979 | farsb | 0.041085489 | 0.087018819 | 0.265294461 | 0.405710239 | 0.211888538 | target selection |
| ENSDART00000016303 | irx2a | 4.96098E-05 | -0.381386746 | 0.064410059 | 0.752093904 | 0.989751717 | target selection |
| ENSDART00000016360 | si:ch73-199e17.1 | 0.028708281 | 2.690029092 | 3.363503415 | 3.69470671 | 3.529091333 | target selection |

| | | | | | | | |
|---|---|---|---|---|---|---|---|
| ENSDART00000016370 | dirc2 | 0.000512152 | 0.134905484 | 0.442850685 | 0.565287267 | 0.464631841 | target selection |
| ENSDART00000016597 | nfkbiab | 0.038559823 | 0.207474067 | 0.316720279 | 0.269395099 | 0.411969915 | target selection |
| ENSDART00000016710 | scrn3 | 0.001328724 | -0.247629441 | -0.171036354 | 0.44401532 | -0.052873668 | target selection |
| ENSDART00000016860 | ppp2r1bb | 0.015469128 | 0.018499492 | 0.284521267 | 0.321334616 | 0.193996267 | target selection |
| ENSDART00000017299 | tdg.1 | 0.043370331 | 0.102853523 | 0.194463285 | 0.348171907 | 0.196594799 | target selection |
| ENSDART00000018150 | neurod6b | 0.012342358 | 0.484226112 | 1.695956927 | 1.935318885 | 1.555476315 | target selection |
| ENSDART00000018498 | helz2 | 0.00076674 | -0.063748361 | -0.192163226 | 1.151139617 | 0.013207133 | target selection |
| ENSDART00000018735 | dnaja2l | 0.000177237 | -0.20745831 | 0.012628814 | 0.350147509 | 0.311991137 | target selection |
| ENSDART00000019045 | ebp | 0.003165422 | -0.186359255 | 0.947675675 | 1.363611199 | 0.55228711 | target selection |
| ENSDART00000019199 | rab39ba | 0.013286981 | -0.125963686 | 0.3041991 | 0.44382958 | 0.206562916 | target selection |
| ENSDART00000019617 | rsad2 | 0.008246433 | -0.056061565 | -0.489318095 | 2.739834278 | -0.378754222 | target selection |
| ENSDART00000019658 | nacad | 8.67551E-05 | -0.254606168 | 0.44010123 | 0.744298293 | 0.29398429 | target selection |
| ENSDART00000019925 | GNB4 | 0.000718633 | 0.24039241 | 0.43881347 | 0.966289057 | 1.074168682 | target selection |
| ENSDART00000020122 | ywhah | 0.000166651 | -0.243048068 | -0.041853394 | 0.44002204 | 0.422069749 | target selection |
| ENSDART00000020621 | mapk4 | 0.001185711 | -0.137443479 | 0.345820304 | 0.800085717 | 0.474998646 | target selection |
| ENSDART00000020970 | pgm2 | 0.021002038 | 0.182379442 | 0.224260514 | 0.427220163 | 0.369397058 | target selection |
| ENSDART00000021341 | kif3ca | 0.001627738 | 0.127548246 | 0.488322423 | 0.730937873 | 0.492487971 | target selection |
| ENSDART00000021693 | ank2a | 0.031838485 | 0.088061869 | 0.283890991 | 0.488712819 | 0.41975176 | target selection |
| ENSDART00000022270 | arhgap33 | 0.034698678 | 0.060975731 | 0.144058893 | 0.360327899 | 0.271763038 | target selection |
| ENSDART00000022909 | klhl18 | 0.00115719 | -0.153412936 | 0.073357396 | 0.397749012 | 0.249762723 | target selection |
| ENSDART00000023278 | fads2 | 0.00304733 | -0.280275959 | 0.609167416 | 1.126645574 | 0.470269511 | target selection |
| ENSDART00000023944 | lmnl3 | 0.001714837 | -0.134674249 | 0.538751787 | 0.920098245 | 0.528719758 | target selection |
| ENSDART00000023959 | arntl1a | 7.43514E-05 | 0.340904931 | 1.030627189 | 1.45255612 | 0.975945139 | target selection |
| ENSDART00000024135 | tubb2 | 7.10385E-05 | -0.097026879 | 1.085064176 | 1.27895467 | 1.027950073 | target selection |
| ENSDART00000025852 | tnni2b.1 | 0.000911246 | -0.577234038 | 4.833234821 | 5.215670785 | 3.899605393 | target selection |
| ENSDART00000026800 | kifap3b | 0.001775429 | 0.203180282 | 0.588115089 | 0.640508934 | 0.450843691 | target selection |
| ENSDART00000026992 | sox4a | 3.6604E-05 | 0.168735758 | 0.553498244 | 0.871364123 | 0.630648218 | target selection |
| ENSDART00000027158 | psmd3 | 0.034155477 | 0.079773681 | 0.202742016 | 0.46801517 | 0.212038089 | target selection |
| ENSDART00000027379 | bicral | 0.012896166 | 0.180958951 | 0.201052129 | 0.429766407 | 0.252075745 | target selection |
| ENSDART00000027598 | tpm3 | 0.022054847 | 0.692886998 | 2.316316343 | 2.715886041 | 1.570432549 | target selection |
| ENSDART00000027718 | fxr2 | 0.020107844 | 0.070260526 | 0.251047986 | 0.370057736 | 0.114198992 | target selection |
| ENSDART00000027758 | rtn1b | 0.000135341 | -0.626628578 | 0.207141964 | 0.617559397 | 0.433726227 | target selection |
| ENSDART00000028607 | chd6 | 0.008909863 | 0.00488027 | -0.079639497 | 0.533277243 | 0.378096362 | target selection |
| ENSDART00000029121 | usp5 | 0.005099354 | 0.186533139 | 0.264124698 | 0.696851044 | 0.548402162 | target selection |
| ENSDART00000031470 | pafah1b1b | 0.001868725 | -0.06625793 | 0.19164974 | 0.422888398 | 0.388967525 | target selection |
| ENSDART00000033724 | fabp3 | 0.001967837 | 0.043357927 | 0.829727862 | 0.980692721 | 0.446444515 | target selection |
| ENSDART00000034216 | dync1h1 | 0.038721816 | 0.15631552 | 0.099035755 | 0.755817679 | 0.554910929 | target selection |
| ENSDART00000034914 | pvalb3 | 0.006583055 | -0.375270417 | 3.922610155 | 4.415489459 | 3.238850832 | target selection |
| ENSDART00000035245 | spire2 | 9.16648E-05 | 0.223570717 | 0.667119992 | 0.874236758 | 0.626613708 | target selection |
| ENSDART00000035409 | zc2hc1a | 0.000588633 | 0.100393521 | 0.424633369 | 0.532716266 | 0.492034687 | target selection |
| ENSDART00000036926 | vangl1 | 0.000398071 | -0.329590873 | -0.033233324 | 0.383553313 | 0.3451937 | target selection |
| ENSDART00000038391 | pkz | 0.008061109 | 0.411244814 | 0.129537507 | 1.595328762 | 0.021626224 | target selection |
| ENSDART00000040557 | CRIP2 (1 of many) | 0.000349374 | -0.132882678 | 0.562285777 | 0.768710088 | 0.441120913 | target selection |
| ENSDART00000042134 | dock7 | 0.014840867 | 0.063646619 | 0.209946461 | 0.48316367 | 0.274325565 | target selection |
| ENSDART00000042162 | tm7sf2 | 0.008621509 | 0.1050645 | 0.758038931 | 0.952561873 | 0.4137868 | target selection |
| ENSDART00000042218 | pafah1b1a | 0.024122965 | 0.049322983 | 0.127134272 | 0.313096774 | 0.220746894 | target selection |
| ENSDART00000042255 | rab6bb | 0.000105127 | 0.249992007 | 1.135776594 | 1.498200114 | 1.307756366 | target selection |
| ENSDART00000043932 | atp2a1 | 0.046659681 | 1.138744452 | 4.285725335 | 5.558096993 | 3.994336477 | target selection |
| ENSDART00000044000 | plxna3 | 0.000326965 | 0.522678231 | 0.518528602 | 0.862237135 | 0.748340859 | target selection |
| ENSDART00000044057 | 43346 | 0.00053559 | -0.17670354 | 0.196164293 | 0.478730018 | 0.43476862 | target selection |
| ENSDART00000044453 | ano5a | 0.00350062 | -0.027551778 | 0.223592536 | 0.548330121 | 0.427829524 | target selection |
| ENSDART00000044949 | syt16 | 4.96098E-05 | -0.001457693 | -0.053066568 | 0.72824965 | 0.634879457 | target selection |
| ENSDART00000045391 | srgap2 | 0.005041829 | 0.042169336 | 0.118632088 | 0.377794988 | 0.307955564 | target selection |
| ENSDART00000045410 | thy1 | 8.488E-05 | 0.906274294 | 2.922050483 | 2.917431913 | 2.440516966 | target selection |
| ENSDART00000046678 | pak2b | 0.02432669 | 0.067537468 | 0.265866932 | 0.367938524 | 0.197303091 | target selection |
| ENSDART00000046933 | sult1st5 | 0.009503355 | 0.218426293 | 0.869949792 | 0.854237199 | 0.702293309 | target selection |
| ENSDART00000047082 | gdap1l1 | 0.00058454 | -0.237605119 | 0.285676986 | 0.669438597 | 0.59731396 | target selection |
| ENSDART00000047143 | specc1 | 0.039415449 | 0.250947915 | 0.397558549 | 0.627196335 | 0.309975843 | target selection |
| ENSDART00000047409 | myh14 | 0.014779327 | 0.151853201 | 0.372826224 | 0.643058289 | 0.555183408 | target selection |
| ENSDART00000048110 | six4b | 0.002795018 | 0.757934328 | 2.322346312 | 2.776043581 | 1.88012445 | target selection |
| ENSDART00000049194 | gpr37b | 2.18017E-05 | -0.562015213 | 0.012994785 | 0.671207906 | 0.757619967 | target selection |
| ENSDART00000051231 | gnb2 | 0.002814147 | -0.235263329 | 0.124417594 | 0.575032632 | 0.495502612 | target selection |
| ENSDART00000051234 | tnika | 0.001841908 | 0.106489865 | 0.189584243 | 0.309030435 | 0.511149463 | target selection |
| ENSDART00000051697 | evla | 0.003779087 | -0.055749697 | 0.354222037 | 0.437976697 | 0.322467243 | target selection |
| ENSDART00000052124 | fam49al | 0.000100528 | -0.409057809 | -0.177161374 | 0.563238983 | 0.47605027 | target selection |
| ENSDART00000052511 | hnrnpa0l | 0.02613431 | 0.098563881 | 0.214926285 | 0.391405743 | 0.211573162 | target selection |
| ENSDART00000053284 | bcl9 | 0.003126045 | -0.057531558 | 0.216352129 | 0.432379429 | 0.260365529 | target selection |
| ENSDART00000053869 | slc44a2 | 0.000322013 | 0.03477078 | 0.96666703 | 1.3453814 | 0.899279007 | target selection |
| ENSDART00000054664 | tnnc1b | 0.019014873 | -0.083935925 | 3.176389417 | 3.430200861 | 1.884280525 | target selection |
| ENSDART00000054691 | uba1 | 0.014095383 | 0.10035699 | 0.287144805 | 0.420196482 | 0.237625478 | target selection |
| ENSDART00000055340 | fus | 0.000159901 | -0.011490396 | 0.383733399 | 0.662780299 | 0.416441518 | target selection |
| ENSDART00000055380 | tubb5 | 2.92142E-06 | 2.169554793 | 3.899556169 | 4.316772272 | 3.57490014 | target selection |
| ENSDART00000055756 | tbc1d12a | 0.00448854 | -0.033893992 | 0.095679716 | 0.342628218 | 0.232107662 | target selection |
| ENSDART00000056286 | h1f0 | 0.001739475 | -0.326358805 | 0.273543898 | 0.626230484 | 0.593578182 | target selection |
| ENSDART00000056544 | tox4a | 0.02114031 | 0.131870382 | 0.296259606 | 0.350040714 | 0.249645538 | target selection |

| | | | | | | | |
|---|---|---|---|---|---|---|---|
| ENSDART00000056795 | hectd3 | 0.004611386 | 0.187585997 | 0.34309988 | 0.435708832 | 0.360997812 | target selection |
| ENSDART00000057584 | slc1a4 | 0.00134801 | 0.291595567 | 1.233259249 | 1.405286425 | 1.089173123 | target selection |
| ENSDART00000058574 | NA | 0.032955534 | 3.231975259 | 2.942133944 | 3.360676782 | 3.717330017 | target selection |
| ENSDART00000059586 | spegb | 4.52852E-05 | 0.60633626 | 1.046592342 | 1.773590673 | 1.534849845 | target selection |
| ENSDART00000059756 | ralba | 0.014582204 | 0.050810042 | 0.323679093 | 0.529596851 | 0.259207353 | target selection |
| ENSDART00000060561 | csdc2a | 0.006063577 | -0.084744858 | 0.609682067 | 0.574951056 | 0.285883083 | target selection |
| ENSDART00000060766 | rab11a | 0.001380734 | 0.179968247 | 0.479437298 | 0.674000894 | 0.468733557 | target selection |
| ENSDART00000061417 | si:ch211-245h14.1 | 0.00918017 | 0.537591116 | 0.459433616 | 1.387404638 | -0.014689392 | target selection |
| ENSDART00000061470 | mtss1la | 0.002100653 | -0.342228031 | 0.053961203 | 0.113929451 | 0.018495568 | target selection |
| ENSDART00000061523 | il17a/f3 | 0.017627165 | 0.228114616 | 0.421937628 | 0.621201213 | 0.769032572 | target selection |
| ENSDART00000061955 | myl13 | 0.006605056 | 0.400602979 | 3.991209963 | 4.290518336 | 2.830575983 | target selection |
| ENSDART00000062551 | cyp51 | 0.0002611 | 0.018772193 | 0.850105191 | 1.080881624 | 0.554137375 | target selection |
| ENSDART00000062560 | zgc:77784 | 0.054165556 | 0.130172136 | 0.171187774 | 0.614861949 | 0.368542156 | target selection |
| ENSDART00000063938 | mast1a | 0.000314801 | 0.148358744 | 0.555318149 | 1.032634027 | 1.008672267 | target selection |
| ENSDART00000065097 | dpysl3 | 0.000311162 | -0.247117182 | 0.32883927 | 0.696704704 | 0.442557487 | target selection |
| ENSDART00000065600 | sc5d | 0.009204947 | -0.12560715 | 0.619273704 | 0.802648489 | 0.437253012 | target selection |
| ENSDART00000066177 | tuba2 | 7.26748E-05 | -0.341980556 | 0.290211936 | 0.695639518 | 0.704284087 | target selection |
| ENSDART00000066391 | csnk1e | 0.000593122 | 0.01172114 | 0.595315231 | 0.792117229 | 0.433182971 | target selection |
| ENSDART00000066975 | impdh1b | 0.000512556 | 0.195914466 | 0.339716801 | 0.56808711 | 0.477613078 | target selection |
| ENSDART00000067066 | parp6b | 1.45694E-05 | -0.443166413 | 0.101161009 | 0.402103535 | 0.516387314 | target selection |
| ENSDART00000067327 | abhd2b | 0.0021138 | 0.159640353 | 0.698443956 | 0.611964225 | 0.750739323 | target selection |
| ENSDART00000073583 | islr2 | 0.000634488 | 0.313523936 | 1.12325114 | 1.332162091 | 1.179316934 | target selection |
| ENSDART00000074924 | mbnl1 | 0.094159425 | 0.32889023 | 0.277718787 | 0.642145523 | 0.211541009 | target selection |
| ENSDART00000074979 | rnft2 | 0.02083117 | 0.06664962 | 0.449659345 | 0.588779493 | 0.279626887 | target selection |
| ENSDART00000075112 | clvs2 | 0.003104562 | 0.165980729 | 0.346379383 | 0.75563827 | 0.535875063 | target selection |
| ENSDART00000075260 | inab | 2.82217E-05 | -0.558694355 | 0.275313434 | 1.165237079 | 1.111612471 | target selection |
| ENSDART00000075331 | insm1b | 0.009480138 | 0.180818368 | 0.558805202 | 0.852549514 | 0.856981121 | target selection |
| ENSDART00000075743 | eprs | 0.03427447 | 0.181617497 | 0.208060281 | 0.357036503 | 0.248273691 | target selection |
| ENSDART00000075808 | apbb3 | 0.002539213 | -0.025817957 | 0.161567226 | 0.465828121 | 0.45985711 | target selection |
| ENSDART00000075902 | klhl43 | 0.004256148 | 1.602362948 | 3.319039146 | 3.627754288 | 3.174379945 | target selection |
| ENSDART00000076506 | wisp1a | 0.025227573 | 0.075916017 | 0.908100257 | 1.435223818 | 0.542654062 | target selection |
| ENSDART00000076574 | rtn1a | 0.00434437 | -0.132166015 | 0.374408359 | 0.606191338 | 0.476970779 | target selection |
| ENSDART00000077215 | ppp2r5b | 0.002524433 | 0.185911454 | 0.229029828 | 0.615811771 | 0.611674376 | target selection |
| ENSDART00000077459 | smyd2a | 0.042433381 | 0.109333535 | 0.326579335 | 0.423001533 | 0.365696401 | target selection |
| ENSDART00000077539 | tuba1c | 0.00109551 | -0.211130957 | 0.180769664 | 0.840172721 | 0.834422327 | target selection |
| ENSDART00000078137 | ankrd54 | 0.034178169 | 0.051281727 | 0.23783078 | 0.382868701 | 0.32135773 | target selection |
| ENSDART00000078277 | msmo1 | 0.001153812 | 0.087631699 | 1.109674669 | 1.47165822 | 0.852850391 | target selection |
| ENSDART00000078334 | celsr3 | 0.002605308 | 0.469496099 | 0.512562938 | 0.74195357 | 0.599858827 | target selection |
| ENSDART00000078771 | SBK1 | 0.00198437 | -0.390774004 | 0.13684565 | 0.923517231 | 0.784697628 | target selection |
| ENSDART00000079235 | cd99l2 | 0.000613723 | -0.167082345 | 0.448795434 | 0.91020128 | 0.864265064 | target selection |
| ENSDART00000079283 | tmeff1b | 0.001133915 | 0.060962109 | 0.679659218 | 0.971148806 | 0.712213835 | target selection |
| ENSDART00000079559 | ifit16 | 0.03326116 | 0.716638095 | 0.325468397 | 2.311988055 | -0.283531995 | target selection |
| ENSDART00000079711 | slc25a1a | 0.000122706 | 0.025053389 | 0.916416559 | 1.283110285 | 0.925187833 | target selection |
| ENSDART00000079778 | ifit8 | 0.035967167 | 1.432186723 | 0.097505141 | 4.166007884 | 0.96250098 | target selection |
| ENSDART00000080033 | SSBP2 (1 of many) | 0.034092146 | 0.123939988 | 0.461173497 | 0.600467157 | 0.3572706 | target selection |
| ENSDART00000080042 | rab33a | 0.001631005 | -0.045588719 | 0.341497657 | 0.630733663 | 0.503197845 | target selection |
| ENSDART00000080256 | nefma | 1.75467E-05 | 0.572074143 | 2.009260502 | 2.281701505 | 1.529767044 | target selection |
| ENSDART00000080289 | prrc2c | 0.035944076 | 0.060093623 | 0.047472332 | 0.306363857 | 0.189281148 | target selection |
| ENSDART00000080602 | map7d2b | 0.001304993 | 0.243598476 | 0.831190471 | 1.239754477 | 0.898189276 | target selection |
| ENSDART00000081059 | rps6kb1b | 0.042381041 | 0.156747941 | 0.167680861 | 0.344534054 | 0.174989656 | target selection |
| ENSDART00000081092 | si:dkeyp-77h1.4 | 0.000199258 | -0.383994825 | 0.24121529 | 0.967657356 | 0.94823494 | target selection |
| ENSDART00000081223 | krt5 | 0.004002952 | 2.400508858 | 4.729984226 | 4.862312842 | 5.813921541 | target selection |
| ENSDART00000081272 | gcn1 | 0.02181186 | 0.168820856 | 0.100284898 | 0.363305756 | 0.2232516 | target selection |
| ENSDART00000081601 | cept1a | 0.020882958 | 0.150762086 | 0.296977372 | 0.445683893 | 0.275050142 | target selection |
| ENSDART00000082944 | dock6 | 0.004360669 | 0.129469289 | 0.283374744 | 0.379764071 | 0.290753034 | target selection |
| ENSDART00000083002 | map1aa | 0.00881795 | 0.023824463 | 0.057494175 | 0.8302085 | 0.470304447 | target selection |
| ENSDART00000083212 | fscn1a | 0.001033408 | -0.172521555 | 0.403697769 | 0.865067524 | 0.635846942 | target selection |
| ENSDART00000083605 | tbc1d25 | 0.020097848 | 0.094218097 | 0.226970926 | 0.332123969 | 0.140604089 | target selection |
| ENSDART00000083628 | ddit3 | 0.002192622 | -0.053032847 | 0.505000026 | 0.598711637 | 0.25299651 | target selection |
| ENSDART00000083797 | TBC1D9B | 0.000642503 | 0.273448063 | 0.469124554 | 0.581828184 | 0.40676136 | target selection |
| ENSDART00000084819 | arhgap35b | 0.001659006 | 0.101645005 | 0.291379647 | 0.535032879 | 0.462050203 | target selection |
| ENSDART00000085230 | atl1 | 0.003020568 | -0.283512177 | 0.138626411 | 0.619353197 | 0.442930265 | target selection |
| ENSDART00000085263 | SELENOI | 0.02776376 | 0.036305982 | 0.149651758 | 0.460720877 | 0.164646013 | target selection |
| ENSDART00000085319 | sos2 | 0.038217845 | 0.062155314 | 0.173737983 | 0.464042198 | 0.208637802 | target selection |
| ENSDART00000085388 | bmp3 | 0.004205562 | -0.146774305 | 1.240822845 | 1.494378971 | 0.416428855 | target selection |
| ENSDART00000085453 | cluhb | 5.72005E-05 | -0.484145834 | -0.430773206 | 0.678836708 | -0.017745757 | target selection |
| ENSDART00000085684 | ttll11 | 0.000375747 | -0.16810055 | 0.020727161 | 0.348458395 | 0.408691778 | target selection |
| ENSDART00000085719 | si:ch211-10a23.2 | 0.001067718 | 0.141906082 | 0.927202925 | 1.15606105 | 0.59243708 | target selection |
| ENSDART00000086301 | irge4 | 0.015651339 | 0.875091617 | 0.681146857 | 1.737500297 | 0.803175592 | target selection |
| ENSDART00000086333 | jarid2a | 0.000531966 | -0.109318405 | 0.11715985 | 0.373609719 | 0.35216542 | target selection |
| ENSDART00000086409 | dync1i1 | 0.002016009 | -0.021724393 | 0.427961112 | 0.551451734 | 0.438467635 | target selection |
| ENSDART00000086434 | tmcc2 | 0.000793313 | 0.022092982 | 0.338134833 | 0.850580716 | 0.840264518 | target selection |
| ENSDART00000086537 | NA | 0.022370255 | 0.422133332 | 0.379254853 | 0.651471983 | 0.472158305 | target selection |
| ENSDART00000086946 | mov10b.1 | 0.016426365 | 0.41644893 | 0.305142124 | 1.89370555 | 0.04709269 | target selection |
| ENSDART00000087105 | myom2a | 0.029772573 | 2.177324505 | 3.608086766 | 3.587707417 | 2.779461648 | target selection |

| | | | | | | | |
|---|---|---|---|---|---|---|---|
| ENSDART00000087107 | eif4g1a | 0.01673556 | 0.252655247 | -0.051267066 | 0.413833734 | 0.330964133 | target selection |
| ENSDART00000087115 | rims1b | 0.002016009 | -0.155072254 | 0.041830315 | 0.378907446 | 0.287621284 | target selection |
| ENSDART00000087204 | dusp3a | 0.000778506 | -0.327203373 | -0.053264752 | 0.592884224 | 0.622404395 | target selection |
| ENSDART00000088364 | kif1aa | 0.000306344 | -0.06544832 | 0.34113245 | 0.707853532 | 0.648494399 | target selection |
| ENSDART00000088653 | prss12 | 0.001247242 | -0.641371677 | 0.068962263 | 0.215098805 | 0.069158882 | target selection |
| ENSDART00000089076 | dot1l | 0.009173096 | 0.094529933 | 0.165735937 | 0.448240264 | 0.463978672 | target selection |
| ENSDART00000089079 | mpnd | 0.001644836 | 0.049906779 | 0.784892334 | 0.951903848 | 0.691834891 | target selection |
| ENSDART00000089141 | fsd1 | 0.000907455 | 0.140911421 | 0.537063785 | 0.66216846 | 0.489473142 | target selection |
| ENSDART00000089445 | agap1 | 0.028468759 | 0.111728293 | 0.12652166 | 0.321279752 | 0.223152778 | target selection |
| ENSDART00000090596 | fgf12b | 0.000227894 | -0.346267349 | -0.034488788 | 0.704892235 | 0.660670447 | target selection |
| ENSDART00000090748 | pcdh1g9 | 0.004897476 | 0.151679401 | 0.129768355 | 0.681696549 | 0.559518569 | target selection |
| ENSDART00000090874 | kcnh7 | 0.000589961 | 0.089634054 | 0.066677874 | 0.800119794 | 0.739010611 | target selection |
| ENSDART00000091241 | si:ch73-22o12.1 | 0.003818266 | -0.053659471 | 0.086463787 | 0.414689493 | 0.463272551 | target selection |
| ENSDART00000091644 | abi1b | 0.016742251 | 0.181595456 | 0.359192672 | 0.565695572 | 0.32038751 | target selection |
| ENSDART00000092270 | r3hdm4 | 0.003016249 | 0.109330042 | 0.341009284 | 0.430965659 | 0.322117566 | target selection |
| ENSDART00000092690 | srebf2 | 0.003411856 | -0.007030288 | 0.441941306 | 0.773565631 | 0.436893785 | target selection |
| ENSDART00000092948 | peli1b | 0.025378177 | 0.088983174 | 0.130189996 | 0.342697853 | 0.254903056 | target selection |
| ENSDART00000098057 | ftr19 | 0.021658197 | 0.135898929 | 0.299355243 | 0.876453981 | -0.013001669 | target selection |
| ENSDART00000098284 | ftr14 | 0.009877168 | 0.428599203 | 0.304151818 | 1.741077228 | -0.09285898 | target selection |
| ENSDART00000099977 | mex3c | 0.051405603 | 0.161233198 | 0.169039948 | 0.509657606 | 0.094152947 | target selection |
| ENSDART00000100310 | dbn1 | 0.015825859 | 0.014767176 | 0.221674025 | 0.550020524 | 0.324902581 | target selection |
| ENSDART00000100332 | fgf12b | 0.00038813 | -0.220479512 | 0.029676005 | 0.518232207 | 0.657400596 | target selection |
| ENSDART00000100619 | zgc:158803 | 0.028599369 | 0.343521186 | 0.388812734 | 0.518620429 | 0.45576524 | target selection |
| ENSDART00000101038 | tmie | 0.000195011 | 0.304653896 | 1.158765103 | 1.299122704 | 1.103412502 | target selection |
| ENSDART00000101282 | bcr | 0.005519422 | 0.172989029 | 0.500540591 | 0.708792261 | 0.385177654 | target selection |
| ENSDART00000101537 | mex3b | 0.000847723 | -0.20648165 | 0.480244705 | 0.713185315 | 0.514833911 | target selection |
| ENSDART00000101789 | flot2b | 0.000781013 | 0.047673595 | 0.464453756 | 0.590584487 | 0.461504068 | target selection |
| ENSDART00000102011 | G3BP2 (1 of many) | 0.015925333 | 0.183053703 | 0.292940696 | 0.414353657 | 0.391197687 | target selection |
| ENSDART00000102212 | tdp2a | 0.010054985 | 1.019634105 | 0.831370615 | 2.830207945 | 0.310574015 | target selection |
| ENSDART00000102305 | cspg5a | 9.03292E-05 | -0.238742988 | 0.181941504 | 0.751580581 | 0.677972326 | target selection |
| ENSDART00000102384 | sesn2 | 0.029981898 | 0.221656214 | 0.707312613 | 0.831938385 | 0.650662976 | target selection |
| ENSDART00000102520 | palm1a | 0.000475692 | 0.447081887 | 0.96607967 | 1.214937327 | 0.924772733 | target selection |
| ENSDART00000102539 | st8sia5 | 0.002696406 | 0.153973405 | 0.592846857 | 1.061043515 | 0.777183018 | target selection |
| ENSDART00000102665 | aste1a | 0.002793888 | 0.478587949 | 0.328068557 | 2.324198666 | 0.218780766 | target selection |
| ENSDART00000102712 | tgm2a | 0.024073602 | 0.064390317 | 0.421175856 | 0.197637738 | 0.730671181 | target selection |
| ENSDART00000102715 | tuba8l3 | 0.000489924 | 1.032352614 | 1.505768874 | 1.918452626 | 1.697934809 | target selection |
| ENSDART00000102868 | etnk2 | 0.002660549 | -0.119773861 | 0.158574082 | 0.41890552 | 0.249052751 | target selection |
| ENSDART00000103588 | mxa | 0.022415732 | 0.174383297 | -0.012993595 | 1.447594397 | 0.019311963 | target selection |
| ENSDART00000103622 | irf7 | 0.002828188 | -0.049332142 | -0.2878598 | 1.533925917 | -0.275107455 | target selection |
| ENSDART00000103815 | stmn2a | 4.96098E-05 | 0.140137036 | 1.176009379 | 1.538144395 | 1.355356024 | target selection |
| ENSDART00000103922 | atat1 | 0.015462868 | 0.145026536 | 0.665460259 | 0.937418253 | 0.683690675 | target selection |
| ENSDART00000104299 | cnp | 1.98934E-05 | 0.814960421 | 1.713687269 | 2.20652149 | 1.77606709 | target selection |
| ENSDART00000104336 | tnnc2 | 0.0274914 | 0.703924131 | 2.455705453 | 2.232729877 | 1.561721699 | target selection |
| ENSDART00000104478 | fxr1 | 0.013415803 | 0.018053797 | 0.098113697 | 0.377704225 | 0.105513892 | target selection |
| ENSDART00000104523 | arntl1b | 0.000255127 | -0.158144123 | 0.59291526 | 1.113237486 | 0.52944604 | target selection |
| ENSDART00000104720 | si:ch1073-385f13.3 | 0.037756042 | 0.258133799 | 0.237616235 | 1.745458762 | -0.119672944 | target selection |
| ENSDART00000104730 | ifit14 | 0.018324408 | 0.457651023 | 0.403391968 | 2.518119653 | 0.537556431 | target selection |
| ENSDART00000105405 | cirbpb | 0.023596757 | 0.001082658 | 0.185977938 | 0.46521688 | -0.018730312 | target selection |
| ENSDART00000105694 | adprm | 0.001249299 | 0.302167589 | 0.521514455 | 0.908630272 | 0.271324554 | target selection |
| ENSDART00000105873 | cry4 | 0.001698814 | -0.404336974 | 0.122755184 | 0.671823255 | 0.515326773 | target selection |
| ENSDART00000106120 | ywhag2 | 0.014767815 | 0.097311782 | 0.334968358 | 0.584303472 | 0.612363287 | target selection |
| ENSDART00000108796 | stox2b | 0.00058454 | 0.143615187 | 0.758742406 | 1.120462456 | 0.794663921 | target selection |
| ENSDART00000109288 | myo16 | 0.003478587 | 0.115485168 | 0.089369719 | 0.360073499 | 0.534996054 | target selection |
| ENSDART00000109615 | tmem255a | 0.000634043 | 0.198392818 | 0.721353886 | 0.92551731 | 0.71998369 | target selection |
| ENSDART00000109750 | rap2aa | 0.015651953 | 0.293487605 | 0.482823125 | 0.662590656 | 0.396980886 | target selection |
| ENSDART00000109973 | tp53bp1 | 0.000154883 | 0.061139928 | 0.539512942 | 0.657824217 | 0.222950524 | target selection |
| ENSDART00000110512 | ybx1 | 0.014352201 | 0.187678935 | 0.306636132 | 0.543417204 | 0.157462635 | target selection |
| ENSDART00000110696 | mxb | 0.001413354 | -0.714840475 | -2.62990329 | 3.629462021 | -0.04990443 | target selection |
| ENSDART00000110964 | bag6 | 0.014908663 | 0.132774646 | 0.150367292 | 0.354477336 | 0.199439987 | target selection |
| ENSDART00000110994 | sqlea | 0.000140513 | 0.510664466 | 1.875874264 | 2.415595993 | 1.866281513 | target selection |
| ENSDART00000111278 | sorcs2 | 0.000322759 | -0.061568527 | 0.240045859 | 0.726875042 | 0.744539967 | target selection |
| ENSDART00000111303 | rfx7 | 0.00161581 | 0.075232989 | 0.085195342 | 0.459783838 | 0.727409902 | target selection |
| ENSDART00000111454 | UBA6 | 0.018326914 | -0.095626975 | 0.120665542 | 0.496662318 | 0.328619297 | target selection |
| ENSDART00000111475 | pcdh7a | 0.008009641 | 0.109274008 | 0.268985487 | 0.465377244 | 0.354118271 | target selection |
| ENSDART00000111480 | bcorl1 | 0.017081311 | 0.013917394 | 0.173170994 | 0.344992278 | 0.232054818 | target selection |
| ENSDART00000111561 | zmp:0000000735 | 0.000689533 | 0.220285812 | 0.110420777 | 1.549961519 | 0.06406158 | target selection |
| ENSDART00000111641 | morn4 | 0.00308528 | 0.104472779 | 0.38320597 | 0.615725038 | 0.556227667 | target selection |
| ENSDART00000111671 | mri1 | 0.001380734 | 1.003264298 | 0.870246845 | 1.443034807 | 1.04475049 | target selection |
| ENSDART00000111748 | pcdhb | 0.000529346 | 0.251557806 | 0.39162054 | 0.85552916 | 0.748922663 | target selection |
| ENSDART00000112032 | ARHGAP22 (1 of many) | 0.001967837 | 0.396989069 | 0.894459004 | 1.264783984 | 1.011214691 | target selection |
| ENSDART00000112075 | NA | 0.034693937 | 0.743523521 | 0.909798042 | 1.147105824 | 0.978802209 | target selection |
| ENSDART00000112312 | lrch2 | 0.015854118 | 0.101952948 | 0.206054969 | 0.497048672 | 0.404162087 | target selection |
| ENSDART00000112313 | wdr43 | 0.033970998 | 2.967338106 | 2.621161906 | 3.405726008 | 2.58425126 | target selection |
| ENSDART00000112438 | si:ch73-335m24.5 | 0.002769129 | -0.12977304 | -0.024649938 | 0.356284768 | 0.481799029 | target selection |
| ENSDART00000112460 | dolk | 0.02235203 | 0.234400054 | 0.274257795 | 0.504999751 | 0.504335897 | target selection |

| | | | | | | | |
|---|---|---|---|---|---|---|---|
| ENSDART00000112598 | otud4 | 0.006973136 | 0.000834797 | 0.118841749 | 0.673957038 | 0.019221004 | target selection |
| ENSDART00000112694 | fam171a2b | 0.001676296 | -0.001835186 | 0.12088862 | 0.541529951 | 0.657778888 | target selection |
| ENSDART00000112728 | tmem175 | 0.004215641 | -0.092618029 | 0.36107371 | 0.426939782 | 0.221686367 | target selection |
| ENSDART00000112883 | fmnl2b | 0.000169466 | -0.095811604 | 0.162122972 | 0.557976718 | 0.595265469 | target selection |
| ENSDART00000113087 | hmx1 | 0.000694227 | -0.389387565 | 0.021377927 | 0.234018679 | 0.270888749 | target selection |
| ENSDART00000113097 | hsd17b7 | 0.001028061 | -0.488876774 | 0.314769091 | 0.517151777 | 0.148788215 | target selection |
| ENSDART00000113418 | igsf3 | 0.023363274 | 0.097020516 | 0.19405944 | 0.607620623 | 0.491282072 | target selection |
| ENSDART00000114919 | si:dkey-85k7.12 | 0.012263789 | 0.382806755 | -0.226494664 | 2.101494432 | -0.132340955 | target selection |
| ENSDART00000115023 | PARG | 0.014732696 | 0.013352577 | 0.067569536 | 0.312910159 | 0.18617429 | target selection |
| ENSDART00000121837 | efs | 0.000534872 | 0.29165502 | 0.378275961 | 0.750399498 | 0.505660243 | target selection |
| ENSDART00000121981 | smarce1 | 0.04263061 | 0.060968412 | 0.140256121 | 0.341840216 | 0.044607056 | target selection |
| ENSDART00000121984 | ssbp3b | 0.013312578 | -0.092095029 | 0.005321722 | 0.831022729 | 0.629434393 | target selection |
| ENSDART00000122133 | mkrn2os.1 | 0.028489994 | 0.117189294 | 0.106487037 | 0.442579902 | 0.198380082 | target selection |
| ENSDART00000122170 | smc5 | 0.003077552 | 0.002327719 | -0.01045929 | 0.55091638 | 0.063311494 | target selection |
| ENSDART00000122681 | CR848841.1 | 0.000177237 | 0.726082565 | 1.593466942 | 1.807590182 | 1.450515621 | target selection |
| ENSDART00000122700 | tenm3 | 0.01323139 | 0.157516404 | 0.267363723 | 0.981020214 | 1.113087141 | target selection |
| ENSDART00000122742 | rorcb | 0.000390105 | -0.773981248 | 0.298893511 | 0.803866913 | 0.643975269 | target selection |
| ENSDART00000122803 | usp9 | 0.023717938 | 0.139967851 | 0.094990222 | 0.369664537 | 0.232490119 | target selection |
| ENSDART00000122889 | myhz1.3 | 0.011565935 | 0.065159767 | 3.686062072 | 4.139837605 | 2.765333679 | target selection |
| ENSDART00000122891 | tmem97 | 0.01179775 | 0.209697033 | 0.237651064 | 0.43725776 | 0.414256644 | target selection |
| ENSDART00000122924 | CABZ01089151.1 | 0.020552635 | 0.66835416 | 0.574150269 | 1.297855118 | 0.40982894 | target selection |
| ENSDART00000123063 | fgf18a | 0.006439631 | 0.196370642 | 0.632761225 | 0.668396752 | 0.413800422 | target selection |
| ENSDART00000123136 | FP236812.4 | 0.090688964 | 0.346817394 | 0.374427758 | 0.928731714 | 0.426600457 | target selection |
| ENSDART00000123338 | kif3a | 0.021156666 | 0.128582487 | 0.488909245 | 0.490973182 | 0.361344362 | target selection |
| ENSDART00000123518 | tuba1b | 0.00416486 | 0.343846477 | 0.74266829 | 1.086564506 | 0.883480954 | target selection |
| ENSDART00000123590 | si:ch211-91p5.3 | 0.006291033 | 0.174508046 | -0.021173312 | 0.906803099 | -0.35666968 | target selection |
| ENSDART00000123607 | cyfip2 | 0.009681558 | 0.234079405 | 0.294840867 | 0.917405124 | 0.783474958 | target selection |
| ENSDART00000123648 | pcdh1a3 | 6.48692E-06 | -0.182322903 | 0.28663259 | 1.138716215 | 1.048655391 | target selection |
| ENSDART00000124040 | insm1a | 1.45694E-05 | 0.50962443 | 1.551047216 | 1.738068624 | 1.104992333 | target selection |
| ENSDART00000124085 | usp18 | 0.007028791 | 0.058285571 | 0.001224867 | 0.967078356 | -0.263223461 | target selection |
| ENSDART00000124220 | nptxrb | 2.18017E-05 | -0.324541852 | -0.000122151 | 0.687818829 | 0.894343302 | target selection |
| ENSDART00000124346 | fn1a | 0.000390105 | 1.458758212 | 2.264590948 | 2.662817487 | 2.142590168 | target selection |
| ENSDART00000124440 | CABZ01029822.1 | 0.001076608 | 0.125194338 | 0.049659662 | 0.670452016 | 0.712162088 | target selection |
| ENSDART00000124485 | pcdh2ab8 | 0.036614379 | 0.093884444 | 0.182265582 | 0.466276394 | 0.290720698 | target selection |
| ENSDART00000125281 | ngfra | 0.001456887 | -0.140591626 | 0.234351775 | 0.618758909 | 0.552257695 | target selection |
| ENSDART00000125590 | adam22 | 0.000133356 | -0.13203003 | 0.438612977 | 0.776065364 | 0.829224171 | target selection |
| ENSDART00000126029 | dhx58 | 0.002259461 | 0.173247162 | -0.097510604 | 2.186112068 | 0.050096401 | target selection |
| ENSDART00000126148 | zmp:0000000912 | 0.027039876 | 0.164571908 | 0.167838799 | 1.993878391 | 0.463546785 | target selection |
| ENSDART00000126253 | fam124b | 0.007297303 | -0.802465606 | 0.105938283 | 0.208441414 | 0.082147827 | target selection |
| ENSDART00000126299 | gdi1 | 0.001249299 | -0.113565456 | 0.226089497 | 0.711067572 | 0.636383115 | target selection |
| ENSDART00000126542 | ppp1r14ba | 6.23756E-05 | 0.329398236 | 1.23193061 | 1.655895613 | 1.416335991 | target selection |
| ENSDART00000126588 | rac3a | 0.001030626 | -0.258496234 | 0.207987177 | 0.575325222 | 0.581016326 | target selection |
| ENSDART00000126873 | myhz2 | 0.000888975 | -2.902978535 | 4.975016398 | 5.62978424 | 4.2721239 | target selection |
| ENSDART00000126991 | ascl1b | 0.003805801 | 0.538037435 | 0.289776062 | 1.627554047 | -0.154081652 | target selection |
| ENSDART00000127214 | tmtc4 | 0.003138731 | -0.125359905 | 0.4024138 | 0.47318852 | 0.32908352 | target selection |
| ENSDART00000127717 | FO704622.1 | 0.044011406 | 0.358399682 | 0.167598174 | 1.49594356 | -0.017716595 | target selection |
| ENSDART00000127854 | ccni | 0.023810358 | 3.365173646 | 3.708027214 | 3.915718651 | 4.117994882 | target selection |
| ENSDART00000127971 | TMEM184B (1 of many) | 0.008188761 | 0.084681099 | 0.512647767 | 0.678333061 | 0.362195553 | target selection |
| ENSDART00000128174 | foxp1b | 0.015925333 | 0.060807795 | 0.283454627 | 0.694455914 | 0.748409918 | target selection |
| ENSDART00000128457 | ephb1 | 0.008075493 | 0.374973875 | 0.620484905 | 1.032855939 | 0.805207117 | target selection |
| ENSDART00000128659 | pcdh2ab10 | 0.033001298 | 0.122813964 | 0.17598472 | 0.479887931 | 0.226887021 | target selection |
| ENSDART00000128742 | CABZ01118678.1 | 0.000370918 | 0.234931395 | 0.748418816 | 0.975879993 | 0.726138214 | target selection |
| ENSDART00000128760 | pacs2 | 0.002951261 | -0.00152054 | 0.063293894 | 0.531537781 | 0.626023304 | target selection |
| ENSDART00000128965 | bcar1 | 0.029418485 | 0.218283788 | 0.254312241 | 0.324642744 | 0.284302999 | target selection |
| ENSDART00000128978 | CR391998.1 | 0.012467303 | 1.262775633 | 1.767194773 | 1.40829971 | 2.265912659 | target selection |
| ENSDART00000129210 | cry1ba | 0.000444051 | -0.306284135 | 0.315253029 | 0.635836839 | 0.449599168 | target selection |
| ENSDART00000129643 | aplnrb | 0.039820338 | -0.038448881 | 0.28030901 | 0.821411842 | 0.34582741 | target selection |
| ENSDART00000129704 | abhd17aa | 0.001225897 | -0.089448965 | -0.057797962 | 0.474531723 | 0.44635249 | target selection |
| ENSDART00000130546 | maptb | 0.000182178 | 0.390340472 | 1.128769701 | 1.393133544 | 1.028418073 | target selection |
| ENSDART00000130554 | isg15 | 0.009167819 | 0.469884875 | 0.09681696 | 3.101380991 | 0.025074196 | target selection |
| ENSDART00000130569 | st8sia5 | 0.00179529 | -0.025137728 | 0.545403948 | 0.761405173 | 0.616910757 | target selection |
| ENSDART00000131431 | sulf2a | 0.000307309 | 0.379800676 | 0.895838573 | 1.99747824 | 2.182238849 | target selection |
| ENSDART00000131721 | fam65b | 0.003648083 | -0.06007994 | 0.061772028 | 0.37892142 | 0.335909594 | target selection |
| ENSDART00000131944 | tagln3b | 0.009948051 | 0.184610888 | 0.818287372 | 1.023242059 | 0.555937355 | target selection |
| ENSDART00000132175 | nadl1.1 | 0.000119513 | 0.709405923 | 0.881560036 | 1.238414697 | 1.046443747 | target selection |
| ENSDART00000132206 | si:dkey-242g16.2 | 0.027067133 | 0.572333551 | 0.964289734 | 1.608250495 | 0.992663233 | target selection |
| ENSDART00000132761 | arhgdig | 0.00124178 | -3.192123378 | 0.713824666 | 1.228568907 | 0.922752193 | target selection |
| ENSDART00000133131 | uba1 | 0.002246206 | 0.147030579 | 0.239356262 | 0.529136799 | 0.293288779 | target selection |
| ENSDART00000133466 | CR847503.1 | 0.009263143 | -0.023677254 | 0.459363169 | 0.719903439 | 0.430626169 | target selection |
| ENSDART00000133473 | CU207343.1 | 0.024547153 | 0.284146912 | 1.104685019 | 1.557001573 | 0.994606199 | target selection |
| ENSDART00000133851 | nyap2b | 6.48692E-06 | -0.344969094 | 0.36816036 | 0.857787268 | 0.825724613 | target selection |
| ENSDART00000134044 | vdac3 | 0.00042933 | -0.155270833 | 0.146773494 | 0.378912201 | 0.402649801 | target selection |
| ENSDART00000134052 | si:dkey-17e16.8 | 0.048722402 | 0.112444287 | 0.152961651 | 0.336755528 | 0.189687147 | target selection |
| ENSDART00000134514 | smarca4a | 0.00733086 | 0.020398578 | 0.110031025 | 0.339984244 | 0.255390677 | target selection |
| ENSDART00000134729 | BX936308.1 | 3.77986E-05 | -0.246065702 | 0.716060471 | 1.529870623 | 1.562652796 | target selection |

| | | | | | | | |
|---|---|---|---|---|---|---|---|
| ENSDART00000134801 | ccdc92 | 0.00194329 | 0.127931369 | 0.677319918 | 0.797788944 | 0.697201329 | target selection |
| ENSDART00000135125 | serp2 | 0.018167343 | 4.40213E-05 | 0.384051284 | 0.40863701 | 0.667539121 | target selection |
| ENSDART00000135147 | si:ch211-218d20.15 | 0.001847721 | -0.111515958 | 0.398515484 | 0.749278715 | 0.887643411 | target selection |
| ENSDART00000135230 | chd7 | 0.009588901 | 0.169349437 | 0.346809261 | 0.458651128 | 0.318074781 | target selection |
| ENSDART00000135248 | anxa13l | 0.005796202 | 2.875521493 | 3.590436866 | 3.752911284 | 3.4448321 | target selection |
| ENSDART00000135399 | phf11 | 0.025606413 | -0.105908519 | 0.331215825 | 2.01559672 | 0.351931859 | target selection |
| ENSDART00000135821 | mfsd6b | 0.000245464 | 0.610408738 | 1.551908888 | 1.729395905 | 1.289342978 | target selection |
| ENSDART00000135989 | cep170aa | 0.001998468 | 0.062401976 | 0.336049788 | 0.4713762 | 0.353973795 | target selection |
| ENSDART00000136103 | ubap2a | 0.017101809 | 0.173955465 | 0.276885256 | 0.421817829 | 0.278699527 | target selection |
| ENSDART00000136141 | kcnq3 | 1.15646E-05 | -0.157652998 | 0.519381195 | 1.385651499 | 1.123132968 | target selection |
| ENSDART00000136439 | bcl7a | 0.000926509 | 0.000990732 | 0.374886928 | 0.452965996 | 0.230358497 | target selection |
| ENSDART00000137228 | si:rp71-80o10.4 | 0.016583954 | -0.005305454 | 1.094390245 | 1.175846782 | 0.577915575 | target selection |
| ENSDART00000137624 | eef2a.2 | 0.029418485 | 2.619194541 | 2.67605773 | 3.551783602 | 3.00375243 | target selection |
| ENSDART00000137903 | BX640512.2 | 0.003143898 | -0.158278671 | -0.01949183 | 1.915495669 | -0.120027797 | target selection |
| ENSDART00000137918 | bbc3 | 0.000309757 | 0.101268325 | 0.767629492 | 0.853650102 | 0.865966102 | target selection |
| ENSDART00000138081 | bcl6ab | 0.000589104 | 0.948060654 | 0.690527167 | 0.988530239 | 1.012261919 | target selection |
| ENSDART00000138321 | parp12b | 0.041240008 | 0.255975647 | 0.107710029 | 0.742615255 | 0.386793554 | target selection |
| ENSDART00000138759 | clta | 0.003329747 | -0.330987711 | -0.247185854 | 1.110067577 | 0.98649153 | target selection |
| ENSDART00000139299 | zgc:162928 | 0.01695739 | 0.559235647 | 0.580335587 | 1.258831581 | 1.081287075 | target selection |
| ENSDART00000139387 | atp1b1b | 0.003609059 | -0.121937317 | 0.340318468 | 0.40085255 | 0.510906972 | target selection |
| ENSDART00000139702 | casp3a | 0.000935288 | 0.559260909 | 1.157211853 | 1.445414898 | 1.128298777 | target selection |
| ENSDART00000139863 | dcbld2 | 0.015156186 | 0.210663919 | 0.284613411 | 0.515685864 | 0.299926066 | target selection |
| ENSDART00000139917 | myha | 0.009294776 | 0.315818594 | 4.091377824 | 5.948901319 | 4.44901869 | target selection |
| ENSDART00000140230 | zgc:123068 | 0.02114031 | 0.653257046 | 0.043755773 | 2.027002482 | 0.024521261 | target selection |
| ENSDART00000140255 | si:dkey-12h9.6 | 0.007190928 | 0.01002873 | 0.192183301 | 0.434559619 | 0.43853246 | target selection |
| ENSDART00000140284 | zbtb16a | 0.028958693 | 2.407975491 | 2.022869932 | 2.341949009 | 2.679474288 | target selection |
| ENSDART00000140301 | TTC9 | 0.001356985 | 0.099722728 | 0.582192427 | 0.813799221 | 0.513650171 | target selection |
| ENSDART00000140586 | col1a1a | 0.05696964 | 0.136049595 | 0.737681947 | 1.130025782 | 0.5982928 | target selection |
| ENSDART00000140908 | gse1 | 0.021503672 | 0.266945251 | 0.781532641 | 1.123032019 | 0.70398488 | target selection |
| ENSDART00000140940 | cluhb | 1.88889E-05 | -1.005907144 | -1.058846619 | 1.005581701 | -0.029667711 | target selection |
| ENSDART00000141237 | si:dkeyp-113d7.10 | 0.010693512 | -0.138953924 | 0.055420663 | 0.3781204 | 0.217616879 | target selection |
| ENSDART00000141278 | si:ch211-14c7.2 | 0.002167167 | 0.247729936 | 0.220022891 | 0.550240269 | 0.375370237 | target selection |
| ENSDART00000141338 | enc1 | 0.008595135 | 0.132252305 | 0.206249243 | 0.415692262 | 0.451357063 | target selection |
| ENSDART00000141367 | si:ch211-238n5.4 | 0.007639785 | -0.027130317 | 0.13157615 | 0.368504617 | 0.417326096 | target selection |
| ENSDART00000141444 | abcg4a | 0.000281821 | -0.060112999 | 0.269266665 | 0.721083006 | 0.688157505 | target selection |
| ENSDART00000141446 | coro1cb | 0.00025494 | -0.332235831 | 0.115994662 | 0.453423489 | 0.231540088 | target selection |
| ENSDART00000141750 | tnnt3b | 0.003035847 | -1.89549374 | 3.616832877 | 4.325157956 | 3.265344737 | target selection |
| ENSDART00000141876 | ftr30 | 0.039458126 | 0.380222613 | 0.277973219 | 1.408194744 | 0.300573986 | target selection |
| ENSDART00000141974 | acsl3b | 0.002176351 | -0.154671373 | 0.25230778 | 0.484219802 | 0.119435773 | target selection |
| ENSDART00000141981 | loxhd1a | 0.006545343 | 0.329259022 | 0.257582221 | 0.952686722 | 1.148017375 | target selection |
| ENSDART00000142155 | myh14 | 0.030748502 | 0.116679043 | 0.189831513 | 0.30854982 | 0.263648695 | target selection |
| ENSDART00000142377 | MEGF9 (1 of many) | 0.000470359 | -0.489828408 | -0.163053003 | 0.463102805 | 0.22210349 | target selection |
| ENSDART00000142919 | ssh1b | 0.007672783 | 0.196097873 | 0.173857746 | 0.465900153 | 0.503515831 | target selection |
| ENSDART00000143697 | si:dkeyp-69e1.8 | 0.006141198 | 0.277815064 | 0.526114648 | 0.670410394 | 0.498924627 | target selection |
| ENSDART00000143819 | VASH1 | 0.001175355 | 0.033504925 | 0.324442057 | 0.595830734 | 0.534870749 | target selection |
| ENSDART00000143999 | tnnt3b | 0.007788648 | 0.546685869 | 4.375596584 | 5.084902064 | 3.992962258 | target selection |
| ENSDART00000144342 | ppp1r14ba | 9.11161E-05 | 0.017178827 | 0.781919568 | 1.244415225 | 1.189050173 | target selection |
| ENSDART00000144681 | anxa13l | 0.000360396 | 0.688342481 | 1.628944564 | 2.065542903 | 1.40538163 | target selection |
| ENSDART00000144737 | srsf3b | 0.029418485 | 0.361810279 | 0.317106803 | 0.304841622 | 0.541582711 | target selection |
| ENSDART00000144766 | nars | 0.003162373 | 0.166803353 | 0.38562143 | 0.598983821 | 0.308867409 | target selection |
| ENSDART00000145065 | NA | 0.00387273 | 4.897906418 | 2.757476262 | 4.60015098 | 5.019792283 | target selection |
| ENSDART00000145124 | dact3a | 0.021766754 | 0.112856383 | 0.770301052 | 1.625107526 | 0.355230326 | target selection |
| ENSDART00000145230 | eef1db | 0.009491292 | 0.569560585 | 0.627404265 | 1.018116547 | 0.789247315 | target selection |
| ENSDART00000145494 | kcnq3 | 5.06657E-05 | -0.150620266 | 0.575528836 | 1.458464727 | 1.126016522 | target selection |
| ENSDART00000145605 | camk2d1 | 0.020107844 | 0.045519482 | 0.325117172 | 0.151136219 | 0.629830606 | target selection |
| ENSDART00000145886 | osbpl3b | 0.023840313 | 0.13475307 | 0.290875362 | 0.450588606 | 0.199350453 | target selection |
| ENSDART00000146106 | nfil3-5 | 5.17084E-05 | 0.032256315 | 1.067374086 | 1.300867348 | 1.052274713 | target selection |
| ENSDART00000146107 | ywhae1 | 0.007536007 | -0.007595908 | 0.375656099 | 0.665999115 | 0.632685234 | target selection |
| ENSDART00000146180 | csrnp2 | 0.003710792 | 0.237147162 | 0.374829692 | 0.697707124 | 0.402573932 | target selection |
| ENSDART00000146247 | zc2hc1a | 0.010853056 | 0.181680565 | 0.536623189 | 0.750568991 | 0.668481158 | target selection |
| ENSDART00000146400 | st8sia5 | 0.004793697 | 0.352731191 | 0.917371468 | 1.067350167 | 0.852063315 | target selection |
| ENSDART00000146517 | dpysl2b | 0.000112531 | 0.345677828 | 1.022911519 | 1.287775094 | 0.946528154 | target selection |
| ENSDART00000146563 | magixa | 0.011667422 | 0.075133988 | 0.355594726 | 0.57567448 | 0.363486006 | target selection |
| ENSDART00000146708 | adar | 0.009288398 | -0.042581428 | -0.004444765 | 0.40535861 | 0.024953366 | target selection |
| ENSDART00000146776 | tspan2b | 0.000107854 | 0.412627689 | 1.44398444 | 1.603529521 | 0.88938646 | target selection |
| ENSDART00000146872 | emid1 | 0.002495957 | 0.014325328 | 0.393704295 | 0.644359098 | 0.692252594 | target selection |
| ENSDART00000147474 | stmn2b | 2.74697E-05 | 0.410261377 | 1.543598694 | 2.044768785 | 1.667647819 | target selection |
| ENSDART00000147481 | phf11 | 0.071106895 | 2.519105146 | 2.056646778 | 4.198159431 | 2.169035406 | target selection |
| ENSDART00000147609 | shc1 | 4.71668E-05 | -0.131375988 | 0.113051171 | 0.56388659 | 0.681815315 | target selection |
| ENSDART00000147625 | pik3r3b | 0.00053559 | -0.257748291 | 0.021181096 | 0.371087748 | 0.412468691 | target selection |
| ENSDART00000147794 | nt5c2l1 | 0.034100602 | 2.1295515 | 2.541950381 | 3.263642166 | 1.818207515 | target selection |
| ENSDART00000148249 | plppr4b | 1.75467E-05 | -0.24256353 | -0.068311603 | 1.098973668 | 1.360337893 | target selection |
| ENSDART00000148280 | prr18 | 0.000193263 | 0.103510486 | 0.905956065 | 1.016662607 | 0.752186459 | target selection |
| ENSDART00000148296 | tnni1d | 0.009288398 | 0.552273501 | 5.279876604 | 5.419924358 | 3.332159231 | target selection |
| ENSDART00000148475 | cspg5a | 2.33589E-05 | 0.090085539 | 0.726711302 | 1.793409667 | 1.632985142 | target selection |

| | | | | | | | |
|---|---|---|---|---|---|---|---|
| ENSDART00000148661 | stat2 | 0.006978095 | 0.11967996 | -0.016653911 | 1.354000143 | 0.023510931 | target selection |
| ENSDART00000148927 | tgfb2 | 0.017007145 | 0.452467242 | 0.589301589 | 0.923363405 | 0.590115295 | target selection |
| ENSDART00000149041 | reep1 | 0.002149847 | -0.249037096 | 0.218537351 | 0.449912583 | 0.370752516 | target selection |
| ENSDART00000149237 | xirp2a | 0.062966527 | 0.190889632 | 0.253883647 | 0.37076536 | 0.19112506 | target selection |
| ENSDART00000149351 | scd | 5.17084E-05 | -0.095125933 | 1.291914984 | 1.511394876 | 0.539585936 | target selection |
| ENSDART00000149684 | si:dkeyp-52c3.2 | 0.035317591 | 2.125770421 | 2.315774699 | 2.351460593 | 2.938761267 | target selection |
| ENSDART00000149730 | stox2a | 0.007236331 | 0.026475093 | 0.234017219 | 0.506304198 | 0.292160739 | target selection |
| ENSDART00000149768 | kif21a | 0.000349997 | -0.021643242 | 0.361851849 | 0.686688714 | 0.546979573 | target selection |
| ENSDART00000149777 | setx | 0.006824859 | 0.020956021 | 0.009824369 | 0.394668282 | 0.077312111 | target selection |
| ENSDART00000149878 | otud4 | 0.000390311 | 0.242358714 | 0.196542793 | 0.87579795 | 0.138730183 | target selection |
| ENSDART00000149956 | apc | 0.012706968 | 0.195183663 | 0.26527048 | 0.385323133 | 0.453680151 | target selection |
| ENSDART00000150128 | foxp1b | 0.020184087 | 0.347545443 | 0.556370195 | 1.342974258 | 1.269825931 | target selection |
| ENSDART00000150148 | slc7a6 | 7.6604E-05 | 0.22923155 | 0.720543979 | 1.05061766 | 1.171565717 | target selection |
| ENSDART00000150362 | si:ch211-197k17.3 | 0.00308528 | -0.145582952 | 0.205377076 | 0.711396584 | 0.681320306 | target selection |
| ENSDART00000150863 | nrp2a | 0.0002611 | 0.543372402 | 0.197731133 | 0.582446066 | 0.760458596 | target selection |
| ENSDART00000151044 | baz2ba | 0.006344596 | 2.371956879 | 3.808025242 | 3.802702579 | 3.968575995 | target selection |
| ENSDART00000151195 | 43166 | 0.003646166 | -0.18277274 | 0.240052097 | 0.74148711 | 0.799803253 | target selection |
| ENSDART00000151717 | 43166 | 0.000507818 | 0.010098896 | 0.251138543 | 0.521642437 | 0.401235935 | target selection |
| ENSDART00000151899 | ldlrb | 0.001252059 | -0.013482381 | 0.739364147 | 1.30137615 | 0.56832803 | target selection |
| ENSDART00000152181 | mapta | 0.000943511 | -0.246945515 | 0.298203412 | 0.945051693 | 0.777575562 | target selection |
| ENSDART00000152284 | si:ch73-301j1.1 | 0.003849593 | -0.68804704 | -0.624114046 | 1.993753311 | -0.48476616 | target selection |
| ENSDART00000152489 | irf3 | 0.006723268 | -0.152502705 | 0.050924712 | 1.421424509 | -0.375867622 | target selection |
| ENSDART00000152636 | clocka | 0.010610046 | 0.008631834 | 0.196030905 | 0.593132542 | 0.426491394 | target selection |
| ENSDART00000152678 | ifit10 | 0.037152675 | 0.50194473 | 0.092674262 | 4.442632616 | 1.169305685 | target selection |
| ENSDART00000153013 | eef1db | 0.018003813 | 0.150654853 | 0.305331563 | 0.633860114 | 0.458033815 | target selection |
| ENSDART00000153087 | bmf2 | 0.016249513 | 0.285623957 | 0.323465948 | 0.348774195 | 0.627801645 | target selection |
| ENSDART00000153212 | adgra1a | 0.000596344 | 0.27394507 | 0.382231602 | 0.735064793 | 0.705504548 | target selection |
| ENSDART00000153217 | eef1db | 9.24337E-05 | 0.666551397 | 1.483640099 | 2.024242417 | 1.323561336 | target selection |
| ENSDART00000153289 | nos1apb | 0.003126045 | -0.207119032 | 0.371227589 | 0.722590517 | 0.685079944 | target selection |
| ENSDART00000153377 | eef1db | 1.1419E-05 | 0.533867077 | 0.950085432 | 1.385417409 | 1.0336151 | target selection |
| ENSDART00000153510 | si:ch73-160l8.2 | 0.032861605 | 0.056068509 | 0.128148598 | 1.161251796 | 0.352706117 | target selection |
| ENSDART00000153562 | ppfia3 | 0.004813371 | -0.022654827 | 0.232304266 | 0.485406891 | 0.444415736 | target selection |
| ENSDART00000153657 | trim3a | 0.002721779 | 0.068235605 | 0.356875561 | 0.585605002 | 0.467600874 | target selection |
| ENSDART00000153700 | BX088707.1 | 0.01037926 | 0.029406694 | 0.3559423 | 0.787563577 | 0.447534651 | target selection |
| ENSDART00000154045 | glg1a | 0.024649569 | 0.211468857 | 0.293330382 | 0.40088818 | 0.351496464 | target selection |
| ENSDART00000154393 | helz2 | 0.000250511 | -0.348999688 | -0.710690348 | 2.383802665 | -0.188541475 | target selection |
| ENSDART00000154719 | hip1 | 0.001124982 | 0.155954348 | 0.357624902 | 0.536540352 | 0.421858349 | target selection |
| ENSDART00000154978 | si:ch211-284f22.3 | 0.034457087 | 0.111486547 | 0.17394927 | 0.442535968 | 0.352663532 | target selection |
| ENSDART00000155094 | si:ch211-286c4.6 | 0.018611393 | 0.931010056 | 1.14507062 | 1.165043802 | 1.805632584 | target selection |
| ENSDART00000155128 | meis3 | 0.088742714 | 1.719560674 | 1.869294835 | 2.789896441 | 2.118152659 | target selection |
| ENSDART00000155190 | maptb | 0.000235238 | 1.096239137 | 1.804532446 | 2.232737802 | 1.717568221 | target selection |
| ENSDART00000155267 | si:ch211-149e23.4 | 0.033633212 | 0.305598533 | 0.378888393 | 0.64366915 | 0.422691793 | target selection |
| ENSDART00000155368 | si:ch73-95l15.3 | 0.006605202 | 1.516126674 | 2.634881995 | 2.849448463 | 2.215584039 | target selection |
| ENSDART00000155464 | ftr73 | 0.005040201 | 0.436406361 | 0.364310996 | 2.049125852 | 0.40110848 | target selection |
| ENSDART00000155610 | tdg.2 | 0.007955678 | 0.198987795 | 0.279928785 | 0.39077816 | 0.364649362 | target selection |
| ENSDART00000155723 | si:ch211-24o10.6 | 0.041249708 | -0.255651916 | 0.580162906 | 3.993132482 | 0.654326546 | target selection |
| ENSDART00000156295 | myhb | 0.001660423 | -2.03199835 | 4.211906798 | 4.339398917 | 2.940685436 | target selection |
| ENSDART00000156535 | vav3b | 2.2034E-05 | -0.155549402 | 0.298871549 | 0.871724785 | 0.959064908 | target selection |
| ENSDART00000156546 | plcxd1 | 0.074703162 | 0.33889602 | 0.161133476 | 0.572020307 | 0.368311625 | target selection |
| ENSDART00000156577 | si:ch211-130m23.5 | 0.001555418 | -0.007378589 | 0.400588111 | 0.725075783 | 0.606240178 | target selection |
| ENSDART00000156690 | ftr88 | 0.038938505 | 0.478641753 | 0.356185618 | 1.950902073 | 0.248066147 | target selection |
| ENSDART00000156765 | soga3a | 0.005830767 | 0.515035881 | 0.806381678 | 1.137120698 | 0.812578505 | target selection |
| ENSDART00000156792 | kif1aa | 0.004591248 | -0.016449727 | 0.304055121 | 0.524314954 | 0.462526752 | target selection |
| ENSDART00000156806 | rps6kl1 | 0.014022061 | 0.03495508 | 0.325476415 | 0.37135553 | 0.249571068 | target selection |
| ENSDART00000157058 | si:dkey-276j7.1 | 0.003257025 | -0.114061291 | 0.35744712 | 0.794710779 | 0.605587625 | target selection |
| ENSDART00000157407 | nacad | 0.00067643 | -0.349281215 | 0.532726398 | 0.824860187 | 0.457523633 | target selection |
| ENSDART00000157475 | coro7 | 0.000780014 | 0.174942149 | 0.053392758 | 0.483358888 | 0.70362148 | target selection |
| ENSDART00000157663 | reln | 0.0021138 | -0.012850615 | -0.065342924 | 0.75286937 | 0.74278543 | target selection |
| ENSDART00000157682 | kat2a | 0.0058872 | 0.094563424 | 0.264392397 | 0.483214757 | 0.33498107 | target selection |
| ENSDART00000157817 | lrrfip1b | 0.000829135 | -0.412751624 | 0.086992217 | 0.168101768 | 0.241775802 | target selection |
| ENSDART00000157920 | BX664622.3 | 0.008776907 | 0.309964523 | 0.2387983 | 0.211520656 | 0.43648414 | target selection |
| ENSDART00000157958 | nav3 | 0.000514549 | 0.289857115 | 0.341415382 | 0.725222646 | 0.845169813 | target selection |
| ENSDART00000158072 | gpc2 | 0.000503033 | 0.147140386 | 0.761040446 | 1.023252339 | 0.612693915 | target selection |
| ENSDART00000158098 | gpr132b | 0.000239262 | -0.335128962 | -0.097182083 | 0.890467313 | 0.848088338 | target selection |
| ENSDART00000158532 | gnb2 | 0.000292971 | -0.185937539 | 0.268112673 | 0.644184639 | 0.701859814 | target selection |
| ENSDART00000158533 | nup50 | 0.010377142 | 0.24455835 | 0.550142856 | 0.78129311 | 0.523300254 | target selection |
| ENSDART00000158563 | wu:fj29h11 | 0.00145023 | 0.059907274 | -0.166860874 | 1.78519307 | 0.017403627 | target selection |
| ENSDART00000158825 | actc1b | 0.011976464 | 0.015935591 | 2.858633574 | 3.729694641 | 2.583924646 | target selection |
| ENSDART00000158837 | dync1h1 | 0.002663982 | 0.179298196 | 0.363338197 | 0.829945857 | 0.598133568 | target selection |
| ENSDART00000159252 | abca3b | 0.000342238 | 0.181429492 | 0.428459166 | 0.640212151 | 0.464148107 | target selection |
| ENSDART00000159316 | mvda | 0.005340573 | 0.177821409 | 1.524901044 | 1.830059782 | 0.927790622 | target selection |
| ENSDART00000159454 | CABZ01044277.1 | 0.01993115 | 0.109016927 | 0.066686129 | 0.335359626 | 0.279394938 | target selection |
| ENSDART00000159673 | dync1i2b | 0.004377247 | 0.025252757 | 0.176262587 | 0.413723725 | 0.300516643 | target selection |
| ENSDART00000159782 | agtpbp1 | 0.000322759 | -0.044706055 | 0.173624177 | 0.473061984 | 0.407831162 | target selection |
| ENSDART00000160107 | CU639413.1 | 0.012966692 | 0.54252914 | 0.409259037 | 1.419043143 | 0.130740048 | target selection |

| | | | | | | | |
|---|---|---|---|---|---|---|---|
| ENSDART00000160154 | si:dkey-238i5.2 | 0.000470962 | -0.200329967 | 0.657530632 | 1.151706484 | 1.107326351 | target selection |
| ENSDART00000160288 | FP236812.6 | 0.067256516 | 0.238382616 | 0.543365019 | 0.866982901 | 0.48841948 | target selection |
| ENSDART00000160444 | si:dkey-95j14.1 | 0.010110237 | -0.172108125 | 0.025439256 | 0.48271623 | 0.286285124 | target selection |
| ENSDART00000160571 | mxe | 0.005032085 | 0.057431509 | -0.237146132 | 2.037062599 | -0.095948656 | target selection |
| ENSDART00000160600 | cecr5 | 0.002653527 | -0.047380691 | 0.036242917 | 0.854821049 | 0.34511846 | target selection |
| ENSDART00000160783 | rorcb | 0.000119513 | -0.657257878 | 0.184912673 | 0.679992874 | 0.484278813 | target selection |
| ENSDART00000160901 | FMNL1 (1 of many) | 0.003432504 | 1.814810207 | 1.982271858 | 2.471309982 | 2.117776194 | target selection |
| ENSDART00000161048 | psmf1 | 0.005446855 | -0.002898908 | -0.064926331 | 0.496331974 | -0.135580746 | target selection |
| ENSDART00000161081 | satb1a | 0.000922422 | -0.172286726 | 0.059256047 | 0.306569639 | 0.367466245 | target selection |
| ENSDART00000161153 | GAN | 0.001865237 | -0.076211127 | 0.283155249 | 0.536837618 | 0.420868128 | target selection |
| ENSDART00000161407 | tnika | 0.002828188 | -0.172678134 | 0.105967748 | 0.465016741 | 0.501980476 | target selection |
| ENSDART00000161566 | b4galt6 | 0.026706504 | 0.155163411 | 0.348743299 | 0.367166825 | 0.307075635 | target selection |
| ENSDART00000161693 | cetn2 | 0.000119513 | -0.42442122 | 0.157217987 | 0.592113306 | 0.77779027 | target selection |
| ENSDART00000162055 | mark1 | 0.005808489 | -0.053725407 | 0.099849875 | 0.364172927 | 0.499618468 | target selection |
| ENSDART00000162459 | scarb2a | 0.014234373 | 0.204566318 | 0.711810949 | 0.821282796 | 0.534817262 | target selection |
| ENSDART00000162529 | zgc:172139 | 0.021599545 | 0.398363371 | 0.414482162 | 0.745277151 | 0.505205904 | target selection |
| ENSDART00000162617 | ppp2cb | 0.012327284 | 0.050413288 | 0.287682373 | 0.42325509 | 0.191699941 | target selection |
| ENSDART00000162761 | CU984600.2 | 0.008281487 | 0.225517614 | -0.442281881 | 2.62985535 | 0.023562363 | target selection |
| ENSDART00000162838 | agrn | 0.003446605 | 0.183430669 | 0.084636207 | 0.451853257 | 0.572854924 | target selection |
| ENSDART00000162915 | dclk2b | 0.000530154 | 0.022821815 | 0.633802822 | 1.144009661 | 0.962163073 | target selection |
| ENSDART00000163526 | mansc1 | 0.000511051 | 0.060022776 | 0.316271416 | 0.600176682 | 0.64715281 | target selection |
| ENSDART00000163680 | hmgcs1 | 0.000409858 | 0.250779539 | 0.717953967 | 2.106714123 | 1.525845318 | target selection |
| ENSDART00000163741 | pwwp2b | 0.008900663 | 0.088274726 | 0.335331052 | 0.428666425 | 0.318380002 | target selection |
| ENSDART00000163930 | znfx1 | 0.000555101 | 0.194042843 | 0.097168789 | 1.523223234 | 0.201252376 | target selection |
| ENSDART00000164038 | SAMD14 | 0.00059038 | -0.096477793 | 0.217514922 | 0.565397199 | 0.769428937 | target selection |
| ENSDART00000164055 | cap2 | 0.01463785 | 0.292123593 | 0.450008213 | 0.705800498 | 0.580755871 | target selection |
| ENSDART00000164298 | RNF122 | 0.025378177 | 0.037145597 | 0.153249015 | 0.432199178 | 0.405886045 | target selection |
| ENSDART00000164585 | mvda | 0.003470173 | -0.034972011 | 1.767545823 | 1.673143467 | 0.929274175 | target selection |
| ENSDART00000164612 | MYH7 (1 of many) | 0.021678375 | 0.128311883 | 3.589180748 | 4.208970977 | 3.051349831 | target selection |
| ENSDART00000164647 | slc12a2 | 0.001104902 | -0.231168401 | 0.356367716 | 0.84043135 | 0.654792579 | target selection |
| ENSDART00000164809 | si:ch73-233f7.5 | 9.56612E-05 | 0.260792098 | 0.492638431 | 0.828767158 | 0.768289306 | target selection |
| ENSDART00000164853 | cnp | 0.01179775 | 1.119904973 | 1.074260191 | 2.53120974 | 1.692544703 | target selection |
| ENSDART00000165066 | eif2ak2 | 0.000768304 | 0.145136508 | 0.290335878 | 1.008511718 | 0.199274727 | target selection |
| ENSDART00000165199 | mapre2 | 0.003404085 | 0.262487966 | 1.260089875 | 1.425017722 | 0.706527865 | target selection |
| ENSDART00000165230 | map4k4 | 0.006674249 | -0.058109942 | 0.098582575 | 0.424133469 | 0.341075529 | target selection |
| ENSDART00000165318 | thsd7bb | 0.000400742 | -0.173014975 | 0.179455909 | 1.051340912 | 1.022924373 | target selection |
| ENSDART00000165484 | jpt1a | 0.001390124 | 0.363755224 | 1.1796439 | 1.211655345 | 0.942198247 | target selection |
| ENSDART00000165656 | mxd3 | 6.23756E-05 | 0.030444632 | 0.928898979 | 1.014066924 | 0.787179484 | target selection |
| ENSDART00000165824 | setdb1b | 0.057995641 | 0.086359323 | 0.217956862 | 0.320141564 | 0.188338727 | target selection |
| ENSDART00000165877 | purg | 0.000570625 | -0.159238984 | -0.087175961 | 1.048436006 | 0.831059287 | target selection |
| ENSDART00000165987 | DST | 0.053793435 | 0.207553536 | 0.19383587 | 0.337156934 | 0.233508941 | target selection |
| ENSDART00000166174 | PARP12 | 0.001662054 | 0.236086028 | 0.041618587 | 1.350827793 | 0.15615804 | target selection |
| ENSDART00000166224 | smyhc2 | 0.001834127 | 1.194842223 | 5.367086788 | 6.852910143 | 4.718916943 | target selection |
| ENSDART00000166242 | NA | 0.084522001 | 0.20194611 | 0.241162188 | 0.512911525 | 0.246398136 | target selection |
| ENSDART00000166268 | YTHDC2 | 0.005459224 | 0.015267861 | -0.127181664 | 0.395706164 | 0.317645369 | target selection |
| ENSDART00000166580 | pak1 | 0.001850572 | 0.254456198 | 0.505135694 | 0.67991388 | 0.523148639 | target selection |
| ENSDART00000166607 | RAPGEF4 (1 of many) | 0.000589961 | -0.233842979 | 0.163001528 | 0.421327636 | 0.212581301 | target selection |
| ENSDART00000166634 | rnf213b | 0.005353377 | 0.322643691 | -0.072602668 | 2.242716781 | -0.029575481 | target selection |
| ENSDART00000166693 | NA | 0.006591698 | 0.184413566 | -0.110459993 | 1.581805418 | 0.290855458 | target selection |
| ENSDART00000166756 | mfsd11 | 0.024397386 | 0.314593735 | 0.230184492 | 0.795700169 | 0.565917277 | target selection |
| ENSDART00000167128 | pargl | 0.012921442 | 0.159346244 | 0.133314978 | 0.619065571 | -0.065913899 | target selection |
| ENSDART00000167219 | pcdh1g26 | 0.001884118 | -0.056136761 | 0.084950376 | 0.428942573 | 0.447913813 | target selection |
| ENSDART00000167226 | pycr1b | 0.004907334 | 0.1083633 | 0.252664475 | 0.730734657 | 0.419591074 | target selection |
| ENSDART00000167322 | si:ch73-233f7.3 | 0.009706658 | -0.088252023 | 0.25632847 | 0.562157447 | 0.392412002 | target selection |
| ENSDART00000167388 | vps33a | 0.033661318 | 0.155328285 | 0.245633599 | 0.343008348 | 0.290252576 | target selection |
| ENSDART00000167613 | hmgcs1 | 0.006978095 | -0.061825835 | 0.154559892 | 0.893983325 | 0.609306628 | target selection |
| ENSDART00000167696 | si:ch73-233f7.4 | 0.012726801 | 0.264747026 | 0.489460093 | 0.625111029 | 0.721718733 | target selection |
| ENSDART00000167947 | CABZ01079480.1 | 0.056544045 | 0.418778383 | 0.234022686 | 1.236946888 | 0.318728833 | target selection |
| ENSDART00000168157 | mkrn2os.2 | 0.002390041 | 0.50354684 | 0.675600001 | 1.137753613 | 0.994948824 | target selection |
| ENSDART00000168167 | rtcb | 0.011929276 | 0.060457492 | 0.250042841 | 0.449158496 | 0.303788278 | target selection |
| ENSDART00000168241 | tubb2b | 0.000133028 | -0.194997441 | 0.285540167 | 0.888424822 | 0.706942534 | target selection |
| ENSDART00000168483 | si:dkey-16p21.7 | 0.019918813 | 0.12768447 | 0.414452748 | 0.453712173 | 0.342486315 | target selection |
| ENSDART00000168542 | arf3a | 0.004338147 | 0.034708072 | 0.783187174 | 0.81852048 | 0.521503928 | target selection |
| ENSDART00000168559 | ttf2 | 0.00064709 | -3.902752419 | 0.990904721 | 1.020977359 | 0.830250106 | target selection |
| ENSDART00000168822 | pabpc4 | 0.000648976 | 1.120879979 | 1.597763229 | 2.131974577 | 1.975353129 | target selection |
| ENSDART00000168876 | si:dkey-81h8.1 | 0.000251123 | -0.18803818 | 0.217329319 | 0.889109393 | 1.046963425 | target selection |
| ENSDART00000168899 | pcdh1g33 | 0.000227894 | -0.406478127 | -0.012838877 | 0.30409891 | 0.306864798 | target selection |
| ENSDART00000168918 | pcdh1gc6 | 0.013984224 | 0.501599317 | 0.826475212 | 1.380988505 | 1.066751642 | target selection |
| ENSDART00000169105 | pcdh1g29 | 1.36144E-05 | -0.285141513 | 0.143203693 | 0.405619816 | 0.452362051 | target selection |
| ENSDART00000169130 | ftr64 | 0.001382674 | 0.270335706 | 0.025822717 | 1.271602536 | 0.119170332 | target selection |
| ENSDART00000169187 | ptpro | 0.001151232 | 0.079888282 | 0.61716785 | 1.250280006 | 1.189522062 | target selection |
| ENSDART00000169341 | nrp1a | 0.010684329 | 0.186448987 | 0.415165621 | 0.563193133 | 0.501195382 | target selection |
| ENSDART00000169648 | NA | 0.003509558 | 0.47490981 | 0.626028668 | 1.036581019 | 0.748938091 | target selection |
| ENSDART00000169667 | ptpro | 1.98934E-05 | -0.163271096 | 0.493460364 | 0.820001756 | 0.696103761 | target selection |
| ENSDART00000169823 | pnpla3 | 0.017079861 | 0.41574021 | 0.387767111 | 0.775977498 | 0.378524498 | target selection |

| | | | | | | | |
|---|---|---|---|---|---|---|---|
| ENSDART00000169845 | pargl | 0.003678598 | 0.069840572 | -0.083042796 | 0.787500652 | -0.250417109 | target selection |
| ENSDART00000170184 | fgd1 | 0.00148037 | 0.020796397 | 0.102196495 | 0.582360556 | 0.487921156 | target selection |
| ENSDART00000170341 | apba1a | 0.002945703 | -0.229826313 | -0.069954525 | 0.548704484 | 0.456234995 | target selection |
| ENSDART00000170405 | CARTPT (1 of many) | 0.010911523 | 1.790273718 | 2.206490975 | 2.680185278 | 2.041251968 | target selection |
| ENSDART00000170571 | dmtn | 0.016814523 | 0.084482265 | 0.234365609 | 0.453911363 | 0.30033427 | target selection |
| ENSDART00000170874 | phldb1b | 0.048701514 | 0.7459988 | 0.426405219 | 0.977480972 | 0.780660286 | target selection |
| ENSDART00000170951 | PXDN | 0.08148621 | 0.459355707 | 0.64772414 | 0.996368549 | 0.498632459 | target selection |
| ENSDART00000171021 | rab3ip | 0.002818463 | 0.020022814 | 0.162794734 | 0.729696081 | 0.694579303 | target selection |
| ENSDART00000171178 | nlrc3l | 0.007264025 | 0.061705987 | 0.093907561 | 0.374016309 | -0.032278843 | target selection |
| ENSDART00000171244 | ssh2b | 0.005083479 | -0.244446722 | -0.139215381 | 0.546963377 | 0.2356201 | target selection |
| ENSDART00000171335 | map4k4 | 0.002682809 | 0.106316233 | 0.336083411 | 0.398966135 | 0.337674059 | target selection |
| ENSDART00000171354 | nt5c2l1 | 0.075016406 | 0.533832785 | 0.693578932 | 1.620342483 | 0.507169588 | target selection |
| ENSDART00000171382 | epb41l3a | 0.001372006 | -0.440465797 | 0.150336785 | 0.618974575 | 0.751984906 | target selection |
| ENSDART00000171433 | tnni1d | 0.008010071 | 0.40952122 | 3.032354782 | 4.009528497 | 2.823338021 | target selection |
| ENSDART00000171624 | Metazoa_SRP | 0.007467603 | 1.040618727 | 2.218519951 | 2.250692356 | 1.867352594 | target selection |
| ENSDART00000171871 | cbfb | 0.02115372 | 0.41950513 | 0.366762581 | 0.672082991 | 0.461647583 | target selection |
| ENSDART00000172022 | dpysl4 | 0.001494008 | 0.270457415 | 0.912051797 | 1.185681882 | 0.874375314 | target selection |
| ENSDART00000173011 | GRXCR1 | 0.000822037 | 0.401804428 | 0.903685752 | 1.561252834 | 1.150514684 | target selection |
| ENSDART00000173052 | map7d2b | 0.000808461 | 0.261433378 | 1.055987677 | 1.048743858 | 0.7752628 | target selection |
| ENSDART00000001313 | rimbp2 | 0.004394152 | -0.29201819 | -0.480722019 | -0.187717263 | 0.029819479 | brain innervation |
| ENSDART00000001805 | csmd2 | 0.014563941 | -0.646636229 | -0.876947398 | -0.300952212 | -0.072675683 | brain innervation |
| ENSDART00000002691 | tspan7b | 0.000775407 | -0.507607245 | -0.383440036 | -0.192864148 | 0.037733122 | brain innervation |
| ENSDART00000002908 | olfm1a | 0.000941975 | -0.612571688 | -0.410284408 | -0.115653646 | 0.220963821 | brain innervation |
| ENSDART00000003825 | cplx2l | 6.48692E-06 | -1.121107993 | -0.523528158 | 0.430342309 | 0.533514858 | brain innervation |
| ENSDART00000004416 | lrp1ba | 1.45694E-05 | -0.680071532 | -0.933155051 | -0.315048731 | 0.19015716 | brain innervation |
| ENSDART00000004548 | barhl1b | 0.009605408 | -2.472825095 | -1.41656094 | -0.268019579 | 0.230183047 | brain innervation |
| ENSDART00000004717 | igf1 | 0.001566135 | -0.688444818 | -0.421253017 | -0.11318285 | 0.107184312 | brain innervation |
| ENSDART00000005086 | atp1a1b | 0.029368278 | -0.434454878 | -0.290341007 | -0.096388542 | -0.096973501 | brain innervation |
| ENSDART00000005929 | ppp3ca | 0.003189261 | -0.536938929 | -0.189476468 | 0.016656951 | 0.377870736 | brain innervation |
| ENSDART00000006380 | tbx3a | 0.004106215 | -0.244591708 | -0.352646993 | -0.144447405 | 0.06961498 | brain innervation |
| ENSDART00000006435 | gpr27 | 0.005338552 | -0.291018325 | -0.294873264 | -0.134468137 | 0.025589784 | brain innervation |
| ENSDART00000006474 | glra4b | 2.84805E-05 | -0.56482245 | -0.497893783 | 0.107334542 | 0.502166045 | brain innervation |
| ENSDART00000006602 | pde4a | 0.001271118 | -0.284660623 | -0.353015908 | -0.053068214 | 0.108985293 | brain innervation |
| ENSDART00000006612 | tbr1b | 2.95145E-05 | -1.166029945 | -1.008207372 | 0.617040877 | 1.169470893 | brain innervation |
| ENSDART00000006619 | rbpms2b | 3.29362E-06 | -0.996572205 | -0.617915271 | -0.145310289 | 0.143289699 | brain innervation |
| ENSDART00000007208 | lrrc4bb | 0.000199258 | -0.502121898 | -0.528156819 | 0.019633372 | 0.067706598 | brain innervation |
| ENSDART00000007308 | wnt10a | 0.000994591 | -1.246823676 | -1.58888825 | -0.322358389 | 0.209755977 | brain innervation |
| ENSDART00000007531 | slit2 | 0.008856124 | -0.281096395 | -0.311205466 | -0.174560483 | 0.038385548 | brain innervation |
| ENSDART00000007584 | snap25a | 0.002371664 | -0.520469397 | -0.457109198 | -0.208714288 | -0.117586032 | brain innervation |
| ENSDART00000007806 | zbtb16a | 0.004364661 | -0.634450415 | -0.731920804 | 0.190781828 | -0.364381887 | brain innervation |
| ENSDART00000007972 | dlgap4b | 0.009649948 | -0.346497047 | -0.558852799 | -0.093125576 | -0.08715572 | brain innervation |
| ENSDART00000008594 | tmem178 | 0.000118072 | -0.626672748 | -0.469011917 | -0.272377327 | 0.052520712 | brain innervation |
| ENSDART00000008711 | gys1 | 0.002746026 | -0.668450567 | -0.607630136 | -0.28295897 | -0.009086585 | brain innervation |
| ENSDART00000008840 | otofa | 0.003654787 | -0.788283647 | -0.975123326 | -0.33101193 | -0.149384457 | brain innervation |
| ENSDART00000009691 | scml4 | 0.001274316 | -0.348233268 | -0.325522885 | -0.113421585 | 0.115178638 | brain innervation |
| ENSDART00000009892 | gabbr1a | 9.33581E-05 | -0.618262887 | -0.683782394 | 0.013008531 | 0.484024166 | brain innervation |
| ENSDART00000010119 | eef1a2 | 7.10385E-05 | -1.586399577 | -1.404358062 | -0.455450934 | 0.174130308 | brain innervation |
| ENSDART00000010199 | fam219ab | 0.022425099 | -0.329176512 | -0.152550548 | -0.14317011 | 0.01066054 | brain innervation |
| ENSDART00000012391 | cabp1a | 0.002341037 | -0.440359676 | -0.517676368 | -0.117661538 | 0.040314239 | brain innervation |
| ENSDART00000012546 | ctbp2a | 0.01793082 | -0.410954451 | -0.209595307 | -0.078015176 | -0.146761341 | brain innervation |
| ENSDART00000012862 | isl2a | 3.28594E-05 | -0.663813396 | 0.011664777 | 0.746201858 | 1.074694305 | brain innervation |
| ENSDART00000013228 | cacna1aa | 8.82236E-05 | -0.425294008 | -0.48842278 | 0.10428355 | 0.42170802 | brain innervation |
| ENSDART00000013311 | grm6a | 5.17084E-05 | -0.718441456 | -0.89272341 | -0.019297194 | 0.441642652 | brain innervation |
| ENSDART00000014897 | srgap1b | 0.002585408 | -0.562468132 | -0.53652608 | -0.21984937 | 0.24230424 | brain innervation |
| ENSDART00000015279 | rtn4rl1a | 0.001754036 | -0.620130019 | -0.765846328 | -0.205912996 | 0.07424375 | brain innervation |
| ENSDART00000015418 | irf2bpl | 0.001028061 | -0.517127602 | -0.432851187 | -0.112506589 | -0.180160421 | brain innervation |
| ENSDART00000015629 | stxbp1a | 0.003413489 | -0.407999477 | -0.254916105 | -0.074938098 | 0.048338374 | brain innervation |
| ENSDART00000016535 | kcns3a | 0.000529346 | -1.299949965 | -1.227726419 | -0.464780559 | 0.150117362 | brain innervation |
| ENSDART00000016916 | gria4b | 0.002016009 | -0.27171917 | -0.41554348 | -0.145968175 | 0.021630803 | brain innervation |
| ENSDART00000017229 | ncam1a | 0.001161091 | -0.359842421 | -0.143522661 | 0.045009487 | 0.122759016 | brain innervation |
| ENSDART00000017309 | ca16b | 0.003700234 | -0.395585068 | -0.492018229 | -0.260071918 | -0.046674115 | brain innervation |
| ENSDART00000017679 | ppp2r2ca | 0.000199258 | -0.437371106 | -0.275733569 | 0.22042039 | 0.361372826 | brain innervation |
| ENSDART00000018351 | zgc:65851 | 8.05022E-05 | -1.669295979 | -1.346342078 | -0.126748353 | 0.319422369 | brain innervation |
| ENSDART00000019140 | rorab | 0.000701449 | -0.499015892 | -0.249117065 | -0.271786747 | 0.004902877 | brain innervation |
| ENSDART00000019294 | si:dkeyp-75b4.9 | 7.89787E-05 | -0.789027106 | -0.841731007 | -0.221050872 | 0.359721563 | brain innervation |
| ENSDART00000019573 | zgc:65894 | 0.000144061 | -0.648921378 | -0.373996254 | 0.357740851 | 0.502243837 | brain innervation |
| ENSDART00000020174 | dynll2b | 0.013618441 | -0.376289244 | -0.003912141 | -0.039237308 | 0.00732876 | brain innervation |
| ENSDART00000020569 | creld1b | 0.022767452 | -0.396401062 | -0.06913614 | 0.015512001 | 0.011512871 | brain innervation |
| ENSDART00000021231 | slmapb | 5.24617E-05 | -0.557559725 | -0.496297491 | -0.195575587 | 0.049672527 | brain innervation |
| ENSDART00000022010 | hivep2b | 0.004657196 | -0.404683257 | -0.428225913 | -0.085969125 | 0.061919909 | brain innervation |
| ENSDART00000022042 | scn8aa | 2.1371E-05 | -0.64882134 | -0.631929626 | 0.169338069 | 0.591046475 | brain innervation |
| ENSDART00000022768 | ak5 | 8.39504E-05 | -0.529999471 | -0.599207091 | -0.317508372 | 0.131326788 | brain innervation |
| ENSDART00000022866 | pisd | 0.0088045 | -0.555411298 | -0.214837183 | -0.086260422 | -0.186182517 | brain innervation |
| ENSDART00000022976 | kctd16b | 1.37519E-05 | -0.982447595 | -1.281976718 | -0.566394426 | 0.110166911 | brain innervation |
| ENSDART00000023038 | dacha | 0.000322945 | -0.511894205 | -0.418991667 | -0.004522839 | 0.112876213 | brain innervation |

| | | | | | | | |
|---|---|---|---|---|---|---|---|
| ENSDART00000023210 | trim13 | 0.005927492 | -0.37704188 | -0.363410115 | -0.204332616 | 0.075535016 | brain innervation |
| ENSDART00000024778 | robo3 | 0.006011043 | -0.327155095 | -0.251732012 | -0.124129181 | 0.058627004 | brain innervation |
| ENSDART00000025031 | pou4f1 | 1.45694E-05 | -0.37458842 | -0.169318799 | 0.774588614 | 1.160187758 | brain innervation |
| ENSDART00000025256 | igfbp2b | 7.89787E-05 | -0.742463234 | -0.107964006 | 0.616733017 | 0.74050304 | brain innervation |
| ENSDART00000025494 | hprt1l | 0.007216648 | -0.364550689 | -0.385937863 | -0.083973479 | -0.129534044 | brain innervation |
| ENSDART00000025583 | fgf8a | 0.004531744 | -0.058331217 | -0.819626176 | 0.05604093 | 0.237151994 | brain innervation |
| ENSDART00000025997 | dip2cb | 0.000892442 | -0.352423277 | -0.192868324 | 0.083423296 | 0.161945092 | brain innervation |
| ENSDART00000026152 | asap2a | 0.002746972 | -0.416411014 | -0.084267369 | -0.025927141 | 0.001189582 | brain innervation |
| ENSDART00000026378 | slc25a6 | 8.89795E-05 | -0.475274891 | -0.471356749 | -0.229323143 | 0.038491163 | brain innervation |
| ENSDART00000026401 | TMEM178B (1 of many) | 0.010150208 | -0.387053719 | -0.214500403 | -0.049460199 | 0.011667684 | brain innervation |
| ENSDART00000027345 | tmem59l | 4.21243E-05 | -0.559867163 | -0.384910477 | -0.005343137 | 0.202377241 | brain innervation |
| ENSDART00000027398 | kcna2a | 2.54845E-05 | -1.558634612 | -1.413069345 | -0.244650937 | 0.296687437 | brain innervation |
| ENSDART00000027463 | hmx4 | 0.005683277 | -0.340009377 | -0.236809718 | -0.106768059 | 0.00156649 | brain innervation |
| ENSDART00000028285 | pgbd5 | 0.001885682 | -0.471148265 | -0.169774685 | 0.263561856 | 0.22277577 | brain innervation |
| ENSDART00000028338 | scamp5a | 0.008022596 | -0.358452184 | -0.354788931 | -0.092970627 | 0.058203665 | brain innervation |
| ENSDART00000028390 | fgf12a | 6.48692E-06 | -0.792701948 | -0.713373555 | -0.153423419 | 0.185286633 | brain innervation |
| ENSDART00000028895 | negr1 | 0.018126285 | -0.362135851 | -0.256191375 | -0.15121136 | -0.084875523 | brain innervation |
| ENSDART00000029457 | sh2d3ca | 0.000340312 | -0.605491696 | -0.301886794 | 0.051143961 | 0.055429229 | brain innervation |
| ENSDART00000030409 | asap1b | 0.006111083 | -0.387126717 | -0.32480613 | -0.179905202 | -0.016544261 | brain innervation |
| ENSDART00000030773 | foxo3a | 0.000701449 | -0.522674186 | -0.346284342 | -0.025883071 | 0.086783781 | brain innervation |
| ENSDART00000031091 | vsnl1a | 1.75467E-05 | -0.845008079 | -0.77079299 | -0.390375819 | 0.072718281 | brain innervation |
| ENSDART00000031167 | tfap2d | 0.000251123 | -0.967562253 | -0.629795841 | 0.638492839 | 1.047009841 | brain innervation |
| ENSDART00000031546 | chrna6 | 2.82487E-05 | -0.487671995 | -0.420962215 | 0.210734418 | 0.46609085 | brain innervation |
| ENSDART00000031937 | diras1a | 0.00058454 | -0.474540918 | -0.418703997 | -0.179232074 | -0.012563291 | brain innervation |
| ENSDART00000032161 | galnt14 | 0.009427844 | -0.529482639 | -0.362873656 | -0.264119496 | -0.01842007 | brain innervation |
| ENSDART00000033361 | ttyh3b | 0.000150409 | -0.3186879 | -0.037617104 | 0.269211477 | 0.394612295 | brain innervation |
| ENSDART00000033657 | grm6b | 0.000670763 | -0.379593806 | -0.511743403 | -0.159835887 | 0.100793439 | brain innervation |
| ENSDART00000034790 | pcp4l1 | 0.000799851 | -0.408039344 | -0.390896541 | -0.213514281 | -0.035791454 | brain innervation |
| ENSDART00000034849 | grin1b | 0.002361523 | -0.325252553 | -0.609188282 | -0.185270574 | 0.122354295 | brain innervation |
| ENSDART00000036015 | ryr1b | 0.000999216 | -0.116788998 | -0.165312146 | 0.372637833 | 0.693396813 | brain innervation |
| ENSDART00000037126 | eno2 | 4.91848E-05 | -0.315372643 | -0.196767149 | 0.300750061 | 0.433471501 | brain innervation |
| ENSDART00000037265 | olfm1b | 0.001442301 | -0.595000689 | -1.108211527 | -0.18099665 | 0.18250463 | brain innervation |
| ENSDART00000037848 | dpp6b | 0.004843495 | -0.469010511 | -0.191315433 | -0.026922326 | -0.081540125 | brain innervation |
| ENSDART00000038294 | tp53inp1 | 0.025035616 | -0.610161281 | 0.029061275 | -0.033185195 | -0.028504863 | brain innervation |
| ENSDART00000038740 | galnt9 | 1.45694E-05 | -0.705450847 | -0.542663541 | -0.098391024 | 0.177029625 | brain innervation |
| ENSDART00000040049 | camk2d2 | 0.002828188 | -0.441100303 | -0.359209869 | -0.184359733 | 0.062709499 | brain innervation |
| ENSDART00000040184 | tenm1 | 0.015961295 | -0.25843006 | -0.314319075 | -0.175496746 | -0.005818614 | brain innervation |
| ENSDART00000040278 | efna2a | 0.000117841 | -0.820985512 | -0.748851824 | -0.238813858 | 0.346525856 | brain innervation |
| ENSDART00000040456 | cdc42bpab | 0.001587567 | -0.484798472 | -0.258449967 | -0.177779388 | 0.139329121 | brain innervation |
| ENSDART00000040500 | tspan9a | 0.025495825 | -0.374305977 | -0.544164723 | -0.123124308 | -0.153915406 | brain innervation |
| ENSDART00000040537 | gjd1a | 0.000656024 | -0.421902855 | -0.214222628 | 0.04546907 | 0.177131603 | brain innervation |
| ENSDART00000040669 | sphkap | 0.000260159 | -0.388395859 | -0.454597188 | 0.021563144 | 0.051812339 | brain innervation |
| ENSDART00000041714 | atp6v0a1b | 0.001487934 | -0.327889206 | -0.302694525 | -0.070700098 | 0.131627227 | brain innervation |
| ENSDART00000041740 | ubl7a | 0.000296518 | -0.21096997 | -0.119372658 | 0.298633763 | 0.574535573 | brain innervation |
| ENSDART00000041800 | epha8 | 0.002472108 | -0.01929649 | -0.346405332 | 0.164805877 | 0.748357168 | brain innervation |
| ENSDART00000041820 | lingo1a | 0.017075627 | -0.3685918 | -0.267056385 | -0.206204269 | -0.004686852 | brain innervation |
| ENSDART00000041869 | grin1a | 0.006472926 | -0.314741638 | -0.554584424 | -0.238417502 | 0.088185899 | brain innervation |
| ENSDART00000042083 | gria4a | 0.0035639 | -0.246573777 | -0.413439839 | -0.038291419 | 0.265641743 | brain innervation |
| ENSDART00000042683 | cadpsb | 2.18017E-05 | -0.417914059 | -0.394489719 | 0.149869704 | 0.381243736 | brain innervation |
| ENSDART00000042972 | srpk1b | 0.001312564 | -0.590887937 | -0.595753709 | -0.259675811 | -0.090086557 | brain innervation |
| ENSDART00000042984 | epha6 | 0.019981901 | -4.335346398 | -2.736601299 | -1.023821359 | 0.479123476 | brain innervation |
| ENSDART00000043312 | srsf5a | 0.000184045 | -0.406457659 | -0.349179029 | 0.276259838 | -0.003669678 | brain innervation |
| ENSDART00000043823 | osbpl10b | 0.014034602 | -0.615302858 | -0.415836047 | -0.293972471 | 0.154234814 | brain innervation |
| ENSDART00000043857 | irx5a | 8.26035E-05 | -0.492824084 | -0.561776224 | 0.072886283 | 0.234832625 | brain innervation |
| ENSDART00000043924 | mpp6b | 0.002539705 | -0.45376092 | -0.273823891 | -0.178237916 | -0.157850955 | brain innervation |
| ENSDART00000044241 | KCNQ2 (1 of many) | 1.2193E-05 | -0.793205862 | -0.43808803 | 0.589879296 | 0.84351704 | brain innervation |
| ENSDART00000044276 | dip2bb | 0.000804407 | -0.182549034 | -0.085935994 | 0.236910154 | 0.379645319 | brain innervation |
| ENSDART00000044371 | tox | 0.000891702 | -0.382184105 | -0.377592563 | -0.085511133 | 0.05287602 | brain innervation |
| ENSDART00000044423 | magi1b | 0.000170813 | -0.521004582 | -0.785194432 | 0.042021364 | 0.52355612 | brain innervation |
| ENSDART00000044678 | GABRA2 (1 of many) | 0.020491546 | -0.18505593 | -0.452041699 | -0.058625322 | 0.015850219 | brain innervation |
| ENSDART00000045086 | prkceb | 0.014103468 | -0.397671937 | -0.107653926 | 0.08012307 | 0.031299867 | brain innervation |
| ENSDART00000045628 | irx6a | 0.000781013 | -0.424070635 | -0.239579413 | 0.17038084 | 0.276181325 | brain innervation |
| ENSDART00000045861 | slc43a2a | 0.000311245 | -0.304475968 | -0.378976343 | 0.111239312 | 0.188992557 | brain innervation |
| ENSDART00000045888 | tkta | 0.001494578 | -0.540178562 | -0.404491163 | -0.145350137 | 0.055411831 | brain innervation |
| ENSDART00000046211 | lnx2a | 0.039476911 | -0.592157981 | -0.322543056 | -0.148137638 | -0.128059395 | brain innervation |
| ENSDART00000046438 | kcnk2b | 0.000775831 | -1.164610191 | -1.614380279 | -0.460863171 | 0.258871322 | brain innervation |
| ENSDART00000046530 | rab42a | 0.000219336 | -0.542354056 | -0.349553972 | -0.104901238 | 0.132258211 | brain innervation |
| ENSDART00000047399 | mmp24 | 4.91848E-05 | -0.337136574 | -0.281370104 | 0.467116358 | 0.594580786 | brain innervation |
| ENSDART00000047569 | igf2b | 5.24617E-05 | -0.538912754 | -0.46572679 | -0.194582206 | 0.102619195 | brain innervation |
| ENSDART00000048855 | mtus1b | 0.002977128 | -0.449558058 | -0.427741832 | 0.166302333 | 0.000701193 | brain innervation |
| ENSDART00000048994 | pbx3b | 0.000910662 | -0.545498331 | -0.428509616 | 0.129533791 | 0.149586816 | brain innervation |
| ENSDART00000049177 | rab6ba | 2.09905E-05 | -0.532131764 | -0.115244957 | 0.417564602 | 0.557714845 | brain innervation |
| ENSDART00000049462 | rab15 | 0.000457208 | -0.425089968 | -0.185335259 | 0.21479615 | 0.289096284 | brain innervation |
| ENSDART00000049992 | syt9b | 2.01848E-05 | -0.672988403 | -0.530192885 | 0.095421952 | 0.328394957 | brain innervation |
| ENSDART00000050037 | chrnb3b | 6.48692E-06 | -0.414911692 | -0.004263798 | 0.444800143 | 0.950014126 | brain innervation |

| | | | | | | | |
|---|---|---|---|---|---|---|---|
| ENSDART00000050077 | sdcbp | 0.000304005 | -0.585127298 | -0.412438306 | -0.175403243 | -0.033038502 | brain innervation |
| ENSDART00000050303 | b3gat2 | 0.001365289 | -0.439065182 | -0.488507748 | -0.270928467 | 0.095807034 | brain innervation |
| ENSDART00000050308 | calm1b | 0.011264952 | -0.359535363 | -0.241046962 | -0.074464366 | 0.015414816 | brain innervation |
| ENSDART00000050311 | rltpr | 0.000161203 | -0.433814029 | -0.24205503 | 0.339832334 | 0.433192877 | brain innervation |
| ENSDART00000050445 | trim2a | 0.004134049 | -0.427473932 | -0.288810529 | -0.127175492 | 0.041370255 | brain innervation |
| ENSDART00000050762 | phactr3b | 0.00849815 | -0.629592387 | -0.363049766 | -0.266715209 | -0.083768273 | brain innervation |
| ENSDART00000051182 | arhgap4b | 0.003900528 | -0.41225839 | -0.320516515 | -0.225939723 | -0.055826953 | brain innervation |
| ENSDART00000051693 | irx4a | 0.000591404 | -1.596922819 | -2.014349322 | 0.164610846 | 0.924604512 | brain innervation |
| ENSDART00000052113 | lingo1b | 0.029426879 | -0.337763041 | -0.148396867 | -0.080123582 | -0.033848116 | brain innervation |
| ENSDART00000052346 | gnao1b | 0.000112893 | -0.534691372 | -0.39418636 | -0.143300564 | 0.218777626 | brain innervation |
| ENSDART00000052802 | calb2b | 3.76305E-05 | -1.334486374 | -1.407708939 | -0.475837952 | 0.218368357 | brain innervation |
| ENSDART00000054137 | igfbp5b | 0.000573039 | -0.49960111 | -0.321727345 | 0.242454388 | 0.365417817 | brain innervation |
| ENSDART00000054452 | dlgap1b | 0.000993187 | -0.55212867 | -0.830902632 | -0.186012386 | 0.281181047 | brain innervation |
| ENSDART00000054552 | cdh8 | 0.000956736 | -0.482196874 | -0.526834841 | -0.273874965 | 0.0505098 | brain innervation |
| ENSDART00000054833 | rgs11 | 5.42724E-05 | -0.582556718 | -0.789821251 | -0.427532995 | 0.127688375 | brain innervation |
| ENSDART00000054989 | fscn1b | 0.000261639 | -0.529299226 | -0.417661269 | 0.066759573 | 0.494675778 | brain innervation |
| ENSDART00000055038 | rybpa | 0.00042906 | -0.189184893 | -0.138456944 | 0.268105644 | 0.373370341 | brain innervation |
| ENSDART00000055269 | gng13b | 1.11583E-05 | -0.70656822 | -0.547882014 | 0.352208471 | 0.531882213 | brain innervation |
| ENSDART00000055936 | isl2b | 4.91848E-05 | -1.527471977 | -1.312125436 | 0.470803449 | 0.853726063 | brain innervation |
| ENSDART00000056138 | igsf8 | 0.001004207 | -0.41438699 | -0.276734989 | -0.097715498 | 0.052342087 | brain innervation |
| ENSDART00000056328 | elovl4b | 0.005164205 | -0.472600721 | -0.194626997 | -0.10087467 | 0.110456628 | brain innervation |
| ENSDART00000056369 | cadm2a | 0.04906434 | -0.177754586 | -0.327047473 | -0.068598077 | -0.03429359 | brain innervation |
| ENSDART00000056639 | faim2a | 0.008797915 | -0.345615546 | -0.285647372 | -0.169343793 | 0.000937504 | brain innervation |
| ENSDART00000056671 | brinp2 | 8.25087E-05 | -0.564206507 | -0.55260615 | -0.353392227 | 0.031594069 | brain innervation |
| ENSDART00000056885 | CU929046.1 | 2.74697E-05 | -0.462145697 | -0.331087748 | 0.285634599 | 0.389113052 | brain innervation |
| ENSDART00000057369 | igfbp5a | 0.010190417 | -0.503493701 | -0.12557648 | 0.011432007 | -0.035083178 | brain innervation |
| ENSDART00000058736 | grm4 | 8.05022E-05 | -0.475368296 | -0.757934708 | -0.299132614 | 0.099413724 | brain innervation |
| ENSDART00000058829 | scrt1b | 0.000688325 | -0.466509454 | -0.564612343 | -0.054278048 | 0.250358448 | brain innervation |
| ENSDART00000058877 | rap2ab | 8.2583E-05 | -0.786170126 | -0.835369537 | -0.021239814 | 0.42494962 | brain innervation |
| ENSDART00000059179 | nptxra | 0.000962944 | -0.572157626 | -0.495039684 | -0.356374901 | -0.015079326 | brain innervation |
| ENSDART00000059228 | vil1 | 0.006605202 | -0.324461409 | -0.136026611 | -0.08718702 | 0.103993243 | brain innervation |
| ENSDART00000059631 | BX936415.1 | 0.005303358 | -0.988751505 | -1.218737364 | -0.343900834 | 0.242173034 | brain innervation |
| ENSDART00000059841 | si:ch211-257p13.3 | 1.18742E-05 | -0.713606094 | -0.675183925 | 0.07538087 | 0.401418787 | brain innervation |
| ENSDART00000059869 | adra2a | 0.000142155 | -1.042718178 | -0.710015995 | -0.232479367 | 0.220270649 | brain innervation |
| ENSDART00000060051 | fgf14 | 0.028223243 | -0.493332703 | -0.385375543 | -0.154886482 | -0.098910242 | brain innervation |
| ENSDART00000060160 | calb2a | 1.97197E-05 | -1.240040436 | -1.237007367 | -0.232768326 | 0.397108945 | brain innervation |
| ENSDART00000060625 | lgi3 | 6.84125E-05 | -0.698992604 | -0.869112792 | -0.10986795 | 0.409134057 | brain innervation |
| ENSDART00000062697 | gfra2a | 0.003719587 | -0.450482933 | -0.554254218 | -0.069449306 | -0.130685043 | brain innervation |
| ENSDART00000062874 | atp1b3b | 0.000182199 | -0.415562805 | -0.319287363 | -0.078895256 | 0.137689897 | brain innervation |
| ENSDART00000062887 | disp2 | 2.923E-05 | -0.441631902 | -0.296469901 | 0.444242177 | 0.781646932 | brain innervation |
| ENSDART00000063783 | itm2ca | 0.000867732 | -0.362486731 | -0.282712517 | -0.07258841 | 0.153576522 | brain innervation |
| ENSDART00000063825 | sprn | 0.013488739 | -0.451586162 | -0.272184034 | -0.06406637 | 0.092302795 | brain innervation |
| ENSDART00000064672 | NA | 5.42724E-05 | -1.010331454 | -0.756315191 | -0.126164495 | 0.278123254 | brain innervation |
| ENSDART00000064833 | mafaa | 0.002047801 | -0.957052225 | -1.143383392 | -0.379735148 | 0.067354966 | brain innervation |
| ENSDART00000064902 | ssbp4 | 0.000263251 | -0.349245142 | -0.241492171 | 0.147485085 | 0.214650497 | brain innervation |
| ENSDART00000065159 | zgc:158291 | 3.4928E-05 | -1.165646756 | -0.784542684 | 0.443310678 | 0.751149417 | brain innervation |
| ENSDART00000065372 | kcnj3b | 9.03292E-05 | -0.605628976 | -0.778191304 | -0.104344748 | 0.360074992 | brain innervation |
| ENSDART00000065599 | cadm1a | 0.004110692 | -0.54681119 | -0.778847768 | -0.187580903 | 0.116530415 | brain innervation |
| ENSDART00000065929 | hs6st3b | 0.000935288 | -0.878348455 | -0.942691234 | -0.255367675 | 0.253670478 | brain innervation |
| ENSDART00000066192 | glra2 | 0.00133046 | -0.5075286 | -0.700662657 | -0.376893165 | 0.421840665 | brain innervation |
| ENSDART00000066294 | cdk5r1b | 7.6604E-05 | -0.511562763 | -0.495240798 | 0.085364893 | 0.188592346 | brain innervation |
| ENSDART00000066372 | id4 | 4.11188E-05 | -0.719733282 | -0.518590484 | 0.441847928 | 0.655214602 | brain innervation |
| ENSDART00000066839 | slc35g2b | 7.55931E-05 | -0.671846523 | -0.750415444 | -0.309110396 | 0.04424778 | brain innervation |
| ENSDART00000067514 | rbpms2a | 4.11188E-05 | -1.175610868 | -1.156235961 | 0.045281157 | 0.593799851 | brain innervation |
| ENSDART00000067531 | syn2a | 8.82236E-05 | -0.50370548 | -0.414636803 | -0.227904446 | 0.132370635 | brain innervation |
| ENSDART00000073919 | kcnc1b | 2.71726E-05 | -0.758440974 | -0.784989596 | -0.33092052 | 0.322314215 | brain innervation |
| ENSDART00000073950 | olfm1a | 7.10385E-05 | -0.852827208 | -0.587563633 | 0.013637392 | 0.435090193 | brain innervation |
| ENSDART00000074036 | rcvrna | 0.035967167 | -0.553376367 | -0.06531313 | -0.11758335 | -0.07626787 | brain innervation |
| ENSDART00000074362 | pcdh18b | 0.00033039 | -0.440881831 | -0.447646282 | -0.227758863 | 0.176890552 | brain innervation |
| ENSDART00000074543 | hs3st4 | 0.007620644 | -0.453389419 | -0.572006435 | -0.310659668 | 0.002760951 | brain innervation |
| ENSDART00000074678 | chrnb3a | 4.11188E-05 | -0.683036397 | -0.585209438 | -0.069659384 | 0.207417172 | brain innervation |
| ENSDART00000074698 | opn3 | 0.003138642 | -0.589287895 | -0.235981184 | 0.224725485 | -0.032745726 | brain innervation |
| ENSDART00000074718 | spire1b | 0.006352299 | -0.446651583 | -0.478750814 | -0.155393912 | -0.07838085 | brain innervation |
| ENSDART00000074997 | CU302436.3 | 0.001679054 | -0.857583363 | -0.990723788 | -0.187854306 | 0.199234775 | brain innervation |
| ENSDART00000075123 | pcp4a | 4.91848E-05 | -0.598752351 | -0.632528063 | -0.40265776 | -0.022235336 | brain innervation |
| ENSDART00000075278 | atp1b4 | 0.008495106 | -0.758712842 | -0.43098592 | -0.240547748 | -0.077197984 | brain innervation |
| ENSDART00000075340 | eef1a1b | 2.23813E-05 | -0.66677551 | -0.473919347 | -0.344351157 | 0.106841151 | brain innervation |
| ENSDART00000075749 | ppp2r2ca | 0.000117007 | -0.492014929 | -0.268923214 | 0.328067214 | 0.425314748 | brain innervation |
| ENSDART00000075918 | pcmtd2 | 0.00393919 | -0.427264647 | -0.257111837 | -0.149394946 | 0.062644737 | brain innervation |
| ENSDART00000076946 | PDE4DIP | 0.000299141 | -0.575491599 | -0.478988093 | -0.210265098 | -0.157443712 | brain innervation |
| ENSDART00000077216 | astn1 | 0.002342395 | -0.280184115 | -0.378831195 | -0.109389357 | 0.054556021 | brain innervation |
| ENSDART00000077222 | ldlrad4b | 0.002766629 | -0.327819788 | -0.475734059 | -0.025614815 | 0.028486052 | brain innervation |
| ENSDART00000078079 | pcnxl2 | 0.000511051 | -0.518593139 | -0.325199094 | 0.020646332 | 0.403616211 | brain innervation |
| ENSDART00000078115 | sdhda | 0.002274284 | -0.602799581 | -0.148637611 | 0.027660422 | 0.175615096 | brain innervation |
| ENSDART00000078232 | cdh10a | 0.005048068 | -0.495858439 | -0.491434243 | -0.331011431 | 0.033804718 | brain innervation |

| | | | | | | | |
|---|---|---|---|---|---|---|---|
| ENSDART00000078561 | SPTBN4 | 0.000403881 | -0.239897718 | -0.33102434 | 0.293671093 | 0.445086753 | brain innervation |
| ENSDART00000078856 | dlg3 | 0.001948843 | -0.31041276 | -0.361138693 | -0.191257864 | 0.041004354 | brain innervation |
| ENSDART00000079092 | si:dkey-261i16.5 | 0.004123458 | -0.526813533 | -0.563690076 | -0.369398724 | 0.061616577 | brain innervation |
| ENSDART00000079138 | ptenb | 0.013692958 | -0.337433521 | -0.277401842 | -0.145004066 | -0.034401548 | brain innervation |
| ENSDART00000079144 | ptenb | 0.00889295 | -0.303152595 | -0.238217531 | -0.001915727 | -0.031063744 | brain innervation |
| ENSDART00000079443 | GABRA2 (1 of many) | 0.000263077 | -0.623319426 | -0.542833879 | -0.157704672 | 0.138036393 | brain innervation |
| ENSDART00000079840 | rorca | 0.006111083 | -0.65562057 | -0.064052995 | 0.161319682 | 0.103804992 | brain innervation |
| ENSDART00000080064 | CR847953.1 | 0.004084205 | 0.050009503 | -0.040451977 | 0.259802571 | 0.438904414 | brain innervation |
| ENSDART00000080486 | ywhag1 | 1.75467E-05 | -1.02392497 | -0.686234935 | -0.010129168 | 0.2177797 | brain innervation |
| ENSDART00000080808 | six3a | 0.000687981 | -0.437705937 | -0.430575558 | -0.140593988 | -0.07114268 | brain innervation |
| ENSDART00000080927 | snap25b | 0.001998235 | -0.418114286 | -0.385475041 | -0.102955296 | 0.076752829 | brain innervation |
| ENSDART00000081140 | CT990561.1 | 0.000401616 | -0.348057632 | -0.380510005 | -0.109063338 | 0.180807023 | brain innervation |
| ENSDART00000082301 | myrip | 0.000993187 | -0.137038516 | -0.238384017 | 0.084718693 | 0.41131665 | brain innervation |
| ENSDART00000082745 | EMB | 0.000245083 | -0.651102521 | -0.703010006 | -0.074268357 | 0.192993872 | brain innervation |
| ENSDART00000082842 | JPH3 (1 of many) | 0.000165745 | -0.540363483 | -0.227235849 | 0.34555956 | 0.43863208 | brain innervation |
| ENSDART00000083040 | hs3st2 | 0.000442216 | -1.059711916 | -1.469328404 | -0.190665179 | 0.30791737 | brain innervation |
| ENSDART00000083066 | asphd2 | 0.000463544 | -0.487940941 | -0.316881347 | -0.126036033 | 0.003992278 | brain innervation |
| ENSDART00000083569 | oaz2b | 2.90186E-05 | -0.447559352 | -0.669337328 | -0.007273423 | 0.348858251 | brain innervation |
| ENSDART00000083670 | CABZ01041604.1 | 0.025197766 | -0.549505735 | -0.427354636 | -0.238335089 | -0.140824262 | brain innervation |
| ENSDART00000084014 | cplx1 | 2.54845E-05 | -0.741807214 | -0.862001833 | -0.211893904 | 0.449532765 | brain innervation |
| ENSDART00000084119 | si:ch1073-44g3.1 | 0.001946729 | -0.384776638 | -0.202587046 | 0.01646479 | 0.134112994 | brain innervation |
| ENSDART00000084264 | adcy2a | 0.000976529 | -0.574108823 | -0.494423944 | 0.049230433 | 0.113035845 | brain innervation |
| ENSDART00000084354 | cpeb3 | 0.004463264 | -0.1650884 | -0.603027317 | 0.172363697 | 0.2398868 | brain innervation |
| ENSDART00000084355 | zgc:165481 | 0.000683159 | -0.665978496 | -0.883055142 | -0.199024252 | -0.082238068 | brain innervation |
| ENSDART00000084730 | zgc:162160 | 1.57453E-05 | -0.758204424 | -0.82447566 | -0.164736838 | 0.207922006 | brain innervation |
| ENSDART00000084890 | si:ch211-284e13.4 | 0.000766042 | -0.124688162 | -0.262617592 | 0.237393976 | 0.369141192 | brain innervation |
| ENSDART00000085210 | cacna1ha | 0.001618544 | -0.413195046 | -0.405391107 | -0.236177845 | 0.006546496 | brain innervation |
| ENSDART00000085253 | mid1 | 0.001365289 | -0.357661223 | -0.330080838 | -0.002902011 | 0.078651096 | brain innervation |
| ENSDART00000085472 | grm2a | 0.000638552 | -0.506007863 | -0.654883229 | -0.268518199 | 0.087304206 | brain innervation |
| ENSDART00000085565 | capn15 | 0.002791482 | -0.382220369 | -0.426351921 | -0.021041552 | 0.033591442 | brain innervation |
| ENSDART00000085573 | rgs7bpa | 0.003138642 | -0.750985181 | -0.701592527 | -0.219110071 | -0.146877341 | brain innervation |
| ENSDART00000085743 | AL935194.1 | 0.002127025 | -0.774745358 | -0.68914542 | -0.32626184 | 0.039334462 | brain innervation |
| ENSDART00000086617 | gabbr2 | 1.88318E-05 | -0.577324776 | -0.926125922 | -0.433004274 | 0.052999603 | brain innervation |
| ENSDART00000086797 | adgrl3.1 | 0.00088836 | -0.114053477 | -0.068166121 | 0.465121947 | 0.57391191 | brain innervation |
| ENSDART00000087118 | xylt1 | 0.004031897 | -0.272537601 | -0.39007956 | -0.067871098 | 0.053209242 | brain innervation |
| ENSDART00000087148 | cbln4 | 0.000138972 | -0.802680954 | -0.769511703 | -0.146403482 | 0.304907793 | brain innervation |
| ENSDART00000087191 | mark4a | 0.000877772 | -0.415170271 | -0.372344968 | 0.042679549 | 0.108705644 | brain innervation |
| ENSDART00000087280 | cacnb3a | 0.000177237 | -0.323388935 | -0.465191463 | 0.268224123 | 0.533679905 | brain innervation |
| ENSDART00000087426 | bcl11aa | 3.5708E-05 | -0.367520296 | -0.334252656 | 0.232372171 | 0.511638417 | brain innervation |
| ENSDART00000087441 | GFOD1 | 0.020470078 | -0.421601041 | -0.219799435 | -0.160457067 | 0.058971995 | brain innervation |
| ENSDART00000087586 | c2cd4a | 0.000100528 | -0.882018106 | -1.482044104 | -0.368555759 | 0.13043866 | brain innervation |
| ENSDART00000087857 | unc5db | 0.000103226 | -0.464548804 | -0.771046574 | -0.267052184 | 0.0901494 | brain innervation |
| ENSDART00000088159 | nrxn1a | 0.00486688 | -0.55423697 | -0.307440551 | 0.007663974 | -0.036725507 | brain innervation |
| ENSDART00000088178 | nrxn1a | 0.000255127 | -0.424942944 | -0.357890595 | -0.01594532 | 0.150075885 | brain innervation |
| ENSDART00000088199 | zgc:162707 | 0.004813371 | -0.481388546 | -0.51007365 | -0.179950315 | -0.039369231 | brain innervation |
| ENSDART00000088249 | hcn4l | 0.001402519 | -0.487910369 | -0.340719397 | 0.02727434 | 0.039415999 | brain innervation |
| ENSDART00000088833 | si:ch73-233f7.1 | 0.00031728 | -1.214678988 | -0.552820604 | -0.229157577 | 0.532052231 | brain innervation |
| ENSDART00000089574 | tub | 0.003777754 | -0.338880988 | -0.401453479 | -0.101730157 | -0.038037712 | brain innervation |
| ENSDART00000089699 | prrt1 | 0.000415146 | -0.603201784 | -0.702306626 | -0.280611263 | -0.047540659 | brain innervation |
| ENSDART00000089867 | ppp2r2cb | 0.018790846 | -0.452836704 | -0.44451904 | -0.189867818 | -0.082146078 | brain innervation |
| ENSDART00000090019 | zeb2b | 0.016845819 | -0.694941899 | -0.428168304 | -0.296660745 | 0.024845807 | brain innervation |
| ENSDART00000090306 | xpr1a | 0.001093301 | -0.158299732 | -0.301120383 | 0.179396195 | 0.424625304 | brain innervation |
| ENSDART00000090397 | kiaa1549la | 4.91848E-05 | -0.446832614 | -0.575803117 | -0.232865535 | 0.10265337 | brain innervation |
| ENSDART00000090844 | zgc:153018 | 0.006825939 | -0.414733046 | -0.080390611 | -0.214556251 | -0.032786964 | brain innervation |
| ENSDART00000091004 | pcdh1a | 0.000227894 | -0.630879138 | -0.627674521 | -0.227167517 | 0.17418403 | brain innervation |
| ENSDART00000091489 | ppp1r9bb | 0.000117869 | -0.563706295 | -0.346664158 | -0.080794638 | 0.037246954 | brain innervation |
| ENSDART00000091664 | apc2 | 0.011436776 | -0.58155429 | -0.22615315 | 0.136652181 | 0.006999355 | brain innervation |
| ENSDART00000091726 | fam78ba | 4.91848E-05 | -0.542782218 | -0.34207093 | 0.322203074 | 0.384590985 | brain innervation |
| ENSDART00000091818 | tulp4b | 4.52852E-05 | -0.531343855 | -0.277019809 | 0.128892269 | 0.551121944 | brain innervation |
| ENSDART00000091955 | nrxn2b | 0.004348675 | -0.363238969 | -0.38605701 | -0.167716958 | 0.046918181 | brain innervation |
| ENSDART00000092182 | ppm1la | 0.000340312 | -0.411282562 | -0.137536548 | 0.156346356 | 0.193635075 | brain innervation |
| ENSDART00000092250 | btbd11a | 0.005509984 | -0.250819977 | -0.42148278 | -0.144893414 | 0.093892491 | brain innervation |
| ENSDART00000092257 | PLD5 | 5.26677E-05 | -0.97635635 | -0.828911686 | 0.040708181 | 0.36058098 | brain innervation |
| ENSDART00000092665 | srebf1 | 0.004619636 | -0.417658309 | -0.274360919 | 0.114636817 | 0.022214824 | brain innervation |
| ENSDART00000093310 | celf5a | 0.002632159 | -0.268154664 | -0.455004312 | 0.014815598 | 0.070405312 | brain innervation |
| ENSDART00000097330 | dnm1b | 4.69744E-05 | -0.51283412 | -0.480365631 | 0.101619237 | 0.531582232 | brain innervation |
| ENSDART00000097695 | cntnap3 | 0.004847659 | -0.160493697 | -0.529313782 | -0.088115665 | 0.248028839 | brain innervation |
| ENSDART00000097935 | si:dkey-226m8.10 | 0.000254175 | -0.692305988 | -0.817395111 | -0.44855289 | 0.050175879 | brain innervation |
| ENSDART00000098082 | GJD2 (1 of many) | 7.52983E-05 | -0.795483498 | -0.746707283 | 0.031300804 | 0.456345915 | brain innervation |
| ENSDART00000098311 | KCNJ4 | 0.008555913 | -0.486217707 | -0.434414335 | -0.027097558 | 0.056787682 | brain innervation |
| ENSDART00000098667 | camk2b1 | 0.000104487 | -0.575252799 | -0.701444666 | -0.414382214 | 0.077165391 | brain innervation |
| ENSDART00000098727 | svopa | 0.0001406 | -0.499297278 | -0.394465259 | 0.04106692 | 0.031976449 | brain innervation |
| ENSDART00000098840 | ralgps1 | 0.000405977 | -0.525377749 | -0.528705894 | -0.140236147 | 0.156796103 | brain innervation |
| ENSDART00000098859 | neurod6a | 0.000108364 | -0.523659161 | -0.308368621 | 0.583701544 | 0.979244768 | brain innervation |
| ENSDART00000099019 | tmem91 | 2.12106E-05 | -0.661722152 | -0.699132453 | -0.1920109 | -0.057277853 | brain innervation |

| | | | | | | | |
|---|---|---|---|---|---|---|---|
| ENSDART00000099049 | NA | 0.000503033 | -0.472159773 | -0.420532241 | -0.250637325 | -0.02437969 | brain innervation |
| ENSDART00000099283 | dalrd3 | 0.001372006 | -0.272196688 | -0.367831508 | -0.108158836 | 0.07067938 | brain innervation |
| ENSDART00000099325 | si:dkey-27p18.5 | 3.38347E-05 | -0.821017867 | -0.681399645 | -0.140207831 | 0.113491694 | brain innervation |
| ENSDART00000099607 | slc6a17 | 0.006359324 | -0.493487825 | -0.358976735 | -0.156955324 | 0.010739948 | brain innervation |
| ENSDART00000099872 | slc17a6b | 0.000184957 | -0.645728405 | -0.242319052 | 0.108083805 | 0.273480994 | brain innervation |
| ENSDART00000099934 | kcnc1a | 0.023562909 | -0.987934197 | -1.510396524 | -0.387995788 | 0.170469595 | brain innervation |
| ENSDART00000100022 | H2AFX (1 of many) | 0.004391006 | -0.512580891 | -0.39296931 | -0.220510282 | -0.111022185 | brain innervation |
| ENSDART00000100103 | acss2l | 0.006451154 | -0.342716857 | -0.308956489 | -0.115247207 | 0.030743328 | brain innervation |
| ENSDART00000100322 | kcnh5b | 4.91848E-05 | -0.532935362 | -0.444322241 | -0.174846578 | 0.219995697 | brain innervation |
| ENSDART00000100444 | fam19a5a | 8.2583E-05 | -0.63242802 | -0.536601669 | -0.265992198 | 0.12320759 | brain innervation |
| ENSDART00000100596 | pcdh1a6 | 0.018324408 | -0.483816366 | -0.257584823 | -0.06041051 | 0.00168094 | brain innervation |
| ENSDART00000100658 | esrra | 8.74195E-05 | -0.571475474 | -0.395296451 | 0.169003635 | 0.212368184 | brain innervation |
| ENSDART00000100743 | cttnbp2 | 0.001176693 | -0.40355339 | -0.391763552 | 0.009167842 | 0.081098 | brain innervation |
| ENSDART00000101070 | dachd | 0.00434437 | -0.296270747 | -0.351582482 | -0.130791564 | -0.009393292 | brain innervation |
| ENSDART00000101134 | khdrbs2 | 0.003070422 | -0.661640499 | -0.469301828 | -0.265343985 | -0.029039929 | brain innervation |
| ENSDART00000101477 | emp1 | 0.0002611 | -0.241905322 | -0.037268798 | 0.203715295 | 0.377786296 | brain innervation |
| ENSDART00000101603 | kidins220b | 0.004633928 | -0.01985848 | -0.087358266 | 0.173253678 | 0.374974923 | brain innervation |
| ENSDART00000101627 | IGLON5 | 0.000503033 | -0.423175903 | -0.742308046 | -0.313681293 | 0.102626627 | brain innervation |
| ENSDART00000101631 | satb1b | 0.000172812 | -0.360112936 | -0.384116906 | 0.134197196 | 0.481946288 | brain innervation |
| ENSDART00000101653 | CU639469.1 | 2.18017E-05 | -0.529716959 | -0.505372118 | -0.050085906 | 0.195515782 | brain innervation |
| ENSDART00000101658 | ppp1r1b | 0.001434846 | -0.639757199 | -0.267805539 | -0.098160681 | -0.228099263 | brain innervation |
| ENSDART00000102368 | grin1a | 4.69744E-05 | -0.57807298 | -0.694484674 | -0.374619925 | 0.025284051 | brain innervation |
| ENSDART00000102455 | gucy1a3 | 0.001644836 | -0.52073984 | -0.506169014 | -0.27571082 | 0.101070807 | brain innervation |
| ENSDART00000102562 | ankrd10b | 0.000614271 | -0.585231519 | -0.124121182 | 0.139197554 | 0.179030577 | brain innervation |
| ENSDART00000102782 | gria2a | 0.000180202 | -0.472482927 | -0.595061919 | -0.203521695 | 0.050111807 | brain innervation |
| ENSDART00000103043 | nsfa | 0.000872721 | -0.472055845 | -0.418381067 | -0.313792962 | -0.037512478 | brain innervation |
| ENSDART00000103151 | dlgap3 | 0.004847659 | -0.304185948 | -0.430889474 | -0.103996234 | 0.188728565 | brain innervation |
| ENSDART00000103407 | tmem245 | 0.001938223 | -0.353429601 | -0.331655767 | -0.089201446 | 0.066435984 | brain innervation |
| ENSDART00000103750 | fam131bb | 0.000282529 | -0.555314971 | -0.519589863 | -0.143114917 | 0.156539518 | brain innervation |
| ENSDART00000104135 | gabra5 | 1.15646E-05 | -0.398056951 | -0.510180941 | 0.262181183 | 0.751061332 | brain innervation |
| ENSDART00000104188 | igsf21b | 0.000907455 | -0.476667576 | -0.274511883 | -0.073843861 | 0.013583331 | brain innervation |
| ENSDART00000104289 | rab3ab | 0.00027551 | -0.744433679 | -0.710299004 | -0.400971594 | -0.021261155 | brain innervation |
| ENSDART00000104327 | vsx1 | 0.00488592 | -0.248525466 | -0.619641572 | -0.034766195 | -0.028538746 | brain innervation |
| ENSDART00000104751 | mxi1 | 0.002350825 | -0.384587031 | -0.210909301 | 0.22179917 | 0.104111801 | brain innervation |
| ENSDART00000104895 | rgs7a | 1.96612E-05 | -0.76773786 | -0.54211529 | -0.053731925 | 0.298916153 | brain innervation |
| ENSDART00000104933 | eepd1 | 0.033087133 | -0.181346781 | -0.362247495 | -0.01782264 | 0.011720383 | brain innervation |
| ENSDART00000104950 | atp1a3a | 5.18727E-05 | -0.469887861 | -0.514207298 | -0.064097938 | 0.181102676 | brain innervation |
| ENSDART00000105484 | si:ch211-216b21.2 | 0.000149497 | -0.960990997 | -0.619204359 | -0.135410073 | 0.236742995 | brain innervation |
| ENSDART00000105588 | pcdh1b | 0.006658285 | -0.246217756 | -0.446101289 | -0.039720059 | 0.09005245 | brain innervation |
| ENSDART00000105743 | ntng1a | 0.000844413 | -0.262343031 | -0.343435025 | -0.06227485 | 0.248931143 | brain innervation |
| ENSDART00000105753 | olfm3a | 0.008447718 | -0.365975807 | -0.465135675 | -0.139223772 | 0.043379292 | brain innervation |
| ENSDART00000105774 | ek1 | 0.000784035 | -0.422572147 | -0.621215995 | -0.219935537 | -0.023014864 | brain innervation |
| ENSDART00000105932 | si:dkeyp-110e4.11 | 6.00847E-05 | -0.775118699 | -0.662012678 | -0.111191342 | 0.348802036 | brain innervation |
| ENSDART00000106166 | rx1 | 0.002317622 | -1.001905061 | -0.358282402 | 0.191592029 | -0.031411117 | brain innervation |
| ENSDART00000106619 | nrn1a | 0.000319291 | -1.225032386 | -1.272422646 | -0.361032349 | 0.032061464 | brain innervation |
| ENSDART00000108535 | gabrb1 | 0.000145808 | -1.10767094 | -1.249599593 | -0.238929019 | 0.417524871 | brain innervation |
| ENSDART00000108729 | adamtsl2 | 0.000110744 | -0.842317462 | -0.70962753 | 0.040474933 | 0.841307736 | brain innervation |
| ENSDART00000108736 | adam12 | 0.000825994 | -0.319460786 | -0.316431452 | 0.445307645 | 0.87790832 | brain innervation |
| ENSDART00000108808 | frem2a | 6.41137E-06 | -0.381822233 | -0.539670066 | 0.662155423 | 0.974116851 | brain innervation |
| ENSDART00000108818 | arhgap32a | 0.00053265 | -0.576365369 | -0.434660146 | -0.258610983 | 0.095307278 | brain innervation |
| ENSDART00000109029 | map6a | 0.002736161 | -0.330392405 | -0.354970167 | -0.034172445 | 0.05340091 | brain innervation |
| ENSDART00000109040 | gpr158a | 0.001331938 | -0.331785981 | -0.632662732 | -0.236983197 | 0.105308258 | brain innervation |
| ENSDART00000109243 | sema4bb | 0.000494919 | -0.591195487 | -0.421350422 | -0.25044186 | -0.118505649 | brain innervation |
| ENSDART00000109356 | klf7a | 0.000177237 | -0.7063233 | -0.659659661 | -0.044315321 | 0.157900055 | brain innervation |
| ENSDART00000109507 | RAP1GDS1 | 0.00047864 | -0.399835735 | -0.351308804 | 0.076766568 | 0.119897865 | brain innervation |
| ENSDART00000109511 | si:ch211-186j3.6 | 0.020586406 | -0.164289524 | -0.357536961 | -0.081167433 | 0.02827477 | brain innervation |
| ENSDART00000109568 | pip4k2ab | 0.008492808 | -0.31785183 | -0.37156543 | -0.209120418 | 0.064881056 | brain innervation |
| ENSDART00000109833 | zmat3 | 0.001153184 | -0.462380241 | -0.737159777 | -0.151865097 | 0.090633241 | brain innervation |
| ENSDART00000110004 | frmpd3 | 0.000503033 | -0.530320903 | -0.511652794 | -0.387982186 | 0.071931622 | brain innervation |
| ENSDART00000110061 | spock2 | 0.000177237 | -0.746309396 | -0.624301954 | -0.158480844 | 0.116789254 | brain innervation |
| ENSDART00000110126 | cacng3b | 0.000230274 | -0.394325343 | -0.354117689 | -0.012321564 | 0.185829622 | brain innervation |
| ENSDART00000110136 | cntnap5b | 0.000354446 | -0.584561492 | -0.553235513 | -0.330365654 | 0.070638148 | brain innervation |
| ENSDART00000110279 | si:dkey-183c6.8 | 0.000114847 | -0.455316373 | -0.174866477 | 0.555526708 | 0.790514442 | brain innervation |
| ENSDART00000110409 | frmd5 | 0.001654051 | -0.554751162 | -0.59840736 | -0.384150671 | 0.071106799 | brain innervation |
| ENSDART00000110503 | adam11 | 1.36144E-05 | -0.835498408 | -0.874358127 | 0.055154913 | 0.615136144 | brain innervation |
| ENSDART00000110590 | VSTM2B | 0.004543629 | -0.57346103 | -0.627600936 | -0.34222959 | -0.040178972 | brain innervation |
| ENSDART00000110679 | lrfn5b | 2.98725E-05 | -0.958299909 | -0.447121499 | 0.131848444 | 0.317540452 | brain innervation |
| ENSDART00000110734 | FAM163A | 0.001644836 | -0.371556039 | -0.423320211 | -0.272260914 | 0.021575603 | brain innervation |
| ENSDART00000110751 | tiam1a | 3.21969E-05 | -0.473694473 | -0.373415926 | 0.157725394 | 0.265421677 | brain innervation |
| ENSDART00000111131 | elfn1b | 0.001021091 | -1.012125284 | -0.755320746 | -0.301892543 | 0.27076153 | brain innervation |
| ENSDART00000111146 | si:ch211-26b3.4 | 0.000598789 | -0.414256231 | -0.420417531 | -0.178854692 | 0.040407002 | brain innervation |
| ENSDART00000111156 | pdp1 | 0.000503033 | -0.641430349 | -0.338633085 | -0.014499951 | 0.034332362 | brain innervation |
| ENSDART00000111203 | tagln3a | 0.000536544 | -0.530904757 | -0.489741378 | -0.221881281 | -0.037658487 | brain innervation |
| ENSDART00000111301 | luzp2 | 0.000592988 | -0.629816567 | -0.548346899 | 0.040106065 | 0.088332731 | brain innervation |
| ENSDART00000111438 | mgea5 | 0.002109252 | -0.307275123 | -0.024811963 | 0.160874404 | 0.159343333 | brain innervation |

| | | | | | | | |
|---|---|---|---|---|---|---|---|
| ENSDART00000111506 | lrrc75ba | 4.11188E-05 | -0.649545281 | -0.408918781 | -0.064365089 | 0.060080923 | brain innervation |
| ENSDART00000111535 | elfn1a | 0.001460323 | -0.273088167 | -0.336173663 | -0.097567542 | 0.139029668 | brain innervation |
| ENSDART00000111639 | rereb | 0.003012303 | -0.168137061 | -0.390988508 | -0.080335114 | 0.086629083 | brain innervation |
| ENSDART00000111642 | brinp3a.1 | 7.55931E-05 | -0.710663483 | -0.69660402 | -0.30207639 | -0.147819903 | brain innervation |
| ENSDART00000111948 | sez6l2 | 0.000199502 | -0.5142749 | -0.140866125 | 0.323444655 | 0.419947542 | brain innervation |
| ENSDART00000112003 | adgrb1a | 0.00037001 | -0.32135813 | -0.631154823 | -0.317944877 | 0.097147786 | brain innervation |
| ENSDART00000112170 | unm_hu7912 | 0.000340312 | -0.404814593 | -0.573062573 | -0.192411138 | 0.071188401 | brain innervation |
| ENSDART00000112370 | ano11 | 1.45694E-05 | -1.080551268 | -1.39069517 | -0.608655336 | 0.046438031 | brain innervation |
| ENSDART00000112414 | rapgef5a | 0.002098034 | -0.533059964 | -0.605979704 | -0.027243653 | 0.007460247 | brain innervation |
| ENSDART00000112484 | nlgn2b | 2.18017E-05 | -0.707313853 | -0.645741453 | -0.193542068 | 0.120518294 | brain innervation |
| ENSDART00000112546 | pkn3 | 0.017988416 | -0.415654116 | -0.225897422 | -0.124759806 | -0.079978695 | brain innervation |
| ENSDART00000112671 | bicd1a | 0.000246292 | -0.391567366 | -0.28876668 | -0.040397452 | 0.195532163 | brain innervation |
| ENSDART00000112895 | FNDC10 | 3.90156E-05 | -0.46103084 | -0.039905964 | 0.379016706 | 0.57822448 | brain innervation |
| ENSDART00000113081 | gpr158b | 4.77268E-06 | -0.777838913 | -0.859733549 | -0.109236002 | 0.425659764 | brain innervation |
| ENSDART00000113171 | nlgn1 | 0.000370918 | -0.287475058 | -0.532070389 | -0.095767589 | 0.228264127 | brain innervation |
| ENSDART00000113502 | si:dkey-84j12.1 | 4.60467E-05 | -0.363232597 | -0.10430598 | 0.492332906 | 0.665079551 | brain innervation |
| ENSDART00000113511 | lzts1 | 0.000109579 | -1.206756297 | -0.713910989 | -0.061680884 | 0.474255612 | brain innervation |
| ENSDART00000113551 | trim2b | 0.0245375 | -0.50417548 | -0.207383112 | -0.112448816 | 0.043074002 | brain innervation |
| ENSDART00000113773 | cdc42bpab | 0.002362892 | -0.428966416 | -0.296406465 | 0.003528598 | 0.157857062 | brain innervation |
| ENSDART00000113796 | cacnb3b | 1.63409E-06 | -1.378695476 | -1.717513295 | -0.09050578 | 0.468842834 | brain innervation |
| ENSDART00000113864 | faxca | 0.000400485 | -0.531664025 | -0.191322961 | -0.005943785 | 0.107092327 | brain innervation |
| ENSDART00000113924 | lrrc7 | 0.000228698 | -0.537461037 | -0.585108204 | -0.303180727 | -0.024642369 | brain innervation |
| ENSDART00000113985 | mctp1a | 0.000375747 | -0.392181593 | -0.510456941 | -0.185251742 | 0.106694132 | brain innervation |
| ENSDART00000114010 | slitrk3a | 0.006522424 | -0.361859007 | -0.532282315 | -0.225491041 | 0.106517685 | brain innervation |
| ENSDART00000114267 | map6b | 2.98725E-05 | -0.567896811 | -0.403955591 | 0.492985119 | 0.85461741 | brain innervation |
| ENSDART00000114432 | ntng2a | 0.000596344 | -0.38146173 | -0.278136767 | -0.282818481 | 0.289761799 | brain innervation |
| ENSDART00000114448 | CHST8 | 0.000248091 | -0.44164787 | -0.464005415 | 0.047368929 | 0.426791372 | brain innervation |
| ENSDART00000114677 | si:ch73-62l21.1 | 0.007349533 | -0.323827666 | -0.531978643 | -0.239685061 | 0.043755156 | brain innervation |
| ENSDART00000114723 | rapgefl1 | 0.020836881 | -0.431713581 | -0.430345279 | -0.085299154 | -0.113952199 | brain innervation |
| ENSDART00000114954 | rapgef5b | 0.002739168 | -0.637101425 | -0.566396383 | -0.284127257 | 0.00714868 | brain innervation |
| ENSDART00000114959 | cdh24a | 0.004579864 | -0.51153845 | -0.93145045 | -0.229616461 | 0.228723625 | brain innervation |
| ENSDART00000115027 | tmem151bb | 0.000172675 | -0.856524713 | -0.708298022 | 0.291447963 | 0.495918048 | brain innervation |
| ENSDART00000115089 | zgc:171482 | 0.003451532 | -0.437080228 | -0.507022457 | -0.285662093 | 0.021798354 | brain innervation |
| ENSDART00000115141 | cacnb3b | 0.000257119 | -1.10918155 | -1.536403603 | -0.203181521 | 0.418821499 | brain innervation |
| ENSDART00000115260 | si:ch211-113g11.6 | 0.005572285 | -0.558167113 | -1.193520332 | -0.037233796 | 0.533309069 | brain innervation |
| ENSDART00000121460 | prdm8b | 0.00076978 | -0.383342673 | -0.361617984 | -0.117709411 | 0.067402978 | brain innervation |
| ENSDART00000121708 | pcsk1nl | 0.000169466 | -0.536353064 | -0.421583117 | -0.086670577 | -0.06684806 | brain innervation |
| ENSDART00000121874 | nfasca | 0.002178187 | -0.536181619 | -0.366142839 | -0.19557692 | -0.058427118 | brain innervation |
| ENSDART00000121913 | kctd12b | 0.015495815 | -0.424186634 | -0.205906213 | -0.149727383 | -0.132895958 | brain innervation |
| ENSDART00000122037 | rbfox1 | 0.000487998 | -0.419101241 | -0.392215043 | 0.102043249 | 0.360818016 | brain innervation |
| ENSDART00000122041 | nrcama | 0.012857389 | -0.49531831 | -0.707479994 | -0.094699246 | 0.009754423 | brain innervation |
| ENSDART00000122099 | dynll2a | 0.004627449 | -0.463212519 | -0.169502826 | -0.013591394 | 0.001709702 | brain innervation |
| ENSDART00000122101 | tbx2b | 0.001190205 | -0.056468629 | 0.017248499 | 0.19422425 | 0.382938949 | brain innervation |
| ENSDART00000122389 | elmod1 | 3.77986E-05 | -0.62893484 | -0.673357812 | -0.159560027 | 0.32261476 | brain innervation |
| ENSDART00000122574 | ppp1r3aa | 0.000793313 | -0.153287429 | -0.684598529 | 0.022675976 | 0.569399664 | brain innervation |
| ENSDART00000122905 | rac3b | 0.000254175 | -0.501948401 | -0.363291367 | -0.120049924 | 0.166932504 | brain innervation |
| ENSDART00000123839 | arrb1 | 0.000512152 | -0.44263739 | -0.489445552 | -0.281663489 | 0.083496974 | brain innervation |
| ENSDART00000123851 | fut8b | 0.013533882 | -0.379186004 | -0.673089848 | -0.195485297 | 0.141395943 | brain innervation |
| ENSDART00000123868 | dthd1 | 0.020324061 | -0.700427011 | -0.013611963 | -0.073941373 | -0.012612052 | brain innervation |
| ENSDART00000124112 | pou4f2 | 2.95145E-05 | -1.368026557 | -1.308261738 | 0.344262724 | 0.908586045 | brain innervation |
| ENSDART00000124331 | si:ch1073-174d20.1 | 0.000107168 | -0.948877018 | -0.570444361 | 0.133772468 | 0.53293925 | brain innervation |
| ENSDART00000124534 | mbnl2 | 0.001000183 | -0.407254222 | 0.078580914 | -0.044925118 | 0.120266311 | brain innervation |
| ENSDART00000124710 | dlg5a | 0.003770761 | -0.356924394 | -0.265850728 | -0.130233283 | 0.096340819 | brain innervation |
| ENSDART00000124751 | kcnip3b | 2.63402E-05 | -1.29606503 | -1.383048631 | -0.490749215 | -0.039127508 | brain innervation |
| ENSDART00000124827 | lgi2a | 0.00013528 | -0.389885576 | -0.462014195 | -0.106597692 | 0.145509175 | brain innervation |
| ENSDART00000125045 | dscama | 0.001138119 | -0.36310557 | -0.486996676 | -0.080151729 | 0.074065642 | brain innervation |
| ENSDART00000125074 | kcnab2b | 0.001012163 | -0.745923366 | -1.091983107 | -0.420367552 | 0.000775062 | brain innervation |
| ENSDART00000125284 | nlgn2a | 0.000673063 | -0.210046545 | -0.366872132 | 0.090424981 | 0.163948503 | brain innervation |
| ENSDART00000125432 | esrrd | 0.007748762 | -0.825323711 | -0.251243419 | -0.11920169 | -0.154136212 | brain innervation |
| ENSDART00000125450 | gpc1a | 1.08487E-05 | -0.40101998 | -0.317779496 | 0.568583174 | 0.472068591 | brain innervation |
| ENSDART00000125536 | appb | 0.000152384 | -0.473539223 | -0.363304525 | 0.075744005 | 0.226987466 | brain innervation |
| ENSDART00000125563 | homer3b | 0.030075636 | -0.366344126 | -0.72350131 | -0.2232749 | 0.112901448 | brain innervation |
| ENSDART00000125638 | tbx5b | 0.032861605 | -0.961612456 | -0.494056529 | -0.317150736 | -0.11065371 | brain innervation |
| ENSDART00000125733 | chst1 | 0.032601457 | -1.779723555 | -0.640531075 | -0.420086911 | -0.327548455 | brain innervation |
| ENSDART00000125759 | si:dkey-22o12.2 | 0.004657196 | -0.433356276 | -0.386370833 | -0.195481295 | -0.118191078 | brain innervation |
| ENSDART00000126640 | RHOBTB3 | 0.006456063 | -0.934122492 | -0.733675156 | -0.213973967 | -0.180952113 | brain innervation |
| ENSDART00000126744 | nrg3b | 0.000370918 | -0.484773132 | -0.477903604 | -0.212248622 | 0.011007147 | brain innervation |
| ENSDART00000126766 | erbb4b | 0.001365263 | -0.378985194 | -0.566156911 | -0.016423806 | 0.214915468 | brain innervation |
| ENSDART00000126945 | adgrb3 | 0.006975472 | -0.310129925 | -0.752937386 | -0.198981749 | -0.020103794 | brain innervation |
| ENSDART00000127047 | tusc5a | 2.01848E-05 | -1.04192916 | -0.556828029 | 0.377559818 | 0.604205052 | brain innervation |
| ENSDART00000127136 | slco5a1b | 0.039811839 | -0.145489914 | -0.411558687 | 0.007460079 | -0.021875568 | brain innervation |
| ENSDART00000127286 | si:ch73-140j24.4 | 6.73259E-05 | -0.467040467 | -0.569237917 | -0.172853738 | 0.202142114 | brain innervation |
| ENSDART00000128226 | slc6a11b | 6.9873E-05 | -0.388946235 | -0.51506191 | -0.246391039 | 0.364300065 | brain innervation |
| ENSDART00000128271 | si:dkey-114c15.7 | 0.003693464 | -0.608254547 | -0.511435956 | -0.075108717 | 0.021034155 | brain innervation |
| ENSDART00000128302 | stk35 | 0.014339104 | -0.274652044 | -0.340334252 | -0.095489621 | 0.00314331 | brain innervation |

| | | | | | | | |
|---|---|---|---|---|---|---|---|
| ENSDART00000128690 | agap3 | 0.009899954 | -0.218144691 | -0.339443752 | 0.012538674 | 0.008782978 | brain innervation |
| ENSDART00000128705 | ndrg3a | 0.000135332 | -0.684036937 | -0.148509308 | 0.416765356 | 0.500717941 | brain innervation |
| ENSDART00000128794 | gpm6ab | 0.000948605 | -0.753205656 | -0.206704843 | 0.388986514 | 0.547688547 | brain innervation |
| ENSDART00000128839 | fez1 | 0.000920284 | -0.463224643 | -0.323499809 | -0.160691051 | 0.039189596 | brain innervation |
| ENSDART00000129055 | nrg2b | 0.00309942 | -0.525587506 | -0.415280837 | -0.153878493 | 0.206094594 | brain innervation |
| ENSDART00000129248 | clstn1 | 0.000984416 | -0.435055335 | -0.404808779 | -0.072836605 | 0.145009995 | brain innervation |
| ENSDART00000129511 | robo2 | 6.11958E-05 | -0.431501059 | -0.512946091 | -0.075617078 | 0.250070953 | brain innervation |
| ENSDART00000129559 | btbd3b | 0.000261206 | -0.741839183 | -0.720572279 | -0.364214548 | 0.15940566 | brain innervation |
| ENSDART00000129819 | si:ch211-151p13.8 | 0.000362487 | -0.747854488 | -0.665734365 | -0.328239319 | -0.007548317 | brain innervation |
| ENSDART00000129834 | nat16 | 0.000882713 | -0.654571013 | -0.560511117 | -0.238004357 | -0.212203603 | brain innervation |
| ENSDART00000129866 | pcsk2 | 0.000314849 | -0.608807617 | -0.601188495 | -0.114458443 | 0.092104737 | brain innervation |
| ENSDART00000129910 | tmem240a | 0.000995127 | -0.628384792 | -0.334786051 | 0.214415995 | 0.163191044 | brain innervation |
| ENSDART00000129924 | hmgb3a | 0.001276923 | -0.441161001 | -0.289637174 | 0.049371785 | 0.069315625 | brain innervation |
| ENSDART00000130007 | GABRA2 (1 of many) | 7.10385E-05 | -0.583670012 | -0.533882123 | -0.170964807 | 0.218833255 | brain innervation |
| ENSDART00000130103 | kctd16a | 1.73574E-05 | -0.864638484 | -0.862435538 | 0.056246994 | 0.316558654 | brain innervation |
| ENSDART00000130125 | gnao1a | 0.000604807 | -0.213616659 | -0.159180808 | 0.167107834 | 0.384555308 | brain innervation |
| ENSDART00000130310 | epb41l3a | 0.00059038 | -0.373985055 | -0.254132145 | 0.194766582 | 0.26124696 | brain innervation |
| ENSDART00000130573 | atp6ap1la | 0.001946729 | -0.419066331 | -0.227150493 | -0.139216735 | 0.064701891 | brain innervation |
| ENSDART00000131075 | dusp23b | 0.033109676 | -0.349967426 | -0.590231643 | 0.026826234 | -0.132825162 | brain innervation |
| ENSDART00000131126 | bcl6a | 0.005688069 | -0.096649799 | -0.044696983 | 0.295011601 | 0.436349263 | brain innervation |
| ENSDART00000131143 | sept5b | 0.000106562 | -0.480326507 | -0.332562488 | 0.283118689 | 0.526750102 | brain innervation |
| ENSDART00000131579 | ptprdb | 7.68811E-06 | -0.516674931 | -0.642051878 | 0.273217974 | 0.528486925 | brain innervation |
| ENSDART00000131714 | mbnl2 | 0.008848885 | -0.406601788 | -0.032543807 | -0.091713267 | 0.092369986 | brain innervation |
| ENSDART00000131736 | kcnq2a | 2.08005E-05 | -0.749729315 | -0.34933354 | 0.575885203 | 0.723193716 | brain innervation |
| ENSDART00000132027 | spock2 | 0.000219336 | -0.61060119 | -0.435727551 | -0.163124474 | 0.204372495 | brain innervation |
| ENSDART00000132123 | thsd7ba | 0.001248576 | -0.490674171 | -0.551076996 | -0.028899921 | -0.007032812 | brain innervation |
| ENSDART00000132278 | rnf144ab | 0.012297973 | -0.480742338 | -0.392838261 | -0.157627213 | -0.112649454 | brain innervation |
| ENSDART00000132294 | fut8a | 0.003138642 | -0.131220853 | -0.116575506 | 0.125377151 | 0.342318079 | brain innervation |
| ENSDART00000132342 | kiaa1549lb | 0.011357239 | -0.130776123 | -0.485143208 | -0.118054479 | 0.086040071 | brain innervation |
| ENSDART00000132471 | csmd2 | 6.2134E-05 | -0.459001483 | -0.911607393 | -0.371722747 | 0.062349627 | brain innervation |
| ENSDART00000132543 | klhdc8b | 0.004230106 | -0.423366405 | -0.574887213 | -0.139148992 | -0.034877401 | brain innervation |
| ENSDART00000132583 | nrxn2a | 0.001331938 | -0.296541609 | -0.578930697 | -0.085512446 | 0.138469202 | brain innervation |
| ENSDART00000132685 | kcnn1b | 0.00155515 | -1.625205585 | -1.495755896 | -0.13719571 | 0.28532188 | brain innervation |
| ENSDART00000132691 | adgrb3 | 0.007781888 | -0.413275118 | -0.450913383 | -0.19602447 | 0.067293319 | brain innervation |
| ENSDART00000132930 | pcbp3 | 0.000849348 | -0.89548835 | -0.955377646 | -0.326630607 | 0.075971616 | brain innervation |
| ENSDART00000133092 | st6galnac5a | 0.000503033 | -0.798487612 | -0.408716424 | -0.156113795 | 0.220161008 | brain innervation |
| ENSDART00000133223 | COQ10A | 0.001579476 | -0.381890626 | -0.12435669 | 0.152677483 | 0.108453147 | brain innervation |
| ENSDART00000133330 | fam214b | 0.006799542 | -0.307417712 | -0.32820559 | -0.047756819 | 0.003374146 | brain innervation |
| ENSDART00000133385 | lrp1ba | 0.000155554 | -0.614935587 | -0.929322267 | -0.360685611 | 0.193952764 | brain innervation |
| ENSDART00000133504 | dab2ipb | 0.019374625 | -0.309610082 | -0.535236809 | -0.062134515 | 0.075905091 | brain innervation |
| ENSDART00000133956 | phactr3a | 0.000149865 | -0.466148377 | -0.040906797 | 0.400008422 | 0.572153181 | brain innervation |
| ENSDART00000133985 | kcnd1 | 0.000248125 | -1.187342293 | -1.392518344 | -0.747443213 | -0.002522665 | brain innervation |
| ENSDART00000134035 | cntnap5b | 0.005466942 | -1.01200416 | -1.154440105 | -0.403471674 | 0.139198923 | brain innervation |
| ENSDART00000134261 | dnm1b | 1.07133E-05 | -0.433650537 | -0.451510325 | 0.138870574 | 0.465815695 | brain innervation |
| ENSDART00000134307 | dclk1a | 0.001654051 | -1.448196855 | -0.41548572 | -0.010990365 | -0.586401585 | brain innervation |
| ENSDART00000134475 | gdap1 | 0.004885039 | -0.459369959 | -0.408869462 | -0.283212336 | -0.008346648 | brain innervation |
| ENSDART00000134564 | srsf5b | 0.01711161 | -0.27445076 | -0.31745813 | -0.113899297 | -0.080138136 | brain innervation |
| ENSDART00000134715 | elmod1 | 5.02856E-05 | -1.115600356 | -1.115324879 | -0.105456948 | 0.485174818 | brain innervation |
| ENSDART00000134722 | bcl2l13 | 0.00372102 | -0.594192147 | -0.544774096 | -0.184876076 | -0.124896832 | brain innervation |
| ENSDART00000134816 | slc2a3a | 0.00067643 | -0.406786727 | -0.444029943 | -0.052958142 | 0.129218418 | brain innervation |
| ENSDART00000134832 | rbpms2b | 0.003328957 | -1.576691045 | -2.099790887 | -0.701820765 | 0.187979586 | brain innervation |
| ENSDART00000134855 | efr3bb | 0.003310987 | -0.263384976 | -0.381748192 | -0.1453091 | 0.002074194 | brain innervation |
| ENSDART00000134973 | pcnxl2 | 7.10385E-05 | -0.772294262 | -0.54715302 | -0.00593982 | 0.313205165 | brain innervation |
| ENSDART00000135052 | slc8a4a | 0.003870205 | -0.449869312 | -0.676457859 | -0.284767037 | 0.064060964 | brain innervation |
| ENSDART00000135284 | vsnl1a | 3.95966E-05 | -0.802778535 | -0.782046001 | -0.520311896 | -0.04073792 | brain innervation |
| ENSDART00000135475 | cadm2a | 0.000283607 | -0.589549243 | -0.825400299 | -0.33568035 | 0.065563209 | brain innervation |
| ENSDART00000135513 | rgs6 | 0.001782169 | -0.368608418 | -0.140264684 | -0.010741291 | 0.084818744 | brain innervation |
| ENSDART00000135602 | si:ch73-290k24.6 | 0.008047992 | -0.456455854 | -0.524968049 | -0.128888416 | -0.002715321 | brain innervation |
| ENSDART00000135624 | FAM163A | 0.011945865 | -0.714036121 | -0.427639458 | -0.142310427 | 0.163241634 | brain innervation |
| ENSDART00000135692 | UNC13A | 0.006141198 | -0.209420413 | -0.513653337 | -0.217224189 | 0.093101705 | brain innervation |
| ENSDART00000135702 | rgs11 | 0.000709831 | -0.576481143 | -0.793334647 | -0.383725367 | 0.048951277 | brain innervation |
| ENSDART00000135715 | fam184a | 0.000322759 | -0.344643213 | -0.481218539 | -0.282456424 | 0.057932244 | brain innervation |
| ENSDART00000135764 | si:dkey-70p6.1 | 0.000167073 | -0.54777588 | -0.724047584 | 0.062614493 | 0.292769909 | brain innervation |
| ENSDART00000135768 | lrp1ba | 4.91848E-05 | -0.685782746 | -0.945707299 | -0.285013804 | 0.195853428 | brain innervation |
| ENSDART00000135811 | atp2b3a | 0.000535509 | -0.361290503 | -0.436904447 | -0.166639435 | 0.060424957 | brain innervation |
| ENSDART00000136123 | kdm6bb | 0.024284969 | -0.32427763 | -0.244056677 | -0.170049978 | -0.028150527 | brain innervation |
| ENSDART00000136154 | gpr158a | 0.014511767 | -0.522434846 | -0.680761714 | -0.064607745 | -0.081447185 | brain innervation |
| ENSDART00000136233 | TENM3 | 2.33589E-05 | -0.571502608 | -0.593065001 | 0.242250375 | 0.436842415 | brain innervation |
| ENSDART00000136269 | rimbp2 | 0.016247803 | -0.196271668 | -0.410552217 | -0.160743106 | 0.064316711 | brain innervation |
| ENSDART00000136488 | zmat4a | 0.008573382 | -0.690962776 | -0.637104526 | -0.28999313 | -0.088561559 | brain innervation |
| ENSDART00000136578 | syn1 | 4.91848E-05 | -0.141221338 | -0.191909782 | 0.363564902 | 0.481540125 | brain innervation |
| ENSDART00000136708 | rbpms2b | 1.12792E-05 | -1.024689498 | -0.80090545 | -0.103078191 | 0.159293341 | brain innervation |
| ENSDART00000136754 | cdh13 | 0.000634488 | -0.587587145 | -0.627840519 | -0.172128457 | 0.142910694 | brain innervation |
| ENSDART00000136759 | prdm13 | 0.00326136 | -0.225066145 | -0.413438688 | -0.129520305 | 0.036146962 | brain innervation |
| ENSDART00000137037 | iqsec3b | 0.006778044 | -0.405090433 | -0.576572578 | -0.30180595 | 0.030413937 | brain innervation |

| | | | | | | | |
|---|---|---|---|---|---|---|---|
| ENSDART00000137353 | klhl17 | 0.00053559 | -0.368352542 | -0.314849207 | -0.039056928 | 0.155772163 | brain innervation |
| ENSDART00000137355 | tet3 | 0.033062691 | -0.245273707 | -0.407219331 | -0.050664281 | -0.06569376 | brain innervation |
| ENSDART00000137756 | hecw2a | 3.29362E-06 | -0.728856656 | -1.003560302 | -0.22088311 | 0.383324077 | brain innervation |
| ENSDART00000137889 | ank1b | 0.002868986 | -0.447220804 | -0.737203628 | 0.205089912 | 0.418795119 | brain innervation |
| ENSDART00000137900 | grin2ab | 2.59942E-05 | -0.452993272 | -0.616161489 | 0.576592075 | 1.050321835 | brain innervation |
| ENSDART00000138048 | pbx3b | 0.007841242 | -0.599655595 | -0.602159282 | -0.228775798 | -0.019503135 | brain innervation |
| ENSDART00000138294 | rab41 | 0.000446102 | -0.246606124 | -0.179576466 | 0.274617961 | 0.438688085 | brain innervation |
| ENSDART00000138540 | ttyh1 | 5.26677E-05 | -0.563796304 | -0.437755604 | -0.092270149 | 0.165178407 | brain innervation |
| ENSDART00000138695 | ank1b | 0.000591348 | -0.498943842 | -0.553539638 | -0.03799849 | 0.33423854 | brain innervation |
| ENSDART00000138834 | mab21l3 | 0.00420635 | -0.591171056 | -0.506133859 | -0.165242719 | -0.104286764 | brain innervation |
| ENSDART00000138850 | cacna2d2b | 0.008428026 | -0.226630579 | -0.433395437 | -0.06209639 | 0.11961013 | brain innervation |
| ENSDART00000139035 | si:ch211-215c18.3 | 1.2512E-05 | -1.205532305 | -0.512285433 | 0.437440349 | 0.984798646 | brain innervation |
| ENSDART00000139178 | slc25a14 | 0.001193104 | -0.405048408 | -0.151867564 | -0.020130795 | 0.082212943 | brain innervation |
| ENSDART00000139265 | si:ch211-12e13.12 | 0.00026634 | -0.4797238 | -0.27645528 | 0.604278273 | 0.83384752 | brain innervation |
| ENSDART00000139279 | rbpms2b | 0.001175355 | -0.774960547 | -0.830410704 | 0.12323811 | 0.508122144 | brain innervation |
| ENSDART00000139448 | GALNTL6 | 0.000882263 | -0.494238764 | -0.383984121 | -0.129586789 | 0.079582323 | brain innervation |
| ENSDART00000139475 | samd10b | 0.001021091 | -0.569533963 | -0.524261904 | -0.125238267 | -0.046566236 | brain innervation |
| ENSDART00000139493 | olfm1b | 0.002701643 | -0.400757951 | -0.50188464 | -0.148663949 | 0.07973127 | brain innervation |
| ENSDART00000139514 | ano11 | 0.001023352 | -1.063704055 | -1.17363318 | -0.628817004 | -0.055546726 | brain innervation |
| ENSDART00000139795 | syt2a | 7.6604E-05 | -0.704240682 | -0.905957314 | -0.438675098 | 0.054000022 | brain innervation |
| ENSDART00000139841 | gabrb1 | 0.000278329 | -0.439523493 | -0.431067027 | -0.18481289 | 0.263500104 | brain innervation |
| ENSDART00000139954 | BX005364.1 | 0.014034602 | -0.374788931 | -0.616534875 | -0.121001872 | 0.068005074 | brain innervation |
| ENSDART00000140161 | necab2 | 2.33589E-05 | -0.820451686 | -0.597146212 | 0.232199679 | 0.59216621 | brain innervation |
| ENSDART00000140365 | slc12a5b | 0.002585408 | -0.506920714 | -0.535232712 | -0.246533716 | 0.015331251 | brain innervation |
| ENSDART00000140436 | map2 | 0.004423084 | -0.405015536 | -0.195771527 | -0.230781013 | 0.041022881 | brain innervation |
| ENSDART00000140476 | atp6v0a2a | 0.000363306 | -0.389741082 | 0.011917076 | 0.174652763 | 0.265809184 | brain innervation |
| ENSDART00000140575 | map4l | 0.032473859 | -0.647123698 | -0.438384786 | -0.184748643 | -0.024632978 | brain innervation |
| ENSDART00000140739 | lrrc4bb | 0.029601456 | -0.565179268 | -0.464872302 | -0.152447222 | -0.079782637 | brain innervation |
| ENSDART00000140827 | bcl11ab | 0.000602987 | -0.293093875 | -0.330078462 | 0.796150714 | 1.216507448 | brain innervation |
| ENSDART00000141086 | robo2 | 0.010303026 | -0.237929535 | -0.432116468 | -0.02855362 | 0.131332743 | brain innervation |
| ENSDART00000141103 | gfra4a | 0.0021138 | -0.446789167 | -0.472299982 | -0.170582645 | 0.04427624 | brain innervation |
| ENSDART00000141752 | asphd1 | 0.007785924 | -0.390703626 | -0.359372328 | -0.291984856 | 0.011586147 | brain innervation |
| ENSDART00000141779 | fndc5a | 0.000119544 | -1.597130885 | -1.939288998 | -0.836516829 | 0.004414498 | brain innervation |
| ENSDART00000141917 | ank1a | 3.03667E-05 | -1.167872922 | -1.059135645 | -0.600851313 | -0.041118915 | brain innervation |
| ENSDART00000141958 | cdh24a | 0.005763869 | -0.328998631 | -0.596758779 | -0.216164592 | 0.127161048 | brain innervation |
| ENSDART00000142061 | si:dkey-32e23.4 | 0.001867076 | -0.369490073 | -0.293305321 | 0.04108125 | 0.132687244 | brain innervation |
| ENSDART00000142087 | foxp2 | 0.000565675 | -0.522019738 | -0.4194261 | 0.065656049 | 0.27497631 | brain innervation |
| ENSDART00000142122 | cplx2l | 0.000340549 | -1.429903189 | -0.89072761 | 0.133226492 | 0.114783163 | brain innervation |
| ENSDART00000142140 | ppfia2 | 0.001332606 | -0.37587085 | -0.48411577 | 0.218087957 | 0.20796301 | brain innervation |
| ENSDART00000142141 | grin2aa | 0.000227894 | -0.499492643 | -0.789969038 | -0.312539264 | 0.08079802 | brain innervation |
| ENSDART00000142489 | slco5a1b | 0.001879617 | -0.38890353 | -0.329924623 | -0.120653024 | -0.00920305 | brain innervation |
| ENSDART00000142524 | chrnb2a | 9.33064E-06 | -0.869143617 | -0.99350722 | -0.040693199 | 0.426386933 | brain innervation |
| ENSDART00000142653 | grm1a | 0.003262292 | -0.469462455 | -0.467735907 | -0.088400916 | 0.178280203 | brain innervation |
| ENSDART00000143152 | cpne5a | 0.000387877 | -0.664046359 | -0.825602355 | -0.316663455 | 0.050239809 | brain innervation |
| ENSDART00000143286 | lnx1 | 0.005464372 | -0.74422542 | -0.772813048 | -0.069056217 | 0.13158415 | brain innervation |
| ENSDART00000143410 | EMB | 2.1041E-05 | -1.826210099 | -1.592080235 | -0.486327336 | 0.178400158 | brain innervation |
| ENSDART00000143938 | sema3ga | 0.000230274 | 0.034230149 | -0.448431565 | 0.102011548 | 0.447499442 | brain innervation |
| ENSDART00000143969 | BX247868.1 | 6.23756E-05 | -1.179119979 | -1.430780681 | -0.233116942 | 0.342359274 | brain innervation |
| ENSDART00000144050 | NA | 0.025646401 | -0.572877113 | -0.701246998 | -0.196460214 | -0.040487474 | brain innervation |
| ENSDART00000144186 | pou6f2 | 7.05055E-05 | -0.474463335 | -0.465571046 | 0.092941552 | 0.436081081 | brain innervation |
| ENSDART00000144215 | pcnxl2 | 1.45694E-05 | -0.697249288 | -0.448197444 | 0.040412938 | 0.382373048 | brain innervation |
| ENSDART00000144361 | ank1b | 2.46332E-05 | -0.731673078 | -0.825040806 | 0.086374477 | 0.310723345 | brain innervation |
| ENSDART00000144894 | bcl11ba | 3.52427E-05 | -0.656989822 | -0.679320488 | 0.181338486 | 0.623847694 | brain innervation |
| ENSDART00000145096 | fam219ab | 0.007936937 | -0.366547695 | -0.337752539 | -0.191344563 | -0.015845187 | brain innervation |
| ENSDART00000145108 | MFF (1 of many) | 0.004863057 | -0.387524941 | -0.341112286 | -0.251149344 | -0.013627184 | brain innervation |
| ENSDART00000145114 | ptprsa | 0.011196176 | -0.133739633 | -0.353332073 | -0.057919239 | 0.041724305 | brain innervation |
| ENSDART00000145434 | gpr52 | 0.010169943 | -0.413465096 | -0.575440094 | -0.276023467 | -0.017738845 | brain innervation |
| ENSDART00000145489 | ano11 | 0.000184045 | -1.065264341 | -1.355279668 | -0.34443399 | 0.075205433 | brain innervation |
| ENSDART00000145616 | adarb1b | 8.45506E-05 | -0.549552658 | -0.593167659 | -0.056719481 | 0.082749855 | brain innervation |
| ENSDART00000145775 | mpp4a | 0.007561534 | -0.462462268 | -0.206618846 | -0.102159961 | 0.056169031 | brain innervation |
| ENSDART00000145777 | adgrb3 | 0.018611393 | -0.131394962 | -0.455026919 | -0.13584478 | 0.067074927 | brain innervation |
| ENSDART00000145809 | prkag3a | 0.029695868 | -4.153185554 | -1.353616268 | -0.923701192 | -0.043754788 | brain innervation |
| ENSDART00000145835 | tmx3 | 0.000403881 | -0.913440216 | 0.029499449 | 0.308787653 | 0.34181057 | brain innervation |
| ENSDART00000146055 | ptchd4 | 6.99374E-05 | -0.534340079 | -0.556417786 | 0.177414073 | 0.199962898 | brain innervation |
| ENSDART00000146120 | luzp2 | 0.000370918 | -0.491780393 | -0.439244253 | 0.071649151 | 0.105029542 | brain innervation |
| ENSDART00000146132 | si:ch211-254n4.3 | 0.000799784 | -0.407187074 | -0.522964952 | -0.215771966 | 0.042955183 | brain innervation |
| ENSDART00000146530 | epha6 | 0.000442465 | -0.320688922 | -0.528894361 | -0.185025461 | 0.089827402 | brain innervation |
| ENSDART00000146785 | tmem163b | 2.46332E-05 | -0.68540579 | -0.770110571 | -0.115396875 | 0.356833449 | brain innervation |
| ENSDART00000147001 | si:dkeyp-72h1.1 | 0.00059038 | -0.667988087 | -0.713655804 | -0.172938308 | 0.232582151 | brain innervation |
| ENSDART00000147105 | tdh | 0.003138642 | -0.645059373 | -0.498416454 | -0.202501341 | -0.068176647 | brain innervation |
| ENSDART00000147188 | trpm3 | 0.001151232 | -0.625119804 | -0.737554924 | -0.108574358 | 0.285413653 | brain innervation |
| ENSDART00000147218 | efna3a | 0.00377485 | -0.679676128 | -0.850928012 | -0.177104508 | -0.025688419 | brain innervation |
| ENSDART00000147464 | ryr3 | 4.24705E-05 | -0.823352364 | -0.716491782 | 0.05941278 | 0.608108075 | brain innervation |
| ENSDART00000147483 | ephb6 | 0.000402225 | -0.346946617 | -0.195722816 | 0.204471928 | 0.396713027 | brain innervation |
| ENSDART00000147699 | mical3b | 0.021599545 | 0.112525626 | 0.108456465 | 0.226887131 | 0.765699718 | brain innervation |

| | | | | | | | |
|---|---|---|---|---|---|---|---|
| ENSDART00000147799 | ndrg3b | 0.015070633 | -0.751369578 | -0.184479497 | 0.170135881 | -0.060187322 | brain innervation |
| ENSDART00000147831 | ACAP2 (1 of many) | 0.000728778 | -0.265239588 | -0.086744823 | 0.171263201 | 0.342225898 | brain innervation |
| ENSDART00000147998 | plxna2 | 0.007335615 | -0.136413264 | -0.347150332 | -0.077394761 | 0.05783618 | brain innervation |
| ENSDART00000148055 | si:ch211-67e16.11 | 0.000248091 | -0.532227459 | -0.535880084 | -0.161015502 | 0.246252593 | brain innervation |
| ENSDART00000148107 | SRCIN1 | 5.24617E-05 | -0.358492187 | -0.594267639 | -0.130298738 | 0.102935385 | brain innervation |
| ENSDART00000148120 | ephb2a | 5.7178E-05 | -0.204590684 | -0.196760275 | 0.359222548 | 0.385663441 | brain innervation |
| ENSDART00000148199 | stxbp1a | 0.002936628 | -0.541285454 | -0.573795052 | -0.292066803 | -0.020658402 | brain innervation |
| ENSDART00000148405 | si:ch211-113g11.6 | 0.005927492 | -0.63339569 | -0.861477951 | -0.083811842 | 0.259008105 | brain innervation |
| ENSDART00000148465 | slc1a1 | 3.38347E-05 | -0.751249091 | -0.22594806 | 0.456533505 | 0.753777978 | brain innervation |
| ENSDART00000148580 | BX005108.2 | 0.002349998 | -0.474486107 | -0.575204555 | -0.246024325 | -0.012613897 | brain innervation |
| ENSDART00000148590 | coro2ba | 0.007621679 | -0.3608902 | -0.079968694 | -0.141782791 | 0.048751465 | brain innervation |
| ENSDART00000148653 | tmem145 | 0.019023689 | -0.454635973 | -0.21216688 | -0.078241248 | 0.035128268 | brain innervation |
| ENSDART00000148662 | kcnc3a | 0.003848394 | -0.800710128 | -1.106624714 | -0.368118634 | 0.013489304 | brain innervation |
| ENSDART00000148900 | kcnc2 | 0.002497064 | -0.819802341 | -0.972756522 | -0.275729768 | 0.337871594 | brain innervation |
| ENSDART00000149032 | oat | 0.006013085 | -0.368235431 | -0.40939103 | -0.17630927 | -0.009811502 | brain innervation |
| ENSDART00000149158 | dmxl2 | 0.001023352 | -0.239475556 | -0.38571718 | -0.12120231 | 0.150747303 | brain innervation |
| ENSDART00000149175 | pou4f3 | 3.33896E-05 | -1.520089487 | -1.231379123 | 0.357488642 | 0.743891028 | brain innervation |
| ENSDART00000149276 | lzts3b | 0.009414182 | -0.642406039 | -0.470964243 | -0.143114993 | -0.054979203 | brain innervation |
| ENSDART00000149283 | zgc:112001 | 0.000242559 | -0.707617666 | -0.492468931 | 0.006563399 | 0.231921747 | brain innervation |
| ENSDART00000149335 | kcna1a | 7.55931E-05 | -0.92291358 | -0.966385056 | -0.49989391 | -0.001249229 | brain innervation |
| ENSDART00000149367 | si:ch73-290k24.5 | 0.009217803 | -0.213839599 | -0.466705261 | -0.018322662 | 0.130528894 | brain innervation |
| ENSDART00000149425 | SLC25A22 (1 of many) | 0.000236845 | -0.428786809 | -0.626535356 | -0.125118952 | 0.12464047 | brain innervation |
| ENSDART00000149443 | pcdh19 | 0.000181327 | -0.547618951 | -0.594468642 | -0.345755383 | 0.11053464 | brain innervation |
| ENSDART00000149552 | FP325120.1 | 0.00320961 | -0.456523138 | -0.25356238 | -0.125718739 | 0.024031302 | brain innervation |
| ENSDART00000149618 | mid1 | 0.002670678 | -0.406831126 | -0.160364287 | 0.131354752 | 0.136351442 | brain innervation |
| ENSDART00000149666 | dock3 | 0.004782655 | -0.065088232 | -0.425598201 | -0.009057049 | 0.161364952 | brain innervation |
| ENSDART00000149685 | gnao1a | 0.000111916 | -0.290173142 | -0.321971232 | 0.118032615 | 0.301423091 | brain innervation |
| ENSDART00000149740 | si:ch211-149b19.2 | 0.000439231 | -0.851560484 | -0.688459713 | 0.123433485 | 0.442881876 | brain innervation |
| ENSDART00000149859 | gldn | 0.020986271 | -0.670802099 | -0.540937826 | -0.034323087 | -0.157128438 | brain innervation |
| ENSDART00000149947 | bcl2b | 0.000305619 | -0.663523207 | -0.706570637 | -0.112082166 | -0.007869569 | brain innervation |
| ENSDART00000149992 | si:dkey-231j24.3 | 0.002102775 | -0.792591394 | -0.543368341 | 0.065178142 | 0.204406436 | brain innervation |
| ENSDART00000150036 | adam23a | 3.77986E-05 | -0.62166293 | -0.515362417 | 0.016147076 | 0.303932596 | brain innervation |
| ENSDART00000150068 | sox6 | 0.001331938 | -0.111707858 | -0.047191704 | 0.355284385 | 0.700337568 | brain innervation |
| ENSDART00000150146 | smarcd3b | 0.000677616 | -0.318612181 | -0.278781268 | 0.723080439 | 0.640333876 | brain innervation |
| ENSDART00000150839 | fam57bb | 0.005203275 | -0.627521697 | -0.387137362 | 0.25664183 | 0.060693872 | brain innervation |
| ENSDART00000150949 | nfyc | 0.013743075 | -0.535274529 | -0.301463031 | -0.212807295 | 0.074596031 | brain innervation |
| ENSDART00000150955 | bmpr2a | 0.012462623 | -0.667543016 | -0.436786519 | -0.136898455 | 0.053141678 | brain innervation |
| ENSDART00000151252 | myo10l1 | 3.38347E-05 | -0.09485067 | -0.323915115 | 0.494165668 | 0.855040148 | brain innervation |
| ENSDART00000151299 | scn8ab | 1.02827E-05 | -0.339946061 | -0.372438378 | 0.28752079 | 0.526264643 | brain innervation |
| ENSDART00000151571 | ahdc1 | 0.000666959 | -0.753523402 | -0.774930454 | -0.265860844 | -0.025283973 | brain innervation |
| ENSDART00000151648 | ntng2b | 0.000275413 | -0.44606387 | -0.406664301 | -0.175135105 | 0.111487092 | brain innervation |
| ENSDART00000151672 | si:ch211-276a17.5 | 0.026710645 | -0.716987346 | -0.427751363 | -0.201345778 | -0.037618143 | brain innervation |
| ENSDART00000151674 | pcdh9 | 0.000894529 | -0.204552278 | -0.440082629 | -0.022873178 | 0.221939437 | brain innervation |
| ENSDART00000151685 | lhfpl4a | 0.000326965 | -0.450299252 | -0.419390296 | -0.093062661 | 0.17106537 | brain innervation |
| ENSDART00000151698 | syt3 | 0.000157185 | -0.446756628 | -0.536044497 | -0.128625894 | 0.106551311 | brain innervation |
| ENSDART00000151842 | lrrn3a | 2.01848E-05 | -0.579197629 | -0.548770022 | -0.143313453 | 0.249550285 | brain innervation |
| ENSDART00000151853 | btbd10a | 0.000362175 | -0.220795221 | -0.048228521 | 0.169228662 | 0.338815973 | brain innervation |
| ENSDART00000151904 | foxn3 | 0.000466769 | -0.402174688 | -0.533042351 | -0.141780798 | 0.022724604 | brain innervation |
| ENSDART00000151915 | ptchd1 | 6.75287E-05 | -0.713611001 | -0.66822522 | 0.042208909 | 0.328774722 | brain innervation |
| ENSDART00000151920 | si:ch211-140m22.7 | 0.011455954 | -0.371308279 | -0.473110833 | -0.103450206 | 0.052557108 | brain innervation |
| ENSDART00000151921 | dnmt3ab | 0.001890284 | -0.131014471 | -0.104325638 | 0.10563751 | 0.340310238 | brain innervation |
| ENSDART00000151970 | si:ch211-248l17.3 | 0.000139914 | -0.372087768 | -0.337523565 | 0.171433041 | 0.720267147 | brain innervation |
| ENSDART00000152011 | DOCK4 (1 of many) | 0.000227894 | -0.107148853 | -0.150156526 | 0.321338768 | 0.540658105 | brain innervation |
| ENSDART00000152603 | egln2 | 0.018920791 | -0.344914112 | -0.184325703 | -0.029073268 | 0.070142585 | brain innervation |
| ENSDART00000152924 | stxbp4 | 0.002497064 | 0.009805274 | -0.210366163 | 0.385226099 | 0.296155125 | brain innervation |
| ENSDART00000152973 | RNF157 | 0.002439392 | -0.237155894 | -0.35583425 | -0.110722293 | 0.097030733 | brain innervation |
| ENSDART00000153022 | AL935194.1 | 0.000106562 | -0.839883385 | -0.952071296 | -0.323659944 | -0.119202215 | brain innervation |
| ENSDART00000153146 | rbfox3a | 0.000155554 | -0.546449898 | -0.43963695 | 0.174623158 | 0.484406984 | brain innervation |
| ENSDART00000153284 | tbkbp1 | 0.000630366 | -0.504212532 | -0.489626186 | -0.240468452 | 0.072471561 | brain innervation |
| ENSDART00000153514 | si:ch211-121j5.4 | 0.006293989 | -0.438676742 | -0.189457468 | -0.187818697 | -0.117536785 | brain innervation |
| ENSDART00000153704 | anks1ab | 3.38532E-05 | -0.440221806 | -0.75898091 | -0.133742531 | 0.137005152 | brain innervation |
| ENSDART00000153730 | CU856222.1 | 0.000116387 | -0.553584022 | -0.481855955 | -0.007954528 | 0.343506731 | brain innervation |
| ENSDART00000153824 | napba | 0.002994106 | -0.634929885 | -0.324273151 | -0.096308603 | 0.10005339 | brain innervation |
| ENSDART00000153828 | atp2b3b | 8.36884E-05 | -0.474522045 | -0.66566516 | -0.285016921 | 0.075707306 | brain innervation |
| ENSDART00000153897 | ndnfl | 8.26035E-05 | -0.781136652 | -1.071097643 | -0.053351937 | 0.577095222 | brain innervation |
| ENSDART00000154039 | cntnap2a | 0.000510711 | -0.318660467 | -0.524836596 | -0.23230439 | 0.054653452 | brain innervation |
| ENSDART00000154217 | tapbpl | 6.43148E-05 | -0.790387635 | -1.110046917 | -0.234167162 | 0.243016032 | brain innervation |
| ENSDART00000154253 | VAMP1 (1 of many) | 4.24705E-05 | -0.590038106 | -0.991557836 | -0.153916319 | 0.292977881 | brain innervation |
| ENSDART00000154256 | gabbr1a | 0.000825994 | -0.701390177 | -1.072254559 | -0.049084095 | 0.380533583 | brain innervation |
| ENSDART00000154333 | camkva | 0.000142247 | -0.731914874 | -0.620456244 | -0.460647205 | -0.026116956 | brain innervation |
| ENSDART00000154501 | cntnap1 | 0.000112893 | -0.715173489 | -0.84270145 | -0.359147123 | 0.122603648 | brain innervation |
| ENSDART00000154667 | SORCS3 | 0.000413975 | -0.864119649 | -0.84034251 | -0.370097899 | 0.056888282 | brain innervation |
| ENSDART00000154748 | sv2bb | 0.000171543 | -0.593359268 | -0.537099772 | -0.285465428 | 0.087161334 | brain innervation |
| ENSDART00000154754 | grik1b | 0.004364661 | -0.541985174 | -0.220416154 | -0.114773141 | -0.128231092 | brain innervation |
| ENSDART00000154811 | shank1 | 0.000776925 | -0.390024258 | -0.720748027 | -0.113172669 | 0.256359072 | brain innervation |

| | | | | | | | |
|---|---|---|---|---|---|---|---|
| ENSDART00000154869 | cacna1ha | 7.43198E-05 | -0.561980799 | -0.554587419 | -0.200655166 | 0.115415453 | brain innervation |
| ENSDART00000155184 | lcorl | 0.021296835 | -0.199231763 | -0.444480785 | -0.032192793 | -0.024389475 | brain innervation |
| ENSDART00000155397 | soat2 | 0.016354858 | -0.618559497 | -0.31486995 | -0.194806138 | -0.125519338 | brain innervation |
| ENSDART00000155494 | CR384099.1 | 0.00148037 | -0.702495416 | -0.755760921 | -0.229440848 | 0.245420937 | brain innervation |
| ENSDART00000155502 | sez6b | 0.000937142 | -0.382545228 | -0.653854879 | -0.149982953 | 0.168281376 | brain innervation |
| ENSDART00000155563 | abca5 | 0.003355787 | -0.240041791 | -0.35443471 | -0.098648169 | 0.068480985 | brain innervation |
| ENSDART00000155592 | pcdh17 | 0.000461528 | -0.283490769 | -0.422406754 | -0.268768577 | 0.186259856 | brain innervation |
| ENSDART00000155658 | rapgefl1 | 0.004578126 | -0.418362292 | -0.397338899 | 0.000660896 | -0.001869225 | brain innervation |
| ENSDART00000155666 | znf576.1 | 0.015256508 | -0.372721752 | -0.23410815 | -0.134923201 | -0.013577883 | brain innervation |
| ENSDART00000155757 | si:dkey-16p21.8 | 0.008752524 | -0.383053569 | -0.315569666 | -0.222717784 | 0.028079211 | brain innervation |
| ENSDART00000155840 | nyap2a | 0.000605791 | -0.526663005 | -0.654892136 | 0.27374848 | 0.170536216 | brain innervation |
| ENSDART00000155843 | cntnap2a | 8.92064E-05 | -0.263075507 | -0.343897302 | 0.676718804 | 0.99202798 | brain innervation |
| ENSDART00000155848 | CU929046.2 | 0.004951436 | -0.462340311 | -0.419607593 | -0.082173547 | -0.018364502 | brain innervation |
| ENSDART00000155866 | zbtb16b | 8.488E-05 | -0.434457287 | -0.595844526 | 0.044569606 | 0.015262284 | brain innervation |
| ENSDART00000155935 | clstn3 | 0.000159813 | -0.324952015 | -0.256429293 | 0.203826088 | 0.321978764 | brain innervation |
| ENSDART00000155949 | grm8b | 0.001007682 | -0.426657348 | -0.505637682 | -0.163630603 | 0.069819734 | brain innervation |
| ENSDART00000156012 | si:dkey-35i13.1 | 1.41905E-06 | -1.182557557 | -0.949918392 | -0.338171836 | -0.056213778 | brain innervation |
| ENSDART00000156341 | fam117ba | 7.05055E-05 | -0.500099563 | -0.265235629 | 0.202099115 | 0.368226992 | brain innervation |
| ENSDART00000156375 | kcnq2b | 0.000178718 | -0.284091354 | -0.197722227 | 0.27148365 | 0.512045099 | brain innervation |
| ENSDART00000156380 | CR848757.1 | 2.98725E-05 | -0.431298815 | -0.377014166 | 0.404962464 | 0.865655337 | brain innervation |
| ENSDART00000156393 | AL772314.1 | 6.26753E-05 | -0.272422966 | -0.157258449 | 0.218420805 | 0.368190657 | brain innervation |
| ENSDART00000156570 | slc6a17 | 0.002913561 | -0.448593137 | -0.394227085 | -0.032159276 | 0.074756396 | brain innervation |
| ENSDART00000156705 | si:dkeyp-72g9.4 | 0.000811547 | -0.743143803 | -0.697928004 | -0.454613917 | -0.077239927 | brain innervation |
| ENSDART00000156820 | si:dkey-7j14.5 | 4.77268E-06 | -0.971869146 | -0.757334904 | 0.08249331 | 0.800607437 | brain innervation |
| ENSDART00000156965 | osbpl5 | 5.8648E-05 | -0.452639629 | -0.228065482 | 0.296781507 | 0.346469129 | brain innervation |
| ENSDART00000156982 | si:ch211-195b15.8 | 0.000596344 | -0.203622983 | -0.158924419 | -0.050797883 | 0.351311882 | brain innervation |
| ENSDART00000157018 | stmn3 | 4.91848E-05 | -1.008205118 | -1.406964646 | -0.116902659 | 0.621837811 | brain innervation |
| ENSDART00000157125 | vsnl1b | 2.15557E-05 | -0.832964446 | -0.803942924 | -0.29707501 | 0.109485311 | brain innervation |
| ENSDART00000157171 | si:ch211-168d23.3 | 0.027638894 | -0.276296251 | -0.552181073 | -0.115547916 | -0.061680907 | brain innervation |
| ENSDART00000157237 | si:dkey-56f14.7 | 0.013233841 | -0.377561364 | -0.203140811 | -0.202521092 | -0.065412084 | brain innervation |
| ENSDART00000157265 | trim13 | 0.022539179 | -0.831738752 | -0.32341984 | -0.102064486 | -0.071474505 | brain innervation |
| ENSDART00000157302 | si:dkey-7j14.5 | 1.31948E-06 | -0.795010515 | -0.583416344 | 0.327377123 | 0.951280694 | brain innervation |
| ENSDART00000157428 | pik3r3b | 0.001317626 | -0.522784766 | -0.44851499 | -0.152608244 | 0.028231585 | brain innervation |
| ENSDART00000157449 | clstn2 | 0.000466769 | -0.339354151 | -0.60478304 | 0.06475081 | 0.303731917 | brain innervation |
| ENSDART00000157454 | ncam1a | 0.005875165 | -0.363879899 | -0.114400367 | 0.050687773 | 0.064981174 | brain innervation |
| ENSDART00000157518 | hspa12a | 5.66718E-05 | -0.767789618 | -0.701014156 | 0.093464144 | 0.364648116 | brain innervation |
| ENSDART00000157769 | pcdh1gc5 | 0.001135528 | -0.403524411 | -0.226721593 | 0.040583925 | 0.029815654 | brain innervation |
| ENSDART00000157788 | CABZ01031894.1 | 0.004627449 | -0.75935498 | -0.746140313 | -0.275189537 | -0.221209558 | brain innervation |
| ENSDART00000158213 | akap12b | 0.000529346 | -1.00724688 | -1.146388756 | -0.599808188 | 0.009805796 | brain innervation |
| ENSDART00000158355 | bsk146 | 0.000219336 | -0.973617487 | -0.7163723 | -0.116416923 | 0.102394026 | brain innervation |
| ENSDART00000158429 | tmem42b | 0.001002974 | -0.433937172 | -0.428108528 | -0.048957595 | 0.127616146 | brain innervation |
| ENSDART00000158489 | zgc:77058 | 1.34007E-05 | -0.495737975 | -0.228595493 | 0.141443601 | 0.440592752 | brain innervation |
| ENSDART00000158500 | atp8b2 | 0.000419956 | -0.533001796 | -0.553651375 | -0.17260282 | 0.179842571 | brain innervation |
| ENSDART00000158618 | cpne4b | 4.49433E-05 | -1.041271467 | -0.75854166 | 0.438341768 | 1.045239509 | brain innervation |
| ENSDART00000158671 | sult4a1 | 8.51245E-05 | -0.462609111 | -0.368663113 | -0.083169387 | 0.19862288 | brain innervation |
| ENSDART00000158723 | sncb | 0.008362897 | -0.257429657 | -0.430720868 | -0.069889176 | 0.08182323 | brain innervation |
| ENSDART00000158936 | grin1a | 0.000250511 | -0.976292061 | -1.003442352 | -0.317903979 | 0.15163936 | brain innervation |
| ENSDART00000159104 | atp1b2a | 0.000804628 | -0.488650552 | -0.417675156 | -0.129746035 | 0.084213141 | brain innervation |
| ENSDART00000159249 | oxr1a | 0.000445792 | -0.748854688 | -0.518408458 | -0.301236362 | -0.153948362 | brain innervation |
| ENSDART00000159291 | podxl2 | 0.000184045 | -0.45367951 | -0.207714244 | 0.333702352 | 0.497299671 | brain innervation |
| ENSDART00000159589 | rtn4rl1b | 0.00013806 | -0.618605611 | -0.334861623 | -0.03219779 | 0.419833076 | brain innervation |
| ENSDART00000159601 | serpini1 | 0.000352454 | -0.44709139 | -0.357709199 | -0.198408132 | 0.076823185 | brain innervation |
| ENSDART00000159627 | scn4ba | 0.000451042 | -1.651719386 | -1.747876787 | -0.730948262 | -0.207716251 | brain innervation |
| ENSDART00000159690 | gpr75 | 0.002568245 | -0.483974147 | -0.377125661 | 0.037666989 | 0.123067934 | brain innervation |
| ENSDART00000159693 | NYAP1 | 0.001144712 | -0.548984567 | -0.516611795 | -0.021366781 | 0.000605996 | brain innervation |
| ENSDART00000159727 | pcdh11 | 0.000494919 | -0.294947297 | -0.365638864 | -0.089140913 | 0.17419873 | brain innervation |
| ENSDART00000159805 | CABZ01085924.1 | 0.000283607 | -0.392684277 | -0.456709342 | 0.004929566 | 0.18758642 | brain innervation |
| ENSDART00000159853 | FQ311890.1 | 0.000463544 | -0.276177481 | -0.432407927 | -0.032078728 | 0.27665545 | brain innervation |
| ENSDART00000159942 | kcnc3b | 0.000847723 | -0.794025304 | -1.361623092 | -0.389946342 | 0.009706478 | brain innervation |
| ENSDART00000160018 | DUSP26 | 0.00184704 | -0.660306321 | -0.709450564 | -0.129656337 | 0.007184288 | brain innervation |
| ENSDART00000160138 | sobpa | 0.003891676 | -0.321225255 | -0.895188795 | -0.069068022 | 0.312612038 | brain innervation |
| ENSDART00000160350 | IQSEC1 (1 of many) | 0.012665885 | -0.73025949 | -0.724161984 | -0.227945297 | -0.158150827 | brain innervation |
| ENSDART00000160464 | pcdh1gb2 | 0.037611012 | -0.570850707 | -0.31567681 | -0.173184848 | -0.165810233 | brain innervation |
| ENSDART00000160466 | ndrg4 | 0.001418732 | -0.797536516 | -0.447807808 | -0.265830716 | -0.169347229 | brain innervation |
| ENSDART00000160469 | si:ch211-271b14.1 | 0.005629236 | -0.816113506 | -0.956386746 | -0.419124903 | -0.031051269 | brain innervation |
| ENSDART00000160662 | CABZ01056578.1 | 0.001955985 | -0.678580777 | -0.38089804 | -0.298190661 | -0.130306976 | brain innervation |
| ENSDART00000160789 | SDK2 (1 of many) | 0.006197253 | -0.531586731 | -0.76443238 | -0.126364502 | 0.194536592 | brain innervation |
| ENSDART00000160809 | agap1 | 0.000470359 | -0.12094537 | -0.11171461 | 0.245949776 | 0.322867446 | brain innervation |
| ENSDART00000160811 | b3gat1a | 0.000188707 | -0.509506437 | -0.522178413 | -0.230803048 | 0.152925891 | brain innervation |
| ENSDART00000160881 | pcdh1g30 | 0.001490782 | -0.53895262 | -0.083348601 | 0.159825551 | 0.241016465 | brain innervation |
| ENSDART00000160885 | stbd1 | 0.007301761 | -0.425801769 | -0.465748469 | -0.195945725 | 0.034690374 | brain innervation |
| ENSDART00000161003 | cpne9 | 0.001482057 | -1.056006542 | -1.193405404 | -0.117014879 | 0.290018934 | brain innervation |
| ENSDART00000161115 | CR382281.1 | 0.004084205 | -1.148281272 | -0.70950654 | -0.315399276 | 0.210191426 | brain innervation |
| ENSDART00000161137 | scn1bb | 2.29102E-05 | -0.455645009 | -0.546610376 | 0.183837534 | 0.407926687 | brain innervation |
| ENSDART00000161264 | ntrk3b | 0.000108364 | -0.768053413 | -0.724776652 | -0.30616772 | 0.183034577 | brain innervation |

| | | | | | | | |
|---|---|---|---|---|---|---|---|
| ENSDART00000161436 | pcdh7b | 5.5105E-05 | -0.687665671 | -0.773740554 | -0.245502692 | 0.12030382 | brain innervation |
| ENSDART00000161507 | zgc:153426 | 1.98934E-05 | -1.880708803 | -0.117657621 | 0.865737326 | 1.072245785 | brain innervation |
| ENSDART00000161532 | zfyve9b | 0.000512152 | -0.607365928 | -0.871459147 | -0.090833114 | 0.138377783 | brain innervation |
| ENSDART00000161561 | ANK1 (1 of many) | 0.001021091 | -0.90876402 | -0.980007711 | -0.299110873 | -0.111310605 | brain innervation |
| ENSDART00000161567 | KCNT1 (1 of many) | 0.000451042 | -0.564631347 | -0.618941642 | -0.212804629 | 0.112863315 | brain innervation |
| ENSDART00000161646 | sema6bb | 0.009956893 | -0.22063759 | -0.362707192 | -0.160882628 | 0.032299092 | brain innervation |
| ENSDART00000161700 | st3gal2 | 0.000191101 | -0.330022994 | -0.187434223 | 0.066957229 | 0.392836726 | brain innervation |
| ENSDART00000161725 | map7d1a | 0.007973168 | -0.696007823 | -0.432281688 | 0.042931769 | 0.13512701 | brain innervation |
| ENSDART00000161992 | PAQR9 | 0.016559516 | -0.193753966 | -0.524789375 | -0.118624882 | 0.042415903 | brain innervation |
| ENSDART00000162002 | grm1b | 0.001159496 | -0.568603213 | -0.750999293 | -0.21404532 | -0.094504234 | brain innervation |
| ENSDART00000162023 | plppr2a | 0.001528476 | 0.019417247 | -0.576626688 | 0.082766084 | 0.317388761 | brain innervation |
| ENSDART00000162133 | tenm4 | 0.007324183 | -0.287462387 | -0.412164757 | -0.184969779 | 0.049525758 | brain innervation |
| ENSDART00000162277 | grip1 | 5.26677E-05 | -0.456620328 | -0.2169797 | 0.146006741 | 0.157130316 | brain innervation |
| ENSDART00000162381 | dkk3a | 0.000417493 | -0.551126502 | -0.361561955 | -0.324385943 | 0.074151181 | brain innervation |
| ENSDART00000162601 | scn2b | 0.000115668 | -0.529498392 | -0.64570335 | -0.359290818 | 0.039638437 | brain innervation |
| ENSDART00000162670 | slc8a1b | 0.001547111 | -0.269852688 | -0.529256481 | -0.096682296 | 0.048355338 | brain innervation |
| ENSDART00000162675 | trim2b | 0.004332449 | -0.409757052 | -0.631223683 | -0.07653124 | 0.093922127 | brain innervation |
| ENSDART00000162714 | pcdh10b | 0.00041526 | -0.340908137 | -0.430434424 | -0.069086695 | 0.178421467 | brain innervation |
| ENSDART00000162850 | irx3a | 0.000148815 | -0.694816328 | -0.301932125 | 0.187764734 | 0.208405543 | brain innervation |
| ENSDART00000162984 | cacnb2a | 0.000644738 | -0.567328475 | -0.858575333 | 0.120461387 | 0.550245867 | brain innervation |
| ENSDART00000163006 | magi1b | 1.36917E-05 | -0.558734199 | -0.633598853 | 0.045195354 | 0.572492467 | brain innervation |
| ENSDART00000163096 | gpr22b | 4.91848E-05 | -1.106969479 | -1.366483735 | -0.599957035 | 0.051341663 | brain innervation |
| ENSDART00000163216 | prune2 | 0.005893002 | -0.414992243 | -0.329741076 | -0.094156508 | 0.003976732 | brain innervation |
| ENSDART00000163355 | gria2a | 0.001252059 | -0.37007276 | -0.57694131 | -0.293854127 | 0.240567837 | brain innervation |
| ENSDART00000163519 | CABZ01050166.1 | 0.000257591 | -0.484053035 | -0.532447227 | -0.143157501 | 0.191505756 | brain innervation |
| ENSDART00000163529 | FAM126B | 0.002016009 | -0.519426512 | -0.370293712 | 0.022402011 | 0.032038169 | brain innervation |
| ENSDART00000163539 | scrt1a | 7.10385E-05 | -0.576475687 | -0.187048661 | 0.063865515 | 0.398969662 | brain innervation |
| ENSDART00000163612 | gabrb4 | 0.000645779 | -0.445185281 | -0.610684009 | -0.211332691 | 0.127493005 | brain innervation |
| ENSDART00000163616 | homer1b | 0.002838981 | -0.487597805 | -0.239525084 | -0.239441952 | -0.083482271 | brain innervation |
| ENSDART00000163622 | kirrel3l | 0.013286981 | -0.615947415 | -0.493751402 | -0.207879997 | 0.127527609 | brain innervation |
| ENSDART00000163635 | erbb4b | 0.011458427 | -0.423431984 | -0.772483288 | -0.24962409 | 0.07826798 | brain innervation |
| ENSDART00000163669 | cplx2 | 0.011375495 | -0.402497387 | -0.424820216 | -0.158797592 | 0.046498934 | brain innervation |
| ENSDART00000163677 | NA | 0.000160913 | -0.639905731 | -0.179601491 | 0.106036319 | 0.34272584 | brain innervation |
| ENSDART00000163791 | acss2l | 0.000960572 | -0.694063426 | -0.500876422 | -0.115488475 | -0.045115535 | brain innervation |
| ENSDART00000163867 | gnb1b | 0.000919184 | -0.511340157 | -0.236747213 | 0.117718298 | 0.111335872 | brain innervation |
| ENSDART00000163897 | lgi1b | 0.000144591 | -0.358188713 | -0.274223364 | -0.009680681 | 0.361584537 | brain innervation |
| ENSDART00000163903 | kcna2b | 5.02856E-05 | -1.041609083 | -1.150732339 | -0.278604653 | 0.240702332 | brain innervation |
| ENSDART00000163951 | plppr1 | 0.00335169 | -0.393500272 | -0.316896098 | -0.148613826 | 0.078198855 | brain innervation |
| ENSDART00000164114 | grb2a | 0.000322945 | -0.50918281 | -0.254702947 | 0.293231191 | 0.639895729 | brain innervation |
| ENSDART00000164121 | mboat2b | 0.01376164 | -0.365653488 | -0.229675996 | -0.118902893 | -0.108744854 | brain innervation |
| ENSDART00000164163 | si:dkey-202l22.3 | 3.90156E-05 | -0.676171429 | -1.127391316 | 0.216542828 | 0.638796234 | brain innervation |
| ENSDART00000164178 | prrt2 | 7.10385E-05 | -0.547052118 | -0.283537507 | 0.297793332 | 0.499767197 | brain innervation |
| ENSDART00000164190 | ksr2 | 0.000228698 | -0.449890716 | -0.587945856 | -0.131638865 | 0.158651331 | brain innervation |
| ENSDART00000164326 | si:ch73-119p20.1 | 8.83909E-05 | -0.36329758 | -0.388299747 | 0.050179382 | 0.305653209 | brain innervation |
| ENSDART00000164361 | gcgra | 0.001631005 | -0.404304946 | -0.520606663 | -0.179928762 | -0.009148827 | brain innervation |
| ENSDART00000164597 | si:ch73-127m5.1 | 0.000190025 | -0.712165198 | -0.239448184 | 0.59846121 | 0.690834997 | brain innervation |
| ENSDART00000164879 | slc8a2b | 0.004027736 | -0.298961891 | -0.426517442 | -0.175774312 | 0.049460205 | brain innervation |
| ENSDART00000164982 | cdh4 | 5.24617E-05 | -0.415355609 | -0.489911794 | 0.041403227 | 0.334639192 | brain innervation |
| ENSDART00000164989 | si:ch211-121j5.4 | 0.005899482 | -2.541662275 | -0.597024815 | -0.157115726 | -0.615748868 | brain innervation |
| ENSDART00000165004 | gria3b | 0.001296944 | -0.410652366 | -0.638891942 | -0.274205904 | 0.132979921 | brain innervation |
| ENSDART00000165058 | rims2a | 0.004379828 | -0.759006031 | -0.607036719 | -0.257997898 | -0.111766909 | brain innervation |
| ENSDART00000165082 | ppp1r1b | 0.000811535 | -0.592550768 | -0.433249399 | 0.132918309 | 0.038849346 | brain innervation |
| ENSDART00000165108 | jph3 | 0.008210772 | -0.254573235 | -0.426555215 | 0.040987467 | 0.08534723 | brain innervation |
| ENSDART00000165158 | iqsec3a | 0.002139835 | -0.287263936 | -0.339420666 | -0.160997893 | 0.073213089 | brain innervation |
| ENSDART00000165400 | slc1a2b | 0.01293062 | -0.437547383 | -0.373563248 | -0.176045295 | -0.026222695 | brain innervation |
| ENSDART00000165420 | si:dkey-161j23.5 | 0.000212569 | -0.418752064 | -0.09080604 | 0.112517016 | 0.202865451 | brain innervation |
| ENSDART00000165443 | zgc:153615 | 4.64299E-05 | -0.585165411 | -0.400257189 | -0.265863973 | -0.020005469 | brain innervation |
| ENSDART00000165609 | barhl2 | 0.000401616 | -0.31272019 | -0.374018735 | 0.126371762 | 0.224852149 | brain innervation |
| ENSDART00000165638 | pax10 | 0.001292159 | -0.459647113 | -0.417992165 | -0.200180992 | -0.120970793 | brain innervation |
| ENSDART00000165654 | atp1b2a | 5.13907E-05 | -0.572829136 | -0.493634763 | -0.065881564 | 0.075479325 | brain innervation |
| ENSDART00000165659 | epb41l3a | 0.027884112 | -0.249067696 | -0.637410719 | -0.072556493 | -0.093775549 | brain innervation |
| ENSDART00000165698 | pbx1a | 0.000272902 | -0.205212048 | -0.330154536 | 0.047550564 | 0.321002436 | brain innervation |
| ENSDART00000165744 | kif1b | 0.001276988 | -0.018685925 | -0.126032816 | 0.340319561 | 0.347744455 | brain innervation |
| ENSDART00000165757 | pax6b | 0.019981901 | -1.030621281 | -0.736504391 | -0.510180699 | -0.112500923 | brain innervation |
| ENSDART00000165785 | pcdh10a | 0.001354936 | -0.2607204 | -0.385620436 | -0.040787194 | 0.144409091 | brain innervation |
| ENSDART00000166114 | sema3ab | 0.01892315 | -0.726704016 | -0.305486009 | -0.172433457 | -0.053903823 | brain innervation |
| ENSDART00000166351 | nkrf | 0.010292856 | -0.332043429 | -0.213944917 | -0.117297764 | -0.03436019 | brain innervation |
| ENSDART00000166470 | CU855947.1 | 0.009503355 | -0.432778868 | -0.336734093 | -0.219445916 | -0.079633712 | brain innervation |
| ENSDART00000166502 | satb2 | 1.1419E-05 | -0.540440017 | -0.158642442 | 1.216799418 | 1.496340928 | brain innervation |
| ENSDART00000166508 | fdft1 | 0.007609288 | -0.467410589 | -0.45021858 | 0.057865823 | -0.125768039 | brain innervation |
| ENSDART00000166533 | zgc:123181 | 2.15557E-05 | -0.682536387 | -0.078871471 | 0.212564109 | 0.407498801 | brain innervation |
| ENSDART00000166575 | ppp3ca | 0.000363739 | -0.493387865 | -0.574192875 | 0.281204144 | 0.517021392 | brain innervation |
| ENSDART00000167306 | ldb2b | 0.003805801 | -0.838228207 | -0.415457403 | -0.325734782 | 0.110687477 | brain innervation |
| ENSDART00000167324 | ebf3a | 4.91848E-05 | -0.585714789 | -0.243731213 | 0.563349846 | 0.843370215 | brain innervation |
| ENSDART00000167464 | GNAZ | 0.003016249 | -0.341892906 | -0.155068478 | 0.102581405 | 0.079027974 | brain innervation |

| | | | | | | | |
|---|---|---|---|---|---|---|---|
| ENSDART00000167726 | RYR2 | 5.17084E-05 | -0.554726037 | -0.413134766 | 0.099586067 | 0.289722828 | brain innervation |
| ENSDART00000167995 | napba | 0.006669949 | -0.341681302 | -0.334450461 | -0.176497327 | -0.027366162 | brain innervation |
| ENSDART00000168038 | edil3a | 7.05055E-05 | -0.585018977 | -0.697545399 | -0.17495912 | 0.33520597 | brain innervation |
| ENSDART00000168121 | lzts3a | 0.020009895 | -0.348828928 | -0.543883063 | -0.164874094 | 0.093345065 | brain innervation |
| ENSDART00000168270 | CABZ01048053.1 | 0.000157185 | -0.515099411 | -0.533680852 | 0.225344787 | 0.362581791 | brain innervation |
| ENSDART00000168396 | FO904869.1 | 0.008573911 | -0.490668054 | -0.607434677 | -0.194530972 | 0.12309886 | brain innervation |
| ENSDART00000168453 | slc43a2a | 0.000169466 | -0.36850619 | -0.532753158 | 0.198651765 | 0.248433931 | brain innervation |
| ENSDART00000168568 | CU861477.1 | 0.002569884 | -0.399340079 | -0.694343635 | -0.183948068 | 0.054030168 | brain innervation |
| ENSDART00000168633 | CABZ01085700.1 | 5.17084E-05 | -0.53401794 | -0.080184443 | 0.667052745 | 0.803861054 | brain innervation |
| ENSDART00000168729 | fam196ab | 0.003860077 | -0.54578901 | -0.555866325 | -0.405243034 | 0.063979844 | brain innervation |
| ENSDART00000168749 | nlgn3a | 0.001942462 | -0.417078437 | -0.490224096 | -0.240516544 | -0.086053557 | brain innervation |
| ENSDART00000168805 | kcnk9 | 0.002102587 | -0.553605245 | -0.952161118 | -0.308244106 | 0.289642563 | brain innervation |
| ENSDART00000168885 | nrg1 | 0.000322759 | -0.78621851 | -0.719452399 | 0.167065606 | 0.289901187 | brain innervation |
| ENSDART00000168969 | htr7c | 0.004472999 | -0.617759768 | -0.791263551 | -0.091828465 | 0.038777953 | brain innervation |
| ENSDART00000169081 | mtmr7b | 0.000669667 | -0.302136082 | -0.184242994 | 0.142322853 | 0.09943817 | brain innervation |
| ENSDART00000169127 | adgrb3 | 0.004610119 | -0.338634159 | -0.316792657 | -0.154400064 | 0.089681207 | brain innervation |
| ENSDART00000169243 | znf384l | 0.012912717 | -0.366831524 | -0.600767675 | -0.247311028 | 0.205515141 | brain innervation |
| ENSDART00000169552 | cdh13 | 0.000225609 | -0.778914955 | -0.463881372 | -0.06287293 | 0.145633136 | brain innervation |
| ENSDART00000169626 | kcnc1a | 0.003703617 | -0.804732723 | -1.394467436 | -0.226498418 | 0.073837621 | brain innervation |
| ENSDART00000169768 | slc8a3 | 0.004351059 | -0.133596734 | -0.60366341 | -0.159093114 | 0.187240856 | brain innervation |
| ENSDART00000169948 | GRM7 | 0.005631068 | -0.415342332 | -0.376852993 | -0.226859206 | -0.023200649 | brain innervation |
| ENSDART00000170089 | epha4b | 0.011264004 | -0.586426265 | -0.698463168 | -0.228203192 | -0.016930318 | brain innervation |
| ENSDART00000170165 | cep97 | 0.016618446 | -4.277597338 | -0.154895502 | -0.337498335 | -0.461056374 | brain innervation |
| ENSDART00000170396 | apbb2b | 2.95145E-05 | -0.997027217 | -0.745617105 | 0.597402109 | 1.180709512 | brain innervation |
| ENSDART00000170423 | jakmip3 | 0.002756705 | -0.243674673 | -0.387799515 | -0.202768181 | 0.090619484 | brain innervation |
| ENSDART00000170453 | slc20a1b | 0.003686476 | -0.460714141 | -0.247481668 | -0.017046699 | -0.023964599 | brain innervation |
| ENSDART00000170460 | nrxn3b | 0.003312965 | -0.2276052 | -0.615850363 | -0.134586161 | -0.003465474 | brain innervation |
| ENSDART00000170510 | RIMBP2 (1 of many) | 0.000415146 | -0.653406892 | -0.568163429 | -0.174788334 | -0.007324529 | brain innervation |
| ENSDART00000170569 | syt12 | 1.45694E-05 | -0.747896951 | -0.906145295 | -0.362441574 | -0.047384497 | brain innervation |
| ENSDART00000170620 | ctxn1 | 0.006828168 | -1.120908397 | -0.319127099 | 0.144625495 | 0.239695278 | brain innervation |
| ENSDART00000170680 | ptprdb | 1.22823E-05 | -0.822263801 | -1.170454394 | 0.28663796 | 0.697055522 | brain innervation |
| ENSDART00000170700 | nacc1b | 0.000772038 | -0.295710389 | -0.416714615 | 0.013950982 | 0.110349783 | brain innervation |
| ENSDART00000170839 | pcdh1g26 | 7.10385E-05 | -0.648364026 | -0.34360777 | 0.683964001 | 0.607743239 | brain innervation |
| ENSDART00000170854 | gphnb | 2.98725E-05 | -0.471165414 | -0.533169869 | -0.144401896 | 0.220388206 | brain innervation |
| ENSDART00000170875 | BX663610.1 | 0.008814649 | -0.528067918 | -0.63401091 | -0.217086155 | -0.025338796 | brain innervation |
| ENSDART00000170888 | pkma | 0.009639463 | -0.382873513 | -0.359191464 | -0.202639045 | 0.04775771 | brain innervation |
| ENSDART00000170932 | rims2a | 0.000444888 | -0.482409185 | -0.599060629 | -0.273352321 | -0.056203652 | brain innervation |
| ENSDART00000170952 | pvrl2l | 0.000405848 | -0.402141005 | -0.731362921 | -0.043297784 | 0.281757642 | brain innervation |
| ENSDART00000171013 | tenm4 | 0.009203641 | -0.557784731 | -0.616815207 | -0.086642376 | -0.101260046 | brain innervation |
| ENSDART00000171073 | tox2 | 0.012896891 | -0.713271279 | -0.691854818 | -0.08505138 | -0.233730891 | brain innervation |
| ENSDART00000171114 | si:dkey-18j18.3 | 0.00335169 | -5.054012109 | -5.054146995 | -1.846245188 | 0.223162792 | brain innervation |
| ENSDART00000171288 | magi2b | 0.003659486 | -0.2674353 | -0.525609031 | -0.078206648 | -0.00532511 | brain innervation |
| ENSDART00000171306 | stxbp1a | 0.001277839 | -0.614795684 | -0.539858263 | -0.126841305 | 0.057169963 | brain innervation |
| ENSDART00000171359 | asic1b | 0.000948363 | -0.583045866 | -0.669491195 | -0.186449925 | 0.212036898 | brain innervation |
| ENSDART00000171490 | PCDH8 | 2.78074E-05 | -0.518696681 | -0.628285802 | -0.168709969 | 0.408565003 | brain innervation |
| ENSDART00000171494 | ssbp2 | 0.000598084 | -0.298958691 | -0.315587827 | -0.028952479 | 0.152638316 | brain innervation |
| ENSDART00000171506 | NA | 6.00847E-05 | -0.792662919 | -0.4993043 | 0.08528346 | 0.180572156 | brain innervation |
| ENSDART00000171571 | FAM184A (1 of many) | 0.001237101 | -0.84950254 | -1.068290057 | 0.155587747 | 0.361779982 | brain innervation |
| ENSDART00000171594 | mef2aa | 0.010231208 | -0.649844171 | -0.685014218 | -0.301258241 | -0.003741614 | brain innervation |
| ENSDART00000171674 | abhd8a | 0.000204711 | -0.482604068 | -0.462135418 | 0.077091696 | 0.23737745 | brain innervation |
| ENSDART00000171704 | soul4 | 0.004085991 | -0.335441282 | -0.335430337 | -0.104813252 | -0.062286587 | brain innervation |
| ENSDART00000171951 | slit2 | 0.003148896 | -0.546372344 | -0.56571293 | -0.270916548 | -0.035288929 | brain innervation |
| ENSDART00000172232 | sv2a | 0.00145694 | -0.325501969 | -0.478402861 | -0.246153159 | 0.031695064 | brain innervation |
| ENSDART00000172465 | gnb1b | 0.003905267 | -0.429380032 | -0.139749463 | 0.050389396 | 0.071415749 | brain innervation |
| ENSDART00000172552 | dyrk2 | 0.007164707 | -1.00865602 | -4.097564958 | -0.48472737 | -0.263450257 | brain innervation |
| ENSDART00000172624 | chn1 | 0.001420911 | -0.429491719 | -0.379515323 | -0.144891996 | -0.16828013 | brain innervation |
| ENSDART00000172634 | ryr3 | 0.003072278 | -1.057085419 | -1.586723574 | -0.163110923 | 0.331797887 | brain innervation |
| ENSDART00000172638 | gria3a | 0.000581789 | -0.308989617 | -0.39028425 | -0.157465183 | 0.147319123 | brain innervation |
| ENSDART00000172645 | unc5da | 0.000935288 | -0.375739047 | -0.576943357 | -0.183677467 | 0.03667029 | brain innervation |
| ENSDART00000172664 | kirrel3l | 0.000274768 | -0.579597094 | -0.334224423 | -0.00269739 | 0.256102934 | brain innervation |
| ENSDART00000173017 | atp2b2 | 0.002838732 | -0.485284755 | -0.507921276 | -0.213758166 | 0.050306247 | brain innervation |
| ENSDART00000173060 | rph3ab | 4.91848E-05 | -0.917567753 | -0.595167162 | 0.673778236 | 0.835535825 | brain innervation |
| ENSDART00000173108 | gpc3 | 2.23813E-05 | -0.979170892 | -0.757011653 | 0.064815285 | 0.413031736 | brain innervation |
| ENSDART00000173126 | klhl4 | 0.012397761 | -0.765151123 | -0.487744119 | -0.047554388 | -0.018081928 | brain innervation |
| ENSDART00000173192 | pcdh1b | 0.001151232 | -0.448371004 | -0.996454455 | -0.461950735 | 0.179661768 | brain innervation |
| ENSDART00000173210 | kcnab2a | 6.7807E-05 | -0.633080178 | -0.760216594 | -0.295624932 | -0.02803528 | brain innervation |
| ENSDART00000173228 | slc6a11b | 0.000233744 | -1.626653664 | -2.205227269 | -0.432079303 | 0.774903159 | brain innervation |
| ENSDART00000173301 | sfxn5b | 0.004367955 | -0.333126218 | -0.098925577 | -0.061350927 | 0.22562903 | brain innervation |
| ENSDART00000173421 | lingo2a | 0.000589104 | -0.727375414 | -0.584905703 | -0.169839849 | 0.100856447 | brain innervation |
| ENSDART00000000804 | slc8a1b | 0.0088045 | -0.216070414 | -0.730017061 | -0.564883211 | -0.309532287 | downregulated during regeneration - early |
| ENSDART00000000876 | nr4a1 | 0.000894056 | -2.262372442 | -2.399082592 | -2.145415905 | -1.543723254 | downregulated during regeneration - early |
| ENSDART00000001907 | slc16a3 | 0.026714705 | -0.228617593 | -0.852511558 | -0.452127784 | -0.215137636 | downregulated during regeneration - early |
| ENSDART00000003008 | gad1b | 0.009594186 | -0.375298425 | -0.380219055 | -0.456528647 | -0.347996362 | downregulated during regeneration - early |
| ENSDART00000003548 | znf385a | 0.001069696 | -0.544762633 | -0.635515273 | -0.316063497 | -0.128464507 | downregulated during regeneration - early |
| ENSDART00000003736 | anos1b | 0.02243441 | -0.470963792 | -0.442368787 | -0.444507654 | -0.318716779 | downregulated during regeneration - early |

| | | | | | | | |
|---|---|---|---|---|---|---|---|
| ENSDART00000004034 | hpca | 0.000804628 | -0.538662149 | -0.391929378 | -0.242660916 | -0.195867715 | downregulated during regeneration - early |
| ENSDART00000004626 | sec62 | 0.03450478 | -0.335700502 | -0.25907023 | -0.14783183 | -0.155829549 | downregulated during regeneration - early |
| ENSDART00000005299 | hsd17b12a | 0.040876822 | -0.345584522 | -0.261321345 | -0.169519073 | -0.263954396 | downregulated during regeneration - early |
| ENSDART00000005724 | ncanb | 0.007781565 | -0.308688969 | -0.586976696 | -0.410319756 | -0.132925594 | downregulated during regeneration - early |
| ENSDART00000005738 | slitrk2 | 0.002539213 | -0.40615183 | -0.499760308 | -0.343598039 | -0.061449053 | downregulated during regeneration - early |
| ENSDART00000005842 | fgf1a | 0.001013659 | -0.526483511 | -0.462706743 | -0.482028176 | -0.1224057 | downregulated during regeneration - early |
| ENSDART00000006417 | pgm1 | 0.001451795 | -0.367653769 | -0.345273975 | -0.348210636 | -0.086795004 | downregulated during regeneration - early |
| ENSDART00000007401 | MAP3K13 | 0.019500385 | -0.231843501 | -0.363439198 | -0.331818795 | -0.269324908 | downregulated during regeneration - early |
| ENSDART00000007522 | anos1a | 0.001275399 | -0.500742525 | -0.407287946 | -0.526812004 | -0.211300468 | downregulated during regeneration - early |
| ENSDART00000007630 | nhlh2 | 0.010056242 | -0.28794611 | -0.324673306 | -0.385652145 | -0.261092419 | downregulated during regeneration - early |
| ENSDART00000007778 | grik1a | 0.008380474 | -0.423674769 | -0.190804556 | -0.271114709 | -0.0631136 | downregulated during regeneration - early |
| ENSDART00000008287 | pgam1a | 0.011948793 | -0.270559913 | -0.416576315 | -0.324017463 | -0.168254728 | downregulated during regeneration - early |
| ENSDART00000009337 | eno1a | 0.014647445 | -0.396364992 | -0.428345122 | -0.288646494 | -0.130833414 | downregulated during regeneration - early |
| ENSDART00000009569 | slc12a5b | 0.001340966 | -0.543274129 | -0.639831062 | -0.420973005 | -0.072437192 | downregulated during regeneration - early |
| ENSDART00000009653 | kcna1b | 0.000806652 | -0.51649998 | -0.917921415 | -0.602149268 | -0.420615578 | downregulated during regeneration - early |
| ENSDART00000009777 | glra3 | 0.000683221 | -0.385455571 | -0.554614528 | -0.516083394 | -0.22262256 | downregulated during regeneration - early |
| ENSDART00000009938 | tcf12 | 0.025246927 | -0.385864267 | -0.351608927 | -0.283149928 | -0.323464659 | downregulated during regeneration - early |
| ENSDART00000010257 | fam73a | 0.046446554 | -0.203941778 | -0.403469011 | -0.199714259 | -0.045216291 | downregulated during regeneration - early |
| ENSDART00000010452 | zgc:91860 | 0.008679209 | -0.346186809 | -0.437676548 | -0.463741509 | -0.203120402 | downregulated during regeneration - early |
| ENSDART00000011287 | aqp4 | 0.010984291 | -0.584037248 | -0.688442177 | -0.670108654 | -0.488547149 | downregulated during regeneration - early |
| ENSDART00000011627 | irx7 | 0.004741095 | -0.27830639 | -0.509115666 | -0.233145912 | -0.150983114 | downregulated during regeneration - early |
| ENSDART00000011936 | ccdc106a | 0.067939329 | -0.452156269 | -0.26003137 | -0.330808493 | -0.261299438 | downregulated during regeneration - early |
| ENSDART00000012673 | gnb3a | 0.000657061 | -0.611271729 | -0.360057677 | -0.265096652 | -0.439594606 | downregulated during regeneration - early |
| ENSDART00000013003 | tfap2b | 0.008192536 | -0.299930992 | -0.578681033 | -0.25971747 | -0.279510711 | downregulated during regeneration - early |
| ENSDART00000013117 | syt5b | 0.003648083 | -0.473395784 | -0.439637374 | -0.372750952 | -0.393185839 | downregulated during regeneration - early |
| ENSDART00000013360 | ppp1r3cb | 0.004590888 | -0.357586146 | -0.346668382 | -0.405594995 | -0.235559662 | downregulated during regeneration - early |
| ENSDART00000014668 | pcsk1 | 0.025485875 | -0.444651661 | -0.49391188 | -0.327964124 | -0.406864512 | downregulated during regeneration - early |
| ENSDART00000014726 | tp53i11b | 0.03271715 | -0.378677803 | -0.267454179 | -0.193372925 | -0.148965425 | downregulated during regeneration - early |
| ENSDART00000015333 | gbx2 | 0.065341577 | -0.202064704 | -0.378779754 | -0.261598975 | -0.176372699 | downregulated during regeneration - early |
| ENSDART00000015956 | efna1b | 0.054157717 | -0.412330789 | -0.211680079 | -0.214205067 | -0.214663513 | downregulated during regeneration - early |
| ENSDART00000016181 | ndrg3a | 0.015357816 | -0.359622862 | -0.274997666 | -0.226177976 | -0.143111599 | downregulated during regeneration - early |
| ENSDART00000016983 | spon1a | 0.001261708 | -0.566333419 | -0.54864414 | -0.425205045 | -0.214533594 | downregulated during regeneration - early |
| ENSDART00000017259 | fgf13a | 0.058775394 | -0.403933655 | -0.684393894 | -0.638256176 | -0.336496098 | downregulated during regeneration - early |
| ENSDART00000017551 | slc6a1b | 0.011705967 | -0.387372781 | -0.441576322 | -0.34448646 | -0.23699282 | downregulated during regeneration - early |
| ENSDART00000017599 | rem1 | 0.048973345 | -0.151492945 | -0.383680013 | -0.100323724 | -0.098680467 | downregulated during regeneration - early |
| ENSDART00000017763 | CABZ01071180.1 | 0.028986846 | -0.429470778 | -0.866508186 | -0.658991937 | -0.375481088 | downregulated during regeneration - early |
| ENSDART00000017774 | cacng5a | 0.002016009 | -0.685385238 | -0.470575642 | -0.348446742 | -0.159039188 | downregulated during regeneration - early |
| ENSDART00000018501 | opn4.1 | 0.0088045 | -0.494086134 | -0.180456491 | -0.369290026 | -0.100666897 | downregulated during regeneration - early |
| ENSDART00000018654 | rnd1b | 0.003262292 | -0.711104866 | -0.668463449 | -0.311306501 | -0.458881777 | downregulated during regeneration - early |
| ENSDART00000019748 | lin7a | 0.004883865 | -0.334365216 | -0.254636287 | -0.400398823 | -0.216397551 | downregulated during regeneration - early |
| ENSDART00000019905 | fndc4b | 0.000292971 | -0.870561937 | -0.746602431 | -0.973680622 | -0.402474596 | downregulated during regeneration - early |
| ENSDART00000019949 | ndrg2 | 0.015961295 | -0.366275522 | -0.294111898 | -0.252300979 | -0.169375948 | downregulated during regeneration - early |
| ENSDART00000020054 | opcml | 0.001389805 | -0.412711216 | -0.540582472 | -0.347997034 | -0.057254512 | downregulated during regeneration - early |
| ENSDART00000020249 | dusp5 | 0.033661318 | -0.842223428 | -0.88819591 | -1.001072274 | -0.721604289 | downregulated during regeneration - early |
| ENSDART00000020741 | aldoaa | 0.004379828 | -0.416493866 | -0.444493146 | -0.351916406 | -0.154405211 | downregulated during regeneration - early |
| ENSDART00000021605 | LRRC4C | 0.002327436 | -0.456880748 | -0.587108705 | -0.299621227 | -0.131466817 | downregulated during regeneration - early |
| ENSDART00000021609 | gad2 | 0.02923464 | -0.376538118 | -0.375877531 | -0.287892544 | -0.231277899 | downregulated during regeneration - early |
| ENSDART00000022290 | mdh1aa | 0.00652818 | -0.386096556 | -0.387245145 | -0.361974171 | -0.162628891 | downregulated during regeneration - early |
| ENSDART00000022549 | atp1b3a | 0.005736694 | -0.438514495 | -0.341784188 | -0.389598734 | -0.186751495 | downregulated during regeneration - early |
| ENSDART00000022579 | GABRG3 | 0.002100653 | -0.524427233 | -0.895243712 | -0.602147004 | -0.152848624 | downregulated during regeneration - early |
| ENSDART00000022625 | nrarpb | 0.033808649 | -0.295040721 | -0.303522559 | -0.284578601 | -0.228026779 | downregulated during regeneration - early |
| ENSDART00000023562 | CABZ01041610.1 | 0.000144061 | -1.012987922 | -1.132303657 | -0.526387248 | -0.210938966 | downregulated during regeneration - early |
| ENSDART00000024304 | per3 | 0.008088213 | -0.478794008 | -0.23341732 | -0.352073188 | -0.024889246 | downregulated during regeneration - early |
| ENSDART00000024832 | stat5a | 0.004077075 | -0.212236095 | -0.408877789 | -0.375060924 | -0.185569558 | downregulated during regeneration - early |
| ENSDART00000025466 | slc18a2 | 0.042155205 | -0.576881609 | -0.281211576 | -0.258781682 | -0.379077041 | downregulated during regeneration - early |
| ENSDART00000026174 | dgkh | 0.000322759 | -0.6307495 | -1.007774241 | -0.588261808 | -0.107017931 | downregulated during regeneration - early |
| ENSDART00000026765 | slc18a3a | 0.00377171 | -0.342520426 | -0.454539851 | -0.359489571 | -0.310297658 | downregulated during regeneration - early |
| ENSDART00000026766 | aldocb | 0.007676315 | -0.483629021 | -0.540701781 | -0.406696036 | -0.135568984 | downregulated during regeneration - early |
| ENSDART00000027465 | cacna2d4b | 0.014868435 | -0.370016882 | -0.141565087 | -0.247577188 | 0.068079685 | downregulated during regeneration - early |
| ENSDART00000028500 | nxn | 0.008517342 | -0.356492001 | -0.304319321 | -0.337794036 | -0.002781587 | downregulated during regeneration - early |
| ENSDART00000029703 | kcnh1a | 1.36144E-05 | -0.531746722 | -1.366001544 | -0.565478925 | 0.09193536 | downregulated during regeneration - early |
| ENSDART00000029981 | ppp3cb | 0.001261708 | -0.453839737 | -0.543662557 | -0.385030051 | 0.007551117 | downregulated during regeneration - early |
| ENSDART00000030811 | cables2b | 0.004679084 | -0.198257533 | -0.480718843 | -0.449759807 | -0.060172476 | downregulated during regeneration - early |
| ENSDART00000031139 | slc24a4b | 0.00106989 | -0.756345109 | -1.188291659 | -0.474432561 | -0.06484409 | downregulated during regeneration - early |
| ENSDART00000031265 | rtn4r | 0.006344596 | -0.753039937 | -0.836146377 | -0.430012914 | -0.160533624 | downregulated during regeneration - early |
| ENSDART00000032695 | asic4a | 0.021143724 | -0.405928595 | -0.689262684 | -0.527033258 | -0.378352566 | downregulated during regeneration - early |
| ENSDART00000033248 | fam107b | 0.004517445 | -0.588148397 | -0.439287754 | -0.362004691 | -0.311865235 | downregulated during regeneration - early |
| ENSDART00000033362 | gatad2b | 0.00857344 | -0.40244942 | -0.286045422 | -0.214483951 | -0.084688348 | downregulated during regeneration - early |
| ENSDART00000034421 | cdk14 | 0.046595201 | -0.37810724 | -0.738234108 | -0.33600573 | -0.064327168 | downregulated during regeneration - early |
| ENSDART00000034737 | cpne8 | 0.000212569 | -0.307817154 | -0.810818352 | -0.660194536 | 0.005050706 | downregulated during regeneration - early |
| ENSDART00000034883 | mcf2a | 0.007383608 | -0.149015362 | -0.337699782 | -0.173487291 | 0.057006752 | downregulated during regeneration - early |
| ENSDART00000036050 | rs1a | 0.003706014 | -0.391328986 | -0.392177441 | -0.469736828 | -0.271292581 | downregulated during regeneration - early |
| ENSDART00000036472 | zgc:110852 | 0.003559691 | -0.482143709 | -0.509026678 | -0.613205563 | -0.094495965 | downregulated during regeneration - early |
| ENSDART00000036997 | camk2n1a | 0.004142036 | -0.377223565 | -0.40737699 | -0.242898936 | -0.23919335 | downregulated during regeneration - early |
| ENSDART00000037007 | tpi1a | 0.002762116 | -0.595344218 | -0.412573356 | -0.319655345 | -0.136192131 | downregulated during regeneration - early |

| | | | | | | | |
|---|---|---|---|---|---|---|---|
| ENSDART00000037922 | slc6a8 | 0.004497123 | -0.298142259 | -0.526146924 | -0.491663812 | -0.167137822 | downregulated during regeneration - early |
| ENSDART00000038290 | crhb | 0.005277161 | -0.533237668 | -0.584543834 | -0.601107267 | -0.51443145 | downregulated during regeneration - early |
| ENSDART00000039277 | lhfpl3 | 0.000949477 | -0.493635642 | -0.41740998 | -0.318633213 | -0.189800628 | downregulated during regeneration - early |
| ENSDART00000039571 | camk2a | 0.001363478 | -0.838498146 | -0.905374301 | -0.593078838 | -0.446882891 | downregulated during regeneration - early |
| ENSDART00000040086 | pacsin1a | 0.019909357 | -0.311462824 | -0.286213879 | -0.425986821 | -0.031983095 | downregulated during regeneration - early |
| ENSDART00000040275 | kcnj11l | 0.033377037 | -0.790164361 | -0.507671216 | -0.85342057 | -0.51870362 | downregulated during regeneration - early |
| ENSDART00000040502 | trpc5a | 0.004129329 | -0.568729425 | -1.014519921 | -0.522696272 | -0.18638518 | downregulated during regeneration - early |
| ENSDART00000040672 | mecp2 | 0.047533941 | -0.242382035 | -0.413395534 | -0.149398689 | -0.148226048 | downregulated during regeneration - early |
| ENSDART00000041388 | cacng2a | 0.015156186 | -0.088843708 | -0.391878721 | -0.181623249 | -0.021159404 | downregulated during regeneration - early |
| ENSDART00000041503 | slc4a4a | 0.010983086 | -0.308542931 | -0.316588825 | -0.297229141 | -0.158105481 | downregulated during regeneration - early |
| ENSDART00000041861 | syt1a | 0.033833708 | -0.326937152 | -0.373717465 | -0.346529457 | -0.191055943 | downregulated during regeneration - early |
| ENSDART00000042386 | unm_sa1261 | 0.008933267 | -0.368338169 | -0.259611498 | -0.383565927 | -0.183992374 | downregulated during regeneration - early |
| ENSDART00000043058 | TENM2 | 0.005410022 | -0.559723778 | -0.553182426 | -0.524601572 | -0.11140967 | downregulated during regeneration - early |
| ENSDART00000043180 | gria3b | 0.000788653 | -0.628974157 | -0.919920549 | -0.567970281 | 0.039022237 | downregulated during regeneration - early |
| ENSDART00000043945 | NA | 0.013000886 | -0.527606114 | -0.345502165 | -0.38556129 | -0.085040536 | downregulated during regeneration - early |
| ENSDART00000044294 | fryb | 0.002449444 | -0.196859444 | -0.540115494 | -0.186847147 | -0.085192364 | downregulated during regeneration - early |
| ENSDART00000044733 | NPBWR2 | 0.007828891 | -0.533926323 | -0.49881296 | -0.385102988 | -0.276547602 | downregulated during regeneration - early |
| ENSDART00000044735 | gria1b | 0.029927553 | -0.281408167 | -0.37749512 | -0.41239512 | -0.179683873 | downregulated during regeneration - early |
| ENSDART00000044896 | camk2d2 | 0.00145694 | -0.471410347 | -0.378784897 | -0.323522916 | -0.120871605 | downregulated during regeneration - early |
| ENSDART00000044986 | rnd1a | 0.003670956 | -0.894483212 | -0.275620607 | -0.461458 | -0.242681728 | downregulated during regeneration - early |
| ENSDART00000045616 | gabbr1b | 0.008210772 | -0.290531763 | -0.52893984 | -0.416723119 | -0.192345482 | downregulated during regeneration - early |
| ENSDART00000045842 | rcan3 | 0.002351134 | -0.531033223 | -0.615136042 | -0.34287969 | -0.068102336 | downregulated during regeneration - early |
| ENSDART00000046498 | sema3fa | 0.004560464 | -0.424113863 | -0.318396861 | -0.293026202 | -0.101692261 | downregulated during regeneration - early |
| ENSDART00000046542 | igf1rb | 0.011264004 | -0.265329623 | -0.432225627 | -0.135218314 | -0.147354602 | downregulated during regeneration - early |
| ENSDART00000046663 | camta1b | 0.022220725 | -0.175908238 | -0.543108898 | -0.289499765 | -0.001152787 | downregulated during regeneration - early |
| ENSDART00000047416 | slc4a8 | 0.004215641 | -0.393460236 | -0.539636776 | -0.542467324 | -0.330988764 | downregulated during regeneration - early |
| ENSDART00000047541 | bach1b | 0.039802589 | -0.343772395 | -0.510091534 | -0.468889885 | -0.333515696 | downregulated during regeneration - early |
| ENSDART00000048036 | gem | 0.018018574 | -0.746612275 | -0.695957849 | -0.479539177 | -0.330372498 | downregulated during regeneration - early |
| ENSDART00000048365 | syt6b | 0.014002312 | -0.461087231 | -0.40037241 | -0.458496339 | -0.153902769 | downregulated during regeneration - early |
| ENSDART00000048432 | dlg4a | 0.00231226 | -0.380839229 | -0.483618741 | -0.31591813 | -0.034480778 | downregulated during regeneration - early |
| ENSDART00000048775 | mbd3b | 0.027491007 | -0.291847963 | -0.361950456 | -0.358913558 | -0.222238531 | downregulated during regeneration - early |
| ENSDART00000048819 | rassf2a | 0.006222767 | -0.204366055 | -0.399787028 | -0.377471637 | -0.118487631 | downregulated during regeneration - early |
| ENSDART00000048893 | pcbp3 | 0.004826213 | -0.37198127 | -0.476664366 | -0.345261768 | -0.190932807 | downregulated during regeneration - early |
| ENSDART00000049036 | zgc:92275 | 0.008658551 | -0.447578375 | -0.84153571 | -0.52710535 | -0.386325614 | downregulated during regeneration - early |
| ENSDART00000049240 | tob1a | 0.008289796 | -0.553621268 | -0.528794995 | -0.629368997 | -0.384085742 | downregulated during regeneration - early |
| ENSDART00000050018 | cnksr1 | 0.010245667 | 0.043033394 | -0.349786212 | -0.053406139 | -0.094004348 | downregulated during regeneration - early |
| ENSDART00000050559 | sh3rf1 | 0.018714496 | -0.402839399 | -0.313691549 | -0.2479685 | -0.17125224 | downregulated during regeneration - early |
| ENSDART00000051723 | si:ch211-193k19.1 | 0.001416864 | -0.524075663 | -0.333228258 | -0.259244771 | -0.088981638 | downregulated during regeneration - early |
| ENSDART00000052083 | fjx1 | 0.030276337 | -0.061315226 | -0.667932816 | -0.149604151 | 0.07387804 | downregulated during regeneration - early |
| ENSDART00000052620 | npy | 0.000988999 | -0.456788991 | -0.449905237 | -0.529362885 | -0.310086201 | downregulated during regeneration - early |
| ENSDART00000052989 | ache | 0.010699261 | -0.369275035 | -0.306496087 | -0.342496448 | -0.099646921 | downregulated during regeneration - early |
| ENSDART00000053750 | acsl2 | 7.10385E-05 | -0.784570064 | -0.626588121 | -0.673780261 | -0.449722468 | downregulated during regeneration - early |
| ENSDART00000054674 | mtnr1aa | 0.014726917 | -0.46530953 | -0.508091587 | -0.367618326 | -0.477932737 | downregulated during regeneration - early |
| ENSDART00000054736 | bhlhe23 | 0.004928313 | -0.369485488 | -0.514619457 | -0.390416516 | -0.190127988 | downregulated during regeneration - early |
| ENSDART00000054790 | zmp:0000001069 | 0.000507818 | -0.970537582 | -1.254323928 | -0.841864806 | -0.072822598 | downregulated during regeneration - early |
| ENSDART00000055019 | ndufa4 | 0.000829971 | -0.243803126 | -0.403243102 | -0.444723985 | -0.066106721 | downregulated during regeneration - early |
| ENSDART00000055134 | ogfr | 0.011591036 | -0.383562939 | -0.286066309 | -0.21229445 | -0.172842524 | downregulated during regeneration - early |
| ENSDART00000055262 | cdk5r1a | 0.023933397 | -0.345552765 | -0.416237328 | -0.217352163 | -0.195826689 | downregulated during regeneration - early |
| ENSDART00000055264 | CA10 (1 of many) | 0.032319936 | -0.387267589 | -0.307777286 | -0.264643025 | -0.127280824 | downregulated during regeneration - early |
| ENSDART00000055465 | si:ch211-149k23.9 | 0.051826156 | -0.23663044 | -0.394558494 | -0.331258304 | -0.323817823 | downregulated during regeneration - early |
| ENSDART00000055890 | znf385c | 0.011543549 | -0.397413591 | -0.444425008 | -0.419820845 | -0.297765515 | downregulated during regeneration - early |
| ENSDART00000056035 | PMM1 | 0.019075692 | -0.318342084 | -0.457172695 | -0.421149458 | -0.159828821 | downregulated during regeneration - early |
| ENSDART00000056721 | ldhd | 0.006978095 | -0.445248015 | -0.387879357 | -0.435006793 | -0.179899207 | downregulated during regeneration - early |
| ENSDART00000056810 | drd1b | 0.002481856 | -0.453660686 | -0.522513213 | -0.435699921 | -0.247093083 | downregulated during regeneration - early |
| ENSDART00000057258 | slc12a5a | 0.009167819 | -0.776929597 | -2.750292921 | -0.618148385 | -0.058229169 | downregulated during regeneration - early |
| ENSDART00000057422 | pacsin1a | 0.016789533 | -0.590114235 | -0.673330755 | -0.518895915 | -0.206589128 | downregulated during regeneration - early |
| ENSDART00000058415 | zmp:0000001075 | 0.017815971 | -0.428240629 | -0.582010832 | -0.289943093 | -0.116590577 | downregulated during regeneration - early |
| ENSDART00000058706 | fosaa | 0.004423084 | -1.574082598 | -1.225695241 | -1.240461281 | -1.154909276 | downregulated during regeneration - early |
| ENSDART00000059446 | znf385b | 0.027304375 | -0.260906147 | -0.315001382 | -0.161772411 | -0.110285684 | downregulated during regeneration - early |
| ENSDART00000059489 | prmt8b | 0.00072061 | -0.385132635 | -0.375733065 | -0.525036565 | 0.065426538 | downregulated during regeneration - early |
| ENSDART00000059984 | deptor | 0.033009621 | -0.356805464 | -0.334367455 | -0.324987791 | -0.320690594 | downregulated during regeneration - early |
| ENSDART00000060056 | tpi1b | 0.004694 | -0.330637537 | -0.371028005 | -0.31545087 | -0.122811105 | downregulated during regeneration - early |
| ENSDART00000060321 | RAMP1 | 0.003256496 | -0.401526014 | -0.370225897 | -0.385056936 | -0.044799362 | downregulated during regeneration - early |
| ENSDART00000060356 | dgkh | 0.001359425 | -0.649006381 | -0.810232451 | -0.629585196 | -0.212393687 | downregulated during regeneration - early |
| ENSDART00000060946 | sgsm1b | 0.010421156 | -0.173061984 | -0.355540157 | -0.215401885 | -0.063998625 | downregulated during regeneration - early |
| ENSDART00000062150 | zgc:77752 | 0.038990469 | -0.401367403 | -0.150322453 | -0.162634546 | -0.096457711 | downregulated during regeneration - early |
| ENSDART00000062603 | cadm1b | 0.040704846 | -0.182938767 | -0.538377364 | -0.267052947 | -0.164493851 | downregulated during regeneration - early |
| ENSDART00000062736 | coasy | 0.038825302 | -0.31852882 | -0.407402551 | -0.404610538 | -0.370618301 | downregulated during regeneration - early |
| ENSDART00000063551 | ppm1e | 0.011668571 | -0.4375512 | -0.67599947 | -0.28655858 | -0.160579724 | downregulated during regeneration - early |
| ENSDART00000063703 | si:dkey-71h2.2 | 0.004477335 | -0.547517928 | -0.696070326 | -0.446988451 | -0.222080286 | downregulated during regeneration - early |
| ENSDART00000063704 | crip3 | 0.023735552 | -0.295432006 | -0.264072284 | -0.393391872 | -0.087537117 | downregulated during regeneration - early |
| ENSDART00000063706 | fndc4a | 1.36144E-05 | -0.898991439 | -1.194204845 | -0.829294946 | -0.104966589 | downregulated during regeneration - early |
| ENSDART00000063779 | efhd1 | 0.000461528 | -0.527832136 | -0.620416333 | -0.244685479 | -0.230757866 | downregulated during regeneration - early |
| ENSDART00000063816 | kcnk3a | 0.00834001 | -0.460164643 | -0.659361375 | -0.48206205 | -0.175491486 | downregulated during regeneration - early |
| ENSDART00000064241 | nrxn3a | 0.014683084 | -0.593329637 | -0.86602502 | -0.45366796 | -0.193651725 | downregulated during regeneration - early |

| | | | | | | | |
|---|---|---|---|---|---|---|---|
| ENSDART00000064403 | nptnb | 0.050632772 | -0.282191976 | -0.43563203 | -0.351497687 | -0.147485108 | downregulated during regeneration - early |
| ENSDART00000064866 | prkab1a | 0.004813371 | -0.370484412 | -0.353103565 | -0.349388814 | -0.328698623 | downregulated during regeneration - early |
| ENSDART00000065361 | etv5b | 0.019698661 | -0.645686026 | -0.347502384 | -0.293939032 | -0.317696675 | downregulated during regeneration - early |
| ENSDART00000065366 | st6gal1 | 0.005057779 | -0.520593433 | -0.711298232 | -0.60540202 | -0.072634403 | downregulated during regeneration - early |
| ENSDART00000065507 | plppr2b | 0.00275056 | -0.22717931 | -0.353326082 | -0.20008884 | 0.028481334 | downregulated during regeneration - early |
| ENSDART00000066269 | arl4d | 0.002710576 | -0.431926082 | -0.429953919 | -0.294676093 | -0.224713874 | downregulated during regeneration - early |
| ENSDART00000066290 | UTS2R | 0.02114031 | -0.720663714 | -0.982454784 | -0.512077429 | -0.545080013 | downregulated during regeneration - early |
| ENSDART00000066386 | shisa9a | 0.017384242 | -0.462438855 | -0.347473048 | -0.310604934 | -0.078066291 | downregulated during regeneration - early |
| ENSDART00000066896 | syt1a | 0.019811481 | -0.255336633 | -0.349305824 | -0.330024338 | -0.178637525 | downregulated during regeneration - early |
| ENSDART00000067190 | tspan9b | 0.000694227 | -0.605960388 | -0.476716761 | -0.251844268 | -0.266924971 | downregulated during regeneration - early |
| ENSDART00000067211 | gpr37l1b | 0.009682631 | -0.390917116 | -0.36533905 | -0.206436073 | -0.113142613 | downregulated during regeneration - early |
| ENSDART00000067733 | zgc:77838 | 0.041639821 | -0.192172111 | -0.379046905 | -0.170230593 | -0.131538824 | downregulated during regeneration - early |
| ENSDART00000067741 | cacng6b | 0.039457626 | -2.498204907 | -1.377375045 | -1.107680653 | -1.191335449 | downregulated during regeneration - early |
| ENSDART00000067762 | MYO1D | 0.006648495 | -0.207118869 | -0.403579734 | -0.286614596 | -0.314212256 | downregulated during regeneration - early |
| ENSDART00000073588 | kcnj11 | 0.051199639 | -0.465815764 | -0.671487298 | -0.425150531 | -0.293255153 | downregulated during regeneration - early |
| ENSDART00000073617 | opn4xa | 0.013394015 | -0.377358091 | -0.474333233 | -0.655013246 | -0.269116892 | downregulated during regeneration - early |
| ENSDART00000074685 | glrbb | 0.001927288 | -0.387255204 | -0.421576892 | -0.370807969 | -0.094075713 | downregulated during regeneration - early |
| ENSDART00000074838 | kcnk3b | 0.00778428 | -0.808453975 | -0.995140212 | -0.30053552 | -0.689829672 | downregulated during regeneration - early |
| ENSDART00000074936 | gabrr2a | 0.040026502 | -0.375623148 | -0.534637844 | -0.431940406 | -0.359950744 | downregulated during regeneration - early |
| ENSDART00000075510 | ngb | 0.004883865 | -0.534714702 | -0.310443771 | -0.511019459 | -0.235764625 | downregulated during regeneration - early |
| ENSDART00000075935 | vtnb | 0.011844748 | -0.377853158 | -0.359520721 | -0.399371834 | -0.233041975 | downregulated during regeneration - early |
| ENSDART00000075940 | mtnr1ba | 0.03016157 | -0.423445838 | -1.139676296 | -0.754844062 | -0.657118397 | downregulated during regeneration - early |
| ENSDART00000075974 | ism2b | 0.049434429 | -0.752954546 | -0.582959852 | -0.548810475 | -0.377147389 | downregulated during regeneration - early |
| ENSDART00000075993 | crtc1b | 0.069455159 | -0.280354395 | -0.577500657 | -0.194615785 | -0.207111517 | downregulated during regeneration - early |
| ENSDART00000076215 | CR735102.1 | 0.0256797 | -0.360410997 | -0.514821634 | -0.137026978 | -0.257442941 | downregulated during regeneration - early |
| ENSDART00000077664 | atp2b1a | 0.015884799 | -0.283300071 | -0.323367416 | -0.218529441 | -0.07251725 | downregulated during regeneration - early |
| ENSDART00000077805 | gria2a | 0.012308945 | -0.63140941 | -0.839709321 | -0.362303249 | -0.053629448 | downregulated during regeneration - early |
| ENSDART00000077809 | cyp26c1 | 0.003805801 | -0.796873586 | -0.763455549 | -0.70044496 | -0.582141171 | downregulated during regeneration - early |
| ENSDART00000077823 | lrit3a | 0.035313407 | -0.481897882 | -0.48516269 | -0.481656788 | -0.455604814 | downregulated during regeneration - early |
| ENSDART00000078249 | kcnc3a | 0.004480141 | -0.908848026 | -1.288714191 | -0.644595688 | 0.010477706 | downregulated during regeneration - early |
| ENSDART00000078795 | ahcyl1 | 0.001402519 | -0.360916144 | -0.460734164 | -0.414619806 | -0.201451999 | downregulated during regeneration - early |
| ENSDART00000079035 | rap1gap | 0.009758715 | -0.57187807 | -0.701331328 | -0.299987324 | -0.289210087 | downregulated during regeneration - early |
| ENSDART00000079341 | plch1 | 0.009734608 | -0.457888719 | -0.43639001 | -0.345543016 | -0.155469708 | downregulated during regeneration - early |
| ENSDART00000079454 | vamp2 | 0.027826876 | -0.197978353 | -0.366012835 | -0.326357547 | -0.1060827 | downregulated during regeneration - early |
| ENSDART00000079597 | vps36 | 0.031565534 | -0.205754956 | -0.373190829 | -0.317459289 | -0.134792851 | downregulated during regeneration - early |
| ENSDART00000080430 | gfra2b | 9.16648E-05 | -0.62088032 | -0.576945607 | -0.392707521 | -0.144958503 | downregulated during regeneration - early |
| ENSDART00000081129 | cdk15 | 4.91848E-05 | -0.731126626 | -0.682923022 | -0.544281357 | -0.253398129 | downregulated during regeneration - early |
| ENSDART00000081797 | sash1b | 0.031235096 | -0.368433045 | -0.365106826 | -0.357216383 | -0.178826864 | downregulated during regeneration - early |
| ENSDART00000081926 | CU856539.1 | 0.01098705 | -0.250902601 | -0.402416425 | -0.31567263 | -0.091643419 | downregulated during regeneration - early |
| ENSDART00000081978 | KCNJ6 | 0.003763847 | -0.364130454 | -0.389723131 | -0.355694518 | -0.126078935 | downregulated during regeneration - early |
| ENSDART00000081990 | strip2 | 0.005385838 | -0.530994895 | -0.582233551 | -0.263527962 | -0.22210065 | downregulated during regeneration - early |
| ENSDART00000082012 | gsk3aa | 0.056003609 | -0.41333003 | -0.370003942 | -0.288947953 | -0.203102626 | downregulated during regeneration - early |
| ENSDART00000082142 | EFEMP1 (1 of many) | 0.081611118 | -1.76159347 | -0.849874708 | -0.584181957 | -1.080912576 | downregulated during regeneration - early |
| ENSDART00000082438 | dlgap2a | 0.01013158 | -0.366533513 | -0.540400426 | -0.257659814 | -0.037294062 | downregulated during regeneration - early |
| ENSDART00000082604 | galnt18b | 0.073840455 | -0.214631898 | -0.351577935 | -0.246678211 | -0.218706176 | downregulated during regeneration - early |
| ENSDART00000083100 | NA | 0.001483339 | -0.514905306 | -0.64333228 | -0.306814422 | -0.178642671 | downregulated during regeneration - early |
| ENSDART00000083407 | b4galnt4a | 0.056841831 | -0.141610106 | -0.312267637 | -0.129787335 | -0.08625756 | downregulated during regeneration - early |
| ENSDART00000083416 | gabrd | 0.001981601 | -0.516993342 | -0.41812722 | -0.415766575 | -0.292897654 | downregulated during regeneration - early |
| ENSDART00000083427 | slc25a29 | 0.004497123 | -0.338331576 | -0.636979798 | -0.361185623 | -0.191915619 | downregulated during regeneration - early |
| ENSDART00000083453 | slc32a1 | 0.006674249 | -0.393731273 | -0.378330473 | -0.304655077 | -0.16825712 | downregulated during regeneration - early |
| ENSDART00000083788 | CU633832.1 | 0.021355398 | -0.745373175 | -0.699527481 | -0.49975457 | -0.170614876 | downregulated during regeneration - early |
| ENSDART00000084024 | sv2c | 0.003671994 | -0.354928985 | -0.65582646 | -0.477522685 | -0.220826118 | downregulated during regeneration - early |
| ENSDART00000084381 | sybu | 0.003015186 | -0.763177411 | -0.608942869 | -0.49552138 | -0.188366829 | downregulated during regeneration - early |
| ENSDART00000084530 | coro2ba | 0.034451982 | -0.558160234 | -0.686474796 | -0.264666518 | -0.204619183 | downregulated during regeneration - early |
| ENSDART00000084803 | asic2 | 0.006230689 | -0.588195088 | -0.477145292 | -0.317477641 | -0.106481463 | downregulated during regeneration - early |
| ENSDART00000085121 | sdk2b | 0.030596624 | -0.222566721 | -0.365719278 | -0.147657392 | -0.012041014 | downregulated during regeneration - early |
| ENSDART00000085438 | rps6ka5 | 0.005012104 | -0.245466951 | -0.387017016 | -0.309352273 | -0.119154274 | downregulated during regeneration - early |
| ENSDART00000085522 | hspb6 | 0.048389146 | -0.63697673 | -0.382982575 | -0.524822831 | -0.392317705 | downregulated during regeneration - early |
| ENSDART00000086117 | kcnab2b | 0.011382754 | -0.362026581 | -0.574025944 | -0.45188395 | -0.145196155 | downregulated during regeneration - early |
| ENSDART00000086176 | nckap1 | 0.036431263 | -0.438983658 | -0.533344195 | -0.439721489 | -0.241138066 | downregulated during regeneration - early |
| ENSDART00000086181 | cabp7b | 0.018426524 | -0.513778995 | -0.579329622 | -0.46209057 | -0.104843884 | downregulated during regeneration - early |
| ENSDART00000087196 | zgc:153240 | 0.004497123 | -0.381380177 | -0.392146036 | -0.277166885 | -0.130863015 | downregulated during regeneration - early |
| ENSDART00000087565 | eva1a | 0.005375255 | -0.378976356 | -0.384921525 | -0.250389955 | -0.168669364 | downregulated during regeneration - early |
| ENSDART00000087654 | adcy6a | 0.006256994 | -0.242190723 | -0.49624198 | -0.145481294 | -0.053244608 | downregulated during regeneration - early |
| ENSDART00000088146 | ensab | 0.029418485 | -0.270112525 | -0.281607362 | -0.344659286 | -0.084313447 | downregulated during regeneration - early |
| ENSDART00000088240 | sypb | 0.022549623 | -0.232689126 | -0.445901517 | -0.375392134 | -0.149058325 | downregulated during regeneration - early |
| ENSDART00000088270 | yjefn3 | 0.006522424 | -0.490178998 | -0.350279664 | -0.399921095 | -0.213170988 | downregulated during regeneration - early |
| ENSDART00000088881 | git2a | 0.019698661 | -0.368814104 | -0.568589271 | -0.278482365 | -0.109167728 | downregulated during regeneration - early |
| ENSDART00000088908 | srgap1a | 0.001679054 | -0.173831935 | -0.499543132 | -0.183148374 | 0.021193005 | downregulated during regeneration - early |
| ENSDART00000089033 | lingo3a | 0.002706926 | -0.421566075 | -0.428623374 | -0.267183911 | -0.077777224 | downregulated during regeneration - early |
| ENSDART00000089126 | trhde.2 | 0.031143163 | -0.409075032 | -0.405058445 | -0.383617798 | -0.218104015 | downregulated during regeneration - early |
| ENSDART00000089748 | rorb | 0.030623213 | -0.330969943 | -0.539847879 | -0.425468683 | -0.138099942 | downregulated during regeneration - early |
| ENSDART00000089967 | cacna1bb | 0.093941409 | -0.202836628 | -0.60611593 | -0.329318476 | -0.282649478 | downregulated during regeneration - early |
| ENSDART00000090252 | atp2b3a | 0.003519185 | -0.293678426 | -0.382019678 | -0.276234519 | -0.034568861 | downregulated during regeneration - early |
| ENSDART00000090335 | hipk2 | 0.016509232 | -0.356266924 | -0.261702878 | -0.185362868 | -0.057591831 | downregulated during regeneration - early |

| | | | | | | | |
|---|---|---|---|---|---|---|---|
| ENSDART00000090580 | si:dkey-215k6.1 | 0.007297303 | -0.353341621 | -0.406611115 | -0.330167398 | -0.116132596 | downregulated during regeneration - early |
| ENSDART00000090669 | pleca | 0.078237713 | -0.254130273 | -0.426440212 | -0.236918819 | -0.12154508 | downregulated during regeneration - early |
| ENSDART00000091151 | nell2b | 0.000442465 | -0.408104365 | -0.533629775 | -0.525750199 | -0.236120482 | downregulated during regeneration - early |
| ENSDART00000091271 | prkg2l | 0.030415718 | -0.404320096 | -0.412762377 | -0.544649422 | -0.030408858 | downregulated during regeneration - early |
| ENSDART00000091416 | cntn3a.1 | 0.007676315 | -0.478191534 | -0.24621663 | -0.434975173 | -0.073764559 | downregulated during regeneration - early |
| ENSDART00000091472 | kcnv2b | 0.007620644 | -0.549604515 | -0.416623158 | -0.289225353 | -0.099467789 | downregulated during regeneration - early |
| ENSDART00000091532 | ndnf | 0.009217803 | -0.379105533 | -0.357674977 | -0.332342823 | -0.25490154 | downregulated during regeneration - early |
| ENSDART00000091612 | dab2ipa | 0.015157115 | -0.32378487 | -0.312766229 | -0.160534626 | -0.215968275 | downregulated during regeneration - early |
| ENSDART00000091683 | alkbh5 | 0.020455943 | -0.400075016 | -0.448742364 | -0.161106699 | -0.178285375 | downregulated during regeneration - early |
| ENSDART00000092013 | tmtc1 | 0.011155618 | -0.379958165 | -0.299353758 | -0.293525457 | -0.123093715 | downregulated during regeneration - early |
| ENSDART00000092051 | CABZ01081780.1 | 0.001500572 | -0.497076796 | -0.683547714 | -0.447049069 | -0.155316145 | downregulated during regeneration - early |
| ENSDART00000092183 | lrrc3b | 0.025950704 | -0.469734111 | -0.850621377 | -0.496353665 | -0.22903801 | downregulated during regeneration - early |
| ENSDART00000092356 | neto1 | 0.016069216 | -0.404606374 | -0.302098716 | -0.383296194 | -0.272667962 | downregulated during regeneration - early |
| ENSDART00000092493 | ptprt | 0.007621679 | -0.175495642 | -0.444255461 | -0.308001112 | -0.058909837 | downregulated during regeneration - early |
| ENSDART00000092647 | cers1 | 0.035286858 | -0.35221666 | -0.292832737 | -0.390981877 | -0.135220693 | downregulated during regeneration - early |
| ENSDART00000092691 | pea15 | 0.000272535 | -0.578361557 | -0.742039167 | -0.342285902 | -0.180997926 | downregulated during regeneration - early |
| ENSDART00000093236 | TULP3 | 0.019409502 | -0.257622072 | -0.196815312 | -0.318097587 | -0.153320475 | downregulated during regeneration - early |
| ENSDART00000097685 | LINGO3 (1 of many) | 0.000592646 | -0.608919806 | -0.847171007 | -0.317394062 | -0.126526617 | downregulated during regeneration - early |
| ENSDART00000097770 | gc3 | 0.022797575 | -0.462816548 | -0.433977737 | -0.284956472 | -0.124147363 | downregulated during regeneration - early |
| ENSDART00000098424 | trib2 | 0.000168321 | -0.755876906 | -0.434835466 | -0.414057266 | -0.366344874 | downregulated during regeneration - early |
| ENSDART00000098639 | cntn5 | 0.018537916 | -0.259764888 | -0.493723855 | -0.530432667 | -0.172558087 | downregulated during regeneration - early |
| ENSDART00000098668 | abcc8b | 0.045086702 | -0.217525237 | -0.529877646 | -0.409188595 | -0.288785069 | downregulated during regeneration - early |
| ENSDART00000099202 | igsf11 | 0.016255169 | -0.183727476 | -0.31314564 | -0.202768681 | -0.036422441 | downregulated during regeneration - early |
| ENSDART00000099235 | rnf44 | 0.003130236 | -0.234703995 | -0.37643632 | -0.256248531 | 0.012942846 | downregulated during regeneration - early |
| ENSDART00000099869 | slc17a7b | 0.012556905 | -0.359089552 | -0.531552161 | -0.360552754 | 0.022409885 | downregulated during regeneration - early |
| ENSDART00000100074 | pbx3a | 0.034362521 | -0.314908702 | -0.751877287 | -0.359257757 | -0.077459645 | downregulated during regeneration - early |
| ENSDART00000100223 | zgc:91860 | 0.02057139 | -0.412727201 | -0.311203152 | -0.280203316 | -0.243367869 | downregulated during regeneration - early |
| ENSDART00000100290 | napbb | 0.006047456 | -0.403034353 | -0.312472409 | -0.341096083 | -0.127811116 | downregulated during regeneration - early |
| ENSDART00000100327 | nptx1l | 0.010245667 | -0.348659958 | -0.433920996 | -0.393251042 | -0.286427391 | downregulated during regeneration - early |
| ENSDART00000100639 | chrm4a | 0.001996892 | -0.541979173 | -0.464947812 | -0.239039654 | -0.216630506 | downregulated during regeneration - early |
| ENSDART00000100681 | ncam2 | 0.01695739 | -0.42007768 | -0.748629002 | -0.327699832 | -0.08364547 | downregulated during regeneration - early |
| ENSDART00000101231 | syt7b | 0.02142158 | -0.85705254 | -2.266705376 | -0.529986528 | 0.010282974 | downregulated during regeneration - early |
| ENSDART00000101948 | GJA9 (1 of many) | 0.063141225 | -0.638779519 | -0.432316335 | -0.423080448 | -0.46097286 | downregulated during regeneration - early |
| ENSDART00000102445 | clasp1a | 0.056020682 | -0.444419251 | -0.511465397 | -0.218813787 | -0.372606163 | downregulated during regeneration - early |
| ENSDART00000102461 | rgs8 | 0.000273326 | -0.536939066 | -0.575929959 | -0.463779418 | -0.057720257 | downregulated during regeneration - early |
| ENSDART00000102790 | glrba | 0.002147221 | -0.477875363 | -0.33299196 | -0.36632817 | -0.172063586 | downregulated during regeneration - early |
| ENSDART00000102881 | fam43b | 0.029887325 | -0.413239873 | -0.283572592 | -0.358392622 | -0.273570929 | downregulated during regeneration - early |
| ENSDART00000103532 | kcnh5a | 0.002023943 | -0.481841968 | -0.667651405 | -0.403869416 | -0.116032422 | downregulated during regeneration - early |
| ENSDART00000103549 | skib | 0.029718853 | -0.170611371 | -0.408422527 | -0.193614247 | -0.134954723 | downregulated during regeneration - early |
| ENSDART00000103640 | hey1 | 0.023203992 | -0.385659584 | -0.747117397 | -0.392306381 | -0.416692063 | downregulated during regeneration - early |
| ENSDART00000103646 | kcng2 | 0.000233744 | -0.538883844 | -0.646084515 | -0.39646557 | 0.008271529 | downregulated during regeneration - early |
| ENSDART00000103894 | cacng8b | 3.32082E-05 | -0.589374624 | -0.680390231 | -0.379441318 | -0.105621743 | downregulated during regeneration - early |
| ENSDART00000103980 | crip2 | 0.004627449 | -0.517107368 | -0.364195263 | -0.546778441 | -0.267901424 | downregulated during regeneration - early |
| ENSDART00000104008 | dtnbp1b | 0.006824859 | -0.424284075 | -0.457257716 | -0.424822836 | -0.199214695 | downregulated during regeneration - early |
| ENSDART00000104027 | rims3 | 0.000442465 | -0.345163808 | -0.70584123 | -0.314103014 | -0.01029762 | downregulated during regeneration - early |
| ENSDART00000104293 | pex5la | 8.79987E-05 | -0.678333589 | -0.710207201 | -0.577955484 | -0.480904778 | downregulated during regeneration - early |
| ENSDART00000104481 | slc17a7a | 0.009884056 | -0.380361502 | -0.377656856 | -0.259723773 | -0.186490981 | downregulated during regeneration - early |
| ENSDART00000104722 | cdk5r2a | 0.016942898 | -0.371112934 | -0.317321269 | -0.137896092 | -0.229143186 | downregulated during regeneration - early |
| ENSDART00000104828 | gtpbp2b | 0.002166891 | -0.421398738 | -0.385176207 | -0.273058228 | -0.083654585 | downregulated during regeneration - early |
| ENSDART00000105477 | lrrtm2 | 0.001866651 | -0.540265042 | -0.543880814 | -0.470126158 | -0.070951006 | downregulated during regeneration - early |
| ENSDART00000105866 | si:ch73-213k20.5 | 0.001847721 | -0.515754761 | -0.874847543 | -0.892891327 | -0.392973916 | downregulated during regeneration - early |
| ENSDART00000106096 | drd1a | 0.032267182 | -0.279465728 | -0.508847304 | -0.2349577 | -0.295766526 | downregulated during regeneration - early |
| ENSDART00000106186 | prex2 | 0.010817469 | -0.375401587 | -0.276106515 | -0.274390577 | -0.295066887 | downregulated during regeneration - early |
| ENSDART00000108507 | CSMD3 | 0.008588572 | -0.201722796 | -0.640032982 | -0.257154173 | 0.075603806 | downregulated during regeneration - early |
| ENSDART00000108574 | gramd1bb | 0.016102005 | -0.232214076 | -0.405839915 | -0.354417408 | -0.144413075 | downregulated during regeneration - early |
| ENSDART00000108814 | nrip2 | 0.024870229 | -0.213998541 | -0.302896779 | -0.276048404 | -0.173620142 | downregulated during regeneration - early |
| ENSDART00000108959 | RNF208 | 0.003573129 | -0.615021133 | -0.757500788 | -0.372077015 | -0.309696014 | downregulated during regeneration - early |
| ENSDART00000108990 | pex5lb | 0.015010849 | -0.323313621 | -0.539021494 | -0.34249137 | -0.198103063 | downregulated during regeneration - early |
| ENSDART00000109014 | GJC1 | 0.052919095 | -0.38880893 | -0.552238053 | -0.31502822 | -0.300328618 | downregulated during regeneration - early |
| ENSDART00000109044 | grm8a | 0.000415801 | -0.470031444 | -0.746629562 | -0.459544685 | -0.031467357 | downregulated during regeneration - early |
| ENSDART00000109224 | CCKBR (1 of many) | 0.012371829 | -0.56783642 | -0.51463677 | -0.498556224 | -0.286398879 | downregulated during regeneration - early |
| ENSDART00000109257 | RASGRF1 | 7.10385E-05 | -0.570969407 | -0.754386918 | -0.426488142 | -0.098297531 | downregulated during regeneration - early |
| ENSDART00000109308 | si:ch73-60h1.1 | 0.05270735 | -0.222101504 | -0.315217201 | -0.340339223 | -0.206722759 | downregulated during regeneration - early |
| ENSDART00000109420 | kcnip1b | 2.98725E-05 | -0.712749849 | -0.701262469 | -0.816924162 | -0.561177799 | downregulated during regeneration - early |
| ENSDART00000109486 | gprc5ba | 0.001396814 | -0.628238673 | -0.569388482 | -0.508289237 | -0.226450363 | downregulated during regeneration - early |
| ENSDART00000109537 | snphb | 0.000529346 | -0.466057723 | -0.636087511 | -0.318006337 | -0.040021342 | downregulated during regeneration - early |
| ENSDART00000109732 | amigo1 | 0.034874709 | -0.336374535 | -0.536025942 | -0.442802906 | -0.174075192 | downregulated during regeneration - early |
| ENSDART00000109990 | abhd15a | 0.035729768 | -0.466192052 | -0.318128979 | -0.316342009 | -0.301397533 | downregulated during regeneration - early |
| ENSDART00000110016 | ube2ql1 | 0.022726575 | -0.274241146 | -0.513839036 | -0.485900432 | -0.131799131 | downregulated during regeneration - early |
| ENSDART00000110080 | aatka | 0.025606413 | -0.15444172 | -0.531359629 | -0.399426039 | -0.066680489 | downregulated during regeneration - early |
| ENSDART00000110092 | mdga1 | 0.024397386 | -0.307586742 | -0.496736277 | -0.237358341 | -0.048338544 | downregulated during regeneration - early |
| ENSDART00000110383 | map7a | 0.000639961 | -0.540958501 | -0.669475763 | -0.490700501 | -0.102714716 | downregulated during regeneration - early |
| ENSDART00000110432 | kazna | 0.080551232 | -0.433352738 | -0.292283864 | -0.217890679 | -0.318143053 | downregulated during regeneration - early |
| ENSDART00000110447 | slitrk3b | 0.024397386 | -0.707318961 | -0.905108657 | -0.279729387 | -0.217159944 | downregulated during regeneration - early |
| ENSDART00000110547 | ANKRD34A | 0.001359425 | -0.459666228 | -0.579840169 | -0.409587984 | -0.250229823 | downregulated during regeneration - early |

| | | | | | | | |
|---|---|---|---|---|---|---|---|
| ENSDART00000110789 | esyt2b | 0.006322758 | -0.389945342 | -0.213760019 | -0.33183342 | -0.035634516 | downregulated during regeneration - early |
| ENSDART00000110821 | ttc19 | 0.010842 | -0.356526006 | -0.209564592 | -0.274076967 | -0.065932254 | downregulated during regeneration - early |
| ENSDART00000110854 | elk1 | 0.059586248 | -0.317422178 | -0.365267467 | -0.144808984 | -0.160403171 | downregulated during regeneration - early |
| ENSDART00000110879 | magi2b | 0.020773351 | -0.318025437 | -0.557603079 | -0.245924365 | -0.15150652 | downregulated during regeneration - early |
| ENSDART00000110974 | zdhhc12b | 0.011929276 | -0.321463982 | -0.336778233 | -0.359206693 | -0.209775575 | downregulated during regeneration - early |
| ENSDART00000110976 | col19a1 | 0.036381892 | -0.113310455 | -0.554338462 | -0.336015886 | -0.135833904 | downregulated during regeneration - early |
| ENSDART00000111055 | kcna4 | 0.004463264 | -0.237309303 | -0.448087799 | -0.436530246 | -0.275701844 | downregulated during regeneration - early |
| ENSDART00000111080 | adgrb3 | 0.02511369 | -0.692483265 | -0.604853517 | -0.34586076 | -0.181234915 | downregulated during regeneration - early |
| ENSDART00000111111 | nlgn4b | 0.015044537 | -0.558022607 | -0.771924705 | -0.300759795 | -0.167214796 | downregulated during regeneration - early |
| ENSDART00000111234 | gdpd5a | 0.025523567 | -0.267734676 | -0.323349922 | -0.30731905 | -0.184332435 | downregulated during regeneration - early |
| ENSDART00000111261 | cdh24b | 0.001978324 | -0.501530291 | -0.712397312 | -0.362295258 | -0.168930493 | downregulated during regeneration - early |
| ENSDART00000111323 | AMIGO3 (1 of many) | 0.023492759 | -0.274114526 | -0.487248865 | -0.174614423 | -0.21591611 | downregulated during regeneration - early |
| ENSDART00000111456 | rbm10 | 0.014371639 | -0.2435311 | -0.244375112 | -0.319750026 | -0.111931812 | downregulated during regeneration - early |
| ENSDART00000111509 | CABZ01072096.1 | 0.079186307 | -0.19271783 | -0.37213696 | -0.269302922 | -0.183488656 | downregulated during regeneration - early |
| ENSDART00000111571 | rap1gap2b | 0.001659006 | -0.543266541 | -0.656699315 | -0.515603594 | -0.222139298 | downregulated during regeneration - early |
| ENSDART00000111636 | DTX4 (1 of many) | 2.01848E-05 | -1.011696111 | -1.148570348 | -0.809918586 | -0.692193186 | downregulated during regeneration - early |
| ENSDART00000111656 | gpr78a | 0.000190025 | -0.727925057 | -0.963113505 | -0.708360358 | -0.204121228 | downregulated during regeneration - early |
| ENSDART00000111759 | zmp:0000000794 | 0.02020282 | -0.428447438 | -0.466968786 | -0.39807432 | -0.23111401 | downregulated during regeneration - early |
| ENSDART00000111823 | GRIK3 | 0.003171283 | -0.298812703 | -0.633195231 | -0.313411843 | -0.167732562 | downregulated during regeneration - early |
| ENSDART00000111841 | BX248501.1 | 0.00748053 | -0.587056195 | -1.220156186 | -0.870300177 | -0.776464497 | downregulated during regeneration - early |
| ENSDART00000112106 | fam155a | 0.010271025 | -0.397020883 | -0.419375873 | -0.258307362 | -0.051593808 | downregulated during regeneration - early |
| ENSDART00000112296 | si:ch211-152n14.4 | 0.042906616 | -3.416151322 | -1.061691573 | -0.948841593 | -2.14718796 | downregulated during regeneration - early |
| ENSDART00000112299 | gpr37a | 0.027970823 | -0.365323083 | -0.2884934 | -0.319868677 | -0.150652035 | downregulated during regeneration - early |
| ENSDART00000112301 | trhde.1 | 0.014638227 | -0.264986832 | -0.332160295 | -0.182975887 | -0.056935825 | downregulated during regeneration - early |
| ENSDART00000112441 | tlr7 | 0.00039492 | -0.49393668 | -0.654080017 | -0.57399465 | -0.158959649 | downregulated during regeneration - early |
| ENSDART00000112646 | CNNM1 | 0.000705938 | -0.428104026 | -0.544400388 | -0.592385966 | -0.267977747 | downregulated during regeneration - early |
| ENSDART00000112655 | nfasca | 0.000530673 | -0.501256457 | -0.441261682 | -0.289256026 | -0.116172993 | downregulated during regeneration - early |
| ENSDART00000112735 | fam19a1a | 0.000783264 | -0.689638571 | -0.889095001 | -0.657230812 | -0.483951597 | downregulated during regeneration - early |
| ENSDART00000112743 | nmbb | 0.023560515 | -0.255890888 | -1.177861162 | -0.767211345 | -0.814299265 | downregulated during regeneration - early |
| ENSDART00000113004 | CABZ01068356.1 | 0.025858283 | -0.41749624 | -0.503262627 | -0.597219984 | -0.132734415 | downregulated during regeneration - early |
| ENSDART00000113058 | gpsm2 | 0.017032668 | -0.158548143 | -0.583640034 | -0.403831438 | -0.106682634 | downregulated during regeneration - early |
| ENSDART00000113162 | pi4kaa | 0.016474673 | -0.145989753 | -0.358429295 | -0.235575632 | -0.007193852 | downregulated during regeneration - early |
| ENSDART00000113196 | si:ch211-157b11.14 | 0.011627674 | -0.299973945 | -0.39773998 | -0.291303207 | -0.069868393 | downregulated during regeneration - early |
| ENSDART00000113280 | FRMD5 (1 of many) | 0.004407873 | -0.439670364 | -0.372637017 | -0.247112316 | -0.218940122 | downregulated during regeneration - early |
| ENSDART00000113454 | chgb | 0.034869075 | -0.341137392 | -0.331982228 | -0.294276744 | -0.206705528 | downregulated during regeneration - early |
| ENSDART00000114083 | mcf2la | 0.003137832 | -0.277595318 | -0.392280599 | -0.224516886 | 0.025969597 | downregulated during regeneration - early |
| ENSDART00000114117 | necab1 | 0.017561897 | -0.468499392 | -0.405674747 | -0.46743135 | -0.222378141 | downregulated during regeneration - early |
| ENSDART00000114322 | rusc1 | 0.045517543 | -0.055969033 | -0.296840251 | -0.124744669 | -0.048299568 | downregulated during regeneration - early |
| ENSDART00000114442 | fndc5b | 0.001847721 | -0.498984664 | -0.658119927 | -0.45669978 | -0.18817484 | downregulated during regeneration - early |
| ENSDART00000114660 | opn8b | 0.000645779 | -0.615308217 | -0.695080703 | -0.641485332 | -0.112308149 | downregulated during regeneration - early |
| ENSDART00000114711 | sorcs1 | 0.000116387 | -0.736062495 | -0.837966489 | -0.594713883 | -0.208228471 | downregulated during regeneration - early |
| ENSDART00000114748 | vip | 0.00097129 | -0.375915262 | -0.449108971 | -0.514462327 | -0.417228927 | downregulated during regeneration - early |
| ENSDART00000114750 | gabrb2 | 0.00023578 | -0.56997023 | -0.610789713 | -0.475013901 | -0.067415538 | downregulated during regeneration - early |
| ENSDART00000115058 | pbxip1a | 0.00153043 | -0.476509078 | -0.442420659 | -0.523364891 | -0.310557938 | downregulated during regeneration - early |
| ENSDART00000115118 | tns1a | 5.24617E-05 | -0.409165389 | -0.800200161 | -0.457588882 | -0.074958883 | downregulated during regeneration - early |
| ENSDART00000115138 | rapgef4 | 0.007642045 | -0.335306767 | -0.551146979 | -0.593390846 | -0.144257686 | downregulated during regeneration - early |
| ENSDART00000115161 | reps2 | 0.075212446 | -0.084468471 | -0.327140094 | -0.166340372 | -0.127803804 | downregulated during regeneration - early |
| ENSDART00000115278 | cx47.1 | 0.000962944 | -0.445735222 | -0.554690047 | -0.399294096 | -0.241214414 | downregulated during regeneration - early |
| ENSDART00000115365 | rassf10a | 0.008044263 | -0.525931221 | -0.509933384 | -0.256813636 | -0.202027084 | downregulated during regeneration - early |
| ENSDART00000121457 | lbh | 0.006197253 | -0.556077202 | -0.419617758 | -0.340252817 | -0.354555634 | downregulated during regeneration - early |
| ENSDART00000121496 | gpr153 | 0.022523327 | -0.22286471 | -0.331603234 | -0.24476212 | -0.082544732 | downregulated during regeneration - early |
| ENSDART00000121503 | cplx3b | 0.010056242 | -0.427974445 | -0.393867199 | -0.470572672 | -0.340193244 | downregulated during regeneration - early |
| ENSDART00000121684 | nat8l | 0.000470359 | -0.712971152 | -0.758652511 | -0.423007329 | -0.36098028 | downregulated during regeneration - early |
| ENSDART00000121826 | bean1 | 0.000596344 | -0.425046714 | -0.600778843 | -0.413029577 | -0.180932047 | downregulated during regeneration - early |
| ENSDART00000121872 | mast3b | 0.013889645 | -0.277342703 | -0.556828614 | -0.261994844 | -0.168334337 | downregulated during regeneration - early |
| ENSDART00000122081 | sybu | 0.000261206 | -0.707212841 | -0.524661917 | -0.600466578 | -0.221891711 | downregulated during regeneration - early |
| ENSDART00000122429 | klf15 | 0.006222767 | -0.341190152 | -0.616050077 | -0.578895446 | -0.337948098 | downregulated during regeneration - early |
| ENSDART00000122454 | citb | 0.001304993 | -0.619522503 | -1.363556906 | -0.83905374 | -0.294671991 | downregulated during regeneration - early |
| ENSDART00000122654 | rnd1a | 0.045949377 | -0.834540827 | -0.662552714 | -0.915729348 | -0.640521788 | downregulated during regeneration - early |
| ENSDART00000122682 | cabp1b | 0.009667225 | -0.294760944 | -0.547217913 | -0.530251845 | -0.289128709 | downregulated during regeneration - early |
| ENSDART00000122747 | tmem30aa | 0.01889397 | -0.229416393 | -0.299526834 | -0.201671762 | -0.059032826 | downregulated during regeneration - early |
| ENSDART00000123359 | nlgn1 | 0.001676296 | -0.188766802 | -0.577714477 | -0.350080443 | 0.032932651 | downregulated during regeneration - early |
| ENSDART00000123534 | cacna1g | 0.007400402 | -0.255311686 | -0.53660486 | -0.469326401 | -0.279013734 | downregulated during regeneration - early |
| ENSDART00000123797 | pcp4b | 0.000180202 | -0.598256523 | -0.489822031 | -0.568163011 | -0.081816834 | downregulated during regeneration - early |
| ENSDART00000123844 | si:dkey-77f5.3 | 0.019467351 | -4.277427291 | -0.495308507 | -0.670171681 | -0.698447042 | downregulated during regeneration - early |
| ENSDART00000123950 | PRIMA1 | 0.001128218 | -0.578840887 | -0.634067967 | -0.550475282 | -0.159964437 | downregulated during regeneration - early |
| ENSDART00000124298 | klhl29 | 0.013613126 | -0.069964723 | -0.409007114 | -0.117673983 | 0.023652831 | downregulated during regeneration - early |
| ENSDART00000124480 | adgrl1a | 0.006467002 | -0.174945207 | -0.479872966 | -0.185768365 | 0.06038548 | downregulated during regeneration - early |
| ENSDART00000124582 | napbb | 0.006047456 | -0.403034353 | -0.312472409 | -0.341096083 | -0.127811116 | downregulated during regeneration - early |
| ENSDART00000124662 | rtn4a | 0.027810244 | -0.503868889 | -0.627866078 | -0.609450822 | -0.190620657 | downregulated during regeneration - early |
| ENSDART00000124708 | gabra6b | 0.028541039 | -0.347504052 | -0.445856563 | -0.426704654 | -0.236544239 | downregulated during regeneration - early |
| ENSDART00000124876 | VSTM2B | 0.000962944 | -0.453041699 | -0.513162057 | -0.416504873 | -0.137923272 | downregulated during regeneration - early |
| ENSDART00000125344 | skilb | 0.010360832 | -0.241964121 | -0.627384358 | -0.493621211 | -0.368838752 | downregulated during regeneration - early |
| ENSDART00000125531 | plppr5a | 0.001540499 | -0.485992962 | -0.468686144 | -0.487868254 | -0.311270154 | downregulated during regeneration - early |
| ENSDART00000125766 | camkvb | 0.000261206 | -0.518737667 | -0.560273381 | -0.364168693 | -0.104855004 | downregulated during regeneration - early |

| | | | | | | | |
|---|---|---|---|---|---|---|---|
| ENSDART00000125900 | hs3st3b1b | 0.005535018 | -0.2288214 | -0.294028167 | -0.23141286 | 0.023426889 | downregulated during regeneration - early |
| ENSDART00000126300 | shisa8b | 0.024559038 | -0.373408846 | -0.464508355 | -0.412634146 | -0.258826308 | downregulated during regeneration - early |
| ENSDART00000126314 | pcdh8 | 0.024963085 | -0.25826413 | -0.454008614 | -0.231067015 | -0.062706758 | downregulated during regeneration - early |
| ENSDART00000126559 | rap1gap2a | 0.025646401 | -0.313212246 | -0.347504992 | -0.446138134 | -0.231468434 | downregulated during regeneration - early |
| ENSDART00000126739 | tspan5a | 0.010049723 | -0.28156215 | -0.24229244 | -0.35515584 | -0.059588327 | downregulated during regeneration - early |
| ENSDART00000126845 | ldb1a | 0.021507957 | -0.250500011 | -0.460439315 | -0.246081535 | -0.230460175 | downregulated during regeneration - early |
| ENSDART00000126966 | esrrga | 0.017679749 | -0.248566713 | -0.265310961 | -0.321308408 | -0.070523791 | downregulated during regeneration - early |
| ENSDART00000127062 | egr3 | 0.052026135 | -1.907539966 | -1.048685041 | -1.405658043 | -1.087307214 | downregulated during regeneration - early |
| ENSDART00000127144 | cacna2d2a | 0.013891126 | -0.206435603 | -0.394190367 | -0.167540436 | 0.000520565 | downregulated during regeneration - early |
| ENSDART00000127167 | fam217b | 0.004391006 | -0.253255723 | -0.39768374 | -0.389011868 | -0.208058743 | downregulated during regeneration - early |
| ENSDART00000127219 | FAM155A (1 of many) | 0.037495039 | -0.247003441 | -0.575960705 | -0.390291652 | -0.269667553 | downregulated during regeneration - early |
| ENSDART00000127236 | syt13 | 7.44856E-05 | -0.555582206 | -0.558950845 | -0.738809627 | -0.549910893 | downregulated during regeneration - early |
| ENSDART00000127426 | ERBB4 (1 of many) | 0.006128742 | -0.100134914 | -0.701677799 | -0.360350675 | -0.397963746 | downregulated during regeneration - early |
| ENSDART00000127428 | peli3 | 0.049273759 | -0.233448958 | -0.50115077 | -0.383840384 | -0.13792159 | downregulated during regeneration - early |
| ENSDART00000127467 | pygmb | 0.000803251 | -0.413201582 | -0.432147367 | -0.560055907 | -0.134010249 | downregulated during regeneration - early |
| ENSDART00000127506 | si:ch73-380n15.2 | 0.004844187 | -0.654968439 | -0.466259045 | -0.336236181 | -0.320196074 | downregulated during regeneration - early |
| ENSDART00000127513 | znf385d | 0.079618695 | -0.506377127 | -0.828614957 | -0.387596815 | -0.346943547 | downregulated during regeneration - early |
| ENSDART00000128198 | scn12aa | 0.032530799 | -0.298887564 | -0.309751416 | -0.23551982 | -0.15404656 | downregulated during regeneration - early |
| ENSDART00000128223 | CHCHD5 | 0.048304569 | -0.405489857 | -0.5400056 | -0.416697607 | -0.210022955 | downregulated during regeneration - early |
| ENSDART00000128224 | FO904873.1 | 0.031307521 | -0.267293261 | -0.580808884 | -0.319669262 | -0.131136583 | downregulated during regeneration - early |
| ENSDART00000128807 | gpr22a | 0.000870998 | -0.50401992 | -0.32421881 | -0.337086094 | -0.059033497 | downregulated during regeneration - early |
| ENSDART00000128959 | zgc:113223 | 0.022614665 | -1.993126278 | -0.750526857 | -1.059101824 | -0.610323514 | downregulated during regeneration - early |
| ENSDART00000129058 | rheb | 0.002834268 | -0.430188972 | -0.312082399 | -0.301953935 | -0.04976624 | downregulated during regeneration - early |
| ENSDART00000129202 | DDX17 | 0.027985198 | -0.231356945 | -0.275266303 | -0.31249155 | -0.24318473 | downregulated during regeneration - early |
| ENSDART00000129498 | mef2d | 0.03679078 | -0.188451231 | -0.381445449 | -0.263610552 | -0.197973462 | downregulated during regeneration - early |
| ENSDART00000130215 | TMEM179 (1 of many) | 0.028527553 | -0.219747105 | -0.315705451 | -0.194562459 | -0.061125473 | downregulated during regeneration - early |
| ENSDART00000130393 | tlr7 | 0.010379183 | -0.687566787 | -0.954352742 | -0.823581986 | -0.270839916 | downregulated during regeneration - early |
| ENSDART00000130881 | gabrr2a | 0.025544321 | -0.366410023 | -0.492281459 | -0.409834 | -0.272101123 | downregulated during regeneration - early |
| ENSDART00000130891 | camk4 | 0.000560128 | -0.554565795 | -0.502410128 | -0.607057653 | -0.157488548 | downregulated during regeneration - early |
| ENSDART00000130977 | mtcl1 | 0.038066989 | -0.152016911 | -0.472594538 | -0.24239267 | -0.004366794 | downregulated during regeneration - early |
| ENSDART00000131134 | si:ch211-206a7.2 | 0.008848885 | -0.226522125 | -0.472315068 | -0.371394997 | -0.101330709 | downregulated during regeneration - early |
| ENSDART00000131204 | asap1b | 0.001304993 | -0.546891821 | -0.455253903 | -0.26913051 | -0.092257648 | downregulated during regeneration - early |
| ENSDART00000131323 | si:dkey-193c22.1 | 0.006648495 | -0.375872197 | -0.187390024 | -0.399004983 | -0.194977152 | downregulated during regeneration - early |
| ENSDART00000131361 | kcnip3b | 0.008595135 | -0.353450144 | -1.035812483 | -0.692091767 | -0.071599751 | downregulated during regeneration - early |
| ENSDART00000131478 | kcnab1a | 0.005028852 | -0.495628945 | -0.276495434 | -0.300290472 | -0.132892856 | downregulated during regeneration - early |
| ENSDART00000131506 | cdh6 | 0.022814815 | -0.336043184 | -0.313603925 | -0.315311457 | -0.136816058 | downregulated during regeneration - early |
| ENSDART00000131866 | trhra | 0.014686352 | -0.77311942 | -0.482590954 | -0.518111078 | -0.348001173 | downregulated during regeneration - early |
| ENSDART00000132387 | pvalb6 | 2.98725E-05 | -0.612280143 | -0.883743713 | -0.424014361 | -0.028285642 | downregulated during regeneration - early |
| ENSDART00000132411 | si:ch1073-184j22.2 | 0.001677265 | -0.765110378 | -1.0433591 | -0.590491476 | -0.153408732 | downregulated during regeneration - early |
| ENSDART00000132522 | syt13 | 0.015571917 | -0.475976654 | -0.458152818 | -0.514134367 | -0.475986362 | downregulated during regeneration - early |
| ENSDART00000132582 | map4l | 0.000829135 | -0.700979622 | -0.493129551 | -0.385069171 | -0.268022699 | downregulated during regeneration - early |
| ENSDART00000132767 | pxdc1a | 0.019460106 | -0.518773739 | -0.198850875 | -0.436310873 | -0.094674576 | downregulated during regeneration - early |
| ENSDART00000132829 | inadl | 0.020169482 | -0.343759067 | -0.387749769 | -0.321781319 | -0.246918999 | downregulated during regeneration - early |
| ENSDART00000132838 | ank1a | 7.52983E-05 | -1.15020902 | -1.58246422 | -0.789622662 | -0.12782961 | downregulated during regeneration - early |
| ENSDART00000132911 | vwc2l | 0.004885993 | -0.573360674 | -0.839884971 | -0.564408735 | 0.020047377 | downregulated during regeneration - early |
| ENSDART00000133028 | mus81 | 0.049533299 | -0.28108498 | -0.562830421 | -0.307741383 | -0.072240912 | downregulated during regeneration - early |
| ENSDART00000133099 | zgc:123105 | 0.003047993 | -0.457591599 | -0.29947281 | -0.397070368 | -0.177964487 | downregulated during regeneration - early |
| ENSDART00000133325 | nell2b | 0.000401619 | -0.450880659 | -0.507896343 | -0.516937003 | -0.181021434 | downregulated during regeneration - early |
| ENSDART00000133487 | fosb | 0.034512759 | -0.882572207 | -1.273781011 | -0.772506712 | -0.914864674 | downregulated during regeneration - early |
| ENSDART00000133644 | magi2a | 0.001489209 | -0.426844627 | -0.586214992 | -0.344780913 | 0.065236587 | downregulated during regeneration - early |
| ENSDART00000133701 | slc6a1a | 0.017494941 | -0.417718543 | -0.444992936 | -0.335539494 | -0.221449117 | downregulated during regeneration - early |
| ENSDART00000133735 | cacng7b | 0.004879285 | -0.52911361 | -0.351880161 | -0.311790544 | -0.261905914 | downregulated during regeneration - early |
| ENSDART00000133743 | camkk1a | 0.001676296 | -0.520213698 | -0.784879077 | -0.322164805 | -0.513359014 | downregulated during regeneration - early |
| ENSDART00000133802 | inpp5jb | 0.006314264 | -0.26851081 | -0.522364408 | -0.244805361 | -0.012462884 | downregulated during regeneration - early |
| ENSDART00000134053 | usp2a | 0.031101324 | -0.375529791 | -0.273036858 | -0.320691469 | -0.159380591 | downregulated during regeneration - early |
| ENSDART00000134186 | cplx4b | 0.027048639 | -0.392479495 | -0.543085477 | -0.51593407 | -0.267696813 | downregulated during regeneration - early |
| ENSDART00000134431 | pcbp3 | 0.001578304 | -0.497499314 | -0.525712825 | -0.323624866 | -0.140513929 | downregulated during regeneration - early |
| ENSDART00000134489 | htr2cl1 | 0.010885977 | -0.503209957 | -0.409297224 | -0.43743207 | -0.200430987 | downregulated during regeneration - early |
| ENSDART00000134518 | cpa6 | 0.014368587 | -0.382405278 | -0.418058887 | -0.443147645 | -0.210306107 | downregulated during regeneration - early |
| ENSDART00000134661 | gstcd | 0.067079 | -0.989748846 | -0.61234101 | -0.452964111 | -0.446444607 | downregulated during regeneration - early |
| ENSDART00000135206 | ppp6r2b | 0.014582204 | -0.281142732 | -0.364406934 | -0.331381396 | -0.106942863 | downregulated during regeneration - early |
| ENSDART00000135481 | si:ch211-67n3.9 | 0.025747062 | -0.264301646 | -0.482759797 | -0.356089399 | -0.148916978 | downregulated during regeneration - early |
| ENSDART00000135897 | tmeff2a | 0.000235661 | -0.507501095 | -0.570530323 | -0.389897465 | -0.071849112 | downregulated during regeneration - early |
| ENSDART00000135965 | plch1 | 0.00810207 | -0.574356968 | -0.567600235 | -0.421755163 | -0.154028312 | downregulated during regeneration - early |
| ENSDART00000136200 | cbln2b | 0.007124918 | -0.48916567 | -0.352468034 | -0.442788192 | -0.14816736 | downregulated during regeneration - early |
| ENSDART00000136255 | otofa | 0.000894529 | -0.67545742 | -0.815938232 | -0.602392654 | -0.103624276 | downregulated during regeneration - early |
| ENSDART00000136390 | hcn2b | 0.012583499 | -0.288020893 | -0.351077892 | -0.312560003 | -0.049583355 | downregulated during regeneration - early |
| ENSDART00000136497 | CR318624.1 | 0.022425099 | -0.290569983 | -0.906819744 | -0.441990313 | 0.029701559 | downregulated during regeneration - early |
| ENSDART00000136559 | ano8a | 0.000185329 | -0.477178876 | -0.572871399 | -0.340924619 | -0.093243478 | downregulated during regeneration - early |
| ENSDART00000136592 | ckbb | 0.021710741 | -0.985379774 | -1.268435214 | -0.989267529 | -0.749991465 | downregulated during regeneration - early |
| ENSDART00000136654 | grapb | 0.038386468 | -0.34841027 | -0.721289328 | -0.501261917 | -0.353877694 | downregulated during regeneration - early |
| ENSDART00000136726 | magi3b | 0.006204545 | -0.381646177 | -0.461208385 | -0.226303301 | -0.080158554 | downregulated during regeneration - early |
| ENSDART00000136733 | glra1 | 0.010376336 | -0.259893278 | -0.432961012 | -0.409765299 | -0.248913141 | downregulated during regeneration - early |
| ENSDART00000136789 | tjp2a | 0.004813371 | -0.247782376 | -0.511487885 | -0.433268095 | -0.195886379 | downregulated during regeneration - early |
| ENSDART00000136805 | rbm10 | 0.044578922 | -4.539752918 | -3.15582054 | -1.19980751 | -2.046490722 | downregulated during regeneration - early |

| | | | | | | | |
|---|---|---|---|---|---|---|---|
| ENSDART00000136853 | fam163b | 0.012342358 | -0.426911899 | -0.715978676 | -0.456698496 | -0.087825906 | downregulated during regeneration - early |
| ENSDART00000137021 | cabp4 | 0.005059311 | -0.36160439 | -0.600623701 | -0.526609212 | -0.179412794 | downregulated during regeneration - early |
| ENSDART00000137038 | asic2 | 0.002292785 | -0.373156822 | -0.72567768 | -0.446804249 | -0.152680828 | downregulated during regeneration - early |
| ENSDART00000137214 | gpm6bb | 0.003631812 | -0.410929898 | -0.351705205 | -0.352631557 | -0.244368267 | downregulated during regeneration - early |
| ENSDART00000137287 | pebp1 | 0.030268752 | -0.410574808 | -0.266170824 | -0.343257245 | -0.128567003 | downregulated during regeneration - early |
| ENSDART00000137293 | dia1b | 0.011346243 | -0.377972932 | -0.365130624 | -0.231479469 | -0.228625621 | downregulated during regeneration - early |
| ENSDART00000137424 | c1qtnf4 | 0.013968714 | -0.387372103 | -0.312978581 | -0.242907328 | -0.243606392 | downregulated during regeneration - early |
| ENSDART00000137443 | ANK1 (1 of many) | 0.009480138 | -0.657062014 | -0.810024547 | -0.456200062 | -0.254232075 | downregulated during regeneration - early |
| ENSDART00000137505 | vamp1 | 0.002192481 | -0.431791589 | -0.393372103 | -0.433651273 | -0.142134375 | downregulated during regeneration - early |
| ENSDART00000137648 | si:dkeyp-123h10.2 | 0.001203555 | -0.55097741 | -0.596976893 | -0.447594093 | -0.275058949 | downregulated during regeneration - early |
| ENSDART00000137680 | BX663503.1 | 0.042893923 | -0.232606929 | -0.407355209 | -0.255389687 | -0.079162056 | downregulated during regeneration - early |
| ENSDART00000137728 | st6gal1 | 0.00818074 | -0.677667407 | -0.699420985 | -0.736040173 | -0.083670402 | downregulated during regeneration - early |
| ENSDART00000137848 | palmda | 0.050492107 | -0.364561265 | -0.248835039 | -0.231952958 | -0.127331926 | downregulated during regeneration - early |
| ENSDART00000138139 | zgc:153157 | 0.006145665 | -0.590921601 | -0.369202687 | -0.339560998 | -0.377004198 | downregulated during regeneration - early |
| ENSDART00000138140 | SLC4A5 (1 of many) | 0.02741598 | -0.322967614 | -0.430634585 | -0.36463151 | -0.15311364 | downregulated during regeneration - early |
| ENSDART00000138185 | gad1b | 0.050170102 | -0.304744145 | -0.418178899 | -0.358818296 | -0.231950915 | downregulated during regeneration - early |
| ENSDART00000138477 | BX571945.1 | 0.007291871 | -0.597559877 | -0.845904193 | -0.62766334 | -0.501592231 | downregulated during regeneration - early |
| ENSDART00000138687 | ppp1r9a | 0.010965837 | -0.410087357 | -0.337226328 | -0.257566135 | -0.006942521 | downregulated during regeneration - early |
| ENSDART00000138733 | cdh12a | 0.010404332 | -0.402719414 | -0.467471363 | -0.247597318 | -0.111519505 | downregulated during regeneration - early |
| ENSDART00000138755 | dclk1a | 0.013281778 | -0.464513032 | -0.326162873 | -0.241704881 | -0.01162875 | downregulated during regeneration - early |
| ENSDART00000138820 | lrtm2a | 0.053649719 | -0.205456304 | -0.473378158 | -0.285055681 | -0.252340958 | downregulated during regeneration - early |
| ENSDART00000138822 | SSBP4 (1 of many) | 0.003854547 | -0.268549166 | -0.417362634 | -0.271649078 | -0.060566364 | downregulated during regeneration - early |
| ENSDART00000138890 | kcnd2 | 0.006198931 | -0.399063976 | -0.347742148 | -0.33423415 | -0.044907554 | downregulated during regeneration - early |
| ENSDART00000139033 | lrp1bb | 0.016464653 | -0.300129828 | -0.380241873 | -0.239929923 | -0.108887683 | downregulated during regeneration - early |
| ENSDART00000139191 | trpc1 | 0.008989451 | -0.495478264 | -0.440776327 | -0.289497882 | -0.094213915 | downregulated during regeneration - early |
| ENSDART00000139196 | kif1ab | 0.024103105 | -0.338072705 | -0.653527626 | -0.314444638 | -0.193117382 | downregulated during regeneration - early |
| ENSDART00000139560 | ppp3r1a | 0.023218149 | -0.205912354 | -0.312380575 | -0.194151002 | 0.010760735 | downregulated during regeneration - early |
| ENSDART00000139568 | nsmfa | 0.033226863 | -0.224590353 | -0.433156538 | -0.157442583 | -0.141070436 | downregulated during regeneration - early |
| ENSDART00000139569 | zgc:92658 | 5.24617E-05 | -0.993075003 | -0.842810673 | -0.747118722 | -0.361693255 | downregulated during regeneration - early |
| ENSDART00000139608 | upf3a | 0.003570588 | -0.436308546 | -0.475063889 | -0.33694709 | -0.215591369 | downregulated during regeneration - early |
| ENSDART00000139633 | pip4k2ab | 0.018433591 | -0.336773688 | -0.358059228 | -0.306910553 | -0.048531465 | downregulated during regeneration - early |
| ENSDART00000139937 | frmpd1a | 0.008579097 | -0.391315372 | -0.461157908 | -0.442405648 | -0.041073617 | downregulated during regeneration - early |
| ENSDART00000140028 | lrrc4.1 | 0.00194329 | -0.491138395 | -0.422849217 | -0.374271959 | -0.216352385 | downregulated during regeneration - early |
| ENSDART00000140220 | si:ch211-168k14.2 | 0.057257329 | -0.180662482 | -0.408562234 | -0.217555172 | -0.18329178 | downregulated during regeneration - early |
| ENSDART00000140632 | bcl2l13 | 0.00107971 | -0.49849375 | -0.354616708 | -0.335413941 | -0.223271321 | downregulated during regeneration - early |
| ENSDART00000140650 | ankrd6b | 0.004329931 | -0.31619699 | -0.471630097 | -0.512160686 | -0.029088129 | downregulated during regeneration - early |
| ENSDART00000140995 | mfsd2b | 0.018126285 | -0.430905256 | -0.255202968 | -0.49728314 | -0.134050367 | downregulated during regeneration - early |
| ENSDART00000141177 | slc24a4a | 8.45072E-05 | -0.524899894 | -0.691002096 | -0.394086808 | -0.038902918 | downregulated during regeneration - early |
| ENSDART00000141634 | grid2 | 0.003104562 | -0.260805209 | -0.581973695 | -0.28493372 | -0.055775802 | downregulated during regeneration - early |
| ENSDART00000142010 | ncam1b | 0.027293911 | -0.299560612 | -0.317330716 | -0.349579067 | -0.156694047 | downregulated during regeneration - early |
| ENSDART00000142013 | penka | 0.001690789 | -0.474299096 | -0.531588241 | -0.656476493 | -0.472875583 | downregulated during regeneration - early |
| ENSDART00000142129 | tacr2 | 0.0004696 | -0.694450519 | -0.993924228 | -0.826737427 | -0.21380073 | downregulated during regeneration - early |
| ENSDART00000142171 | syt6a | 0.011695536 | -0.424476082 | -0.649670345 | -0.435293945 | -0.206511692 | downregulated during regeneration - early |
| ENSDART00000142223 | hivep3b | 0.013948629 | -0.312228644 | -0.441058379 | -0.334712198 | -0.18438466 | downregulated during regeneration - early |
| ENSDART00000142397 | samd12 | 0.034654408 | -0.674160268 | -0.821942052 | -0.657095298 | -0.360701575 | downregulated during regeneration - early |
| ENSDART00000142413 | BX897741.1 | 0.02739836 | -0.396693848 | -0.590528792 | -0.411359985 | -0.422415067 | downregulated during regeneration - early |
| ENSDART00000142654 | BX324123.2 | 0.001948843 | -0.377689742 | -0.458973085 | -0.444383379 | -0.275709525 | downregulated during regeneration - early |
| ENSDART00000142748 | UNC13A | 0.007676315 | -0.222825276 | -0.520698317 | -0.335972829 | 0.134560543 | downregulated during regeneration - early |
| ENSDART00000142806 | stxbp5a | 0.013774152 | -0.250986358 | -0.382479123 | -0.333968332 | -0.143471828 | downregulated during regeneration - early |
| ENSDART00000142810 | gabra3 | 0.005380857 | -0.405359169 | -0.863749916 | -0.692169745 | -0.647424845 | downregulated during regeneration - early |
| ENSDART00000142837 | enpp5 | 0.001384619 | -0.408425134 | -0.376571863 | -0.363824939 | -0.214479077 | downregulated during regeneration - early |
| ENSDART00000142975 | si:ch1073-70f20.1 | 0.018752626 | -0.443514404 | -0.337920446 | -0.430300513 | -0.297428401 | downregulated during regeneration - early |
| ENSDART00000142976 | kcnip3a | 0.00059038 | -0.515340349 | -0.554488278 | -0.50826115 | -0.165828333 | downregulated during regeneration - early |
| ENSDART00000143382 | il1rapl2 | 0.002636211 | -0.355933029 | -0.71045348 | -0.38678508 | -0.068375826 | downregulated during regeneration - early |
| ENSDART00000143554 | ampd2b | 0.018018574 | -0.328956412 | -0.382404642 | -0.421715131 | -0.209511328 | downregulated during regeneration - early |
| ENSDART00000143582 | si:ch211-236p22.1 | 0.000480969 | -0.593497691 | -1.183006832 | -0.39659144 | -0.180756629 | downregulated during regeneration - early |
| ENSDART00000143703 | flrt3 | 0.013070438 | -0.315208699 | -0.338016477 | -0.216002726 | -0.160149445 | downregulated during regeneration - early |
| ENSDART00000143784 | doc2d | 0.00429824 | -0.282456786 | -0.449882775 | -0.341326778 | 0.036564808 | downregulated during regeneration - early |
| ENSDART00000143787 | tnr | 0.054368123 | -0.299938525 | -0.646994379 | -0.19048806 | -0.113023917 | downregulated during regeneration - early |
| ENSDART00000143867 | syngap1b | 0.009560651 | -0.339959547 | -0.514759454 | -0.488418852 | -0.182310837 | downregulated during regeneration - early |
| ENSDART00000143878 | arhgef9a | 0.001216477 | -0.465269949 | -0.707003765 | -0.492066921 | -0.306386806 | downregulated during regeneration - early |
| ENSDART00000143911 | si:dkey-105e17.1 | 0.001995188 | -0.529795865 | -0.492017338 | -0.300236892 | -0.145540976 | downregulated during regeneration - early |
| ENSDART00000143919 | gabbr1b | 0.013735417 | -0.121288245 | -0.415278076 | -0.279488234 | -0.175446101 | downregulated during regeneration - early |
| ENSDART00000144175 | ccdc136b | 0.006621714 | -0.408583735 | -0.440451575 | -0.345784282 | -0.125541648 | downregulated during regeneration - early |
| ENSDART00000144255 | magixb | 0.023143967 | -0.339150091 | -0.441125122 | -0.366876369 | -0.172380242 | downregulated during regeneration - early |
| ENSDART00000144297 | ubac2 | 0.017079861 | -0.266999312 | -0.798888734 | -0.707434326 | -0.564642121 | downregulated during regeneration - early |
| ENSDART00000144600 | bin1a | 0.005414718 | -0.647409952 | -0.602565457 | -0.5226216 | -0.498459302 | downregulated during regeneration - early |
| ENSDART00000144995 | MFSD3 | 0.030278232 | -0.419691404 | -0.388696945 | -0.281399212 | -0.247251894 | downregulated during regeneration - early |
| ENSDART00000145220 | arvcfa | 0.000248952 | -0.706915106 | -0.825791875 | -0.448746575 | -0.046016635 | downregulated during regeneration - early |
| ENSDART00000145226 | si:ch211-233h19.2 | 0.050170102 | -0.372287835 | -0.175787817 | -0.260726103 | -0.131614613 | downregulated during regeneration - early |
| ENSDART00000145342 | cbln1 | 0.014767146 | -0.285689309 | -0.291683977 | -0.350650667 | -0.126158592 | downregulated during regeneration - early |
| ENSDART00000145364 | sybu | 0.039389579 | -0.525383816 | -0.345930455 | -0.451355227 | -0.158271762 | downregulated during regeneration - early |
| ENSDART00000145387 | shank2 | 0.001007682 | -0.271198688 | -0.514736215 | -0.289482704 | -0.011228429 | downregulated during regeneration - early |
| ENSDART00000145413 | si:ch211-253p2.2 | 0.01662249 | -0.612329709 | -0.583952587 | -0.545467055 | -0.273784221 | downregulated during regeneration - early |
| ENSDART00000145615 | cacna1i | 0.003514334 | -0.38207214 | -0.804209146 | -0.404812647 | -0.085276411 | downregulated during regeneration - early |

| | | | | | | | |
|---|---|---|---|---|---|---|---|
| ENSDART00000145916 | lingo4a | 0.003535619 | -0.858684072 | -0.829176543 | -0.734684966 | -0.796722528 | downregulated during regeneration - early |
| ENSDART00000145979 | col19a1 | 0.019480943 | -0.158274269 | -0.337487778 | -0.291137211 | -0.179712512 | downregulated during regeneration - early |
| ENSDART00000146113 | arhgef9b | 0.000103226 | -0.6095368 | -0.587211831 | -0.441711189 | -0.048741645 | downregulated during regeneration - early |
| ENSDART00000146227 | slc24a3 | 0.000387884 | -0.627503774 | -0.581196675 | -0.447342274 | -0.279205796 | downregulated during regeneration - early |
| ENSDART00000146284 | kcnh3 | 0.00027551 | -0.880077839 | -1.089282019 | -0.771605014 | -0.433474862 | downregulated during regeneration - early |
| ENSDART00000146321 | fibcd1 | 0.012665885 | -0.340172452 | -0.594838596 | -0.369380512 | -0.071609744 | downregulated during regeneration - early |
| ENSDART00000146327 | sgk1 | 0.004289396 | -0.407553517 | -0.497547741 | -0.442772363 | -0.391494952 | downregulated during regeneration - early |
| ENSDART00000146380 | map6d1 | 0.003805801 | -0.622612519 | -0.482084178 | -0.312516702 | -0.212427733 | downregulated during regeneration - early |
| ENSDART00000146422 | samd12 | 0.010712161 | -0.651610173 | -1.082672704 | -0.694390946 | -0.535477884 | downregulated during regeneration - early |
| ENSDART00000146424 | bin1a | 0.007279599 | -0.478145779 | -0.444599263 | -0.2729609 | -0.137965745 | downregulated during regeneration - early |
| ENSDART00000146712 | ccdc85a | 0.014638227 | -0.396306457 | -0.805958759 | -0.536293039 | -0.356615708 | downregulated during regeneration - early |
| ENSDART00000147308 | amigo1 | 0.02740918 | -0.366406631 | -0.580638611 | -0.462226604 | -0.20619589 | downregulated during regeneration - early |
| ENSDART00000147473 | pcsk1 | 0.010143774 | -0.408472737 | -0.345735504 | -0.351470668 | -0.372482226 | downregulated during regeneration - early |
| ENSDART00000147658 | bhlhe22 | 0.01205445 | -0.302235414 | -0.31899187 | -0.305987775 | -0.102192149 | downregulated during regeneration - early |
| ENSDART00000147868 | vwc2 | 0.003299709 | -0.776498099 | -0.767193994 | -0.55435848 | -0.286994867 | downregulated during regeneration - early |
| ENSDART00000148073 | shisa7b | 0.034090908 | -0.212233425 | -0.392255987 | -0.298336106 | -0.168085028 | downregulated during regeneration - early |
| ENSDART00000148564 | si:ch211-236p22.1 | 0.00081324 | -0.699364761 | -0.998683731 | -0.566423622 | -0.06995253 | downregulated during regeneration - early |
| ENSDART00000148576 | pygmb | 0.002081425 | -0.380093922 | -0.425025096 | -0.447938233 | -0.092571282 | downregulated during regeneration - early |
| ENSDART00000148930 | lingo4b | 0.038693778 | -0.229730557 | -0.308881186 | -0.199125951 | -0.339156928 | downregulated during regeneration - early |
| ENSDART00000148971 | ctbp2a | 0.015781384 | -0.618298464 | -0.594645977 | -0.414478241 | -0.261581238 | downregulated during regeneration - early |
| ENSDART00000149079 | sagb | 0.045816167 | -0.468101765 | -0.230897218 | -0.419750319 | -0.182809331 | downregulated during regeneration - early |
| ENSDART00000149297 | slc20a2 | 0.01613952 | -0.247675699 | -0.319353289 | -0.309560982 | -0.222732695 | downregulated during regeneration - early |
| ENSDART00000149411 | atxn1b | 0.052229213 | -0.356760798 | -0.298446699 | -0.334747495 | -0.251003673 | downregulated during regeneration - early |
| ENSDART00000149608 | nexn | 0.041923827 | -0.437154987 | -0.289194529 | -0.346094746 | -0.307861675 | downregulated during regeneration - early |
| ENSDART00000149914 | aatka | 0.000313682 | -0.270082025 | -0.541170375 | -0.440872523 | -0.175726567 | downregulated during regeneration - early |
| ENSDART00000149993 | pygmb | 0.000820449 | -0.443639651 | -0.459929444 | -0.472379886 | -0.083989126 | downregulated during regeneration - early |
| ENSDART00000150228 | chata | 0.006385907 | -0.431975216 | -0.532525907 | -0.558690348 | -0.371717362 | downregulated during regeneration - early |
| ENSDART00000150353 | mdh1aa | 0.015618075 | -0.527537579 | -0.6816935 | -0.262947 | -0.041325795 | downregulated during regeneration - early |
| ENSDART00000151127 | thraa | 0.039890399 | -0.798583389 | -0.802364861 | -0.351487338 | -0.589241182 | downregulated during regeneration - early |
| ENSDART00000151200 | mmp17a | 0.001404149 | -0.370431866 | -1.068890772 | -0.706081815 | -0.179291377 | downregulated during regeneration - early |
| ENSDART00000151229 | kcnj5 | 0.032556649 | -0.403592435 | -0.680204447 | -0.664541848 | -0.399506515 | downregulated during regeneration - early |
| ENSDART00000151358 | rgl3a | 0.000209364 | -0.536133566 | -0.674550597 | -0.735844208 | -0.535493428 | downregulated during regeneration - early |
| ENSDART00000151423 | c1ql3b | 0.004627449 | -0.428342943 | -0.592751887 | -0.458841393 | -0.097254864 | downregulated during regeneration - early |
| ENSDART00000151567 | sgk494b | 0.013519377 | -0.371209064 | -0.282108451 | -0.446431854 | -0.117181288 | downregulated during regeneration - early |
| ENSDART00000151627 | si:ch73-362m14.2 | 0.002433479 | -1.029772234 | -1.504293907 | -1.467613461 | -0.562650292 | downregulated during regeneration - early |
| ENSDART00000152069 | chst2b | 0.01938191 | -0.329630849 | -0.401663244 | -0.360860216 | -0.363893389 | downregulated during regeneration - early |
| ENSDART00000152371 | usp2a | 0.034504752 | -0.286860533 | -0.506577502 | -0.337595735 | 0.001938619 | downregulated during regeneration - early |
| ENSDART00000152376 | CLSTN2 (1 of many) | 0.008176596 | -0.314646809 | -0.323760148 | -0.422303692 | -0.021838531 | downregulated during regeneration - early |
| ENSDART00000152409 | mpp3b | 0.050836368 | -0.399859226 | -0.270912664 | -0.49906829 | -0.114613724 | downregulated during regeneration - early |
| ENSDART00000152531 | chrna10a | 0.027468632 | -0.39848157 | -0.348304481 | -0.425468269 | -0.274269467 | downregulated during regeneration - early |
| ENSDART00000152556 | FAM13C | 0.014686352 | -0.458278075 | -0.375744323 | -0.341944162 | -0.232139437 | downregulated during regeneration - early |
| ENSDART00000152580 | nlgn4a | 0.013502603 | -0.383612645 | -0.391621152 | -0.368243921 | -0.24740854 | downregulated during regeneration - early |
| ENSDART00000152858 | rcan2 | 0.017079861 | -0.344037465 | -0.451039286 | -0.208970338 | -0.321499331 | downregulated during regeneration - early |
| ENSDART00000153167 | hlfb | 0.010966965 | -0.470839666 | -0.326611575 | -0.235752692 | -0.264047019 | downregulated during regeneration - early |
| ENSDART00000153187 | thrab | 0.001402519 | -0.362356661 | -0.551240408 | -0.403932333 | -0.192068079 | downregulated during regeneration - early |
| ENSDART00000153247 | si:ch211-120g10.1 | 0.002147221 | -0.520249419 | -0.414424265 | -0.454570577 | -0.107523144 | downregulated during regeneration - early |
| ENSDART00000153256 | cacna1g | 0.001006824 | -0.53708851 | -0.717133052 | -0.415346006 | -0.243050395 | downregulated during regeneration - early |
| ENSDART00000153391 | otofa | 0.002551958 | -0.790883428 | -1.357026899 | -0.581227909 | -0.190433713 | downregulated during regeneration - early |
| ENSDART00000153591 | si:dkey-56f14.7 | 0.023379519 | -0.35474859 | -0.1685547 | -0.219665386 | -0.050831688 | downregulated during regeneration - early |
| ENSDART00000153673 | kcnb1 | 0.00060684 | -0.632196562 | -0.836186932 | -0.481100002 | -0.109364665 | downregulated during regeneration - early |
| ENSDART00000153925 | fut9a | 0.002016009 | -0.379466906 | -0.312742341 | -0.333889875 | -0.027131911 | downregulated during regeneration - early |
| ENSDART00000153937 | prkd1 | 0.021615739 | -0.431650217 | -0.382419593 | -0.495097396 | -0.310742086 | downregulated during regeneration - early |
| ENSDART00000154132 | nptx2a | 0.03060143 | -0.534082272 | -0.549346998 | -0.399917367 | -0.21269489 | downregulated during regeneration - early |
| ENSDART00000154148 | camk1a | 9.45963E-05 | -0.513931344 | -0.665527159 | -0.499415187 | -0.020106333 | downregulated during regeneration - early |
| ENSDART00000154187 | fat3a | 0.00879177 | -0.16861177 | -0.487078475 | -0.285722941 | -0.100061338 | downregulated during regeneration - early |
| ENSDART00000154228 | sgip1a | 0.019808116 | -0.210460002 | -0.370824353 | -0.364544583 | -0.177769156 | downregulated during regeneration - early |
| ENSDART00000154238 | AL928908.3 | 0.098059417 | -0.284715118 | -0.465774511 | -0.246230328 | -0.209644064 | downregulated during regeneration - early |
| ENSDART00000154437 | dyrk1ab | 0.025500103 | -0.405599591 | -0.293606249 | -0.153631672 | -0.200586701 | downregulated during regeneration - early |
| ENSDART00000154539 | slc17a7b | 0.013310088 | -0.194590302 | -0.417792811 | -0.270877884 | 0.095165091 | downregulated during regeneration - early |
| ENSDART00000154679 | ypel2b | 0.000167073 | -0.575722509 | -0.671947843 | -0.397167717 | -0.03983803 | downregulated during regeneration - early |
| ENSDART00000154682 | CR847531.1 | 0.002649033 | -0.541034447 | -0.676553431 | -0.554824044 | -0.149530967 | downregulated during regeneration - early |
| ENSDART00000154732 | ptprt | 0.007520653 | -0.128635692 | -0.441586772 | -0.224155866 | -0.045463061 | downregulated during regeneration - early |
| ENSDART00000154760 | CU302436.3 | 0.000164423 | -0.931829111 | -0.859047685 | -0.631190961 | -0.184130931 | downregulated during regeneration - early |
| ENSDART00000154778 | map1ab | 0.016443396 | -0.607759134 | -0.711103734 | -0.339973576 | -0.303457691 | downregulated during regeneration - early |
| ENSDART00000154783 | BX511311.5 | 0.001907584 | -0.420893047 | -0.406058694 | -0.342057398 | -0.24848029 | downregulated during regeneration - early |
| ENSDART00000154916 | p4htm | 0.025523567 | -0.322717782 | -0.242652877 | -0.246255125 | -0.141036056 | downregulated during regeneration - early |
| ENSDART00000155108 | si:ch211-153l6.6 | 0.024999915 | -0.396073461 | -0.324628235 | -0.330118035 | -0.333690214 | downregulated during regeneration - early |
| ENSDART00000155229 | rxfp3.3b | 0.002524433 | -0.899836922 | -0.961252446 | -0.631060044 | -0.38629797 | downregulated during regeneration - early |
| ENSDART00000155243 | BX571762.2 | 0.000109677 | -0.58394173 | -0.66558831 | -0.494788274 | -0.139509269 | downregulated during regeneration - early |
| ENSDART00000155346 | tbc1d24 | 0.000529346 | -0.444989153 | -0.328605075 | -0.340554332 | -0.035933185 | downregulated during regeneration - early |
| ENSDART00000155480 | ptprnb | 0.001147964 | -0.501074747 | -0.734898939 | -0.55867421 | -0.198202123 | downregulated during regeneration - early |
| ENSDART00000155496 | gal3st2 | 0.005444477 | -0.347287801 | -0.358994611 | -0.502209455 | 0.011977936 | downregulated during regeneration - early |
| ENSDART00000155539 | CR786577.1 | 0.00335169 | -0.296712547 | -0.710798753 | -0.341045728 | 0.032686751 | downregulated during regeneration - early |
| ENSDART00000155786 | adgrl1a | 0.019467351 | -0.190147203 | -0.479741899 | -0.292479007 | 0.056973675 | downregulated during regeneration - early |
| ENSDART00000155821 | kcnt2 | 0.00366084 | -1.064051569 | -1.353278896 | -1.209879321 | -0.444718597 | downregulated during regeneration - early |

| | | | | | | | |
|---|---|---|---|---|---|---|---|
| ENSDART00000155853 | gprc5ba | 0.000668535 | -0.568480744 | -0.680176078 | -0.476445621 | -0.089942946 | downregulated during regeneration - early |
| ENSDART00000155865 | tmtops2a | 0.015325737 | -0.317973372 | -0.37404155 | -0.204729003 | -0.178609254 | downregulated during regeneration - early |
| ENSDART00000156019 | phlpp1 | 0.010699261 | -0.378155747 | -0.257656641 | -0.150573304 | -0.126584813 | downregulated during regeneration - early |
| ENSDART00000156280 | si:ch211-195b15.7 | 0.04283071 | -0.343611395 | -0.333351498 | -0.20091334 | -0.160599804 | downregulated during regeneration - early |
| ENSDART00000156340 | si:ch211-167b20.8 | 0.018211877 | -0.291865267 | -0.289158159 | -0.304691219 | -0.111151074 | downregulated during regeneration - early |
| ENSDART00000156351 | zfat | 0.027293911 | -0.386268069 | -0.362611932 | -0.387180541 | -0.301395116 | downregulated during regeneration - early |
| ENSDART00000156418 | itpk1b | 0.001814914 | -0.567310725 | -0.6808899 | -0.20043499 | -0.215676742 | downregulated during regeneration - early |
| ENSDART00000156447 | ap2a1 | 0.048872862 | -0.168026562 | -0.400633541 | -0.149036642 | -0.01932092 | downregulated during regeneration - early |
| ENSDART00000156473 | CR627483.1 | 0.003693464 | -0.446874417 | -0.944979531 | -0.551426346 | -0.288797786 | downregulated during regeneration - early |
| ENSDART00000156574 | si:ch211-270g19.5 | 0.033836445 | -0.160521866 | -0.305818991 | -0.352315858 | -0.223463705 | downregulated during regeneration - early |
| ENSDART00000156615 | amigo3 | 0.011421096 | -0.174881747 | -0.348495554 | -0.328942663 | 0.002458748 | downregulated during regeneration - early |
| ENSDART00000156741 | BX663503.2 | 0.000389264 | -0.41351136 | -0.477289266 | -0.358517546 | -0.050787842 | downregulated during regeneration - early |
| ENSDART00000156760 | grik4 | 0.048815993 | -0.347451056 | -0.634709172 | -0.373537369 | -0.211041818 | downregulated during regeneration - early |
| ENSDART00000156864 | CR376745.2 | 0.003637926 | -0.465037351 | -0.777375633 | -0.748299595 | -0.606506499 | downregulated during regeneration - early |
| ENSDART00000156877 | prss56 | 0.031769506 | -1.154037343 | -0.806544973 | -0.688753663 | -0.465249106 | downregulated during regeneration - early |
| ENSDART00000156923 | plekhd1 | 0.012263789 | -0.640317774 | -0.396125765 | -0.428766409 | -0.43244988 | downregulated during regeneration - early |
| ENSDART00000156928 | nrxn3a | 0.007397439 | 0.020925857 | -0.341808434 | -0.239366202 | 0.052782357 | downregulated during regeneration - early |
| ENSDART00000157043 | rtn4r | 0.022328328 | -0.358605066 | -0.3593825 | -0.41712234 | -0.136427457 | downregulated during regeneration - early |
| ENSDART00000157124 | CT033825.1 | 0.001443362 | -0.560424235 | -0.809708864 | -0.680777647 | -0.144372819 | downregulated during regeneration - early |
| ENSDART00000157129 | slc12a5a | 0.000109677 | -0.482400538 | -0.74419089 | -0.401518291 | -0.07447909 | downregulated during regeneration - early |
| ENSDART00000157195 | ncam2 | 0.001019802 | -0.590792178 | -0.719154845 | -0.40764277 | 0.002619526 | downregulated during regeneration - early |
| ENSDART00000157330 | necab3 | 0.001497317 | -0.311593638 | -0.396628677 | -0.274835563 | -0.023399638 | downregulated during regeneration - early |
| ENSDART00000157414 | plxna4 | 0.033833708 | -0.256858488 | -0.391015088 | -0.285048615 | -0.106219383 | downregulated during regeneration - early |
| ENSDART00000157535 | unc13c | 0.004469286 | -0.230182256 | -0.566319347 | -0.291866986 | -0.102048139 | downregulated during regeneration - early |
| ENSDART00000157659 | camta1b | 0.01608096 | -0.574265168 | -0.690891494 | -0.316064045 | -0.339000781 | downregulated during regeneration - early |
| ENSDART00000157852 | crtac1b | 0.042224659 | -0.401370563 | -0.59197436 | -0.358347223 | -0.176621049 | downregulated during regeneration - early |
| ENSDART00000157943 | TENM2 | 0.003686025 | -0.301363214 | -0.846881931 | -0.344978545 | 0.012452 | downregulated during regeneration - early |
| ENSDART00000158169 | cacna2d2a | 0.001267982 | -0.273970943 | -0.601526504 | -0.288906256 | -0.138856816 | downregulated during regeneration - early |
| ENSDART00000158204 | ntrk3a | 0.003654787 | -0.448230073 | -0.669954401 | -0.450935682 | -0.114414424 | downregulated during regeneration - early |
| ENSDART00000158259 | lrtm1 | 0.053954169 | -0.264679565 | -0.523836662 | -0.478235847 | -0.361630227 | downregulated during regeneration - early |
| ENSDART00000158294 | ppp2r2cb | 0.007536007 | -0.427479199 | -0.698226267 | -0.466339082 | -0.260646218 | downregulated during regeneration - early |
| ENSDART00000158358 | stk40 | 0.042731689 | -0.305579471 | -0.394234001 | -0.179882029 | -0.355392322 | downregulated during regeneration - early |
| ENSDART00000158515 | NA | 0.009800471 | -0.601694587 | -0.246037227 | -0.439227399 | -0.290329932 | downregulated during regeneration - early |
| ENSDART00000158540 | PRKCA (1 of many) | 0.021044962 | -0.338994629 | -0.305121271 | -0.237610526 | -0.160203254 | downregulated during regeneration - early |
| ENSDART00000158820 | golga7bb | 0.029287312 | -0.441549718 | -0.650879189 | -0.491036037 | -0.164497612 | downregulated during regeneration - early |
| ENSDART00000158919 | lpar1 | 0.053989613 | -0.789836287 | -0.638879508 | -0.710773837 | -0.465534721 | downregulated during regeneration - early |
| ENSDART00000159087 | drd2b | 0.000503033 | -0.654775306 | -0.766659414 | -0.766987138 | -0.309487495 | downregulated during regeneration - early |
| ENSDART00000159163 | c2cd2l | 0.008021107 | -0.400916824 | -0.28913261 | -0.285175907 | -0.160469096 | downregulated during regeneration - early |
| ENSDART00000159165 | b3galt2 | 0.001631005 | -0.421223634 | -0.395995108 | -0.436113716 | -0.390524412 | downregulated during regeneration - early |
| ENSDART00000159368 | myo19 | 0.038545701 | -0.061571264 | -0.562765185 | -0.263758975 | -0.061266965 | downregulated during regeneration - early |
| ENSDART00000159493 | baiap2a | 0.031365399 | -0.408759075 | -0.584635172 | -0.205977447 | -0.074404323 | downregulated during regeneration - early |
| ENSDART00000159523 | BX248501.1 | 0.045817107 | -0.802081931 | -1.281225474 | -1.178391388 | -0.841643515 | downregulated during regeneration - early |
| ENSDART00000159604 | mbd3b | 0.065285449 | -0.404570692 | -0.312781624 | -0.318828454 | -0.248466452 | downregulated during regeneration - early |
| ENSDART00000159608 | fstl5 | 0.043959869 | -0.296329015 | -0.522282128 | -0.291629941 | -0.144161607 | downregulated during regeneration - early |
| ENSDART00000159745 | PCP4L1 (1 of many) | 7.10385E-05 | -0.632453626 | -0.700162017 | -0.354659005 | -0.133179733 | downregulated during regeneration - early |
| ENSDART00000159769 | pam | 0.003805801 | -0.468861179 | -0.403502349 | -0.282541701 | -0.241769832 | downregulated during regeneration - early |
| ENSDART00000159916 | tns1a | 0.00347249 | -0.270694335 | -0.694531751 | -0.372872148 | -0.014128978 | downregulated during regeneration - early |
| ENSDART00000160236 | fras1 | 0.023812224 | -0.444181856 | -0.638830186 | -0.533612869 | -0.242512624 | downregulated during regeneration - early |
| ENSDART00000160328 | zgc:109949 | 0.017825781 | -0.362552375 | -0.659300829 | -0.471802707 | -0.430381288 | downregulated during regeneration - early |
| ENSDART00000160337 | si:ch73-299h12.8 | 0.056707482 | -0.912707831 | -0.414120081 | -0.53567861 | -0.405510308 | downregulated during regeneration - early |
| ENSDART00000160542 | cabp2a | 0.013233841 | -0.310654407 | -0.386778791 | -0.172317372 | -0.340396034 | downregulated during regeneration - early |
| ENSDART00000160555 | sema6e | 0.029524772 | -0.284106568 | -0.324928323 | -0.250029521 | -0.168860793 | downregulated during regeneration - early |
| ENSDART00000160652 | iqsec3a | 0.002605308 | -0.564945044 | -0.609250827 | -0.334399494 | -0.124185292 | downregulated during regeneration - early |
| ENSDART00000160927 | cers1 | 0.004232418 | -0.519571193 | -0.339170129 | -0.296459142 | -0.066758323 | downregulated during regeneration - early |
| ENSDART00000161121 | oxct1a | 0.011687043 | -0.18384202 | -0.521566935 | -0.230939432 | -0.260776637 | downregulated during regeneration - early |
| ENSDART00000161156 | kcnj9 | 0.000230274 | -0.647175508 | -0.597438502 | -0.398278727 | -0.23455974 | downregulated during regeneration - early |
| ENSDART00000161250 | grin1b | 0.009000055 | -0.242810387 | -0.349766254 | -0.246157691 | 0.067747546 | downregulated during regeneration - early |
| ENSDART00000161272 | dclk1a | 0.011155635 | -0.780154894 | -0.444639927 | -0.375433616 | -0.486681237 | downregulated during regeneration - early |
| ENSDART00000161297 | lrrn3b | 0.008563 | -0.369968613 | -0.470809526 | -0.36045804 | -0.046069156 | downregulated during regeneration - early |
| ENSDART00000161414 | meis2a | 0.012130485 | -0.352024462 | -0.259692709 | -0.258698057 | -0.07620107 | downregulated during regeneration - early |
| ENSDART00000161533 | gfra1b | 0.017199091 | -0.396684985 | -0.507239408 | -0.32341327 | -0.285757426 | downregulated during regeneration - early |
| ENSDART00000161698 | spock1 | 0.008495106 | -0.400537237 | -0.522818259 | -0.36383883 | -0.3494177 | downregulated during regeneration - early |
| ENSDART00000161708 | si:ch73-29l19.1 | 0.048892164 | -0.183985257 | -0.352166832 | -0.310452263 | -0.095145467 | downregulated during regeneration - early |
| ENSDART00000161735 | flrt1a | 0.026767073 | -0.150942569 | -0.355176624 | -0.261636355 | -0.097731299 | downregulated during regeneration - early |
| ENSDART00000161798 | si:ch211-188f17.1 | 0.022416821 | -0.190236489 | -0.379885925 | -0.177844783 | -0.0015826 | downregulated during regeneration - early |
| ENSDART00000161828 | si:rp71-1h20.9 | 0.000291847 | -1.646379844 | -1.190834107 | -1.032421214 | -1.471047696 | downregulated during regeneration - early |
| ENSDART00000161938 | si:dkeyp-9d4.3 | 0.040982425 | -0.230392301 | -0.461066854 | -0.432261332 | -0.276495229 | downregulated during regeneration - early |
| ENSDART00000162063 | ppifb | 0.003061973 | -0.192515925 | -0.861096392 | -0.286677305 | -0.512048401 | downregulated during regeneration - early |
| ENSDART00000162222 | SHC3 | 0.03131236 | -0.255397657 | -0.359792108 | -0.178043549 | -0.323533675 | downregulated during regeneration - early |
| ENSDART00000162228 | grip2a | 0.010667361 | -0.209580983 | -0.468643062 | -0.36247099 | -0.147880595 | downregulated during regeneration - early |
| ENSDART00000162331 | si:dkey-22o22.2 | 0.015731546 | -0.434953721 | -0.633123757 | -0.287732073 | -0.108064019 | downregulated during regeneration - early |
| ENSDART00000162387 | zbtb38 | 0.015925333 | -0.362042869 | -0.285747988 | -0.199364681 | -0.27814593 | downregulated during regeneration - early |
| ENSDART00000162403 | napgb | 0.030278232 | -0.42647327 | -0.557407706 | -0.383093342 | -0.343712014 | downregulated during regeneration - early |
| ENSDART00000162595 | camk2g1 | 0.032809386 | -0.045798893 | -0.36296086 | -0.180481912 | 0.032536235 | downregulated during regeneration - early |
| ENSDART00000162683 | trappc9 | 0.067505731 | -0.370182614 | -0.24071592 | -0.273499446 | -0.133452933 | downregulated during regeneration - early |

| | | | | | | | |
|---|---|---|---|---|---|---|---|
| ENSDART00000162711 | cnksr2b | 0.026706504 | -0.20949159 | -0.28214605 | -0.418907028 | -0.082150791 | downregulated during regeneration - early |
| ENSDART00000162732 | NA | 0.000536544 | -0.497798966 | -0.517625334 | -0.441909965 | -0.132483146 | downregulated during regeneration - early |
| ENSDART00000162804 | KCNT1 (1 of many) | 0.004678129 | -0.406408577 | -0.48233323 | -0.378259831 | 0.110845983 | downregulated during regeneration - early |
| ENSDART00000162857 | nr4a3 | 0.003361003 | -1.482016444 | -1.152680943 | -0.606031454 | -1.193862677 | downregulated during regeneration - early |
| ENSDART00000162886 | il1rapl1b | 0.006723268 | -0.535131005 | -0.60836138 | -0.536159373 | -0.086785386 | downregulated during regeneration - early |
| ENSDART00000162916 | si:ch211-177d9.1 | 0.017970224 | -0.561077771 | -0.777629972 | -0.458457407 | -0.043076344 | downregulated during regeneration - early |
| ENSDART00000163106 | cntnap5l | 0.001234581 | -0.230959418 | -0.661142608 | -0.469168415 | -0.076363144 | downregulated during regeneration - early |
| ENSDART00000163380 | si:dkey-22o22.2 | 0.002605308 | -0.500713087 | -0.482673389 | -0.426086075 | -0.212897339 | downregulated during regeneration - early |
| ENSDART00000163491 | reep6 | 0.023753852 | -1.146193953 | -0.500723418 | -0.964434532 | -0.007386228 | downregulated during regeneration - early |
| ENSDART00000163523 | sez6l | 0.009922627 | -0.314616095 | -0.257996779 | -0.399166894 | -0.159678617 | downregulated during regeneration - early |
| ENSDART00000163568 | kcnc1a | 0.003649921 | -0.672590813 | -0.97944076 | -0.78875743 | -0.1747006 | downregulated during regeneration - early |
| ENSDART00000163597 | CABZ01029366.1 | 0.009204947 | -0.380235128 | -0.253799187 | -0.219703335 | -0.156720283 | downregulated during regeneration - early |
| ENSDART00000163793 | slitrk6 | 0.012145748 | -0.404346501 | -0.384982132 | -0.24261489 | -0.144542921 | downregulated during regeneration - early |
| ENSDART00000163892 | ldha | 0.068646243 | -0.444310122 | -0.296598203 | -0.299673411 | -0.138453042 | downregulated during regeneration - early |
| ENSDART00000163952 | zgc:110045 | 0.056216274 | -0.36736834 | -0.55642828 | -0.201308362 | -0.286026212 | downregulated during regeneration - early |
| ENSDART00000164015 | zgc:66483 | 0.030464157 | -0.233929761 | -0.339147918 | -0.342271085 | -0.145997916 | downregulated during regeneration - early |
| ENSDART00000164129 | asic4b | 2.97728E-05 | -0.92148029 | -1.144921239 | -0.577321683 | -0.304835302 | downregulated during regeneration - early |
| ENSDART00000164506 | dlg1l | 0.011264952 | -0.401430092 | -0.615035177 | -0.528577315 | -0.282785588 | downregulated during regeneration - early |
| ENSDART00000164581 | galr2b | 0.012570369 | -0.842297369 | -0.875318855 | -0.561173438 | -0.495794792 | downregulated during regeneration - early |
| ENSDART00000164621 | ndrg4 | 0.023203992 | -0.368994826 | -0.243279912 | -0.209623349 | -0.144971204 | downregulated during regeneration - early |
| ENSDART00000164623 | ptn | 0.025195087 | -0.285740219 | -0.301123559 | -0.395585979 | -0.169574914 | downregulated during regeneration - early |
| ENSDART00000164693 | CT583672.1 | 0.003678598 | -0.40975231 | -0.465896033 | -0.295925202 | -0.161007007 | downregulated during regeneration - early |
| ENSDART00000164733 | 43358 | 0.015015416 | -0.353566412 | -0.328984592 | -0.353599969 | -0.269459941 | downregulated during regeneration - early |
| ENSDART00000164844 | rictora | 0.015981762 | -0.308713132 | -0.348726405 | -0.242009414 | -0.189747444 | downregulated during regeneration - early |
| ENSDART00000164890 | pxdc1a | 0.02057139 | -0.553855303 | -0.268569574 | -0.515208528 | -0.163588555 | downregulated during regeneration - early |
| ENSDART00000164979 | ARHGAP44 (1 of many) | 0.017944789 | -0.298736282 | -0.611440452 | -0.190082373 | -0.04950856 | downregulated during regeneration - early |
| ENSDART00000165115 | adcy3a | 0.026979418 | -0.345722559 | -0.440118918 | -0.380340005 | -0.11083624 | downregulated during regeneration - early |
| ENSDART00000165124 | si:ch73-213k20.5 | 0.021047643 | -0.366310901 | -0.428402723 | -0.397554725 | -0.148030091 | downregulated during regeneration - early |
| ENSDART00000165292 | nsmfb | 0.006359324 | -0.267190978 | -0.447188182 | -0.263868703 | -0.115307671 | downregulated during regeneration - early |
| ENSDART00000165370 | nxph2b | 0.001996892 | -0.350612818 | -0.344729721 | -0.406615862 | -0.05363573 | downregulated during regeneration - early |
| ENSDART00000165411 | NA | 0.000228491 | -0.599585032 | -0.699498667 | -0.398460986 | -0.09881633 | downregulated during regeneration - early |
| ENSDART00000165423 | abcc8b | 0.052957916 | -0.169557865 | -0.54863705 | -0.322687745 | -0.280461618 | downregulated during regeneration - early |
| ENSDART00000165448 | rims2b | 0.01901087 | -0.321612136 | -0.490793078 | -0.295423658 | 0.030315799 | downregulated during regeneration - early |
| ENSDART00000165743 | gabrg2 | 0.002854235 | -0.320626657 | -0.470320689 | -0.258833602 | 0.01857369 | downregulated during regeneration - early |
| ENSDART00000165898 | gbe1b | 0.008897545 | -0.311269447 | -0.395300505 | -0.303280984 | -0.17462698 | downregulated during regeneration - early |
| ENSDART00000165903 | slmapa | 0.006879176 | -5.974312545 | -3.688098031 | -4.583524263 | -0.943141088 | downregulated during regeneration - early |
| ENSDART00000165912 | si:ch73-380n15.2 | 0.000461528 | -0.521676191 | -0.515528635 | -0.335328887 | -0.345110245 | downregulated during regeneration - early |
| ENSDART00000165979 | sncgb | 0.004112749 | -0.522103986 | -0.516181722 | -0.379782599 | -0.140861461 | downregulated during regeneration - early |
| ENSDART00000166025 | irs2a | 0.03450478 | -0.31567938 | -0.330763757 | -0.333387068 | -0.269837586 | downregulated during regeneration - early |
| ENSDART00000166027 | trpc1 | 0.019215781 | -0.303725684 | -0.42864481 | -0.334157939 | -0.231516525 | downregulated during regeneration - early |
| ENSDART00000166042 | vipr2 | 0.00847573 | -0.341756427 | -0.277003119 | -0.379732317 | -0.038627099 | downregulated during regeneration - early |
| ENSDART00000166135 | zbtb47b | 0.004311817 | -0.402709337 | -0.409852132 | -0.474957603 | -0.394147628 | downregulated during regeneration - early |
| ENSDART00000166148 | gabra1 | 3.38347E-05 | -0.578656795 | -0.788703398 | -0.59715214 | -0.072809071 | downregulated during regeneration - early |
| ENSDART00000166308 | cib2 | 0.004624842 | -0.434731659 | -0.335262372 | -0.470437684 | -0.240772493 | downregulated during regeneration - early |
| ENSDART00000166531 | pcdh15b | 0.014638227 | -0.215971807 | -0.468745045 | -0.336826359 | 0.084344716 | downregulated during regeneration - early |
| ENSDART00000166681 | frrs1l | 0.005203275 | -0.4975582 | -0.505651734 | -0.29386319 | -0.136005256 | downregulated during regeneration - early |
| ENSDART00000166730 | slitrk4 | 0.026033323 | -0.433556166 | -0.537605748 | -0.487948345 | -0.301835783 | downregulated during regeneration - early |
| ENSDART00000166889 | nptna | 0.001225897 | -0.194507678 | -0.523176341 | -0.288117341 | 0.166715743 | downregulated during regeneration - early |
| ENSDART00000166957 | purba | 0.055560475 | -0.28509761 | -0.466147452 | -0.146018067 | -0.191067595 | downregulated during regeneration - early |
| ENSDART00000167052 | etv1 | 0.01072353 | -0.327639473 | -0.469002174 | -0.32895701 | -0.22464313 | downregulated during regeneration - early |
| ENSDART00000167164 | MPV17L | 0.002297766 | -0.408335699 | -0.461800821 | -0.169486828 | -0.082955483 | downregulated during regeneration - early |
| ENSDART00000167451 | NA | 0.03168371 | -0.147201966 | -0.622186255 | -0.321157937 | -0.196934609 | downregulated during regeneration - early |
| ENSDART00000167502 | CR855860.1 | 0.030485193 | -0.377799337 | -0.542912736 | -0.423594114 | -0.115674527 | downregulated during regeneration - early |
| ENSDART00000167664 | atxn1a | 0.004580336 | -0.182914727 | -0.487043114 | -0.17911118 | -0.100136551 | downregulated during regeneration - early |
| ENSDART00000167847 | eef1a1l2 | 0.028902271 | -0.649781637 | -0.501730111 | -0.557800044 | -0.257462283 | downregulated during regeneration - early |
| ENSDART00000168089 | cyp27a7 | 0.044240808 | -0.742372496 | -0.705345887 | -0.793489907 | -0.543448957 | downregulated during regeneration - early |
| ENSDART00000168310 | acap3b | 0.015781384 | -0.431660294 | -0.622927148 | -0.474200982 | -0.250587028 | downregulated during regeneration - early |
| ENSDART00000168518 | asic4b | 2.97728E-05 | -0.92148029 | -1.144921239 | -0.577321683 | -0.304835302 | downregulated during regeneration - early |
| ENSDART00000168534 | NPFFR2 (1 of many) | 0.012462623 | -0.514269049 | -0.40572501 | -0.533037042 | -0.336902595 | downregulated during regeneration - early |
| ENSDART00000168631 | cacna1hb | 0.008579097 | -0.565978595 | -0.722362792 | -0.435741841 | -0.029665329 | downregulated during regeneration - early |
| ENSDART00000168718 | chrm5a | 0.033210482 | -0.32077931 | -0.748869105 | -0.317792694 | -0.07500967 | downregulated during regeneration - early |
| ENSDART00000168754 | cacnb2a | 0.028142339 | -0.312716955 | -0.298709564 | -0.211334034 | -0.110938146 | downregulated during regeneration - early |
| ENSDART00000168821 | CDH22 | 0.019500385 | -0.242258311 | -0.624670054 | -0.303879761 | -0.070304381 | downregulated during regeneration - early |
| ENSDART00000168920 | nbeab | 0.028147746 | -0.251742312 | -0.363594185 | -0.174304201 | -0.037829821 | downregulated during regeneration - early |
| ENSDART00000169040 | si:ch211-235e9.8 | 0.001532808 | -0.436326332 | -0.357257361 | -0.23322888 | -0.084450729 | downregulated during regeneration - early |
| ENSDART00000169108 | kcnj12a | 0.013360877 | -0.480725794 | -0.616905664 | -0.509410819 | -0.355199716 | downregulated during regeneration - early |
| ENSDART00000169136 | CABZ01007222.1 | 0.002957541 | -0.363042867 | -0.477045243 | -0.379599521 | -0.116458215 | downregulated during regeneration - early |
| ENSDART00000169548 | rgs7b | 0.024602158 | -0.179117493 | -0.337814246 | -0.235695562 | -0.091038288 | downregulated during regeneration - early |
| ENSDART00000169609 | tefb | 0.032368681 | -0.156114581 | -0.555141888 | -0.301378906 | -0.24132076 | downregulated during regeneration - early |
| ENSDART00000169687 | dlg1 | 0.023714388 | -0.200796553 | -0.39742906 | -0.281934456 | -0.007509175 | downregulated during regeneration - early |
| ENSDART00000169746 | ampd2a | 0.000935288 | -0.446564291 | -0.523779729 | -0.426555345 | -0.117905151 | downregulated during regeneration - early |
| ENSDART00000169764 | syt1a | 0.009024763 | -0.535633412 | -0.482595993 | -0.403145231 | -0.029612735 | downregulated during regeneration - early |
| ENSDART00000169810 | stub1 | 0.029463211 | -0.352758303 | -0.24873956 | -0.212157701 | -0.100812458 | downregulated during regeneration - early |
| ENSDART00000170362 | CABZ01088490.1 | 0.001365289 | -0.602203241 | -0.75748097 | -0.591873288 | -0.034015136 | downregulated during regeneration - early |
| ENSDART00000170709 | st8sia1 | 0.007584916 | -0.358435626 | -0.336878724 | -0.21402261 | -0.093884674 | downregulated during regeneration - early |

| | | | | | | | |
|---|---|---|---|---|---|---|---|
| ENSDART00000170993 | afmid | 0.002016009 | -0.391958822 | -0.311185654 | -0.48659874 | -0.071120804 | downregulated during regeneration - early |
| ENSDART00000170998 | tnrc6c2 | 0.033092405 | -0.257719977 | -0.319066529 | -0.317919253 | -0.242876098 | downregulated during regeneration - early |
| ENSDART00000171014 | ptprfa | 0.004256148 | -0.231763582 | -0.479834972 | -0.30404021 | -0.129420632 | downregulated during regeneration - early |
| ENSDART00000171041 | tceb3 | 0.028527553 | -0.063048764 | -0.305499385 | -0.056078786 | -0.065276347 | downregulated during regeneration - early |
| ENSDART00000171182 | FO704914.1 | 0.000532548 | -0.372200999 | -0.770608409 | -0.45129487 | -0.014659406 | downregulated during regeneration - early |
| ENSDART00000171215 | slc6a3 | 0.001490782 | -0.667608441 | -0.754783262 | -0.446986408 | -0.343120625 | downregulated during regeneration - early |
| ENSDART00000171426 | pdzph1 | 0.00721984 | -0.416069913 | -0.411889039 | -0.457121209 | -0.221062124 | downregulated during regeneration - early |
| ENSDART00000171496 | CDK18 | 0.00320961 | -0.371495713 | -0.374286185 | -0.433391037 | -0.282164654 | downregulated during regeneration - early |
| ENSDART00000171523 | CABZ01089030.1 | 0.001040281 | -0.303576632 | -0.51118876 | -0.528168137 | -0.25789403 | downregulated during regeneration - early |
| ENSDART00000171683 | GABRG1 | 0.028772568 | -0.455034367 | -0.392373746 | -0.437520168 | -0.270886274 | downregulated during regeneration - early |
| ENSDART00000171696 | sergef | 0.00024376 | -0.83692277 | -0.920324583 | -0.711315475 | -0.040508735 | downregulated during regeneration - early |
| ENSDART00000171711 | gpsm1a | 0.009294776 | -0.376386863 | -0.294884117 | -0.277096957 | -0.079488929 | downregulated during regeneration - early |
| ENSDART00000171728 | FQ323156.1 | 0.000261206 | -0.3925045 | -0.548422409 | -0.578285827 | -0.21374206 | downregulated during regeneration - early |
| ENSDART00000171743 | sypa | 0.000648054 | -0.458751426 | -0.455626528 | -0.334063629 | -0.142465032 | downregulated during regeneration - early |
| ENSDART00000171749 | TMEM150A (1 of many) | 0.00159523 | -0.745537318 | -0.560172793 | -0.666392254 | -0.39318407 | downregulated during regeneration - early |
| ENSDART00000171762 | arhgef11 | 0.023596757 | -0.293734421 | -0.490220654 | -0.20280274 | 0.049910063 | downregulated during regeneration - early |
| ENSDART00000171777 | syt7b | 0.000155554 | -0.587310767 | -0.899783938 | -0.612333236 | -0.170082745 | downregulated during regeneration - early |
| ENSDART00000171868 | sgut1 | 0.013366886 | -0.438334002 | -0.252211506 | -0.468859348 | -0.232780566 | downregulated during regeneration - early |
| ENSDART00000171935 | brpf3a | 0.010653756 | -0.333420883 | -0.26685281 | -0.204490748 | -0.091216638 | downregulated during regeneration - early |
| ENSDART00000171948 | zgc:172282 | 0.000409858 | -0.447105187 | -0.64749442 | -0.36124568 | -0.081452365 | downregulated during regeneration - early |
| ENSDART00000172045 | zgc:73340 | 0.004702472 | -0.432558368 | -0.571519732 | -0.606453829 | -0.389736033 | downregulated during regeneration - early |
| ENSDART00000172128 | sez6l | 0.001036975 | -0.457515145 | -0.649880208 | -0.297003963 | -0.214224661 | downregulated during regeneration - early |
| ENSDART00000172149 | sh3rf2 | 0.000935288 | -0.359308533 | -0.521296357 | -0.501584275 | -0.200324498 | downregulated during regeneration - early |
| ENSDART00000172218 | nsmfb | 0.001185822 | -0.786748472 | -1.480880504 | -0.428989171 | -0.523180434 | downregulated during regeneration - early |
| ENSDART00000172267 | KCNV1 | 0.026772289 | -0.730291266 | -0.683938113 | -0.423486953 | -0.317970814 | downregulated during regeneration - early |
| ENSDART00000172310 | zbtb4 | 0.013310088 | -0.365422353 | -0.277101341 | -0.237436935 | -0.247791483 | downregulated during regeneration - early |
| ENSDART00000172329 | CABZ01084963.1 | 0.005531203 | -0.076477014 | -0.479017439 | -0.153366972 | -0.215592031 | downregulated during regeneration - early |
| ENSDART00000172336 | cabp2a | 0.007145684 | -0.40143904 | -0.491969769 | -0.23184904 | -0.50001549 | downregulated during regeneration - early |
| ENSDART00000172337 | rhot1a | 0.009491292 | -3.878236672 | -5.686158747 | -0.665991717 | -0.822715826 | downregulated during regeneration - early |
| ENSDART00000172373 | ompa | 0.040539001 | -0.333379791 | -0.33389964 | -0.383107757 | -0.229559723 | downregulated during regeneration - early |
| ENSDART00000172496 | add3a | 0.056985155 | -0.173242509 | -0.314301562 | -0.183893521 | -0.110146899 | downregulated during regeneration - early |
| ENSDART00000172661 | nrgnb | 0.0006387 | -0.441646104 | -0.659637875 | -0.590118296 | -0.403632798 | downregulated during regeneration - early |
| ENSDART00000172689 | drd4a | 0.00510387 | -0.269065132 | -0.433761644 | -0.356034882 | -0.090259185 | downregulated during regeneration - early |
| ENSDART00000172862 | kcnj14 | 0.044172739 | -0.439592228 | -0.323667837 | -0.482461949 | -0.197270237 | downregulated during regeneration - early |
| ENSDART00000173109 | nrtn | 0.012342358 | -0.527610722 | -0.443827628 | -0.46689713 | -0.285016979 | downregulated during regeneration - early |
| ENSDART00000173113 | si:ch211-129p13.1 | 0.001328724 | -0.464452233 | -0.696136912 | -0.478469763 | -0.206239945 | downregulated during regeneration - early |
| ENSDART00000173119 | pcdh11 | 0.000993187 | -0.634370446 | -0.732192902 | -0.531501202 | -0.25060062 | downregulated during regeneration - early |
| ENSDART00000173133 | utrn | 0.064203588 | -0.11206335 | -0.324597481 | -0.223274112 | -0.197393841 | downregulated during regeneration - early |
| ENSDART00000173134 | BX571952.1 | 0.000520606 | -0.551010684 | -0.633352911 | -0.460069257 | -0.004126181 | downregulated during regeneration - early |
| ENSDART00000173386 | cacna1da | 0.028175613 | -0.09786225 | -0.344688682 | -0.169985603 | 0.048564501 | downregulated during regeneration - early |
| ENSDART00000000069 | slc9a3r1a | 0.024869861 | -0.019828621 | -0.165096913 | -0.579587833 | -0.318451026 | downregulated during regeneration - mid |
| ENSDART00000000192 | ptpn4b | 0.005203881 | -0.253123991 | -0.355280061 | -0.416236309 | -0.192007 | downregulated during regeneration - mid |
| ENSDART00000002029 | fkbp8 | 0.030637152 | -0.081020923 | -0.107047698 | -0.329759128 | -0.094897987 | downregulated during regeneration - mid |
| ENSDART00000003170 | mid1ip1l | 0.009183287 | -0.359873821 | -0.329776238 | -0.564188118 | -0.321447922 | downregulated during regeneration - mid |
| ENSDART00000003335 | snx12 | 0.02794941 | -0.102084433 | -0.181980514 | -0.47779818 | -0.16493026 | downregulated during regeneration - mid |
| ENSDART00000003465 | gipc2 | 0.02229367 | -0.142551263 | -0.216969239 | -0.344609382 | -0.269869981 | downregulated during regeneration - mid |
| ENSDART00000003939 | syngr1a | 0.024987601 | 0.053450312 | -0.149997984 | -0.371497084 | -0.066149387 | downregulated during regeneration - mid |
| ENSDART00000004075 | uqcc2 | 0.023840313 | -0.167833305 | -0.222493804 | -0.340331706 | -0.224913678 | downregulated during regeneration - mid |
| ENSDART00000004241 | inhbaa | 0.001317626 | 0.208890493 | -0.32024378 | -0.370155553 | -0.220482209 | downregulated during regeneration - mid |
| ENSDART00000004420 | rab4a | 0.034580091 | -0.104588348 | -0.223085036 | -0.330744686 | -0.154422823 | downregulated during regeneration - mid |
| ENSDART00000005573 | tmem237b | 0.009950411 | 0.093826402 | -0.187405072 | -0.456194249 | -0.210768209 | downregulated during regeneration - mid |
| ENSDART00000005590 | churc1 | 0.018990086 | -0.030881068 | -0.24423848 | -0.319974325 | -0.284115834 | downregulated during regeneration - mid |
| ENSDART00000005733 | tma16 | 0.047982796 | -0.013223994 | -0.144098168 | -0.425909208 | -0.168461973 | downregulated during regeneration - mid |
| ENSDART00000005957 | lrit1a | 0.025420775 | -0.250152343 | -0.282879742 | -0.392873609 | -0.351289311 | downregulated during regeneration - mid |
| ENSDART00000006927 | use1 | 0.036974502 | -0.314698173 | -0.230168127 | -0.342013543 | -0.20782604 | downregulated during regeneration - mid |
| ENSDART00000007021 | atp6v1ba | 0.031629016 | -0.168038972 | -0.268033099 | -0.367112615 | -0.152327119 | downregulated during regeneration - mid |
| ENSDART00000008302 | insra | 0.048141934 | -0.066772744 | -0.178139649 | -0.319689592 | -0.098012672 | downregulated during regeneration - mid |
| ENSDART00000009740 | smad7 | 0.030913639 | -0.082405078 | -0.352637537 | -0.492507075 | -0.066549718 | downregulated during regeneration - mid |
| ENSDART00000010378 | myo3b | 0.013233841 | 0.121341422 | -0.396947151 | -0.677290933 | -0.232688387 | downregulated during regeneration - mid |
| ENSDART00000011149 | fam185a | 0.051285619 | -0.225870361 | -0.305876809 | -0.396323932 | -0.356813487 | downregulated during regeneration - mid |
| ENSDART00000011229 | sub1b | 0.026680132 | -0.082257398 | -0.179031151 | -0.405374566 | -0.271111897 | downregulated during regeneration - mid |
| ENSDART00000011317 | ntm | 0.027578317 | -0.235410973 | -0.452806913 | -0.580240928 | -0.252707536 | downregulated during regeneration - mid |
| ENSDART00000012229 | fkbp1b | 0.049049155 | -0.199888531 | -0.218085861 | -0.490853212 | -0.25173597 | downregulated during regeneration - mid |
| ENSDART00000012376 | gabrr1 | 0.006630005 | -0.444264456 | -0.314539223 | -0.677749333 | -0.339326443 | downregulated during regeneration - mid |
| ENSDART00000012478 | mmadhc | 0.028172785 | -0.097599671 | -0.214733382 | -0.311326383 | -0.186284094 | downregulated during regeneration - mid |
| ENSDART00000012938 | phgdh | 0.077650287 | -0.315221747 | -0.241109028 | -0.448879252 | -0.249449553 | downregulated during regeneration - mid |
| ENSDART00000013411 | cahz | 0.037241402 | -0.282657507 | -0.146895201 | -0.382063619 | -0.133810291 | downregulated during regeneration - mid |
| ENSDART00000013839 | tmbim4 | 0.056276304 | -0.093382601 | -0.138429676 | -0.303111951 | -0.191824583 | downregulated during regeneration - mid |
| ENSDART00000014021 | slc25a39 | 0.007324183 | -0.175830036 | -0.233948732 | -0.630252381 | -0.299226836 | downregulated during regeneration - mid |
| ENSDART00000014031 | dpf2 | 0.020424071 | 0.062076591 | -0.192206303 | -0.356876024 | -0.045146421 | downregulated during regeneration - mid |
| ENSDART00000014632 | katnb1 | 0.009776962 | -0.1660519 | -0.366507385 | -0.386560357 | -0.02476169 | downregulated during regeneration - mid |
| ENSDART00000015710 | snrkb | 0.033001298 | -0.234771599 | -0.444259361 | -0.383002363 | -0.24868656 | downregulated during regeneration - mid |
| ENSDART00000015755 | rasl11b | 0.004084205 | -0.21929532 | -0.630137444 | -0.720127948 | -0.404206999 | downregulated during regeneration - mid |
| ENSDART00000015951 | bsg | 0.037560436 | -0.039046362 | -0.213267965 | -0.489263219 | -0.308938953 | downregulated during regeneration - mid |
| ENSDART00000016099 | CASKIN2 | 0.026969746 | -0.196661177 | -0.318487444 | -0.498724654 | -0.242993173 | downregulated during regeneration - mid |

| | | | | | | | |
|---|---|---|---|---|---|---|---|
| ENSDART00000016803 | grpel1 | 0.047568271 | -0.106418203 | -0.12256904 | -0.467582595 | -0.298130125 | downregulated during regeneration - mid |
| ENSDART00000018047 | zgc:112294 | 0.033754546 | -0.129085452 | -0.359907528 | -0.863079854 | -0.32905376 | downregulated during regeneration - mid |
| ENSDART00000018117 | ppp1r14aa | 0.009559586 | -0.10674343 | -0.179038105 | -0.425069279 | -0.232436913 | downregulated during regeneration - mid |
| ENSDART00000018972 | zgc:92818 | 0.010173835 | 0.012004445 | -0.040419062 | -0.784852086 | -0.38349701 | downregulated during regeneration - mid |
| ENSDART00000019053 | faima | 0.060563605 | -0.147906168 | -0.161120397 | -0.359996788 | -0.164587924 | downregulated during regeneration - mid |
| ENSDART00000021260 | sept8b | 0.002390041 | -0.246039495 | -0.120094514 | -0.518439508 | -0.250786208 | downregulated during regeneration - mid |
| ENSDART00000021346 | arl3l2 | 0.060063095 | -0.339022452 | -0.304754125 | -0.490852638 | -0.247921956 | downregulated during regeneration - mid |
| ENSDART00000022393 | si:dkeyp-57f11.2 | 0.005040201 | 0.163453067 | -0.122007101 | -0.459561011 | 0.032532197 | downregulated during regeneration - mid |
| ENSDART00000022688 | tob1b | 0.00387273 | -0.393522131 | -0.302261171 | -0.612204875 | -0.372656833 | downregulated during regeneration - mid |
| ENSDART00000022998 | ANO2 (1 of many) | 0.026969746 | 0.005158406 | -0.146342409 | -0.548931715 | -0.196350546 | downregulated during regeneration - mid |
| ENSDART00000023588 | guca1a | 0.012263789 | 0.527717677 | -0.52699324 | -1.299472888 | -0.244316084 | downregulated during regeneration - mid |
| ENSDART00000023686 | ankrd33ab | 0.007529247 | -0.23872945 | -0.400254403 | -1.062972805 | -0.551724897 | downregulated during regeneration - mid |
| ENSDART00000023806 | zgc:110319 | 0.067739819 | -0.195130618 | -0.122813174 | -0.337219496 | -0.098833156 | downregulated during regeneration - mid |
| ENSDART00000024136 | gngt2a | 0.005454674 | 0.175894019 | 0.078473268 | -0.788482627 | -0.299650763 | downregulated during regeneration - mid |
| ENSDART00000024331 | glsb | 0.018752626 | 0.029280722 | -0.254060441 | -0.66092512 | -0.487127171 | downregulated during regeneration - mid |
| ENSDART00000024619 | gorasp1a | 0.010796842 | -0.145027203 | -0.174354089 | -0.499227113 | -0.186036756 | downregulated during regeneration - mid |
| ENSDART00000025046 | ppp1caa | 0.031235096 | 0.04938914 | -0.175273742 | -0.396680839 | -0.049267892 | downregulated during regeneration - mid |
| ENSDART00000025414 | slc2a1a | 0.029003347 | 0.059681786 | -0.155207235 | -0.469818878 | -0.137194664 | downregulated during regeneration - mid |
| ENSDART00000025962 | gyg1a | 0.016443396 | -0.132586486 | -0.231554909 | -0.320101499 | -0.261622046 | downregulated during regeneration - mid |
| ENSDART00000026692 | ubtd1a | 0.001139558 | -0.10869502 | -0.342281946 | -0.985605367 | -0.704345293 | downregulated during regeneration - mid |
| ENSDART00000027050 | cnga3b | 0.004666996 | -0.139313288 | -0.277577971 | -1.175814494 | -0.697938349 | downregulated during regeneration - mid |
| ENSDART00000027454 | si:ch211-207i1.2 | 0.016936477 | -0.470697042 | -0.923472632 | -1.21436549 | -0.514660002 | downregulated during regeneration - mid |
| ENSDART00000028048 | necap1 | 0.01084285 | -0.117195999 | -0.341344035 | -0.478671947 | -0.252467934 | downregulated during regeneration - mid |
| ENSDART00000029459 | gipr | 0.007928614 | -0.120368109 | -0.334116608 | -0.559985722 | -0.423861096 | downregulated during regeneration - mid |
| ENSDART00000029774 | tmem55bb | 0.04854903 | -0.03621571 | -0.196356812 | -0.337053328 | -0.094153869 | downregulated during regeneration - mid |
| ENSDART00000030885 | uckl1a | 0.010869905 | 0.183203205 | -0.212616494 | -0.705475026 | -0.013040484 | downregulated during regeneration - mid |
| ENSDART00000030920 | gid8a | 0.037565208 | -0.116648875 | -0.277182294 | -0.355798783 | -0.233597587 | downregulated during regeneration - mid |
| ENSDART00000031426 | skilb | 0.000177771 | -0.234186907 | -0.553407151 | -0.565936684 | -0.420869627 | downregulated during regeneration - mid |
| ENSDART00000031638 | slc48a1a | 0.029570651 | -0.121591299 | -0.156663294 | -0.340136476 | -0.182759405 | downregulated during regeneration - mid |
| ENSDART00000031650 | hsp70l | 0.005097739 | -0.846526969 | -1.057845455 | -1.357523269 | -1.215078404 | downregulated during regeneration - mid |
| ENSDART00000032459 | aqp1a.1 | 0.010333906 | -0.275584647 | -0.402261221 | -0.531838868 | -0.410567004 | downregulated during regeneration - mid |
| ENSDART00000033325 | slc25a24 | 0.027461561 | -0.429859875 | -0.309149544 | -0.52291133 | -0.286046915 | downregulated during regeneration - mid |
| ENSDART00000034441 | tcp11l2 | 0.019467351 | 0.020775808 | -0.208761169 | -0.796222831 | -0.248557177 | downregulated during regeneration - mid |
| ENSDART00000034705 | ntmt1 | 0.034977529 | -0.020729663 | -0.187405157 | -0.44452011 | -0.329716724 | downregulated during regeneration - mid |
| ENSDART00000036373 | cfap206 | 0.011264004 | 0.026049802 | 0.014543106 | -0.421468514 | -0.063922477 | downregulated during regeneration - mid |
| ENSDART00000037065 | sccpdhb | 0.030676359 | -0.135500672 | -0.263394229 | -0.35527789 | -0.147169147 | downregulated during regeneration - mid |
| ENSDART00000038924 | sult1st1 | 0.028350112 | 0.0423301 | -0.129133641 | -0.497646461 | -0.249352869 | downregulated during regeneration - mid |
| ENSDART00000039295 | lrrfip1a | 0.001138119 | 0.201172846 | -0.052744603 | -0.60255637 | -0.234396047 | downregulated during regeneration - mid |
| ENSDART00000039485 | gabarapl2 | 0.031009754 | -0.111002032 | -0.162035427 | -0.331707696 | -0.213225245 | downregulated during regeneration - mid |
| ENSDART00000040116 | tnrc5 | 0.037247868 | -0.132560784 | -0.181210054 | -0.368693498 | -0.22290886 | downregulated during regeneration - mid |
| ENSDART00000040346 | efr3ba | 0.045439062 | -0.15754636 | -0.295932904 | -0.334562517 | -0.212090743 | downregulated during regeneration - mid |
| ENSDART00000041417 | camk1b | 0.063189142 | -0.259356662 | -0.222277099 | -0.408590245 | -0.120593834 | downregulated during regeneration - mid |
| ENSDART00000041443 | igsf21a | 0.078639351 | -0.224595347 | -0.278632007 | -0.419145006 | -0.220026937 | downregulated during regeneration - mid |
| ENSDART00000041707 | unc119a | 0.031766969 | -0.21008147 | -0.190962771 | -0.5453967 | -0.153347487 | downregulated during regeneration - mid |
| ENSDART00000042276 | nxph1 | 0.036299878 | -0.320960824 | -0.220089718 | -0.50268354 | -0.230788559 | downregulated during regeneration - mid |
| ENSDART00000042572 | ablim1b | 0.036021217 | 0.023130185 | -0.084172657 | -0.818233794 | -0.360713717 | downregulated during regeneration - mid |
| ENSDART00000042599 | dennd6aa | 0.041657173 | -0.0532753 | -0.029202145 | -0.371955611 | -0.136277272 | downregulated during regeneration - mid |
| ENSDART00000043492 | trappc6bl | 0.010042504 | -0.12751818 | -0.21743347 | -0.476970052 | -0.302538631 | downregulated during regeneration - mid |
| ENSDART00000043507 | ciarta | 0.063631477 | -0.034891069 | -0.156171585 | -0.423543489 | -0.111653268 | downregulated during regeneration - mid |
| ENSDART00000043801 | cabp5b | 0.010073304 | -0.44296443 | -0.237302409 | -0.884658288 | -0.372660272 | downregulated during regeneration - mid |
| ENSDART00000043855 | dclk2a | 0.001528476 | -0.104441255 | -0.395008642 | -0.62766162 | -0.184810585 | downregulated during regeneration - mid |
| ENSDART00000043953 | mfsd2b | 0.011605032 | -0.226277331 | -0.20295362 | -0.451440771 | 0.020277702 | downregulated during regeneration - mid |
| ENSDART00000045303 | tmprss9 | 0.052101742 | -0.116725282 | -0.194770382 | -0.365140883 | -0.064254559 | downregulated during regeneration - mid |
| ENSDART00000045374 | smad3a | 0.043076195 | -0.063329624 | -0.157712113 | -0.303211349 | -0.108940629 | downregulated during regeneration - mid |
| ENSDART00000046066 | capn1a | 0.045050212 | -0.174433704 | -0.141559786 | -0.304523077 | -0.134358035 | downregulated during regeneration - mid |
| ENSDART00000046951 | ptpn11b | 0.011788375 | -0.12533711 | -0.212829251 | -0.391615008 | -0.186327808 | downregulated during regeneration - mid |
| ENSDART00000047362 | msra | 0.006658285 | -0.031803809 | -0.196438308 | -0.422002476 | -0.295991402 | downregulated during regeneration - mid |
| ENSDART00000049885 | si:dkey-172j4.3 | 0.036535058 | -0.146868316 | -0.089398841 | -0.317501099 | -0.03734398 | downregulated during regeneration - mid |
| ENSDART00000050202 | rcan3 | 0.007786427 | -0.240590006 | -0.307380747 | -0.423427653 | -0.130410568 | downregulated during regeneration - mid |
| ENSDART00000050271 | hexb | 0.004813371 | 0.237342774 | -0.182676311 | -0.489143261 | -0.233593029 | downregulated during regeneration - mid |
| ENSDART00000050750 | rrm2b | 0.073861488 | -0.324758311 | -0.221575036 | -0.394470152 | -0.266012642 | downregulated during regeneration - mid |
| ENSDART00000051566 | zgc:101016 | 0.024284969 | 0.072400267 | -0.278957969 | -0.822407258 | -0.192676389 | downregulated during regeneration - mid |
| ENSDART00000052065 | si:rp71-39b20.4 | 0.019747468 | -0.313076871 | -0.443642109 | -0.50831344 | -0.2481846 | downregulated during regeneration - mid |
| ENSDART00000052318 | mdka | 0.007797716 | -0.324471926 | -0.37363799 | -0.54050807 | -0.370260207 | downregulated during regeneration - mid |
| ENSDART00000052322 | zgc:110699 | 0.012719677 | -0.266905411 | -0.248249923 | -0.482205646 | -0.223841905 | downregulated during regeneration - mid |
| ENSDART00000053367 | hmgn3 | 0.044180927 | -0.174648563 | -0.165335262 | -0.310027237 | -0.181225286 | downregulated during regeneration - mid |
| ENSDART00000053494 | anks4b | 0.048286815 | -0.562445249 | -0.270911364 | -0.580434775 | -0.289114333 | downregulated during regeneration - mid |
| ENSDART00000054007 | slc8a4b | 0.027048639 | -0.011734485 | -0.299949751 | -0.451416872 | -0.315264472 | downregulated during regeneration - mid |
| ENSDART00000054408 | gsg1l | 0.001512737 | -0.338795387 | -0.334881942 | -0.642215658 | -0.428166696 | downregulated during regeneration - mid |
| ENSDART00000054472 | tll1 | 0.030023854 | -0.241128673 | -0.208136 | -0.494254448 | -0.202017577 | downregulated during regeneration - mid |
| ENSDART00000054760 | zgc:162144 | 0.018655612 | -0.154581168 | -0.586579914 | -0.580706302 | -0.278802593 | downregulated during regeneration - mid |
| ENSDART00000055186 | atp5j2 | 0.01462956 | -0.168904246 | -0.309514947 | -0.449394402 | -0.201772036 | downregulated during regeneration - mid |
| ENSDART00000055611 | isca2 | 0.041099023 | -0.127743819 | -0.187288303 | -0.524298231 | -0.307747827 | downregulated during regeneration - mid |
| ENSDART00000055817 | PIGG | 0.023931815 | -0.041747749 | -0.262998575 | -0.390921965 | -0.198616638 | downregulated during regeneration - mid |
| ENSDART00000056213 | pik3r1 | 0.029029749 | -0.34639661 | -0.306212461 | -0.436349985 | -0.247937167 | downregulated during regeneration - mid |

| | | | | | | | |
|---|---|---|---|---|---|---|---|
| ENSDART00000056734 | setd7 | 0.028411307 | -0.044426299 | -0.050437779 | -0.332150365 | -0.191170985 | downregulated during regeneration - mid |
| ENSDART00000056735 | rgs20 | 0.035861975 | -0.263112929 | -0.317514258 | -0.646869905 | -0.425880318 | downregulated during regeneration - mid |
| ENSDART00000056927 | egln1a | 0.069201585 | -0.373304639 | -0.331810821 | -0.548972472 | -0.202283187 | downregulated during regeneration - mid |
| ENSDART00000057918 | si:ch211-147h1.4 | 0.006788162 | 0.058919084 | -0.340722361 | -0.680330981 | -0.211102009 | downregulated during regeneration - mid |
| ENSDART00000057957 | itm2cb | 0.004277486 | -0.410626815 | -0.396999775 | -0.657592832 | -0.426965619 | downregulated during regeneration - mid |
| ENSDART00000058255 | bbs5 | 0.033037209 | -0.153890395 | -0.351139568 | -0.380364475 | -0.12338798 | downregulated during regeneration - mid |
| ENSDART00000058339 | ap3s2 | 0.051751157 | -0.205243617 | -0.191498501 | -0.306412801 | -0.176311273 | downregulated during regeneration - mid |
| ENSDART00000058470 | pik3r1 | 0.020653282 | -0.272621258 | -0.391826992 | -0.565271638 | -0.177820971 | downregulated during regeneration - mid |
| ENSDART00000058628 | ccsapb | 0.038389384 | -0.109779777 | -0.249497929 | -0.321372328 | -0.06107705 | downregulated during regeneration - mid |
| ENSDART00000058785 | fam210ab | 0.009653548 | -0.074078069 | -0.284264661 | -0.891770658 | -0.393378695 | downregulated during regeneration - mid |
| ENSDART00000058936 | scamp5b | 0.046680552 | -0.278094691 | -0.317619737 | -0.403892079 | -0.134676622 | downregulated during regeneration - mid |
| ENSDART00000060001 | pnp6 | 0.007828891 | -0.139557872 | -0.321502493 | -0.386186078 | -0.030509602 | downregulated during regeneration - mid |
| ENSDART00000060015 | chka | 0.008595135 | -0.144459192 | -0.274568124 | -0.456317122 | -0.171918553 | downregulated during regeneration - mid |
| ENSDART00000060184 | chka | 0.004120762 | -0.101617548 | -0.321325171 | -0.420229119 | -0.136541498 | downregulated during regeneration - mid |
| ENSDART00000060532 | zgc:110796 | 0.029635773 | -0.139659209 | -0.284517331 | -0.33969202 | -0.16468656 | downregulated during regeneration - mid |
| ENSDART00000061106 | bhlhe41 | 0.000553147 | 0.043056173 | -0.423617996 | -0.754959095 | -0.246095493 | downregulated during regeneration - mid |
| ENSDART00000061265 | rnf141 | 0.032782977 | -0.12379903 | -0.215585747 | -0.342006011 | -0.212187704 | downregulated during regeneration - mid |
| ENSDART00000061745 | inpp4ab | 0.000385909 | -0.337306716 | -0.94773917 | -0.831022148 | -0.394471236 | downregulated during regeneration - mid |
| ENSDART00000062402 | tpd52l1 | 0.005726323 | -0.180007689 | -0.341056198 | -0.772074089 | -0.368132956 | downregulated during regeneration - mid |
| ENSDART00000062556 | sod2 | 0.00889295 | -0.183508685 | -0.287491323 | -0.424543766 | -0.237833299 | downregulated during regeneration - mid |
| ENSDART00000062935 | heca | 0.065650579 | -0.371152131 | -0.569989768 | -0.691744059 | -0.51577268 | downregulated during regeneration - mid |
| ENSDART00000063835 | otx5 | 0.082497078 | -0.207695123 | -0.226833273 | -0.417236259 | -0.126675877 | downregulated during regeneration - mid |
| ENSDART00000063944 | tmem30ab | 0.018132587 | -0.052321591 | -0.186777496 | -0.351757869 | -0.167369044 | downregulated during regeneration - mid |
| ENSDART00000064738 | atpif1b | 0.005335064 | -0.12884804 | -0.286001583 | -0.530012586 | -0.266598201 | downregulated during regeneration - mid |
| ENSDART00000064842 | padi2 | 0.046738515 | -0.110475283 | -0.200123181 | -0.394420172 | -0.195593685 | downregulated during regeneration - mid |
| ENSDART00000064913 | fto | 0.054038459 | -0.098234146 | -0.428383355 | -0.257568512 | -0.239748235 | downregulated during regeneration - mid |
| ENSDART00000065143 | unc119b | 0.020870612 | -0.086346577 | -0.282642438 | -0.651558775 | -0.265879719 | downregulated during regeneration - mid |
| ENSDART00000065467 | dedd1 | 0.0321404 | -0.00326722 | -0.100306912 | -0.35251663 | -0.041277003 | downregulated during regeneration - mid |
| ENSDART00000065755 | gpn3 | 0.095709295 | -0.172157475 | -0.155442383 | -0.381057864 | -0.19097226 | downregulated during regeneration - mid |
| ENSDART00000066256 | vti1a | 0.042541047 | -0.147124368 | -0.216073091 | -0.331785573 | -0.195609352 | downregulated during regeneration - mid |
| ENSDART00000066259 | kcnk1a | 0.055503316 | -0.289802216 | -0.227947241 | -0.67177 | -0.392395516 | downregulated during regeneration - mid |
| ENSDART00000066380 | ca7 | 0.023602236 | -0.236035328 | -0.017625204 | -0.765902599 | -0.026455411 | downregulated during regeneration - mid |
| ENSDART00000066382 | aqp8a.1 | 0.025161694 | -0.164652843 | -0.325293072 | -0.493634593 | -0.109118528 | downregulated during regeneration - mid |
| ENSDART00000066655 | mybl1 | 0.005666275 | -0.255810306 | -0.487313899 | -0.72749049 | -0.668391364 | downregulated during regeneration - mid |
| ENSDART00000066963 | atp6v1f | 0.047327566 | -0.089791813 | -0.169571399 | -0.304106913 | -0.234245815 | downregulated during regeneration - mid |
| ENSDART00000067005 | bcat1 | 0.046674561 | -0.064708558 | -0.22197643 | -0.301840187 | -0.097404482 | downregulated during regeneration - mid |
| ENSDART00000067078 | plekhg5a | 0.081075709 | -0.147306721 | -0.274518319 | -0.555343847 | -0.255866929 | downregulated during regeneration - mid |
| ENSDART00000067448 | acat1 | 0.021507957 | -0.217922513 | -0.251881532 | -0.396031362 | -0.209577052 | downregulated during regeneration - mid |
| ENSDART00000067510 | crabp1a | 0.029243935 | -0.186487239 | -0.41656224 | -0.606281006 | -0.420918895 | downregulated during regeneration - mid |
| ENSDART00000067764 | stk17a | 0.02229764 | -0.185636748 | -0.198053553 | -0.515379898 | -0.253038095 | downregulated during regeneration - mid |
| ENSDART00000073694 | smu1b | 0.006932921 | -0.023572641 | -0.696487436 | -0.934479138 | -0.393819626 | downregulated during regeneration - mid |
| ENSDART00000073932 | NA | 0.003072278 | 0.121194935 | -0.376988949 | -0.788016576 | -0.265683315 | downregulated during regeneration - mid |
| ENSDART00000074117 | aspa | 0.007828891 | -0.032383232 | -0.304124913 | -0.397629458 | -0.358898864 | downregulated during regeneration - mid |
| ENSDART00000075070 | hsf2 | 0.014034602 | -0.066635095 | -0.276496416 | -0.474097609 | -0.186498896 | downregulated during regeneration - mid |
| ENSDART00000075421 | sord | 0.017496481 | -0.008165085 | -0.251228216 | -0.437538687 | -0.226988953 | downregulated during regeneration - mid |
| ENSDART00000075513 | aqp9b | 0.023071688 | -0.181865878 | -0.358464178 | -0.54013493 | -0.322928595 | downregulated during regeneration - mid |
| ENSDART00000076929 | prkg2 | 0.055091796 | -0.322610383 | -0.221985796 | -0.523406711 | -0.304941411 | downregulated during regeneration - mid |
| ENSDART00000077619 | b3gat1b | 0.017905694 | -0.266663197 | -0.223974588 | -0.393683459 | -0.125347427 | downregulated during regeneration - mid |
| ENSDART00000077724 | gnb5b | 0.032645202 | -0.216593155 | -0.279473354 | -0.485943487 | -0.092161444 | downregulated during regeneration - mid |
| ENSDART00000077839 | atf7b | 0.014656899 | -0.232485852 | -0.196508033 | -0.391624214 | -0.206346003 | downregulated during regeneration - mid |
| ENSDART00000078014 | poldip2 | 0.028918785 | 0.012504595 | -0.191194187 | -0.45353852 | -0.221096745 | downregulated during regeneration - mid |
| ENSDART00000078226 | mtnr1bb | 0.021380755 | -0.285812097 | -0.289316068 | -0.599481549 | -0.276891324 | downregulated during regeneration - mid |
| ENSDART00000078311 | zgc:154093 | 0.050880744 | -0.170397489 | -0.310689773 | -0.421136702 | -0.191616401 | downregulated during regeneration - mid |
| ENSDART00000078336 | klc3 | 0.011790336 | -0.15725421 | -0.267140148 | -0.672772286 | -0.331888789 | downregulated during regeneration - mid |
| ENSDART00000078838 | rab3aa | 0.031838485 | -0.225658115 | -0.303506805 | -0.374028659 | -0.138563861 | downregulated during regeneration - mid |
| ENSDART00000078908 | usp1 | 0.022814815 | -0.163356677 | -0.229514103 | -0.329514373 | -0.251997637 | downregulated during regeneration - mid |
| ENSDART00000079104 | ndufs6 | 0.003180081 | -0.146008815 | -0.377028896 | -0.443194118 | -0.327081615 | downregulated during regeneration - mid |
| ENSDART00000079629 | ppm1nb | 0.004971122 | -0.279926594 | -0.345062767 | -0.462173038 | -0.171806172 | downregulated during regeneration - mid |
| ENSDART00000079686 | zmp:0000001103 | 0.006141198 | 0.143973249 | -0.889323937 | -0.759747318 | -0.487193938 | downregulated during regeneration - mid |
| ENSDART00000079810 | si:dkey-222b8.4 | 0.001538399 | -0.076664011 | -0.231174428 | -0.530241167 | -0.088965105 | downregulated during regeneration - mid |
| ENSDART00000079866 | slc30a9 | 0.045584509 | -0.061810697 | -0.211371943 | -0.307919743 | -0.172076689 | downregulated during regeneration - mid |
| ENSDART00000080389 | fam13a | 0.004095858 | 0.342651645 | -0.065971229 | -0.119440295 | 0.116639155 | downregulated during regeneration - mid |
| ENSDART00000081214 | selt1a | 0.041966249 | -0.31898529 | -0.284714031 | -0.513594635 | -0.412445291 | downregulated during regeneration - mid |
| ENSDART00000081325 | dynll1 | 0.0066372 | -0.057380796 | -0.1118203 | -0.516198015 | -0.112248637 | downregulated during regeneration - mid |
| ENSDART00000081611 | cgnb | 0.001086517 | -0.200219306 | -0.438466186 | -1.067503957 | -0.17391972 | downregulated during regeneration - mid |
| ENSDART00000081761 | bin1b | 0.028778753 | -0.023096846 | -0.153451242 | -0.459181763 | -0.132136814 | downregulated during regeneration - mid |
| ENSDART00000081781 | PLEKHG3 | 0.034279564 | -0.076813883 | -0.173391418 | -0.382548311 | -0.034437681 | downregulated during regeneration - mid |
| ENSDART00000082066 | atpv0e2 | 0.002085662 | -0.268294846 | -0.377237748 | -0.604078331 | -0.39998077 | downregulated during regeneration - mid |
| ENSDART00000082523 | impa2 | 0.00795396 | -0.208830066 | -0.398914852 | -0.498863304 | -0.267214166 | downregulated during regeneration - mid |
| ENSDART00000082622 | fsd1l | 0.03087552 | -0.046506283 | -0.340407639 | -0.41095417 | -0.201162325 | downregulated during regeneration - mid |
| ENSDART00000082937 | fscn2a | 0.054349324 | -0.244556215 | -0.436229092 | -0.571250568 | -0.199107259 | downregulated during regeneration - mid |
| ENSDART00000084007 | TULP2 | 0.035633594 | -0.184892554 | -0.234742374 | -0.492430755 | -0.167640019 | downregulated during regeneration - mid |
| ENSDART00000084011 | cplx4a | 0.024776472 | -0.220118176 | -0.343953082 | -0.589002173 | -0.338126511 | downregulated during regeneration - mid |
| ENSDART00000084135 | ubtd1a | 0.002016009 | -0.22864625 | -0.383233305 | -0.947657938 | -0.688823144 | downregulated during regeneration - mid |
| ENSDART00000084353 | tbc1d10ab | 0.051425296 | -0.174770874 | -0.349891521 | -0.619429045 | -0.244339697 | downregulated during regeneration - mid |

| | | | | | | | |
|---|---|---|---|---|---|---|---|
| ENSDART00000084417 | timm17b | 0.014386448 | -0.134748392 | -0.15578708 | -0.404232794 | -0.18557959 | downregulated during regeneration - mid |
| ENSDART00000084792 | prosc | 0.057325828 | -0.109541047 | -0.121094609 | -0.31852643 | -0.202300262 | downregulated during regeneration - mid |
| ENSDART00000084965 | cep104 | 0.022501885 | -0.196545786 | -0.225873142 | -0.456327943 | -0.208454995 | downregulated during regeneration - mid |
| ENSDART00000085252 | pqlc3 | 0.004793697 | -0.117366098 | -0.27675705 | -0.43476147 | -0.253093536 | downregulated during regeneration - mid |
| ENSDART00000085277 | pfkmb | 0.010085498 | 0.010653815 | -0.414332737 | -0.69351419 | -0.144457554 | downregulated during regeneration - mid |
| ENSDART00000085693 | gpm6bb | 0.009450076 | -0.236628216 | -0.325741908 | -0.400571921 | -0.218025638 | downregulated during regeneration - mid |
| ENSDART00000085716 | mtmr10 | 0.000364305 | 0.168573884 | -0.120117023 | -0.772748173 | -0.357648365 | downregulated during regeneration - mid |
| ENSDART00000086263 | mettl7a | 0.05857751 | -0.420009286 | -0.158826062 | -0.791334734 | -0.334080113 | downregulated during regeneration - mid |
| ENSDART00000086619 | prkca | 0.027625402 | -0.184045571 | -0.418285198 | -0.5357326 | -0.310736719 | downregulated during regeneration - mid |
| ENSDART00000086753 | dapk2a | 0.010335438 | 0.129419785 | -0.339935803 | -1.219284243 | -0.369104907 | downregulated during regeneration - mid |
| ENSDART00000086994 | nat15 | 0.031838485 | -0.144704703 | -0.149678764 | -0.337458722 | -0.152039739 | downregulated during regeneration - mid |
| ENSDART00000087070 | abcc5 | 0.0132651 | 0.063506972 | -0.209331316 | -0.331966374 | -0.070006315 | downregulated during regeneration - mid |
| ENSDART00000087300 | gabrb3 | 0.027218185 | -0.106002759 | -0.319333142 | -0.574890474 | -0.11105969 | downregulated during regeneration - mid |
| ENSDART00000087570 | BRSK2 (1 of many) | 0.039389579 | -0.035673495 | -0.289130617 | -0.495870531 | -0.115059951 | downregulated during regeneration - mid |
| ENSDART00000087643 | tesk2 | 0.017205213 | -0.250604524 | -0.077411225 | -0.349635246 | -0.097357509 | downregulated during regeneration - mid |
| ENSDART00000088027 | ssx2ipb | 0.061068477 | -0.254273665 | -0.300537366 | -0.659689526 | -0.365610708 | downregulated during regeneration - mid |
| ENSDART00000089042 | kcnh4b | 0.028142339 | -0.293363138 | -0.305961812 | -0.46205676 | -0.207459851 | downregulated during regeneration - mid |
| ENSDART00000089161 | CCDC181 | 0.032175544 | -0.238708752 | -0.276570098 | -0.364394143 | -0.212879757 | downregulated during regeneration - mid |
| ENSDART00000089526 | otc | 0.028981739 | -0.228950714 | -0.50691522 | -0.941930968 | -0.587900491 | downregulated during regeneration - mid |
| ENSDART00000090079 | synm | 0.001644836 | -0.113622285 | -0.399027729 | -0.70355851 | -0.251171145 | downregulated during regeneration - mid |
| ENSDART00000090484 | tecpr1a | 0.027062614 | -0.079583722 | -0.244227266 | -0.30265493 | -0.135390116 | downregulated during regeneration - mid |
| ENSDART00000090611 | sh3gl2a | 0.003738261 | -0.3620314 | -0.426828792 | -0.627861098 | -0.256916064 | downregulated during regeneration - mid |
| ENSDART00000091183 | erfl3 | 0.034693937 | -0.064022984 | -0.222171088 | -0.380622737 | -0.084018803 | downregulated during regeneration - mid |
| ENSDART00000091452 | TULP2 | 0.003694199 | -0.183522253 | -0.233522919 | -0.588056807 | -0.14814607 | downregulated during regeneration - mid |
| ENSDART00000091615 | iffo1a | 0.045036938 | -0.255558776 | -0.319129649 | -0.402938235 | -0.1114809 | downregulated during regeneration - mid |
| ENSDART00000091620 | atp8a1 | 0.025084869 | -0.02813322 | -0.262306637 | -0.342910163 | -0.163047911 | downregulated during regeneration - mid |
| ENSDART00000091707 | dbpa | 0.017575469 | -0.158262702 | -0.327851995 | -0.357551076 | -0.129469939 | downregulated during regeneration - mid |
| ENSDART00000091729 | mlc1 | 0.000239262 | -0.413445193 | -0.441446959 | -0.597560289 | -0.400243567 | downregulated during regeneration - mid |
| ENSDART00000092114 | ERBB4 (1 of many) | 0.013394015 | -0.124937285 | -0.553537291 | -0.521885412 | -0.338309306 | downregulated during regeneration - mid |
| ENSDART00000092416 | rabl2 | 0.018433591 | -0.237808365 | -0.250658742 | -0.360133928 | -0.144079351 | downregulated during regeneration - mid |
| ENSDART00000093093 | coro2bb | 0.051450895 | -0.315281988 | -0.251087256 | -0.471013192 | -0.151819548 | downregulated during regeneration - mid |
| ENSDART00000093331 | rreb1a | 0.018126285 | -0.024901133 | -0.155149005 | -0.517234878 | -0.050769944 | downregulated during regeneration - mid |
| ENSDART00000097338 | napaa | 0.005374512 | -0.21867082 | -0.198889828 | -0.441263528 | -0.268359175 | downregulated during regeneration - mid |
| ENSDART00000097466 | fam169aa | 0.080339729 | -0.25560307 | -0.389662586 | -0.539552997 | -0.303655 | downregulated during regeneration - mid |
| ENSDART00000098263 | kctd9a | 0.018790846 | -0.029411295 | -0.237424393 | -0.359070767 | -0.190010537 | downregulated during regeneration - mid |
| ENSDART00000098648 | gc2 | 0.024271936 | -0.195652835 | -0.378422178 | -0.402398473 | -0.153861921 | downregulated during regeneration - mid |
| ENSDART00000099180 | elovl8a | 0.019558454 | -0.351208319 | -0.254455279 | -0.702580214 | -0.196165774 | downregulated during regeneration - mid |
| ENSDART00000099244 | CDHR2 | 0.017384242 | -0.127835174 | -0.277679177 | -0.519160965 | -0.069385603 | downregulated during regeneration - mid |
| ENSDART00000099392 | irgq2 | 0.055536456 | -0.080759981 | -0.21229475 | -0.382096269 | -0.042741954 | downregulated during regeneration - mid |
| ENSDART00000100131 | si:ch211-242e8.1 | 0.00099671 | 0.043404066 | -0.450710823 | -0.466087638 | 0.073034701 | downregulated during regeneration - mid |
| ENSDART00000100194 | msi2b | 0.036147745 | -0.174290948 | -0.286012891 | -0.448717518 | -0.171818835 | downregulated during regeneration - mid |
| ENSDART00000100287 | grk7a | 0.01613952 | -0.29429259 | -0.34179807 | -0.766271921 | -0.26689268 | downregulated during regeneration - mid |
| ENSDART00000100667 | skia | 0.021156666 | -0.180939747 | -0.108825609 | -0.328725059 | -0.076853639 | downregulated during regeneration - mid |
| ENSDART00000100762 | inpp4ab | 0.000398071 | -0.155677368 | -0.41981961 | -0.539565797 | -0.358079101 | downregulated during regeneration - mid |
| ENSDART00000100869 | ppp3r1b | 0.04032679 | -0.102347685 | -0.214447853 | -0.31663745 | -0.211814655 | downregulated during regeneration - mid |
| ENSDART00000100885 | nrn1la | 0.001326538 | -0.520024314 | -0.30927413 | -0.690569972 | -0.461401545 | downregulated during regeneration - mid |
| ENSDART00000101576 | tmem230b | 0.009263143 | 0.001436175 | -0.176978383 | -0.445323711 | -0.253116022 | downregulated during regeneration - mid |
| ENSDART00000102898 | zgc:158258 | 0.013286981 | -0.205000336 | -0.263168834 | -0.54350319 | -0.257007642 | downregulated during regeneration - mid |
| ENSDART00000102969 | spock3 | 0.026589151 | -0.0552616 | -0.214726057 | -0.592493725 | -0.285702337 | downregulated during regeneration - mid |
| ENSDART00000103070 | cdk17 | 0.006385907 | 0.018864851 | -0.23502559 | -0.366641807 | -0.137520544 | downregulated during regeneration - mid |
| ENSDART00000103267 | fam212ab | 0.059492509 | -0.276107898 | -0.473876533 | -0.614056911 | -0.421775209 | downregulated during regeneration - mid |
| ENSDART00000103293 | ndufa5 | 0.014686352 | -0.136898371 | -0.38263765 | -0.527827076 | -0.351287298 | downregulated during regeneration - mid |
| ENSDART00000103450 | lactbl1b | 0.002609113 | -0.310148031 | -0.346227281 | -0.611014657 | -0.245438949 | downregulated during regeneration - mid |
| ENSDART00000103487 | zgc:195001 | 0.006385907 | -0.335765581 | -0.344851478 | -0.707965666 | -0.31393888 | downregulated during regeneration - mid |
| ENSDART00000103628 | btbd6a | 0.003515537 | -0.278201553 | -0.439382236 | -0.547351526 | -0.157769223 | downregulated during regeneration - mid |
| ENSDART00000104234 | gjd2b | 0.009000055 | -0.226334837 | -0.451484904 | -0.506629051 | -0.204050045 | downregulated during regeneration - mid |
| ENSDART00000104279 | znf516 | 0.018737197 | -0.151066862 | -0.30868422 | -0.587761132 | -0.246031248 | downregulated during regeneration - mid |
| ENSDART00000104298 | ndufb5 | 0.062748344 | -0.141095269 | -0.212773777 | -0.353488244 | -0.122828248 | downregulated during regeneration - mid |
| ENSDART00000104307 | eml1 | 0.005712911 | 0.215532654 | -0.343903678 | -0.604837308 | -0.271121591 | downregulated during regeneration - mid |
| ENSDART00000104487 | cox4i2 | 0.004567663 | 0.027176247 | -0.218042175 | -0.688187338 | -0.440124172 | downregulated during regeneration - mid |
| ENSDART00000104536 | chsy1 | 0.002085662 | 0.021777196 | -0.175630111 | -0.644756219 | -0.079001152 | downregulated during regeneration - mid |
| ENSDART00000104674 | camk1db | 0.011089782 | -0.318172003 | -0.339895825 | -0.49572727 | -0.229789777 | downregulated during regeneration - mid |
| ENSDART00000104708 | abhd12 | 0.030797346 | -0.134054463 | -0.304895231 | -0.501902497 | -0.262631127 | downregulated during regeneration - mid |
| ENSDART00000105485 | si:dkey-78l4.14 | 0.009178848 | -0.103336903 | -0.24562657 | -0.381336593 | -0.194197896 | downregulated during regeneration - mid |
| ENSDART00000105545 | arl3 | 0.035698906 | -0.028403306 | -0.158533169 | -0.38830015 | -0.193573844 | downregulated during regeneration - mid |
| ENSDART00000105561 | si:dkey-42p8.3 | 0.005648279 | -0.126414242 | -0.359710173 | -0.388006213 | -0.105772793 | downregulated during regeneration - mid |
| ENSDART00000105602 | elovl6 | 0.064667433 | -0.283618771 | -0.171237628 | -0.388002017 | -0.174233768 | downregulated during regeneration - mid |
| ENSDART00000105667 | si:dkey-121a11.3 | 0.07695913 | -0.369671493 | -0.58967553 | -0.842073302 | -0.52980672 | downregulated during regeneration - mid |
| ENSDART00000105681 | cdc14ab | 0.008133889 | -0.104501589 | -0.239167071 | -0.39502699 | -0.259112662 | downregulated during regeneration - mid |
| ENSDART00000105974 | pvalb9 | 0.005388807 | -0.398960737 | -0.428084025 | -0.618672138 | -0.278532192 | downregulated during regeneration - mid |
| ENSDART00000108943 | shroom2a | 0.0263983 | 0.03299057 | -0.127271939 | -0.499446332 | -0.229050052 | downregulated during regeneration - mid |
| ENSDART00000108989 | adamts14 | 0.018280387 | -0.185318243 | -0.40325002 | -0.417840329 | -0.182643804 | downregulated during regeneration - mid |
| ENSDART00000109023 | fmnl3 | 0.0083599 | -0.186687955 | -0.312847961 | -0.533074314 | -0.280720309 | downregulated during regeneration - mid |
| ENSDART00000109037 | cdk5r2b | 0.061835261 | -0.48847507 | -0.44246865 | -0.592226601 | -0.418268347 | downregulated during regeneration - mid |
| ENSDART00000109314 | arhgap10 | 0.008289796 | -0.214061464 | -0.171864796 | -0.398074347 | -0.037252057 | downregulated during regeneration - mid |

| | | | | | | | |
|---|---|---|---|---|---|---|---|
| ENSDART00000109440 | adamts9 | 0.042577687 | -0.162680626 | -0.21784206 | -0.326391297 | -0.145144869 | downregulated during regeneration - mid |
| ENSDART00000109485 | gal3st3 | 0.008289796 | -0.139388785 | -0.282505485 | -0.388230748 | -0.163229068 | downregulated during regeneration - mid |
| ENSDART00000109567 | nhsa | 0.005353377 | -0.05015222 | -0.331046598 | -0.348641258 | -0.190382203 | downregulated during regeneration - mid |
| ENSDART00000109570 | NA | 0.009680262 | -0.315520739 | -0.355339215 | -1.137398742 | -0.355379016 | downregulated during regeneration - mid |
| ENSDART00000109604 | palmdb | 0.002896578 | -0.228743892 | -0.477529612 | -0.779896236 | -0.307691368 | downregulated during regeneration - mid |
| ENSDART00000109876 | sun1 | 0.020586406 | -0.255190881 | -0.339472925 | -0.466317308 | -0.169389668 | downregulated during regeneration - mid |
| ENSDART00000110033 | abhd8b | 0.063929589 | -0.069420963 | -0.144372686 | -0.35601389 | -0.098305508 | downregulated during regeneration - mid |
| ENSDART00000110041 | lrrc38a | 0.006744186 | -0.167375954 | -0.314913504 | -0.608219505 | -0.293673506 | downregulated during regeneration - mid |
| ENSDART00000110431 | si:ch73-280o22.2 | 0.030699866 | -0.128270202 | -0.267395445 | -0.366965125 | -0.13375727 | downregulated during regeneration - mid |
| ENSDART00000110571 | fam150a | 0.015571917 | -0.089437558 | -0.463371816 | -0.643350488 | -0.328581938 | downregulated during regeneration - mid |
| ENSDART00000110606 | abhd16a | 0.001679054 | 0.034068276 | -0.290276411 | -0.385494504 | -0.249441823 | downregulated during regeneration - mid |
| ENSDART00000111021 | si:dkeyp-14d3.1 | 0.005053949 | -0.147889699 | -0.589075251 | -0.655456899 | -0.257575305 | downregulated during regeneration - mid |
| ENSDART00000111271 | CABZ01080074.2 | 0.007536007 | 0.091182245 | -0.411344825 | -0.551440399 | -0.297376381 | downregulated during regeneration - mid |
| ENSDART00000111343 | lmf2a | 0.022218278 | -0.264313993 | -0.256081583 | -0.425329386 | -0.228346273 | downregulated during regeneration - mid |
| ENSDART00000111536 | rmdn2 | 0.042891 | -0.153020266 | -0.402861584 | -0.346146607 | -0.222410217 | downregulated during regeneration - mid |
| ENSDART00000111625 | ninl | 0.004129329 | -0.236310585 | -0.326810382 | -0.437556672 | -0.154099243 | downregulated during regeneration - mid |
| ENSDART00000111666 | npdc1a | 0.037722848 | -0.090316422 | -0.266587539 | -0.30135563 | -0.100212297 | downregulated during regeneration - mid |
| ENSDART00000111717 | fscn2b | 0.015420308 | -0.120313044 | -0.274854028 | -0.483029563 | -0.127989104 | downregulated during regeneration - mid |
| ENSDART00000111993 | si:dkey-19b23.15 | 0.001555418 | -0.211537347 | -0.357802737 | -0.511679441 | -0.364770311 | downregulated during regeneration - mid |
| ENSDART00000112156 | si:ch211-253b8.5 | 0.014985328 | -0.087783514 | -0.192500909 | -0.49330541 | -0.306875488 | downregulated during regeneration - mid |
| ENSDART00000112768 | ubald1a | 0.0312149 | -0.163342676 | -0.260286427 | -0.366999537 | -0.147535942 | downregulated during regeneration - mid |
| ENSDART00000112967 | rilp | 0.001611738 | -0.073950343 | -0.174960248 | -0.511756745 | -0.145692497 | downregulated during regeneration - mid |
| ENSDART00000113089 | slc4a11 | 0.060667717 | -0.230777933 | -0.44932083 | -0.596234565 | -0.271629672 | downregulated during regeneration - mid |
| ENSDART00000113362 | ip6k1 | 0.017686321 | -0.059673843 | -0.326658711 | -0.456047384 | -0.047258149 | downregulated during regeneration - mid |
| ENSDART00000113376 | fam13b | 0.00601663 | -0.247096949 | -0.341873515 | -0.575011995 | -0.179280023 | downregulated during regeneration - mid |
| ENSDART00000113384 | lyrm4 | 0.026927975 | -0.082514668 | -0.167513155 | -0.434528553 | -0.231621665 | downregulated during regeneration - mid |
| ENSDART00000113649 | impg1a | 0.03238504 | -0.10562067 | -0.311339049 | -0.59002435 | -0.135488682 | downregulated during regeneration - mid |
| ENSDART00000113799 | si:ch211-132b12.7 | 0.001099064 | 0.091573373 | -0.269264305 | -0.847675231 | -0.371595689 | downregulated during regeneration - mid |
| ENSDART00000113853 | prokr1a | 0.062959243 | -0.220461946 | -0.109814734 | -0.599222392 | -0.17974305 | downregulated during regeneration - mid |
| ENSDART00000113912 | ntm | 0.012936971 | -0.178976918 | -0.363265579 | -0.513893339 | -0.211182567 | downregulated during regeneration - mid |
| ENSDART00000114118 | CU639468.1 | 0.004971122 | -0.073130118 | -0.36036163 | -0.387238652 | -0.163804242 | downregulated during regeneration - mid |
| ENSDART00000114134 | wrb | 0.049748908 | -0.123951898 | -0.095408141 | -0.313002762 | -0.1622087 | downregulated during regeneration - mid |
| ENSDART00000114172 | pde6d | 0.006969648 | -0.201449776 | -0.351770435 | -0.511138243 | -0.216254622 | downregulated during regeneration - mid |
| ENSDART00000114182 | fyco1b | 0.017131781 | 0.10137681 | -0.156409007 | -0.465781329 | -0.01243848 | downregulated during regeneration - mid |
| ENSDART00000114533 | mn1a | 0.064969544 | -0.109536494 | -0.207833595 | -0.685216226 | -0.20481073 | downregulated during regeneration - mid |
| ENSDART00000114705 | gprc5bb | 0.016311478 | -0.132340316 | -0.338695197 | -0.459834622 | -0.091865364 | downregulated during regeneration - mid |
| ENSDART00000114719 | frmpd1a | 0.0134351 | -0.29565145 | -0.338039747 | -0.498795038 | -0.059125706 | downregulated during regeneration - mid |
| ENSDART00000114746 | lrrc58a | 0.018924903 | -0.133537399 | 0.014548786 | -0.622798481 | -0.259773873 | downregulated during regeneration - mid |
| ENSDART00000115221 | adat2 | 0.007676315 | 0.066031062 | -1.495132242 | -2.125701316 | -0.144340549 | downregulated during regeneration - mid |
| ENSDART00000115224 | pvrl1a | 0.012357429 | -0.001527 | -0.239159739 | -0.512986688 | -0.121351808 | downregulated during regeneration - mid |
| ENSDART00000115343 | atp8b4 | 0.007152296 | -0.126263785 | -0.363733117 | -0.385610912 | -0.00148052 | downregulated during regeneration - mid |
| ENSDART00000115370 | mettl22 | 0.044172739 | -0.250044414 | -0.320129908 | -0.572576042 | -0.437533988 | downregulated during regeneration - mid |
| ENSDART00000121545 | brms1 | 0.048520479 | -0.198338884 | -0.120854433 | -0.325603911 | -0.103470514 | downregulated during regeneration - mid |
| ENSDART00000121598 | phf10 | 0.052326315 | -0.237213765 | -0.309795055 | -0.326418269 | -0.213703564 | downregulated during regeneration - mid |
| ENSDART00000121952 | h2afy2 | 0.039334382 | -0.217094069 | -0.401231455 | -0.429907398 | -0.128476741 | downregulated during regeneration - mid |
| ENSDART00000122059 | scoca | 0.051450895 | -0.009226259 | -0.04540809 | -0.349436614 | -0.115738429 | downregulated during regeneration - mid |
| ENSDART00000122716 | PDE1C | 0.014686352 | -0.224024171 | -0.483764938 | -0.500061121 | -0.40054702 | downregulated during regeneration - mid |
| ENSDART00000122929 | bbs4 | 0.023936488 | -0.14534671 | -0.308002271 | -0.496501405 | -0.252523436 | downregulated during regeneration - mid |
| ENSDART00000123040 | pfkpb | 0.018655612 | -0.26169445 | -0.241526904 | -0.415842652 | -0.206785479 | downregulated during regeneration - mid |
| ENSDART00000123360 | susd5 | 0.005867979 | -0.191753051 | -0.73208098 | -0.853142314 | -0.526763596 | downregulated during regeneration - mid |
| ENSDART00000123450 | KCNB2 (1 of many) | 0.014634638 | -0.271043423 | -0.632402343 | -0.828557597 | -0.168958818 | downregulated during regeneration - mid |
| ENSDART00000123505 | rab11fip4a | 0.004897476 | -0.154164113 | -0.404988879 | -0.479622862 | -0.179766383 | downregulated during regeneration - mid |
| ENSDART00000123820 | tmem135 | 0.049596963 | -0.104475369 | -0.143585965 | -0.370543783 | -0.122253976 | downregulated during regeneration - mid |
| ENSDART00000123970 | mntb | 0.011725523 | -0.115861458 | -0.101461538 | -0.360694649 | -0.01567692 | downregulated during regeneration - mid |
| ENSDART00000124140 | zgc:101566 | 0.000727221 | -0.242404826 | -0.290163127 | -0.636219456 | -0.416307006 | downregulated during regeneration - mid |
| ENSDART00000124290 | MFN1 | 0.012467303 | -0.108610875 | -0.185612855 | -0.471855683 | -0.083062873 | downregulated during regeneration - mid |
| ENSDART00000124329 | cbx8a | 0.01903005 | -0.13552578 | -0.23364206 | -0.31740894 | -0.19855692 | downregulated during regeneration - mid |
| ENSDART00000124335 | smim4 | 0.017820373 | -0.115055175 | -0.371982424 | -0.338416379 | -0.248500803 | downregulated during regeneration - mid |
| ENSDART00000124562 | zgc:152977 | 0.011844097 | -0.424781995 | -0.464146326 | -0.707134675 | -0.367623104 | downregulated during regeneration - mid |
| ENSDART00000124676 | sv2ba | 0.018790846 | -0.120324523 | -0.401787287 | -0.561843381 | -0.191682717 | downregulated during regeneration - mid |
| ENSDART00000124945 | ubap1lb | 0.007333969 | 0.21207026 | -0.004652141 | -0.773302877 | -0.344881822 | downregulated during regeneration - mid |
| ENSDART00000125299 | plk2a | 0.086996216 | -0.143584568 | -0.170521478 | -0.297471862 | -0.183865098 | downregulated during regeneration - mid |
| ENSDART00000125397 | kri1 | 0.069160004 | -0.318299605 | -0.278223015 | -0.439932558 | -0.332258967 | downregulated during regeneration - mid |
| ENSDART00000125466 | alpi.2 | 0.052273162 | -0.05537717 | -0.356245119 | -0.658920603 | -0.289506352 | downregulated during regeneration - mid |
| ENSDART00000125561 | atp6ap2 | 0.008210772 | -0.203601896 | -0.343976801 | -0.371684174 | -0.265876436 | downregulated during regeneration - mid |
| ENSDART00000125864 | camkk1b | 0.039389579 | -0.169636507 | -0.512096721 | -0.591294824 | -0.14296099 | downregulated during regeneration - mid |
| ENSDART00000125979 | tshz3a | 0.02052819 | -0.321412605 | -0.323547538 | -0.394821529 | -0.291421679 | downregulated during regeneration - mid |
| ENSDART00000126076 | NA | 0.012675019 | -0.016912473 | -0.350302068 | -0.401962349 | -0.001679825 | downregulated during regeneration - mid |
| ENSDART00000126084 | ctdspl3 | 0.009272964 | -0.310396194 | -0.278185547 | -0.408791973 | -0.234117517 | downregulated during regeneration - mid |
| ENSDART00000126282 | nr1d1 | 0.008088213 | -0.1014969 | -0.191656676 | -0.494430716 | -0.13667305 | downregulated during regeneration - mid |
| ENSDART00000126661 | rhobtb1 | 0.007785924 | -0.165202139 | -0.331565823 | -0.522361525 | -0.299461153 | downregulated during regeneration - mid |
| ENSDART00000126842 | flj11011l | 0.051190638 | -0.120546184 | -0.161649138 | -0.295902143 | -0.182318736 | downregulated during regeneration - mid |
| ENSDART00000126916 | mob2b | 0.004538993 | 0.26836075 | -0.004670217 | -0.398977994 | -0.072830738 | downregulated during regeneration - mid |
| ENSDART00000126928 | si:dkey-73n8.3 | 0.012465298 | -0.354837143 | 0.244926012 | -3.843926683 | -0.176933271 | downregulated during regeneration - mid |
| ENSDART00000127131 | mtmr10 | 0.016195506 | 0.05904867 | -0.240178 | -0.607174209 | -0.120957306 | downregulated during regeneration - mid |

| | | | | | | | |
|---|---|---|---|---|---|---|---|
| ENSDART00000127216 | fam161a | 0.056105466 | -0.082566753 | -0.243407902 | -0.44855597 | -0.155296925 | downregulated during regeneration - mid |
| ENSDART00000127274 | sema7a | 0.056867894 | -0.198678021 | -0.353988759 | -0.478218296 | -0.427230313 | downregulated during regeneration - mid |
| ENSDART00000127331 | cdc37l1 | 0.022102317 | -0.115815783 | -0.102908381 | -0.388723888 | -0.245749713 | downregulated during regeneration - mid |
| ENSDART00000127398 | CR753886.1 | 0.031973637 | -0.159141993 | -0.332294072 | -0.331878243 | -0.244595408 | downregulated during regeneration - mid |
| ENSDART00000127568 | ankha | 0.014386448 | -0.068226585 | -0.359480026 | -0.686453521 | -0.222767571 | downregulated during regeneration - mid |
| ENSDART00000127656 | cln6b | 0.003337702 | -0.363769389 | -0.208045101 | -0.481848131 | -0.19203643 | downregulated during regeneration - mid |
| ENSDART00000128696 | CABZ01035108.1 | 0.038217845 | -0.30766878 | -0.382099096 | -0.769557831 | -0.298631085 | downregulated during regeneration - mid |
| ENSDART00000128698 | rnf34b | 0.019215781 | -0.122706526 | -0.390373676 | -0.658840165 | -0.257572429 | downregulated during regeneration - mid |
| ENSDART00000128721 | rcvrn3 | 0.023143967 | -0.045382046 | -0.143627209 | -0.579607896 | -0.393091134 | downregulated during regeneration - mid |
| ENSDART00000128722 | RGS9BP | 0.039271588 | -0.328845343 | -0.481472863 | -0.560905879 | -0.19840165 | downregulated during regeneration - mid |
| ENSDART00000128859 | prlra | 0.029917459 | -0.153277376 | -0.232075961 | -0.477332275 | -0.11182176 | downregulated during regeneration - mid |
| ENSDART00000128975 | slc25a32a | 0.003209414 | 0.048183005 | -0.130483176 | -0.493409396 | -0.216291506 | downregulated during regeneration - mid |
| ENSDART00000129156 | adcy7 | 0.012672965 | -0.081832737 | -0.232314587 | -0.506962474 | -0.282923342 | downregulated during regeneration - mid |
| ENSDART00000129308 | ACVR1C | 0.02188897 | -0.083617591 | -0.230914728 | -0.630759756 | -0.255735924 | downregulated during regeneration - mid |
| ENSDART00000129751 | si:ch211-105f12.2 | 0.063999698 | -0.375204291 | -0.254543259 | -0.612224773 | -0.289815804 | downregulated during regeneration - mid |
| ENSDART00000130093 | c10h21orf59 | 0.010893084 | -0.163395921 | -0.484168269 | -0.87971069 | -0.649928123 | downregulated during regeneration - mid |
| ENSDART00000130142 | ube2w | 0.018144005 | 0.018587872 | -0.041621436 | -0.37919808 | -0.07627037 | downregulated during regeneration - mid |
| ENSDART00000130366 | si:dkey-21c1.4 | 0.040026502 | -0.116466944 | -0.239339578 | -0.392905857 | -0.311667635 | downregulated during regeneration - mid |
| ENSDART00000130537 | spag1a | 0.016180105 | -0.063362594 | -0.300759627 | -0.420912866 | -0.152669564 | downregulated during regeneration - mid |
| ENSDART00000130818 | ptprz1b | 0.007396212 | -0.275892935 | -0.352115142 | -0.541325565 | -0.202924453 | downregulated during regeneration - mid |
| ENSDART00000131206 | 43160 | 0.02794941 | -0.087431315 | -0.336892252 | -0.48450384 | -0.18585378 | downregulated during regeneration - mid |
| ENSDART00000131286 | erlin2 | 0.045086702 | -0.000462383 | -0.183644452 | -0.434826094 | -0.105981059 | downregulated during regeneration - mid |
| ENSDART00000131349 | mob2b | 0.003887497 | 0.337219743 | -0.092777728 | -0.774689204 | -0.397735432 | downregulated during regeneration - mid |
| ENSDART00000131517 | si:ch211-232m10.6 | 0.039816981 | -0.306074293 | -0.295884968 | -0.441790198 | -0.281067097 | downregulated during regeneration - mid |
| ENSDART00000131627 | kcnn3 | 0.030198325 | -0.085165392 | -0.23467025 | -0.387311243 | -0.108636046 | downregulated during regeneration - mid |
| ENSDART00000131848 | BX004774.2 | 0.004739684 | 0.065629354 | -0.435237015 | -1.040450944 | -0.818745042 | downregulated during regeneration - mid |
| ENSDART00000131973 | lrmp | 0.011245854 | -0.070199137 | -0.469517154 | -1.098854311 | -0.136597848 | downregulated during regeneration - mid |
| ENSDART00000132591 | si:ch211-191i18.4 | 0.016122246 | 0.099974449 | -0.068457558 | -0.479286849 | -0.192940747 | downregulated during regeneration - mid |
| ENSDART00000132744 | ppcdc | 0.004289396 | 0.019198695 | -0.177358944 | -0.471740583 | -0.31681582 | downregulated during regeneration - mid |
| ENSDART00000133035 | syt5a | 0.045873551 | -0.188232763 | -0.271065673 | -0.434073976 | -0.186984299 | downregulated during regeneration - mid |
| ENSDART00000133039 | tac3a | 0.010923556 | -0.2410584 | -0.489343566 | -0.63037654 | -0.413025711 | downregulated during regeneration - mid |
| ENSDART00000133220 | CR788254.2 | 0.033633212 | -0.013907422 | -0.231082485 | -0.739605077 | -0.487191429 | downregulated during regeneration - mid |
| ENSDART00000133242 | syne2b | 0.0092524 | 0.14436825 | -0.456480635 | -0.844135801 | -0.206154738 | downregulated during regeneration - mid |
| ENSDART00000133583 | si:ch211-117m20.4 | 0.037722848 | -0.073043862 | -0.13648785 | -0.437013406 | -0.146443055 | downregulated during regeneration - mid |
| ENSDART00000133613 | AL954694.1 | 0.044914837 | -0.189304139 | -0.336191594 | -0.52116431 | -0.341767633 | downregulated during regeneration - mid |
| ENSDART00000133661 | tasp1 | 0.035967167 | -0.129454637 | -0.177592146 | -0.299309238 | -0.129141605 | downregulated during regeneration - mid |
| ENSDART00000133867 | arfip2a | 0.037668797 | -0.237325336 | -0.320204428 | -0.424850477 | -0.153128941 | downregulated during regeneration - mid |
| ENSDART00000133869 | kcnk1a | 0.066233336 | -0.389144165 | -0.38978007 | -0.667699511 | -0.286776021 | downregulated during regeneration - mid |
| ENSDART00000134190 | sort1b | 0.011533823 | -0.233086943 | -0.116629656 | -0.586941063 | -0.191255985 | downregulated during regeneration - mid |
| ENSDART00000134399 | slkb | 0.002060335 | -0.338857379 | -0.632619838 | -0.770465733 | -0.089077643 | downregulated during regeneration - mid |
| ENSDART00000134697 | pik3ip1 | 0.001209453 | -0.252912627 | -0.195771257 | -0.573002676 | -0.212029641 | downregulated during regeneration - mid |
| ENSDART00000134773 | zfand5a | 0.01494686 | -0.170709233 | -0.221425066 | -0.980962271 | -0.517481995 | downregulated during regeneration - mid |
| ENSDART00000134864 | gigyf1b | 0.0256797 | -0.032906085 | -0.141419247 | -0.36943011 | -0.170431533 | downregulated during regeneration - mid |
| ENSDART00000135118 | efr3ba | 0.060997845 | -0.168363836 | -0.303586815 | -0.356914459 | -0.264543553 | downregulated during regeneration - mid |
| ENSDART00000135374 | ptpn21 | 0.062431525 | -0.060719081 | -0.163907746 | -0.321158981 | -0.096688483 | downregulated during regeneration - mid |
| ENSDART00000135436 | jtb | 0.036479303 | -0.083914092 | -0.13827795 | -0.410566521 | -0.208466854 | downregulated during regeneration - mid |
| ENSDART00000135479 | fstl5 | 0.046057741 | -0.187703631 | -0.312045142 | -0.413295205 | -0.11029682 | downregulated during regeneration - mid |
| ENSDART00000135510 | syne2b | 0.002913561 | -0.011968335 | -0.47360345 | -0.918119469 | -0.289310189 | downregulated during regeneration - mid |
| ENSDART00000135690 | lactbl1a | 0.016724084 | -0.207283355 | -0.397072585 | -0.513557165 | -0.348206521 | downregulated during regeneration - mid |
| ENSDART00000135738 | zgc:194990 | 0.007620644 | 0.067005172 | -0.387370648 | -0.731552205 | -0.192679748 | downregulated during regeneration - mid |
| ENSDART00000135860 | rnf41l | 0.023081255 | -0.353446179 | -0.473576771 | -0.616364395 | -0.289592085 | downregulated during regeneration - mid |
| ENSDART00000135919 | amd1 | 0.013040781 | -0.131714213 | -0.276160556 | -0.537901531 | -0.163925689 | downregulated during regeneration - mid |
| ENSDART00000136208 | ncam1b | 0.026842776 | -0.292790209 | -0.308372802 | -0.421218735 | -0.19466435 | downregulated during regeneration - mid |
| ENSDART00000136647 | ext1c | 0.014963248 | -0.283149695 | -0.275883116 | -0.465218527 | -0.113484828 | downregulated during regeneration - mid |
| ENSDART00000136770 | si:dkey-246e1.3 | 0.017642395 | -0.264481758 | -0.308595158 | -0.554881445 | -0.406535962 | downregulated during regeneration - mid |
| ENSDART00000136771 | dnajc5ga | 0.026478876 | -0.246041931 | -0.267357646 | -0.352933326 | -0.221144382 | downregulated during regeneration - mid |
| ENSDART00000137047 | scoca | 0.043980855 | -0.178388629 | -0.216665672 | -0.49140437 | -0.218641566 | downregulated during regeneration - mid |
| ENSDART00000137185 | cnga3b | 0.013354221 | -0.145832564 | -0.301563517 | -1.136000666 | -0.708190865 | downregulated during regeneration - mid |
| ENSDART00000137236 | hpcal1 | 0.014842483 | -0.065446201 | -0.487196869 | -0.639863396 | -0.431847778 | downregulated during regeneration - mid |
| ENSDART00000137292 | tmem106bb | 0.004741095 | -0.265825137 | -0.29210537 | -0.411370487 | -0.25270276 | downregulated during regeneration - mid |
| ENSDART00000137364 | si:dkey-33c12.3 | 0.0001406 | 0.174067673 | -0.534549963 | -0.496919129 | 0.189169246 | downregulated during regeneration - mid |
| ENSDART00000137633 | si:ch211-194e15.5 | 0.009667225 | -0.00297372 | -0.215384681 | -0.930894546 | -0.593142383 | downregulated during regeneration - mid |
| ENSDART00000137817 | rap1gap2a | 0.029185454 | -0.351211604 | -0.330279371 | -0.569882036 | -0.258477103 | downregulated during regeneration - mid |
| ENSDART00000137896 | fibpb | 0.035286858 | -0.042394017 | -0.144008268 | -0.458312875 | -0.18906893 | downregulated during regeneration - mid |
| ENSDART00000137899 | tmem9 | 0.056822582 | -0.365732748 | -0.460492017 | -0.592427521 | -0.282793654 | downregulated during regeneration - mid |
| ENSDART00000138232 | arl13a | 0.018790846 | -0.123428499 | -0.297389057 | -0.827109894 | -0.393133456 | downregulated during regeneration - mid |
| ENSDART00000138270 | copz2 | 0.022463985 | -0.133761169 | -0.127273242 | -0.35316272 | -0.169760508 | downregulated during regeneration - mid |
| ENSDART00000138541 | rpgrip1 | 0.043421442 | -0.096214383 | -0.271667846 | -0.617065899 | -0.286110235 | downregulated during regeneration - mid |
| ENSDART00000138658 | si:ch211-117m20.4 | 0.082561098 | -0.301116828 | -0.255854166 | -0.65213458 | -0.329202684 | downregulated during regeneration - mid |
| ENSDART00000138793 | si:ch211-203k16.3 | 0.033480582 | -0.114803625 | -0.341143223 | -0.894722004 | -0.064858026 | downregulated during regeneration - mid |
| ENSDART00000138866 | pbxip1a | 0.000760227 | -0.45481847 | -0.358362517 | -0.587330976 | -0.331438505 | downregulated during regeneration - mid |
| ENSDART00000138922 | elovl8a | 0.097148471 | -0.299886352 | -0.255106336 | -0.59316691 | -0.339691401 | downregulated during regeneration - mid |
| ENSDART00000139042 | si:ch211-132b12.7 | 0.002274284 | 0.072730857 | -0.322841214 | -0.765244522 | -0.309126323 | downregulated during regeneration - mid |
| ENSDART00000139102 | dbpb | 0.001993937 | 0.113164671 | -0.364559303 | -0.423196921 | -0.137052116 | downregulated during regeneration - mid |
| ENSDART00000139310 | bcat2 | 0.032825581 | -0.180902608 | -0.445596686 | -0.579108461 | -0.343900677 | downregulated during regeneration - mid |

| | | | | | | | |
|---|---|---|---|---|---|---|---|
| ENSDART00000139461 | spock3 | 0.043085704 | -0.122253969 | -0.220033244 | -0.389207603 | -0.090203867 | downregulated during regeneration - mid |
| ENSDART00000139477 | trim110 | 0.030645263 | -0.21821102 | -0.289321848 | -0.373728133 | -0.159590708 | downregulated during regeneration - mid |
| ENSDART00000139715 | dpm3 | 0.028357685 | -0.11194386 | -0.125527265 | -0.41579896 | -0.188378465 | downregulated during regeneration - mid |
| ENSDART00000139900 | diaph3 | 0.028408144 | 0.052388836 | -0.258521684 | -0.529199016 | -0.186043701 | downregulated during regeneration - mid |
| ENSDART00000140145 | ajap1 | 0.019585067 | -0.237572949 | -0.464416488 | -0.62912768 | -0.262821389 | downregulated during regeneration - mid |
| ENSDART00000140375 | zfyve27 | 0.02776376 | -0.214217108 | -0.191210439 | -0.353015645 | -0.21311838 | downregulated during regeneration - mid |
| ENSDART00000140553 | frmpd1a | 0.012556905 | -0.279082186 | -0.31970424 | -0.46726564 | -0.019879501 | downregulated during regeneration - mid |
| ENSDART00000141031 | BX004774.2 | 0.016675166 | -0.096197487 | -0.455215086 | -0.947274727 | -0.721728098 | downregulated during regeneration - mid |
| ENSDART00000141157 | ablim3 | 0.029525765 | -0.090063728 | -0.09247473 | -0.384522771 | 0.048083962 | downregulated during regeneration - mid |
| ENSDART00000141340 | si:dkey-183n20.15 | 0.02850112 | -0.434296772 | -0.549401302 | -1.33032419 | -0.748989345 | downregulated during regeneration - mid |
| ENSDART00000141493 | ANO2 (1 of many) | 0.006900346 | 0.056182997 | -0.29662332 | -0.7863422 | -0.307277237 | downregulated during regeneration - mid |
| ENSDART00000141734 | hivep2a | 0.017814842 | -0.218148846 | -0.337091836 | -0.481607769 | -0.327040294 | downregulated during regeneration - mid |
| ENSDART00000142157 | arl15a | 0.011135896 | -0.037401347 | -0.185890543 | -0.49114445 | -0.348659966 | downregulated during regeneration - mid |
| ENSDART00000142454 | si:dkey-121a11.3 | 0.046134939 | -0.218109071 | -0.329830559 | -0.334040533 | -0.225452629 | downregulated during regeneration - mid |
| ENSDART00000142543 | si:dkey-206f10.1 | 0.009178848 | -0.205833854 | -0.070068511 | -0.826534181 | -0.287245824 | downregulated during regeneration - mid |
| ENSDART00000142605 | sorbs2b | 0.003262292 | -0.090898219 | -0.353328393 | -0.576838002 | -0.278787769 | downregulated during regeneration - mid |
| ENSDART00000142731 | rce1b | 0.030361713 | -0.103631801 | -0.326325127 | -0.391135117 | -0.061782364 | downregulated during regeneration - mid |
| ENSDART00000143165 | tsc22d1 | 0.012984014 | -0.098364585 | -0.21923408 | -0.39849137 | 0.017570634 | downregulated during regeneration - mid |
| ENSDART00000143199 | wasf3a | 0.033862985 | -0.216537889 | -0.280788628 | -0.398676762 | -0.269991874 | downregulated during regeneration - mid |
| ENSDART00000143245 | nptnb | 0.00231226 | -0.0813348 | -0.106325128 | -0.477835951 | -0.112902967 | downregulated during regeneration - mid |
| ENSDART00000143503 | usp21 | 0.010231208 | -0.183997919 | -0.332937745 | -0.642449016 | -0.356635014 | downregulated during regeneration - mid |
| ENSDART00000143625 | map7d2b | 0.042433381 | 0.025364857 | -1.606530561 | -4.429840485 | -0.88152256 | downregulated during regeneration - mid |
| ENSDART00000143662 | ankrd33aa | 0.01899428 | -0.118189532 | -0.449119978 | -0.599045615 | 0.004982045 | downregulated during regeneration - mid |
| ENSDART00000143829 | nxnl1 | 0.034874709 | -0.184554354 | -0.325799787 | -0.486032926 | -0.185522737 | downregulated during regeneration - mid |
| ENSDART00000143874 | akna | 0.021937256 | -0.124615674 | -0.323808831 | -0.482322372 | -0.23639171 | downregulated during regeneration - mid |
| ENSDART00000144592 | sik2a | 0.005427451 | -0.05071398 | -0.135507163 | -0.424760133 | 0.101311014 | downregulated during regeneration - mid |
| ENSDART00000144802 | dfnb31b | 0.012605558 | -0.206535765 | -0.346317212 | -0.484783097 | -0.018383317 | downregulated during regeneration - mid |
| ENSDART00000144804 | mxra7 | 0.020724346 | -0.254910086 | -0.321688262 | -0.500299705 | -0.323281006 | downregulated during regeneration - mid |
| ENSDART00000144890 | bokb | 0.052287778 | -0.312629523 | -0.278040732 | -0.730243904 | -0.348692502 | downregulated during regeneration - mid |
| ENSDART00000145035 | saga | 0.025257904 | -0.094875139 | -0.251791894 | -0.561974573 | -0.363496236 | downregulated during regeneration - mid |
| ENSDART00000145258 | glsb | 0.012086831 | -0.010345949 | -0.147193332 | -0.750348982 | -0.227873062 | downregulated during regeneration - mid |
| ENSDART00000145545 | dclk1a | 0.028108682 | -0.21717792 | -0.314020567 | -0.407893866 | -0.042242501 | downregulated during regeneration - mid |
| ENSDART00000145728 | sdhc | 0.056867894 | -0.035025769 | -0.113602993 | -0.325388824 | -0.192371035 | downregulated during regeneration - mid |
| ENSDART00000145834 | si:ch211-233a24.2 | 0.017986602 | -0.054480296 | -0.065552868 | -0.373189 | -0.049333125 | downregulated during regeneration - mid |
| ENSDART00000145894 | zgc:195245 | 0.033020938 | -0.325743953 | -0.471221888 | -0.661613866 | -0.246097731 | downregulated during regeneration - mid |
| ENSDART00000146008 | camkvl | 0.050814856 | -0.251523364 | -0.299024254 | -0.462517021 | -0.14005057 | downregulated during regeneration - mid |
| ENSDART00000146962 | STX3 (1 of many) | 0.041930419 | -0.081644241 | -0.246049482 | -0.328837917 | -0.149895857 | downregulated during regeneration - mid |
| ENSDART00000147368 | osbp2 | 0.027388792 | -0.137018463 | -0.22987814 | -0.404444797 | -0.125629393 | downregulated during regeneration - mid |
| ENSDART00000147422 | arhgap23b | 0.009929644 | -0.106264079 | -0.33507693 | -0.670558797 | -0.201207875 | downregulated during regeneration - mid |
| ENSDART00000147502 | grhprb | 0.004106215 | 0.278026338 | -0.217638113 | -0.512645749 | -0.092734666 | downregulated during regeneration - mid |
| ENSDART00000147504 | bzw2 | 0.002177213 | 0.031120244 | -0.513458671 | -0.643056049 | -0.325765553 | downregulated during regeneration - mid |
| ENSDART00000147884 | syne2b | 0.0066372 | 0.046598878 | -0.495566729 | -1.074400394 | -0.335549303 | downregulated during regeneration - mid |
| ENSDART00000147903 | lrit1a | 0.028192085 | -0.179060864 | -0.15522738 | -0.479781004 | -0.346169989 | downregulated during regeneration - mid |
| ENSDART00000148039 | snx19a | 0.037712225 | -0.086070054 | -0.238060141 | -0.431834798 | -0.166151211 | downregulated during regeneration - mid |
| ENSDART00000148066 | znf395b | 0.020342933 | -0.110739145 | -0.366906743 | -0.460042969 | -0.183105049 | downregulated during regeneration - mid |
| ENSDART00000148106 | mef2aa | 0.026582721 | -0.081330082 | -0.55524128 | -0.609035118 | -0.185252531 | downregulated during regeneration - mid |
| ENSDART00000148175 | si:ch1073-440b2.1 | 0.004197991 | -0.235967394 | -0.243649374 | -0.449710659 | -0.316169237 | downregulated during regeneration - mid |
| ENSDART00000148246 | pole4 | 0.028322957 | -0.080211224 | -0.333184545 | -0.567162517 | -0.350297708 | downregulated during regeneration - mid |
| ENSDART00000148353 | usf2 | 0.006964046 | -0.107987138 | -0.265954127 | -0.405201274 | -0.200197809 | downregulated during regeneration - mid |
| ENSDART00000148536 | GK3P | 0.017775184 | -0.308304163 | -0.269790809 | -0.586409039 | -0.39358576 | downregulated during regeneration - mid |
| ENSDART00000148537 | roraa | 0.01612441 | -0.160828233 | -0.221872095 | -0.45696721 | -0.083100906 | downregulated during regeneration - mid |
| ENSDART00000148639 | ubxn2a | 0.015578607 | -0.292754646 | -0.267419804 | -0.368360861 | -0.228619688 | downregulated during regeneration - mid |
| ENSDART00000148831 | fgf11b | 0.048355827 | -0.269342716 | -0.341549392 | -0.474527478 | -0.280873475 | downregulated during regeneration - mid |
| ENSDART00000148958 | fam150a | 0.006385907 | -0.081185595 | -0.392189222 | -0.650641125 | -0.329645149 | downregulated during regeneration - mid |
| ENSDART00000148982 | bin1b | 0.017473198 | 0.009431427 | -0.214307092 | -0.49686004 | -0.050411548 | downregulated during regeneration - mid |
| ENSDART00000149029 | sagb | 0.030697622 | -0.046274749 | -0.449576755 | -0.950069775 | -0.043925247 | downregulated during regeneration - mid |
| ENSDART00000149339 | si:ch211-175f12.2 | 0.046869499 | -0.297380973 | -0.332932663 | -0.389357991 | -0.204952824 | downregulated during regeneration - mid |
| ENSDART00000149352 | si:ch211-163l21.11 | 0.045582714 | -0.266781177 | -0.351609906 | -0.560319486 | -0.211979525 | downregulated during regeneration - mid |
| ENSDART00000149429 | boka | 0.007908312 | -0.220682727 | -0.305502243 | -0.692638969 | -0.393597854 | downregulated during regeneration - mid |
| ENSDART00000149512 | CU467655.1 | 0.058548676 | -0.320227651 | -0.180971804 | -0.440965145 | -0.207148298 | downregulated during regeneration - mid |
| ENSDART00000149634 | atp6ap2 | 0.075143936 | -0.108949021 | -0.143807338 | -0.457436101 | -0.199272375 | downregulated during regeneration - mid |
| ENSDART00000149935 | nt5c3a | 0.022408352 | 0.05115392 | -0.273408486 | -0.497911915 | -0.317644766 | downregulated during regeneration - mid |
| ENSDART00000150045 | adamts18 | 0.038201053 | -0.187059316 | -0.292648437 | -0.628764119 | 0.030985495 | downregulated during regeneration - mid |
| ENSDART00000150789 | USMG5 | 0.058128769 | -0.285153636 | -0.258859464 | -0.457141251 | -0.093210696 | downregulated during regeneration - mid |
| ENSDART00000151013 | ensaa | 0.001153184 | -0.338295629 | -0.414678183 | -0.638369138 | -0.341312384 | downregulated during regeneration - mid |
| ENSDART00000151022 | si:ch211-191i18.4 | 0.046753078 | -0.318627462 | -0.467051418 | -0.723047751 | -0.473341501 | downregulated during regeneration - mid |
| ENSDART00000151203 | rgs9a | 0.049667416 | -0.190499153 | -0.256482975 | -0.462262315 | -0.086082277 | downregulated during regeneration - mid |
| ENSDART00000151386 | CU659670.1 | 0.044578949 | -0.061309281 | -0.197320939 | -0.510180812 | -0.129307388 | downregulated during regeneration - mid |
| ENSDART00000151419 | tgfa | 0.049309412 | -0.147139604 | -0.1316737 | -0.368157435 | -0.244071655 | downregulated during regeneration - mid |
| ENSDART00000151670 | fbxl16 | 0.055597767 | -0.100117774 | -0.155204519 | -0.627708744 | -0.129502503 | downregulated during regeneration - mid |
| ENSDART00000151808 | rcvrn3 | 0.013770716 | -0.028816037 | -0.156203513 | -0.580402071 | -0.415774036 | downregulated during regeneration - mid |
| ENSDART00000152005 | ppp1r13ba | 0.024276272 | -0.032736325 | -0.117362615 | -0.454836475 | -0.009028338 | downregulated during regeneration - mid |
| ENSDART00000152172 | BX248418.1 | 0.005026826 | -0.267252802 | -0.476860817 | -0.659952516 | -0.030823789 | downregulated during regeneration - mid |
| ENSDART00000152689 | oplah | 0.01992766 | -0.018001137 | -0.164313071 | -0.461321001 | -0.221934453 | downregulated during regeneration - mid |
| ENSDART00000152768 | si:ch211-207i1.2 | 0.036445549 | -0.201357893 | -0.51029626 | -0.665018804 | -0.121583536 | downregulated during regeneration - mid |

| | | | | | | | |
|---|---|---|---|---|---|---|---|
| ENSDART00000152968 | si:dkey-21c1.1 | 0.005983812 | 0.134479312 | -0.180316529 | -0.580606373 | -0.202289334 | downregulated during regeneration - mid |
| ENSDART00000153086 | kctd17 | 0.063822766 | -0.109500243 | -0.169719356 | -0.312398256 | -0.16098927 | downregulated during regeneration - mid |
| ENSDART00000153124 | CU570682.1 | 0.019075692 | -0.274154453 | -0.462025745 | -0.49154679 | -0.26836126 | downregulated during regeneration - mid |
| ENSDART00000153190 | vash2 | 0.019554708 | -0.078136066 | -0.226923996 | -0.395295378 | -0.160874636 | downregulated during regeneration - mid |
| ENSDART00000153442 | rxfp2a | 0.01711161 | -0.191591927 | -0.398184153 | -0.707111514 | -0.312025768 | downregulated during regeneration - mid |
| ENSDART00000153773 | cipca | 0.004341144 | -0.044231438 | -0.31334692 | -0.44667945 | -0.17971329 | downregulated during regeneration - mid |
| ENSDART00000154085 | gipr | 0.032143116 | -0.077819916 | -0.236822644 | -0.445528895 | -0.377678023 | downregulated during regeneration - mid |
| ENSDART00000154627 | eogt | 0.014130752 | -0.136366673 | -0.382886458 | -0.560103122 | -0.203176718 | downregulated during regeneration - mid |
| ENSDART00000154917 | ciartb | 0.004147933 | -0.07665894 | -0.17887759 | -0.516654864 | -0.219843882 | downregulated during regeneration - mid |
| ENSDART00000155078 | CU466278.1 | 0.067207866 | -0.496260032 | -0.275828398 | -0.937323619 | -0.494833793 | downregulated during regeneration - mid |
| ENSDART00000155088 | ankrd33ab | 0.019678793 | -0.091123111 | -0.280595773 | -2.122583018 | -0.908625438 | downregulated during regeneration - mid |
| ENSDART00000155324 | pdpk1a | 0.051252746 | -0.082713691 | -0.135052421 | -0.317218269 | -0.180842854 | downregulated during regeneration - mid |
| ENSDART00000155458 | ubap1la | 0.036933769 | -0.102679722 | -0.709773263 | -0.512828077 | -0.333708251 | downregulated during regeneration - mid |
| ENSDART00000155580 | ccdc32 | 0.028277545 | -0.137851793 | -0.182759184 | -0.331717766 | -0.143672128 | downregulated during regeneration - mid |
| ENSDART00000155839 | CR759918.1 | 0.036715052 | -0.263859971 | -0.262088445 | -0.379170984 | -0.118091172 | downregulated during regeneration - mid |
| ENSDART00000155859 | larp4ab | 0.007335615 | -0.193577441 | -0.330790279 | -0.534105504 | -0.201385609 | downregulated during regeneration - mid |
| ENSDART00000155956 | ssx2ipa | 0.040205407 | -0.036908945 | -0.252732501 | -0.388773092 | -0.156731785 | downregulated during regeneration - mid |
| ENSDART00000156008 | adcy1b | 0.021755167 | -0.126277791 | -0.148680925 | -0.963041725 | -0.495906031 | downregulated during regeneration - mid |
| ENSDART00000156051 | dhx32b | 0.010222513 | -0.034036848 | -0.386771049 | -0.600165189 | -0.131769991 | downregulated during regeneration - mid |
| ENSDART00000156169 | ube2o | 0.055683104 | -0.089939623 | -0.17956897 | -0.346977584 | -0.218736801 | downregulated during regeneration - mid |
| ENSDART00000156179 | sema7a | 0.031092135 | -0.273960356 | -0.559537035 | -0.479585038 | -0.478067901 | downregulated during regeneration - mid |
| ENSDART00000156211 | adcyap1r1b | 0.029377262 | -0.120918148 | -0.186017 | -0.540439336 | -0.277725276 | downregulated during regeneration - mid |
| ENSDART00000156247 | CU468915.1 | 0.04446722 | -0.147323171 | -0.162953088 | -0.662599642 | -0.420048082 | downregulated during regeneration - mid |
| ENSDART00000156365 | BX248121.1 | 0.044459756 | -0.069755444 | -0.2828198 | -0.35442143 | -0.173325265 | downregulated during regeneration - mid |
| ENSDART00000156438 | tmem138 | 0.003538125 | -0.131418974 | -0.291816206 | -0.603920899 | -0.471900931 | downregulated during regeneration - mid |
| ENSDART00000156492 | tiam2a | 0.033593902 | -0.024981814 | -0.236489206 | -0.397107093 | -0.120407246 | downregulated during regeneration - mid |
| ENSDART00000156527 | crhr1 | 0.045012134 | -0.121975919 | -0.246139891 | -0.376184792 | -0.120567191 | downregulated during regeneration - mid |
| ENSDART00000156568 | CR387996.1 | 0.023717938 | -0.565799723 | -0.534585499 | -0.855400153 | -0.72349809 | downregulated during regeneration - mid |
| ENSDART00000156757 | CU633486.1 | 0.075604115 | -0.302968456 | -0.206059 | -0.531920994 | -0.165463059 | downregulated during regeneration - mid |
| ENSDART00000156831 | si:dkeyp-47f9.4 | 0.018916133 | -0.215799622 | -0.290204625 | -0.428110244 | -0.249285093 | downregulated during regeneration - mid |
| ENSDART00000156918 | si:dkey-16p21.8 | 0.016886751 | -0.278111877 | -0.271868386 | -0.501824009 | -0.172182623 | downregulated during regeneration - mid |
| ENSDART00000156967 | itcha | 0.025939466 | -0.048958922 | -0.233534262 | -0.334119802 | -0.057332431 | downregulated during regeneration - mid |
| ENSDART00000157066 | ppp1r16b | 0.028214992 | -0.047627869 | -0.115324195 | -0.408403631 | -0.014573035 | downregulated during regeneration - mid |
| ENSDART00000157437 | park2 | 0.017079861 | -0.135617767 | -0.306295495 | -0.445890787 | -0.195890902 | downregulated during regeneration - mid |
| ENSDART00000157515 | FO704641.1 | 0.005096692 | -0.135830256 | -0.41280018 | -0.451790561 | -0.015210245 | downregulated during regeneration - mid |
| ENSDART00000157570 | asmt | 0.002360359 | -0.467907647 | -0.512776687 | -0.801096178 | -0.257600065 | downregulated during regeneration - mid |
| ENSDART00000157755 | cds1 | 0.00817514 | -0.167833858 | -0.281185983 | -0.514312915 | -0.176231692 | downregulated during regeneration - mid |
| ENSDART00000157979 | plppr5b | 0.003299709 | -0.141986763 | -0.474240998 | -0.437275413 | -0.217953336 | downregulated during regeneration - mid |
| ENSDART00000158055 | kcnh6a | 0.064248443 | -0.288242124 | -0.323669014 | -0.559331748 | -0.259833446 | downregulated during regeneration - mid |
| ENSDART00000158077 | arr3b | 0.063631477 | -0.13602706 | -0.148118881 | -0.527427286 | -0.215489656 | downregulated during regeneration - mid |
| ENSDART00000158202 | CABZ01078449.1 | 0.039683894 | -0.388078647 | -0.185889564 | -2.923902226 | 0.109460724 | downregulated during regeneration - mid |
| ENSDART00000158289 | taok1b | 0.039389579 | -0.136932712 | -0.168799291 | -0.390684145 | -0.122927658 | downregulated during regeneration - mid |
| ENSDART00000158301 | CR749168.4 | 0.010360832 | -0.284512048 | -0.514337485 | -0.519239658 | -0.283197555 | downregulated during regeneration - mid |
| ENSDART00000158338 | CNNM1 | 0.001487078 | -0.362878072 | -0.490740729 | -0.646651234 | -0.29213891 | downregulated during regeneration - mid |
| ENSDART00000158395 | dclk2a | 0.01163651 | -0.152411223 | -0.291700953 | -0.710292221 | -0.218291454 | downregulated during regeneration - mid |
| ENSDART00000158440 | arhgap12b | 0.041157695 | -0.039244537 | -0.138433678 | -0.355110347 | -0.091493676 | downregulated during regeneration - mid |
| ENSDART00000158587 | casd1 | 0.047492859 | -0.21622109 | -0.133405356 | -0.572868553 | -0.255662771 | downregulated during regeneration - mid |
| ENSDART00000158692 | CABZ01112401.1 | 0.000824931 | 0.113542035 | -0.344191992 | -0.461207823 | -0.256682679 | downregulated during regeneration - mid |
| ENSDART00000158854 | tpd52l1 | 0.028449452 | -0.063123021 | -0.257733521 | -0.65813611 | -0.260747428 | downregulated during regeneration - mid |
| ENSDART00000158917 | abhd10a | 0.008288008 | -0.199951388 | -0.281143798 | -0.472212357 | -0.246558895 | downregulated during regeneration - mid |
| ENSDART00000158962 | dpf2 | 0.022330586 | 0.023344289 | -0.160855631 | -0.651177712 | -0.029134577 | downregulated during regeneration - mid |
| ENSDART00000159061 | si:ch211-185a18.2 | 0.068645885 | -0.270017947 | -0.418794122 | -0.535787315 | -0.21407671 | downregulated during regeneration - mid |
| ENSDART00000159123 | btbd17b | 0.02583682 | -0.172708316 | -0.362184438 | -0.675325658 | -0.245697841 | downregulated during regeneration - mid |
| ENSDART00000159178 | hspa8 | 0.00434437 | -1.810725395 | -0.905134344 | -6.819365873 | -1.947204217 | downregulated during regeneration - mid |
| ENSDART00000159581 | CT583723.1 | 0.081780045 | -0.203564175 | -0.159816156 | -0.458224986 | -0.16588667 | downregulated during regeneration - mid |
| ENSDART00000159654 | CR847844.2 | 0.019450344 | 0.025394661 | -0.340096605 | -0.548288987 | -0.346027376 | downregulated during regeneration - mid |
| ENSDART00000159670 | abtb1 | 0.013224152 | -0.10462458 | -0.155472174 | -0.372376774 | -0.233981977 | downregulated during regeneration - mid |
| ENSDART00000159733 | CABZ01063757.1 | 0.004296618 | 0.129910694 | -0.203491407 | -0.855433405 | -0.227320323 | downregulated during regeneration - mid |
| ENSDART00000159950 | si:ch1073-450f2.1 | 0.040770952 | -0.285454261 | -0.252056938 | -0.418320337 | -0.16839092 | downregulated during regeneration - mid |
| ENSDART00000159987 | si:dkey-65b13.13 | 0.035889612 | -0.209652039 | -0.259703439 | -0.637719895 | -0.234430743 | downregulated during regeneration - mid |
| ENSDART00000160115 | per1a | 0.005681638 | 0.041503641 | -0.272737679 | -0.476359218 | -0.15653808 | downregulated during regeneration - mid |
| ENSDART00000160134 | hibadha | 0.01949596 | -0.191259291 | -0.221221329 | -0.647202373 | -0.244609156 | downregulated during regeneration - mid |
| ENSDART00000160175 | SYNDIG1 | 0.040346146 | -0.028604943 | -0.23293463 | -0.464850771 | -0.153309965 | downregulated during regeneration - mid |
| ENSDART00000160242 | TYMP | 0.002200182 | -0.668893964 | -0.374517624 | -0.757876583 | -0.434642219 | downregulated during regeneration - mid |
| ENSDART00000160284 | pclob | 0.022160325 | -0.164621492 | -0.643566861 | -0.569524507 | -0.247739035 | downregulated during regeneration - mid |
| ENSDART00000160400 | gchfr | 0.016130084 | -0.373470826 | -0.390793782 | -0.620398999 | -0.352985133 | downregulated during regeneration - mid |
| ENSDART00000160439 | CABZ01037298.1 | 0.009820514 | -0.12483018 | -0.201319697 | -0.520978425 | -0.240731645 | downregulated during regeneration - mid |
| ENSDART00000160450 | CU571069.1 | 0.000313682 | -0.391785076 | -0.855515025 | -0.904047805 | -0.612021184 | downregulated during regeneration - mid |
| ENSDART00000160468 | ubl3a | 0.003138642 | -0.157200321 | -0.291255908 | -0.684230474 | -0.436379037 | downregulated during regeneration - mid |
| ENSDART00000160495 | si:dkey-184p18.2 | 0.011672204 | 0.380936601 | -0.937356475 | -0.364392869 | -0.267264411 | downregulated during regeneration - mid |
| ENSDART00000160538 | arhgap21a | 0.006013085 | -0.416942655 | -0.374699279 | -1.045101409 | -0.721453587 | downregulated during regeneration - mid |
| ENSDART00000160562 | impg2b | 0.011993283 | 0.024461351 | -0.26080833 | -0.706384589 | -0.340668028 | downregulated during regeneration - mid |
| ENSDART00000160700 | dnm1l | 0.023931815 | -0.370704077 | -0.348232697 | -0.495814147 | -0.141373029 | downregulated during regeneration - mid |
| ENSDART00000160742 | clmn | 0.002103359 | -0.341601235 | -0.332225059 | -0.489517534 | -0.288919784 | downregulated during regeneration - mid |
| ENSDART00000160753 | GANAB (1 of many) | 0.023696289 | 0.004666803 | -0.095016348 | -0.32432452 | -0.161685037 | downregulated during regeneration - mid |

| | | | | | | | |
|---|---|---|---|---|---|---|---|
| ENSDART00000160841 | si:ch211-248g20.5 | 0.022501885 | -0.153486145 | -0.342330243 | -0.525669956 | -0.178412263 | downregulated during regeneration - mid |
| ENSDART00000160866 | cry2 | 0.002361523 | -0.114688269 | -0.279009524 | -0.604836383 | -0.332518261 | downregulated during regeneration - mid |
| ENSDART00000161059 | tusc2a | 0.013390976 | 0.068801861 | -0.121312821 | -0.423616648 | -0.242714442 | downregulated during regeneration - mid |
| ENSDART00000161261 | lmbrd2a | 0.083564138 | -0.185598793 | -0.153033552 | -0.360733538 | -0.113007589 | downregulated during regeneration - mid |
| ENSDART00000161266 | CU928046.1 | 0.025523567 | -0.122276569 | -0.195535625 | -0.336584046 | -0.188368371 | downregulated during regeneration - mid |
| ENSDART00000161397 | atg13 | 0.021797231 | -0.137451879 | -0.18705676 | -0.405447689 | -0.269252338 | downregulated during regeneration - mid |
| ENSDART00000161440 | limch1b | 0.039972057 | -0.008873535 | -0.193340096 | -0.430561872 | -0.157445684 | downregulated during regeneration - mid |
| ENSDART00000161652 | si:dkeyp-72e1.9 | 0.040887007 | -0.109528412 | -0.231594686 | -0.432617543 | -0.203491956 | downregulated during regeneration - mid |
| ENSDART00000161755 | CU681842.1 | 0.03135437 | -0.157328162 | -0.245310878 | -0.348302227 | -0.161138988 | downregulated during regeneration - mid |
| ENSDART00000161773 | KCNJ11 (1 of many) | 0.040962132 | -0.422252277 | -0.360338989 | -0.607728772 | -0.332578335 | downregulated during regeneration - mid |
| ENSDART00000161823 | ubl3a | 0.020595702 | -0.089124518 | -0.239047703 | -0.471864878 | -0.290785384 | downregulated during regeneration - mid |
| ENSDART00000161840 | CT574575.1 | 0.006228736 | -0.29915223 | -0.372107241 | -0.764083264 | -0.340151107 | downregulated during regeneration - mid |
| ENSDART00000161882 | bzw2 | 0.010880494 | 0.083435949 | -0.31200966 | -0.5490146 | -0.324311149 | downregulated during regeneration - mid |
| ENSDART00000162070 | tmem9 | 0.005875165 | -0.208225931 | -0.361514875 | -0.511868133 | -0.393713678 | downregulated during regeneration - mid |
| ENSDART00000162493 | kif5ab | 0.000645779 | -0.358862317 | -0.407894033 | -0.585721315 | -0.34361757 | downregulated during regeneration - mid |
| ENSDART00000162664 | 5S_rRNA | 0.014771508 | -0.438943374 | -1.186726175 | -0.920371372 | -0.548334869 | downregulated during regeneration - mid |
| ENSDART00000162669 | slc4a5 | 0.010442784 | -0.13383932 | -0.22542524 | -0.430682825 | -0.287279235 | downregulated during regeneration - mid |
| ENSDART00000162827 | si:dkey-92j12.5 | 0.030149141 | -0.013228451 | -0.265522895 | -0.385437174 | -0.080127062 | downregulated during regeneration - mid |
| ENSDART00000162858 | lmbrd1 | 0.053770748 | -0.243655467 | -0.23291492 | -0.332336473 | -0.114530821 | downregulated during regeneration - mid |
| ENSDART00000162875 | rogdi | 0.014149547 | -0.224891588 | -0.27706391 | -0.36191309 | -0.159630193 | downregulated during regeneration - mid |
| ENSDART00000162940 | mob2b | 0.021735616 | 0.020755037 | -0.2829937 | -0.541864256 | -0.204837959 | downregulated during regeneration - mid |
| ENSDART00000163093 | lrp12 | 0.015282229 | -0.07747456 | -0.233717712 | -0.371900586 | -0.129203418 | downregulated during regeneration - mid |
| ENSDART00000163250 | mef2cb | 0.003671994 | 0.361224616 | -0.300957895 | -0.57227951 | -0.080214844 | downregulated during regeneration - mid |
| ENSDART00000163352 | impg2b | 0.015981762 | -0.050961847 | -0.309849144 | -0.745225088 | -0.302394708 | downregulated during regeneration - mid |
| ENSDART00000163445 | zgc:109982 | 0.01612441 | 0.178949347 | -0.334063834 | -0.837985378 | -0.307869575 | downregulated during regeneration - mid |
| ENSDART00000163521 | si:ch1073-83n3.2 | 0.019433758 | -0.135028227 | -0.352956778 | -0.389044273 | -0.105210001 | downregulated during regeneration - mid |
| ENSDART00000163908 | rnasekb | 0.014026672 | -0.236636314 | -0.265732137 | -0.418465587 | -0.246757364 | downregulated during regeneration - mid |
| ENSDART00000163935 | med30 | 0.026243123 | -0.198525489 | -0.227576976 | -0.505162548 | -0.348410732 | downregulated during regeneration - mid |
| ENSDART00000163976 | CABZ01069287.1 | 0.008492808 | 0.247964945 | -0.272161916 | -0.664208233 | -0.362164102 | downregulated during regeneration - mid |
| ENSDART00000163998 | rps6ka3a | 0.02187407 | 0.01131102 | -0.11493059 | -0.372925255 | -0.200625555 | downregulated during regeneration - mid |
| ENSDART00000164566 | akt3a | 0.001867076 | 0.045255643 | -0.432188612 | -0.498284714 | -0.321504061 | downregulated during regeneration - mid |
| ENSDART00000164650 | pdk3b | 0.093881144 | -0.133813888 | -0.219494437 | -0.474289524 | -0.225858079 | downregulated during regeneration - mid |
| ENSDART00000164695 | arl3l1 | 0.005956006 | -0.208201897 | -0.292220291 | -0.610724074 | -0.376046469 | downregulated during regeneration - mid |
| ENSDART00000164773 | CABZ01072036.1 | 0.021489826 | -0.106672866 | -0.494009284 | -0.613761036 | -0.476277849 | downregulated during regeneration - mid |
| ENSDART00000164792 | cfap74 | 0.02603196 | -0.15899724 | -0.307564837 | -0.482816939 | -0.375147114 | downregulated during regeneration - mid |
| ENSDART00000164855 | crebl2 | 0.013882038 | -0.127345071 | -0.2122205 | -0.432997567 | -0.264872881 | downregulated during regeneration - mid |
| ENSDART00000164983 | anapc15 | 0.022231819 | -0.036185378 | -0.198619926 | -0.408385072 | -0.233599439 | downregulated during regeneration - mid |
| ENSDART00000165018 | cdc42se2 | 0.019981901 | -0.053295614 | -0.137825766 | -0.478383249 | -0.349419762 | downregulated during regeneration - mid |
| ENSDART00000165049 | impg2b | 0.011108721 | -0.010489295 | -0.310856751 | -0.727501593 | -0.334825186 | downregulated during regeneration - mid |
| ENSDART00000165065 | uqcr10 | 0.046716104 | -0.222678585 | -0.318669731 | -0.412855116 | -0.201529958 | downregulated during regeneration - mid |
| ENSDART00000165308 | me2 | 0.026211285 | -0.16075229 | -0.327815721 | -0.451710296 | -0.323851706 | downregulated during regeneration - mid |
| ENSDART00000165333 | si:ch211-207l14.1 | 0.024829185 | -0.144575592 | -0.337549275 | -0.714109722 | -0.444634079 | downregulated during regeneration - mid |
| ENSDART00000165425 | aak1a | 0.033202686 | -0.009756971 | -0.269537246 | -0.43668389 | -0.03607957 | downregulated during regeneration - mid |
| ENSDART00000165542 | si:dkeyp-72e1.9 | 0.014624499 | 0.115590472 | -0.201422352 | -0.774050016 | -0.351374892 | downregulated during regeneration - mid |
| ENSDART00000165557 | BX088524.3 | 0.018703026 | -0.275785171 | -0.331787981 | -0.469187171 | -0.127928431 | downregulated during regeneration - mid |
| ENSDART00000166177 | impg2b | 0.000397106 | 0.006257085 | -0.447154942 | -0.735592553 | -0.255871486 | downregulated during regeneration - mid |
| ENSDART00000166241 | inpp5b | 0.040796459 | -0.146961161 | -0.217731387 | -0.46875304 | -0.117265313 | downregulated during regeneration - mid |
| ENSDART00000166313 | thrb | 0.0223553 | -0.193382381 | -0.415286369 | -0.400030228 | -0.136855946 | downregulated during regeneration - mid |
| ENSDART00000166374 | si:dkey-31f5.11 | 0.05364057 | -0.246069029 | -0.257528931 | -0.445065012 | -0.224168767 | downregulated during regeneration - mid |
| ENSDART00000166463 | cnot6b | 0.033934614 | -0.099106983 | -0.163429858 | -0.300924183 | -0.105283807 | downregulated during regeneration - mid |
| ENSDART00000166515 | si:dkeyp-57d7.4 | 0.006203091 | -0.099733783 | -0.492859873 | -1.003446185 | -0.404973513 | downregulated during regeneration - mid |
| ENSDART00000166650 | bsg | 0.017988416 | -0.333252581 | -0.445374368 | -0.502358398 | -0.336861466 | downregulated during regeneration - mid |
| ENSDART00000166659 | lrrn2 | 0.002931511 | -0.321629641 | -0.404637061 | -0.652990901 | -0.235544369 | downregulated during regeneration - mid |
| ENSDART00000166714 | impg2b | 0.033226863 | -0.072742859 | -0.391473223 | -0.740015258 | -0.360324281 | downregulated during regeneration - mid |
| ENSDART00000166843 | tom1 | 0.02794941 | -0.282793851 | -0.146845245 | -0.347923357 | -0.109554429 | downregulated during regeneration - mid |
| ENSDART00000167117 | si:ch1073-469d17.2 | 0.075212446 | -0.226355895 | -0.436402154 | -0.568359696 | -0.320345187 | downregulated during regeneration - mid |
| ENSDART00000167179 | asf1ba | 0.000512281 | -0.325814646 | -0.52479315 | -0.773177729 | -0.605632897 | downregulated during regeneration - mid |
| ENSDART00000167391 | arhgap21a | 0.004364661 | -0.385966805 | -0.235029359 | -0.527723306 | -0.308128762 | downregulated during regeneration - mid |
| ENSDART00000167570 | actr3b | 0.000292971 | -0.361119463 | -0.449811578 | -0.822009016 | -0.639517901 | downregulated during regeneration - mid |
| ENSDART00000167612 | rnf34a | 0.030702335 | -0.125673437 | -0.210904428 | -0.512991238 | -0.24247481 | downregulated during regeneration - mid |
| ENSDART00000167660 | pja2 | 0.003557275 | -0.078344887 | -0.326712885 | -0.382150009 | -0.239022195 | downregulated during regeneration - mid |
| ENSDART00000167948 | hcn1 | 0.022898139 | -0.135796002 | -0.504326543 | -0.514845616 | -0.152463147 | downregulated during regeneration - mid |
| ENSDART00000167977 | kcnh4b | 0.021960856 | -0.815041676 | -0.830195363 | -1.200187818 | -0.588368652 | downregulated during regeneration - mid |
| ENSDART00000168036 | rdh8b | 0.047560325 | -0.188576333 | -0.341040166 | -0.496499443 | -0.09323063 | downregulated during regeneration - mid |
| ENSDART00000168160 | pip5k1cb | 0.007934853 | -0.047296457 | -0.437985038 | -0.478356963 | -0.284131261 | downregulated during regeneration - mid |
| ENSDART00000168531 | irf2bp2b | 0.089399502 | -0.128403316 | -0.188582728 | -0.404745176 | -0.138034767 | downregulated during regeneration - mid |
| ENSDART00000168616 | ppa1a | 0.022223646 | 0.095923853 | -0.258115212 | -0.556184684 | -0.293684673 | downregulated during regeneration - mid |
| ENSDART00000168639 | cry1ab | 0.023297649 | -0.18716056 | -0.280815639 | -0.387838008 | -0.275846695 | downregulated during regeneration - mid |
| ENSDART00000168850 | taok1b | 0.052592075 | -0.044686095 | -0.116318692 | -0.429811181 | -0.132610503 | downregulated during regeneration - mid |
| ENSDART00000169363 | CR678435.1 | 0.029303695 | -0.089321474 | -0.319658973 | -0.354063558 | -0.259523127 | downregulated during regeneration - mid |
| ENSDART00000169367 | CABZ01061884.1 | 0.037612568 | -0.298329532 | -0.377577791 | -0.54544765 | -0.245416552 | downregulated during regeneration - mid |
| ENSDART00000169376 | PLEKHG5 (1 of many) | 0.004338147 | -0.003560342 | -0.365335228 | -0.516938003 | -0.057649582 | downregulated during regeneration - mid |
| ENSDART00000169484 | amph | 0.012342358 | -0.176945265 | -0.219628366 | -0.476094931 | -0.135317446 | downregulated during regeneration - mid |
| ENSDART00000169511 | ppp2r2bb | 0.005388807 | -0.347621729 | -0.40572046 | -0.501873649 | -0.158073428 | downregulated during regeneration - mid |
| ENSDART00000169733 | klhl23 | 0.013248249 | -0.117061422 | -0.244154323 | -0.447638501 | -0.20478162 | downregulated during regeneration - mid |

| | | | | | | | |
|---|---|---|---|---|---|---|---|
| ENSDART00000169828 | rlbp1a | 0.024411518 | -0.278789297 | -0.290042852 | -0.461967142 | -0.22894081 | downregulated during regeneration - mid |
| ENSDART00000169932 | ccdc142 | 0.032850794 | -0.157730988 | -0.223081294 | -0.3402281 | -0.055715693 | downregulated during regeneration - mid |
| ENSDART00000169943 | CABZ01084447.1 | 0.000529346 | -0.382296742 | -0.397697491 | -0.572266726 | -0.473542202 | downregulated during regeneration - mid |
| ENSDART00000170012 | mapre3b | 0.020911664 | -0.205130823 | -0.301066074 | -0.42167523 | -0.136278286 | downregulated during regeneration - mid |
| ENSDART00000170208 | limch1b | 0.017620686 | -0.042803542 | -0.148322889 | -0.621537079 | -0.273084471 | downregulated during regeneration - mid |
| ENSDART00000170285 | def8 | 0.009606339 | -0.050118345 | -0.13948188 | -0.345534099 | -0.220243255 | downregulated during regeneration - mid |
| ENSDART00000170376 | mpp7a | 0.034023228 | -0.022565767 | -0.302686812 | -0.467837523 | -0.093897739 | downregulated during regeneration - mid |
| ENSDART00000170546 | wdr17 | 0.062857343 | -0.20142732 | -0.213426713 | -0.433783757 | -0.087288031 | downregulated during regeneration - mid |
| ENSDART00000170575 | nfat5b | 0.018611393 | -0.001493923 | -0.392122738 | -0.566909919 | -0.288747412 | downregulated during regeneration - mid |
| ENSDART00000170583 | 43167 | 0.022767452 | -0.037993536 | -0.377054474 | -0.284528801 | -0.103767971 | downregulated during regeneration - mid |
| ENSDART00000170695 | lrit1b | 0.041328295 | -0.349980517 | -0.215317539 | -0.688918418 | -0.304938883 | downregulated during regeneration - mid |
| ENSDART00000170762 | slc44a5b | 0.022987263 | -0.083921908 | -0.392138039 | -0.663427101 | -0.045247688 | downregulated during regeneration - mid |
| ENSDART00000170856 | smu1b | 0.004570574 | 0.353209488 | -0.339997639 | -0.798035862 | -0.487523616 | downregulated during regeneration - mid |
| ENSDART00000170955 | fn3krp | 0.064000341 | -0.183544623 | -0.156282547 | -0.395153052 | -0.231090781 | downregulated during regeneration - mid |
| ENSDART00000171202 | clrn1 | 0.023596757 | -0.342498627 | -0.174544267 | -0.411984605 | -0.164186516 | downregulated during regeneration - mid |
| ENSDART00000171237 | kcnj2b | 0.002963545 | -0.406552715 | -0.524353506 | -0.661059283 | -0.393570981 | downregulated during regeneration - mid |
| ENSDART00000171320 | dcp2 | 0.018930059 | -0.072540161 | -0.060086595 | -0.320231159 | -0.153613562 | downregulated during regeneration - mid |
| ENSDART00000171393 | efr3a | 0.008676069 | 0.028329226 | -0.20834437 | -0.54511994 | -0.110508442 | downregulated during regeneration - mid |
| ENSDART00000171639 | prkacbb | 0.016570302 | -0.216859972 | -0.333993863 | -0.380485937 | -0.232933535 | downregulated during regeneration - mid |
| ENSDART00000171823 | cdc14ab | 0.07124152 | -0.17216976 | -0.3147465 | -0.385013143 | -0.20236803 | downregulated during regeneration - mid |
| ENSDART00000172076 | hook1 | 0.057207579 | -0.040243255 | -0.111929437 | -0.520350092 | -0.090216969 | downregulated during regeneration - mid |
| ENSDART00000172114 | calm3a | 0.016086191 | -0.154349916 | -0.209222212 | -0.409530683 | -0.13969668 | downregulated during regeneration - mid |
| ENSDART00000172166 | NDUFC1 | 0.016665414 | -0.092745693 | -0.325133407 | -0.417118078 | -0.256060161 | downregulated during regeneration - mid |
| ENSDART00000172190 | ajap1 | 0.020568703 | -0.155573908 | -0.463634934 | -0.533953643 | -0.131876486 | downregulated during regeneration - mid |
| ENSDART00000172294 | ctps1b | 0.001175355 | -0.150552652 | -0.426760874 | -0.523567714 | -0.150809326 | downregulated during regeneration - mid |
| ENSDART00000172662 | fam96a | 0.013435197 | -0.029501914 | -0.191990056 | -0.34693921 | -0.272376792 | downregulated during regeneration - mid |
| ENSDART00000172811 | ubap1lb | 0.017531533 | 0.254294803 | -0.127702895 | -0.996786994 | -0.222342304 | downregulated during regeneration - mid |
| ENSDART00000172944 | FO681390.1 | 0.035889612 | -0.061230657 | -0.175829366 | -0.634551101 | -0.148743958 | downregulated during regeneration - mid |
| ENSDART00000173056 | arap3 | 0.010245667 | -0.20638853 | -0.271205037 | -0.427111789 | -0.285607029 | downregulated during regeneration - mid |
| ENSDART00000173169 | pclob | 0.015925333 | -0.241926883 | -0.695024268 | -0.666850982 | -0.322920156 | downregulated during regeneration - mid |
| ENSDART00000173400 | FP236327.1 | 0.017516593 | -0.107319508 | -0.233503644 | -0.613884847 | -0.314094176 | downregulated during regeneration - mid |
| ENSDART00000173430 | pfklb | 0.002058842 | -0.150367408 | -0.216221217 | -0.597100494 | -0.348773684 | downregulated during regeneration - mid |
| ENSDART00000003042 | mdkb | 0.000235661 | -0.457982371 | -0.488604373 | -0.629508021 | -0.485178616 | downregulated during regeneration - late |
| ENSDART00000003612 | NA | 0.0080829 | -0.145765054 | -0.499207076 | -0.35686816 | -0.607333409 | downregulated during regeneration - late |
| ENSDART00000004903 | rdh10b | 0.01613952 | -0.385872948 | -0.19159484 | -0.565090954 | -0.486336464 | downregulated during regeneration - late |
| ENSDART00000005053 | slc12a4 | 0.020066139 | -0.135764977 | -0.039802706 | -0.399029267 | -0.999098872 | downregulated during regeneration - late |
| ENSDART00000007122 | guca1b | 0.03198451 | -0.373174817 | -0.243090423 | -0.518795688 | -0.641476933 | downregulated during regeneration - late |
| ENSDART00000008010 | pdk2a | 0.008401701 | -0.070635847 | 0.148907691 | -0.31523766 | -0.171233724 | downregulated during regeneration - late |
| ENSDART00000008326 | pon2 | 0.003003196 | 0.005358778 | -0.010211923 | -0.591626082 | -0.490299784 | downregulated during regeneration - late |
| ENSDART00000008607 | ttyh2l | 0.001115539 | 0.127438148 | -0.071060271 | -0.346686037 | -0.327410338 | downregulated during regeneration - late |
| ENSDART00000009952 | zfand5a | 0.001528476 | -0.376361025 | -0.47427013 | -1.180526792 | -1.063292908 | downregulated during regeneration - late |
| ENSDART00000010046 | rhpn2 | 0.005024347 | -0.531285649 | -0.280012714 | -0.367908087 | -0.481376216 | downregulated during regeneration - late |
| ENSDART00000011135 | kita | 0.005077923 | -0.1368269 | -0.266279657 | -0.502022945 | -0.592221276 | downregulated during regeneration - late |
| ENSDART00000011519 | slc6a1l | 0.002718393 | 0.105782424 | -0.162202998 | -0.466558128 | -0.662799909 | downregulated during regeneration - late |
| ENSDART00000014843 | bdnf | 0.000108354 | -0.572929745 | -0.434797241 | -0.517838864 | -0.969054184 | downregulated during regeneration - late |
| ENSDART00000015095 | uts1 | 0.00114562 | -0.304915972 | -0.432455801 | -0.651436618 | -0.693973757 | downregulated during regeneration - late |
| ENSDART00000017485 | sf3b6 | 0.015609101 | -0.19155437 | -0.273140548 | -0.376775786 | -0.432460535 | downregulated during regeneration - late |
| ENSDART00000018054 | trh | 0.001423412 | -0.369432982 | -0.400024601 | -0.784012545 | -0.865751572 | downregulated during regeneration - late |
| ENSDART00000018347 | cab39l1 | 0.011046859 | -0.259625217 | -0.162823278 | -0.312896931 | -0.294075261 | downregulated during regeneration - late |
| ENSDART00000018886 | ghra | 0.006658285 | -0.12746236 | -0.080311259 | -0.403757478 | -0.551820215 | downregulated during regeneration - late |
| ENSDART00000019766 | tgfb3 | 0.020798309 | -0.14127581 | -0.182290633 | -0.363811528 | -0.271524635 | downregulated during regeneration - late |
| ENSDART00000020999 | angptl1a | 0.068605693 | -0.447558553 | -0.389827093 | -0.516507166 | -1.101603289 | downregulated during regeneration - late |
| ENSDART00000021121 | stx5al | 0.020719498 | -0.079551416 | -0.239231712 | -0.554538789 | -0.564757113 | downregulated during regeneration - late |
| ENSDART00000021168 | rxrga | 0.00269825 | 0.040179684 | -0.194377683 | -0.381512289 | -0.377392107 | downregulated during regeneration - late |
| ENSDART00000022051 | gins1 | 0.011046859 | -0.049060202 | -0.07395231 | -0.409999594 | -0.285702931 | downregulated during regeneration - late |
| ENSDART00000022581 | rab22a | 0.018963501 | -0.072513798 | -0.08434934 | -0.299709625 | -0.230388715 | downregulated during regeneration - late |
| ENSDART00000023613 | her6 | 0.030622517 | -0.335042933 | -0.234567638 | -0.250120727 | -0.59712361 | downregulated during regeneration - late |
| ENSDART00000024328 | slc34a2a | 0.032189514 | -1.294382652 | -0.588426598 | -0.664087867 | -1.114341851 | downregulated during regeneration - late |
| ENSDART00000026085 | ptges | 0.014800178 | -0.133597623 | -0.186036004 | -0.449634161 | -0.455296325 | downregulated during regeneration - late |
| ENSDART00000026316 | sema3gb | 0.023200703 | -0.148485986 | -0.183297493 | -0.3621349 | -0.465943338 | downregulated during regeneration - late |
| ENSDART00000027417 | zgc:171704 | 0.00039492 | 0.227240571 | -0.665719796 | -1.161293822 | -1.438167938 | downregulated during regeneration - late |
| ENSDART00000029946 | ube2b | 0.013774152 | -0.262228145 | -0.117966704 | -0.377845713 | -0.176574464 | downregulated during regeneration - late |
| ENSDART00000030579 | crhbp | 0.001133915 | 0.109353477 | -0.053663997 | -0.382440888 | -0.742296052 | downregulated during regeneration - late |
| ENSDART00000031727 | vamp8 | 0.000787471 | 0.086516161 | -0.020853386 | -0.34297459 | -0.531441391 | downregulated during regeneration - late |
| ENSDART00000032393 | ginm1 | 0.012470284 | -0.054454012 | -0.092963972 | -0.327748186 | -0.224389181 | downregulated during regeneration - late |
| ENSDART00000032502 | nebl | 0.026706504 | -0.405575453 | -0.348273521 | -0.638746976 | -0.69843843 | downregulated during regeneration - late |
| ENSDART00000034638 | ccdc28a | 0.018433591 | -0.059055282 | -0.067802136 | -0.368600818 | -0.274995246 | downregulated during regeneration - late |
| ENSDART00000035031 | sgk1 | 0.002559781 | 0.036456511 | -0.228295468 | -0.323414575 | -0.393803665 | downregulated during regeneration - late |
| ENSDART00000035067 | abhd2a | 0.008989451 | -0.280282686 | -0.122469223 | -0.369410687 | -0.218167862 | downregulated during regeneration - late |
| ENSDART00000038310 | ormdl3 | 0.010195593 | -0.300841314 | -0.124106092 | -0.535837175 | -0.413488994 | downregulated during regeneration - late |
| ENSDART00000038674 | tmem230a | 0.005028852 | -0.312877262 | -0.360088684 | -0.444507782 | -0.485519275 | downregulated during regeneration - late |
| ENSDART00000039399 | cavin2a | 0.000442465 | -0.454722967 | -0.463350309 | -0.727916614 | -0.482344794 | downregulated during regeneration - late |
| ENSDART00000042123 | cx52.6 | 0.003425348 | -0.622226461 | -0.079417665 | -0.692157173 | -0.611068819 | downregulated during regeneration - late |
| ENSDART00000042307 | fam60a | 0.000248952 | -0.454119179 | -0.522927402 | -0.624149133 | -1.020018948 | downregulated during regeneration - late |
| ENSDART00000042481 | phf23a | 0.019414194 | -0.371442433 | -0.230558793 | -0.302265692 | -0.28776115 | downregulated during regeneration - late |

| | | | | | | | |
|---|---|---|---|---|---|---|---|
| ENSDART00000043226 | guca1c | 0.014712033 | -0.531402053 | -0.370893475 | -0.804093243 | -0.754648782 | downregulated during regeneration - late |
| ENSDART00000044426 | si:dkey-240h12.4 | 0.016178224 | 0.077516947 | -0.039107616 | -0.22879164 | -0.507759534 | downregulated during regeneration - late |
| ENSDART00000044860 | maff | 0.002310171 | 0.057535561 | 0.039208376 | -0.366680053 | -0.441007369 | downregulated during regeneration - late |
| ENSDART00000045126 | lama5 | 0.007419702 | -0.044158589 | 0.007727115 | -0.324426294 | -0.496259172 | downregulated during regeneration - late |
| ENSDART00000046360 | rhoua | 0.001489209 | -0.234114921 | -0.11992892 | -0.459682602 | -0.396381281 | downregulated during regeneration - late |
| ENSDART00000046716 | cited2 | 0.010493088 | -0.252125812 | -0.225219779 | -0.427941506 | -0.423861644 | downregulated during regeneration - late |
| ENSDART00000047378 | sst3 | 0.024602158 | -0.489863502 | -0.562162931 | -0.649979789 | -0.586308909 | downregulated during regeneration - late |
| ENSDART00000049075 | add3a | 0.023840313 | -0.25124279 | -0.167771329 | -0.330721354 | -0.242087591 | downregulated during regeneration - late |
| ENSDART00000050230 | tspan3a | 0.014404044 | -0.260999172 | -0.146975695 | -0.364155673 | -0.280149491 | downregulated during regeneration - late |
| ENSDART00000051197 | c10h21orf59 | 0.043628495 | -0.175790498 | -0.1331203 | -0.516409592 | -0.442586367 | downregulated during regeneration - late |
| ENSDART00000051357 | zmat5 | 0.04367746 | -0.126081267 | -0.164432805 | -0.376616148 | -0.245219161 | downregulated during regeneration - late |
| ENSDART00000051516 | tacr1a | 0.03254446 | -0.234436772 | -0.252170036 | -0.371966451 | -0.377528767 | downregulated during regeneration - late |
| ENSDART00000051546 | rps6ka3a | 0.004239143 | -0.063638799 | 0.031915539 | -0.376231824 | -0.261540289 | downregulated during regeneration - late |
| ENSDART00000051655 | snrnp27 | 0.019324618 | -0.149284802 | -0.196914421 | -0.340354698 | -0.296345743 | downregulated during regeneration - late |
| ENSDART00000051948 | si:dkey-17m8.2 | 0.029473595 | -2.249269734 | -0.678959768 | -1.034434829 | -1.318620097 | downregulated during regeneration - late |
| ENSDART00000052029 | cart3 | 0.000305619 | -0.527262177 | -0.518524777 | -0.63363087 | -0.567396556 | downregulated during regeneration - late |
| ENSDART00000052423 | spry2 | 0.008879163 | -0.373633249 | -0.274093568 | -0.317163264 | -0.534996547 | downregulated during regeneration - late |
| ENSDART00000052703 | nucb2b | 0.005154611 | -0.072514611 | -0.317049949 | -0.490530665 | -0.459714357 | downregulated during regeneration - late |
| ENSDART00000052912 | pcdh20 | 0.016905836 | -0.38541457 | -0.110151808 | -0.462368809 | -0.238274084 | downregulated during regeneration - late |
| ENSDART00000053310 | tmem18 | 0.023272159 | -0.091641733 | -0.057752607 | -0.353980434 | -0.264448619 | downregulated during regeneration - late |
| ENSDART00000053463 | mgll | 0.063259686 | -0.18197879 | -0.150318561 | -0.320930933 | -0.180307684 | downregulated during regeneration - late |
| ENSDART00000053773 | lsm6 | 0.013538515 | -0.104076864 | -0.094751595 | -0.350900229 | -0.413521857 | downregulated during regeneration - late |
| ENSDART00000053916 | mtnr1ab | 0.032098855 | -0.333808774 | -0.363646472 | -0.291230509 | -0.411800208 | downregulated during regeneration - late |
| ENSDART00000054386 | qdprb1 | 0.066779801 | -0.130677564 | -0.146959168 | -0.522581238 | -0.268035824 | downregulated during regeneration - late |
| ENSDART00000054462 | smim19 | 0.016203475 | -0.090105146 | -0.174385509 | -0.37090077 | -0.289350438 | downregulated during regeneration - late |
| ENSDART00000054867 | aup1 | 0.017245517 | -0.021467306 | -0.053295505 | -0.309019838 | -0.217937602 | downregulated during regeneration - late |
| ENSDART00000055139 | col9a3 | 0.000444888 | 0.340683061 | 0.20859259 | -0.192921312 | -1.195377277 | downregulated during regeneration - late |
| ENSDART00000055706 | her15.1 | 0.021144178 | -1.483608056 | -1.154441068 | -0.698110478 | -1.436427969 | downregulated during regeneration - late |
| ENSDART00000055709 | her2 | 0.030146589 | -0.984524994 | -0.833253569 | -0.717745512 | -1.233522642 | downregulated during regeneration - late |
| ENSDART00000055779 | ggact.2 | 0.030279926 | -0.390632373 | -0.197002813 | -0.565987089 | -0.546417063 | downregulated during regeneration - late |
| ENSDART00000056939 | zgc:85858 | 0.037793991 | -0.254903494 | -0.357469282 | -0.615277126 | -0.499477333 | downregulated during regeneration - late |
| ENSDART00000056996 | sfrp5 | 0.011389892 | 0.077859934 | 0.01622227 | -0.244583159 | -1.042810978 | downregulated during regeneration - late |
| ENSDART00000057124 | tefa | 0.007560757 | -0.17959692 | 0.074135039 | -0.330628318 | -0.262330789 | downregulated during regeneration - late |
| ENSDART00000057125 | tefa | 0.002939873 | -0.190705988 | 0.094171259 | -0.306461854 | -0.301060608 | downregulated during regeneration - late |
| ENSDART00000057439 | parla | 0.056801891 | -0.763945698 | -0.13237506 | -0.4980372 | -0.373141189 | downregulated during regeneration - late |
| ENSDART00000057519 | zgc:194209 | 0.001633797 | 0.014343488 | -0.12072846 | -0.449908342 | -0.453116382 | downregulated during regeneration - late |
| ENSDART00000057689 | bag3 | 0.030198325 | -0.537315577 | -0.216588912 | -0.658528557 | -0.795247472 | downregulated during regeneration - late |
| ENSDART00000057865 | ier3ip1 | 0.003879543 | -0.032004312 | -0.136273649 | -0.345681209 | -0.364367077 | downregulated during regeneration - late |
| ENSDART00000058667 | RDH13 (1 of many) | 0.0088045 | -0.138024992 | -0.086121321 | -0.690305733 | -0.540517763 | downregulated during regeneration - late |
| ENSDART00000059478 | lrrc32 | 0.00518047 | -0.083635057 | -0.196945688 | -0.286292563 | -0.79945825 | downregulated during regeneration - late |
| ENSDART00000059550 | lrrc51 | 0.005486754 | -1.204791639 | -0.640574252 | -1.049736423 | -1.0433072 | downregulated during regeneration - late |
| ENSDART00000060174 | jagn1a | 0.011982948 | -0.416430339 | -0.038483688 | -0.018273384 | -0.314381472 | downregulated during regeneration - late |
| ENSDART00000060304 | dhrs13a.3 | 0.001013659 | -0.00917183 | 0.036729289 | -0.280144118 | -0.407439325 | downregulated during regeneration - late |
| ENSDART00000061007 | mt2 | 0.000122194 | -0.024087588 | -0.437804286 | -0.890429121 | -0.686083783 | downregulated during regeneration - late |
| ENSDART00000061117 | rrbp1b | 0.019354723 | -0.094086695 | 0.04423759 | -0.324207933 | -0.60750175 | downregulated during regeneration - late |
| ENSDART00000061261 | cx43 | 0.004463264 | -0.064274202 | -0.132171143 | -0.392515504 | -0.502400185 | downregulated during regeneration - late |
| ENSDART00000061435 | hsbp1b | 0.014624499 | -0.02310974 | -0.083751559 | -0.317854229 | -0.256774332 | downregulated during regeneration - late |
| ENSDART00000062066 | si:dkey-177p2.6 | 0.005438298 | -0.250914133 | -0.251912097 | -0.422547135 | -0.291399654 | downregulated during regeneration - late |
| ENSDART00000062143 | zgc:77650 | 0.004824695 | -0.109887255 | -0.133378465 | -0.376117658 | -0.331935654 | downregulated during regeneration - late |
| ENSDART00000062181 | RALGDS | 0.019466285 | -0.386705371 | -0.180287867 | -0.473841885 | -0.479084169 | downregulated during regeneration - late |
| ENSDART00000062518 | gstr | 0.027807337 | -0.173225499 | -0.241367808 | -0.232576695 | -0.339957813 | downregulated during regeneration - late |
| ENSDART00000063625 | gpx3 | 0.000679804 | 0.093413749 | 0.050792899 | -0.431389541 | -0.288399492 | downregulated during regeneration - late |
| ENSDART00000063804 | wu:fj39g12 | 0.000278198 | -1.205265642 | -1.354971991 | -1.517691105 | -1.318916155 | downregulated during regeneration - late |
| ENSDART00000064111 | faub | 0.021419678 | 0.024926044 | -0.061766338 | -0.402186419 | -0.29870419 | downregulated during regeneration - late |
| ENSDART00000064376 | sod1 | 0.030292962 | -0.248477917 | -0.212122084 | -0.427881094 | -0.390078569 | downregulated during regeneration - late |
| ENSDART00000064878 | gxylt2 | 0.010154159 | -0.088947943 | -0.180342387 | -0.380560197 | -0.388684923 | downregulated during regeneration - late |
| ENSDART00000066703 | rdh10a | 0.019015879 | -0.287660525 | -0.158056129 | -0.356819377 | -0.262167994 | downregulated during regeneration - late |
| ENSDART00000067239 | guca1g | 0.003508363 | -0.979413995 | -1.042352228 | -1.597708164 | -1.275542792 | downregulated during regeneration - late |
| ENSDART00000067362 | cart2 | 0.014638227 | -0.749355761 | -0.384853037 | -0.719287733 | -0.625188951 | downregulated during regeneration - late |
| ENSDART00000067446 | slc38a4 | 0.000237691 | -0.251589327 | -0.331078479 | -0.606665517 | -0.538910759 | downregulated during regeneration - late |
| ENSDART00000074100 | osgn1 | 0.000510711 | 0.255332714 | 0.013078065 | -0.613186841 | -0.487873824 | downregulated during regeneration - late |
| ENSDART00000074380 | tsga10 | 0.026816553 | -0.358572407 | -0.32335115 | -0.582555062 | -0.41305108 | downregulated during regeneration - late |
| ENSDART00000074458 | ptpmt1 | 0.006314435 | -0.138560892 | -0.279060294 | -0.535667209 | -0.518833125 | downregulated during regeneration - late |
| ENSDART00000075150 | bmp4 | 0.0114419 | -0.321654174 | -0.192558055 | -0.354985644 | -0.520317573 | downregulated during regeneration - late |
| ENSDART00000075499 | si:dkey-283b1.7 | 0.001565956 | -0.753135255 | -0.256638623 | -0.866666345 | -0.691414461 | downregulated during regeneration - late |
| ENSDART00000075519 | aldh1a2 | 0.047619389 | -0.136704985 | -0.161016939 | -0.528203419 | -0.420522124 | downregulated during regeneration - late |
| ENSDART00000075663 | cracr2b | 0.004668635 | 0.080855847 | 0.057965052 | -0.927246946 | -0.842015409 | downregulated during regeneration - late |
| ENSDART00000076066 | lin37 | 0.005026727 | -0.271630989 | -0.17922011 | -0.496068109 | -0.578879729 | downregulated during regeneration - late |
| ENSDART00000077008 | alox5ap | 0.0014833 | 0.122223757 | -0.082901221 | -0.550940123 | -0.70030614 | downregulated during regeneration - late |
| ENSDART00000077080 | PTP4A3 (1 of many) | 0.006896678 | 0.077833133 | 0.004217905 | -0.463336754 | -0.666773375 | downregulated during regeneration - late |
| ENSDART00000077411 | cxcl12b | 0.001960123 | -0.236947042 | -0.2488008 | -0.593903488 | -0.559818867 | downregulated during regeneration - late |
| ENSDART00000077951 | pcolce2b | 0.013245259 | -0.119540012 | -0.232226227 | -0.447813206 | -1.286575617 | downregulated during regeneration - late |
| ENSDART00000078148 | smc1a | 0.041808931 | -3.399991944 | -0.069804776 | -1.248223526 | -0.569589215 | downregulated during regeneration - late |
| ENSDART00000078181 | SLC3A2 (1 of many) | 0.020491546 | -0.23816401 | -0.122270906 | -0.56307324 | -1.283472558 | downregulated during regeneration - late |
| ENSDART00000078449 | itih6 | 0.002591094 | -0.23325998 | -0.04381884 | -0.494745518 | -1.555031348 | downregulated during regeneration - late |

| | | | | | | | |
|---|---|---|---|---|---|---|---|
| ENSDART00000078858 | si:ch73-86n18.1 | 0.003028992 | 0.106229924 | -0.155735751 | -0.58770094 | -0.854457693 | downregulated during regeneration - late |
| ENSDART00000078916 | smim7 | 0.014686352 | -0.217464923 | -0.109941617 | -0.350272436 | -0.25934653 | downregulated during regeneration - late |
| ENSDART00000079112 | calca | 0.031235096 | -0.439449486 | -0.32190484 | -0.601830596 | -0.728154291 | downregulated during regeneration - late |
| ENSDART00000079879 | si:dkey-91i10.3 | 0.060601627 | -0.580141681 | -0.553279281 | -0.678150475 | -0.52463511 | downregulated during regeneration - late |
| ENSDART00000080014 | rps8b | 0.012554187 | -0.261943359 | -0.309301007 | -0.481979191 | -0.409711249 | downregulated during regeneration - late |
| ENSDART00000081946 | zgc:112332 | 0.003060275 | -0.051059621 | 0.027246103 | -0.289131429 | -0.652053172 | downregulated during regeneration - late |
| ENSDART00000082151 | uqcrh | 0.037614097 | -0.212671975 | -0.358890471 | -0.337939592 | -0.281374001 | downregulated during regeneration - late |
| ENSDART00000083367 | prcp | 0.014634638 | -0.058674879 | -0.090308681 | -0.37460188 | -0.414832363 | downregulated during regeneration - late |
| ENSDART00000084055 | fzd7a | 0.04742464 | -0.350921417 | -0.318886271 | -0.299513467 | -0.86880486 | downregulated during regeneration - late |
| ENSDART00000084131 | fam160b2 | 0.009592472 | 0.036570039 | -0.082247362 | -0.429951116 | -0.30533058 | downregulated during regeneration - late |
| ENSDART00000084598 | vimp | 0.005671256 | -0.003252442 | -0.076004385 | -0.324006724 | -0.304272734 | downregulated during regeneration - late |
| ENSDART00000084861 | cish | 0.004022079 | -0.446878588 | -0.18032932 | -1.508647694 | -1.506136587 | downregulated during regeneration - late |
| ENSDART00000085294 | tnfrsf9a | 0.001754036 | -0.468498688 | -0.352709139 | -0.665957795 | -1.403982945 | downregulated during regeneration - late |
| ENSDART00000085442 | mut | 0.009834259 | -0.094284216 | -0.181855717 | -0.336054518 | -0.342951233 | downregulated during regeneration - late |
| ENSDART00000086051 | mecom | 0.04049764 | -0.550304416 | -0.298852875 | -0.293768303 | -1.334615988 | downregulated during regeneration - late |
| ENSDART00000086905 | nrn1lb | 0.000133356 | -0.455174799 | -0.417502528 | -0.690990621 | -0.656036806 | downregulated during regeneration - late |
| ENSDART00000087339 | cdon | 0.026243123 | -0.358409006 | -0.160970657 | -0.495687672 | -0.865355598 | downregulated during regeneration - late |
| ENSDART00000088342 | cytip | 0.00140081 | 0.001918005 | -0.070156162 | -0.471206178 | -0.369058853 | downregulated during regeneration - late |
| ENSDART00000089015 | zbtb7a | 0.00191624 | -0.478666207 | -0.203919761 | -0.738823143 | -0.713706626 | downregulated during regeneration - late |
| ENSDART00000089339 | dph7 | 0.018858047 | -0.317220664 | -0.267955931 | -0.402220665 | -0.245139947 | downregulated during regeneration - late |
| ENSDART00000089342 | cfap126 | 0.014365121 | -0.196227391 | -0.233968621 | -0.449699376 | -0.526724945 | downregulated during regeneration - late |
| ENSDART00000091124 | aifm3 | 0.010447806 | -0.238811768 | -0.279253276 | -0.598306352 | -0.861256088 | downregulated during regeneration - late |
| ENSDART00000091156 | tacc1 | 0.012895641 | -0.166984032 | -0.162668339 | -0.396514006 | -0.330157066 | downregulated during regeneration - late |
| ENSDART00000091252 | spata13 | 0.001728321 | 0.108055641 | -0.023595371 | -0.309799561 | -0.91906317 | downregulated during regeneration - late |
| ENSDART00000093000 | PLEKHB1 | 0.010802023 | -0.366311468 | -0.143931789 | -0.403311135 | -0.393296015 | downregulated during regeneration - late |
| ENSDART00000098045 | gas1b | 0.005181375 | -0.225161666 | -0.17230943 | -0.331649886 | -0.52065088 | downregulated during regeneration - late |
| ENSDART00000098575 | trim110 | 0.020470078 | -0.538539785 | -0.074004136 | -0.613819609 | -0.257802554 | downregulated during regeneration - late |
| ENSDART00000099089 | ggcx | 0.031769506 | -0.059935848 | -0.149142859 | -0.333945915 | -0.254577798 | downregulated during regeneration - late |
| ENSDART00000099208 | asph | 0.026872157 | -0.288641006 | -0.271409423 | -0.460024196 | -0.387402543 | downregulated during regeneration - late |
| ENSDART00000099476 | fam174b | 0.003432504 | -0.249603276 | -0.126773683 | -0.376574062 | -0.35637543 | downregulated during regeneration - late |
| ENSDART00000099528 | spry4 | 0.000910711 | -1.005535524 | -0.789062042 | -0.760146662 | -1.480463024 | downregulated during regeneration - late |
| ENSDART00000100286 | fgfr4 | 0.004958321 | -0.349487842 | -0.279142199 | -0.557520873 | -0.83628708 | downregulated during regeneration - late |
| ENSDART00000100386 | mstnb | 0.032308002 | -0.232479836 | -0.19741782 | -0.546065599 | -0.487217319 | downregulated during regeneration - late |
| ENSDART00000100605 | ttc32 | 0.004111621 | -0.362267617 | -0.196641209 | -0.477779827 | -0.443477008 | downregulated during regeneration - late |
| ENSDART00000102125 | schip1 | 0.014880314 | -0.065634739 | -0.008431238 | -0.356995513 | -0.237294266 | downregulated during regeneration - late |
| ENSDART00000103586 | hdac9b | 0.003730132 | -0.63802499 | -0.480501706 | -0.6152094 | -0.590506452 | downregulated during regeneration - late |
| ENSDART00000103785 | ggact.3 | 0.044966319 | -1.209900002 | -0.367470534 | -1.050388105 | -0.531293993 | downregulated during regeneration - late |
| ENSDART00000103795 | ggact.1 | 0.008574428 | 0.007439004 | -0.050660849 | -0.406911538 | -0.573615848 | downregulated during regeneration - late |
| ENSDART00000104043 | dkk3b | 0.003678598 | 0.124905707 | -0.440538945 | -0.566305744 | -1.06504734 | downregulated during regeneration - late |
| ENSDART00000104257 | lpin1 | 0.011195192 | -0.263973453 | -0.162876624 | -0.357103753 | -0.3626544 | downregulated during regeneration - late |
| ENSDART00000104545 | avpr2ab | 0.033298538 | -0.127228111 | -0.325246207 | -0.456539735 | -1.288721053 | downregulated during regeneration - late |
| ENSDART00000104592 | pm20d1.2 | 0.003337853 | -0.022497234 | -0.114097556 | -0.391118153 | -0.365853437 | downregulated during regeneration - late |
| ENSDART00000105323 | zgc:162255 | 0.003678598 | -0.250927131 | -0.470845634 | -0.44137647 | -0.557741796 | downregulated during regeneration - late |
| ENSDART00000105813 | krt222 | 0.028543949 | -0.522679169 | -0.470823148 | -0.650543043 | -0.842913393 | downregulated during regeneration - late |
| ENSDART00000105903 | lsm5 | 0.003228981 | 0.135971292 | -0.078983757 | -0.333257253 | -0.331997667 | downregulated during regeneration - late |
| ENSDART00000109124 | rspo2 | 0.005414718 | 0.077521406 | 0.271943255 | -0.228972258 | -0.854369599 | downregulated during regeneration - late |
| ENSDART00000109546 | unc5b | 0.002533609 | -0.076752071 | -0.101810869 | -0.423560106 | -0.292578205 | downregulated during regeneration - late |
| ENSDART00000109853 | zmp:0000000801 | 0.002881312 | -0.4485216 | -0.462342097 | -0.503236916 | -0.960013508 | downregulated during regeneration - late |
| ENSDART00000110544 | znf219 | 0.031344075 | -0.234403747 | -0.210413301 | -0.211690001 | -0.32688667 | downregulated during regeneration - late |
| ENSDART00000111165 | best2 | 0.019487475 | -0.15773889 | -0.249582303 | -0.643700083 | -1.151697377 | downregulated during regeneration - late |
| ENSDART00000111767 | si:ch211-39k3.2 | 0.001551573 | -0.094085509 | -0.095643794 | -0.600534706 | -0.396103136 | downregulated during regeneration - late |
| ENSDART00000111799 | HEPACAM (1 of many) | 0.022218395 | -0.300776869 | -0.301240305 | -0.377807505 | -0.329578876 | downregulated during regeneration - late |
| ENSDART00000111905 | BX649498.1 | 0.031814592 | -0.175767299 | -0.206924623 | -0.494153858 | -0.425985895 | downregulated during regeneration - late |
| ENSDART00000112333 | cnnm2b | 0.00144644 | -0.413171039 | -0.296279679 | -0.539640998 | -0.529524127 | downregulated during regeneration - late |
| ENSDART00000112579 | scg2b | 0.00638925 | -0.360046965 | -0.210825369 | -0.351760362 | -0.440018453 | downregulated during regeneration - late |
| ENSDART00000112845 | zgc:195173 | 0.001259259 | -0.156651959 | -0.182049019 | -0.580426063 | -0.462296521 | downregulated during regeneration - late |
| ENSDART00000112856 | TNC (1 of many) | 0.024409241 | -0.33689758 | -0.285587425 | -0.51132075 | -1.127367477 | downregulated during regeneration - late |
| ENSDART00000113197 | prok1 | 0.007586659 | -0.431580361 | -0.477066939 | -0.390033973 | -0.689815046 | downregulated during regeneration - late |
| ENSDART00000113734 | chrdl2 | 0.03875014 | -0.47545279 | -0.359333121 | -0.57438139 | -0.417914769 | downregulated during regeneration - late |
| ENSDART00000113859 | crispld1a | 0.000346997 | 0.006653123 | -0.124334074 | -0.47278833 | -1.109435686 | downregulated during regeneration - late |
| ENSDART00000114319 | si:ch211-76l23.7 | 0.037291854 | -0.274578898 | -0.196596974 | -0.473622321 | -1.111584541 | downregulated during regeneration - late |
| ENSDART00000114676 | gstm.2 | 0.00889295 | 0.102339801 | -0.045272058 | -0.466318631 | -0.673317054 | downregulated during regeneration - late |
| ENSDART00000115356 | rabif | 0.025960292 | -0.104235333 | -0.157538528 | -0.394672329 | -0.327606577 | downregulated during regeneration - late |
| ENSDART00000116415 | NA | 0.01907307 | -0.367308157 | -0.297245306 | -0.576085374 | -0.866023911 | downregulated during regeneration - late |
| ENSDART00000121226 | FO704882.1 | 0.007797716 | -0.227266262 | -0.409349874 | -0.661498137 | -0.983304677 | downregulated during regeneration - late |
| ENSDART00000121675 | angptl1a | 0.033012979 | -0.785726694 | -0.739578659 | -0.711456 | -1.277545302 | downregulated during regeneration - late |
| ENSDART00000121716 | FAM107A | 0.022475824 | -0.353665396 | -0.22740498 | -0.416694229 | -0.314761483 | downregulated during regeneration - late |
| ENSDART00000121817 | fbln7 | 0.002545259 | 0.143978228 | 0.156275351 | -0.420940219 | -1.064416584 | downregulated during regeneration - late |
| ENSDART00000122359 | si:dkey-164f24.2 | 0.005230737 | -0.316246733 | -0.171154186 | -0.508268616 | -0.361228089 | downregulated during regeneration - late |
| ENSDART00000122519 | st8sia6 | 0.009263888 | -0.258839848 | -0.191246071 | -0.329756561 | -0.383108642 | downregulated during regeneration - late |
| ENSDART00000122966 | hapln1a | 0.031344075 | -5.328261829 | -0.041096146 | -1.913311737 | -3.425165696 | downregulated during regeneration - late |
| ENSDART00000123282 | si:dkey-7i4.1 | 0.03479782 | -0.131885839 | -0.288104676 | -0.328002164 | -0.299371464 | downregulated during regeneration - late |
| ENSDART00000124762 | hsp70.1 | 0.013341759 | -0.061931986 | -0.652822267 | -0.746588848 | -0.837604985 | downregulated during regeneration - late |
| ENSDART00000125116 | tnfaip8l3 | 0.008910267 | -0.29259694 | -0.291950956 | -0.382018775 | -0.337072666 | downregulated during regeneration - late |
| ENSDART00000125691 | sntb1 | 0.003107952 | -0.296403963 | -0.433378655 | -0.538134366 | -0.484556917 | downregulated during regeneration - late |

| | | | | | | | |
|---|---|---|---|---|---|---|---|
| ENSDART00000125923 | cxcl14 | 0.013886694 | -0.090480689 | -0.043715373 | -0.409912244 | -0.49269465 | downregulated during regeneration - late |
| ENSDART00000126870 | ndufa4l2b | 0.005353377 | -0.364506533 | -0.363911454 | -0.576856062 | -0.429052776 | downregulated during regeneration - late |
| ENSDART00000127093 | CR749162.1 | 0.045674994 | -1.06520168 | -0.24719149 | -0.825954556 | -0.565925266 | downregulated during regeneration - late |
| ENSDART00000127536 | cbx7b | 0.007396728 | -0.401352835 | -0.168978702 | -0.391659396 | -0.597817617 | downregulated during regeneration - late |
| ENSDART00000128011 | tnfsf10l3 | 0.014034602 | -0.207145482 | -0.351713028 | -0.904110327 | -0.837055548 | downregulated during regeneration - late |
| ENSDART00000128014 | dnah2 | 0.046252381 | -0.407226928 | -0.498892771 | -0.604204962 | -0.453086208 | downregulated during regeneration - late |
| ENSDART00000128127 | slc38a3b | 0.005596862 | -0.302545777 | -0.299963192 | -0.361680425 | -0.612800784 | downregulated during regeneration - late |
| ENSDART00000128149 | fam32a | 0.003797938 | -0.07314666 | -0.18816053 | -0.368263865 | -0.361807904 | downregulated during regeneration - late |
| ENSDART00000129214 | COX7A2 (1 of many) | 0.01962183 | -0.085157083 | -0.364093354 | -0.62455009 | -0.475756885 | downregulated during regeneration - late |
| ENSDART00000129597 | sorbs3 | 0.003535619 | 0.016863628 | 0.072993125 | -0.314882072 | -0.546339357 | downregulated during regeneration - late |
| ENSDART00000129710 | crtac1a | 0.003137832 | -0.277382094 | 0.025987329 | -0.337990061 | -0.608225673 | downregulated during regeneration - late |
| ENSDART00000130264 | rpz | 0.050182269 | -0.422979934 | -0.28718615 | -0.432905099 | -0.242626605 | downregulated during regeneration - late |
| ENSDART00000131027 | TMEM216 | 0.033109676 | -0.171407049 | -0.12306379 | -0.494331831 | -0.331123076 | downregulated during regeneration - late |
| ENSDART00000131582 | cbx3b | 0.014192021 | -0.219219 | -0.174464705 | -0.354832647 | -0.341719477 | downregulated during regeneration - late |
| ENSDART00000132915 | serpine3 | 0.007368463 | -0.093377161 | -0.220423221 | -0.800809367 | -1.077921671 | downregulated during regeneration - late |
| ENSDART00000132995 | ak3 | 0.000854936 | 0.087221034 | 0.050666468 | -0.189471235 | -0.418943401 | downregulated during regeneration - late |
| ENSDART00000133496 | CU467110.1 | 0.0321077 | -0.265746647 | -0.189978673 | -0.362219245 | -0.151144337 | downregulated during regeneration - late |
| ENSDART00000133891 | FP015789.1 | 0.00918017 | -0.349747936 | -0.468679484 | -0.676764109 | -0.710268862 | downregulated during regeneration - late |
| ENSDART00000134202 | ccka | 0.012864775 | -0.373763887 | -0.443030746 | -0.635815918 | -0.444990754 | downregulated during regeneration - late |
| ENSDART00000135403 | nuak2 | 0.022463379 | -0.203159013 | -0.012266437 | -0.34684763 | -0.520334493 | downregulated during regeneration - late |
| ENSDART00000135556 | lonrf1 | 0.00148037 | -0.330548829 | 0.12250287 | -0.445965292 | -0.332349033 | downregulated during regeneration - late |
| ENSDART00000136049 | syt10 | 0.00992455 | -0.382437751 | -0.322146795 | -0.465356993 | -0.383104059 | downregulated during regeneration - late |
| ENSDART00000136084 | mfhas1 | 0.005371229 | -0.229093837 | -0.123648609 | -0.428888075 | -0.346098599 | downregulated during regeneration - late |
| ENSDART00000136190 | si:ch211-133l5.7 | 0.008776907 | 0.046455816 | -0.284643729 | -0.455348095 | -0.512893839 | downregulated during regeneration - late |
| ENSDART00000136538 | cish | 0.003012303 | -0.292880454 | -0.45612089 | -0.971812918 | -1.16494411 | downregulated during regeneration - late |
| ENSDART00000138308 | gpr186 | 0.002651594 | -1.473279373 | -0.953004046 | -1.169373422 | -1.314238919 | downregulated during regeneration - late |
| ENSDART00000138661 | tspan11 | 0.00016317 | -0.312645804 | -0.301551652 | -0.855364566 | -0.684847545 | downregulated during regeneration - late |
| ENSDART00000139174 | fam46c | 0.014064789 | -0.16066992 | -0.125475027 | -0.118453643 | -0.616919987 | downregulated during regeneration - late |
| ENSDART00000139701 | zfand5a | 0.001239881 | -0.357581428 | -0.278083252 | -0.905903537 | -0.782247372 | downregulated during regeneration - late |
| ENSDART00000140779 | fundc1 | 0.026565285 | -1.365145935 | -0.380101108 | -0.950161557 | -1.079798871 | downregulated during regeneration - late |
| ENSDART00000140828 | CU179758.1 | 0.006872094 | -0.432947507 | -0.14249918 | -0.435392677 | -0.381420161 | downregulated during regeneration - late |
| ENSDART00000141030 | pde4ca | 0.026489121 | -0.308097694 | -0.281644648 | -0.391359015 | -0.416436171 | downregulated during regeneration - late |
| ENSDART00000141345 | si:ch211-133l5.7 | 0.002874771 | 0.067098833 | -0.266407921 | -0.458017208 | -0.557765253 | downregulated during regeneration - late |
| ENSDART00000141437 | sdc2 | 0.011417724 | 0.001568967 | 0.007072686 | -0.196613876 | -0.36025161 | downregulated during regeneration - late |
| ENSDART00000141487 | pvrl3b | 0.003733102 | -0.333456043 | -0.273917158 | -0.488003853 | -0.734425178 | downregulated during regeneration - late |
| ENSDART00000141671 | socs2 | 0.000634801 | -0.321775893 | -0.091083829 | -1.001751456 | -1.186845404 | downregulated during regeneration - late |
| ENSDART00000141920 | si:ch211-224l10.4 | 0.059256345 | -1.330425532 | -0.617989075 | -0.986788428 | -0.796603249 | downregulated during regeneration - late |
| ENSDART00000142112 | vma21 | 0.030307206 | -0.082665202 | -0.082534524 | -0.180302218 | -0.34518071 | downregulated during regeneration - late |
| ENSDART00000142196 | fam167ab | 0.003623005 | -0.920556961 | -0.832834787 | -1.167437341 | -1.267796279 | downregulated during regeneration - late |
| ENSDART00000142735 | si:ch211-13f8.1 | 0.003979354 | -0.324319655 | -0.241821619 | -0.588481479 | -0.554389586 | downregulated during regeneration - late |
| ENSDART00000142851 | itih6 | 0.001733341 | -0.217828278 | -0.246312318 | -4.361209214 | -4.055941297 | downregulated during regeneration - late |
| ENSDART00000143200 | kcnma1a | 0.037170694 | -0.226894558 | -0.334282966 | -0.480968248 | -0.397036214 | downregulated during regeneration - late |
| ENSDART00000143840 | gstk1 | 0.026706504 | 0.057517576 | -0.140839441 | -0.201267582 | -0.834065155 | downregulated during regeneration - late |
| ENSDART00000144067 | cabp5a | 0.012049052 | -0.477283062 | -0.255392436 | -0.467866686 | -0.403996112 | downregulated during regeneration - late |
| ENSDART00000145282 | sepp1a | 0.000461528 | -0.462981641 | -0.396376841 | -0.662591735 | -0.92136404 | downregulated during regeneration - late |
| ENSDART00000145558 | si:ch211-237l4.6 | 0.040053922 | -0.183280044 | -0.231325216 | -0.614649778 | -0.510830341 | downregulated during regeneration - late |
| ENSDART00000145762 | dnaaf1 | 0.000461528 | -0.407466904 | -0.372899622 | -0.590562671 | -0.565268948 | downregulated during regeneration - late |
| ENSDART00000146472 | kcng4a | 0.010687727 | -0.174962319 | -0.077658154 | -0.410630934 | -0.442662471 | downregulated during regeneration - late |
| ENSDART00000146892 | abcg2d | 0.014710738 | 0.141142908 | -0.029073312 | -0.499348256 | -1.743973024 | downregulated during regeneration - late |
| ENSDART00000147068 | olfml2ba | 0.013360877 | -0.192176717 | 0.036009408 | -0.274084705 | -0.587185898 | downregulated during regeneration - late |
| ENSDART00000147201 | slc5a7a | 0.014511734 | -0.334790717 | -0.440092664 | -0.539424926 | -0.607892581 | downregulated during regeneration - late |
| ENSDART00000147582 | agmo | 0.00862856 | -0.172065266 | -0.060434285 | -0.409289881 | -0.408628944 | downregulated during regeneration - late |
| ENSDART00000147854 | mfap5 | 2.23813E-05 | -0.362752481 | -0.424642186 | -0.795916329 | -0.74190928 | downregulated during regeneration - late |
| ENSDART00000148456 | abcb4 | 0.020586406 | -0.551564057 | -0.079040965 | -0.313501093 | -1.124934266 | downregulated during regeneration - late |
| ENSDART00000149222 | pglyrp2 | 0.005895019 | -0.019258678 | -0.204426336 | -0.652532049 | -2.674820692 | downregulated during regeneration - late |
| ENSDART00000149574 | ca14 | 0.025606413 | -0.121794187 | -0.18625006 | -0.561537732 | -0.325393218 | downregulated during regeneration - late |
| ENSDART00000149652 | AL928685.4 | 0.043205592 | -0.221690272 | -0.265792933 | -0.407449839 | -0.293257945 | downregulated during regeneration - late |
| ENSDART00000149659 | ezh1 | 0.003513906 | 0.045526141 | 0.005041587 | -0.251417621 | -0.327846848 | downregulated during regeneration - late |
| ENSDART00000149689 | mob1bb | 0.002365371 | -0.2705299 | -0.18213785 | -0.463988827 | -0.361129298 | downregulated during regeneration - late |
| ENSDART00000149816 | hspb1 | 0.017271665 | -0.566005876 | -0.156876752 | -0.219080206 | -0.313669761 | downregulated during regeneration - late |
| ENSDART00000150386 | si:ch73-6k14.2 | 0.000164423 | -0.397499259 | -0.321996107 | -1.565555621 | -1.132350175 | downregulated during regeneration - late |
| ENSDART00000152068 | AL590149.1 | 0.016178224 | -0.322191556 | -0.083241136 | -0.42500156 | -0.283301232 | downregulated during regeneration - late |
| ENSDART00000152145 | si:ch211-207i1.2 | 0.024547153 | -1.105453902 | -1.700204988 | -2.013950338 | -1.361589232 | downregulated during regeneration - late |
| ENSDART00000152253 | kcnj13 | 0.002332929 | -0.175334575 | -0.22735068 | -0.506366898 | -0.552648836 | downregulated during regeneration - late |
| ENSDART00000152942 | jcada | 0.004739684 | -0.053928395 | -0.076278472 | -0.332447962 | -0.296311136 | downregulated during regeneration - late |
| ENSDART00000154141 | BX649398.2 | 0.000216813 | 0.278828233 | 0.21540553 | -0.861481785 | -0.785329833 | downregulated during regeneration - late |
| ENSDART00000155838 | luzp1 | 0.061460285 | -0.38633981 | -0.211709262 | -0.146984907 | -0.314685818 | downregulated during regeneration - late |
| ENSDART00000156256 | angpt4 | 0.024383427 | -0.447666401 | -0.531623063 | -0.584960614 | -0.899124282 | downregulated during regeneration - late |
| ENSDART00000156509 | fam150bb | 0.026883176 | -0.200699582 | -0.03489316 | -0.310837396 | -0.503245808 | downregulated during regeneration - late |
| ENSDART00000157338 | gstz1 | 0.062825179 | -0.391223456 | -0.14748942 | -0.590184545 | -0.319150237 | downregulated during regeneration - late |
| ENSDART00000158260 | c1galt1a | 0.000459643 | 0.184684125 | 0.138397385 | -0.286164045 | -0.509177074 | downregulated during regeneration - late |
| ENSDART00000159588 | rnf24 | 0.016665414 | -0.199572807 | -0.336306858 | -0.290748993 | -0.299485771 | downregulated during regeneration - late |
| ENSDART00000159752 | sort1a | 0.002661298 | -0.095510148 | -0.060577332 | -0.362899297 | -0.357946523 | downregulated during regeneration - late |
| ENSDART00000159908 | camk1db | 0.001174125 | -0.548571048 | -0.478436516 | -0.873630088 | -0.641909911 | downregulated during regeneration - late |
| ENSDART00000160305 | si:zfos-943e10.1 | 4.9299E-05 | -0.732433247 | -0.018482138 | -0.2347001 | -0.520119378 | downregulated during regeneration - late |

| | | | | | | | |
|---|---|---|---|---|---|---|---|
| ENSDART00000160425 | LAMP1 | 0.002170917 | -0.291224845 | -0.711008477 | -0.768820578 | -0.726430297 | downregulated during regeneration - late |
| ENSDART00000160431 | dhrs13l1 | 0.026954719 | -0.306310555 | -0.206975564 | -0.316329966 | -0.293060295 | downregulated during regeneration - late |
| ENSDART00000160707 | smx5 | 0.001677265 | -0.006041688 | -0.141079427 | -0.427667941 | -0.488718653 | downregulated during regeneration - late |
| ENSDART00000160829 | mhc1zda | 0.014447102 | -0.410179214 | -0.276613879 | -0.418038837 | -0.322538004 | downregulated during regeneration - late |
| ENSDART00000160955 | slc38a3b | 0.012986721 | -0.475545538 | -0.367921084 | -0.471714594 | -0.850494481 | downregulated during regeneration - late |
| ENSDART00000160956 | oaz1b | 0.012598032 | -0.133855172 | -0.165932835 | -0.380106363 | -0.375655267 | downregulated during regeneration - late |
| ENSDART00000161525 | zgc:171704 | 0.000191101 | 0.267822787 | -0.368878081 | -1.0813007 | -1.211102226 | downregulated during regeneration - late |
| ENSDART00000161932 | npr3 | 0.00058454 | 0.255355781 | 0.183557252 | -0.357151542 | -1.063769364 | downregulated during regeneration - late |
| ENSDART00000162010 | zgc:92140 | 0.011458427 | -0.256831407 | -0.245054913 | -0.345912399 | -0.346929099 | downregulated during regeneration - late |
| ENSDART00000162359 | sorbs1 | 0.007620644 | -0.104497249 | -0.20837462 | -0.418151878 | -0.317718588 | downregulated during regeneration - late |
| ENSDART00000162564 | CT027980.1 | 0.000781831 | -0.351165538 | -0.178285414 | -0.533647461 | -0.500417084 | downregulated during regeneration - late |
| ENSDART00000162587 | chd5 | 0.003143898 | -0.534129477 | -1.024077887 | -0.70517204 | -0.958580386 | downregulated during regeneration - late |
| ENSDART00000163305 | fxyd6l | 0.013876441 | -0.209100518 | -0.188713869 | -0.620315584 | -0.44198331 | downregulated during regeneration - late |
| ENSDART00000163394 | dgke | 0.019521843 | -0.154264184 | -0.257555682 | -0.346312178 | -0.3002364 | downregulated during regeneration - late |
| ENSDART00000163565 | plcl2 | 0.005353377 | -0.007453137 | -0.119106713 | -0.251934099 | -0.322673112 | downregulated during regeneration - late |
| ENSDART00000164729 | SBSPON | 0.014479917 | -0.213183167 | -0.151008123 | -0.380923154 | -0.573582751 | downregulated during regeneration - late |
| ENSDART00000164791 | fkbp3 | 0.030292962 | -0.300438451 | -0.231059822 | -0.287267966 | -0.464270989 | downregulated during regeneration - late |
| ENSDART00000165290 | cyb5a | 0.002005251 | 0.19405994 | 0.064877953 | -0.350823438 | -0.576368917 | downregulated during regeneration - late |
| ENSDART00000165570 | rgs3a | 0.01962183 | -0.252183908 | -0.25277418 | -0.390798302 | -0.347427277 | downregulated during regeneration - late |
| ENSDART00000166105 | frem1a | 0.02305984 | -0.140882698 | -0.093532592 | -0.525495905 | -1.018483252 | downregulated during regeneration - late |
| ENSDART00000166449 | pik3r3a | 0.049250284 | -0.698577911 | -0.3007013 | -0.248394691 | -0.547713549 | downregulated during regeneration - late |
| ENSDART00000166527 | ftr87 | 0.023542744 | -0.180960216 | -0.104077306 | -0.471569991 | -0.343688512 | downregulated during regeneration - late |
| ENSDART00000166829 | NA | 0.030198325 | -0.799969413 | -0.359199835 | -0.712167572 | -2.039500454 | downregulated during regeneration - late |
| ENSDART00000167330 | CU179758.1 | 0.003900528 | -0.385159517 | -0.273959547 | -0.55992419 | -0.56237529 | downregulated during regeneration - late |
| ENSDART00000167506 | scg2a | 0.003405768 | -0.457333464 | -0.29124415 | -0.144155502 | -0.510044484 | downregulated during regeneration - late |
| ENSDART00000167824 | timp4.3 | 0.01037926 | -0.389539203 | -0.013931581 | -0.399789503 | -0.590224284 | downregulated during regeneration - late |
| ENSDART00000167963 | nrg1 | 0.037106045 | -0.278608256 | -0.388339035 | -0.420420553 | -0.389329729 | downregulated during regeneration - late |
| ENSDART00000168201 | golim4a | 0.009167687 | -0.134763119 | -0.005515007 | -0.245821925 | -1.172515705 | downregulated during regeneration - late |
| ENSDART00000168228 | tmem184a | 0.016284951 | -0.3188994 | -0.465602708 | -0.713259634 | -0.580015552 | downregulated during regeneration - late |
| ENSDART00000168278 | si:ch211-276i12.4 | 0.002327436 | -0.631335043 | -0.270754342 | -0.641142033 | -0.623314164 | downregulated during regeneration - late |
| ENSDART00000168750 | smarca1 | 0.005353377 | -0.162085202 | -0.116464254 | -0.382463649 | -0.380530416 | downregulated during regeneration - late |
| ENSDART00000169228 | vat1l | 0.00721984 | 0.089392965 | -0.09841303 | -0.250056266 | -0.442720544 | downregulated during regeneration - late |
| ENSDART00000169380 | slc25a43 | 0.005026727 | -0.226077688 | 0.088930874 | -0.355915914 | -0.508574161 | downregulated during regeneration - late |
| ENSDART00000170381 | CR855389.1 | 0.046578416 | -0.538290002 | -0.347216121 | -0.566215377 | -0.407287476 | downregulated during regeneration - late |
| ENSDART00000171006 | hpcal4 | 0.014069453 | -0.435402713 | -0.198074705 | -0.466665783 | -0.338540363 | downregulated during regeneration - late |
| ENSDART00000171417 | si:ch73-299h12.3 | 0.004952764 | -0.719381589 | -0.760232748 | -0.826233659 | -0.863901262 | downregulated during regeneration - late |
| ENSDART00000171815 | abcc8b | 0.014959825 | 0.00284442 | -0.044162256 | -0.758748776 | -0.514944979 | downregulated during regeneration - late |
| ENSDART00000171824 | si:ch211-227e10.2 | 0.024734469 | -0.663768919 | -0.251157559 | -0.496668883 | -0.404241953 | downregulated during regeneration - late |
| ENSDART00000172019 | CU694219.1 | 0.010271025 | -0.358471879 | -0.088786066 | -0.423165049 | -1.570028426 | downregulated during regeneration - late |
| ENSDART00000172357 | tmem132a | 0.051805764 | -0.246648072 | -0.102389273 | -0.224742481 | -0.519178406 | downregulated during regeneration - late |
| ENSDART00000172672 | angptl2b | 0.010046682 | -0.321678324 | -0.001662718 | -0.330394771 | -0.841768465 | downregulated during regeneration - late |
| ENSDART00000173237 | mid2 | 0.032645202 | -0.115282743 | -0.310001722 | -0.367171215 | -0.341782639 | downregulated during regeneration - late |

**Table S3. Ingenuity Pathway Analysis of temporally clustered, differentially expressed genes.**

| Ingenuity Canonical Pathways | -log(p-value) | Ratio | cluster | genes | collapsed pathways |
|---|---|---|---|---|---|
| EIF2 Signaling | 23.8 | 0.163 | growth toward the midline | RPL36A,RPS27,RPS23,RPS18,RPL39,Rpl22l1,RPL35A,RPS4X,RPL7,RPS11,RPS28,RPS7,RPL35,MAP2K2,RPS13,UBA52,RPS9,MRAS,RPL21,RPS17,RPL18,RPS24,RPL4,RPL30,RPL12,RPL23,RPS21,RPL10A,RPL27,RPL15,RPL8,PIK3CB,RPS15A,RPS25,RPL38,RPSA | EIF2 Signaling |
| mTOR Signaling | 12 | 0.114 | growth toward the midline | PRKCQ,PLD3,RPS27,RHOC,RPS23,RPS18,RPS21,RPS4X,RPS11,RPS28,RPS7,HMOX1,RHOG,RPS13,PRKCD,RPS9,MRAS,RPS25,RPS15A,PIK3CB,RPS17,RPS24,RPSA | mTOR Signaling |
| Regulation of eIF4 and p70S6K Signaling | 9.52 | 0.115 | growth toward the midline | RPS27,RPS23,RPS18,RPS21,RPS4X,RPS11,RPS28,RPS7,MAP2K2,RPS13,RPS9,MRAS,RPS25,PIK3CB,RPS15A,RPS17,RPS24,RPSA | mTOR Signaling |
| Phagosome Maturation | 6.48 | 0.0946 | growth toward the midline | TUBB4B,TCIRG1,ATP6AP1,CTSZ,CTSV,CTSA,ATP6V1H,TUBA8,NCF2,CTSB,CTSC,NAPA,BET1L,ATP6V0E1 | Phagosome Maturation |
| IL-8 Signaling | 5.69 | 0.0761 | growth toward the midline | PRKCQ,PLD3,RHOC,CSTB,BCL2L1,HMOX1,ITGAM,RHOG,MAP2K2,PRKCD,NCF2,ITGAV,MRAS,PIK3CB,MMP9 | IL-8 Signaling |
| IL-3 Signaling | 4 | 0.0964 | growth toward the midline | JAK1,PRKCQ,MAP2K2,PRKCD,MRAS,PIK3CB,STAT3,INPP5D | IL-3 Signaling |
| EIF2 Signaling | 9.84 | 0.0995 | midline crossing | EIF2AK1,ATF3,EIF3H,RRAS,HRAS,TRIB3,EIF2S3,EIF2S1,EIF2S2,RPL9,EIF3M,EIF3F,RRAS2,EIF3B,EIF4A1,EIF3I,ATF4,RPL5,AGO1,RPS12,ACTC1,EIF3L | EIF2 Signaling |
| Regulation of eIF4 and p70S6K Signaling | 9.15 | 0.115 | midline crossing | EIF3H,RRAS,HRAS,EIF2S3,EIF2S1,MAPK11,EIF2S2,EIF4EBP1,EIF3M,EIF3F,RRAS2,EIF3B,EIF4A1,MKNK1,EIF3I,RPS12,AGO1,EIF3L | mTOR Signaling |
| Germ Cell-Sertoli Cell Junction Signaling | 7.65 | 0.0983 | midline crossing | SRC,RRAS,TUBB4B,RAC1,ITGA6,ILK,TUBA4A,MAPK9,HRAS,GSN,LIMK1,RRAS2,TUBA3E,JUP,ACTC1,ACTG1,CTNND1 | RAS/MAPK signaling |
| Agrin Interactions at Neuromuscular Junction | 7.31 | 0.159 | midline crossing | SRC,RRAS2,JUN,RRAS,RAC1,ITGA6,MAPK9,HRAS,ACTC1,ACTG1,ITGB3 | agrin interactions |
| Virus Entry via Endocytic Pathways | 6.93 | 0.118 | midline crossing | SRC,AP2M1,RRAS,HLA-B,ITGA6,ABL1,RAC1,HRAS,ITGB3,RRAS2,FLNC,ACTC1,ACTG1 | endocytic pathways |
| Integrin Signaling | 6.87 | 0.0822 | midline crossing | RAP2B,RAPGEF1,SRC,RRAS,ARF1,RAC1,ITGA6,ILK,ABL1,HRAS,TNK2,GSN,ITGB3,CAPNS1,RRAS2,ACTC1,ACTG1,MYL12A | integrin signaling |
| Epithelial Adherens Junction Signaling | 6.28 | 0.0959 | midline crossing | SRC,RAPGEF1,MYL6,RRAS,TUBB4B,RAC1,TUBA4A,HRAS,RRAS2,JUP,TUBA3E,ACTC1,ACTG1,CTNND1 | adhesive junctions |
| Sertoli Cell-Sertoli Cell Junction Signaling | 5.96 | 0.0843 | midline crossing | SRC,RRAS,TUBB4B,RAC1,ILK,TUBA4A,MAPK9,HRAS,MAPK11,JUN,RRAS2,JUP,TUBA3E,ACTC1,ACTG1 | adhesive junctions |
| Paxillin Signaling | 5.93 | 0.106 | midline crossing | SRC,RRAS2,RRAS,ARF1,RAC1,ITGA6,MAPK9,HRAS,MAPK11,ACTC1,ACTG1,ITGB3 | adhesive junctions |
| Estrogen-Dependent Breast Cancer Signaling | 5.69 | 0.125 | midline crossing | SRC,RRAS2,JUN,RRAS,CREB3,DHRS11,HSD17B7,HRAS,ATF4,STAT5B | RAS/MAPK signaling |
| Molecular Mechanisms of Cancer | 5.36 | 0.0558 | midline crossing | RAP2B,RAPGEF1,SRC,ARHGEF4,RRAS,RAC1,APAF1,ABL1,MAPK9,HRAS,BAX,E2F3,MAPK11,FAS,RB1,JUN,RRAS2,CASP9,NFKBIA,ARHGEF2,SMAD1,CTNND1 | RAS/MAPK signaling |
| Remodeling of Epithelial Adherens Junctions | 5.33 | 0.13 | midline crossing | SRC,NME1,TUBB4B,MAPRE1,TUBA4A,TUBA3E,ACTC1,ACTG1,CTNND1 | adhesive junctions |
| Prostate Cancer Signaling | 4.93 | 0.103 | midline crossing | RB1,NFKBIA,CASP9,RRAS2,HSP90AB1,RRAS,CREB3,ABL1,HRAS,ATF4 | RAS/MAPK signaling |
| UVC-Induced MAPK Signaling | 4.91 | 0.163 | midline crossing | SRC,RRAS2,JUN,RRAS,MAPK9,HRAS,MAPK11 | RAS/MAPK signaling |
| NRF2-mediated Oxidative Stress Response | 4.85 | 0.0725 | midline crossing | USP14,RRAS,PRDX1,MAPK9,HRAS,DNAJC21,JUN,RRAS2,VCP,ATF4,TXN,FKBP5,ACTC1,ACTG1 | NRF2-mediated Oxidative Stress Response |
| Role of Tissue Factor in Cancer | 4.73 | 0.0887 | midline crossing | SRC,RRAS2,RRAS,RAC1,ITGA6,HRAS,STAT5B,MAPK11,F3,ITGB3,LIMK1 | RAS/MAPK signaling |
| mTOR Signaling | 4.65 | 0.0697 | midline crossing | EIF3H,RRAS,RAC1,FKBP1A,HRAS,EIF4EBP1,EIF3M,EIF3F,RRAS2,EIF3B,EIF4A1,EIF3I,RPS12,EIF3L | mTOR Signaling |
| Apoptosis Signaling | 4.38 | 0.1 | midline crossing | CAPNS1,NFKBIA,CASP9,RRAS2,RRAS,APAF1,HRAS,BAX,FAS | apoptosis signaling |
| Myc Mediated Apoptosis Signaling | 4.38 | 0.114 | midline crossing | CASP9,RRAS2,RRAS,APAF1,MAPK9,HRAS,BAX,FAS | apoptosis signaling |
| Caveolar-mediated Endocytosis Signaling | 4.33 | 0.113 | midline crossing | SRC,FLNC,HLA-B,ITGA6,ABL1,ACTC1,ACTG1,ITGB3 | endocytic pathways |
| Death Receptor Signaling | 4.27 | 0.0968 | midline crossing | NFKBIA,CASP9,APAF1,PARP8,PARP12,ACTC1,ACTG1,FAS,LIMK1 | apoptosis signaling |
| tRNA Charging | 4.14 | 0.154 | midline crossing | RARS,GARS,TARS,SARS,IARS,QARS | tRNA Charging |
| ILK Signaling | 4.12 | 0.066 | midline crossing | FN1,MYL6,FERMT2,CREB3,ILK,MAPK9,ITGB3,JUN,FLNC,ATF4,RSU1,ACTC1,ACTG1 | integrin signaling |
| ATM Signaling | 4.09 | 0.0918 | midline crossing | NFKBIA,GADD45B,JUN,CREB3,ABL1,MAPK9,ATF4,TDP1,MAPK11 | ATM Signaling |
| Superpathway of Cholesterol Biosynthesis | 13.1 | 0.393 | target selection | MVD,SQLE,EBP,ACAT2,DHCR24,HSD17B7,MSMO1,TM7SF2,HMGCS1,SC5D,CYP51A1 | cholesterol biosynthesis |
| Cholesterol Biosynthesis I | 11.7 | 0.615 | target selection | SQLE,EBP,DHCR24,HSD17B7,MSMO1,TM7SF2,SC5D,CYP51A1 | cholesterol biosynthesis |
| Cholesterol Biosynthesis II (via 24,25-dihydrolanosterol) | 11.7 | 0.615 | target selection | SQLE,EBP,DHCR24,HSD17B7,MSMO1,TM7SF2,SC5D,CYP51A1 | cholesterol biosynthesis |
| Cholesterol Biosynthesis III (via Desmosterol) | 11.7 | 0.615 | target selection | SQLE,EBP,DHCR24,HSD17B7,MSMO1,TM7SF2,SC5D,CYP51A1 | cholesterol biosynthesis |
| Actin Cytoskeleton Signaling | 9.06 | 0.0837 | target selection | MYH4,FN1,SOS2,HRAS,FGD1,MYH7,BCAR1,RAC3,SSH1,APC,PIK3R3,SHC1,PAK1,CYFIP2,FGF18,PAK2,ARHGAP35,ACTC1,MYL3 | actin cytoskeleton signaling |
| Breast Cancer Regulation by Stathmin1 | 8.94 | 0.0878 | target selection | TUBA1B,PPP2R5B,SOS2,TUBB2A,TUBA4A,HRAS,TUBB,PPP1R14B,TUBB2B,PIK3R3,SHC1,PPP2CB,PAK1,CAMK2D,TUBA1A,GNB2,TUBA1C,PPP2R1B | axon guidance |
| Axonal Guidance Signaling | 8.59 | 0.0574 | target selection | TUBA1B,PLXNA3,NRP2,BMP3,SOS2,TUBB2A,TUBA4A,ADAM22,HRAS,L1CAM,DPYSL5,TUBB,BCAR1,RAC3,TUBB2B,PIK3R3,SHC1,PAK1,EPHB1,TUBA1A,PAK2,GNB2,SRGAP2,TUBA1C,MYL3,NRP1 | axon guidance |
| Epithelial Adherens Junction Signaling | 7.57 | 0.0959 | target selection | MYH4,TUBA1B,TUBB2A,TUBA4A,HRAS,MYH7,TUBB,APC,TUBB2B,TUBA1A,TGFB2,TUBA1C,ACTC1,MYL3 | adhesive junctions |
| Germ Cell-Sertoli Cell Junction Signaling | 7.47 | 0.0867 | target selection | TUBA1B,TUBB2A,TUBA4A,HRAS,TUBB,RAC3,BCAR1,TUBB2B,PIK3R3,PAK1,TUBA1A,PAK2,TGFB2,TUBA1C,ACTC1 | adhesive junctions |
| Remodeling of Epithelial Adherens Junctions | 7.27 | 0.145 | target selection | TUBA1B,TUBA1A,TUBB2A,TUBA4A,TUBA1C,TUBB,ACTC1,APC,TUBB2B,Mapre2 | adhesive junctions |
| 14-3-3-mediated Signaling | 6.35 | 0.0916 | target selection | PIK3R3,TUBA1B,CBL,TUBA1A,YWHAH,TUBB2A,TUBA4A,HRAS,VIM,TUBA1C,TUBB,TUBB2B | 14-3-3-mediated signaling |
| Zymosterol Biosynthesis | 6.21 | 0.667 | target selection | HSD17B7,MSMO1,TM7SF2,CYP51A1 | cholesterol biosynthesis |
| ERK/MAPK Signaling | 5.88 | 0.07 | target selection | YWHAH,PPP2R5B,SOS2,HRAS,PPP1R14B,RAC3,BCAR1,PIK3R3,SHC1,PPP2CB,PAK1,PAK2,STAT1,PPP2R1B | ERK/MAPK signaling |
| Gap Junction Signaling | 5.27 | 0.0667 | target selection | PIK3R3,CSNK1E,TUBA1B,DBN1,TUBA1A,TUBB2A,SOS2,TUBA4A,HRAS,TUBA1C,TUBB,ACTC1,TUBB2B | gap junction signaling |
| Phagosome Maturation | 4.99 | 0.0743 | target selection | DYNC1H1,TUBA1B,Dync1i2,TUBA1A,VPS33A,DYNC1I1,TUBB2A,TUBA4A,TUBA1C,TUBB,TUBB2B | phagasome maturation |
| PI3K/AKT Signaling | 4.86 | 0.08 | target selection | PIK3R3,SHC1,RPS6KB1,PPP2CB,NFKBIA,YWHAH,SOS2,PPP2R5B,HRAS,PPP2R1B | PI3/AKT signaling |
| PTEN Signaling | 4.24 | 0.0756 | target selection | PIK3R3,SHC1,RPS6KB1,CBL,YWHAH,SOS2,HRAS,BCAR1,RAC3 | PTEN signaling |
| Integrin Signaling | 4.07 | 0.0548 | target selection | PIK3R3,SHC1,PAK1,ARF3,PAK2,SOS2,RALB,HRAS,ITGA10,BCAR1,ACTC1,RAC3 | integrin signaling |
| Ephrin B Signaling | 4.06 | 0.0959 | target selection | EPHB1,PAK1,CBL,ABI1,GNB2,HRAS,RAC3 | ephrin B signaling |
| Glutamate Receptor Signaling | 12.2 | 0.263 | brain innervation | GRIN1,GRM2,GRIN2A,GRM8,GRM1,SLC1A1,GRIP1,HOMER3,GRM4,GNB1,GRM6,SLC1A2,HOMER1,GRIA3,GRIK1 | glutamate receptor signaling |
| Synaptic Long Term Depression | 8.98 | 0.115 | brain innervation | GRM2,CACNA2D2,GUCY1A1,GRM8,GRM1,RYR2,CACNA1H,CACNB3,GRM4,GNAZ,CACNG3,IGF1,GNAO1,PPM1L,RYR3,GRM6,PRKCE,PPP2R2C,CACNB2,GRIA3 | synaptic LTD |
| Calcium Signaling | 8.44 | 0.102 | brain innervation | RAP2A,GRIN1,GRIN2A,CACNA2D2,CHRNA6,RYR2,MEF2A,SLC8A3,CACNA1H,SLC8A2,CACNB3,ATP2B2,CACNG3,ATP2B3,RYR3,CACNB2,CHRNB3,SLC8A1,PPP3CA,GRIA3,GRIK1 | calcium signaling |
| CREB Signaling in Neurons | 8.22 | 0.0991 | brain innervation | GRIN1,ADCY2,GRIN2A,GRM2,CACNA2D2,GRM8,GRM1,CACNA1H,GNG13,CACNB3,GRM4,GNAZ,PIK3R3,GNB1,CACNG3,GNAO1,GRM6,PRKCE,CACNB2,GRIK1,GRIA3 | CREB signaling |
| GABA Receptor Signaling | 7.85 | 0.147 | brain innervation | GABRA5,ADCY2,CACNA2D2,CACNA1H,GABBR1,GPHN,CACNB3,CACNG3,DNM1,NSF,GABBR2,GABRB1,CACNB2,GABRA2 | GABA receptor signaling |
| Opioid Signaling Pathway | 6.52 | 0.0823 | brain innervation | GRIN1,ADCY2,GRIN2A,CACNA2D2,RYR2,RGS7,CACNA1H,CACNB3,RGS6,RAC3,SLC12A5,GNB1,CACNG3,ARRB1,GNAO1,RYR3,PRKCE,RGS11,CACNB2,PPP3CA | opiod signaling |
| Axonal Guidance Signaling | 6.25 | 0.0618 | brain innervation | EPHB2,GNG13,SEMA6B,ADAM11,PLXNA2,GNB1,EPHB6,NTNG1,IGF1,NTNG2,UNC5D,PRKCE,ADAM23,TUBB4A,ROBO2,PPP3CA,EFNA2,SLIT2,GNAZ,ROBO3,RAC3,PIK3R3,SEMA3A,EPHA6,ADAM12,WNT10A,GNAO1,LINGO1 | axon guidance |
| Netrin Signaling | 5.96 | 0.154 | brain innervation | CACNG3,CACNA2D2,RYR2,RYR3,CACNA1H,UNC5D,CACNB2,CACNB3,RAC3,PPP3CA | netrin signaling |
| Role of NFAT in Cardiac Hypertrophy | 4.69 | 0.0731 | brain innervation | ADCY2,CACNA2D2,MEF2A,CACNA1H,SLC8A3,GNG13,SLC8A2,CACNB3,PIK3R3,GNB1,CACNG3,IGF1,PRKCE,CACNB2,SLC8A1,PPP3CA | NFAT signaling |
| nNOS Signaling in Skeletal Muscle Cells | 4.64 | 0.171 | brain innervation | CACNG3,CACNA2D2,RYR2,RYR3,CACNA1H,CACNB2,CACNB3 | nNOS signaling |
| Chondroitin Sulfate Biosynthesis | 4.57 | 0.14 | brain innervation | CHST1,SULT4A1,XYLT1,HS3ST2,B3GAT1,HS6ST3,HS3ST4,B3GAT2 | glycosaminoglycan biosynthesis |
| G Beta Gamma Signaling | 4.48 | 0.0965 | brain innervation | CACNG3,GNB1,ADCY2,CACNA2D2,GNAO1,CACNA1H,GNG13,PRKCE,CACNB2,CACNB3,GNAZ | GPCR signaling |
| Dermatan Sulfate Biosynthesis | 4.46 | 0.136 | brain innervation | CHST1,SULT4A1,XYLT1,HS3ST2,B3GAT1,HS6ST3,HS3ST4,B3GAT2 | glycosaminoglycan biosynthesis |
| Gαi Signaling | 4.28 | 0.0917 | brain innervation | GNB1,GABBR2,ADCY2,GRM2,ADRA2A,GRM8,RGS7,GRM6,GNG13,GABBR1,GRM4 | GPCR signaling |
| α-Adrenergic Signaling | 4 | 0.103 | brain innervation | GNB1,ADCY2,GYS1,ADRA2A,SLC8A3,GNG13,PRKCE,SLC8A2,SLC8A1 | adrenergic signaling |

| | | | | | |
|---|---|---|---|---|---|
| GABA Receptor Signaling | 17.8 | 0.242 | downregulated during regeneration - early | CACNA1G,CACNA1I,CACNG6,SLC32A1,AP2A1,CACNA1D,ADCY3,GABBR1,GABRB2,CACNG2,GAD2,GABRG2,GPR37,GABRG1,CACNA1B,GABRA6,SLC6A1,CACNG7,CACNB2,GABRA1,CACNG8,GABRD,CACNA2D4 | GABA receptor signaling |
| Opioid Signaling Pathway | 13.6 | 0.119 | downregulated during regeneration - early | CACNA1I,AP2A1,CAMK4,SLC12A5,CACNG2,GNB3,CAMK2A,PPP3R1,CACNG7,CACNB2,CACNG8,CACNA2D4,PRKD1,PPP3CA,CACNA1G,GRIN1,CACNG6,CACNA1D,ADCY3,RPS6KA5,NPBWR2,FOSB,CACNA1B,KCNJ5,KCNJ9,PENK,RGS8,KCNJ6,CAMK2G | opiod signaling |
| Calcium Signaling | 12.8 | 0.126 | downregulated during regeneration - early | CACNA1I,CAMK4,ATP2B1,GRIA1,CACNG2,CAMK2A,PPP3R1,CACNG7,CACNB2,CACNG8,CACNA2D4,PPP3CA,CACNA1G,CACNG6,GRIN1,CACNA1D,TRPC1,CHRNA10,CACNA1B,ATP2B3,MEF2D,CAMKK1,RCAN3,SLC8A1,GRIA3,CAMK2G | calcium signaling |
| Role of NFAT in Cardiac Hypertrophy | 10.6 | 0.11 | downregulated during regeneration - early | CACNA1G,CACNA1I,CACNG6,CACNA1D,CAMK4,ADCY3,PLCH1,CACNG2,CAMK2A,GNB3,CACNA1B,MEF2D,PPP3R1,IGF1R,CACNG7,RCAN3,IRS2,CACNB2,CACNG8,SLC8A1,CACNA2D4,PRKD1,PPP3CA,CAMK2G | NFAT signaling |
| CREB Signaling in Neurons | 10.1 | 0.108 | downregulated during regeneration - early | CACNA1G,CACNA1I,CACNG6,GRIN1,CACNA1D,CAMK4,GRID2,GRIA1,ADCY3,PLCH1,CACNG2,GNB3,CAMK2A,GRIK4,CACNA1B,CACNG7,IRS2,CACNB2,CACNG8,CACNA2D4,PRKD1,CAMK2G,GRIA3 | CREB signaling |
| nNOS Signaling in Skeletal Muscle Cells | 9.36 | 0.268 | downregulated during regeneration - early | CACNA1I,CACNA1G,CACNG2,CACNG6,CACNA1D,CAMK4,CACNA1B,CACNG7,CACNB2,CACNG8,CACNA2D4 | nNOS signaling |
| Dopamine-DARPP32 Feedback in cAMP Signaling | 8.9 | 0.116 | downregulated during regeneration - early | KCNJ12,GRIN1,CACNA1D,CAMK4,PPP1R3C,ADCY3,DRD2,PLCH1,KCNJ11,DRD1,KCNJ14,KCNJ5,PPP3R1,CAMKK1,KCNJ9,DRD4,KCNJ6,PRKD1,PPP3CA | dopamine signaling |
| Corticotropin Releasing Hormone Signaling | 8.39 | 0.122 | downregulated during regeneration - early | CACNA1G,CACNA1I,CACNG6,CACNA1D,CAMK4,GUCY2D,ADCY3,CRH,CACNG2,CACNA1B,MEF2D,NR4A1,CACNG7,CACNB2,CACNG8,CACNA2D4,PRKD1 | corticotropin-releasing hormone signaling |
| Netrin Signaling | 8.14 | 0.185 | downregulated during regeneration - early | CACNA1I,CACNA1G,CACNG2,CACNG6,CACNA1D,CACNA1B,PPP3R1,CACNG7,CACNB2,CACNG8,CACNA2D4,PPP3CA | netrin signaling |
| G Beta Gamma Signaling | 7.91 | 0.132 | downregulated during regeneration - early | CACNA1I,CACNA1G,CACNG6,CACNA1D,CACNG2,GNB3,CACNA1B,KCNJ5,KCNJ9,CACNG7,CACNB2,KCNJ6,CACNG8,CACNA2D4,PRKD1 | GPCR signaling |
| Synaptic Long Term Depression | 7.68 | 0.103 | downregulated during regeneration - early | CACNA1G,CACNA1I,CACNG6,CACNA1D,GUCY2D,GRID2,GRIA1,CRH,PLCH1,CACNG2,CACNA1B,IGF1R,CACNG7,CACNB2,CACNG8,CACNA2D4,PRKD1,GRIA3 | synaptic LTD |
| GNRH Signaling | 7.26 | 0.103 | downregulated during regeneration - early | CACNA1G,CACNA1I,CACNG6,CACNA1D,CAMK4,MAP3K13,ADCY3,CACNG2,GNB3,CAMK2A,CACNA1B,CACNG7,CACNB2,CACNG8,CACNA2D4,PRKD1,CAMK2G | GNRH signaling |
| CCR5 Signaling in Macrophages | 7.14 | 0.137 | downregulated during regeneration - early | CACNA1I,CACNA1G,CACNG6,CACNA1D,CAMK4,CACNG2,GNB3,CACNA1B,CACNG7,CACNB2,CACNG8,CACNA2D4,PRKD1 | CCRG signaling |
| GPCR-Mediated Nutrient Sensing in Enteroendocrine Cells | 7.14 | 0.125 | downregulated during regeneration - early | CACNA1I,CACNA1G,CACNG6,CACNA1D,ADCY3,RAPGEF4,PLCH1,CACNG2,CACNA1B,CACNG7,CACNB2,CACNG8,CACNA2D4,PRKD1 | GPCR signaling |
| PKCθ Signaling in T Lymphocytes | 6.73 | 0.101 | downregulated during regeneration - early | CACNA1G,CACNA1I,CACNG6,CACNA1D,MAP3K13,CACNG2,CAMK2A,CACNA1B,PPP3R1,CACNG7,IRS2,CACNB2,CACNG8,CACNA2D4,PPP3CA,CAMK2G | |
| FcγRIIB Signaling in B Lymphocytes | 6.21 | 0.139 | downregulated during regeneration - early | CACNA1I,CACNA1G,CACNG2,CACNG6,CACNA1D,CACNA1B,CACNG7,CACNB2,IRS2,CACNG8,CACNA2D4 | |
| Type II Diabetes Mellitus Signaling | 5.44 | 0.0909 | downregulated during regeneration - early | CACNA1G,CACNG2,CACNA1I,CACNG6,CACNA1D,PRKAB1,CACNA1B,CACNG7,CACNB2,IRS2,CACNG8,CACNA2D4,PRKD1,KCNJ11 | |
| Androgen Signaling | 5.29 | 0.0949 | downregulated during regeneration - early | CACNA1I,CACNA1G,CACNG6,CACNA1D,CAMK4,CACNG2,GNB3,CACNA1B,CACNG7,CACNB2,CACNG8,CACNA2D4,PRKD1 | |
| Gap Junction Signaling | 4.89 | 0.0769 | downregulated during regeneration - early | TJP2,GUCY2D,GRIA1,ADCY3,GJA9,DRD2,GJC1,PLCH1,LPAR1,DRD1,PPP3R1,IRS2,PRKD1,PPP3CA,GRIA3 | gap junction signaling |
| Glutamate Receptor Signaling | 4.7 | 0.14 | downregulated during regeneration - early | GRIN1,CAMK4,GNB3,GRIK4,SLC17A7,GRID2,GRIA1,GRIA3 | glutamate receptor signaling |
| Synaptic Long Term Potentiation | 4.36 | 0.0902 | downregulated during regeneration - early | GRIN1,CAMK2A,CAMK4,GRIA1,PPP1R3C,PPP3R1,PLCH1,PPP3CA,PRKD1,CAMK2G,GRIA3 | synaptic LTD |
| cAMP-mediated signaling | 4.1 | 0.0658 | downregulated during regeneration - early | CAMK4,VIPR2,CHRM4,ADCY3,GABBR1,RAPGEF4,DRD2,CHRM5,CAMK2A,LPAR1,DRD1,PPP3R1,DRD4,PPP3CA,CAMK2G | cAMP-mediated signaling |
| Gustation Pathway | 4.08 | 0.0779 | downregulated during regeneration - early | CACNA1I,CACNA1G,CACNG2,CACNG6,CACNA1D,GNB3,CACNA1B,ADCY3,CACNG7,CACNB2,CACNG8,CACNA2D4 | |
| Cellular Effects of Sildenafil (Viagra) | 4.08 | 0.084 | downregulated during regeneration - early | SLC4A5,CACNG2,CACNG6,CACNA1D,CAMK4,GPR37,GUCY2D,ADCY3,CACNG7,CACNG8,PLCH1 | |
| Neuroinflammation Signaling Pathway | 4.08 | 0.0579 | downregulated during regeneration - early | GRIN1,GRIA1,GABBR1,GABRB2,GAD2,GABRG2,GABRG1,KCNJ5,PPP3R1,GABRA6,KCNJ9,SLC6A1,TLR7,IRS2,KCNJ6,GABRA1,GABRD,PPP3CA | neuroinflammation signaling |
| Phototransduction Pathway | 4.74 | 0.132 | downregulated during regeneration - late | PRKACB,GNB5,GUCY2F,SAG,RGS9BP,GNGT2,PDE6D | phototransduction |

**Table S4. Differentially accesible sequences with distance to nearest annotated genes and nearest differentially-expressed genes.**

| new_seqnames | chr | start | end | width | strand | overall_distance | overall_gene | overall_DE_distance | overall_DE_gene | timepoint |
|---|---|---|---|---|---|---|---|---|---|---|
| **chr1:27509-28008** | chr1 | 27509 | 28008 | 500 | * | 70 | eed | 15791 | cep97 | differentially open at 2dpi |
| **chr1:185007-185506** | chr1 | 185007 | 185506 | 500 | * | 11536 | grtp1a | -79577 | cenpe | differentially open at 2dpi |
| **chr1:58446298-58446797** | chr1 | 58446298 | 58446797 | 500 | * | -7231 | lpar2b | 133113 | mfap4 | differentially open at 2dpi |
| **chr1:58479677-58480176** | chr1 | 58479677 | 58480176 | 500 | * | -8 | thumpd1 | 166491 | mfap4 | differentially open at 2dpi |
| **chr1:58542861-58543360** | chr1 | 58542861 | 58543360 | 500 | * | 1938 | olfm2b | 229675 | mfap4 | differentially open at 2dpi |
| **chr1:58608084-58608583** | chr1 | 58608084 | 58608583 | 500 | * | 104 | parn | 294899 | mfap4 | differentially open at 2dpi |
| **chr2:51352621-51353120** | chr2 | 51352621 | 51353120 | 500 | * | 2 | ftr67 | 698402 | dad1 | differentially open at 2dpi |
| **chr2:52763439-52764389** | chr2 | 52763439 | 52764389 | 951 | * | 28200 | tle2c | -336627 | scn12aa | differentially open at 2dpi |
| **chr2:54965275-54965774** | chr2 | 54965275 | 54965774 | 500 | * | -970 | ankrd12 | -56216 | acss2l | differentially open at 2dpi |
| **chr2:56491466-56491965** | chr2 | 56491466 | 56491965 | 500 | * | 40208 | gpx4b | 40208 | gpx4b | differentially open at 2dpi |
| **chr3:52202-52701** | chr3 | 52202 | 52701 | 500 | * | -30528 | mhc1zfa | -3902825 | plbd1 | differentially open at 2dpi |
| **chr3:1454919-1455418** | chr3 | 1454919 | 1455418 | 500 | * | -7 | wbp2nl | -2500109 | plbd1 | differentially open at 2dpi |
| **chr3:1463472-1463971** | chr3 | 1463472 | 1463971 | 500 | * | 8547 | wbp2nl | -2491555 | plbd1 | differentially open at 2dpi |
| **chr3:4122604-4123103** | chr3 | 4122604 | 4123103 | 500 | * | 128600 | art4 | 167577 | plbd1 | differentially open at 2dpi |
| **chr3:4123577-4124076** | chr3 | 4123577 | 4124076 | 500 | * | 129572 | art4 | 168549 | plbd1 | differentially open at 2dpi |
| **chr3:5139540-5140039** | chr3 | 5139540 | 5140039 | 500 | * | 17995 | tefb | 227365 | txn2 | differentially open at 2dpi |
| **chr3:11625609-11626108** | chr3 | 11625609 | 11626108 | 500 | * | 32071 | hlfa | -7.76E+05 | ccnf | differentially open at 2dpi |
| **chr3:32328635-32329134** | chr3 | 32328635 | 32329134 | 500 | * | -32035 | trpm4a | 49820 | rras | differentially open at 2dpi |
| **chr3:36970639-36971138** | chr3 | 36970639 | 36971138 | 500 | * | 32 | psmc3ip | 673091 | baxa | differentially open at 2dpi |
| **chr3:38598193-38598692** | chr3 | 38598193 | 38598692 | 500 | * | 175698 | si:dkey-106c17.2 | -8.28E+05 | aldoaa | differentially open at 2dpi |
| **chr3:55222028-55222527** | chr3 | 55222028 | 55222527 | 500 | * | -1291 | si:dkey-188h10.3 | -483963 | rhbdf1a | differentially open at 2dpi |
| **chr3:55995907-55996406** | chr3 | 55995907 | 55996406 | 500 | * | -40111 | srl | -511125 | rac1b | differentially open at 2dpi |
| **chr3:57474388-57474887** | chr3 | 57474388 | 57474887 | 500 | * | -14839 | cyth1a | -14839 | cyth1a | differentially open at 2dpi |
| **chr3:59285313-59285812** | chr3 | 59285313 | 59285812 | 500 | * | -106203 | nptx1l | -106203 | nptx1l | differentially open at 2dpi |
| **chr3:60369415-60369914** | chr3 | 60369415 | 60369914 | 500 | * | -4502 | anks4b | -4502 | anks4b | differentially open at 2dpi |
| **chr3:61682115-61682614** | chr3 | 61682115 | 61682614 | 500 | * | 168342 | srebf1 | 168342 | srebf1 | differentially open at 2dpi |
| **chr4:9945910-9946409** | chr4 | 9945910 | 9946409 | 500 | * | 35542 | sbf1 | 35542 | sbf1 | differentially open at 2dpi |
| **chr4:68244163-68245059** | chr4 | 68244163 | 68245059 | 897 | * | 407814 | znf1117 | -40932543 | fam19a5a | differentially open at 2dpi |
| **chr5:1090761-1091632** | chr5 | 1090761 | 1091632 | 872 | * | 11308 | mbd2 | -608799 | dhx33 | differentially open at 2dpi |
| **chr5:11919877-11920376** | chr5 | 11919877 | 11920376 | 500 | * | -95059 | zgc:171242 | 104502 | ksr2 | differentially open at 2dpi |
| **chr5:37564957-37565456** | chr5 | 37564957 | 37565456 | 500 | * | -10354 | gucy2d | 79835 | slc12a9 | differentially open at 2dpi |
| **chr5:37904932-37905431** | chr5 | 37904932 | 37905431 | 500 | * | 1646 | atp1b2b | -260141 | slc12a9 | differentially open at 2dpi |
| **chr5:50861109-50861608** | chr5 | 50861109 | 50861608 | 500 | * | 64942 | rasgrf2b | 323059 | homer1b | differentially open at 2dpi |
| **chr5:52934307-52934806** | chr5 | 52934307 | 52934806 | 500 | * | -354 | taf6 | 1157357 | ccnb1 | differentially open at 2dpi |
| **chr5:53882442-53882941** | chr5 | 53882442 | 53882941 | 500 | * | 25298 | ssna1 | 209221 | ccnb1 | differentially open at 2dpi |
| **chr5:64184151-64184650** | chr5 | 64184151 | 64184650 | 500 | * | -43 | si:ch211-236k19.4 | 8691 | zgc:101858 | differentially open at 2dpi |
| **chr5:66928807-66929306** | chr5 | 66928807 | 66929306 | 500 | * | 53220 | zbtb20 | -136024 | si:dkey-251i10.2 | differentially open at 2dpi |
| **chr5:71139273-71139772** | chr5 | 71139273 | 71139772 | 500 | * | 71080 | fam78ab | -2869087 | ank1a | differentially open at 2dpi |
| **chr5:71403458-71404338** | chr5 | 71403458 | 71404338 | 881 | * | -6085 | ddx54 | -3133463 | ank1a | differentially open at 2dpi |
| **chr6:327105-327604** | chr6 | 327105 | 327604 | 500 | * | 5604 | pde6ha | 92121 | grap2b | differentially open at 2dpi |
| **chr6:526191-526690** | chr6 | 526191 | 526690 | 500 | * | -34639 | zgc:92360 | -106965 | grap2b | differentially open at 2dpi |
| **chr6:11900072-11900571** | chr6 | 11900072 | 11900571 | 500 | * | 129106 | wdsub1 | 129106 | wdsub1 | differentially open at 2dpi |
| **chr6:25271538-25272037** | chr6 | 25271538 | 25272037 | 500 | * | -1 | kyat3 | 553745 | barhl2 | differentially open at 2dpi |
| **chr6:58980501-58981000** | chr6 | 58980501 | 58981000 | 500 | * | 26 | mars | -210349 | soat2 | differentially open at 2dpi |
| **chr6:59759972-59760471** | chr6 | 59759972 | 59760471 | 500 | * | -2546 | zgc:65895 | 392387 | slc32a1 | differentially open at 2dpi |
| **chr6:59865579-59866078** | chr6 | 59865579 | 59866078 | 500 | * | -34759 | ddx3b | 286781 | slc32a1 | differentially open at 2dpi |
| **chr7:5304709-5305208** | chr7 | 5304709 | 5305208 | 500 | * | -51551 | vangl2 | -413506 | ltb4r | differentially open at 2dpi |
| **chr7:6944553-6945429** | chr7 | 6944553 | 6945429 | 877 | * | 130125 | actn3b | 315381 | pfdn2 | differentially open at 2dpi |
| **chr7:7351882-7352381** | chr7 | 7351882 | 7352381 | 500 | * | 90394 | nit1 | -91760 | pfdn2 | differentially open at 2dpi |
| **chr7:10318592-10319091** | chr7 | 10318592 | 10319091 | 500 | * | -1169 | zfand6 | 1493865 | mex3b | differentially open at 2dpi |
| **chr7:24647925-24648424** | chr7 | 24647925 | 24648424 | 500 | * | 37590 | krcp | 37590 | krcp | differentially open at 2dpi |
| **chr7:40885503-40886002** | chr7 | 40885503 | 40886002 | 500 | * | 5600 | scrib | 173085 | ncf2 | differentially open at 2dpi |
| **chr7:69217869-69218368** | chr7 | 69217869 | 69218368 | 500 | * | 6310 | ctdnep1b | 65170 | slc1a1 | differentially open at 2dpi |
| **chr7:73160434-73160933** | chr7 | 73160434 | 73160933 | 500 | * | -64881 | cbln6 | -1522829 | cacnb2a | differentially open at 2dpi |
| **chr7:73948002-73948501** | chr7 | 73948002 | 73948501 | 500 | * | -50472 | cldnd1a | -2310397 | cacnb2a | differentially open at 2dpi |
| **chr7:73951811-73952310** | chr7 | 73951811 | 73952310 | 500 | * | -54280 | cldnd1a | -2314205 | cacnb2a | differentially open at 2dpi |
| **chr7:74078097-74078596** | chr7 | 74078097 | 74078596 | 500 | * | 8340 | ppp1cbl | -2440491 | cacnb2a | differentially open at 2dpi |
| **chr9:5470172-5470671** | chr9 | 5470172 | 5470671 | 500 | * | 19075 | tnfsf13b | -1623689 | si:dkey-4e7.3 | differentially open at 2dpi |
| **chr9:44213691-44214190** | chr9 | 44213691 | 44214190 | 500 | * | 217825 | ube2e3 | -477399 | znf385b | differentially open at 2dpi |
| **chr10:153604-154103** | chr10 | 153604 | 154103 | 500 | * | 14006 | erg | -33955 | ets2 | differentially open at 2dpi |
| **chr10:384344-384843** | chr10 | 384344 | 384843 | 500 | * | 39623 | zgc:171929 | -173891 | psmg1 | differentially open at 2dpi |
| **chr10:443696-444195** | chr10 | 443696 | 444195 | 500 | * | -72955 | npffr1l3 | -233243 | psmg1 | differentially open at 2dpi |
| **chr10:1799621-1800120** | chr10 | 1799621 | 1800120 | 500 | * | 2 | epb41l4a | -1589167 | psmg1 | differentially open at 2dpi |
| **chr10:5135268-5135767** | chr10 | 5135268 | 5135767 | 500 | * | -20 | zgc:113274 | -278421 | nfil3 | differentially open at 2dpi |
| **chr10:5211717-5212216** | chr10 | 5211717 | 5212216 | 500 | * | -22074 | sptlc1 | -201973 | nfil3 | differentially open at 2dpi |
| **chr10:6381773-6382272** | chr10 | 6381773 | 6382272 | 500 | * | -55 | zgc:114200 | -569140 | il6st | differentially open at 2dpi |
| **chr10:13701874-13702373** | chr10 | 13701874 | 13702373 | 500 | * | 112604 | cntfr | 659549 | lpar1 | differentially open at 2dpi |
| **chr11:165810-166309** | chr11 | 165810 | 166309 | 500 | * | -12993 | zgc:172352 | 103162 | slc11a2 | differentially open at 2dpi |
| **chr11:12802692-12803447** | chr11 | 12802692 | 12803447 | 756 | * | 230 | zgpat | 291573 | elovl1b | differentially open at 2dpi |
| **chr11:20623277-20623776** | chr11 | 20623277 | 20623776 | 500 | * | 230478 | slc6a14 | 353656 | cdh4 | differentially open at 2dpi |
| **chr11:45076061-45076560** | chr11 | 45076061 | 45076560 | 500 | * | 17205 | sos1 | 337578 | tk1 | differentially open at 2dpi |
| **chr12:3830775-3831274** | chr12 | 3830775 | 3831274 | 500 | * | -3675 | nif3l1 | 19675 | cabp5a | differentially open at 2dpi |
| **chr12:16832439-16832938** | chr12 | 16832439 | 16832938 | 500 | * | 904 | pank1b | 246466 | lipf | differentially open at 2dpi |
| **chr12:34189941-34190440** | chr12 | 34189941 | 34190440 | 500 | * | 17907 | socs3b | 144084 | birc5a | differentially open at 2dpi |
| **chr12:38575744-38576243** | chr12 | 38575744 | 38576243 | 500 | * | -57150 | abca5 | 193924 | rpl38 | differentially open at 2dpi |
| **chr12:48551037-48551536** | chr12 | 48551037 | 48551536 | 500 | * | 11658 | tex47 | 244390 | lrrc20 | differentially open at 2dpi |
| **chr13:1001301-1001800** | chr13 | 1001301 | 1001800 | 500 | * | 25 | wdr92 | -1782 | pno1 | differentially open at 2dpi |
| **chr13:1484577-1485076** | chr13 | 1484577 | 1485076 | 500 | * | -206298 | bag2 | -206298 | bag2 | differentially open at 2dpi |
| **chr13:6989718-6990217** | chr13 | 6989718 | 6990217 | 500 | * | 30213 | LOC795051 | -933230 | hells | differentially open at 2dpi |
| **chr13:51544130-51544629** | chr13 | 51544130 | 51544629 | 500 | * | 25604 | klc1b | 73078 | lbh | differentially open at 2dpi |
| **chr13:51623489-51623988** | chr13 | 51623489 | 51623988 | 500 | * | -6280 | lbh | -6280 | lbh | differentially open at 2dpi |
| **chr14:181543-182042** | chr14 | 181543 | 182042 | 500 | * | 21534 | gpc2 | 31590 | mcm7 | differentially open at 2dpi |
| **chr14:24633380-24633879** | chr14 | 24633380 | 24633879 | 500 | * | 103294 | pde6a | 646233 | cplx2 | differentially open at 2dpi |
| **chr14:51841744-51842243** | chr14 | 51841744 | 51842243 | 500 | * | -6172 | rpl26 | -13465 | atp6v0e1 | differentially open at 2dpi |

| | | | | | | | | | | |
|---|---|---|---|---|---|---|---|---|---|---|
| **chr15:22879-23378** | chr15 | 22879 | 23378 | 500 | * | -266 | bace1 | 320244 | limk1a | differentially open at 2dpi |
| **chr15:116690-117189** | chr15 | 116690 | 117189 | 500 | * | -12598 | cyp2y3 | 226432 | limk1a | differentially open at 2dpi |
| **chr15:201627-202126** | chr15 | 201627 | 202126 | 500 | * | -14330 | LOC100002960 | 141496 | limk1a | differentially open at 2dpi |
| **chr15:1569009-1569954** | chr15 | 1569009 | 1569954 | 946 | * | -159 | ift80 | -234 | smc4 | differentially open at 2dpi |
| **chr15:20593110-20593609** | chr15 | 20593110 | 20593609 | 500 | * | -220 | tnfaip1 | 387086 | usp2a | differentially open at 2dpi |
| **chr15:44093163-44093662** | chr15 | 44093163 | 44093662 | 500 | * | 23567 | zgc:112998 | 63611 | lrrc51 | differentially open at 2dpi |
| **chr15:45801381-45801880** | chr15 | 45801381 | 45801880 | 500 | * | 22858 | lrfn1 | 176107 | sagb | differentially open at 2dpi |
| **chr15:47041964-47042463** | chr15 | 47041964 | 47042463 | 500 | * | -16574 | sik3 | 1416691 | sagb | differentially open at 2dpi |
| **chr15:47515380-47515879** | chr15 | 47515380 | 47515879 | 500 | * | -118293 | cep57 | 1890107 | sagb | differentially open at 2dpi |
| **chr16:1384691-1385190** | chr16 | 1384691 | 1385190 | 500 | * | 94195 | sim1a | 647278 | irx1a | differentially open at 2dpi |
| **chr16:2582464-2582963** | chr16 | 2582464 | 2582963 | 500 | * | -2223 | acot22 | -261727 | adcy3a | differentially open at 2dpi |
| **chr16:20249721-20250220** | chr16 | 20249721 | 20250220 | 500 | * | -17526 | ahr1a | -165515 | hdac9b | differentially open at 2dpi |
| **chr16:48595152-48595651** | chr16 | 48595152 | 48595651 | 500 | * | 76547 | ext1a | -136770 | eif3ha | differentially open at 2dpi |
| **chr16:54619065-54619564** | chr16 | 54619065 | 54619564 | 500 | * | 15369 | pklr | 1540191 | trip13 | differentially open at 2dpi |
| **chr17:487548-488047** | chr17 | 487548 | 488047 | 500 | * | -29921 | chrm5a | -372235 | gchfr | differentially open at 2dpi |
| **chr17:1395288-1395787** | chr17 | 1395288 | 1395787 | 500 | * | -6587 | dio3a | 113774 | wars | differentially open at 2dpi |
| **chr17:8489676-8490175** | chr17 | 8489676 | 8490175 | 500 | * | 45616 | ctbp2a | 45616 | ctbp2a | differentially open at 2dpi |
| **chr17:28689744-28690243** | chr17 | 28689744 | 28690243 | 500 | * | -211 | strn3 | 104643 | g2e3 | differentially open at 2dpi |
| **chr17:43711273-43711772** | chr17 | 43711273 | 43711772 | 500 | * | 38413 | ahsa1b | -670253 | npc2 | differentially open at 2dpi |
| **chr17:51613348-51613847** | chr17 | 51613348 | 51613847 | 500 | * | 20204 | nol10 | 478212 | adi1 | differentially open at 2dpi |
| **chr18:272145-272644** | chr18 | 272145 | 272644 | 500 | * | -12073 | larp6a | -746281 | pkma | differentially open at 2dpi |
| **chr18:5548260-5548759** | chr18 | 5548260 | 5548759 | 500 | * | 16453 | slc24a5 | -89086 | mibp2 | differentially open at 2dpi |
| **chr18:6627792-6628291** | chr18 | 6627792 | 6628291 | 500 | * | 76073 | fam168a | 469717 | utp15 | differentially open at 2dpi |
| **chr18:7072291-7072790** | chr18 | 7072291 | 7072790 | 500 | * | -110 | calub | 25219 | utp15 | differentially open at 2dpi |
| **chr18:8700770-8701269** | chr18 | 8700770 | 8701269 | 500 | * | -79132 | si:ch211-220f12.1 | -712952 | sema3ab | differentially open at 2dpi |
| **chr18:41520738-41521237** | chr18 | 41520738 | 41521237 | 500 | * | -964 | ccnl1a | -964 | ccnl1a | differentially open at 2dpi |
| **chr18:50489493-50489992** | chr18 | 50489493 | 50489992 | 500 | * | 37722 | cd276 | 37722 | cd276 | differentially open at 2dpi |
| **chr19:19931582-19932081** | chr19 | 19931582 | 19932081 | 500 | * | -7732 | jazf1a | 590331 | mpp6b | differentially open at 2dpi |
| **chr19:20512302-20512801** | chr19 | 20512302 | 20512801 | 500 | * | 9611 | mpp6b | 9611 | mpp6b | differentially open at 2dpi |
| **chr19:25880479-25880978** | chr19 | 25880479 | 25880978 | 500 | * | -112 | rpa3 | -193838 | rims3 | differentially open at 2dpi |
| **chr20:6486555-6487054** | chr20 | 6486555 | 6487054 | 500 | * | -16897 | mep1b | -1202574 | ism2b | differentially open at 2dpi |
| **chr20:15652847-15653346** | chr20 | 15652847 | 15653346 | 500 | * | -148 | jun | -148 | jun | differentially open at 2dpi |
| **chr20:15656516-15657393** | chr20 | 15656516 | 15657393 | 878 | * | -3684 | si:dkey-86e18.2 | 3710 | jun | differentially open at 2dpi |
| **chr20:15687329-15687828** | chr20 | 15687329 | 15687828 | 500 | * | -13254 | faslg | 34334 | jun | differentially open at 2dpi |
| **chr20:17982452-17982951** | chr20 | 17982452 | 17982951 | 500 | * | 142181 | cdh2 | -281083 | aqp4 | differentially open at 2dpi |
| **chr20:20868444-20868943** | chr20 | 20868444 | 20868943 | 500 | * | 37340 | ppp1r13bb | 53391 | ckbb | differentially open at 2dpi |
| **chr20:38079431-38079930** | chr20 | 38079431 | 38079930 | 500 | * | -49304 | angel2 | -50019 | galnt14 | differentially open at 2dpi |
| **chr20:39316562-39317061** | chr20 | 39316562 | 39317061 | 500 | * | -149 | cyp39a1 | -31215 | rcan2 | differentially open at 2dpi |
| **chr20:54830661-54831160** | chr20 | 54830661 | 54831160 | 500 | * | -18186 | ppp1r35 | 540969 | rps29 | differentially open at 2dpi |
| **chr21:697792-698291** | chr21 | 697792 | 698291 | 500 | * | -838 | oaz1b | 395532 | lhfpl2a | differentially open at 2dpi |
| **chr21:4375223-4375722** | chr21 | 4375223 | 4375722 | 500 | * | -183 | dolk | 283687 | lrrc8aa | differentially open at 2dpi |
| **chr21:11235062-11235561** | chr21 | 11235062 | 11235561 | 500 | * | -110 | rtkn2b | -128574 | dnajc21 | differentially open at 2dpi |
| **chr21:39416988-39417487** | chr21 | 39416988 | 39417487 | 500 | * | -224 | mntb | 104537 | dynll2b | differentially open at 2dpi |
| **chr22:20971838-20972337** | chr22 | 20971838 | 20972337 | 500 | * | -46618 | fkbp8 | 49628 | ssbp4 | differentially open at 2dpi |
| **chr22:37094885-37095384** | chr22 | 37094885 | 37095384 | 500 | * | -109487 | si:dkey-37m8.11 | -601092 | psmd1 | differentially open at 2dpi |
| **chr23:1280885-1281384** | chr23 | 1280885 | 1281384 | 500 | * | -145829 | psmb11b | 715726 | ogfr | differentially open at 2dpi |
| **chr23:2174238-2174737** | chr23 | 2174238 | 2174737 | 500 | * | -147271 | prdm5 | -307710 | tcp1 | differentially open at 2dpi |
| **chr23:14677630-14678129** | chr23 | 14677630 | 14678129 | 500 | * | -30018 | nkain4 | 216449 | birc7 | differentially open at 2dpi |
| **chr23:16765279-16766037** | chr23 | 16765279 | 16766037 | 759 | * | 296 | fkbp1ab | -9815 | sdcbp2 | differentially open at 2dpi |
| **chr23:39321018-39321517** | chr23 | 39321018 | 39321517 | 500 | * | 26749 | bmf1 | -132751 | src | differentially open at 2dpi |
| **chr23:45941247-45941746** | chr23 | 45941247 | 45941746 | 500 | * | 28838 | ednrab | 920679 | atp1b2a | differentially open at 2dpi |
| **chr25:136191-136690** | chr25 | 136191 | 136690 | 500 | * | -45643 | zgc:114188 | 2772371 | scamp2 | differentially open at 2dpi |
| **chr25:222688-223187** | chr25 | 222688 | 223187 | 500 | * | 626 | zgc:92481 | 2685873 | scamp2 | differentially open at 2dpi |
| **chr25:10696492-10696991** | chr25 | 10696492 | 10696991 | 500 | * | 37 | ap3s2 | 246291 | zgc:110339 | differentially open at 2dpi |
| **chrUn_KN149790v1:100748-101247** | chrUn_KN149790v1 | 100748 | 101247 | 500 | * | -6772 | zgc:171422 | NA | NA | differentially open at 2dpi |
| **chrUn_KN150361v1:39209-39708** | chrUn_KN150361v1 | 39209 | 39708 | 500 | * | NA | NA | NA | NA | differentially open at 2dpi |
| **chrUn_KN150600v1:341-840** | chrUn_KN150600v1 | 341 | 840 | 500 | * | NA | NA | NA | NA | differentially open at 2dpi |
| **chr2:15088215-15088714** | chr2 | 15088215 | 15088714 | 500 | * | -100384 | leprot | 1076793 | ebna1bp2 | differentially closed at 2dpi |
| **chr5:61868726-61869225** | chr5 | 61868726 | 61869225 | 500 | * | -57698 | abr | -1.11E+06 | ncf1 | differentially closed at 2dpi |
| **chr7:5468715-5469214** | chr7 | 5468715 | 5469214 | 500 | * | -103604 | arhgef11 | -577512 | ltb4r | differentially closed at 2dpi |
| **chr20:15442099-15442598** | chr20 | 15442099 | 15442598 | 500 | * | 177367 | prdx6 | -210896 | jun | differentially closed at 2dpi |
| **chr20:18383573-18384072** | chr20 | 18383573 | 18384072 | 500 | * | 30430 | psma8 | 120037 | aqp4 | differentially closed at 2dpi |
| **chr21:24198964-24199463** | chr21 | 24198964 | 24199463 | 500 | * | 282640 | cadm1a | 282640 | cadm1a | differentially closed at 2dpi |
| **chr22:17120757-17121529** | chr22 | 17120757 | 17121529 | 773 | * | 26763 | frem1b | -460764 | polr2eb | differentially closed at 2dpi |
| **chr22:30060403-30060902** | chr22 | 30060403 | 30060902 | 500 | * | -79 | si:dkey-286j15.1 | -86975 | dusp5 | differentially closed at 2dpi |
| **chr24:16096606-16097105** | chr24 | 16096606 | 16097105 | 500 | * | -305731 | sema5a | -593269 | cbln2b | differentially closed at 2dpi |
| **chr6:59759972-59760471** | chr6 | 59759972 | 59760471 | 500 | * | -2546 | zgc:65895 | 392387 | slc32a1 | differentially open at 12dpi |
| **chr6:59899089-59900027** | chr6 | 59899089 | 59900027 | 939 | * | -1029 | ddx3b | 253051 | slc32a1 | differentially open at 12dpi |
| **chr7:6944553-6945429** | chr7 | 6944553 | 6945429 | 877 | * | 130125 | actn3b | 315381 | pfdn2 | differentially open at 12dpi |
| **chr9:17415700-17416199** | chr9 | 17415700 | 17416199 | 500 | * | -5523 | zgc:101559 | -15126 | rgcc | differentially open at 12dpi |
| **chr15:253918-254417** | chr15 | 253918 | 254417 | 500 | * | 115 | sept4b | 89204 | limk1a | differentially open at 12dpi |
| **chr17:43296504-43297003** | chr17 | 43296504 | 43297003 | 500 | * | 141105 | plk4 | -255485 | npc2 | differentially closed at 12dpi |
| **chr2:81394-81893** | chr2 | 81394 | 81893 | 500 | * | 31014 | zgc:153913 | -1514947 | exosc4 | differentially open at 12dpi |
| **chr2:19780119-19781338** | chr2 | 19780119 | 19781338 | 1220 | * | -66386 | pimr141 | 265831 | elovl1a | differentially open at 12dpi |
| **chr2:19781474-19781973** | chr2 | 19781474 | 19781973 | 500 | * | -65390 | pimr141 | 266827 | elovl1a | differentially open at 12dpi |
| **chr2:19787237-19788192** | chr2 | 19787237 | 19788192 | 956 | * | -59400 | pimr141 | 272817 | elovl1a | differentially open at 12dpi |
| **chr2:19813598-19814097** | chr2 | 19813598 | 19814097 | 500 | * | -33266 | pimr141 | 298951 | elovl1a | differentially open at 12dpi |
| **chr2:19817067-19817998** | chr2 | 19817067 | 19817998 | 932 | * | -29582 | pimr141 | 302635 | elovl1a | differentially open at 12dpi |
| **chr2:19818257-19818756** | chr2 | 19818257 | 19818756 | 500 | * | -28608 | pimr141 | 303609 | elovl1a | differentially open at 12dpi |
| **chr2:19822778-19823647** | chr2 | 19822778 | 19823647 | 870 | * | -23902 | pimr141 | 308315 | elovl1a | differentially open at 12dpi |
| **chr2:19826215-19827613** | chr2 | 19826215 | 19827613 | 1399 | * | -20200 | pimr141 | 312017 | elovl1a | differentially open at 12dpi |
| **chr3:55093026-55093806** | chr3 | 55093026 | 55093806 | 781 | * | 52958 | mafga | -355101 | rhbdf1a | differentially open at 12dpi |
| **chr3:55095105-55095604** | chr3 | 55095105 | 55095604 | 500 | * | 51020 | mafga | -357039 | rhbdf1a | differentially open at 12dpi |
| **chr3:55100242-55101294** | chr3 | 55100242 | 55101294 | 1053 | * | 45606 | mafga | -362453 | rhbdf1a | differentially open at 12dpi |
| **chr3:55105417-55106689** | chr3 | 55105417 | 55106689 | 1273 | * | 40321 | mafga | -367738 | rhbdf1a | differentially open at 12dpi |
| **chr4:28604846-28605345** | chr4 | 28604846 | 28605345 | 500 | * | -126528 | mir430a-18 | -1293028 | fam19a5a | differentially open at 12dpi |
| **chr5:47142933-47143815** | chr5 | 47142933 | 47143815 | 883 | * | -36609 | cox7c | -1237825 | hapln1a | differentially open at 12dpi |

| | | | | | | | | | | |
|---|---|---|---|---|---|---|---|---|---|---|
| **chr7:52730398-52730897** | chr7 | 52730398 | 52730897 | 500 | * | -38212 | LOC557301 | -447240 | tcf12 | differentially open at 12dpi |
| **chr7:52740095-52740594** | chr7 | 52740095 | 52740594 | 500 | * | -47908 | LOC557301 | -456936 | tcf12 | differentially open at 12dpi |
| **chr7:52742377-52743183** | chr7 | 52742377 | 52743183 | 807 | * | -50344 | LOC557301 | -459372 | tcf12 | differentially open at 12dpi |
| **chr7:52748837-52749336** | chr7 | 52748837 | 52749336 | 500 | * | -56650 | LOC557301 | -465678 | tcf12 | differentially open at 12dpi |
| **chr7:54788736-54789235** | chr7 | 54788736 | 54789235 | 500 | * | -22586 | snai3 | 543078 | galns | differentially open at 12dpi |
| **chr7:54790698-54791197** | chr7 | 54790698 | 54791197 | 500 | * | -20624 | snai3 | 541116 | galns | differentially open at 12dpi |
| **chr8:34228765-34229586** | chr8 | 34228765 | 34229586 | 822 | * | -152153 | pbx3b | -152153 | pbx3b | differentially open at 12dpi |
| **chr8:34229943-34230784** | chr8 | 34229943 | 34230784 | 842 | * | -153341 | pbx3b | -153341 | pbx3b | differentially open at 12dpi |
| **chr8:34235505-34236004** | chr8 | 34235505 | 34236004 | 500 | * | -158731 | pbx3b | -158731 | pbx3b | differentially open at 12dpi |
| **chr8:34239947-34240446** | chr8 | 34239947 | 34240446 | 500 | * | -163173 | pbx3b | -163173 | pbx3b | differentially open at 12dpi |
| **chr8:34243228-34243727** | chr8 | 34243228 | 34243727 | 500 | * | -166455 | pbx3b | -166455 | pbx3b | differentially open at 12dpi |
| **chr8:34244143-34244642** | chr8 | 34244143 | 34244642 | 500 | * | -167369 | pbx3b | -167369 | pbx3b | differentially open at 12dpi |
| **chr8:34846434-34847288** | chr8 | 34846434 | 34847288 | 855 | * | 49679 | hpdb | -769838 | pbx3b | differentially open at 12dpi |
| **chr8:34851982-34852936** | chr8 | 34851982 | 34852936 | 955 | * | 55277 | hpdb | -775436 | pbx3b | differentially open at 12dpi |
| **chr8:34857277-34857776** | chr8 | 34857277 | 34857776 | 500 | * | 60344 | hpdb | -780503 | pbx3b | differentially open at 12dpi |
| **chr8:34874225-34874724** | chr8 | 34874225 | 34874724 | 500 | * | 77292 | hpdb | -797451 | pbx3b | differentially open at 12dpi |
| **chr10:16847552-16848051** | chr10 | 16847552 | 16848051 | 500 | * | 62555 | stoml2 | 304222 | slc27a6 | differentially open at 12dpi |
| **chr10:16848134-16848633** | chr10 | 16848134 | 16848633 | 500 | * | 61973 | stoml2 | 304804 | slc27a6 | differentially open at 12dpi |
| **chr10:16848644-16849143** | chr10 | 16848644 | 16849143 | 500 | * | 61463 | stoml2 | 305314 | slc27a6 | differentially open at 12dpi |
| **chr10:16851015-16852861** | chr10 | 16851015 | 16852861 | 1847 | * | 58419 | stoml2 | 308358 | slc27a6 | differentially open at 12dpi |
| **chr10:16858621-16859391** | chr10 | 16858621 | 16859391 | 771 | * | 51351 | stoml2 | 315426 | slc27a6 | differentially open at 12dpi |
| **chr10:16880577-16881974** | chr10 | 16880577 | 16881974 | 1398 | * | 29081 | stoml2 | 337696 | slc27a6 | differentially open at 12dpi |
| **chr11:17430700-17431898** | chr11 | 17430700 | 17431898 | 1199 | * | 148053 | fam19a4b | -1279472 | magi1b | differentially open at 12dpi |
| **chr11:17432488-17432987** | chr11 | 17432488 | 17432987 | 500 | * | 146614 | fam19a4b | -1278033 | magi1b | differentially open at 12dpi |
| **chr11:17432997-17433496** | chr11 | 17432997 | 17433496 | 500 | * | 146106 | fam19a4b | -1277525 | magi1b | differentially open at 12dpi |
| **chr11:17433633-17434603** | chr11 | 17433633 | 17434603 | 971 | * | 145234 | fam19a4b | -1276653 | magi1b | differentially open at 12dpi |
| **chr13:18988177-18988676** | chr13 | 18988177 | 18988676 | 500 | * | -164595 | eif3s10 | 807947 | eif4e1c | differentially open at 12dpi |
| **chr14:19788805-19789543** | chr14 | 19788805 | 19789543 | 739 | * | -70165 | fmr1 | 1928907 | ssrp1a | differentially open at 12dpi |
| **chr14:19792068-19792567** | chr14 | 19792068 | 19792567 | 500 | * | -67021 | fmr1 | 1925763 | ssrp1a | differentially open at 12dpi |
| **chr14:19792901-19794038** | chr14 | 19792901 | 19794038 | 1138 | * | -65869 | fmr1 | 1924611 | ssrp1a | differentially open at 12dpi |
| **chr14:19796029-19797437** | chr14 | 19796029 | 19797437 | 1409 | * | -62606 | fmr1 | 1921348 | ssrp1a | differentially open at 12dpi |
| **chr14:19810584-19811525** | chr14 | 19810584 | 19811525 | 942 | * | -48285 | fmr1 | 1907027 | ssrp1a | differentially open at 12dpi |
| **chr15:14830321-14830820** | chr15 | 14830321 | 14830820 | 500 | * | 13985 | or129-1 | -556208 | zgc:162193 | differentially open at 12dpi |
| **chr16:26006060-26007021** | chr16 | 26006060 | 26007021 | 962 | * | -91798 | irge4 | 352429 | arhgef1b | differentially open at 12dpi |
| **chr16:26013553-26014547** | chr16 | 26013553 | 26014547 | 995 | * | -99308 | irge4 | 344919 | arhgef1b | differentially open at 12dpi |
| **chr16:26053213-26053712** | chr16 | 26053213 | 26053712 | 500 | * | -85665 | prss59.1 | 305507 | arhgef1b | differentially open at 12dpi |
| **chr16:26056249-26056748** | chr16 | 26056249 | 26056748 | 500 | * | -82629 | prss59.1 | 302471 | arhgef1b | differentially open at 12dpi |
| **chr16:26058671-26059170** | chr16 | 26058671 | 26059170 | 500 | * | -80207 | prss59.1 | 300049 | arhgef1b | differentially open at 12dpi |
| **chr16:26065441-26065940** | chr16 | 26065441 | 26065940 | 500 | * | -73437 | prss59.1 | 293279 | arhgef1b | differentially open at 12dpi |
| **chr16:26067798-26068703** | chr16 | 26067798 | 26068703 | 906 | * | -70877 | prss59.1 | 290719 | arhgef1b | differentially open at 12dpi |
| **chr17:45219176-45219675** | chr17 | 45219176 | 45219675 | 500 | * | 27546 | capn3a | 184145 | tmem206 | differentially open at 12dpi |
| **chr17:45222066-45222565** | chr17 | 45222066 | 45222565 | 500 | * | 30436 | capn3a | 181255 | tmem206 | differentially open at 12dpi |
| **chr17:46968319-46968818** | chr17 | 46968319 | 46968818 | 500 | * | -38468 | zgc:103755 | 1242885 | smoc1 | differentially open at 12dpi |
| **chr17:46977515-46978014** | chr17 | 46977515 | 46978014 | 500 | * | -29272 | zgc:103755 | 1233689 | smoc1 | differentially open at 12dpi |
| **chr17:46978496-46978995** | chr17 | 46978496 | 46978995 | 500 | * | -28290 | zgc:103755 | 1232707 | smoc1 | differentially open at 12dpi |
| **chr17:46981453-46982699** | chr17 | 46981453 | 46982699 | 1247 | * | -24960 | zgc:103755 | 1229377 | smoc1 | differentially open at 12dpi |
| **chr20:4437575-4438074** | chr20 | 4437575 | 4438074 | 500 | * | -68077 | ipcef1 | 846406 | ism2b | differentially open at 12dpi |
| **chr21:25809834-25810333** | chr21 | 25809834 | 25810333 | 500 | * | -44733 | mettl27 | -225121 | rpl23a | differentially open at 12dpi |
| **chr21:25823531-25824030** | chr21 | 25823531 | 25824030 | 500 | * | -58429 | mettl27 | -211425 | rpl23a | differentially open at 12dpi |
| **chr21:44162938-44163437** | chr21 | 44162938 | 44163437 | 500 | * | 45439 | fgf1b | -205396 | camk2a | differentially open at 12dpi |
| **chr22:24497163-24497662** | chr22 | 24497163 | 24497662 | 500 | * | -35194 | wdr47b | -275554 | rgs2 | differentially open at 12dpi |
| **chr22:24500122-24501449** | chr22 | 24500122 | 24501449 | 1328 | * | -31820 | wdr47b | -278928 | rgs2 | differentially open at 12dpi |
| **chr22:24503566-24505214** | chr22 | 24503566 | 24505214 | 1649 | * | -28216 | wdr47b | -282532 | rgs2 | differentially open at 12dpi |
| **chr23:12684435-12685935** | chr23 | 12684435 | 12685935 | 1501 | * | -166455 | snx21 | -274391 | phactr3a | differentially open at 12dpi |
| **chr23:16244775-16246161** | chr23 | 16244775 | 16246161 | 1387 | * | -85259 | mrgbp | -122666 | eef1a2 | differentially open at 12dpi |
| **chr24:16118773-16120428** | chr24 | 16118773 | 16120428 | 1656 | * | -282987 | sema5a | -616013 | cbln2b | differentially open at 12dpi |
| **chr24:16126674-16127173** | chr24 | 16126674 | 16127173 | 500 | * | -275663 | sema5a | -623337 | cbln2b | differentially open at 12dpi |
| **chrUn_KN149692v1:4089-5235** | chrUn_KN149692v1 | 4089 | 5235 | 1147 | * | NA | NA | NA | NA | differentially open at 12dpi |
| **chrUn_KN149692v1:5249-6134** | chrUn_KN149692v1 | 5249 | 6134 | 886 | * | NA | NA | NA | NA | differentially open at 12dpi |
| **chrUn_KN149692v1:10411-11564** | chrUn_KN149692v1 | 10411 | 11564 | 1154 | * | NA | NA | NA | NA | differentially open at 12dpi |
| **chrUn_KN149692v1:13822-14321** | chrUn_KN149692v1 | 13822 | 14321 | 500 | * | NA | NA | NA | NA | differentially open at 12dpi |
| **chrUn_KN149692v1:16713-17212** | chrUn_KN149692v1 | 16713 | 17212 | 500 | * | NA | NA | NA | NA | differentially open at 12dpi |
| **chrUn_KN149692v1:20397-21478** | chrUn_KN149692v1 | 20397 | 21478 | 1082 | * | NA | NA | NA | NA | differentially open at 12dpi |

**Table S5. Differentially expressed transcripts that encode transcription factors with and without known motifs.**

| Ensembl ID | zebrafish gene symbol | padj | LFC: 2dpi-0dpi | LFC: 4dpi-0dpi | LFC: 7dpi-0dpi | LFC: 12dpi-0dpi | cluster name |
|---|---|---|---|---|---|---|---|
| ENSDART00000140760 | e2f7 | 0.044060134 | 0.798632306 | 0.496792942 | -0.026056819 | -0.454809271 | growth toward the midline |
| ENSDART00000056005 | ascl1a | 0.032093457 | 1.787445127 | 1.283325532 | 0.749862779 | 0.792492648 | growth toward the midline |
| ENSDART00000128488 | e2f8 | 0.003382652 | 3.852661131 | -0.929008264 | 3.252279997 | 2.01178299 | growth toward the midline |
| ENSDART00000010248 | mitfb | 0.042577687 | 0.302243319 | 0.237275907 | 0.106756262 | -0.337810921 | growth toward the midline |
| ENSDART00000157487 | tfec | 0.001790465 | 0.534318298 | 0.423012313 | 0.205933849 | -0.313187686 | growth toward the midline |
| ENSDART00000114288 | foxp4 | 0.028339664 | 0.17723361 | 0.325948274 | 0.017102596 | -0.613519733 | growth toward the midline |
| ENSDART00000158466 | creb3l2 | 0.011352496 | 0.073267533 | 0.091531466 | -0.07087733 | -0.32121815 | growth toward the midline |
| ENSDART00000164766 | tfec | 0.012357429 | 0.976052476 | 1.18639141 | 0.809154671 | -1.643283392 | growth toward the midline |
| ENSDART00000142584 | alx1 | 0.047364517 | 1.361674144 | 1.279023188 | 1.723053191 | -0.115335088 | growth toward the midline |
| ENSDART00000081468 | ccdc79 | 0.001875423 | 0.331555315 | 0.505423262 | 0.318137438 | -0.103372006 | growth toward the midline |
| ENSDART00000155066 | atf5b | 0.027985198 | 0.303223783 | 0.449779577 | 0.245269838 | -0.118029433 | growth toward the midline |
| ENSDART00000160644 | rela | 0.030637152 | 0.027046368 | 0.222531925 | 0.299056597 | -0.13648552 | growth toward the midline |
| ENSDART00000146767 | fosl1a | 0.046117855 | 0.841007094 | 1.285901651 | 0.733422298 | 0.173089455 | growth toward the midline |
| ENSDART00000152378 | tgif1 | 0.003122803 | 0.194631245 | 0.419803811 | 0.14379228 | -0.205607529 | growth toward the midline |
| ENSDART00000051549 | tp53 | 0.029170799 | 0.110640197 | 0.281712522 | 0.180041895 | -0.162598759 | growth toward the midline |
| ENSDART00000013409 | prmt3 | 0.000336326 | 0.583250552 | 0.486757337 | 0.420985526 | -0.028812313 | growth toward the midline |
| ENSDART00000104519 | stat3 | 0.004480141 | 0.6458865 | 0.459098575 | 0.438501049 | 0.201812376 | growth toward the midline |
| ENSDART00000058277 | znf800b | 0.018079865 | 0.363600272 | 0.378301813 | 0.156623376 | -0.047842936 | growth toward the midline |
| ENSDART00000103982 | nfe2l1a | 0.007828891 | 0.652859432 | 0.675562602 | 0.38648833 | -0.18398816 | growth toward the midline |
| ENSDART00000093279 | spi1b | 0.002640644 | 0.891535687 | 0.563855895 | 0.100587929 | -0.531177734 | growth toward the midline |
| ENSDART00000036729 | spi1b | 0.004358158 | 0.910663058 | 0.717004711 | 0.095052493 | -0.117318513 | growth toward the midline |
| ENSDART00000008373 | fosl1a | 0.001576361 | 0.78271954 | 0.570182098 | 0.226810387 | -0.332448721 | growth toward the midline |
| ENSDART00000093199 | tead3b | 0.016523707 | 1.081516431 | 0.86043279 | 0.478204597 | -0.215799128 | growth toward the midline |
| ENSDART00000062702 | cebpb | 0.018433591 | 0.973724375 | 0.549827472 | 0.384246832 | -0.627629236 | growth toward the midline |
| ENSDART00000056457 | mitfa | 0.005353377 | 0.929754118 | 0.38849893 | 0.30836176 | -0.311729197 | growth toward the midline |
| ENSDART00000080854 | stat3 | 0.03033326 | 0.856010749 | 0.444560766 | 0.379796349 | -0.326021696 | growth toward the midline |
| ENSDART00000017185 | tbx20 | 0.035967167 | 0.559146809 | 0.370265655 | -0.151461394 | -0.656442573 | growth toward the midline |
| ENSDART00000086053 | PRDM16 | 0.037061149 | 0.751033854 | 0.474975066 | 0.399761078 | -0.264438466 | growth toward the midline |
| ENSDART00000052366 | cebpa | 0.035286858 | 0.847474055 | 0.814939712 | 0.318232429 | -0.173654256 | growth toward the midline |
| ENSDART00000062854 | nfe2l2a | 0.008134819 | 0.155734951 | 0.146836561 | -0.112595282 | -0.207726174 | growth toward the midline |
| ENSDART00000016135 | nfe2l3 | 0.02142158 | 0.030439041 | 0.342226134 | 0.430712572 | 0.086440986 | midline crossing |
| ENSDART00000141068 | sox11b | 8.49E-05 | 1.611838536 | 2.177304965 | 2.010624009 | 1.194765222 | midline crossing |
| ENSDART00000149074 | sall1b | 2.18E-05 | 1.974229239 | 2.883794957 | 2.506155499 | 1.735121972 | midline crossing |
| ENSDART00000138821 | nfil3 | 0.000101984 | 1.376309909 | 2.121391468 | 1.99718084 | 1.39511595 | midline crossing |
| ENSDART00000153227 | nfe2l1b | 0.013323448 | 0.688085235 | 1.301360444 | 1.138008541 | 0.729999096 | midline crossing |
| ENSDART00000082434 | tgif1 | 1.75E-05 | 0.822910382 | 1.229522702 | 1.441745222 | 0.912580325 | midline crossing |
| ENSDART00000104317 | klf7b | 0.000976528 | 0.389436838 | 1.156291561 | 1.140404424 | 0.652703479 | midline crossing |
| ENSDART00000099849 | arntl2 | 9.24E-05 | 0.982355145 | 1.78511579 | 1.609893892 | 1.255639962 | midline crossing |
| ENSDART00000123263 | phf20a | 0.000701449 | 3.204242437 | 4.684552705 | 5.028378766 | 3.758919167 | midline crossing |
| ENSDART00000124040 | insm1a | 1.46E-05 | 0.50962443 | 1.551047216 | 1.738068624 | 1.104992333 | midline crossing |
| ENSDART00000159652 | sox4b | 0.008897727 | 0.767109202 | 1.080655119 | 1.389017231 | 0.976257694 | midline crossing |
| ENSDART00000062576 | thyn1 | 0.006623227 | 0.902996379 | 0.741810201 | 0.792519989 | 0.219218083 | midline crossing |
| ENSDART00000098285 | atf5a | 0.018496153 | 0.558593719 | 0.754458044 | 0.445954377 | 0.160619888 | midline crossing |
| ENSDART00000063912 | jun | 0.002341904 | 1.166146477 | 1.429328618 | 1.29995447 | 0.480560375 | midline crossing |
| ENSDART00000022060 | atf3 | 2.20E-05 | 2.958540036 | 3.348918067 | 2.852908333 | 1.811501419 | midline crossing |
| ENSDART00000110040 | sox11a | 8.49E-05 | 1.573346229 | 1.723237215 | 1.23195588 | 0.781584101 | midline crossing |
| ENSDART00000141941 | zfpm2a | 0.046610115 | 3.791965294 | 3.402723828 | 3.788725401 | 3.249505128 | midline crossing |
| ENSDART00000161610 | tcf3b | 0.04032679 | 0.066966853 | 0.197082202 | 0.107652202 | -0.03222147 | midline crossing |
| ENSDART00000012791 | sp8a | 0.049626195 | 1.57251014 | 2.331511764 | 2.00847442 | 1.039704847 | midline crossing |
| ENSDART00000122628 | junba | 0.016136147 | 0.280813702 | 0.583374691 | 0.41181471 | 0.073998168 | midline crossing |
| ENSDART00000024872 | creb3l3l | 0.003959852 | 0.316205692 | 0.628018553 | 0.525420097 | 0.254183225 | midline crossing |
| ENSDART00000123559 | znf521 | 0.000648976 | 1.710422257 | 2.255410725 | 1.850379402 | 0.814044199 | midline crossing |
| ENSDART00000147849 | klf6a | 0.000870446 | 1.130982604 | 1.870693821 | 1.403514717 | 0.445060502 | midline crossing |
| ENSDART00000076161 | hoxb5b | 0.013675681 | 2.652791903 | 3.351280664 | 2.618897772 | 0.864721852 | midline crossing |
| ENSDART00000033494 | klf6a | 0.000170861 | 1.084550888 | 1.797692613 | 1.625562666 | 0.460444617 | midline crossing |
| ENSDART00000077484 | zhx2a | 0.005803283 | 0.117201305 | 0.356185793 | 0.299966576 | 0.024327323 | midline crossing |
| ENSDART00000055609 | atf4b | 0.027440234 | 0.138344185 | 0.51758353 | 0.353617358 | 0.087935611 | midline crossing |
| ENSDART00000138081 | bcl6ab | 0.000589104 | 0.948060654 | 0.690527167 | 0.988530239 | 1.012261919 | midline crossing |
| ENSDART00000140284 | zbtb16a | 0.028958693 | 2.407975491 | 2.022869932 | 2.341949009 | 2.679474288 | midline crossing |
| ENSDART00000132119 | max | 0.028918785 | 0.339991432 | 0.197083488 | 0.14057043 | 0.318686745 | midline crossing |
| ENSDART00000139763 | stat5a | 0.018355029 | 3.442434617 | 3.964844196 | 3.417693165 | 3.322184683 | midline crossing |
| ENSDART00000109552 | baz2ba | 0.004360669 | 0.627489244 | 0.525669422 | 0.712410774 | 0.393101067 | midline crossing |
| ENSDART00000029843 | vezf1a | 0.001561003 | 0.660507751 | 0.776194217 | 0.884683465 | 0.428803142 | midline crossing |
| ENSDART00000041877 | csrnp1a | 0.000166716 | 1.299872398 | 1.369159722 | 1.350463811 | 0.717287101 | midline crossing |
| ENSDART00000170865 | nme2b.1 | 0.007383951 | 0.635993303 | 0.679448572 | 0.689008761 | 0.355584206 | midline crossing |
| ENSDART00000135381 | six4a | 0.04980099 | 2.521510715 | 2.3671408 | 2.587075649 | 1.691172374 | midline crossing |
| ENSDART00000018743 | phf20a | 0.027269932 | 0.312029532 | 0.419745206 | 0.325238046 | 0.318832686 | midline crossing |
| ENSDART00000163794 | wt1b | 0.02934736 | 0.454982734 | 0.330694147 | 0.317167947 | 0.300420298 | midline crossing |
| ENSDART00000066317 | foxn2b | 0.033486585 | -0.194754205 | -0.013664444 | 0.221561118 | -0.080707177 | target selection |
| ENSDART00000158598 | si:ch211-232b12.5 | 0.004900248 | -0.304573383 | 0.212713124 | 0.452361247 | 0.038090441 | target selection |

| | | | | | | | |
|---|---|---|---|---|---|---|---|
| ENSDART00000154418 | relb | 0.010687727 | -0.043359819 | 0.083083005 | 0.26390054 | -0.071618881 | target selection |
| ENSDART00000114473 | trafd1 | 0.018765627 | -0.080068432 | 0.005557339 | 0.252422771 | -0.057760312 | target selection |
| ENSDART00000123197 | relb | 0.019118248 | -0.261655378 | -0.003226565 | 0.486237661 | -0.265984214 | target selection |
| ENSDART00000152489 | irf3 | 0.006723268 | -0.152502705 | 0.050924712 | 1.421424509 | -0.375867622 | target selection |
| ENSDART00000141792 | stat1b | 0.033143681 | -1.394223287 | -0.705218411 | 2.324563925 | -0.328164239 | target selection |
| ENSDART00000103622 | irf7 | 0.002828188 | -0.049332142 | -0.2878598 | 1.533925917 | -0.275107455 | target selection |
| ENSDART00000148661 | stat2 | 0.006978095 | 0.11967996 | -0.016653911 | 1.354000143 | 0.023510931 | target selection |
| ENSDART00000005720 | stat1a | 0.010250585 | 0.035880276 | 0.045195542 | 0.615137639 | 0.020671748 | target selection |
| ENSDART00000156935 | pbx4 | 0.039411387 | 0.960146191 | 0.964981474 | 3.716226941 | 3.606625876 | target selection |
| ENSDART00000151044 | baz2ba | 0.006344596 | 2.371956879 | 3.808025242 | 3.802702579 | 3.968575995 | target selection |
| ENSDART00000023959 | arntl1a | 7.44E-05 | 0.340904931 | 1.030627189 | 1.45255612 | 0.975945139 | target selection |
| ENSDART00000104523 | arntl1b | 0.000255127 | -0.158144123 | 0.59291526 | 1.113237486 | 0.52944604 | target selection |
| ENSDART00000083628 | ddit3 | 0.002192622 | -0.053032847 | 0.505000026 | 0.598711637 | 0.25299651 | target selection |
| ENSDART00000110512 | ybx1 | 0.014352201 | 0.187678935 | 0.306636132 | 0.543417204 | 0.157462635 | target selection |
| ENSDART00000135647 | si:ch211-113e8.10 | 0.041164305 | -0.117833263 | -0.003423122 | 0.202369294 | 0.16110756 | target selection |
| ENSDART00000025575 | clocka | 0.041666185 | -0.204691736 | 0.089686311 | 0.498135587 | 0.316118853 | target selection |
| ENSDART00000152636 | clocka | 0.010610046 | 0.008631834 | 0.196030905 | 0.593132542 | 0.426491394 | target selection |
| ENSDART00000089076 | dot1l | 0.009173096 | 0.094529933 | 0.165735937 | 0.448240264 | 0.463978672 | target selection |
| ENSDART00000013605 | zbtb20 | 0.00075553 | 0.021191929 | 0.070670156 | 0.462529396 | 0.345025042 | target selection |
| ENSDART00000165877 | purg | 0.000570625 | -0.159238984 | -0.087175961 | 1.048436006 | 0.831059287 | target selection |
| ENSDART00000000280 | stat1b | 0.037541166 | -0.005066017 | 0.035352481 | 0.819259146 | 0.412054224 | target selection |
| ENSDART00000087450 | klf13 | 0.013792445 | -0.22166629 | -0.059571459 | 0.622443971 | 0.288396102 | target selection |
| ENSDART00000146180 | csrnp2 | 0.003710792 | 0.237147162 | 0.374829692 | 0.697707124 | 0.402573932 | target selection |
| ENSDART00000026992 | sox4a | 3.66E-05 | 0.168735758 | 0.553498244 | 0.871364123 | 0.630648218 | target selection |
| ENSDART00000048110 | six4b | 0.002795018 | 0.757934328 | 2.322346312 | 2.776043581 | 1.88012445 | target selection |
| ENSDART00000013961 | mycla | 0.015752787 | 0.091841991 | 0.503579515 | 0.855574629 | 0.294011787 | target selection |
| ENSDART00000165656 | mxd3 | 6.24E-05 | 0.030444632 | 0.928898979 | 1.014066924 | 0.787179484 | target selection |
| ENSDART00000018150 | neurod6b | 0.012342358 | 0.484226112 | 1.695956927 | 1.935318885 | 1.555476315 | target selection |
| ENSDART00000084819 | arhgap35b | 0.001659006 | 0.101645005 | 0.291379647 | 0.535032879 | 0.462050203 | target selection |
| ENSDART00000075331 | insm1b | 0.009480138 | 0.180818368 | 0.558805202 | 0.852549514 | 0.856981121 | target selection |
| ENSDART00000123147 | zbtb8a | 0.026040305 | -0.281752517 | 0.027773484 | 0.518421131 | 0.366031095 | target selection |
| ENSDART00000018945 | meis3 | 0.00431294 | -0.263611366 | 0.389783427 | 1.01804147 | 0.898863275 | target selection |
| ENSDART00000111014 | myca | 0.024986086 | -0.198726525 | 0.073855205 | 0.316369866 | 0.312548093 | target selection |
| ENSDART00000092690 | srebf2 | 0.003411856 | -0.007030288 | 0.441941306 | 0.773565631 | 0.436893785 | target selection |
| ENSDART00000104712 | hmg20a | 0.038745219 | 0.003064442 | 0.120221197 | 0.366113134 | 0.275609671 | target selection |
| ENSDART00000033755 | homezb | 0.043076195 | -0.059847052 | 0.060728596 | 0.214721864 | 0.105513871 | target selection |
| ENSDART00000047724 | zeb1b | 0.032175544 | -0.128023017 | -0.020882446 | 0.202453179 | 0.103960026 | target selection |
| ENSDART00000085728 | klf8 | 0.014069453 | -0.229536876 | -0.061888352 | 0.204863755 | 0.114771443 | target selection |
| ENSDART00000037516 | znf827 | 0.032136194 | 0.117251719 | -0.001942433 | 0.235271462 | 0.223866066 | target selection |
| ENSDART00000150128 | foxp1b | 0.020184087 | 0.347545443 | 0.556370195 | 1.342974258 | 1.269825931 | target selection |
| ENSDART00000128174 | foxp1b | 0.015925333 | 0.060807795 | 0.283454627 | 0.694455914 | 0.748409918 | target selection |
| ENSDART00000060193 | thap3 | 0.039451639 | 0.219009088 | 0.189913345 | 0.546438828 | 0.599142566 | target selection |
| ENSDART00000154730 | tsc22d1 | 0.044459756 | 0.125146937 | 0.071380598 | 0.346511952 | 0.250641998 | target selection |
| ENSDART00000114246 | adnpb | 0.022770625 | -0.221340948 | -0.183838281 | -0.013868823 | 0.044049817 | brain innervation |
| ENSDART00000109356 | klf7a | 0.000177237 | -0.7063233 | -0.659659661 | -0.044315321 | 0.157900055 | brain innervation |
| ENSDART00000144186 | pou6f2 | 7.05E-05 | -0.474463335 | -0.465571046 | 0.092941552 | 0.436081081 | brain innervation |
| ENSDART00000124112 | pou4f2 | 2.95E-05 | -1.368026557 | -1.308261738 | 0.344262724 | 0.908586045 | brain innervation |
| ENSDART00000051693 | irx4a | 0.000591404 | -1.596922819 | -2.014349322 | 0.164610846 | 0.924604512 | brain innervation |
| ENSDART00000149175 | pou4f3 | 3.34E-05 | -1.520089487 | -1.231379123 | 0.357488642 | 0.743891028 | brain innervation |
| ENSDART00000055936 | isl2b | 4.92E-05 | -1.527471977 | -1.312125436 | 0.470803449 | 0.853726063 | brain innervation |
| ENSDART00000023038 | dacha | 0.000322945 | -0.511894205 | -0.418991667 | -0.004522839 | 0.112876213 | brain innervation |
| ENSDART00000142087 | foxp2 | 0.000565675 | -0.522019738 | -0.4194261 | 0.065656049 | 0.27497631 | brain innervation |
| ENSDART00000100508 | ebf1b | 0.010170292 | -0.252122108 | -0.356560297 | -0.077233007 | 0.218128715 | brain innervation |
| ENSDART00000162485 | pax6a | 0.013416969 | -0.383059238 | -0.321515523 | -0.107334897 | 0.077147684 | brain innervation |
| ENSDART00000112529 | znf319a | 0.023015173 | -0.271600655 | -0.228932764 | -0.01869534 | 0.01981331 | brain innervation |
| ENSDART00000165609 | barhl2 | 0.000401616 | -0.31272019 | -0.374018735 | 0.126371762 | 0.224852149 | brain innervation |
| ENSDART00000010896 | isl1 | 0.002539705 | -0.281634356 | -0.275090348 | 0.147086866 | 0.205392134 | brain innervation |
| ENSDART00000145072 | neurod4 | 0.04301836 | -0.171174556 | -0.165007281 | 0.082300141 | 0.101934405 | brain innervation |
| ENSDART00000165774 | pax6a | 0.007156543 | -0.382942423 | -0.218376905 | 0.094924321 | 0.282463268 | brain innervation |
| ENSDART00000101316 | znf618 | 0.019075692 | -0.33316659 | -0.306975689 | -0.091658088 | 0.079150867 | brain innervation |
| ENSDART00000064833 | mafaa | 0.002047801 | -0.957052225 | -1.143383392 | -0.379735148 | 0.067354966 | brain innervation |
| ENSDART00000004548 | barhl1b | 0.009605408 | -2.472825095 | -1.41656094 | -0.268019579 | 0.230183047 | brain innervation |
| ENSDART00000006380 | tbx3a | 0.004106215 | -0.244591708 | -0.352646993 | -0.144447405 | 0.06961498 | brain innervation |
| ENSDART00000101070 | dachd | 0.00434437 | -0.296270747 | -0.351582482 | -0.130791564 | -0.009393292 | brain innervation |
| ENSDART00000151904 | foxn3 | 0.000466769 | -0.402174688 | -0.533042351 | -0.141780798 | 0.022724604 | brain innervation |
| ENSDART00000124751 | kcnip3b | 2.63E-05 | -1.29606503 | -1.383048631 | -0.490749215 | -0.039127508 | brain innervation |
| ENSDART00000151571 | ahdc1 | 0.000666959 | -0.753523402 | -0.774930454 | -0.265860844 | -0.025283973 | brain innervation |
| ENSDART00000080808 | six3a | 0.000687981 | -0.437705937 | -0.430575558 | -0.140593988 | -0.07114268 | brain innervation |
| ENSDART00000171594 | mef2aa | 0.010231208 | -0.649844171 | -0.685014218 | -0.301258241 | -0.003741614 | brain innervation |
| ENSDART00000062982 | foxo3b | 0.005646833 | -0.232285352 | -0.219543276 | -0.013673206 | 0.18814406 | brain innervation |
| ENSDART00000009691 | scml4 | 0.001274316 | -0.348233268 | -0.325522885 | -0.113421585 | 0.115178638 | brain innervation |
| ENSDART00000160784 | satb1b | 0.006456063 | -0.404843174 | -0.932859181 | 0.352272227 | 0.695966701 | brain innervation |
| ENSDART00000101631 | satb1b | 0.000172812 | -0.360112936 | -0.384116906 | 0.134197196 | 0.481946288 | brain innervation |

| | | | | | | | |
|---|---|---|---|---|---|---|---|
| ENSDART00000144894 | bcl11ba | 3.52E-05 | -0.656989822 | -0.679320488 | 0.181338486 | 0.623847694 | brain innervation |
| ENSDART00000165698 | pbx1a | 0.000272902 | -0.205212048 | -0.330154536 | 0.047550564 | 0.321002436 | brain innervation |
| ENSDART00000081170 | cux1a | 0.000900234 | -0.092953946 | -0.2283391 | 0.138990685 | 0.296890549 | brain innervation |
| ENSDART00000160675 | dpf1 | 0.00210146 | -0.344267461 | -0.462495684 | 0.239034332 | 0.46811192 | brain innervation |
| ENSDART00000045628 | irx6a | 0.000781013 | -0.424070635 | -0.239579413 | 0.17038084 | 0.276181325 | brain innervation |
| ENSDART00000048994 | pbx3b | 0.000910662 | -0.545498331 | -0.428509616 | 0.129533791 | 0.149586816 | brain innervation |
| ENSDART00000077157 | six3b | 0.008086296 | -0.273810969 | -0.17356253 | 0.033591817 | 0.098241195 | brain innervation |
| ENSDART00000136245 | cux1a | 0.013366886 | -0.163227222 | -0.237382603 | -0.00302824 | 0.164419575 | brain innervation |
| ENSDART00000154646 | bcl11ba | 0.001677265 | -0.533670184 | -0.516802708 | 0.152594691 | 0.570166001 | brain innervation |
| ENSDART00000153150 | myt1la | 0.001328724 | -0.378588425 | -0.355041474 | 0.131237693 | 0.393706803 | brain innervation |
| ENSDART00000166565 | tshz3b | 0.011919383 | -0.202679362 | -0.107608077 | 0.106987356 | 0.28761259 | brain innervation |
| ENSDART00000013148 | pou3f1 | 0.00849815 | -0.360437586 | -0.322601739 | -0.060819622 | 0.211329038 | brain innervation |
| ENSDART00000162945 | sox5 | 0.023596757 | -0.177385379 | -0.174200373 | -0.003535976 | 0.152149827 | brain innervation |
| ENSDART00000171091 | zeb2b | 0.008482497 | -0.344020449 | -0.404143226 | -0.222307379 | 0.140992606 | brain innervation |
| ENSDART00000170631 | ebf1a | 0.016255169 | -0.245696473 | -0.326200552 | -0.132641619 | 0.115308885 | brain innervation |
| ENSDART00000156464 | prr12b | 0.021599545 | -0.068684116 | -0.215586059 | 0.031477455 | 0.130187301 | brain innervation |
| ENSDART00000167440 | zfhx3 | 0.018342429 | -0.067144102 | -0.226395351 | 0.21751863 | 0.266797189 | brain innervation |
| ENSDART00000167538 | lcor | 0.031258003 | -0.151917658 | -0.14482463 | 0.130691961 | 0.223379898 | brain innervation |
| ENSDART00000165223 | pbx1b | 0.015188807 | -0.243991197 | -0.377373898 | -0.18693501 | 0.141565589 | brain innervation |
| ENSDART00000172410 | jazf1a | 0.045787549 | -0.354446023 | -0.388940674 | -0.011638795 | -0.018894552 | brain innervation |
| ENSDART00000022010 | hivep2b | 0.004657196 | -0.404683257 | -0.428225913 | -0.085969125 | 0.061919909 | brain innervation |
| ENSDART00000138048 | pbx3b | 0.007841242 | -0.599655595 | -0.602159282 | -0.228775798 | -0.019503135 | brain innervation |
| ENSDART00000137355 | tet3 | 0.033062691 | -0.245273707 | -0.407219331 | -0.050664281 | -0.06569376 | brain innervation |
| ENSDART00000104790 | znf292b | 0.044988705 | -0.062768954 | -0.233087007 | -0.040681021 | 0.023710275 | brain innervation |
| ENSDART00000136729 | ebf1b | 0.021027344 | -2.79991744 | -3.068779235 | -0.720206417 | 0.313112936 | brain innervation |
| ENSDART00000132589 | mef2d | 0.028030725 | -0.20910625 | -0.209165039 | -0.020444617 | 0.107291243 | brain innervation |
| ENSDART00000003884 | mynn | 0.033340088 | -0.108806257 | -0.247003557 | 0.197551703 | 0.151198862 | brain innervation |
| ENSDART00000104327 | vsx1 | 0.00488592 | -0.248525466 | -0.619641572 | -0.034766195 | -0.028538746 | brain innervation |
| ENSDART00000084035 | znf532 | 0.047492859 | -0.199573234 | -0.524026601 | -0.06012509 | 0.154210784 | brain innervation |
| ENSDART00000134064 | mxd1 | 0.015571917 | -0.125104323 | -0.265030232 | -0.047451672 | 0.092856449 | brain innervation |
| ENSDART00000136759 | prdm13 | 0.00326136 | -0.225066145 | -0.413438688 | -0.129520305 | 0.036146962 | brain innervation |
| ENSDART00000166540 | kcnip3b | 0.047816671 | -0.988549414 | -0.704807394 | -0.278733631 | 0.119291619 | brain innervation |
| ENSDART00000030773 | foxo3a | 0.000701449 | -0.522674186 | -0.346284342 | -0.025883071 | 0.086783781 | brain innervation |
| ENSDART00000111842 | sall1a | 0.033435193 | -0.3702182 | -0.245271728 | -0.043742586 | 0.007554245 | brain innervation |
| ENSDART00000166976 | esrrb | 0.030820243 | -0.481593528 | -0.114040699 | 0.229959907 | 0.235353141 | brain innervation |
| ENSDART00000151575 | thraa | 0.013354221 | -0.36498113 | -0.584870078 | 0.329302923 | -0.150628947 | brain innervation |
| ENSDART00000007806 | zbtb16a | 0.004364661 | -0.634450415 | -0.731920804 | 0.190781828 | -0.364381887 | brain innervation |
| ENSDART00000125158 | znf574 | 0.029152083 | -0.158249462 | -0.223654122 | 0.188172427 | 0.054732788 | brain innervation |
| ENSDART00000011890 | zbtb22a | 0.03843558 | -0.42030539 | -0.536466631 | 0.147549186 | 0.05725602 | brain innervation |
| ENSDART00000155866 | zbtb16b | 8.49E-05 | -0.434457287 | -0.595844526 | 0.044569606 | 0.015262284 | brain innervation |
| ENSDART00000001795 | sp3a | 0.0407578 | -0.269877105 | -0.305437708 | -0.047306155 | -0.033496274 | brain innervation |
| ENSDART00000109493 | tshz2 | 0.008648063 | -0.247840671 | 0.10394662 | 0.365597315 | 0.467215515 | brain innervation |
| ENSDART00000083296 | ZNF423 | 0.00814358 | -0.278315489 | 0.051691519 | 0.407184973 | 0.489807123 | brain innervation |
| ENSDART00000058685 | zfpm2a | 0.011667422 | -0.080421638 | -0.032548615 | 0.512175645 | 0.757319343 | brain innervation |
| ENSDART00000110069 | CABZ01075131.1 | 0.008675958 | -0.117445692 | -0.062527677 | 0.307574487 | 0.429328576 | brain innervation |
| ENSDART00000131126 | bcl6a | 0.005688069 | -0.096649799 | -0.044696983 | 0.295011601 | 0.436349263 | brain innervation |
| ENSDART00000111303 | rfx7 | 0.00161581 | 0.075232989 | 0.085195342 | 0.459783838 | 0.727409902 | brain innervation |
| ENSDART00000122101 | tbx2b | 0.001190205 | -0.056468629 | 0.017248499 | 0.19422425 | 0.382938949 | brain innervation |
| ENSDART00000065139 | gfi1ab | 0.023335678 | -0.252440054 | -0.123023724 | 0.269699656 | 0.68808671 | brain innervation |
| ENSDART00000150938 | sox6 | 0.001331938 | -0.111707858 | -0.047191704 | 0.355284385 | 0.700337568 | brain innervation |
| ENSDART00000012862 | isl2a | 3.29E-05 | -0.663813396 | 0.011664777 | 0.746201858 | 1.074694305 | brain innervation |
| ENSDART00000158634 | ebf3a | 0.006418819 | -0.930015824 | -0.218309076 | 0.836197649 | 1.139950385 | brain innervation |
| ENSDART00000166502 | satb2 | 1.14E-05 | -0.540440017 | -0.158642442 | 1.216799418 | 1.496340928 | brain innervation |
| ENSDART00000025031 | pou4f1 | 1.46E-05 | -0.37458842 | -0.169318799 | 0.774588614 | 1.160187758 | brain innervation |
| ENSDART00000016303 | irx2a | 4.96E-05 | -0.381386746 | 0.064410059 | 0.752093904 | 0.989751717 | brain innervation |
| ENSDART00000098859 | neurod6a | 0.000108364 | -0.523659161 | -0.308368621 | 0.583701544 | 0.979244768 | brain innervation |
| ENSDART00000031167 | tfap2d | 0.000251123 | -0.967562253 | -0.629795841 | 0.638492839 | 1.047009841 | brain innervation |
| ENSDART00000006612 | tbr1b | 2.95E-05 | -1.166029945 | -1.008207372 | 0.617040877 | 1.169470893 | brain innervation |
| ENSDART00000157855 | ebf3a | 0.000670771 | -0.765871102 | -0.436598509 | 0.580406091 | 0.942511549 | brain innervation |
| ENSDART00000167324 | ebf3a | 4.92E-05 | -0.585714789 | -0.243731213 | 0.563349846 | 0.843370215 | brain innervation |
| ENSDART00000128504 | si:dkey-106g10.7 | 0.03328728 | 0.018091706 | 0.0404047 | 0.309180984 | 0.459844792 | brain innervation |
| ENSDART00000165955 | zhx3 | 0.013572162 | -0.011624436 | -0.109014479 | 0.013312322 | 0.351264725 | brain innervation |
| ENSDART00000092357 | sgsm2 | 0.018543094 | -0.379055867 | -0.375730796 | -0.097265456 | -0.053118339 | brain innervation |
| ENSDART00000170400 | ahdc1 | 0.034543013 | -2.952674589 | -2.088515533 | 0.893679477 | 1.203421878 | brain innervation |
| ENSDART00000150949 | nfyc | 0.013743075 | -0.535274529 | -0.301463031 | -0.212807295 | 0.074596031 | brain innervation |
| ENSDART00000166351 | nkrf | 0.010292856 | -0.332043429 | -0.213944917 | -0.117297764 | -0.03436019 | brain innervation |
| ENSDART00000090019 | zeb2b | 0.016845819 | -0.694941899 | -0.428168304 | -0.296660745 | 0.024845807 | brain innervation |
| ENSDART00000018603 | tbx4 | 0.04873911 | -0.302508569 | -0.188111077 | -0.131947864 | 0.000377602 | brain innervation |
| ENSDART00000171642 | dpf1 | 0.005353377 | -0.380689139 | 0.060927985 | 0.320062308 | 0.358114425 | brain innervation |
| ENSDART00000113087 | hmx1 | 0.000694227 | -0.389387565 | 0.021377927 | 0.234018679 | 0.270888749 | brain innervation |
| ENSDART00000162474 | irx1a | 0.002931603 | -1.132797992 | -0.349993381 | 0.581252213 | 0.55366573 | brain innervation |
| ENSDART00000162850 | irx3a | 0.000148815 | -0.694816328 | -0.301932125 | 0.187764734 | 0.208405543 | brain innervation |
| ENSDART00000172990 | znf628 | 0.031889344 | -0.174846517 | -0.081526828 | 0.094662541 | 0.071436879 | brain innervation |

| | | | | | | | |
|---|---|---|---|---|---|---|---|
| ENSDART00000112756 | ZNF608 | 0.004908881 | -0.246715323 | -0.122479962 | 0.171371612 | 0.114220238 | brain innervation |
| ENSDART00000090226 | znf319b | 0.009551702 | -0.155602419 | -0.092883138 | 0.255379558 | 0.168359048 | brain innervation |
| ENSDART00000053782 | scrt2 | 0.003310987 | -0.252422437 | -0.133353661 | 0.196438899 | 0.160308382 | brain innervation |
| ENSDART00000100658 | esrra | 8.74E-05 | -0.571475474 | -0.395296451 | 0.169003635 | 0.212368184 | brain innervation |
| ENSDART00000008906 | znf503 | 0.001473115 | -0.367880648 | -0.389528031 | 0.161538025 | 0.184463 | brain innervation |
| ENSDART00000164488 | mbnl2 | 0.0337199 | -3.245293256 | -3.314362259 | 0.902121491 | 1.413765252 | brain innervation |
| ENSDART00000130163 | nr2f2 | 0.012656704 | -0.280168769 | -0.057880311 | 0.350863039 | 0.468160546 | brain innervation |
| ENSDART00000074833 | rx3 | 0.010379183 | -0.279918529 | -0.301637969 | 0.143576872 | 0.172520697 | brain innervation |
| ENSDART00000006737 | sox5 | 0.038386468 | -0.253791611 | -0.278700286 | 0.184042389 | 0.275951824 | brain innervation |
| ENSDART00000109347 | nacc2 | 0.002373876 | -0.303011569 | -0.338345203 | 0.399887844 | 0.337648971 | brain innervation |
| ENSDART00000133959 | clockb | 0.010583972 | -0.261160651 | -0.146340415 | 0.224249915 | 0.15359596 | brain innervation |
| ENSDART00000108775 | ubp1 | 0.021499635 | -0.192682892 | -0.115240547 | 0.10453418 | 0.097860664 | brain innervation |
| ENSDART00000092665 | srebf1 | 0.004619636 | -0.417658309 | -0.274360919 | 0.114636817 | 0.022214824 | brain innervation |
| ENSDART00000104751 | mxi1 | 0.002350825 | -0.384587031 | -0.210909301 | 0.22179917 | 0.104111801 | brain innervation |
| ENSDART00000131714 | mbnl2 | 0.008848885 | -0.406601788 | -0.032543807 | -0.091713267 | 0.092369986 | brain innervation |
| ENSDART00000124534 | mbnl2 | 0.001000183 | -0.407254222 | 0.078580914 | -0.044925118 | 0.120266311 | brain innervation |
| ENSDART00000155666 | znf576.1 | 0.015256508 | -0.372721752 | -0.23410815 | -0.134923201 | -0.013577883 | brain innervation |
| ENSDART00000047541 | bach1b | 0.039802589 | -0.343772395 | -0.510091534 | -0.468889885 | -0.333515696 | downregulated during regeneration - early |
| ENSDART00000007630 | nhlh2 | 0.010056242 | -0.28794611 | -0.324673306 | -0.385652145 | -0.261092419 | downregulated during regeneration - early |
| ENSDART00000129498 | mef2d | 0.03679078 | -0.188451231 | -0.381445449 | -0.263610552 | -0.197973462 | downregulated during regeneration - early |
| ENSDART00000055890 | znf385c | 0.011543549 | -0.397413591 | -0.444425008 | -0.419820845 | -0.297765515 | downregulated during regeneration - early |
| ENSDART00000024832 | stat5a | 0.004077075 | -0.212236095 | -0.408877789 | -0.375060924 | -0.185569558 | downregulated during regeneration - early |
| ENSDART00000082346 | tfap2a | 0.038054224 | -0.238141907 | -0.307957291 | -0.242159487 | -0.230339237 | downregulated during regeneration - early |
| ENSDART00000013003 | tfap2b | 0.008192536 | -0.299930992 | -0.578681033 | -0.25971747 | -0.279510711 | downregulated during regeneration - early |
| ENSDART00000167052 | etv1 | 0.01072353 | -0.327639473 | -0.469002174 | -0.32895701 | -0.22464313 | downregulated during regeneration - early |
| ENSDART00000157659 | camta1b | 0.01608096 | -0.574265168 | -0.690891494 | -0.316064045 | -0.339000781 | downregulated during regeneration - early |
| ENSDART00000169609 | tefb | 0.032368681 | -0.156114581 | -0.555141888 | -0.301378906 | -0.24132076 | downregulated during regeneration - early |
| ENSDART00000103640 | hey1 | 0.023203992 | -0.385659584 | -0.747117397 | -0.392306381 | -0.416692063 | downregulated during regeneration - early |
| ENSDART00000089748 | rorb | 0.030623213 | -0.330969943 | -0.539847879 | -0.425468683 | -0.138099942 | downregulated during regeneration - early |
| ENSDART00000122429 | klf15 | 0.006222767 | -0.341190152 | -0.616050077 | -0.578895446 | -0.337948098 | downregulated during regeneration - early |
| ENSDART00000153187 | thrab | 0.001402519 | -0.362356661 | -0.551240408 | -0.403932333 | -0.192068079 | downregulated during regeneration - early |
| ENSDART00000054736 | bhlhe23 | 0.004928313 | -0.369485488 | -0.514619457 | -0.390416516 | -0.190127988 | downregulated during regeneration - early |
| ENSDART00000125344 | skilb | 0.010360832 | -0.241964121 | -0.627384358 | -0.493621211 | -0.368838752 | downregulated during regeneration - early |
| ENSDART00000131361 | kcnip3b | 0.008595135 | -0.353450144 | -1.035812483 | -0.692091767 | -0.071599751 | downregulated during regeneration - early |
| ENSDART00000026017 | bhlhe40 | 0.046057741 | -0.186426622 | -0.329757373 | -0.286559283 | -0.071751826 | downregulated during regeneration - early |
| ENSDART00000131134 | si:ch211-206a7.2 | 0.008848885 | -0.226522125 | -0.472315068 | -0.371394997 | -0.101330709 | downregulated during regeneration - early |
| ENSDART00000046663 | camta1b | 0.022220725 | -0.175908238 | -0.543108898 | -0.289499765 | -0.001152787 | downregulated during regeneration - early |
| ENSDART00000172199 | zmat4a | 0.029447477 | -0.170025595 | -0.220098279 | -0.198411401 | 0.018178403 | downregulated during regeneration - early |
| ENSDART00000142223 | hivep3b | 0.013948629 | -0.312228644 | -0.441058379 | -0.334712198 | -0.18438466 | downregulated during regeneration - early |
| ENSDART00000147658 | bhlhe22 | 0.01205445 | -0.302235414 | -0.31899187 | -0.305987775 | -0.102192149 | downregulated during regeneration - early |
| ENSDART00000126966 | esrrga | 0.017679749 | -0.248566713 | -0.265310961 | -0.321308408 | -0.070523791 | downregulated during regeneration - early |
| ENSDART00000127353 | nr1d2b | 0.027085654 | -0.29103008 | -0.243688493 | -0.346507775 | -0.100908835 | downregulated during regeneration - early |
| ENSDART00000040672 | mecp2 | 0.047533941 | -0.242382035 | -0.413395534 | -0.149398689 | -0.148226048 | downregulated during regeneration - early |
| ENSDART00000057644 | lhx4 | 0.048389146 | -0.267668007 | -0.268543602 | -0.190125613 | -0.119751289 | downregulated during regeneration - early |
| ENSDART00000089923 | znf652 | 0.026337068 | -0.30560984 | -0.322535846 | -0.127141181 | -0.092457617 | downregulated during regeneration - early |
| ENSDART00000157890 | tcf7l1b | 0.035686017 | -4.555123239 | -4.673271453 | -1.829433189 | -0.732249154 | downregulated during regeneration - early |
| ENSDART00000136488 | zmat4a | 0.008573382 | -0.690962776 | -0.637104526 | -0.28999313 | -0.088561559 | downregulated during regeneration - early |
| ENSDART00000164349 | e2f4 | 0.042377891 | -0.246759183 | -0.226331986 | -0.10688884 | -0.101679787 | downregulated during regeneration - early |
| ENSDART00000003548 | znf385a | 0.001069696 | -0.544762633 | -0.635515273 | -0.316063497 | -0.128464507 | downregulated during regeneration - early |
| ENSDART00000033362 | gatad2b | 0.00857344 | -0.40244942 | -0.286045422 | -0.214483951 | -0.084688348 | downregulated during regeneration - early |
| ENSDART00000026865 | l3mbtl1a | 0.016106242 | -0.339631378 | -0.338161449 | -0.153879513 | -0.055902641 | downregulated during regeneration - early |
| ENSDART00000037879 | crx | 0.018920791 | -0.254014844 | -0.310425293 | -0.22793154 | -0.020194538 | downregulated during regeneration - early |
| ENSDART00000135443 | kdm5ba | 0.039457626 | -0.212935098 | -0.210173481 | -0.205976005 | -0.095029648 | downregulated during regeneration - early |
| ENSDART00000165757 | pax6b | 0.019981901 | -1.030621281 | -0.736504391 | -0.510180699 | -0.112500923 | downregulated during regeneration - early |
| ENSDART00000059446 | znf385b | 0.027304375 | -0.260906147 | -0.315001382 | -0.161772411 | -0.110285684 | downregulated during regeneration - early |
| ENSDART00000165120 | purab | 0.041240008 | -0.246648902 | -0.188751457 | -0.157104845 | -0.035763175 | downregulated during regeneration - early |
| ENSDART00000009938 | tcf12 | 0.025246927 | -0.385864267 | -0.351608927 | -0.283149928 | -0.323464659 | downregulated during regeneration - early |
| ENSDART00000048775 | mbd3b | 0.027491007 | -0.291847963 | -0.361950456 | -0.358913558 | -0.222238531 | downregulated during regeneration - early |
| ENSDART00000156351 | zfat | 0.027293911 | -0.386268069 | -0.362611932 | -0.387180541 | -0.301395116 | downregulated during regeneration - early |
| ENSDART00000166135 | zbtb47b | 0.004311817 | -0.402709337 | -0.409852132 | -0.474957603 | -0.394147628 | downregulated during regeneration - early |
| ENSDART00000055706 | her15.1 | 0.021144178 | -1.483608056 | -1.154441068 | -0.698110478 | -1.436427969 | downregulated during regeneration - early |
| ENSDART00000133487 | fosb | 0.034512759 | -0.882572207 | -1.273781011 | -0.772506712 | -0.914864674 | downregulated during regeneration - early |
| ENSDART00000000876 | nr4a1 | 0.000894056 | -2.262372442 | -2.399082592 | -2.145415905 | -1.543723254 | downregulated during regeneration - early |
| ENSDART00000151127 | thraa | 0.039890399 | -0.798583389 | -0.802364861 | -0.351487338 | -0.589241182 | downregulated during regeneration - early |
| ENSDART00000162857 | nr4a3 | 0.003361003 | -1.482016444 | -1.152680943 | -0.606031454 | -1.193862677 | downregulated during regeneration - early |
| ENSDART00000153167 | hlfb | 0.010966965 | -0.470839666 | -0.326611575 | -0.235752692 | -0.264047019 | downregulated during regeneration - early |
| ENSDART00000065361 | etv5b | 0.019698661 | -0.645686026 | -0.347502384 | -0.293939032 | -0.317696675 | downregulated during regeneration - early |
| ENSDART00000162387 | zbtb38 | 0.015925333 | -0.362042869 | -0.285747988 | -0.199364681 | -0.27814593 | downregulated during regeneration - early |
| ENSDART00000172310 | zbtb4 | 0.013310088 | -0.365422353 | -0.277101341 | -0.237436935 | -0.247791483 | downregulated during regeneration - early |
| ENSDART00000054020 | hivep3b | 0.042576139 | 0.041000622 | -0.364922266 | -0.147208921 | -0.149490959 | downregulated during regeneration - mid |
| ENSDART00000169283 | znf644b | 0.046753078 | 0.013670999 | -0.206539721 | -0.202072284 | -0.100289754 | downregulated during regeneration - mid |
| ENSDART00000161387 | tcf12 | 0.008531138 | 0.370378706 | 0.009395421 | -0.374363269 | -0.152945712 | downregulated during regeneration - mid |
| ENSDART00000127099 | nr2e3 | 0.030676359 | 0.281603482 | 0.002911001 | -0.560613973 | -0.033774044 | downregulated during regeneration - mid |

| | | | | | | | |
|---|---|---|---|---|---|---|---|
| ENSDART00000163250 | mef2cb | 0.003671994 | 0.361224616 | -0.300957895 | -0.57227951 | -0.080214844 | downregulated during regeneration - mid |
| ENSDART00000131731 | mef2ca | 0.006008543 | 0.234535922 | -0.250149626 | -0.381098175 | -0.047609857 | downregulated during regeneration - mid |
| ENSDART00000113286 | phf19 | 0.035367252 | 0.32344819 | -0.020838317 | -0.493273441 | -0.244699917 | downregulated during regeneration - mid |
| ENSDART00000093331 | rreb1a | 0.018126285 | -0.024901133 | -0.155149005 | -0.517234878 | -0.050769944 | downregulated during regeneration - mid |
| ENSDART00000104279 | znf516 | 0.018737197 | -0.151066862 | -0.30868422 | -0.587761132 | -0.246031248 | downregulated during regeneration - mid |
| ENSDART00000143165 | tsc22d1 | 0.012984014 | -0.098364585 | -0.21923408 | -0.39849137 | 0.017570634 | downregulated during regeneration - mid |
| ENSDART00000148537 | roraa | 0.01612441 | -0.160828233 | -0.221872095 | -0.45696721 | -0.083100906 | downregulated during regeneration - mid |
| ENSDART00000100667 | skia | 0.021156666 | -0.180939747 | -0.108825609 | -0.328725059 | -0.076853639 | downregulated during regeneration - mid |
| ENSDART00000123970 | mntb | 0.011725523 | -0.115861458 | -0.101461538 | -0.360694649 | -0.01567692 | downregulated during regeneration - mid |
| ENSDART00000077839 | atf7b | 0.014656899 | -0.232485852 | -0.196508033 | -0.391624214 | -0.206346003 | downregulated during regeneration - mid |
| ENSDART00000165710 | gpbp1l1 | 0.036571732 | -0.164201122 | -0.182753901 | -0.290062625 | -0.125147623 | downregulated during regeneration - mid |
| ENSDART00000124740 | ncoa2 | 0.048038072 | -0.028796998 | -0.187850383 | -0.230877254 | 0.045612327 | downregulated during regeneration - mid |
| ENSDART00000164082 | znf609a | 0.041293307 | 0.094336402 | -0.129509301 | -0.187095832 | 0.037446717 | downregulated during regeneration - mid |
| ENSDART00000045374 | smad3a | 0.043076195 | -0.063329624 | -0.157712113 | -0.303211349 | -0.108940629 | downregulated during regeneration - mid |
| ENSDART00000166313 | thrb | 0.0223553 | -0.193382381 | -0.415286369 | -0.400030228 | -0.136855946 | downregulated during regeneration - mid |
| ENSDART00000031426 | skilb | 0.000177771 | -0.234186907 | -0.553407151 | -0.565936684 | -0.420869627 | downregulated during regeneration - mid |
| ENSDART00000148106 | mef2aa | 0.026582721 | -0.081330082 | -0.55524128 | -0.609035118 | -0.185252531 | downregulated during regeneration - mid |
| ENSDART00000091707 | dbpa | 0.017575469 | -0.158262702 | -0.327851995 | -0.357551076 | -0.129469939 | downregulated during regeneration - mid |
| ENSDART00000148066 | znf395b | 0.020342933 | -0.110739145 | -0.366906743 | -0.460042969 | -0.183105049 | downregulated during regeneration - mid |
| ENSDART00000075070 | hsf2 | 0.014034602 | -0.066635095 | -0.276496416 | -0.474097609 | -0.186498896 | downregulated during regeneration - mid |
| ENSDART00000148353 | usf2 | 0.006964046 | -0.107987138 | -0.265954127 | -0.405201274 | -0.200197809 | downregulated during regeneration - mid |
| ENSDART00000061106 | bhlhe41 | 0.000553147 | 0.043056173 | -0.423617996 | -0.754959095 | -0.246095493 | downregulated during regeneration - mid |
| ENSDART00000139102 | dbpb | 0.001993937 | 0.113164671 | -0.364559303 | -0.423196921 | -0.137052116 | downregulated during regeneration - mid |
| ENSDART00000100181 | sall3b | 0.034764006 | -0.170454734 | -0.209016072 | -0.285639823 | -0.232809645 | downregulated during regeneration - mid |
| ENSDART00000141734 | hivep2a | 0.017814842 | -0.218148846 | -0.337091836 | -0.481607769 | -0.327040294 | downregulated during regeneration - mid |
| ENSDART00000170575 | nfat5b | 0.018611393 | -0.001493923 | -0.392122738 | -0.566909919 | -0.288747412 | downregulated during regeneration - mid |
| ENSDART00000143874 | akna | 0.021937256 | -0.124615674 | -0.323808831 | -0.482322372 | -0.23639171 | downregulated during regeneration - mid |
| ENSDART00000127157 | hlfa | 0.048701514 | -0.089881453 | -0.212609643 | -0.346930267 | -0.147102401 | downregulated during regeneration - mid |
| ENSDART00000126282 | nr1d1 | 0.008088213 | -0.1014969 | -0.191656676 | -0.494430716 | -0.13667305 | downregulated during regeneration - mid |
| ENSDART00000078529 | kin | 0.03060143 | -1.437736057 | -0.295663012 | -0.493580296 | -4.734617185 | downregulated during regeneration - late |
| ENSDART00000055709 | her2 | 0.030146589 | -0.984524994 | -0.833253569 | -0.717745512 | -1.233522642 | downregulated during regeneration - late |
| ENSDART00000023613 | her6 | 0.030622517 | -0.335042933 | -0.234567638 | -0.250120727 | -0.59712361 | downregulated during regeneration - late |
| ENSDART00000086051 | mecom | 0.04049764 | -0.550304416 | -0.298852875 | -0.293768303 | -1.334615988 | downregulated during regeneration - late |
| ENSDART00000110544 | znf219 | 0.031344075 | -0.234403747 | -0.210413301 | -0.211690001 | -0.32688667 | downregulated during regeneration - late |
| ENSDART00000053367 | hmgn3 | 0.044180927 | -0.174648563 | -0.165335262 | -0.310027237 | -0.181225286 | downregulated during regeneration - late |
| ENSDART00000089015 | zbtb7a | 0.00191624 | -0.478666207 | -0.203919761 | -0.738823143 | -0.713706626 | downregulated during regeneration - late |
| ENSDART00000164855 | crebl2 | 0.013882038 | -0.127345071 | -0.2122205 | -0.432997567 | -0.264872881 | downregulated during regeneration - late |
| ENSDART00000066655 | mybl1 | 0.005666275 | -0.255810306 | -0.487313899 | -0.72749049 | -0.668391364 | downregulated during regeneration - late |
| ENSDART00000125174 | nr1i2 | 0.028902271 | -0.056601149 | -0.085618729 | -0.264588441 | -0.326132843 | downregulated during regeneration - late |
| ENSDART00000167844 | mafk | 0.014095383 | -0.00863672 | 0.25450476 | -0.276014619 | -0.283559487 | downregulated during regeneration - late |
| ENSDART00000057124 | tefa | 0.007560757 | -0.17959692 | 0.074135039 | -0.330628318 | -0.262330789 | downregulated during regeneration - late |
| ENSDART00000057125 | tefa | 0.002939873 | -0.190705988 | 0.094171259 | -0.306461854 | -0.301060608 | downregulated during regeneration - late |
| ENSDART00000172251 | creb3l1 | 0.029283678 | -0.33540662 | 0.193573394 | -4.078076569 | -0.842213331 | downregulated during regeneration - late |
| ENSDART00000128815 | stra13 | 0.002651628 | 0.38711598 | 0.144426216 | -0.403589602 | -0.227357691 | downregulated during regeneration - late |
| ENSDART00000005263 | usf1l | 0.019856195 | 0.061508869 | 0.080355717 | -0.137096705 | -0.208096843 | downregulated during regeneration - late |
| ENSDART00000044860 | maff | 0.002310171 | 0.057535561 | 0.039208376 | -0.366680053 | -0.441007369 | downregulated during regeneration - late |
| ENSDART00000078781 | znf706 | 0.034124552 | 0.015897963 | -0.052514643 | -0.230153549 | -0.188792592 | downregulated during regeneration - late |

**Table S6. Differentially expressed transcripts that encode transcription factors with known motifs.**

| Ensembl ID | zebrafish gene symbo | padj | LFC: 2dpi-0dpi | LFC: 4dpi-0dpi | LFC: 7dpi-0dpi | LFC: 12dpi-0dpi | cluster name |
|---|---|---|---|---|---|---|---|
| ENSDART00000140760 | e2f7 | 0.044060134 | 0.798632306 | 0.496792942 | -0.026056819 | -0.454809271 | growth toward the midline |
| ENSDART00000056005 | ascl1a | 0.032093457 | 1.787445127 | 1.283325532 | 0.749862779 | 0.792492648 | growth toward the midline |
| ENSDART00000128488 | e2f8 | 0.003382652 | 3.852661131 | -0.929008264 | 3.252279997 | 2.01178299 | growth toward the midline |
| ENSDART00000132119 | max | 0.028918785 | 0.339991432 | 0.197083488 | 0.14057043 | 0.318686745 | growth toward the midline |
| ENSDART00000163794 | wt1b | 0.02934736 | 0.454982734 | 0.330694147 | 0.317167947 | 0.300420298 | growth toward the midline |
| ENSDART00000033494 | klf6a | 0.000170861 | 1.084550888 | 1.797692613 | 1.625562666 | 0.460444617 | growth toward the midline |
| ENSDART00000147849 | klf6a | 0.000870446 | 1.130982604 | 1.870693821 | 1.403514717 | 0.445060502 | growth toward the midline |
| ENSDART00000076161 | hoxb5b | 0.013675681 | 2.652791903 | 3.351280664 | 2.618897772 | 0.864721852 | growth toward the midline |
| ENSDART00000122628 | junba | 0.016136147 | 0.280813702 | 0.583374691 | 0.41181471 | 0.073998168 | growth toward the midline |
| ENSDART00000146767 | fosl1a | 0.046117855 | 0.841007094 | 1.285901651 | 0.733422298 | 0.173089455 | growth toward the midline |
| ENSDART00000161610 | tcf3b | 0.04032679 | 0.066966853 | 0.197082202 | 0.107652202 | -0.03222147 | growth toward the midline |
| ENSDART00000051549 | tp53 | 0.029170799 | 0.110640197 | 0.281712522 | 0.180041895 | -0.162598759 | growth toward the midline |
| ENSDART00000164711 | NFATC2 (1 of many) | 0.017032668 | 0.774520656 | 0.760515741 | 0.734404519 | 0.027133082 | growth toward the midline |
| ENSDART00000063912 | jun | 0.002341904 | 1.166146477 | 1.429328618 | 1.29995447 | 0.480560375 | growth toward the midline |
| ENSDART00000022060 | atf3 | 2.20E-05 | 2.958540036 | 3.348918067 | 2.852908333 | 1.811501419 | growth toward the midline |
| ENSDART00000110040 | sox11a | 8.49E-05 | 1.573346229 | 1.723237215 | 1.23195588 | 0.781584101 | growth toward the midline |
| ENSDART00000104519 | stat3 | 0.004480141 | 0.6458865 | 0.459098575 | 0.438501049 | 0.201812376 | growth toward the midline |
| ENSDART00000142584 | alx1 | 0.047364517 | 1.361674144 | 1.279023188 | 1.723053191 | -0.115335088 | growth toward the midline |
| ENSDART00000135381 | six4a | 0.04980099 | 2.521510715 | 2.3671408 | 2.587075649 | 1.691172374 | growth toward the midline |
| ENSDART00000012791 | sp8a | 0.049626195 | 1.57251014 | 2.331511764 | 2.00847442 | 1.039704847 | growth toward the midline |
| ENSDART00000160644 | rela | 0.030637152 | 0.027046368 | 0.222531925 | 0.299056597 | -0.13648552 | growth toward the midline |
| ENSDART00000010248 | mitfb | 0.042577687 | 0.302243319 | 0.237275907 | 0.106756262 | -0.337810921 | growth toward the midline |
| ENSDART00000157487 | tfec | 0.001790465 | 0.534318298 | 0.423012313 | 0.205933849 | -0.313187686 | growth toward the midline |
| ENSDART00000164766 | tfec | 0.012357429 | 0.976052476 | 1.18639141 | 0.809154671 | -1.643283392 | growth toward the midline |
| ENSDART00000158466 | creb3l2 | 0.011352496 | 0.073267533 | 0.091531466 | -0.07087733 | -0.32121815 | growth toward the midline |
| ENSDART00000152378 | tgif1 | 0.003122803 | 0.194631245 | 0.419803811 | 0.14379228 | -0.205607529 | growth toward the midline |
| ENSDART00000062702 | cebpb | 0.018433591 | 0.973724375 | 0.549827472 | 0.384246832 | -0.627629236 | growth toward the midline |
| ENSDART00000056457 | mitfa | 0.005353377 | 0.929754118 | 0.38849893 | 0.30836176 | -0.311729197 | growth toward the midline |
| ENSDART00000080854 | stat3 | 0.03033326 | 0.856010749 | 0.444560766 | 0.379796349 | -0.326021696 | growth toward the midline |
| ENSDART00000093279 | spi1b | 0.002640644 | 0.891535687 | 0.563855895 | 0.100587929 | -0.531177734 | growth toward the midline |
| ENSDART00000036729 | spi1b | 0.004358158 | 0.910663058 | 0.717004711 | 0.095052493 | -0.117318513 | growth toward the midline |
| ENSDART00000008373 | fosl1a | 0.001576361 | 0.78271954 | 0.570182098 | 0.226810387 | -0.332448721 | growth toward the midline |
| ENSDART00000093199 | tead3b | 0.016523707 | 1.081516431 | 0.86043279 | 0.478204597 | -0.215799128 | growth toward the midline |
| ENSDART00000017185 | tbx20 | 0.035967167 | 0.559146809 | 0.370265655 | -0.151461394 | -0.656442573 | growth toward the midline |
| ENSDART00000052366 | cebpa | 0.035286858 | 0.847474055 | 0.814939712 | 0.318232429 | -0.173654256 | growth toward the midline |

| | | | | | | | |
|---|---|---|---|---|---|---|---|
| ENSDART00000005263 | usf1l | 0.019856195 | 0.061508869 | 0.080355717 | -0.137096705 | -0.208096843 | growth toward the midline |
| ENSDART00000062854 | nfe2l2a | 0.008134819 | 0.155734951 | 0.146836561 | -0.112595282 | -0.207726174 | growth toward the midline |
| ENSDART00000138081 | bcl6ab | 0.000589104 | 0.948060654 | 0.690527167 | 0.988530239 | 1.012261919 | midline crossing |
| ENSDART00000104317 | klf7b | 0.000976528 | 0.389436838 | 1.156291561 | 1.140404424 | 0.652703479 | midline crossing |
| ENSDART00000124040 | insm1a | 1.46E-05 | 0.50962443 | 1.551047216 | 1.738068624 | 1.104992333 | midline crossing |
| ENSDART00000048110 | six4b | 0.002795018 | 0.757934328 | 2.322346312 | 2.776043581 | 1.88012445 | midline crossing |
| ENSDART00000159652 | sox4b | 0.008897727 | 0.767109202 | 1.080655119 | 1.389017231 | 0.976257694 | midline crossing |
| ENSDART00000092690 | srebf2 | 0.003411856 | -0.007030288 | 0.441941306 | 0.773565631 | 0.436893785 | midline crossing |
| ENSDART00000033755 | homezb | 0.043076195 | -0.059847052 | 0.060728596 | 0.214721864 | 0.105513871 | midline crossing |
| ENSDART00000018945 | meis3 | 0.00431294 | -0.263611366 | 0.389783427 | 1.01804147 | 0.898863275 | midline crossing |
| ENSDART00000075331 | insm1b | 0.009480138 | 0.180818368 | 0.558805202 | 0.852549514 | 0.856981121 | midline crossing |
| ENSDART00000150128 | foxp1b | 0.020184087 | 0.347545443 | 0.556370195 | 1.342974258 | 1.269825931 | midline crossing |
| ENSDART00000128174 | foxp1b | 0.015925333 | 0.060807795 | 0.283454627 | 0.694455914 | 0.748409918 | midline crossing |
| ENSDART00000023959 | arntl1a | 7.44E-05 | 0.340904931 | 1.030627189 | 1.45255612 | 0.975945139 | midline crossing |
| ENSDART00000026992 | sox4a | 3.66E-05 | 0.168735758 | 0.553498244 | 0.871364123 | 0.630648218 | midline crossing |
| ENSDART00000083628 | ddit3 | 0.002192622 | -0.053032847 | 0.505000026 | 0.598711637 | 0.25299651 | midline crossing |
| ENSDART00000110512 | ybx1 | 0.014352201 | 0.187678935 | 0.306636132 | 0.543417204 | 0.157462635 | midline crossing |
| ENSDART00000141068 | sox11b | 8.49E-05 | 1.611838536 | 2.177304965 | 2.010624009 | 1.194765222 | midline crossing |
| ENSDART00000138821 | nfil3 | 0.000101984 | 1.376309909 | 2.121391468 | 1.99718084 | 1.39511595 | midline crossing |
| ENSDART00000082434 | tgif1 | 1.75E-05 | 0.822910382 | 1.229522702 | 1.441745222 | 0.912580325 | midline crossing |
| ENSDART00000139763 | stat5a | 0.018355029 | 3.442434617 | 3.964844196 | 3.417693165 | 3.322184683 | midline crossing |
| ENSDART00000055609 | atf4b | 0.027440234 | 0.138344185 | 0.51758353 | 0.353617358 | 0.087935611 | midline crossing |
| ENSDART00000152636 | clocka | 0.010610046 | 0.008631834 | 0.196030905 | 0.593132542 | 0.426491394 | target selection |
| ENSDART00000166976 | esrrb | 0.030820243 | -0.481593528 | -0.114040699 | 0.229959907 | 0.235353141 | target selection |
| ENSDART00000066317 | foxn2b | 0.033486585 | -0.194754205 | -0.013664444 | 0.221561118 | -0.080707177 | target selection |
| ENSDART00000104523 | arntl1b | 0.000255127 | -0.158144123 | 0.59291526 | 1.113237486 | 0.52944604 | target selection |
| ENSDART00000133959 | clockb | 0.010583972 | -0.261160651 | -0.146340415 | 0.224249915 | 0.15359596 | target selection |
| ENSDART00000047724 | zeb1b | 0.032175544 | -0.128023017 | -0.020882446 | 0.202453179 | 0.103960026 | target selection |
| ENSDART00000085728 | klf8 | 0.014069453 | -0.229536876 | -0.061888352 | 0.204863755 | 0.114771443 | target selection |
| ENSDART00000013605 | zbtb20 | 0.00075553 | 0.021191929 | 0.070670156 | 0.462529396 | 0.345025042 | target selection |
| ENSDART00000087450 | klf13 | 0.013792445 | -0.22166629 | -0.059571459 | 0.622443971 | 0.288396102 | target selection |
| ENSDART00000000280 | stat1b | 0.037541166 | -0.005066017 | 0.035352481 | 0.819259146 | 0.412054224 | target selection |
| ENSDART00000151575 | thraa | 0.013354221 | -0.36498113 | -0.584870078 | 0.329302923 | -0.150628947 | target selection |
| ENSDART00000154418 | relb | 0.010687727 | -0.043359819 | 0.083083005 | 0.26390054 | -0.071618881 | target selection |
| ENSDART00000123197 | relb | 0.019118248 | -0.261655378 | -0.003226565 | 0.486237661 | -0.265984214 | target selection |
| ENSDART00000152489 | irf3 | 0.006723268 | -0.152502705 | 0.050924712 | 1.421424509 | -0.375867622 | target selection |
| ENSDART00000141792 | stat1b | 0.033143681 | -1.394223287 | -0.705218411 | 2.324563925 | -0.328164239 | target selection |
| ENSDART00000103622 | irf7 | 0.002828188 | -0.049332142 | -0.2878598 | 1.533925917 | -0.275107455 | target selection |

| | | | | | | | |
|---|---|---|---|---|---|---|---|
| ENSDART00000148661 | stat2 | 0.006978095 | 0.11967996 | -0.016653911 | 1.354000143 | 0.023510931 | target selection |
| ENSDART00000005720 | stat1a | 0.010250585 | 0.035880276 | 0.045195542 | 0.615137639 | 0.020671748 | target selection |
| ENSDART00000156935 | pbx4 | 0.039411387 | 0.960146191 | 0.964981474 | 3.716226941 | 3.606625876 | target selection |
| ENSDART00000025575 | clocka | 0.041666185 | -0.204691736 | 0.089686311 | 0.498135587 | 0.316118853 | target selection |
| ENSDART00000125158 | znf574 | 0.029152083 | -0.158249462 | -0.223654122 | 0.188172427 | 0.054732788 | target selection |
| ENSDART00000092665 | srebf1 | 0.004619636 | -0.417658309 | -0.274360919 | 0.114636817 | 0.022214824 | target selection |
| ENSDART00000104751 | mxi1 | 0.002350825 | -0.384587031 | -0.210909301 | 0.22179917 | 0.104111801 | target selection |
| ENSDART00000111014 | myca | 0.024986086 | -0.198726525 | 0.073855205 | 0.316369866 | 0.312548093 | brain innervation |
| ENSDART00000083296 | ZNF423 | 0.00814358 | -0.278315489 | 0.051691519 | 0.407184973 | 0.489807123 | brain innervation |
| ENSDART00000171642 | dpf1 | 0.005353377 | -0.380689139 | 0.060927985 | 0.320062308 | 0.358114425 | brain innervation |
| ENSDART00000113087 | hmx1 | 0.000694227 | -0.389387565 | 0.021377927 | 0.234018679 | 0.270888749 | brain innervation |
| ENSDART00000162474 | irx1a | 0.002931603 | -1.132797992 | -0.349993381 | 0.581252213 | 0.55366573 | brain innervation |
| ENSDART00000162850 | irx3a | 0.000148815 | -0.694816328 | -0.301932125 | 0.187764734 | 0.208405543 | brain innervation |
| ENSDART00000053782 | scrt2 | 0.003310987 | -0.252422437 | -0.133353661 | 0.196438899 | 0.160308382 | brain innervation |
| ENSDART00000166834 | RFX7 (1 of many) | 0.009820514 | -0.017124727 | -0.055450521 | 0.18457609 | 0.267891323 | brain innervation |
| ENSDART00000111303 | rfx7 | 0.00161581 | 0.075232989 | 0.085195342 | 0.459783838 | 0.727409902 | brain innervation |
| ENSDART00000131126 | bcl6a | 0.005688069 | -0.096649799 | -0.044696983 | 0.295011601 | 0.436349263 | brain innervation |
| ENSDART00000065139 | gfi1ab | 0.023335678 | -0.252440054 | -0.123023724 | 0.269699656 | 0.68808671 | brain innervation |
| ENSDART00000012862 | isl2a | 3.29E-05 | -0.663813396 | 0.011664777 | 0.746201858 | 1.074694305 | brain innervation |
| ENSDART00000025031 | pou4f1 | 1.46E-05 | -0.37458842 | -0.169318799 | 0.774588614 | 1.160187758 | brain innervation |
| ENSDART00000016303 | irx2a | 4.96E-05 | -0.381386746 | 0.064410059 | 0.752093904 | 0.989751717 | brain innervation |
| ENSDART00000130163 | nr2f2 | 0.012656704 | -0.280168769 | -0.057880311 | 0.350863039 | 0.468160546 | brain innervation |
| ENSDART00000122101 | tbx2b | 0.001190205 | -0.056468629 | 0.017248499 | 0.19422425 | 0.382938949 | brain innervation |
| ENSDART00000150068 | sox6 | 0.001331938 | -0.111707858 | -0.047191704 | 0.355284385 | 0.700337568 | brain innervation |
| ENSDART00000062982 | foxo3b | 0.005646833 | -0.232285352 | -0.219543276 | -0.013673206 | 0.18814406 | brain innervation |
| ENSDART00000030773 | foxo3a | 0.000701449 | -0.522674186 | -0.346284342 | -0.025883071 | 0.086783781 | brain innervation |
| ENSDART00000100658 | esrra | 8.74E-05 | -0.571475474 | -0.395296451 | 0.169003635 | 0.212368184 | brain innervation |
| ENSDART00000108775 | ubp1 | 0.021499635 | -0.192682892 | -0.115240547 | 0.10453418 | 0.097860664 | brain innervation |
| ENSDART00000074833 | rx3 | 0.010379183 | -0.279918529 | -0.301637969 | 0.143576872 | 0.172520697 | brain innervation |
| ENSDART00000006737 | sox5 | 0.038386468 | -0.253791611 | -0.278700286 | 0.184042389 | 0.275951824 | brain innervation |
| ENSDART00000011890 | zbtb22a | 0.03843558 | -0.42030539 | -0.536466631 | 0.147549186 | 0.05725602 | brain innervation |
| ENSDART00000001795 | sp3a | 0.0407578 | -0.269877105 | -0.305437708 | -0.047306155 | -0.033496274 | brain innervation |
| ENSDART00000018603 | tbx4 | 0.04873911 | -0.302508569 | -0.188111077 | -0.131947864 | 0.000377602 | brain innervation |
| ENSDART00000003884 | mynn | 0.033340088 | -0.108806257 | -0.247003557 | 0.197551703 | 0.151198862 | brain innervation |
| ENSDART00000104327 | vsx1 | 0.00488592 | -0.248525466 | -0.619641572 | -0.034766195 | -0.028538746 | brain innervation |
| ENSDART00000165223 | pbx1b | 0.015188807 | -0.243991197 | -0.377373898 | -0.18693501 | 0.141565589 | brain innervation |
| ENSDART00000167440 | zfhx3 | 0.018342429 | -0.067144102 | -0.226395351 | 0.21751863 | 0.266797189 | brain innervation |
| ENSDART00000167538 | lcor | 0.031258003 | -0.151917658 | -0.14482463 | 0.130691961 | 0.223379898 | brain innervation |

| | | | | | | | |
|---|---|---|---|---|---|---|---|
| ENSDART00000013148 | pou3f1 | 0.00849815 | -0.360437586 | -0.322601739 | -0.060819622 | 0.211329038 | brain innervation |
| ENSDART00000109356 | klf7a | 0.000177237 | -0.7063233 | -0.659659661 | -0.044315321 | 0.157900055 | brain innervation |
| ENSDART00000138048 | pbx3b | 0.007841242 | -0.599655595 | -0.602159282 | -0.228775798 | -0.019503135 | brain innervation |
| ENSDART00000136245 | cux1a | 0.013366886 | -0.163227222 | -0.237382603 | -0.00302824 | 0.164419575 | brain innervation |
| ENSDART00000170631 | ebf1a | 0.016255169 | -0.245696473 | -0.326200552 | -0.132641619 | 0.115308885 | brain innervation |
| ENSDART00000064833 | mafaa | 0.002047801 | -0.957052225 | -1.143383392 | -0.379735148 | 0.067354966 | brain innervation |
| ENSDART00000004548 | barhl1b | 0.009605408 | -2.472825095 | -1.41656094 | -0.268019579 | 0.230183047 | brain innervation |
| ENSDART00000006380 | tbx3a | 0.004106215 | -0.244591708 | -0.352646993 | -0.144447405 | 0.06961498 | brain innervation |
| ENSDART00000151904 | foxn3 | 0.000466769 | -0.402174688 | -0.533042351 | -0.141780798 | 0.022724604 | brain innervation |
| ENSDART00000080808 | six3a | 0.000687981 | -0.437705937 | -0.430575558 | -0.140593988 | -0.07114268 | brain innervation |
| ENSDART00000171594 | mef2aa | 0.010231208 | -0.649844171 | -0.685014218 | -0.301258241 | -0.003741614 | brain innervation |
| ENSDART00000136729 | ebf1b | 0.021027344 | -2.79991744 | -3.068779235 | -0.720206417 | 0.313112936 | brain innervation |
| ENSDART00000162945 | sox5 | 0.023596757 | -0.177385379 | -0.174200373 | -0.003535976 | 0.152149827 | brain innervation |
| ENSDART00000132589 | mef2d | 0.028030725 | -0.20910625 | -0.209165039 | -0.020444617 | 0.107291243 | brain innervation |
| ENSDART00000165698 | pbx1a | 0.000272902 | -0.205212048 | -0.330154536 | 0.047550564 | 0.321002436 | brain innervation |
| ENSDART00000144894 | bcl11ba | 3.52E-05 | -0.656989822 | -0.679320488 | 0.181338486 | 0.623847694 | brain innervation |
| ENSDART00000006612 | tbr1b | 2.95E-05 | -1.166029945 | -1.008207372 | 0.617040877 | 1.169470893 | brain innervation |
| ENSDART00000154646 | bcl11ba | 0.001677265 | -0.533670184 | -0.516802708 | 0.152594691 | 0.570166001 | brain innervation |
| ENSDART00000160675 | dpf1 | 0.00210146 | -0.344267461 | -0.462495684 | 0.239034332 | 0.46811192 | brain innervation |
| ENSDART00000144186 | pou6f2 | 7.05E-05 | -0.474463335 | -0.465571046 | 0.092941552 | 0.436081081 | brain innervation |
| ENSDART00000124112 | pou4f2 | 2.95E-05 | -1.368026557 | -1.308261738 | 0.344262724 | 0.908586045 | brain innervation |
| ENSDART00000051693 | irx4a | 0.000591404 | -1.596922819 | -2.014349322 | 0.164610846 | 0.924604512 | brain innervation |
| ENSDART00000149175 | pou4f3 | 3.34E-05 | -1.520089487 | -1.231379123 | 0.357488642 | 0.743891028 | brain innervation |
| ENSDART00000055936 | isl2b | 4.92E-05 | -1.527471977 | -1.312125436 | 0.470803449 | 0.853726063 | brain innervation |
| ENSDART00000165609 | barhl2 | 0.000401616 | -0.31272019 | -0.374018735 | 0.126371762 | 0.224852149 | brain innervation |
| ENSDART00000010896 | isl1 | 0.002539705 | -0.281634356 | -0.275090348 | 0.147086866 | 0.205392134 | brain innervation |
| ENSDART00000100508 | ebf1b | 0.010170292 | -0.252122108 | -0.356560297 | -0.077233007 | 0.218128715 | brain innervation |
| ENSDART00000081170 | cux1a | 0.000900234 | -0.092953946 | -0.2283391 | 0.138990685 | 0.296890549 | brain innervation |
| ENSDART00000162485 | pax6a | 0.013416969 | -0.383059238 | -0.321515523 | -0.107334897 | 0.077147684 | brain innervation |
| ENSDART00000165774 | pax6a | 0.007156543 | -0.382942423 | -0.218376905 | 0.094924321 | 0.282463268 | brain innervation |
| ENSDART00000142087 | foxp2 | 0.000565675 | -0.522019738 | -0.4194261 | 0.065656049 | 0.27497631 | brain innervation |
| ENSDART00000048994 | pbx3b | 0.000910662 | -0.545498331 | -0.428509616 | 0.129533791 | 0.149586816 | brain innervation |
| ENSDART00000077157 | six3b | 0.008086296 | -0.273810969 | -0.17356253 | 0.033591817 | 0.098241195 | brain innervation |
| ENSDART00000040672 | mecp2 | 0.047533941 | -0.242382035 | -0.413395534 | -0.149398689 | -0.148226048 | downregulated during regeneration - early |
| ENSDART00000037879 | crx | 0.018920791 | -0.254014844 | -0.310425293 | -0.22793154 | -0.020194538 | downregulated during regeneration - early |
| ENSDART00000122429 | klf15 | 0.006222767 | -0.341190152 | -0.616050077 | -0.578895446 | -0.337948098 | downregulated during regeneration - early |
| ENSDART00000166313 | thrb | 0.0223553 | -0.193382381 | -0.415286369 | -0.400030228 | -0.136855946 | downregulated during regeneration - early |
| ENSDART00000089748 | rorb | 0.030623213 | -0.330969943 | -0.539847879 | -0.425468683 | -0.138099942 | downregulated during regeneration - early |

| | | | | | | | |
|---|---|---|---|---|---|---|---|
| ENSDART00000000876 | nr4a1 | 0.000894056 | -2.262372442 | -2.399082592 | -2.145415905 | -1.543723254 | downregulated during regeneration - early |
| ENSDART00000153187 | thrab | 0.001402519 | -0.362356661 | -0.551240408 | -0.403932333 | -0.192068079 | downregulated during regeneration - early |
| ENSDART00000054736 | bhlhe23 | 0.004928313 | -0.369485488 | -0.514619457 | -0.390416516 | -0.190127988 | downregulated during regeneration - early |
| ENSDART00000167052 | etv1 | 0.01072353 | -0.327639473 | -0.469002174 | -0.32895701 | -0.22464313 | downregulated during regeneration - early |
| ENSDART00000026017 | bhlhe40 | 0.046057741 | -0.186426622 | -0.329757373 | -0.286559283 | -0.071751826 | downregulated during regeneration - early |
| ENSDART00000131134 | si:ch211-206a7.2 | 0.008848885 | -0.226522125 | -0.472315068 | -0.371394997 | -0.101330709 | downregulated during regeneration - early |
| ENSDART00000024832 | stat5a | 0.004077075 | -0.212236095 | -0.408877789 | -0.375060924 | -0.185569558 | downregulated during regeneration - early |
| ENSDART00000091707 | dbpa | 0.017575469 | -0.158262702 | -0.327851995 | -0.357551076 | -0.129469939 | downregulated during regeneration - early |
| ENSDART00000126966 | esrrga | 0.017679749 | -0.248566713 | -0.265310961 | -0.321308408 | -0.070523791 | downregulated during regeneration - early |
| ENSDART00000127353 | nr1d2b | 0.027085654 | -0.29103008 | -0.243688493 | -0.346507775 | -0.100908835 | downregulated during regeneration - early |
| ENSDART00000047541 | bach1b | 0.039802589 | -0.343772395 | -0.510091534 | -0.468889885 | -0.333515696 | downregulated during regeneration - early |
| ENSDART00000007630 | nhlh2 | 0.010056242 | -0.28794611 | -0.324673306 | -0.385652145 | -0.261092419 | downregulated during regeneration - early |
| ENSDART00000129498 | mef2d | 0.03679078 | -0.188451231 | -0.381445449 | -0.263610552 | -0.197973462 | downregulated during regeneration - early |
| ENSDART00000164349 | e2f4 | 0.042377891 | -0.246759183 | -0.226331986 | -0.10688884 | -0.101679787 | downregulated during regeneration - early |
| ENSDART00000157890 | tcf7l1b | 0.035686017 | -4.555123239 | -4.673271453 | -1.829433189 | -0.732249154 | downregulated during regeneration - early |
| ENSDART00000165757 | pax6b | 0.019981901 | -1.030621281 | -0.736504391 | -0.510180699 | -0.112500923 | downregulated during regeneration - early |
| ENSDART00000169609 | tefb | 0.032368681 | -0.156114581 | -0.555141888 | -0.301378906 | -0.24132076 | downregulated during regeneration - early |
| ENSDART00000103640 | hey1 | 0.023203992 | -0.385659584 | -0.747117397 | -0.392306381 | -0.416692063 | downregulated during regeneration - early |
| ENSDART00000013003 | tfap2b | 0.008192536 | -0.299930992 | -0.578681033 | -0.25971747 | -0.279510711 | downregulated during regeneration - early |
| ENSDART00000082346 | tfap2a | 0.038054224 | -0.238141907 | -0.307957291 | -0.242159487 | -0.230339237 | downregulated during regeneration - early |
| ENSDART00000147658 | bhlhe22 | 0.01205445 | -0.302235414 | -0.31899187 | -0.305987775 | -0.102192149 | downregulated during regeneration - early |
| ENSDART00000057644 | lhx4 | 0.048389146 | -0.267668007 | -0.268543602 | -0.190125613 | -0.119751289 | downregulated during regeneration - early |
| ENSDART00000009938 | tcf12 | 0.025246927 | -0.385864267 | -0.351608927 | -0.283149928 | -0.323464659 | downregulated during regeneration - early |
| ENSDART00000065361 | etv5b | 0.019698661 | -0.645686026 | -0.347502384 | -0.293939032 | -0.317696675 | downregulated during regeneration - early |
| ENSDART00000055709 | her2 | 0.030146589 | -0.984524994 | -0.833253569 | -0.717745512 | -1.233522642 | downregulated during regeneration - early |
| ENSDART00000055706 | her15.1 | 0.021144178 | -1.483608056 | -1.154441068 | -0.698110478 | -1.436427969 | downregulated during regeneration - early |
| ENSDART00000133487 | fosb | 0.034512759 | -0.882572207 | -1.273781011 | -0.772506712 | -0.914864674 | downregulated during regeneration - early |
| ENSDART00000151127 | thraa | 0.039890399 | -0.798583389 | -0.802364861 | -0.351487338 | -0.589241182 | downregulated during regeneration - early |
| ENSDART00000153167 | hlfb | 0.010966965 | -0.470839666 | -0.326611575 | -0.235752692 | -0.264047019 | downregulated during regeneration - early |
| ENSDART00000086051 | mecom | 0.04049764 | -0.550304416 | -0.298852875 | -0.293768303 | -1.334615988 | downregulated during regeneration - early |
| ENSDART00000023613 | her6 | 0.030622517 | -0.335042933 | -0.234567638 | -0.250120727 | -0.59712361 | downregulated during regeneration - early |
| ENSDART00000125174 | nr1i2 | 0.028902271 | -0.056601149 | -0.085618729 | -0.264588441 | -0.326132843 | downregulated during regeneration - late |
| ENSDART00000089015 | zbtb7a | 0.00191624 | -0.478666207 | -0.203919761 | -0.738823143 | -0.713706626 | downregulated during regeneration - late |
| ENSDART00000044860 | maff | 0.002310171 | 0.057535561 | 0.039208376 | -0.366680053 | -0.441007369 | downregulated during regeneration - late |
| ENSDART00000167844 | mafk | 0.014095383 | -0.00863672 | 0.25450476 | -0.276014619 | -0.283559487 | downregulated during regeneration - late |
| ENSDART00000057124 | tefa | 0.007560757 | -0.17959692 | 0.074135039 | -0.330628318 | -0.262330789 | downregulated during regeneration - late |
| ENSDART00000057125 | tefa | 0.002939873 | -0.190705988 | 0.094171259 | -0.306461854 | -0.301060608 | downregulated during regeneration - late |
| ENSDART00000163250 | mef2cb | 0.003671994 | 0.361224616 | -0.300957895 | -0.57227951 | -0.080214844 | downregulated during regeneration - late |

| | | | | | | | |
|---|---|---|---|---|---|---|---|
| ENSDART00000131731 | mef2ca | 0.006008543 | 0.234535922 | -0.250149626 | -0.381098175 | -0.047609857 | downregulated during regeneration - late |
| ENSDART00000161387 | tcf12 | 0.008531138 | 0.370378706 | 0.009395421 | -0.374363269 | -0.152945712 | downregulated during regeneration - late |
| ENSDART00000127099 | nr2e3 | 0.030676359 | 0.281603482 | 0.002911001 | -0.560613973 | -0.033774044 | downregulated during regeneration - late |
| ENSDART00000066655 | mybl1 | 0.005666275 | -0.255810306 | -0.487313899 | -0.72749049 | -0.668391364 | downregulated during regeneration - late |
| ENSDART00000172251 | creb3l1 | 0.029283678 | -0.33540662 | 0.193573394 | -4.078076569 | -0.842213331 | downregulated during regeneration - late |
| ENSDART00000170575 | nfat5b | 0.018611393 | -0.001493923 | -0.392122738 | -0.566909919 | -0.288747412 | downregulated during regeneration - late |
| ENSDART00000148106 | mef2aa | 0.026582721 | -0.081330082 | -0.55524128 | -0.609035118 | -0.185252531 | downregulated during regeneration - late |
| ENSDART00000075070 | hsf2 | 0.014034602 | -0.066635095 | -0.276496416 | -0.474097609 | -0.186498896 | downregulated during regeneration - late |
| ENSDART00000148353 | usf2 | 0.006964046 | -0.107987138 | -0.265954127 | -0.405201274 | -0.200197809 | downregulated during regeneration - late |
| ENSDART00000061106 | bhlhe41 | 0.000553147 | 0.043056173 | -0.423617996 | -0.754959095 | -0.246095493 | downregulated during regeneration - late |
| ENSDART00000139102 | dbpb | 0.001993937 | 0.113164671 | -0.364559303 | -0.423196921 | -0.137052116 | downregulated during regeneration - late |
| ENSDART00000077839 | atf7b | 0.014656899 | -0.232485852 | -0.196508033 | -0.391624214 | -0.206346003 | downregulated during regeneration - late |
| ENSDART00000045374 | smad3a | 0.043076195 | -0.063329624 | -0.157712113 | -0.303211349 | -0.108940629 | downregulated during regeneration - late |
| ENSDART00000093331 | rreb1a | 0.018126285 | -0.024901133 | -0.155149005 | -0.517234878 | -0.050769944 | downregulated during regeneration - late |
| ENSDART00000148537 | roraa | 0.01612441 | -0.160828233 | -0.221872095 | -0.45696721 | -0.083100906 | downregulated during regeneration - late |
| ENSDART00000123970 | mntb | 0.011725523 | -0.115861458 | -0.101461538 | -0.360694649 | -0.01567692 | downregulated during regeneration - late |
| ENSDART00000127157 | hlfa | 0.048701514 | -0.089881453 | -0.212609643 | -0.346930267 | -0.147102401 | downregulated during regeneration - late |
| ENSDART00000126282 | nr1d1 | 0.008088213 | -0.1014969 | -0.191656676 | -0.494430716 | -0.13667305 | downregulated during regeneration - late |

**Table S7. Transcription factors that are differentially expressed during regeneration.**

| ENSEMBL ID | zebrafish gene symbol | cluster order | HGNC symbol | motif status | DNA binding domain | #motifs |
|---|---|---|---|---|---|---|
| ENSDART00000128815 | stra13 | 1 | CENPX | n | Unknown | 0 |
| ENSDART00000098285 | atf5a | 1 | ATF5 | n | bZIP | 0 |
| ENSDART00000062576 | thyn1 | 1 | THYN1 | n | Unknown | 0 |
| ENSDART00000013409 | prmt3 | 1 | PRMT3 | n | C2H2 ZF | 0 |
| ENSDART00000058277 | znf800b | 1 | ZNF800 | n | C2H2 ZF | 0 |
| ENSDART00000103982 | nfe2l1a | 1 | NFE2L1 | n | bZIP | 0 |
| ENSDART00000081468 | ccdc79 | 1 | TERB1 | n | bZIP | 0 |
| ENSDART00000155066 | atf5b | 1 | ATF5 | n | bZIP | 0 |
| ENSDART00000114288 | foxp4 | 1 | FOXP4 | n | Forkhead | 0 |
| ENSDART00000086053 | PRDM16 | 1 | PRDM16 | n | C2H2 ZF | 0 |
| ENSDART00000140760 | e2f7 | 1 | E2F7 | y | E2F | 1 |
| ENSDART00000056005 | ascl1a | 1 | ASCL1 | y | bHLH | 1 |
| ENSDART00000128488 | e2f8 | 1 | E2F8 | y | E2F | 1 |
| ENSDART00000132119 | max | 1 | MAX | y | bHLH | 4 |
| ENSDART00000163794 | wt1b | 1 | WT1 | y | C2H2 ZF | 8 |
| ENSDART00000033494 | klf6a | 1 | KLF6 | y | C2H2 ZF | 39 |
| ENSDART00000147849 | klf6a | 1 | KLF6 | y | C2H2 ZF | 39 |
| ENSDART00000076161 | hoxb5b | 1 | HOXB5 | y | Homeodomain | 1 |
| ENSDART00000122628 | junba | 1 | JUNB | y | bZIP | 10 |
| ENSDART00000146767 | fosl1a | 1 | FOSL1 | y | bZIP | 6 |
| ENSDART00000161610 | tcf3b | 1 | TCF3 | y | bHLH | 3 |
| ENSDART00000051549 | tp53 | 1 | TP53 | y | p53 | 3 |
| ENSDART00000164711 | NFATC2 | 1 | NFATC2 | y | Rel | 1 |
| ENSDART00000063912 | jun | 1 | JUN | y | bZIP | 12 |
| ENSDART00000022060 | atf3 | 1 | ATF3 | y | bZIP | 1 |
| ENSDART00000110040 | sox11a | 1 | SOX11 | y | HMG/Sox | 1 |
| ENSDART00000104519 | stat3 | 1 | STAT3 | y | STAT | 2 |
| ENSDART00000142584 | alx1 | 1 | ALX1 | y | Homeodomain | 1 |
| ENSDART00000135381 | six4a | 1 | SIX4 | y | Homeodomain | 24 |
| ENSDART00000012791 | sp8a | 1 | SP8 | y | C2H2 ZF | 1 |
| ENSDART00000160644 | rela | 1 | RELA | y | Rel | 1 |
| ENSDART00000010248 | mitfb | 1 | MITF | y | bHLH | 2 |
| ENSDART00000157487 | tfec | 1 | TFEC | y | bHLH | 1 |
| ENSDART00000164766 | tfec | 1 | TFEC | y | bHLH | 1 |
| ENSDART00000158466 | creb3l2 | 1 | CREB3L2 | y | bZIP | 1 |
| ENSDART00000152378 | tgif1 | 1 | TGIF1 | y | Homeodomain | 1 |
| ENSDART00000062702 | cebpb | 1 | CEBPB | y | bZIP | 2 |
| ENSDART00000056457 | mitfa | 1 | MITF | y | bHLH | 2 |
| ENSDART00000080854 | stat3 | 1 | STAT3 | y | STAT | 2 |
| ENSDART00000093279 | spi1b | 1 | SPIB | y | Ets | 1 |
| ENSDART00000036729 | spi1b | 1 | SPIB | y | Ets | 1 |
| ENSDART00000008373 | fosl1a | 1 | FOSL1 | y | bZIP | 6 |
| ENSDART00000093199 | tead3b | 1 | TEAD3 | y | TEA | 1 |
| ENSDART00000017185 | tbx20 | 1 | TBX20 | y | T-box | 1 |
| ENSDART00000052366 | cebpa | 1 | CEBPA | y | bZIP | 4 |
| ENSDART00000005263 | usf1l | 1 | USF1 | y | bHLH | 2 |
| ENSDART00000062854 | nfe2l2a | 1 | NFE2L2 | y | bZIP | 2 |
| ENSDART00000016135 | nfe2l3 | 2 | NFE2L3 | n | bZIP | 0 |
| ENSDART00000123263 | phf20a | 2 | PHF20 | n | AT hook | 0 |
| ENSDART00000146180 | csrnp2 | 2 | CSRNP2 | n | Unknown | 0 |
| ENSDART00000013961 | mycla | 2 | MYCL | n | bHLH | 0 |
| ENSDART00000018150 | neurod6b | 2 | NEUROD6 | n | bHLH | 0 |
| ENSDART00000165656 | mxd3 | 2 | MXD3 | n | bHLH | 0 |
| ENSDART00000114473 | trafd1 | 2 | TRAFD1 | n | C2H2 ZF | 0 |
| ENSDART00000158598 | si:ch211-232b12.5 | 2 | ZHX3 | n | #N/A | 0 |
| ENSDART00000151044 | baz2ba | 2 | BAZ2B | n | MBD | 0 |
| ENSDART00000141941 | zfpm2a | 2 | ZFPM2 | n | C2H2 ZF | 0 |
| ENSDART00000109552 | baz2ba | 2 | BAZ2B | n | MBD | 0 |
| ENSDART00000029843 | vezf1a | 2 | VEZF1 | n | C2H2 ZF | 0 |
| ENSDART00000041877 | csrnp1a | 2 | CSRNP1 | n | Unknown | 0 |
| ENSDART00000170865 | nme2b.1 | 2 | NME2 | n | Unknown | 0 |
| ENSDART00000018743 | phf20a | 2 | PHF20 | n | AT hook | 0 |
| ENSDART00000099849 | arntl2 | 2 | ARNTL2 | n | bHLH | 0 |
| ENSDART00000149074 | sall1b | 2 | SALL1 | n | C2H2 ZF | 0 |
| ENSDART00000153227 | nfe2l1b | 2 | NFE2L1 | n | bZIP | 0 |
| ENSDART00000024872 | creb3l3l | 2 | CREB3L3 | n | bZIP | 0 |
| ENSDART00000123559 | znf521 | 2 | ZNF521 | n | C2H2 ZF | 0 |
| ENSDART00000077484 | zhx2a | 2 | ZHX2 | n | Homeodomain | 0 |
| ENSDART00000138081 | bcl6ab | 2 | BCL6 | y | C2H2 ZF | 1 |
| ENSDART00000104317 | klf7b | 2 | KLF7 | y | C2H2 ZF | 38 |
| ENSDART00000124040 | insm1a | 2 | INSM1 | y | C2H2 ZF | 1 |
| ENSDART00000048110 | six4b | 2 | SIX4 | y | Homeodomain | 24 |
| ENSDART00000159652 | sox4b | 2 | SOX4 | y | HMG/Sox | 1 |
| ENSDART00000092690 | srebf2 | 2 | SREBF2 | y | bHLH | 1 |
| ENSDART00000033755 | homezb | 2 | HOMEZ | y | Homeodomain | 1 |
| ENSDART00000018945 | meis3 | 2 | MEIS3 | y | Homeodomain | 1 |
| ENSDART00000075331 | insm1b | 2 | INSM1 | y | C2H2 ZF | 1 |
| ENSDART00000150128 | foxp1b | 2 | FOXP1 | y | Forkhead | 2 |
| ENSDART00000128174 | foxp1b | 2 | FOXP1 | y | Forkhead | 2 |
| ENSDART00000023959 | arntl1a | 2 | ARNTL | y | bHLH | 1 |
| ENSDART00000026992 | sox4a | 2 | SOX4 | y | HMG/Sox | 1 |
| ENSDART00000083628 | ddit3 | 2 | DDIT3 | y | bZIP | 1 |
| ENSDART00000110512 | ybx1 | 2 | YBX1 | y | CSD | 4 |
| ENSDART00000141068 | sox11b | 2 | SOX11 | y | HMG/Sox | 1 |
| ENSDART00000138821 | nfil3 | 2 | NFIL3 | y | bZIP | 1 |
| ENSDART00000082434 | tgif1 | 2 | TGIF1 | y | Homeodomain | 1 |
| ENSDART00000139763 | stat5a | 2 | STAT5A | y | STAT | 1 |
| ENSDART00000055609 | atf4b | 2 | ATF4 | y | bZIP | 1 |
| ENSDART00000172990 | znf628 | 3 | ZNF628 | n | C2H2 ZF | 0 |
| ENSDART00000112756 | ZNF608 | 3 | ZNF608 | n | C2H2 ZF | 0 |

| DE transcript cluster | DETF cluster order | total TFs | TFs with motifs |
|---|---|---|---|
| 1clust5 | 1 | 40 | 31 |
| 2clust4 | 1,2 | 76 | 48 |
| 3clust2 | 3 | 41 | 18 |
| 4clust6 | 4 | 75 | 51 |
| 5clust7 | 5 | 62 | 34 |
| 6clust3 | 6 | 41 | 23 |
| 7clust1 | 6 | 41 | 23 |

| | | | | | | |
|---|---|---|---|---|---|---|
| ENSDART00000090226 | znf319b | 3 | ZNF319 | n | C2H2 ZF | 0 |
| ENSDART00000098859 | neurod6a | 3 | NEUROD6 | n | bHLH | 0 |
| ENSDART00000031167 | tfap2d | 3 | TFAP2D | n | AP-2 | 0 |
| ENSDART00000166502 | satb2 | 3 | SATB2 | n | CUT; Homeodomain | 0 |
| ENSDART00000167324 | ebf3a | 3 | EBF3 | n | EBF1 | 0 |
| ENSDART00000157855 | ebf3a | 3 | EBF3 | n | EBF1 | 0 |
| ENSDART00000158634 | ebf3a | 3 | EBF3 | n | EBF1 | 0 |
| ENSDART00000128504 | si:dkey-106g10.7 | 3 | C11orf95 | n | #N/A | 0 |
| ENSDART00000109493 | tshz2 | 3 | TSHZ2 | n | C2H2 ZF | 0 |
| ENSDART00000058685 | zfpm2a | 3 | ZFPM2 | n | C2H2 ZF | 0 |
| ENSDART00000110069 | CABZ01075131.1 | 3 | CSRNP3 | n | Unknown | 0 |
| ENSDART00000166565 | tshz3b | 3 | TSHZ3 | n | C2H2 ZF | 0 |
| ENSDART00000123147 | zbtb8a | 3 | ZBTB8A | n | C2H2 ZF | 0 |
| ENSDART00000084819 | arhgap35b | 3 | ARHGAP35 | n | Unknown | 0 |
| ENSDART00000104712 | hmg20a | 3 | HMG20A | n | HMG/Sox | 0 |
| ENSDART00000135647 | si:ch211-113e8.10 | 3 | ZFP91 | n | C2H2 ZF | 0 |
| ENSDART00000165877 | purg | 3 | PURG | n | Unknown | 0 |
| ENSDART00000089076 | dot1l | 3 | DOT1L | n | AT hook | 0 |
| ENSDART00000140284 | zbtb16a | 3 | ZBTB16 | n | C2H2 ZF | 0 |
| ENSDART00000165955 | zhx3 | 3 | ZHX3 | n | #N/A | 0 |
| ENSDART00000037516 | znf827 | 3 | ZNF827 | n | C2H2 ZF | 0 |
| ENSDART00000060193 | thap3 | 3 | THAP3 | n | THAP finger | 0 |
| ENSDART00000154730 | tsc22d1 | 3 | TSC22D1 | n | Unknown | 0 |
| ENSDART00000152636 | clocka | 3 | CLOCK | y | bHLH | 1 |
| ENSDART00000166976 | esrrb | 3 | ESRRB | y | Nuclear receptor | 3 |
| ENSDART00000066317 | foxn2b | 3 | FOXN2 | y | Forkhead | 11 |
| ENSDART00000104523 | arntl1b | 3 | ARNTL | y | bHLH | 1 |
| ENSDART00000133959 | clockb | 3 | CLOCK | y | bHLH | 1 |
| ENSDART00000047724 | zeb1b | 3 | ZEB1 | y | C2H2 ZF; Homeodomain | 3 |
| ENSDART00000085728 | klf8 | 3 | KLF8 | y | C2H2 ZF | 38 |
| ENSDART00000013605 | zbtb20 | 3 | ZBTB20 | y | C2H2 ZF | 2 |
| ENSDART00000087450 | klf13 | 3 | KLF13 | y | C2H2 ZF | 1 |
| ENSDART00000000280 | stat1b | 3 | STAT1 | y | STAT | 4 |
| ENSDART00000151575 | thraa | 3 | THRA | y | Nuclear receptor | 15 |
| ENSDART00000154418 | relb | 3 | RELB | y | Rel | 1 |
| ENSDART00000123197 | relb | 3 | RELB | y | Rel | 1 |
| ENSDART00000152489 | irf3 | 3 | IRF3 | y | IRF | 1 |
| ENSDART00000141792 | stat1b | 3 | STAT1 | y | STAT | 4 |
| ENSDART00000103622 | irf7 | 3 | IRF7 | y | IRF | 1 |
| ENSDART00000148661 | stat2 | 3 | STAT2 | y | STAT | 1 |
| ENSDART00000005720 | stat1a | 3 | STAT1 | y | STAT | 4 |
| ENSDART00000156935 | pbx4 | 3 | PBX4 | y | Homeodomain | 15 |
| ENSDART00000025575 | clocka | 3 | CLOCK | y | bHLH | 1 |
| ENSDART00000125158 | znf574 | 3 | ZNF574 | y | C2H2 ZF | 2 |
| ENSDART00000092665 | srebf1 | 3 | SREBF1 | y | bHLH | 1 |
| ENSDART00000104751 | mxi1 | 3 | MXI1 | y | bHLH | 1 |
| ENSDART00000101316 | znf618 | 4 | ZNF618 | n | C2H2 ZF | 0 |
| ENSDART00000112529 | znf319a | 4 | ZNF319 | n | C2H2 ZF | 0 |
| ENSDART00000172410 | jazf1a | 4 | JAZF1 | n | C2H2 ZF | 0 |
| ENSDART00000089923 | znf652 | 4 | ZNF652 | n | C2H2 ZF | 0 |
| ENSDART00000101070 | dachd | 4 | DACH2 | n | Unknown | 0 |
| ENSDART00000114246 | adnpb | 4 | ADNP2 | n | Homeodomain | 0 |
| ENSDART00000023038 | dacha | 4 | DACH2 | n | Unknown | 0 |
| ENSDART00000045628 | irx6a | 4 | IRX6 | n | Homeodomain | 0 |
| ENSDART00000145072 | neurod4 | 4 | NEUROD4 | n | bHLH | 0 |
| ENSDART00000109347 | nacc2 | 4 | NACC2 | n | Unknown | 0 |
| ENSDART00000008906 | znf503 | 4 | ZNF503 | n | C2H2 ZF | 0 |
| ENSDART00000155866 | zbtb16b | 4 | ZBTB16 | n | C2H2 ZF | 0 |
| ENSDART00000164488 | mbnl2 | 4 | MBNL2 | n | CCCH ZF | 0 |
| ENSDART00000160784 | satb1b | 4 | SATB1 | n | CUT; Homeodomain | 0 |
| ENSDART00000101631 | satb1b | 4 | SATB1 | n | CUT; Homeodomain | 0 |
| ENSDART00000137355 | tet3 | 4 | TET3 | n | CxxC | 0 |
| ENSDART00000007806 | zbtb16a | 4 | ZBTB16 | n | C2H2 ZF | 0 |
| ENSDART00000046663 | camta1b | 4 | CAMTA1 | n | CG-1 | 0 |
| ENSDART00000104790 | znf292b | 4 | ZNF292 | n | C2H2 ZF | 0 |
| ENSDART00000084035 | znf532 | 4 | ZNF532 | n | C2H2 ZF | 0 |
| ENSDART00000134064 | mxd1 | 4 | MXD1 | n | bHLH | 0 |
| ENSDART00000136759 | prdm13 | 4 | PRDM13 | n | C2H2 ZF | 0 |
| ENSDART00000156464 | prr12b | 4 | PRR12 | n | AT hook | 0 |
| ENSDART00000153150 | myt1la | 4 | MYT1L | n | C2H2 ZF | 0 |
| ENSDART00000171091 | zeb2b | 4 | ZEB2 | n | C2H2 ZF; Homeodomain | 0 |
| ENSDART00000131714 | mbnl2 | 4 | MBNL2 | n | CCCH ZF | 0 |
| ENSDART00000124534 | mbnl2 | 4 | MBNL2 | n | CCCH ZF | 0 |
| ENSDART00000164643 | ZNF609 (1 of many) | 4 | ZNF609 | n | C2H2 ZF | 0 |
| ENSDART00000155666 | znf576.1 | 4 | ZNF576 | n | C2H2 ZF | 0 |
| ENSDART00000111014 | myca | 4 | MYC | y | bHLH | 4 |
| ENSDART00000083296 | ZNF423 | 4 | ZNF423 | y | C2H2 ZF | 1 |
| ENSDART00000171642 | dpf1 | 4 | DPF1 | y | C2H2 ZF | 2 |
| ENSDART00000113087 | hmx1 | 4 | HMX1 | y | Homeodomain | 1 |
| ENSDART00000162474 | irx1a | 4 | IRX1 | y | Homeodomain | 17 |
| ENSDART00000162850 | irx3a | 4 | IRX3 | y | Homeodomain | 17 |
| ENSDART00000053782 | scrt2 | 4 | SCRT2 | y | C2H2 ZF | 1 |
| ENSDART00000166834 | RFX7 (1 of many) | 4 | RFX7 | y | RFX | 14 |
| ENSDART00000111303 | rfx7 | 4 | RFX7 | y | RFX | 14 |
| ENSDART00000131126 | bcl6a | 4 | BCL6 | y | C2H2 ZF | 1 |
| ENSDART00000065139 | gfi1ab | 4 | GFI1 | y | C2H2 ZF | 1 |
| ENSDART00000012862 | isl2a | 4 | ISL2 | y | Homeodomain | 1 |
| ENSDART00000025031 | pou4f1 | 4 | POU4F1 | y | Homeodomain; POU | 1 |
| ENSDART00000016303 | irx2a | 4 | IRX2 | y | Homeodomain | 18 |
| ENSDART00000130163 | nr2f2 | 4 | NR2F2 | y | Nuclear receptor | 1 |
| ENSDART00000122101 | tbx2b | 4 | TBX2 | y | T-box | 1 |
| ENSDART00000150068 | sox6 | 4 | SOX6 | y | HMG/Sox | 1 |
| ENSDART00000062982 | foxo3b | 4 | FOXO3 | y | Forkhead | 2 |

| | | | | | | |
|---|---|---|---|---|---|---|
| ENSDART00000030773 | foxo3a | 4 | FOXO3 | y | Forkhead | 2 |
| ENSDART00000100658 | esrra | 4 | ESRRA | y | Nuclear receptor | 2 |
| ENSDART00000108775 | ubp1 | 4 | UBP1 | y | Grainyhead | 8 |
| ENSDART00000074833 | rx3 | 4 | RAX | y | Homeodomain | 1 |
| ENSDART00000006737 | sox5 | 4 | SOX5 | y | HMG/Sox | 1 |
| ENSDART00000011890 | zbtb22a | 4 | ZBTB22 | y | C2H2 ZF | 4 |
| ENSDART00000001795 | sp3a | 4 | SP3 | y | C2H2 ZF | 1 |
| ENSDART00000018603 | tbx4 | 4 | TBX4 | y | T-box | 1 |
| ENSDART00000003884 | mynn | 4 | MYNN | y | C2H2 ZF | 1 |
| ENSDART00000104327 | vsx1 | 4 | VSX1 | y | Homeodomain | 1 |
| ENSDART00000165223 | pbx1b | 4 | PBX1 | y | Homeodomain | 1 |
| ENSDART00000167440 | zfhx3 | 4 | ZFHX3 | y | C2H2 ZF; Homeodomain | 2 |
| ENSDART00000167538 | lcor | 4 | LCOR | y | Pipsqueak | 4 |
| ENSDART00000013148 | pou3f1 | 4 | POU3F1 | y | Homeodomain; POU | 1 |
| ENSDART00000109356 | klf7a | 4 | KLF7 | y | C2H2 ZF | 38 |
| ENSDART00000138048 | pbx3b | 4 | PBX3 | y | Homeodomain | 1 |
| ENSDART00000136245 | cux1a | 4 | CUX1 | y | CUT; Homeodomain | 1 |
| ENSDART00000170631 | ebf1a | 4 | EBF1 | y | EBF1 | 3 |
| ENSDART00000064833 | mafaa | 4 | MAFA | y | bZIP | 25 |
| ENSDART00000004548 | barhl1b | 4 | BARHL1 | y | Homeodomain | 1 |
| ENSDART00000006380 | tbx3a | 4 | TBX3 | y | T-box | 14 |
| ENSDART00000151904 | foxn3 | 4 | FOXN3 | y | Forkhead | 11 |
| ENSDART00000080808 | six3a | 4 | SIX3 | y | Homeodomain | 1 |
| ENSDART00000171594 | mef2aa | 4 | MEF2A | y | MADS box | 3 |
| ENSDART00000136729 | ebf1b | 4 | EBF1 | y | EBF1 | 3 |
| ENSDART00000162945 | sox5 | 4 | SOX5 | y | HMG/Sox | 1 |
| ENSDART00000132589 | mef2d | 4 | MEF2D | y | MADS box | 1 |
| ENSDART00000165698 | pbx1a | 4 | PBX1 | y | Homeodomain | 1 |
| ENSDART00000144894 | bcl11ba | 4 | BCL11B | y | C2H2 ZF | 3 |
| ENSDART00000006612 | tbr1b | 4 | TBR1 | y | T-box | 1 |
| ENSDART00000154646 | bcl11ba | 4 | BCL11B | y | C2H2 ZF | 3 |
| ENSDART00000160675 | dpf1 | 4 | DPF1 | y | C2H2 ZF | 2 |
| ENSDART00000144186 | pou6f2 | 4 | POU6F2 | y | Homeodomain; POU | 1 |
| ENSDART00000124112 | pou4f2 | 4 | POU4F2 | y | Homeodomain; POU | 1 |
| ENSDART00000051693 | irx4a | 4 | IRX4 | y | Homeodomain | 17 |
| ENSDART00000149175 | pou4f3 | 4 | POU4F3 | y | Homeodomain; POU | 1 |
| ENSDART00000055936 | isl2b | 4 | ISL2 | y | Homeodomain | 1 |
| ENSDART00000165609 | barhl2 | 4 | BARHL2 | y | Homeodomain | 1 |
| ENSDART00000010896 | isl1 | 4 | ISL1 | y | Homeodomain | 10 |
| ENSDART00000100508 | ebf1b | 4 | EBF1 | y | EBF1 | 3 |
| ENSDART00000081170 | cux1a | 4 | CUX1 | y | CUT; Homeodomain | 1 |
| ENSDART00000162485 | pax6a | 4 | PAX6 | y | Homeodomain; Paired box | 1 |
| ENSDART00000165774 | pax6a | 4 | PAX6 | y | Homeodomain; Paired box | 1 |
| ENSDART00000142087 | foxp2 | 4 | FOXP2 | y | Forkhead | 1 |
| ENSDART00000048994 | pbx3b | 4 | PBX3 | y | Homeodomain | 1 |
| ENSDART00000077157 | six3b | 4 | SIX3 | y | Homeodomain | 1 |
| ENSDART00000022010 | hivep2b | 5 | HIVEP2 | n | C2H2 ZF | 0 |
| ENSDART00000136488 | zmat4a | 5 | ZMAT4 | n | C2H2 ZF | 0 |
| ENSDART00000151571 | ahdc1 | 5 | AHDC1 | n | AT hook | 0 |
| ENSDART00000059446 | znf385b | 5 | ZNF385B | n | C2H2 ZF | 0 |
| ENSDART00000165120 | purab | 5 | PURA | n | Unknown | 0 |
| ENSDART00000172199 | zmat4a | 5 | ZMAT4 | n | C2H2 ZF | 0 |
| ENSDART00000166540 | kcnip3b | 5 | KCNIP3 | n | Unknown | 0 |
| ENSDART00000111842 | sall1a | 5 | SALL1 | n | C2H2 ZF | 0 |
| ENSDART00000150949 | nfyc | 5 | NFYC | n | Unknown | 0 |
| ENSDART00000166351 | nkrf | 5 | NKRF | n | Unknown | 0 |
| ENSDART00000090019 | zeb2b | 5 | ZEB2 | n | C2H2 ZF; Homeodomain | 0 |
| ENSDART00000009691 | scml4 | 5 | SCML4 | n | AT hook | 0 |
| ENSDART00000170400 | ahdc1 | 5 | AHDC1 | n | AT hook | 0 |
| ENSDART00000124751 | kcnip3b | 5 | KCNIP3 | n | Unknown | 0 |
| ENSDART00000003548 | znf385a | 5 | ZNF385A | n | C2H2 ZF | 0 |
| ENSDART00000092357 | sgsm2 | 5 | SGSM2 | n | BED ZF | 0 |
| ENSDART00000026865 | l3mbtl1a | 5 | L3MBTL1 | n | C2H2 ZF | 0 |
| ENSDART00000033362 | gatad2b | 5 | GATAD2B | n | GATA | 0 |
| ENSDART00000078529 | kin | 5 | KIN | n | C2H2 ZF | 0 |
| ENSDART00000162857 | nr4a3 | 5 | NR4A3 | n | Nuclear receptor | 0 |
| ENSDART00000110544 | znf219 | 5 | ZNF219 | n | C2H2 ZF | 0 |
| ENSDART00000156351 | zfat | 5 | ZFAT | n | C2H2 ZF | 0 |
| ENSDART00000166135 | zbtb47b | 5 | ZBTB47 | n | C2H2 ZF | 0 |
| ENSDART00000048775 | mbd3b | 5 | MBD3 | n | MBD | 0 |
| ENSDART00000157659 | camta1b | 5 | CAMTA1 | n | CG-1 | 0 |
| ENSDART00000142223 | hivep3b | 5 | HIVEP3 | n | C2H2 ZF | 0 |
| ENSDART00000055890 | znf385c | 5 | ZNF385C | n | C2H2 ZF | 0 |
| ENSDART00000135443 | kdm5ba | 5 | KDM5B | n | ARID/BRIGHT | 0 |
| ENSDART00000162387 | zbtb38 | 5 | ZBTB38 | n | C2H2 ZF | 0 |
| ENSDART00000172310 | zbtb4 | 5 | ZBTB4 | n | C2H2 ZF | 0 |
| ENSDART00000040672 | mecp2 | 5 | MECP2 | y | MBD; AT hook | 2 |
| ENSDART00000037879 | crx | 5 | CRX | y | Homeodomain | 1 |
| ENSDART00000122429 | klf15 | 5 | KLF15 | y | C2H2 ZF | 6 |
| ENSDART00000166313 | thrb | 5 | THRB | y | Nuclear receptor | 17 |
| ENSDART00000089748 | rorb | 5 | RORB | y | Nuclear receptor | 1 |
| ENSDART00000000876 | nr4a1 | 5 | NR4A1 | y | Nuclear receptor | 1 |
| ENSDART00000153187 | thrab | 5 | THRA | y | Nuclear receptor | 15 |
| ENSDART00000054736 | bhlhe23 | 5 | BHLHE23 | y | bHLH | 1 |
| ENSDART00000167052 | etv1 | 5 | ETV1 | y | Ets | 1 |
| ENSDART00000026017 | bhlhe40 | 5 | BHLHE40 | y | bHLH | 2 |
| ENSDART00000131134 | si:ch211-206a7.2 | 5 | FOXO6 | y | Forkhead | 1 |
| ENSDART00000024832 | stat5a | 5 | STAT5A | y | STAT | 1 |
| ENSDART00000091707 | dbpa | 5 | DBP | y | bZIP | 1 |
| ENSDART00000126966 | esrrga | 5 | ESRRG | y | Nuclear receptor | 1 |
| ENSDART00000127353 | nr1d2b | 5 | NR1D2 | y | Nuclear receptor | 7 |
| ENSDART00000047541 | bach1b | 5 | BACH1 | y | bZIP | 1 |
| ENSDART00000007630 | nhlh2 | 5 | NHLH2 | y | bHLH | 21 |

| | | | | | | |
|---|---|---|---|---|---|---|
| ENSDART00000129498 | mef2d | 5 | MEF2D | y | MADS box | 1 |
| ENSDART00000164349 | e2f4 | 5 | E2F4 | y | E2F | 1 |
| ENSDART00000157890 | tcf7l1b | 5 | TCF7L1 | y | HMG/Sox | 1 |
| ENSDART00000165757 | pax6b | 5 | PAX6 | y | Homeodomain; Paired box | 1 |
| ENSDART00000169609 | tefb | 5 | TEF | y | bZIP | 1 |
| ENSDART00000103640 | hey1 | 5 | HEY1 | y | bHLH | 1 |
| ENSDART00000013003 | tfap2b | 5 | TFAP2B | y | AP-2 | 1 |
| ENSDART00000082346 | tfap2a | 5 | TFAP2A | y | AP-2 | 5 |
| ENSDART00000147658 | bhlhe22 | 5 | BHLHE22 | y | bHLH | 1 |
| ENSDART00000057644 | lhx4 | 5 | LHX4 | y | Homeodomain | 1 |
| ENSDART00000009938 | tcf12 | 5 | TCF12 | y | bHLH | 1 |
| ENSDART00000065361 | etv5b | 5 | ETV5 | y | Ets | 1 |
| ENSDART00000055709 | her2 | 5 | HES5 | y | bHLH | 1 |
| ENSDART00000055706 | her15.1 | 5 | HES5 | y | bHLH | 1 |
| ENSDART00000133487 | fosb | 5 | FOSB | y | bZIP | 3 |
| ENSDART00000151127 | thraa | 5 | THRA | y | Nuclear receptor | 15 |
| ENSDART00000153167 | hlfb | 5 | HLF | y | bZIP | 2 |
| ENSDART00000086051 | mecom | 5 | MECOM | y | C2H2 ZF | 1 |
| ENSDART00000023613 | her6 | 5 | HES1 | y | bHLH | 1 |
| ENSDART00000031426 | skilb | 6 | SKIL | n | Unknown | 0 |
| ENSDART00000148066 | znf395b | 6 | ZNF395 | n | C2H2 ZF | 0 |
| ENSDART00000125344 | skilb | 6 | SKIL | n | Unknown | 0 |
| ENSDART00000131361 | kcnip3b | 6 | KCNIP3 | n | Unknown | 0 |
| ENSDART00000054020 | hivep3b | 6 | HIVEP3 | n | C2H2 ZF | 0 |
| ENSDART00000169283 | znf644b | 6 | ZNF644 | n | C2H2 ZF | 0 |
| ENSDART00000100667 | skia | 6 | SKI | n | Unknown | 0 |
| ENSDART00000100181 | sall3b | 6 | SALL3 | n | C2H2 ZF | 0 |
| ENSDART00000141734 | hivep2a | 6 | HIVEP2 | n | C2H2 ZF | 0 |
| ENSDART00000124740 | ncoa2 | 6 | NCOA2 | n | bHLH | 0 |
| ENSDART00000164082 | znf609a | 6 | ZNF609 | n | C2H2 ZF | 0 |
| ENSDART00000143165 | tsc22d1 | 6 | TSC22D1 | n | Unknown | 0 |
| ENSDART00000143874 | akna | 6 | AKNA | n | AT hook | 0 |
| ENSDART00000104279 | znf516 | 6 | ZNF516 | n | C2H2 ZF | 0 |
| ENSDART00000165710 | gpbp1l1 | 6 | GPBP1L1 | n | Unknown | 0 |
| ENSDART00000053367 | hmgn3 | 6 | HMGN3 | n | HMG/Sox | 0 |
| ENSDART00000164855 | crebl2 | 6 | CREBL2 | n | bZIP | 0 |
| ENSDART00000113286 | phf19 | 6 | PHF1 | n | Unknown | 0 |
| ENSDART00000078781 | znf706 | 6 | ZNF706 | n | C2H2 ZF | 0 |
| ENSDART00000125174 | nr1i2 | 6 | NR1I2 | y | Nuclear receptor | 5 |
| ENSDART00000089015 | zbtb7a | 6 | ZBTB7A | y | C2H2 ZF | 2 |
| ENSDART00000044860 | maff | 6 | MAFF | y | bZIP | 2 |
| ENSDART00000167844 | mafk | 6 | MAFK | y | bZIP | 2 |
| ENSDART00000057124 | tefa | 6 | TEF | y | bZIP | 1 |
| ENSDART00000057125 | tefa | 6 | TEF | y | bZIP | 1 |
| ENSDART00000163250 | mef2cb | 6 | MEF2C | y | MADS box | 1 |
| ENSDART00000131731 | mef2ca | 6 | MEF2C | y | MADS box | 1 |
| ENSDART00000161387 | tcf12 | 6 | TCF12 | y | bHLH | 1 |
| ENSDART00000127099 | nr2e3 | 6 | NR2E3 | y | Nuclear receptor | 1 |
| ENSDART00000066655 | mybl1 | 6 | MYBL1 | y | Myb/SANT | 1 |
| ENSDART00000172251 | creb3l1 | 6 | CREB3L1 | y | bZIP | 1 |
| ENSDART00000170575 | nfat5b | 6 | NFAT5 | y | Rel | 1 |
| ENSDART00000148106 | mef2aa | 6 | MEF2A | y | MADS box | 3 |
| ENSDART00000075070 | hsf2 | 6 | HSF2 | y | HSF | 1 |
| ENSDART00000148353 | usf2 | 6 | USF2 | y | bHLH | 2 |
| ENSDART00000061106 | bhlhe41 | 6 | BHLHE41 | y | bHLH | 1 |
| ENSDART00000139102 | dbpb | 6 | DBP | y | bZIP | 1 |
| ENSDART00000077839 | atf7b | 6 | ATF7 | y | bZIP | 1 |
| ENSDART00000045374 | smad3a | 6 | SMAD3 | y | SMAD | 2 |
| ENSDART00000093331 | rreb1a | 6 | RREB1 | y | C2H2 ZF | 1 |
| ENSDART00000148537 | roraa | 6 | RORA | y | Nuclear receptor | 1 |
| ENSDART00000123970 | mntb | 6 | MNT | y | bHLH | 1 |
| ENSDART00000127157 | hlfa | 6 | HLF | y | bZIP | 2 |
| ENSDART00000126282 | nr1d1 | 6 | NR1D1 | y | Nuclear receptor | 7 |

**Table S8. Gene ontology (GO) analysis of putative Jun transcriptional targets.**

| GO.ID | Term | Annotated | Significant | Expected | pvalue | Genes |
|---|---|---|---|---|---|---|
| GO:0003824 | catalytic activity | 5689 | 404 | 355.24 | 0.0414 | abca1a;abcb4;abcf1;abcg2c;abhd14a;abhd4;abl1;acadl;acadvl;ache;acsl2;acsl4a;acsl4b;acss1;adam15;adam23a;adam8a;adam8b;adat2;adhfe1;adkb;adss;agpat4;ahcy;ahcyl1;ahcyl2;ak2;alad;aldh4a1;aldoaa;aldoca;alg6;alpk2;ampd2b;ankib1a;apobec2b;arih2;ash1l;asns;asph;atic;atp1a1b;atp2a1;atp2b2;atp6ap1b;atp6ap1la;bcat2;blmh;btk;bub1bb;cad;camk1ga;camk1gb;camk2b2;camkmt;casp9;cbsa;cdc14ab;cdc42bpab;cdc42l;cdk1;cdk2;cdkn1a;cenpe;cers2a;cers4b;ckba;clk2a;clpp;cmtr1;cndp2;colgalt2;cracr2ab;csnk1a1;csnk1e;csnk1g1;ctbp2a;cth;cthl;ctsba;ctsc;ctsz;cyb5r1;dad1;dalrd3;dclk1b;ddx54;dgat1b;dgkab;dhfr;dhx32b;dhx33;dhx57;dip2ba;dnase1l4.1;dpf2;dpf2l;dph6;dpysl2b;dpysl4;dus2;dusp27;dusp6;dync1h1;dyrk1ab;ebp;ech1;eef1a1l2;eef2b;ehhadh;eif2ak1;eif2s3;eif3ha;eif4a1b;elovl1b;entpd5b;ern2;esd;etfdh;fahd2a;fam213b;fdps;fkbp1aa;fkbp5;fkbp9;ftsj3;fuca1.1;fuca1.2;galm;galns;gars;gba;gbp1;GDPGP1;ggh;gig2o;glrx;glud1a;gna12a;gnpda2;gnsb;gpx4a;GSK3B (1 of many);gstm.1;gstm.2;gtpbp1;guk1a;hadhb;hectd3;hibadha;hmbsa;hmox1a;hpdl;hspa14;hunk;iars;icmt;idi1;ifi30;ifit14;ifit16;ilk;impa1;impdh1b;impdh2;inpp4aa;ip6k1;ippk;isg20;jak1;kat5a;kif1b;kif20bb;kif26ab;kif26bb;kif3ca;kif3cb;kif4;kif5c;klc1a;klc3;klc4;klhl21;ksr2;lgmn;lgsn;limk1a;lipf;loxl4;lrsam1;lypla1;lypla2;manba;mapk4;mapk9;marcksb;mcm2;mcm5;mcm7;megf8;melk;mettl1;METTL18;mettl2a;mical1;mknk1;moxd1;mri1;msra;mthfd1l;mtm1;mtmr7a;myo10l3;myo18aa;myo19;myo1ea;nars;nat16;nek12;nme2b.1;nme4;nme7;npl;nsdhl;nsun5;nt5c2l1;nudt4b;nudt5;nudt9;oat;otud5a;OTUD7A;oxnad1;pafah1b3;paox1;parp2;parp4;parp8;pbk;pcsk1;pde4ca;pde4cb;pde9a;pdia6;pdk2b;pfkmb;pgam1a;pgm1;pgm2;pgm5;pik3cb;pip5kl1;pitrm1;pkn1a;pla2g15;plcxd1;pld3;plk1;pmt;pnpla7a;pnpla8;polr1c;polr2eb;polr3gla;ppcdc;ppil1;ppp5c;ppt1;prkx;prmt3;prpsap1;prss16;psen2;psma5;psma6l;psmb1;psmb2;psmc1a;ptges;ptp4a2b;ptp4a3;ptpdc1b;ptpn21;ptpn23a;pygmb;pyroxd2;qars;rab10;rab11bb;rab13;rab20;rab32a;rab3ab;rab41;rab42a;rab43;rabggtb;rabl2;rac1a;ran;rap1b;rap2b;rars;rbbp9;rhbdl3;rhoca;rhot1a;rnaseka;RNF14 (1 of many);rnf19b;rnpep;rps6kb1b;rrad;rragca;rras;rras2;rrm2;rtca;sae1;sardh;sars;sccpdhb;scpep1;sec14l8;sept3;setdb1a;setdb1b;sgk494a;sh3pxd2b;shmt1;si:ch1073-296d18.1;si:ch211-117c9.1;si:ch211-188f17.1;si:ch211-195b13.1;si:ch211-195b15.8;si:ch211-282j22.3;si:ch211-63o20.7;si:ch73-127m5.1;si:dkey-16p21.7;si:dkey-204f11.64;si:dkey-222f8.3;si:dkey-226m8.10;si:dkey-30c15.17;si:dkey-32e23.4;si:dkey-9i23.14;sik1;SLA (1 of many);slc3a2a;smarca5;smarcad1b;smox;smpd1;sod3b;spast;spra;spsb4a;src;srpk1a;st3gal3b;stambpl1;steap4;sult5a1;sult6b1;suox;tars;tcirg1b;tgm1l1;tmprss9;top1mt;tpte;trmt61a;tuba2;tuba8l;tubb4b;txn;tyr;uap1l1;ube2c;ube2e1;uck2a;uck2b;ugdh;umps;uqcrq;usp18;vcp;vcpip1;wars;wdsub1;wee1;zak;zdhhc12b;zgc:101663;zgc:103438;zgc:110269;zgc:112285;zgc:152830;zgc:153031;zgc:158659;zgc:162396;zgc:171704;zgc:198419;zgc:77752;zgc:92066 |
| GO:0016787 | hydrolase activity | 2346 | 180 | 146.49 | 0.03182 | abca1a;abcb4;abcf1;abcg2c;abhd14a;abhd4;ache;adam15;adam23a;adam8a;adam8b;adat2;ahcy;ahcyl1;ahcyl2;ampd2b;apobec2b;atic;atp1a1b;atp2a1;atp2b2;atp6ap1b;atp6ap1la;blmh;cad;casp9;cdc14ab;cdc42l;cenpe;clpp;cndp2;cracr2ab;ctsba;ctsc;ctsz;ddx54;dhx32b;dhx33;dhx57;dnase1l4.1;dpysl2b;dpysl4;dusp27;dusp6;dync1h1;eef1a1l2;eef2b;eif2s3;eif3ha;eif4a1b;entpd5b;ern2;esd;fuca1.1;fuca1.2;galns;gba;gbp1;ggh;gna12a;gnpda2;gnsb;gtpbp1;hspa14;iars;idi1;impa1;inpp4aa;isg20;kif1b;kif20bb;kif26ab;kif26bb;kif3ca;kif3cb;kif4;kif5c;klc1a;klc3;klc4;lgmn;lipf;lypla1;lypla2;manba;mcm2;mcm5;mcm7;megf8;mthfd1l;mtm1;mtmr7a;myo10l3;myo18aa;myo19;myo1ea;nt5c2l1;nudt4b;nudt5;nudt9;otud5a;OTUD7A;pafah1b3;pcsk1;pde4ca;pde4cb;pde9a;plcxd1;pnpla7a;pnpla8;ppp5c;ppt1;prss16;psen2;psma5;psma6l;psmb1;psmb2;psmc1a;ptp4a2b;ptp4a3;ptpdc1b;ptpn21;ptpn23a;rab10;rab11bb;rab13;rab20;rab32a;rab3ab;rab41;rab42a;rab43;rabl2;rac1a;ran;rap1b;rap2b;rbbp9;rhbdl3;rhoca;rhot1a;rnaseka;rnpep;rrad;rragca;rras;rras2;scpep1;sec14l8;sept3;si:ch211-195b15.8;si:ch211-282j22.3;si:ch73-127m5.1;si:dkey-204f11.64;si:dkey-222f8.3;si:dkey-226m8.10;si:dkey-30c15.17;si:dkey-32e23.4;si:dkey-9i23.14;smarca5;smarcad1b;smpd1;spast;stambpl1;tcirg1b;tmprss9;tpte;tuba2;tuba8l;tubb4b;usp18;vcp;vcpip1;zgc:103438;zgc:110269;zgc:112285;zgc:152830;zgc:171704;zgc:77752 |
| GO:0005525 | GTP binding | 518 | 47 | 32.35 | 0.0064 | adss;anxa6;arf1;ARL3 (1 of many);arl8a;arl8bb;cdc42l;cracr2ab;eef1a1l2;eef2b;eif2s3;gbp1;gna12a;gnl3;gtpbp1;gtpbp4;megf8;rab10;rab11bb;rab13;rab20;rab32a;rab3ab;rab41;rab42a;rab43;rabl2;rac1a;ran;rap1b;rap2b;rhoca;rhot1a;rrad;rragca;rras;rras2;sec14l8;sept3;si:dkey-32e23.4;si:dkey-98f17.5;si:dkeyp-69e1.8;tuba2;tuba8l;tubb4b;zgc:171704;zgc:77650 |
| GO:0008289 | lipid binding | 384 | 40 | 23.98 | 0.02919 | acbd5a;adap2;aida;anxa6;apoeb;col4a3bpa;col4a3bpb;cpne3;cpne9;esyt1a;fabp11a;fabp7a;fermt2;hs1bp3;melk;osbpl1a;pacsin3;picalmb;prom1b;pxdc1a;rasal1b;rbp1;rbp2a;rbp7b;rlbp1a;sdc4;sdcbp2;sh3pxd2b;si:ch211-195b13.1;si:dkey-231j24.3;snx12;snx19a;sptbn1;sptbn2;syt4;syt5a;syt6a;tex2;vcp;vps36 |
| GO:0003924 | GTPase activity | 347 | 37 | 21.67 | 0.00103 | cdc42l;cracr2ab;eef1a1l2;eef2b;eif2s3;gbp1;gna12a;gtpbp1;megf8;rab10;rab11bb;rab13;rab20;rab32a;rab3ab;rab41;rab42a;rab43;rabl2;rac1a;ran;rap1b;rap2b;rhoca;rhot1a;rrad;rragca;rras;rras2;sec14l8;sept3;si:dkey-204f11.64;si:dkey-32e23.4;tuba2;tuba8l;tubb4b;zgc:171704 |

| | | | | | |
|---|---|---|---|---|---|
| GO:0016887 | ATPase activity | 315 | 28 | 19.67 | 0.03947 | abca1a;abcb4;abcf1;abcg2c;atp1a1b;atp2a1;atp2b2;atp6ap1b;atp6ap1la;cenpe;ddx54;dhx32b;dhx33;dhx57;eif4a1b;hspa14;kif1b;kif20bb;kif3ca;kif3cb;kif4;kif5c;psmc1a;si:dkey-226m8.10;smarca5;spast;tcirg1b;vcp |
| GO:0008017 | microtubule binding | 179 | 25 | 11.18 | 0.00647 | camsap2a;camsap3;ccdc88b;ccsapb;cenpe;clasp2;dpysl2b;eml1;jakmip3;kif1b;kif20bb;kif26ab;kif26bb;kif3ca;kif3cb;kif4;kif5c;map6b;mapre1b;mapre3b;nusap1;pafah1b1a;prc1b;si:dkey-32e23.4;spast |
| GO:0005085 | guanyl-nucleotide exchange factor activity | 165 | 21 | 10.3 | 0.04798 | arhgef10lb;arhgef2;arhgef33;arhgef4;cyth1a;dennd5b;dennd6aa;dock11;dock9b;kalrna;mcf2b;mcf2l2;mcf2la;rabif;rapgef1a;rapgef4;rasgef1bb;sec61b;sh2d3ca;si:dkey-38p12.3;tiam2a |
| GO:0004725 | protein tyrosine phosphatase activity | 112 | 12 | 6.99 | 0.04675 | cdc14ab;dusp6;mtm1;mtmr7a;ptp4a2b;ptp4a3;ptpdc1b;ptpn21;ptpn23a;si:ch211-195b15.8;tpte;zgc:77752 |
| GO:0019843 | rRNA binding | 32 | 11 | 2 | 0.00013 | cirbpb;nop53;ppan;rpl12;rpl23;rpl5b;rpl8;rpl9;rps18;rps4x;rps9 |
| GO:0003743 | translation initiation factor activity | 61 | 11 | 3.81 | 0.00125 | eif2s1a;eif2s1b;eif2s2;eif2s3;eif3ba;eif3ha;eif3i;eif3m;eif4a1b;eif4g2a;eif6 |
| GO:0000149 | SNARE binding | 90 | 11 | 5.62 | 0.01428 | napga;snap25a;stx11a;stx11b.1;stx11b.2;stx12;stx3a;stx4;syt4;syt5a;syt6a |
| GO:0051082 | unfolded protein binding | 87 | 11 | 5.43 | 0.01927 | cct4;cct8;hsp90aa1.2;hsp90ab1;hspa14;pfdn1;pfdn2;pfdn4;ssuh2.2;ssuh2.4;tcp1 |
| GO:0005089 | Rho guanyl-nucleotide exchange factor activity | 86 | 10 | 5.37 | 0.0414 | arhgef10lb;arhgef2;arhgef33;arhgef4;kalrna;mcf2b;mcf2l2;mcf2la;si:dkey-38p12.3;tiam2a |
| GO:0008138 | protein tyrosine/serine/threonine phosphatase activity | 57 | 9 | 3.56 | 0.01616 | cdc14ab;dusp27;dusp6;ptp4a2b;ptp4a3;ptpdc1b;si:ch211-195b15.8;tpte;zgc:77752 |
| GO:0005484 | SNAP receptor activity | 34 | 8 | 2.12 | 0.00095 | sec22bb;snap25a;stx11a;stx11b.1;stx11b.2;stx12;stx3a;stx4 |
| GO:0003746 | translation elongation factor activity | 27 | 7 | 1.69 | 0.00107 | eef1a1l2;eef1b2;eef1db;eef1g;eef2b;eif5a2;gtpbp1 |
| GO:0004715 | non-membrane spanning protein tyrosine kinase activity | 41 | 7 | 2.56 | 0.01262 | abl1;btk;jak1;melk;SLA (1 of many);src;wee1 |
| GO:0031492 | nucleosomal DNA binding | 44 | 7 | 2.75 | 0.01831 | CU459186.1;hmgn3;si:ch211-113a14.12;si:dkey-108k21.14;si:dkey-261m9.12;zgc:153405;zgc:173552 |
| GO:0017049 | GTP-Rho binding | 18 | 6 | 1.12 | 0.00057 | anln;cdc42ep1a;cdc42ep2;cdc42ep5;pkn1a;zgc:154093 |
| GO:0030170 | pyridoxal phosphate binding | 43 | 6 | 2.69 | 0.04941 | cbsa;cth;cthl;oat;pygmb;shmt1 |
| GO:0043022 | ribosome binding | 23 | 5 | 1.44 | 0.0448 | c1qbp;eif2s1a;eif2s1b;eif5a2;eif6 |
| GO:0051010 | microtubule plus-end binding | 9 | 4 | 0.56 | 0.00148 | clasp2;mapre1b;mapre3b;pafah1b1a |
| GO:0044325 | ion channel binding | 21 | 4 | 1.31 | 0.03861 | cbarpb;rims1b;scn1bb;si:ch211-126j24.1 |
| GO:0031386 | protein tag | 10 | 3 | 0.62 | 0.02092 | rps27a;sumo2b;uba52 |
| GO:0070840 | dynein complex binding | 10 | 3 | 0.62 | 0.02092 | cenpf;dctn1b;pafah1b1a |
| GO:0008474 | palmitoyl-(protein) hydrolase activity | 11 | 3 | 0.69 | 0.02745 | lypla1;lypla2;ppt1 |
| GO:0008199 | ferric iron binding | 12 | 3 | 0.75 | 0.03494 | zgc:109934;zgc:198419;zgc:92066 |
| GO:0003962 | cystathionine gamma-synthase activity | 2 | 2 | 0.12 | 0.0039 | cth;cthl |
| GO:0004123 | cystathionine gamma-lyase activity | 2 | 2 | 0.12 | 0.0039 | cth;cthl |
| GO:0004146 | dihydrofolate reductase activity | 2 | 2 | 0.12 | 0.0039 | dhfr;zgc:153031 |
| GO:0004614 | phosphoglucomutase activity | 2 | 2 | 0.12 | 0.0039 | pgm1;pgm5 |
| GO:0004814 | arginine-tRNA ligase activity | 3 | 2 | 0.19 | 0.0112 | dalrd3;rars |
| GO:0004560 | alpha-L-fucosidase activity | 3 | 2 | 0.19 | 0.0112 | fuca1.1;fuca1.2 |
| GO:0003938 | IMP dehydrogenase activity | 3 | 2 | 0.19 | 0.0112 | impdh1b;impdh2 |
| GO:0046592 | polyamine oxidase activity | 3 | 2 | 0.19 | 0.0112 | paox1;smox |
| GO:0043515 | kinetochore binding | 3 | 2 | 0.19 | 0.0112 | clasp2;spdl1 |
| GO:0046933 | proton-transporting ATP synthase activity, rotational mechanism | 4 | 2 | 0.25 | 0.02148 | atp6ap1b;atp6ap1la |
| GO:0008097 | 5S rRNA binding | 4 | 2 | 0.25 | 0.02148 | nop53;rpl5b |
| GO:0005159 | insulin-like growth factor receptor binding | 5 | 2 | 0.31 | 0.03432 | igf1;socs2 |
| GO:0005528 | FK506 binding | 5 | 2 | 0.31 | 0.03432 | fkbp1aa;fkbp5 |
| GO:0004332 | fructose-bisphosphate aldolase activity | 5 | 2 | 0.31 | 0.03432 | aldoaa;aldoca |
| GO:0008022 | protein C-terminus binding | 5 | 2 | 0.31 | 0.03432 | pdzd11;sae1 |
| GO:0015232 | heme transporter activity | 6 | 2 | 0.37 | 0.04938 | flvcr1;flvcr2b |
| GO:0051011 | microtubule minus-end binding | 6 | 2 | 0.37 | 0.04938 | camsap2a;camsap3 |
| GO:0003708 | retinoic acid receptor activity | 6 | 2 | 0.37 | 0.04938 | nr1d4b;rarab |

**Table S9. Gene ontology analysis of regeneration-associated genes with peak expression during initial axon growth toward the midline (Fig. 1C).**

| GO.ID | Term | Annotated | Significant | Expected | pvalue | Genes |
|---|---|---|---|---|---|---|
| GO:0023052 | signaling | 4223 | 838 | 754.01 | 0.03835 | [illegible]1a;adgrl3.1;adm2a;adora1b;adra2a;afap1;agap1;agrn;aida;ajuba;akap12b;akt3a;aldh1a2;alg5;angptl1a;angptl2b;ank1a;ank1b;ank2a;ankrd6b;anxa6;ap2m1a;apaf1;apba1a;apc;apc2;aplnrb;appb;arap3;arhgap10;arhgap11a;arhgap12b;arhgap21a;arhgap22;arhgap23b;arhgap32a;arhgap33;arhgap35b;arhgap4b;arhgdia;arhgdig;arhgef10lb;arhgef11;arhgef2;arhgef33;arhgef4;arhgef9a;arhgef9b;arr3b;arrb1;atl1;avpr2ab;baxa;baxb;bbc3;bbs4;bcar1;bcl2b;bcl2l1;bcl6a;bcl9;bcr;bdnf;bhlhe41;bmp3;bmp4;bmpr2a;bnip3lb;bnip4;btk;BX897691.1;cab39l;cab39l1;CABZ01081780.1;cacna1aa;cacna1bb;cacna1g;cacna1ha;cacna1hb;cacna1i;cacnb1;cacnb2a;cacnb3a;cacnb3b;cacng2a;cacng3b;cacng5a;cacng7b;cacng8b;cad;cadpsb;calb2a;calb2b;calca;calm1b;calm3a;camk4;cart2;cart3;cbl;cby1;ccdc136b;ccdc28a;ccka;ccr9a;cd276;cd63;cdc42bpab;cdc42se2;cdkn1a;cdon;cds1;celsr3;cetn2;chata;chn1;chrm4a;chrm5a;chrna10a;chrna6;chrnb2a;chrnb3a;chrnb3b;chsy1;cib2;cish;citb;clstn1;clstn2;clstn3;cmklr1;cnksr2b;cntfr;col1a1a;cplx2;cplx2l;cplx3b;cplx4a;cplx4b;CR382281.1;creb3l3l;crhb;crhbp;crhr1;crlf1a;csnk1e;CT030188.1;CT990561.1;cx43;cxcr3.3;cyth1a;dab2ipa;dab2ipb;dact3a;dclk1a;dclk1b;dclk2a;dclk2b;ddit3;dedd1;def8;deptor;dgke;dgkh;dicp3.1;dicp3.3;diras1a;dkk3a;dkk3b;dlg1l;dlg3;dlg4a;dlgap1b;dlgap2a;dlgap3;dlgap4b;dlgap5;dmxl2;dock11;dock3;DOCK4 (1 of many);dock6;dock7;dock9b;dot1l;drd1a;drd1b;drd2b;drd4a;dusp5;efna1b;efna2a;efna3a;ek1;enpp1;epha4b;epha6;ephb1;ephb2a;ephb6;erbb4b;erlin2;esrra;esrrga;f3a;fam13b;fas;fermt2;fgd1;fgf11b;fgf12a;fgf12b;fgf13a;fgf14;fgf18a;fgf1a;fgf6a;fgf8a;fgfr4;fjx1;fndc5b;foxo3a;fscn1a;fto;fzd2;fzd7a;gabbr1a;gabbr1b;gabbr2;gabra1;gabra3;gabra5;gabra6b;gabrb1;gabrb2;gabrb3;gabrb4;gabrd;gabrg2;gabrr1;gabrr2a;gadd45ba;gadd45ga;galr2b;gatad2b;gc2;gcgra;gck;gdi1;gem;ghra;gid8a;gipr;gjd2b;glra1;glra2;glra3;glra4b;glrba;glrbb;gnao1a;gnao1b;GNAZ;gnb1b;gnb2;gnb3a;gnb5b;gng13b;gng5;gngt2a;gpc1a;gpc1b;gpc2;gpc3;gpr132b;gpr153;gpr158a;gpr158b;gpr186;gpr22a;gpr22b;gpr27;gpr37a;gpr37b;gpr37l1b;gpr52;gpr75;gpr78a;gpr84;gprc5ba;gprc5bb;gpx4b;grb2a;gria1b;gria2a;gria3a;gria3b;gria4a;gria4b;grid2;grik1a;grik1b;GRIK3;grik4;grin1a;grin1b;grin2aa;grin2ab;grk7a;grm1a;grm1b;grm2a;grm4;grm6a;grm6b;grm8a;grm8b;hbegfa;hbegfb;hcar1-4;hcn1;hdr;her15.1;her2;her6;hey1;hgfb;hmgn3;hmha1a;homer1b;homer3b;hrasa;hrasb;hrh3;hsp90ab1;htr2cl1;htr7c;hunk;id4;igf1;igf1rb;igf2b;il17a/f1;il17a/f3;il1rapl1b;il1rapl2;il4r.1;il6st;ilk;impa1;impa2;inhbaa;inpp5b;insra;iqgap2;iqsec3a;iqsec3b;irs2a;itga10;itga6a;itgae.2;itgav;itgb3b;itgb7;itpr1a;jagn1a;jak1;jun;jupa;kalrna;kat2a;kcnab1a;kcnab2b;kcnh1a;kcnh3;kcnh4b;kcnh5a;kcnh5b;kcnk3a;kcnk3b;kcnq3;kctd13;kctd9a;kita;klhl24b;klhl6;ksr2;lepa;lepr;lims1;lin7a;lpar1;magi2a;map2k2b;MAP3K13;map4k4;mapk11;mapk4;mapk9;mark4a;mast1a;mast3b;mcf2a;mcf2l2;mcf2la;mdka;mdkb;mecom;melk;mgll;mibp2;mknk1;mknk2a;mstnb;mtnr1aa;mtnr1ab;mtnr1ba;mtnr1bb;myo10l1; |
| GO:0034220 | ion transmembrane transport | 780 | 243 | 139.27 | 0.00029 | ankha;anxa6;aplnrb;aqp1a.1;asic1b;asic2;asic4a;asic4b;atp1a1b;atp1a3a;atp1b1b;atp1b2a;atp1b3a;atp1b3b;atp2a1;atp2b1a;atp2b2;atp2b3a;atp2b3b;atp6ap1b;atp6ap1la;atp6v0a1b;atp6v0a2a;atp6v0e1;atp6v1ba;atp6v1f;atp6v1h;atpv0e2;best2;CABZ01081780.1;cacna1aa;cacna1bb;cacna1da;cacna1g;cacna1ha;cacna1hb;cacna1i;cacna2d2a;cacna2d2b;cacna2d4b;cacnb1;cacnb2a;cacnb3a;cacnb3b;cacng2a;cacng3b;cacng5a;cacng6b;cacng7b;cacng8b;cbarpb;chrna10a;chrna6;chrnb2a;chrnb3a;chrnb3b;clcn7;clic4;clic5a;cnga3b;cox4i2;COX7A2 (1 of many);fam155a;fxyd6l;gabra1;gabra3;gabra5;gabra6b;gabrb1;gabrb2;gabrb3;gabrb4;gabrd;gabrg2;gabrr1;gabrr2a;gem;gjd2b;glra1;glra2;glra3;glra4b;glrba;glrbb;gria2a;gria3a;gria3b;gria4a;gria4b;grid2;grik1a;grik1b;GRIK3;grik4;grin1a;grin1b;grin2aa;grin2ab;hcn1;hcn2b;hcn4l;hecw2a;htr2cl1;itpr1a;jph3;kcna1a;kcna1b;kcna2a;kcna2b;kcna4;kcnab1a;kcnab2a;kcnab2b;kcnb1;kcnc1a;kcnc1b;kcnc2;kcnc3a;kcnc3b;kcnd1;kcnd2;kcng2;kcng4a;kcnh1a;kcnh3;kcnh4b;kcnh5a;kcnh5b;kcnh6a;kcnh7;kcnj11;kcnj11l;kcnj12a;kcnj13;kcnj14;kcnj2b;kcnj3b;kcnj5;KCNJ6;kcnj9;kcnk10a;kcnk10b;kcnk1a;kcnk2b;kcnk3a;kcnk3b;kcnk9;kcnma1a;kcnn1b;kcnn3;kcnq2a;kcnq2b;kcnq3;kcns3a;kcnt2;KCNV1;kcnv2b;klhl24b;lrrc8aa;lrrc8c;mcoln1a;ndufa4;piezo2a.2;ppifb;rem1;rhag;rrad;ryr1b;RYR2;ryr3;scn12aa;scn1bb;scn1lab;scn8aa;scn8ab;sfxn5b;shank1;shank2;shisa7b;shisa8b;si:ch1073-450f2.1;si:ch211-140m22.7;si:ch73-335m24.5;si:ch73-380n15.2;si:rp71-39b20.4;slc12a2;slc12a4;slc12a5a;slc12a5b;slc12a7a;slc12a9;slc15a4;slc16a3;slc16a8;slc17a7a;slc17a7b;slc18a3a;slc20a2;slc24a3;slc24a4a;slc24a4b;slc24a5;slc25a14;slc25a29;slc25a32a;slc30a8;slc30a9;slc32a1;slc33a1;slc34a2a;slc38a3b;slc38a4;slc39a1;slc43a2a;slc4a2b;slc4a4a;slc4a8;slc5a6a;slc6a15;SLC7A1 (1 of many);slc7a7;slc8a1b;slc8a2b;slc8a3;slc8a4a;slc8a4b;slc9a5;snap25a;snap25b;steap4;tcirg1b;tmem163b;tmem175;tpte;trpc1;trpc5a;trpm3;trpv1;ttyh1;ttyh2l;ttyh3b;vdac3;zgc:162160 |

| | | | | | | |
|---|---|---|---|---|---|---|
| GO:0006468 | protein phosphorylation | 1204 | 213 | 214.97 | 0.04144 | aak1a;aatka;abl1;acsl4a;ACVR1C;adcyap1b;adgrg1;aida;akt3a;alpk2;ankrd6b;aplnrb;atg13;bcr;bmp3;bmp4;bmpr2a;bsk146;btk;bub1bb;cab39l;cab39l1;camk1a;camk1b;camk1db;camk2a;camk2b1;camk2d1;camk2d2;camk2g1;camk2n1a;camk4;camkk1a;camkk1b;camkva;camkvb;camkvl;cart2;cart3;cbl;ccna2;ccnb1;ccng1;ccni;ccnjl;cdc42bpab;cdk1;cdk14;cdk15;cdk17;CDK18;cdk5r1a;cdk5r1b;cdk5r2a;cdk5r2b;cdkn1a;cds1;csnk1e;CT990561.1;dapk2a;dclk1a;dclk1b;dclk2a;deptor;dicp3.1;dicp3.3;dusp5;dyrk1ab;dyrk2;eif2ak1;eif2ak2;ek1;enpp1;epha4b;epha6;ephb1;ephb2a;ephb6;erbb4b;ern2;fgfr4;gadd45ba;gadd45ga;gc2;gcn1;ghra;gprc5ba;gprc5bb;grk7a;gsk3aa;GSK3B (1 of many);hipk2;hunk;igf1rb;il6st;ilk;inhbaa;inpp5jb;insra;jak1;kcnh1a;kcnh3;kcnh4b;kcnh5a;kcnh5b;kita;ksr2;lepr;limk1a;map2k2b;MAP3K13;map4k4;mapk11;mapk4;mapk9;mark1;mark4a;mast1a;mast3b;melk;mknk1;mknk2a;mstnb;mylk5;myo3b;nek12;nek6;ntrk3a;ntrk3b;nuak1b;nuak2;oxsr1a;pak1;pak2b;pbk;pdk2a;pdk2b;pdk3b;pdpk1a;pebp1;pik3cb;pim1;pim3;pimr138;pkn3;pkz;plk1;plk2a;ppp2r5b;prkacbb;prkca;prkcdb;prkceb;prkcq;prkd1;prkg2;prkx;prnpb;prok1;qars;rap1b;riok1;rps6ka3a;rps6ka5;rps6kb1b;rps6kl1;sgk1;sgk494a;sgk494b;sh2d3ca;sh3bp5b;sh3rf1;sh3rf2;si:ch1073-184j22.2;si:ch211-117c9.1;si:ch211-147h1.4;si:ch211-191i18.4;si:ch211-195b13.1;si:ch211-195b15.8;si:ch211-255i3.4;si:ch73-60h1.1;si:ch73-62l21.1;si:dkey-16p21.7;si:dkey-17m8.1;si:dkey-240h12.4;sik1;sik2a;SLA (1 of many);slkb;smarca4a;snrkb;spegb;spry2;spry4;src;srpk1b;stk17a;stk35;stk40;taok1b;tesk2;tgfb2;tgfb3;tnika;tnk2b;trib2;trib3;unc119a;unc119b;unm_sa1261;wee1;zak;zgc:158659 |
| GO:0050790 | regulation of catalytic activity | 883 | 180 | 157.66 | 0.0051 | acap3b;adcyap1b;agap1;agap3;aida;alox5ap;anos1a;anos1b;appb;arap3;arhgap10;arhgap12b;arhgap21a;arhgap23b;arhgap32a;arhgap33;arhgap4b;arhgdia;arhgdig;asap1b;asap2a;atg13;atp1b1b;atp1b2a;atp1b3a;atp1b3b;aup1;baxa;baxb;bbc3;bcl2l1;bcr;birc7;calm1b;calm3a;camk2n1a;cap2;cart2;cart3;cbl;ccna2;ccnb1;ccng1;ccni;ccnjl;cdk5r1a;cdk5r1b;cdk5r2a;cdk5r2b;cdkn1a;chn1;cish;cst14a.2;cst14b.1;ctsba;dab2ipa;dab2ipb;deptor;DOCK4 (1 of many);dusp5;dynll1;dynll2a;dynll2b;elmod1;ensaa;ensab;esco2;gadd45ba;gadd45ga;gchfr;gcn1;gdi1;git2a;gprc5ba;gprc5bb;gpsm1a;gpsm2;grpel1;guca1a;guca1b;guca1c;guca1g;hdr;hip1;hmha1a;igf1rb;iqgap2;map2k2b;MAP3K13;map4k4;mcm2;ncf2;nek12;oaz1b;oaz2b;oxsr1a;pafah1b1a;pak1;pak2b;pcsk1nl;phactr3a;phactr3b;pik3r1;pik3r3a;pik3r3b;plxna2;ppifb;ppp1r14aa;ppp1r14ba;ppp1r1b;ppp1r3cb;ppp2r2bb;ppp2r2ca;ppp2r2cb;ppp2r5b;prkab1a;prnpb;prok1;prpsap1;psmd1;psmd3;qars;rap1b;rap1gap;rap1gap2a;rap1gap2b;RAP1GDS1;rasgef1bb;rcan1a;rcan2;rcan3;rgl3a;rgs6;rgs8;rictora;rxfp2a;sec23b;serpine3;serpinh1a;serpini1;sgsm1b;sh3bp5b;sh3pxd2b;si:ch1073-184j22.2;si:ch211-191i18.4;si:ch211-195b11.3;si:ch211-195b15.7;si:ch211-195b15.8;si:ch211-255i3.4;si:ch211-270g19.5;si:dkey-17m8.1;si:dkey-203a12.8;si:dkey-242g16.2;si:dkey-77g12.1;slkb;smarca4a;socs2;spry2;spry4;srgap1a;srgap1b;srgap2;stxbp5a;syngap1b;taok1b;tbc1d10ab;tbc1d12a;tbc1d17;tbc1d25;tbc1d4;tiam1a;timp2b;timp4.3;tnika;trappc6bl;trib2;trib3;unc119a;unc119b;zak |
| GO:0007010 | cytoskeleton organization | 787 | 180 | 140.52 | 0.04594 | abl1;ablim1b;ablim3;ache;actr3b;add3a;afap1;ajuba;anln;ap1ar;apc;apc2;arhgef10lb;arhgef11;atat1;baiap2a;bcar1;CABZ01044277.1;CABZ01118678.1;calm1b;calm3a;camsap3;cap2;capza1a;cbfb;ccdc88b;ccsapb;cct5;cetn2;cfap206;cfl1;clasp2;clstn1;cnn3a;coro1cb;coro2ba;coro2bb;CU639469.1;dbn1;dchs1b;diaph3;dlg1l;dmtn;dpysl2b;dpysl3;DST;dusp27;dync1li2;efs;eml1;epb41b;epb41l3a;espnla;evla;fermt2;flncb;fmnl2a;fmnl2b;fmnl3;frmd5;fscn1a;fscn1b;fscn2a;fscn2b;GAN;grip2a;gsna;havcr1;hip1;hook1;inpp5jb;iqsec3a;iqsec3b;katnb1;kif11;kif4;lima1a;lrmp;map1aa;map1ab;map2;map4l;map6a;map6b;map6d1;map7a;map7d1a;map7d2b;mapre1b;mapre2;mapre3b;maptb;mark1;mark4a;mast1a;mef2aa;mef2d;mical3b;myom2a;nckap1;nusap1;pacsin1a;pafah1b1a;pafah1b1b;pfn2;pgm5;pleca;pls3;ppp1r9a;ppp1r9bb;prc1b;rac3a;rac3b;racgap1;rhobtb1;rhoca;rhogb;rhoua;rictora;rnd1a;rnd1b;sdc4;sdcbp;sdcbp2;sept15;sept3;sept5a;sept5b;shroom2a;si:ch211-114n24.6;si:ch211-207i1.2;si:ch73-199e17.1;si:dkey-56f14.7;si:dkeyp-57f11.2;sorbs1;sorbs2b;sorbs3;spag1a;specc1;specc1lb;spire1b;spire2;ssh1b;ssh2b;stmn2a;stmn2b;stmn3;stmn4;stmn4l;sun1;syne2b;tacc1;tmod2;tmsb;tmsb2;tnnt2c;tnnt3b;tpm3;triobpa;tuba1a;tuba1b;tuba1c;tuba2;tuba8l;tuba8l3;tuba8l4;tubb2;tubb2b;tubb4b;tubb5;vangl2;vcp;vil1;wasf3a;xirp2a;zgc:152977;zgc:153426;zgc:154093;zgc:65894;zgc:86764 |

| | | | | | | |
|---|---|---|---|---|---|---|
| GO:0048666 | neuron development | 570 | 177 | 101.77 | 0.01571 | ache;adcy1b;adcyap1b;agrn;alcamb;aldoaa;amigo1;anos1a;apc;appb;arl3l1;arl3l2;asah1b;ascl1a;atat1;atp2b1a;bbs4;bbs5;bdnf;bsg;CABZ01118678.1;cad;camsap3;capn1a;casp3a;casp9;ccsapb;cdh4;chd7;clstn1;cnp;cntfr;cntn2;cntn3a.1;col19a1;ctbp2a;cxcl12b;cyfip2;diras1a;dpysl2b;dpysl3;dpysl4;dpysl5a;dscama;e2f8;efna1b;efna2a;epha4b;etv1;etv5b;ewsr1b;fryb;fus;gldn;gpm6ab;gprc5ba;gprc5bb;grip1;grna;hcn1;hecw2a;hmx1;il6st;inab;inpp5jb;insm1a;irx4a;isl2a;islr2;kcnc3a;kctd13;kif1b;kif3a;kif5ab;kif5c;lgmn;lingo1a;lingo1b;llph;map1aa;map1ab;map2;map4k4;map4l;maptb;mcoln1a;mecp2;ncam1a;ncam1b;nckap1;ndrg4;nefma;nexn;nfasca;ninl;nptna;nptnb;nrg1;nrp1a;nrp2a;nrxn2a;nsfa;nyap2a;nyap2b;olfm1a;pacsin1a;pak1;pcdh18b;plppr4b;pls3;plxna2;ppp1r9a;ppp1r9bb;prmt8b;ptprfa;rab10;rac1a;rb1;rbpms2a;rbpms2b;robo2;robo3;robo4;rpgrip1;rtn4a;rtn4r;scn8aa;sema3fa;sema3ga;sema3gb;sema4aa;sema4bb;sema6bb;sema6e;sema7a;shank1;shank2;si:ch1073-450f2.1;si:ch211-113g11.6;si:ch211-129c21.1;si:dkey-16p21.7;skia;skib;skilb;slc33a1;slit2;slitrk2;slitrk3a;slitrk3b;slitrk4;slitrk6;snap25b;sox11a;sox11b;srgap2;ssh1b;ssh2b;stat3;stmn2a;stmn2b;stmn3;stmn4;stmn4l;tbx2b;tenm1;TENM2;tenm3;tenm4;thrb;tmem106bb;tmem59l;tnika;tnr;uba1;vangl2;zfyve27;zgc:77784 |
| GO:0007268 | chemical synaptic transmission | 352 | 147 | 62.85 | 0.00147 | adcyap1b;apba1a;bdnf;CABZ01081780.1;cacna1aa;cacna1bb;cacnb1;cacnb2a;cacnb3a;cacnb3b;cacng2a;cacng3b;cacng5a;cacng7b;cacng8b;cadpsb;calb2a;calb2b;chata;chrm4a;chrm5a;chrna10a;chrna6;chrnb2a;chrnb3a;chrnb3b;clstn1;clstn2;clstn3;cplx2;cplx2l;cplx3b;cplx4a;cplx4b;CR382281.1;dlg1l;dlg3;dlg4a;dlgap1b;dlgap2a;drd4a;gabra1;gabra3;gabra5;gabra6b;gabrb1;gabrb2;gabrb3;gabrb4;gabrd;gabrg2;gabrr1;gabrr2a;glra1;glra2;glra3;glra4b;glrba;glrbb;grik1a;grik1b;GRIK3;grik4;grin2aa;grin2ab;grm1a;grm1b;grm2a;grm4;grm6a;grm6b;grm8a;grm8b;hcn1;htr2cl1;htr7c;kat2a;kcnq3;lin7a;napaa;napab;napba;napbb;neto1;nlgn1;nlgn2a;nlgn2b;nlgn3a;nptx2a;nrxn2a;nsmfa;nsmfb;otofa;pdzd11;penka;plcl2;ppfia3;ppp1r9bb;rimbp2;rims1b;rims2a;rims2b;rims3;sdcbp;sdcbp2;shank1;shank2;shisa7b;shisa8b;si:ch1073-450f2.1;si:ch73-380n15.2;slc12a4;slc12a5a;slc12a5b;slc12a7a;slc17a6b;slc17a7a;slc17a7b;snap25a;snap25b;stx11a;stx11b.2;stx2a;stx4;sv2a;sv2ba;sv2bb;sv2c;syn1;syn2a;sypa;sypb;syt10;syt11b;syt12;syt13;syt1a;syt2a;syt3;syt4;syt5a;syt5b;syt6a;syt6b;syt7b;syt9a;syt9b |
| GO:0031175 | neuron projection development | 443 | 143 | 79.1 | 0.00597 | adcy1b;adcyap1b;agrn;alcamb;aldoaa;amigo1;anos1a;apc;appb;arl3l1;arl3l2;asah1b;ascl1a;bdnf;bsg;CABZ01118678.1;cad;camsap3;capn1a;casp3a;casp9;ccsapb;clstn1;cnp;cntfr;cntn2;cntn3a.1;col19a1;cxcl12b;cyfip2;dpysl2b;dpysl3;dpysl4;dpysl5a;dscama;e2f8;efna1b;efna2a;epha4b;etv5b;fryb;fus;gpm6ab;gprc5ba;gprc5bb;grip1;grna;hecw2a;il6st;inab;inpp5jb;isl2a;islr2;kcnc3a;kctd13;kif1b;kif3a;kif5ab;kif5c;lgmn;lingo1a;llph;map1aa;map1ab;map2;map4k4;map4l;maptb;mecp2;ncam1a;ncam1b;nckap1;nefma;nexn;nfasca;nptna;nptnb;nrp1a;nrp2a;nrxn2a;nsfa;nyap2a;nyap2b;olfm1a;pacsin1a;pak1;pcdh18b;plppr4b;pls3;plxna2;ppp1r9a;ppp1r9bb;prmt8b;ptprfa;rab10;rac1a;rb1;rbpms2a;rbpms2b;robo2;robo3;robo4;rtn4a;rtn4r;scn8aa;sema3fa;sema3ga;sema3gb;sema4aa;sema4bb;sema6bb;sema6e;sema7a;shank1;shank2;si:ch1073-450f2.1;si:ch211-113g11.6;si:ch211-129c21.1;si:dkey-16p21.7;slc33a1;slit2;slitrk2;slitrk3a;slitrk3b;slitrk4;slitrk6;srgap2;ssh1b;ssh2b;stat3;stmn2a;stmn2b;stmn3;stmn4;stmn4l;tenm3;tmem106bb;tmem59l;tnika;tnr;uba1;vangl2;zfyve27;zgc:77784 |
| GO:0007417 | central nervous system development | 519 | 127 | 92.67 | 0.00406 | adcyap1b;adgrg1;agrn;agtpbp1;alcamb;aldh1a2;aldoaa;amigo1;anos1a;apc;appb;ascl1a;ascl1b;asphd1;atp1a1b;atp1a3a;bcl11aa;bcl11ab;bdnf;cad;capn1a;ccsapb;cenpf;chn1;cnnm2b;cntn2;CU639469.1;CU861477.1;cxcl12b;cyp26c1;dpysl2b;drd4a;dync1h1;eif3ha;enpp1;etv5b;ewsr1b;fgf8a;fn1a;gbx2;glra1;h2afy2;hapln1a;her15.1;her6;hgfb;hmgcs1;hsbp1b;igf1rb;igf2b;insm1a;irx7;josd2;jun;katnb1;kcnb1;kcnh1a;kctd13;kirrel3l;lgi1b;lzic;map2;marcksa;marcksb;marcksl1a;mdkb;meis3;msi2b;ncam1a;ncanb;nlrc3l;nono;nrg1;nsfa;ntrk3a;otud4;pafah1b1a;pax6b;pcdh10b;pcdh19;pebp1;pou4f2;pou4f3;prdm8b;prmt8b;prnpb;psen2;psmd3;ptpro;ptprsa;pycr1b;qars;rbfox1;robo2;rps15a;rps4x;rtn4a;sez6l2;sfrp5;shank1;shank2;SHC3;si:ch1073-450f2.1;si:ch73-119p20.1;six3a;slc33a1;slc7a7;slc9a3r1a;slit2;smarca4a;smc1a;sox11a;sox11b;spry4;syngap1b;tacc1;tbx2b;tgif1;thraa;tpp1;usp21;usp24;usp28;vangl2;vcana;vwc2;ywhag1 |

| | | | | | | |
|---|---|---|---|---|---|---|
| GO:0016477 | cell migration | 546 | 119 | 97.49 | 0.02508 | akap12b;alcamb;ankrd6b;anos1a;anxa6;apc;apc2;aplnrb;apoc1;arhgap4b;arrb1;atp1a1b;bcar1;ccr9a;chd7;cmklr1;cntn2;ctnnd1;cxcl12b;cxcl14;cxcr3.3;dmxl2;dpysl3;dscama;efs;esrra;fermt2;fgf8a;fmnl3;fn1a;fscn1a;fscn1b;fscn2a;fscn2b;fzd2;fzd7a;git2a;gnb1b;gpc3;grip2a;hgfb;hsp90ab1;igf1rb;igsf11;ildr1b;inpp5d;insm1a;isl2a;itgav;itgb3b;itgb7;jagn1a;kcnj13;kita;klf6a;lama5;megf8;meis3;mmp14b;mmp9;mt2;nckap1;nrg1;nrp1a;nrp2a;nsmfa;nsmfb;nusap1;pak1;pik3cb;plekhg5a;plxna2;prcp;ptges;ptpn23a;pum3;rac1a;rac3a;rac3b;rap1b;rcc2;rhobtb1;rhoca;rnd1a;rnd1b;robo4;rtn4a;rxfp3.3b;sdc2;sdc4;sema3fa;sema3ga;sema3gb;sema4aa;sema4bb;sema6bb;sema6e;sema7a;si:ch211-113g11.6;si:ch211-129c21.1;si:ch211-215c18.3;si:dkey-56f14.7;slc24a5;slit2;smarca4a;srgap1a;srgap1b;srgap2;sst6;stat3;tbx2b;thy1;tmie;tmsb;tmsb2;vangl2;zbtb4;zeb2b;zgc:158659 |
| GO:0048812 | neuron projection morphogenesis | 351 | 115 | 62.67 | 0.01301 | adcy1b;agrn;alcamb;aldoaa;anos1a;apc;appb;arl3l1;arl3l2;asah1b;bdnf;bsg;cad;capn1a;casp3a;casp9;clstn1;cntn2;cntn3a.1;col19a1;cxcl12b;cyfip2;dpysl2b;dpysl3;dpysl4;dpysl5a;dscama;e2f8;efna1b;efna2a;epha4b;etv5b;fus;grna;hecw2a;inab;isl2a;islr2;kcnc3a;kctd13;kif1b;kif3a;kif5ab;kif5c;llph;map1aa;map1ab;map4k4;mecp2;ncam1a;ncam1b;nckap1;nexn;nfasca;nptna;nptnb;nrp1a;nrp2a;nrxn2a;nsfa;nyap2a;nyap2b;olfm1a;pacsin1a;pak1;pcdh18b;plppr4b;pls3;plxna2;prmt8b;ptprfa;rac1a;rb1;rbpms2a;rbpms2b;robo2;robo3;robo4;rtn4a;rtn4r;scn8aa;sema3fa;sema3ga;sema3gb;sema4aa;sema4bb;sema6bb;sema6e;sema7a;shank1;shank2;si:ch1073-450f2.1;si:ch211-113g11.6;si:ch211-129c21.1;si:dkey-16p21.7;slc33a1;slit2;slitrk2;slitrk3a;slitrk3b;slitrk4;slitrk6;srgap2;ssh1b;ssh2b;stat3;tenm3;tmem106bb;tmem59l;tnika;tnr;uba1;vangl2;zfyve27;zgc:77784 |
| GO:0006412 | translation | 426 | 113 | 76.06 | 0.0002 | aimp1;ajuba;atp6v0e1;cirbpb;cpeb1a;cpeb3;dalrd3;disp2;dph6;dre-mir-21-1;dre-mir-21-2;eef1a1b;eef1a1l2;eef1a2;eef1b2;eef1db;eef1g;eef2a.2;eef2b;eif2ak1;eif2ak2;eif2s1a;eif2s1b;eif2s2;eif2s3;eif3ba;eif3f;eif3ha;eif3i;eif3m;eif3s6ip;eif4a1b;eif4ebp1;eif4g1a;eif4g2a;eif5a;eif5a2;eif6;eprs;farsb;faub;fxr1;fxr2;gars;gcn1;gemin5;gtpbp1;gtpbp2b;hspa14;iars;igf2bp3;lin28a;mov10b.1;nars;nck2a;nhp2;pkz;qars;rars;rpl10a;rpl12;rpl18;rpl21;rpl22;rpl22l1;rpl23;rpl23a;rpl27;rpl29;rpl30;rpl34;rpl35;rpl35a;rpl36a;rpl38;rpl39;rpl4;RPL41;rpl5a;rpl5b;rpl8;rpl9;rps11;rps12;rps13;rps15a;rps17;rps18;rps21;rps23;rps24;rps27.2;rps28;rps29;rps3a;rps4x;rps6kb1b;rps7;rps8a;rps8b;rps9;rpsa;rwdd1;sars;shmt1;tars;tnrc6c2;uba52;ube2e1;upf3a;uqcc2;wars;ybx1 |
| GO:0007017 | microtubule-based process | 400 | 98 | 71.42 | 0.01987 | apc;apc2;arhgef11;atat1;CABZ01044277.1;CABZ01118678.1;calm1b;calm3a;camsap3;ccdc88b;ccsapb;cenpe;cetn2;cfap206;clasp2;clstn1;CU639469.1;dchs1b;dctn1b;dnah2;DST;dync1h1;dync1i1;dync1i2a;dync1i2b;dync1li2;dynll1;dynll2a;dynll2b;eml1;hook1;inpp5jb;ippk;katnb1;kif11;kif1aa;kif1ab;kif1b;kif20a;kif20bb;kif21a;kif3a;kif3ca;kif3cb;kif4;kif5ab;kif5c;lrmp;map1aa;map1ab;map2;map4l;map6a;map6b;map6d1;map7a;map7d1a;map7d2b;mapre1b;mapre2;mapre3b;maptb;mark1;mark4a;nusap1;pafah1b1a;pafah1b1b;prc1b;racgap1;rhot1a;si:ch211-114n24.6;si:ch211-235e9.8;si:ch211-257p13.3;si:ch73-199e17.1;spag1a;stmn2a;stmn2b;stmn3;stmn4;stmn4l;tacc1;tuba1a;tuba1b;tuba1c;tuba2;tuba8l;tuba8l3;tuba8l4;tubb2;tubb2b;tubb4b;tubb5;vangl2;vbp1;vcp;zgc:153426;zgc:65894;zgc:86764 |
| GO:0007409 | axonogenesis | 295 | 95 | 52.67 | 0.04257 | adcy1b;agrn;alcamb;aldoaa;anos1a;apc;appb;arl3l1;arl3l2;asah1b;bsg;capn1a;casp3a;casp9;clstn1;cntn2;cntn3a.1;col19a1;cxcl12b;cyfip2;dpysl2b;dpysl3;dpysl5a;dscama;e2f8;efna1b;efna2a;epha4b;etv5b;fus;grna;isl2a;islr2;kcnc3a;kctd13;kif1b;kif3a;kif5ab;kif5c;map1aa;map1ab;mecp2;ncam1a;ncam1b;nexn;nfasca;nptna;nptnb;nrp1a;nrp2a;nrxn2a;nsfa;olfm1a;pak1;plppr4b;pls3;plxna2;ptprfa;rb1;rbpms2a;rbpms2b;robo2;robo3;robo4;rtn4a;rtn4r;scn8aa;sema3fa;sema3ga;sema3gb;sema4aa;sema4bb;sema6bb;sema6e;sema7a;si:ch211-113g11.6;si:ch211-129c21.1;si:dkey-16p21.7;slc33a1;slit2;slitrk2;slitrk3a;slitrk3b;slitrk4;slitrk6;ssh1b;ssh2b;stat3;tenm3;tmem59l;tnr;uba1;vangl2;zfyve27;zgc:77784 |
| GO:0007264 | small GTPase mediated signal transduction | 379 | 93 | 67.67 | 0.00016 | akap12b;arhgap32a;arhgap33;arhgdia;arhgdig;arhgef10lb;arhgef11;arhgef2;arhgef33;arhgef4;arhgef9a;arhgef9b;bcr;ccdc28a;cdc42se2;chn1;cyth1a;dock11;dock3;DOCK4 (1 of many);dock6;dock7;dock9b;fgd1;gdi1;grb2a;iqsec3a;iqsec3b;kalrna;kctd13;ksr2;mcf2a;mcf2l2;mcf2la;plekhg5a;prex2;rab10;rab11a;rab15;rab20;rab22a;rab32a;rab33a;rab34a;rab34b;rab39ba;rab3aa;rab3ab;rab41;rab42a;rab43;rab4a;rab6ba;rab6bb;rabif;rabl2;rac1a;rac3a;rac3b;ralba;ralgps1;rap1b;rap1gap;rap1gap2a;rap1gap2b;rapgef1a;rapgef4;rapgef5a;rapgef5b;rapgefl1;rasal1b;rasgef1bb;RASGRF1;rgl3a;rhobtb1;rhoca;rhogb;rhot1a;rhoua;rnd1a;rnd1b;robo4;sh2d3ca;si:dkey-38p12.3;sos2;spata13;spry2;spry4;tiam1a;tiam2a;vav3b;zgc:154093;zgc:91909 |

| | | | | | | |
|---|---|---|---|---|---|---|
| GO:0007420 | brain development | 373 | 88 | 66.6 | 0.00193 | adcyap1b;agtpbp1;alcamb;aldh1a2;aldoaa;amigo1;anos1a;apc;ascl1a;ascl1b;asphd1;atp1a1b;atp1a3a;bcl11aa;bcl11ab;bdnf;capn1a;ccsapb;cenpf;cnnm2b;CU639469.1;cxcl12b;cyp26c1;dpysl2b;eif3ha;enpp1;etv5b;fgf8a;fn1a;gbx2;h2afy2;her15.1;her6;hgfb;igf2b;irx7;jun;katnb1;kcnb1;kcnh1a;kctd13;kirrel3l;lgi1b;marcksa;marcksb;mdkb;meis3;msi2b;nrg1;nsfa;otud4;pafah1b1a;pax6b;pcdh10b;pcdh19;pebp1;pou4f2;pou4f3;prmt8b;psen2;psmd3;ptpro;ptprsa;pycr1b;qars;rbfox1;rps15a;rps4x;rtn4a;sez6l2;sfrp5;shank1;shank2;si:ch1073-450f2.1;si:ch73-119p20.1;six3a;slc9a3r1a;sox11a;sox11b;spry4;syngap1b;tacc1;tbx2b;tgif1;thraa;vangl2;vwc2;ywhag1 |
| GO:0023014 | signal transduction by protein phosphorylation | 326 | 67 | 58.21 | 0.03029 | acsl4a;adcyap1b;aida;ankrd6b;bmp3;bmp4;cab39l;cab39l1;cart2;cart3;cds1;CT990561.1;dicp3.1;dicp3.3;dusp5;epha4b;epha6;ephb1;ephb2a;ephb6;erbb4b;gadd45ba;gadd45ga;igf1rb;ilk;inhbaa;insra;kcnh1a;kcnh3;kcnh4b;kcnh5a;kcnh5b;kita;map2k2b;MAP3K13;map4k4;mapk11;mapk4;mapk9;mstnb;nek12;ntrk3a;ntrk3b;oxsr1a;pak1;pak2b;pebp1;prok1;qars;rap1b;sh3rf1;sh3rf2;si:ch1073-184j22.2;si:ch211-191i18.4;si:ch211-195b15.8;si:dkey-17m8.1;slkb;smarca4a;spry2;spry4;taok1b;tgfb2;tgfb3;tnika;trib2;trib3;zak |
| GO:0050804 | modulation of chemical synaptic transmission | 110 | 59 | 19.64 | 0.01314 | adcyap1b;bdnf;CABZ01081780.1;cacng2a;cacng3b;cacng5a;cacng7b;cacng8b;calb2a;calb2b;clstn1;clstn2;clstn3;cplx2;cplx2l;cplx3b;cplx4a;cplx4b;grik1a;grik1b;GRIK3;grik4;grin2aa;grin2ab;grm1a;grm1b;grm2a;grm4;grm6a;grm6b;grm8a;grm8b;kat2a;napaa;napab;napba;napbb;neto1;nlgn1;nlgn2a;nlgn2b;nlgn3a;nptx2a;nsmfa;nsmfb;plcl2;ppp1r9bb;rims1b;rims2a;rims2b;rims3;shank1;shank2;shisa7b;shisa8b;si:ch1073-450f2.1;sypa;sypb;syt12 |
| GO:0051726 | regulation of cell cycle | 519 | 58 | 92.67 | 0.00636 | adcyap1b;anapc15;apc;brinp2;brinp3a.1;bub1bb;cables2b;ccna2;ccnb1;ccng1;ccni;ccnjl;ccsapb;cdk1;cdk15;cdkn1a;cetn2;chmp4bb;dlg1l;dot1l;dync1li2;e2f3;e2f8;etv5b;gadd45ba;gadd45ga;hsp90ab1;id4;insm1a;jun;junba;kif11;mdka;mdm4;pea15;pebp1;pim3;pimr138;ppp1r13ba;prc1b;rab11fip4a;rb1;rpl23a;rpl35;rpl36a;rpl7;rps15a;rps18;rps29;rps7;rps8a;si:ch211-147h1.4;si:dkey-25o16.4;si:dkey-79d12.5;spdl1;ssrp1a;tp53bp1;zwilch |
| GO:0006816 | calcium ion transport | 169 | 57 | 30.17 | 0.00343 | anxa6;aplnrb;atp2a1;atp2b1a;atp2b2;atp2b3a;atp2b3b;CABZ01081780.1;cacna1aa;cacna1bb;cacna1da;cacna1g;cacna1ha;cacna1hb;cacna1i;cacna2d2a;cacna2d2b;cacna2d4b;cacnb1;cacnb2a;cacnb3a;cacnb3b;cacng2a;cacng3b;cacng5a;cacng6b;cacng7b;cacng8b;cbarpb;fam155a;gem;homer1b;homer3b;htr2cl1;itpr1a;jph3;mcoln1a;psen2;rem1;rrad;ryr1b;RYR2;ryr3;si:ch211-270g19.5;si:ch73-335m24.5;slc24a3;slc24a4a;slc24a4b;slc24a5;slc8a1b;slc8a2b;slc8a3;slc8a4a;slc8a4b;trpc1;trpc5a;trpv1 |
| GO:0042391 | regulation of membrane potential | 157 | 56 | 28.03 | 0.00309 | adcyap1b;cacna1g;cacna1ha;cacna1hb;cacna1i;chrna10a;chrna6;chrnb2a;chrnb3a;chrnb3b;gabra1;gabra3;gabra5;gabra6b;gabrb1;gabrb2;gabrb3;gabrb4;gabrd;gabrg2;gabrr1;gabrr2a;glra1;glra2;glra3;glra4b;glrba;glrbb;grin2aa;grin2ab;grm1a;grm1b;kcnab1a;kcnab2b;kcnh1a;kcnh3;kcnh4b;kcnh5a;kcnh5b;kcnh6a;kcnh7;kcnt2;piezo2a.2;rims1b;rims2a;rims2b;rims3;scn12aa;scn1bb;scn1lab;scn8aa;scn8ab;shank1;shank2;si:ch1073-450f2.1;si:ch73-380n15.2 |
| GO:0007015 | actin filament organization | 253 | 55 | 45.17 | 0.03905 | actr3b;add3a;ap1ar;arhgef10lb;arhgef11;baiap2a;bcar1;cap2;capza1a;cct5;cfl1;dbn1;dlg1l;dpysl3;efs;espnla;evla;fmnl3;fscn1a;fscn1b;fscn2a;fscn2b;gsna;hip1;lima1a;mical3b;myom2a;pfn2;pls3;ppp1r9a;ppp1r9bb;rac3a;rac3b;rhobtb1;rhoca;rhogb;rhoua;rnd1a;rnd1b;sdc4;shroom2a;sorbs1;sorbs2b;sorbs3;spire1b;spire2;ssh1b;ssh2b;tmod2;tmsb;tmsb2;tpm3;vangl2;vil1;zgc:154093 |
| GO:0070588 | calcium ion transmembrane transport | 146 | 48 | 26.07 | 0.00014 | anxa6;aplnrb;atp2a1;atp2b1a;atp2b2;atp2b3a;atp2b3b;CABZ01081780.1;cacna1aa;cacna1bb;cacna1da;cacna1g;cacna1ha;cacna1hb;cacna1i;cacna2d2a;cacna2d2b;cacna2d4b;cacnb1;cacnb2a;cacnb3a;cacnb3b;cacng2a;cacng3b;cacng5a;cacng6b;cacng7b;cacng8b;cbarpb;fam155a;gem;htr2cl1;itpr1a;jph3;mcoln1a;rem1;rrad;ryr1b;RYR2;ryr3;si:ch73-335m24.5;slc24a3;slc24a4a;slc24a4b;slc24a5;trpc1;trpc5a;trpv1 |
| GO:0007269 | neurotransmitter secretion | 71 | 41 | 12.68 | 0.01159 | cadpsb;cplx2;cplx2l;cplx3b;cplx4a;cplx4b;lin7a;napaa;napab;napba;napbb;otofa;pdzd11;ppfia3;rims1b;rims2a;rims2b;rims3;snap25a;snap25b;stx11a;stx11b.2;stx2a;stx4;syn1;syn2a;syt10;syt11b;syt12;syt13;syt1a;syt2a;syt3;syt4;syt5a;syt5b;syt6a;syt6b;syt7b;syt9a;syt9b |
| GO:0035335 | peptidyl-tyrosine dephosphorylation | 112 | 31 | 20 | 0.00656 | ca16b;cdc14ab;dusp23b;DUSP26;dusp3a;dusp5;mtmr7a;mtmr7b;ptenb;ptp4a1;ptp4a2b;ptp4a3;ptpdc1b;ptpmt1;ptpn11b;ptpn21;ptpn23a;ptpn4b;ptprdb;ptprfa;ptprnb;ptpro;ptprsa;ptprt;ptprz1b;si:ch1073-184j22.2;si:ch211-195b15.8;ssh1b;ssh2b;tpte;zgc:77752 |
| GO:0007018 | microtubule-based movement | 156 | 31 | 27.85 | 0.01777 | arhgef11;ccsapb;cenpe;cfap206;clstn1;dctn1b;dnah2;dync1h1;dync1i1;dync1i2a;dync1i2b;dync1li2;ippk;kif11;kif1aa;kif1ab;kif1b;kif20a;kif20bb;kif21a;kif3a;kif3ca;kif3cb;kif4;kif5ab;kif5c;pafah1b1a;pafah1b1b;rhot1a;si:ch211-235e9.8;si:ch211-257p13.3 |
| GO:0003143 | embryonic heart tube morphogenesis | 113 | 30 | 20.18 | 0.04242 | apc;atp1a1b;atp6ap1b;atp6v1f;bbs4;bbs5;bmp4;bmpr2a;bnip3lb;cacnb2a;camk2a;camk2g1;cenpf;chd7;fabp3;fn1a;fzd2;gpr22a;mef2cb;ndrg4;pak1;pkp2;scn12aa;skia;skib;slc8a4a;smarca4a;tbx2b;tbx5b;trib3 |
| GO:0035249 | synaptic transmission, glutamatergic | 50 | 29 | 8.93 | 0.0001 | adcyap1b;cacng2a;cacng3b;cacng5a;cacng7b;cacng8b;grik1a;grik1b;GRIK3;grik4;grm1a;grm1b;grm2a;grm4;grm6a;grm6b;grm8a;grm8b;napaa;napab;napba;napbb;nptx2a;shank1;shank2;si:ch1073-450f2.1;slc17a6b;slc17a7a;slc17a7b |

| | | | | | | |
|---|---|---|---|---|---|---|
| GO:0051480 | regulation of cytosolic calcium ion concentration | 166 | 28 | 29.64 | 0.00462 | adcyap1b;aplnrb;atp2b1a;atp2b2;atp2b3a;atp2b3b;calb2a;calb2b;ccr9a;cmklr1;cxcr3.3;fam155a;grm1a;grm1b;htr2cl1;jph3;mcoln1a;pvalb2;pvalb3;pvalb6;pvalb9;rxfp3.3b;ryr1b;RYR2;ryr3;slc8a4a;trpc1;trpc5a |
| GO:0007626 | locomotory behavior | 85 | 28 | 15.18 | 0.01321 | abhd12;adgrl3.1;atp1a3a;bag3;chata;fhl1a;flnca;flncb;gdpd5a;gprc5ba;gprc5bb;htr2cl1;mecp2;nr1d1;pleca;psmd3;ryr1b;scn1lab;scn8aa;slc18a2;slc1a2b;snap25a;snap25b;sncb;sncgb;syngap1b;tpp1;vcp |
| GO:0050770 | regulation of axonogenesis | 90 | 27 | 16.07 | 0.04238 | arl3l1;arl3l2;casp3a;casp9;clstn1;cntn2;dpysl2b;epha4b;pak1;plxna2;rbpms2a;rbpms2b;robo2;sema3fa;sema3ga;sema3gb;sema4aa;sema4bb;sema6bb;sema6e;sema7a;si:ch211-113g11.6;si:ch211-129c21.1;slit2;ssh1b;ssh2b;zfyve27 |
| GO:1902476 | chloride transmembrane transport | 69 | 24 | 12.32 | 0.00057 | best2;clcn7;clic4;clic5a;gabra1;gabra3;gabra5;gabra6b;gabrg2;glra1;glra2;glra3;glra4b;glrba;glrbb;slc12a2;slc12a4;slc12a5a;slc12a5b;slc12a7a;slc12a9;ttyh1;ttyh2l;ttyh3b |
| GO:0050806 | positive regulation of synaptic transmission | 42 | 22 | 7.5 | 0.00014 | adcyap1b;CABZ01081780.1;cacng2a;cacng3b;cacng5a;cacng7b;cacng8b;calb2a;calb2b;clstn1;clstn2;clstn3;grin2aa;grin2ab;rims1b;rims2a;rims2b;rims3;shank1;shank2;si:ch1073-450f2.1;syt12 |
| GO:0060078 | regulation of postsynaptic membrane potential | 66 | 22 | 11.78 | 0.00055 | adcyap1b;chrna10a;chrna6;chrnb2a;chrnb3a;chrnb3b;gabra1;gabra3;gabra5;gabra6b;gabrg2;glra1;glrba;glrbb;grin2aa;grin2ab;grm1a;grm1b;shank1;shank2;si:ch1073-450f2.1;si:ch73-380n15.2 |
| GO:0007623 | circadian rhythm | 64 | 22 | 11.43 | 0.01789 | arntl1a;arntl2;ciarta;ciartb;cipca;clocka;htr7c;id4;impdh2;mitfa;nfil3;nfil3-5;npas2;npy;nr1d1;nr1d4b;otx5;per1a;per3;prok1;scn1lab;si:ch211-132b12.7 |
| GO:0048167 | regulation of synaptic plasticity | 38 | 21 | 6.78 | 0.00617 | CABZ01081780.1;calb2a;calb2b;grin2aa;grin2ab;kat2a;neto1;nsmfa;nsmfb;rims1b;rims2a;rims2b;rims3;shank1;shank2;shisa7b;shisa8b;si:ch1073-450f2.1;sypa;sypb;syt12 |
| GO:0042176 | regulation of protein catabolic process | 81 | 21 | 14.46 | 0.03176 | ankib1a;apc2;csnk1e;gpc3;hecw2a;hsp90ab1;itcha;oaz2b;psmc1a;psmc1b;psmd1;psmd3;RNF14 (1 of many);stub1;timp2b;timp4.3;trib2;trib3;uqcc2;vcp;vip |
| GO:0045666 | positive regulation of neuron differentiation | 62 | 21 | 11.07 | 0.03898 | adcyap1b;ascl1a;ascl1b;brinp2;brinp3a.1;cntfr;cntn2;gprc5ba;gprc5bb;il6st;llph;mdkb;pacsin1a;plxna2;shank1;shank2;si:ch1073-450f2.1;si:ch211-194k22.8;tcf12;tenm3;zfyve27 |
| GO:0002181 | cytoplasmic translation | 59 | 20 | 10.53 | 0.00381 | atp6v0e1;cpeb1a;cpeb3;dph6;eif2s3;eif3ba;eif3ha;eif3i;eif3m;eif3s6ip;hspa14;lin28a;rpl29;rpl35a;rpl8;rpl9;rps21;rps28;rwdd1;ube2e1 |
| GO:0045765 | regulation of angiogenesis | 73 | 20 | 13.03 | 0.03705 | adgrb1a;adgrb3;aplnrb;dab2ipa;dab2ipb;iars;jcada;mt2;nme2b.1;prkd1;prok1;ptenb;sars;sox11b;tars;tmem184a;unc5b;VASH1;vash2;yjefn3 |
| GO:0042074 | cell migration involved in gastrulation | 70 | 19 | 12.5 | 0.02761 | ankrd6b;apoc1;ctnnd1;cxcl12b;dscama;esrra;fzd7a;git2a;gpc3;itgav;megf8;mmp14b;ptges;rac1a;stat3;tbx2b;vangl2;zbtb4;zeb2b |
| GO:0031018 | endocrine pancreas development | 61 | 19 | 10.89 | 0.03783 | adamts9;apc;ascl1b;c2cd4a;fscn1a;fzd2;gck;insm1a;insm1b;irx3a;kcnip3b;kcnj11;lepr;pax6b;sept15;slc30a8;sox4b;srebf1;zgc:153115 |
| GO:0043065 | positive regulation of apoptotic process | 70 | 19 | 12.5 | 0.04987 | apc;baxa;baxb;bbc3;bmf2;bnip3lb;bnip4;casp3a;casp9;dab2ipa;dab2ipb;esco2;fas;hdr;hip1;kctd13;melk;osgn1;ppifb |
| GO:0031101 | fin regeneration | 59 | 18 | 10.53 | 0.01234 | aldh1a2;chd4a;col1a1a;ctsba;cx43;esco2;hapln1a;hspd1;igf1rb;igf2b;inhbaa;kdm6bb;krt5;mmp9;nrg1;smarca4a;tubb5;vdac3 |
| GO:0051017 | actin filament bundle assembly | 55 | 17 | 9.82 | 0.00536 | add3a;arhgef10lb;arhgef11;baiap2a;dpysl3;espnla;fscn1a;fscn1b;fscn2a;fscn2b;lima1a;pfn2;pls3;rhoca;rnd1a;rnd1b;sdc4 |
| GO:0019226 | transmission of nerve impulse | 24 | 16 | 4.29 | 0.01099 | cacna1g;cacna1ha;cacna1hb;cacna1i;cacng2a;cacng3b;cacng5a;cacng7b;cacng8b;kcnab1a;kcnab2b;nfasca;scn12aa;scn1lab;scn8aa;scn8ab |
| GO:0017158 | regulation of calcium ion-dependent exocytosis | 21 | 16 | 3.75 | 0.01951 | cacna1g;cacna1ha;cacna1hb;cacna1i;napaa;napab;napba;napbb;rims1b;rims2a;rims2b;rims3;scamp5a;syt10;syt9a;syt9b |
| GO:0009636 | response to toxic substance | 63 | 16 | 11.25 | 0.02054 | abcb4;alox5ap;blmh;cat;eif2ak2;esd;fam213b;gpx3;gpx4b;gstk1;loxhd1a;pebp1;pkz;prdx1;sod1;sod2 |
| GO:0030705 | cytoskeleton-dependent intracellular transport | 56 | 15 | 10 | 0.00097 | ccdc88b;clstn1;dctn1b;hook1;kif1aa;kif1ab;kif1b;kif3a;kif5ab;kif5c;map6a;map6b;pafah1b1a;rhot1a;si:ch211-235e9.8 |
| GO:0071526 | semaphorin-plexin signaling pathway | 47 | 15 | 8.39 | 0.00323 | nrp1a;nrp2a;plxna2;plxna3;plxna4;sema3fa;sema3ga;sema3gb;sema4aa;sema4bb;sema6bb;sema6e;sema7a;si:ch211-113g11.6;si:ch211-129c21.1 |
| GO:0070121 | Kupffer's vesicle development | 52 | 15 | 9.28 | 0.03473 | atp6ap1b;atp6v1f;bbs4;bbs5;cx43;fgf8a;gpr22a;inpp5b;itgav;nkd3;rab11a;rab3ip;sdc2;tcp1;unc119b |
| GO:0009266 | response to temperature stimulus | 52 | 15 | 9.28 | 0.04259 | bhlhe41;cirbpb;clocka;cry1ba;cry2;dnaja2l;hsf2;hsp70.1;hsp70l;hspa14;hspb1;pebp1;trpm3;trpv1;ucp3 |
| GO:0031110 | regulation of microtubule polymerization or depolymerization | 31 | 14 | 5.54 | 0.00032 | apc;apc2;CABZ01118678.1;camsap3;clasp2;inpp5jb;mapre1b;mapre2;mapre3b;stmn2a;stmn2b;stmn3;stmn4;stmn4l |
| GO:0046928 | regulation of neurotransmitter secretion | 20 | 14 | 3.57 | 0.00346 | cplx2;cplx2l;cplx3b;cplx4a;cplx4b;napaa;napab;napba;napbb;rims1b;rims2a;rims2b;rims3;syt12 |
| GO:0006096 | glycolytic process | 35 | 14 | 6.25 | 0.01863 | aldoaa;aldoca;aldocb;eno1a;eno2;gck;pfklb;pfkmb;pfkpb;pgam1a;pkma;scn1lab;tpi1a;tpi1b |
| GO:0060216 | definitive hemopoiesis | 48 | 14 | 8.57 | 0.03709 | angptl1a;angptl2b;camk4;cbl;dkc1;hdac6;hsp70l;mbd3b;pbx1a;rpl22l1;rps24;sae1;scml4;tfec |
| GO:0046854 | phosphatidylinositol phosphorylation | 48 | 14 | 8.57 | 0.03709 | cish;impa1;impa2;pi4k2a;pi4kaa;pik3cb;pik3r1;pik3r3a;pik3r3b;pip4k2ab;pip5k1cb;pip5kl1;socs2;zgc:158659 |
| GO:0006414 | translational elongation | 42 | 13 | 7.5 | 0.03805 | dph6;eef1a1b;eef1a1l2;eef1a2;eef1b2;eef1db;eef1g;eef2a.2;eef2b;eif5a;eif5a2;gtpbp1;gtpbp2b |

| | | | | | | |
|---|---|---|---|---|---|---|
| GO:0015991 | ATP hydrolysis coupled proton transport | 35 | 12 | 6.25 | 0.01482 | atp1a1b;atp1a3a;atp6ap1b;atp6ap1la;atp6v0a1b;atp6v0a2a;atp6v0e1;atp6v1ba;atp6v1f;atp6v1h;atpv0e2;tcirg1b |
| GO:0048048 | embryonic eye morphogenesis | 36 | 12 | 6.43 | 0.02238 | cct3;hs6st2;mfap2;plxna2;sfrp1a;sfrp5;smarca4a;sox11a;sox11b;sox4a;sox4b;vangl2 |
| GO:0055075 | potassium ion homeostasis | 28 | 12 | 5 | 0.03705 | atp1a1b;atp1a3a;atp1b1b;atp1b2a;atp1b3a;atp1b3b;slc12a2;slc12a4;slc12a5a;slc12a5b;slc12a7a;slc12a9 |
| GO:0033334 | fin morphogenesis | 51 | 12 | 9.11 | 0.03729 | apc;bmp4;col1a1a;cx43;fras1;frem1a;frem2a;grip2a;lama5;mmp9;pi4kaa;tll1 |
| GO:0048791 | calcium ion-regulated exocytosis of neurotransmitter | 19 | 11 | 3.39 | 0.00011 | rims1b;rims2a;rims2b;rims3;syt11b;syt1a;syt2a;syt4;syt5a;syt5b;syt7b |
| GO:0048488 | synaptic vesicle endocytosis | 21 | 11 | 3.75 | 0.00032 | dnm1b;dnm3a;nlgn1;nlgn2a;nlgn2b;nlgn3a;si:ch211-233a24.2;syt1a;syt2a;syt5a;syt5b |
| GO:0007019 | microtubule depolymerization | 23 | 11 | 4.11 | 0.01108 | apc;apc2;CABZ01118678.1;camsap3;ccsapb;katnb1;stmn2a;stmn2b;stmn3;stmn4;stmn4l |
| GO:0048843 | negative regulation of axon extension involved in axon guidance | 31 | 11 | 5.54 | 0.01472 | robo2;sema3fa;sema3ga;sema3gb;sema4aa;sema4bb;sema6bb;sema6e;sema7a;si:ch211-113g11.6;si:ch211-129c21.1 |
| GO:0048013 | ephrin receptor signaling pathway | 33 | 11 | 5.89 | 0.03808 | CT990561.1;efna1b;efna2a;efna3a;ek1;epha4b;epha6;ephb1;ephb2a;ephb6;si:dkeyp-9d4.3 |
| GO:0008286 | insulin receptor signaling pathway | 36 | 11 | 6.43 | 0.03981 | cish;enpp1;foxo3a;igf1rb;insra;irs2a;pik3r1;pik3r3a;pik3r3b;si:ch211-284e13.4;socs2 |
| GO:0097120 | receptor localization to synapse | 22 | 11 | 3.93 | 0.04254 | cacng2a;cacng3b;cacng5a;cacng7b;cacng8b;dlg1l;dlg3;dlg4a;gphnb;kif5ab;kif5c |
| GO:2000311 | regulation of AMPA receptor activity | 19 | 10 | 3.39 | 0.00062 | cacng2a;cacng3b;cacng5a;cacng7b;cacng8b;shank1;shank2;shisa7b;shisa8b;si:ch1073-450f2.1 |
| GO:0050919 | negative chemotaxis | 30 | 10 | 5.36 | 0.03058 | sema3fa;sema3ga;sema3gb;sema4aa;sema4bb;sema6bb;sema6e;sema7a;si:ch211-113g11.6;si:ch211-129c21.1 |
| GO:0051968 | positive regulation of synaptic transmission, glutamatergic | 18 | 9 | 3.21 | 0.00187 | adcyap1b;cacng2a;cacng3b;cacng5a;cacng7b;cacng8b;shank1;shank2;si:ch1073-450f2.1 |
| GO:0048813 | dendrite morphogenesis | 23 | 9 | 4.11 | 0.01117 | hecw2a;prmt8b;rbpms2a;rbpms2b;shank1;shank2;si:ch1073-450f2.1;tenm3;tmem106bb |
| GO:0060291 | long-term synaptic potentiation | 20 | 9 | 3.57 | 0.01457 | CABZ01081780.1;calb2a;calb2b;grin2aa;grin2ab;shank1;shank2;si:ch1073-450f2.1;syt12 |
| GO:0044772 | mitotic cell cycle phase transition | 83 | 9 | 14.82 | 0.03204 | anapc15;bub1bb;cdk1;chmp4bb;clasp2;ensaa;lzts1;rb1;spdl1 |
| GO:0031103 | axon regeneration | 22 | 9 | 3.93 | 0.03219 | ascl1a;cnp;cntfr;cntn2;dpysl2b;epha4b;il6st;lgmn;lingo1a |
| GO:0070593 | dendrite self-avoidance | 13 | 8 | 2.32 | 0.00056 | bsg;cntn3a.1;dscama;nexn;nptna;nptnb;robo3;robo4 |
| GO:0032922 | circadian regulation of gene expression | 13 | 8 | 2.32 | 0.00056 | ciarta;ciartb;clocka;nfil3-5;nr1d1;otx5;per1a;per3 |
| GO:0000470 | maturation of LSU-rRNA | 18 | 8 | 3.21 | 0.00626 | ftsj3;las1l;nhp2;rpl10a;rpl35;rpl7;rsl1d1;snu13b |
| GO:2000300 | regulation of synaptic vesicle exocytosis | 10 | 8 | 1.79 | 0.01115 | napaa;napab;napba;napbb;rims1b;rims2a;rims2b;rims3 |
| GO:0043200 | response to amino acid | 24 | 8 | 4.29 | 0.01204 | glra1;glra2;glra3;glra4b;glrba;glrbb;rragca;sesn2 |
| GO:0006171 | cAMP biosynthetic process | 26 | 8 | 4.64 | 0.01761 | adcy1b;adcy2a;adcy3a;adcy6a;adcy7;cap2;rxfp2a;si:dkey-206f10.1 |
| GO:0006817 | phosphate ion transport | 17 | 8 | 3.04 | 0.0318 | ankha;slc17a7a;slc17a7b;slc20a1a;slc20a1b;slc20a2;slc25a14;slc34a2a |
| GO:1901385 | regulation of voltage-gated calcium channel activity | 12 | 8 | 2.14 | 0.03849 | cacnb1;cacnb2a;cacnb3a;cacnb3b;cbarpb;gem;rem1;rrad |
| GO:0006884 | cell volume homeostasis | 23 | 8 | 4.11 | 0.03986 | lrrc8aa;lrrc8c;slc12a2;slc12a4;slc12a5a;slc12a5b;slc12a7a;slc12a9 |
| GO:0046847 | filopodium assembly | 13 | 7 | 2.32 | 0.00092 | arhgef9b;dpysl3;fmnl3;gpm6ab;robo4;srgap2;trpc1 |
| GO:0007157 | heterophilic cell-cell adhesion via plasma membrane cell adhesion molecules | 12 | 7 | 2.14 | 0.00196 | igsf21a;igsf21b;pvrl2l;si:ch73-22o12.1;tenm1;TENM2;tenm3 |
| GO:0048168 | regulation of neuronal synaptic plasticity | 10 | 7 | 1.79 | 0.00275 | neto1;nsmfa;nsmfb;shisa7b;shisa8b;sypa;sypb |
| GO:0045879 | negative regulation of smoothened signaling pathway | 14 | 7 | 2.5 | 0.0061 | cdon;gatad2b;gpc3;sox11a;sox11b;sox4a;sox4b |
| GO:0009648 | photoperiodism | 15 | 7 | 2.68 | 0.00626 | arntl1a;arntl1b;arntl2;clocka;npas2;per1a;per3 |
| GO:0070654 | sensory epithelium regeneration | 15 | 7 | 2.68 | 0.00969 | apobec2b;ascl1a;fzd2;hbegfa;insm1a;mdka;tgif1 |
| GO:1903670 | regulation of sprouting angiogenesis | 22 | 7 | 3.93 | 0.01969 | aplnrb;iars;jcada;mt2;sars;sox11b;tars |
| GO:0043113 | receptor clustering | 14 | 7 | 2.5 | 0.02054 | agrn;dlg1l;dlg3;dlg4a;gphnb;grna;slc9a3r1a |
| GO:0055064 | chloride ion homeostasis | 17 | 7 | 3.04 | 0.02107 | calca;slc12a2;slc12a4;slc12a5a;slc12a5b;slc12a7a;slc12a9 |
| GO:0046686 | response to cadmium ion | 17 | 7 | 3.04 | 0.02107 | baxa;gadd45ba;hsp70.1;hsp70l;jun;mt2;sod2 |
| GO:0099590 | neurotransmitter receptor internalization | 17 | 7 | 3.04 | 0.03224 | cacng2a;cacng3b;cacng5a;cacng7b;cacng8b;dnm1b;dnm3a |
| GO:0000027 | ribosomal large subunit assembly | 19 | 7 | 3.39 | 0.0394 | mrto4;nop53;ppan;rpl12;rpl23a;rpl5a;rpl5b |
| GO:0051932 | synaptic transmission, GABAergic | 10 | 6 | 1.79 | 0.00092 | gabra1;gabra3;gabra5;gabra6b;gabrg2;plcl2 |
| GO:0060012 | synaptic transmission, glycinergic | 9 | 6 | 1.61 | 0.00165 | glra1;glra2;glra3;glra4b;glrba;glrbb |
| GO:0051597 | response to methylmercury | 10 | 6 | 1.79 | 0.0035 | baxa;gadd45ba;jun;mt2;sod1;sod2 |
| GO:0030032 | lamellipodium assembly | 16 | 6 | 2.86 | 0.0112 | ablim1b;ablim3;dmtn;robo4;ssh1b;ssh2b |
| GO:0061386 | closure of optic fissure | 13 | 6 | 2.32 | 0.0176 | sfrp1a;sfrp5;sox11a;sox11b;sox4a;sox4b |
| GO:0008345 | larval locomotory behavior | 13 | 6 | 2.32 | 0.0176 | psmd3;ryr1b;scn8aa;slc1a2b;sncb;sncgb |

| | | | | | | |
|---|---|---|---|---|---|---|
| GO:0036376 | sodium ion export across plasma membrane | 13 | 6 | 2.32 | 0.0176 | atp1a1b;atp1a3a;atp1b1b;atp1b2a;atp1b3a;atp1b3b |
| GO:0030007 | cellular potassium ion homeostasis | 13 | 6 | 2.32 | 0.0176 | atp1a1b;atp1a3a;atp1b1b;atp1b2a;atp1b3a;atp1b3b |
| GO:0098970 | postsynaptic neurotransmitter receptor diffusion trapping | 13 | 6 | 2.32 | 0.0176 | cacng2a;cacng3b;cacng5a;cacng7b;cacng8b;gphnb |
| GO:0043280 | positive regulation of cysteine-type endopeptidase activity involved in apoptotic process | 24 | 6 | 4.29 | 0.02063 | baxa;baxb;bbc3;esco2;hdr;hip1 |
| GO:0006883 | cellular sodium ion homeostasis | 14 | 6 | 2.5 | 0.02623 | atp1a1b;atp1a3a;atp1b1b;atp1b2a;atp1b3a;atp1b3b |
| GO:0099054 | presynapse assembly | 9 | 6 | 1.61 | 0.03181 | bdnf;mecp2;nlgn1;nlgn2a;nlgn2b;nlgn3a |
| GO:0007196 | adenylate cyclase-inhibiting G-protein coupled glutamate receptor signaling pathway | 5 | 5 | 0.89 | 0.00018 | grm4;grm6a;grm6b;grm8a;grm8b |
| GO:0045956 | positive regulation of calcium ion-dependent exocytosis | 7 | 5 | 1.25 | 0.00276 | cacna1g;cacna1ha;cacna1hb;cacna1i;scamp5a |
| GO:0098700 | neurotransmitter loading into synaptic vesicle | 7 | 5 | 1.25 | 0.00435 | slc17a6b;slc17a7a;slc17a7b;slc18a2;slc32a1 |
| GO:0042984 | regulation of amyloid precursor protein biosynthetic process | 7 | 5 | 1.25 | 0.00568 | itm2ca;itm2cb;necab1;necab2;necab3 |
| GO:0002138 | retinoic acid biosynthetic process | 8 | 5 | 1.43 | 0.00568 | apc;hmx4;rbp1;rdh10a;rdh10b |
| GO:0000160 | phosphorelay signal transduction system | 8 | 5 | 1.43 | 0.00627 | kcnh1a;kcnh3;kcnh4b;kcnh5a;kcnh5b |
| GO:0031023 | microtubule organizing center organization | 48 | 5 | 8.57 | 0.03199 | calm1b;calm3a;cetn2;clasp2;pafah1b1a |
| GO:0030878 | thyroid gland development | 11 | 5 | 1.96 | 0.03228 | bcl2l1;fgf8a;thraa;thrab;thrb |
| GO:0034625 | fatty acid elongation, monounsaturated fatty acid | 12 | 5 | 2.14 | 0.04738 | elovl1a;elovl1b;elovl4b;elovl6;elovl8a |
| GO:0034626 | fatty acid elongation, polyunsaturated fatty acid | 12 | 5 | 2.14 | 0.04738 | elovl1a;elovl1b;elovl4b;elovl6;elovl8a |
| GO:0046855 | inositol phosphate dephosphorylation | 12 | 5 | 2.14 | 0.04738 | impa1;impa2;inpp5b;inpp5jb;ptenb |
| GO:0019367 | fatty acid elongation, saturated fatty acid | 12 | 5 | 2.14 | 0.04738 | elovl1a;elovl1b;elovl4b;elovl6;elovl8a |
| GO:0098943 | neurotransmitter receptor transport, postsynaptic endosome to lysosome | 12 | 5 | 2.14 | 0.04738 | cacng2a;cacng3b;cacng5a;cacng7b;cacng8b |
| GO:0010807 | regulation of synaptic vesicle priming | 4 | 4 | 0.71 | 0.00101 | napaa;napab;napba;napbb |
| GO:0035494 | SNARE complex disassembly | 4 | 4 | 0.71 | 0.00101 | napaa;napab;napba;napbb |
| GO:0045162 | clustering of voltage-gated sodium channels | 6 | 4 | 1.07 | 0.0112 | gldn;ndrg4;nsfa;snap25b |
| GO:0021634 | optic nerve formation | 6 | 4 | 1.07 | 0.0112 | klf6a;klf7b;smarca4a;tuba1a |
| GO:0006177 | GMP biosynthetic process | 8 | 4 | 1.43 | 0.0197 | gmps;hprt1l;impdh1b;impdh2 |
| GO:0007158 | neuron cell-cell adhesion | 7 | 4 | 1.25 | 0.02245 | nlgn1;nlgn2a;nlgn2b;nlgn3a |
| GO:0097104 | postsynaptic membrane assembly | 7 | 4 | 1.25 | 0.02245 | nlgn1;nlgn2a;nlgn2b;nlgn3a |
| GO:0097105 | presynaptic membrane assembly | 7 | 4 | 1.25 | 0.02245 | nlgn1;nlgn2a;nlgn2b;nlgn3a |
| GO:0021772 | olfactory bulb development | 7 | 4 | 1.25 | 0.02245 | anos1a;enpp1;msi2b;ptprsa |
| GO:0051965 | positive regulation of synapse assembly | 7 | 4 | 1.25 | 0.02245 | clstn1;clstn2;clstn3;lrrc4bb |
| GO:0046548 | retinal rod cell development | 7 | 4 | 1.25 | 0.02245 | rpgrip1;sox11a;sox11b;tbx2b |
| GO:0003208 | cardiac ventricle morphogenesis | 9 | 4 | 1.61 | 0.03185 | fgf8a;fhl1a;ilk;nrg1 |
| GO:0007026 | negative regulation of microtubule depolymerization | 8 | 4 | 1.43 | 0.03864 | apc;apc2;CABZ01118678.1;camsap3 |
| GO:0060999 | positive regulation of dendritic spine development | 8 | 4 | 1.43 | 0.03864 | llph;shank1;shank2;si:ch1073-450f2.1 |
| GO:0043652 | engulfment of apoptotic cell | 8 | 4 | 1.43 | 0.03864 | havcr1;rac3a;rac3b;rhobtb1 |
| GO:0006855 | drug transmembrane transport | 8 | 4 | 1.43 | 0.03864 | abcg2c;abcg2d;slc18a2;slc18a3a |
| GO:0051591 | response to cAMP | 9 | 4 | 1.61 | 0.04273 | enpp1;jun;junba;pebp1 |
| GO:0070884 | regulation of calcineurin-NFAT signaling cascade | 3 | 3 | 0.54 | 0.00569 | rcan1a;rcan2;rcan3 |
| GO:0060509 | Type I pneumocyte differentiation | 3 | 3 | 0.54 | 0.00569 | thraa;thrab;thrb |
| GO:1901842 | negative regulation of high voltage-gated calcium channel activity | 3 | 3 | 0.54 | 0.00569 | gem;rem1;rrad |
| GO:0010996 | response to auditory stimulus | 4 | 3 | 0.71 | 0.0197 | kcnma1a;otofa;wrb |
| GO:0036368 | cone photoresponse recovery | 4 | 3 | 0.71 | 0.0197 | eml1;rcvrn3;rcvrna |
| GO:0035385 | Roundabout signaling pathway | 4 | 3 | 0.71 | 0.0197 | robo2;robo3;zgc:77784 |
| GO:0016199 | axon midline choice point recognition | 4 | 3 | 0.71 | 0.0197 | adcy1b;robo2;robo3 |

| | | | | | | |
|---|---|---|---|---|---|---|
| GO:0000447 | endonucleolytic cleavage in ITS1 to separate SSU-rRNA from 5.8S rRNA and LSU-rRNA from tricistronic rRNA transcript (SSU-rRNA, 5.8S rRNA, LSU-rRNA) | 4 | 3 | 0.71 | 0.0197 | abt1;rps21;rpsa |
| GO:0019388 | galactose catabolic process | 6 | 3 | 1.07 | 0.03186 | galm;pgm1;pgm5 |
| GO:0071376 | cellular response to corticotropin-releasing hormone stimulus | 5 | 3 | 0.89 | 0.04274 | crhr1;nr4a1;nr4a3 |
| GO:0042264 | peptidyl-aspartic acid hydroxylation | 5 | 3 | 0.89 | 0.04274 | asph;asphd1;asphd2 |
| GO:0010998 | regulation of translational initiation by eIF2 alpha phosphorylation | 5 | 3 | 0.89 | 0.04274 | eif2ak1;eif2ak2;pkz |
| GO:2000463 | positive regulation of excitatory postsynaptic potential | 5 | 3 | 0.89 | 0.04274 | shank1;shank2;si:ch1073-450f2.1 |
| GO:0097107 | postsynaptic density assembly | 5 | 3 | 0.89 | 0.04274 | shank1;shank2;si:ch1073-450f2.1 |
| GO:0097264 | self proteolysis | 5 | 3 | 0.89 | 0.04274 | TENM2;tenm3;tenm4 |
| GO:1904825 | protein localization to microtubule plus-end | 5 | 3 | 0.89 | 0.04274 | mapre1b;mapre2;mapre3b |
| GO:0045634 | regulation of melanocyte differentiation | 2 | 2 | 0.36 | 0.03187 | ctbp2a;hipk2 |
| GO:0051012 | microtubule sliding | 2 | 2 | 0.36 | 0.03187 | pafah1b1a;pafah1b1b |
| GO:0045905 | positive regulation of translational termination | 2 | 2 | 0.36 | 0.03187 | eif5a;eif5a2 |
| GO:0048025 | negative regulation of mRNA splicing, via spliceosome | 2 | 2 | 0.36 | 0.03187 | rbmx;ybx1 |
| GO:0038026 | reelin-mediated signaling pathway | 2 | 2 | 0.36 | 0.03187 | dab2ipa;dab2ipb |
| GO:1904059 | regulation of locomotor rhythm | 2 | 2 | 0.36 | 0.03187 | nr1d1;scn1lab |
| GO:0044210 | 'de novo' CTP biosynthetic process | 2 | 2 | 0.36 | 0.03187 | ctps1a;ctps1b |
| GO:0036071 | N-glycan fucosylation | 2 | 2 | 0.36 | 0.03187 | fut8a;fut8b |
| GO:0002154 | thyroid hormone mediated signaling pathway | 2 | 2 | 0.36 | 0.03187 | thraa;thrb |
| GO:0035024 | negative regulation of Rho protein signal transduction | 2 | 2 | 0.36 | 0.03187 | akap12b;kctd13 |
| GO:0030948 | negative regulation of vascular endothelial growth factor receptor signaling pathway | 2 | 2 | 0.36 | 0.03187 | dab2ipa;dab2ipb |
| GO:0008295 | spermidine biosynthetic process | 2 | 2 | 0.36 | 0.03187 | amd1;srm |
| GO:0001778 | plasma membrane repair | 2 | 2 | 0.36 | 0.03187 | anxa6;dysf |
| GO:0006452 | translational frameshifting | 2 | 2 | 0.36 | 0.03187 | eif5a;eif5a2 |
| GO:0033578 | protein glycosylation in Golgi | 2 | 2 | 0.36 | 0.03187 | fut8a;fut8b |
| GO:0021766 | hippocampus development | 2 | 2 | 0.36 | 0.03187 | enpp1;pebp1 |
| GO:0048240 | sperm capacitation | 2 | 2 | 0.36 | 0.03187 | abhd2a;pebp1 |
| GO:0009098 | leucine biosynthetic process | 2 | 2 | 0.36 | 0.03187 | bcat1;bcat2 |
| GO:0009099 | valine biosynthetic process | 2 | 2 | 0.36 | 0.03187 | bcat1;bcat2 |
| GO:0051969 | regulation of transmission of nerve impulse | 2 | 2 | 0.36 | 0.03187 | kcnab1a;kcnab2b |
| GO:0009449 | gamma-aminobutyric acid biosynthetic process | 2 | 2 | 0.36 | 0.03187 | gad1b;gad2 |
| GO:0006696 | ergosterol biosynthetic process | 2 | 2 | 0.36 | 0.03187 | acat2;fdft1 |
| GO:0015820 | leucine transport | 2 | 2 | 0.36 | 0.03187 | slc6a15;slc6a17 |
| GO:0000461 | endonucleolytic cleavage to generate mature 3'-end of SSU-rRNA from (SSU-rRNA, 5.8S rRNA, LSU-rRNA) | 2 | 2 | 0.36 | 0.03187 | rps21;rpsa |
| GO:0035553 | oxidative single-stranded RNA demethylation | 2 | 2 | 0.36 | 0.03187 | alkbh5;fto |
| GO:0060290 | transdifferentiation | 2 | 2 | 0.36 | 0.03187 | insm1a;insm1b |

**Table S10. RNA sample quality control.**

| Sample Name | Nanodrop | | | | | Bioanalyzer | | | | | | Qubit | |
|---|---|---|---|---|---|---|---|---|---|---|---|---|---|
| | 260/280 | 260/230 | Nanodrop Conc (ng/ul) | Dilution factor | Nano final (ng/ul) | RIN | From (bp) | To (bp) | Total average size | size distribution in CV (%) | Bionalyzer Conc. (ng/ul) | Qubit ng/ul | Total Qubit ug |
| 0RNA_1 | 2.18 | 0.42 | 2.6 | 20 | 52 | 8.7 | 179 | 1328 | 382 | 35.6 | 14.952 | 52.2 | 1.044 |
| 0RNA_2 | 2.39 | 1.15 | 2.5 | 20 | 50 | 8.6 | 156 | 1351 | 372 | 33.7 | 38.755 | 61 | 1.22 |
| 0RNA_3 | 1.65 | 0.8 | 2.8 | 20 | 56 | 8.6 | 149 | 1358 | 360 | 29.5 | 23.6 | 53.2 | 1.064 |
| 2RNA_1 | 1.65 | 1.68 | 4 | 20 | 80 | 8.6 | 182 | 1345 | 373 | 33.6 | 56.448 | 96.6 | 1.932 |
| 2RNA_2 | 1.72 | 0.49 | 6.5 | 10 | 65 | 7.8 | 171 | 1354 | 388 | 37.9 | 18.571 | 48.6 | 0.972 |
| 2RNA_3 | 2.11 | 2.01 | 1.96 | 20 | 39.2 | 8.6 | 174 | 1328 | 363 | 30.9 | 62.109 | 49.6 | 0.992 |
| 4RNA_1 | 1.76 | 0.35 | 3.8 | 20 | 76 | 8.2 | 171 | 1321 | 380 | 38.5 | 13.475 | 79 | 1.58 |
| 4RNA_2 | 2.1 | 2.11 | 2.12 | 20 | 42.4 | 8.5 | 170 | 1353 | 385 | 38.9 | 82.079 | 60.2 | 1.204 |
| 4RNA_3 | 1.88 | 1.03 | 3.33 | 20 | 66.6 | 7.7 | 180 | 1323 | 410 | 40.3 | 41.509 | 45.6 | 0.912 |
| 7RNA_1 | 1.93 | 0.38 | 48.9 | 20 | 48.9 | 7.8 | 141 | 1331 | 404 | 42.2 | 16.036 | 62.2 | 1.244 |
| 7RNA_2 | 1.9 | 0.73 | 58.3 | 20 | 58.3 | 8.3 | 183 | 1366 | 413 | 40.5 | 29.565 | 57.8 | 1.156 |
| 7RNA_3 | 1.99 | 0.36 | 57.3 | 20 | 57.3 | 8.6 | 183 | 1341 | 398 | 39.2 | 14.112 | 53.2 | 1.064 |
| 12RNA_1 | 1.77 | 0.15 | 4.9 | 10 | 49 | 8.6 | 186 | 1340 | 382 | 37 | 5.55 | 45 | 0.99 |
| 12RNA_2 | 1.91 | 0.68 | 49.9 | 20 | 49.9 | 8 | 173 | 1337 | 387 | 38 | 25.84 | 51.4 | 1.028 |
| 12RNA_3 | 2.18 | 1.7 | 5.72 | 10 | 57.2 | 8.7 | 187 | 1335 | 400 | 37.6 | 63.92 | 67.2 | 1.344 |

**Table S11. ATAC-seq library sample quality control.**

| Sample Name | Nanodrop | | | Bioanalyzer | | | | Qubit | |
|---|---|---|---|---|---|---|---|---|---|
| | 260/280 | 260/230 | Nanodrop Conc (ng/ul) | Bio Avg fragment | Molarity (nmol/L) | Average size | Bionalyzer Conc. (ng/ul) | Qubit ng/ul | Total Qubit ug |
| 0dpiATAC_1 | 2.05 | 1.86 | 30.3 | 384 | 48.4 | 338 | 4.327 | 16.8 | 252 |
| 0dpiATAC_2 | 2.08 | 0.8 | 23.7 | 503 | 20.2 | 536 | 6.111 | 10.5 | 157.5 |
| 0dpiATAC_3 | 1.89 | 1.75 | 61.2 | 396 | 49.4 | 301 | 32.282 | 27 | 405 |
| 2dpiATAC_1 | 1.86 | 1.45 | 75.6 | 359 | 69.5 | 351 | 5.156 | 25.4 | 381 |
| 2dpiATAC_2 | 1.93 | 1.38 | 45.7 | 513 | 10.2 | 587 | 3.37 | 10.4 | 156 |
| 2dpiATAC_3 | 2.11 | 2.01 | 39.2 | 352 | 56.4 | 339 | 9.158 | 18 | 270 |
| 4dpiATAC_1 | 1.81 | 0.61 | 50.71 | 403 | 18.6 | 361 | 7.845 | 11.3 | 169.5 |
| 4dpiATAC_2 | 2.12 | 0.56 | 48.12 | 397 | 27.3 | 363 | 61.829 | 6.62 | 99.3 |
| 4dpiATAC_3 | 1.86 | 1.74 | 37.9 | 365 | 53.7 | 286 | 5.698 | 23.2 | 348 |
| 7dpiATAC_2 | 1.9 | 0.73 | 58.3 | 492 | 17.3 | 573 | 6.33 | 11.3 | 169.5 |
| 7dpiATAC_3 | 1.91 | 2.06 | 39.6 | 367 | 48.1 | 307 | 4.975 | 16.9 | 253.5 |
| 12dpiATAC_1 | 2.02 | 1.82 | 21.5 | 467 | 33.7 | 485 | 30.625 | 17.2 | 258 |
| 12dpiATAC_2 | 1.92 | 0.7 | 56.44 | 363 | 63.9 | 350 | 6.925 | 28 | 420 |
| 12dpiATAC_3 | 2.12 | 0.66 | 56.5 | 364 | 85.4 | 326 | 6.165 | 26 | 390 |

**Table S12. PCR primers for ATAC-seq libraries based on Nextera indices .**

| Sample Name | Primer Sequences | Index |
|---|---|---|
| 0dpiATAC_1 | **CAAGCAGAAGACGGCATACGAGATTCGCCTTAGTCTCGTGGGCTCGGAGATGT** | Ad2.1_TAAGGCGA |
| 0dpiATAC_2 | **CAAGCAGAAGACGGCATACGAGATTTCTGCCTGTCTCGTGGGCTCGGAGATGT** | Ad2.3_AGGCAGAA |
| 0dpiATAC_3 | **CAAGCAGAAGACGGCATACGAGATGTAGAGAGGTCTCGTGGGCTCGGAGATGT** | Ad2.7_CTCTCTAC |
| 2dpiATAC_1 | **CAAGCAGAAGACGGCATACGAGATCAGCCTCGGTCTCGTGGGCTCGGAGATGT** | Ad2.10_CGAGGCTG |
| 2dpiATAC_2 | **CAAGCAGAAGACGGCATACGAGATTGCCTCTTGTCTCGTGGGCTCGGAGATGT** | Ad2.11_AAGAGGCA |
| 2dpiATAC_3 | **CAAGCAGAAGACGGCATACGAGATTCCTCTACGTCTCGTGGGCTCGGAGATGT** | Ad2.12_GTAGAGGA |
| 4dpiATAC_1 | **CAAGCAGAAGACGGCATACGAGATGCTCAGGAGTCTCGTGGGCTCGGAGATGT** | Ad2.4_TCCTGAGC |
| 4dpiATAC_2 | **CAAGCAGAAGACGGCATACGAGATAGGAGTCCGTCTCGTGGGCTCGGAGATGT** | Ad2.5_GGACTCCT |
| 4dpiATAC_3 | **CAAGCAGAAGACGGCATACGAGATCATGCCTAGTCTCGTGGGCTCGGAGATGT** | Ad2.6_TAGGCATG |
| 7dpiATAC_2 | **CAAGCAGAAGACGGCATACGAGATCCTCTCTGGTCTCGTGGGCTCGGAGATGT** | Ad2.8_CAGAGAGG |
| 7dpiATAC_3 | **CAAGCAGAAGACGGCATACGAGATAGCGTAGCGTCTCGTGGGCTCGGAGATGT** | Ad2.9_GCTACGCT |
| 12dpiATAC_1 | **CAAGCAGAAGACGGCATACGAGATATCACGACGTCTCGTGGGCTCGGAGATGT** | Ad2.13_GTCGTGAT |
| 12dpiATAC_2 | **CAAGCAGAAGACGGCATACGAGATACAGTGGTGTCTCGTGGGCTCGGAGATGT** | Ad2.14_ACCACTGT |
| 12dpiATAC_3 | **CAAGCAGAAGACGGCATACGAGATCAGATCCAGTCTCGTGGGCTCGGAGATGT** | Ad2.15_TGGATCTG |
